# Acquired stress resilience through bacteria-to-nematode horizontal gene transfer

**DOI:** 10.1101/2023.08.20.554039

**Authors:** Taruna Pandey, Chinmay Kalluraya, Bingying Wang, Ting Xu, Xinya Huang, Shouhong Guang, Matthew D. Daugherty, Dengke K. Ma

**Affiliations:** Cardiovascular Research Institute and Department of Physiology, University of California San Francisco, San Francisco, USA; Department of Molecular Biology, University of California, San Diego, San Diego, USA; The USTC RNA Institute, Ministry of Education Key Laboratory for Membraneless Organelles & Cellular Dynamics, School of Life Sciences, Division of Life Sciences and Medicine, Department of Obstetrics and Gynecology, The First Affiliated Hospital of USTC, Biomedical Sciences and Health Laboratory of Anhui Province, University of Science and Technology of China, Hefei, Anhui, China; Innovative Genomics Institute, University of California, Berkeley, USA

## Abstract

Natural selection drives acquisition of organismal resilience traits to protect against adverse environments. Horizontal gene transfer (HGT) is an important evolutionary mechanism for the acquisition of novel traits, including metazoan acquisition of functions in immunity, metabolism, and reproduction via interdomain HGT (iHGT) from bacteria. We report that the nematode gene *rml-3*, which was acquired by iHGT from bacteria, enables exoskeleton resilience and protection against environmental toxins in *C. elegans*. Phylogenetic analysis reveals that diverse nematode RML-3 proteins form a single monophyletic clade most highly similar to bacterial enzymes that biosynthesize L-rhamnose to build cell wall polysaccharides. *C. elegans rml-3* is regulated in developing seam cells by heat stress and stress-resistant dauer stage. Importantly, *rml-3* deficiency impairs cuticle integrity, barrier functions and organismal stress resilience, phenotypes that are rescued by exogenous L-rhamnose. We propose that iHGT of an ancient bacterial *rml-3* homolog enables L-rhamnose biosynthesis in nematodes that facilitates cuticle integrity and organismal resilience in adaptation to environmental stresses during evolution. These findings highlight the remarkable contribution of iHGT on metazoan evolution that is conferred by the domestication of bacterial genes.

## Introduction

Genetic information is primarily transmitted vertically from parents to offspring through sexual or asexual reproduction. However, genes can also be transmitted from one organism to a non-offspring organism via horizontal gene transfer (HGT). While HGT is widespread among bacteria, interdomain HGT (iHGT) between bacteria and eukaryotes is less common. In particular, iHGT from bacteria into metazoans is expected to be very rare, as long-term functional acquisition requires a transferred bacterial gene to integrate into a metazoan germline, followed by subsequent gene expression and a selective benefit to the recipient metazoan during evolution. Despite this expected rarity, cases of iHGT have been strongly suggested by phylogenetic analysis for several lineage-specific metazoan functions (Zhaxybayeva & Doolittle, 2011; Husnik & McCutcheon, 2018; Dunning Hotopp, 2011; Crisp *et al*, 2015). Examples include metazoan domestication of bacterial genes that allow nutrient intake, metabolic functions, reproductive behaviors and immune functions that confer defense against pathogens and predators (Metcalf *et al*, 2014; Wybouw *et al*, 2014; Chou *et al*, 2015; Verster *et al*, 2019; Morehouse *et al*, 2020; Bernheim *et al*, 2021; Wheeler *et al*, 2013; Ferguson *et al*, 2011; Luan *et al*, 2015; Schönknecht *et al*, 2013; Scholl *et al*, 2003; Danchin *et al*, 2010; Acuña *et al*, 2012; Mayer *et al*, 2011; Pauchet & Heckel, 2013).

Nematodes are the most abundant species of metazoans on Earth and crucial members of terrestrial ecosystems (van den Hoogen *et al*, 2019; Haag *et al*, 2018; Sommer & Bumbarger, 2012). In nature, nematodes occupy diverse ecological niches and consequently evolved many unique strategies to survive under conditions of adverse environmental stresses. For example, nematodes can deploy remarkable plasticity by shifting to the dauer stage of reproductive arrest during unfavorable environmental conditions and entering anhydrobiosis to survive extreme desiccation (Fielenbach & Antebi, 2008; McGill *et al*, 2015). Nematode genomes encode many enzymes that function to detoxify environmental toxins and genetic programs that enhance resilience to specific environmental stresses (Wang *et al*, 2022; Rodriguez *et al*, 2013). Additionally, nematodes withstand harsh environments physically aided by the exoskeleton, with specialized layers of cuticle that forms the barrier between the animal and its environment (Johnstone, 1994). In the cuticle, extracellular matrix biomolecules can be extensively cross-linked (e.g. by collagen) and modified by various types of lipid- and glycan-conjugates to enable cuticle integrity and barrier functions.

Nematodes are also unique among known metazoans in that they encode an L-rhamnose biosynthetic pathway. This pathway is widespread in prokaryotes and plants, where L-rhamnose forms the constituent of lipopolysaccharide (LPS) of diverse bacteria (Mistou *et al*, 2016), the cell wall and specialized metabolites of plants (Jiang *et al*, 2021), and the cell wall of pathogenic fungi (Martinez *et al*, 2012). Strikingly, among metazoans, the production of L-rhamnose has only been described in nematodes through the activities of four separate enzymes, RML-1 through RML-4, which are expressed during formation of the *C. elegans* cuticle (Feng *et al*, 2016). Nonetheless, the evolutionary origins and functional relevance of key enzymes in the nematode-specific L-rhamnose pathway remained unresolved.

In this work, we used phylogenetic methods to demonstrate that one of the key enzymes of the nematode L-rhamnose biosynthetic pathway, RML-3, arose from an iHGT event from bacteria. We identified RML-3 protein sequences in a diverse range of nematodes that form a single highly supported monophyletic clade most similar to bacterial dTDP-4-dehydrorhamnose-3-5-epimerase enzymes based on multiple phylogenetic reconstruction methods. In the model organism nematode *C. elegans*, we found that *rml-3* is expressed and dynamically regulated by stress in developing seam cells and hypoderm. Functional analyses revealed the critical roles of RML-3 in cuticle integrity, supporting the evolutionarily acquired L-rhamnose biosynthetic capacity in nematodes and exoskeleton stress resilience through iHGT from bacteria.

## Results

### Nematodes acquired a bacteria-derived gene in the L-rhamnose biosynthetic pathway

To understand the atypical phylogenetic distribution of the L-rhamnose biosynthetic pathway across the tree of life, we searched for patterns of conservation among each constituent enzyme. In plants and fungi, L-rhamnose is produced via the UDP pathway, while in bacteria and nematodes, the RML pathway leads to the biosynthesis of L-rhamnose (Fig. 1A) (Giraud & Naismith, 2000). Notably, RML-3 is the only enzyme in the L-rhamnose pathway that is found in nematodes but appears to be absent in other eukaryotes. To learn more about the evolutionary origins of RML-3, we performed a BLASTP search to obtain all available homologous sequences from the RefSeq database using the *C. elegans* RML-3 protein sequence (accession NP_509046.1). Outside nematodes, the next most closely-related sequences ordered by the BLASTP bit-score (S) probability (*e*-value) were identified in bacterial and archaeal genomes (Supplemental table S1). In the highly conserved bacterial homologs, RML-3 is a dTDP-4-dehydrorhamnose 3,5-epimerase that catalyzes the third step in L-rhamnose biosynthesis from glucose-1-phosphate (Fig. 1A) (Mistou *et al*, 2016). To increase our sampling of nematodes, we also queried the non-redundant (NR), whole-genome sequencing (WGS), and transcriptome (TSA) databases. These searches revealed RML-3 homologs in a wide range of species in the *Chromadoria* clade of nematodes, which includes both free-living (e.g. *C. elegans*) as well as parasitic (e.g. *Enterobius vermicularis*) nematodes (Fig. S1, Supplemental table S1) (Smythe *et al*, 2019). In contrast, we found no RML-3 homologs in well-sequenced members of the *Dorylaimia* clade of nematodes (e.g. *Trichinella spiralis*) (Fig. S1). Based on this, we infer that nematode RML-3 was horizontally acquired from bacteria after the divergence of the *Chromadoria* and *Dorylaimia* clades, which is estimated to have occurred >400 million years ago (Smythe *et al*, 2019; Rota-Stabelli *et al*, 2013).

**Fig. 1.**
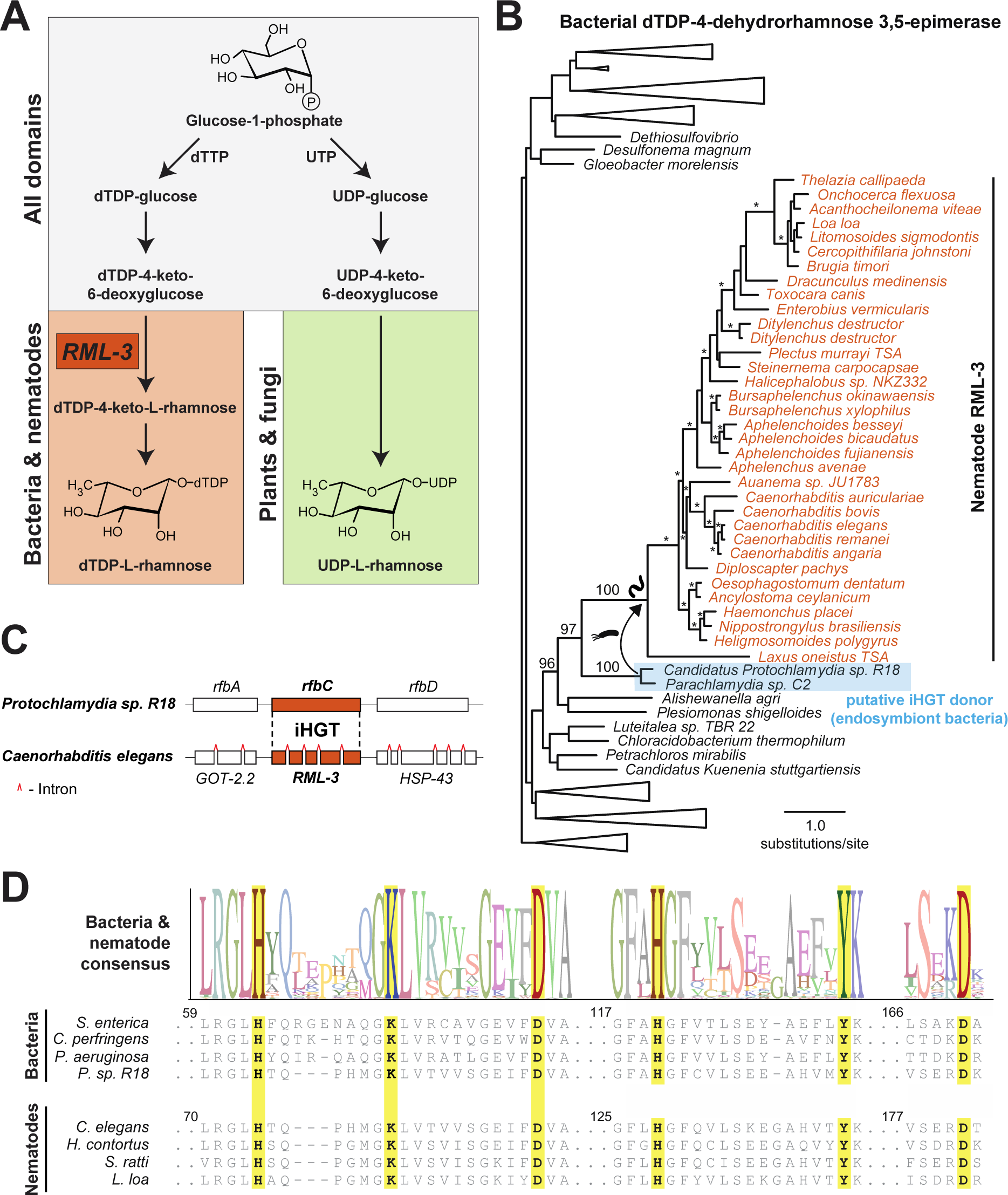
Nematodes acquired an enzyme required for L-rhamnose sugar biosynthesis by iHGT. **(A)** L-rhamnose biosynthetic pathway with reaction intermediates and differences in the conservation of enzymes across domains of life. The presence of homologs of the *C. elegans* RML-3 enzyme and production of dTDP-L-rhamnose is unique to bacteria and nematodes (orange). **(B)** Maximum-likelihood phylogenetic tree of RML-3 proteins and bacterial homologs. Relevant bootstrap branch support values greater than 90 are indicated by asterisks or numbers. Scale bar shows the estimated divergence in amino acid changes per residue. A complete phylogenetic tree with support values is found in Supplemental File 2. **(C)** Schematic genomic locus architecture of RML-3 homologs in *C. elegans* (RML-3) and *C. perfringens* (*rfbC*). Red carets within *C. elegans* genes represent introns. **(D)** Consensus sequence and positions of the predicted catalytic residues (yellow) for RML-3 homologs shown in (**B**). Four bacterial sequences are shown, including from three representative bacteria (*Salmonella enterica*, *Clostridium perfringens*, and *Pseudomonas aeruginosa*) and an example from the clade of putative iHGT donor bacteria, *Candidatus Protochlamydia sp. R18,* as well as four representative nematode sequences. Amino acid numbering corresponds to the *S. enterica* and *C. elegans* proteins respectively.

To confirm the bacterial origin of nematode RML-3, we reconstructed maximum likelihood (ML) phylogenetic tree using available RML-3 protein sequences from prokaryotes and metazoans (Fig. 1B, Table S1, Supplemental files). Our phylogenetic analyses reveal that nematode RML-3 forms a single monophyletic clade with high branch support, nested within a larger group of bacterial sequences. The monophyly of the nematode clade is maintained irrespective of the choice of ML phylogenetic reconstruction software or substitution model (Fig. S2, Supplemental files). The most closely-related bacterial species in all analyses are members of the *Chlamydiales* order of bacteria, which are obligate intracellular bacteria (Bayramova *et al*, 2018; Elwell *et al*, 2016) (Fig. 1B, Fig. S2). These data suggest that intracellular *Chlamydiales* bacteria were likely the “donor” species for RML-3, although caution should be taken given our inference that this gene transfer event occurred >400 million years ago (Rota-Stabelli *et al*, 2013) and there has likely been subsequent evolution that has taken place in bacterial and nematode genomes that may complicate inferences of an exact “donor”.

Over the period of *rml-3* gene domestication, nematodes acquired four introns within the horizontally transferred *rml-3* coding region (Fig. 1C). Nematode RML-3 homologs also show similar patterns of amino acid conservation as seen in bacterial enzymes, including retention of critical catalytic residues (Fig. 1D) (Giraud & Naismith, 2000), which is consistent with the described epimerase catalytic activity (Feng *et al*, 2016) of *C. elegans* RML-3. Importantly, RML-3 lacks similarity to the enzyme that catalyzes the epimerase reaction in plants and fungi, which is part of a bi-functional or tri-functional enzyme that catalyzes the last two steps in the biosynthetic pathway that results in UDP-L-rhamnose (Fig. 1A). Together, these data indicate that nematodes acquired an essential enzyme in L-rhamnose biosynthesis by iHGT from bacteria.

### *rml-3* expression pattern and regulation during development, dauer and stress

To gain a deeper understanding of the fascinating evolutionary acquisition of RML-3 from bacteria through iHGT, and its physiological relevance in modern nematodes, we characterized the expression, regulation and biological functions of RML-3 in the tractable model organism nematode *C. elegans*.

We first generated an integrated transgene-based transcriptional reporter, *rml-3*p::GFP, and CRISPR-mediated GFP knock-in (KI) alleles at the endogenous *rml-3* locus (Fig. 2, Fig. S3). The *rml-3* transcriptional reporter showed specific *rml-3* expression in the hypoderm (primarily in the seam cells, and less abundantly in the Hyp7) during larval development, starting from larval L1 through L4 and young adult (24 hrs post L4) stages (Figs. 2A-2D). Seam cells, also referred to as lateral hypodermal cells, share functions with the major hypodermis in secreting biomolecules to form the cuticle in *C. elegans* (Johnstone, 1994). In dauer animals, we observed *rml-3* expression predominantly in the hypodermal syncytium as seam cells was dramatically shrunk (Fig. 2E). The CRISPR GFP knock-in alleles showed identical expression patterns as the transcriptional reporter during late embryonic and larval development, albeit with lower overall fluorescence intensity of the knock-in reporters (Fig. S3). In mature adult animals (> 48 hrs post L4), we observed no detectable *rml-3* expression from either transcriptional or endogenous *rml-3* knock-in GFP reporters.

**Fig. 2.**
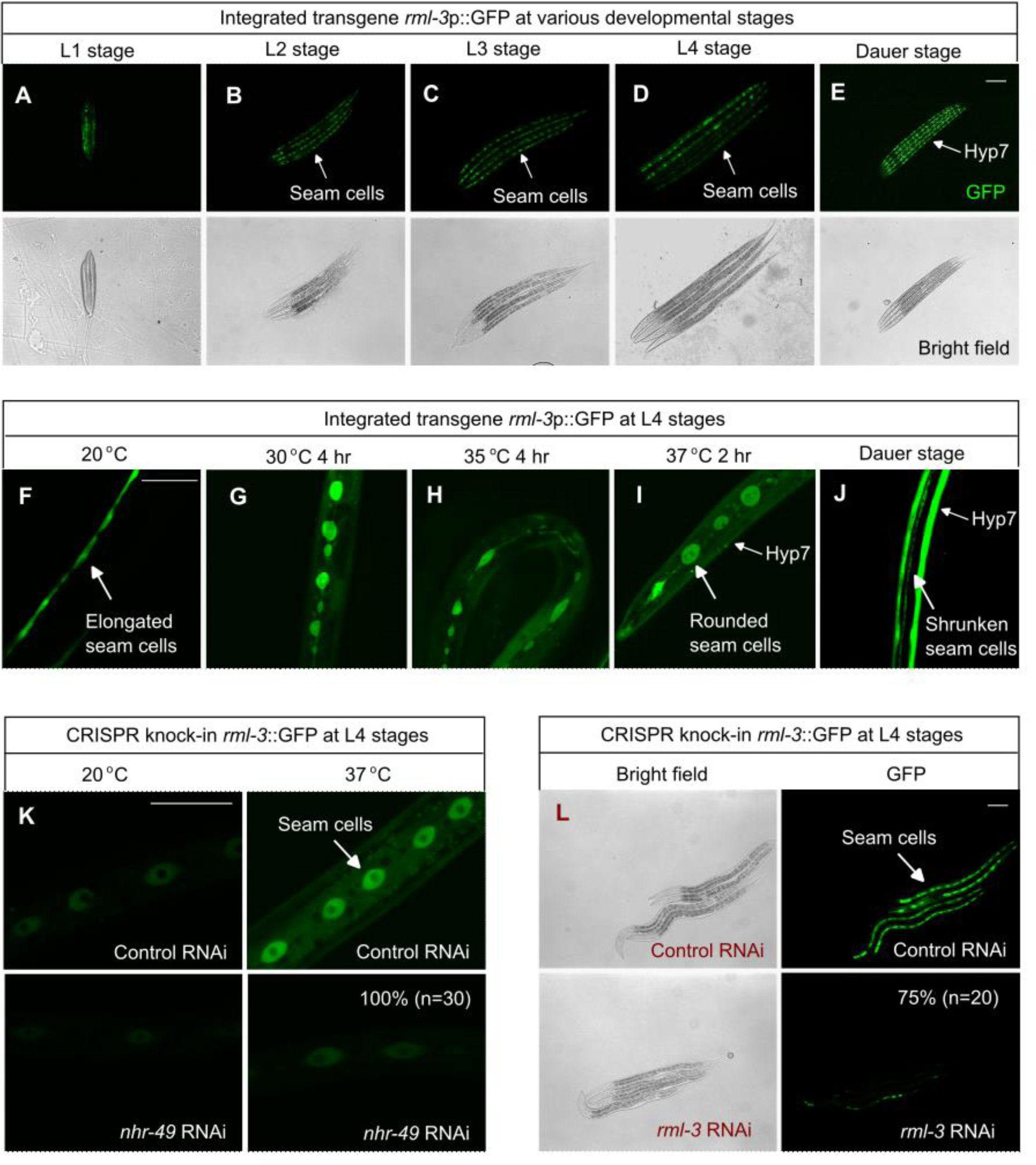
*C. elegans rml-3* expression pattern and regulation in development, dauer, and stress responses. **(A-E)**, Transcriptional GFP reporters show *rml-3* expression patterns during larval development and in dauer-stage animals. **(F-J)**, Transcriptional reporters show *rml-3* expression patterns after heat stress and heat-induced dauer formation. **(K)**, CRISPR-mediated RML-3::GFP KI reporters show *rml-3* expression in seam cells and induction by heat (37 °C for 2 hrs) in an NHR-49-dependent manner. **(L)** *rml-3* RNAi abolishes RML-3::GFP expression in KI reporters. Scale bars: 100 µm.

To examine the regulation of *rml-3* by environmental stress, we exposed *rml-3*p::GFP animals to various types of stress (hypoxia, anoxia, hypothermia, freezing, heat and osmotic shock). We found that only heat shock at various degrees and durations caused markedly increased expression of *rml-3* in seam cells, accompanied with rounded seam cell morphology (Figs. 2F-2J). We used RNAi of several candidate genes (*hsf-1, hif-1, nhr-49, skn-1, daf-16, sta-2*) encoding known stress-responding transcription factors to probe the mechanism of *rml-3* regulation by heat shock (Brunquell *et al*, 2016; Ma *et al*, 2015; Rodriguez *et al*, 2013; Jiang *et al*, 2018). We did not observe the apparent blocking effects of RNAi against *hsf-1*, which encodes the canonical heat shock factor in *C. elegans*. Among other candidate transcription factor-encoding genes tested, we observed that *nhr-49* (nuclear hormone receptor) RNAi strongly suppressed *rml-3* up-regulation by heat (Fig. 2K). Taken together, our expression analyses reveal specific baseline expression of *rml-3* in the developing seam cells, rapid up-regulation by heat stress, and prominent expression in the dauer hypoderm. As NHR-49 regulates pleiotropic stress-dependent genetic programs (Taubert *et al*, 2008; Vozdek *et al*, 2018; Naim *et al*, 2021; Dasgupta *et al*, 2020), we sought to probe specific biological functions of RML-3 using *rml-3* RNAi, which strongly decreased endogenous *rml-3*::GFP expression in the CRISPR KI strain (Fig. 2L).

### *rml-3* RNAi impairs cuticle integrity and epithelial barrier functions

The *C. elegans* cuticle is structured as dorsal and ventral regions covering the broad dorsal and ventral hypodermis and the narrow lateral regions overlying the seam cells (Johnstone, 1994). The cuticle is synthesized at the end of embryogenesis and in five cycles prior to each larval stage. The seam cell and hypodermal expression pattern of *rml-3* led us to examine the functional roles of RML-3 in cuticle formation and integrity. Toward this, we tested the effect of *rml-3* RNAi using a GFP reporter of COL-19, a secreted collagen protein critical for exoskeleton structure of the cuticle (Thein *et al*, 2003; Zhang *et al*, 2021, 2020). The COL-19::GFP reporter is abundant along the annular furrows and lateral alae in the cuticle under normal conditions (Fig. 3A). RNAi of several genes involved in collagen synthesis, including *dpy-7, dpy-11, sqt-1, rol-6, bli-1* have been previously reported to cause disruption, branching, and amorphous state changes in lateral alae and annuli leading to morphological defects labeled by COL-19::GFP (Thein *et al*, 2003). We found that both *rml-1* and *rml-3* RNAi caused amorphous lateral alae and disrupted annuli, frequently accompanied by severe hole-like cuticle defects, as revealed in COL-19::GFP animals (Figs. 3A-3B).

**Fig. 3.**
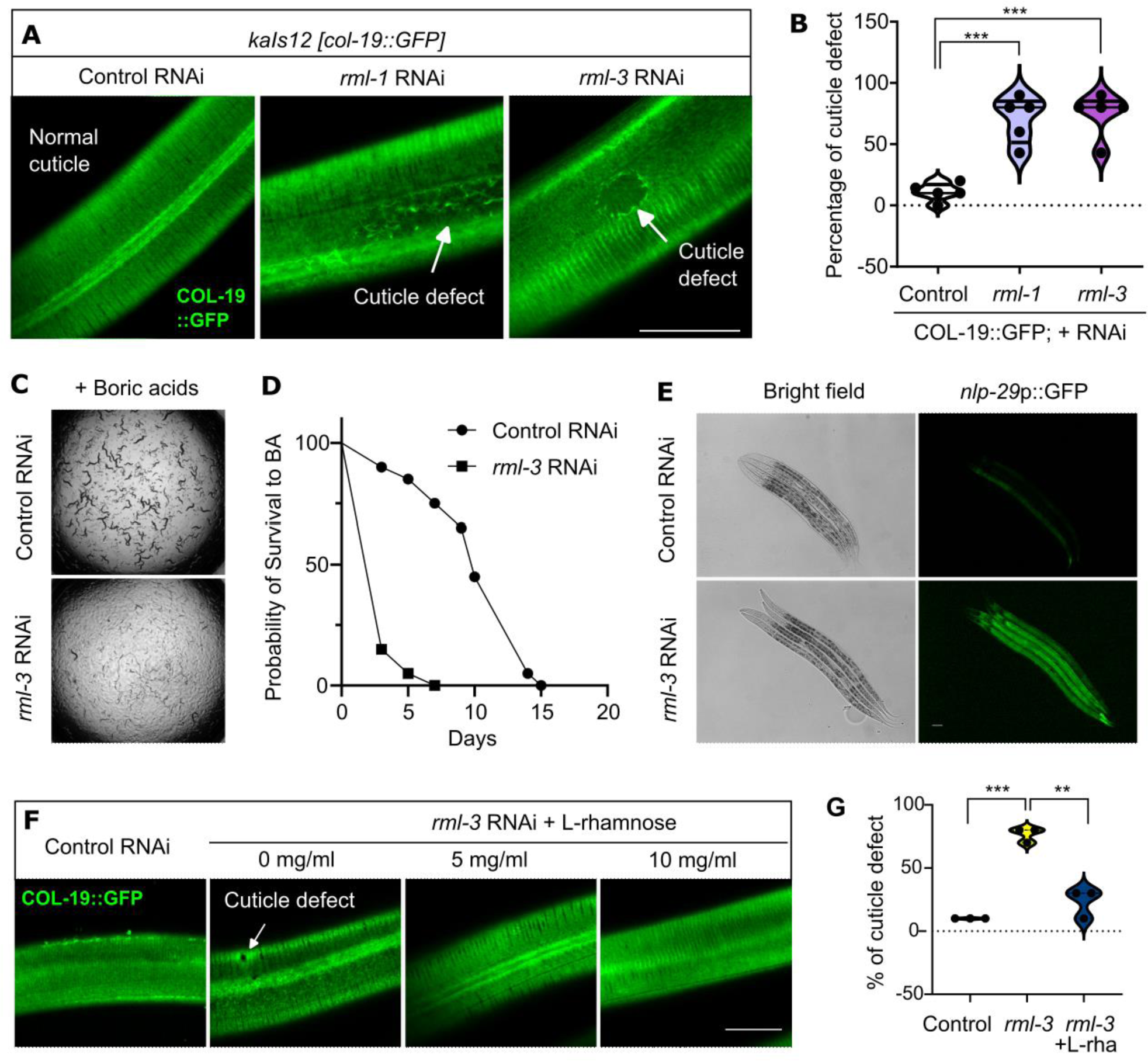
*rml-3* RNAi impairs cuticle integrity and epithelial barrier functions in *C. elegans*. **(A)**, Representative confocal images showing cuticle hole defects caused by RNAi against *rml-1* or *rml-3* visualized by the collagen COL-19::GFP reporter. **(B)**, Quantification of cuticle defects. *** indicates *P* < 0.001 (N = 5 independent experiments). **(C)**, Representative wild-field images showing boric acid (BA) sensitivity of animals treated with RNAi against *rml-3.* **(D)**, Survival curves showing accelerated death of animals (n > 50) treated with RNAi against *rml-3* under BA supplementation conditions. **(E)**, Representative epifluorescence images showing up-regulation of *nlp-29*p::GFP caused by RNAi against *rml-3*. **(F)**, Cuticle defects of animals with *rml-3* RNAi dose-dependently rescued by exogenous L-rhamnose. **(G)**, Quantification of rescue by L-rhamnose of the cuticle defect in animals with RNAi against *rml-3*. *** indicates *P* < 0.001 (N = 3 independent experiments). Scale bars: 50 µm.

The integrity of the cuticle is vital for barrier functions and protection against environmental toxins in *C. elegans*. Mutations of *rml-2*, also known as *bus-5*, led to impaired cuticle-barrier functions and sensitivity to boric acid (BA), a chemical present in nature and as an insecticide that imposes organismal toxicity (Gravato-Nobre *et al*, 2005; Xiong *et al*, 2017; Kiesche-Nesselrodt & Hooser, 1990). Hence, we next assessed the BA sensitivity of animals treated with *rml-3* RNAi. We found that a low-dose exposure to BA (6 mM in medium) caused robust developmental delay in animals with *rml-3* RNAi (Fig. 3C). Moreover, a high-dose exposure to BA (20 mM) caused a dramatic reduction in the survival of animals with *rml-3* RNAi compared to control (Fig. 3D), demonstrating an essential role of RML-3 in protection against the BA toxin.

To further characterize the effect of *rml-3* RNAi on cuticle integrity, we used an integrated transcriptional reporter *nlp-29*p::GFP. *nlp-29* encodes an antimicrobial peptide reported to increase in expression under conditions of hypodermal infection, wounds or disrupted cuticle in *C. elegans* (Sinner *et al*, 2021; Pujol *et al*, 2008; Chandler & Choe, 2022). We found that *rml-3* RNAi strongly increased *nlp-29*p::GFP expression both during late larval development (L4) and in young adults (24 hrs post L4). Consistently, RNAi against *rml-2* (also known as *bus-5*) also markedly increased *nlp-29*p::GFP expression (Fig. S4). Using a different collagen GFP reporter DPY-7::GFP, we observed additional cuticle defects in animals treated with *rml-3* RNAi characterized by DPY-7::GFP aggregation and abnormal alae (Fig. S5A). Given the striking similarity and homology of RML-3 to bacterial enzymes for L-rhamnose biosynthesis, we reasoned that *rml-3* RNAi caused cuticle defects resulting from reduced biosynthesis of L-rhamnose. If so, defects caused by *rml-3* RNAi could be rescued by supplementation with L-rhamnose, which can be converted to dTDP-L-rhanmose by ubiquitously expressed nucleotidyltransferases (Li *et al*, 2022). Indeed, we found that exogenously provided L-rhamnose partially rescued the hole-like cuticle defect of animals with *rml-3* RNAi (Figs. 3F, 3G). Additionally, *rml-3* RNAi caused reduced resistance to heat stress (Figs. S5B-5C), a phenotype also rescued by exogenous L-rhamnose (Fig. S5C). These results indicate that *rml-3* is crucial for the L-rhamnose biosynthetic pathway, which in turn is necessary to maintain proper cuticle organization and stress resilience against adverse environmental stresses.

### RML-3 is essential for larval development and haploinsufficient for BA resistance

We next sought to determine the null phenotype of *rml-3* since RNAi does not eliminate gene functions. Using CRISPR/Cas9-mediated deletions, we obtained two independent alleles to remove the entire coding region of *rml-3* (Fig. 4A). We found these homozygous deletion alleles caused identical larval L2-L3 developmental arrest phenotypes (Fig. 4B). From 20 morphologically normal F1 self-progeny of heterozygous deletion mutants, the F2 progeny segregated with the expected 1:3 Mendelian ratio for the L2-L3 larval arrest phenotype (Fig. 4C). From the self-progeny of heterozygous deletion mutants, we genotyped 20 randomly selected larval-arrested animals and found 100% of them carry homozygous deletions (Fig. 4D). Supplementation of L-rhamnose partially rescued the larval arrest phenotype of the *rml-3* homozygous mutants (Fig. 4B). Given the larval arrest phenotype of *rml-3* homozygotes, we further examined the BA sensitivity and *nlp-29*p::GFP phenotypes of *rml-3* heterozygous deletion mutants. Compared to wild type, *rml-3*/+ animals showed activation of *nlp-29*p::GFP and markedly reduced survival under BA treatment (Figs. 4E, 4F). Together, these results demonstrate that RML-3 is functionally important for progression through larval development through L-rhamnose production, while heterozygous *rml-3* causes a less severe developmental phenotype but marked defects in cuticle-barrier functions, leading to BA sensitivity and *nlp-29*p::GFP activation.

**Fig. 4.**
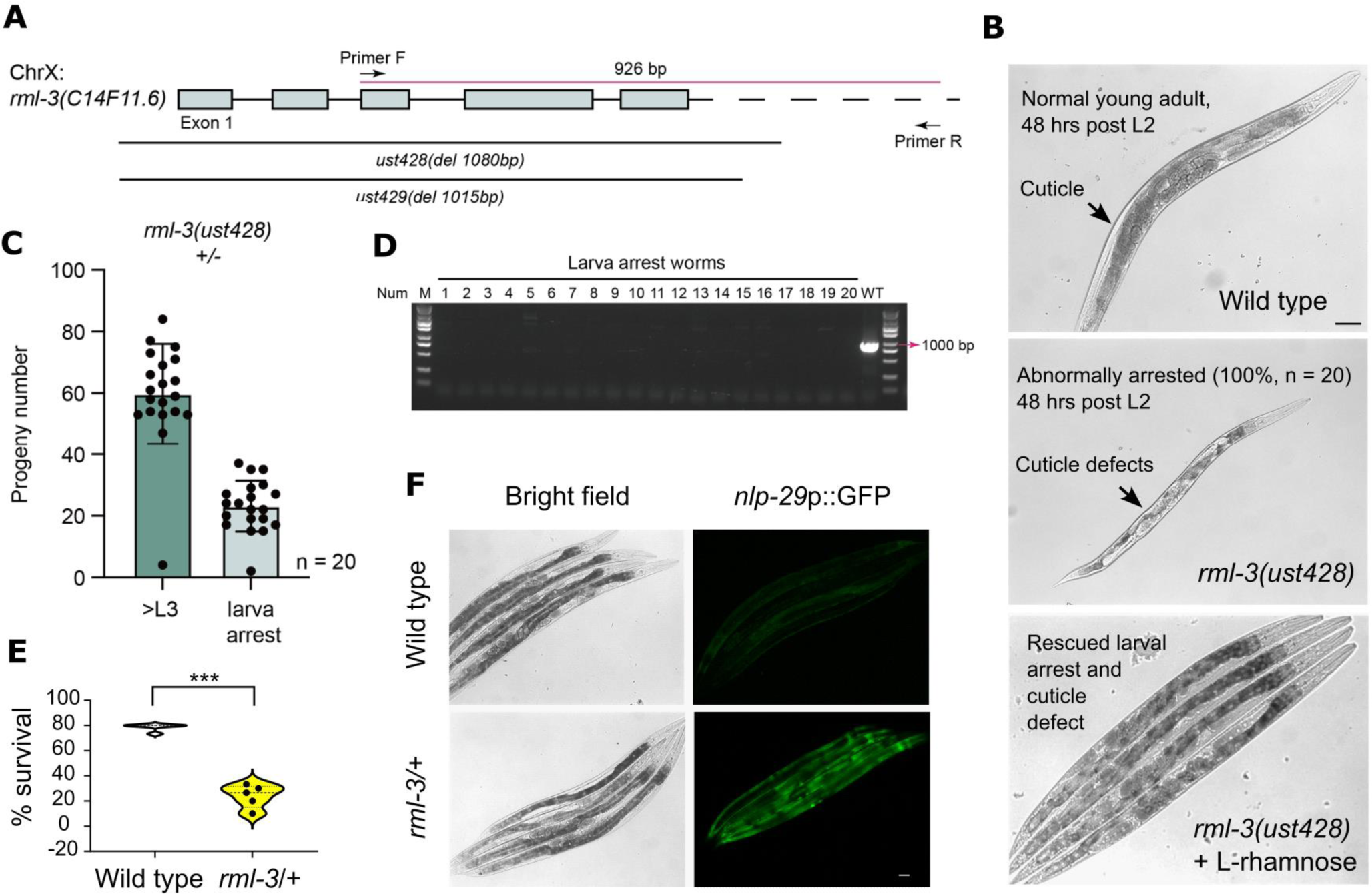
RML-3 ensures normal development with intact cuticle and promotes organismal resilience to environmental stress. **(A)**, Schematic showing CRISPR-mediated deletions of *rml-3.* **(B)**, Representative bright-field images showing developmental arrest of animals with homozygous deletions of *rml-3* and cuticle defects. Shown is also rescued development by exogenous supplementation of L-rhamnose (10 mg/ml in medium). **(C)**, Quantification of the number of progeny that can progress beyond L3 or become arrested from 20 *rml-3*/+ heterozygous parents. The ratio of the wild type vs arrested larvae corresponds to the Mendelian 3:1 ratio. **(D)**, PCR genotyping of 20 arrested larvae confirming *rml-3* homozygosity. **(E)**, *rml-3* heterozygous deletion mutants showing reduced survival under BA. **(F)**, Representative bright-field and epifluorescence images showing *rml-3* heterozygous mutants with up-regulated *nlp-29*p::GFP reporter expression. Scale bars: 50 µm.

## Discussion

Surviving unfavorable environmental conditions and stress have conferred nematodes with remarkable phenotypic plasticity that is reflected in their genome evolution. Unlike common modes of genome evolution based on germline DNA mutations and organismal trait selection, rare iHGT can facilitate metazoan evolution by incorporating existing functional genomic material from bacteria into the genome of an animal. Here, we characterized the evolutionary origin and functional consequences of a critical L-rhamnose biosynthesis enzyme that nematodes acquired from bacteria via iHGT. Leveraging the increased availability of genome sequence data and multiple ML-based phylogenetic reconstruction methods, we can unequivocally show that RML-3 was transferred to nematodes followed by gene domestication that involved the evolution of introns and conservation of critical amino acid residues. This gene is now represented across a wide range of nematodes, including diverse members of the well-sampled *Rhabditina, Tylenchina*, and *Spirurina* clades of nematodes.

How did a single iHGT event facilitate the biosynthesis of L-rhamnose in nematodes? The enzymes that catalyze the first two steps of the pathway shown in Fig. 1A, RML-1 (nucleotidyltransferase) and RML-2 (dehydratase) in nematodes, have homologs widely present in all domains of life, where they can contribute to other sugar biosynthetic pathways (Mistou *et al*, 2016; Martinez *et al*, 2012). Interestingly, the distribution of RML-4 homologs within eukaryotes is itself polyphyletic and is even found in eukaryotic viruses (Parakkottil Chothi *et al*, 2010), raising the possibility that it has been horizontally transferred among eukaryotes, possibly in a virus-mediated manner (Fig. S6). Regardless of the mechanism of RML-4 origin in nematodes, it is clear that RML-1, RML-2, and RML-4 in nematodes are closely related to eukaryotic enzymes that perform similar functions. The acquisition of RML-3 from bacteria completes the biosynthetic pathway, enabling nematodes to generate L-rhamnose uniquely among known metazoans.

The identity of the bacterial “donor” species for RML-3, and the general mechanisms of iHGT, are poorly understood, owing to the high frequency of gene transfer within bacteria and the rarity of iHGT. Our phylogenetic analyses suggest that the bacterial “donors” of nematode-specific RML-3 belong to members of the *Chlamydiales* order of bacteria, which are obligate intracellular bacteria and stable symbionts of diverse organisms (Bayramova *et al*, 2018; Elwell *et al*, 2016). These data indicate that endosymbiosis may have facilitated the iHGT of RML-3 into nematodes, consistent with several other instances of iHGT from endoymbiont bacteria into their host species (Dunning Hotopp *et al*, 2007; Sloan *et al*, 2014; Kondo *et al*, 2002). Although RML-3 homologs in bacteria, including in *Chlamydiales* (Fig. 1C), often resides in a chromosomal operon with other rhamnose biosynthetic enzymes, it is also possible that factors that facilitate bacterial HGT, including plasmids and other mobile genetic elements, may have contributed to iHGT. In addition, recent data indicate that virus-like or transposon-mediated gene transfer may occur in nematodes, suggesting that these genetic elements could be an additional contributing factor to this evolutionarily important mechanism of functional innovation (Moore *et al*, 2021; Dennis *et al*, 2012; Widen *et al*, 2023).

How do RML-3 and the acquired capacity to generate L-rhamnose contribute to fitness benefit in nematodes? Our expression and functional analyses of *rml-3* in *C. elegans* demonstrate its critical roles in conferring cuticle integrity and organismal resilience against environmental stresses. As a naturally occurring sugar widely distributed among bacteria and plants, L-rhamnose can act as a versatile building block in composite biomolecules, including rhamnolipids, extracellular polysaccharides, and glycosylated flagella (Schirm *et al*, 2004; Wild *et al*, 1997; Gao *et al*, 2001). These L-rhamnose conjugated biomolecules with altered physical chemical properties can protect organisms against environmental stresses, such as those from pathogen infection or toxin infiltration. Given the striking cuticle defect and organismal sensitivity to BA in RML-3 deficient *C. elegans*, we propose that L-rhamnose is conjugated to biomolecule target(s) that are critical for cuticle integrity and protection against environmental stresses. While the complete loss of RML-3 causes larval arrest with severe cuticle defects, the partial loss of RML-3 allows larval developmental progression, yet with impaired cuticle leading to BA sensitivity and activation of a cuticle stress-responding pathway for *nlp-29* up-regulation. The L-rhamnose conjugated biomolecules critical for cuticle integrity remain to be identified and investigated. Leading candidates include extracellular biomolecules such as collagen, cuticulin, and peptidoglycan, which have been previously implicated in conferring cuticle integrity and resilience to stress (Johnstone, 1994; Sandhu *et al*, 2021; Ewald *et al*, 2015; Li Zheng *et al*, 2020).

In summary, our study reveals the evolutionary origin and organismal role of an L-rhamnose biosynthetic enzyme RML-3 that was acquired from bacteria via iHGT and domesticated in nematodes. With the role in conferring resilience to nematodes against environmental stresses, *rml-3* adds to the growing repertoire of HGTs from microbial species in the *C. elegans* and nematode genomes, including our recent discovery of cyanide detoxification enzymes in *C. elegans* evolutionarily acquired from ancient green algae (Wang *et al*, 2022). Our work supports the emerging notion that the domestication of bacterial genes by iHGT is more widespread in shaping metazoan evolution than currently appreciated and can provide diverse functional capabilities to specific metazoan lineages because of constant and ubiquitous exposure to bacteria. Although rare, iHGT events have had a profound impact on metazoan evolution, as highlighted by the essential roles of RML-3 in modern nematodes to enable exoskeleton integrity, organismal resilience and survival under adverse environmental conditions.

## Materials and Methods

### Phylogenetic analysis

*C. elegans* RML-3 (accession NP_509046.1) was used to query the NCBI RefSeq protein database using BLASTp (Altschul *et al*, 1990) with an e-value cutoff of 1e-5 to obtain the top 5000 eukaryotic and prokaryotic RML-3 homologs. Additional nematode sequences were obtained by querying the NCBI non-redundant (NR) database using BLASTp for all nematodes or the transcriptome shotgun assembly (TSA) database using tBLASTn for nematodes outside the *Rhabditina*, *Tylenchina*, and *Spirulina* clades that were already well covered from protein searches. Resulting sequences are listed in Supplemental File 1. Resulting sequences were aligned using Clustal Omega (Sievers & Higgins, 2014) using two iterations of refinement. Incomplete sequences and poorly aligning proteins were removed from subsequent analyses. To eliminate closely related sequences and reduce the total sequence number, sequences with >95% identity were reduced to a single unique sequence using CD-HIT with a 0.95 sequence identity cutoff (Fu *et al*, 2012). Following realignment with Clustal Omega, alignment positions in which >90% of all sequences had a gap were removed.

IQ-TREE (Nguyen *et al*, 2015) phylogenies were generated using the “-bb 1000-alrt 1000” commands for the generation of 1000 ultrafast bootstrap and SH-aLRT support values. The best fitting substitution model was determined by ModelFinder (Kalyaanamoorthy *et al*, 2017) using the “-m AUTO” command or the substitution model was specified as shown in Fig. S2 using the “-m” command. FastTree (Price *et al*, 2009) (version 2.1.11) phylogenies were generated using 20 rate categories. All Newick-formatted phylogenetic trees generated from IQ-TREE and FastTree, with support values, can be found in Supplementary File 2. Protein phylogenies were visualized as unrooted trees using FigTree (http://tree.bio.ed.ac.uk/software/figtree/).

RML-4 database searches were performed using residues 335-631 of *C. elegans* RML-4 (accession NP_001040727.1). Homologs were obtained by querying the RefSeq (bacteria and eukaryotes) or NR (viruses) databases, using an e-value cutoff of 1e-5.

Alignments were generated and refined as above, using the same 95% sequence identity cutoff to remove near-redundant sequences. A maximum likelihood phylogenetic tree was generated using IQ-TREE and ModelFinder as above. A complete Newick-formatted phylogenetic tree with support values can be found in Supplementary File 2. Eukaryotic RML-1 homologs were identified using *C. elegans* RML-1 (accession NP_499842.1) and searching against the eukaryotic RefSeq database using BLASTP.

### *C. elegans* culture and transgene construction

All the *C. elegans* strains used in this study were maintained in accordance with the standard laboratory procedures unless otherwise stated (Brenner, 1974). The worms were grown on an OP50 *Escherichia coli* strain at 20 °C. Embryos were isolated through sodium hypochlorite treatment. RNAi exposure was performed according to the standard procedure (Kamath & Ahringer, 2003). Genotypes of strains used are: N2 Bristol strain (wild type), *rml-3*(*ust428*), *rml-3*(*ust429*), *rml-3*(*ust430*[*rml-3::gfp::3xflag*]), *bus-5*(*e3133*), *kaIs12* [*col-19*::*GFP*], *dmaIs145* [*rml-3*p::*GFP*; *unc-54*p::*mCherry*], *qxIs722* [*dpy-7*p::*dpy-7*::*SfGFP* (single copy)], *frIs7* [*nlp-29*p::*GFP* + *col-12*p::*dsRed*].

For constructing *rml-3* transcriptional reporter, GFP-3’UTR was PCR amplified using primers (5’-AGCTTGCATGCCTGCAGGTCG-3’ and 5’-AAGGGCCCGTACGGCCGACTA-3’) and plasmid pPD95.75 (Plasmid #1494-Addgene) as the template. Then, *rml-3* gene-specific primers (Forward 5’-taggtcaaaagtgctggtggc-3’; Reverse 5’-cgacctgcaggcatgcaagcttattcaatgcgagtcaggcaagaa-3’, with sequence immediately upstream of the protein-coding region so that 5’UTR is part of the transgene) were amplified by PCR using a genomic DNA template. Nested PCR was used to generate the gene promoter-GFP fusion product. The fusion product was injected at 20 ng/µl with unc-54::mCherry co-injection marker at 10 ng/µl. The integration of the extrachromosomal arrays was performed through ultraviolet irradiation and backcrossed for three to six times.

### CRISPR/Cas9-mediated GFP knock-in and gene deletion of *rml-3*

For CRISPR knock-in at the endogenous *rml-3* locus with 3xFLAG::GFP, a 3xFLAG::GFP region was PCR amplified with the primers 5’-ATGGACTACAAAGACCATGACGG-3’ and 5′-AGCTCCACCTCCACCTCCTTTG-3′ from the genomic DNA of 3xFLAG::GFP::RPOA-2. A 1.5 kb homologous left arm was PCR amplified with the primers 5′-GGGTAACGCCAGCACGTGTGCGTACTTCGGTCGTGAAATT-3′ and 5’-TCATGGTCTTTGTAGTCCATAGTAGGATGCGACATTATTCAAT-3’. A 1.5 kb homologous right arm was PCR amplified with the primers 5’-AAGGAGGTGGAGGTGGAGCTATGTCGCATCCTACTCCAG-3’ and 5’-CAGCGGATAACAATTTCACAGCGGTGCATCTTCACATTA-3’. The backbone was PCR amplified from the plasmid pCFJ151 with the primers 5’-CACACGTGCTGGCGTTACC-3’ and 5’-TGTGAAATTGTTATCCGCTGG-3’. All these fragments were joined together by a Gibson assembly to form the 3xFLAG::gfp::*rml-3* plasmid with the ClonExpress MultiS One Step Cloning Kit (Vazyme Biotech). This plasmid was coinjected into N2 animals with three sgRNA expression vectors: *rml-3*_sgRNA#1, *rml-3*_sgRNA#2, *rml-3*_sgRNA#3, 5 ng/μl pCFJ90, and 50 ng/μl Cas9 II expressing plasmid.

To construct sgRNA expression plasmids for CRISPR-mediated *rml-3* deletion, the 20 bp *unc-119* sgRNA guide sequence in the pU6::unc-119 sgRNA(F + E) vector was replaced with different sgRNA guide sequences. Addgene plasmid #47549 was used to express the Cas9 II protein. A plasmid mixture containing 30 ng/μl of each of the three sgRNA expression vectors, 50 ng/μl Cas9 II expressing plasmid, and 5 ng/μl pCFJ90 was co-injected into animals. The deletion mutants were screened by PCR amplification and confirmed by sequencing.

### Epifluorescence microscopy

The epifluorescence compound microscope (Leica DM5000 B Automated Upright Microscope System with 10× and 20× objective lens) and confocal microscope (Leica TCS SPE with a 63× objective lens) were used to capture fluorescence images. Briefly, synchronous worm population was used for imaging at different developmental stages of indicated genotypes or treated with different RNAi that were randomly picked and anesthetized with 10 mM sodium azide in M9 solution (Sigma-Aldrich) on 2% agar pads on slides. The control and treatment groups were imaged under identical settings and conditions. For the thermal stress assay, the worms were subjected to heat stress at various temperatures (30, 35 and 37 °C) with respect to control worms at 20 °C prior imaging. The percentage of cuticle damage was scored in accordance to disruption in the worm cuticles with respect to the control worms (N > 50). The rescue of cuticle damage was tested post supplementation of worms with 5-10 mg/ml of L-rhamnose (Fisher, 10030-85-0) and the vehicle control (M9 buffer).

### Boric acid sensitivity and rescue assay

BA (Fisher, S25202A) dissolved at 6 mM in the Nematode Growth Medium (NGM) was used for developmental delay studies and 20 mM for survival analysis. For developmental effect studies, control and *rml-3* RNAi-treated L1 were exposed to BA and then imaged on the 6^th^ day of adulthood post L4 stages. For survival analysis, the strains were exposed to BA from the L1 larval stage and the percentage survival was scored either through the last worm surviving with RNAi treatment or post 3 days of exposure in mutants with their respective controls. 10 mg/ml L-rhamnose dissolved in the M9 buffer spread on NGM was used for the rescue assay.

### Statistics and reproducibility

All the data in the current manuscript were analyzed using GraphPad Prism 9.2.0 Software (Graphpad, San Diego, CA) and presented as means ± S.D. unless otherwise specified, with significance *P* values calculated by unpaired two-sided *t*-tests (comparisons between two groups), one-way or two-way ANOVA (comparisons across more than two groups) and adjusted with Bonferroni’s corrections. The lifespan assay was quantified using Kaplan–Meier lifespan analysis, and p values were calculated using the log-rank test.

## Acknowledgments

Some strains were provided by the *Caenorhabditis* Genetics Center (CGC), which is funded by the NIH Office of Research Infrastructure Programs (P40 OD010440). The work was supported by NIH grants (R35GM139618 to D.K.M., R35GM133633 to M.D.D.), UCSF PBBR New Frontier Research (D.K.M.), UCSF BARI Investigator Award

(D.K.M.), Pew Scholars Program (D.K.M., M.D.D.), and Burroughs Welcome Investigators in the Pathogenesis of Infectious Disease Program (M.D.D.).

## Author contributions

T.P., B.W. and D.K.M. designed, performed and analyzed the *C. elegans* experiments, contributed to project conceptualization and wrote the manuscript. T.X., X.H., and S.G. contributed to the generation of CRISPR alleles and experimental characterization. C.K. and M.D.D. performed phylogenetic analysis for the discovery of *rml-3* as originated by iHGT and contributed to project conceptualization and wrote the manuscript. D.K.M. and

M.D.D. supervised the project.

## Competing interests

The authors declare no competing interests.

## Materials & Correspondences

Correspondence and material requests should be addressed to Dengke K. Ma, Ph. D. and Matthew Daugherty, Ph. D.

---

**Fig. S1.**
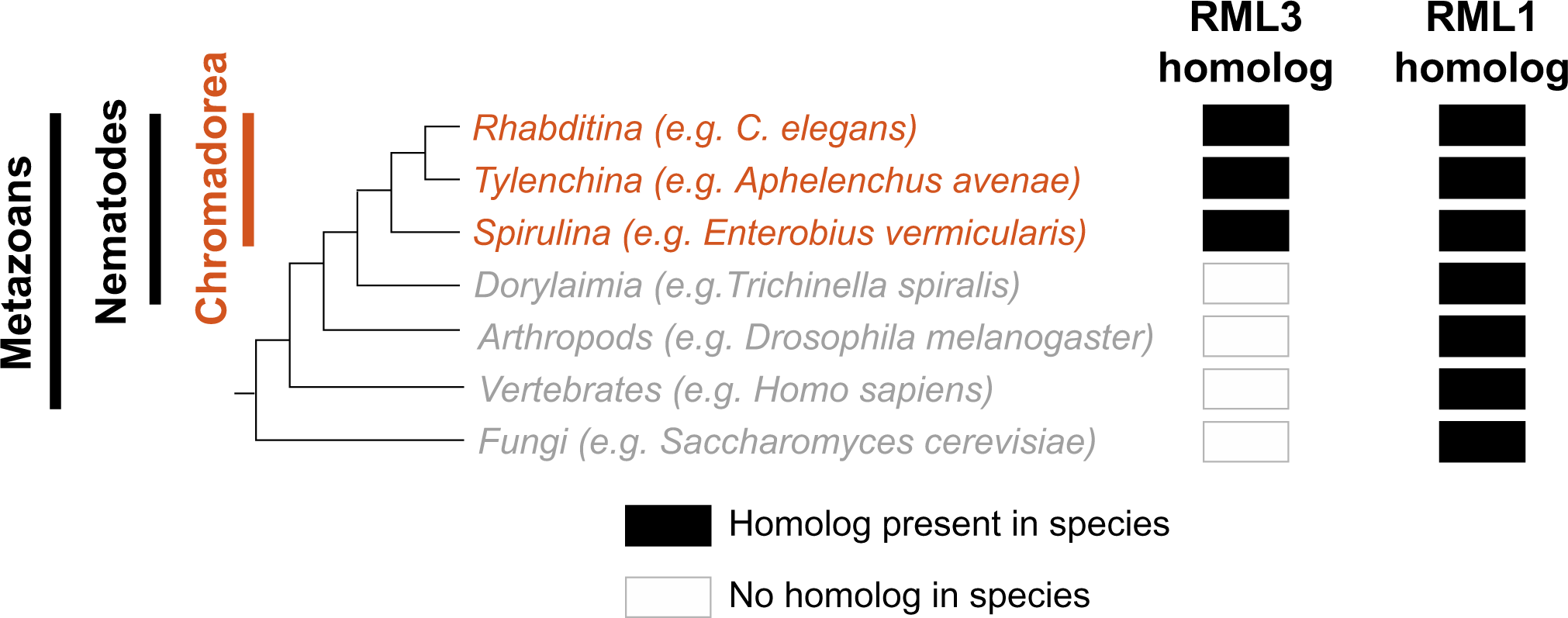
Phylogenomic distribution of RML-1 and RML-3 enzymes in eukaryotes. The phylogenetic tree of major clades of species queried for homologs of RML-1 and RML-3 homologs. The presence of homologous protein sequences was determined by BLASTp searches respectively using *C. elegans* RML-3 (accession NP_509046.1) or RML-1 (accession NP_499842.1). Filled black boxes indicate the presence of a homologous protein in the non-redundant (NR) database. In all cases, the protein identity was >49% and the e-value was <1e-150. Unfilled boxes indicate that there is no homologous protein with an e-value <0.01 when searching for the indicated homolog.

**Fig. S2.**
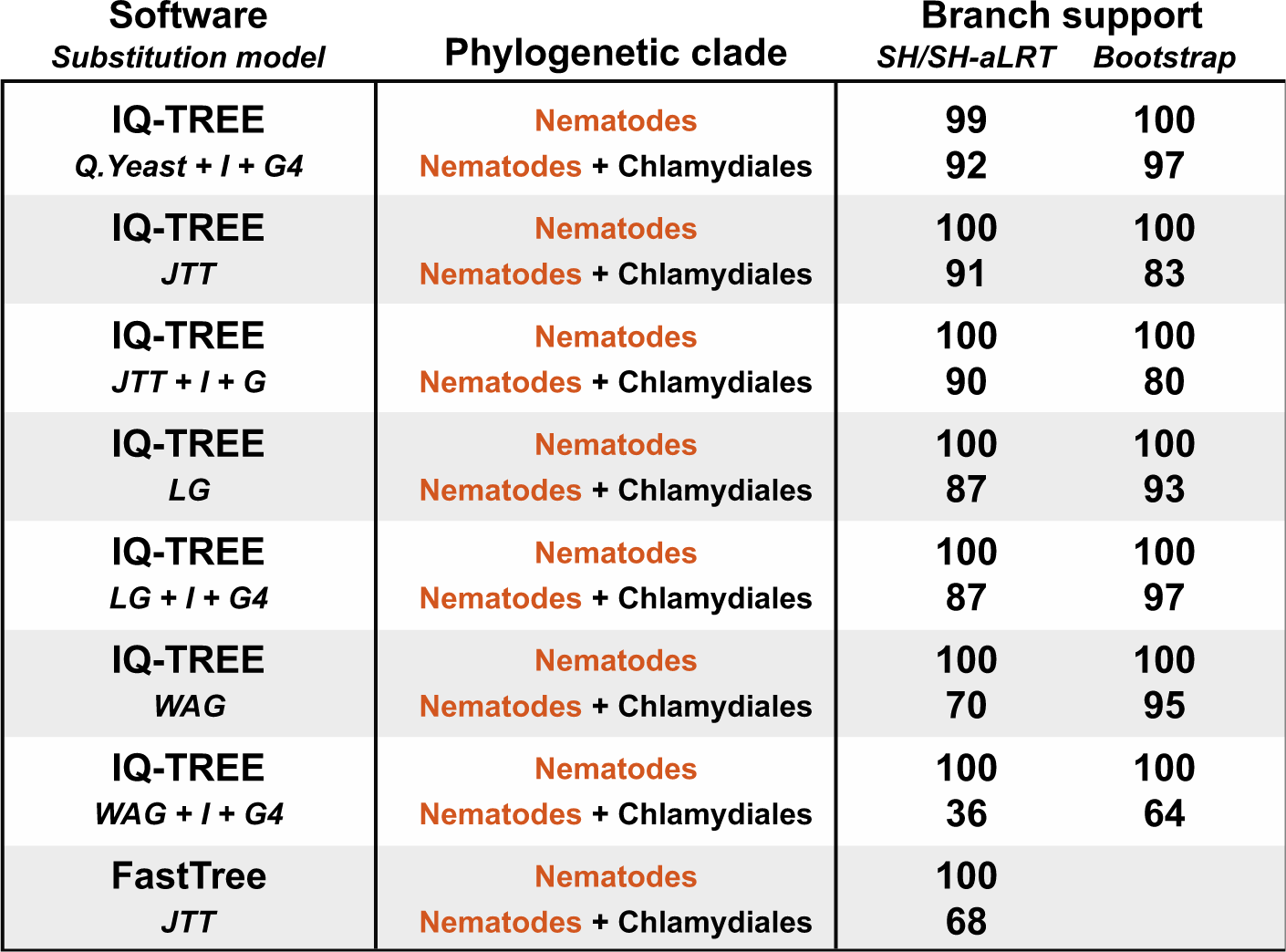
Phylogenetic inferences are robust to different software and substitution models. Phylogenetic trees of RML-3 protein homologs were generated with different indicated parameters (see Methods). An asterisk indicates the best fitting substitution model. For each analysis, branch support values for monophyletic clades are shown. Complete phylogenetic trees support values for all analyses are found in Supplemental File 2.

**Fig. S3.**
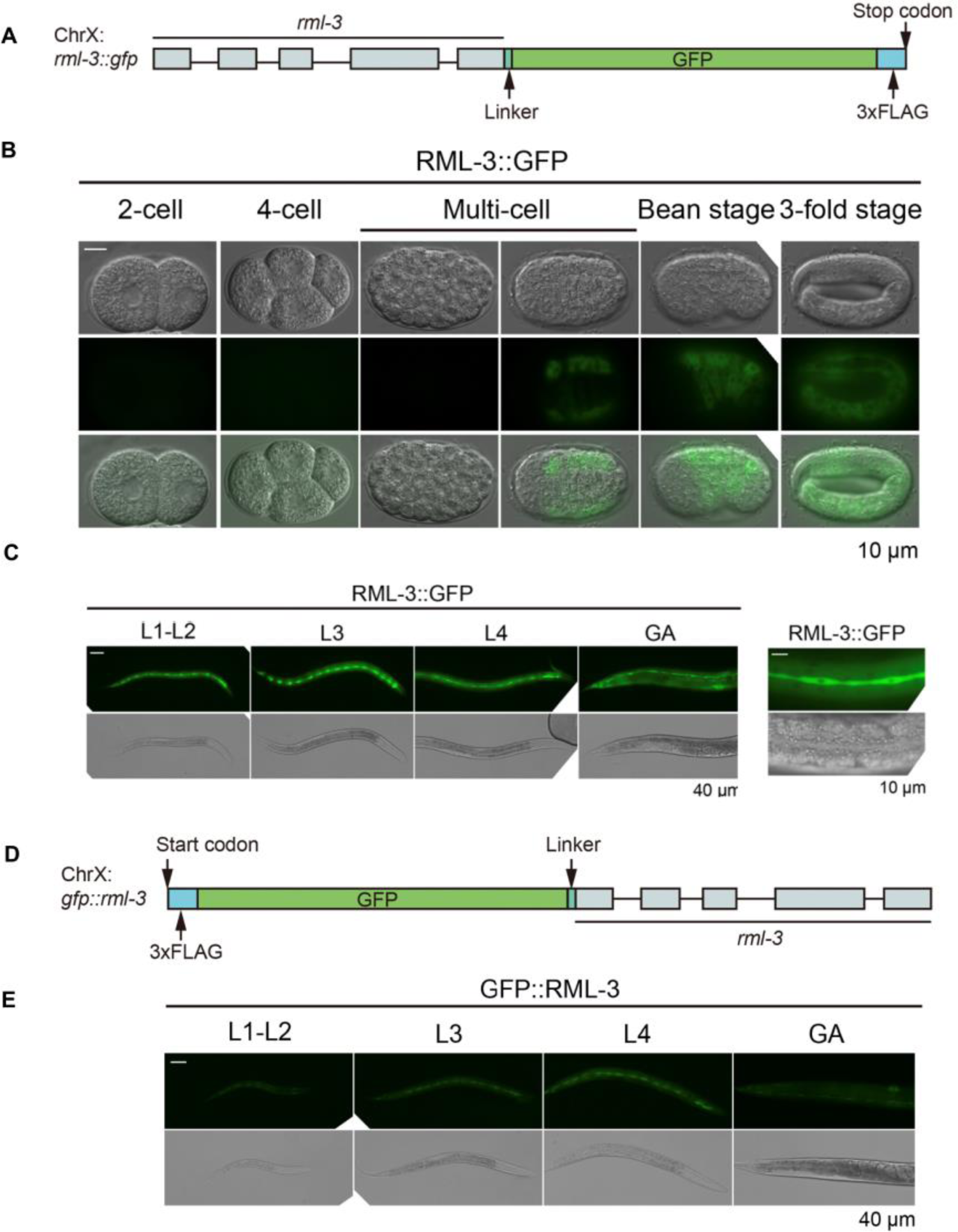
RML-3 expression pattern by CRISPR GFP knock-in at either end of RML-3. (A), Schematic for the design of CRISPR GFP knock-in at the C-terminal end of RML-3. (B), Representative brightfield and epifluorescence images showing expression of *rml-3* in developing seam cells and hypoderm during late embryonic development. (C) Representative bright-field and epifluorescence images showing expression of *rml-3* in developing seam cells and hypoderm during larval development. (D), Schematic for the design of CRISPR GFP knock-in at the N-terminal end of RML-3. (E) Representative brightfield and epifluorescence images showing expression of *rml-3* in developing seam cells during larval development.

**Fig. S4.**
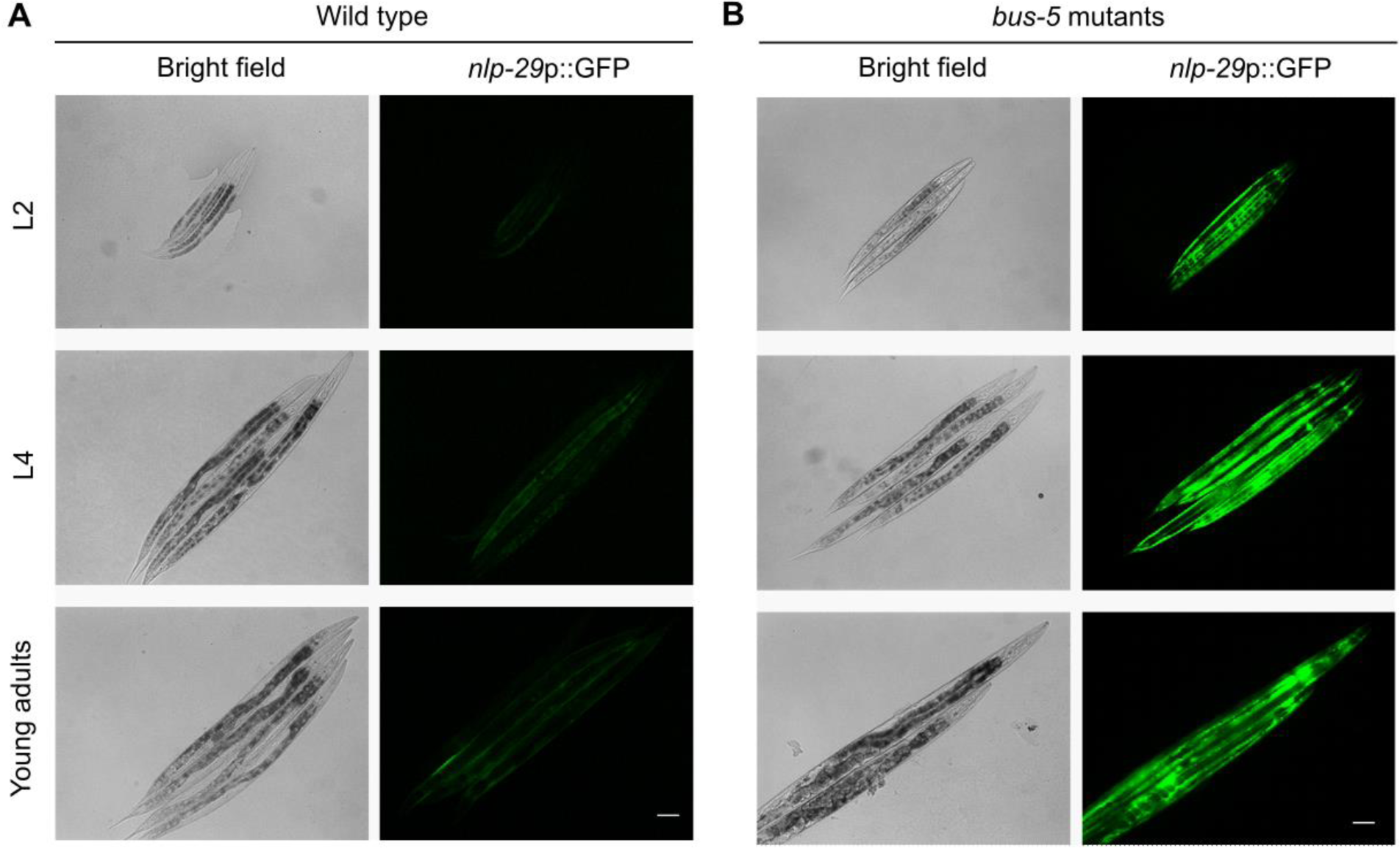
Mutation of *rml-2* (aka *bus-5*) causes constitutive *nlp-29*p::GFP up-regulation. (A), Representative epifluorescence images showing baseline *nlp-29*p::GFP abundance in wild type. (B), Representative epifluorescence images showing *nlp-29*p::GFP up-regulation with RNAi against *rml-2*. Scale bars: 50 µm.

**Fig. S5.**
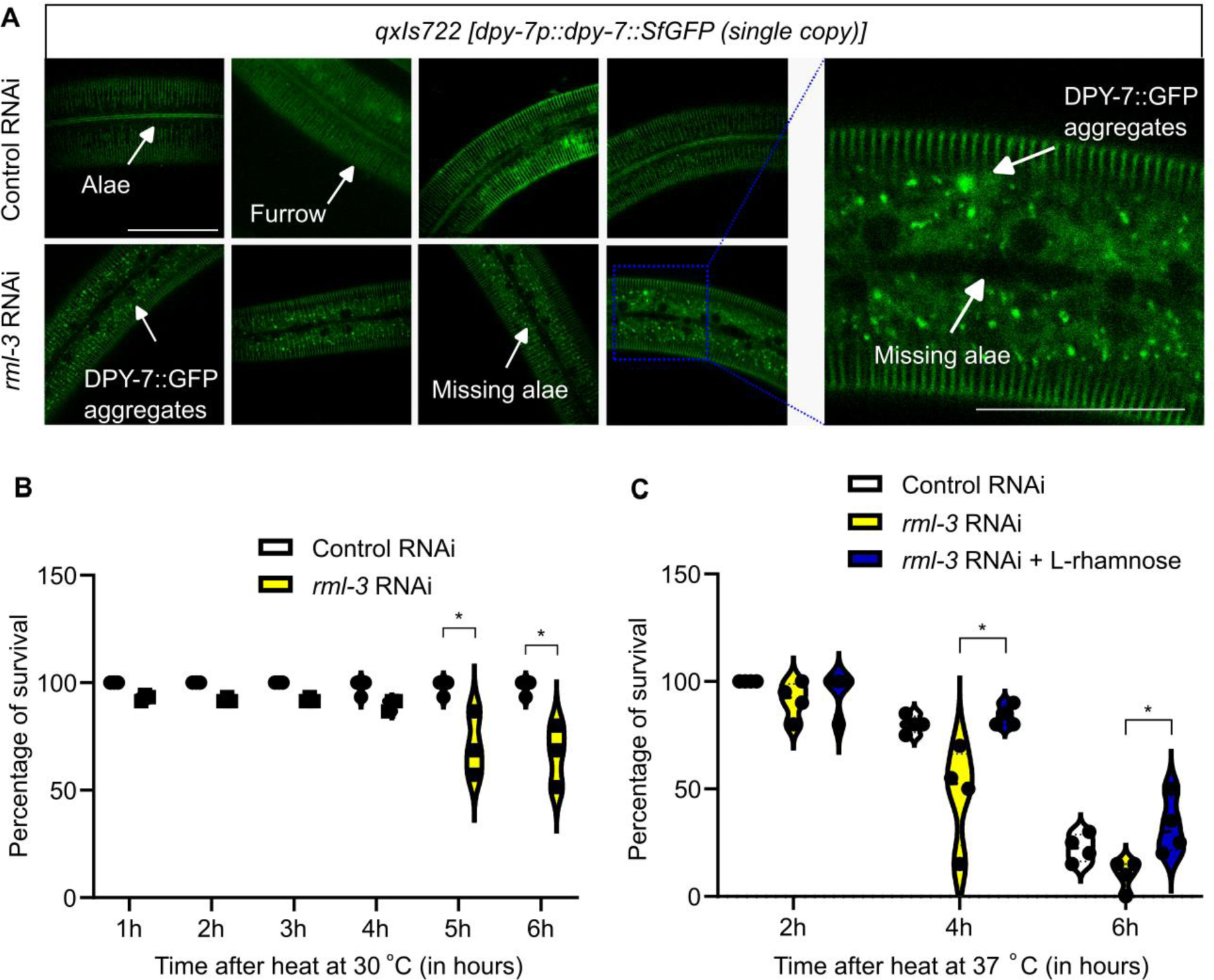
Additional phenotypes caused by *rml-3* RNAi. (A) Shown are representative confocal fluorescence images indicating cuticle defects, including collagen reporter DPY-7::GFP aggregates and missing DPY-7::GFP+ alae in animals with *rml-3* RNAi. Scale bars: 50 µm. **(B)** Survival analysis of control and *rml-3* RNAi-treated animals subject to 30 °C heat stress. * indicates *P* < 0.05 (N = 3 independent experiments, n > 50 animals per group). **(C)** Survival analysis of control and *rml-3* RNAi treated animals subject to 37 °C heat stress. * indicates *P* < 0.05 (N = 3 independent experiments, n > 50 animals per group).

**Fig. S6.**
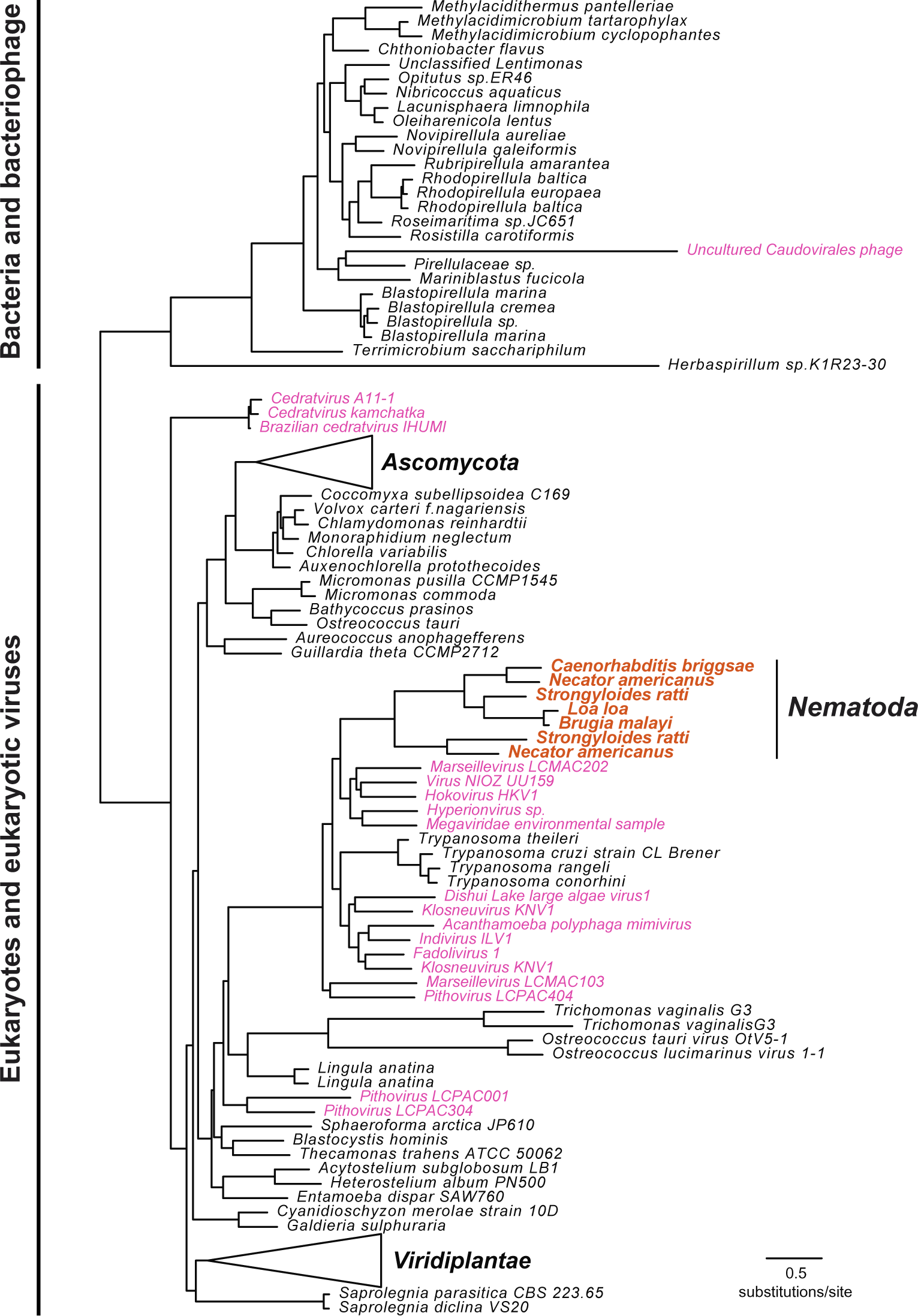
The phylogenetic tree of homologs of *C. elegans* RML-4. Maximum-likelihood phylogenetic tree of RML-4 homologs found in eukaryotes, bacteria, and viruses. Nematodes are labeled in orange and bacteriophage and eukaryotic viruses are labeled in pink. Major clades of proteins from plants (*Viridiplantae*) and fungi (*Ascomycota*) are collapse for clarity. Scale bar shows the estimated divergence in amino acid changes per residue. A complete phylogenetic tree with support values is found in Supplemental File 2.

## Notes

### Competing Interest Statement

The authors have declared no competing interest.

