## Supplemental file 2 for "Acquired stress resilience through bacteria-to-nematode horizontal gene transfer"

**Supplementary File 2: IQTREE and FastTree output phylogenetic trees.**

For IQTREE, the unrooted maximum likelihood tree is shown. For each node, results from 1000 iteration SH-aLRT and 1000 iteration ultrafast bootstraps are shown as support values respectively. For FastTree, the unrooted maximum likelihood tree is shown, with SH support values shown for each node.

**Nematode RML-3, with bacterial and eukaryotic homologs**

**Shown in Figures 1B and Figure S2**

ML tree method: IQ-TREE

Sequences: 2787

Substitution model: Q.yeast+I+G4 (best fit model)

(WP_063501760.1-Escherichia-coli:0.0523445588,(WP_044173167.1-Phytobacter-massiliensis:0.0544515715,(((((((WP_203451401.1-Lelliottia-sp.-RWM.1:0.0910168800,WP_185654835.1-Citrobacter-braakii:0.1690791729)90.8/100:0.1168326875,WP_183170804.1-Atlantibacter-sp.-RC6:0.3407300252)96.1/100:0.1530446137,(WP_160868038.1-Pantoea-sp.-Taur:0.5086026935,WP_008499438.1-Enterobacter-cloacae-complex:0.2721634168)87.7/97:0.0783903965)69/96:0.0195382880,((WP_090085547.1-Escherichia-coli:0.0868115874,WP_234083964.1-Enterobacter-quasiroggenkampii:0.1954541908)20.2/72:0.0348379820,WP_103283990.1-Citrobacter:0.2076658856)99.8/100:0.2475929171)36.2/99:0.0685006909,(((((((((((((((((((((((((((WP_249615874.1-Craterilacuibacter-sp.-RT1T:0.6745229249,(WP_163316497.1-Crenobacter-sedimenti:0.2155468047,WP_160797365.1-Craterilacuibacter-sinensis:0.1731389183)95.7/100:0.2559231906)35.4/48:0.0388442776,WP_014088374.1-Pseudogulbenkiania-sp.-NH8B:0.3299089325)92.1/49:0.0986335907,((WP_174873379.1-Vogesella-oryzae:0.1372931471,WP_137008594.1-Aquitalea-aquatilis:0.5425248380)56.3/91:0.1548975789,WP_147289921.1-Crenobacter-cavernae:0.1811870100)90.8/94:0.1212588360)0/30:0.0000025742,WP_008952446.1-Pseudogulbenkiania-ferrooxidans:0.3681145219)86.7/46:0.0564018644,((((((((((((((WP_004123988.1-Rhizobium-freirei:0.1383810781,WP_095433322.1-Rhizobium-sp.-11515TR:0.0512369911)82.3/92:0.0315107693,((WP_183712806.1-unclassified-Rhizobium:0.0439946095,WP_015341698.1-Rhizobium:0.0514817490)86.8/89:0.0179287079,WP_106999541.1-Rhizobium-sp.-JAB6:0.0655669286)0/88:0.0000029994)95/99:0.0273258544,((WP_183765619.1-unclassified-Rhizobium:0.0478044179,WP_120667780.1-unclassified-Rhizobium:0.0791047962)52.3/100:0.0378621217,WP_037195771.1-Rhizobium-lusitanum:0.0418659193)84.2/90:0.0218230656)0/72:0.0000025228,WP_184706966.1-Rhizobium-lusitanum:0.1335264371)50.7/85:0.0099295552,WP_092719658.1-Rhizobium-multihospitium:0.0448132876)51.3/87:0.0155943646,WP_174074493.1-Agrobacterium-rhizogenes:0.1308669762)85.5/100:0.0297746476,WP_168880117.1-Rhizobium-sp.-P28RR-XV:0.0576110226)81.6/99:0.0179230318,WP_112341687.1-Rhizobium-tropici:0.0465522351)86.1/99:0.0237995622,WP_199146672.1-Rhizobium-sp.-ASV8:0.2524046380)66.7/99:0.0078698581,WP_145321829.1-unclassified-Rhizobium:0.1330356815)99.6/100:0.2221874207,(WP_043579071.1-Chromobacterium:0.0670379294,WP_071113424.1-Chromobacterium-sphagni:0.1569426063)98.7/100:0.2086229206)95.1/100:0.1102769289,WP_020932001.1-Cycloclasticus-zancles:0.5207673219)68.9/78:0.0372337612,(((WP_189531325.1-Paludibacterium-paludis:0.7123583222,(WP_094838100.1-Bordetella-genomosp.-4:0.2508691360,WP_160351730.1-Bordetella-sp.-15P40C-2:0.3340198254)99.5/100:0.3436434478)86.1/85:0.0940728308,(((WP_213263532.1-Burkholderia-pyrrocinia:0.1800164859,WP_060279477.1-Burkholderia-cepacia-complex:0.1914705872)97.9/100:0.2125621570,(((WP_208101977.1-Burkholderia-sp.-D-99:0.0424988997,WP_205714023.1-Burkholderia-cepacia:0.0404688023)75/100:0.0095887424,WP_174919663.1-Burkholderia-metallica:0.0266784593)80.6/100:0.0354284464,WP_077021900.1-Burkholderia-cenocepacia:0.1331520265)99.5/100:0.3079733433)78.7/100:0.0425158726,(WP_159673755.1-Andreprevotia-sp.-IGB-42:0.2428970558,WP_084093192.1-Andreprevotia-lacus:0.1106347901)99.7/100:0.2966246155)91.6/98:0.1019529640)72.7/83:0.0220820474,((WP_043620211.1-Chromobacterium-piscinae:0.2554030589,(WP_148717329.1-Chitinolyticbacter-meiyuanensis:1.0970351210,WP_045052286.1-Chromobacterium-violaceum:0.2493911425)87.7/88:0.1745943458)26.9/58:0.0752920122,WP_103900886.1-Chromobacterium-alticapitis:0.3885915273)96.6/92:0.1831046355)88/97:0.0653651260)74.6/82:0.0391200629,(((((((((((((WP_250059255.1-Halomonas-sp.-ATCH28:0.4867801219,WP_179153381.1-Chromohalobacter-salexigens:0.2748845526)89.9/95:0.1256430706,WP_250939176.1-unclassified-Larsenimonas:0.3740555140)82.2/91:0.0692944698,WP_106478747.1-Phytohalomonas-tamaricis:0.3938988647)79.5/90:0.0636519251,(WP_218634960.1-Marinobacter-arenosus:0.1654358706,WP_240716217.1-Halomonas-neptunia:0.1361390309)98.1/100:0.2210226627)97.4/88:0.1510806617,(((WP_225237063.1-Massilia-oculi:0.4355987778,WP_147937383.1-Massilia-arenae:0.1814459544)97.1/100:0.1913696827,(WP_192051557.1-unclassified-Massilia:0.1042292928,WP_136222466.1-Massilia-sp.-Mn16-1_5:0.1811348427)79.8/100:0.1128521718)94.3/80:0.1289095939,(WP_209394989.1-Rhodoferax-sp.-AJA081-3:0.3902123248,WP_094844257.1-Bordetella-genomosp.-11:0.2454467320)88.8/100:0.0808813139)92.1/66:0.0763321641)0/35:0.0000024050,(((((((((((((WP_138213821.1-Pseudomonas-nitroreducens:0.0500461660,WP_017517140.1-Pseudomonas-nitroreducens:0.0292394915)100/100:0.4652570464,WP_085721622.1-Pseudomonas:0.2107858178)64.1/63:0.0201101004,WP_104727848.1-Pseudomonas-oleovorans:0.2007223008)62/87:0.0430047666,((WP_177138191.1-Hydrogenophaga-aromaticivorans:0.1874589609,(WP_236964528.1-Hydrogenophaga-sp.-SL48:0.0867001780,(WP_226493830.1-Hydrogenophaga-taeniospiralis:0.0531275570,WP_210824367.1-Hydrogenophaga-aromaticivorans:0.0853349243)79.2/100:0.0728034319)98.1/100:0.1307777404)36.9/77:0.0421127460,WP_133157403.1-Hydrogenophaga-pseudoflava:0.1998627143)94.1/100:0.1038404383)82.9/93:0.0619037350,(WP_042945119.1-Pseudomonas-fluorescens:0.0622013847,WP_181643258.1-Pseudomonas-sivasensis:0.1066062021)99.6/100:0.2024495086)48.7/49:0.0455500424,(WP_169965430.1-unclassified-Pseudomonas:0.0780347997,WP_047278503.1-Pseudomonas-lundensis:0.0183069244)99.8/100:0.1931251309)80.1/51:0.0413136485,(WP_065086507.1-Pseudomonas:0.3716406730,WP_205489922.1-Pseudomonas-lactucae:0.1999562185)70.1/67:0.0098964513)89.8/81:0.0486894293,(WP_184594840.1-Pseudomonas-nitritireducens:0.2711442740,WP_192381882.1-Pseudomonas-sp.-PDM20:0.1373810190)99.7/100:0.2005491199)83/67:0.0321618321,((((WP_191471912.1-Pseudomonas-asiatica:0.2696821539,(WP_043318452.1-Pseudomonas-sp.-PI1:0.0555946385,WP_043268312.1-Pseudomonas:0.0443009474)45.4/100:0.0286790421)94.2/100:0.0943309573,WP_089390899.1-Pseudomonas-delhiensis:0.1685867410)85.7/100:0.0391841115,WP_076426141.1-Pseudomonas-alcaligenes:0.2073187183)90.5/100:0.0823132448,WP_233297479.1-Pseudomonas:0.2168366012)43.1/92:0.0339044864)81.8/47:0.0306060180,WP_214501190.1-Pseudomonas-sp.-RC3H12:0.3904882495)43.8/54:0.0327047720,WP_155457065.1-Massilia-eburnea:0.2722636254)67.9/4:0.0213451107,(((WP_029241333.1-Pseudomonas:0.0938103052,((WP_054996663.1-Pseudomonas-ficuserectae:0.0575385677,(WP_007252556.1-Pseudomonas-syringae-group:0.0655358129,WP_025390602.1-Pseudomonas-syringae:0.0328183422)15.9/89:0.0239777768)93.7/100:0.0603869641,WP_125830676.1-Pseudomonas-sp.-v388:0.0972075183)88.4/100:0.0580377573)91.8/100:0.1112214288,((WP_016973326.1-Pseudomonas-tolaasii:0.1472501590,WP_099548838.1-Pseudomonas-sp.-2995-3:0.1435365808)85.9/100:0.0475548730,(WP_177015116.1-Pseudomonas-costantinii:0.0755912858,WP_141063556.1-Pseudomonas-fluorescens:0.1305621148)91.6/100:0.0851432568)92.5/100:0.0988048250)99.4/100:0.1895519652,(WP_058488093.1-Pseudomonas:0.2225656578,WP_043256200.1-Pseudomonas-knackmussii:0.1965854333)87.8/98:0.0864868741)81.2/91:0.0414642703)91.2/11:0.0506779036,(((WP_020564242.1-Methylosarcina-fibrata:0.3130094059,((WP_180152801.1-Parapusillimonas-granuli:0.2096389189,(WP_088146851.1-Achromobacter-denitrificans:0.1417375397,WP_105555808.1-Achromobacter-sp.-MYb9:0.1118605380)97.6/100:0.1333099091)80.2/84:0.0453218866,(((WP_074973890.1-Pseudomonas-otitidis:0.2227304549,WP_068803504.1-Immundisolibacter-cernigliae:0.3190910862)69.1/98:0.1201751113,WP_011111299.1-Nitrosomonas-europaea:0.2744539011)0/31:0.0075491897,WP_046857648.1-Pseudomonas-sp.-CCOS-191:0.2721606273)71.8/82:0.0310456148)88.8/79:0.0703549584)75.5/56:0.0186165922,(WP_101152997.1-Pseudomonas-sp.-Choline-3u-10:0.1705392649,((WP_046617631.1-Stutzerimonas-stutzeri:0.0962144429,WP_135265705.1-Stutzerimonas-stutzeri:0.0761020789)89.2/100:0.0840700335,WP_003299494.1-Stutzerimonas-stutzeri:0.1283028735)88/100:0.0482905620)93.2/100:0.0751271215)88.6/50:0.0286248463,((WP_209872004.1-Pseudomonas-sp.-BP8:0.2781194068,WP_066093302.1-Hydrogenophaga-crassostreae:0.2165173410)63.4/16:0.0975262525,(WP_125865093.1-Stutzerimonas-stutzeri:0.2155529992,WP_200290630.1-Stutzerimonas-balearica:0.1872025839)89.9/97:0.0949953618)49.9/7:0.0203007208)63.7/6:0.0135284900)88.1/61:0.0264758827)87.5/75:0.0256092886,((WP_076518421.1-Achromobacter-sp.-MFA1-R4:0.1285986559,(WP_104142565.1-Achromobacter-spanius:0.0273244365,WP_120157261.1-Achromobacter-sp.-B7:0.0740545384)83.9/98:0.0418613802)91.8/98:0.0996849092,WP_158658301.1-Achromobacter-sp.-AONIH1:0.5327759592)95/99:0.1282366797)0/51:0.0000029594,(WP_077733980.1-Paenalcaligenes-hominis:0.0653098058,WP_167661700.1-Paenalcaligenes-hominis:0.0335335449)100/100:0.4138938893)88.6/72:0.0285115606,((((((WP_136867012.1-Guyparkeria-sp.-SB14A:0.5222351137,WP_217963608.1-Candidimonas-humi:0.3904334745)8.4/55:0.0658146059,WP_102075761.1-Pollutimonas-subterranea:0.2339494636)89.8/60:0.1162791076,(((((WP_220354719.1-Rhizobium-cauense:0.3369530860,WP_016557527.1-Rhizobium-grahamii:0.1698055115)88.9/100:0.0690325477,((WP_184450570.1-Rhizobium-sp.-BK538:0.3045061545,(WP_072373837.1-Rhizobium-tibeticum:0.1284844483,WP_007537558.1-Rhizobium-mesoamericanum:0.0717594040)76/100:0.0616296767)60.4/99:0.0336240894,WP_018896487.1-Rhizobium-sp.-2MFCol3.1:0.3154751402)57.2/99:0.0251316547)85.2/100:0.0682365076,(WP_153273632.1-Rhizobium-grahamii:0.3313930808,WP_233136718.1-Agrobacterium-vaccinii:0.3658630484)19.5/33:0.0366104874)99.2/100:0.2584117254,(((WP_057247240.1-Duganella-sp.-Root198D2:0.3337379007,WP_212592286.1-Janthinobacterium-sp.-TND4EL3:0.2947652092)83.9/86:0.0746909642,WP_102071501.1-Pollutimonas-nitritireducens:0.4027557137)37.4/21:0.0371077705,WP_047550623.1-Methylotenera-sp.-G11:0.3087114597)85/29:0.0555429820)79.6/10:0.0487471129,WP_227683402.1-Pusillimonas-sp.-MFBS29:0.3813041036)57.7/10:0.0586003022)75.5/49:0.0310190978,WP_216129682.1-Azospira-inquinata:0.5547535987)73.8/55:0.0156998402,(WP_019141909.1-Noviherbaspirillum-massiliense:0.2706841525,WP_146330669.1-Noviherbaspirillum-sp.-UKPF54:0.2991118381)88.8/93:0.0875965751)83.7/77:0.0528248519,(((((WP_129151777.1-Achromobacter-aloeverae:0.1365160252,WP_088599355.1-unclassified-Achromobacter:0.0898955392)97.2/100:0.1466675920,WP_057654143.1-Bordetella-sp.-N:0.2393382154)90.4/100:0.1069612629,WP_086067427.1-Bordetella:0.2643110105)84.3/100:0.0684424188,(WP_066641528.1-Bordetella-sp.-H567:0.1130757426,(WP_086058993.1-Bordetella-genomosp.-9:0.2611179258,WP_094856450.1-Bordetella-genomosp.-10:0.2098949841)93.8/100:0.1393610342)87.2/100:0.1033875140)90.1/99:0.0967653045,WP_091467897.1-Methylophilus-rhizosphaerae:0.5674597275)92.7/92:0.1064448959)83.6/69:0.0460242188)15.1/9:0.0165988338,(((WP_088420173.1-Pseudomonas-nitroreducens:0.1731980144,WP_254475170.1-unclassified-Pseudomonas:0.0912626586)86.4/99:0.0394534799,WP_205338011.1-Pseudomonas-sp.-PDNC002:0.0886782295)97.8/100:0.1622946323,(WP_033955816.1-Pseudomonas-aeruginosa:0.0213139522,WP_003153917.1-Pseudomonas-aeruginosa-group:0.0650521404)100/100:0.3030200199)94.6/59:0.1089316908)63.2/5:0.0077949983,((WP_230502293.1-Janthinobacterium-sp.-FW305-128:0.6861201704,WP_249918475.1-Pseudomonas-sp.-R3.Fl:0.1969182178)85.2/60:0.1333547632,((WP_092579398.1-Achromobacter-sp.-NFACC18-2:0.6276898681,WP_223462047.1-Pseudomonas-sp.-A-R-26:0.2899570269)47.8/88:0.0591960124,WP_058422493.1-Pseudomonas:0.2756079235)84.1/95:0.0800274243)86.2/20:0.0657724544)72.6/10:0.0140392995,((((((((((((((((WP_188630571.1-Oceanisphaera-marina:0.1931467531,WP_104485062.1-Oceanisphaera-arctica:0.1533664451)77.3/100:0.0291148234,WP_204795415.1-Oceanisphaera-litoralis:0.0400038888)99.8/100:0.2833616679,((WP_072282603.1-Syntrophotalea-acetylenivorans:0.2167158945,((WP_092056970.1-Desulfuromonas-acetexigens:0.2371245983,WP_043770486.1-Desulfobulbus-propionicus:0.3449033778)66.3/91:0.0879452006,(WP_028317018.1-Desulfobulbus-elongatus:0.5769646736,WP_148894253.1-Geothermobacter-ehrlichii:0.4352748059)30/12:0.1050187332)29.7/10:0.0620094139)43.8/10:0.0170570413,WP_155876508.1-Desulfuromonas-sp.-AOP6:0.4653966360)77.6/100:0.1323532751)48.5/98:0.0648784464,WP_072282412.1-Syntrophotalea-acetylenivorans:0.3432181242)87.7/70:0.0846948673,((WP_040200185.1-Geoalkalibacter-subterraneus:0.3455485634,WP_040097704.1-Geoalkalibacter-ferrihydriticus:0.3380373407)38.6/94:0.0771293924,WP_066727622.1-Desulfuromonas-sp.-DDH964:0.4342186458)83.8/68:0.0632684987)50.2/76:0.0339177943,((WP_137423090.1-Desulfoglaeba-alkanexedens:0.5254827943,WP_027714505.1-Desulfuromonas-sp.-TF:0.3676050484)80.7/60:0.0834765075,WP_053550565.1-Desulfuromonas-soudanensis:0.4089582664)87.6/69:0.0854928108)52.9/57:0.0179711973,((((((((((((((WP_089129802.1-Tolypothrix-sp.-NIES-4075:0.1232338937,WP_096563566.1-Scytonema-sp.-NIES-4073:0.1615331257)87.5/89:0.0598074985,WP_048866388.1-Scytonema-tolypothrichoides:0.1147221193)41.4/89:0.0283634937,WP_096563464.1-Scytonema-sp.-NIES-4073:0.1480063931)99.4/100:0.2173285725,WP_096557790.1-Nostoc-sp.-NIES-4103:0.2390442227)69.7/95:0.0437722883,((((WP_009344618.1-Cylindrospermopsis-raciborskii:0.1075137022,WP_240868223.1-Cylindrospermopsis-raciborskii:0.0299169184)99.8/100:0.2094840155,WP_096570274.1-Sphaerospermopsis:0.0685417226)35.2/96:0.0131659899,WP_220611028.1-Sphaerospermopsis-torques-reginae:0.0774019763)98.1/100:0.1209710706,((WP_237989437.1-Nostoc-sp.-UHCC-0870:0.1031575549,WP_190559626.1-Anabaena-sphaerica:0.1342592314)75.3/100:0.0595310711,WP_206263219.1-Nostoc-sp.-UHCC-0702:0.0928344771)86.9/98:0.0640779624)88.9/99:0.0332737861)75.5/84:0.0165375968,((((WP_015128325.1-Calothrix-sp.-PCC-7507:0.1631512041,WP_017653186.1-Fortiea-contorta:0.2441354749)5.7/89:0.0626228832,((WP_193199005.1-Nostoc-sp.-MG11:0.0654282048,WP_162396505.1-Nostoc-sp.-B-2019:0.0603212323)94.8/100:0.0927350695,WP_214439445.1-Atlanticothrix-silvestris:0.1944102428)43.5/99:0.0816234986)82.1/97:0.0413288541,(((WP_045870014.1-unclassified-Tolypothrix:0.0376848035,WP_190656031.1-Nostoc-linckia:0.0461691382)98.2/100:0.1020331240,(WP_190743070.1-Tolypothrix-sp.-FACHB-123:0.0612869481,WP_190545010.1-Calothrix:0.0469988688)97.1/100:0.0950713098)44.9/100:0.0460374310,WP_086758595.1-Nostoc-sp.-106C:0.0929087427)32.1/98:0.0375088281)25.2/83:0.0400263871,WP_016866333.1-Fischerella:0.1270750687)81.7/81:0.0350973832)85.5/99:0.0800398213,(WP_044881819.1-Neochlamydia-sp.-EPS4:0.0824724288,WP_166155173.1-unclassified-Neochlamydia:0.0000021042)100/100:0.7095182401)31.8/74:0.0804981249,(WP_015193936.1-Stanieria-cyanosphaera:0.0572192681,WP_096384422.1-Stanieria-sp.-NIES-3757:0.1000706370)99.8/100:0.2602151143)84.1/93:0.0491785722,WP_015143663.1-Pleurocapsa-sp.-PCC-7327:0.1784603055)90/87:0.0975223016,((((((((WP_106232846.1-Pleurocapsa-sp.-CCALA-161:0.2354206612,WP_229639079.1-Waterburya-agarophytonicola:0.0357027502)81.3/100:0.0394163239,WP_096721927.1-Chondrocystis-sp.-NIES-4102:0.1278782074)99.2/100:0.2029513914,WP_006510035.1-Xenococcus-sp.-PCC-7305:0.2506681856)91.5/100:0.0920357496,WP_036478304.1-Myxosarcina-sp.-GI1:0.2563641529)9.3/98:0.0333804183,WP_012595695.1-Rippkaea-orientalis:0.2922793925)38.6/54:0.0439287365,WP_264322079.1-Zarconia-navalis:0.2820186171)79.9/54:0.0319292298,((WP_007308862.1-Crocosphaera-watsonii:0.1206243058,WP_107668138.1-Cyanothece-sp.-BG0011:0.0471302674)4.6/85:0.0095525460,(WP_008278255.1-Crocosphaera-chwakensis:0.0852380112,WP_009545547.1-Crocosphaera-subtropica:0.0289983088)54.7/61:0.0112151747)99.7/98:0.2097416694)88.1/97:0.0309205230,(WP_106909014.1-Microcystis-sp.-MC19:0.3733248178,(WP_124977990.1-Aphanothece-sacrum:0.3099064593,WP_013324143.1-Gloeothece-verrucosa:0.1975421851)0/33:0.0100327574)81.1/46:0.0226702887)81.3/48:0.0409827311)53.3/72:0.0433362527,((((((((WP_102179481.1-Fischerella-thermalis:0.0434734272,((WP_096675774.1-Fischerella-sp.-NIES-4106:0.0670595509,WP_053458209.1-Hapalosiphon-sp.-MRB220:0.0441170096)73/99:0.0086849031,(WP_131119525.1-Westiellopsis-prolifica:0.0668915812,WP_016860528.1-Fischerella-muscicola:0.0349936407)84.6/100:0.0186333426)96.1/99:0.0980316491)99.7/100:0.2725144552,WP_190825375.1-Iningainema-tapete:0.3756802957)23.1/19:0.0167528349,((WP_015202791.1-Crinalium-epipsammum:0.3117797353,WP_035154029.1-Calothrix-sp.-336/3:0.5166651894)74.7/97:0.1057901000,((WP_099102643.1-Nostoc-sp.-Peltigera-malacea-cyanobiont-DB3992:0.1162879840,((WP_190950926.1-Nostoc-punctiforme:0.0624618769,(WP_221704991.1-Desmonostoc-muscorum:0.0636953715,WP_096537650.1-Nostoc-linckia:0.0243652083)74.8/98:0.0091534323)64/96:0.0347817706,WP_190959861.1-Nostoc:0.0251729371)97.5/99:0.1039671887)99.8/100:0.1596967742,((((((WP_193938462.1-Cuspidothrix-issatschenkoi:0.2240587824,((WP_096666215.1-Dolichospermum-compactum:0.0811660693,(WP_190387835.1-Aphanizomenon:0.0812122809,WP_168466156.1-Aphanizomenon:0.0279881115)47.3/99:0.0218147222)82/83:0.0193855610,WP_015080416.1-Anabaena-sp.-90:0.0414918331)70.9/83:0.0175206889)86.1/52:0.0261476280,(((WP_190370804.1-Anabaena-sp.-FACHB-1237:0.2001797899,(WP_190392428.1-Dolichospermum-sp.-FACHB-1091:0.0178756844,(WP_028089231.1-Dolichospermum-circinale:0.0257286763,WP_137908967.1-Nostocales:0.0649030060)99.2/100:0.0797612613)98.7/100:0.1234173682)41.5/95:0.0384869130,WP_168537172.1-Anabaena-sp.-UHCC-0253:0.0927295428)83/43:0.0203448398,WP_168502193.1-Anabaena-sp.-UHCC-0187:0.0098013740)69.8/43:0.0065989120)100/100:0.1821422076,(WP_010998622.1-Nostocaceae:0.0840694517,WP_011320078.1-Nostocaceae:0.0798218971)75.3/100:0.0252288020)93.8/100:0.0671279145,((WP_015138800.1-Nostoc-sp.-PCC-7524:0.0996959867,WP_066425233.1-Anabaena-sp.-4-3:0.1269491299)97.7/100:0.0936497500,(WP_199337214.1-Nostoc-sp.-FACHB-152:0.1629389528,(WP_239729951.1-Nodularia-sphaerocarpa:0.0906521530,WP_173403323.1-Nodularia-spumigena:0.0660043689)92.2/100:0.0615490727)79.6/96:0.0331052669)72.9/93:0.0105916052)81.6/97:0.0248198214,WP_199337786.1-Nostoc-sp.-FACHB-280:0.1867258476)94.4/100:0.0859227288,(WP_096624129.1-Calothrix-sp.-NIES-3974:0.1022119240,WP_095721999.1-Calothrix-elsteri:0.1355440601)98.9/100:0.1588154067)78.3/100:0.0275543488)91.6/100:0.0664967802)12.1/17:0.0337113359)97.4/100:0.1082308321,(WP_194029353.1-Lusitaniella-coriacea:0.3206882046,WP_190493486.1-unclassified-Microcoleus:0.3161475384)0/20:0.0627228396)72.6/15:0.0152984429,(((((WP_224410051.1-Oscillatoria-salina:0.2926063363,WP_250123015.1-Chroococcidiopsis-sp.-CCMEE-29:0.2094541333)92.9/96:0.1042277531,WP_190531554.1-Coleofasciculus-sp.-FACHB-542:0.2402057712)79.4/90:0.0335570070,((WP_233747769.1-Leptodesmis-sichuanensis:0.3010054754,(WP_192216758.1-Microcoleus-sp.-FACHB-SPT15:0.2209242542,WP_224345144.1-Kovacikia-minuta:0.1422305145)41.4/86:0.0271967083)80.9/88:0.0203771319,(WP_223049852.1-Leptothermofonsia-sichuanensis:0.0932125653,WP_073549891.1-Chroogloeocystis-siderophila:0.2431847030)81.5/99:0.0513957863)91.2/90:0.0457317156)0/17:0.0000029500,WP_008178350.1-Moorena-producens:0.3428695885)36.1/11:0.0157860788,((WP_193873404.1-unclassified-Coleofasciculus:0.1831204964,WP_254010234.1-Symplocastrum-sp.-BBK-W-15:0.1741507034)85.4/27:0.0528486971,(WP_190666400.1-Coleofasciculus-sp.-FACHB-501:0.1595793427,WP_190525322.1-Coleofasciculus-sp.-FACHB-T130:0.0830077927)97.1/100:0.1021617247)66/17:0.0106147459)87/76:0.0249268482)79.8/26:0.0415136878,(((WP_015170385.1-Geitlerinema-sp.-PCC-7407:0.3544917154,WP_017661508.1-Baaleninema-simplex:0.3869448567)80.7/88:0.0694923937,(WP_226578226.1-Microseira-wollei:0.2784323351,((WP_073595522.1-Phormidium-ambiguum:0.1006108138,WP_193982193.1-Phormidium-sp.-LEGE-05292:0.0937492605)99.3/100:0.1402660326,(WP_249064905.1-Argonema-antarcticum:0.0702428096,WP_190468950.1-unclassified-Planktothrix:0.1163910422)79.3/97:0.0220279686)86.9/97:0.0544424372)70.3/99:0.0218141256)82.7/59:0.0311550131,WP_007355666.1-Kamptonema:0.1960960693)57.2/38:0.0357045097)41.1/17:0.0117186994,((((((((((WP_202221768.1-Okeania-sp.-KiyG1:0.0615265299,WP_094672483.1-Hydrocoleum-sp.-CS-953:0.0618939476)66.7/100:0.0224403387,WP_124146462.1-Okeania-hirsuta:0.0083616457)99.6/100:0.2326709400,WP_071516849.1-Geitlerinema-sp.-PCC-9228:0.3195738180)49.7/99:0.0725965595,WP_008052766.1-Limnospira-indica:0.3835726155)71.4/33:0.0448574430,(WP_009783380.1-Lyngbya:0.1167056070,WP_046277942.1-Limnoraphis-robusta:0.0523373679)97.2/100:0.1020454491)91.2/65:0.0676920908,(((((WP_236507253.1-Tychonema-sp.-BBK16:0.0524341890,WP_096828456.1-Tychonema-bourrellyi:0.0883998507)91.3/100:0.0457141532,WP_242724542.1-Microcoleus-vaginatus:0.0443314157)80.6/100:0.0149705865,WP_015174083.1-Oscillatoria-nigro-viridis:0.0246013744)84.9/99:0.0237805932,(WP_194009709.1-unclassified-Tychonema:0.0492870198,WP_265235823.1-Lyngbya-sp.-CCAP-1446/10:0.0543128261)87.4/100:0.0280364287)75.4/83:0.0300409146,WP_193973976.1-Microcoleus-sp.-LEGE-07076:0.0384748045)99.8/100:0.1421141737)30.7/5:0.0354405067,(WP_190877489.1-unclassified-Planktothricoides:0.1950747298,((WP_083623900.1-Planktothrix-serta:0.0500611559,WP_072717475.1-Planktothrix:0.0372990142)35.2/36:0.0238213001,(WP_079676747.1-Planktothrix-sp.-PCC-11201:0.0336862489,(WP_026797127.1-Planktothrix:0.0359321488,WP_214559879.1-Planktothrix-agardhii:0.0530238102)44.3/99:0.0371873645)98.2/100:0.0962356072)99/100:0.1365203706)87.8/7:0.0477759442)0/51:0.0000022809,((WP_261197618.1-unclassified-Laspinema:0.0530960748,(WP_207087219.1-Phormidium-pseudopriestleyi:0.0434300606,WP_261204505.1-Laspinema-sp.-D2d:0.0544815631)84.3/100:0.0279517556)96.9/100:0.1186015433,WP_200665545.1-Oxynema-sp.-CENA135:0.3214903543)90.9/84:0.0671158592)66/68:0.0211867044,WP_069965800.1-unclassified-Desertifilum:0.4311400755)81.9/19:0.0373595608,(WP_017715496.1-Oscillatoria-sp.-PCC-10802:0.2007612936,(WP_190665495.1-Microcoleus-sp.-FACHB-672:0.0849799087,WP_261220767.1-Ancylothrix-sp.-D3o:0.2563667272)96.4/99:0.1032074000)92.2/99:0.0656612597)70.4/16:0.0101127942)83.1/22:0.0485735957,((((WP_204137747.1-Halomicronema-sp.-CCY15110:0.0665511722,WP_039724568.1-Leptolyngbya-sp.-LK:0.1043012589)99.7/100:0.2577873724,WP_023076112.1-Leptolyngbya-sp.-Heron-Island-J:0.3221164201)91.9/97:0.1528585554,WP_017324474.1-Synechococcus-sp.-PCC-7336:0.4214376477)81.9/97:0.0485691845,WP_190772446.1-unclassified-Leptolyngbya:0.2503164355)86.2/92:0.1025589883)72.5/85:0.0516870646)78.8/100:0.1506771006,WP_066347484.1-Geminocystis-sp.-NIES-3708:0.7625782708)43.4/34:0.0218482110,(((WP_205222281.1-Desulfogranum-marinum:0.3712404544,WP_205242362.1-Desulfobulbus-alkaliphilus:0.2006644647)87/96:0.0849833553,WP_267928740.1-Desulfolithobacter-dissulfuricans:0.1685983647)98.7/96:0.1594302248,((WP_250680171.1-Geomonas-sp.-Red32:0.3078240040,WP_199381999.1-Geomesophilobacter-sediminis:0.2781354371)82.6/100:0.0668948609,WP_205218910.1-Desulforhopalus-vacuolatus:0.5324116691)88.3/98:0.0895119473)83.3/32:0.0344371374)93.7/87:0.0700234533,((WP_015405619.1-Desulfocapsa-sulfexigens:0.3581375541,(((WP_163339734.1-Desulfopila-sp.-IMCC35008:0.4563504488,WP_136805251.1-Desulfosediminicola-flagellatus:0.3979738086)38.7/100:0.0193220888,WP_073615262.1-Desulfopila-aestuarii:0.3711652911)96.7/100:0.1723945482,(WP_228856481.1-Desulfomarina-profundi:0.5150116148,WP_183349429.1-Desulfoprunum-benzoelyticum:0.2318024137)83.1/100:0.0681784127)78.1/100:0.0660226201)89.1/92:0.0536446590,WP_028582690.1-Desulfogranum-japonicum:0.4995613220)69.9/70:0.0509514948)74.9/48:0.0361042587)77.4/61:0.0234195981,(WP_103114677.1-Geothermobacter-hydrogeniphilus:0.4705639859,WP_221248821.1-Desulfuromonas-versatilis:0.3628631657)88.2/87:0.1307598204)88.9/92:0.0463680314,((((((WP_212718705.1-unclassified-Synechocystis:0.0386918983,WP_228015966.1-Synechocystis-salina:0.1375277627)84.6/100:0.0490518341,WP_010873214.1-unclassified-Synechocystis:0.0463004471)99.9/100:0.5038957096,WP_204103763.1-Spirulina-major:0.6784122066)87.7/97:0.1561330711,(WP_084055900.1-Desulfacinum-hydrothermale:0.4763718295,WP_123289658.1-Desulfosoma-caldarium:0.3857487110)97.4/100:0.2584223418)82.2/94:0.0579154207,((((WP_116346290.1-Cupriavidus-taiwanensis:0.1779792991,WP_124685047.1-Cupriavidus-pauculus:0.3025560562)100/100:0.5542777152,(WP_220807781.1-Noviherbaspirillum-aridicola:0.7616284987,WP_119020982.1-Halomonas-sp.-JS92-SW72:0.3291780502)87.1/98:0.1970799765)81/94:0.0983404105,((WP_200222041.1-Achromobacter-xylosoxidans:0.2182579946,WP_259251071.1-Achromobacter-sp.-JUb104:0.2173804240)89/100:0.1061988351,(((((WP_111826915.1-Achromobacter-sp.-HZ01:0.1621678863,WP_102775120.1-Achromobacter:0.0256159171)100/100:0.2036839425,(WP_129525645.1-Achromobacter-agilis:0.0425148621,(WP_129240460.1-Achromobacter-veterisilvae:0.0568245187,WP_043544564.1-Achromobacter-sp.-RTa:0.0399082658)87/100:0.0254321581)48.3/98:0.0189462869)87.1/100:0.0831466709,(((WP_175208175.1-Achromobacter-anxifer:0.0773967373,(((WP_251864322.1-Achromobacter-sp.-Marseille-Q4962:0.0978179743,WP_164793551.1-Achromobacter-insolitus:0.0431068893)97.7/95:0.0757275588,WP_207472676.1-Achromobacter-sp.-SD115:0.0131361353)93.9/95:0.0471208384,WP_125471167.1-Achromobacter:0.0226263634)96.2/97:0.0543829883)78.9/92:0.0163963012,WP_175164188.1-Achromobacter-aegrifaciens:0.0550679319)91.4/96:0.0492673945,WP_088158656.1-Achromobacter-xylosoxidans:0.0800840207)99.7/100:0.1377720465)58.1/87:0.0083241221,WP_174717299.1-Achromobacter-denitrificans:0.1240794645)91/93:0.0472075743,(WP_088588627.1-Achromobacter-marplatensis:0.2273085908,WP_169273406.1-Achromobacter-sp.-Bel:0.1194360844)96.5/100:0.1264726732)78.6/81:0.0427682279)94.6/100:0.1292722358)88.5/81:0.0726241391,(WP_099797666.1-Thermostichus-lividus:0.6802402058,WP_196167318.1-Pseudomonas-monteilii:0.3819962466)93.9/97:0.2452140974)72.7/78:0.0201199017)92.6/93:0.0431783994,(((((((((WP_238511579.1-Cupriavidus-gilardii:0.3542294307,WP_150984056.1-Cupriavidus-basilensis:0.6454741836)80.8/82:0.0963934784,WP_224042884.1-Cupriavidus-respiraculi:0.4239128880)77.2/77:0.0573970132,WP_116600700.1-Cupriavidus-alkaliphilus:0.2386160968)13.1/36:0.0276019478,(WP_018008101.1-Cupriavidus-neocaledonicus:0.0678962724,WP_183298446.1-Cupriavidus-alkaliphilus:0.0760197289)99.9/100:0.2008017230)69.2/60:0.0249724409,(((WP_226989104.1-Desulfuromonas-sp.-TF:0.5800724865,WP_227459030.1-Cupriavidus-pauculus:0.4354952904)94.7/94:0.2514095179,((WP_108623287.1-Orrella-marina:0.6380093097,WP_243478771.1-Orrella-sp.-f23:0.4765569265)92.4/99:0.2516760023,WP_227181580.1-Leeia-sp.-TBRC-13508:0.3436990143)38.6/86:0.0388442169)86.2/84:0.0879237852,WP_213445339.1-Variovorax-sp.-PCZ-1:0.5157819655)64.1/82:0.1030954171)94/83:0.1063811914,WP_136680209.1-Neptunomonas-sp.-XY-337:0.3141576902)82.7/44:0.0325235775,(((((((((((WP_207000528.1-Trinickia-sp.-DHG64:0.1777380356,WP_124149903.1-Paraburkholderia-dinghuensis:0.1522493243)71.5/100:0.0676586341,(WP_266250364.1-Paraburkholderia-megapolitana:0.2299685196,WP_026121090.1-Paraburkholderia-kururiensis:0.1507528691)53.1/100:0.0711648990)94.1/100:0.1002092638,(((((((((((WP_059459219.1-Burkholderia-vietnamiensis:0.3215306602,WP_121276956.1-Trinickia-fusca:0.1693748939)91.5/99:0.0738103314,WP_136892825.1-Trinickia-sp.-7GSK02:0.0426373474)83.2/99:0.0375457434,WP_201650988.1-Paraburkholderia-sabiae:0.0499643379)26.1/90:0.0165020371,WP_205970438.1-Paraburkholderia-sp.-Tr-20389:0.0508356987)83/97:0.0234827686,WP_054928771.1-Paraburkholderia-caribensis:0.0452921628)85.5/95:0.0262457358,((WP_205976236.1-Paraburkholderia-sp.-Se-20369:0.3159375603,WP_206951398.1-Trinickia-sp.-4D114:0.0844134132)97.1/93:0.1186969708,WP_059889870.1-Burkholderia-stagnalis:0.2829476359)20.7/29:0.0204086937)75.6/87:0.0182946336,WP_209935850.1-Paraburkholderia-sp.-LEh10:0.1257135417)86.4/88:0.0283199291,(((((WP_060092900.1-Burkholderia-vietnamiensis:0.0313805628,WP_226113708.1-Burkholderia-sp.-AU33545:0.0568578826)99.7/100:0.2016771902,WP_069272519.1-Burkholderia-cepacia:0.3222719405)12.6/14:0.0450501009,((WP_175791145.1-Burkholderia-ambifaria:0.1300695144,WP_059842355.1-Burkholderia-multivorans:0.0480727138)74.5/96:0.0157338906,WP_105772332.1-Burkholderia-multivorans:0.0463898685)50.9/95:0.0195678893)58.7/12:0.0112939218,(WP_006496666.1-Burkholderia-cenocepacia:0.0406061926,WP_059716874.1-Burkholderia-ubonensis:0.0470207641)91.7/100:0.0372265629)79.2/13:0.0147805550,(WP_168792792.1-Paraburkholderia-aromaticivorans:0.1046492980,(WP_004544228.1-Burkholderia:0.0660757241,WP_059583574.1-pseudomallei-group:0.0417869478)75/99:0.0287397105)82.5/95:0.0296717390)71.7/11:0.0172273409)85.4/17:0.0265735721,(WP_060156927.1-Burkholderia-ubonensis:0.1365824468,WP_061168975.1-Caballeronia-hypogeia:0.1303859258)21.5/8:0.0082857344)91.6/95:0.0381315717,((WP_201138986.1-Paraburkholderia-domus:0.0621691783,WP_266262001.1-Paraburkholderia-madseniana:0.0325780026)98.9/100:0.1349673547,WP_009889585.1-Burkholderia-thailandensis:0.1323401906)84.6/100:0.0312066516)86.2/98:0.0666501794,WP_134195781.1-Paraburkholderia-rhizosphaerae:0.3924976871)81.4/92:0.0420815489)90.5/92:0.0772655435,WP_038711716.1-Burkholderia-sp.-lig30:0.3536239059)7.4/22:0.0171668711,WP_179404315.1-Burkholderia-guangdongensis:0.2272235230)95.7/78:0.1141842772,((WP_063494936.1-Paraburkholderia-phytofirmans:0.1024422389,WP_254318368.1-Paraburkholderia-sp.-CNPSo-3272:0.0000020715)63.1/78:0.0440173117,WP_042262232.1-Paraburkholderia-heleia:0.0432871104)98.5/97:0.1601869742)68/95:0.0583625609,WP_056136218.1-Massilia-sp.-Root133:0.2837679785)93.9/98:0.1062734577,((WP_115680924.1-Cupriavidus-taiwanensis:0.3409488141,WP_018313502.1-Cupriavidus-sp.-UYPR2.512:0.4016557100)39.7/10:0.0200414780,(WP_135702508.1-Cupriavidus-oxalaticus:0.2101290387,(WP_071014919.1-Cupriavidus-sp.-USMAHM13:0.3150832802,WP_174426450.1-Cupriavidus-basilensis:0.5548767341)90.1/100:0.1405292583)70.9/9:0.0240865362)85.3/92:0.0490651878)92.8/99:0.0602307710,(WP_228215938.1-Aromatoleum-toluclasticum:0.3216590577,WP_199264031.1-Desulfobulbus-oligotrophicus:0.4546166253)75.6/80:0.0681096452)74.2/86:0.0184035969,((WP_183165579.1-Azomonas-macrocytogenes:0.2542240465,(((WP_102836461.1-Stutzerimonas-stutzeri:0.0388747026,(WP_256071125.1-Stutzerimonas-stutzeri:0.0472428697,WP_045423249.1-Pseudomonadaceae:0.0385290150)83.7/98:0.0176319047)95.7/99:0.0715490373,(WP_255961233.1-Stutzerimonas-stutzeri:0.0387001725,WP_025243649.1-Stutzerimonas-stutzeri:0.0651611462)3.5/98:0.0047506899)98.9/96:0.0786955012,(WP_122077451.1-Pseudomonadaceae:0.0460461514,((WP_037024448.1-Pseudomonas-saudiphocaensis:0.0931962031,WP_181084609.1-Stutzerimonas-stutzeri:0.0103486117)90.2/100:0.0410372433,(WP_011911332.1-Stutzerimonas-stutzeri:0.0593441058,(((WP_230358311.1-Pseudomonas-sp.-NP21570:0.0235221690,(WP_193680295.1-Pseudomonas-lopnurensis:0.0614520648,WP_200589398.1-Stutzerimonas-stutzeri:0.0313314397)63.6/100:0.0463377002)90.2/98:0.0331458146,(WP_237255234.1-Pseudomonas-oligotrophica:0.0947015065,WP_043295157.1-Stutzerimonas-degradans:0.0258793817)96.6/99:0.0582089783)75.5/88:0.0096049117,WP_120997453.1-_Pseudomonas:0.0465636202)89.1/90:0.0295378514)86.8/97:0.0285458565)79.3/94:0.0195989119)0/89:0.0000028039)64.9/96:0.0380198077)56.2/83:0.0584969039,WP_028238613.1-Stutzerimonas-azotifigens:0.1013760868)98.8/100:0.1436290016)89.1/87:0.0432883682,((WP_250024752.1-Cupriavidus-campinensis:0.2011069384,WP_094797980.1-Bordetella-genomosp.-5:0.3279380476)82.3/65:0.1218712540,WP_226921657.1-Kinneretia-sp.-XES5:0.4095442014)67.8/56:0.0435746990)49.1/44:0.0154146355)88.9/65:0.0604761122,((((((((((((((WP_209862330.1-Pseudomonas-sp.-BP8:0.7600178688,(WP_054891693.1-Pseudomonas:0.1089018016,WP_236238170.1-Pseudomonas-faucium:0.2000225027)6.1/35:0.0336705209)71/95:0.0220468973,(WP_158461286.1-Pseudomonas-fluorescens:0.3276369945,(WP_200141208.1-unclassified-Pseudomonas:0.2732000535,WP_029612831.1-Pseudomonas-parafulva:0.0294216553)94.6/100:0.0924487111)63.9/97:0.0511837043)91.7/96:0.0579020460,(((WP_217887155.1-Pseudomonas-xanthosomae:0.0470786494,WP_217840807.1-Pseudomonas-fakonensis:0.0540417441)73.6/100:0.0193123755,WP_186654015.1-Pseudomonas-xantholysinigenes:0.3122317619)86.2/99:0.0543754082,(WP_201176141.1-Pseudomonas-sp.-S31:0.0272069993,WP_261240966.1-unclassified-Pseudomonas:0.0735221926)99.2/100:0.1502529378)77.8/82:0.0228176005)81.8/78:0.0286248727,WP_225786622.1-Pseudomonas:0.1103630673)49.7/89:0.0440050316,WP_110605176.1-Pseudomonas-soli:0.3814745150)94.1/100:0.0876802300,((WP_181290220.1-Pseudomonas-brassicacearum:0.2124434962,((((WP_201004194.1-Pseudomonas-cichorii:0.1098692898,WP_176569640.1-Pseudomonas-eucalypticola:0.1837860635)90.9/90:0.0658724712,(WP_219063768.1-Pseudomonas-sp.-UMAB-08:0.3043984769,((WP_222540780.1-Pseudomonas-sp.-DR208:0.1654777788,(WP_253509974.1-Pseudomonas-rhodesiae:0.1600523359,WP_093202821.1-Pseudomonas-yamanorum:0.0375971780)86.6/100:0.0539705552)70.6/81:0.0163810176,((WP_065875813.1-Pseudomonas:0.0612384583,(WP_033898696.1-Pseudomonas-fluorescens-group:0.0489248686,(WP_186687394.1-Pseudomonas-sp.-SWRI18:0.0729915658,WP_169378974.1-Pseudomonas:0.0221869268)81.5/100:0.0337993012)62.2/100:0.0154852107)98.9/100:0.1019045761,WP_105697656.1-Pseudomonas-poae:0.0562928979)76.1/100:0.0153343068)45/81:0.0521840202)89.2/100:0.0602842055)85.5/90:0.0487293118,WP_043192328.1-Pseudomonas-rhizosphaerae:0.3751296895)56.8/65:0.0123088805,(WP_249678359.1-Pseudomonas-sp.-PIA16:0.2331864442,WP_268799018.1-Pseudomonas-sp.-BSw22131:0.2264080189)68.3/78:0.0241285663)87.6/93:0.0767675515)79.9/99:0.0540367023,WP_108096457.1-Pseudomonas-sp.-GV071:0.1614517777)50.6/30:0.0853046991)92.8/100:0.0597272980,WP_252273535.1-Pseudomonas-sp.-YIM-B01952:0.1436343659)83.1/100:0.0320946105,WP_143488038.1-Pseudomonas-mangiferae:0.1988381089)83/38:0.0452389041,(((((WP_058067682.1-Pseudomonas-sp.-TTU2014-080ASC:0.2017003158,(WP_077525673.1-Pseudomonas-resinovorans:0.1619953568,WP_137821430.1-Pseudomonas-sp.-D-2018:0.1056621640)56.4/30:0.0383195961)96.3/96:0.0789253486,(WP_003453405.1-Pseudomonas:0.1690253759,(WP_016495240.1-Pseudomonas-resinovorans:0.0723766739,WP_044873077.1-Pseudomonas-sp.-LFM046:0.0896520517)32.3/97:0.0298579869)72.7/27:0.0137212536)77.3/25:0.0187691585,WP_028626762.1-Pseudomonas-resinovorans:0.0617266286)31.7/10:0.0091278419,(WP_120654068.1-Pseudomonas-sp.-DY-1:0.1129781370,WP_151138079.1-Pseudomonas-lalkuanensis:0.0555497110)84.7/97:0.0183629072)87/22:0.0371721053,(WP_192399936.1-Pseudomonas-sp.-PDM15:0.2185657076,WP_226282979.1-Pseudomonas-lalkuanensis:0.0843792130)97.6/100:0.1251024585)50.8/36:0.0398853377)86.8/26:0.0554557239,((((WP_090448460.1-Pseudomonas-benzenivorans:0.2883123929,(WP_025164100.1-Pseudomonas-taeanensis:0.0806358868,WP_147170838.1-Pseudomonas-sp.-SJZ079:0.0066682806)94.8/100:0.0715313438)0/42:0.0000028841,((WP_208707982.1-Pseudomonas-sp.-LPB0260:0.0387968869,WP_255840290.1-Pseudomonas-benzenivorans:0.0512182579)92.8/100:0.0596949073,((WP_131198269.1-Pseudomonas-dryadis:0.1741433174,(WP_131178052.1-Pseudomonas-daroniae:0.0851062817,(WP_084307530.1-Pseudomonas-flavescens:0.0961861129,(WP_249909557.1-Pseudomonas-fulva:0.0616105204,(WP_074889242.1-Pseudomonas-argentinensis:0.0358907876,WP_027910132.1-Pseudomonas-sp.-URMO17WK12_I4:0.0623409187)38.9/96:0.0185436300)85.4/99:0.0498180411)75.1/96:0.0347661607)88/96:0.0526930745)95.8/100:0.1111944236,(WP_233685494.1-Pseudomonas-anguilliseptica:0.0798974005,WP_257778987.1-Pseudomonas-sp.-32.2.56:0.0000023102)99.5/100:0.1435501947)73.3/68:0.0327960917)69.2/55:0.0387021126)86.9/60:0.0502188657,(WP_069518708.1-Pseudomonas:0.1942953116,((((((((WP_119694090.1-Pseudomonas:0.0636204129,WP_106738315.1-Pseudomonas-mendocina:0.0676081074)92/100:0.0486432728,WP_150606948.1-Pseudomonas-oleovorans:0.0765859958)89.3/98:0.0482319888,WP_024309956.1-Pseudomonas-sp.-P818:0.0563788865)64.2/74:0.0160727414,WP_090337209.1-Pseudomonas-chengduensis:0.0287177663)0/74:0.0004605111,WP_143497094.1-Pseudomonas-sp.-ALS1131:0.0503997344)63.8/31:0.0226977316,WP_074939507.1-Pseudomonas-composti:0.0293150647)86.3/99:0.0231219737,WP_021444205.1-Pseudomonas-sp.-EGD-AK9:0.0702154164)89.2/99:0.0344999227,WP_159970704.1-Pseudomonas-sp.-8Z:0.1013037979)94.2/100:0.0815504961)16.9/18:0.0198616238)89.5/27:0.0390040769,WP_238040454.1-Pseudomonas-sp.-MMS21-TM103:0.1604877630)73.1/100:0.0289946280)95.2/90:0.0805700247,(WP_220803742.1-Pseudomonas-sp.-NCCP-436:0.2079556251,(WP_002810067.1-Nitrosococcus-oceani:0.0934528513,WP_013221233.1-Nitrosococcus-watsonii:0.0637340256)100/100:0.4303344552)83.1/96:0.0285170089)74.5/36:0.0164103207,WP_090225506.1-Pseudomonas-kuykendallii:0.1639632446)88.4/37:0.0667979823,WP_025281494.1-Ectothiorhodospira-haloalkaliphila:0.6932076884)7.1/92:0.0244841153,WP_226440716.1-Quatrionicoccus-australiensis:0.5658079831)53.7/86:0.0364934425)85.6/51:0.0239604223,((((WP_006216420.1-Achromobacter-piechaudii:0.1675112816,WP_219991400.1-Achromobacter-sp.-ES-001:0.1238567903)91.1/100:0.0681954882,WP_183017169.1-Achromobacter-sp.-UMC46:0.0718400845)81.6/71:0.0425816737,WP_100503206.1-Achromobacter-ruhlandii:0.1975986089)57.6/71:0.0746674424,WP_175192416.1-Achromobacter-deleyi:0.0537756403)100/100:0.4265683973)0/39:0.0000022516)94.4/89:0.0395584774)90.2/88:0.0469541094,(((((((WP_239190817.1-Candidatus-Nitrotoga-sp.-HW29:0.0765877864,WP_239797721.1-Candidatus-Nitrotoga-arctica:0.0396396951)68.6/68:0.0098918057,WP_239287644.1-Candidatus-Nitrotoga-sp.-1052:0.0278197344)0/65:0.0018322075,WP_239235002.1-Candidatus-Nitrotoga-sp.-BS:0.0811041087)100/100:0.3992175916,((WP_237889684.1-Dissulfurimicrobium-hydrothermale:0.4470741664,WP_009207329.1-Sulfuricella-denitrificans:0.5155776004)47.5/42:0.0255876467,WP_237247350.1-Sideroxyarcus-emersonii:0.2743569652)69.9/66:0.0904317220)58.9/91:0.0518472941,((((((((((((WP_065305794.1-Janthinobacterium-psychrotolerans:0.1750548517,WP_034778721.1-Janthinobacterium-lividum:0.0485691546)53.6/100:0.0036514351,WP_035822645.1-Janthinobacterium-sp.-RA13:0.0860769327)76.2/100:0.0108614207,WP_152282105.1-Janthinobacterium-violaceinigrum:0.0266454723)77.5/100:0.0086615568,WP_183742413.1-unclassified-Janthinobacterium:0.0630818763)75.4/100:0.0083924383,(((((WP_139090581.1-Janthinobacterium-lividum:0.0878140566,WP_230539136.1-Janthinobacterium-sp.-FW305-129:0.0531226817)87.3/94:0.0196239449,WP_094442213.1-Janthinobacterium-sp.-PC23-8:0.0844828151)36/94:0.0086323694,WP_215403374.1-Janthinobacterium-sp.-JC611:0.0324044157)74.2/100:0.0113442222,WP_102124214.1-Janthinobacterium-sp.-ROICE36:0.0448243205)76.8/100:0.0086969062,WP_096238059.1-Janthinobacterium-svalbardensis:0.0256588845)77.1/100:0.0088163525)99.5/100:0.1470489209,WP_023495308.1-Methyloglobulus-morosus:0.2479989203)80.7/96:0.0316935926,(WP_099917421.1-Massilia-psychrophila:0.1743240624,(WP_205319084.1-Glaciimonas-sp.-PAMC28666:0.0965820457,WP_092396716.1-Collimonas-sp.-OK412:0.1657994199)74.7/98:0.0219538832)89.1/92:0.0387172044)0/65:0.0045491004,WP_101482628.1-unclassified-Janthinobacterium:0.2741631847)90.8/95:0.0808785623,WP_250479419.1-unclassified-Caballeronia:0.5331910383)83.2/88:0.0399128490,((WP_162020603.1-Undibacterium-crateris:0.1944932199,WP_186884883.1-Undibacterium:0.0674746492)80/99:0.0395019719,WP_186861877.1-Undibacterium-griseum:0.0564797141)100/100:0.2093053566)78.5/71:0.0292297178,((WP_247737070.1-Thauera-aromatica:0.0681541486,WP_039805139.1-Azotobacter-chroococcum:0.2255689665)98.7/100:0.1952434399,((WP_195874647.1-Herminiimonas-contaminans:0.1772634374,WP_041297045.1-Janthinobacterium-sp.-Marseille:0.1207196305)80.5/100:0.0376600394,(WP_076590920.1-Herminiimonas-arsenitoxidans:0.2190701603,(WP_206604468.1-Herminiimonas-fonticola:0.0562369173,WP_188420499.1-Oxalicibacterium-solurbis:0.2384648469)48.5/99:0.0202455689)86.2/100:0.0549901671)98.8/100:0.1490105516)54.7/61:0.0210497241)65.2/54:0.0625919713,WP_217900122.1-Noviherbaspirillum-humi:0.2906283308)89/100:0.0979261322)75.3/65:0.0476718656,((WP_006529211.1-Gloeocapsa-sp.-PCC-73106:0.4851246539,WP_047549861.1-Methylotenera-sp.-G11:0.2513850012)71.4/82:0.0394202953,WP_217345508.1-Noviherbaspirillum-sp.-L7-7A:0.3531005163)75.3/76:0.0293245490)79.8/61:0.0408282231,WP_054689510.1-Desulfosarcina-cetonica:0.4251739241)91.1/92:0.0677890503)83.2/53:0.0286555609,((((((((((WP_198111724.1-Inhella-proteolytica:0.5993864825,(WP_037482909.1-Sphaerotilus-natans:0.2779218994,(WP_028996355.1-Azohydromonas-australica:0.0986296855,WP_066337818.1-Azohydromonas-lata:0.1325567634)93.9/100:0.1304215879)85.8/100:0.1217313304)55.8/91:0.0708558997,WP_241396682.1-Pelomonas-sp.-CA6:0.1402187659)97.7/99:0.2561988810,((WP_074865046.1-Atopomonas-hussainii:0.3259858606,WP_019561187.1-Caldimonas-manganoxidans:0.3146473522)73.5/88:0.0468072439,WP_123210144.1-Stagnimonas-aquatica:1.1703702273)85.8/89:0.0820251203)29.2/89:0.0490857081,WP_047193087.1-Schlegelella-brevitalea:0.3682935871)80.1/98:0.0915154560,((((WP_132974196.1-Thiogranum-longum:0.3474278611,WP_201344172.1-Thiohalobacter-sp.-COW1:0.1890450634)100/100:0.5398619700,(WP_234070476.1-Myxococcus-stipitatus:0.3272779935,WP_120555541.1-Corallococcus-aberystwythensis:0.3996736102)98.3/100:0.3543685063)39/28:0.0688523284,WP_110265962.1-Sinimarinibacterium-flocculans:0.5764950362)68/57:0.1315835291,(((((WP_169107474.1-Comamonas-suwonensis:0.0671488545,WP_219343559.1-Comamonas-sp.-Y33R10-2:0.1109630441)94.6/100:0.0678482207,(WP_003073420.1-Comamonas:0.0926434383,((WP_211457163.1-Comamonas-sp.-PE63:0.0680721572,WP_148729134.1-Comamonas-sp.-Z3:0.0579584259)67.8/94:0.0088986963,WP_034391098.1-Comamonas-thiooxydans:0.0225574995)73/94:0.0114811739)89.8/100:0.0362893269)59.4/95:0.0184447788,WP_218241177.1-Comamonas-fluminis:0.1579575277)99.5/100:0.2616924410,(WP_012203208.1-Delftia-acidovorans:0.3135846526,WP_058879441.1-Comamonas-kerstersii:0.1749923399)41.7/99:0.0546814414)88.5/100:0.1034048295,WP_066475960.1-Comamonas-terrae:0.1727653932)97.4/100:0.2716443492)75.8/25:0.1385256952)88.3/24:0.0940427735,(WP_164156863.1-Sandarakinorhabdus-rubra:0.3697250007,((WP_070114691.1-Acidithiobacillus-caldus:0.4328830665,((WP_009566950.1-Acidithiobacillus:0.0550505397,WP_215870853.1-unclassified-Acidithiobacillus:0.1083838457)75.7/100:0.0724554098,(WP_101536718.1-Acidithiobacillus-sp.-SH:0.2396725211,WP_163095688.1-Acidithiobacillus-ferrianus:0.0484263989)49.4/83:0.0587202520)90.7/99:0.1296270389)9.2/98:0.1153732748,WP_152145882.1-Alkalilimnicola-sp.-S0819:0.5550070063)70.2/99:0.1359156300)94.2/90:0.1450846870)12/60:0.0193132540,((WP_108647113.1-Polynucleobacter-rarus:0.0313317946,WP_084281813.1-Polynucleobacter-kasalickyi:0.0555653990)100/100:0.7816104940,((WP_215371371.1-Polynucleobacter-sp.-MWH-UH24A:0.2740668633,WP_173942040.1-Polynucleobacter-antarcticus:0.2241756326)87.4/100:0.1007737597,((((WP_128113711.1-Polynucleobacter-necessarius:0.1325653492,WP_215385554.1-Polynucleobacter-sp.-AP-Kolm-20A-A1:0.0979077936)91.7/100:0.0875839968,WP_216177372.1-Polynucleobacter-sp.-AP-Feld-500C-C5:0.2323694162)83.9/96:0.0445427733,WP_114639791.1-Polynucleobacter-necessarius:0.1588313657)90.6/87:0.0824324780,((WP_216254756.1-Polynucleobacter-sp.-AP-Melu-500A-A1:0.0684587065,WP_216218975.1-Polynucleobacter-sp.-AP-Latsch-80-C2:0.0520330306)98.7/100:0.1184587730,((((WP_215319389.1-Polynucleobacter-paneuropaeus:0.1273480812,(WP_215352354.1-Polynucleobacter-sp.-MWH-P3-07-1:0.0587874910,WP_215380622.1-Polynucleobacter-paludilacus:0.0479367468)47.9/100:0.0429668391)95.3/100:0.0758047176,WP_062307935.1-Polynucleobacter-sinensis:0.0713763025)35.6/89:0.0217562012,(((((((WP_215376847.1-Polynucleobacter-asymbioticus:0.0345583360,WP_215328794.1-Polynucleobacter-sp.-AP-Sving-400A-A2:0.0699508041)97.4/100:0.0696303505,WP_173959551.1-Polynucleobacter-arcticus:0.0395077795)92.2/100:0.0324265048,WP_216225867.1-Polynucleobacter-sp.-MG-27-Goln-C1:0.1130759888)41.4/100:0.0084178104,(WP_216168716.1-Polynucleobacter-bastaniensis:0.1101632866,WP_216178894.1-Polynucleobacter-sp.-80A-SIGWE:0.0188987263)88.1/98:0.0222457089)86.2/88:0.0299170011,WP_215299358.1-Polynucleobacter-sp.-AP-Nino-20-G2:0.1187960338)86.9/85:0.0271459459,(WP_216268685.1-Polynucleobacter-sp.-Nonnen-W13:0.0642266603,(WP_215334312.1-Polynucleobacter-sp.-AP-Ainpum-60-G11:0.0591437901,WP_251374783.1-Polynucleobacter-corsicus:0.0427721878)74.6/95:0.0077158640)87.8/78:0.0166630887)0/83:0.0000023349,WP_216197047.1-Polynucleobacter-sp.-UK-Mo-2m-Kol15:0.1004575057)1.1/34:0.0158439764)88.2/100:0.0334526762,WP_076024025.1-Polynucleobacter-sphagniphilus:0.1143597877)70.2/89:0.0092565682)96.1/99:0.0812647963)11.2/72:0.0536502357)97.6/97:0.3169307154)96.1/98:0.2559810959)80.8/87:0.0589144154,((WP_160809027.1-Nitrosomonas-sp.-HPC101:0.7005971010,WP_154206467.1-Ralstonia-pickettii:0.1950417348)63.9/98:0.0799810235,WP_253536546.1-Ralstonia-sp.-21YRMH01-3:0.5833306562)74.6/87:0.0677432599)0/74:0.0012806306,(((WP_036252969.1-Methylobacter-sp.-BBA5.1:0.3784741098,((((WP_019918777.1-Methyloversatilis-discipulorum:0.0675889290,WP_018229738.1-Methyloversatilis-universalis:0.0695979704)89.6/100:0.1216125366,WP_183409942.1-Litorivivens-lipolytica:0.6977137625)19.3/90:0.0776719112,(WP_101941536.1-Uliginosibacterium-sp.-TH139:0.1399585037,WP_095523004.1-Candidatus-Dactylopiibacterium-carminicum:0.1401437201)97/100:0.2089499358)89.1/100:0.1086012467,WP_101249932.1-Telmatospirillum-siberiense:0.7743019905)23.9/67:0.0717953506)92.5/78:0.0869245220,((((WP_109041087.1-Azospira-sp.-I13:0.1522273740,WP_014237023.1-Azospira-oryzae:0.1788858093)97/100:0.1732783436,WP_067106371.1-Methylobacillus-sp.-MM3:0.2964742439)56.9/96:0.0594163255,WP_201673630.1-Ramlibacter-monticola:0.3073597392)80.3/92:0.0365792226,WP_212785890.1-Ferrigenium-kumadai:0.3255755081)70.9/84:0.0606475153)79.8/66:0.0302136842,(((((WP_074796726.1-Nitrosospira-briensis:0.0957174340,(WP_090882638.1-Nitrosovibrio-sp.-Nv6:0.0772102495,WP_097065525.1-Nitrosovibrio-sp.-Nv4:0.0446388692)90.9/100:0.0773746196)99.7/100:0.2853520215,(WP_220807643.1-Noviherbaspirillum-aridicola:0.4339262192,WP_122244186.1-Vandammella-animalimorsus:0.2690812364)88.5/97:0.1096784668)70.3/77:0.0603325953,(WP_223986959.1-Cupriavidus-pampae:0.0706756648,((WP_011517306.1-Cupriavidus-metallidurans:0.0648652632,WP_019451253.1-Cupriavidus-sp.-BIS7:0.0688806374)64/86:0.0068609158,WP_082079449.1-Cupriavidus-basilensis:0.1055048675)81.8/86:0.0378667124)99.8/100:0.2446193829)74.4/76:0.0343073952,((((((WP_106935816.1-unclassified-Variovorax:0.3551970351,(WP_126747644.1-Variovorax-sp.-DXTD-1:0.2251965601,((WP_093207264.1-Variovorax-sp.-YR750:0.1573360952,WP_206177103.1-Variovorax-sp.-RKNM96:0.1420589962)26.5/36:0.0326230903,WP_068630051.1-Variovorax-sp.-PAMC-28711:0.3522577604)54.5/27:0.0159958381)82/50:0.0517014361)48.2/80:0.0170098444,WP_219210863.1-Variovorax-boronicumulans:0.2543588387)70.9/37:0.0417905954,((WP_192194832.1-Variovorax-sp.-38R:0.2851420576,WP_160553990.1-Xylophilus-rhododendri:0.2236228535)56.8/77:0.0647431189,WP_093441791.1-Variovorax-sp.-770b2:0.2021073943)92.8/78:0.0806180542)91.7/97:0.0763634822,(WP_169420802.1-Ramlibacter-agri:0.3087996500,WP_166067558.1-Diaphorobacter-sp.-HDW4B:0.3116120359)76.4/58:0.0335266182)76.5/56:0.0168069611,(WP_026436726.1-Acidovorax-sp.-JHL-9:0.2543301013,WP_086926522.1-Acidovorax-carolinensis:0.2796650628)41.3/76:0.0730536889)87.4/92:0.0313390139,((WP_174484170.1-methanotrophic-endosymbiont-of-Bathymodiolus-puteoserpentis:0.3434927518,WP_137890779.1-Ramlibacter-sp.-2FC:0.1526622303)72.2/40:0.0182064931,WP_137918285.1-Hydrogenophaga-sp.-2FB:0.2561141420)74.9/43:0.0179970762)73.5/64:0.0128821030)76.8/70:0.0165087597,(WP_057673948.1-Curvibacter-sp.-PAE-UM:0.2475301405,WP_056898839.1-Pseudorhodoferax-sp.-Leaf274:0.3845968614)42.6/77:0.0259099531)83.6/80:0.0420135483)88/65:0.0366098210)98.2/75:0.0892042496,((WP_014706296.1-Methylophaga-nitratireducenticrescens:0.0000025690,WP_136552334.1-Methylophaga-sp.-SB9B:0.0996306934)100/100:0.6547476414,(WP_153584563.1-Caenimonas-koreensis:0.4011544584,(WP_007185580.1-Hydrocarboniphaga-effusa:0.0954825362,WP_007185565.1-Hydrocarboniphaga-effusa:0.1101757935)99.2/100:0.2716184871)89.4/100:0.1817269957)79/83:0.0537680912)76.1/67:0.0138798727)87.9/80:0.0419250738,WP_054772520.1-Methylogaea-oryzae:0.3187719672)86.1/68:0.0491296607,(((WP_227390214.1-Halomonas-malpeensis:0.8798216076,WP_104228621.1-Solimonas-fluminis:0.4144045054)90.3/45:0.1596539209,(WP_083560927.1-Oceanococcus-atlanticus:0.4556613150,(WP_046292890.1-Pandoraea-oxalativorans:0.1196348996,WP_150719538.1-Pandoraea-capi:0.1200677981)100/100:0.4576396230)44.8/43:0.0957803414)0/32:0.0000021235,WP_153720048.1-Spiribacter-sp.-C176:0.4194636027)95.4/70:0.1606258797)85/60:0.0285408821,(((WP_205202385.1-Azonexus-hydrophilus:0.2925102737,WP_173581083.1-Azonexus-fungiphilus:0.3725392321)31.6/85:0.0297055741,WP_121459656.1-Azonexus-fungiphilus:0.4059188044)41.2/82:0.1024536716,((WP_117174481.1-Rhodoferax-lacus:0.3288297425,((WP_094482496.1-Rhodoferax-sp.-TH121:0.1243081144,WP_105262671.1-Rhodoferax-sp.-TS-BS-61-7:0.0436170434)83.8/100:0.0532904403,WP_214121631.1-Curvibacter-sp.-CHRR-16:0.3606100171)91.1/100:0.1679790251)42.1/47:0.0446878773,(WP_066884254.1-Dechloromonas:0.1393096374,WP_027458229.1-Dechloromonas-agitata:0.1531845116)95.5/100:0.1337807213)84.8/91:0.0864709269)67.3/67:0.0706196420)40.5/58:0.0137212565,((((WP_176062241.1-Anaeromyxobacter-diazotrophicus:0.2529031501,WP_248361410.1-Anaeromyxobacter-oryzae:0.3325866963)98.3/100:0.2406575040,WP_258826205.1-Massilia-agri:0.5435527905)69.1/92:0.0399424020,(((WP_018882239.1-unclassified-Thioalkalivibrio:0.5819001775,WP_125198734.1-Guyparkeria-sp.-SCN-R1:0.2418782948)85.1/99:0.0944596543,(((WP_026296529.1-Thioalkalivibrio-sp.-ALJ2:0.1733900947,WP_019569965.1-Thioalkalivibrio-sp.-ALE11:0.1086322614)88/100:0.0322331115,WP_028484075.1-Thioalkalivibrio-sp.-ALE17:0.0550856876)73.4/100:0.0112628554,WP_018169543.1-Thioalkalivibrio-sp.-ALMg9:0.0425685991)97.7/100:0.1490003394)52.6/97:0.0410789175,WP_250072676.1-Guyparkeria-hydrothermalis:0.2824231802)99.3/99:0.2102805065)94/78:0.1070561512,((((WP_167682126.1-Rhodocyclus-gracilis:0.4443397102,(WP_153115862.1-Rhodocyclus-tenuis:0.0571336835,WP_200385811.1-Rhodocyclus-tenuis:0.0327364437)94.9/100:0.1204932428)88.9/100:0.1095993303,WP_201212349.1-Rhodocyclus-purpureus:0.1949381819)97.9/99:0.1781331659,WP_263768642.1-Propionivibrio-sp.-SG131:0.4168655719)41.3/73:0.0313721882,WP_153110537.1-Propionivibrio-limicola:0.3740182657)75.8/79:0.0345971798)76.7/70:0.0855743204)40/28:0.0272789608,(((((((((WP_269022943.1-Methylomonas-sp.-MP1:0.1101865160,(WP_140911374.1-Methylomonas-koyamae:0.2682605872,(WP_071159776.1-Methylomonas-sp.-LWB:0.2221181433,((WP_020485671.1-Methylomonas-sp.-MK1:0.2587811163,(WP_256607223.1-Methylomonas-sp.-WSC-7:0.1025786903,(WP_205450646.1-Methylomonas-sp.-EFPC1:0.1095876922,WP_064036612.1-Methylomonas-methanica:0.0000024053)0/90:0.0031959897)91.6/100:0.0963627183)90.1/79:0.0903454591,WP_036273427.1-Methylomonas:0.1689785408)0/55:0.0000023926)89.1/100:0.0883222139)96.7/99:0.1310991096)77.2/68:0.0574593212,(WP_196437371.1-Methylomonas-sp.-LL1:0.1678008434,(WP_150047608.1-Methylomonas-rhizoryzae:0.4316287826,WP_066983474.1-Methylomonas-lenta:0.1132490772)87.4/41:0.0992609941)83.4/43:0.0642658625)97.6/85:0.1822419176,(WP_148817865.1-Pigmentiphaga-aceris:0.4922407024,WP_004074545.1-Desulfobacter-postgatei:0.2873589115)78.7/85:0.1038708391)61/60:0.0441050146,(((WP_238617884.1-Ectothiorhodospira-shaposhnikovii:0.1719793859,WP_025282248.1-Ectothiorhodospira-haloalkaliphila:0.1650642833)86/100:0.0873301529,WP_238622182.1-unclassified-Ectothiorhodospira:0.1739508043)93.2/91:0.0828699868,(WP_256068944.1-Stutzerimonas-stutzeri:0.2742822606,((((((((((((WP_060482506.1-Pseudomonas-sp.-NBRC-111119:0.2524244829,WP_159409308.1-Pseudomonas-putida:0.1149406341)91.4/52:0.0728240928,(WP_182365213.1-Pseudomonas:0.1551636044,((WP_110607173.1-Pseudomonas-fulva:0.0175371027,WP_088523134.1-unclassified-Pseudomonas:0.0670240367)95.4/100:0.0499871014,(WP_167339957.1-Pseudomonas-fulva:0.0385857420,WP_186476580.1-Pseudomonas-promysalinigenes:0.0574029895)80.9/97:0.0319466352)51/80:0.0129237319)78.1/16:0.0239661940)0/10:0.0000029259,(WP_201214098.1-Pseudomonas-sp.-S36:0.0275962463,WP_054890059.1-Pseudomonas:0.0809503195)96.8/99:0.0635268458)74.1/16:0.0173713800,WP_267821885.1-Pseudomonas-putida:0.1129978768)68.4/17:0.0054798271,WP_182365203.1-Pseudomonas-juntendi:0.1091523412)28.3/20:0.0094148099,WP_115272924.1-Pseudomonas-putida:0.2182161290)0/15:0.0000024513,(WP_167277242.1-Pantoea-sp.-Tr-811:0.0241746411,WP_103449615.1-Pseudomonas-putida:0.0720067324)98.6/100:0.0760240275)63.6/16:0.0165197595,((WP_196129978.1-Pseudomonas-fulva:0.1189393523,WP_201164743.1-Pseudomonas-sp.-S60:0.0494002275)98.4/100:0.1240670263,(((WP_174142546.1-Pseudomonas-putida:0.1883247447,WP_027907237.1-Pseudomonas:0.0493548916)73.6/94:0.0162908327,WP_075044077.1-Pseudomonas-putida:0.0584242510)66.2/93:0.0332282539,(WP_023379297.1-Pseudomonas-sp.-VLB120:0.1235256436,WP_202675963.1-Pseudomonas-sp.-SK2:0.0518418552)98.4/100:0.0962111820)88.1/100:0.0565730724)91.8/93:0.0430592835)84.1/16:0.0234005864,(WP_046489762.1-Pseudomonas-veronii:0.1520635418,(WP_059395284.1-Pseudomonas-putida:0.1182881990,(WP_009685980.1-Pseudomonas:0.1055104850,WP_024086460.1-Pseudomonas:0.0805921884)86.8/100:0.0607552115)82.2/84:0.0471921181)46.3/15:0.0712476840)79.7/35:0.0247601221,(WP_029885231.1-Pseudomonas:0.0314640833,(WP_230064532.1-Pseudomonas-sp.-CIP-10:0.0272558601,WP_197863301.1-Pseudomonas-putida:0.0539111435)95.5/84:0.0590323430)18/43:0.0061828659)95.5/85:0.0818283503,WP_060394622.1-Pseudomonas-entomophila:0.1481077395)96.8/100:0.1083170344,WP_183165449.1-Azomonas-macrocytogenes:0.1595609223)84.7/98:0.0491193466)76.8/65:0.0394188763)89.9/57:0.0528820978)91/28:0.0664510922,(WP_018865194.1-Thioalkalivibrio-sp.-ARh3:0.2340891464,WP_048706771.1-Thauera-humireducens:0.4491536958)91.3/86:0.0736174257)56.4/24:0.0134301602,((WP_019964198.1-Pannonibacter-phragmitetus:0.9335774393,WP_138632053.1-Pseudoalteromonas-spongiae:0.4166995748)84.6/95:0.0910434532,WP_064030819.1-Methylomonas-koyamae:0.3335695174)89.4/55:0.0860898069)61.8/24:0.0115958089,(WP_168987795.1-Azoarcus-taiwanensis:0.3072454838,WP_013164002.1-Desulfurivibrio-alkaliphilus:0.4422642427)83.8/76:0.0731248243)94.4/52:0.0774310039,(((WP_136386324.1-Pseudothauera-rhizosphaerae:0.3065744598,WP_114649035.1-Pseudothauera-hydrothermalis:0.1206550928)36.5/44:0.0359599215,WP_187718084.1-Thauera-sp.-CAU-1555:0.1713037369)98.9/100:0.2059454806,WP_026685886.1-Azovibrio-restrictus:0.2565941157)79.9/57:0.0827404590)73.3/94:0.0158264080,(WP_066098770.1-Halothiobacillus-sp.-LS2:0.1879174747,WP_070073009.1-Acidihalobacter-aeolianus:0.2670761590)92.7/99:0.0915978188)92.6/84:0.0851884931)83.7/66:0.0548893714)96.1/84:0.1304691139,(WP_044615503.1-Gynuella-sunshinyii:0.7167619746,((WP_139871681.1-Candidatus-Methylopumilus-universalis:0.3284521791,(WP_139929800.1-Candidatus-Methylopumilus-universalis:0.4224623387,WP_140005423.1-Candidatus-Methylopumilus-planktonicus:0.1164681980)90.6/100:0.1505320298)99/100:0.3559569124,WP_058575749.1-unclassified-Guyparkeria:0.3441682273)4.6/75:0.0297361795)71.6/93:0.1102116605)95.5/96:0.1253408214)88.4/92:0.0683421771)97.1/55:0.1489387807,(((((((((((WP_236369955.1-Aeromonas-veronii:0.0624510504,WP_113736183.1-Aeromonas-caviae:0.0529851366)22.8/75:0.0192355374,((WP_042019775.1-Aeromonas-sobria:0.0716556917,WP_139495680.1-Aeromonas-veronii:0.0329925256)95.5/100:0.0747308455,(WP_098969895.1-Aeromonas-sp.-CU5:0.0649314935,WP_159163188.1-Aeromonas-veronii:0.0685422764)89.8/100:0.0480496152)85.7/100:0.0437386729)86.4/97:0.0375735698,(((((WP_080988732.1-Aeromonas-caviae:0.0575303266,WP_108536035.1-Aeromonas-sp.-HMWF014:0.0562757631)92/100:0.0482210546,(WP_252615985.1-Aeromonas-hydrophila:0.0745494598,WP_041208446.1-Aeromonas-jandaei:0.3293934066)72/97:0.0202902068)85.8/97:0.0252289076,WP_214003767.1-Aeromonas-caviae:0.0705200936)64.9/97:0.0084441815,(WP_113739100.1-Aeromonas-veronii:0.0777079643,(WP_226032029.1-Aeromonas-hydrophila:0.0341984489,WP_139459904.1-Aeromonas-veronii:0.1924887216)80.5/97:0.0208961370)75.5/96:0.0139695295)31.2/98:0.0206110582,WP_143237783.1-Aeromonas-caviae:0.0686304958)94.3/100:0.0791548105)93.1/99:0.0650356344,WP_201035193.1-Aeromonas-media:0.1797455952)86.7/88:0.0578570025,(WP_082969903.1-Aeromonas-dhakensis:0.1664204789,WP_104454419.1-Aeromonas-sp.-ASNIH2:0.0740514262)76.6/87:0.0250283225)81.9/97:0.0406571840,(WP_220777726.1-Shewanella-hafniensis:0.0437392970,WP_055647506.1-Shewanella-sp.-Sh95:0.0396134940)95/100:0.0720261958)87.1/98:0.0635717758,(WP_263173612.1-Shewanella-sp.-SM73:0.1943075482,((WP_263191039.1-Shewanella-sp.-SM87:0.0908481693,WP_263255112.1-Shewanella-sp.-SM32:0.0592493869)79.8/100:0.0208277101,(WP_263178828.1-Shewanella-sp.-SM101:0.0546380551,WP_109248233.1-Shewanella-algae:0.0932929382)94.4/100:0.0556958309)87.3/100:0.0497793985)75.7/98:0.0266578623)95.9/98:0.1066342762,((WP_248067241.1-unclassified-Shewanella:0.1344972455,WP_148505453.1-Vibrio-cholerae:0.1315712646)87.5/100:0.0787212373,(WP_218558465.1-Shewanella-putrefaciens:0.1014669299,WP_011919582.1-Shewanella-putrefaciens:0.0446083290)100/100:0.1924532286)77.3/87:0.0529952795)84.8/60:0.0501042311,((((WP_147247296.1-Brenneria-salicis:0.0460658436,WP_172289906.1-unclassified-Brenneria:0.1519238550)83.8/100:0.0542033999,WP_039356706.1-Pectobacterium-actinidiae:0.1194087529)82.5/78:0.1399369446,((WP_198997094.1-Pectobacterium-carotovorum:0.0301825820,(WP_165707548.1-Pectobacterium:0.0465688996,WP_174866005.1-Pectobacterium-polaris:0.0341093983)99/100:0.0867056725)82/78:0.0249300707,WP_145962313.1-Pectobacterium-peruviense:0.0703004798)97.5/100:0.1588658992)93.6/100:0.1389414142,(WP_208149206.1-Shewanella-algae:0.4405344427,(WP_219142360.1-Aeromonas-sanarellii:0.2489627535,(((WP_045500143.1-Vibrio-hyugaensis:0.2491582227,WP_045419977.1-Vibrio-campbellii:0.1132815661)52/100:0.0481961563,WP_025533544.1-Vibrio-parahaemolyticus:0.0917311979)94.6/100:0.0597313698,WP_011788809.1-Shewanella-sp.-W3-18-1:0.1010190183)0/43:0.0020849535)19.8/42:0.0325655321)98/100:0.2104659844)80.5/67:0.0484983987)86.6/28:0.0506193106,((((WP_073350970.1-Aeromonas-hydrophila:0.1731438756,WP_190284026.1-Aeromonas-hydrophila:0.0524755640)78.9/100:0.0341120772,WP_241344946.1-Aeromonas-sp.-MR16:0.1094027111)89.4/100:0.0858614654,WP_244081328.1-Aeromonas-caviae:0.3503655476)68.3/19:0.0669578405,WP_076573677.1-Aeromonas-sp.-RU39B:0.4497339443)55.1/13:0.0266924828)84.9/73:0.0719989469,(((((WP_050718733.1-Aeromonas-tecta:0.1945697890,WP_100859741.1-Aeromonas-lusitana:0.0501484220)77.3/100:0.0123542244,WP_042654929.1-Aeromonas-encheleia:0.0714523289)96.6/100:0.0776034440,WP_077098334.1-Aeromonas-hydrophila:0.0699981840)76.3/100:0.0185620133,WP_043167431.1-Aeromonas-hydrophila:0.1472523506)65.7/98:0.0119713001,WP_252420256.1-Aeromonas-taiwanensis:0.1112943103)99.1/100:0.1680992933)90.9/45:0.0743948272)72.9/61:0.0189136235,WP_223952962.1-Aeromonas-caviae:0.2645442797)87.3/78:0.0410520221,(WP_108652490.1-Dongshaea-marina:0.3594367724,((WP_026470910.1-Alkanindiges-illinoisensis:0.2985377990,(WP_248147450.1-Moraxella-osloensis:0.4787565796,WP_092690500.1-Acinetobacter-kyonggiensis:0.3001306333)91.3/100:0.1620378797)100/100:0.5863501491,WP_134696535.1-Aeromonas-taiwanensis:0.8135080323)92.8/96:0.2304414950)48.2/64:0.0517591206)82.8/91:0.0294529243,(WP_208005626.1-Oceanisphaera-pacifica:0.1297904416,(WP_087034853.1-Oceanisphaera-profunda:0.0862189300,WP_087034873.1-Oceanisphaera-profunda:0.1713335146)91.7/100:0.0808441773)99.7/100:0.1892809717)62.5/77:0.0276739478,((WP_097405671.1-Kosakonia:0.0584279230,WP_128482629.1-Kosakonia-cowanii:0.2367970240)91.9/79:0.0937254151,WP_161518249.1-Kosakonia-sp.-MH5:0.0000020117)100/100:0.3380202455)86.6/66:0.0542905065,((WP_234664799.1-Aeromonas-allosaccharophila:0.2639274126,(((((((WP_245131329.1-Aeromonas-veronii:0.1914851145,((WP_101347551.1-Aeromonas-sobria:0.1008562423,WP_201955411.1-Aeromonas-veronii:0.0770414904)96.5/100:0.1241478877,(WP_136501435.1-Aeromonas-veronii:0.2518614277,(WP_268029297.1-Aeromonas-hydrophila:0.1608013804,WP_201942265.1-Aeromonas-veronii:0.1975460379)95.6/100:0.1465036127)81.7/100:0.0601206230)85.5/97:0.0622921197)81.5/12:0.0384848761,WP_210550341.1-Aeromonas-sp.-SrichE-2G:0.4508381533)0/5:0.0000021953,(WP_225159845.1-Aeromonas-caviae:0.1466004522,(WP_163135231.1-Aeromonas-rivipollensis:0.0442612790,WP_241279715.1-Aeromonas-sp.-MR19:0.0422947000)100/100:0.3850630732)84.2/96:0.0632159366)40.4/12:0.0200227544,(((WP_213320225.1-Aeromonas-caviae:0.0815057666,WP_203764723.1-Aeromonas-caviae:0.1274710470)98.3/100:0.1908597434,(((WP_257692500.1-Aeromonas-caviae:0.2476140280,WP_113719982.1-Aeromonas-salmonicida:0.2042783225)69.1/93:0.0353920314,WP_069554789.1-Aeromonas-sp.-DNP9:0.2220866899)87.5/90:0.0782175854,((WP_227095090.1-Aeromonas-veronii:0.1314142428,WP_152109808.1-Plesiomonas-shigelloides:0.0804388280)89.7/100:0.0666295882,WP_168366848.1-Dickeya-zeae:0.2889971368)89.8/100:0.0643557804)79.5/96:0.0439396710)79.2/53:0.0426996849,((WP_050666161.1-Aeromonas-schubertii:0.0854072301,WP_059168611.1-Aeromonas-salmonicida:0.1700474692)69.6/100:0.0290029029,WP_210600017.1-Aeromonas-sp.-MrichA-1:0.0727761639)87.5/100:0.0534992295)88.2/99:0.0546387374)83.6/89:0.0392644767,(((((WP_148612677.1-Aeromonas-sobria:0.0513688081,((WP_201355655.1-Aeromonas-hydrophila:0.0617412331,WP_201951905.1-Aeromonas-veronii:0.0433682032)71.9/98:0.0087609977,WP_171268810.1-Aeromonas-media:0.0360880958)4.1/84:0.0205852391)100/100:0.2055678482,(WP_042063659.1-Aeromonas-allosaccharophila:0.0473293195,WP_256684552.1-Aeromonas-salmonicida:0.0443968631)28.4/82:0.0193236274)23/98:0.0115003642,(((WP_212539243.1-Aeromonas-salmonicida:0.0439751661,((WP_187590672.1-Aeromonas-hydrophila:0.1698763281,WP_017782281.1-Aeromonas-dhakensis:0.0361139885)60.4/100:0.0194012917,WP_060586794.1-Aeromonas-schubertii:0.0693460129)99.4/100:0.1198530324)88.5/100:0.0363324206,WP_201904077.1-Aeromonas-veronii:0.0769768835)75.8/100:0.0261061419,WP_206874356.1-Aeromonas-veronii:0.0131428065)75.2/98:0.0138862683)96.4/99:0.1346128351,WP_042649704.1-Aeromonas-media:0.1258251453)14.2/78:0.0572976842,WP_139406182.1-Aeromonas-veronii:0.3924930611)77.5/87:0.0542093149)81.1/89:0.0341029987,(WP_253626250.1-Aeromonas-hydrophila:0.1407073849,((WP_224211331.1-Aeromonas-enteropelogenes:0.2848931987,WP_082041448.1-Aeromonas-eucrenophila:0.3435143752)81.7/97:0.0564008690,(((WP_268010485.1-Aeromonas-dhakensis:0.0979970391,WP_098985072.1-Aeromonas-sp.-CA23:0.0655243843)97.8/100:0.1530016388,(WP_181268965.1-Aeromonas-veronii:0.0529086758,(WP_129146711.1-Aeromonas-caviae:0.0337421521,(WP_081843667.1-Aeromonas-taiwanensis:0.0258007339,WP_139744464.1-Aeromonas-media:0.0563261656)84.1/100:0.0289516424)76.4/100:0.0206310471)88/100:0.1182389953)95.5/100:0.1997835554,WP_200561028.1-unclassified-Erwinia:0.5100832059)90.2/98:0.1457258606)88.8/76:0.0638712932)92.9/75:0.0806393011)82.5/51:0.0440093492,((((WP_268421127.1-Aeromonas-media:0.0942429931,WP_111910157.1-Aeromonas:0.0297854850)80.4/90:0.0263932935,WP_128343177.1-Aeromonas-caviae:0.0613864714)95.6/100:0.0777830542,WP_268037199.1-Aeromonas-dhakensis:0.1520659059)29.3/92:0.0192444593,(WP_139733908.1-Aeromonas-veronii:0.0171865511,((WP_234579942.1-Aeromonas-salmonicida:0.0993117164,(WP_234676856.1-Aeromonas-rivipollensis:0.0220662582,WP_269382839.1-Aeromonas-hydrophila:0.1145953870)32/100:0.0105720702)36.1/81:0.0099756238,WP_222134508.1-Vibrio-cholerae:0.2185655726)75.8/81:0.0197571640)98.8/99:0.1183811261)86.4/97:0.0374973471)76.8/65:0.0321946974)72.8/58:0.0251882773,((((WP_216990617.1-Aeromonas-sp.-FDAARGOS-1409:0.2147793730,(((((WP_059168599.1-Aeromonas-salmonicida:0.0807593985,(WP_033115905.1-Aeromonas-lacus:0.0737764728,((WP_223920301.1-Aeromonas-caviae:0.0500334619,WP_226058924.1-Shewanella-sp.-SR1:0.1702200210)75/98:0.0207483017,(WP_058394234.1-Aeromonas-salmonicida:0.0433103761,WP_257690725.1-Aeromonas-veronii:0.0670330589)80.5/97:0.0285766367)45.7/97:0.0158837092)94.8/100:0.0739333378)95.9/100:0.0999860604,(((WP_223945983.1-Aeromonas-caviae:0.1636958959,(((WP_265458754.1-Aeromonas-salmonicida:0.1096348619,WP_125597700.1-Aeromonas-salmonicida:0.0300225918)70.3/99:0.0053145329,(WP_180911080.1-Aeromonas-caviae:0.0533722596,WP_264818242.1-Aeromonas-caviae:0.0353547504)91.9/100:0.0269856265)0/97:0.0000023797,WP_163128050.1-Aeromonas-allosaccharophila:0.1203379403)75.2/97:0.0112701067)74.6/52:0.0078593531,(WP_040098646.1-Aeromonas:0.1281992989,((WP_263309349.1-Aeromonas-caviae:0.0549273212,(WP_041217344.1-Aeromonas-hydrophila:0.0553324326,WP_255912544.1-Aeromonas-sp.-JL9:0.0443311571)75/83:0.0092519993)73.5/83:0.0084441535,WP_257686696.1-Aeromonas-veronii:0.0560933315)72.4/100:0.0143427653)75.9/100:0.0186306796)69.5/51:0.0088493177,((WP_139750984.1-Aeromonas-media:0.0399165884,WP_224430864.1-Aeromonas-rivuli:0.0334376130)75.4/99:0.0264510779,WP_041996113.1-Aeromonas-bivalvium:0.0725981345)98.1/99:0.0744504666)85/40:0.0363414326)73.3/50:0.0203852349,WP_265776268.1-Aeromonas-veronii:0.1736455114)91.2/96:0.0927274128,WP_236563281.1-Aeromonas-veronii:0.2355319997)56.8/34:0.0097199965,(((((WP_201942221.1-Aeromonas-veronii:0.2529063120,WP_159143398.1-Aeromonas-veronii:0.0178088806)81.1/40:0.0259119026,(WP_268854930.1-Aeromonas-enteropelogenes:0.1513778457,(((WP_203131090.1-Aeromonas-jandaei:0.0731307256,WP_111901368.1-Aeromonas-veronii:0.0607058235)38.9/17:0.0106443330,WP_139692403.1-Aeromonas-sobria:0.0405697000)72.3/9:0.0104835607,WP_042657830.1-Aeromonas:0.0737555764)76.5/88:0.0087969750)18.5/64:0.0149232188)0/39:0.0000026656,WP_223380193.1-Aeromonas-enteropelogenes:0.0456460722)99/99:0.1260870086,(WP_236321701.1-Aeromonas-veronii:0.0994750847,WP_257119864.1-Aeromonas-hydrophila:0.0264447964)84.2/99:0.0471534887)86.7/74:0.0427358643,(WP_257707542.1-Aeromonas-caviae:0.0782823700,WP_182951616.1-Aeromonas-sp.-WP2-W18-CRE-05:0.0832185732)99.3/100:0.1749299765)0/82:0.0000020725)71.1/35:0.0268358883)86.3/97:0.0372986279,((WP_064338398.1-Aeromonas-veronii:0.2478452814,(((WP_042864139.1-Aeromonas-caviae:0.0504612588,WP_225159881.1-Aeromonas-caviae:0.0551843562)87.8/100:0.0565460399,(WP_244056308.1-Aeromonas-caviae:0.1817491663,WP_202046159.1-Aeromonas-media:0.1661511626)92.2/93:0.1568417393)92.1/90:0.0934413577,(WP_045790994.1-Aeromonas-hydrophila:0.2070306460,((((((WP_247163170.1-Escherichia-coli:0.0242144595,WP_038644959.1-Pantoea:0.1102202616)98.9/100:0.1282280599,(WP_252422582.1-Aeromonas-taiwanensis:0.0219923394,WP_263719954.1-Aeromonas-caviae:0.1216215928)78.1/90:0.0603727208)87.2/98:0.0525370203,WP_033131297.1-Aeromonas-aquatica:0.0280259315)95.8/88:0.0603420967,WP_241344952.1-Aeromonas-sp.-MR16:0.0174893459)87.9/87:0.0317928643,WP_160838676.1-Aeromonas-caviae:0.1070073261)48.4/32:0.0200257224,((WP_049048056.1-Aeromonas-hydrophila:0.1943349680,WP_167569591.1-Aeromonas-veronii:0.1503138194)58.6/63:0.0568368726,(WP_116547128.1-Plesiomonas-shigelloides:0.1250627092,(((WP_139496062.1-Escherichia-coli:0.0880465739,WP_215269322.1-Aeromonas-dhakensis:0.0525133545)31/88:0.0087562698,WP_201996661.1-Aeromonas-veronii:0.0503460479)74.3/67:0.0133191084,WP_235463836.1-Aeromonas-australiensis:0.0206234829)64.1/40:0.0246092195)74.3/48:0.0201522802)57.2/87:0.0386315000)99.3/100:0.1571443425)75.5/8:0.0175308675)81.1/81:0.0334016731)39.2/88:0.0247551653,(WP_223932214.1-Aeromonas-caviae:0.0166261046,WP_161506954.1-Aeromonas-media:0.1814340198)99.9/100:0.2266762350)94.6/95:0.0733272894)60.3/43:0.0124201239,(WP_100646402.1-Aeromonas-veronii:0.0733732893,((WP_252601295.1-Aeromonas-dhakensis:0.0272494409,WP_139432874.1-Aeromonas-dhakensis:0.1993608705)91.4/100:0.0452577610,WP_106553824.1-Aeromonas-hydrophila:0.0829133133)90.6/100:0.0428811790)99.8/100:0.2208101072)88.8/81:0.0643912792,WP_265468300.1-Aeromonas-salmonicida:0.1974113660)78.9/78:0.0344586149)93.7/68:0.0741650929)82.9/25:0.0336538044,((WP_047682696.1-Xenorhabdus:0.0759353248,WP_099124588.1-Xenorhabdus-stockiae:0.1208121587)98.4/100:0.2143879468,(WP_074024273.1-Xenorhabdus-eapokensis:0.1068767034,WP_217470569.1-Photorhabdus-akhurstii:0.0935984289)88.8/100:0.0755824821)97.4/100:0.1301290092)82.4/37:0.0325479992,(((WP_151054911.1-Cellvibrio-sp.-KY-YJ-3:0.3250422782,WP_078044787.1-Cellvibrio-sp.-PSBB023:0.2995873076)72/95:0.0422276338,(WP_012489013.1-Cellvibrio-japonicus:0.2158498929,WP_087465811.1-Cellvibrio-sp.-PSBB006:0.2702801500)84.4/95:0.0498177157)99.2/100:0.1949371182,((WP_048221111.1-Citrobacter:0.2031357177,WP_263807164.1-Salmonella-enterica:0.2561634986)97.5/100:0.2769884079,WP_206164314.1-Citrobacter:0.3091514388)88.4/53:0.0900944138)28.6/21:0.0207127590)92.2/31:0.0944472419,(((((((WP_172912938.1-Pantoea-stewartii:0.1864238660,WP_145394131.1-Pantoea-sp.-SJZ147:0.1859040958)99/100:0.3474376429,(WP_086868392.1-Kosakonia-pseudosacchari:0.7666899186,WP_061356323.1-Escherichia-coli:1.0816013966)83.1/94:0.2648308282)74.7/91:0.1678174962,WP_039304623.1-Cedecea-neteri:0.5145896156)87.9/82:0.1380722191,(WP_043158915.1-Aeromonas-hydrophila:0.6420703435,WP_153752706.1-unclassified-Citrobacter:0.8649930790)38.1/89:0.1119843663)41.4/61:0.0545684908,((((((WP_071788415.1-Franconibacter-pulveris:0.6890790329,(WP_073514120.1-Vibrio-parahaemolyticus:0.0958009017,WP_032481483.1-Vibrio:0.0128133528)100/100:1.4559339840)20.6/33:0.1020236878,(WP_020232007.1-Escherichia-coli:0.3507782459,(WP_228338545.1-Enterobacter-hormaechei:0.1816692995,(WP_109845273.1-Enterobacter:0.0772546803,WP_150146763.1-Enterobacter-roggenkampii:0.0507777992)87.9/100:0.1221761084)98.8/100:0.3142877553)47.6/45:0.0620406361)96.9/83:0.2323779128,((((((((((((((((((((((((((((((((((((WP_051234278.1-Halomonas-halocynthiae:1.0071051911,WP_110648656.1-Salinicola-peritrichatus:0.4541532134)91.6/100:0.2357690654,WP_254581577.1-Marinobacterium-sedimentorum:0.5519513408)96.7/97:0.2819014148,(((((WP_081193716.1-Halomonas-sp.-BC1:0.4153279305,WP_035566812.1-Halomonas:0.2401450927)56.4/65:0.0852564197,(((((((WP_224850395.1-Marinobacter-nauticus:0.2201842920,WP_110630630.1-Salinicola-salarius:0.1114045125)88.5/95:0.0567509625,(WP_193085952.1-Halomonas-sp.-FME65:0.1627117102,((WP_159342670.1-Halomonas-meridiana:0.4246760436,WP_253554573.1-Halomonas-sp.-DN3:0.2555807844)67.1/100:0.0326741820,WP_102589198.1-Halomonas-urumqiensis:0.1380703870)91.9/100:0.0573326622)38.8/64:0.0203911749)87.3/47:0.0502655291,((WP_089675781.1-Halomonas-aquamarina:0.0676078790,WP_062362595.1-Halomonas-axialensis:0.0268500356)37.1/92:0.0182098718,(WP_253557393.1-unclassified-Halomonas:0.0632712508,WP_089676106.1-Halomonas-aquamarina:0.0361516907)0/63:0.0000026991)76.4/84:0.0114622410)81.1/53:0.0255257787,WP_027961801.1-Halomonas-halodenitrificans:0.0357985878)86.4/53:0.0276575948,WP_079551663.1-Halomonas-subglaciescola:0.0898117060)91.9/55:0.0462044520,WP_195687443.1-Halomonas-sp.-KAO:0.0295135590)86.5/72:0.0451278280,(WP_193343673.1-Halomonas-elongata:0.0839377858,WP_064701650.1-Halomonas-caseinilytica:0.0171945702)99.2/100:0.1459985358)93.8/98:0.0951264559)85.7/63:0.0500457961,((((WP_158774340.1-Cobetia-sp.-L2A1:0.0883798894,(WP_144727127.1-Cobetia-crustatorum:0.0428388050,WP_088744253.1-Cobetia-sp.-QF-1:0.0804184994)70.9/73:0.0082017352)87.8/78:0.0368767520,WP_158774127.1-Cobetia-sp.-L2A1:0.0372639477)79.4/82:0.0174558966,(WP_077373671.1-Cobetia:0.0704915201,((WP_213115235.1-Cobetia-sp.-MC34:0.0516419054,WP_200017180.1-Cobetia-sp.-cqz5-12:0.0774046233)91.7/100:0.0750939571,WP_225346840.1-Cobetia-amphilecti:0.0536609318)83.4/100:0.0378639770)92.6/100:0.0411072681)98.7/100:0.1910389164,(WP_017842145.1-Methylotuvimicrobium-buryatense:0.5224886731,(WP_251978361.1-unclassified-Salinicola:0.1004335103,WP_189443129.1-Salinicola-rhizosphaerae:0.1492788483)96.2/100:0.1608644769)80.2/56:0.0715495872)92.8/59:0.1114371601)89.5/48:0.0470515754,(WP_149282777.1-Halomonas-sp.-Y2R2:0.0862227201,WP_073436013.1-Halomonas-cupida:0.1350343456)89.9/60:0.0705242097)96.2/55:0.0799228574,(WP_248234415.1-Marinobacter-bryozoorum:0.1234100551,(WP_228223924.1-Marinobacter-nauticus:0.0860693522,WP_098418752.1-Marinobacter-sp.-LV10MA510-1:0.1772476311)87.6/99:0.0918511644)96.1/99:0.1032417116)76.3/42:0.0137793337)37.7/12:0.0233048563,WP_021818206.1-Halomonas-huangheensis:0.1681497082)92.6/57:0.0672899465,((((((WP_066801876.1-Moraxella-oblonga:0.1751782358,(WP_197128702.1-Psychrobacter-sp.-NZS113:0.2580138933,WP_201616712.1-Psychrobacter-immobilis:0.2277581160)98.5/100:0.2904941024)95.3/100:0.2741875077,WP_036548095.1-Nitrincola-lacisaponensis:0.5673528125)82.2/92:0.0700599226,((WP_041157945.1-Halomonas-sp.-KHS3:0.0861121712,(WP_120386672.1-Halomonas-alkaliphila:0.1753318347,(WP_141441391.1-Halomonas-titanicae:0.1033013351,WP_226251132.1-unclassified-Halomonas:0.0874244369)54/97:0.0158251598)83.1/96:0.0816386370)98.9/100:0.1426184085,WP_228219791.1-Halomonas-meridiana:0.1298381955)91.4/100:0.0841230652)93/90:0.0533254345,(((((WP_030071911.1-Halomonas-alkaliantarctica:0.0624725210,WP_095603285.1-Halomonas-humidisoli:0.0323674495)63.4/72:0.0243291315,WP_253485683.1-Halomonas-sp.-XH26:0.0384099921)29.5/71:0.0156896158,WP_038485954.1-Halomonas-campaniensis:0.0262974119)98.6/100:0.0995680440,WP_162218282.1-Halomonas-alkaliphila:0.0390268440)93.8/100:0.0513687558,(WP_078087027.1-unclassified-Halomonas:0.0319571851,WP_096655487.1-Halomonas-nigrificans:0.0597610052)73.4/54:0.0102446720)99.1/100:0.1396926809)90.5/84:0.0694504875,(WP_127061154.1-Halomonas-nanhaiensis:0.2652999842,(WP_027958907.1-unclassified-Halomonas:0.0495031523,WP_064236373.1-unclassified-Halomonas:0.0712138076)98.7/100:0.1510358357)0/5:0.0000027866)73.8/6:0.0195881863,WP_096281135.1-Halomonas-sp.-JB37:0.3053378105)89.7/48:0.0509518853)86.5/47:0.0895880431,WP_179929642.1-Halomonas-salicampi:0.6177684015)94.6/100:0.1222612403,WP_138986566.1-Neptunomonas-concharum:0.4284754459)86.1/94:0.0707528459,(WP_015904383.1-Desulforapulum-autotrophicum:0.4148734078,WP_111960646.1-Desulfobacter-hydrogenophilus:0.4439104643)96.1/97:0.2959137535)82.9/93:0.0480548601,((((((WP_230291920.1-Croceicoccus-sp.-Ery5:0.7947536811,(WP_116367876.1-Parahaliea-mediterranea:0.0929037466,WP_148063811.1-Parahaliea-aestuarii:0.1448714512)95.7/100:0.2945600500)93.4/89:0.2886415789,WP_129645432.1-Steroidobacter-agariperforans:0.4784238604)94.4/89:0.2995765579,((((((((WP_036226274.1-Mesoaciditoga-lauensis:0.3291262752,((WP_077198159.1-Thermosipho-affectus:0.1205623845,(WP_012057290.1-Thermosipho-melanesiensis:0.2521334013,(WP_127352844.1-Caldicellulosiruptor-changbaiensis:0.0981654914,WP_014043315.1-Caldicellulosiruptor-acetigenus:0.0276839986)100/100:0.6525247352)57/26:0.0563235443)97.4/100:0.1466734345,WP_047753768.1-Kosmotoga-pacifica:0.2385501360)78.5/33:0.0264830197)15.8/33:0.0505350507,((((((WP_117708548.1-Fusobacterium-mortiferum:0.0000021696,WP_204716698.1-Fusobacterium-mortiferum:0.1004963955)80.4/100:0.0269938047,WP_115270465.1-Fusobacterium-necrogenes:0.0963329209)54.7/100:0.0271135553,WP_204716813.1-Fusobacterium-mortiferum:0.6608422887)83.3/87:0.0862066069,(((WP_130889691.1-Fusobacterium-ulcerans:0.0581749929,((WP_005950923.1-Fusobacterium-varium:0.0458166730,WP_070580525.1-Fusobacterium:0.0812083296)41.8/99:0.0162310973,WP_227372109.1-Fusobacterium-ulcerans:0.0491615247)84.2/99:0.0221172265)79.8/98:0.0157771096,(WP_241635364.1-Fusobacterium-gastrosuis:0.1665988719,(WP_074518138.1-Fusobacterium-necrophorum:0.1156716765,(WP_211495349.1-Fusobacterium-necrophorum:0.1856461194,WP_032890075.1-Fusobacterium-nucleatum:1.0794974520)84.4/96:0.1166439841)95.1/96:0.1070031230)98/96:0.1050476985)85.9/95:0.0398405244,WP_261793436.1-Fusobacterium-varium:0.3894976500)88/89:0.0757415461)96.6/100:0.2170392083,((WP_147004474.1-Leptotrichia-wadei:0.0350002269,WP_018499460.1-Leptotrichia-wadei:0.0466690210)94.9/100:0.1049085371,(WP_068155536.1-Leptotrichia-sp.-oral-taxon-847:0.0929916312,(WP_099992503.1-Fusobacterium-pseudoperiodonticum:0.1759911857,(WP_211617428.1-Leptotrichia-sp.-oral-taxon-218:0.0194115369,WP_094080762.1-Leptotrichia-sp.-oral-taxon-498:0.0656087095)95.3/100:0.0726667175)73.3/100:0.0562722486)93.6/100:0.0899285220)99.4/100:0.3338559215)89.4/93:0.1250506804,((WP_013388736.1-Ilyobacter-polytropus:0.7480293774,WP_173194267.1-Paenibacillus-frigoriresistens:0.3978883387)49.5/63:0.0893068450,WP_012548442.1-Dictyoglomus-thermophilum:0.6182267388)26.2/20:0.0146952513)98.2/100:0.1914475686)97.7/100:0.1564117751,(WP_099829233.1-Mesotoga-sp.-Brook.08.105.5.1:0.2311063414,WP_099830786.1-Mesotoga-sp.-Brook.08.105.5.1:0.2884406179)99.3/100:0.2380655837)49.5/79:0.0995103393,(((((((((((((((((((((WP_054742706.1-Cellulosilyticum-ruminicola:0.1247947915,(WP_177671917.1-Zhenhengia-yiwuensis:0.0625570007,WP_053982981.1-Niameybacter-massiliensis:0.1522084571)98.5/100:0.1724311071)59.3/99:0.0126863800,WP_069999280.1-Cellulosilyticum-sp.-I15G10I2:0.3957013825)99.1/100:0.2174370050,WP_152890015.1-Clostridium-tarantellae:0.1897663594)89.9/35:0.0690214455,(WP_195972826.1-Clostridium-thermobutyricum:0.1423172388,(WP_209795868.1-Clostridium-moniliforme:0.0893117455,WP_194191032.1-Clostridium-chrysemydis:0.1689281827)93.5/100:0.0747203645)35.4/84:0.0266926044)37.6/15:0.0141865357,WP_040194942.1-Clostridium-culturomicium:0.0975783657)79.8/15:0.0215554944,((WP_073337580.1-Clostridium-grantii:0.2092788235,WP_072832886.1-Clostridium-collagenovorans:0.0400049721)82.9/81:0.0271665286,WP_072892386.1-Clostridium-fallax:0.1689603404)86.8/34:0.0272585972)96.6/51:0.0552033023,(((((((((WP_094369216.1-Romboutsia-weinsteinii:0.2497031855,WP_021359979.1-Clostridioides-difficile:0.5142105453)12.7/96:0.0509230146,(WP_074918830.1-Terrisporobacter-glycolicus:0.0985243714,WP_195929149.1-Clostridium-sp.-1001270J_160509_D11:0.0517015083)84.6/100:0.0408892598)89.9/96:0.0539412082,((WP_092726456.1-Romboutsia-lituseburensis:0.2616088650,WP_258242885.1-Romboutsia-lituseburensis:0.0468657210)98.4/100:0.1540363075,(WP_057560648.1-Paeniclostridium-sordellii:0.1266799913,((WP_057580750.1-Paeniclostridium-sordellii:0.1672870559,WP_057538715.1-Paeniclostridium-sordellii:0.0625068415)78.8/99:0.0180113833,WP_195252007.1-Romboutsia-sp.-1001713B170207_170306_H8:0.0779451224)84.9/99:0.0208165957)55.5/96:0.0196655531)85/100:0.0265899871)85.8/93:0.0374710249,((((((WP_066874757.1-Clostridium-mediterraneense:0.2876780192,WP_216313757.1-Clostridium-gasigenes:0.0616816076)86.4/99:0.0495831798,(WP_029452249.1-Clostridium-algidicarnis:0.1433279165,WP_216183125.1-Clostridium-gasigenes:0.0410551482)64.8/100:0.0509321865)93.6/91:0.0749983109,WP_066677671.1-Clostridium-septicum:0.0689412734)73.9/82:0.0119355993,(WP_095406848.1-Romboutsia-maritimum:0.2049793557,WP_238903698.1-Clostridium-sp.-YIM-B02506:0.1014800840)78.9/20:0.0179997219)33.4/12:0.0199355389,WP_197096717.1-Paeniclostridium-sordellii:0.1248224547)81.5/17:0.0244252818,(((WP_195959433.1-Clostridium-tertium:0.1032953188,WP_195429094.1-Clostridium-sp.-D46t1_190503_E9:0.0263640677)70.3/100:0.0259275337,WP_047001949.1-Clostridium-sp.-C8:0.0893396973)81.7/92:0.0214753171,(WP_066890539.1-Clostridium-nigeriense:0.0684169840,WP_016207773.1-Clostridium-sartagoforme:0.0527569972)42.5/19:0.0095635092)94.7/99:0.0461999234)81.1/79:0.0303308022)85/83:0.0352291014,WP_163195303.1-Clostridium-thermarum:0.1858642830)74.2/47:0.0116383947,(WP_055666468.1-Desnuesiella-massiliensis:0.2947118830,(((((WP_055265352.1-Clostridium:0.0459899051,(WP_186859336.1-Clostridium-sp.-NSJ-6:0.0816666428,(WP_055265304.1-Clostridium-disporicum:0.0873078361,(WP_154821366.1-Clostridium-saudiense:0.0154610373,WP_195245504.1-Clostridium-celatum:0.0923032804)79.6/100:0.0397412489)65.2/99:0.0442539553)85.4/99:0.0724913356)91.1/100:0.1036289411,WP_186834444.1-Eubacteriales:0.1771344970)95.1/100:0.0858065619,((((WP_235068992.1-Turicibacter-sp.-TJ11:0.0863309681,WP_212726030.1-Turicibacter-bilis:0.0548125878)40.3/29:0.0078501611,WP_195933776.1-Turicibacter-sanguinis:0.0372722152)83/29:0.0219663214,WP_155223812.1-Turicibacter-sanguinis:0.0378704160)80.8/62:0.0336021016,WP_161831114.1-Turicibacter-sp.-TS3:0.0468603617)98.9/100:0.1527399330)57.4/75:0.0318247332,WP_198610231.1-Clostridium-perfringens:0.0975789240)92.4/99:0.0498992327,WP_055258173.1-Sarcina-ventriculi:0.1128285581)87.8/95:0.0482365086)69.1/30:0.0309642147)68.3/45:0.0199805516,WP_133628539.1-Fonticella-tunisiensis:0.1049643307)0/45:0.0000022117,WP_055069574.1-Clostridium-massiliamazoniense:0.2610886046)92.5/55:0.0400636289,(((WP_072725426.1-Tepidibacter-thalassicus:0.2095604652,WP_095404508.1-Romboutsia-maritimum:0.5666262235)95.1/79:0.1382665609,WP_072889493.1-Tepidibacter-formicigenes:0.0545158735)93.2/78:0.0812724531,WP_066825124.1-Clostridium-tepidiprofundi:0.1606927723)88.4/78:0.0473877202)74.8/46:0.0107662196)76.6/32:0.0097223564,((((((((((((((WP_169105633.1-Clostridium-sp.-SM-530-WT-3G:0.0669627074,WP_077866663.1-Clostridium-saccharobutylicum:0.3389109333)10.4/91:0.0274902507,WP_027636628.1-Clostridium-butyricum:0.1324073473)91.8/91:0.0521520871,(((((WP_009169246.1-unclassified-Clostridium:0.1688698059,WP_077843022.1-Clostridium-beijerinckii:0.2152086023)77.2/100:0.0309495733,WP_041896941.1-Clostridium-beijerinckii:0.2928116821)92.3/100:0.0986067608,WP_069680896.1-Clostridium-taeniosporum:0.3300899206)49.9/99:0.0520908845,WP_160359894.1-Clostridium-chromiireducens:0.4283070439)92/91:0.0703706199,(WP_265859166.1-Clostridium-botulinum:0.1573832721,((WP_154531076.1-Inconstantimicrobium-porci:0.2052171310,WP_009169009.1-Clostridium-sp.-DL-VIII:0.0254628528)87.7/99:0.0702230593,WP_077856313.1-Clostridium-beijerinckii:0.0000026823)94.4/99:0.1052733988)52.7/26:0.0440322200)62.8/21:0.0151273131)74.7/36:0.0162258115,WP_022750218.1-Clostridium-saccharobutylicum:0.0775799383)10.2/32:0.0083942504,WP_226126241.1-Clostridium-algoriphilum:0.1907156926)86.3/96:0.0215059197,(WP_185652834.1-Clostridium-sp.-DJ247:0.0321871932,WP_185652641.1-Clostridium-sp.-DJ247:0.0983554622)89.6/100:0.0331052242)84.9/96:0.0190634590,WP_077848125.1-Clostridium-puniceum:0.2187284174)72.9/24:0.0089649030,(WP_015391310.1-Clostridium-saccharoperbutylacetonicum:0.0828642518,(WP_077846315.1-Clostridium-puniceum:0.0714623484,WP_238915641.1-Clostridium-sp.-YIM-B02555:0.0884587936)83.8/100:0.0273373345)96.3/100:0.0496592132)58.5/22:0.0041828099,((WP_261829154.1-Clostridium-sp.-TW13:0.1175917323,WP_224033465.1-Clostridium-gelidum:0.1583344172)0/71:0.0000023041,(WP_077855754.1-Clostridium-beijerinckii:0.0580358706,WP_041894754.1-Clostridium-beijerinckii:0.0388244007)99.4/100:0.1110326423)87.3/77:0.0227852342)99.6/98:0.1449259420,WP_216390068.1-Clostridium-sp.-MSJ-8:0.3427556597)45/100:0.0307438331,WP_090010800.1-Clostridium-sp.-DSM-8431:0.1416102575)84.4/99:0.0610895077,(WP_106059304.1-Clostridium-vincentii:0.1524480958,WP_143316579.1-Clostridium:0.0305322360)79.4/100:0.0254241075)92.5/98:0.0588538003,WP_055184115.1-Clostridium-paraputrificum:0.0883196509)91.8/85:0.0517555670,WP_261829490.1-Clostridium-sp.-TW13:0.3007829506)74.8/36:0.0115127575)79.5/54:0.0142072961,WP_072990516.1-Clostridium-cavendishii:0.3827685975)0/41:0.0039954690,WP_148566871.1-Acetobacterium-paludosum:0.6498608372)83.2/73:0.0496919042,(((WP_216288229.1-Clostridium-psychrophilum:0.2058316826,WP_216137569.1-Clostridium-estertheticum:0.0458654133)5.1/99:0.0083588290,WP_220704225.1-Clostridium-estertheticum:0.0557741539)93.7/100:0.0666625104,WP_220321605.1-Clostridium-tagluense:0.0979836720)96.8/100:0.1130428364)92.7/85:0.0915981118,(WP_142413383.1-Hathewaya-massiliensis:0.3272915267,WP_035130905.1-Clostridium-sulfidigenes:0.3684293830)99.4/98:0.2400619894)86.9/81:0.0571317926,((WP_079421587.1-Clostridium-oryzae:0.5768652933,WP_226126924.1-Clostridium-algoriphilum:0.5276058077)65.4/94:0.0787904302,(((WP_227847393.1-Clostridioides-sp.-ZZV15-6598:0.1598328135,WP_227846117.1-Clostridioides-sp.-ES-S-0145-01:0.0412199693)95.7/100:0.0623688132,(WP_227842647.1-unclassified-Clostridioides:0.0420149062,WP_003420982.1-Clostridioides-difficile:0.1311848794)83.8/100:0.0314025889)84.3/100:0.0389460995,WP_038810653.1-Clostridioides-difficile:0.0443651665)100/100:0.4152290641)84.2/94:0.0788083342)86.8/80:0.0578698094,WP_099190526.1-Tepidibacter-mesophilus:0.2909472711)85.4/81:0.0545485211,(WP_112093115.1-Methanothermobacter-tenebrarum:0.4338573241,WP_013413362.1-Methanothermus-fervidus:0.3750257517)97.3/98:0.3094377735)75/60:0.0296526849,((WP_111570784.1-Halanaerobium-saccharolyticum:0.1113263552,(WP_089717478.1-Halanaerobium-congolense:0.0783693976,WP_089862761.1-Halanaerobium-salsuginis:0.0390458019)60.9/97:0.0573012627)99.9/100:0.4756079680,WP_072903743.1-Hathewaya-proteolytica:0.6985645427)48.5/56:0.0358849722)91.6/65:0.0648358713,(((WP_088538685.1-Methanobrevibacter-sp.-87.7:0.1012766963,WP_016359256.1-Methanobrevibacter-sp.-AbM4:0.1579016080)94.9/100:0.1435362042,WP_067148181.1-Methanobrevibacter-olleyae:0.1795878478)100/100:0.2646249866,(((WP_013644504.1-Methanobacterium-lacus:0.2970486490,WP_054834988.1-Methanobrevibacter-arboriphilus:0.3126053512)89.5/100:0.0837379121,(WP_011406138.1-Methanosphaera:0.2526890167,(WP_112123902.1-Methanosphaera-sp.-BMS:0.1554419325,(WP_069592847.1-Methanosphaera-sp.-WGK6:0.0453925756,WP_095608065.1-Methanosphaera-cuniculi:0.1643232792)97.2/100:0.1325934250)12.6/91:0.0437588502)98.3/100:0.1482148714)53.9/74:0.0068736831,(WP_248612240.1-Methanobacterium-alcaliphilum:0.2076416101,((WP_071907175.1-Methanobacterium-congolense:0.0383406171,WP_209584978.1-Methanobacterium-aggregans:0.0548814195)99.5/100:0.1860089089,(WP_069583666.1-Methanobacterium:0.2591036455,WP_048190917.1-Methanobacterium-sp.-SMA-27:0.2203789093)84.5/96:0.0956086016)58.5/96:0.0206511204)83.1/93:0.0379120359)85.7/86:0.0395237688)97.9/70:0.0961504105)73.6/49:0.0122096581,((((((WP_068555806.1-Thermotalea-metallivorans:0.2023497828,WP_073236503.1-Desulforamulus-putei:0.1603187696)74.4/100:0.0424041799,WP_095133872.1-Anaeromicrobium-sediminis:0.3017878941)86.3/57:0.0450698262,WP_035164300.1-Caloranaerobacter-azorensis:0.1068327613)63.7/58:0.0330497649,WP_090442301.1-Natronincola-peptidivorans:0.2629298838)94.4/96:0.0787224411,(((((WP_252130283.1-Clostridium-sp.-SYSU_GA19001:0.5127397676,WP_202749105.1-Clostridium-sp.-YIM-B02515:0.1641818888)98.2/98:0.1860438734,((((WP_104410444.1-Clostridium-algidicarnis:0.0657047327,WP_185162668.1-Clostridium-algidicarnis:0.0983827111)100/100:0.2866518111,((WP_090038599.1-Clostridium-frigidicarnis:0.2258354631,WP_209459870.1-Youngiibacter-multivorans:0.3117783952)87.3/100:0.0871038174,(WP_164955553.1-Clostridium-estertheticum:0.2695493176,(WP_125002390.1-Clostridium-tagluense:0.2621409184,(WP_226136064.1-Clostridium-estertheticum:0.0366878232,WP_216146875.1-Clostridium-frigoris:0.0999763669)97.9/100:0.1594548569)76.5/99:0.0547427280)95.2/99:0.1362459324)42.1/98:0.0373712354)47/82:0.0592064506,(WP_234123145.1-Clostridium-hydrogenum:0.2986476149,((WP_084114600.1-Clostridium-acidisoli:0.1476303784,(WP_015614219.1-Clostridium-pasteurianum:0.0296677338,(WP_010235680.1-Clostridium-arbusti:0.0174290807,WP_003443690.1-Clostridium-pasteurianum:0.0732444964)91.7/100:0.0410763663)94.2/100:0.0519899809)47.8/100:0.0195571116,WP_026884435.1-Clostridium-akagii:0.1412773223)89.2/100:0.1205230927)52.3/84:0.0551449823)29.6/92:0.0312065011,WP_216462460.1-Clostridium-bornimense:0.2282606747)88.9/99:0.0642821233)85.9/85:0.0486644325,(((WP_165412483.1-Clostridium-estertheticum:0.0949787311,WP_216302578.1-unclassified-Clostridium:0.0140112352)99.9/100:0.3036920134,WP_216457288.1-Clostridium-simiarum:0.2051445515)63.9/80:0.0451296623,(WP_204598070.1-Clostridium-pascui:0.2128023126,WP_027632043.1-Clostridium-hydrogeniformans:0.2134764827)41.5/91:0.0133800283)95.5/100:0.1016880222)16.3/29:0.0426704537,((((WP_066398085.1-Cytobacillus-eiseniae:0.4682603452,(WP_076307499.1-Paenibacillus-odorifer:0.0885453326,((WP_042140168.1-Paenibacillus-sp.-FSL-P4-0081:0.0182628269,WP_036721118.1-Paenibacillus-sp.-FSL-R7-277:0.1025661464)84.5/100:0.0507331313,WP_074087222.1-Paenibacillus-sp.-P32E:0.0506167509)89.4/100:0.1289265260)96.4/100:0.1550765179)35.6/76:0.0587273447,((WP_056034132.1-Paenibacillus-sp.-Leaf72:0.1328937770,WP_197936429.1-Paenibacillus-sp.-GSMTC-2017:0.2348022616)89.1/100:0.1009843857,WP_090798944.1-Paenibacillus-sp.-GP183:0.3749439796)61.6/81:0.0787720582)24.7/31:0.0582528273,WP_262684732.1-Paenibacillus-sp.-WQ-127069:0.2621380817)85.3/99:0.1203891186,WP_036935152.1-Pseudobacteroides-cellulosolvens:0.2600665625)90.8/97:0.1399435917)82.3/94:0.0579933049,((WP_069974893.1-Geosporobacter-ferrireducens:0.6421679070,(WP_223038819.1-Crassaminicella-profunda:0.1347848625,(WP_148809648.1-Crassaminicella-thermophila:0.0867891829,WP_218283043.1-Crassaminicella-sp.-143-21:0.4694792124)88.6/100:0.0557418899)86.7/100:0.0936382962)90.4/100:0.1363344585,WP_250858284.1-Oceanirhabdus-seepicola:0.3365897305)99/100:0.2506480328)70.4/11:0.0116143021)86.3/25:0.0591470502,((WP_009529486.1-Peptoanaerobacter-stomatis:0.2126112457,WP_068913445.1-Criibacterium-bergeronii:0.3213098584)99.7/100:0.3641472765,WP_116670296.1-Methanobrevibacter-woesei:0.4455185911)89.8/62:0.1029038302)0/22:0.0000022810)85.5/22:0.0598345251,(WP_038266426.1-Peptoclostridium-litorale:0.6874113645,WP_160686059.1-Clostridium-sp.-C2-6-12:0.3805128061)90.1/91:0.1125004989)80.1/17:0.0458292006,(((((((WP_058965344.1-Fournierella-massiliensis:0.2640191210,WP_185752312.1-Butyrivibrio-sp.-AE2032:0.5662822751)87.9/86:0.1284287394,WP_155202396.1-Ruthenibacterium-lactatiformans:0.2982609199)92/100:0.1584354703,(WP_186886939.1-Anaerofilum-hominis:0.2313829748,WP_102049807.1-Pygmaiobacter-massiliensis:0.3833311586)88.3/84:0.1117878618)98.4/100:0.3090099477,((((((WP_072827923.1-Fibrobacter-sp.-UWB12:0.0481859365,WP_088630837.1-unclassified-Fibrobacter:0.0436569191)39.1/65:0.0409692261,WP_073304542.1-Fibrobacter:0.2767277088)99.8/99:0.2277195932,(WP_073155408.1-Fibrobacter:0.0817284809,WP_073188830.1-Fibrobacter-sp.-UWP2:0.0193726497)48/84:0.0310912574)39/33:0.0224119855,WP_097034884.1-Fibrobacter-sp.-UWT3:0.0358510466)81.3/29:0.0236537747,WP_073056819.1-unclassified-Fibrobacter:0.0284922998)99.7/100:0.3054769104,WP_256135338.1-Bittarella-massiliensis:0.4126814957)86.7/72:0.1733050723)81.3/52:0.0772873319,(WP_266162911.1-Caproicibacterium-sp.-BJN0003:0.2930068790,WP_220321633.1-Clostridium-tagluense:0.5222322813)91.8/55:0.1377115225)89.2/57:0.0775029782,(WP_105204772.1-Neobittarella-massiliensis:0.2108203447,WP_022475313.1-Huintestinicola-butyrica:0.4504349822)97.6/93:0.1825165913)54.8/45:0.0292583671,((((WP_091684654.1-Anaerocolumna-aminovalerica:0.3326996520,(WP_033168390.1-Clostridium-sp.-KNHs205:0.1305997225,((WP_073588199.1-Anaerocolumna-xylanovorans:0.0487709586,WP_185256524.1-Anaerocolumna-chitinilytica:0.2151421220)72/100:0.0669820322,WP_073279502.1-Anaerocolumna-jejuensis:0.1736526549)51.6/100:0.0721794017)99.4/100:0.2641416459)95.9/100:0.1909223973,((((WP_253721803.1-Treponema-sp.-OMZ-787:0.9090353727,WP_005539496.1-Johnsonella-ignava:0.2165172196)89.2/94:0.1255631864,(WP_060930762.1-Lachnoanaerobaculum-saburreum:0.3351198908,(((((((((((((((WP_024837362.1-Lacrimispora:0.1474885472,WP_077612318.1-Clostridium-sp.-Marseille-P2415:0.0496646101)0/99:0.0012618148,(WP_024293365.1-Lachnospiraceae:0.0518856923,WP_092242699.1-Lacrimispora:0.0376850072)94.6/100:0.0548901392)88.4/99:0.0357771529,((WP_124065139.1-Clostridium-sp.-E02:0.1393625166,WP_117415169.1-Clostridium-indicum:0.0639150914)77.6/95:0.0152017103,WP_143321893.1-Clostridium-sp.-HBUAS56010:0.0455909922)77.1/95:0.0165267555)17.5/66:0.0161472084,WP_138313639.1-Hungatella-hathewayi:0.0422401270)83.6/94:0.0211564500,WP_186424215.1-Lacrimispora-celerecrescens:0.2289928278)91.5/94:0.0480290506,((WP_227052204.1-Anaerostipes-caccae:0.0467760376,WP_137330070.1-Anaerostipes-rhamnosivorans:0.1267032780)99/100:0.1315810518,WP_006781439.1-Hungatella-hathewayi:0.1219311992)89.9/100:0.0422228452)71.4/89:0.0109086948,(WP_231062061.1-Lacrimispora-sp.-NSJ-141:0.1443974716,WP_249304640.1-Qiania-dongpingensis:0.1020580133)95.9/99:0.1009458270)85.3/88:0.0311219627,(WP_215718428.1-Anaerostipes-hadrus:0.0188106031,WP_075760385.1-Anaerostipes-sp.-494a:0.0795774056)99.5/100:0.1189235251)68.5/81:0.0096991258,WP_118541387.1-unclassified-Clostridium:0.1154478459)0/74:0.0000020769,(WP_024736308.1-Enterocloster-asparagiformis:0.0562151253,WP_066545911.1-Clostridium-sp.-AT4:0.0527555573)90.6/90:0.0312469390)60.8/63:0.0084190069,WP_215700061.1-Clostridium-sp.-MCC353:0.0863901215)85.7/95:0.0202572016,(((WP_045093538.1-Clostridium-sp.-FS41:0.0796735034,WP_025485467.1-Lachnospiraceae:0.0846387300)69.2/100:0.0259064919,((WP_154471253.1-Clostridium-porci:0.1938141759,WP_178313416.1-Lacrimispora-sp.-210928-DFI.3.58:0.0075640865)30.4/99:0.0243474303,WP_002577448.1-Enterocloster:0.1058677968)83.9/99:0.0266394194)67.5/94:0.0175783594,((((WP_087223935.1-Lachnoclostridium-sp.-An14:0.0588741397,WP_101876396.1-Lachnoclostridium-edouardi:0.0747240831)90/100:0.0412121983,WP_118243934.1-Clostridium-sp.-AM58-1XD:0.0900775865)71.8/95:0.0117498456,WP_118742831.1-Clostridium-sp.-AM28-20LB:0.0443595074)88.4/95:0.0180580710,WP_008399284.1-unclassified-Clostridium:0.0263124829)90.9/97:0.0290288183)70.6/80:0.0178471908)89.2/99:0.0366392157,(WP_003497689.1-_Clostridium:0.0730166462,WP_125142063.1-Clostridium-transplantifaecale:0.0544439894)84.7/100:0.0356708569)73.5/94:0.0354026537,WP_118555425.1-Lachnotalea-sp.-AF33-28:0.3608835537)32.4/85:0.0231940088,WP_251497048.1-Otoolea-sp.-DSM-111138:0.1877979381)88.2/100:0.1031818509)93.5/100:0.1292351751)89.6/93:0.0941125424,((((((((((((((((WP_035764434.1-Butyrivibrio-sp.-NC2002:0.1677021526,((WP_026526812.1-Butyrivibrio-sp.-VCD2006:0.2324400842,WP_026491547.1-unclassified-Butyrivibrio:0.0233447356)65.4/100:0.0543811026,WP_044913869.1-Butyrivibrio-sp.-WCE2006:0.0345467197)93.1/100:0.0815587839)74.1/11:0.0648655485,WP_091822189.1-Butyrivibrio-sp.-ob235:0.0658804068)100/100:0.2710903610,WP_092060841.1-Butyrivibrio-sp.-INlla14:0.0188380726)85.3/11:0.0556715488,WP_013279951.1-Butyrivibrio-proteoclasticus:0.0449994863)83.4/11:0.0205869445,(((((WP_022768176.1-unclassified-Butyrivibrio:0.0339430355,WP_022766545.1-Butyrivibrio-sp.-XPD2006:0.0810417224)99.3/100:0.1486521170,WP_034446055.1-Butyrivibrio-sp.-AE2032:0.0977818428)89.1/97:0.0689120889,WP_024867291.1-Butyrivibrio-sp.-FCS014:0.0420979080)86.7/58:0.0438721038,(WP_071175335.1-Butyrivibrio:0.1082955497,WP_026513668.1-Butyrivibrio-sp.-LB2008:0.0660181586)83.3/58:0.0611516026)70.5/97:0.0341930617,WP_026663187.1-Butyrivibrio-proteoclasticus:0.1060323537)86.6/79:0.0251416720)0/11:0.0000020709,WP_026652320.1-Butyrivibrio-proteoclasticus:0.0651270785)99.8/100:0.2381970767,((WP_019160740.1-Blautia:0.0924844078,WP_173738571.1-Blautia-schinkii:0.0306466840)85.1/99:0.0329136509,(WP_158421413.1-Blautia:0.0455848953,(WP_148445023.1-Eubacteriales:0.0421161469,WP_187566223.1-Blautia-faecis:0.0733139047)93/100:0.0525648453)78.2/88:0.0306460121)97.1/99:0.1099007549)95.9/86:0.1035407999,WP_178047055.1-Fusicatenibacter-sp.-CLA-AA-H277:0.1585351883)80.8/77:0.0294488336,(WP_230400100.1-Novisyntrophococcus-fermenticellae:0.3086696268,WP_109624131.1-Murimonas-intestini:0.1173858000)37.6/29:0.0433525430)0/16:0.0000029163,((WP_173751851.1-Blautia-caecimuris:0.0676383143,WP_117997894.1-Blautia-obeum:0.0189464596)99/100:0.1058655983,(((WP_158367551.1-Lachnospiraceae:0.0495189093,WP_235247881.1-Faecalicatena-contorta:0.1151970994)67.2/99:0.0584891025,(WP_251390531.1-Mediterraneibacter-sp.-DSM-111629:0.0633506538,WP_073962323.1-_Ruminococcus:0.0390246918)79.7/99:0.0379342825)78.7/86:0.0241475954,WP_070089756.1-Merdimonas-faecis:0.0499909249)86.3/88:0.0308240520)55/81:0.0389685420)84.2/85:0.0411463495,WP_132379639.1-Muricomes-intestini:0.1514239331)95.6/93:0.0658445199,WP_087156947.1-Lachnoclostridium-sp.-An181:0.0740947203)79.1/91:0.0248314157,(WP_073994331.1-Parasporobacterium-paucivorans:0.2172978502,(WP_099466994.1-Konateibacter-massiliensis:0.1129894896,WP_151145296.1-Candidatus-Galacturonibacter-soehngenii:0.0774541138)95.7/100:0.1346822370)70.3/63:0.1011351475)46.4/52:0.0301039454,WP_115483253.1-Anaerosacchariphilus-polymeriproducens:0.1568135279)40.2/79:0.0546547560,WP_158351687.1-Diplocloster-modestus:0.3723363942)55.1/95:0.0686200203,WP_158346166.1-Lachnospiraceae:0.4116510643)96.3/95:0.1718659365)67.5/89:0.0266700565,WP_226914794.1-Intestinibacter-bartlettii:0.4708358285)90/92:0.0740935444)67.1/68:0.0463853194,((((WP_073273890.1-Anaerocolumna-jejuensis:0.2206192974,WP_033167869.1-Clostridium-sp.-KNHs205:0.2178821438)44.3/100:0.0763446783,((WP_010268185.1-Paenibacillus-senegalensis:0.2686176991,WP_155699987.1-Paenibacillus-psychroresistens:0.0925400234)15.7/60:0.0323129367,(WP_202751253.1-Clostridium-sp.-YIM-B02515:0.0842774497,WP_090040610.1-Clostridium-frigidicarnis:0.0446257844)44.2/85:0.0125489917)90.6/86:0.1040405420)98.9/100:0.2575835532,((WP_249305773.1-Jutongia-hominis:0.3367152662,(WP_055944671.1-unclassified-Clostridium:0.0557931492,WP_216564272.1-Eubacterium-sp.-MSJ-13:0.0379243167)78.1/100:0.0474202441)41/76:0.0321016291,WP_022463459.1-Jutongia-huaianensis:0.0986925143)99.8/100:0.3460261414)78.3/91:0.0719450857,WP_058352784.1-Acetivibrio-ethanolgignens:0.4980023554)95.6/97:0.1377645396)92.4/94:0.1429332924,((WP_213236459.1-Fusibacter-paucivorans:1.0531234500,(((((((((((WP_160564586.1-Apibacter-sp.-B2912:0.1704598339,(WP_055424523.1-Apibacter-mensalis:0.1434623641,(WP_160568957.1-unclassified-Apibacter:0.0169893909,WP_166004639.1-Apibacter-sp.-B2966:0.0733845158)98.3/100:0.1354995118)80.1/98:0.0988815137)97.4/100:0.1814480229,(WP_105245553.1-Apibacter-adventoris:0.3156979281,WP_146261858.1-Apibacter-muscae:0.1803423525)38.1/86:0.0822697618)99.9/100:0.4295173211,(WP_084015419.1-Moheibacter-sediminis:0.5713650363,WP_182042771.1-Moheibacter-lacus:0.4272443450)94.8/98:0.2115951679)33.4/90:0.0461963250,(WP_068596492.1-Vaginella-massiliensis:0.2320431112,((WP_230477342.1-Faecalibacter-bovis:0.1643857424,WP_125349156.1-Empedobacter-falsenii:0.1480669229)69.7/92:0.0481357893,(WP_125113420.1-Algoriella-xinjiangensis:0.1246142758,WP_092908829.1-Algoriella-xinjiangensis:0.0524195806)93.4/100:0.0779351432)96.8/92:0.1961741502)97.9/100:0.2567270383)94.9/95:0.2582137662,(WP_014783622.1-Aequorivita-sublithincola:0.5401825085,((((WP_230143398.1-unclassified-Pedobacter:0.1616603844,WP_215166022.1-Pedobacter-sp.-ISL-64:0.1393013199)92.3/99:0.1231344854,WP_133260761.1-Pedobacter-changchengzhani:0.1932879596)87.5/89:0.0757812546,((WP_056872627.1-Pedobacter-sp.-Leaf194:0.1360335679,((WP_123203876.1-Pedobacter-jejuensis:0.1102720200,WP_188416210.1-Pedobacter-mendelii:0.0454655198)11.5/92:0.0204306097,WP_056087060.1-Pedobacter-sp.-Leaf176:0.0793251246)75/100:0.0210780281)97.2/97:0.0866037543,WP_109929147.1-Pedobacter-paludis:0.0659389014)81.3/87:0.0416876382)80.1/75:0.0578233562,WP_199156598.1-Pedobacter-sp.-ASV2:0.1349471418)97.8/94:0.3352888855)98.5/99:0.3926886948)93.9/99:0.2074949240,(((WP_107809536.1-Elizabethkingia-sp.-YR214:0.2286814363,WP_219971314.1-Chryseobacterium-sp.-LJ756:0.2177167586)86.3/79:0.0880903700,WP_076395614.1-Chryseobacterium:0.0728891777)83.7/79:0.1118217696,WP_088467981.1-Chryseobacterium-sp.-VAUSW3:0.1646217117)99.8/100:0.3684059384)0/25:0.0002828117,WP_111126400.1-Mesonia-sp.-K7:1.2807827174)97.6/95:0.2629994940,((((WP_205722928.1-Fulvivirga-lutea:0.8220885361,(((((((((((WP_053470876.1-unclassified-Flavobacterium:0.2435912747,WP_095952323.1-Flavobacterium-sp.-ACN6:0.0540990354)42.5/97:0.0115087118,WP_056200692.1-Flavobacterium-sp.-Root186:0.0507073201)84.5/97:0.0177017603,WP_115889823.1-Flavobacterium-cutihirudinis:0.0726677033)45.1/98:0.0086680900,WP_198857739.1-Flavobacterium-sp.-IB48:0.0544870004)80.3/100:0.0135844510,(WP_269237486.1-Flavobacterium-sp.-IMCC34775:0.1195004748,WP_163393939.1-Flavobacterium-limi:0.0505789316)0/100:0.0031320050)34.9/95:0.0167801001,WP_264205577.1-Flavobacterium-sp.-LS1R49:0.0366742952)89.4/100:0.0250636700,((WP_173968237.1-Flavobacterium-collinsii:0.1052628427,WP_264525312.1-Flavobacterium-sp.-N502536:0.0686163202)92.5/100:0.0488038119,WP_070907856.1-Flavobacterium-tructae:0.1258294765)75.8/87:0.0114076904)76.3/87:0.0145152910,WP_091091738.1-Flavobacterium-gillisiae:0.1468403455)98.5/98:0.1008517258,WP_248907802.1-Flavobacterium-sp.-K5-23:0.1842534549)83.7/96:0.0472863176,(((WP_116761981.1-Flavobacterium-laiguense:0.0644993200,WP_143377960.1-Flavobacterium-sp.-ZT3R18:0.0619432914)97.4/100:0.0956973241,((WP_144071503.1-Flavobacterium-franklandianum:0.0896277338,WP_133277350.1-Flavobacterium-nackdongense:0.1249226357)41.8/97:0.0407891515,WP_194311471.1-Flavobacterium-soyangense:0.1761122780)77.9/95:0.0458902163)58.1/92:0.0415662607,WP_103806521.1-Flavobacterium-alvei:0.1327629682)84/95:0.0509132553)98.7/98:0.2835249118,(WP_140000461.1-Flavobacterium-microcysteis:0.3644796537,(((WP_194640005.1-Flavobacterium-sp.-HJJ:0.1517712348,WP_073246022.1-Flavobacterium-flevense:0.0875383131)69.5/100:0.0302643163,WP_100843428.1-Flavobacterium-sp.-5:0.1617556270)66.1/91:0.0805837897,WP_140487657.1-Flavobacterium-sp.-GSA192:0.0638990734)99.6/100:0.2928424721)94/100:0.1883918329)93.9/99:0.2246987214)91.1/90:0.1851478063,WP_057952567.1-Salinivirga-cyanobacteriivorans:0.9396184052)42.2/72:0.0839813628,(WP_116872776.1-Terrimonas-sp.:0.7350091755,(WP_004155558.1-Microscilla-marina:0.5897142487,WP_101358295.1-Raineya-orbicola:0.3504033721)98.9/100:0.4918736418)85.6/89:0.1606219666)90.2/92:0.1392688174,(((((WP_233378072.1-Maribellus-sp.-CM-23:0.0956484539,WP_247978154.1-Maribellus-sp.-YY47:0.0990585869)94.8/100:0.1715188555,(WP_167609329.1-Maribellus-sediminis:0.2567740517,(WP_045033348.1-Draconibacterium-sediminis:0.0391336709,WP_163345719.1-Draconibacterium-halophilum:0.1176547493)99.2/100:0.2155864049)92.8/100:0.1642544756)99.9/100:0.4819821375,(((WP_106543043.1-Prolixibacter-denitrificans:0.1685489563,WP_025865917.1-Prolixibacter-bellariivorans:0.0305786223)74.1/78:0.0090358769,WP_153632016.1-Prolixibacter-sp.-SD074:0.0745048522)99.9/99:0.3966531788,((WP_159517929.1-Sunxiuqinia-indica:0.1678149311,WP_053182762.1-Sunxiuqinia-dokdonensis:0.1512923696)88.2/100:0.0770093077,WP_133466511.1-Sunxiuqinia-elliptica:0.2472464434)97.8/100:0.2114154214)39.4/79:0.1132364291)94.5/88:0.1317483126,(WP_068701711.1-Paludibacter-jiangxiensis:0.6925337631,(((WP_101263396.1-Labilibaculum-filiforme:0.0670573196,WP_245870435.1-Labilibaculum-manganireducens:0.0641241641)97.4/99:0.1373081906,(WP_152734748.1-Marinifilum-sp.-N1E240:0.1118326020,(WP_171595603.1-Marinifilum-caeruleilacunae:0.1056043750,(WP_268189532.1-Marinifilum-sp.-D737:0.0863473456,WP_054720444.1-Marinifilum-fragile:0.0403516450)86.5/100:0.0317041504)84.8/100:0.0374265901)91.6/100:0.0836065198)38.3/93:0.0599501433,WP_127342456.1-Ancylomarina-longa:0.2113640681)98.6/100:0.3324157792)73.6/11:0.0292768994)72.5/11:0.0525555319,((WP_011500232.1-Methanococcoides-burtonii:0.3761593755,WP_135611779.1-Methanococcoides-sp.-AM1:0.2818947661)53.5/97:0.0892419624,(((((WP_128503182.1-Methanosarcina-sp.-MSH10X1:0.2544857454,WP_011033115.1-Methanosarcina-mazei:0.0978419428)28.5/67:0.0487537802,WP_048134921.1-Methanosarcina-sp.-1.H.T.1A.1:0.0518948400)93.1/93:0.1041430347,WP_048167026.1-Methanosarcina:0.1427542975)31.8/71:0.0532783492,WP_048177163.1-Methanosarcina-sp.-MTP4:0.1315249865)69.1/76:0.0893544054,(WP_048137589.1-Methanosarcina-horonobensis:0.2270715720,((WP_048184631.1-Methanosarcina-siciliae:0.0845808705,WP_011023680.1-Methanosarcina-acetivorans:0.0660671978)37.3/89:0.0142133999,WP_048173663.1-Methanosarcina-siciliae:0.0737948862)96.1/100:0.1409688768)72.4/92:0.0919211834)79.8/92:0.1440741821)99.8/99:0.4237817828)91.1/81:0.1375985628)57.7/43:0.1363111620)95.1/100:0.2319529156,(((WP_085659180.1-Prosthecochloris-sp.-HL-130-GSB:0.6747417422,WP_038563105.1-Draconibacterium-orientale:0.4428289041)96.9/100:0.2663646942,(WP_040596643.1-Prevotella-stercorea:0.1424772119,WP_207153370.1-Prevotella-herbatica:0.1365843046)99.8/100:0.4221287550)28.8/84:0.0762865733,(WP_223927321.1-Prevotella-lacticifex:0.3176326404,WP_009010825.1-Prevotella-sp.-C561:0.2775496319)100/100:0.7364159343)87.1/100:0.2266062437)80.1/99:0.1597891572,((((((WP_013633636.1-Pseudopedobacter-saltans:0.2171924603,WP_154287284.1-Pedobacter-puniceum:0.2433534277)88.7/100:0.1221329364,WP_160842504.1-Pedobacter-sp.-HMF7647:0.0991359729)100/100:0.3405745681,(WP_258137696.1-Mucilaginibacter-sp.-BJC16-A38:0.2975936006,(WP_259068220.1-Mucilaginibacter-sp.-X4EP1:0.0958759287,WP_144912900.1-Mucilaginibacter-frigoritolerans:0.1104050821)89.8/100:0.0985102468)96.1/98:0.1759101099)16.9/97:0.0171027389,WP_074492796.1-Mucilaginibacter-polytrichastri:0.2835628751)99.5/100:0.4323003527,WP_163286637.1-Cryomorpha-ignava:0.8673106713)37.4/99:0.1599238810,WP_130092062.1-Brumimicrobium-glaciale:0.8659447487)90.3/100:0.1535965483)85.3/99:0.2039111270,WP_078711972.1-Anaerorhabdus-furcosa:0.9414994300)84.6/79:0.1895706285)73.7/78:0.0774378681,WP_138788932.1-Paenibacillus-thermoaerophilus:0.8086773205)93.5/98:0.1598369406)88.6/91:0.0893901575)97.8/63:0.1527014964)74.3/17:0.0293620626,WP_119970050.1-Clostridium-fermenticellae:0.4153571758)94.4/90:0.1732058071)98.2/99:0.2974351012,((((((WP_018154793.1-Methanothermococcus-thermolithotrophicus:0.2250188643,WP_048201153.1-Methanocaldococcus-bathoardescens:0.0884050427)64.4/100:0.0796231317,(((WP_011869065.1-Methanococcus-maripaludis:0.1283524901,WP_013998786.1-Methanococcus-maripaludis:0.0868909543)79/100:0.0602645130,WP_181493664.1-Methanococcus-maripaludis:0.1066177489)29.9/89:0.0149783687,WP_181504225.1-Methanococcus-maripaludis:0.0848580093)100/100:0.5125318493)23.6/78:0.0507360991,(WP_052562942.1-Candidatus-Brocadia-sinica:0.6325419327,WP_173298646.1-Thermanaeromonas-sp.-C210:0.4594063180)84.5/90:0.1097329462)80.5/89:0.0713999720,((WP_119360783.1-Meiothermus-luteus:0.2625686280,WP_027877894.1-Meiothermus-cerbereus:0.0782224412)84.8/97:0.0958814968,WP_208013758.1-Meiothermus-sp.-CFH-77666:0.0866634355)100/100:1.1312456110)89.1/88:0.1384685943,((((((WP_010868297.1-Pyrococcus-abyssi:0.1309177761,((WP_048103006.1-Aciduliprofundum-boonei:0.4522956748,WP_148120035.1-Methanofervidicoccus-sp.-A16:0.2253239516)86/97:0.0417342589,WP_088862340.1-Thermococcus-celer:0.1260713146)74.6/48:0.0144397185)86.6/46:0.0255464529,((((((((((((WP_237702072.1-Thermococcus-sp.-AM4:0.0514929613,WP_223211916.1-Thermococcus-henrietii:0.0797683320)96.7/100:0.0823696231,WP_014733609.1-Pyrococcus-sp.-ST04:0.1802222659)88.2/99:0.0301974796,(WP_139679991.1-Thermococcus-indicus:0.0461034199,WP_148882943.1-Thermococcus-aciditolerans:0.0559033983)87.8/99:0.0322037223)18.6/20:0.0196639489,WP_058938040.1-Thermococcus-celericrescens:0.0409674719)69.7/79:0.0104017881,(WP_088865724.1-Thermococcus-barossii:0.0894842895,(WP_167711671.1-unclassified-Thermococcus:0.0360263365,WP_232460978.1-Thermococcus-radiotolerans:0.0703401054)73.9/30:0.0076508915)75.4/19:0.0110220121)81.8/92:0.0250026338,WP_258085050.1-Thermococcus-sp.-813A4:0.0432383965)84.1/81:0.0195627467,WP_088181368.1-Thermococcus-sp.-5-4:0.0888789369)92.8/100:0.0459170856,(WP_152879125.1-Thermococcus-sp.-101-C5:0.0707132001,WP_015850310.1-Thermococcus-sibiricus:0.0760175329)72.2/100:0.0102174559)85.6/100:0.0215641501,WP_209475361.1-Thermococcus-stetteri:0.1357646318)45.2/45:0.0106691812,WP_253304977.1-Thermococcus-aggregans:0.0312991277)87.9/69:0.0364781125,WP_068664737.1-Thermococcus-piezophilus:0.0806626555)95.2/75:0.0631135507,((WP_167913028.1-Thermococcus-sp.-21S7:0.1135148873,WP_048150691.1-Palaeococcus-ferrophilus:0.0368212181)97.6/100:0.0721531594,((((WP_088855609.1-Thermococcus-siculi:0.0782761287,WP_087036111.1-Thermococcus-litoralis:0.0502080087)82.3/99:0.0263746296,WP_167888701.1-Thermococcus-sp.-MV5:0.1089281623)90.9/87:0.0319259036,(WP_013468354.1-Thermococcus-barophilus:0.0773626222,WP_013906489.1-Pyrococcus-yayanosii:0.0303523987)84.1/92:0.0187910090)87/67:0.0212585944,WP_048053121.1-Pyrococcus-horikoshii:0.0390262213)81.2/56:0.0194852773)74.9/46:0.0135559963)44.1/38:0.0114071098)99.1/100:0.1059149910,WP_007044577.1-Methanotorris-formicicus:0.0901726255)66.3/75:0.0461225976,WP_013798345.1-Methanotorris-igneus:0.0535634654)55.8/91:0.1094515424,WP_013866394.1-Methanothermococcus-okinawensis:0.2064572132)99.8/100:0.4399220028,((((WP_245533116.1-Coprothermobacter-platensis:0.8763617963,WP_011996389.1-Coxiella-burnetii:1.2505635840)27.7/98:0.0885481430,WP_022856039.1-Thermodesulfobacterium:0.3097464985)87/98:0.1339681775,WP_150071182.1-Thermodesulfobacterium-sp.-TA1:0.3597861026)92.1/98:0.2034496533,(((WP_029522925.1-Persephonella-sp.-KM09-Lau-8:0.1674593766,(WP_028950938.1-Sulfurihydrogenibium-subterraneum:0.2489266269,((WP_132526207.1-Phorcysia-thermohydrogeniphila:0.1263932366,WP_013638952.1-Desulfurobacterium-thermolithotrophum:0.0792475832)85.6/100:0.0479467412,WP_089322203.1-Desulfurobacterium-atlanticum:0.2291539490)95/100:0.1112659877)79.6/99:0.0794321977)92.8/96:0.0892533855,WP_096999769.1-Persephonella-hydrogeniphila:0.1678035013)42.8/92:0.0444627185,WP_200674641.1-Persephonella-atlantica:0.0683538920)94.7/100:0.1453311886)97.3/89:0.1512405752)91.7/99:0.1291614055)98.6/100:0.2921779882,WP_236098952.1-Dethiosulfovibrio:1.0549855005)29.1/15:0.0597088285)97.3/96:0.3421494649,(WP_207681674.1-Desulfonema-magnum:0.8646662916,WP_230840800.1-Gloeobacter-morelensis:0.5267130873)78.3/75:0.1653423008)86.7/12:0.0743319317,(((((GJNO01008138.1-Laxus-oneistus-TSA:1.6507792794,(((((((((((VDM95900.1-Thelazia-callipaeda:0.3119090286,(((VDO27950.1-Onchocerca-flexuosa:0.2242933673,VBB33646.1-Acanthocheilonema-viteae:0.0962874912)15.7/21:0.0258530188,((XP_003135693.1-Loa-loa:0.1102151431,VDK73328.1-Litomosoides-sigmodontis:0.0674712033)38.4/66:0.0143958789,CAG9536712.1-Cercopithifilaria-johnstoni:0.0393662524)83.2/65:0.0269143739)10.6/20:0.0148966715,VDO31788.1-Brugia-timori:0.1641352242)96.9/98:0.2187766345)100/100:0.4616674927,VDN56932.1-Dracunculus-medinensis:0.5045194324)74.9/77:0.0874149080,KHN87800.1-Toxocara-canis:0.2824752330)53.3/61:0.1036896202,VDD97390.1-Enterobius-vermicularis:0.5517100587)94.8/100:0.2035105484,(KAI1709308.1-Ditylenchus-destructor:0.1690504740,KAI1730870.1-Ditylenchus-destructor:0.0376035336)99.8/100:0.2914364086)17.9/49:0.0711411996,(GGJS01006545.1-Plectus-murrayi-TSA:0.6584294055,TKR72632.1-Steinernema-carpocapsae:0.3224179695)27.5/43:0.0379858213)44.5/57:0.1183720692,KAE9552274.1-Halicephalobus-sp.-NKZ332:0.4383678547)69/97:0.1343830308,((CAD5228290.1-Bursaphelenchus-okinawaensis:0.0610592956,CAD5235627.1-Bursaphelenchus-xylophilus:0.0446259342)38.5/94:0.0819192493,((KAI6206096.1-Aphelenchoides-besseyi:0.1954376922,KAI6191010.1-Aphelenchoides-bicaudatus:0.1155325429)83.3/100:0.0738483382,KAI6234710.1-Aphelenchoides-fujianensis:0.1344057156)93.1/100:0.1317418730)96.9/100:0.1578508655)77.2/75:0.0712849705,KAH7695359.1-Aphelenchus-avenae:0.3766737272)99.1/98:0.2352186585,((CAI4229259.1-Auanema-sp.-JU1783:0.6465422238,(CAD6189320.1-Caenorhabditis-auriculariae:0.5200530845,(CAB3400093.1-Caenorhabditis-bovis:0.3042574833,((NP_509046.1-Caenorhabditis-elegans:0.0407013651,XP_003117635.1-Caenorhabditis-remanei:0.0402376241)32/99:0.0600744522,CAI5454992.1-Caenorhabditis-angaria:0.0395951203)82.3/100:0.0634928507)94/100:0.1561275005)98/99:0.2424326328)76.1/98:0.0524686565,PAV65180.1-Diploscapter-pachys:0.3355302451)85.9/98:0.0599666196)74.2/97:0.0374795442,((KHJ99538.1-Oesophagostomum-dentatum:0.0748055514,EPB80542.1-Ancylostoma-ceylanicum:0.0586690475)96.6/99:0.1194078876,((VDO65958.1-Haemonchus-placei:0.1862341911,VDL70484.1-Nippostrongylus-brasiliensis:0.1125841801)86.1/100:0.0338198585,VDO95824.1-Heligmosomoides-polygyrus:0.1007419991)98.7/100:0.1788872663)89.4/99:0.1703033503)98.1/96:0.5043341697)99.2/100:1.0726470190,(WP_042281872.1-Candidatus-Protochlamydia-sp.-R18:0.1851424164,WP_068471358.1-Parachlamydia-sp.-C2:0.2245671509)99.7/100:0.9645236777)91.8/97:0.3890208115,(WP_008983868.1-Alishewanella-agri:1.3574273966,WP_265845290.1-Plesiomonas-shigelloides:1.2615176453)78.8/98:0.1694439122)88.9/96:0.2404393281,((WP_239493201.1-Luteitalea-sp.-TBR-22:0.5718532741,WP_058867107.1-Chloracidobacterium-thermophilum:0.6371233056)92.1/91:0.3088579222,(WP_161826415.1-Petrachloros-mirabilis:0.6331588631,WP_099324654.1-Candidatus-Kuenenia-stuttgartiensis:0.9647277100)77.7/48:0.1258703631)31.1/25:0.1106401820)94.9/47:0.1772923886,((((WP_255135177.1-Gracilimonas-sp.-CAU-1638:0.4912740604,WP_020401568.1-Gracilimonas-tropica:0.4677759238)99.8/100:0.8238085503,WP_198243301.1-methane-oxidizing-endosymbiont-of-Gigantopelta-aegis:0.5474632265)93/75:0.2360690039,((((((((((((((((WP_176021856.1-Stenotrophomonas-sp.-NA06056:0.0613842756,WP_049440708.1-Stenotrophomonas-maltophilia:0.0672602702)99.1/100:0.2496036800,WP_132999175.1-Luteimonas-arsenica:0.2791350627)11.3/87:0.1126119110,WP_078758981.1-Lysobacter-spongiicola:0.4786649330)95.7/93:0.1745614092,WP_131151171.1-Dyella:0.2082535656)94.8/66:0.0906972969,(WP_209622196.1-Frateuria-flava:0.1015562827,WP_091339503.1-Frateuria-terrea:0.0663746690)93.6/100:0.0532724018)62.2/65:0.0287835718,WP_201315287.1-Dyella-sp.-EPa41:0.1996667471)53.8/32:0.0079584922,WP_253565187.1-Dyella-sp.-Sa:0.1321705226)78/44:0.0221688194,(WP_063674027.1-Dyella-thiooxydans:0.1877726982,(WP_188800319.1-Dyella-caseinilytica:0.2334914339,WP_192554051.1-Dyella-acidiphila:0.1720110739)93.3/100:0.1186484172)88.6/55:0.0647271492)90.4/57:0.0469937866,((((((WP_214557828.1-unclassified-Rhodanobacter:0.0493138710,WP_068095172.1-unclassified-Rhodanobacter:0.0403917309)58/100:0.0276529966,(WP_008437080.1-Rhodanobacter-thiooxydans:0.0602985030,(WP_063107523.1-Rhodanobacter-thiooxydans:0.0436517805,WP_211005527.1-Rhodanobacter-sp.-B2A1Ga4:0.0495802459)50.7/100:0.0092483678)54.4/100:0.0241771238)96/100:0.0721958888,WP_072762453.1-Rhodanobacter-sp.-OK091:0.0147859224)72.9/43:0.0113516190,((WP_077439480.1-Rhodanobacter-sp.-C01:0.1521380172,WP_158752649.1-Dyella-sp.-S184:0.0332048623)54.6/31:0.0192359585,WP_077445946.1-Rhodanobacter-sp.-C05:0.0701220804)87.1/31:0.0292629441)76.8/26:0.0280957589,WP_184416890.1-Rhodanobacter-sp.-MP7CTX1:0.0711921140)93.5/97:0.0542475192,WP_015446789.1-Rhodanobacter-denitrificans:0.0624681820)95.6/100:0.0685983596)82.3/49:0.0364683757,(((WP_168608661.1-Oleiagrimonas-citrea:0.0813335644,WP_113064358.1-Oleiagrimonas-sp.-MCCC-1A03011:0.0317215439)89.7/100:0.0828277017,WP_043099571.1-Oleiagrimonas-soli:0.0882208761)99.8/100:0.2003303863,((WP_233843762.1-Dyella-sp.-2HG41-7:0.0510249375,WP_188793403.1-Dyella-nitratireducens:0.1004693897)54.4/77:0.0470356895,WP_126673574.1-Dyella-dinghuensis:0.0836321942)91.2/100:0.0845103072)82.3/57:0.0691094214)52.2/26:0.0152817028,(((WP_214570707.1-unclassified-Dyella:0.2208657935,((WP_130619635.1-Dyella-amyloliquefaciens:0.1158689959,WP_199044928.1-Dyella-sp.-ASV24:0.0941571756)76.6/94:0.0245686972,(WP_266169930.1-Dyella-sp.-RXD159:0.1032756433,(WP_114846719.1-Dyella-tabacisoli:0.0686444407,WP_090451016.1-Dyella-sp.-OK004:0.0934815349)88.4/90:0.0458779409)20.7/68:0.0162987717)74/90:0.0123334258)86.9/94:0.0315717495,(WP_111981026.1-Dyella-jiangningensis:0.2318934764,WP_266183414.1-Dyella-sp.-RBB1W86:0.1131492715)0/34:0.0341509132)88.5/94:0.0538382732,WP_204636097.1-Dyella-kyungheensis:0.2038204836)82.5/57:0.0372538474)79.6/53:0.0428296865,(WP_049621948.1-Frateuria-defendens:0.1555789612,WP_179481164.1-unclassified-Rhodanobacter:0.1624829408)79.8/72:0.0430216827)69.6/41:0.0336804597,WP_019467068.1-Dyella-japonica:0.3313880773)100/100:0.3743306497,(WP_205000141.1-Parashewanella-hymeniacidonis:0.0812494395,WP_129946484.1-Shewanella-sp.-OPT22:0.0757403178)100/100:1.0871419340)83.7/50:0.1185179428,(((((WP_256181981.1-Cloacibacillus-evryensis:0.2626958120,WP_087368268.1-Cloacibacillus-sp.-An23:0.2305388393)100/100:0.7670868002,WP_141198172.1-Persicimonas-caeni:0.7054972862)96.2/94:0.3247699421,(((WP_084578244.1-Sporomusa-malonica:0.6250160606,WP_131919244.1-Heliophilum-fasciatum:0.3897249175)26.9/96:0.0921991179,(WP_090940141.1-Pelosinus-propionicus:0.3869397195,WP_240986382.1-Acididesulfobacillus-acetoxydans:0.3855368906)80.8/92:0.0777376390)47.3/50:0.0919889146,WP_092486361.1-Desulfoscipio-geothermicus:0.1704336265)60.4/92:0.1043316595)75/93:0.0723024604,((((WP_246588317.1-Pelorhabdus-rhamnosifermentans:0.2900543654,(WP_092074275.1-Dendrosporobacter-quercicolus:0.3717070031,(WP_145090815.1-Sporomusa-sp.-KB1:0.0726039004,WP_093793074.1-Sporomusa-acidovorans:0.0864487414)95.9/100:0.1128518379)83.7/56:0.0816923594)48.9/20:0.0703312260,(((WP_007952170.1-Pelosinus-fermentans:0.1434469622,WP_007939097.1-Pelosinus:0.1694334512)95.7/100:0.1383605918,((WP_132081660.1-Anaerospora-hongkongensis:0.1558437700,WP_132083235.1-Anaerospora-hongkongensis:0.2584393528)96.4/100:0.1565660789,(WP_094603851.1-Sporomusa-silvacetica:0.1628350864,WP_144352952.1-Sporomusa-termitida:0.1982006362)88.1/100:0.0991521983)68.6/42:0.0763567653)0.5/12:0.0112946077,((WP_054258662.1-Propionispora-sp.-2/2-37:0.3054561466,(WP_054258683.1-Propionispora-sp.-2/2-37:0.2110740316,(WP_149735423.1-Propionispora-hippei:0.0903830944,WP_091744506.1-Propionispora-vibrioides:0.2240418998)79.6/97:0.0606261644)89.6/96:0.0884943380)23.6/10:0.0353662804,WP_188400350.1-Sporomusa-sp.-GT1:0.4431205832)29.9/7:0.0382757406)36.8/11:0.0398301748)83.5/93:0.0648968114,WP_110954047.1-Massilibacillus-massiliensis:0.4063628408)23.2/25:0.0179035369,WP_188399481.1-Sporomusa-sp.-GT1:0.4022191565)88.8/100:0.0855280144)94.9/93:0.1569240032,((((((WP_200343816.1-Halochromatium-glycolicum:0.2182702108,WP_201222000.1-Halochromatium-roseum:0.1882654739)94.5/100:0.1480759047,(WP_200237937.1-Thiohalocapsa-halophila:0.0878390177,WP_058558062.1-Thiohalocapsa-sp.-ML1:0.3572530304)97.4/100:0.1982247218)98.6/100:0.1910249129,WP_182584206.1-Thiospirillum-jenense:0.6074257691)0/58:0.0689597017,(WP_200286492.1-Rhabdochromatium-marinum:0.1950997107,WP_009150811.1-Thiorhodovibrio-sp.-970:0.0953714614)98/100:0.2859937442)97.1/100:0.1594707182,(((((WP_200207675.1-Chromatium-weissei:0.3135158140,WP_200154756.1-Chromatium-okenii:0.1043675825)99.3/100:0.2127759028,(((WP_200158060.1-Allochromatium-vinosum:0.0955971157,WP_200376120.1-Thiocystis-violacea:0.1522504843)89.6/100:0.0705828238,(WP_164456073.1-Thiorhodococcus-minor:0.1284910796,(WP_164652045.1-Thiorhodococcus-mannitoliphagus:0.0657381923,WP_200330387.1-Thiocystis-violacea:0.1364938699)23.1/8:0.0219399247)96.8/100:0.1008626174)7.1/9:0.0228556283,((WP_043749566.1-Imhoffiella-purpurea:0.1050502848,WP_007039504.1-Thiorhodococcus-drewsii:0.0897195632)93.1/100:0.0739883149,(WP_201092787.1-Thiocystis-minor:0.1060952736,WP_132978621.1-Thiobaca-trueperi:0.1416648574)84.5/100:0.0277810486)0/7:0.0000021415)85.1/20:0.0394528629)85.2/98:0.0407473672,(WP_200388819.1-Thiocapsa-imhoffii:0.2410855146,((WP_007191850.1-Thiocapsa-marina:0.0585795760,WP_232409912.1-Thiocapsa-bogorovii:0.0359727941)0/4:0.0115192624,WP_120796078.1-Thiocapsa-rosea:0.0386118022)96.9/100:0.1195116172)94.9/100:0.1116871821)92.9/100:0.0995239863,(WP_089729519.1-Candidatus-Thiosymbion-oneisti:0.5578944080,(WP_100918325.1-Candidatus-Thiodictyon-syntrophicum:0.1158653725,WP_020502667.1-Lamprocystis-purpurea:0.1494259009)88.9/100:0.0781283988)49.7/94:0.0857668946)93.5/100:0.1088631076,(WP_141730858.1-Oligoflexus-tunisiensis:0.6588498431,WP_101758446.1-Oceanicoccus-sp.-KOV_DT_Chl:0.6388769659)95/98:0.2414949528)81.9/97:0.0506418841)73.5/97:0.0988749911,WP_018175268.1-unclassified-Thioalkalivibrio:0.5941871572)62/88:0.1024830636)85/83:0.0848360738)42.7/50:0.1068862311,WP_008811140.1-Parasutterella-excrementihominis:0.9450917345)75.5/40:0.0234264864)82.2/36:0.0618390509,(((((((((((((((((WP_094934028.1-Enterobacter-cloacae-complex:0.9046170433,WP_016795256.1-Vibrio-cyclitrophicus:0.2414487047)89.8/94:0.1508417040,WP_039473409.1-Vibrio-sinaloensis:0.4407213629)85/94:0.0848696868,(WP_010368546.1-Pseudoalteromonas-piscicida:0.6323323227,WP_086590801.1-Vibrio-parahaemolyticus:0.5986514770)80.4/94:0.0876475885)94.4/100:0.1472148762,((WP_061013169.1-Vibrionaceae:0.3577847383,WP_155662549.1-Aliivibrio-fischeri:0.1952023767)90.1/91:0.1286364622,WP_171328835.1-Vibrio-splendidus:0.3619809833)73.1/46:0.0328086981)71.4/59:0.0986127749,((((WP_057065012.1-Citrobacter:1.0242616636,WP_001100815.1-Escherichia-coli:0.4006226973)93.7/100:0.2865650272,WP_207379479.1-Shewanella-sedimentimangrovi:0.2255806097)24.3/45:0.1065149544,WP_269408820.1-Pseudoalteromonas-shioyasakiensis:0.3759130201)56.6/54:0.0450094292,WP_016537220.1-Cedecea-davisae:0.3180321150)36.3/76:0.0337542199)54/77:0.0297238912,(WP_195504469.1-Citrobacter-farmeri:0.4144443890,WP_064554809.1-Buttiauxella-noackiae:0.4294866456)99.5/99:0.3670502448)71.1/94:0.0598023445,(WP_240304790.1-Vibrio-campbellii:0.7027132594,WP_004741399.1-Vibrio:0.3733926564)93.4/99:0.2061785263)92.3/100:0.1412229172,WP_216735790.1-Hahella-sp.-HN01:0.4213616920)97.9/99:0.2101355192,WP_127021653.1-Rheinheimera-mangrovi:0.6228725931)77.2/90:0.0368730002,(WP_132584255.1-Rheinheimera-sp.-D18:0.4940028222,(((((WP_131912015.1-Celerinatantimonas-diazotrophica:0.3386025392,((WP_092385961.1-Halopseudomonas-salegens:0.2947999372,WP_036859756.1-Porticoccus-hydrocarbonoclasticus:0.3582208537)85.7/99:0.1289675066,((WP_076722556.1-Marinobacter-lutaoensis:0.3129788119,(WP_106695101.1-Marinobacter-shengliensis:0.1007896829,(WP_222473125.1-Marinobacter-daepoensis:0.1018703565,(WP_092030923.1-Marinobacter-sp.-DSM-26671:0.1710116243,WP_148609564.1-Marinobacter-sp.-BW6:0.1189864673)29.2/97:0.0412891118)0/93:0.0000029531)96.4/100:0.1347539011)35.6/99:0.0456351575,WP_036130839.1-Marinobacter-nitratireducens:0.1946288485)66.7/99:0.0728349363)81.1/100:0.0667408883)83.8/83:0.0674310498,(WP_027875186.1-Spongiibacter-marinus:0.2239402782,WP_204995064.1-Spongiibacter-marinus:0.1373486999)86.4/100:0.0641205352)86.1/83:0.0470994597,WP_168451636.1-Spongiibacter-thalassae:0.0628940628)42.4/82:0.0526013086,WP_062382560.1-Zhongshania-aliphaticivorans:0.0940869916)71.1/83:0.0856580439,((WP_184464382.1-Zhongshania-antarctica:0.0649198032,WP_198570926.1-Spongiibacter-nanhainus:0.1233504838)74.5/63:0.0088746655,WP_103683243.1-Marortus-luteolus:0.0441716972)84.4/65:0.0732528355)89.7/96:0.0949428614)85.9/90:0.0684407353)96.1/98:0.0976126077,((((((((((((((((((WP_171588903.1-Pelistega-europaea:0.5022634193,(WP_087653467.1-Klebsiella/Raoultella-group:0.2123210254,WP_028357569.1-Brackiella-oedipodis:0.3690421333)67.3/99:0.0719633266)78.2/98:0.0476498666,WP_154751639.1-unclassified-Erwinia:0.3593217863)86.7/73:0.0512449864,(((WP_032441370.1-Klebsiella-pneumoniae:0.1741109777,(WP_015583911.1-Raoultella-ornithinolytica:0.0858633359,WP_265776798.1-Klebsiella-oxytoca:0.2253752422)55.4/100:0.0681337042)90.7/100:0.0877443689,(WP_029991521.1-Tatumella-ptyseos:0.1082364433,(WP_212707699.1-Tatumella-sp.-JGM130:0.0627058825,WP_087487466.1-Tatumella-citrea:0.0843970387)68.7/99:0.0201586453)83.3/100:0.0399026147)88/99:0.0929553008,WP_186921353.1-Klebsiella-quasipneumoniae:0.2702294043)27.1/51:0.0217645154)74.1/55:0.0138108067,WP_140470594.1-Ewingella-americana:0.4076485278)77/79:0.0324523676,((WP_040666911.1-Neisseria-wadsworthii:0.1397011855,WP_135034416.1-Neisseria:0.2512632286)92/100:0.0927266919,(((WP_214236207.1-Rosenbergiella-sp.-S61:0.1002317178,WP_241647140.1-Rosenbergiella-sp.-JB07:0.1838375437)57.7/100:0.0530936790,WP_214195533.1-Rosenbergiella-collisarenosi:0.1354464178)95.2/100:0.1337789293,WP_101506252.1-Erwinia-sp.-B116:0.2032777238)59.9/100:0.0653406670)79.4/98:0.0368744894)85.3/78:0.0313320750,(WP_142430937.1-Klebsiella-oxytoca:0.2230498302,(WP_239614184.1-Raoultella-ornithinolytica:0.1634022353,((WP_032699180.1-Raoultella:0.0760000711,WP_228399925.1-Raoultella:0.0452601802)73/100:0.0205369142,WP_004138711.1-Klebsiella:0.0260876816)93/100:0.0521009496)81.9/70:0.0265039104)83.4/66:0.0406602670)84.2/60:0.0199535866,((((WP_228398973.1-Insectihabitans-xujianqingii:0.1421374123,WP_074823218.1-Pragia-fontium:0.1328787056)95/100:0.1132457788,WP_130971788.1-Hafnia-alvei:0.1843776659)83.5/94:0.0407098539,WP_165428828.1-Rouxiella-badensis:0.1513406672)74.1/93:0.0090053486,(WP_142514871.1-Klebsiella-huaxiensis:0.2684195677,((WP_112167140.1-Klebsiella:0.1950498728,WP_200519240.1-Klebsiella-michiganensis:0.1415273540)92.3/100:0.0790019963,WP_137713144.1-Brenneria-rubrifaciens:0.1677615163)55.8/93:0.0109444134)76.2/27:0.0166750021)88.8/28:0.0206373635)0/25:0.0000024053,((WP_131063651.1-Raoultella-planticola:0.0317083196,WP_163980548.1-Raoultella-ornithinolytica:0.0553512965)98.9/100:0.1206730059,(WP_148250181.1-Raoultella-terrigena:0.1393614957,WP_032686784.1-Raoultella-planticola:0.0441933393)96.3/99:0.0818322705)79.6/56:0.0291683472)79.1/26:0.0147969508,(WP_193937755.1-Klebsiella-grimontii:0.2603884494,(WP_024360036.1-Klebsiella:0.0891054789,WP_234284225.1-Klebsiella-oxytoca:0.1037900156)91.9/100:0.0782087710)89.6/25:0.0539493074)0/21:0.0000027663,(WP_024553491.1-Franconibacter-helveticus:0.5967632198,(((((((WP_136167279.1-Brenneria-sp.-CFCC-11842:0.0649606575,WP_264089117.1-unclassified-Brenneria:0.1489955772)66.4/87:0.0567117949,WP_109053612.1-Brenneria-roseae:0.1409841412)71.7/78:0.0205794181,(((((WP_180742451.1-Pectobacterium-sp.-CFBP8739:0.3687651929,((((((((((WP_047710158.1-Pectobacterium-atrosepticum:0.0830496126,WP_198482318.1-Pectobacterium-parmentieri:0.0527911790)33.2/88:0.0127883535,WP_033071760.1-Pectobacterium-parmentieri:0.0437904818)92.3/99:0.0571508796,WP_226305805.1-Pectobacterium-versatile:0.0292321767)83/99:0.0328729112,((WP_039280054.1-Pectobacterium-brasiliense:0.0477101853,WP_039517132.1-Pectobacterium-brasiliense:0.0463605755)86.6/87:0.0301167352,WP_039346077.1-Pectobacterium-fontis:0.0438479983)74.4/83:0.0091575573)91.6/97:0.0385765258,WP_103972204.1-Pectobacterium:0.0228478745)77.2/33:0.0104919126,WP_205558955.1-Pectobacterium-brasiliense:0.0809436884)0/23:0.0000023899,(WP_151242707.1-Pectobacterium-carotovorum:0.0579839793,WP_116186106.1-Pectobacterium-aquaticum:0.0617113017)0.2/85:0.0099332819)84.2/95:0.0304141524,WP_119888241.1-Pectobacterium:0.0711877986)80.5/81:0.0199141602,WP_205946867.1-Pectobacterium:0.0807412072)70.7/78:0.0105364323,WP_233976643.1-Pectobacterium-versatile:0.0386981979)0/71:0.0000028786)79.9/21:0.0190066995,((((WP_180777316.1-Pectobacterium-aroidearum:0.0847645697,WP_107169883.1-Pectobacterium:0.0638194345)85/100:0.0245647889,WP_039504461.1-Pectobacterium:0.0314486967)84.4/99:0.0192220074,(WP_233968669.1-Pectobacterium-polaris:0.0776208111,WP_048259384.1-Pectobacterium-peruviense:0.0194865065)91/97:0.0402078964)77.5/87:0.0104140316,WP_138255615.1-Pectobacterium:0.0304983924)76.4/90:0.0117590197)81/21:0.0196939783,WP_109411993.1-Pectobacterium-polaris:0.0715214383)0/21:0.0000023403,((WP_249680869.1-Pectobacterium-polaris:0.0643472020,WP_125232508.1-Pectobacterium-versatile:0.0692207767)76.5/97:0.0197707916,WP_005974084.1-Pectobacterium-wasabiae:0.0419681633)86.2/96:0.0220882978)96.2/84:0.0741623644,(WP_132452860.1-Samsonia-erythrinae:0.1639837260,WP_130637964.1-Pectobacterium-versatile:0.0881139318)44.1/98:0.0265819776)87.4/95:0.0595094794)97.8/100:0.1425819881,((WP_121576612.1-Brenneria-alni:0.1274999862,WP_095845203.1-Gibbsiella-quercinecans:0.1249130567)73.7/70:0.0090884876,((WP_214648782.1-Pectobacterium-carotovorum:0.0509576001,((WP_261846379.1-Pectobacterium-carotovorum:0.0838998120,WP_121590317.1-Brenneria-goodwinii:0.0091059261)93.7/100:0.0405467562,(WP_182099415.1-Pectobacterium-aroidearum:0.0504681929,WP_039482612.1-Pectobacterium:0.0398840159)72.4/100:0.0121970917)4.4/91:0.0096639839)72.9/98:0.0088135765,WP_249228983.1-Brenneria-tiliae:0.0649157418)93.2/99:0.0516794312)75.1/66:0.0312624349)0/53:0.0000028987,(((((WP_168640580.1-Dickeya-sp.-CFBP-2040:0.1583963259,(WP_027712673.1-Dickeya-chrysanthemi:0.0606406315,WP_012770533.1-Dickeya:0.0488928561)74.3/99:0.0096659851)75/98:0.0098270892,(WP_012883965.1-Dickeya-parazeae:0.0958614220,WP_236306746.1-Dickeya-zeae:0.0488671708)79.6/100:0.0190625137)74/98:0.0078583306,((WP_038918362.1-Dickeya:0.0423365939,WP_123251184.1-Dickeya-undicola:0.0650270596)49.5/98:0.0094320658,WP_022632763.1-Dickeya-solani:0.0370886025)90.3/99:0.0312687333)98.5/100:0.1344590895,(WP_148877341.1-Serratia-marcescens:0.0972234744,WP_193158894.1-Serratia-marcescens:0.0722303707)96/100:0.1074653929)84.5/98:0.0437397405,WP_062791489.1-Serratia-plymuthica:0.1025447666)85.4/88:0.0287179631)85.6/74:0.0327767344,WP_096000273.1-Klebsiella-pneumoniae:0.2838694404)33.3/89:0.0262703833,WP_248464278.1-Pectobacterium-versatile:0.2366086192)54.4/91:0.0322410963)40/48:0.0192569720)91.4/95:0.0623511909,(((((WP_267333169.1-unclassified-Gilliamella:0.2434623805,WP_267361838.1-Gilliamella-sp.-B2772:0.2184330324)53.8/93:0.0152571739,WP_086361229.1-Gilliamella:0.1859522491)88.4/90:0.0398131924,((WP_065574597.1-Gilliamella-apicola:0.0816933654,WP_065569248.1-Gilliamella-apicola:0.0513814282)71.7/100:0.0111777095,WP_065610260.1-Gilliamella-apicola:0.0585996557)97.2/100:0.0941961712)77.8/46:0.0242717372,(((((WP_086319877.1-Gilliamella-apis:0.0287558223,WP_198214961.1-Gilliamella-sp.-M0320:0.0792924325)87.5/99:0.0466664448,WP_039129858.1-Gilliamella:0.0342906947)74.5/99:0.0149021559,(WP_065737496.1-Gilliamella-apicola:0.0485043413,WP_267331332.1-unclassified-Gilliamella:0.0899287761)97/100:0.0791020304)94.9/100:0.0672892864,(WP_065587409.1-Gilliamella-apicola:0.0876332362,WP_065580031.1-Gilliamella-apicola:0.0954931226)90.4/100:0.0443626673)0/17:0.0000025093,(WP_065579324.1-Gilliamella-apis:0.1165874221,WP_160432071.1-Gilliamella-sp.-Pas-s25:0.1645245610)43.3/71:0.0306607653)91/99:0.0593944579)96/89:0.1673128600,(WP_247177500.1-Escherichia-coli:0.7956148762,YP_009985672.1-Escherichia-phage-muut:0.3058123043)99.2/99:0.5237835251)83.2/98:0.0854430748)65.2/81:0.0767566963,(WP_246757812.1-Erwinia-phyllosphaerae:0.2856282701,WP_232759612.1-unclassified-Pantoea:0.2040448189)89.6/98:0.1101791129)55.8/80:0.0285172252,((WP_146378871.1-Cronobacter-turicensis:0.0854999458,WP_032180402.1-Escherichia-coli:0.1373168805)99.9/100:0.2917433590,((WP_126816534.1-Enterobacter-mori:0.1257280299,WP_049841125.1-Klebsiella-sp.-RIT-PI-d:0.5906421017)27.5/68:0.0585650060,WP_256879438.1-Escherichia-albertii:0.1467318229)91.1/82:0.0874983796)86.5/86:0.0770881800)87.7/92:0.0890452698,WP_253460089.1-Erwinia-persicina:0.4120192625)40/82:0.0719407426,WP_100256321.1-Reinekea-forsetii:0.9491083540)0/80:0.0000026463,((WP_168836617.1-Vibrio-agarilyticus:0.1769381449,(WP_076537303.1-Shewanella-sp.-UCD-KL21:0.0313624390,WP_102531614.1-Shewanella-sp.-10N.286.52.B9:0.0721128966)100/100:0.8582660746)90.2/92:0.1417875195,(WP_137671336.1-Agarivorans-sp.-Toyoura001:0.0589888414,WP_221077378.1-Agarivorans-albus:0.0847204931)89.6/100:0.1248139093)95.6/93:0.1385788933)92.8/100:0.0591972890,((((WP_261844981.1-Amphritea-ceti:0.5962057910,(((WP_269124033.1-Marinomonas-sp.-15G1-11:0.0352152691,WP_191602238.1-Marinomonas-algicola:0.1264604152)98.9/100:0.2133495365,WP_048693477.1-Catenovulum-maritimum:0.3096312887)83.2/100:0.0791562979,WP_228191388.1-Marinomonas-communis:0.2088309152)78.9/86:0.0421510562)20.6/15:0.0405438671,WP_067204681.1-Marinomonas-aquimarina:0.0372802792)69.9/15:0.0162204822,WP_055461762.1-Marinomonas-fungiae:0.1003016268)87.1/67:0.0642575688,WP_067216963.1-Marinomonas-gallaica:0.0634117397)97.9/100:0.1720559803)91/98:0.0828606969,((WP_145010177.1-Pseudomonas-psychrotolerans:0.0246100865,WP_077172422.1-Pseudomonas-psychrotolerans:0.0596995847)100/100:0.3416285286,WP_084334293.1-Pseudomonas-indica:0.1466500639)91.2/100:0.0910651498)90.3/98:0.0889911855)80.2/80:0.0535761695,(((((WP_138320019.1-Thalassotalea-litorea:0.2307376477,WP_136736185.1-Thalassotalea-mangrovi:0.1873068676)99.8/100:0.4537074133,WP_114327629.1-Candidatus-Colwellia-aromaticivorans:0.1862230865)85.6/100:0.0934083289,WP_101108574.1-Psychromonas-sp.-psych-6C06:0.3256397653)76/91:0.0561773556,((WP_016794198.1-Vibrio-cyclitrophicus:0.0645281283,WP_102424249.1-Vibrio-cyclitrophicus:0.0601914525)78.4/100:0.0341340439,(WP_261903470.1-Vibrio-fortis:0.1194783788,WP_240207223.1-Vibrio-sp.-CyArs1:0.0611142635)88.8/100:0.0618334484)98.1/100:0.1851705238)42.5/88:0.0790992292,WP_148513713.1-Vibrio:0.4076234229)10.4/33:0.0604852909)89.3/50:0.0932468469,(WP_067584343.1-Endozoicomonas-ascidiicola:0.3081460949,((WP_260491935.1-Pseudoalteromonas-flavipulchra:0.2874831717,(WP_039494249.1-Pseudoalteromonas-flavipulchra:0.1263940358,WP_055022295.1-Pseudoalteromonas-sp.-P1-26:0.1079582344)53.4/100:0.0647734909)80.5/100:0.0379039169,(WP_149602089.1-Pseudoalteromonas-distincta:0.0558872541,WP_248518672.1-Pseudoalteromonas-sp.-2CM32C:0.0374959939)97.6/100:0.1492878649)99.1/100:0.3134339914)10.5/24:0.0753474794)78.3/76:0.0590652171,WP_150896168.1-Vibrio-fortis:0.2941611949)83.5/78:0.0861540749,WP_199145406.1-Citrobacter-sp.-ASV29:0.5112506661)97.8/100:0.2718625243,(WP_133132051.1-Legionella-sp.-W10-070:0.4425218262,(((((((((((((WP_027223798.1-Legionella-pneumophila:0.0813165489,WP_106142999.1-Legionella-pneumophila:0.0454906672)63.8/35:0.0040156687,(WP_061513621.1-Legionella:0.0237934557,WP_027226296.1-Legionella-pneumophila:0.0709211374)99.5/94:0.0897525588)76.7/38:0.0120836174,WP_044499121.1-Legionella-pneumophila:0.0400708085)36.5/38:0.0087141132,WP_265717942.1-Legionella-sp.-PATHC032:0.0749824990)71.3/100:0.0253708122,WP_027220106.1-Legionella-pneumophila:0.0212664918)84.7/100:0.0260440053,WP_027266778.1-Legionella-pneumophila:0.0466729869)96/100:0.1085078944,(WP_058494182.1-Legionella-worsleiensis:0.1931745759,(WP_127106294.1-Legionella-sp.-km535:0.0545431460,WP_058474153.1-Legionella-quateirensis:0.0417405805)89.7/100:0.0548245440)93.4/100:0.0952281568)92.3/100:0.0900693106,WP_031567420.1-Legionella-wadsworthii:0.2759475394)74.7/99:0.0351439892,WP_045097199.1-Legionella-fallonii:0.1683112277)88.5/100:0.0363695522,(WP_232890831.1-unclassified-Legionella:0.0597121128,(WP_058459148.1-Legionella-bozemanae:0.0343268229,WP_058518697.1-Legionella-parisiensis:0.0653149449)83.1/100:0.0356530782)99.4/100:0.1349885275)60.4/96:0.0328150202,WP_058513550.1-Legionella-santicrucis:0.1555572216)91.1/92:0.0481158776,WP_058463758.1-Legionella-cincinnatiensis:0.0286822745)0/10:0.0558394122,WP_128130294.1-Legionella-sainthelensi:0.0203326565)99.2/100:0.4707314017)99.1/100:0.4134552730)81.1/43:0.0609184340,WP_062153084.1-Beggiatoa-leptomitoformis:0.7463373514)86.4/39:0.0894210558)81.3/81:0.0594412732)92.1/27:0.0962642713)76.7/9:0.0385123360,(((KAI6244242.1-Aphelenchoides-fujianensis:0.8293499526,KAI6170547.1-Aphelenchoides-bicaudatus:1.1293167862)90.6/100:0.2663464285,WP_077329105.1-Virgibacillus-siamensis:0.7403863214)10.3/97:0.1085333403,WP_110520589.1-Herpetosiphon-llansteffanensis:1.2865468460)94.3/63:0.1776366754)98.6/91:0.3187247746)83.6/98:0.0984359544,((((((WP_125841544.1-Stutzerimonas-stutzeri:0.5439317227,((WP_262731117.1-Alcanivorax-marinus:0.2678035901,(WP_106232428.1-Halomonas-ventosae:0.1009236644,WP_228234339.1-Alcanivorax-marinus:0.2755846716)84.5/100:0.0535744788)81.9/100:0.0311859920,WP_175070852.1-Halomonas-taeanensis:0.1620065414)3.4/80:0.0930099058)84/100:0.1166946032,(WP_019018778.1-Halomonas-lutea:0.4198801029,WP_092999033.1-Thiohalomonas-denitrificans:0.1592843576)93.2/100:0.1934159075)87.3/100:0.0744657496,((WP_146294265.1-Euhalothece-natronophila:0.3452835243,WP_159299499.1-Spiribacter-sp.-SSL99:0.3092175834)90.3/100:0.1328576876,WP_011630054.1-Alkalilimnicola-ehrlichii:0.2397045888)82.2/100:0.0905280290)86.9/100:0.0616759211,WP_127477393.1-Sulfurivermis-fontis:0.4211974998)84.6/100:0.0712661309,WP_028988655.1-Thermithiobacillus-tepidarius:0.3650256932)92.9/100:0.1273805531,WP_069857109.1-Desulfoplanes-formicivorans:0.5182620946)77.6/99:0.0665609685)91.8/97:0.1098929772,WP_156872703.1-Ketobacter-sp.-MCCC-1A13808:0.6670036176)29.1/74:0.0697394384,(((WP_151033206.1-Cellvibrio-sp.-KY-GH-1:0.1078145067,WP_049631514.1-Cellvibrio-sp.-pealriver:0.1056627750)83.7/100:0.1215311634,WP_062059887.1-Cellvibrio-sp.-OA-2007:0.0491891254)100/100:0.5109950759,WP_178968040.1-Neptunomonas-phycophila:0.4733590864)93.1/96:0.2488971413)79.6/89:0.0602216205)94.2/95:0.1182867638,WP_255857717.1-Bacterioplanoides-sp.-SCSIO-12839:0.5514490241)93.6/92:0.1254170514,(WP_248714967.1-Agarivorans-sp.-TSD2052:0.0941565080,WP_152781274.1-Agarivorans-sp.-B2Z047:0.1051268824)99.9/100:0.2385389220)62.1/11:0.0273416650,((WP_220103444.1-Neiella-holothuriorum:0.1505388607,WP_087505717.1-Neiella-marina:0.2117947966)96.6/100:0.1222342418,((WP_028862352.1-Psychromonas-aquimarina:0.2803373611,WP_187408568.1-Saccharophagus-sp.-K07:0.2133245753)85.6/100:0.0715000745,(WP_251261538.1-Echinimonas-agarilytica:0.2164497465,(WP_096084688.1-Agaribacterium-haliotis:0.5123312969,WP_188151970.1-Teredinibacter-waterburyi:0.3015343712)93.2/92:0.1839090783)44/90:0.0309327560)95.4/92:0.1396840266)78/82:0.0726992602)66.8/13:0.0933017178,((WP_142941196.1-Aliikangiella-marina:0.4398060230,(WP_081148784.1-Cognaticolwellia-beringensis:0.0576207131,WP_007634918.1-Paraglaciecola-psychrophila:0.0552761901)99.8/100:0.3096075046)12.4/57:0.0464174311,WP_068546663.1-Thalassotalea-crassostreae:0.3314149637)91/99:0.0903811718)65.3/12:0.0320838487,((((((WP_167868984.1-Shewanella-sp.-WXL01:0.6143063192,(WP_222525605.1-Ferrimonas-balearica:0.1862036422,WP_013344594.1-Ferrimonas-balearica:0.2237138734)91.8/100:0.1860802318)78.8/90:0.1384775465,WP_084884043.1-Vibrio-sp.-qd031:0.1474114715)72.6/69:0.0338095350,((WP_193630602.1-Shewanella-benthica:0.2895188067,WP_159656123.1-Vibrio-atypicus:0.0641451215)79.4/98:0.0916906515,(WP_269599241.1-Salinivibrio-kushneri:0.0852480624,WP_136483218.1-Vibrio-sp.-H11:0.0967065426)95/100:0.0965119394)74.6/76:0.0484818231)72.8/78:0.0285275201,(((((WP_047049276.1-Vibrio-mexicanus:0.2731855083,WP_264874461.1-Vibrio-agarivorans:0.1307609991)84/68:0.0320835776,(WP_022943097.1-Psychromonas-hadalis:0.2271728947,WP_020335761.1-Vibrio-natriegens:0.1196525519)83/84:0.0460873140)61.6/62:0.0233555531,WP_047874860.1-Photobacterium-aphoticum:0.3358051942)4.6/57:0.0091692712,(((WP_050648448.1-Vibrio-sp.-J2-4:0.2020770947,WP_205390674.1-Vibrio:0.1144780074)68.1/72:0.0146622938,WP_140076235.1-Vibrio-parahaemolyticus:0.1851983396)85.1/46:0.0358953066,(WP_012143346.1-Shewanella-sediminis:0.3376898081,(WP_167678613.1-Shewanella-aestuarii:0.2426061629,((((WP_188955407.1-Shewanella-ulleungensis:0.0756894691,WP_137222113.1-Shewanella-sp.-MEBiC00475:0.0868523314)86.9/26:0.0491356346,((((WP_138577097.1-unclassified-Pseudoalteromonas:0.1426656682,(WP_264390127.1-Pseudoalteromonas-sp.-A3:0.0347886761,(WP_096739047.1-Pseudoalteromonas-atlantica:0.0528779019,WP_248462211.1-Pseudoalteromonas-sp.-2CM37A:0.0241260869)90.2/100:0.0288561650)77.4/98:0.0167877839)87.3/93:0.0392883486,WP_182693438.1-Pseudoalteromonas-sp.-SR45-5:0.0790129730)32.5/67:0.0225288189,WP_256848444.1-Pseudoalteromonas-carrageenovora:0.0348433941)63.5/67:0.0064103978,WP_126005793.1-Pseudoalteromonas-sp.-Xi13:0.0395513393)97.7/99:0.1062963914)31.2/20:0.0477820322,WP_155694061.1-Psychrosphaera-haliotis:0.1753885721)85.2/25:0.0409251020,(WP_113818410.1-Psychromonas-sp.-B3M02:0.0828881655,(WP_025564776.1-Psychromonas-sp.-SP041:0.0704963178,WP_166372414.1-Psychromonas-sp.-SA13A:0.0178024165)76.8/100:0.0686800602)95.2/100:0.0966085535)73.2/25:0.0124157289)82/96:0.0320487058)68.6/28:0.0122838063)73.9/51:0.0159242884)88.7/85:0.0225747575,WP_164649367.1-Vibrio-astriarenae:0.1323415740)78.4/88:0.0184412413)0/74:0.0000022080,(WP_016959527.1-Enterovibrio-norvegicus:0.0789089042,WP_229491984.1-Enterovibrio-norvegicus:0.1868730176)99.2/100:0.1437938247)81.8/96:0.0501868389,WP_093326971.1-Thalassotalea-agarivorans:0.1558028637)95.9/100:0.1158645317)87.7/16:0.0471703202,((((((WP_160153838.1-Microbulbifer-sp.-ALW1:0.4084185643,((WP_263600466.1-Microbulbifer-agarilyticus:0.1875874271,WP_078085574.1-Microbulbifer-mangrovi:0.1197039929)38.9/87:0.0150289434,WP_029249715.1-Microbulbifer-agarilyticus:0.1432893621)86.1/87:0.0806809895)97.2/100:0.1692223355,(((WP_085356321.1-Neisseria-dumasiana:0.2732201029,(WP_101133177.1-Neisseria-meningitidis:0.0836936084,(WP_005552781.1-Aggregatibacter-actinomycetemcomitans:0.2859200621,(WP_127202078.1-Neisseria-meningitidis:0.1595081579,WP_237092367.1-Neisseria-sp.-ZJ104:0.1986289668)86.2/100:0.0784516707)18/37:0.0114560868)83.7/87:0.0738208612)89.9/100:0.2148216573,WP_031579865.1-Ruminobacter-sp.-RM87:0.6802295028)95.1/99:0.1890004133,((WP_138624780.1-Pseudoalteromonas-sp.-S4492:0.3198621512,WP_130050482.1-Pseudoalteromonas-sp.-MEBiC-03485:0.2212188090)85.2/100:0.1271890154,(WP_189610147.1-Saccharospirillum-salsuginis:0.4374639655,WP_111640782.1-Marinimicrobium-alkaliphilum:0.4978221601)75.9/100:0.0899587405)72.4/100:0.1645753725)43.3/94:0.0501276847)87.3/91:0.0590551401,WP_186505417.1-Neptunicella-marina:0.2729313395)91/59:0.0973935924,(WP_102798355.1-Bowmanella-denitrificans:0.1042826048,WP_214344498.1-Bowmanella-yangjiangensis:0.0438236441)99.9/100:0.2446520388)73.5/50:0.0302299537,(WP_009839692.1-Pseudoalteromonas-tunicata:0.8444126273,(WP_124026287.1-Alteromonas-sediminis:0.2058055402,WP_100655941.1-Alteromonas-flava:0.3283416402)96.6/99:0.1753309017)85.1/28:0.0676770679)75.9/22:0.0254923329,((WP_118961222.1-Colwellia-sp.-RSH04:0.5989324733,(WP_216048042.1-Aliiglaciecola-lipolytica:0.2576616519,WP_057181266.1-Colwellia:0.3267565761)64.8/27:0.0198785226)65/23:0.0616483939,(WP_131095783.1-Alteromonas-sp.-KUL42:0.1041662445,(((((((WP_100972994.1-Alteromonas-sp.-MB-3u-76:0.1319803718,WP_269520955.1-Alteromonas-sp.-BMJM2:0.0666718874)72.7/79:0.0399947510,(((((((WP_156078883.1-Alteromonas:0.0858142787,(WP_044058190.1-Alteromonas-australica:0.0549198142,WP_159580202.1-Alteromonas-sp.-KUL106:0.0274433029)74.6/88:0.0071446681)21.1/60:0.0092692257,WP_014977514.1-Alteromonas-macleodii:0.0455118539)0/59:0.0000021352,WP_239220085.1-Alteromonas-sp.-MmMcT2-2:0.0350436690)83.3/66:0.0087689676,(WP_063457834.1-Alteromonas-stellipolaris:0.0446480002,(WP_202940767.1-Alteromonas-macleodii:0.0294484597,WP_061095937.1-Alteromonas-macleodii:0.0558207381)82.2/63:0.0167650636)76.7/59:0.0083329164)0/57:0.0000027170,(WP_202262575.1-unclassified-Alteromonas:0.0660067138,WP_187447473.1-Alteromonas-sp.-BZK5:0.0271638500)0/81:0.0075224253)92.2/91:0.0400482354,WP_057795858.1-Alteromonas-stellipolaris:0.0463444499)80.3/66:0.0167207864,WP_163106912.1-Alteromonas-genovensis:0.0572745795)84.6/73:0.0238059824)0/60:0.0000021033,(WP_071816068.1-Alteromonas-sp.-V450:0.1244427417,WP_198816918.1-Alteromonas-sp.-IB21:0.0301051135)87.1/100:0.0284636796)95.3/78:0.0458491721,WP_163085646.1-Alteromonas-profundi:0.0540694072)70.1/68:0.0069742480,(WP_130638173.1-Alteromonas-sp.-KUL49:0.0773732978,WP_115592514.1-Alteromonas-aestuariivivens:0.0635246717)77.7/86:0.0194847396)91.9/99:0.0395592278,((WP_265616768.1-Alteromonas-sp.-ASW11-7:0.1346107671,WP_216028209.1-Alteromonas-sp.-C1M14:0.0706660698)74.1/46:0.0194359536,((WP_265607972.1-Alteromonas-sp.-ASW11-130:0.0970070159,WP_119499891.1-Alteromonas-sp.-RKMC-009:0.0715605044)63.7/81:0.0358417359,(WP_216026947.1-Aestuariibacter-sp.-A3R04:0.0824978861,(WP_070176406.1-Alteromonas-lipolytica:0.0477092466,WP_073325432.1-Marisediminitalea-aggregata:0.0660747182)93.6/100:0.0634096854)83.9/69:0.0271900280)65.4/48:0.0132886138)95.3/61:0.0512340723)70.6/55:0.0229372992,(WP_262993134.1-Alteromonas-sp.-ASW11-19:0.1010851641,((WP_218354448.1-Alteromonas-lipotrueiana:0.1173164950,WP_195811108.1-Salinimonas-marina:0.0802809733)75.8/92:0.0272368410,WP_018984642.1-Salinimonas-chungwhensis:0.1522580670)87.7/98:0.0355898303)88.1/57:0.0406257929)91.5/85:0.0559530956)89.9/81:0.0662497512)85.6/25:0.0480731480)99.3/100:0.1557014829)79.6/15:0.0435075026,(WP_250459773.1-Microbulbifer-sp.-GX-H0434:0.1599812219,WP_152451766.1-Microbulbifer-sp.-THAF38:0.2462717515)96.2/100:0.1356740534)82.3/93:0.0905320856,((((WP_262601261.1-Endozoicomonas-euniceicola:0.1392500518,WP_066015073.1-Endozoicomonas-atrinae:0.1909160810)95.9/100:0.1438294728,WP_257287521.1-Endozoicomonas-sp.-SESOKO2:0.1417391207)88.9/100:0.1170120533,(WP_067585802.1-Endozoicomonas-ascidiicola:0.1953310155,(WP_020580625.1-Endozoicomonas-elysicola:0.1270282365,WP_257293945.1-Endozoicomonas-sp.-YOMI1:0.1795196842)93.3/100:0.1043250811)99.4/100:0.3080466951)94.7/96:0.2016730930,WP_116686321.1-Pelagibaculum-spongiae:0.4677412044)98.7/100:0.3391103780)94.2/97:0.1645704054,((WP_047883806.1-Photobacterium-ganghwense:0.1955279221,WP_146157089.1-Photobacterium-rosenbergii:0.2736873211)89.4/30:0.1200403684,(WP_187463805.1-Photobacterium-sp.-BZF1:0.0810595720,WP_039467548.1-Photobacterium-gaetbulicola:0.2098623455)3.7/75:0.0419152304)67.1/24:0.0884761458)1.1/6:0.0338187993,((WP_188026786.1-Tolumonas-osonensis:0.0332640876,WP_024870911.1-Tolumonas-lignilytica:0.1766425999)88.4/100:0.0935261391,WP_105076797.1-Pantoea-ananatis:0.6427703960)54.2/39:0.0559976572)92.1/13:0.0860827919,((((WP_007466453.1-Photobacterium-marinum:0.3615023049,WP_187508308.1-Pantoea-agglomerans:0.2218518060)55.7/55:0.0779826585,((WP_214216788.1-Rosenbergiella-epipactidis:0.0907668651,WP_241623482.1-Rosenbergiella-epipactidis:0.0238811192)99/100:0.1919806464,(WP_192411534.1-Pantoea-sp.-PNT02:0.0861064960,((WP_125311258.1-Pantoea-dispersa:0.1094423674,((WP_150019269.1-unclassified-Pantoea:0.0962649364,WP_140917215.1-Pantoea-deleyi:0.0568663534)38.7/94:0.0082164277,WP_069025179.1-Pantoea-agglomerans:0.0095945695)96/97:0.0485295967)85.6/89:0.0348451825,WP_097116658.1-Pantoea-sp.-GL120224-02:0.0010234949)87.3/89:0.0746019638)96.7/100:0.1608116184)92.3/98:0.1048548072)72.9/29:0.0209816054,(WP_269294270.1-Pantoea-sp.-LMR881:0.0666932742,WP_215846277.1-Pantoea-sp.-EnvD:0.1043054378)94.9/100:0.0913296069)69.5/28:0.0241587705,((WP_144135068.1-Kosakonia-radicincitans:0.4574364948,((WP_088208149.1-Enterobacter-cancerogenus:0.1525463303,WP_121932958.1-Enterobacter-hormaechei:0.1350223797)86/100:0.0645740608,WP_023330866.1-Enterobacter:0.0591981281)93/100:0.1359026690)89/51:0.1080645921,WP_120453320.1-Kalamiella-piersonii:0.1444703962)51.8/11:0.0357981619)97.2/94:0.1128596505)92.1/23:0.0730835288,((WP_120356561.1-Alginatibacterium-sediminis:0.5506360500,((WP_075989065.1-Vibrio-fluvialis:0.0516251378,WP_044366945.1-Vibrio-fluvialis:0.0517160341)34.5/32:0.0293823170,(WP_076657081.1-unclassified-Vibrio:0.0926282761,WP_239929047.1-Vibrio-kanaloae:0.0483783344)98.5/100:0.1511130582)86.1/95:0.0917054619)32.6/32:0.0861123956,WP_182744595.1-Shewanella-sp.-SR44-4:0.4336972219)65/28:0.0359557768)79/27:0.0313292404,((WP_020431327.1-Vibrio-fluvialis:0.1577974184,WP_055453311.1-Vibrio-fluvialis:0.0635025976)85.8/100:0.0581243887,(WP_114765998.1-Vibrio-rhodolitus:0.1895371145,WP_182108830.1-Vibrio-marinisediminis:0.1070546540)94.2/100:0.1472619932)30.2/84:0.0586273202)94.7/25:0.0791438951,(((WP_211647400.1-Photobacterium-sp.-GJ3:0.0508461653,WP_120511428.1-Photobacterium-salinisoli:0.0784954971)93.9/100:0.1022449714,(((WP_200225058.1-Rahnella-aceris:0.1628847239,WP_195814554.1-Rahnella:0.1508998153)88.5/100:0.0661855919,WP_169403689.1-Rouxiella-aceris:0.0946998021)99.5/100:0.2210473679,((WP_101201417.1-Paraglaciecola-sp.-MB-3u-78:0.3104790606,WP_101064017.1-Moritella-sp.-Urea-trap-13:0.2972590449)41/90:0.0607623565,WP_172533652.1-Vibrio-aestuarianus:0.4421699817)76.3/67:0.0689263067)0/12:0.0000021333)95.3/99:0.0954401567,((((WP_170899585.1-Vibrio-sp.-3-2-1:0.1594625256,WP_017221010.1-Moritella-dasanensis:0.1840135260)87.2/100:0.0559571842,WP_107284330.1-Photobacterium-lipolyticum:0.1581917062)93.3/100:0.0878835756,WP_159197271.1-Escherichia-coli:0.1528182270)82.8/99:0.0482370987,(WP_124938567.1-Vibrio-viridaestus:0.2280870801,WP_088133934.1-Vibrio-gazogenes:0.3434559819)58.9/55:0.0333624430)52.4/38:0.0400113457)75.1/43:0.0246089716)0/17:0.0000021487,((((((WP_239941760.1-Vibrio-mediterranei:0.1565716575,(WP_252012797.1-Vibrio-sp.-SCSIO-43140:0.1837825260,WP_252012765.1-Vibrio-sp.-SCSIO-43140:0.1766428725)77.4/97:0.0348748236)0/48:0.0280177074,((WP_062462511.1-Vibrio-mediterranei:0.1406246158,WP_171392669.1-Vibrio:0.1860291412)71.3/73:0.0443001404,(WP_102532726.1-Vibrio-sp.-10N.286.48.B7:0.1197127647,WP_250617847.1-Vibrio-sp.-S4B1:0.1201165338)90.3/100:0.0934986731)40.7/38:0.0369106372)87.5/98:0.0917072251,(WP_221070344.1-Vibrio-alfacsensis:0.1487401578,WP_011189464.1-Desulfotalea-psychrophila:0.6441600341)74.8/54:0.0860147176)79/48:0.0517917497,((WP_050928752.1-Yersinia-enterocolitica:0.3971184788,WP_012883389.1-Dickeya-parazeae:0.3436733599)32.5/85:0.0592775393,((WP_035339894.1-Dickeya:0.0335127280,WP_125258312.1-Dickeya-lacustris:0.1178802246)99.7/100:0.3022320432,WP_042868619.1-Dickeya:0.1135762565)93.3/100:0.1358579459)95.5/99:0.1513988925)69.8/22:0.0286825701,WP_102533425.1-Vibrio-breoganii:0.2176301734)74.1/41:0.0242224163,WP_193290728.1-Vibrio-parahaemolyticus:0.4432386185)72.6/21:0.0180691083)86.5/25:0.0345706039,(WP_261894433.1-Vibrio-porteresiae:0.2193468791,WP_234495031.1-Vibrio-maritimus:0.3197831579)84.7/39:0.1016072583)74.6/18:0.0143275955,(((WP_038175370.1-Vibrio-pacinii:0.2157111304,((WP_102941572.1-Vibrio-diazotrophicus:0.1733928398,WP_193262143.1-Vibrio-parahaemolyticus:0.4053366937)81.7/99:0.0413490039,WP_171749982.1-Vibrio-sp.-RE88:0.2949580284)77.7/80:0.0357702210)0/71:0.0037389019,(WP_075764060.1-Photobacterium-proteolyticum:0.1975395915,WP_239411815.1-Photobacterium-sp.-OFAV2-7:0.2391054261)92.4/32:0.1011004173)84.7/27:0.0604247960,((WP_190336189.1-Vibrio-sp.-S9_S30:0.0941871785,WP_022612811.1-Vibrio-nigripulchritudo:0.1931049216)92.6/100:0.0865270592,(((WP_102481006.1-Vibrio-breoganii:0.2153573785,(((WP_258489575.1-Vibrio-alginolyticus:0.1787315059,WP_152797327.1-Salinivibrio-sp.-VYel1:0.3991523705)42/80:0.0542561202,(WP_261874894.1-Vibrio-rarus:0.2347472551,((WP_045031875.1-Photobacterium-phosphoreum:0.0381636329,WP_107213458.1-Photobacterium-phosphoreum:0.0451381452)98.9/100:0.1839976390,WP_138522257.1-Pseudoalteromonas-galatheae:0.4690007733)75.4/56:0.0249581070)84.3/56:0.0664403482)79.8/46:0.0607367238,(WP_017031451.1-Vibrio-breoganii:0.0738747231,WP_248975945.1-Shewanella-kaireitica:0.2780798082)90.4/54:0.0802743103)74.4/84:0.0312962674)79.4/95:0.0395784136,((((((WP_011638275.1-Shewanella-frigidimarina:0.1502284138,WP_223418564.1-Shewanella-inventionis:0.2843005382)73.4/100:0.0478747983,(WP_220774251.1-Shewanella-glacialipiscicola:0.0830509461,WP_182751750.1-Shewanella-sp.-SG44-2:0.1906536062)94.4/100:0.1161501914)88.3/100:0.0783770004,(WP_197173106.1-Pseudoalteromonas-sp.-NZS11_1:0.1983687851,WP_182748079.1-Pseudoalteromonas-sp.-SR41-1:0.2650192683)90.3/99:0.0887868466)83.2/98:0.0364670734,(((WP_069589493.1-Salinivibrio-sp.-DV:0.1215517063,WP_077662143.1-Salinivibrio-proteolyticus:0.0208965290)80.3/100:0.0320681669,((WP_077659434.1-Salinivibrio-kushneri:0.1783760118,WP_077659452.1-Salinivibrio-kushneri:0.0445331869)74.1/100:0.0341464021,WP_269578450.1-Salinivibrio-kushneri:0.1149571481)86.8/100:0.0553718684)100/100:0.3692287271,(WP_041156219.1-Vibrio-mytili:0.1516889015,(WP_047480300.1-Vibrio-campbellii:0.0853285609,WP_045494469.1-Vibrio-hyugaensis:0.0535123255)99.3/100:0.2118960082)98.2/99:0.2137851928)75.7/30:0.0295390299)49.7/29:0.0137405383,(WP_166378370.1-Pseudoalteromonas-sp.-Z9A4:0.2310670072,WP_012549058.1-Aliivibrio-salmonicida:0.2641471033)80.3/97:0.0385538304)77.7/29:0.0469402046,WP_005395829.1-Vibrio-diabolicus:0.3643530004)0/29:0.0064707777)76.5/30:0.0370203428,(((((WP_261298106.1-Shewanella-sp.-SP1S2-7:0.1332077083,WP_167678467.1-Shewanella-aestuarii:0.1276022014)99.9/100:0.3768106213,WP_248994245.1-Shewanella-gaetbuli:0.3098658852)88.5/100:0.1492091207,WP_212599444.1-Shewanella-sp.-M16:0.3389803251)93.6/99:0.1503435076,WP_261923504.1-Shewanella-sp.-NFH-SH190041:0.2392046413)90.1/99:0.1147238850,WP_102523418.1-Vibrio-tapetis:0.2292642523)92.7/30:0.1021014286)77.5/29:0.0344661935)79.7/22:0.0271959336)83.5/26:0.0266851482)37.5/59:0.0390674368,WP_014230611.1-Vibrio-sp.-EJY3:0.2545773333)63.9/11:0.0244841345,((((((((((((WP_102457123.1-Vibrio-breoganii:0.1777449972,WP_077647682.1-Salinivibrio-sp.-IB643:0.2257137094)79.7/99:0.0364846418,((((((WP_061023752.1-Vibrio-splendidus:0.0987035685,WP_220768829.1-Shewanella-sp.-MBTL60-007:0.1895599628)77.9/100:0.0717137161,(WP_248991952.1-Shewanella-profunda:0.1896854523,WP_110456309.1-Shewanella-algidipiscicola:0.2406025194)33.2/21:0.0684849744)68.9/11:0.0276722131,(WP_182754472.1-Shewanella-sp.-SG41-3:0.2037538797,WP_059744187.1-Shewanella-frigidimarina:0.3081660562)64.1/13:0.0544395427)68/47:0.0237570256,WP_204999274.1-Parashewanella-hymeniacidonis:0.2986472093)51.1/99:0.0229490054,WP_222242368.1-Pseudoalteromonas-piscicida:0.5931296649)71.3/70:0.0326662379,(((WP_243025268.1-Vibrio-cincinnatiensis:0.7721452216,WP_047110243.1-Vibrio-vulnificus:0.3464516489)71.8/99:0.0658062741,WP_102313365.1-Vibrio-cyclitrophicus:0.2204779189)88.8/98:0.0918909684,WP_031827242.1-Vibrio-parahaemolyticus:0.4637913623)89.5/69:0.0936052284)73.9/93:0.0284667300)91.2/95:0.0613122567,(((WP_017020702.1-Aliivibrio-logei:0.2192712015,WP_145570128.1-Vibrio-tasmaniensis:0.1926195108)61.4/86:0.0395586654,(WP_005597034.1-Vibrio-scophthalmi:0.2410575413,((WP_108176621.1-Vibrio-splendidus:0.0590660613,WP_239918548.1-Vibrio-sp.-Isolate22:0.0473610344)80.1/100:0.0452526220,(WP_243579549.1-Vibrio-splendidus:0.3412024584,WP_135385147.1-Vibrio-tasmaniensis:0.2327242757)71.6/99:0.0448397430)55.9/98:0.0419202517)79.5/87:0.0349573705)85/100:0.0584750081,WP_238130349.1-Vibrio-cincinnatiensis:0.2613614176)51.7/95:0.0246772395)87.3/93:0.0308155261,(WP_171343915.1-Vibrio-cyclitrophicus:0.1163218918,((WP_048661380.1-Vibrio-crassostreae:0.1153530119,WP_194091044.1-Vibrio-hibernica:0.3846982070)0/77:0.0000020057,WP_193846925.1-Vibrio-sp.-OPT10:0.1336481199)86.8/93:0.0632267261)82.2/83:0.0361428308)78/76:0.0127419514,(WP_161753058.1-Vibrio-toranzoniae:0.1949624686,(WP_102248986.1-Vibrio-tasmaniensis:0.2645853210,WP_135445135.1-Vibrio-tasmaniensis:0.1129295364)97.2/100:0.1369745130)87.1/99:0.0544646757)74.1/83:0.0158197483,(((WP_268674123.1-Vibrio-scophthalmi:0.1289949419,WP_102475754.1-Shewanella-sp.-10N.286.52.C2:0.3052619472)81.8/96:0.0506188944,(WP_012551417.1-Aliivibrio-salmonicida:0.0810724451,WP_012276595.1-Shewanella-halifaxensis:0.1360806444)97.8/100:0.1460508168)86.3/81:0.0724672915,(((((WP_102420589.1-Vibrio-cyclitrophicus:0.1379402334,WP_017085924.1-Vibrio-splendidus:0.2308416275)72.4/79:0.0264238417,WP_132940416.1-Vibrio-crassostreae:0.1936748450)12.7/58:0.0608990962,WP_102457030.1-Vibrio-cyclitrophicus:0.1726925283)76.3/77:0.0384392139,WP_102459739.1-Vibrio-lentus:0.2148722468)68.2/67:0.0310674007,WP_102554090.1-Vibrio-splendidus:0.1744294316)91.2/66:0.0736741336)82/62:0.0398974343)75.5/58:0.0154897374,(((WP_063656223.1-Aliivibrio-fischeri:0.1662842218,((WP_155675904.1-Aliivibrio-fischeri:0.1223285589,WP_130314193.1-Vibrio-vulnificus:0.2386991967)74/81:0.0277640306,WP_017105169.1-Vibrio:0.1993394535)87.6/82:0.0502374223)84.8/68:0.0413363658,((WP_137008959.1-Vibrio-kanaloae:0.1874993742,WP_135353160.1-Aliivibrio-fischeri:0.1168181435)88.7/80:0.0773948103,WP_248638627.1-Vibrio-crassostreae:0.2183606147)23.1/22:0.0219855094)0/17:0.0000028449,(WP_076646094.1-Vibrio-sp.-10N.222.47.A9:0.1415667250,WP_102363333.1-Vibrio-cyclitrophicus:0.1716254613)95.7/93:0.1285086824)86.6/64:0.0350119107)92.4/80:0.0587608139,((((WP_059021773.1-Vibrio-coralliirubri:0.0836187881,WP_135380937.1-Vibrio-tasmaniensis:0.1393682225)80.8/99:0.0407427957,WP_102276040.1-Vibrio-lentus:0.1713138346)70.4/88:0.0207384069,((WP_102571993.1-Vibrio:0.2004666878,WP_248388778.1-Vibrio-cyclitrophicus:0.2124496789)73.4/17:0.0295511888,(WP_102298256.1-Vibrio-cyclitrophicus:0.1526798210,WP_265133188.1-Vibrio-splendidus:0.1384624742)53.1/99:0.0389433126)73.3/16:0.0342877966)73.3/15:0.0346568356,((((((WP_155682312.1-Aliivibrio-fischeri:0.1316578227,WP_012532661.1-Aliivibrio-fischeri:0.1731001184)56.1/73:0.0300909116,(((WP_108162474.1-Vibrio-splendidus:0.0433314449,WP_004742905.1-Vibrio-tubiashii:0.0709391556)99.2/100:0.1492366897,(((WP_102503160.1-Vibrio-splendidus:0.1863537497,WP_155677763.1-Aliivibrio-fischeri:0.1805432939)86.7/76:0.0887629300,WP_016768872.1-Vibrio-cyclitrophicus:0.1786816616)74.1/71:0.0380624324,WP_017069285.1-Vibrio-crassostreae:0.1150983016)75.3/71:0.0645454210)85.3/84:0.0757014221,(WP_258428294.1-Vibrio-sp.-RM-69-4:0.1526058213,WP_102428030.1-Vibrio-sp.-10N.261.54.E10:0.2170456516)68.3/69:0.0367578691)79.2/21:0.0434656170)89.4/30:0.0689513557,(WP_193834777.1-Vibrio-sp.-OPT20:0.0530909026,(WP_029865567.1-Vibrio-parahaemolyticus:0.0757846368,WP_213967907.1-Vibrio-alginolyticus:0.0574734536)98.5/100:0.1329533179)97/100:0.1304902665)37.3/43:0.0138285780,((WP_130049361.1-Aliivibrio-finisterrensis:0.1152688131,WP_019826652.1-Vibrio-splendidus:0.1982253532)52.7/96:0.0382950098,WP_260256886.1-Vibrio-sp.-NC2:0.1673968908)76.1/91:0.0296209027)86.6/93:0.0508774839,(WP_017063118.1-Vibrio:0.1816742851,(WP_004733311.1-Vibrio-splendidus:0.1516285292,((WP_009843931.1-Vibrio-sp.-AND4:0.1406438417,(WP_102353859.1-Vibrio-cyclitrophicus:0.1755366748,WP_171315631.1-Vibrio-splendidus:0.1680227387)85.8/100:0.0771668643)65.7/20:0.0231157040,(WP_102285079.1-Vibrio-lentus:0.1395343104,WP_235060641.1-Vibrio-sp.-SS-MA-C1-2:0.3398716046)85.3/97:0.0641297877)71.4/20:0.0225550007)0/95:0.0026298939)84.4/79:0.0475264962)44.5/75:0.0347445768,WP_182796407.1-Vibrio-sp.-SG41-7:0.2657275795)76.7/11:0.0218556249)74.4/79:0.0614653850)52.6/69:0.0072972945,((((((((WP_017027407.1-Vibrio-breoganii:0.2069080660,WP_102443327.1-Vibrio-breoganii:0.1301787696)22.9/90:0.0300294864,WP_172380549.1-Vibrio-sp.-Vb339:0.1986016793)87/91:0.0722685636,WP_108189654.1-Vibrio-splendidus:0.2100747225)62.7/79:0.0282875090,WP_063660043.1-Aliivibrio-fischeri:0.2032532124)75.5/76:0.0612766542,WP_203433235.1-Shewanella-sp.-LZH-2:0.3344724320)0/41:0.0000025668,((WP_065600047.1-Aliivibrio-sp.-1S128:0.1667663293,WP_155682326.1-Aliivibrio-fischeri:0.1002402794)91.3/99:0.0843554973,(WP_102385704.1-Vibrio-cyclitrophicus:0.1610055512,WP_239937663.1-Vibrio-chagasii:0.2145940950)77/82:0.0832209455)0/67:0.0000020306)84.7/73:0.0336915177,((WP_102482947.1-Vibrio-splendidus:0.1363035973,WP_102394203.1-Vibrio-sp.-10N.222.54.B6:0.1848816268)79.2/98:0.0544658790,(((WP_138589904.1-Pseudoalteromonas-aurantia:0.2434473140,(WP_218529053.1-Shewanella-sp.-NIFS-20-20:0.1640674059,WP_225483056.1-Vibrio-vulnificus:0.1157375144)87.1/100:0.0718222466)59.6/85:0.0821211105,WP_016784063.1-Vibrio:0.1424802960)54.8/82:0.0531376125,WP_017449116.1-Vibrio-parahaemolyticus:0.2850935463)28.8/28:0.0341924785)78.6/28:0.0295160732)75.8/32:0.0165662836,(((WP_135455718.1-Vibrio-echinoideorum:0.1978220731,((((WP_161672404.1-Vibrio-toranzoniae:0.0841989046,WP_017040654.1-Vibrio-genomosp.-F10:0.2741074484)80/91:0.0441693890,((WP_162023628.1-Shewanella-sp.-Arc9-LZ:0.1164097470,WP_248997855.1-Shewanella-marinintestina:0.0583491027)98.2/100:0.1412578863,WP_017094106.1-Vibrio-splendidus:0.1396230340)69/92:0.0309136240)86.4/98:0.0539564251,WP_137025093.1-Vibrio-kanaloae:0.2168969227)47.9/31:0.0423885701,(WP_137027437.1-Vibrio-kanaloae:0.0439256778,WP_108117598.1-Vibrio-splendidus:0.0950104273)92.2/100:0.0619140207)0/16:0.0110375293)72.6/31:0.0492921116,WP_077679840.1-Vibrio-kanaloae:0.1907761696)24.7/78:0.0157128064,(WP_140342839.1-Vibrio-parahaemolyticus:0.2720516170,WP_135445143.1-Vibrio-tasmaniensis:0.2427077384)85.6/39:0.0711928736)75.9/32:0.0115214600)88.8/81:0.0298635298)82.3/86:0.0268744535,(WP_243584123.1-Vibrio-splendidus:0.1943601311,(((WP_102462022.1-Vibrio-splendidus:0.0867756592,WP_187463684.1-Photobacterium-sp.-BZF1:0.0906847595)91/100:0.1068104019,WP_259574191.1-Shewanella-baltica:0.4042629242)58.6/81:0.0529103784,WP_029788868.1-Vibrio:0.3906229006)58.6/39:0.0552461566)91.4/97:0.1155398127)50.4/12:0.0121357453,WP_087845049.1-Photobacterium-malacitanum:0.3362974611)74.6/10:0.0095115471,(((((((((((((WP_193255073.1-Vibrio-navarrensis:0.0412082457,WP_258469755.1-Vibrio-alginolyticus:0.1613999390)96.3/100:0.1269412420,((WP_009847952.1-Vibrio:0.2982743232,WP_233894267.1-Vibrio-kanaloae:0.1958018045)88.1/100:0.1071324574,(WP_048661352.1-Vibrio-crassostreae:0.0873049356,WP_114634227.1-Vibrio-splendidus:0.0902468294)67.5/100:0.0671063865)96.9/100:0.1554994288)5.7/60:0.0219792571,((((((WP_041510674.1-Shewanella:0.0305592111,(((WP_045569365.1-Vibrio:0.1083510031,WP_039437343.1-Vibrio-navarrensis:0.0558738440)95.2/100:0.1297505358,WP_176299354.1-Vibrio-sp.-05-20-BW147:0.2131573088)90.1/100:0.1212303298,(WP_140233673.1-Shewanella-polaris:0.3208400979,WP_192022372.1-Shewanella-sp.-WPAGA9:0.2977933556)46.9/100:0.0616646087)97.2/100:0.2070871474)19.8/98:0.0240050547,WP_187575439.1-Shewanella-algae:0.1082030464)94.1/99:0.0908685886,(WP_208158397.1-Shewanella-algae:0.1110006544,WP_208157589.1-Shewanella-algae:0.0590963641)81.2/100:0.0410488131)86.1/99:0.0756433303,WP_045460498.1-Vibrio-hyugaensis:0.2523086663)80.6/99:0.0577313873,WP_220051788.1-Shewanella-sp.-FJAT-51860:0.2025576864)75/94:0.0289517063,WP_239850080.1-Shewanella-sp.-Isolate8:0.1500816060)75.4/70:0.0520171094)70.4/57:0.0320542573,WP_118894008.1-Vibrio-vulnificus:0.2223341341)76.8/28:0.0322500678,(WP_239872404.1-Shewanella-insulae:0.2092594674,WP_238097025.1-Vibrio-cincinnatiensis:0.2157715312)12.9/49:0.0399560414)79.6/71:0.0239114538,(WP_106651046.1-Shewanella-sp.-Pdp11:0.2011427763,WP_206625521.1-Vibrio-vulnificus:0.2422704938)77.5/88:0.0700262701)86.6/70:0.0515293341,WP_102986644.1-Vibrio-diazotrophicus:0.2142707483)70.1/43:0.0388786696,((WP_102563398.1-Vibrio-sp.-10N.286.49.B3:0.1262587719,WP_120030433.1-Vibrio-sinensis:0.1659965552)62.5/92:0.0533040153,(((WP_011497090.1-Shewanella-denitrificans:0.4510223999,(WP_259522409.1-Shewanella-baltica:0.1602328420,WP_173243873.1-Shewanella-sp.-DC2-4:0.2040519719)88/100:0.0803522149)86.2/93:0.0617857087,(WP_011716450.1-Shewanella-sp.-ANA-3:0.4670115422,WP_069440067.1-Pseudoalteromonas-tetraodonis:0.3718450293)72.8/70:0.0429462504)61.5/54:0.0302859560,WP_130066657.1-Aliivibrio-finisterrensis:0.2895887197)70.2/39:0.0353287791)86.1/48:0.0587057058)77.5/80:0.0317102232,((WP_238813939.1-Vibrio-vulnificus:0.1499490539,(((((WP_108083222.1-Vibrio-sp.-ZF-223:0.1740287209,(WP_102532836.1-Vibrio-sp.-10N.286.48.B7:0.1707961317,WP_254101081.1-Alteromonas-sp.-LMIT007:0.1841582097)93.8/100:0.0846336717)92.1/97:0.0762186154,(WP_171342182.1-Vibrio-cyclitrophicus:0.2040991039,WP_161972844.1-Enterovibrio-baiacu:0.2119263097)76.5/98:0.0527273753)26.4/95:0.0619189350,WP_121852788.1-Parashewanella-spongiae:0.3546112129)71/97:0.0300058290,((WP_103194250.1-Vibrio-vulnificus:0.0622587130,WP_158146950.1-Vibrio-fluvialis:0.1654633741)87.2/100:0.0615660846,WP_233927909.1-Vibrio-fluvialis:0.2442310286)83.8/100:0.0655126720)77.1/96:0.0343403313,WP_112353352.1-Shewanella-benthica:0.3594871812)82.2/98:0.0419708124)82.1/98:0.0549545081,WP_039841338.1-Vibrio-harveyi-group:0.2763649669)68.2/92:0.0527360449)83.2/58:0.0403103197,(((((WP_233983986.1-Vibrio-fluvialis:0.0794527203,WP_099031861.1-Pseudoalteromonas-sp.-GCY:0.2239649731)19.5/57:0.0295396408,WP_144217665.1-Shewanella-algae:0.3195535481)78.7/94:0.0359289951,(((WP_208844179.1-Pseudoalteromonas-xiamenensis:0.1866826407,WP_072616325.1-Vibrio-vulnificus:0.2902383510)82.4/95:0.0681364564,WP_233976434.1-Vibrio:0.2181809007)81.7/92:0.0348696512,WP_229630939.1-Vibrio-coralliilyticus:0.3206564745)75.8/88:0.0204305060)83.5/28:0.0261346063,(((WP_158107962.1-Vibrio-furnissii:0.1680531445,WP_185890410.1-Vibrio-vulnificus:0.2325319521)67.7/72:0.0579832891,WP_225486226.1-Vibrio-vulnificus:0.2586226666)14.7/14:0.0437270597,(WP_143680610.1-Vibrio-furnissii:0.1545026659,(WP_086585081.1-Vibrio-parahaemolyticus:0.0406725158,WP_217557493.1-Vibrio-metschnikovii:0.0790513431)98.4/92:0.1213484072)54.2/36:0.0248194397)73.3/20:0.0148102925)74.9/39:0.0265576937,WP_069533349.1-Vibrio-parahaemolyticus:0.2590510347)82.1/49:0.0314725817)74.3/8:0.0105574000,((((WP_211915530.1-Vibrio-sp.-B1REV9:0.1568891302,WP_017420706.1-Vibrio-vulnificus:0.1538080829)81/68:0.0398400920,(WP_165312521.1-Vibrio-ziniensis:0.1752727806,(WP_192928291.1-Vibrio-vulnificus:0.0383906900,WP_132698543.1-Vibrio-harveyi-group:0.1679977174)96.9/100:0.1260840673)73.4/14:0.0727300506)0/12:0.0000028359,(WP_213972370.1-Vibrio-alginolyticus:0.2645166968,WP_017024716.1-Vibrio-rumoiensis:0.2241712292)87.2/97:0.0890033486)79.2/14:0.0363982080,((WP_068331709.1-Photobacterium-jeanii:0.2573002656,(WP_031414678.1-Vibrio-parahaemolyticus:0.1635191836,WP_248468589.1-Shewanella-sp.-1CM18E:0.2665363886)75.3/98:0.0875139157)81.2/88:0.0807887306,WP_206974914.1-Vibrio-parahaemolyticus:0.2326988492)19.1/14:0.0164599800)77.5/14:0.0235377819)0/9:0.0000028361,((WP_172561824.1-Vibrio-furnissii:0.1597767444,WP_023404414.1-Vibrio-halioticoli:0.3603318851)72.8/93:0.0299067655,(((WP_217542775.1-Vibrio-metschnikovii:0.0644406106,WP_113593871.1-Vibrio:0.1350446535)89.4/100:0.0783247344,WP_020328057.1-Vibrio-fluvialis:0.1824665557)77.6/83:0.0549592439,WP_238819535.1-Vibrio-furnissii:0.1458855790)65.7/31:0.0228256049)93.4/32:0.0821381723)93.8/41:0.0460445085,((((((WP_046223474.1-Vibrio-sp.-ECSMB14106:0.1423720938,WP_102562373.1-Vibrio-breoganii:0.1886261485)81.1/98:0.0480834502,((((WP_102546929.1-Vibrio-breoganii:0.2031663065,WP_102329601.1-Vibrio-breoganii:0.3646336327)54.8/86:0.0528427596,WP_102461639.1-Vibrio-breoganii:0.1836605188)79.8/86:0.0597142072,(WP_141346804.1-Vibrio-inusitatus:0.2521052795,WP_102282524.1-Vibrio-cyclitrophicus:0.2337798185)80.5/92:0.0612064447)56.4/92:0.0204170307,WP_135455667.1-Vibrio-echinoideorum:0.1700120662)94.8/97:0.0959970199)67.3/97:0.0183512515,WP_218382835.1-Photobacterium-sp.-WH24:0.2425538032)93.3/96:0.0812238254,((WP_020332409.1-Vibrio-fluvialis:0.0960350557,WP_096629284.1-Salinivibrio-sp.-YCSC6:0.2571463573)93.3/100:0.1103172840,(((WP_047463293.1-Vibrio-fluvialis:0.0786662020,WP_193167845.1-Vibrio-navarrensis:0.4936451515)73.6/99:0.0692760855,WP_148560072.1-Vibrio-mimicus:0.2540647712)83.7/95:0.0507645558,(WP_102446429.1-Vibrio-breoganii:0.2597857494,WP_193154481.1-Vibrio-navarrensis:0.2726397270)71.2/95:0.0447838841)74.6/91:0.0219211760)90.9/96:0.0670567473)83.3/86:0.0236695825,((WP_067416136.1-Enterovibrio-coralii:0.1473579457,WP_149593442.1-Pseudoalteromonas-sp.-FUC4:0.2729266941)79.7/100:0.0496320409,WP_102038478.1-Shewanella-sp.-GutCb:0.3648011872)85.2/98:0.0562895917)75.7/72:0.0166871442,((((WP_102455158.1-Vibrio-breoganii:0.2042491683,(WP_188011373.1-Vibrio-vulnificus:0.3132258457,WP_102451772.1-Vibrio-breoganii:0.2071981652)84.4/100:0.1211244472)50.8/100:0.0874415745,WP_065296243.1-Vibrio-natriegens:0.1774879542)86.2/88:0.0639402185,((WP_163929979.1-Paraferrimonas-sp.-SM1919:0.2435209269,(WP_213886549.1-Vibrio-alginolyticus:0.3403362739,WP_158173598.1-Vibrio-alginolyticus:0.0528703130)90.4/56:0.1266779836)87.9/56:0.0684131446,(WP_172855536.1-Vibrio-coralliilyticus:0.3128828181,(WP_234235026.1-Shewanella-sp.-AS16:0.2812595379,WP_088905011.1-Shewanella-marisflavi:0.3166794091)81.3/95:0.0605351253)83.9/93:0.0508257129)61.5/10:0.0089984310)95.9/42:0.1066720183,((WP_107253996.1-Photobacterium-indicum:0.2649328698,WP_102395415.1-Enterovibrio-norvegicus:0.1678733356)88.5/100:0.0857017313,WP_045069930.1-Photobacterium-leiognathi:0.2423943332)0/64:0.0184951561)66.1/10:0.0262245695)0/8:0.0000022940)89.5/13:0.0364538082)85.6/12:0.0265954063)82.5/49:0.0190840564,((WP_048899071.1-Photobacterium-swingsii:0.1794538853,((((WP_172851426.1-Vibrio-coralliilyticus:0.2282474177,WP_182742106.1-Shewanella-sp.-SG44-6:0.3710968968)86.2/100:0.0961708103,WP_108186198.1-Vibrio-splendidus:0.1719192914)91.4/100:0.0931498003,((WP_248368760.1-Vibrio-sp.-1CM8B:0.1349901595,WP_161680902.1-Proteus:0.5812150416)80.6/84:0.0949035522,(WP_046209502.1-Vibrio:0.2138017709,(WP_061039794.1-Vibrio-coralliirubri:0.0549334059,(WP_044365158.1-Vibrio-fluvialis:0.4745901736,WP_025523961.1-Vibrio-parahaemolyticus:0.2193992056)91.2/93:0.1217761096)88.1/89:0.1160504339)41.5/80:0.0337009832)42.9/64:0.0174698438)89.9/65:0.0860177110,(WP_139685731.1-Vibrio-tasmaniensis:0.2047962192,WP_248386338.1-Vibrio-sp.-ED004:0.2766047352)72.9/58:0.0283002335)80.9/57:0.0418080713)51.1/83:0.0889206709,WP_103155729.1-Vibrio-vulnificus:0.1506193658)83.8/51:0.0480473218)89.2/78:0.0375122372,((((((WP_232600401.1-Photobacterium-carnosum:0.2329166461,(WP_232603643.1-Photobacterium-carnosum:0.1843467067,WP_232615715.1-Photobacterium-carnosum:0.1100552763)41.9/99:0.0379810602)74.1/99:0.0353398935,WP_060997345.1-Photobacterium-aquimaris:0.1790834265)82.1/94:0.0446858741,WP_065173140.1-Photobacterium-kishitanii:0.2214975421)14.6/59:0.0480361605,WP_058428281.1-Pseudoalteromonas:0.3504538862)78.4/74:0.0552350556,(WP_080158029.1-Photobacterium-piscicola:0.3197186648,(WP_107295906.1-Photobacterium-phosphoreum:0.2729234262,WP_065194916.1-Photobacterium-phosphoreum:0.1915492674)79.4/93:0.0528620667)81.5/88:0.0474249190)87.1/71:0.0379310901,((((WP_237041393.1-Photobacterium-damselae:0.2159202770,WP_161129723.1-Photobacterium-lucens:0.2138670426)71.6/65:0.0256492427,WP_148560094.1-Vibrio-mimicus:0.2505022959)73/56:0.0598790459,((WP_267741605.1-Vibrio-cholerae:0.1753019210,(WP_172567459.1-Vibrio-fluvialis:0.1478992558,WP_199354891.1-Vibrio-cholerae:0.0453198943)96.2/100:0.1518026510)0/78:0.0449500636,WP_102137258.1-Vibrio-cholerae:0.1309169720)87.5/84:0.0894043083)84.8/94:0.0508885489,((WP_237042700.1-Photobacterium-damselae:0.1936194757,((WP_148519618.1-Vibrio-cholerae:0.0356058872,(WP_167570263.1-Aeromonas-veronii:0.0878862072,WP_000864158.1-Vibrio-metoecus:0.0461024763)92.9/100:0.0407455981)85.7/100:0.0242106088,WP_148520854.1-Vibrio-cholerae:0.0466686822)97.3/100:0.1096955439)79/35:0.0387284396,((((WP_249572008.1-Vibrio-cholerae:0.1516913777,(((WP_000864153.1-Vibrio-cholerae:0.0991939454,(WP_084988554.1-Vibrio:0.1834657134,WP_206634697.1-Vibrio-vulnificus:0.2738304449)93.4/100:0.1286376803)74.3/15:0.0255387086,(WP_176284769.1-Photobacterium-damselae:0.3186168948,WP_088132129.1-Vibrio-cholerae:0.1270847418)79.1/98:0.0688265741)0/6:0.0000026290,WP_152138288.1-Plesiomonas-shigelloides:0.2748682348)83.5/44:0.0517468703)0/14:0.0088075543,WP_045618469.1-Vibrio-vulnificus:0.2507221350)73.4/70:0.0175215124,WP_255122387.1-Vibrio-cholerae:0.1682254409)77.4/83:0.0220680573,WP_129120576.1-Veronia-nyctiphanis:0.3407521005)76.5/31:0.0137296482)87.4/41:0.0399981948)85.3/34:0.0233887618)0/11:0.0000027148)66.1/20:0.0100242796,((((((WP_155660100.1-Aliivibrio-fischeri:0.1437706303,WP_095457151.1-Vibrio:0.1368389362)86.1/100:0.0644572637,(WP_252020469.1-Vibrio-sp.-SCSIO-43136:0.1507173360,WP_243584106.1-Vibrio-splendidus:0.2415729737)81.5/100:0.0474107562)91.6/100:0.0826013756,WP_166696422.1-Providencia-rettgeri:0.1902460029)88.5/100:0.1069390009,((((WP_076542412.1-Shewanella-sp.-UCD-KL21:0.4296435930,WP_232581009.1-Photobacterium-phosphoreum:1.2473398977)82.4/69:0.2389090051,(WP_108452928.1-Enterobacter-cloacae:0.5399227214,(WP_133040538.1-Shewanella-fodinae:0.4013939064,WP_237309237.1-Vibrio:0.4130996619)21.4/74:0.0551019648)74.2/98:0.0424838885)83/52:0.0741281213,WP_047876260.1-Photobacterium-aphoticum:0.3012365566)0/15:0.0356533983,(WP_070968050.1-Vibrio-sonorensis:0.3038776269,WP_180567593.1-Oceanospirillales:0.2101904044)77.3/60:0.0638428091)0/4:0.0299937224)65.3/47:0.0528189485,WP_220058598.1-Shewanella-sp.-FJAT-53555:0.4731754341)82.1/21:0.0435352573,((((WP_095430409.1-Vibrio-vulnificus:0.0980818705,WP_114994956.1-Grimontia-hollisae:0.2796397459)93.5/100:0.1399413653,WP_017007824.1-Enterovibrio-norvegicus:0.2113667170)88.8/90:0.0973644596,WP_028024862.1-Enterovibrio-calviensis:0.3183637350)79.6/70:0.0516722090,((((((WP_033540599.1-Shewanella:0.2356053502,WP_031815865.1-Vibrio-parahaemolyticus:0.1889654163)18.6/98:0.0481845876,WP_258516089.1-Vibrio-alginolyticus:0.1806128484)29.7/79:0.0770920948,((WP_064489977.1-Vibrio-parahaemolyticus:0.0624039000,WP_201976964.1-Aeromonas-enteropelogenes:0.1761961066)94.4/100:0.1384441132,WP_224657569.1-Vibrio-vulnificus:0.2089021984)71.5/97:0.0726575992)83/90:0.0540735213,WP_193250619.1-Vibrio-parahaemolyticus:0.1992650589)88.9/99:0.0719362887,((((WP_232612305.1-Photobacterium-phosphoreum:0.5218666265,WP_136989485.1-Vibrio-sp.-F13:0.3634747416)88.5/100:0.1523445482,WP_138627230.1-Pseudoalteromonas-sp.-S4741:0.2319038294)84/77:0.0747410847,(WP_211678393.1-Moritella-sp.-36:0.3808454078,WP_269377025.1-Vibrio-sinaloensis:0.2768721815)79/75:0.0592727624)69.7/85:0.0570943215,WP_039484180.1-Vibrio-sinaloensis:0.3189227744)75.2/69:0.0638705320)0/34:0.0000024688,((WP_188234612.1-Vibrio-sp.-Y2-5:0.0746180969,WP_057556016.1-Vibrio-cholerae:0.0423295947)100/100:0.3259009844,(WP_123015137.1-Vibrio-zhugei:0.0464594117,WP_077313846.1-Vibrio-palustris:0.0453855464)99.3/100:0.2343299934)86/55:0.0795467167)80.1/38:0.0447731748)73.3/65:0.0271432824)86.3/16:0.0357893336)62.3/9:0.0164382757,(WP_017008437.1-Enterovibrio-norvegicus:0.1843704344,((WP_188008910.1-Grimontia-hollisae:0.2125801362,(WP_251878143.1-Grimontia-sp.-020920N:0.2590871356,WP_102448490.1-Vibrio-splendidus:0.2154812100)34.6/45:0.0424470478)81.7/82:0.0455061160,(WP_261835819.1-Vibrio-ishigakensis:0.1612522721,WP_017221048.1-Moritella-dasanensis:0.2387908497)96.7/100:0.1764209158)75/86:0.0327252494)91.4/89:0.0752648599)76.4/43:0.0168239958,(((((WP_230432956.1-Plesiomonas-shigelloides:0.1431985480,WP_222763364.1-Vibrio-fluvialis:0.1818784318)81.1/100:0.0431813026,((WP_013365656.1-_Enterobacter:0.3449566595,WP_214579582.1-Enterobacter-asburiae:0.1131700963)79.3/67:0.0565584026,((WP_258988101.1-Scandinavium-hiltneri:0.0923337032,WP_181620050.1-Enterobacter-sp.-RHBSTW-00994:0.0900558490)95.7/100:0.0892695194,WP_193129600.1-Klebsiella-aerogenes:0.0976942823)78.6/100:0.0483456382)99.9/100:0.2773134427)24.2/50:0.0074483078,(WP_171138527.1-Vibrio-plantisponsor:0.1135861052,WP_102966424.1-Vibrio:0.1142695721)98.9/100:0.1555593363)87.4/47:0.0533428921,(WP_117235674.1-Vibrio-maerlii:0.1754604819,(WP_103882085.1-Vibrio-hangzhouensis:0.0770308827,WP_265676699.1-Vibrio-sp.-ZSDZ65:0.1818176823)94.8/100:0.1002067998)85.2/96:0.0542687454)74.1/51:0.0276522301,WP_251878087.1-Grimontia-sp.-020920N:0.2111344473)43.1/33:0.0309930748)75.7/19:0.0251630698,(((((((WP_261600599.1-Photobacterium-sp.-TY1-4:0.1283986322,WP_268679810.1-Vibrio-barjaei:0.2030246896)81.6/64:0.0352639755,(((WP_112404745.1-Vibrio-diazotrophicus:0.0833757538,WP_202661182.1-Vibrio-fluvialis:0.0663595865)92.9/100:0.0858571377,WP_045898906.1-Enterobacteriaceae:0.2363167316)47.7/59:0.0236759347,(((WP_263100201.1-Kosakonia-sp.-ML.JS2a:0.0437835667,WP_200550470.1-Kosakonia-sp.-LAM2021:0.1202786325)97.7/100:0.1594734839,((WP_015854950.1-Musicola-paradisiaca:0.0590205582,WP_192457620.1-Musicola-keenii:0.1062505118)100/100:0.3397860448,(WP_094109047.1-Lonsdalea-iberica:0.1495667712,(WP_026742149.1-Lonsdalea-quercina:0.0481293379,WP_094117837.1-Lonsdalea-britannica:0.0443977715)96.5/100:0.1024234675)79.3/64:0.0293405560)51.1/62:0.0556564137)92.6/100:0.1155641507,(WP_038178862.1-Vibrio-rhizosphaerae:0.1517897557,(WP_107187026.1-Photobacterium-kishitanii:0.2842426259,WP_046218796.1-Photobacterium-halotolerans:0.2827250286)20.6/60:0.0558848452)90.8/100:0.1084935441)71.8/60:0.0537251693)78/62:0.0310186620)91.1/98:0.0574539646,(WP_107294031.1-Photobacterium-phosphoreum:0.0593483309,WP_107276976.1-Photobacterium-sp.-NCIMB-13483:0.0833364983)99.5/100:0.1474501881)49.7/94:0.0302802861,((WP_094127815.1-Vibrio-sp.-V09_P4A23P171:0.0909256191,WP_199867560.1-Vibrio-anguillarum:0.1839121108)95.6/100:0.1318423233,WP_268809473.1-Vibrio-parahaemolyticus:0.2001647840)71.1/90:0.0655359338)87.3/76:0.0513156440,((((((WP_192015074.1-Photobacterium-arenosum:0.0963921489,WP_088876367.1-Vibrio-mediterranei:0.3997149678)83.2/98:0.0615439922,(WP_022590465.1-Vibrio-nigripulchritudo:0.2582188878,WP_045977433.1-Vibrio-neptunius:0.3192303484)88.7/98:0.1026639682)80.9/98:0.0354469906,WP_036979211.1-Providencia-alcalifaciens:0.3620031986)69.4/75:0.0392182213,WP_006646466.1-Photobacterium-sp.-SKA34:0.2488628867)87.3/97:0.0656629918,((WP_045039948.1-Photobacterium-iliopiscarium:0.2678097378,((WP_065176337.1-Photobacterium-aquimaris:0.0959660415,WP_107199726.1-Photobacterium-phosphoreum:0.1070498793)89.3/100:0.0580031391,WP_023931546.1-Photobacterium-leiognathi:0.1753753543)84.5/84:0.0628195357)83.1/98:0.0632871952,WP_068898547.1-Enterovibrio-pacificus:0.2248556093)90.7/75:0.0716004829)76.9/18:0.0200912458,WP_065208126.1-Photobacterium-phosphoreum:0.2261253873)74.5/16:0.0277533907)45.2/12:0.0151729474,((WP_208891742.1-Vibrio-anguillarum:0.2234656561,(WP_148533341.1-Vibrio-cholerae:0.1080102419,WP_107294203.1-Photobacterium-sp.-GB-50:0.2708285876)26.8/83:0.0173822892)91.3/64:0.0731091726,((((WP_000864156.1-Vibrio-cholerae:0.0945670235,WP_000864152.1-Vibrio-cholerae:0.1445410916)91.9/99:0.0807214636,WP_102786270.1-Vibrio-cholerae:0.0800432867)69.8/74:0.0524292756,(WP_000864154.1-Vibrio-cholerae:0.1392176157,WP_211689658.1-Moritella-sp.-28:0.5144959647)17.5/20:0.0106333481)77.1/26:0.0260973962,(((WP_175242188.1-Vibrio-cholerae:0.1378021969,WP_095464062.1-Vibrio-cholerae:0.1178577209)85.6/19:0.0568855322,WP_095478449.1-Vibrio:0.1084997295)60.3/16:0.0192332591,WP_149560528.1-Vibrio-cholerae:0.1530074495)83.1/82:0.0419181945)0/21:0.0000023546)91.3/36:0.0745025029)66.9/12:0.0134725390,(((((((((((WP_232575020.1-Photobacterium-carnosum:0.1643186088,WP_232600381.1-Photobacterium-carnosum:0.1443271746)55/91:0.0219460880,((WP_232585319.1-Photobacterium-iliopiscarium:0.1162752327,WP_107228425.1-Photobacterium-phosphoreum:0.3188574799)19/64:0.0216324754,(WP_232582982.1-Photobacterium-phosphoreum:0.0733423231,((WP_256383785.1-Photobacterium-toruni:0.0625683967,WP_232578536.1-Photobacterium-carnosum:0.0721520230)0/83:0.0000027835,WP_107228404.1-Photobacterium-phosphoreum:0.1266017698)93.1/100:0.0569195132)90.4/100:0.0692294552)91.4/69:0.0703031434)89.4/63:0.0601713164,(WP_045045404.1-Photobacterium-kishitanii:0.0555071310,(WP_045030671.1-Photobacterium-phosphoreum:0.0743651059,WP_107264793.1-Photobacterium-phosphoreum:0.0318707891)59.2/99:0.0272881216)91.1/100:0.0748083836)86.6/83:0.0595118559,WP_053987023.1-Photobacterium-leiognathi:0.1331867267)89.5/76:0.0550077486,((WP_151182432.1-Photobacterium-damselae:0.1595853705,WP_017004373.1-Enterovibrio-norvegicus:0.2958202498)55/69:0.0451074275,(WP_106261939.1-Photobacterium-damselae:0.1447033855,WP_115426223.1-Photobacterium-damselae:0.0963490201)70.5/77:0.0413893333)74.7/64:0.0304391415)82.1/46:0.0288189483,WP_086367281.1-Photobacterium-damselae:0.0545098557)91.7/61:0.0737254860,(((WP_238087945.1-Photobacterium-damselae:0.0771529466,(WP_065172005.1-Photobacterium-damselae:0.1309503484,WP_107185088.1-Photobacterium-leiognathi:0.1087811789)81.2/100:0.0302209186)73.7/99:0.0321800302,WP_151170663.1-Photobacterium-damselae:0.0804713252)70.7/98:0.0402581819,WP_045063675.1-Photobacterium-leiognathi:0.1322984369)0/55:0.0000022444)83.1/58:0.0410778599,WP_237041371.1-Photobacterium-damselae:0.1344739549)25.9/66:0.0224267055,WP_107293979.1-Photobacterium-phosphoreum:0.2391454648)84/75:0.0375185631,(WP_087853896.1-Photobacterium-andalusiense:0.0578344610,WP_107274737.1-Photobacterium-phosphoreum:0.1545309904)99.2/100:0.1473540672)0/74:0.0059249751,((WP_069212356.1-Vibrio-cholerae:0.0488949445,WP_202641945.1-Vibrio-fluvialis:0.1614726837)94.2/100:0.1064418736,(WP_199351799.1-Vibrio-cholerae:0.1163544032,(WP_107213932.1-Photobacterium-phosphoreum:0.1923356684,WP_065173166.1-Photobacterium-kishitanii:0.1939569199)84.6/81:0.0818063243)69.9/65:0.0653934525)88.8/28:0.0738399733)86.3/21:0.0512270251)87.6/25:0.0522601484)76.9/18:0.0359868968,(((WP_011771697.1-Psychromonas-ingrahamii:0.1149538996,WP_182690147.1-Psychromonas-sp.-SR45-3:0.2477274192)84.7/100:0.1163629928,WP_101040106.1-Psychromonas-sp.-MB-3u-54:0.3825484822)82.6/88:0.1352864451,WP_161119826.1-Photobacterium-lucens:0.1694274333)76.4/88:0.0608997717)99.5/83:0.2286198684,((WP_162556830.1-Proteus:0.1165652564,WP_161779122.1-Proteus-sp.-G2675:0.1232245781)99.3/100:0.3265256276,WP_236366479.1-Aeromonas-veronii:0.3450022967)29/25:0.0295047006)83.4/84:0.0519557114)88.6/96:0.0863079898,WP_102949229.1-Aeromonas-caviae:0.5196577501)65.8/49:0.0225228106,WP_225633057.1-Aeromonas-enteropelogenes:0.3429640479)76.7/53:0.0430933332,WP_097364011.1-Edwardsiella-tarda:0.5851066773)76/94:0.0464097953)94/89:0.0890397474,(WP_001100804.1-Enterobacteriaceae:0.2861205322,(WP_213789144.1-Enterobacter-hormaechei:0.0086844441,((WP_148242178.1-Enterobacter-asburiae:0.0693642257,WP_237994110.1-Enterobacter-sp.-DRP3:0.0259031448)34.4/99:0.0074898083,WP_237424876.1-Enterobacter-cloacae-complex-sp.-ECC445:0.0630449839)88.1/100:0.0635521554)99.9/100:0.3151359022)73.4/28:0.0990778376)73.5/23:0.0182908360,(WP_189493202.1-Formosimonas-limnophila:0.3785908173,WP_133620003.1-Hydromonas-duriensis:0.3680046118)99.6/100:0.3165033653)83.9/24:0.0413859524)93.9/27:0.1032642017,WP_100272167.1-Citrobacter-portucalensis:0.3355183142)74.2/26:0.0198242542,((WP_194857598.1-Citrobacter-freundii:0.5340852570,((WP_141032926.1-Escherichia-coli:0.1972660959,WP_196390204.1-Enterobacter-mori:0.0621769929)15.9/63:0.0418805945,(WP_252032207.1-Enterobacter-asburiae:0.0682390224,WP_111966211.1-Enterobacter-cloacae-complex:0.0382790164)81.6/86:0.0372524713)91/86:0.1890886065)99.5/100:0.3480444239,((WP_105274811.1-Escherichia:0.3250893993,WP_064512195.1-Buttiauxella-gaviniae:0.4331892846)88.6/100:0.1119402543,WP_172410167.1-Escherichia-coli:0.4482413437)90.1/91:0.1179796385)83.3/46:0.0749950196)82.3/21:0.0437602674,((((WP_108569539.1-unclassified-Pantoea:0.2991226039,WP_028018520.1-Enterobacter-cloacae-complex:1.0972999624)85.7/100:0.2372487850,WP_250341216.1-Pantoea-eucrina:0.2935729517)89.6/87:0.1822815527,((WP_234505032.1-Kluyvera-intermedia:0.6307076566,WP_230863267.1-Enterobacter-cloacae:0.7155112103)63.8/100:0.1232407651,(WP_029399168.1-Escherichia-coli:0.5053743798,((WP_034496584.1-Buttiauxella-agrestis:0.5736811009,WP_164716793.1-Cedecea-lapagei:0.5310975779)89.8/100:0.2037819651,WP_050862535.1-Enterobacter:0.6023190505)72.5/100:0.0372420386)88.4/100:0.1383730503)92/100:0.1313361164)86.8/87:0.0815795088,((((WP_159282060.1-Rahnella-variigena:0.5987020820,(WP_106902182.1-Citrobacter-freundii:0.2874730111,(WP_126291239.1-Enterobacter-hormaechei:0.2356441328,((((WP_062932819.1-Enterobacter-hormaechei:0.0458088374,WP_252042312.1-Enterobacter-chengduensis:0.0564129185)77.7/100:0.0361511468,WP_048999124.1-Enterobacter:0.0565949126)71.2/76:0.0091722191,WP_063144508.1-Enterobacter-cloacae:0.0533936221)85.6/75:0.0365423998,WP_049008289.1-Enterobacter-cloacae:0.0641579802)82.2/85:0.0621646173)96.2/98:0.1873951928)60.3/100:0.0972142189)80.2/98:0.0483651449,WP_035600513.1-Edwardsiella-tarda:0.4368968424)75.5/56:0.0778577781,((WP_202782010.1-Enterobacter-asburiae:0.1055860240,WP_144817260.1-Pseudescherichia-sp.-L3:0.0575989151)92.5/100:0.2073952509,WP_125117341.1-Aeromonas-caviae:0.5128904015)91/55:0.1340175699)31.7/55:0.0382580935,WP_269535377.1-Citrobacter-sedlakii:0.4461430105)0/24:0.0026220818)60.6/22:0.0201442179)79.1/24:0.0258574107,(((((((((((((WP_151711904.1-Enterobacter-ludwigii:0.0482693801,(WP_032941356.1-Citrobacter-freundii:0.3037219016,WP_058678686.1-Enterobacter:0.1139198384)90.9/100:0.0859767011)92.3/100:0.0953743888,WP_223560970.1-unclassified-Enterobacter:0.0000023259)98.5/100:0.4568824580,WP_044720521.1-Escherichia-coli:0.6264603596)99.6/100:0.5453410341,(WP_159241675.1-Proteus-terrae:0.3201846332,WP_195697765.1-Providencia-rettgeri:0.2649063549)0/13:0.0516514167)87.8/87:0.0726159150,WP_238563424.1-Providencia-rettgeri:0.1660885167)71.7/87:0.0234480957,WP_165880752.1-Providencia-rettgeri:0.2808333073)83.3/100:0.0476072485,WP_231068620.1-Providencia-huaxiensis:0.2244664280)86.3/97:0.0446603027,(((WP_169342312.1-Providencia-stuartii:0.2602394367,WP_164565644.1-Providencia-rettgeri:0.3394598601)57.3/58:0.0870598545,WP_004924376.1-Providencia-stuartii:0.2564771740)79.6/44:0.0684917204,((WP_187128479.1-Providencia-sp.-JUb39:0.2339516488,WP_193115284.1-Escherichia-coli:0.2514025671)84.3/100:0.0696008762,WP_154625743.1-Providencia-stuartii:0.2374751695)67/58:0.0521439202)0/51:0.0027639171)83.6/95:0.0812697230,WP_211007190.1-Pectobacterium-brasiliense:0.8114776337)87.6/98:0.1001020049,((((WP_090123257.1-Kosakonia-arachidis:0.1169163824,WP_020454293.1-Kosakonia-sp.-SOY2:0.1418094070)86.3/100:0.0577818621,WP_017458471.1-Kosakonia-sacchari:0.2138048477)72.5/100:0.0502598234,(WP_200002836.1-Citrobacter-freundii:0.2455229033,WP_080329661.1-Enterobacter-cloacae-complex:0.2042734882)99.3/100:0.2670711473)93.6/100:0.1668054402,WP_090460354.1-Enterobacter-sp.-kpr-6:0.0793132054)99.7/100:0.3505195472)74.9/78:0.0354823215,(((WP_257124601.1-Aeromonas-veronii:0.5767761929,WP_038258850.1-Xenorhabdus-bovienii:0.4119424936)79.7/59:0.0696230835,(WP_064374050.1-Pantoea:0.4752797231,(WP_036421240.1-Morganella-morganii:0.2416968700,(WP_264998074.1-Morganella-morganii:0.1117454863,WP_186849509.1-Morganella-morganii:0.1297831139)92/100:0.1185491547)93.6/100:0.1815449716)91.6/96:0.1195964051)56.9/27:0.0180616325,((WP_161748440.1-Proteus-sp.-G2658:0.3738849473,WP_049199273.1-Proteus-mirabilis:0.0518301830)86.8/100:0.0816081949,WP_157954980.1-Proteus-terrae:0.1024093545)100/100:0.2776820282)43.4/45:0.0261722808)71.7/46:0.0297423576,(WP_086113286.1-Xenorhabdus-beddingii:0.3704895024,WP_038223662.1-Xenorhabdus-bovienii:0.3555694523)22.1/46:0.0881222671)90.4/79:0.0603883916,(((WP_132257458.1-Paucimonas-lemoignei:0.1028179955,WP_119785517.1-Noviherbaspirillum-sedimenti:0.0813585857)100/100:0.7475117511,(WP_120162698.1-Rahnella-variigena:0.0805331982,WP_183847446.1-Rahnella-inusitata:0.0936920869)99.3/100:0.4262049059)23.4/22:0.0675389414,((WP_115348269.1-Hafnia-alvei:0.2140943133,WP_046458830.1-Obesumbacterium-proteus:0.4413880087)79.9/69:0.0629026128,(WP_130961887.1-Hafnia-paralvei:0.2823846642,WP_061059890.1-Hafnia-paralvei:0.2867732659)78/78:0.0406874004)78.4/68:0.0630489801)87.9/49:0.0999649736)88.4/23:0.0431139658)78/50:0.0290168839,((WP_168756429.1-Aeromonas-hydrophila:0.6186533065,WP_061279316.1-Kluyvera-cryocrescens:0.5754822703)92.8/95:0.1946669927,(((((WP_167863639.1-Erwinia-rhapontici:0.2095871255,WP_039385357.1-Pantoea-eucrina:0.2866901637)88.9/100:0.1265513068,WP_118663751.1-Erwinia-persicina:0.1937446312)69.5/100:0.0962449342,WP_062742814.1-Erwinia-persicina:0.2577274623)83/100:0.0990118423,WP_191149682.1-Erwinia-aphidicola:0.1761838013)75.6/88:0.0908642372,((WP_070133611.1-Candidatus-Erwinia-dacicola:0.3164038105,WP_140919905.1-Insectihabitans-xujianqingii:0.3408031955)46.8/87:0.0411044658,WP_029881813.1-Enterobacter:0.4563704201)81.2/69:0.0606474588)78/79:0.0290164492)81.6/56:0.0383817872)87.5/78:0.0422269065,(((((WP_001100781.1-Escherichia-coli:0.3059340045,WP_208464084.1-Escherichia-coli:0.1264418435)91.2/99:0.1003775650,(WP_262931739.1-Escherichia-albertii:0.0491793800,WP_160188772.1-Escherichia-coli:0.0661213100)95.7/100:0.1162306489)76.5/91:0.0352386747,(WP_021540003.1-Escherichia-coli:0.0232000157,WP_047657564.1-Enterobacteriaceae:0.0638039263)97.7/100:0.1276333941)13.2/36:0.0565809735,WP_021577651.1-Escherichia-coli:0.2337450655)0/71:0.0000022649,(WP_220393175.1-Citrobacter-freundii:0.4192807961,WP_219093375.1-Escherichia-coli:0.2684674271)83.3/86:0.0746489865)23.9/33:0.0317680509)33.5/60:0.0160736881,(((((((((WP_127791218.1-Escherichia-coli:0.1687779313,(WP_025210617.1-Escherichia-coli:0.1717973261,WP_001527244.1-Escherichia-coli:0.1202790380)47.8/92:0.0452456539)71.9/98:0.0378386783,WP_085454021.1-Escherichia-coli:0.2031608995)60.3/92:0.0084730402,(((WP_049002539.1-Citrobacter-freundii:0.1846658414,(WP_249576357.1-Enterobacter-vonholyi:0.0909819828,WP_161617217.1-Yokenella-regensburgei:0.0598521584)92.8/100:0.0783832907)83.3/100:0.0286945078,((WP_079834758.1-Salmonella-enterica:0.0000020111,(WP_079793206.1-Salmonella-enterica:0.1864750570,WP_001100807.1-Salmonella:0.1065671290)99.4/100:0.2316081875)73.9/100:0.0598799183,WP_058109100.1-Salmonella-enterica:0.0427293408)99.6/100:0.1428989564)91.4/100:0.0564020913,WP_032303308.1-Escherichia:0.0906410795)96.2/100:0.1031225842)89.5/96:0.0408849301,(((WP_097421351.1-Escherichia-coli:0.1154472992,(WP_137653986.1-Escherichia-albertii:0.0508375020,WP_001100785.1-Escherichia-coli:0.0578383135)93.1/100:0.0842179510)89.6/100:0.0653278610,WP_053290779.1-Escherichia-coli:0.2063270749)16.7/73:0.0147625515,(WP_157704420.1-Escherichia-coli:0.1567170982,WP_042200020.1-Enterobacteriaceae:0.0864381284)78.8/98:0.0593370559)89.1/80:0.0595461370)76.4/67:0.0224845244,(((((WP_259047621.1-Klebsiella-sp.-BIGb0407:0.2823384783,WP_105963664.1-Escherichia-coli:0.0509583762)86.5/97:0.0664008566,WP_015386517.1-Cronobacter-sakazakii:0.2969155162)48.3/85:0.0361389436,(WP_063610377.1-Escherichia-coli:0.1299074793,(WP_088541377.1-Escherichia-coli:0.0440795857,WP_222691870.1-Escherichia-marmotae:0.0862970028)93.2/100:0.0763997701)77.5/74:0.0316084766)79.5/32:0.0382803076,WP_218070502.1-Escherichia-coli:0.1049732905)81.7/36:0.0446230785,(WP_059222105.1-Escherichia-albertii:0.1957581362,WP_187209312.1-Escherichia-coli:0.1138895465)83.5/92:0.0498798020)81.4/34:0.0395589271)91.7/86:0.0623897893,WP_107179425.1-Escherichia:0.1689183599)0/13:0.0000020598,(((((WP_193360946.1-Enterobacter-cloacae-complex-sp.-P39RS:0.0789790554,WP_182945676.1-Enterobacter-cloacae:0.0355462796)98.7/100:0.1478024678,WP_265185975.1-Enterobacter-mori:0.0962838139)83.2/99:0.0437475227,((WP_032636062.1-Enterobacter-kobei:0.0867631998,WP_138014015.1-Enterobacter-ludwigii:0.0904602229)69.6/91:0.0528962335,(WP_063137673.1-Enterobacter-cloacae-complex:0.0446751479,WP_165464597.1-Enterobacter-cloacae:0.0821345704)80/100:0.0533674516)78.8/99:0.0462066376)84.6/80:0.0519328319,((((WP_096205048.1-Enterobacter:0.0438074828,WP_100165931.1-Enterobacter-mori:0.0456850624)94.7/100:0.0985309466,WP_242172213.1-Enterobacter:0.1366348040)50.4/100:0.0643989692,WP_239609625.1-Lelliottia-amnigena:0.1641438628)88.2/72:0.0613948509,((((((WP_042716137.1-Enterobacter-sp.-B509:0.0234738933,WP_241820523.1-Enterobacter-kobei:0.0988749002)99.2/100:0.1566150474,((((((WP_047742849.1-Enterobacter:0.1059661670,WP_040022520.1-Enterobacter-cloacae:0.1230578510)92.9/100:0.1059251427,(WP_129995405.1-Enterobacter-cloacae:0.0796324133,WP_042099316.1-Escherichia-coli:0.1185943052)91.4/100:0.0932623220)86.1/100:0.0964839926,WP_045339085.1-Enterobacter-cloacae-complex:0.2876574647)86.8/92:0.0630496491,(WP_045338355.1-Enterobacter-hormaechei:0.1999563029,(WP_182115563.1-Enterobacter-kobei:0.0481456654,WP_213381159.1-Enterobacter-cloacae:0.0913231468)91/100:0.1110434339)83.3/65:0.0457381985)78.9/54:0.0376236725,WP_063158410.1-Enterobacter-hormaechei:0.1230002852)82.8/56:0.0367539042,WP_063869545.1-Enterobacter-hormaechei:0.0721037533)38.1/57:0.0272764265)82/58:0.0341948817,((((WP_238085128.1-Pseudescherichia-vulneris:0.0147480771,(((WP_257661844.1-Citrobacter-portucalensis:0.1797598969,WP_193134185.1-Citrobacter-freundii:0.1126978024)91.9/100:0.0589413461,WP_129363899.1-Enterobacter-cloacae:0.0375652177)55.7/79:0.0204211504,WP_032652659.1-Enterobacter-cloacae-complex:0.1345596881)67.1/76:0.0099181891)100/100:0.4052385461,WP_172688853.1-Enterobacterales:0.0948888461)86.5/95:0.0938579883,WP_126546226.1-Enterobacter-quasimori:0.1293356099)20.2/81:0.0664587147,((((WP_232904305.1-Buttiauxella-sp.-W03-F01:0.2705857500,WP_259388919.1-Enterobacter-huaxiensis:0.0670327128)80/43:0.0486409691,WP_230350752.1-Lelliottia-sp.-WAP21:0.1891692461)91.9/100:0.0971856576,(((WP_141109706.1-Escherichia-albertii:0.1194873169,(WP_203360733.1-Citrobacter-sp.-R56:0.0873567939,WP_001572827.1-Escherichia-coli:0.0796401039)61/96:0.0222989845)85/99:0.0483331883,WP_258993080.1-Scandinavium-hiltneri:0.1207465354)88.9/99:0.0737639708,(WP_096122580.1-Escherichia-coli:0.1249456852,WP_162379936.1-Citrobacter-sp.-NCU1:0.2254542460)92.2/96:0.0955474044)75.1/87:0.0251930942)80.4/76:0.0297013055,WP_034915863.1-Erwinia-sp.-9145:0.2359491639)74.5/27:0.0336864601)77.8/34:0.0347981612)65.5/25:0.0103517603,(((WP_214593796.1-Enterobacter-mori:0.0794909432,WP_086532312.1-Enterobacter-ludwigii:0.0861591526)99.8/100:0.2562605708,(WP_126315015.1-Enterobacter-hormaechei:0.0601399897,((WP_042284276.1-Citrobacter-sedlakii:0.4282711097,WP_024193259.1-Escherichia-coli:0.1220113887)81.6/95:0.0432572656,WP_099433281.1-Citrobacter-amalonaticus:0.0345003443)100/100:0.4445104639)68.4/95:0.0425134564)75.9/91:0.0676175537,WP_046888360.1-Enterobacter-cloacae:0.1068987383)75.9/12:0.0212597323)23.2/10:0.0342513619,WP_208283396.1-Enterobacter-sichuanensis:0.1812330171)79/96:0.0229097808,WP_033487312.1-Enterobacter-cloacae-complex:0.0964128915)71.3/85:0.0301190458)77.1/59:0.0238081518)94.6/98:0.0626007387,((((WP_175320495.1-Hafnia-paralvei:0.1469195314,WP_130996599.1-Hafnia-paralvei:0.2652878311)92.9/100:0.1631766237,((WP_171269898.1-Aeromonas-media:0.1395754220,WP_216983019.1-Aeromonas-sp.-FDAARGOS-1415:0.1304310589)98.5/100:0.1884545455,WP_252995952.1-Aeromonas-encheleia:0.3789544870)50.4/50:0.0551230527)66.5/97:0.0659113796,WP_047609274.1-Rahnella-aquatilis:0.3343338679)87.7/91:0.0940758153,((WP_075180894.1-Pantoea-sp.-1.19:0.2864812985,WP_034458444.1-Buttiauxella-noackiae:0.3495142701)0/15:0.0116732947,(WP_105591414.1-Pantoea-coffeiphila:0.0901002916,WP_123334936.1-Erwinia-sp.-JUb26:0.0533123241)100/100:0.3273945224)76.8/15:0.0354274564)84.6/87:0.0468377185)56.1/21:0.0113099539)82.8/85:0.0283787581,(((((WP_001542972.1-Escherichia-coli:0.2961458598,WP_262938691.1-Escherichia-albertii:0.1558182618)0/30:0.0000028728,(WP_089706542.1-Escherichia-coli:0.1042134677,WP_032659356.1-Enterobacter-hormaechei:0.5675660473)74.1/78:0.0396294727)94.1/69:0.0681347993,((WP_095374511.1-Escherichia-coli:0.0000028280,WP_012906377.1-Citrobacter-rodentium:0.1423401123)99.7/100:0.2224572630,WP_096972986.1-Escherichia-coli:0.1405628612)70.2/40:0.0369852269)19.2/21:0.0244145623,WP_259000881.1-Scandinavium-goeteborgense:0.2601880481)75.7/30:0.0192829972,(((WP_109548670.1-Escherichia-coli:0.0710721394,WP_001100790.1-Escherichia:0.0238961958)95/100:0.1053119829,(((WP_061892435.1-Escherichia-coli:0.1460641149,WP_253383865.1-unidentified-bacterial-endosymbiont:0.5182436944)35.9/99:0.0243937761,WP_001100798.1-Enterobacteriaceae:0.1905387922)91.9/95:0.0934503810,WP_181237856.1-Escherichia-marmotae:0.0954707602)75.6/40:0.0347004843)38.9/40:0.0476662595,((WP_146314246.1-Escherichia-coli:0.0517425183,WP_096967326.1-Escherichia-coli:0.0351503417)99.1/100:0.1471666266,WP_181465273.1-Escherichia-fergusonii:0.0935195859)52.5/39:0.0519868243)83.5/80:0.0346177807)75.1/21:0.0121578544)84.8/53:0.0385551185,((((WP_162648791.1-Citrobacter-freundii:0.1951424019,(WP_060683585.1-Citrobacter-portucalensis:0.1637617631,(WP_079798993.1-Salmonella-enterica:0.1250623653,WP_032255798.1-Enterobacteriaceae:0.1157522788)69.7/99:0.0151549893)90.7/99:0.0717199770)62.3/86:0.0507201561,(WP_195994783.1-Citrobacter-portucalensis:0.3316775455,WP_048218074.1-Citrobacter-freundii-complex:0.2638379664)85/99:0.1016834277)95.4/99:0.1054133673,(WP_174633339.1-Escherichia-coli:0.1310401388,WP_072664248.1-Escherichia:0.1580899838)89.5/97:0.0856623179)83.9/85:0.0381053772,WP_249031739.1-Tenebrionicola-larvae:0.4607793089)73/69:0.0155822037)75.9/55:0.0138056530)86.2/60:0.0463966861,(WP_187216555.1-Escherichia-coli:0.2790975869,((((WP_181680936.1-Citrobacter-sp.-RHB25-C09:0.1553388094,(WP_090049757.1-Citrobacter-amalonaticus:0.0418913615,WP_196013128.1-Citrobacter-farmeri:0.0449237678)98.8/100:0.1841858668)79.9/100:0.0503645924,WP_103775088.1-Citrobacter-amalonaticus:0.2728895491)96.3/100:0.1537541622,WP_121265736.1-Enterobacter-sp.-R1-2018:0.3821303390)82.1/92:0.0622937654,WP_042093344.1-Escherichia-coli:0.3461113944)39.9/78:0.0165997484)86.8/87:0.0821104244)72.5/76:0.0201645533,((((((((((((((WP_085037002.1-Cronobacter-sakazakii:0.2912193757,WP_076770125.1-Kosakonia-cowanii:0.2255488121)95.7/100:0.2380226599,WP_216981205.1-Lelliottia-amnigena:0.4526465888)55.9/96:0.0843827646,((WP_227534785.1-Erwinia-rhapontici:0.1916882462,WP_017800228.1-Erwinia-toletana:0.2443519021)85.4/100:0.1150024292,WP_255851625.1-Erwinia-persicina:0.3041779401)70.7/100:0.1148560633)87/100:0.1045137896,(WP_089585421.1-Escherichia-coli:0.0470256049,WP_104705897.1-Escherichia-coli:0.0560915644)94.5/100:0.1479548208)91.9/100:0.1294682966,(WP_001610415.1-Escherichia-coli:0.0623542239,WP_138183825.1-Escherichia-sp.-E4385:0.0274573530)98.3/100:0.2452371697)92.5/100:0.1231060439,WP_033870520.1-Enterobacteriaceae:0.2988040953)79.1/89:0.0560594865,(WP_179154141.1-Citrobacter-freundii:0.4373108982,WP_086539147.1-Citrobacter-freundii:0.3390721016)63.8/88:0.1328340070)84.4/62:0.0595384091,((WP_107192407.1-Escherichia-albertii:0.0351372325,WP_181673375.1-Escherichia-fergusonii:0.0551740850)99.5/100:0.2018303465,(((WP_032214194.1-Escherichia-coli:0.0685175859,WP_059217428.1-Escherichia-albertii:0.0660266755)0/3:0.0000020231,WP_085459683.1-Escherichia-coli:0.1384686099)65.9/19:0.0289920210,(WP_073278747.1-Escherichia-coli:0.1213792934,WP_087893724.1-Enterobacteriaceae:0.0392732996)99.2/100:0.2022312231)97.2/100:0.1919140147)80.8/95:0.0529860136)58.5/15:0.0171524271,WP_267387874.1-Citrobacter-portucalensis:0.4365304558)78.6/20:0.0574485002,((WP_157959988.1-Citrobacter:0.7426886753,WP_170897747.1-Escherichia-coli:0.2059166687)72.2/53:0.0608746842,WP_100029735.1-Escherichia-coli:0.2393843314)79.4/22:0.0669970043)76.2/87:0.0536762350,(WP_248249449.1-Enterobacter-roggenkampii:0.2790249642,WP_032936727.1-Citrobacter-freundii:0.1445656386)99.1/100:0.2615476189)61.2/85:0.0227372091,WP_001100786.1-Escherichia-coli:0.2018391878)82.8/74:0.0419491841,(((WP_103946175.1-Lelliottia-aquatilis:0.1592011854,WP_123427447.1-Lelliottia:0.0897028439)56.4/98:0.0424413266,WP_235147774.1-Enterobacter-cloacae:0.3934361027)97.4/100:0.1432588250,((WP_095844187.1-Escherichia-coli:0.0897403277,(WP_086118650.1-Escherichia-coli:0.0368365135,WP_089538038.1-Escherichia-coli:0.0502229067)70.3/100:0.0153440020)99.5/100:0.2835947841,WP_137650031.1-Escherichia-albertii:0.1711249395)86.1/82:0.0825772306)71.1/65:0.0191962295)4.1/73:0.0046193007,((((WP_016236761.1-Enterobacteriaceae:0.1270627922,WP_024239504.1-Escherichia-coli:0.1887536721)95.7/100:0.1695960658,WP_138098269.1-Jejubacter-calystegiae:0.2279266982)52.5/86:0.1031301514,WP_001297344.1-Escherichia-coli:0.3379252754)0/57:0.0037660856,(WP_153879942.1-Citrobacter-freundii:0.3006491991,((((WP_181624464.1-Citrobacter-freundii:0.3311163974,WP_001515521.1-Escherichia-coli:0.2571465781)93.4/99:0.1299859018,(WP_058662478.1-Enterobacteriaceae:0.0705523924,WP_061709645.1-Pseudenterobacter-timonensis:0.0573926714)100/100:0.3567572877)63.2/12:0.0254727610,(WP_157953065.1-Limnobaculum-parvum:0.3352019506,WP_112164393.1-Rahnella-inusitata:0.3052288993)79.3/19:0.0561446410)89.5/86:0.0762186363,WP_073446180.1-Escherichia-coli:0.3614015606)71.9/75:0.0203993837)76.8/48:0.0226767815)89.8/71:0.0673141828)88.4/75:0.0389501858)84/87:0.0302935534,(((WP_119938542.1-Enterobacter-huaxiensis:0.1588458093,WP_050871166.1-Enterobacter-cloacae-complex:0.0563364989)78.4/93:0.0370942193,(WP_167816599.1-Enterobacter-cloacae:0.3972469200,(WP_262183194.1-Enterobacter-mori:0.2044186245,(WP_063412371.1-Enterobacter-cloacae:0.0579768139,WP_047027451.1-Enterobacter-kobei:0.0745558264)97.2/100:0.1773156228)96.4/100:0.2294920779)97.6/98:0.1905109896)33.1/78:0.0186638844,(WP_228937168.1-Escherichia-marmotae:0.0471087060,WP_040079227.1-Escherichia-coli:0.0576319930)91.2/100:0.0429752606)93.2/95:0.0603639618)92.1/86:0.0830044382,WP_202667543.1-Lelliottia-amnigena:0.2925844177)86.3/94:0.0615349974,WP_171149131.1-Erwinia-sp.-JH02:0.2324604965)98.3/100:0.1707762666,((WP_241647798.1-Rosenbergiella-sp.-JB07:0.4549775420,WP_065370581.1-Kosakonia-sacchari:0.2563569108)82.9/97:0.0583100971,WP_181599127.1-Citrobacter-sp.-RHB36-C18:0.3521450591)66.9/89:0.0149999463)93.8/93:0.1100449338)52.2/65:0.0194886450,(((((((((((WP_182931197.1-Enterobacter-cloacae:0.2454826530,(WP_048284323.1-Pluralibacter-gergoviae:0.2056887325,WP_211776196.1-Klebsiella-pneumoniae:0.0708455342)37.8/17:0.0225103143)68.2/16:0.0109281965,WP_047723323.1-Klebsiella-michiganensis:0.0601713557)75.3/16:0.0093974357,((WP_139968923.1-Klebsiella-pneumoniae:0.1347672451,WP_065799756.1-Klebsiella-pneumoniae:0.0076407396)0/17:0.0000021080,WP_095100910.1-Raoultella-sp.-10-1:0.0916844560)89.6/83:0.0474076772)76.8/66:0.0082278302,WP_104011323.1-Klebsiella-michiganensis:0.0613427922)84.8/84:0.0181144329,WP_149270444.1-Klebsiella-aerogenes:0.0511637148)83.5/72:0.0167671325,(WP_221694709.1-Klebsiella-aerogenes:0.1261432747,WP_213760001.1-Klebsiella-aerogenes:0.0874678387)73.4/21:0.0104742844)32.5/10:0.0074291771,WP_223370324.1-Klebsiella-pneumoniae:0.0596134360)88.5/66:0.0180695540,((WP_128320364.1-Raoultella-planticola:0.0745134730,(WP_236948387.1-Klebsiella-sp.-2019SCSN059:0.0485773868,((WP_114692711.1-Klebsiella-pneumoniae:0.0490916578,WP_259891680.1-Klebsiella-variicola:0.0402393710)0/78:0.0000027340,WP_123755035.1-Raoultella-sp.-BIGb0399:0.0483669907)86.1/83:0.0150199933)75.4/81:0.0090959399)71/26:0.0139488379,WP_046852082.1-Raoultella-planticola:0.0492654077)79.1/25:0.0108990659)76.9/47:0.0092603968,((WP_234537246.1-Raoultella-ornithinolytica:0.0529465543,WP_113990968.1-Raoultella:0.0321515705)97.9/100:0.0605113803,(WP_234579544.1-Raoultella-terrigena:0.0588408935,WP_186601692.1-Klebsiella-quasipneumoniae:0.0251277427)74.6/75:0.0084759480)75.4/69:0.0079056414)93.9/75:0.1022659058,((WP_095100919.1-Raoultella-sp.-10-1:0.0735496028,WP_212709529.1-Tatumella-sp.-JGM130:0.0225385535)98.4/99:0.1228157678,WP_236948353.1-Klebsiella-sp.-2019SCSN059:0.1235609599)86.6/68:0.0509881875)81.2/100:0.0497392148,WP_123822891.1-Klebsiella:0.0437821536)98.4/100:0.1417983462)86.6/95:0.0582892310,(WP_072663757.1-Enterobacteriaceae:0.1002298001,WP_021241446.1-Enterobacter-roggenkampii:0.2834308332)18.9/95:0.0315875430)99.3/100:0.1472328028)92.7/100:0.0738362350,XP_011503919.1-Ceratosolen-solmsi-marchali:0.1241899395);

**Nematode RML-3, with bacterial and eukaryotic homologs**

**Shown in Figure S2**

ML tree method: IQ-TREE

Sequences: 2787

Substitution model: JTT

(WP_063501760.1-Escherichia-coli:0.0326409998,(WP_044173167.1-Phytobacter-massiliensis:0.0273458424,((((((((WP_203451401.1-Lelliottia-sp.-RWM.1:0.0517169976,WP_185654835.1-Citrobacter-braakii:0.0962152451)93.2/100:0.0639096418,WP_183170804.1-Atlantibacter-sp.-RC6:0.1685585195)97/99:0.0792465655,(WP_160868038.1-Pantoea-sp.-Taur:0.2097661252,WP_008499438.1-Enterobacter-cloacae-complex:0.1624193025)75.8/66:0.0416877987)37/59:0.0136354128,((WP_090085547.1-Escherichia-coli:0.0486109380,WP_103283990.1-Citrobacter:0.1444512181)13.4/74:0.0150605413,WP_234083964.1-Enterobacter-quasiroggenkampii:0.1022868304)99.9/100:0.1439968778)64.7/68:0.0268507776,((((((((((((((WP_249615874.1-Craterilacuibacter-sp.-RT1T:0.3072114185,WP_008952446.1-Pseudogulbenkiania-ferrooxidans:0.1698771118)77.8/31:0.0266372988,((((((((((((WP_004123988.1-Rhizobium-freirei:0.0807091700,WP_095433322.1-Rhizobium-sp.-11515TR:0.0293528085)84.5/95:0.0211735065,((WP_183712806.1-unclassified-Rhizobium:0.0299120554,WP_015341698.1-Rhizobium:0.0276454200)67.9/76:0.0111982863,WP_106999541.1-Rhizobium-sp.-JAB6:0.0417416149)0/70:0.0000023250)75.1/98:0.0171141003,WP_184706966.1-Rhizobium-lusitanum:0.0794762685)0/72:0.0000028872,(WP_174074493.1-Agrobacterium-rhizogenes:0.0788293644,((WP_183765619.1-unclassified-Rhizobium:0.0329527185,WP_120667780.1-unclassified-Rhizobium:0.0445803362)83.8/100:0.0255342323,WP_037195771.1-Rhizobium-lusitanum:0.0222443323)48.3/97:0.0097606019)78.9/72:0.0123408524)73.6/74:0.0070328362,WP_092719658.1-Rhizobium-multihospitium:0.0305707884)91.5/100:0.0248878345,WP_168880117.1-Rhizobium-sp.-P28RR-XV:0.0345775346)71.6/96:0.0054973867,WP_112341687.1-Rhizobium-tropici:0.0341787021)89.3/100:0.0182345548,(WP_199146672.1-Rhizobium-sp.-ASV8:0.1400150141,WP_145321829.1-unclassified-Rhizobium:0.0720166325)15.3/90:0.0080672762)99.8/100:0.1020926176,(WP_043579071.1-Chromobacterium:0.0541484718,WP_071113424.1-Chromobacterium-sphagni:0.0740277514)99.9/100:0.1192211745)95.1/100:0.0549859694,((WP_189531325.1-Paludibacterium-paludis:0.3092947492,(WP_043620211.1-Chromobacterium-piscinae:0.1413057076,(WP_103900886.1-Chromobacterium-alticapitis:0.1562853063,WP_045052286.1-Chromobacterium-violaceum:0.1657030909)91.5/100:0.0661254092)91.3/100:0.0704666995)55.7/98:0.0333212182,(WP_020932001.1-Cycloclasticus-zancles:0.2059338834,(((WP_213263532.1-Burkholderia-pyrrocinia:0.0967266873,WP_060279477.1-Burkholderia-cepacia-complex:0.1067502790)99.5/100:0.1131436950,(((WP_208101977.1-Burkholderia-sp.-D-99:0.0292421584,WP_205714023.1-Burkholderia-cepacia:0.0256700242)41.6/100:0.0060791975,WP_174919663.1-Burkholderia-metallica:0.0184627734)87.4/100:0.0342292563,WP_077021900.1-Burkholderia-cenocepacia:0.0661077790)99.9/100:0.1463987704)88.9/100:0.0591322933,(WP_159673755.1-Andreprevotia-sp.-IGB-42:0.1407931026,WP_084093192.1-Andreprevotia-lacus:0.0659414606)98.5/100:0.1006751788)76.4/99:0.0385485778)49.5/81:0.0392408260)84.9/80:0.0287407537)92.9/79:0.0418671286,((WP_174873379.1-Vogesella-oryzae:0.0792034420,WP_137008594.1-Aquitalea-aquatilis:0.2317303549)95.2/100:0.0770277902,WP_147289921.1-Crenobacter-cavernae:0.0872043741)92.6/100:0.0539521339)94.1/81:0.0528394908,WP_014088374.1-Pseudogulbenkiania-sp.-NH8B:0.1683236069)87.4/28:0.0319003937)61.9/28:0.0309369586,(WP_163316497.1-Crenobacter-sedimenti:0.0949607547,WP_160797365.1-Craterilacuibacter-sinensis:0.1177160162)100/100:0.1568132523)45.8/60:0.0218807951,((((((WP_223932214.1-Aeromonas-caviae:0.0843799984,WP_161506954.1-Aeromonas-media:0.0296952922)98.1/100:0.0732021106,(WP_208149206.1-Shewanella-algae:0.2133971000,(WP_219142360.1-Aeromonas-sanarellii:0.1148329308,(((WP_045500143.1-Vibrio-hyugaensis:0.1358757561,WP_045419977.1-Vibrio-campbellii:0.0632782640)63.6/99:0.0333801348,WP_025533544.1-Vibrio-parahaemolyticus:0.0533403303)94.8/100:0.0377911354,WP_011788809.1-Shewanella-sp.-W3-18-1:0.0593919829)85.6/100:0.0251567832)19.6/46:0.0209204160)96.3/100:0.0936976350)90/98:0.0470074958,(((WP_147247296.1-Brenneria-salicis:0.0375404691,WP_172289906.1-unclassified-Brenneria:0.0800341206)82.4/100:0.0200905240,WP_039356706.1-Pectobacterium-actinidiae:0.0800519382)95.4/99:0.0571497675,((WP_198997094.1-Pectobacterium-carotovorum:0.0165814476,(WP_165707548.1-Pectobacterium:0.0279808646,WP_174866005.1-Pectobacterium-polaris:0.0245676063)99.1/100:0.0524853538)84.2/99:0.0175942282,WP_145962313.1-Pectobacterium-peruviense:0.0429931211)99.5/100:0.0971837733)95.4/99:0.0658092624)20.1/33:0.0140239865,(((WP_248067241.1-unclassified-Shewanella:0.0810492672,WP_148505453.1-Vibrio-cholerae:0.0771077943)96.3/100:0.0733170242,(WP_218558465.1-Shewanella-putrefaciens:0.0741989809,WP_011919582.1-Shewanella-putrefaciens:0.0190910057)99/100:0.0749733638)89/99:0.0362299121,(((((WP_050718733.1-Aeromonas-tecta:0.1203980051,WP_100859741.1-Aeromonas-lusitana:0.0339110555)64.7/97:0.0068874056,WP_042654929.1-Aeromonas-encheleia:0.0475222361)98.6/100:0.0521177458,WP_077098334.1-Aeromonas-hydrophila:0.0446885772)79.8/100:0.0153390581,WP_043167431.1-Aeromonas-hydrophila:0.0893562867)40.2/98:0.0133677763,WP_252420256.1-Aeromonas-taiwanensis:0.0580124651)97.3/100:0.0693586139)54.9/81:0.0524780016)92.2/84:0.0386244364,(((WP_073350970.1-Aeromonas-hydrophila:0.0954975205,WP_190284026.1-Aeromonas-hydrophila:0.0338357414)88.1/99:0.0256774800,WP_241344946.1-Aeromonas-sp.-MR16:0.0546017840)91.5/99:0.0352747992,(WP_223952962.1-Aeromonas-caviae:0.1369824572,WP_244081328.1-Aeromonas-caviae:0.1308701112)90.8/97:0.0594664907)92.2/96:0.0456635999)77.8/54:0.0344381207,WP_102949229.1-Aeromonas-caviae:0.1987025992)84.6/84:0.0307680485)74.7/32:0.0281660927,((((((((((((WP_250059255.1-Halomonas-sp.-ATCH28:0.2211651994,WP_179153381.1-Chromohalobacter-salexigens:0.1573112771)90.1/100:0.0553964015,WP_250939176.1-unclassified-Larsenimonas:0.1828269375)57.5/98:0.0524574977,WP_106478747.1-Phytohalomonas-tamaricis:0.1785222565)83.6/98:0.0355170565,((WP_136867012.1-Guyparkeria-sp.-SB14A:0.1909675016,WP_058575749.1-unclassified-Guyparkeria:0.1591453799)92.7/100:0.0606966730,(WP_218634960.1-Marinobacter-arenosus:0.0890342221,WP_240716217.1-Halomonas-neptunia:0.0851016811)96.7/100:0.0877171579)90.9/100:0.0405425556)98.8/81:0.0799413719,((WP_077733980.1-Paenalcaligenes-hominis:0.0350774513,WP_167661700.1-Paenalcaligenes-hominis:0.0294777658)100/100:0.2015205363,(((WP_177138191.1-Hydrogenophaga-aromaticivorans:0.0948998459,WP_133157403.1-Hydrogenophaga-pseudoflava:0.1247796391)89.2/99:0.0377737844,(WP_236964528.1-Hydrogenophaga-sp.-SL48:0.0742160343,(WP_226493830.1-Hydrogenophaga-taeniospiralis:0.0303887692,WP_210824367.1-Hydrogenophaga-aromaticivorans:0.0593218127)66.7/99:0.0346252748)92.9/100:0.0499863795)92.6/99:0.0568809486,WP_065086507.1-Pseudomonas:0.2136272766)24.9/33:0.0110845849)28.4/22:0.0202148084)0/32:0.0000026737,((((((((((((WP_269022943.1-Methylomonas-sp.-MP1:0.0737513761,(WP_196437371.1-Methylomonas-sp.-LL1:0.0965032711,WP_066983474.1-Methylomonas-lenta:0.1002623283)95.7/98:0.0549049611)94.4/98:0.0425545228,(WP_140911374.1-Methylomonas-koyamae:0.1530610606,(WP_071159776.1-Methylomonas-sp.-LWB:0.1307579496,((WP_020485671.1-Methylomonas-sp.-MK1:0.1374811189,(WP_256607223.1-Methylomonas-sp.-WSC-7:0.0578010702,(WP_205450646.1-Methylomonas-sp.-EFPC1:0.0706117919,WP_064036612.1-Methylomonas-methanica:0.0000024344)73.4/100:0.0084578590)95.3/100:0.0522002847)76.1/89:0.0470003245,WP_036273427.1-Methylomonas:0.0854144368)14.8/88:0.0092618962)90.3/100:0.0362044646)86.8/100:0.0453361876)91.4/98:0.0427056014,WP_064030819.1-Methylomonas-koyamae:0.1452423193)35.9/38:0.0307604145,WP_148817865.1-Pigmentiphaga-aceris:0.2332498763)69.2/27:0.0240223889,((WP_019964198.1-Pannonibacter-phragmitetus:0.3534597801,WP_150047608.1-Methylomonas-rhizoryzae:0.1543214871)12.3/89:0.0671390822,WP_138632053.1-Pseudoalteromonas-spongiae:0.2341552234)88.1/51:0.0492392899)91.8/36:0.0543379787,WP_004074545.1-Desulfobacter-postgatei:0.1998915217)68.3/27:0.0283352291,(((WP_168987795.1-Azoarcus-taiwanensis:0.1709681005,WP_018865194.1-Thioalkalivibrio-sp.-ARh3:0.1285679569)84/93:0.0482628843,(WP_256068944.1-Stutzerimonas-stutzeri:0.1147970021,WP_026685886.1-Azovibrio-restrictus:0.1246256084)86/98:0.0470384165)82.8/80:0.0270946661,(((WP_238617884.1-Ectothiorhodospira-shaposhnikovii:0.1074257144,WP_025282248.1-Ectothiorhodospira-haloalkaliphila:0.0952891227)89.8/100:0.0579301544,WP_238622182.1-unclassified-Ectothiorhodospira:0.0949241731)89.3/100:0.0525730807,(((((((((WP_060482506.1-Pseudomonas-sp.-NBRC-111119:0.1394117338,WP_159409308.1-Pseudomonas-putida:0.0699159276)95.2/86:0.0461179360,(((WP_110607173.1-Pseudomonas-fulva:0.0112754473,WP_088523134.1-unclassified-Pseudomonas:0.0410851415)94.2/100:0.0310141715,(WP_167339957.1-Pseudomonas-fulva:0.0287514044,WP_186476580.1-Pseudomonas-promysalinigenes:0.0342433356)75.6/100:0.0196357981)32.1/74:0.0246822471,(WP_201214098.1-Pseudomonas-sp.-S36:0.0182105633,WP_054890059.1-Pseudomonas:0.0520177409)96.4/100:0.0402276452)0/60:0.0000025085)87.8/33:0.0116361607,((WP_182365203.1-Pseudomonas-juntendi:0.0512528082,WP_182365213.1-Pseudomonas:0.0861398279)81.4/82:0.0227323065,WP_267821885.1-Pseudomonas-putida:0.0623678511)75.6/34:0.0082136448)80.8/57:0.0104220588,(WP_167277242.1-Pantoea-sp.-Tr-811:0.0167618891,WP_103449615.1-Pseudomonas-putida:0.0461065904)97.7/100:0.0410516497)66.9/59:0.0076746453,((WP_115272924.1-Pseudomonas-putida:0.0904541729,(WP_046489762.1-Pseudomonas-veronii:0.1030355416,(WP_059395284.1-Pseudomonas-putida:0.0732875109,(WP_009685980.1-Pseudomonas:0.0694728613,WP_024086460.1-Pseudomonas:0.0457161813)92.3/100:0.0327336286)89.3/100:0.0257623118)92.1/88:0.0276634363)89.8/86:0.0262494376,((WP_196129978.1-Pseudomonas-fulva:0.0771160182,WP_201164743.1-Pseudomonas-sp.-S60:0.0256415402)99.2/100:0.0719201722,(((WP_174142546.1-Pseudomonas-putida:0.1122841747,WP_027907237.1-Pseudomonas:0.0302090402)37.7/100:0.0087088307,WP_075044077.1-Pseudomonas-putida:0.0408214490)54.2/100:0.0190849475,(WP_023379297.1-Pseudomonas-sp.-VLB120:0.0764745393,WP_202675963.1-Pseudomonas-sp.-SK2:0.0342480119)98.5/100:0.0591220161)89.7/100:0.0399342963)79.8/99:0.0255340993)79.5/65:0.0131880792)93.4/98:0.0283604726,WP_029885231.1-Pseudomonas:0.0251374489)35.1/84:0.0111184093,(WP_230064532.1-Pseudomonas-sp.-CIP-10:0.0177294669,WP_197863301.1-Pseudomonas-putida:0.0345194539)91.7/100:0.0252358807)97.5/98:0.0581239667,WP_060394622.1-Pseudomonas-entomophila:0.0738350073)96.4/100:0.0605465845,WP_183165449.1-Azomonas-macrocytogenes:0.1054468667)69.8/100:0.0290358213)86.6/98:0.0273034116)49.8/83:0.0098137966)96/78:0.0643625069,(WP_136386324.1-Pseudothauera-rhizosphaerae:0.1041688306,(WP_114649035.1-Pseudothauera-hydrothermalis:0.0743976934,WP_187718084.1-Thauera-sp.-CAU-1555:0.1108205181)94.5/100:0.0650608824)98.9/100:0.1111283157)58.8/42:0.0433250078,WP_070073009.1-Acidihalobacter-aeolianus:0.1794191456)0/31:0.0166174975,(WP_013164002.1-Desulfurivibrio-alkaliphilus:0.1594717191,WP_263768642.1-Propionivibrio-sp.-SG131:0.1838030339)68.7/60:0.0767441919)89.3/84:0.0408427233,(((WP_205202385.1-Azonexus-hydrophilus:0.1431065693,WP_121459656.1-Azonexus-fungiphilus:0.1694026634)34.4/58:0.0378140015,WP_173581083.1-Azonexus-fungiphilus:0.1698110183)95.6/89:0.0640999801,((WP_117174481.1-Rhodoferax-lacus:0.1635162097,((WP_094482496.1-Rhodoferax-sp.-TH121:0.0622724849,WP_105262671.1-Rhodoferax-sp.-TS-BS-61-7:0.0377476751)87.2/100:0.0303152039,WP_214121631.1-Curvibacter-sp.-CHRR-16:0.1968702197)96.5/100:0.0994330869)51.1/24:0.0338319059,(WP_066884254.1-Dechloromonas:0.0800981453,WP_027458229.1-Dechloromonas-agitata:0.0847967697)97/100:0.0625699391)83.8/45:0.0295597106)87.5/66:0.0436063620)97.3/96:0.0741748204,((WP_076518421.1-Achromobacter-sp.-MFA1-R4:0.0585847072,(WP_104142565.1-Achromobacter-spanius:0.0217923693,WP_120157261.1-Achromobacter-sp.-B7:0.0457338250)92.9/100:0.0429586141)98.4/100:0.0895569895,((WP_209394989.1-Rhodoferax-sp.-AJA081-3:0.2207736466,WP_094844257.1-Bordetella-genomosp.-11:0.1204308224)93/77:0.0514001071,WP_166067558.1-Diaphorobacter-sp.-HDW4B:0.1423291698)87.3/77:0.0391312309)69.2/64:0.0330331130)92.5/7:0.0351629103)22.1/4:0.0202641716,((((((((((((WP_102075761.1-Pollutimonas-subterranea:0.1296364524,WP_102071501.1-Pollutimonas-nitritireducens:0.1763566646)84.9/99:0.0482695426,(WP_212592286.1-Janthinobacterium-sp.-TND4EL3:0.1534423590,WP_217963608.1-Candidimonas-humi:0.1773668838)88.8/97:0.0511876050)80.6/97:0.0354061039,WP_047550623.1-Methylotenera-sp.-G11:0.1854748841)87.2/98:0.0380954952,(WP_019141909.1-Noviherbaspirillum-massiliense:0.1214575557,WP_146330669.1-Noviherbaspirillum-sp.-UKPF54:0.1810244557)84.2/49:0.0438976808)12.2/15:0.0204205146,(WP_230502293.1-Janthinobacterium-sp.-FW305-128:0.2661705184,WP_249918475.1-Pseudomonas-sp.-R3.Fl:0.1369225446)93/50:0.0696938166)66.5/11:0.0091685843,(WP_074973890.1-Pseudomonas-otitidis:0.1264821840,WP_068803504.1-Immundisolibacter-cernigliae:0.1605711334)75.7/40:0.0572191643)72.2/49:0.0251904231,(WP_011111299.1-Nitrosomonas-europaea:0.1445693895,WP_046857648.1-Pseudomonas-sp.-CCOS-191:0.1323527364)88.9/54:0.0349561522)41.1/91:0.0179012939,WP_158658301.1-Achromobacter-sp.-AONIH1:0.2972235975)79/30:0.0223672997,((((WP_138213821.1-Pseudomonas-nitroreducens:0.0339048744,WP_017517140.1-Pseudomonas-nitroreducens:0.0181286239)100/100:0.1914429019,WP_085721622.1-Pseudomonas:0.1210491302)80.8/31:0.0339119878,(WP_088146851.1-Achromobacter-denitrificans:0.0817904931,WP_105555808.1-Achromobacter-sp.-MYb9:0.0673264258)98.3/100:0.0756966486)66.6/30:0.0310686155,((WP_029241333.1-Pseudomonas:0.0607482235,((WP_054996663.1-Pseudomonas-ficuserectae:0.0330247294,(WP_007252556.1-Pseudomonas-syringae-group:0.0412121623,WP_025390602.1-Pseudomonas-syringae:0.0196584439)72.1/100:0.0226285619)94/100:0.0405172950,WP_125830676.1-Pseudomonas-sp.-v388:0.0602091080)50.6/100:0.0370235178)96.9/100:0.0766226549,((WP_016973326.1-Pseudomonas-tolaasii:0.0890125829,WP_099548838.1-Pseudomonas-sp.-2995-3:0.0873578946)66.4/98:0.0345880169,(WP_177015116.1-Pseudomonas-costantinii:0.0423699748,WP_141063556.1-Pseudomonas-fluorescens:0.0844400094)96.9/100:0.0511920466)93.2/100:0.0490958588)98.7/100:0.0994414447)85.8/25:0.0186084669)85.6/67:0.0202279750,(((((((WP_191471912.1-Pseudomonas-asiatica:0.1531337542,(WP_043318452.1-Pseudomonas-sp.-PI1:0.0406372296,WP_043268312.1-Pseudomonas:0.0294515309)85.2/100:0.0260345355)94.6/100:0.0498011831,(WP_089390899.1-Pseudomonas-delhiensis:0.0898416968,WP_076426141.1-Pseudomonas-alcaligenes:0.1077106408)14.7/75:0.0242737182)96/99:0.0631032115,WP_233297479.1-Pseudomonas:0.1139122719)66.3/58:0.0188543569,(WP_184594840.1-Pseudomonas-nitritireducens:0.1367509567,WP_192381882.1-Pseudomonas-sp.-PDM20:0.0908756403)99.7/100:0.1134858528)58.5/39:0.0332514681,WP_209872004.1-Pseudomonas-sp.-BP8:0.1704263754)80.2/83:0.0326248625,((WP_057247240.1-Duganella-sp.-Root198D2:0.1852267252,WP_155457065.1-Massilia-eburnea:0.0647698105)95.6/95:0.0614094388,WP_180152801.1-Parapusillimonas-granuli:0.1240226102)90.1/39:0.0518383223)0.4/5:0.0122503677,(((WP_058488093.1-Pseudomonas:0.1126649231,WP_043256200.1-Pseudomonas-knackmussii:0.1029660223)49.3/100:0.0341480335,((WP_088420173.1-Pseudomonas-nitroreducens:0.1034404809,WP_254475170.1-unclassified-Pseudomonas:0.0574978718)84.2/100:0.0334333325,WP_205338011.1-Pseudomonas-sp.-PDNC002:0.0448582123)99.7/100:0.1259158063)90.3/99:0.0404648432,((((WP_042945119.1-Pseudomonas-fluorescens:0.0556082826,WP_181643258.1-Pseudomonas-sivasensis:0.0396975731)99.3/100:0.0952487640,WP_205489922.1-Pseudomonas-lactucae:0.1252474574)35.2/13:0.0288867343,((WP_169965430.1-unclassified-Pseudomonas:0.0361294947,WP_047278503.1-Pseudomonas-lundensis:0.0232426556)99.9/100:0.0840794044,WP_104727848.1-Pseudomonas-oleovorans:0.1250349990)66.4/55:0.0163753543)75.5/58:0.0287123698,WP_214501190.1-Pseudomonas-sp.-RC3H12:0.1946889717)82.5/17:0.0279038176)87/25:0.0380097888)83.5/41:0.0252972771)3.7/36:0.0123317372,WP_066093302.1-Hydrogenophaga-crassostreae:0.2022513951)72.6/21:0.0065751110,((((((WP_066098770.1-Halothiobacillus-sp.-LS2:0.1571590663,WP_223462047.1-Pseudomonas-sp.-A-R-26:0.1531078239)78.3/100:0.0476134972,WP_058422493.1-Pseudomonas:0.1247257275)93.3/100:0.0487627497,((WP_225237063.1-Massilia-oculi:0.2090543395,WP_147937383.1-Massilia-arenae:0.1101325702)98.4/100:0.1192122659,(WP_192051557.1-unclassified-Massilia:0.0937666626,WP_136222466.1-Massilia-sp.-Mn16-1_5:0.0744578939)75.5/100:0.0564188871)96/100:0.0620682325)93.4/97:0.0520823662,(WP_125865093.1-Stutzerimonas-stutzeri:0.1317910750,WP_200290630.1-Stutzerimonas-balearica:0.0944764901)86.1/41:0.0376709199)81/31:0.0123567505,WP_020564242.1-Methylosarcina-fibrata:0.1697926302)75.5/29:0.0187442300,(WP_101152997.1-Pseudomonas-sp.-Choline-3u-10:0.0995144363,((WP_046617631.1-Stutzerimonas-stutzeri:0.0612482087,WP_135265705.1-Stutzerimonas-stutzeri:0.0402571028)94.1/100:0.0601879197,WP_003299494.1-Stutzerimonas-stutzeri:0.0723882309)74.7/100:0.0288179888)96/99:0.0538528934)82.6/25:0.0150515478)74.6/43:0.0088613537)92.6/47:0.0435249292,((((WP_044615503.1-Gynuella-sunshinyii:0.2623120622,(WP_046292890.1-Pandoraea-oxalativorans:0.0754681159,WP_150719538.1-Pandoraea-capi:0.0773478214)99.9/100:0.2164356651)41/89:0.0477872297,WP_092579398.1-Achromobacter-sp.-NFACC18-2:0.2678392085)89.3/88:0.0580307563,WP_227683402.1-Pusillimonas-sp.-MFBS29:0.1560295365)88.6/88:0.0408383814,((((WP_220354719.1-Rhizobium-cauense:0.1763270680,WP_016557527.1-Rhizobium-grahamii:0.1078073608)35/98:0.0364610037,(WP_072373837.1-Rhizobium-tibeticum:0.0712545568,WP_007537558.1-Rhizobium-mesoamericanum:0.0558270206)84.6/100:0.0403917996)91.3/99:0.0408564421,((WP_184450570.1-Rhizobium-sp.-BK538:0.1418190526,WP_153273632.1-Rhizobium-grahamii:0.1733123164)39.7/97:0.0303791767,WP_233136718.1-Agrobacterium-vaccinii:0.1869099118)24.7/97:0.0259182177)74.7/98:0.0361980765,WP_018896487.1-Rhizobium-sp.-2MFCol3.1:0.1276925660)99.7/100:0.1259277983)93.3/88:0.0548763628)86.4/97:0.0335138059,((((((((((((((((WP_072282603.1-Syntrophotalea-acetylenivorans:0.0905469494,(WP_226989104.1-Desulfuromonas-sp.-TF:0.2321445184,WP_227459030.1-Cupriavidus-pauculus:0.2449289471)98.7/92:0.1537664232)37.3/27:0.0192492787,WP_092056970.1-Desulfuromonas-acetexigens:0.1499964757)57.3/49:0.0329259410,WP_155876508.1-Desulfuromonas-sp.-AOP6:0.2194268496)57/22:0.0425379678,WP_148894253.1-Geothermobacter-ehrlichii:0.2234066681)74.5/20:0.0300004974,WP_043770486.1-Desulfobulbus-propionicus:0.2016893121)80.6/8:0.0322361908,(WP_028317018.1-Desulfobulbus-elongatus:0.2258568968,(((WP_167682126.1-Rhodocyclus-gracilis:0.2067087214,(WP_153115862.1-Rhodocyclus-tenuis:0.0356099167,WP_200385811.1-Rhodocyclus-tenuis:0.0190742043)91.3/100:0.0687899310)79.1/100:0.0487151886,WP_201212349.1-Rhodocyclus-purpureus:0.1165658674)95.8/100:0.0938950076,WP_153110537.1-Propionivibrio-limicola:0.1613804584)94.5/99:0.0689896481)60.2/5:0.0295340661)97.8/80:0.0801470256,WP_040097704.1-Geoalkalibacter-ferrihydriticus:0.2082683743)35.2/6:0.0231515984,((((((((((WP_014706296.1-Methylophaga-nitratireducenticrescens:0.0200282718,WP_136552334.1-Methylophaga-sp.-SB9B:0.0452753778)100/100:0.2891698533,WP_216129682.1-Azospira-inquinata:0.2159665704)36.1/79:0.0386705406,WP_204103763.1-Spirulina-major:0.2723261956)48.9/63:0.0308984492,(WP_108623287.1-Orrella-marina:0.2871561963,WP_243478771.1-Orrella-sp.-f23:0.2211129375)91.2/99:0.0678379671)92.8/97:0.0748250734,WP_258826205.1-Massilia-agri:0.2146350506)80.9/89:0.0434393557,((WP_176062241.1-Anaeromyxobacter-diazotrophicus:0.1175481316,WP_248361410.1-Anaeromyxobacter-oryzae:0.1862451714)98.7/100:0.0973232204,(((WP_018882239.1-unclassified-Thioalkalivibrio:0.2663370830,WP_125198734.1-Guyparkeria-sp.-SCN-R1:0.1435259020)89.9/98:0.0666550832,(((WP_026296529.1-Thioalkalivibrio-sp.-ALJ2:0.1134275479,WP_019569965.1-Thioalkalivibrio-sp.-ALE11:0.0651142898)88.5/97:0.0210958641,WP_028484075.1-Thioalkalivibrio-sp.-ALE17:0.0347026646)76.5/97:0.0074037123,WP_018169543.1-Thioalkalivibrio-sp.-ALMg9:0.0270774619)98.4/99:0.0653227352)0/56:0.0325587309,WP_250072676.1-Guyparkeria-hydrothermalis:0.1506739749)99.5/100:0.1231940523)87.9/91:0.0467760702)45.7/31:0.0400733336,((WP_083560927.1-Oceanococcus-atlanticus:0.2184024121,WP_104228621.1-Solimonas-fluminis:0.2144547040)98.8/100:0.1594745108,(WP_033955816.1-Pseudomonas-aeruginosa:0.0154792132,WP_003153917.1-Pseudomonas-aeruginosa-group:0.0378340731)100/100:0.1532269069)58.9/28:0.0348873060)82.2/20:0.0232407792,((WP_220807781.1-Noviherbaspirillum-aridicola:0.3034548996,WP_119020982.1-Halomonas-sp.-JS92-SW72:0.1897562132)80.9/97:0.0933401512,(((((WP_200222041.1-Achromobacter-xylosoxidans:0.1159661282,WP_259251071.1-Achromobacter-sp.-JUb104:0.1320274119)97.6/100:0.0667735003,(WP_088588627.1-Achromobacter-marplatensis:0.1179938922,WP_169273406.1-Achromobacter-sp.-Bel:0.0700935238)96/100:0.0607431351)95.7/94:0.0489505800,WP_174717299.1-Achromobacter-denitrificans:0.0726014672)77.5/76:0.0119114971,((((WP_111826915.1-Achromobacter-sp.-HZ01:0.0866817992,WP_102775120.1-Achromobacter:0.0174436321)99.8/100:0.1126183011,WP_043544564.1-Achromobacter-sp.-RTa:0.0304288562)65.2/70:0.0058480588,WP_129525645.1-Achromobacter-agilis:0.0303309081)85.6/72:0.0180557076,WP_129240460.1-Achromobacter-veterisilvae:0.0348514909)97.1/98:0.0464833245)90.1/78:0.0332951880,((WP_175208175.1-Achromobacter-anxifer:0.0456543134,((((WP_251864322.1-Achromobacter-sp.-Marseille-Q4962:0.0618159155,WP_164793551.1-Achromobacter-insolitus:0.0278344534)97.8/99:0.0472672848,WP_207472676.1-Achromobacter-sp.-SD115:0.0098927362)92.7/98:0.0356338535,WP_125471167.1-Achromobacter:0.0093047683)86.2/98:0.0404432183,WP_175164188.1-Achromobacter-aegrifaciens:0.0349444038)0/3:0.0176966568)39.6/76:0.0320058654,WP_088158656.1-Achromobacter-xylosoxidans:0.0559202602)94/100:0.0511408097)97.3/100:0.0978866454)92.9/31:0.0753908902)90/8:0.0387105925,WP_066727622.1-Desulfuromonas-sp.-DDH964:0.2113786736)0/3:0.0043196242,(WP_002810067.1-Nitrosococcus-oceani:0.0438911198,WP_013221233.1-Nitrosococcus-watsonii:0.0540522997)100/100:0.1986409113)87.7/6:0.0269297342)87.6/28:0.0278853536,(((((((WP_238511579.1-Cupriavidus-gilardii:0.1835862464,WP_224042884.1-Cupriavidus-respiraculi:0.2010389325)83.2/70:0.0631208884,((((WP_116346290.1-Cupriavidus-taiwanensis:0.1183473745,WP_124685047.1-Cupriavidus-pauculus:0.1496750894)100/100:0.2427288820,WP_150984056.1-Cupriavidus-basilensis:0.2130493051)93.4/93:0.0731794498,((WP_018008101.1-Cupriavidus-neocaledonicus:0.0346544706,WP_183298446.1-Cupriavidus-alkaliphilus:0.0511276493)99.6/100:0.1205859785,WP_250479419.1-unclassified-Caballeronia:0.2365420451)73.1/38:0.0266242596)81.9/67:0.0280299289,(((WP_115680924.1-Cupriavidus-taiwanensis:0.1576675538,WP_018313502.1-Cupriavidus-sp.-UYPR2.512:0.1954357559)30.3/59:0.0327271979,WP_135702508.1-Cupriavidus-oxalaticus:0.1263191419)82.3/56:0.0317075327,(WP_071014919.1-Cupriavidus-sp.-USMAHM13:0.1525332232,WP_174426450.1-Cupriavidus-basilensis:0.2589113970)88/79:0.0460681932)87.1/93:0.0305810473)75.1/38:0.0270712181)25.1/23:0.0257806920,(WP_116600700.1-Cupriavidus-alkaliphilus:0.1366310571,WP_228215938.1-Aromatoleum-toluclasticum:0.1697299172)89.5/31:0.0461708293)93.1/47:0.0354168951,(((WP_183165579.1-Azomonas-macrocytogenes:0.1318304221,((((WP_102836461.1-Stutzerimonas-stutzeri:0.0231087159,(WP_256071125.1-Stutzerimonas-stutzeri:0.0290887503,WP_045423249.1-Pseudomonadaceae:0.0232874644)86.5/98:0.0116706449)97.9/100:0.0416648519,WP_255961233.1-Stutzerimonas-stutzeri:0.0233032166)45.6/63:0.0031406393,WP_025243649.1-Stutzerimonas-stutzeri:0.0400675359)97.1/97:0.0447694548,(WP_122077451.1-Pseudomonadaceae:0.0285815414,((WP_037024448.1-Pseudomonas-saudiphocaensis:0.0582374958,WP_181084609.1-Stutzerimonas-stutzeri:0.0057497412)94.8/100:0.0295674789,(((WP_011911332.1-Stutzerimonas-stutzeri:0.0392834421,(WP_237255234.1-Pseudomonas-oligotrophica:0.0522409484,WP_043295157.1-Stutzerimonas-degradans:0.0195059286)69.8/100:0.0294994028)80.1/43:0.0111880283,WP_120997453.1-_Pseudomonas:0.0283430222)40.3/40:0.0098166636,(WP_230358311.1-Pseudomonas-sp.-NP21570:0.0136761150,(WP_193680295.1-Pseudomonas-lopnurensis:0.0385958858,WP_200589398.1-Stutzerimonas-stutzeri:0.0193955169)90.9/98:0.0290669058)89.4/75:0.0213562351)93.7/98:0.0281833920)75.6/79:0.0062886438)0/80:0.0000023811)63.6/96:0.0298334018)93.4/100:0.0432673400,WP_028238613.1-Stutzerimonas-azotifigens:0.0499384698)97.7/100:0.0697683875,WP_136680209.1-Neptunomonas-sp.-XY-337:0.1569534029)85.6/79:0.0294553944)35.6/32:0.0115759436,(WP_084055900.1-Desulfacinum-hydrothermale:0.2151406331,WP_123289658.1-Desulfosoma-caldarium:0.1925304892)99.6/99:0.1783476765)72.8/10:0.0107976081,((((((((WP_209862330.1-Pseudomonas-sp.-BP8:0.2931214274,WP_225786622.1-Pseudomonas:0.0339257208)81/94:0.0348336928,(((((WP_054891693.1-Pseudomonas:0.0659586458,WP_236238170.1-Pseudomonas-faucium:0.1111369570)72.1/98:0.0351632391,(WP_200141208.1-unclassified-Pseudomonas:0.1429226176,WP_029612831.1-Pseudomonas-parafulva:0.0261488860)96.9/100:0.0661193644)21.3/72:0.0165312908,WP_158461286.1-Pseudomonas-fluorescens:0.1617382827)65/98:0.0320256654,WP_110605176.1-Pseudomonas-soli:0.1726597864)65.3/78:0.0385962766,(((WP_217887155.1-Pseudomonas-xanthosomae:0.0341302256,WP_217840807.1-Pseudomonas-fakonensis:0.0301534251)89.3/100:0.0313568509,WP_186654015.1-Pseudomonas-xantholysinigenes:0.1481278966)57.3/96:0.0257392897,(WP_201176141.1-Pseudomonas-sp.-S31:0.0296972350,WP_261240966.1-unclassified-Pseudomonas:0.0341120920)97.9/100:0.0766415605)55.3/90:0.0208206289)78.2/73:0.0198233803)98.4/100:0.0712191193,((WP_181290220.1-Pseudomonas-brassicacearum:0.1247019474,(((WP_201004194.1-Pseudomonas-cichorii:0.0890754720,(WP_219063768.1-Pseudomonas-sp.-UMAB-08:0.1618945828,(WP_222540780.1-Pseudomonas-sp.-DR208:0.1023294509,(((WP_065875813.1-Pseudomonas:0.0364809294,(WP_033898696.1-Pseudomonas-fluorescens-group:0.0295559475,(WP_186687394.1-Pseudomonas-sp.-SWRI18:0.0457407966,WP_169378974.1-Pseudomonas:0.0127048982)81.9/100:0.0224360840)63.1/100:0.0110177033)98.9/100:0.0552996816,(WP_253509974.1-Pseudomonas-rhodesiae:0.0934410672,WP_093202821.1-Pseudomonas-yamanorum:0.0277944256)90.8/99:0.0311942938)33.5/92:0.0112814690,WP_105697656.1-Pseudomonas-poae:0.0420149084)61.3/92:0.0044592136)75.3/95:0.0286348138)61.9/100:0.0371110879)89/83:0.0310763729,((WP_249678359.1-Pseudomonas-sp.-PIA16:0.1034996437,WP_176569640.1-Pseudomonas-eucalypticola:0.1062194953)91.9/84:0.0423707414,WP_268799018.1-Pseudomonas-sp.-BSw22131:0.1384395525)48.9/55:0.0113843145)75.1/68:0.0130097944,WP_043192328.1-Pseudomonas-rhizosphaerae:0.1771023190)72.5/95:0.0393575862)90/100:0.0402200506,WP_108096457.1-Pseudomonas-sp.-GV071:0.0840654623)77.2/96:0.0493731868)86.6/99:0.0292490182,WP_252273535.1-Pseudomonas-sp.-YIM-B01952:0.0947697532)85.3/96:0.0263864464,WP_143488038.1-Pseudomonas-mangiferae:0.1069533534)89/96:0.0309389860,(((((((WP_058067682.1-Pseudomonas-sp.-TTU2014-080ASC:0.1201529432,WP_137821430.1-Pseudomonas-sp.-D-2018:0.0655487375)32.5/63:0.0284777088,WP_077525673.1-Pseudomonas-resinovorans:0.0931868324)95/97:0.0522470676,WP_028626762.1-Pseudomonas-resinovorans:0.0334702382)76.4/49:0.0068062369,(WP_120654068.1-Pseudomonas-sp.-DY-1:0.0711333142,WP_151138079.1-Pseudomonas-lalkuanensis:0.0363619710)86.9/85:0.0169942738)76.6/23:0.0061398430,(WP_016495240.1-Pseudomonas-resinovorans:0.0557605784,WP_044873077.1-Pseudomonas-sp.-LFM046:0.0460153195)87.9/88:0.0291211248)35.6/31:0.0165155682,WP_003453405.1-Pseudomonas:0.0961005619)46.6/88:0.0142123498,(WP_192399936.1-Pseudomonas-sp.-PDM15:0.1109451912,WP_226282979.1-Pseudomonas-lalkuanensis:0.0614545479)98.5/100:0.0688108366)87.4/64:0.0293441742)86.8/96:0.0167892008,((((WP_090448460.1-Pseudomonas-benzenivorans:0.1422188259,(WP_208707982.1-Pseudomonas-sp.-LPB0260:0.0276183090,WP_255840290.1-Pseudomonas-benzenivorans:0.0269147526)96.5/100:0.0457849748)50.5/37:0.0186030102,((WP_238040454.1-Pseudomonas-sp.-MMS21-TM103:0.1038145689,WP_069518708.1-Pseudomonas:0.0930029202)79.9/29:0.0206760794,((WP_131198269.1-Pseudomonas-dryadis:0.0790721134,(WP_131178052.1-Pseudomonas-daroniae:0.0492878477,(WP_084307530.1-Pseudomonas-flavescens:0.0578830880,(WP_249909557.1-Pseudomonas-fulva:0.0375448279,(WP_074889242.1-Pseudomonas-argentinensis:0.0231876334,WP_027910132.1-Pseudomonas-sp.-URMO17WK12_I4:0.0373814953)56.4/98:0.0126425249)82.9/99:0.0243812345)83.9/98:0.0285249486)96.3/100:0.0563123560)97.7/99:0.0666833794,(((((((WP_119694090.1-Pseudomonas:0.0393128211,WP_106738315.1-Pseudomonas-mendocina:0.0431776056)92.4/100:0.0323711522,WP_150606948.1-Pseudomonas-oleovorans:0.0483953831)93.9/98:0.0306677290,WP_024309956.1-Pseudomonas-sp.-P818:0.0351877413)80.8/63:0.0124830350,(WP_143497094.1-Pseudomonas-sp.-ALS1131:0.0329296869,WP_090337209.1-Pseudomonas-chengduensis:0.0191874284)0/63:0.0014482612)85.8/83:0.0157914601,WP_074939507.1-Pseudomonas-composti:0.0192444017)86.2/98:0.0146502409,WP_021444205.1-Pseudomonas-sp.-EGD-AK9:0.0438368677)86.6/96:0.0215891793,WP_159970704.1-Pseudomonas-sp.-8Z:0.0647084760)97.8/99:0.0446755075)76.9/95:0.0147094382)84.9/26:0.0215503411)26.8/23:0.0123963847,(WP_025164100.1-Pseudomonas-taeanensis:0.0479884797,WP_147170838.1-Pseudomonas-sp.-SJZ079:0.0053679940)97.2/100:0.0363340618)72.9/25:0.0105859766,(WP_233685494.1-Pseudomonas-anguilliseptica:0.0382103470,WP_257778987.1-Pseudomonas-sp.-32.2.56:0.0143856971)100/100:0.0911889510)95.8/73:0.0487459590)90.3/100:0.0323388985,(WP_220803742.1-Pseudomonas-sp.-NCCP-436:0.1269641266,WP_090225506.1-Pseudomonas-kuykendallii:0.0771671559)88.3/46:0.0351500639)94.4/79:0.0463559686)85.7/30:0.0130340984,((((WP_213445339.1-Variovorax-sp.-PCZ-1:0.2304158190,WP_226921657.1-Kinneretia-sp.-XES5:0.1386039760)92.2/98:0.0698761655,WP_199264031.1-Desulfobulbus-oligotrophicus:0.1798306124)48.2/45:0.0333758153,(WP_091467897.1-Methylophilus-rhizosphaerae:0.2178829062,WP_226440716.1-Quatrionicoccus-australiensis:0.1996631476)84.2/96:0.0296545376)97/61:0.0756616206,WP_025281494.1-Ectothiorhodospira-haloalkaliphila:0.3112071572)67.6/28:0.0072877235)95/92:0.0293614051)81.5/21:0.0219590147,((((((((((WP_198111724.1-Inhella-proteolytica:0.2626842698,((WP_037482909.1-Sphaerotilus-natans:0.1347447346,(WP_028996355.1-Azohydromonas-australica:0.0581633680,WP_066337818.1-Azohydromonas-lata:0.0792630613)97.7/100:0.0873747897)74.6/100:0.0547974765,WP_241396682.1-Pelomonas-sp.-CA6:0.1296669334)32.1/99:0.0522112815)97.8/100:0.1400660926,WP_220807643.1-Noviherbaspirillum-aridicola:0.2086693114)15.9/94:0.0467173966,(WP_160809027.1-Nitrosomonas-sp.-HPC101:0.2389541779,WP_101249932.1-Telmatospirillum-siberiense:0.2785861903)82.3/97:0.0468081468)90.2/92:0.0456127526,((WP_106935816.1-unclassified-Variovorax:0.1852506678,(((WP_126747644.1-Variovorax-sp.-DXTD-1:0.1111951883,((WP_192194832.1-Variovorax-sp.-38R:0.1603459306,WP_160553990.1-Xylophilus-rhododendri:0.1289064862)22.7/99:0.0331583406,WP_093441791.1-Variovorax-sp.-770b2:0.1122950566)89/99:0.0467077194)82.2/96:0.0215091359,WP_068630051.1-Variovorax-sp.-PAMC-28711:0.1794883566)83.1/95:0.0180347954,(WP_093207264.1-Variovorax-sp.-YR750:0.0829892918,WP_206177103.1-Variovorax-sp.-RKNM96:0.0897599032)88.8/97:0.0304821893)89.4/99:0.0260518376)79.9/98:0.0154500984,WP_219210863.1-Variovorax-boronicumulans:0.1400357930)92.5/99:0.0381853772)83.8/90:0.0216033250,WP_056898839.1-Pseudorhodoferax-sp.-Leaf274:0.1902741296)89.9/90:0.0270777587,((((WP_047193087.1-Schlegelella-brevitalea:0.1806347812,WP_019561187.1-Caldimonas-manganoxidans:0.1733144211)51.1/99:0.0807968462,WP_074865046.1-Atopomonas-hussainii:0.1627711317)94.3/95:0.0644244415,WP_154206467.1-Ralstonia-pickettii:0.1810307683)68.5/91:0.0346115337,WP_057673948.1-Curvibacter-sp.-PAE-UM:0.1249945261)83.8/89:0.0193316026)92/90:0.0302310058,(((WP_253536546.1-Ralstonia-sp.-21YRMH01-3:0.3060105971,WP_169420802.1-Ramlibacter-agri:0.1378791886)30.2/6:0.0262555936,(WP_223986959.1-Cupriavidus-pampae:0.0583690435,((WP_011517306.1-Cupriavidus-metallidurans:0.0449337271,WP_019451253.1-Cupriavidus-sp.-BIS7:0.0454542477)0/30:0.0000023492,WP_082079449.1-Cupriavidus-basilensis:0.0706175714)59/33:0.0166814471)99.9/100:0.1062390930)87.1/6:0.0185734502,((WP_074796726.1-Nitrosospira-briensis:0.0683335879,(WP_090882638.1-Nitrosovibrio-sp.-Nv6:0.0492587133,WP_097065525.1-Nitrosovibrio-sp.-Nv4:0.0305357436)88.5/100:0.0402203948)100/100:0.1907907457,(((((WP_026436726.1-Acidovorax-sp.-JHL-9:0.1395779028,(((((WP_169107474.1-Comamonas-suwonensis:0.0401359191,WP_219343559.1-Comamonas-sp.-Y33R10-2:0.0716160513)93.7/100:0.0404641166,(WP_003073420.1-Comamonas:0.0587817057,((WP_211457163.1-Comamonas-sp.-PE63:0.0437076904,WP_148729134.1-Comamonas-sp.-Z3:0.0363831589)21.8/84:0.0057058321,WP_034391098.1-Comamonas-thiooxydans:0.0149070776)72.7/84:0.0080022713)90.9/100:0.0250345569)81.6/100:0.0180847538,WP_218241177.1-Comamonas-fluminis:0.0870803818)99.6/100:0.1195009164,(WP_012203208.1-Delftia-acidovorans:0.1421574059,(WP_066475960.1-Comamonas-terrae:0.1614022185,WP_058879441.1-Comamonas-kerstersii:0.0805435338)90.1/99:0.0474284828)0/29:0.0295419588)100/100:0.1532128728,WP_086926522.1-Acidovorax-carolinensis:0.1169206916)51/48:0.0298963852)80.8/98:0.0360246585,WP_174484170.1-methanotrophic-endosymbiont-of-Bathymodiolus-puteoserpentis:0.1414319345)83.8/92:0.0445847570,WP_201673630.1-Ramlibacter-monticola:0.2134536124)51.2/5:0.0231744730,WP_137890779.1-Ramlibacter-sp.-2FC:0.0735110882)87.8/5:0.0335984282,WP_137918285.1-Hydrogenophaga-sp.-2FB:0.1185965764)43.9/5:0.0181773475)44/5:0.0163895526)55.3/82:0.0222035663)87.1/81:0.0214082034,(WP_036252969.1-Methylobacter-sp.-BBA5.1:0.2071270362,WP_212785890.1-Ferrigenium-kumadai:0.1508714598)86.9/85:0.0430009862)84.3/76:0.0190309565,(((((WP_230291920.1-Croceicoccus-sp.-Ery5:0.3267468367,WP_164156863.1-Sandarakinorhabdus-rubra:0.1197628179)92.8/100:0.0859497250,((WP_070114691.1-Acidithiobacillus-caldus:0.2199176720,((WP_009566950.1-Acidithiobacillus:0.0382724894,WP_215870853.1-unclassified-Acidithiobacillus:0.0633476222)63.7/100:0.0431750582,(WP_101536718.1-Acidithiobacillus-sp.-SH:0.1410543181,WP_163095688.1-Acidithiobacillus-ferrianus:0.0322840630)61.4/97:0.0322673596)94.8/100:0.0841856412)79.3/99:0.0821386377,WP_152145882.1-Alkalilimnicola-sp.-S0819:0.2214223093)83.1/100:0.0718713987)86.5/99:0.0497661844,WP_122244186.1-Vandammella-animalimorsus:0.1925090591)86.5/85:0.0338854338,(WP_153584563.1-Caenimonas-koreensis:0.1773533491,(WP_007185580.1-Hydrocarboniphaga-effusa:0.0522266246,WP_007185565.1-Hydrocarboniphaga-effusa:0.0702922238)99.7/100:0.1564208943)98.5/99:0.1102293245)85.1/88:0.0303458697,((WP_108647113.1-Polynucleobacter-rarus:0.0236909220,WP_084281813.1-Polynucleobacter-kasalickyi:0.0326390545)100/100:0.3477960672,((WP_215371371.1-Polynucleobacter-sp.-MWH-UH24A:0.1565537634,WP_173942040.1-Polynucleobacter-antarcticus:0.1361260398)93/100:0.0750711462,((((WP_128113711.1-Polynucleobacter-necessarius:0.0940799160,WP_215385554.1-Polynucleobacter-sp.-AP-Kolm-20A-A1:0.0609297997)95.7/100:0.0662297338,WP_216177372.1-Polynucleobacter-sp.-AP-Feld-500C-C5:0.1442111652)85/100:0.0305569174,WP_114639791.1-Polynucleobacter-necessarius:0.0977756052)90.8/100:0.0472026661,((WP_216254756.1-Polynucleobacter-sp.-AP-Melu-500A-A1:0.0474647686,WP_216218975.1-Polynucleobacter-sp.-AP-Latsch-80-C2:0.0388098524)99.2/100:0.0810483107,(((((WP_215319389.1-Polynucleobacter-paneuropaeus:0.0814494473,(WP_215352354.1-Polynucleobacter-sp.-MWH-P3-07-1:0.0388328102,WP_215380622.1-Polynucleobacter-paludilacus:0.0340853626)87.8/100:0.0339517914)97.2/100:0.0495592815,WP_062307935.1-Polynucleobacter-sinensis:0.0473993529)30.7/88:0.0169058909,((((((WP_215376847.1-Polynucleobacter-asymbioticus:0.0235021345,WP_215328794.1-Polynucleobacter-sp.-AP-Sving-400A-A2:0.0484197341)99.4/100:0.0496064330,WP_173959551.1-Polynucleobacter-arcticus:0.0273235208)93.4/99:0.0219031223,WP_216225867.1-Polynucleobacter-sp.-MG-27-Goln-C1:0.0776701312)49.8/96:0.0060768464,(WP_216168716.1-Polynucleobacter-bastaniensis:0.0719900455,WP_216178894.1-Polynucleobacter-sp.-80A-SIGWE:0.0125825957)86.3/100:0.0154814515)88.4/98:0.0232034178,WP_215299358.1-Polynucleobacter-sp.-AP-Nino-20-G2:0.0782581501)88.1/83:0.0191628858,((WP_216268685.1-Polynucleobacter-sp.-Nonnen-W13:0.0397166470,WP_251374783.1-Polynucleobacter-corsicus:0.0323045437)37.9/91:0.0141936861,WP_215334312.1-Polynucleobacter-sp.-AP-Ainpum-60-G11:0.0389248232)76/38:0.0071633532)72.3/27:0.0057518604)19.6/19:0.0146139816,WP_216197047.1-Polynucleobacter-sp.-UK-Mo-2m-Kol15:0.0549425939)88.1/97:0.0292093501,WP_076024025.1-Polynucleobacter-sphagniphilus:0.0745634978)77.6/86:0.0150485252)95.9/97:0.0544852445)70.6/100:0.0382240060)99.6/100:0.1515097937)98.1/97:0.1313652798)83.8/71:0.0270013606)93.4/83:0.0449519487,WP_054772520.1-Methylogaea-oryzae:0.1610163716)23.7/6:0.0171241039)87.3/17:0.0239015375,(WP_217345508.1-Noviherbaspirillum-sp.-L7-7A:0.1731227370,WP_047549861.1-Methylotenera-sp.-G11:0.1670482958)85.9/99:0.0471503173)22.5/4:0.0145899458,(((((((((WP_051234278.1-Halomonas-halocynthiae:0.3748558297,WP_110648656.1-Salinicola-peritrichatus:0.2321011408)97.6/99:0.1692930331,((((((WP_066801876.1-Moraxella-oblonga:0.1119780063,(WP_197128702.1-Psychrobacter-sp.-NZS113:0.1341994207,WP_201616712.1-Psychrobacter-immobilis:0.1245844651)98.8/100:0.1254644248)96.1/100:0.1226020813,WP_036548095.1-Nitrincola-lacisaponensis:0.2609276115)89.3/94:0.0518535569,((WP_041157945.1-Halomonas-sp.-KHS3:0.0510842574,((WP_120386672.1-Halomonas-alkaliphila:0.1131758321,WP_226251132.1-unclassified-Halomonas:0.0524496974)4.2/38:0.0191811116,WP_141441391.1-Halomonas-titanicae:0.0588755519)93.2/82:0.0516021584)99.1/98:0.0943794970,WP_228219791.1-Halomonas-meridiana:0.0811229921)95.9/100:0.0636613955)79.8/89:0.0228377236,(((WP_081193716.1-Halomonas-sp.-BC1:0.1901372526,((((WP_224850395.1-Marinobacter-nauticus:0.1200954966,WP_110630630.1-Salinicola-salarius:0.0632863310)84.4/93:0.0357925434,(((WP_079551663.1-Halomonas-subglaciescola:0.0537794568,WP_195687443.1-Halomonas-sp.-KAO:0.0477319482)85.3/25:0.0201748813,WP_027961801.1-Halomonas-halodenitrificans:0.0237249290)76.2/18:0.0181800174,((WP_089675781.1-Halomonas-aquamarina:0.0422361331,WP_062362595.1-Halomonas-axialensis:0.0176540729)25/11:0.0118444939,(WP_253557393.1-unclassified-Halomonas:0.0364006999,WP_089676106.1-Halomonas-aquamarina:0.0235474995)0/95:0.0000029468)83.7/12:0.0089299239)93.9/25:0.0316781850)73.3/23:0.0082375622,(((WP_159342670.1-Halomonas-meridiana:0.2221020534,WP_253554573.1-Halomonas-sp.-DN3:0.1413052815)74.4/99:0.0228385025,WP_102589198.1-Halomonas-urumqiensis:0.0822445119)87/99:0.0191961533,(((WP_017842145.1-Methylotuvimicrobium-buryatense:0.2378543490,(WP_149282777.1-Halomonas-sp.-Y2R2:0.0631877977,WP_073436013.1-Halomonas-cupida:0.0624427036)75.8/98:0.0457223302)95.2/100:0.0682265694,WP_035566812.1-Halomonas:0.1749904102)90.6/97:0.0504190914,(WP_193343673.1-Halomonas-elongata:0.0522782220,WP_064701650.1-Halomonas-caseinilytica:0.0160372409)99.2/100:0.0797668342)93.4/96:0.0402561429)90.9/94:0.0213457899)89.4/22:0.0270057903,WP_193085952.1-Halomonas-sp.-FME65:0.0713052963)88.9/23:0.0311179625)92.7/93:0.0404000114,((WP_248234415.1-Marinobacter-bryozoorum:0.0734875565,(WP_228223924.1-Marinobacter-nauticus:0.0420724935,WP_098418752.1-Marinobacter-sp.-LV10MA510-1:0.1000926385)92.8/100:0.0550749724)92.1/99:0.0486301353,WP_021818206.1-Halomonas-huangheensis:0.1322318354)74.9/26:0.0485569447)86.2/86:0.0299342939,(WP_127061154.1-Halomonas-nanhaiensis:0.1264488158,(WP_096281135.1-Halomonas-sp.-JB37:0.1611434886,(WP_027958907.1-unclassified-Halomonas:0.0345566802,WP_064236373.1-unclassified-Halomonas:0.0436754232)96.8/100:0.0790149280)66.5/99:0.0382573900)66.5/77:0.0229150224)85.5/66:0.0455260574)84.3/62:0.0261681980,((((((WP_030071911.1-Halomonas-alkaliantarctica:0.0416961667,WP_095603285.1-Halomonas-humidisoli:0.0220237706)86.7/86:0.0172656337,WP_253485683.1-Halomonas-sp.-XH26:0.0245174068)77.2/86:0.0103642793,WP_038485954.1-Halomonas-campaniensis:0.0183562729)98.8/100:0.0644210031,WP_162218282.1-Halomonas-alkaliphila:0.0246330865)96.2/100:0.0402711010,WP_078087027.1-unclassified-Halomonas:0.0230276363)0/95:0.0000024617,WP_096655487.1-Halomonas-nigrificans:0.0405594783)99.3/100:0.0770296783)88/70:0.0415503104,(((((WP_158774340.1-Cobetia-sp.-L2A1:0.0603800148,WP_144727127.1-Cobetia-crustatorum:0.0250055432)15.3/83:0.0104969007,WP_088744253.1-Cobetia-sp.-QF-1:0.0515735056)92/92:0.0261249559,WP_158774127.1-Cobetia-sp.-L2A1:0.0193283533)72.9/92:0.0083319324,(WP_077373671.1-Cobetia:0.0414039945,((WP_213115235.1-Cobetia-sp.-MC34:0.0333282714,WP_200017180.1-Cobetia-sp.-cqz5-12:0.0490204768)94.6/100:0.0438895293,WP_225346840.1-Cobetia-amphilecti:0.0316600183)92.3/100:0.0324150592)91.9/99:0.0289977444)98/98:0.0819606737,(WP_251978361.1-unclassified-Salinicola:0.0615598870,WP_189443129.1-Salinicola-rhizosphaerae:0.0813692320)99.8/100:0.1189625855)91.4/94:0.0713905155)83.2/93:0.0606493741)87/41:0.0447682174,WP_179929642.1-Halomonas-salicampi:0.2288367423)59.5/40:0.0282049814,WP_254581577.1-Marinobacterium-sedimentorum:0.2387202565)94.2/99:0.0780831458,(WP_015904383.1-Desulforapulum-autotrophicum:0.1888624256,WP_111960646.1-Desulfobacter-hydrogenophilus:0.2290535272)82.4/100:0.0896845248)7.7/85:0.0664141262,WP_178968040.1-Neptunomonas-phycophila:0.2675532846)88.5/24:0.0958529643,(((WP_151033206.1-Cellvibrio-sp.-KY-GH-1:0.0732844061,WP_049631514.1-Cellvibrio-sp.-pealriver:0.0595360280)87.7/100:0.0600120435,WP_062059887.1-Cellvibrio-sp.-OA-2007:0.0439570483)99.9/100:0.2113986199,WP_156872703.1-Ketobacter-sp.-MCCC-1A13808:0.2662615484)35.6/22:0.0539422645)86.1/86:0.0571974839,((((((WP_125841544.1-Stutzerimonas-stutzeri:0.2423081091,((WP_262731117.1-Alcanivorax-marinus:0.1255906052,(WP_106232428.1-Halomonas-ventosae:0.0691200183,WP_228234339.1-Alcanivorax-marinus:0.1427798655)55.1/100:0.0342289257)84/99:0.0232168398,WP_175070852.1-Halomonas-taeanensis:0.0890351489)40.4/93:0.0747186362)75.4/99:0.0651762420,(WP_019018778.1-Halomonas-lutea:0.2114569373,WP_092999033.1-Thiohalomonas-denitrificans:0.0877359692)89/100:0.0998860731)87.2/100:0.0343486860,((WP_146294265.1-Euhalothece-natronophila:0.1645140869,WP_159299499.1-Spiribacter-sp.-SSL99:0.1704881761)91.9/100:0.0545118249,WP_011630054.1-Alkalilimnicola-ehrlichii:0.1434010862)84.6/88:0.0475570185)46.9/86:0.0484653494,(WP_127477393.1-Sulfurivermis-fontis:0.2005924644,WP_028988655.1-Thermithiobacillus-tepidarius:0.1881453131)41/92:0.0461891912)97.9/99:0.0943870596,WP_069857109.1-Desulfoplanes-formicivorans:0.2153774016)84.3/79:0.0343425368,((WP_116367876.1-Parahaliea-mediterranea:0.0586347576,WP_148063811.1-Parahaliea-aestuarii:0.0843122302)100/100:0.2107166451,WP_129645432.1-Steroidobacter-agariperforans:0.2585506000)98.1/99:0.1378120232)71.9/38:0.0337904635)13.1/87:0.0385421993,(((((((WP_132974196.1-Thiogranum-longum:0.1486883015,WP_201344172.1-Thiohalobacter-sp.-COW1:0.1181010754)99.4/100:0.1868018989,WP_198243301.1-methane-oxidizing-endosymbiont-of-Gigantopelta-aegis:0.2300789776)80.8/82:0.0850065181,((WP_110520589.1-Herpetosiphon-llansteffanensis:0.4568481686,(WP_239493201.1-Luteitalea-sp.-TBR-22:0.2439824416,WP_058867107.1-Chloracidobacterium-thermophilum:0.2558436519)90.6/100:0.1068435742)31.2/92:0.0716400622,WP_099324654.1-Candidatus-Kuenenia-stuttgartiensis:0.3788634663)93/80:0.0999693792)68.2/24:0.0415561840,(WP_161826415.1-Petrachloros-mirabilis:0.3510439666,WP_062153084.1-Beggiatoa-leptomitoformis:0.2435189606)93.4/43:0.0986661738)93/44:0.0558595420,((((((((((((WP_036226274.1-Mesoaciditoga-lauensis:0.1495141105,((((WP_077198159.1-Thermosipho-affectus:0.0683306854,WP_012057290.1-Thermosipho-melanesiensis:0.1414277453)84.4/100:0.0339965979,(WP_127352844.1-Caldicellulosiruptor-changbaiensis:0.0511552512,WP_014043315.1-Caldicellulosiruptor-acetigenus:0.0280159544)100/100:0.2799320315)87.8/100:0.0462754405,((((((WP_117708548.1-Fusobacterium-mortiferum:0.0000026465,(WP_204716698.1-Fusobacterium-mortiferum:0.0187498471,WP_204716813.1-Fusobacterium-mortiferum:0.3080593964)94.2/95:0.0459191574)70.4/95:0.0215411131,WP_115270465.1-Fusobacterium-necrogenes:0.0583915216)97.8/100:0.0712393731,(((WP_130889691.1-Fusobacterium-ulcerans:0.0381650470,(WP_005950923.1-Fusobacterium-varium:0.0378788375,(WP_070580525.1-Fusobacterium:0.0497248718,WP_227372109.1-Fusobacterium-ulcerans:0.0239172749)31.7/100:0.0115507982)87/100:0.0149621101)45.8/100:0.0137027011,(WP_241635364.1-Fusobacterium-gastrosuis:0.1020336576,(WP_074518138.1-Fusobacterium-necrophorum:0.0679426398,(WP_211495349.1-Fusobacterium-necrophorum:0.1273220448,WP_032890075.1-Fusobacterium-nucleatum:0.4314917283)87.6/99:0.0502588536)94.4/99:0.0590212517)98.6/99:0.0678731324)36.7/96:0.0273480092,WP_261793436.1-Fusobacterium-varium:0.2289457948)92.8/99:0.0511001602)95.5/100:0.1072702118,((WP_147004474.1-Leptotrichia-wadei:0.0248801504,WP_018499460.1-Leptotrichia-wadei:0.0301634363)98.1/100:0.0889606123,((WP_068155536.1-Leptotrichia-sp.-oral-taxon-847:0.0740593074,(WP_211617428.1-Leptotrichia-sp.-oral-taxon-218:0.0134804801,WP_094080762.1-Leptotrichia-sp.-oral-taxon-498:0.0429500329)96/100:0.0566460806)43.9/94:0.0291182969,WP_099992503.1-Fusobacterium-pseudoperiodonticum:0.0970205628)33.7/98:0.0393922743)99.8/100:0.1773381382)84.1/97:0.0685196834,WP_173194267.1-Paenibacillus-frigoriresistens:0.2275063510)70.6/23:0.0367803608,(WP_013388736.1-Ilyobacter-polytropus:0.3377003900,WP_012548442.1-Dictyoglomus-thermophilum:0.2531655199)61/22:0.0269154254)96.3/100:0.0901612540)93.2/100:0.0514449731,WP_047753768.1-Kosmotoga-pacifica:0.1311842653)49.2/99:0.0351089791)97.5/100:0.0918885266,(WP_099829233.1-Mesotoga-sp.-Brook.08.105.5.1:0.1353128714,WP_099830786.1-Mesotoga-sp.-Brook.08.105.5.1:0.1454218752)97/100:0.0932933758)96.4/100:0.1039752626,((((((((((((((((((((((((WP_054742706.1-Cellulosilyticum-ruminicola:0.0707712398,(WP_177671917.1-Zhenhengia-yiwuensis:0.0455765312,WP_053982981.1-Niameybacter-massiliensis:0.0900897047)98.9/100:0.0947345625)86.6/100:0.0342359567,WP_069999280.1-Cellulosilyticum-sp.-I15G10I2:0.1977424897)98.9/68:0.0907487161,WP_055666468.1-Desnuesiella-massiliensis:0.1390511852)4.6/24:0.0175862830,WP_072892386.1-Clostridium-fallax:0.0726474893)92.7/25:0.0408039805,(WP_073337580.1-Clostridium-grantii:0.1160005100,WP_072832886.1-Clostridium-collagenovorans:0.0312552457)78.9/95:0.0161009164)92.1/24:0.0241343225,(((WP_152890015.1-Clostridium-tarantellae:0.1181970295,WP_195972826.1-Clostridium-thermobutyricum:0.0711770329)61.8/58:0.0317859585,(WP_209795868.1-Clostridium-moniliforme:0.0531496412,WP_194191032.1-Clostridium-chrysemydis:0.0953037854)96.3/100:0.0532292799)77.7/43:0.0109946896,WP_040194942.1-Clostridium-culturomicium:0.0564959635)76.3/44:0.0100529046)25.6/48:0.0117696071,(((WP_094369216.1-Romboutsia-weinsteinii:0.1217504581,WP_021359979.1-Clostridioides-difficile:0.2564097048)61.7/94:0.0410815877,(WP_074918830.1-Terrisporobacter-glycolicus:0.0525016153,WP_195929149.1-Clostridium-sp.-1001270J_160509_D11:0.0402161002)84.3/91:0.0190942794)96.4/89:0.0463537782,((((((((WP_092726456.1-Romboutsia-lituseburensis:0.1539095224,WP_258242885.1-Romboutsia-lituseburensis:0.0299500748)99/100:0.0784174339,WP_057560648.1-Paeniclostridium-sordellii:0.0541240397)90/73:0.0295557168,(WP_057580750.1-Paeniclostridium-sordellii:0.1005624127,WP_057538715.1-Paeniclostridium-sordellii:0.0329894596)70.6/99:0.0187451820)35.1/27:0.0138477860,WP_195252007.1-Romboutsia-sp.-1001713B170207_170306_H8:0.0390053438)91.9/71:0.0343010062,(((((WP_055265352.1-Clostridium:0.0365031693,(WP_186859336.1-Clostridium-sp.-NSJ-6:0.0558353135,(WP_055265304.1-Clostridium-disporicum:0.0539317093,(WP_154821366.1-Clostridium-saudiense:0.0072043978,WP_195245504.1-Clostridium-celatum:0.0615627394)65/99:0.0248371835)21/87:0.0274329609)57.7/88:0.0439846527)77/100:0.0584780081,WP_186834444.1-Eubacteriales:0.1037383814)96.4/100:0.0540136623,((WP_235068992.1-Turicibacter-sp.-TJ11:0.0526096853,((WP_161831114.1-Turicibacter-sp.-TS3:0.0486578421,WP_155223812.1-Turicibacter-sanguinis:0.0226603331)75.9/82:0.0151370158,WP_195933776.1-Turicibacter-sanguinis:0.0224084316)54.3/80:0.0060118961)86.7/92:0.0173142839,WP_212726030.1-Turicibacter-bilis:0.0172077872)100/100:0.0824071838)0/91:0.0086574398,(WP_066874757.1-Clostridium-mediterraneense:0.1638280014,WP_198610231.1-Clostridium-perfringens:0.0191982615)66.5/73:0.0444997138)35.1/63:0.0139266496,WP_055258173.1-Sarcina-ventriculi:0.0952850104)96.7/86:0.0581484563)76.6/8:0.0112508788,(((WP_195959433.1-Clostridium-tertium:0.0603960154,WP_195429094.1-Clostridium-sp.-D46t1_190503_E9:0.0176457924)83.5/95:0.0156991464,WP_047001949.1-Clostridium-sp.-C8:0.0530959112)83.5/90:0.0124031815,(WP_066890539.1-Clostridium-nigeriense:0.0434712446,WP_016207773.1-Clostridium-sartagoforme:0.0313069023)46.7/44:0.0057183729)97.3/92:0.0331464430)62/4:0.0127006210,WP_095406848.1-Romboutsia-maritimum:0.1224749870)0/6:0.0000028538,(((WP_029452249.1-Clostridium-algidicarnis:0.0897387854,WP_216183125.1-Clostridium-gasigenes:0.0194268678)52.6/99:0.0311283471,WP_216313757.1-Clostridium-gasigenes:0.0618491610)96.7/100:0.0545722518,((WP_163195303.1-Clostridium-thermarum:0.1230062785,WP_066677671.1-Clostridium-septicum:0.0260999754)81.4/57:0.0147963418,(WP_197096717.1-Paeniclostridium-sordellii:0.0718196517,WP_238903698.1-Clostridium-sp.-YIM-B02506:0.0556388898)57.4/82:0.0204457578)46.4/10:0.0070153293)91/49:0.0180669985)67.9/7:0.0124995790)90.2/61:0.0207962066)92.8/91:0.0247194943,(WP_072990516.1-Clostridium-cavendishii:0.2029022597,((((WP_216288229.1-Clostridium-psychrophilum:0.1205322772,WP_216137569.1-Clostridium-estertheticum:0.0336805614)0/100:0.0000028891,WP_220704225.1-Clostridium-estertheticum:0.0356403493)92.3/100:0.0427433792,WP_220321605.1-Clostridium-tagluense:0.0548026290)98.7/100:0.0786096520,WP_133628539.1-Fonticella-tunisiensis:0.0627828186)73.3/98:0.0111069213)93.3/93:0.0271043012)57.7/87:0.0081820900,((((((((((WP_169105633.1-Clostridium-sp.-SM-530-WT-3G:0.0534714475,WP_027636628.1-Clostridium-butyricum:0.0805418386)56.8/45:0.0302492988,WP_077866663.1-Clostridium-saccharobutylicum:0.1667061673)89/98:0.0217064096,WP_261829154.1-Clostridium-sp.-TW13:0.0596775597)62.7/45:0.0111494444,(((((WP_185652834.1-Clostridium-sp.-DJ247:0.0179544757,WP_185652641.1-Clostridium-sp.-DJ247:0.0628525355)3.8/99:0.0175255237,WP_226126241.1-Clostridium-algoriphilum:0.1275475031)79.8/11:0.0108424448,(((WP_022750218.1-Clostridium-saccharobutylicum:0.0389355365,WP_238915641.1-Clostridium-sp.-YIM-B02555:0.0514510200)7.2/3:0.0118123028,WP_077846315.1-Clostridium-puniceum:0.0596344068)43.4/3:0.0122779444,WP_015391310.1-Clostridium-saccharoperbutylacetonicum:0.0501120206)96.1/23:0.0307970097)72.4/21:0.0057650495,((((((WP_009169246.1-unclassified-Clostridium:0.0937088476,WP_077843022.1-Clostridium-beijerinckii:0.1223836005)87/100:0.0308740369,WP_041896941.1-Clostridium-beijerinckii:0.1607419775)91/100:0.0439259272,WP_069680896.1-Clostridium-taeniosporum:0.1782390660)92.5/100:0.0475514015,WP_216390068.1-Clostridium-sp.-MSJ-8:0.2345732697)53.1/45:0.0324718934,(WP_265859166.1-Clostridium-botulinum:0.0976764180,((WP_154531076.1-Inconstantimicrobium-porci:0.1264365807,WP_009169009.1-Clostridium-sp.-DL-VIII:0.0167853321)75.2/99:0.0457878044,WP_077856313.1-Clostridium-beijerinckii:0.0000020180)96.1/99:0.0547006954)89.8/46:0.0286746716)89.2/93:0.0193089118,((WP_160359894.1-Clostridium-chromiireducens:0.2058484763,WP_077848125.1-Clostridium-puniceum:0.0830517096)92.1/95:0.0491061161,(WP_077855754.1-Clostridium-beijerinckii:0.0359571985,WP_041894754.1-Clostridium-beijerinckii:0.0216829185)99.7/100:0.0832844514)0/25:0.0000027386)30.7/3:0.0132220019)83.6/10:0.0137140809,WP_224033465.1-Clostridium-gelidum:0.0867974392)87.8/34:0.0183293412)97.7/99:0.0790092898,WP_148566871.1-Acetobacterium-paludosum:0.2754088901)74.8/89:0.0363675586,WP_055069574.1-Clostridium-massiliamazoniense:0.1204187331)74/38:0.0121485872,(WP_106059304.1-Clostridium-vincentii:0.0856434588,WP_143316579.1-Clostridium:0.0254119057)94.9/100:0.0364256262)40/41:0.0068733583,WP_090010800.1-Clostridium-sp.-DSM-8431:0.1137681383)86.3/91:0.0124946319,WP_055184115.1-Clostridium-paraputrificum:0.0559354321)62/91:0.0124613813,WP_261829490.1-Clostridium-sp.-TW13:0.1696453043)40.5/90:0.0113523905)91.6/97:0.0317075620,WP_066825124.1-Clostridium-tepidiprofundi:0.0860416653)94/97:0.0434298753,WP_072889493.1-Tepidibacter-formicigenes:0.0312957197)98.3/100:0.0625763442,WP_072725426.1-Tepidibacter-thalassicus:0.1363664268)86.6/99:0.0406032479,WP_095404508.1-Romboutsia-maritimum:0.2435091583)86/86:0.0289728170,((WP_142413383.1-Hathewaya-massiliensis:0.1470936621,WP_072903743.1-Hathewaya-proteolytica:0.2591263341)27.7/82:0.0289942369,WP_035130905.1-Clostridium-sulfidigenes:0.1738686072)98.6/99:0.0910157477)90.5/84:0.0430869174,((((WP_227847393.1-Clostridioides-sp.-ZZV15-6598:0.0940123863,WP_227846117.1-Clostridioides-sp.-ES-S-0145-01:0.0299499723)94.9/100:0.0395301506,(WP_227842647.1-unclassified-Clostridioides:0.0303226834,WP_003420982.1-Clostridioides-difficile:0.0806553095)85.8/95:0.0198279396)72.4/95:0.0227057185,WP_038810653.1-Clostridioides-difficile:0.0338915815)100/100:0.2142054183,WP_226126924.1-Clostridium-algoriphilum:0.2441769276)66.8/76:0.0510629179)75.8/68:0.0271571675,WP_099190526.1-Tepidibacter-mesophilus:0.1115085635)62/87:0.0446101674,WP_079421587.1-Clostridium-oryzae:0.2895090309)83.7/94:0.0589871000,(WP_112093115.1-Methanothermobacter-tenebrarum:0.2200277037,WP_013413362.1-Methanothermus-fervidus:0.1629753887)97.5/100:0.1164341172)56.7/73:0.0266685918,((((((WP_088538685.1-Methanobrevibacter-sp.-87.7:0.0533458691,WP_016359256.1-Methanobrevibacter-sp.-AbM4:0.1052991732)98.3/100:0.0886414781,WP_067148181.1-Methanobrevibacter-olleyae:0.1030665351)85.8/100:0.0345670843,WP_116670296.1-Methanobrevibacter-woesei:0.1952003422)99.8/60:0.1105688489,(WP_248612240.1-Methanobacterium-alcaliphilum:0.1218143286,((WP_071907175.1-Methanobacterium-congolense:0.0190455096,WP_209584978.1-Methanobacterium-aggregans:0.0414476584)99.5/100:0.1010488657,(WP_069583666.1-Methanobacterium:0.1467412091,WP_048190917.1-Methanobacterium-sp.-SMA-27:0.1104094082)91.4/96:0.0480520709)2.3/51:0.0156534520)91/96:0.0298648042)73.3/54:0.0148483552,(WP_011406138.1-Methanosphaera:0.1288297164,(WP_112123902.1-Methanosphaera-sp.-BMS:0.0854812333,(WP_069592847.1-Methanosphaera-sp.-WGK6:0.0290976630,WP_095608065.1-Methanosphaera-cuniculi:0.0936324995)97.2/100:0.0715542741)53.9/89:0.0431048821)96.8/100:0.0615932483)85.9/58:0.0244900868,(WP_013644504.1-Methanobacterium-lacus:0.1777687870,WP_054834988.1-Methanobrevibacter-arboriphilus:0.1447345896)88.9/99:0.0366999500)98.8/59:0.0652788682)83.5/34:0.0229802707,((((((WP_068555806.1-Thermotalea-metallivorans:0.1028413129,WP_073236503.1-Desulforamulus-putei:0.0934933837)91.6/99:0.0352499380,WP_090442301.1-Natronincola-peptidivorans:0.1353986771)81.2/93:0.0244284457,WP_035164300.1-Caloranaerobacter-azorensis:0.0626697145)67.9/93:0.0228916898,WP_095133872.1-Anaeromicrobium-sediminis:0.1553360836)71.2/95:0.0277943884,((((((WP_252130283.1-Clostridium-sp.-SYSU_GA19001:0.2169590627,WP_202749105.1-Clostridium-sp.-YIM-B02515:0.1143570256)99/99:0.0992666755,WP_216462460.1-Clostridium-bornimense:0.1374738549)86.5/16:0.0186268447,(((WP_104410444.1-Clostridium-algidicarnis:0.0529357416,WP_185162668.1-Clostridium-algidicarnis:0.0462054426)100/100:0.1289203887,WP_234123145.1-Clostridium-hydrogenum:0.1660813160)85.3/95:0.0300268344,((WP_084114600.1-Clostridium-acidisoli:0.0867285096,(WP_015614219.1-Clostridium-pasteurianum:0.0248424184,(WP_010235680.1-Clostridium-arbusti:0.0116527662,WP_003443690.1-Clostridium-pasteurianum:0.0467026229)87.3/100:0.0219127700)96.7/100:0.0324763188)0/27:0.0110313695,WP_026884435.1-Clostridium-akagii:0.0889928914)97.9/100:0.0704775730)93/93:0.0301206921)28.8/7:0.0130252947,((WP_090038599.1-Clostridium-frigidicarnis:0.1207927190,WP_209459870.1-Youngiibacter-multivorans:0.1688121213)95.6/100:0.0745609980,((WP_164955553.1-Clostridium-estertheticum:0.1584432941,WP_125002390.1-Clostridium-tagluense:0.1333113192)38.8/98:0.0164286794,(WP_226136064.1-Clostridium-estertheticum:0.0246176013,WP_216146875.1-Clostridium-frigoris:0.0612996117)99.7/100:0.0941645379)98.5/100:0.0913807549)60.8/100:0.0231468158)65.4/27:0.0446987881,((WP_165412483.1-Clostridium-estertheticum:0.0576212953,WP_216302578.1-unclassified-Clostridium:0.0098831605)100/100:0.1654848300,((WP_204598070.1-Clostridium-pascui:0.1147921405,WP_216457288.1-Clostridium-simiarum:0.1217884129)7.8/62:0.0297937272,WP_027632043.1-Clostridium-hydrogeniformans:0.1204251325)22.9/98:0.0178813090)88.9/99:0.0466741948)0/14:0.0282890646,(((((WP_066398085.1-Cytobacillus-eiseniae:0.2121864434,(WP_056034132.1-Paenibacillus-sp.-Leaf72:0.0929982012,WP_197936429.1-Paenibacillus-sp.-GSMTC-2017:0.1165313486)94.1/99:0.0636202576)82.6/96:0.0331295432,(WP_076307499.1-Paenibacillus-odorifer:0.0517521586,((WP_042140168.1-Paenibacillus-sp.-FSL-P4-0081:0.0087629225,WP_036721118.1-Paenibacillus-sp.-FSL-R7-277:0.0634454073)92.6/100:0.0323999916,WP_074087222.1-Paenibacillus-sp.-P32E:0.0303016092)95.7/100:0.0756887044)98.3/100:0.0915358459)88.9/96:0.0472240198,(WP_262684732.1-Paenibacillus-sp.-WQ-127069:0.1274702544,WP_090798944.1-Paenibacillus-sp.-GP183:0.2037776061)42.1/95:0.0276201417)93.3/100:0.0731148479,WP_036935152.1-Pseudobacteroides-cellulosolvens:0.1379573561)85.8/98:0.0558474855,(WP_009529486.1-Peptoanaerobacter-stomatis:0.1116700557,WP_068913445.1-Criibacterium-bergeronii:0.1588158039)100/100:0.2007988775)85.5/85:0.0388831392)85.4/93:0.0360691635)83.3/79:0.0263000596,(((((WP_266162911.1-Caproicibacterium-sp.-BJN0003:0.1252250515,WP_256135338.1-Bittarella-massiliensis:0.2160641410)94.1/100:0.0969184095,WP_220321633.1-Clostridium-tagluense:0.2047789532)90/99:0.0767961746,(WP_105204772.1-Neobittarella-massiliensis:0.1059093482,WP_022475313.1-Huintestinicola-butyrica:0.2408526010)97.1/100:0.0779958960)81.5/99:0.0472401613,(((WP_091684654.1-Anaerocolumna-aminovalerica:0.1581408645,(WP_033168390.1-Clostridium-sp.-KNHs205:0.0787596783,((WP_073588199.1-Anaerocolumna-xylanovorans:0.0304807132,WP_185256524.1-Anaerocolumna-chitinilytica:0.1127450734)55.7/100:0.0382423728,WP_073279502.1-Anaerocolumna-jejuensis:0.0928922166)79.6/100:0.0423183573)99.5/100:0.1372157006)97.2/100:0.0839338517,((((WP_060930762.1-Lachnoanaerobaculum-saburreum:0.1640140947,((((((((((((WP_024837362.1-Lacrimispora:0.0841415690,WP_077612318.1-Clostridium-sp.-Marseille-P2415:0.0326086239)72.5/97:0.0059140053,(WP_024293365.1-Lachnospiraceae:0.0345973130,WP_092242699.1-Lacrimispora:0.0233891397)95.7/100:0.0328134906)88/97:0.0155124880,((WP_124065139.1-Clostridium-sp.-E02:0.0799353744,WP_117415169.1-Clostridium-indicum:0.0421218222)75.9/94:0.0078921655,WP_143321893.1-Clostridium-sp.-HBUAS56010:0.0285977278)87.8/94:0.0148835950)77.4/97:0.0088297156,WP_138313639.1-Hungatella-hathewayi:0.0252485999)89/96:0.0212297814,WP_186424215.1-Lacrimispora-celerecrescens:0.1283924610)95.3/94:0.0445466785,(((((WP_045093538.1-Clostridium-sp.-FS41:0.0411611659,WP_025485467.1-Lachnospiraceae:0.0536568565)91.3/88:0.0176175444,(WP_154471253.1-Clostridium-porci:0.1107451165,WP_178313416.1-Lacrimispora-sp.-210928-DFI.3.58:0.0005047603)87.2/51:0.0170506045)0/31:0.0000027801,WP_002577448.1-Enterocloster:0.0647888877)46.8/49:0.0230149512,WP_066545911.1-Clostridium-sp.-AT4:0.0356507845)71.8/34:0.0055873931,WP_024736308.1-Enterocloster-asparagiformis:0.0351369117)72.4/35:0.0065296698)80.9/84:0.0121812600,((WP_251497048.1-Otoolea-sp.-DSM-111138:0.1329740327,(WP_003497689.1-_Clostridium:0.0413945022,WP_125142063.1-Clostridium-transplantifaecale:0.0420413086)90.4/95:0.0270439620)57.3/35:0.0255484266,((((WP_087223935.1-Lachnoclostridium-sp.-An14:0.0384172547,WP_101876396.1-Lachnoclostridium-edouardi:0.0474176337)93.6/100:0.0266732473,WP_118243934.1-Clostridium-sp.-AM58-1XD:0.0576076568)73.1/21:0.0068760049,WP_118742831.1-Clostridium-sp.-AM28-20LB:0.0289052424)87/21:0.0117778730,WP_008399284.1-unclassified-Clostridium:0.0176790042)0/19:0.0000022193)95.1/89:0.0396322283)40.7/14:0.0068273176,WP_215700061.1-Clostridium-sp.-MCC353:0.0569714750)77.6/31:0.0098781436,WP_118541387.1-unclassified-Clostridium:0.0644477113)43.6/80:0.0153721868,((WP_215718428.1-Anaerostipes-hadrus:0.0114899601,WP_075760385.1-Anaerostipes-sp.-494a:0.0478098909)96.6/100:0.0387253416,((WP_227052204.1-Anaerostipes-caccae:0.0317165329,WP_137330070.1-Anaerostipes-rhamnosivorans:0.0703587549)99.2/100:0.0855812441,WP_006781439.1-Hungatella-hathewayi:0.0773159743)88.6/99:0.0341101587)85.4/92:0.0173290898)58.4/77:0.0276207893,(WP_231062061.1-Lacrimispora-sp.-NSJ-141:0.0698130157,WP_249304640.1-Qiania-dongpingensis:0.0678017438)92.7/98:0.0343715313)94.1/92:0.0528983599,WP_118555425.1-Lachnotalea-sp.-AF33-28:0.1461378691)88.3/99:0.0558644324)91.1/100:0.0772572725,WP_005539496.1-Johnsonella-ignava:0.1752282344)28.3/54:0.0389308428,(((((((((((((((WP_035764434.1-Butyrivibrio-sp.-NC2002:0.0920210005,((WP_026526812.1-Butyrivibrio-sp.-VCD2006:0.1369870311,WP_026491547.1-unclassified-Butyrivibrio:0.0159309604)40.6/96:0.0342785330,WP_044913869.1-Butyrivibrio-sp.-WCE2006:0.0210566817)93.5/100:0.0541493525)0/31:0.0334773846,WP_091822189.1-Butyrivibrio-sp.-ob235:0.0485015646)99.6/100:0.1080925829,(((((((WP_022768176.1-unclassified-Butyrivibrio:0.0196297130,WP_022766545.1-Butyrivibrio-sp.-XPD2006:0.0478965909)99.4/100:0.0800229899,WP_034446055.1-Butyrivibrio-sp.-AE2032:0.0535064152)93.4/100:0.0426756825,WP_024867291.1-Butyrivibrio-sp.-FCS014:0.0245513872)87.8/50:0.0244225272,(WP_071175335.1-Butyrivibrio:0.0570948395,WP_026513668.1-Butyrivibrio-sp.-LB2008:0.0409413835)79.4/50:0.0321637082)82.6/99:0.0227630314,WP_026663187.1-Butyrivibrio-proteoclasticus:0.0604684253)56.9/54:0.0146001961,(WP_013279951.1-Butyrivibrio-proteoclasticus:0.0275636067,WP_092060841.1-Butyrivibrio-sp.-INlla14:0.0420034756)86.1/85:0.0146605438)0/17:0.0000024223,WP_026652320.1-Butyrivibrio-proteoclasticus:0.0371175638)96.9/89:0.0713839667)96.7/100:0.0824170519,((WP_019160740.1-Blautia:0.0546322112,WP_173738571.1-Blautia-schinkii:0.0230179054)88/100:0.0247553963,(WP_158421413.1-Blautia:0.0344909164,(WP_148445023.1-Eubacteriales:0.0269306520,WP_187566223.1-Blautia-faecis:0.0438770313)90.3/100:0.0249146104)68.5/87:0.0168448456)98.1/99:0.0717582104)95.6/96:0.0442605313,WP_178047055.1-Fusicatenibacter-sp.-CLA-AA-H277:0.0860809488)1/40:0.0115309500,WP_109624131.1-Murimonas-intestini:0.0771717535)78.8/96:0.0247876449,(WP_173751851.1-Blautia-caecimuris:0.0446038801,WP_117997894.1-Blautia-obeum:0.0124665422)98.8/100:0.0574637298)68.7/80:0.0287604317,(((WP_158367551.1-Lachnospiraceae:0.0253862070,WP_235247881.1-Faecalicatena-contorta:0.0710904466)84.7/95:0.0379530449,(WP_251390531.1-Mediterraneibacter-sp.-DSM-111629:0.0313535618,WP_073962323.1-_Ruminococcus:0.0297483670)89.6/88:0.0233543655)82.3/64:0.0137278718,WP_070089756.1-Merdimonas-faecis:0.0294861355)78.9/62:0.0161845094)0/25:0.0024606292,WP_230400100.1-Novisyntrophococcus-fermenticellae:0.1511882619)88.3/97:0.0277169638,WP_132379639.1-Muricomes-intestini:0.0881378086)93.8/100:0.0414107396,WP_087156947.1-Lachnoclostridium-sp.-An181:0.0474407100)21.1/46:0.0228110065,(WP_073994331.1-Parasporobacterium-paucivorans:0.1215664804,(WP_099466994.1-Konateibacter-massiliensis:0.0653957594,WP_151145296.1-Candidatus-Galacturonibacter-soehngenii:0.0483323506)97.6/100:0.0792119752)51.7/74:0.0501846923)30.2/75:0.0174218149,WP_115483253.1-Anaerosacchariphilus-polymeriproducens:0.0988900649)24.9/68:0.0373850209,WP_158351687.1-Diplocloster-modestus:0.1873805140)45.2/98:0.0436152714,WP_158346166.1-Lachnospiraceae:0.1844894082)98.1/98:0.1029858613)51.1/89:0.0245639381,WP_226914794.1-Intestinibacter-bartlettii:0.2442288531)88.6/24:0.0395808849)27.5/18:0.0291191379,((((WP_073273890.1-Anaerocolumna-jejuensis:0.1534295633,(((WP_010268185.1-Paenibacillus-senegalensis:0.1691480044,WP_155699987.1-Paenibacillus-psychroresistens:0.0524542641)59.2/58:0.0306455837,WP_090040610.1-Clostridium-frigidicarnis:0.0234398911)85.1/58:0.0285129586,WP_202751253.1-Clostridium-sp.-YIM-B02515:0.0338401031)96/93:0.0649930964)0/57:0.0191412266,WP_033167869.1-Clostridium-sp.-KNHs205:0.1407604627)99.4/100:0.1392938803,(WP_249305773.1-Jutongia-hominis:0.1486179730,(WP_022463459.1-Jutongia-huaianensis:0.0663170298,(WP_055944671.1-unclassified-Clostridium:0.0355019731,WP_216564272.1-Eubacterium-sp.-MSJ-13:0.0276076955)94.9/100:0.0466358109)27.5/99:0.0410118035)99.8/100:0.1480525972)72.2/100:0.0408682707,(WP_253721803.1-Treponema-sp.-OMZ-787:0.3001050313,WP_058352784.1-Acetivibrio-ethanolgignens:0.1712775285)66.7/99:0.0613895279)96.4/99:0.0849093969)75.3/90:0.0925835315)92.4/93:0.0827715234,((((WP_069974893.1-Geosporobacter-ferrireducens:0.2902211097,WP_218283043.1-Crassaminicella-sp.-143-21:0.1877421659)31.8/86:0.0543409886,(WP_223038819.1-Crassaminicella-profunda:0.0842567361,WP_148809648.1-Crassaminicella-thermophila:0.0784001027)72.3/98:0.0237206724)98.7/100:0.1118006302,WP_250858284.1-Oceanirhabdus-seepicola:0.1753769119)81.4/100:0.0393487563,WP_160686059.1-Clostridium-sp.-C2-6-12:0.1968441857)86.9/86:0.0583260971)69.4/34:0.0293091958)95.2/70:0.0557994914)67.1/64:0.0172976617,(WP_111570784.1-Halanaerobium-saccharolyticum:0.0554066449,(WP_089717478.1-Halanaerobium-congolense:0.0485302520,WP_089862761.1-Halanaerobium-salsuginis:0.0258837420)93.7/100:0.0472305422)100/100:0.2195617009)47.6/26:0.0180522421,((((WP_058965344.1-Fournierella-massiliensis:0.1486508772,WP_185752312.1-Butyrivibrio-sp.-AE2032:0.2611197308)49.4/99:0.0540527712,WP_155202396.1-Ruthenibacterium-lactatiformans:0.1538739834)94/100:0.0817557986,(WP_186886939.1-Anaerofilum-hominis:0.1507784629,WP_102049807.1-Pygmaiobacter-massiliensis:0.1716016653)73.4/98:0.0533384573)99.9/100:0.1815817090,WP_038266426.1-Peptoclostridium-litorale:0.2529453351)94.2/5:0.0793034162)87.2/5:0.0273956309,(WP_119970050.1-Clostridium-fermenticellae:0.2279034714,((WP_213236459.1-Fusibacter-paucivorans:0.3983111880,(((((((((((WP_160564586.1-Apibacter-sp.-B2912:0.0995092464,(WP_055424523.1-Apibacter-mensalis:0.0784011727,(WP_160568957.1-unclassified-Apibacter:0.0146416675,WP_166004639.1-Apibacter-sp.-B2966:0.0426602890)98.9/100:0.0798289086)81.4/100:0.0505267011)96.5/100:0.0843798412,(WP_105245553.1-Apibacter-adventoris:0.1698046915,WP_146261858.1-Apibacter-muscae:0.1066900269)85.8/100:0.0489199217)99.9/100:0.1857596735,(WP_084015419.1-Moheibacter-sediminis:0.2787662160,WP_182042771.1-Moheibacter-lacus:0.1988171285)94/100:0.0847615268)9.4/88:0.0390341688,(WP_068596492.1-Vaginella-massiliensis:0.1310949153,((WP_230477342.1-Faecalibacter-bovis:0.0988235730,WP_125349156.1-Empedobacter-falsenii:0.0878302101)82.5/96:0.0320685754,(WP_125113420.1-Algoriella-xinjiangensis:0.0757582849,WP_092908829.1-Algoriella-xinjiangensis:0.0344688792)94.1/100:0.0432324235)98.2/96:0.1007590665)96.8/100:0.1143086182)97/99:0.1189215245,(WP_014783622.1-Aequorivita-sublithincola:0.2127070019,((((WP_230143398.1-unclassified-Pedobacter:0.0966922086,WP_215166022.1-Pedobacter-sp.-ISL-64:0.0739318200)97/100:0.0723014906,WP_133260761.1-Pedobacter-changchengzhani:0.0974378554)90.5/100:0.0526476632,((WP_056872627.1-Pedobacter-sp.-Leaf194:0.0842532345,(WP_123203876.1-Pedobacter-jejuensis:0.0656234075,(WP_188416210.1-Pedobacter-mendelii:0.0255388836,WP_056087060.1-Pedobacter-sp.-Leaf176:0.0454920374)71.1/95:0.0158464725)7/86:0.0147047615)97.4/95:0.0543113714,WP_109929147.1-Pedobacter-paludis:0.0312620324)83.3/84:0.0318415341)80.1/83:0.0468484464,WP_199156598.1-Pedobacter-sp.-ASV2:0.0654239464)99.3/97:0.1625891387)98.7/100:0.1547138555)45.9/45:0.0673087528,WP_111126400.1-Mesonia-sp.-K7:0.4764050016)78.5/98:0.0352380592,(((WP_107809536.1-Elizabethkingia-sp.-YR214:0.1234858446,WP_219971314.1-Chryseobacterium-sp.-LJ756:0.1196601893)91.9/100:0.0496757933,WP_076395614.1-Chryseobacterium:0.0486698563)33.8/48:0.0496968210,WP_088467981.1-Chryseobacterium-sp.-VAUSW3:0.0981216307)99.9/100:0.1711108299)96/97:0.0931237285,(((((WP_233378072.1-Maribellus-sp.-CM-23:0.0466686251,WP_247978154.1-Maribellus-sp.-YY47:0.0741846080)96.9/100:0.1016957512,(WP_167609329.1-Maribellus-sediminis:0.1419075292,(WP_045033348.1-Draconibacterium-sediminis:0.0254746240,WP_163345719.1-Draconibacterium-halophilum:0.0696280325)99.4/100:0.1245195613)93.9/100:0.0728795653)100/100:0.2136712474,((WP_106543043.1-Prolixibacter-denitrificans:0.0785163071,(WP_025865917.1-Prolixibacter-bellariivorans:0.0188100723,WP_153632016.1-Prolixibacter-sp.-SD074:0.0527332725)71.5/76:0.0264745147)100/100:0.2003019855,((WP_159517929.1-Sunxiuqinia-indica:0.0954848212,WP_053182762.1-Sunxiuqinia-dokdonensis:0.0846529825)84.5/100:0.0489415012,WP_133466511.1-Sunxiuqinia-elliptica:0.1244116091)98.4/100:0.1297254471)41.1/93:0.0512240857)87.9/94:0.0631704384,(((WP_101263396.1-Labilibaculum-filiforme:0.0390424451,WP_245870435.1-Labilibaculum-manganireducens:0.0408842232)98.9/100:0.0827057183,(WP_152734748.1-Marinifilum-sp.-N1E240:0.0700893588,(WP_171595603.1-Marinifilum-caeruleilacunae:0.0589700532,(WP_268189532.1-Marinifilum-sp.-D737:0.0474662029,WP_054720444.1-Marinifilum-fragile:0.0299984556)78.5/100:0.0184010845)87.8/100:0.0256320408)89.7/100:0.0436433961)45.2/92:0.0356972125,WP_127342456.1-Ancylomarina-longa:0.1060455291)99.7/100:0.1480260707)74.1/71:0.0484933848,(WP_068701711.1-Paludibacter-jiangxiensis:0.2980536691,((WP_011500232.1-Methanococcoides-burtonii:0.1687819759,WP_135611779.1-Methanococcoides-sp.-AM1:0.1486694591)82.9/100:0.0590969800,((((WP_128503182.1-Methanosarcina-sp.-MSH10X1:0.1390391035,(WP_011033115.1-Methanosarcina-mazei:0.0768885335,WP_048134921.1-Methanosarcina-sp.-1.H.T.1A.1:0.0354059036)34.4/100:0.0355887690)34.7/96:0.0440442237,WP_048167026.1-Methanosarcina:0.0730156685)72.3/69:0.0383759095,WP_048177163.1-Methanosarcina-sp.-MTP4:0.0743448182)18.6/68:0.0452313329,(WP_048137589.1-Methanosarcina-horonobensis:0.1267719378,((WP_048184631.1-Methanosarcina-siciliae:0.0526021031,WP_048173663.1-Methanosarcina-siciliae:0.0524381261)66.1/100:0.0107612437,WP_011023680.1-Methanosarcina-acetivorans:0.0323918674)96/100:0.0625862564)89.2/100:0.0642639458)85.6/100:0.0798010409)97.6/100:0.1244710691)66.3/95:0.0614268312)96.2/96:0.0878808307)10.2/82:0.0425952044,(((WP_205722928.1-Fulvivirga-lutea:0.3285305071,(((((((((((WP_053470876.1-unclassified-Flavobacterium:0.1401842495,WP_095952323.1-Flavobacterium-sp.-ACN6:0.0357558607)58.2/99:0.0072352238,WP_056200692.1-Flavobacterium-sp.-Root186:0.0312489287)85.2/100:0.0116957605,WP_115889823.1-Flavobacterium-cutihirudinis:0.0479421692)29/100:0.0054956190,WP_198857739.1-Flavobacterium-sp.-IB48:0.0363378469)77.5/100:0.0085378691,(WP_269237486.1-Flavobacterium-sp.-IMCC34775:0.0744910178,WP_163393939.1-Flavobacterium-limi:0.0337147443)0/99:0.0000021437)53.4/99:0.0109496962,WP_264205577.1-Flavobacterium-sp.-LS1R49:0.0227330818)91.1/100:0.0181967755,((WP_173968237.1-Flavobacterium-collinsii:0.0673882306,WP_264525312.1-Flavobacterium-sp.-N502536:0.0424630374)92.3/100:0.0279623341,WP_070907856.1-Flavobacterium-tructae:0.0742670789)72.7/86:0.0067397814)76.4/86:0.0082164664,WP_091091738.1-Flavobacterium-gillisiae:0.0893474404)97.9/100:0.0545134886,WP_248907802.1-Flavobacterium-sp.-K5-23:0.1065965954)88.3/98:0.0325617659,(((WP_116761981.1-Flavobacterium-laiguense:0.0373768265,WP_143377960.1-Flavobacterium-sp.-ZT3R18:0.0452216792)97/100:0.0626848635,((WP_144071503.1-Flavobacterium-franklandianum:0.0605607247,WP_133277350.1-Flavobacterium-nackdongense:0.0806162608)49.5/100:0.0235468986,WP_194311471.1-Flavobacterium-soyangense:0.1070422073)83.5/100:0.0256179839)82.1/100:0.0273057887,WP_103806521.1-Flavobacterium-alvei:0.0812102882)88.5/100:0.0391528900)100/98:0.1408413714,(WP_140000461.1-Flavobacterium-microcysteis:0.1748595513,(((WP_194640005.1-Flavobacterium-sp.-HJJ:0.0765434952,WP_073246022.1-Flavobacterium-flevense:0.0617947877)82.7/100:0.0210504174,WP_100843428.1-Flavobacterium-sp.-5:0.0825599247)81.3/99:0.0432944760,WP_140487657.1-Flavobacterium-sp.-GSA192:0.0466098554)99.3/100:0.1325405653)91.4/100:0.0791603716)95.1/99:0.0896512773)87.8/85:0.0832320660,(WP_116872776.1-Terrimonas-sp.:0.3116400632,(WP_004155558.1-Microscilla-marina:0.2415088325,WP_101358295.1-Raineya-orbicola:0.1767770470)99.6/100:0.2048313354)77.3/91:0.0720777390)85.5/88:0.0456225633,WP_057952567.1-Salinivirga-cyanobacteriivorans:0.4216108522)52.2/85:0.0550905675)94.9/100:0.1234815502,((WP_085659180.1-Prosthecochloris-sp.-HL-130-GSB:0.3164515795,WP_038563105.1-Draconibacterium-orientale:0.2148461142)97.3/100:0.1200813505,((WP_040596643.1-Prevotella-stercorea:0.0737287098,WP_207153370.1-Prevotella-herbatica:0.0818019419)99.1/100:0.1805493522,(WP_223927321.1-Prevotella-lacticifex:0.1755414811,WP_009010825.1-Prevotella-sp.-C561:0.1349743635)99.8/100:0.2812723819)82.4/99:0.0581835509)97.6/100:0.1511815581)23.7/91:0.1003606875,(((((WP_013633636.1-Pseudopedobacter-saltans:0.1119797140,WP_154287284.1-Pedobacter-puniceum:0.1389290113)76.4/99:0.0533346409,WP_160842504.1-Pedobacter-sp.-HMF7647:0.0635202534)97.7/99:0.1223792634,(WP_074492796.1-Mucilaginibacter-polytrichastri:0.1356388882,((WP_258137696.1-Mucilaginibacter-sp.-BJC16-A38:0.1893813762,WP_259068220.1-Mucilaginibacter-sp.-X4EP1:0.0374479423)29/39:0.0407903318,WP_144912900.1-Mucilaginibacter-frigoritolerans:0.0555883670)98.7/100:0.1011287767)90.9/100:0.0880052755)97.2/100:0.1423908080,WP_163286637.1-Cryomorpha-ignava:0.3792748057)2.3/94:0.0758841363,WP_130092062.1-Brumimicrobium-glaciale:0.3255242488)66.4/95:0.0681525551)92.5/99:0.0987071722)62.5/34:0.0477255954,(WP_138788932.1-Paenibacillus-thermoaerophilus:0.2677823305,WP_078711972.1-Anaerorhabdus-furcosa:0.3765529364)89.3/34:0.0923993683)97.8/100:0.1190499804)45.5/89:0.0229725493)78.9/5:0.0265560025,(((WP_072827923.1-Fibrobacter-sp.-UWB12:0.0257782608,WP_073304542.1-Fibrobacter:0.1676357426)14/8:0.0148800599,WP_088630837.1-unclassified-Fibrobacter:0.0225230131)95.1/97:0.0791645623,((WP_097034884.1-Fibrobacter-sp.-UWT3:0.0267086210,(WP_073056819.1-unclassified-Fibrobacter:0.0173123105,WP_073155408.1-Fibrobacter:0.0528497971)23.8/43:0.0233210393)72.8/46:0.0180996320,WP_073188830.1-Fibrobacter-sp.-UWP2:0.0250817649)96.4/99:0.0612345461)100/100:0.2481045625)82.9/58:0.0507087570)86/98:0.0373866437,(((((WP_018154793.1-Methanothermococcus-thermolithotrophicus:0.1022204291,WP_048201153.1-Methanocaldococcus-bathoardescens:0.0702704185)90.9/100:0.0437392850,((WP_052562942.1-Candidatus-Brocadia-sinica:0.2738489053,WP_173298646.1-Thermanaeromonas-sp.-C210:0.2211038350)88.1/100:0.0718356661,(((WP_011869065.1-Methanococcus-maripaludis:0.0796976085,WP_013998786.1-Methanococcus-maripaludis:0.0478616243)85.4/100:0.0290332287,WP_181493664.1-Methanococcus-maripaludis:0.0622655546)88.6/100:0.0319072056,WP_181504225.1-Methanococcus-maripaludis:0.0313632825)100/100:0.1694766317)78.4/100:0.0571278658)97.3/100:0.0847142640,(((WP_245533116.1-Coprothermobacter-platensis:0.3817485437,WP_022856039.1-Thermodesulfobacterium:0.1731863160)30.7/35:0.0383335430,((WP_119360783.1-Meiothermus-luteus:0.0780520839,(WP_208013758.1-Meiothermus-sp.-CFH-77666:0.1035651142,WP_027877894.1-Meiothermus-cerbereus:0.0515119338)90.3/100:0.0761891787)100/100:0.4405238292,WP_150071182.1-Thermodesulfobacterium-sp.-TA1:0.1490536875)83.5/29:0.0607581166)94.7/95:0.1035536538,(((((WP_011996389.1-Coxiella-burnetii:0.4982589884,WP_013638952.1-Desulfurobacterium-thermolithotrophum:0.0311358079)24.6/99:0.0117814916,WP_132526207.1-Phorcysia-thermohydrogeniphila:0.0715248503)89.1/100:0.0264841976,WP_089322203.1-Desulfurobacterium-atlanticum:0.1245182497)93.3/100:0.0574506306,(WP_029522925.1-Persephonella-sp.-KM09-Lau-8:0.0962557258,(WP_200674641.1-Persephonella-atlantica:0.0658182245,WP_096999769.1-Persephonella-hydrogeniphila:0.0921855701)94.3/100:0.0551019506)85.1/99:0.0564253625)6.8/86:0.0298911762,WP_028950938.1-Sulfurihydrogenibium-subterraneum:0.1009932586)97.1/100:0.0898213444)86.9/95:0.0680841407)76.5/98:0.0730972379,((WP_010868297.1-Pyrococcus-abyssi:0.0103074428,((((WP_013798345.1-Methanotorris-igneus:0.0305182745,WP_013866394.1-Methanothermococcus-okinawensis:0.1518078706)81.3/100:0.0236645975,WP_007044577.1-Methanotorris-formicicus:0.0591662602)98.8/100:0.0639856027,((((((WP_237702072.1-Thermococcus-sp.-AM4:0.0327113404,WP_223211916.1-Thermococcus-henrietii:0.0542374192)97.6/100:0.0531318312,WP_014733609.1-Pyrococcus-sp.-ST04:0.1087272338)72.9/98:0.0125059658,((((WP_088865724.1-Thermococcus-barossii:0.0569029784,((WP_167711671.1-unclassified-Thermococcus:0.0188078047,WP_088181368.1-Thermococcus-sp.-5-4:0.0628064373)77.5/25:0.0057911179,WP_232460978.1-Thermococcus-radiotolerans:0.0431340037)0/25:0.0000027116)76.3/25:0.0113124040,WP_258085050.1-Thermococcus-sp.-813A4:0.0307835913)81.3/24:0.0133027329,(WP_139679991.1-Thermococcus-indicus:0.0288269712,WP_148882943.1-Thermococcus-aciditolerans:0.0342021129)90.7/99:0.0197296677)76.1/22:0.0055891960,WP_058938040.1-Thermococcus-celericrescens:0.0283083296)80/24:0.0087207089)92.3/99:0.0299750897,(WP_152879125.1-Thermococcus-sp.-101-C5:0.0408377171,WP_015850310.1-Thermococcus-sibiricus:0.0527343720)84.8/99:0.0129335294)86.2/96:0.0122925076,(WP_068664737.1-Thermococcus-piezophilus:0.0533748176,WP_253304977.1-Thermococcus-aggregans:0.0345659436)75.7/22:0.0059195554)83.6/86:0.0134893482,WP_088862340.1-Thermococcus-celer:0.0808869163)79.9/34:0.0199037732)90/57:0.0253888600,(((WP_148120035.1-Methanofervidicoccus-sp.-A16:0.1532634539,(WP_013468354.1-Thermococcus-barophilus:0.0415313487,WP_013906489.1-Pyrococcus-yayanosii:0.0231218112)12.6/36:0.0164667733)76.2/34:0.0068475785,WP_048053121.1-Pyrococcus-horikoshii:0.0299451223)0/9:0.0000027799,(((WP_167913028.1-Thermococcus-sp.-21S7:0.0647925923,WP_048150691.1-Palaeococcus-ferrophilus:0.0246574297)91.9/100:0.0251284237,(WP_167888701.1-Thermococcus-sp.-MV5:0.0447386884,WP_209475361.1-Thermococcus-stetteri:0.0604887994)92.1/48:0.0314867630)40.4/42:0.0122609758,(WP_088855609.1-Thermococcus-siculi:0.0448418567,WP_087036111.1-Thermococcus-litoralis:0.0328904902)93.7/100:0.0335413305)71.9/55:0.0125209463)75.6/12:0.0118944729)97.8/99:0.0695624494)97.5/100:0.0576633590,WP_048103006.1-Aciduliprofundum-boonei:0.1646543252)96.6/100:0.1219263607)99/100:0.1342121888,WP_236098952.1-Dethiosulfovibrio:0.3747630030)91.7/98:0.0749644586)100/100:0.2250983454,((((((((WP_094934028.1-Enterobacter-cloacae-complex:0.3355588583,WP_004741399.1-Vibrio:0.1920873221)71/7:0.0658820645,((WP_010368546.1-Pseudoalteromonas-piscicida:0.2811385301,(WP_039473409.1-Vibrio-sinaloensis:0.2432800766,WP_016795256.1-Vibrio-cyclitrophicus:0.1535138753)89.5/76:0.0658802853)50.2/71:0.0184283408,WP_086590801.1-Vibrio-parahaemolyticus:0.2631176622)82.2/27:0.0686444366)52.3/4:0.0353372263,WP_016537220.1-Cedecea-davisae:0.1065657855)66.7/6:0.0404822648,((WP_269408820.1-Pseudoalteromonas-shioyasakiensis:0.1489846838,WP_207379479.1-Shewanella-sedimentimangrovi:0.1464571484)68.8/99:0.0454549840,((WP_061013169.1-Vibrionaceae:0.1838136249,WP_155662549.1-Aliivibrio-fischeri:0.1080815978)73.9/100:0.0620323531,WP_171328835.1-Vibrio-splendidus:0.1882899775)78.5/5:0.0291955096)95.6/78:0.0697094666)41.6/18:0.0309525790,(WP_195504469.1-Citrobacter-farmeri:0.2171944568,WP_064554809.1-Buttiauxella-noackiae:0.1881232399)99.2/98:0.1646829317)91.7/18:0.0603436359,(WP_001100815.1-Escherichia-coli:0.2211262174,WP_240304790.1-Vibrio-campbellii:0.3309298544)47.6/3:0.0819494013)79.3/3:0.0251870220,WP_057065012.1-Citrobacter:0.4181633420)92.6/98:0.0650296436,WP_216735790.1-Hahella-sp.-HN01:0.1957346517)83.3/80:0.0288636368)99.1/52:0.1167692812,((((WP_127021653.1-Rheinheimera-mangrovi:0.2395219815,WP_101108574.1-Psychromonas-sp.-psych-6C06:0.1522258506)5.1/3:0.0273857360,WP_114327629.1-Candidatus-Colwellia-aromaticivorans:0.1208586131)78.8/3:0.0178878739,((WP_138320019.1-Thalassotalea-litorea:0.1421271296,WP_136736185.1-Thalassotalea-mangrovi:0.0845473457)99.8/100:0.1755098827,WP_148513713.1-Vibrio:0.1616463775)67.2/75:0.0526408436)66/75:0.0465527418,((WP_016794198.1-Vibrio-cyclitrophicus:0.0332622791,WP_102424249.1-Vibrio-cyclitrophicus:0.0396439357)90.7/100:0.0381290475,(WP_261903470.1-Vibrio-fortis:0.0691080299,WP_240207223.1-Vibrio-sp.-CyArs1:0.0249013193)90.4/100:0.0277083231)98.7/100:0.0886450261)88/89:0.0341618425)55.8/5:0.0268886987,(((WP_260491935.1-Pseudoalteromonas-flavipulchra:0.1547538880,WP_039494249.1-Pseudoalteromonas-flavipulchra:0.0607784042)43.5/98:0.0283509548,WP_055022295.1-Pseudoalteromonas-sp.-P1-26:0.0474633364)83.4/98:0.0307902160,(WP_149602089.1-Pseudoalteromonas-distincta:0.0327748329,WP_248518672.1-Pseudoalteromonas-sp.-2CM32C:0.0218818252)99.6/100:0.0932155580)99.2/100:0.1281966992)17.1/7:0.0208920612,WP_150896168.1-Vibrio-fortis:0.1572113917)67.4/16:0.0626086018,((((((((((((((((((WP_171588903.1-Pelistega-europaea:0.2344751472,((((WP_267333169.1-unclassified-Gilliamella:0.1185758519,WP_267361838.1-Gilliamella-sp.-B2772:0.1189218122)75.7/99:0.0088958486,WP_086361229.1-Gilliamella:0.1102467629)74.7/92:0.0226456587,((((((WP_086319877.1-Gilliamella-apis:0.0167852882,WP_198214961.1-Gilliamella-sp.-M0320:0.0491131717)92.1/100:0.0268804418,WP_039129858.1-Gilliamella:0.0219215763)74.3/100:0.0086793700,(WP_065737496.1-Gilliamella-apicola:0.0337903152,WP_267331332.1-unclassified-Gilliamella:0.0493203116)97.1/100:0.0454589948)94.5/100:0.0372762338,(WP_065579324.1-Gilliamella-apis:0.0747852260,(WP_065587409.1-Gilliamella-apicola:0.0508308926,WP_065580031.1-Gilliamella-apicola:0.0546710124)80.3/100:0.0236084510)9.2/8:0.0163536469)60.3/13:0.0153746194,WP_160432071.1-Gilliamella-sp.-Pas-s25:0.0948724579)83.7/98:0.0170095186,((WP_065574597.1-Gilliamella-apicola:0.0475343979,WP_065569248.1-Gilliamella-apicola:0.0342173371)74/100:0.0092929169,WP_065610260.1-Gilliamella-apicola:0.0289075118)98.9/100:0.0543577324)38/44:0.0149939115)88/92:0.0455588115,(WP_247177500.1-Escherichia-coli:0.2977805501,YP_009985672.1-Escherichia-phage-muut:0.1866995115)99.8/100:0.2093587908)94.1/99:0.0732284010)65.9/19:0.0058044676,(WP_024553491.1-Franconibacter-helveticus:0.2298362779,(WP_040666911.1-Neisseria-wadsworthii:0.0667410598,WP_135034416.1-Neisseria:0.1401064938)87.6/99:0.0453990579)70.4/11:0.0136119382)95.5/87:0.0347819059,(WP_154751639.1-unclassified-Erwinia:0.1790195536,WP_140470594.1-Ewingella-americana:0.1792818390)89.2/78:0.0379890429)50.3/49:0.0142730700,(((WP_214236207.1-Rosenbergiella-sp.-S61:0.0367823685,(WP_241647140.1-Rosenbergiella-sp.-JB07:0.0935875246,WP_214195533.1-Rosenbergiella-collisarenosi:0.0791455148)67.1/86:0.0362998770)99.6/99:0.0973980376,(WP_029991521.1-Tatumella-ptyseos:0.0650700859,(WP_212707699.1-Tatumella-sp.-JGM130:0.0363425917,WP_087487466.1-Tatumella-citrea:0.0460910929)26.8/68:0.0077596041)72.6/85:0.0245269180)9.6/47:0.0132299531,(WP_032441370.1-Klebsiella-pneumoniae:0.1056062744,(WP_015583911.1-Raoultella-ornithinolytica:0.0514459251,WP_265776798.1-Klebsiella-oxytoca:0.1307331595)63.7/100:0.0314218996)92.1/99:0.0464940548)96.7/87:0.0671378262)59.3/46:0.0161414569,((WP_087653467.1-Klebsiella/Raoultella-group:0.1289553619,WP_028357569.1-Brackiella-oedipodis:0.1732603599)91.5/93:0.0494962737,WP_101506252.1-Erwinia-sp.-B116:0.0930436675)90.3/80:0.0357584970)94.8/87:0.0319130290,((((WP_131063651.1-Raoultella-planticola:0.0251677066,WP_163980548.1-Raoultella-ornithinolytica:0.0281166814)99/100:0.0704166512,WP_142430937.1-Klebsiella-oxytoca:0.0955629833)73.3/29:0.0069232236,((WP_239614184.1-Raoultella-ornithinolytica:0.0715613155,WP_186921353.1-Klebsiella-quasipneumoniae:0.1217392583)82/86:0.0148707546,((WP_032699180.1-Raoultella:0.0459521516,WP_228399925.1-Raoultella:0.0263225728)82.1/100:0.0138939902,WP_004138711.1-Klebsiella:0.0148463023)95.5/99:0.0345259184)91.7/86:0.0273254047)14.3/56:0.0151589504,(WP_148250181.1-Raoultella-terrigena:0.0782163499,WP_032686784.1-Raoultella-planticola:0.0257736641)95.5/99:0.0527383054)74.7/50:0.0067297649)55.3/22:0.0062097296,((((WP_193937755.1-Klebsiella-grimontii:0.1402601485,(WP_024360036.1-Klebsiella:0.0494218964,WP_234284225.1-Klebsiella-oxytoca:0.0627241511)15.9/92:0.0280520591)12.7/17:0.0242935376,WP_142514871.1-Klebsiella-huaxiensis:0.1254032069)89.2/99:0.0249665670,((((((((WP_136167279.1-Brenneria-sp.-CFCC-11842:0.0361236052,WP_264089117.1-unclassified-Brenneria:0.0791945197)36.8/99:0.0293506113,WP_109053612.1-Brenneria-roseae:0.0745664319)88.1/100:0.0284961910,((((((((WP_180742451.1-Pectobacterium-sp.-CFBP8739:0.1473384995,WP_180777316.1-Pectobacterium-aroidearum:0.0075718877)97.4/99:0.0529327225,WP_039504461.1-Pectobacterium:0.0207882549)41.3/64:0.0124922118,(WP_109411993.1-Pectobacterium-polaris:0.0465950022,(WP_233968669.1-Pectobacterium-polaris:0.0480330517,WP_048259384.1-Pectobacterium-peruviense:0.0120627117)78.5/13:0.0132332855)71.3/8:0.0102023007)77/4:0.0114955521,WP_138255615.1-Pectobacterium:0.0133335535)87.6/9:0.0176439975,((WP_249680869.1-Pectobacterium-polaris:0.0311081041,WP_125232508.1-Pectobacterium-versatile:0.0421693037)75.8/97:0.0072429492,WP_005974084.1-Pectobacterium-wasabiae:0.0288774055)87.1/41:0.0182971920)73.5/23:0.0061832306,WP_233976643.1-Pectobacterium-versatile:0.0236968233)0/2:0.0000024119,(((((((WP_047710158.1-Pectobacterium-atrosepticum:0.0490913150,WP_033071760.1-Pectobacterium-parmentieri:0.0251815828)0/92:0.0124313077,(WP_198482318.1-Pectobacterium-parmentieri:0.0107158350,WP_107169883.1-Pectobacterium:0.0483035714)26.6/7:0.0160696379)80.1/94:0.0366315837,WP_226305805.1-Pectobacterium-versatile:0.0121680414)36.4/5:0.0228272600,((WP_039280054.1-Pectobacterium-brasiliense:0.0283155141,WP_039517132.1-Pectobacterium-brasiliense:0.0273576796)88.4/98:0.0191383776,WP_039346077.1-Pectobacterium-fontis:0.0240683002)75.5/92:0.0072759516)85.5/6:0.0162762754,((WP_205558955.1-Pectobacterium-brasiliense:0.0445991006,(WP_151242707.1-Pectobacterium-carotovorum:0.0324949548,WP_116186106.1-Pectobacterium-aquaticum:0.0389967811)65.2/95:0.0059353616)74.5/91:0.0065972967,WP_103972204.1-Pectobacterium:0.0147885906)77.4/86:0.0059300775)89.6/9:0.0174301224,WP_119888241.1-Pectobacterium:0.0340144765)86.6/56:0.0143802227,WP_205946867.1-Pectobacterium:0.0540793671)76.8/6:0.0059040645)96.1/98:0.0359700921,(WP_132452860.1-Samsonia-erythrinae:0.0967213427,WP_130637964.1-Pectobacterium-versatile:0.0470402979)21.1/29:0.0179842478)84.5/99:0.0182781997)97.3/100:0.0539046279,((WP_121576612.1-Brenneria-alni:0.0773544842,((WP_214648782.1-Pectobacterium-carotovorum:0.0298064491,((WP_261846379.1-Pectobacterium-carotovorum:0.0477477621,WP_121590317.1-Brenneria-goodwinii:0.0060572110)93.8/96:0.0238624834,(WP_182099415.1-Pectobacterium-aroidearum:0.0295952081,WP_039482612.1-Pectobacterium:0.0238773675)74/98:0.0058616205)38.8/65:0.0058734339)0/65:0.0000028940,WP_249228983.1-Brenneria-tiliae:0.0427424362)93.1/89:0.0306702620)75.7/92:0.0100329767,WP_062791489.1-Serratia-plymuthica:0.0716875643)77.7/26:0.0226750938)78.8/32:0.0175538516,WP_095845203.1-Gibbsiella-quercinecans:0.0399536853)36.1/28:0.0093075283,((WP_096000273.1-Klebsiella-pneumoniae:0.1338519262,WP_248464278.1-Pectobacterium-versatile:0.1097555233)73.4/86:0.0104099455,((WP_168640580.1-Dickeya-sp.-CFBP-2040:0.0803548422,((((WP_038918362.1-Dickeya:0.0257041350,WP_123251184.1-Dickeya-undicola:0.0406749893)68.5/100:0.0061939630,WP_022632763.1-Dickeya-solani:0.0229594989)93.7/100:0.0237304793,(WP_012883965.1-Dickeya-parazeae:0.0545090770,WP_236306746.1-Dickeya-zeae:0.0292317277)82.6/80:0.0127592878)74/49:0.0058425859,(WP_027712673.1-Dickeya-chrysanthemi:0.0359313874,WP_012770533.1-Dickeya:0.0295680095)75/65:0.0058852743)76.7/68:0.0134125212)99/100:0.0736535585,(WP_148877341.1-Serratia-marcescens:0.0470963001,WP_193158894.1-Serratia-marcescens:0.0428885690)93/100:0.0448148993)88.2/87:0.0283024877)93.2/67:0.0365946274)91.7/99:0.0354767485,WP_137713144.1-Brenneria-rubrifaciens:0.0838458028)82/42:0.0178786354,(WP_112167140.1-Klebsiella:0.1010386627,WP_200519240.1-Klebsiella-michiganensis:0.0834667681)95.4/100:0.0498523193)70.8/25:0.0061338166)42.2/37:0.0105777877,WP_165428828.1-Rouxiella-badensis:0.0928588290)72.4/23:0.0070796658)81.4/47:0.0191024934,((WP_228398973.1-Insectihabitans-xujianqingii:0.0752918533,WP_074823218.1-Pragia-fontium:0.0761268235)96.5/100:0.0616597626,WP_130971788.1-Hafnia-alvei:0.0943705621)85.5/94:0.0252666058)91.9/98:0.0403667811,((WP_126816534.1-Enterobacter-mori:0.0919832423,WP_256879438.1-Escherichia-albertii:0.0696194461)70.4/99:0.0473309109,WP_049841125.1-Klebsiella-sp.-RIT-PI-d:0.2515628593)84/28:0.0389852711)86.4/24:0.0493870217,(WP_246757812.1-Erwinia-phyllosphaerae:0.1341115155,WP_232759612.1-unclassified-Pantoea:0.1035095018)92.7/100:0.0529126952)50.3/22:0.0264650539,(WP_146378871.1-Cronobacter-turicensis:0.0710261342,WP_032180402.1-Escherichia-coli:0.0543881770)99.7/100:0.1246350560)21.8/92:0.0392142331,WP_253460089.1-Erwinia-persicina:0.1809821466)89.4/98:0.0368666647,((WP_168836617.1-Vibrio-agarilyticus:0.0925301071,(WP_076537303.1-Shewanella-sp.-UCD-KL21:0.0183626560,WP_102531614.1-Shewanella-sp.-10N.286.52.B9:0.0547925230)100/100:0.3652453540)94/100:0.0734821783,(WP_137671336.1-Agarivorans-sp.-Toyoura001:0.0511235346,WP_221077378.1-Agarivorans-albus:0.0317438392)95.3/100:0.0773141960)96.6/100:0.0875265939)80.3/74:0.0134533515,WP_100256321.1-Reinekea-forsetii:0.3804565824)84/85:0.0294320839,(WP_261844981.1-Amphritea-ceti:0.2023591355,(((((WP_269124033.1-Marinomonas-sp.-15G1-11:0.0275087635,WP_191602238.1-Marinomonas-algicola:0.0656563425)98.5/100:0.0867902391,WP_048693477.1-Catenovulum-maritimum:0.1559989306)93.9/100:0.0700077742,WP_067204681.1-Marinomonas-aquimarina:0.0151895195)6.7/21:0.0235653495,(WP_067216963.1-Marinomonas-gallaica:0.0375654122,WP_228191388.1-Marinomonas-communis:0.1285904908)9.8/10:0.0292830412)88.9/93:0.0232476934,WP_055461762.1-Marinomonas-fungiae:0.0438724655)89.8/95:0.0409127164)91.6/98:0.0454700084)71.2/89:0.0413076032,((WP_145010177.1-Pseudomonas-psychrotolerans:0.0171968807,WP_077172422.1-Pseudomonas-psychrotolerans:0.0351442432)100/100:0.1544138364,WP_084334293.1-Pseudomonas-indica:0.0895094640)34.2/94:0.0365482084)93.1/93:0.0513761856,(WP_132584255.1-Rheinheimera-sp.-D18:0.2225278536,((((((WP_131912015.1-Celerinatantimonas-diazotrophica:0.1661447733,((WP_092385961.1-Halopseudomonas-salegens:0.1434421683,WP_036859756.1-Porticoccus-hydrocarbonoclasticus:0.1821173032)93.7/99:0.0830061348,((WP_076722556.1-Marinobacter-lutaoensis:0.1722707066,WP_036130839.1-Marinobacter-nitratireducens:0.0906608233)36/89:0.0453709362,((WP_106695101.1-Marinobacter-shengliensis:0.0507761769,(WP_092030923.1-Marinobacter-sp.-DSM-26671:0.0910122576,WP_148609564.1-Marinobacter-sp.-BW6:0.0721916750)86/84:0.0283665474)75.6/84:0.0126401014,WP_222473125.1-Marinobacter-daepoensis:0.0487957912)95.6/99:0.0827092088)14.5/89:0.0344307299)89/100:0.0435989221)87.3/98:0.0360674628,(WP_027875186.1-Spongiibacter-marinus:0.1237660534,WP_204995064.1-Spongiibacter-marinus:0.0815406521)89.7/100:0.0435829360)91.4/98:0.0329262059,WP_168451636.1-Spongiibacter-thalassae:0.0402902307)85.9/97:0.0448294601,WP_062382560.1-Zhongshania-aliphaticivorans:0.0448536594)98.2/100:0.0811712602,(WP_184464382.1-Zhongshania-antarctica:0.0438788545,WP_103683243.1-Marortus-luteolus:0.0285823709)82.6/99:0.0138815873)73.1/94:0.0243586792,WP_198570926.1-Spongiibacter-nanhainus:0.0486691121)93.4/97:0.0720072087)83.8/100:0.0558237642)94.2/78:0.0678916846,WP_067584343.1-Endozoicomonas-ascidiicola:0.1636492063)77.2/37:0.0450083483)87.8/47:0.0522866370,WP_199145406.1-Citrobacter-sp.-ASV29:0.1838105213)91.4/91:0.0820244937,((((GJNO01008138.1-Laxus-oneistus-TSA:0.4597648951,((((((VDM95900.1-Thelazia-callipaeda:0.1766166755,(((((VDO27950.1-Onchocerca-flexuosa:0.1308379764,VDO31788.1-Brugia-timori:0.0994736060)75.1/83:0.0229806264,XP_003135693.1-Loa-loa:0.0654854242)90.7/81:0.0256032410,CAG9536712.1-Cercopithifilaria-johnstoni:0.0275040254)82/81:0.0119898884,VBB33646.1-Acanthocheilonema-viteae:0.0840234095)79.8/81:0.0143044431,VDK73328.1-Litomosoides-sigmodontis:0.0315388321)99.7/98:0.1235949586)100/100:0.2392624007,(VDD97390.1-Enterobius-vermicularis:0.2846001936,KHN87800.1-Toxocara-canis:0.1510234128)73.3/95:0.0529192104)40.5/93:0.0495570919,VDN56932.1-Dracunculus-medinensis:0.2469715533)95.7/98:0.1111809514,(KAI1709308.1-Ditylenchus-destructor:0.1134384366,KAI1730870.1-Ditylenchus-destructor:0.0279831787)99.8/100:0.1584997908)34.5/56:0.0410921031,GGJS01006545.1-Plectus-murrayi-TSA:0.2858892963)16.5/28:0.0371599343,(((((CAI4229259.1-Auanema-sp.-JU1783:0.2894014401,((KHJ99538.1-Oesophagostomum-dentatum:0.0547892109,EPB80542.1-Ancylostoma-ceylanicum:0.0386429729)95.6/100:0.0660536080,((VDO65958.1-Haemonchus-placei:0.1189897080,VDL70484.1-Nippostrongylus-brasiliensis:0.0809049616)69.3/100:0.0235174739,VDO95824.1-Heligmosomoides-polygyrus:0.0677733630)99.5/100:0.1223140135)97/100:0.1138584323)27.9/90:0.0405405837,PAV65180.1-Diploscapter-pachys:0.1783112557)88.5/98:0.0604851881,(CAD6189320.1-Caenorhabditis-auriculariae:0.2953427724,(CAB3400093.1-Caenorhabditis-bovis:0.1685058560,((NP_509046.1-Caenorhabditis-elegans:0.0333396693,XP_003117635.1-Caenorhabditis-remanei:0.0215388598)61.6/100:0.0341270011,CAI5454992.1-Caenorhabditis-angaria:0.0363610385)91.2/100:0.0398160703)78.9/100:0.0661097909)95.8/99:0.0948291848)99.8/90:0.1362650830,((KAH7695359.1-Aphelenchus-avenae:0.2596147841,((CAD5228290.1-Bursaphelenchus-okinawaensis:0.0450736001,CAD5235627.1-Bursaphelenchus-xylophilus:0.0300450275)16.3/79:0.0466116034,((KAI6206096.1-Aphelenchoides-besseyi:0.1309958495,KAI6191010.1-Aphelenchoides-bicaudatus:0.0751079797)44.4/100:0.0503226240,KAI6234710.1-Aphelenchoides-fujianensis:0.0919604206)97.7/100:0.0966840222)91.7/100:0.0944451369)84.7/100:0.0784604341,KAE9552274.1-Halicephalobus-sp.-NKZ332:0.2494031235)13.1/31:0.0519806401)59.7/53:0.0446129014,TKR72632.1-Steinernema-carpocapsae:0.1622151315)78.9/62:0.0263507083)96.2/73:0.2086321533)99.9/100:0.4126457202,(WP_042281872.1-Candidatus-Protochlamydia-sp.-R18:0.1284870845,WP_068471358.1-Parachlamydia-sp.-C2:0.1058556046)100/100:0.2918667031)91.1/83:0.1365784315,(WP_008983868.1-Alishewanella-agri:0.5213980053,WP_265845290.1-Plesiomonas-shigelloides:0.4232308913)78.9/84:0.0903457975)72/98:0.0679765085,(WP_255135177.1-Gracilimonas-sp.-CAU-1638:0.1982833244,WP_020401568.1-Gracilimonas-tropica:0.1915384971)100/100:0.3195638925)95/85:0.1020032598)58.6/38:0.0849488551,(WP_133132051.1-Legionella-sp.-W10-070:0.2268312079,(((((((((WP_027223798.1-Legionella-pneumophila:0.0291374386,WP_265717942.1-Legionella-sp.-PATHC032:0.0344176054)94/92:0.0278247379,WP_106142999.1-Legionella-pneumophila:0.0340106612)65.2/80:0.0039004391,(WP_061513621.1-Legionella:0.0134951748,WP_027226296.1-Legionella-pneumophila:0.0497404504)99.4/89:0.0589463025)87.2/97:0.0124599742,WP_044499121.1-Legionella-pneumophila:0.0231837561)84.4/97:0.0146691928,WP_027220106.1-Legionella-pneumophila:0.0147719567)84.1/97:0.0171562827,WP_027266778.1-Legionella-pneumophila:0.0311162709)96.9/98:0.0645321449,(WP_031567420.1-Legionella-wadsworthii:0.1539106597,(((WP_232890831.1-unclassified-Legionella:0.0380988776,(WP_058459148.1-Legionella-bozemanae:0.0224244523,WP_058518697.1-Legionella-parisiensis:0.0439187081)89.8/100:0.0286458754)99.6/100:0.0814857007,(WP_058513550.1-Legionella-santicrucis:0.1016186997,(WP_058463758.1-Legionella-cincinnatiensis:0.0192151130,WP_128130294.1-Legionella-sainthelensi:0.0499447929)92/100:0.0317713331)63/99:0.0209952854)88.3/96:0.0250638621,WP_045097199.1-Legionella-fallonii:0.1031870327)65/97:0.0257673630)94.6/68:0.0540128043)93.2/65:0.0577438599,(WP_127106294.1-Legionella-sp.-km535:0.0356624118,WP_058474153.1-Legionella-quateirensis:0.0281963320)93.9/100:0.0414135227)1.6/44:0.0374033373,WP_058494182.1-Legionella-worsleiensis:0.0870148001)99.3/100:0.1812699418)99.8/100:0.1895071854)92.3/26:0.0855402481)75.8/19:0.0252626885,(((((((WP_176021856.1-Stenotrophomonas-sp.-NA06056:0.0437089561,WP_049440708.1-Stenotrophomonas-maltophilia:0.0423912325)98.5/100:0.1153668129,WP_132999175.1-Luteimonas-arsenica:0.1983928977)93.7/100:0.0940272305,WP_078758981.1-Lysobacter-spongiicola:0.1902671533)79.8/100:0.0635461134,((((WP_019467068.1-Dyella-japonica:0.1796124147,WP_204636097.1-Dyella-kyungheensis:0.1137795216)83.3/92:0.0411990711,((((((WP_063674027.1-Dyella-thiooxydans:0.1127482056,WP_253565187.1-Dyella-sp.-Sa:0.0646491105)71.7/100:0.0260523178,(WP_131151171.1-Dyella:0.1666834346,(WP_209622196.1-Frateuria-flava:0.0658187135,WP_091339503.1-Frateuria-terrea:0.0445426386)89.3/100:0.0392124600)86.3/96:0.0214174850)78.3/95:0.0273747900,WP_201315287.1-Dyella-sp.-EPa41:0.0975445081)24.2/36:0.0133914749,(WP_188800319.1-Dyella-caseinilytica:0.1288157049,WP_192554051.1-Dyella-acidiphila:0.1112102567)98.1/100:0.0827478617)52/60:0.0202326412,(WP_049621948.1-Frateuria-defendens:0.1041053576,WP_179481164.1-unclassified-Rhodanobacter:0.0847290893)82.8/81:0.0366627978)37.2/6:0.0232740556,(((((((WP_214557828.1-unclassified-Rhodanobacter:0.0296504077,WP_068095172.1-unclassified-Rhodanobacter:0.0278343875)0/34:0.0190585107,((WP_008437080.1-Rhodanobacter-thiooxydans:0.0370387952,WP_063107523.1-Rhodanobacter-thiooxydans:0.0257194568)33.5/100:0.0090098302,WP_211005527.1-Rhodanobacter-sp.-B2A1Ga4:0.0334959283)80.1/100:0.0166600346)93.6/100:0.0373769801,WP_077445946.1-Rhodanobacter-sp.-C05:0.0460278716)14.3/11:0.0100466564,WP_072762453.1-Rhodanobacter-sp.-OK091:0.0137680301)82.8/4:0.0126746904,(WP_077439480.1-Rhodanobacter-sp.-C01:0.0938793433,WP_158752649.1-Dyella-sp.-S184:0.0265567425)86.9/38:0.0165041808)83.5/8:0.0196515616,WP_184416890.1-Rhodanobacter-sp.-MP7CTX1:0.0383200581)93.7/45:0.0354208606,WP_015446789.1-Rhodanobacter-denitrificans:0.0460788055)91.3/100:0.0310821355)59/50:0.0253312140)0/51:0.0000029079,(((WP_168608661.1-Oleiagrimonas-citrea:0.0535067490,WP_113064358.1-Oleiagrimonas-sp.-MCCC-1A03011:0.0192479920)95.8/100:0.0634128102,WP_043099571.1-Oleiagrimonas-soli:0.0444609348)99.8/100:0.1283017944,((WP_233843762.1-Dyella-sp.-2HG41-7:0.0347316103,WP_188793403.1-Dyella-nitratireducens:0.0636512718)82.6/100:0.0284617953,WP_126673574.1-Dyella-dinghuensis:0.0570753274)90.3/100:0.0471551244)91.7/83:0.0481630073)78.7/95:0.0271337693,(WP_214570707.1-unclassified-Dyella:0.1210999997,((WP_111981026.1-Dyella-jiangningensis:0.1448884187,WP_199044928.1-Dyella-sp.-ASV24:0.0267974653)92.6/68:0.0376272165,(WP_130619635.1-Dyella-amyloliquefaciens:0.0770928384,(WP_266169930.1-Dyella-sp.-RXD159:0.0662629286,(WP_266183414.1-Dyella-sp.-RBB1W86:0.1000653821,(WP_114846719.1-Dyella-tabacisoli:0.0415432349,WP_090451016.1-Dyella-sp.-OK004:0.0646895260)0/92:0.0181707336)0/51:0.0099485657)74.6/54:0.0139902486)42.9/52:0.0062818855)77.3/53:0.0169749681)75.6/62:0.0317794246)97.6/100:0.0939906173)99.8/100:0.1628478009,(((WP_256181981.1-Cloacibacillus-evryensis:0.1804658380,WP_087368268.1-Cloacibacillus-sp.-An23:0.0809003352)100/100:0.3748912150,((((WP_246588317.1-Pelorhabdus-rhamnosifermentans:0.1516723295,(WP_092074275.1-Dendrosporobacter-quercicolus:0.2027108978,(WP_145090815.1-Sporomusa-sp.-KB1:0.0434930093,WP_093793074.1-Sporomusa-acidovorans:0.0641204430)95/100:0.0606933819)95.9/99:0.0664588724)87.5/96:0.0408132343,WP_110954047.1-Massilibacillus-massiliensis:0.2236730990)0/45:0.0039062901,((((WP_007952170.1-Pelosinus-fermentans:0.0920745576,WP_007939097.1-Pelosinus:0.0937414934)90.4/100:0.0494881395,(WP_132081660.1-Anaerospora-hongkongensis:0.0783294282,WP_132083235.1-Anaerospora-hongkongensis:0.1625323887)98.4/100:0.1073969410)71/97:0.0273028394,(WP_188399481.1-Sporomusa-sp.-GT1:0.2065928287,((WP_094603851.1-Sporomusa-silvacetica:0.1029251495,WP_144352952.1-Sporomusa-termitida:0.1175462835)89.9/100:0.0481337256,WP_054258662.1-Propionispora-sp.-2/2-37:0.1828512819)54.7/72:0.0336735025)78.6/84:0.0434133952)82/73:0.0251930720,(WP_054258683.1-Propionispora-sp.-2/2-37:0.1246384306,((WP_149735423.1-Propionispora-hippei:0.0670870712,WP_091744506.1-Propionispora-vibrioides:0.1214998613)62.9/96:0.0246150092,WP_188400350.1-Sporomusa-sp.-GT1:0.2177069951)58.5/64:0.0177203956)96.1/98:0.0713052522)89.6/97:0.0410337029)99.2/100:0.1125292153,((WP_090940141.1-Pelosinus-propionicus:0.1828489711,WP_240986382.1-Acididesulfobacillus-acetoxydans:0.1930342751)93.2/100:0.0694600273,WP_092486361.1-Desulfoscipio-geothermicus:0.1206010318)84.7/99:0.0436098542)86.6/85:0.0504043446)16.8/19:0.0308914213,(WP_084578244.1-Sporomusa-malonica:0.3069066860,WP_131919244.1-Heliophilum-fasciatum:0.1831419184)57/94:0.0547658994)87/27:0.0458015363)0/13:0.0467960457,(((((((WP_200343816.1-Halochromatium-glycolicum:0.1334389054,WP_201222000.1-Halochromatium-roseum:0.1085330609)97.9/100:0.1088118501,(WP_200237937.1-Thiohalocapsa-halophila:0.0503273788,WP_058558062.1-Thiohalocapsa-sp.-ML1:0.1990422097)94.1/100:0.0844623619)97.5/100:0.1149038701,(WP_182584206.1-Thiospirillum-jenense:0.2859198517,(WP_200286492.1-Rhabdochromatium-marinum:0.1152196176,WP_009150811.1-Thiorhodovibrio-sp.-970:0.0583932738)94.6/100:0.0963491924)86/100:0.0744726989)96.1/100:0.0815727667,(((((WP_200207675.1-Chromatium-weissei:0.1650437854,WP_200154756.1-Chromatium-okenii:0.0743140366)98.6/100:0.1008762002,((WP_200158060.1-Allochromatium-vinosum:0.0581078624,WP_200376120.1-Thiocystis-violacea:0.0877580878)91.3/100:0.0418458852,(((WP_043749566.1-Imhoffiella-purpurea:0.0612714680,WP_007039504.1-Thiorhodococcus-drewsii:0.0530021552)88.7/100:0.0285566132,((WP_164456073.1-Thiorhodococcus-minor:0.0830278010,WP_200330387.1-Thiocystis-violacea:0.0685212344)4.4/100:0.0183046689,WP_164652045.1-Thiorhodococcus-mannitoliphagus:0.0375652829)96.3/100:0.0539217600)88.6/100:0.0194187912,(WP_201092787.1-Thiocystis-minor:0.0652261391,WP_132978621.1-Thiobaca-trueperi:0.0856025584)84.3/99:0.0184304075)89.7/98:0.0298050754)55.2/82:0.0169682498)73.5/90:0.0263479211,(WP_200388819.1-Thiocapsa-imhoffii:0.1355857139,((WP_007191850.1-Thiocapsa-marina:0.0375071409,WP_232409912.1-Thiocapsa-bogorovii:0.0222985126)21.7/71:0.0124756522,WP_120796078.1-Thiocapsa-rosea:0.0196048992)98.4/100:0.0785832679)92.6/99:0.0555400026)95.2/99:0.0818264336,(WP_100918325.1-Candidatus-Thiodictyon-syntrophicum:0.0847034223,WP_020502667.1-Lamprocystis-purpurea:0.0908543246)96/100:0.0685755364)94.5/100:0.0679601596,(WP_141730858.1-Oligoflexus-tunisiensis:0.2912552894,WP_101758446.1-Oceanicoccus-sp.-KOV_DT_Chl:0.2972994693)96.3/98:0.1047279323)83.1/87:0.0320161296)50.5/87:0.0383287412,WP_018175268.1-unclassified-Thioalkalivibrio:0.3048346605)77/84:0.0228022073,WP_089729519.1-Candidatus-Thiosymbion-oneisti:0.2345593076)82.8/98:0.0511262087,WP_141198172.1-Persicimonas-caeni:0.3664025334)75.6/26:0.0761539335)94.6/99:0.0790771361,((WP_207681674.1-Desulfonema-magnum:0.3280422195,WP_008811140.1-Parasutterella-excrementihominis:0.3625818357)4.2/36:0.0947670898,WP_230840800.1-Gloeobacter-morelensis:0.2660042024)19.9/25:0.0827800344)92.3/39:0.0620260984)95.4/83:0.0851319160,((WP_110265962.1-Sinimarinibacterium-flocculans:0.2063043817,WP_077329105.1-Virgibacillus-siamensis:0.3554079858)66.9/72:0.0770966172,(WP_234070476.1-Myxococcus-stipitatus:0.1984522284,WP_120555541.1-Corallococcus-aberystwythensis:0.1827494316)99.9/100:0.2122228152)16.6/61:0.0379443220)99.4/100:0.1347970963)95.7/97:0.0824370783)81.8/70:0.0237060562,(((((((((((WP_065305794.1-Janthinobacterium-psychrotolerans:0.1065039044,WP_034778721.1-Janthinobacterium-lividum:0.0294099226)68.3/99:0.0043422436,WP_035822645.1-Janthinobacterium-sp.-RA13:0.0526395234)76.5/99:0.0067720152,WP_152282105.1-Janthinobacterium-violaceinigrum:0.0172396052)76.1/99:0.0056569274,WP_183742413.1-unclassified-Janthinobacterium:0.0401025669)73.9/98:0.0057369921,(((((WP_139090581.1-Janthinobacterium-lividum:0.0618252605,(WP_230539136.1-Janthinobacterium-sp.-FW305-129:0.0252800652,WP_101482628.1-unclassified-Janthinobacterium:0.1956898703)40.5/100:0.0126213960)77.2/94:0.0084861515,WP_094442213.1-Janthinobacterium-sp.-PC23-8:0.0523375707)77.2/94:0.0056633632,WP_215403374.1-Janthinobacterium-sp.-JC611:0.0227652632)75.6/100:0.0057227739,WP_102124214.1-Janthinobacterium-sp.-ROICE36:0.0287944117)74.9/99:0.0057989571,WP_096238059.1-Janthinobacterium-svalbardensis:0.0170596015)75.5/99:0.0056214802)100/99:0.0904642564,(WP_099917421.1-Massilia-psychrophila:0.1032831265,(WP_205319084.1-Glaciimonas-sp.-PAMC28666:0.0583760453,WP_092396716.1-Collimonas-sp.-OK412:0.0965285042)68.3/100:0.0114175888)95.8/100:0.0397124240)67/99:0.0070523118,WP_023495308.1-Methyloglobulus-morosus:0.1262428408)98.3/100:0.0717777317,WP_217900122.1-Noviherbaspirillum-humi:0.1600260154)76/20:0.0164894912,(((WP_162020603.1-Undibacterium-crateris:0.1187712469,WP_186861877.1-Undibacterium-griseum:0.0609573652)50/73:0.0112935433,WP_186884883.1-Undibacterium:0.0329670785)99.7/100:0.1096548676,((((((WP_207000528.1-Trinickia-sp.-DHG64:0.1034757553,WP_124149903.1-Paraburkholderia-dinghuensis:0.0801163770)55.5/100:0.0432674707,(WP_266250364.1-Paraburkholderia-megapolitana:0.1335684112,WP_026121090.1-Paraburkholderia-kururiensis:0.0811938837)68.4/100:0.0471784590)97.2/100:0.0734794389,(WP_179404315.1-Burkholderia-guangdongensis:0.1285680234,((WP_063494936.1-Paraburkholderia-phytofirmans:0.0252035473,(WP_254318368.1-Paraburkholderia-sp.-CNPSo-3272:0.0049414482,WP_042262232.1-Paraburkholderia-heleia:0.0522548670)95.3/97:0.0334860000)99.3/100:0.0833718361,WP_038711716.1-Burkholderia-sp.-lig30:0.1661000815)80.8/89:0.0426823875)86.9/89:0.0288222163)37.2/58:0.0337124209,((((((((((((WP_059459219.1-Burkholderia-vietnamiensis:0.1761736227,WP_121276956.1-Trinickia-fusca:0.0916711602)41.1/100:0.0305605849,WP_134195781.1-Paraburkholderia-rhizosphaerae:0.1935643452)88.1/99:0.0286626389,WP_136892825.1-Trinickia-sp.-7GSK02:0.0365110613)86.6/99:0.0163933942,WP_201650988.1-Paraburkholderia-sabiae:0.0314389679)60.8/82:0.0121399136,WP_205970438.1-Paraburkholderia-sp.-Tr-20389:0.0311841669)82.3/83:0.0119415009,WP_054928771.1-Paraburkholderia-caribensis:0.0311554445)70.1/95:0.0106561132,(((WP_060092900.1-Burkholderia-vietnamiensis:0.0103938985,WP_226113708.1-Burkholderia-sp.-AU33545:0.0481434046)100/100:0.1155390231,(WP_205976236.1-Paraburkholderia-sp.-Se-20369:0.1809333176,WP_206951398.1-Trinickia-sp.-4D114:0.0562064025)96.8/26:0.0726039536)38.4/12:0.0147196173,WP_059889870.1-Burkholderia-stagnalis:0.1601158392)81.9/10:0.0147310923)89.2/20:0.0218067974,(((WP_175791145.1-Burkholderia-ambifaria:0.0849975495,WP_105772332.1-Burkholderia-multivorans:0.0306907834)53.3/29:0.0108237340,WP_059842355.1-Burkholderia-multivorans:0.0286655613)86.4/19:0.0142769580,WP_061168975.1-Caballeronia-hypogeia:0.0769733924)81.6/23:0.0134827470)88.9/18:0.0157985945,((WP_209935850.1-Paraburkholderia-sp.-LEh10:0.0754313044,WP_060156927.1-Burkholderia-ubonensis:0.0833255434)28.6/26:0.0119616150,(WP_006496666.1-Burkholderia-cenocepacia:0.0227254211,WP_059716874.1-Burkholderia-ubonensis:0.0340243912)87.8/100:0.0192822418)45/22:0.0136753447)74.4/36:0.0068220724,((WP_201138986.1-Paraburkholderia-domus:0.0400926970,WP_266262001.1-Paraburkholderia-madseniana:0.0187840605)99.8/100:0.0916395664,WP_009889585.1-Burkholderia-thailandensis:0.0728025779)96.8/100:0.0515705548)90.9/61:0.0213177827,WP_168792792.1-Paraburkholderia-aromaticivorans:0.0564036218)63.4/57:0.0110576060,((WP_004544228.1-Burkholderia:0.0179969336,WP_069272519.1-Burkholderia-cepacia:0.1706123575)86/97:0.0196097856,WP_059583574.1-pseudomallei-group:0.0325991239)85/97:0.0126025421)97.6/96:0.0692993395)35.3/97:0.0320194966,WP_056136218.1-Massilia-sp.-Root133:0.1467997444)91.1/100:0.0567629824,WP_153720048.1-Spiribacter-sp.-C176:0.2605201302)85.7/87:0.0418361981)89.1/60:0.0271941881)6.3/16:0.0155974317,(WP_247737070.1-Thauera-aromatica:0.0501450936,WP_039805139.1-Azotobacter-chroococcum:0.1124316552)99.1/100:0.1056723803)77.2/95:0.0279855250,((WP_195874647.1-Herminiimonas-contaminans:0.1002795722,WP_041297045.1-Janthinobacterium-sp.-Marseille:0.0790334099)87.7/100:0.0315373950,((WP_076590920.1-Herminiimonas-arsenitoxidans:0.1345634604,WP_206604468.1-Herminiimonas-fonticola:0.0363702269)25.3/82:0.0209059681,WP_188420499.1-Oxalicibacterium-solurbis:0.1276700538)61.6/97:0.0264954766)98.6/100:0.0707844385)98.2/95:0.0837954325)57.1/4:0.0147972639,(((((((((((((WP_089129802.1-Tolypothrix-sp.-NIES-4075:0.0648849697,WP_096563566.1-Scytonema-sp.-NIES-4073:0.0959731288)36.1/65:0.0315833868,(WP_048866388.1-Scytonema-tolypothrichoides:0.0698108613,WP_096563464.1-Scytonema-sp.-NIES-4073:0.0983700935)40.7/76:0.0165277717)99.9/100:0.1330158352,((((((((WP_009344618.1-Cylindrospermopsis-raciborskii:0.0621217930,WP_240868223.1-Cylindrospermopsis-raciborskii:0.0202996488)99.5/100:0.0865781395,(WP_096570274.1-Sphaerospermopsis:0.0360074019,WP_220611028.1-Sphaerospermopsis-torques-reginae:0.0519709178)56.9/97:0.0267477346)98.4/100:0.0688444413,WP_190559626.1-Anabaena-sphaerica:0.0610533205)88.9/77:0.0315523343,WP_237989437.1-Nostoc-sp.-UHCC-0870:0.0668102100)77.4/77:0.0244759310,WP_206263219.1-Nostoc-sp.-UHCC-0702:0.0588911148)50.7/83:0.0305462234,WP_096557790.1-Nostoc-sp.-NIES-4103:0.1265989576)84.5/96:0.0444280429,WP_016866333.1-Fischerella:0.0675037228)72.3/92:0.0256800314,(((WP_015128325.1-Calothrix-sp.-PCC-7507:0.0936515786,WP_017653186.1-Fortiea-contorta:0.1486919339)50.2/98:0.0352706637,((WP_193199005.1-Nostoc-sp.-MG11:0.0437743825,WP_162396505.1-Nostoc-sp.-B-2019:0.0377167258)96.7/100:0.0551962028,WP_214439445.1-Atlanticothrix-silvestris:0.1201938663)89.1/100:0.0464967723)83.4/98:0.0249423512,(((WP_045870014.1-unclassified-Tolypothrix:0.0243100789,WP_190656031.1-Nostoc-linckia:0.0313764431)99.4/100:0.0692101086,(WP_190743070.1-Tolypothrix-sp.-FACHB-123:0.0421360561,WP_190545010.1-Calothrix:0.0297926349)97.1/100:0.0623967734)75.3/100:0.0324411057,WP_086758595.1-Nostoc-sp.-106C:0.0460745762)82.9/80:0.0304026881)41.2/70:0.0225243345)87.4/88:0.0391489231)99.3/100:0.0889863802,((((((((((((WP_102179481.1-Fischerella-thermalis:0.0354117374,((WP_096675774.1-Fischerella-sp.-NIES-4106:0.0451244758,WP_053458209.1-Hapalosiphon-sp.-MRB220:0.0253209732)72.9/89:0.0055510767,(WP_131119525.1-Westiellopsis-prolifica:0.0446019165,WP_016860528.1-Fischerella-muscicola:0.0192511573)85.8/99:0.0116666976)97.8/95:0.0525409512)100/100:0.1317972553,WP_250123015.1-Chroococcidiopsis-sp.-CCMEE-29:0.1561973602)19.3/16:0.0197936425,((WP_190825375.1-Iningainema-tapete:0.1715780528,(WP_015202791.1-Crinalium-epipsammum:0.1628180637,WP_035154029.1-Calothrix-sp.-336/3:0.2552651279)89.6/100:0.0494391908)84/96:0.0586746064,((WP_099102643.1-Nostoc-sp.-Peltigera-malacea-cyanobiont-DB3992:0.0748607649,(WP_190950926.1-Nostoc-punctiforme:0.0280312703,((WP_221704991.1-Desmonostoc-muscorum:0.0415012379,WP_096537650.1-Nostoc-linckia:0.0117808983)56.8/100:0.0082685010,WP_190959861.1-Nostoc:0.0318683420)16.7/26:0.0181345730)98.1/100:0.0708007916)99.6/100:0.0805809003,(((((((WP_193938462.1-Cuspidothrix-issatschenkoi:0.1275931553,(WP_190370804.1-Anabaena-sp.-FACHB-1237:0.1263456388,WP_168537172.1-Anabaena-sp.-UHCC-0253:0.0286430427)21.8/65:0.0156056053)88/84:0.0191456343,(((WP_190392428.1-Dolichospermum-sp.-FACHB-1091:0.0121431703,(WP_028089231.1-Dolichospermum-circinale:0.0168908568,WP_137908967.1-Nostocales:0.0403109813)98.1/100:0.0535662678)99.7/100:0.0913708637,((WP_096666215.1-Dolichospermum-compactum:0.0534602333,(WP_190387835.1-Aphanizomenon:0.0527122117,WP_168466156.1-Aphanizomenon:0.0162440110)48.8/98:0.0127045028)59.7/98:0.0136340369,WP_015080416.1-Anabaena-sp.-90:0.0287974940)95.7/99:0.0284617139)30.5/48:0.0056898860,WP_168502193.1-Anabaena-sp.-UHCC-0187:0.0000024436)83/60:0.0171582749)99.9/100:0.1010063035,(WP_010998622.1-Nostocaceae:0.0506333010,WP_011320078.1-Nostocaceae:0.0513139762)83.9/100:0.0166831136)92.4/100:0.0364917120,(WP_239729951.1-Nodularia-sphaerocarpa:0.0496134972,WP_173403323.1-Nodularia-spumigena:0.0503970905)97.7/100:0.0506322600)42.1/28:0.0064846286,(WP_199337786.1-Nostoc-sp.-FACHB-280:0.0951071120,WP_199337214.1-Nostoc-sp.-FACHB-152:0.0962522561)85.5/28:0.0271532510)47.5/44:0.0129640044,(WP_015138800.1-Nostoc-sp.-PCC-7524:0.0675198646,WP_066425233.1-Anabaena-sp.-4-3:0.0741695890)96/100:0.0532621736)97/100:0.0679344762,(WP_096624129.1-Calothrix-sp.-NIES-3974:0.0630067605,WP_095721999.1-Calothrix-elsteri:0.0778898763)99/100:0.0762679994)85.5/100:0.0217631938)88.9/100:0.0346120872)81.7/27:0.0228765539)98.6/79:0.0555095782,((((WP_015170385.1-Geitlerinema-sp.-PCC-7407:0.2013851334,WP_194029353.1-Lusitaniella-coriacea:0.1686027081)88.2/99:0.0348976161,(((WP_233747769.1-Leptodesmis-sichuanensis:0.1673053263,WP_223049852.1-Leptothermofonsia-sichuanensis:0.0839636376)62.9/72:0.0050154586,(WP_192216758.1-Microcoleus-sp.-FACHB-SPT15:0.1299586355,WP_224345144.1-Kovacikia-minuta:0.0896243869)42.5/76:0.0245767224)89.8/92:0.0236860235,WP_073549891.1-Chroogloeocystis-siderophila:0.1411514858)70.6/98:0.0136174690)89/97:0.0236719603,(WP_190666400.1-Coleofasciculus-sp.-FACHB-501:0.0979944474,WP_190525322.1-Coleofasciculus-sp.-FACHB-T130:0.0463261996)98.4/100:0.0582665459)63.6/21:0.0041804049,(WP_193873404.1-unclassified-Coleofasciculus:0.1065261073,WP_254010234.1-Symplocastrum-sp.-BBK-W-15:0.1092572207)88.4/89:0.0341415118)90.2/29:0.0211970833)0/67:0.0000025862,(WP_224410051.1-Oscillatoria-salina:0.1732867191,WP_008178350.1-Moorena-producens:0.1460120742)76.1/71:0.0456215217)91.5/90:0.0267818698,(WP_190493486.1-unclassified-Microcoleus:0.2013458363,(WP_190531554.1-Coleofasciculus-sp.-FACHB-542:0.1493878039,(((WP_069965800.1-unclassified-Desertifilum:0.1941870669,(WP_073595522.1-Phormidium-ambiguum:0.0636325941,WP_193982193.1-Phormidium-sp.-LEGE-05292:0.0537131439)93.4/100:0.0439260400)90.7/89:0.0497269271,(WP_249064905.1-Argonema-antarcticum:0.0466915149,WP_190468950.1-unclassified-Planktothrix:0.0777715427)32.1/99:0.0193157228)87.8/89:0.0324610681,WP_226578226.1-Microseira-wollei:0.1349071974)88.7/94:0.0278917501)82.5/81:0.0188458347)78.6/82:0.0126131472)73/88:0.0067021931,(((((((((WP_202221768.1-Okeania-sp.-KiyG1:0.0378981722,WP_124146462.1-Okeania-hirsuta:0.0148119731)9/99:0.0118180856,WP_094672483.1-Hydrocoleum-sp.-CS-953:0.0317137453)99.8/100:0.1188354223,WP_071516849.1-Geitlerinema-sp.-PCC-9228:0.1627785965)95.5/100:0.0725058850,(WP_009783380.1-Lyngbya:0.0679236285,WP_046277942.1-Limnoraphis-robusta:0.0395212093)94.7/100:0.0548544304)70/97:0.0364533533,((WP_083623900.1-Planktothrix-serta:0.0347756637,WP_072717475.1-Planktothrix:0.0226129288)72.8/100:0.0171386080,(WP_079676747.1-Planktothrix-sp.-PCC-11201:0.0211089933,(WP_026797127.1-Planktothrix:0.0226773616,WP_214559879.1-Planktothrix-agardhii:0.0349540653)53.3/100:0.0235534505)98/100:0.0616459086)97.2/100:0.0654850395)79.4/86:0.0318476473,WP_190877489.1-unclassified-Planktothricoides:0.1332012115)86.2/86:0.0229049613,(WP_007355666.1-Kamptonema:0.1043334527,((((WP_236507253.1-Tychonema-sp.-BBK16:0.0357625230,WP_096828456.1-Tychonema-bourrellyi:0.0556472623)93.6/100:0.0296788173,WP_242724542.1-Microcoleus-vaginatus:0.0284879932)80.8/100:0.0117844423,WP_015174083.1-Oscillatoria-nigro-viridis:0.0181102167)77.5/100:0.0081231322,((WP_194009709.1-unclassified-Tychonema:0.0319032261,WP_265235823.1-Lyngbya-sp.-CCAP-1446/10:0.0365594381)88.2/100:0.0183594305,WP_193973976.1-Microcoleus-sp.-LEGE-07076:0.0419459409)39.3/93:0.0121551328)97.5/100:0.0613355821)89.4/89:0.0444428168)81.8/74:0.0210179916,(((WP_261197618.1-unclassified-Laspinema:0.0413630752,WP_207087219.1-Phormidium-pseudopriestleyi:0.0326905066)24.5/96:0.0137529723,WP_261204505.1-Laspinema-sp.-D2d:0.0265439212)98.7/100:0.0811482169,WP_200665545.1-Oxynema-sp.-CENA135:0.1694006015)89.5/88:0.0330682600)86.3/94:0.0213960735,(WP_017715496.1-Oscillatoria-sp.-PCC-10802:0.1303863297,(WP_190665495.1-Microcoleus-sp.-FACHB-672:0.0498224119,WP_261220767.1-Ancylothrix-sp.-D3o:0.1496110913)97.1/100:0.0546670541)87.3/97:0.0308152826)90.4/95:0.0208372823)90.8/90:0.0241935802,(((((WP_204137747.1-Halomicronema-sp.-CCY15110:0.0422507621,WP_039724568.1-Leptolyngbya-sp.-LK:0.0604331146)99.5/100:0.1303587041,WP_023076112.1-Leptolyngbya-sp.-Heron-Island-J:0.1626620004)95.4/100:0.0769613373,WP_017324474.1-Synechococcus-sp.-PCC-7336:0.2072826316)84.3/100:0.0307530015,WP_190772446.1-unclassified-Leptolyngbya:0.1346714304)94.7/97:0.0744790226,(WP_017661508.1-Baaleninema-simplex:0.1706667586,WP_008052766.1-Limnospira-indica:0.1864543272)92.4/36:0.0548755460)75.4/27:0.0110960350)93.9/32:0.0471729991,(WP_106909014.1-Microcystis-sp.-MC19:0.1878904247,(WP_015193936.1-Stanieria-cyanosphaera:0.0153809558,WP_096384422.1-Stanieria-sp.-NIES-3757:0.0799944765)99.9/100:0.1352630191)63.5/86:0.0537400230)15.4/21:0.0141959927,WP_015143663.1-Pleurocapsa-sp.-PCC-7327:0.1495711902)72.7/19:0.0104250962,((((((((WP_106232846.1-Pleurocapsa-sp.-CCALA-161:0.1313094264,WP_229639079.1-Waterburya-agarophytonicola:0.0270425608)78.6/100:0.0311538054,WP_096721927.1-Chondrocystis-sp.-NIES-4102:0.0674220326)99.9/100:0.1225993421,WP_006510035.1-Xenococcus-sp.-PCC-7305:0.1002416995)96.2/100:0.0682130282,WP_036478304.1-Myxosarcina-sp.-GI1:0.1261644554)88.1/97:0.0362719876,WP_012595695.1-Rippkaea-orientalis:0.1311463641)63.3/33:0.0207208854,(WP_006529211.1-Gloeocapsa-sp.-PCC-73106:0.2279446272,WP_124977990.1-Aphanothece-sacrum:0.1062959420)92.3/28:0.0574506194)15.5/13:0.0295885283,(((WP_007308862.1-Crocosphaera-watsonii:0.0713615654,WP_107668138.1-Cyanothece-sp.-BG0011:0.0284600550)84.9/62:0.0115078863,WP_009545547.1-Crocosphaera-subtropica:0.0175697790)83.7/62:0.0113015312,WP_008278255.1-Crocosphaera-chwakensis:0.0398856704)99.5/99:0.0957043631)19.8/10:0.0197218556,WP_264322079.1-Zarconia-navalis:0.1708079380)87/25:0.0363493087)93/55:0.0402000899,WP_013324143.1-Gloeothece-verrucosa:0.0691284448)95.8/71:0.0488742204)87.1/65:0.0399013558,(((((WP_239190817.1-Candidatus-Nitrotoga-sp.-HW29:0.0517879034,WP_239287644.1-Candidatus-Nitrotoga-sp.-1052:0.0132859046)84.3/100:0.0145864397,WP_239235002.1-Candidatus-Nitrotoga-sp.-BS:0.0458463540)68.3/99:0.0064348768,WP_239797721.1-Candidatus-Nitrotoga-arctica:0.0170664592)100/100:0.1754658819,((WP_237889684.1-Dissulfurimicrobium-hydrothermale:0.2088323501,WP_237247350.1-Sideroxyarcus-emersonii:0.1633030180)85.2/100:0.0492272866,WP_009207329.1-Sulfuricella-denitrificans:0.1969718753)91.2/80:0.0506786452)67.9/60:0.0146433546,(WP_044881819.1-Neochlamydia-sp.-EPS4:0.0405625965,WP_166155173.1-unclassified-Neochlamydia:0.0113493622)100/100:0.2723441174)88.4/64:0.0275198644)88/33:0.0351430966,(WP_054689510.1-Desulfosarcina-cetonica:0.1641264471,WP_066347484.1-Geminocystis-sp.-NIES-3708:0.2790833602)78.4/35:0.0616608540)76.4/37:0.0098622182,(WP_250680171.1-Geomonas-sp.-Red32:0.1260694550,WP_199381999.1-Geomesophilobacter-sediminis:0.1686441929)94.9/95:0.0651632533)91.8/98:0.0397968581,WP_072282412.1-Syntrophotalea-acetylenivorans:0.1790653895)94.1/95:0.0590592792,((((WP_205222281.1-Desulfogranum-marinum:0.1609001284,(WP_205242362.1-Desulfobulbus-alkaliphilus:0.1132845404,WP_267928740.1-Desulfolithobacter-dissulfuricans:0.1259165191)89.7/100:0.0519899814)89.5/100:0.0518075788,WP_028582690.1-Desulfogranum-japonicum:0.1922489566)93.8/100:0.0603507700,(WP_137423090.1-Desulfoglaeba-alkanexedens:0.2412270427,WP_027714505.1-Desulfuromonas-sp.-TF:0.1750121487)85.8/99:0.0336080520)87.2/85:0.0233468576,(((WP_163339734.1-Desulfopila-sp.-IMCC35008:0.2191477815,WP_136805251.1-Desulfosediminicola-flagellatus:0.1743545893)48.4/84:0.0202276598,WP_073615262.1-Desulfopila-aestuarii:0.1732020001)95.8/100:0.0749507143,(WP_228856481.1-Desulfomarina-profundi:0.2418939754,WP_183349429.1-Desulfoprunum-benzoelyticum:0.1214078674)58/100:0.0360697318)94.6/99:0.0509635651)43.4/40:0.0126504510)93.1/65:0.0355490824,(WP_015405619.1-Desulfocapsa-sulfexigens:0.2069924656,(WP_040200185.1-Geoalkalibacter-subterraneus:0.1713309399,WP_053550565.1-Desulfuromonas-soudanensis:0.1703488837)65.8/96:0.0593638279)71.4/31:0.0112100154)74.5/40:0.0099008603,WP_205218910.1-Desulforhopalus-vacuolatus:0.2622654411)39.5/44:0.0086347722,(WP_103114677.1-Geothermobacter-hydrogeniphilus:0.2061561694,WP_221248821.1-Desulfuromonas-versatilis:0.1852216984)96.6/96:0.0944479071)84.7/43:0.0332245321,(((WP_250024752.1-Cupriavidus-campinensis:0.1335502027,WP_094797980.1-Bordetella-genomosp.-5:0.1357607408)94.2/100:0.0727496275,WP_227181580.1-Leeia-sp.-TBRC-13508:0.2328791633)83.3/99:0.0304350084,(WP_048706771.1-Thauera-humireducens:0.2621827793,((((WP_006216420.1-Achromobacter-piechaudii:0.0974562092,WP_219991400.1-Achromobacter-sp.-ES-001:0.0703821726)87.4/97:0.0438985900,WP_183017169.1-Achromobacter-sp.-UMC46:0.0419371307)82.9/60:0.0280652317,WP_100503206.1-Achromobacter-ruhlandii:0.1223960314)84.5/60:0.0467556547,WP_175192416.1-Achromobacter-deleyi:0.0299729908)99.8/100:0.1549732573)54.3/35:0.0391185071)97.4/34:0.0566382159)8.2/25:0.0087551756)82.7/6:0.0250295311,(((WP_212718705.1-unclassified-Synechocystis:0.0000020129,(WP_228015966.1-Synechocystis-salina:0.0828101892,WP_010873214.1-unclassified-Synechocystis:0.0586420973)89.6/100:0.0249394718)100/100:0.2775339631,(WP_099797666.1-Thermostichus-lividus:0.2814167674,WP_196167318.1-Pseudomonas-monteilii:0.1869463122)55.8/96:0.0928724052)91.3/54:0.0662641251,(((((WP_019918777.1-Methyloversatilis-discipulorum:0.0316726797,WP_018229738.1-Methyloversatilis-universalis:0.0533579407)95.4/100:0.0874285671,WP_183409942.1-Litorivivens-lipolytica:0.3158786284)6.2/100:0.0576038164,(WP_101941536.1-Uliginosibacterium-sp.-TH139:0.0804010115,WP_095523004.1-Candidatus-Dactylopiibacterium-carminicum:0.0783411050)96/100:0.0866356399)98.6/100:0.1136090796,(WP_109041087.1-Azospira-sp.-I13:0.0962076762,WP_014237023.1-Azospira-oryzae:0.0901128595)96.3/100:0.0827245219)21/93:0.0333518919,WP_067106371.1-Methylobacillus-sp.-MM3:0.1778653101)91.9/79:0.0501052271)0/43:0.0233434793)88.2/82:0.0283336427,((((WP_129151777.1-Achromobacter-aloeverae:0.0825042998,WP_088599355.1-unclassified-Achromobacter:0.0595562800)98.1/100:0.0936410609,WP_057654143.1-Bordetella-sp.-N:0.1356266657)80.5/99:0.0417936023,WP_086067427.1-Bordetella:0.1683246809)80.4/100:0.0471216390,(WP_066641528.1-Bordetella-sp.-H567:0.0661380933,(WP_086058993.1-Bordetella-genomosp.-9:0.1494555568,WP_094856450.1-Bordetella-genomosp.-10:0.1129614483)96.7/100:0.0809874241)85.5/98:0.0689049566)98.2/100:0.0911664954)90.5/55:0.0372923205)94.6/99:0.0588975395,(WP_188630571.1-Oceanisphaera-marina:0.0919855110,(WP_104485062.1-Oceanisphaera-arctica:0.0850523767,WP_204795415.1-Oceanisphaera-litoralis:0.0421234398)17.8/51:0.0189251245)99.9/100:0.1215284350)89/98:0.0399053672,(WP_134696535.1-Aeromonas-taiwanensis:0.3547530753,(WP_094838100.1-Bordetella-genomosp.-4:0.1479042255,WP_160351730.1-Bordetella-sp.-15P40C-2:0.1693442315)97.4/100:0.1148605285)98.7/95:0.1228644466)90.4/33:0.0516020957,((((WP_132257458.1-Paucimonas-lemoignei:0.0677732039,WP_119785517.1-Noviherbaspirillum-sedimenti:0.0469898910)100/100:0.2236256443,(WP_026470910.1-Alkanindiges-illinoisensis:0.1919309910,(WP_248147450.1-Moraxella-osloensis:0.2351131104,WP_092690500.1-Acinetobacter-kyonggiensis:0.1419868435)80.6/100:0.0778491574)98.9/100:0.1590553762)95.7/100:0.0947110764,((WP_148717329.1-Chitinolyticbacter-meiyuanensis:0.2656880027,WP_123210144.1-Stagnimonas-aquatica:0.4219814412)62.6/100:0.0994207786,(KAI6244242.1-Aphelenchoides-fujianensis:0.3396988172,KAI6170547.1-Aphelenchoides-bicaudatus:0.4263407975)98.6/100:0.1974691498)92.9/98:0.0865122771)94.5/83:0.0715407017,(WP_208005626.1-Oceanisphaera-pacifica:0.0877629089,(WP_087034853.1-Oceanisphaera-profunda:0.0665368924,WP_087034873.1-Oceanisphaera-profunda:0.0729592933)50.8/100:0.0487734108)92.9/100:0.0558485959)88.8/35:0.0365384660)83.1/33:0.0267015638)87.8/51:0.0387690154,WP_108652490.1-Dongshaea-marina:0.1707512168)93/88:0.0553058337,((((((((((((((((((((((WP_236369955.1-Aeromonas-veronii:0.0399601601,WP_113736183.1-Aeromonas-caviae:0.0321814497)72.2/100:0.0125894851,((WP_042019775.1-Aeromonas-sobria:0.0440923485,WP_139495680.1-Aeromonas-veronii:0.0209328942)98.2/100:0.0456588698,(WP_098969895.1-Aeromonas-sp.-CU5:0.0393334962,WP_159163188.1-Aeromonas-veronii:0.0400730382)94.1/100:0.0327028787)89.4/100:0.0261339117)86.1/76:0.0212128597,((((WP_263173612.1-Shewanella-sp.-SM73:0.1184910611,((WP_263191039.1-Shewanella-sp.-SM87:0.0589118020,WP_263255112.1-Shewanella-sp.-SM32:0.0404814280)82.3/93:0.0160517481,(WP_263178828.1-Shewanella-sp.-SM101:0.0348625076,WP_109248233.1-Shewanella-algae:0.0626342262)90.4/100:0.0276998227)83.8/97:0.0300539737)95.7/100:0.0592802661,(WP_220777726.1-Shewanella-hafniensis:0.0219621491,WP_055647506.1-Shewanella-sp.-Sh95:0.0309451979)93.9/100:0.0371993328)86.6/100:0.0299299712,(WP_082969903.1-Aeromonas-dhakensis:0.0919155035,WP_104454419.1-Aeromonas-sp.-ASNIH2:0.0466272142)73.5/72:0.0073730229)81.9/74:0.0335615336,WP_201035193.1-Aeromonas-media:0.0891703909)96.7/99:0.0469541018)92.2/99:0.0372974650,((((WP_080988732.1-Aeromonas-caviae:0.0355801407,WP_108536035.1-Aeromonas-sp.-HMWF014:0.0306118405)93/100:0.0291518412,WP_252615985.1-Aeromonas-hydrophila:0.0617473092)85.7/97:0.0171381261,((WP_113739100.1-Aeromonas-veronii:0.0471597269,WP_226032029.1-Aeromonas-hydrophila:0.0349943885)28.4/66:0.0110826839,(WP_214003767.1-Aeromonas-caviae:0.0303774762,(WP_139459904.1-Aeromonas-veronii:0.0462089497,WP_041208446.1-Aeromonas-jandaei:0.1106867870)99.3/97:0.0716666530)80.1/68:0.0161223884)0/69:0.0000022111)77.4/97:0.0066690475,WP_143237783.1-Aeromonas-caviae:0.0460723597)46.2/98:0.0225802408)96.1/99:0.0551824761,(((WP_268010485.1-Aeromonas-dhakensis:0.0602715471,WP_098985072.1-Aeromonas-sp.-CA23:0.0370427443)98.7/100:0.1030280151,(WP_181268965.1-Aeromonas-veronii:0.0299726348,(WP_129146711.1-Aeromonas-caviae:0.0216596773,(WP_081843667.1-Aeromonas-taiwanensis:0.0173225203,WP_139744464.1-Aeromonas-media:0.0356529366)81.7/100:0.0211555725)46.7/79:0.0174597229)88/79:0.0556688917)87.2/99:0.0597279591,WP_200561028.1-unclassified-Erwinia:0.2449576670)93.1/98:0.0652820523)89.1/98:0.0366634228,(WP_059168599.1-Aeromonas-salmonicida:0.0537315646,(WP_033115905.1-Aeromonas-lacus:0.0383965841,((WP_223920301.1-Aeromonas-caviae:0.0294520439,WP_226058924.1-Shewanella-sp.-SR1:0.1009002102)78.3/100:0.0146722990,(WP_058394234.1-Aeromonas-salmonicida:0.0248462217,WP_257690725.1-Aeromonas-veronii:0.0401358401)78.7/100:0.0176560831)76.7/100:0.0198288834)92.5/100:0.0360552269)76.9/87:0.0347790574)96.6/100:0.0474193641,((WP_139750984.1-Aeromonas-media:0.0270631674,WP_224430864.1-Aeromonas-rivuli:0.0239983871)83/100:0.0233655562,WP_041996113.1-Aeromonas-bivalvium:0.0417132350)95.4/100:0.0441701535)75.7/8:0.0064665351,((WP_216990617.1-Aeromonas-sp.-FDAARGOS-1409:0.1279249759,WP_265776268.1-Aeromonas-veronii:0.0921639595)47.3/85:0.0313577067,WP_040098646.1-Aeromonas:0.0726556975)74.9/8:0.0106830415)0/3:0.0000023760,((WP_263309349.1-Aeromonas-caviae:0.0345989817,(WP_041217344.1-Aeromonas-hydrophila:0.0347963551,WP_255912544.1-Aeromonas-sp.-JL9:0.0288209417)74.4/42:0.0057983926)87.1/42:0.0139805441,WP_257686696.1-Aeromonas-veronii:0.0294686348)84.9/97:0.0139271645)79.3/86:0.0100303553,WP_265458754.1-Aeromonas-salmonicida:0.0637046966)76.7/27:0.0068046734,((WP_180911080.1-Aeromonas-caviae:0.0345199215,WP_264818242.1-Aeromonas-caviae:0.0228355982)92.9/100:0.0174255991,WP_125597700.1-Aeromonas-salmonicida:0.0229537255)0/22:0.0000023854)77.6/69:0.0056730819,WP_163128050.1-Aeromonas-allosaccharophila:0.0661440054)85.9/80:0.0160702994,WP_223945983.1-Aeromonas-caviae:0.0735213313)93.6/94:0.0450021135,WP_265468300.1-Aeromonas-salmonicida:0.0911374436)89.1/83:0.0291805507,(((((WP_236563281.1-Aeromonas-veronii:0.1105539993,((((((((WP_247163170.1-Escherichia-coli:0.0174129135,WP_038644959.1-Pantoea:0.0681128274)99.7/100:0.0754778926,(WP_252422582.1-Aeromonas-taiwanensis:0.0167920594,WP_263719954.1-Aeromonas-caviae:0.0756148201)73.9/99:0.0365281875)90/100:0.0335630002,WP_033131297.1-Aeromonas-aquatica:0.0228667053)94.3/98:0.0361693059,WP_241344952.1-Aeromonas-sp.-MR16:0.0128010968)86.4/98:0.0193559166,WP_160838676.1-Aeromonas-caviae:0.0670971674)91.7/100:0.0313639893,(WP_116547128.1-Plesiomonas-shigelloides:0.0804292880,(((WP_139496062.1-Escherichia-coli:0.0566436877,WP_215269322.1-Aeromonas-dhakensis:0.0351987298)45/98:0.0055549801,WP_201996661.1-Aeromonas-veronii:0.0327644700)82.2/47:0.0112486124,WP_235463836.1-Aeromonas-australiensis:0.0122838844)35.1/47:0.0091033518)86.7/99:0.0227752412)85.3/99:0.0265585420,WP_049048056.1-Aeromonas-hydrophila:0.1049918700)79/96:0.0277737587,WP_167569591.1-Aeromonas-veronii:0.0658934656)96.1/100:0.0602199801)80/89:0.0179929313,WP_076573677.1-Aeromonas-sp.-RU39B:0.1982152625)75.2/88:0.0154918901,(WP_213320225.1-Aeromonas-caviae:0.0599590790,WP_203764723.1-Aeromonas-caviae:0.0551959063)98.6/100:0.0802219560)87.3/79:0.0184686746,(WP_100646402.1-Aeromonas-veronii:0.0419448465,((WP_252601295.1-Aeromonas-dhakensis:0.0209174477,WP_139432874.1-Aeromonas-dhakensis:0.1176300926)92.8/100:0.0299668658,WP_106553824.1-Aeromonas-hydrophila:0.0470370026)90.7/100:0.0299328248)99.9/100:0.0960579682)84.4/77:0.0160953532,((((WP_201942221.1-Aeromonas-veronii:0.1051172029,((((((WP_268854930.1-Aeromonas-enteropelogenes:0.0897985246,WP_223380193.1-Aeromonas-enteropelogenes:0.0296253420)74.4/78:0.0063659606,WP_042657830.1-Aeromonas:0.0401803176)74.5/75:0.0115234854,WP_139692403.1-Aeromonas-sobria:0.0292173848)75.6/26:0.0063249042,WP_203131090.1-Aeromonas-jandaei:0.0464105966)29.3/26:0.0041649850,WP_111901368.1-Aeromonas-veronii:0.0347907117)89.8/85:0.0174866375,WP_159143398.1-Aeromonas-veronii:0.0059926226)94.3/93:0.0393713183)91/97:0.0397339273,((WP_236321701.1-Aeromonas-veronii:0.0622311777,WP_257119864.1-Aeromonas-hydrophila:0.0151471576)89.6/93:0.0459294226,(WP_257707542.1-Aeromonas-caviae:0.0453094865,WP_182951616.1-Aeromonas-sp.-WP2-W18-CRE-05:0.0546437377)98.7/100:0.0701376318)79.5/58:0.0157372270)94.5/98:0.0470844342,(((WP_042864139.1-Aeromonas-caviae:0.0272709023,WP_225159881.1-Aeromonas-caviae:0.0427537018)97.8/99:0.0688648639,WP_045790994.1-Aeromonas-hydrophila:0.1238006494)71.1/47:0.0172230187,(WP_252995952.1-Aeromonas-encheleia:0.1681704020,(WP_244056308.1-Aeromonas-caviae:0.0825866947,WP_202046159.1-Aeromonas-media:0.1102040246)96.6/99:0.0704701670)82.7/51:0.0258784478)96.6/80:0.0546553150)20.6/69:0.0187355839,WP_069554789.1-Aeromonas-sp.-DNP9:0.1012303622)92.4/79:0.0456274156)52.4/75:0.0302290802)75.3/79:0.0251742643,WP_234664799.1-Aeromonas-allosaccharophila:0.1079299864)81.7/66:0.0225348760,((((WP_268421127.1-Aeromonas-media:0.0537031283,WP_111910157.1-Aeromonas:0.0176272421)67.3/100:0.0127735832,WP_128343177.1-Aeromonas-caviae:0.0400085095)96.5/100:0.0432340212,WP_268037199.1-Aeromonas-dhakensis:0.0783093463)70.9/95:0.0109108828,(WP_139733908.1-Aeromonas-veronii:0.0128414274,((WP_234579942.1-Aeromonas-salmonicida:0.0613449047,(WP_234676856.1-Aeromonas-rivipollensis:0.0152732658,WP_269382839.1-Aeromonas-hydrophila:0.0670347568)71.7/80:0.0069870589)12.5/77:0.0085544782,WP_222134508.1-Vibrio-cholerae:0.1238580957)66.8/97:0.0084698822)98.9/100:0.0634268604)87.2/98:0.0214200096)76.5/51:0.0140001827,WP_139406182.1-Aeromonas-veronii:0.2147131616)76/51:0.0168717436,(((WP_245131329.1-Aeromonas-veronii:0.0942851610,WP_210550341.1-Aeromonas-sp.-SrichE-2G:0.2115795278)81.1/83:0.0245815735,(((WP_257692500.1-Aeromonas-caviae:0.1240255602,WP_113719982.1-Aeromonas-salmonicida:0.0834858037)93.7/86:0.0790236443,((WP_050666161.1-Aeromonas-schubertii:0.0467115360,WP_059168611.1-Aeromonas-salmonicida:0.0925712369)69.8/77:0.0287539348,WP_210600017.1-Aeromonas-sp.-MrichA-1:0.0320534892)85.6/87:0.0274113405)81.4/93:0.0288112265,(WP_225159845.1-Aeromonas-caviae:0.0859645329,(WP_163135231.1-Aeromonas-rivipollensis:0.0282077681,WP_241279715.1-Aeromonas-sp.-MR19:0.0279181830)100/100:0.1665367223)69.7/94:0.0343409043)78.6/78:0.0158036190)19.3/77:0.0190830729,((WP_101347551.1-Aeromonas-sobria:0.0517438757,WP_201955411.1-Aeromonas-veronii:0.0522860144)95.5/100:0.0561748929,(WP_136501435.1-Aeromonas-veronii:0.1165683265,(WP_268029297.1-Aeromonas-hydrophila:0.0874256941,WP_201942265.1-Aeromonas-veronii:0.1029817256)97.8/100:0.0919129801)88.2/98:0.0459196164)70.8/90:0.0155881330)89/87:0.0332895518)49.4/41:0.0185964155,(((WP_148612677.1-Aeromonas-sobria:0.0260146396,((WP_201355655.1-Aeromonas-hydrophila:0.0392467178,WP_201951905.1-Aeromonas-veronii:0.0285421160)71.5/100:0.0060184038,WP_171268810.1-Aeromonas-media:0.0250158431)13.7/67:0.0179652026)99.7/100:0.0953931076,((((WP_212539243.1-Aeromonas-salmonicida:0.0315163315,((WP_187590672.1-Aeromonas-hydrophila:0.0954849930,WP_060586794.1-Aeromonas-schubertii:0.0438624457)0/33:0.0136553067,WP_017782281.1-Aeromonas-dhakensis:0.0311815537)99.8/100:0.0785179896)91/92:0.0213802393,WP_201904077.1-Aeromonas-veronii:0.0505488753)87/92:0.0173195454,WP_206874356.1-Aeromonas-veronii:0.0096440981)76.6/90:0.0103342631,(WP_042063659.1-Aeromonas-allosaccharophila:0.0298702962,WP_256684552.1-Aeromonas-salmonicida:0.0286609750)56.7/97:0.0194242301)85.5/91:0.0244533249)91.2/87:0.0370379291,WP_042649704.1-Aeromonas-media:0.0813416827)94.6/89:0.0495023721)89.1/58:0.0261125146,WP_253626250.1-Aeromonas-hydrophila:0.1102889351)25.2/10:0.0060724399,((WP_064338398.1-Aeromonas-veronii:0.1106024570,((WP_227095090.1-Aeromonas-veronii:0.0908058743,WP_152109808.1-Plesiomonas-shigelloides:0.0406005866)92.5/99:0.0544111177,WP_168366848.1-Dickeya-zeae:0.1349160722)94.7/100:0.0565952604)76.9/74:0.0421414384,(WP_224211331.1-Aeromonas-enteropelogenes:0.1270766369,WP_082041448.1-Aeromonas-eucrenophila:0.1825656611)93.9/92:0.0574816534)52.3/58:0.0109480094)94.7/77:0.0524254394)88.9/65:0.0283696061,((((WP_139871681.1-Candidatus-Methylopumilus-universalis:0.1315619013,(WP_139929800.1-Candidatus-Methylopumilus-universalis:0.2074030876,WP_140005423.1-Candidatus-Methylopumilus-planktonicus:0.0667465556)82.8/100:0.0928996843)100/100:0.2245018940,((WP_151054911.1-Cellvibrio-sp.-KY-YJ-3:0.1458389072,WP_078044787.1-Cellvibrio-sp.-PSBB023:0.1577995040)41.8/92:0.0338384407,(WP_012489013.1-Cellvibrio-japonicus:0.1096011481,WP_087465811.1-Cellvibrio-sp.-PSBB006:0.1367885692)58.8/72:0.0328504048)68.3/91:0.0653847384)40.7/90:0.0273911752,((WP_097405671.1-Kosakonia:0.0462173756,WP_161518249.1-Kosakonia-sp.-MH5:0.0493090692)87.5/92:0.0448727122,WP_128482629.1-Kosakonia-cowanii:0.0974655642)99.8/100:0.1263877624)67.2/66:0.0252469176,(((WP_047682696.1-Xenorhabdus:0.0466284155,WP_099124588.1-Xenorhabdus-stockiae:0.0616307476)98.3/100:0.0853023447,(WP_074024273.1-Xenorhabdus-eapokensis:0.0554222894,WP_217470569.1-Photorhabdus-akhurstii:0.0699157837)97.4/100:0.0764047504)89.5/93:0.0522875523,(WP_048221111.1-Citrobacter:0.1202821601,WP_263807164.1-Salmonella-enterica:0.1168011076)100/100:0.1382371493)72.4/84:0.0338771788)87/60:0.0408863576)85.1/80:0.0210600552,((WP_035600513.1-Edwardsiella-tarda:0.2098296944,WP_125117341.1-Aeromonas-caviae:0.2090212416)59/93:0.0366104570,(WP_202782010.1-Enterobacter-asburiae:0.0746752059,WP_144817260.1-Pseudescherichia-sp.-L3:0.0261225921)99.9/100:0.1341518113)93.1/93:0.0378005333)24.6/70:0.0186102267,(((WP_085037002.1-Cronobacter-sakazakii:0.1565895631,WP_076770125.1-Kosakonia-cowanii:0.1060618119)96.6/100:0.1190495396,WP_216981205.1-Lelliottia-amnigena:0.2041871733)94.6/100:0.0843768114,((WP_227534785.1-Erwinia-rhapontici:0.1112317241,WP_017800228.1-Erwinia-toletana:0.1189904458)83.9/100:0.0605943055,WP_255851625.1-Erwinia-persicina:0.1426445584)0/68:0.0289551200)98.2/98:0.1011030191)74.1/70:0.0473650042,((((((((WP_172912938.1-Pantoea-stewartii:0.1097942962,WP_145394131.1-Pantoea-sp.-SJZ147:0.0985682126)98.9/100:0.1469863650,WP_086868392.1-Kosakonia-pseudosacchari:0.3764102769)98.1/60:0.1379499303,((WP_043158915.1-Aeromonas-hydrophila:0.2404643112,WP_153752706.1-unclassified-Citrobacter:0.3199919137)87.9/60:0.0630457921,WP_097364011.1-Edwardsiella-tarda:0.2206876239)14.7/46:0.0439331544)0/49:0.0039171615,((((((((((((((((((WP_167868984.1-Shewanella-sp.-WXL01:0.2860336863,(WP_222525605.1-Ferrimonas-balearica:0.1230903273,WP_013344594.1-Ferrimonas-balearica:0.0971506899)90.3/98:0.0856388808)89.7/73:0.0570111933,((((((((WP_142941196.1-Aliikangiella-marina:0.2303020546,(WP_081148784.1-Cognaticolwellia-beringensis:0.0413533091,WP_007634918.1-Paraglaciecola-psychrophila:0.0309686069)99.6/100:0.1298692495)66.2/83:0.0385262349,WP_068546663.1-Thalassotalea-crassostreae:0.1559768730)75.2/93:0.0461017294,((((((((WP_160153838.1-Microbulbifer-sp.-ALW1:0.1854632869,((WP_263600466.1-Microbulbifer-agarilyticus:0.1142439811,WP_078085574.1-Microbulbifer-mangrovi:0.0654615889)66.6/95:0.0162626497,WP_029249715.1-Microbulbifer-agarilyticus:0.0838539686)94.3/95:0.0729880677)94.1/100:0.0736988726,((WP_138624780.1-Pseudoalteromonas-sp.-S4492:0.1612610331,WP_130050482.1-Pseudoalteromonas-sp.-MEBiC-03485:0.1017627974)55.7/88:0.0635252303,(WP_189610147.1-Saccharospirillum-salsuginis:0.2033184058,WP_111640782.1-Marinimicrobium-alkaliphilum:0.2196832900)81.6/88:0.0524591040)94.5/99:0.0715441155)6.1/69:0.0298415672,WP_118961222.1-Colwellia-sp.-RSH04:0.2583804598)8.3/76:0.0234625199,WP_186505417.1-Neptunicella-marina:0.1549065426)95.5/99:0.0651198643,(WP_102798355.1-Bowmanella-denitrificans:0.0481014735,WP_214344498.1-Bowmanella-yangjiangensis:0.0411543714)100/100:0.1293122549)75.8/98:0.0263751298,(WP_124026287.1-Alteromonas-sediminis:0.1195746796,WP_100655941.1-Alteromonas-flava:0.1681239669)99.2/99:0.1160281364)79.1/86:0.0217965197,((WP_216048042.1-Aliiglaciecola-lipolytica:0.1309059530,(WP_057181266.1-Colwellia:0.1444973073,WP_009839692.1-Pseudoalteromonas-tunicata:0.3766901377)78.6/80:0.0370246007)88.6/13:0.0399803116,(WP_131095783.1-Alteromonas-sp.-KUL42:0.0701611697,(((((((((((((WP_100972994.1-Alteromonas-sp.-MB-3u-76:0.0826108884,WP_269520955.1-Alteromonas-sp.-BMJM2:0.0411423366)54.7/100:0.0261065390,(WP_071816068.1-Alteromonas-sp.-V450:0.0775116616,WP_198816918.1-Alteromonas-sp.-IB21:0.0183722578)83.9/100:0.0188451393)63.2/97:0.0156001627,WP_163106912.1-Alteromonas-genovensis:0.0300643490)81.9/95:0.0158595623,WP_057795858.1-Alteromonas-stellipolaris:0.0305065966)93.2/42:0.0233646003,(WP_202262575.1-unclassified-Alteromonas:0.0406206671,WP_187447473.1-Alteromonas-sp.-BZK5:0.0173357307)75.3/80:0.0052587021)0/24:0.0000022787,(WP_063457834.1-Alteromonas-stellipolaris:0.0284704405,(WP_202940767.1-Alteromonas-macleodii:0.0187743532,WP_061095937.1-Alteromonas-macleodii:0.0335467606)84.1/45:0.0106747414)76.1/34:0.0055371304)77.8/25:0.0056602856,(WP_156078883.1-Alteromonas:0.0526504094,(WP_044058190.1-Alteromonas-australica:0.0345507153,WP_159580202.1-Alteromonas-sp.-KUL106:0.0172370468)75.1/92:0.0052701840)74.4/66:0.0058079413)0/32:0.0000021436,WP_239220085.1-Alteromonas-sp.-MmMcT2-2:0.0228011945)0/33:0.0000027138,WP_014977514.1-Alteromonas-macleodii:0.0286462251)87.7/86:0.0274977553,(WP_130638173.1-Alteromonas-sp.-KUL49:0.0509683574,WP_115592514.1-Alteromonas-aestuariivivens:0.0344913756)87.9/89:0.0204744812)39.2/97:0.0066390942,WP_163085646.1-Alteromonas-profundi:0.0411504265)88.8/95:0.0171257224,(((WP_018984642.1-Salinimonas-chungwhensis:0.0983776026,WP_119499891.1-Alteromonas-sp.-RKMC-009:0.0406383087)59.6/44:0.0275711343,(((WP_265616768.1-Alteromonas-sp.-ASW11-7:0.0905850300,WP_265607972.1-Alteromonas-sp.-ASW11-130:0.0631109027)68.5/44:0.0060095027,WP_216026947.1-Aestuariibacter-sp.-A3R04:0.0595335433)75.6/46:0.0091227424,WP_216028209.1-Alteromonas-sp.-C1M14:0.0570274006)69.1/32:0.0050290685)85.6/81:0.0125383548,(WP_070176406.1-Alteromonas-lipolytica:0.0243126529,WP_073325432.1-Marisediminitalea-aggregata:0.0478090232)96.3/100:0.0379474258)94.9/98:0.0234387906)90.2/88:0.0175173113,(WP_262993134.1-Alteromonas-sp.-ASW11-19:0.0575266207,(WP_218354448.1-Alteromonas-lipotrueiana:0.0650664753,WP_195811108.1-Salinimonas-marina:0.0465508374)85.8/99:0.0418644569)76.3/97:0.0129535269)94.7/90:0.0327655208)92.3/90:0.0342785481)75.8/12:0.0145113035)97.9/94:0.0757286294,((WP_250459773.1-Microbulbifer-sp.-GX-H0434:0.1059949760,WP_152451766.1-Microbulbifer-sp.-THAF38:0.1094949855)94.9/100:0.0699823858,((((WP_262601261.1-Endozoicomonas-euniceicola:0.0896067689,WP_066015073.1-Endozoicomonas-atrinae:0.0912211553)85.5/100:0.0657720518,WP_257287521.1-Endozoicomonas-sp.-SESOKO2:0.0898404932)95.8/100:0.0884696711,(WP_067585802.1-Endozoicomonas-ascidiicola:0.1194122540,(WP_020580625.1-Endozoicomonas-elysicola:0.0898627033,WP_257293945.1-Endozoicomonas-sp.-YOMI1:0.0902931930)91.2/100:0.0577020867)99.2/100:0.1405017190)94.8/100:0.0928063949,WP_116686321.1-Pelagibaculum-spongiae:0.2234988490)99.7/99:0.1945486065)84.7/40:0.0396051427)70.7/45:0.0209866957)88.3/49:0.0528192308,((((((WP_255857717.1-Bacterioplanoides-sp.-SCSIO-12839:0.2352800981,WP_096084688.1-Agaribacterium-haliotis:0.2206110311)46.7/97:0.0701201974,WP_188151970.1-Teredinibacter-waterburyi:0.1522704007)89.7/96:0.0753644333,(WP_028862352.1-Psychromonas-aquimarina:0.1439415510,WP_187408568.1-Saccharophagus-sp.-K07:0.1221317149)22.2/97:0.0312879036)59.4/96:0.0415554150,WP_251261538.1-Echinimonas-agarilytica:0.1065204488)96.4/100:0.0707053595,(WP_220103444.1-Neiella-holothuriorum:0.0886438372,WP_087505717.1-Neiella-marina:0.1175297208)96.7/100:0.0594922647)88.4/93:0.0483642346,(WP_248714967.1-Agarivorans-sp.-TSD2052:0.0467787660,WP_152781274.1-Agarivorans-sp.-B2Z047:0.0733634414)99.8/100:0.1199622888)48.2/37:0.0489310835)94/99:0.0475580610,(WP_155694061.1-Psychrosphaera-haliotis:0.1160066098,WP_093326971.1-Thalassotalea-agarivorans:0.0771814283)46.5/85:0.0114808171)92.6/95:0.0265992903,(((((WP_047049276.1-Vibrio-mexicanus:0.1008184145,WP_193630602.1-Shewanella-benthica:0.1531321805)68/99:0.0333593482,WP_047874860.1-Photobacterium-aphoticum:0.1564078641)61/97:0.0183483647,WP_020335761.1-Vibrio-natriegens:0.0900730156)87.9/89:0.0242741397,(((WP_050648448.1-Vibrio-sp.-J2-4:0.0915430736,WP_264874461.1-Vibrio-agarivorans:0.0912441982)90.1/89:0.0294056323,(((((WP_167678613.1-Shewanella-aestuarii:0.1134992543,WP_137222113.1-Shewanella-sp.-MEBiC00475:0.0229701238)90.7/92:0.0360115482,WP_188955407.1-Shewanella-ulleungensis:0.0502965219)95.9/95:0.0510041496,((((WP_138577097.1-unclassified-Pseudoalteromonas:0.0850868021,(WP_264390127.1-Pseudoalteromonas-sp.-A3:0.0220153937,(WP_096739047.1-Pseudoalteromonas-atlantica:0.0345971176,WP_248462211.1-Pseudoalteromonas-sp.-2CM37A:0.0161832283)89.9/100:0.0186291914)77.6/98:0.0110958259)89.4/79:0.0258341614,WP_182693438.1-Pseudoalteromonas-sp.-SR45-5:0.0489987138)47.6/67:0.0149928411,WP_256848444.1-Pseudoalteromonas-carrageenovora:0.0219369749)74.3/67:0.0062347637,WP_126005793.1-Pseudoalteromonas-sp.-Xi13:0.0249564458)95.5/98:0.0472148745)91.4/92:0.0364140268,WP_022943097.1-Psychromonas-hadalis:0.1137843697)55.6/75:0.0069955511,(WP_113818410.1-Psychromonas-sp.-B3M02:0.0364517033,(WP_025564776.1-Psychromonas-sp.-SP041:0.0405038406,WP_166372414.1-Psychromonas-sp.-SA13A:0.0143555112)96.1/100:0.0535977945)91.9/100:0.0361259276)93/99:0.0364811747)18.8/9:0.0111179682,((WP_012143346.1-Shewanella-sediminis:0.1530347876,WP_140076235.1-Vibrio-parahaemolyticus:0.0823599101)82.7/92:0.0241727397,WP_205390674.1-Vibrio:0.0733346302)76.2/88:0.0193372430)82.9/14:0.0180842408)87.9/99:0.0177371991,WP_164649367.1-Vibrio-astriarenae:0.0691999118)90.6/98:0.0189004948)82.1/18:0.0143267079,(WP_084884043.1-Vibrio-sp.-qd031:0.0925245058,(WP_269599241.1-Salinivibrio-kushneri:0.0587569580,WP_136483218.1-Vibrio-sp.-H11:0.0559605801)97.4/100:0.0671138139)84.7/17:0.0206622875)43.2/16:0.0180968141,WP_159656123.1-Vibrio-atypicus:0.0874986003)91.7/87:0.0379178728)86.4/60:0.0372397393,WP_016959527.1-Enterovibrio-norvegicus:0.0574076343)6.8/40:0.0142443483,WP_229491984.1-Enterovibrio-norvegicus:0.0903439462)89.8/73:0.0604221854,WP_138986566.1-Neptunomonas-concharum:0.2316802443)94.7/98:0.0730869753,(WP_085356321.1-Neisseria-dumasiana:0.1486049723,((WP_101133177.1-Neisseria-meningitidis:0.0394047782,WP_005552781.1-Aggregatibacter-actinomycetemcomitans:0.1578884250)75.9/82:0.0202748820,(WP_127202078.1-Neisseria-meningitidis:0.0921126242,WP_237092367.1-Neisseria-sp.-ZJ104:0.1173117180)68/74:0.0390888883)82.9/71:0.0328524900)99.6/100:0.1354397944)88/99:0.0795729288,WP_211689658.1-Moritella-sp.-28:0.1782789238)97.2/83:0.0823121677,((((((((((((((((((((((((((((WP_188026786.1-Tolumonas-osonensis:0.0169332818,WP_024870911.1-Tolumonas-lignilytica:0.1049316376)100/100:0.1163057164,(WP_187463805.1-Photobacterium-sp.-BZF1:0.0508205994,WP_039467548.1-Photobacterium-gaetbulicola:0.1080932652)73.3/63:0.0183137470)91.7/23:0.0426413235,WP_047883806.1-Photobacterium-ganghwense:0.0921801476)35.7/3:0.0160948860,WP_146157089.1-Photobacterium-rosenbergii:0.1382119899)12.6/6:0.0167053091,WP_239411815.1-Photobacterium-sp.-OFAV2-7:0.1388726732)90.9/50:0.0486204602,(WP_105076797.1-Pantoea-ananatis:0.2933037204,(((((WP_144135068.1-Kosakonia-radicincitans:0.2133808762,((WP_088208149.1-Enterobacter-cancerogenus:0.0791371137,WP_121932958.1-Enterobacter-hormaechei:0.0806208855)26.9/97:0.0397900546,WP_023330866.1-Enterobacter:0.0426531866)96.3/98:0.0760668503)89.6/99:0.0685414492,WP_120453320.1-Kalamiella-piersonii:0.0854470912)83/96:0.0224075654,(WP_269294270.1-Pantoea-sp.-LMR881:0.0440963275,WP_215846277.1-Pantoea-sp.-EnvD:0.0609665201)98.2/100:0.0659123580)74.3/93:0.0256994884,WP_187508308.1-Pantoea-agglomerans:0.1282088635)58.5/65:0.0186535038,((WP_214216788.1-Rosenbergiella-epipactidis:0.0479197233,WP_241623482.1-Rosenbergiella-epipactidis:0.0223370999)99.7/100:0.1078203864,(WP_192411534.1-Pantoea-sp.-PNT02:0.0546521629,(WP_125311258.1-Pantoea-dispersa:0.0813785551,((WP_150019269.1-unclassified-Pantoea:0.0504912380,(WP_069025179.1-Pantoea-agglomerans:0.0057244711,WP_140917215.1-Pantoea-deleyi:0.0446618250)78.7/92:0.0169264964)95.3/58:0.0283909166,WP_097116658.1-Pantoea-sp.-GL120224-02:0.0110666663)69.8/50:0.0052922980)89/58:0.0434134069)96/100:0.0566060065)88.2/67:0.0327040492)95.6/100:0.0785264555)7.8/68:0.0267655150)76.3/86:0.0319525605,(((WP_200225058.1-Rahnella-aceris:0.0889369235,WP_195814554.1-Rahnella:0.0992593960)87.1/100:0.0380795856,WP_169403689.1-Rouxiella-aceris:0.0615049333)99.2/100:0.0905675845,WP_101064017.1-Moritella-sp.-Urea-trap-13:0.1661836520)42.7/87:0.0248598691)91.9/46:0.0459026729,(WP_117235674.1-Vibrio-maerlii:0.1039679782,(WP_103882085.1-Vibrio-hangzhouensis:0.0440967110,WP_265676699.1-Vibrio-sp.-ZSDZ65:0.0969591473)93.7/100:0.0508409202)68.7/69:0.0395368528)0/7:0.0000024215,((WP_192015074.1-Photobacterium-arenosum:0.0913861845,WP_075764060.1-Photobacterium-proteolyticum:0.1396827261)91.4/87:0.0501111197,(((((WP_045045404.1-Photobacterium-kishitanii:0.0467792177,WP_107264793.1-Photobacterium-phosphoreum:0.0215494441)90.9/98:0.0241252349,WP_045030671.1-Photobacterium-phosphoreum:0.0191057459)97.2/98:0.0465453468,(WP_087853896.1-Photobacterium-andalusiense:0.0358829927,WP_107274737.1-Photobacterium-phosphoreum:0.0848030298)98.8/100:0.0593598869)71.1/34:0.0120640924,((WP_107293979.1-Photobacterium-phosphoreum:0.0849135574,WP_107228425.1-Photobacterium-phosphoreum:0.0982190875)55.1/98:0.0432611540,WP_107295906.1-Photobacterium-phosphoreum:0.0988054054)91.1/97:0.0484577960)57.4/33:0.0206617918,WP_065194916.1-Photobacterium-phosphoreum:0.1097057621)77/96:0.0369042630)1.4/6:0.0454745518)90.6/99:0.0311244232,(WP_171138527.1-Vibrio-plantisponsor:0.0613613843,WP_102966424.1-Vibrio:0.0550571933)99.7/100:0.0812481708)82.1/99:0.0154031811,(WP_230432956.1-Plesiomonas-shigelloides:0.0840432607,WP_222763364.1-Vibrio-fluvialis:0.0741617397)89.3/99:0.0370877620)79/99:0.0251951015,((WP_148519618.1-Vibrio-cholerae:0.0235746595,(WP_167570263.1-Aeromonas-veronii:0.0507291228,WP_000864158.1-Vibrio-metoecus:0.0301983202)85.8/100:0.0264300084)74.3/100:0.0100147943,WP_148520854.1-Vibrio-cholerae:0.0315298445)98.4/100:0.0611641918)87.4/98:0.0332331733,(WP_188008910.1-Grimontia-hollisae:0.1126256091,WP_065173140.1-Photobacterium-kishitanii:0.1004734130)97.7/99:0.0629906880)36.6/39:0.0144760671,(WP_017008437.1-Enterovibrio-norvegicus:0.1015131800,WP_251878087.1-Grimontia-sp.-020920N:0.0915623796)85.4/87:0.0196786477)76.4/25:0.0086822898,WP_180567593.1-Oceanospirillales:0.1342150950)82.6/13:0.0135400219,((WP_102137258.1-Vibrio-cholerae:0.0899785860,(WP_172567459.1-Vibrio-fluvialis:0.0601170529,WP_199354891.1-Vibrio-cholerae:0.0488306038)92.1/100:0.0641659483)91.8/100:0.0558552940,(WP_261874894.1-Vibrio-rarus:0.1180033801,WP_102038478.1-Shewanella-sp.-GutCb:0.1742189475)29/93:0.0385658397)80.3/6:0.0146589706)76.3/10:0.0101209482,((WP_067416136.1-Enterovibrio-coralii:0.0692824339,((WP_261894433.1-Vibrio-porteresiae:0.1255520428,(WP_259522409.1-Shewanella-baltica:0.1262159678,WP_173243873.1-Shewanella-sp.-DC2-4:0.0655403525)96.4/100:0.0707360228)73.3/94:0.0136989099,(WP_171342182.1-Vibrio-cyclitrophicus:0.1290828331,WP_161972844.1-Enterovibrio-baiacu:0.0679876276)97.3/99:0.0737384281)87.5/99:0.0359555433)81.4/98:0.0189877934,WP_149593442.1-Pseudoalteromonas-sp.-FUC4:0.1348246438)95/97:0.0380749372)91.9/72:0.0259383117,WP_014230611.1-Vibrio-sp.-EJY3:0.0996227079)74.8/60:0.0075152064,((((((((WP_011771697.1-Psychromonas-ingrahamii:0.0953307598,WP_182690147.1-Psychromonas-sp.-SR45-3:0.0879647629)99.3/100:0.1043081698,((WP_197173106.1-Pseudoalteromonas-sp.-NZS11_1:0.1028122599,WP_182748079.1-Pseudoalteromonas-sp.-SR41-1:0.1335054883)65.7/100:0.0357315370,WP_166378370.1-Pseudoalteromonas-sp.-Z9A4:0.1267223967)76.1/100:0.0437773635)28.4/51:0.0232423475,(WP_011638275.1-Shewanella-frigidimarina:0.1052339820,(WP_220774251.1-Shewanella-glacialipiscicola:0.0565941793,WP_182751750.1-Shewanella-sp.-SG44-2:0.0899536465)95.8/100:0.0673335013)94.1/92:0.0511316987)82/74:0.0304561379,((((WP_017027407.1-Vibrio-breoganii:0.0844007956,WP_108189654.1-Vibrio-splendidus:0.0999018386)87.4/56:0.0295327216,(WP_102448490.1-Vibrio-splendidus:0.0827308290,WP_243579549.1-Vibrio-splendidus:0.1279638039)72.3/87:0.0136038883)55/55:0.0239015969,WP_063660043.1-Aliivibrio-fischeri:0.0904786246)91.5/96:0.0354602943,(WP_061023752.1-Vibrio-splendidus:0.0468060677,(WP_220768829.1-Shewanella-sp.-MBTL60-007:0.0863828636,(WP_110456309.1-Shewanella-algidipiscicola:0.1424425796,WP_204999274.1-Parashewanella-hymeniacidonis:0.1036114263)83/99:0.0592904503)24.7/97:0.0183739566)92/98:0.0412184689)81.5/55:0.0193346662)64.2/33:0.0061942783,(WP_192022372.1-Shewanella-sp.-WPAGA9:0.1288913215,WP_248975945.1-Shewanella-kaireitica:0.1395370909)87.9/99:0.0446751146)76.2/32:0.0221087081,(WP_182754472.1-Shewanella-sp.-SG41-3:0.0904983000,WP_059744187.1-Shewanella-frigidimarina:0.1671690859)82.1/91:0.0269647699)92.1/74:0.0323025978,(((WP_102941572.1-Vibrio-diazotrophicus:0.1081491400,WP_112353352.1-Shewanella-benthica:0.1404432393)82.4/43:0.0227018090,(((((WP_172851426.1-Vibrio-coralliilyticus:0.1125624772,WP_182742106.1-Shewanella-sp.-SG44-6:0.1751785659)85.8/99:0.0389109956,WP_121852788.1-Parashewanella-spongiae:0.1401628026)46.7/96:0.0379643644,(WP_239850080.1-Shewanella-sp.-Isolate8:0.0705096079,WP_069440067.1-Pseudoalteromonas-tetraodonis:0.1896195561)84.3/93:0.0454395681)64.9/78:0.0367242920,(WP_011716450.1-Shewanella-sp.-ANA-3:0.1595672718,WP_206625521.1-Vibrio-vulnificus:0.1025706705)76.5/94:0.0513111378)87.6/96:0.0340469022,WP_108186198.1-Vibrio-splendidus:0.1265356923)50.9/50:0.0280990200)88.8/69:0.0269130091,((WP_095430409.1-Vibrio-vulnificus:0.0557748311,WP_114994956.1-Grimontia-hollisae:0.1445938507)93.3/100:0.0531066323,WP_088876367.1-Vibrio-mediterranei:0.2089264075)85.2/40:0.0323941718)29.6/49:0.0197614279)82.1/68:0.0210358220,((WP_038175370.1-Vibrio-pacinii:0.1089315183,((WP_020332409.1-Vibrio-fluvialis:0.0417412760,WP_096629284.1-Salinivibrio-sp.-YCSC6:0.1347611601)91.2/100:0.0500578390,WP_171749982.1-Vibrio-sp.-RE88:0.1228702779)32/96:0.0192163946)79.2/95:0.0241653015,WP_193262143.1-Vibrio-parahaemolyticus:0.1801322220)37.3/91:0.0229626626)93.1/68:0.0347866392)90.3/81:0.0284405515,(((((((((((WP_155660100.1-Aliivibrio-fischeri:0.0708654077,WP_095457151.1-Vibrio:0.0783610685)90.2/100:0.0320684166,(WP_252020469.1-Vibrio-sp.-SCSIO-43136:0.0881622427,WP_243584106.1-Vibrio-splendidus:0.1304248342)85.1/100:0.0364429127)93.6/100:0.0454532183,(WP_108452928.1-Enterobacter-cloacae:0.2645298548,WP_166696422.1-Providencia-rettgeri:0.0694890499)84.2/32:0.0587103141)85.3/31:0.0354140934,((WP_161129723.1-Photobacterium-lucens:0.1119582848,WP_031827242.1-Vibrio-parahaemolyticus:0.1722938310)0/34:0.0319744695,(WP_084988554.1-Vibrio:0.0990586525,WP_206634697.1-Vibrio-vulnificus:0.1213385519)84.8/100:0.0365122671)89.9/89:0.0567220022)66.3/28:0.0210409191,WP_220058598.1-Shewanella-sp.-FJAT-53555:0.2128955122)85.8/89:0.0213983411,(WP_248991952.1-Shewanella-profunda:0.1194323729,(WP_045031875.1-Photobacterium-phosphoreum:0.0205498089,WP_107213458.1-Photobacterium-phosphoreum:0.0310883402)99.3/100:0.0872112642)86.9/98:0.0264567157)67.4/35:0.0170963488,(((WP_158107962.1-Vibrio-furnissii:0.0904127639,WP_185890410.1-Vibrio-vulnificus:0.1153088625)23.5/88:0.0391737449,(WP_172855536.1-Vibrio-coralliilyticus:0.1528059568,WP_144217665.1-Shewanella-algae:0.1513236082)60.5/93:0.0460079649)94.4/96:0.0498454931,(((WP_223418564.1-Shewanella-inventionis:0.0805417840,WP_011497090.1-Shewanella-denitrificans:0.1605791598)96.5/99:0.0783635073,WP_234235026.1-Shewanella-sp.-AS16:0.1423056210)34/83:0.0229604257,((((WP_243584123.1-Vibrio-splendidus:0.0793698203,WP_259574191.1-Shewanella-baltica:0.1729916002)8/77:0.0324896812,(WP_102462022.1-Vibrio-splendidus:0.0481538241,WP_187463684.1-Photobacterium-sp.-BZF1:0.0489443472)98.7/100:0.0887848151)38.5/82:0.0266630076,WP_163929979.1-Paraferrimonas-sp.-SM1919:0.0916271758)72.4/99:0.0398928164,WP_087845049.1-Photobacterium-malacitanum:0.0956420991)86.4/97:0.0296517656)76.1/92:0.0289702031)84.7/68:0.0184880706)91.1/68:0.0276114201,(((((WP_102546929.1-Vibrio-breoganii:0.1104720454,WP_102282524.1-Vibrio-cyclitrophicus:0.1254255729)69.5/58:0.0398901962,WP_102461639.1-Vibrio-breoganii:0.0864844642)71.2/51:0.0400756059,(WP_135455667.1-Vibrio-echinoideorum:0.0801990127,WP_141346804.1-Vibrio-inusitatus:0.1353265198)23.8/16:0.0114634555)85.1/35:0.0330167623,WP_218382835.1-Photobacterium-sp.-WH24:0.1522962023)85.4/88:0.0283145422,((((((WP_102455158.1-Vibrio-breoganii:0.1026023163,(WP_188011373.1-Vibrio-vulnificus:0.1564704495,WP_102451772.1-Vibrio-breoganii:0.0888561918)96.9/99:0.0698536817)69.6/75:0.0163622881,(WP_243025268.1-Vibrio-cincinnatiensis:0.3542324351,WP_070968050.1-Vibrio-sonorensis:0.1562525022)50.4/87:0.0106825448)77.6/75:0.0207729183,((WP_022590465.1-Vibrio-nigripulchritudo:0.1388322462,WP_045977433.1-Vibrio-neptunius:0.1295100193)89.3/99:0.0520589330,(WP_123015137.1-Vibrio-zhugei:0.0335935278,WP_077313846.1-Vibrio-palustris:0.0243863861)99.3/100:0.1011149336)90.6/95:0.0552635240)56.6/74:0.0293309023,WP_193290728.1-Vibrio-parahaemolyticus:0.1903697898)73.6/45:0.0130306437,((WP_261835819.1-Vibrio-ishigakensis:0.1073312717,WP_017221048.1-Moritella-dasanensis:0.0909608515)95.4/98:0.0604049568,WP_102329601.1-Vibrio-breoganii:0.1415069145)81/29:0.0168799127)74.7/46:0.0109840186,(WP_046218796.1-Photobacterium-halotolerans:0.1368602910,WP_102446429.1-Vibrio-breoganii:0.1177180383)88.3/56:0.0407586172)92.8/52:0.0407957332)87.6/53:0.0240283493)27.7/16:0.0086979856,WP_045069930.1-Photobacterium-leiognathi:0.1238171039)34.1/51:0.0239905731,(((WP_232600401.1-Photobacterium-carnosum:0.0884818480,(WP_060997345.1-Photobacterium-aquimaris:0.0999852139,WP_232615715.1-Photobacterium-carnosum:0.0757301237)42.5/88:0.0464033658)69.1/93:0.0343922325,WP_176284769.1-Photobacterium-damselae:0.1223870842)78.5/84:0.0383514947,((WP_249572008.1-Vibrio-cholerae:0.0675079470,(WP_255122387.1-Vibrio-cholerae:0.0759318393,WP_088132129.1-Vibrio-cholerae:0.0880568226)70.4/70:0.0372771721)91.6/68:0.0371875554,(((WP_000864153.1-Vibrio-cholerae:0.0488900963,WP_251878143.1-Grimontia-sp.-020920N:0.1084626438)84.7/100:0.0403571558,((WP_148560094.1-Vibrio-mimicus:0.1252414196,WP_103155729.1-Vibrio-vulnificus:0.0819166391)31/93:0.0272469749,WP_211678393.1-Moritella-sp.-36:0.1922651544)77.5/93:0.0213242880)68.5/88:0.0123084736,WP_045618469.1-Vibrio-vulnificus:0.0981542552)77.6/97:0.0258844113)52.2/68:0.0249246101)0/62:0.0164579424)89.5/77:0.0352270720,(((WP_190336189.1-Vibrio-sp.-S9_S30:0.0681980746,WP_022612811.1-Vibrio-nigripulchritudo:0.0930554544)68.3/95:0.0163443449,((WP_075989065.1-Vibrio-fluvialis:0.0328546317,WP_044366945.1-Vibrio-fluvialis:0.0254927884)88.4/100:0.0372942031,(WP_076657081.1-unclassified-Vibrio:0.0491384255,WP_239929047.1-Vibrio-kanaloae:0.0344435634)98.2/100:0.0752852517)99.4/100:0.1106962479)61.4/66:0.0246573190,(WP_269377025.1-Vibrio-sinaloensis:0.1382822789,WP_088905011.1-Shewanella-marisflavi:0.1498645841)57.7/64:0.0465134865)85.1/94:0.0275259180)75.9/69:0.0126800875)89.6/85:0.0245228639,(((WP_102457123.1-Vibrio-breoganii:0.0902001676,(WP_108083222.1-Vibrio-sp.-ZF-223:0.1105527282,(WP_102532836.1-Vibrio-sp.-10N.286.48.B7:0.0678716509,WP_254101081.1-Alteromonas-sp.-LMIT007:0.1082181510)83.3/100:0.0265525488)91.1/100:0.0346616708)80.3/97:0.0243091299,(((WP_046223474.1-Vibrio-sp.-ECSMB14106:0.0744116814,WP_102562373.1-Vibrio-breoganii:0.0910927788)91.5/72:0.0411203366,WP_102533425.1-Vibrio-breoganii:0.0864739002)25.5/71:0.0166335257,((WP_102481006.1-Vibrio-breoganii:0.0779660616,WP_258489575.1-Vibrio-alginolyticus:0.1115156079)65.5/88:0.0284857659,WP_017031451.1-Vibrio-breoganii:0.0877582507)94.1/86:0.0406284466)85.8/97:0.0228599012)89.4/92:0.0237392792,((((WP_065600047.1-Aliivibrio-sp.-1S128:0.0869762436,(WP_017063118.1-Vibrio:0.0766752302,WP_005395829.1-Vibrio-diabolicus:0.1568468003)41.1/97:0.0302049622)81.9/85:0.0289805449,(((WP_138589904.1-Pseudoalteromonas-aurantia:0.1083749235,(WP_218529053.1-Shewanella-sp.-NIFS-20-20:0.1035454857,WP_225483056.1-Vibrio-vulnificus:0.0587083549)93.9/100:0.0554615628)86.3/99:0.0281739380,WP_012549058.1-Aliivibrio-salmonicida:0.1218231882)89.9/99:0.0348847909,(WP_077647682.1-Salinivibrio-sp.-IB643:0.0743720008,WP_072616325.1-Vibrio-vulnificus:0.1475342248)90.8/98:0.0425552079)86.6/85:0.0292854466)87.9/51:0.0234674164,WP_203433235.1-Shewanella-sp.-LZH-2:0.1554218674)0/27:0.0000022153,((WP_017449116.1-Vibrio-parahaemolyticus:0.0916869833,(WP_172380549.1-Vibrio-sp.-Vb339:0.0952866947,(WP_102443327.1-Vibrio-breoganii:0.0339866907,WP_261923504.1-Shewanella-sp.-NFH-SH190041:0.1760160174)91.3/100:0.0413246651)95.8/100:0.0688381923)84.2/92:0.0280137504,((WP_165312521.1-Vibrio-ziniensis:0.0959500283,WP_152138288.1-Plesiomonas-shigelloides:0.1005269270)65.5/100:0.0213824463,WP_238819535.1-Vibrio-furnissii:0.0594008221)95.9/100:0.0528218042)85.1/84:0.0226056332)91.9/49:0.0279913454)88.7/52:0.0201161158)81/53:0.0179628780,((WP_017020702.1-Aliivibrio-logei:0.1126702647,WP_194091044.1-Vibrio-hibernica:0.1511662653)83.7/92:0.0499072387,(WP_206974914.1-Vibrio-parahaemolyticus:0.0877654515,WP_058428281.1-Pseudoalteromonas:0.1381348361)88.6/94:0.0518154936)78.5/70:0.0424750364)89.4/27:0.0228126436,(((((WP_268674123.1-Vibrio-scophthalmi:0.0894107104,WP_017085924.1-Vibrio-splendidus:0.0933345260)90.5/100:0.0405890452,(WP_102353859.1-Vibrio-cyclitrophicus:0.0787860519,WP_171315631.1-Vibrio-splendidus:0.1057199718)89/100:0.0458774641)3.1/32:0.0249442502,((WP_041156219.1-Vibrio-mytili:0.1288443845,(WP_047480300.1-Vibrio-campbellii:0.0439997638,WP_045494469.1-Vibrio-hyugaensis:0.0412150653)98.3/100:0.0816660864)96.5/100:0.0986445180,WP_102523418.1-Vibrio-tapetis:0.1121495801)85.4/94:0.0614882010)39/51:0.0290590147,(WP_059021773.1-Vibrio-coralliirubri:0.0656428427,(((((WP_012551417.1-Aliivibrio-salmonicida:0.0693962682,WP_012276595.1-Shewanella-halifaxensis:0.0498028026)37/100:0.0205586887,WP_135385147.1-Vibrio-tasmaniensis:0.1205917725)94.4/100:0.0615936140,WP_102571993.1-Vibrio:0.0583373088)70/94:0.0394982726,WP_265133188.1-Vibrio-splendidus:0.0766104893)66.1/82:0.0152955380,WP_102298256.1-Vibrio-cyclitrophicus:0.0677100294)69.9/51:0.0176357413)84.1/35:0.0169871152)79.9/59:0.0197279993,((((((WP_063656223.1-Aliivibrio-fischeri:0.0952829183,(WP_248638627.1-Vibrio-crassostreae:0.0973852358,WP_102313365.1-Vibrio-cyclitrophicus:0.1755390049)23.6/81:0.0126511250)61.1/68:0.0178013396,(WP_137008959.1-Vibrio-kanaloae:0.0912743010,WP_135353160.1-Aliivibrio-fischeri:0.0755232764)94.6/94:0.0611893830)78.3/66:0.0183059200,((WP_102459739.1-Vibrio-lentus:0.0742499143,WP_102363333.1-Vibrio-cyclitrophicus:0.0962824675)83.6/100:0.0292939989,WP_102554090.1-Vibrio-splendidus:0.1135931554)88.1/99:0.0367513473)85.9/20:0.0303621664,((((WP_232603643.1-Photobacterium-carnosum:0.0872065162,WP_193154481.1-Vibrio-navarrensis:0.1466644776)85.5/100:0.0448114272,WP_130314193.1-Vibrio-vulnificus:0.0688156339)83.8/100:0.0356428206,WP_047110243.1-Vibrio-vulnificus:0.2560128795)91.5/93:0.0429776006,WP_155675904.1-Aliivibrio-fischeri:0.0767665621)81.5/89:0.0178649396)80.1/26:0.0126516506,WP_017105169.1-Vibrio:0.1036004297)86.5/94:0.0209563679,WP_182796407.1-Vibrio-sp.-SG41-7:0.1404861472)86/71:0.0232230989)83.7/68:0.0174836056)92/19:0.0268542683,(((WP_171343915.1-Vibrio-cyclitrophicus:0.0677708614,(WP_161753058.1-Vibrio-toranzoniae:0.1008433244,(WP_102248986.1-Vibrio-tasmaniensis:0.1419993068,WP_135445135.1-Vibrio-tasmaniensis:0.0750620151)97.8/100:0.0589255057)86.7/99:0.0242765043)52.3/97:0.0251347159,WP_248368760.1-Vibrio-sp.-1CM8B:0.1102118159)85.7/74:0.0252142499,(((((WP_155682312.1-Aliivibrio-fischeri:0.0399932864,(WP_046209502.1-Vibrio:0.0972635275,WP_061039794.1-Vibrio-coralliirubri:0.1013544327)97.1/94:0.0729720008)92.6/69:0.0505429409,((((WP_108162474.1-Vibrio-splendidus:0.0267457226,WP_004742905.1-Vibrio-tubiashii:0.0474028436)99.4/100:0.0803854925,((WP_102503160.1-Vibrio-splendidus:0.1428756962,WP_017069285.1-Vibrio-crassostreae:0.0527387795)28.3/72:0.0209492450,WP_016768872.1-Vibrio-cyclitrophicus:0.0961453654)92/88:0.0373182575)81.1/86:0.0254534435,(WP_258428294.1-Vibrio-sp.-RM-69-4:0.0767717420,WP_102428030.1-Vibrio-sp.-10N.261.54.E10:0.1078927252)68.3/14:0.0279407449)66.5/8:0.0285561723,WP_012532661.1-Aliivibrio-fischeri:0.1186668815)13.8/17:0.0217442730)55.4/6:0.0217101888,((WP_130049361.1-Aliivibrio-finisterrensis:0.0468126486,WP_019826652.1-Vibrio-splendidus:0.1100165613)76.3/97:0.0214768901,WP_260256886.1-Vibrio-sp.-NC2:0.0918210290)81.1/94:0.0176369380)77.5/47:0.0215590152,((WP_193834777.1-Vibrio-sp.-OPT20:0.0267340288,(WP_029865567.1-Vibrio-parahaemolyticus:0.0505914317,WP_213967907.1-Vibrio-alginolyticus:0.0320035279)98.6/100:0.0871841840)92.1/100:0.0429082752,WP_004733311.1-Vibrio-splendidus:0.0675870361)92.7/88:0.0451553814)89.8/11:0.0301819679,(((WP_102420589.1-Vibrio-cyclitrophicus:0.0678044250,WP_101201417.1-Paraglaciecola-sp.-MB-3u-78:0.1768056045)79.9/95:0.0336724375,WP_132940416.1-Vibrio-crassostreae:0.0966030892)80.7/71:0.0322691052,WP_155677763.1-Aliivibrio-fischeri:0.1125103126)89.1/71:0.0311988299)82.6/56:0.0141040478)74.9/31:0.0156025776)89.5/5:0.0261197766,((WP_238130349.1-Vibrio-cincinnatiensis:0.1264148891,WP_102276040.1-Vibrio-lentus:0.0759997652)44.1/89:0.0207625188,((WP_139685731.1-Vibrio-tasmaniensis:0.0825240725,WP_248386338.1-Vibrio-sp.-ED004:0.1476904264)90/96:0.0495886771,WP_102457030.1-Vibrio-cyclitrophicus:0.0903379948)69.6/56:0.0299670382)88.2/73:0.0326487231)62.5/30:0.0197809097,WP_130066657.1-Aliivibrio-finisterrensis:0.1400414915)75.2/73:0.0172684082,(WP_048661380.1-Vibrio-crassostreae:0.0528253934,WP_193846925.1-Vibrio-sp.-OPT10:0.0718342120)91.1/99:0.0345741026)83/76:0.0149044092,((((((WP_041510674.1-Shewanella:0.0369778034,WP_187575439.1-Shewanella-algae:0.0497021956)95.9/99:0.0485011345,(WP_208158397.1-Shewanella-algae:0.0536019611,WP_208157589.1-Shewanella-algae:0.0397459416)72.9/99:0.0126818979)96/96:0.0370012338,(WP_161672404.1-Vibrio-toranzoniae:0.0351813902,WP_017040654.1-Vibrio-genomosp.-F10:0.1434689056)89/67:0.0349868255)46.4/55:0.0193526630,WP_137025093.1-Vibrio-kanaloae:0.1119334639)81.1/72:0.0161941403,(((WP_020328057.1-Vibrio-fluvialis:0.0864045627,WP_023404414.1-Vibrio-halioticoli:0.1695720006)0/73:0.0153820470,WP_145570128.1-Vibrio-tasmaniensis:0.1346689992)19.3/48:0.0217474448,WP_102475754.1-Shewanella-sp.-10N.286.52.C2:0.1595997607)83.3/48:0.0193222356)88.8/63:0.0358181724,(((WP_135455718.1-Vibrio-echinoideorum:0.0793284106,WP_048899071.1-Photobacterium-swingsii:0.1214283400)69.3/48:0.0134108757,WP_076646094.1-Vibrio-sp.-10N.222.47.A9:0.1176684275)48.2/78:0.0307948611,(WP_137027437.1-Vibrio-kanaloae:0.0274274242,WP_108117598.1-Vibrio-splendidus:0.0548038033)94/99:0.0456045164)21/52:0.0203922808)59.4/32:0.0130814656)73.2/54:0.0117097344)52.1/54:0.0181652771,((((((((WP_239941760.1-Vibrio-mediterranei:0.0849501928,((WP_062462511.1-Vibrio-mediterranei:0.0695477121,(WP_171392669.1-Vibrio:0.0678432779,WP_234495031.1-Vibrio-maritimus:0.1487714398)82.7/98:0.0398027330)43.5/93:0.0318859266,WP_252012765.1-Vibrio-sp.-SCSIO-43140:0.1062432461)73.4/92:0.0186916311)92.9/96:0.0365021955,((WP_221070344.1-Vibrio-alfacsensis:0.1036413727,WP_102532726.1-Vibrio-sp.-10N.286.48.B7:0.0724462183)53.3/97:0.0270535102,WP_250617847.1-Vibrio-sp.-S4B1:0.0748735771)86.1/92:0.0277445294)44.2/59:0.0083842932,WP_252012797.1-Vibrio-sp.-SCSIO-43140:0.0902883486)92.3/99:0.0437899109,WP_268679810.1-Vibrio-barjaei:0.0998412008)88.1/99:0.0315207412,WP_045898906.1-Enterobacteriaceae:0.1010152959)72.9/91:0.0240602857,(((((((WP_261600599.1-Photobacterium-sp.-TY1-4:0.0700952156,((WP_170899585.1-Vibrio-sp.-3-2-1:0.0890984615,WP_017221010.1-Moritella-dasanensis:0.0882150462)93.5/100:0.0403626700,WP_107284330.1-Photobacterium-lipolyticum:0.0736984613)93.2/98:0.0463830640)84.6/35:0.0218208019,(WP_159197271.1-Escherichia-coli:0.0985656214,(WP_114765998.1-Vibrio-rhodolitus:0.0852911261,WP_182108830.1-Vibrio-marinisediminis:0.0811489770)96/100:0.0563954968)78.5/31:0.0168650536)70.6/69:0.0282436189,((WP_211647400.1-Photobacterium-sp.-GJ3:0.0396334927,WP_120511428.1-Photobacterium-salinisoli:0.0383621590)69.1/100:0.0453121638,WP_172533652.1-Vibrio-aestuarianus:0.2340725331)88.1/92:0.0440338105)41.5/41:0.0125552313,WP_124938567.1-Vibrio-viridaestus:0.1171003769)23.8/13:0.0095967905,WP_088133934.1-Vibrio-gazogenes:0.1864291475)84.2/82:0.0153748243,((WP_112404745.1-Vibrio-diazotrophicus:0.0395268295,WP_202661182.1-Vibrio-fluvialis:0.0503987582)97.8/100:0.0582830295,(((WP_107294031.1-Photobacterium-phosphoreum:0.0412221094,WP_107276976.1-Photobacterium-sp.-NCIMB-13483:0.0453547799)97.6/100:0.0802240933,(WP_047463293.1-Vibrio-fluvialis:0.0874415250,WP_148560072.1-Vibrio-mimicus:0.1284874939)95.3/100:0.0513216270)82.4/67:0.0353915168,((WP_036979211.1-Providencia-alcalifaciens:0.1540714796,WP_101040106.1-Psychromonas-sp.-MB-3u-54:0.1301613143)89.9/96:0.0482642242,WP_138627230.1-Pseudoalteromonas-sp.-S4741:0.1786897522)77.2/45:0.0260157219)57.4/35:0.0227537349)86.4/32:0.0200973922)86.2/33:0.0159098488,(WP_020431327.1-Vibrio-fluvialis:0.0663856431,WP_055453311.1-Vibrio-fluvialis:0.0385733713)98.5/100:0.0623197958)91.4/32:0.0315097945)66.8/33:0.0047433596,(WP_188234612.1-Vibrio-sp.-Y2-5:0.0259180098,WP_057556016.1-Vibrio-cholerae:0.0429387767)100/100:0.1653799858)78.9/24:0.0090196904,(((((((((WP_065208126.1-Photobacterium-phosphoreum:0.1010148056,(WP_232585319.1-Photobacterium-iliopiscarium:0.0763445380,(WP_232582982.1-Photobacterium-phosphoreum:0.0418837424,((WP_256383785.1-Photobacterium-toruni:0.0404842562,WP_232578536.1-Photobacterium-carnosum:0.0458666929)38.9/87:0.0041834273,WP_107228404.1-Photobacterium-phosphoreum:0.0696884129)92.7/100:0.0295632372)90.4/100:0.0354187724)68.7/98:0.0265658593)63.5/90:0.0331766193,WP_232600381.1-Photobacterium-carnosum:0.0506018590)88.9/85:0.0284007254,((((((WP_232575020.1-Photobacterium-carnosum:0.1270276327,WP_086367281.1-Photobacterium-damselae:0.0303192762)84.6/99:0.0265215269,WP_151182432.1-Photobacterium-damselae:0.0974474168)34.9/8:0.0111830471,WP_053987023.1-Photobacterium-leiognathi:0.0650673096)69.8/30:0.0151837966,WP_115426223.1-Photobacterium-damselae:0.0551138226)51.8/4:0.0166159901,((((WP_238087945.1-Photobacterium-damselae:0.0539394829,(WP_065172005.1-Photobacterium-damselae:0.0633398445,WP_107185088.1-Photobacterium-leiognathi:0.0611727215)65.1/93:0.0166301764)87.1/74:0.0306461735,WP_151170663.1-Photobacterium-damselae:0.0222939508)80.4/58:0.0201892020,WP_045063675.1-Photobacterium-leiognathi:0.0650199859)80.6/59:0.0243249345,(WP_106261939.1-Photobacterium-damselae:0.0823997373,WP_237041371.1-Photobacterium-damselae:0.0552227847)66/3:0.0284797942)42.7/4:0.0154825748)85/66:0.0163801546,WP_237042700.1-Photobacterium-damselae:0.0775181392)88.7/63:0.0255051520)38.1/26:0.0169989405,WP_107253996.1-Photobacterium-indicum:0.1725385049)55.9/5:0.0081294147,(((WP_237041393.1-Photobacterium-damselae:0.1064548326,WP_080158029.1-Photobacterium-piscicola:0.1067086422)95.4/100:0.0631309606,((WP_065173166.1-Photobacterium-kishitanii:0.1079099187,WP_102395415.1-Enterovibrio-norvegicus:0.1096341341)49.9/86:0.0205005459,WP_047876260.1-Photobacterium-aphoticum:0.1115133445)54.9/98:0.0211907013)79.9/94:0.0209857601,((WP_045039948.1-Photobacterium-iliopiscarium:0.1254224160,((WP_065176337.1-Photobacterium-aquimaris:0.0560421913,WP_107199726.1-Photobacterium-phosphoreum:0.0655668625)78.6/100:0.0323429904,WP_023931546.1-Photobacterium-leiognathi:0.1020748239)80.6/90:0.0368510242)69.9/98:0.0504634493,WP_107213932.1-Photobacterium-phosphoreum:0.0872607976)92.1/39:0.0441765975)83.8/28:0.0292937345)74.3/5:0.0102430523,((((WP_161119826.1-Photobacterium-lucens:0.1256414176,WP_102786270.1-Vibrio-cholerae:0.0478949448)87.5/47:0.0230794137,(WP_000864154.1-Vibrio-cholerae:0.0479057437,(WP_094127815.1-Vibrio-sp.-V09_P4A23P171:0.0898040641,WP_199867560.1-Vibrio-anguillarum:0.0637400616)98.6/98:0.0906183100)83.1/25:0.0144082012)75.1/23:0.0126190135,(WP_175242188.1-Vibrio-cholerae:0.0545435350,(((WP_199351799.1-Vibrio-cholerae:0.0708170190,WP_149560528.1-Vibrio-cholerae:0.0652020538)76.4/95:0.0335878268,WP_095464062.1-Vibrio-cholerae:0.0598153134)83.3/85:0.0256851783,WP_031579865.1-Ruminobacter-sp.-RM87:0.3952671415)77.1/23:0.0110781470)92.8/28:0.0429635517)4.3/4:0.0092450395,(WP_267741605.1-Vibrio-cholerae:0.0932394651,WP_095478449.1-Vibrio:0.0581627378)91.1/88:0.0277046973)77.1/5:0.0131623569)85.5/5:0.0324148384,(((WP_208891742.1-Vibrio-anguillarum:0.1059594935,WP_148533341.1-Vibrio-cholerae:0.0479254091)82.2/100:0.0336737453,WP_107294203.1-Photobacterium-sp.-GB-50:0.1287488824)70.9/45:0.0343254033,(WP_000864156.1-Vibrio-cholerae:0.0415292946,WP_000864152.1-Vibrio-cholerae:0.0730526422)97.1/94:0.0555382466)53.5/32:0.0063878791)79.9/5:0.0152325281,((((WP_263100201.1-Kosakonia-sp.-ML.JS2a:0.0233516853,WP_200550470.1-Kosakonia-sp.-LAM2021:0.0776040093)98.4/100:0.0787233377,((WP_015854950.1-Musicola-paradisiaca:0.0373754873,WP_192457620.1-Musicola-keenii:0.0748136465)100/100:0.1812900068,(WP_094109047.1-Lonsdalea-iberica:0.0701480128,(WP_026742149.1-Lonsdalea-quercina:0.0337124650,WP_094117837.1-Lonsdalea-britannica:0.0266026569)99.4/100:0.0817076187)42.4/99:0.0182108779)89.8/81:0.0479391332)96.6/100:0.0668638952,(WP_038178862.1-Vibrio-rhizosphaerae:0.1114281491,WP_107187026.1-Photobacterium-kishitanii:0.1313191415)76.5/63:0.0090770762)28.5/20:0.0210817762,(WP_006646466.1-Photobacterium-sp.-SKA34:0.1422409810,(WP_232612305.1-Photobacterium-phosphoreum:0.2259262813,WP_136989485.1-Vibrio-sp.-F13:0.1794600836)95.8/99:0.0891652257)55.1/12:0.0100393753)93.5/80:0.0301351890)87/5:0.0296581455,(((((WP_069212356.1-Vibrio-cholerae:0.0485689820,WP_202641945.1-Vibrio-fluvialis:0.0889792485)93.7/100:0.0621442990,WP_129120576.1-Veronia-nyctiphanis:0.1495392738)84.3/56:0.0369047537,WP_045460498.1-Vibrio-hyugaensis:0.1224612146)41.9/48:0.0265772228,WP_068898547.1-Enterovibrio-pacificus:0.1014552722)85.4/54:0.0214617234,((WP_017004373.1-Enterovibrio-norvegicus:0.1331316481,(((WP_045569365.1-Vibrio:0.0579752497,WP_039437343.1-Vibrio-navarrensis:0.0405735407)98.1/100:0.0712505794,WP_176299354.1-Vibrio-sp.-05-20-BW147:0.1236557439)82.3/100:0.0420931674,WP_140233673.1-Shewanella-polaris:0.1776998157)67.2/93:0.0556615091)88.8/5:0.0425430549,((WP_182744595.1-Shewanella-sp.-SR44-4:0.1525709857,WP_152797327.1-Salinivibrio-sp.-VYel1:0.1455795814)88.8/100:0.0396186531,WP_039484180.1-Vibrio-sinaloensis:0.1175011468)82.3/85:0.0209553035)87.6/4:0.0245996069)92.2/5:0.0261669585)74.8/23:0.0121374912)90.5/71:0.0211672596)89.1/75:0.0247801275,(WP_102986644.1-Vibrio-diazotrophicus:0.0877679971,(((WP_239872404.1-Shewanella-insulae:0.1042818796,WP_238097025.1-Vibrio-cincinnatiensis:0.1126243266)52.8/72:0.0277249587,WP_118894008.1-Vibrio-vulnificus:0.1108367134)0/69:0.0000029573,WP_106651046.1-Shewanella-sp.-Pdp11:0.1430503082)84.5/93:0.0291818549)95.4/90:0.0421027115)89.6/88:0.0235138317,(((((WP_233983986.1-Vibrio-fluvialis:0.0531814313,WP_225486226.1-Vibrio-vulnificus:0.1306873739)72.9/27:0.0103399867,WP_143680610.1-Vibrio-furnissii:0.1102917338)88.5/26:0.0242954316,(((((WP_238813939.1-Vibrio-vulnificus:0.0904780979,(WP_217542775.1-Vibrio-metschnikovii:0.0445955314,WP_113593871.1-Vibrio:0.0704758342)97.2/100:0.0594403280)42.7/24:0.0225622229,((WP_068331709.1-Photobacterium-jeanii:0.1193017565,(WP_031414678.1-Vibrio-parahaemolyticus:0.0982669820,WP_248468589.1-Shewanella-sp.-1CM18E:0.1127829861)88.6/100:0.0445039429)67/45:0.0374879166,WP_077679840.1-Vibrio-kanaloae:0.0633296651)75.3/96:0.0334967299)68.9/29:0.0235938457,(WP_086585081.1-Vibrio-parahaemolyticus:0.0256195140,WP_217557493.1-Vibrio-metschnikovii:0.0539203957)93.7/100:0.0452358582)65.9/26:0.0166185023,WP_172561824.1-Vibrio-furnissii:0.0915810796)91.7/27:0.0298639811,(((((WP_155682326.1-Aliivibrio-fischeri:0.0702344485,WP_135445143.1-Vibrio-tasmaniensis:0.1407018708)72.6/26:0.0106213686,(WP_102385704.1-Vibrio-cyclitrophicus:0.0568407819,WP_239937663.1-Vibrio-chagasii:0.1433649400)93.2/91:0.0416331740)90.1/26:0.0318124218,(WP_005597034.1-Vibrio-scophthalmi:0.1194490351,((((WP_016784063.1-Vibrio:0.0895466761,WP_140342839.1-Vibrio-parahaemolyticus:0.1277685350)0/55:0.0036106444,WP_102394203.1-Vibrio-sp.-10N.222.54.B6:0.0977806662)86.2/20:0.0380524057,(WP_108176621.1-Vibrio-splendidus:0.0452196982,WP_239918548.1-Vibrio-sp.-Isolate22:0.0231761422)98.3/100:0.0524131223)77.1/7:0.0197670871,(WP_135380937.1-Vibrio-tasmaniensis:0.1026023740,WP_248388778.1-Vibrio-cyclitrophicus:0.0989168433)56.3/26:0.0218640987)84/7:0.0220347154)89.1/26:0.0369657520)62.2/8:0.0100501276,WP_102482947.1-Vibrio-splendidus:0.0792716633)74.6/96:0.0256446303,(((WP_162023628.1-Shewanella-sp.-Arc9-LZ:0.0596549765,WP_248997855.1-Shewanella-marinintestina:0.0421286172)97.6/100:0.0623362112,(WP_065296243.1-Vibrio-natriegens:0.0948793112,(WP_009843931.1-Vibrio-sp.-AND4:0.0676443898,WP_017094106.1-Vibrio-splendidus:0.0639598438)50.7/95:0.0098102468)89.2/91:0.0325191400)77.7/86:0.0281229378,(WP_102285079.1-Vibrio-lentus:0.1058823374,((WP_009847952.1-Vibrio:0.1353035666,WP_233894267.1-Vibrio-kanaloae:0.0991303661)78.6/100:0.0535050446,(WP_048661352.1-Vibrio-crassostreae:0.0707093852,WP_114634227.1-Vibrio-splendidus:0.0461606370)91.9/100:0.0630143378)92.7/99:0.0481845111)73.1/78:0.0148266132)88.4/97:0.0334036135)96.2/86:0.0401853051)76/26:0.0063677885)82.1/54:0.0136318823,(((((WP_102563398.1-Vibrio-sp.-10N.286.49.B3:0.0752873487,(WP_064489977.1-Vibrio-parahaemolyticus:0.0506965506,WP_201976964.1-Aeromonas-enteropelogenes:0.0796403869)93.6/100:0.0608040002)87.9/77:0.0426321386,(WP_192928291.1-Vibrio-vulnificus:0.0326396913,WP_132698543.1-Vibrio-harveyi-group:0.0904880501)98.4/100:0.0807048516)21.3/34:0.0229770618,(WP_211915530.1-Vibrio-sp.-B1REV9:0.0743381981,(WP_017420706.1-Vibrio-vulnificus:0.0740007829,WP_017024716.1-Vibrio-rumoiensis:0.1522239831)74.1/36:0.0123273605)80.3/36:0.0190634461)81.4/58:0.0280739912,((((WP_039841338.1-Vibrio-harveyi-group:0.0952973634,WP_193167845.1-Vibrio-navarrensis:0.2114443350)84.5/99:0.0394819112,WP_213972370.1-Vibrio-alginolyticus:0.1412649193)76.5/94:0.0386132327,WP_120030433.1-Vibrio-sinensis:0.0937828881)76.6/5:0.0143437661,((WP_268809473.1-Vibrio-parahaemolyticus:0.1184010825,(WP_193250619.1-Vibrio-parahaemolyticus:0.1283612603,(WP_028024862.1-Enterovibrio-calviensis:0.1843664345,WP_017007824.1-Enterovibrio-norvegicus:0.1007091961)80.8/100:0.0269103146)79.5/51:0.0319952631)76.5/28:0.0143243512,(WP_099031861.1-Pseudoalteromonas-sp.-GCY:0.1145042261,((((WP_033540599.1-Shewanella:0.1204372659,WP_031815865.1-Vibrio-parahaemolyticus:0.1031245327)25.2/66:0.0270989133,WP_258516089.1-Vibrio-alginolyticus:0.0850708176)76/83:0.0196187239,((WP_103194250.1-Vibrio-vulnificus:0.0384590788,WP_158146950.1-Vibrio-fluvialis:0.0900315632)97.1/100:0.0590791315,WP_233927909.1-Vibrio-fluvialis:0.1020138989)95.1/100:0.0530951649)88.1/81:0.0320786713,WP_224657569.1-Vibrio-vulnificus:0.1263073766)31.6/45:0.0092725103)76.5/35:0.0222911576)95.6/89:0.0462185505)81.3/5:0.0166036726)85.9/5:0.0173316630,((WP_069533349.1-Vibrio-parahaemolyticus:0.0911912520,WP_235060641.1-Vibrio-sp.-SS-MA-C1-2:0.1787155844)29.3/6:0.0298400026,WP_229630939.1-Vibrio-coralliilyticus:0.1835311190)77.3/6:0.0165567778)87/30:0.0261487394)84.6/50:0.0154904254,(WP_208844179.1-Pseudoalteromonas-xiamenensis:0.1151653588,WP_233976434.1-Vibrio:0.1226686379)86.8/98:0.0382992166)78.7/70:0.0121665515)92.4/76:0.0367237087,((WP_193255073.1-Vibrio-navarrensis:0.0213414288,WP_258469755.1-Vibrio-alginolyticus:0.0972883449)98.7/100:0.0774960683,(WP_220051788.1-Shewanella-sp.-FJAT-51860:0.0973120550,WP_029788868.1-Vibrio:0.1600802154)32.9/97:0.0285071535)65.7/95:0.0236132755)87.5/100:0.0292195858,(WP_013365656.1-_Enterobacter:0.1467549538,(((WP_258988101.1-Scandinavium-hiltneri:0.0621897380,WP_181620050.1-Enterobacter-sp.-RHBSTW-00994:0.0512712139)95.7/100:0.0588335461,WP_193129600.1-Klebsiella-aerogenes:0.0617361942)93.7/100:0.0443115188,WP_214579582.1-Enterobacter-asburiae:0.0769274099)87.7/100:0.0459008071)100/100:0.1335154866)7.3/62:0.0228309576,WP_161680902.1-Proteus:0.2244644448)44.3/19:0.0204368095,((((((WP_261298106.1-Shewanella-sp.-SP1S2-7:0.0726395509,WP_167678467.1-Shewanella-aestuarii:0.0842959192)100/100:0.1762362467,WP_248994245.1-Shewanella-gaetbuli:0.1603129329)89.7/100:0.0767169482,WP_212599444.1-Shewanella-sp.-M16:0.1768275967)85.1/87:0.0880527279,WP_222242368.1-Pseudoalteromonas-piscicida:0.2072674262)0/36:0.0028134309,((WP_073514120.1-Vibrio-parahaemolyticus:0.0573704437,WP_032481483.1-Vibrio:0.0134209487)100/100:0.4832252971,(WP_205000141.1-Parashewanella-hymeniacidonis:0.0534068616,WP_129946484.1-Shewanella-sp.-OPT22:0.0414024090)100/100:0.3409673079)44.5/43:0.0929473445)94.5/39:0.0681951973,(((((WP_069589493.1-Salinivibrio-sp.-DV:0.0839718029,WP_077662143.1-Salinivibrio-proteolyticus:0.0119297640)84/99:0.0270409642,((WP_077659434.1-Salinivibrio-kushneri:0.1019026508,WP_077659452.1-Salinivibrio-kushneri:0.0262328477)84.7/100:0.0226616292,WP_269578450.1-Salinivibrio-kushneri:0.0676445588)80.9/100:0.0351890970)100/100:0.2061628855,WP_138522257.1-Pseudoalteromonas-galatheae:0.1754744927)83/58:0.0300972594,(WP_044365158.1-Vibrio-fluvialis:0.1889539723,WP_025523961.1-Vibrio-parahaemolyticus:0.1236800582)27.7/65:0.0390388436)79.7/57:0.0335368335,(WP_213886549.1-Vibrio-alginolyticus:0.1336044847,WP_158173598.1-Vibrio-alginolyticus:0.0846715162)82.3/94:0.0217419267)91.5/54:0.0378741303)35.1/37:0.0333576029)84.4/3:0.0270786074,((((WP_076542412.1-Shewanella-sp.-UCD-KL21:0.2099958887,WP_007466453.1-Photobacterium-marinum:0.1659564112)15.5/91:0.0391438442,WP_120356561.1-Alginatibacterium-sediminis:0.2148208414)14.1/30:0.0366589639,WP_237309237.1-Vibrio:0.1982085008)78.2/90:0.0310146115,(WP_133040538.1-Shewanella-fodinae:0.1442833343,WP_232581009.1-Photobacterium-phosphoreum:0.4586076307)83.7/70:0.0430494991)96.8/72:0.0734945946)88.8/92:0.0399998291,(((WP_011189464.1-Desulfotalea-psychrophila:0.2879381199,WP_050928752.1-Yersinia-enterocolitica:0.1674420068)0/21:0.0293914022,WP_012883389.1-Dickeya-parazeae:0.1727261012)91.3/88:0.0551069066,((WP_035339894.1-Dickeya:0.0193245923,WP_125258312.1-Dickeya-lacustris:0.0714778849)100/100:0.1473765377,WP_042868619.1-Dickeya:0.0715794737)95.6/100:0.0586801992)90.7/94:0.0445039565)99/100:0.0949127449,((WP_162556830.1-Proteus:0.0732807012,WP_161779122.1-Proteus-sp.-G2675:0.0697430577)99.8/100:0.1435810243,WP_236366479.1-Aeromonas-veronii:0.1545075020)39.3/76:0.0136819533)99.2/96:0.1152566631)79.8/6:0.0519821849,WP_225633057.1-Aeromonas-enteropelogenes:0.1664075097)66.5/5:0.0127770695,WP_061279316.1-Kluyvera-cryocrescens:0.2578103118)90/5:0.0520007442,((WP_071788415.1-Franconibacter-pulveris:0.2949429810,(WP_020232007.1-Escherichia-coli:0.1678052275,(WP_228338545.1-Enterobacter-hormaechei:0.1140624172,(WP_109845273.1-Enterobacter:0.0412181566,WP_150146763.1-Enterobacter-roggenkampii:0.0378489134)84.6/100:0.0678285221)99.6/100:0.1548127177)78.8/97:0.0647022088)92.4/96:0.0923674223,(WP_189493202.1-Formosimonas-limnophila:0.1556100955,WP_133620003.1-Hydromonas-duriensis:0.1963260357)99.6/100:0.1562731221)58.8/5:0.0417297506)93.5/22:0.0398368515,(((((((((((((WP_151711904.1-Enterobacter-ludwigii:0.0263781735,(WP_032941356.1-Citrobacter-freundii:0.1634839278,WP_058678686.1-Enterobacter:0.0652141321)92.6/98:0.0532734530)51.2/98:0.0393362320,WP_223560970.1-unclassified-Enterobacter:0.0263501766)99.7/98:0.1879448726,WP_044720521.1-Escherichia-coli:0.2333297140)96.3/100:0.1422877913,WP_230863267.1-Enterobacter-cloacae:0.2682046093)85/95:0.1162328763,(WP_029399168.1-Escherichia-coli:0.1993652662,((WP_034496584.1-Buttiauxella-agrestis:0.2560125396,WP_164716793.1-Cedecea-lapagei:0.2364579286)70.5/100:0.0699001650,WP_050862535.1-Enterobacter:0.2622380990)33.2/96:0.0215240884)92.4/100:0.0618503981)85.3/79:0.0590411479,(WP_036421240.1-Morganella-morganii:0.1137213417,(WP_264998074.1-Morganella-morganii:0.0638933816,WP_186849509.1-Morganella-morganii:0.0689873542)92/100:0.0670438705)94.7/100:0.0604290312)88/79:0.0461019064,WP_064374050.1-Pantoea:0.2438720327)29.5/75:0.0273760115,((WP_039304623.1-Cedecea-neteri:0.2474672849,WP_159282060.1-Rahnella-variigena:0.2386903178)84.7/83:0.0482744752,(WP_106902182.1-Citrobacter-freundii:0.1592923672,(WP_126291239.1-Enterobacter-hormaechei:0.1179676942,((((WP_062932819.1-Enterobacter-hormaechei:0.0266702973,WP_252042312.1-Enterobacter-chengduensis:0.0366304393)89.2/100:0.0230403368,WP_048999124.1-Enterobacter:0.0328841861)73.2/64:0.0072974901,WP_063144508.1-Enterobacter-cloacae:0.0305200991)85.4/64:0.0210695122,WP_049008289.1-Enterobacter-cloacae:0.0371683037)83.3/96:0.0456544469)97.1/100:0.0812554102)90.7/93:0.0543344649)92.5/86:0.0467931048)82.8/97:0.0328768980,((((WP_234505032.1-Kluyvera-intermedia:0.2785271113,(WP_257124601.1-Aeromonas-veronii:0.2457314292,WP_211007190.1-Pectobacterium-brasiliense:0.2691016648)58/82:0.0265600860)90.3/81:0.0498599450,((((((WP_159241675.1-Proteus-terrae:0.1384398829,WP_195697765.1-Providencia-rettgeri:0.1352891774)94.2/86:0.0638148139,WP_238563424.1-Providencia-rettgeri:0.0872302363)54/68:0.0280838788,WP_165880752.1-Providencia-rettgeri:0.1320325857)86.3/97:0.0310315935,WP_231068620.1-Providencia-huaxiensis:0.1157123661)86.2/81:0.0204330994,(WP_154625743.1-Providencia-stuartii:0.1009778412,WP_004924376.1-Providencia-stuartii:0.1135274741)94.3/81:0.0499997055)80.9/28:0.0245086443,((WP_169342312.1-Providencia-stuartii:0.1433795901,WP_164565644.1-Providencia-rettgeri:0.1549099574)86.7/47:0.0301288696,(WP_187128479.1-Providencia-sp.-JUb39:0.1272545479,WP_193115284.1-Escherichia-coli:0.0923956059)91.9/99:0.0582432705)31.7/30:0.0288882365)96.9/100:0.0596701800)95/86:0.0518231316,(((WP_038258850.1-Xenorhabdus-bovienii:0.1985655370,WP_086113286.1-Xenorhabdus-beddingii:0.1429945527)76.4/73:0.0460175885,WP_038223662.1-Xenorhabdus-bovienii:0.1598847181)27/60:0.0180976942,WP_206164314.1-Citrobacter:0.2012510558)87.4/78:0.0395557580)29.5/76:0.0188018784,((WP_161748440.1-Proteus-sp.-G2658:0.1606793964,WP_049199273.1-Proteus-mirabilis:0.0446916076)86.7/98:0.0437097413,WP_157954980.1-Proteus-terrae:0.0530896850)99.9/100:0.1359591164)72.8/96:0.0208263605)88.9/90:0.0335075192,((WP_167816599.1-Enterobacter-cloacae:0.2314603684,(WP_262183194.1-Enterobacter-mori:0.1270651269,(WP_063412371.1-Enterobacter-cloacae:0.0404299023,WP_047027451.1-Enterobacter-kobei:0.0394627106)96.8/100:0.0719420221)96.2/100:0.0866729894)84.7/100:0.0501756633,((WP_105274811.1-Escherichia:0.1241396763,WP_172410167.1-Escherichia-coli:0.2270020170)50/55:0.0404481280,WP_064512195.1-Buttiauxella-gaviniae:0.1949548355)93.1/99:0.0604630091)95.2/86:0.0506822023)79.3/38:0.0322945760,(((((((WP_090123257.1-Kosakonia-arachidis:0.0713436433,WP_020454293.1-Kosakonia-sp.-SOY2:0.0807611821)90.5/100:0.0430709829,WP_017458471.1-Kosakonia-sacchari:0.1202672783)53.2/100:0.0313051673,(WP_200002836.1-Citrobacter-freundii:0.1335347640,WP_080329661.1-Enterobacter-cloacae-complex:0.1066339365)99/100:0.1350690334)47.4/99:0.0631403214,WP_090460354.1-Enterobacter-sp.-kpr-6:0.0822191818)98.8/100:0.1053744679,WP_140919905.1-Insectihabitans-xujianqingii:0.1517396664)47.1/97:0.0576643555,WP_070133611.1-Candidatus-Erwinia-dacicola:0.1636910865)74.4/6:0.0363427247,(((WP_194857598.1-Citrobacter-freundii:0.2086666419,(((WP_141032926.1-Escherichia-coli:0.1146253475,WP_196390204.1-Enterobacter-mori:0.0377101122)96.4/100:0.0513799034,WP_111966211.1-Enterobacter-cloacae-complex:0.0263940729)24.8/95:0.0166380344,WP_252032207.1-Enterobacter-asburiae:0.0271268960)99.3/99:0.1234533589)98.8/100:0.1259535741,(WP_168756429.1-Aeromonas-hydrophila:0.2662389923,WP_227390214.1-Halomonas-malpeensis:0.3849823290)89.8/92:0.0712307766)16.7/80:0.0435688362,WP_269535377.1-Citrobacter-sedlakii:0.2079754511)70.6/7:0.0337522505)79.2/4:0.0177662090)89/18:0.0355443922,((((((WP_108569539.1-unclassified-Pantoea:0.1538746935,WP_028018520.1-Enterobacter-cloacae-complex:0.3826612989)93.9/100:0.1183839597,WP_250341216.1-Pantoea-eucrina:0.1142710294)95/99:0.1106065024,(WP_213789144.1-Enterobacter-hormaechei:0.0067730774,((WP_148242178.1-Enterobacter-asburiae:0.0453903039,WP_237994110.1-Enterobacter-sp.-DRP3:0.0170302437)76.1/100:0.0049186887,WP_237424876.1-Enterobacter-cloacae-complex-sp.-ECC445:0.0405553114)90.3/100:0.0397895627)100/100:0.1876468212)88.1/12:0.0480209185,((((WP_137650031.1-Escherichia-albertii:0.0766684617,WP_001100804.1-Enterobacteriaceae:0.1196930737)78.9/98:0.0414919812,(WP_248249449.1-Enterobacter-roggenkampii:0.1159779416,WP_032936727.1-Citrobacter-freundii:0.0918849321)99.6/100:0.1332374344)63.7/96:0.0233364601,WP_001100786.1-Escherichia-coli:0.0773954225)95.4/96:0.0543678405,(((WP_107192407.1-Escherichia-albertii:0.0128546404,WP_181673375.1-Escherichia-fergusonii:0.0402062625)99.1/100:0.0802784679,(((WP_032214194.1-Escherichia-coli:0.0413443006,WP_059217428.1-Escherichia-albertii:0.0370984333)60.6/53:0.0058105134,WP_085459683.1-Escherichia-coli:0.0760989869)40.4/62:0.0294402498,(WP_073278747.1-Escherichia-coli:0.0671899933,WP_087893724.1-Enterobacteriaceae:0.0316511791)99.4/100:0.0941744949)98.5/100:0.1081332674)82.8/78:0.0298021273,((((WP_179154141.1-Citrobacter-freundii:0.1984753652,WP_086539147.1-Citrobacter-freundii:0.1754220857)95.1/86:0.0769259285,WP_033870520.1-Enterobacteriaceae:0.1410883403)76.7/43:0.0163127531,WP_267387874.1-Citrobacter-portucalensis:0.2117317822)76.7/39:0.0266363040,((WP_001610415.1-Escherichia-coli:0.0323297934,WP_138183825.1-Escherichia-sp.-E4385:0.0210039897)99.7/100:0.1092487818,(WP_089585421.1-Escherichia-coli:0.0347365968,WP_104705897.1-Escherichia-coli:0.0294857290)99.8/100:0.1385364690)94.2/94:0.0624394100)80.4/42:0.0333392801)90.4/63:0.0371559437)59.9/25:0.0175332701)88.3/12:0.0253948264,((((WP_157959988.1-Citrobacter:0.2869551378,WP_073446180.1-Escherichia-coli:0.1099159013)91.8/100:0.0596953964,((((WP_095844187.1-Escherichia-coli:0.0599431941,WP_086118650.1-Escherichia-coli:0.0232773312)60.8/99:0.0045773945,WP_089538038.1-Escherichia-coli:0.0242919510)100/100:0.1357238155,WP_153879942.1-Citrobacter-freundii:0.1176536766)0/36:0.0097759465,WP_001297344.1-Escherichia-coli:0.1272394987)87.6/89:0.0421674833)85.6/86:0.0279252707,((WP_016236761.1-Enterobacteriaceae:0.0756715537,WP_024239504.1-Escherichia-coli:0.0927218904)95.7/100:0.0715588677,WP_138098269.1-Jejubacter-calystegiae:0.1183277071)79.2/86:0.0390739266)93.6/94:0.0429563432,(((((WP_100272167.1-Citrobacter-portucalensis:0.1373557357,((WP_058662478.1-Enterobacteriaceae:0.0365853135,WP_061709645.1-Pseudenterobacter-timonensis:0.0372157674)99.5/100:0.1193823881,WP_112164393.1-Rahnella-inusitata:0.1311153937)94.7/97:0.0712880237)75.5/30:0.0365048599,(WP_181624464.1-Citrobacter-freundii:0.1596207967,WP_001515521.1-Escherichia-coli:0.1381630075)90.5/99:0.0501076814)86.1/18:0.0270129073,(((WP_171269898.1-Aeromonas-media:0.0657776587,WP_216983019.1-Aeromonas-sp.-FDAARGOS-1415:0.0832745793)99.6/100:0.1253155230,(WP_120162698.1-Rahnella-variigena:0.0421519181,WP_183847446.1-Rahnella-inusitata:0.0514279767)100/100:0.1935171319)92.6/100:0.0515805321,WP_047609274.1-Rahnella-aquatilis:0.1000063583)93.4/100:0.0583658056)78.6/20:0.0151132186,WP_157953065.1-Limnobaculum-parvum:0.1658923816)86.2/94:0.0422871046,(((((WP_170897747.1-Escherichia-coli:0.1144048456,WP_001100781.1-Escherichia-coli:0.1424206623)86.4/100:0.0436220773,WP_001542972.1-Escherichia-coli:0.0890411132)91.6/98:0.0401616442,(WP_220393175.1-Citrobacter-freundii:0.2087427493,WP_219093375.1-Escherichia-coli:0.1117272196)78.1/92:0.0413873861)64.7/80:0.0236482645,(WP_174633339.1-Escherichia-coli:0.0803352186,WP_072664248.1-Escherichia:0.0712764400)94.7/100:0.0561282454)36.9/78:0.0183867013,(WP_208464084.1-Escherichia-coli:0.0963797502,((WP_262931739.1-Escherichia-albertii:0.0292489028,WP_160188772.1-Escherichia-coli:0.0386732530)98.1/100:0.0686515741,(WP_021540003.1-Escherichia-coli:0.0137200725,WP_047657564.1-Enterobacteriaceae:0.0388336519)97.8/100:0.0633825887)71.5/76:0.0266059804)92.3/88:0.0340801281)73.5/81:0.0194615555)94.2/89:0.0408871293)76/68:0.0088970434)82.5/8:0.0134075357,(((((((WP_202667543.1-Lelliottia-amnigena:0.1491238241,(((WP_103946175.1-Lelliottia-aquatilis:0.0956645363,WP_123427447.1-Lelliottia:0.0421275447)73.2/86:0.0474565339,(WP_235147774.1-Enterobacter-cloacae:0.1158173618,WP_061356323.1-Escherichia-coli:0.4333310199)91.9/90:0.0801893773)87.5/90:0.0567897765,WP_075180894.1-Pantoea-sp.-1.19:0.1576813222)37.8/43:0.0205574500)73.1/40:0.0258318706,WP_021577651.1-Escherichia-coli:0.0885996976)81.7/39:0.0120127467,WP_059222105.1-Escherichia-albertii:0.0980379905)83.7/38:0.0118419506,((((((((WP_127791218.1-Escherichia-coli:0.0891380333,WP_025210617.1-Escherichia-coli:0.0804512799)42.4/87:0.0174866301,WP_001527244.1-Escherichia-coli:0.0674392026)86.5/84:0.0337855787,WP_187209312.1-Escherichia-coli:0.0733253190)80.1/10:0.0241512686,(((WP_096972986.1-Escherichia-coli:0.0753295565,(WP_157704420.1-Escherichia-coli:0.0615699726,WP_042200020.1-Enterobacteriaceae:0.0592256238)71.1/72:0.0109635088)87.7/89:0.0201000747,(WP_137653986.1-Escherichia-albertii:0.0257150260,WP_001100785.1-Escherichia-coli:0.0301717565)99/100:0.0607490786)87.8/49:0.0241422004,(((WP_085454021.1-Escherichia-coli:0.0957804401,WP_105963664.1-Escherichia-coli:0.0499586670)49.3/37:0.0224002933,WP_053290779.1-Escherichia-coli:0.0901693528)59.8/88:0.0195012778,((WP_146314246.1-Escherichia-coli:0.0311099458,WP_096967326.1-Escherichia-coli:0.0214493583)98.6/100:0.0659840495,WP_181465273.1-Escherichia-fergusonii:0.0508795240)89.1/49:0.0283092997)41.7/13:0.0146223456)81.6/17:0.0136044242)69.5/4:0.0069015977,(((WP_063610377.1-Escherichia-coli:0.0564164724,WP_001100798.1-Enterobacteriaceae:0.0850243916)79.4/83:0.0157132126,(WP_088541377.1-Escherichia-coli:0.0240905296,WP_222691870.1-Escherichia-marmotae:0.0488048853)95.9/100:0.0452407077)70.8/80:0.0159741819,WP_218070502.1-Escherichia-coli:0.0739758853)87.6/7:0.0327344311)95.4/46:0.0337730745,(((((((WP_193360946.1-Enterobacter-cloacae-complex-sp.-P39RS:0.0466877140,WP_182945676.1-Enterobacter-cloacae:0.0157041854)99.6/100:0.0777644717,((WP_032636062.1-Enterobacter-kobei:0.0479642114,WP_138014015.1-Enterobacter-ludwigii:0.0510148675)59.6/100:0.0307525007,(WP_063137673.1-Enterobacter-cloacae-complex:0.0338944315,WP_165464597.1-Enterobacter-cloacae:0.0417531884)85.4/99:0.0322299579)84.9/98:0.0270537709)77.4/22:0.0203271729,((WP_232904305.1-Buttiauxella-sp.-W03-F01:0.1385541400,WP_259388919.1-Enterobacter-huaxiensis:0.0322935536)56.4/98:0.0279491560,WP_230350752.1-Lelliottia-sp.-WAP21:0.1077309815)47.7/99:0.0250025313)15/8:0.0121231430,WP_265185975.1-Enterobacter-mori:0.0535917572)94/96:0.0407775644,WP_259000881.1-Scandinavium-goeteborgense:0.1130666083)79.3/58:0.0223355383,((((((((((WP_096205048.1-Enterobacter:0.0273912915,WP_100165931.1-Enterobacter-mori:0.0257002582)97/100:0.0575807863,WP_242172213.1-Enterobacter:0.0584824535)29.9/99:0.0325136924,WP_239609625.1-Lelliottia-amnigena:0.0771216674)95.6/96:0.0426589761,WP_033487312.1-Enterobacter-cloacae-complex:0.0359582651)83.7/74:0.0264720570,WP_126315015.1-Enterobacter-hormaechei:0.0834259718)45/32:0.0182313316,WP_063158410.1-Enterobacter-hormaechei:0.0694365260)57.7/3:0.0033942824,(((WP_042716137.1-Enterobacter-sp.-B509:0.0163362074,WP_241820523.1-Enterobacter-kobei:0.0551074515)96.4/100:0.0535707620,(((WP_214593796.1-Enterobacter-mori:0.0477886700,WP_086532312.1-Enterobacter-ludwigii:0.0533348936)98.2/100:0.0841084734,WP_045339085.1-Enterobacter-cloacae-complex:0.1053280057)52.7/72:0.0333191495,((WP_047742849.1-Enterobacter:0.0644366385,WP_040022520.1-Enterobacter-cloacae:0.0565687967)98.9/100:0.0793050169,(WP_129995405.1-Enterobacter-cloacae:0.0527731519,WP_042099316.1-Escherichia-coli:0.0585292038)85.2/100:0.0268667703)57.2/61:0.0386623100)73.5/99:0.0537606347)88.5/34:0.0289052892,(WP_045338355.1-Enterobacter-hormaechei:0.0848721763,(WP_182115563.1-Enterobacter-kobei:0.0260198102,WP_213381159.1-Enterobacter-cloacae:0.0553536568)98.3/100:0.0577794706)86.3/68:0.0198153631)87.5/30:0.0314230778)84.8/6:0.0230422387,WP_063869545.1-Enterobacter-hormaechei:0.0516758827)84.9/12:0.0222164240,((((WP_034915863.1-Erwinia-sp.-9145:0.1197389657,WP_191149682.1-Erwinia-aphidicola:0.1258795178)39.2/56:0.0198262198,WP_126546226.1-Enterobacter-quasimori:0.0704275172)86.5/52:0.0217080114,((((WP_141109706.1-Escherichia-albertii:0.0756281484,(WP_203360733.1-Citrobacter-sp.-R56:0.0536214968,WP_001572827.1-Escherichia-coli:0.0513964349)0/64:0.0146741916)94.4/100:0.0456668232,WP_258993080.1-Scandinavium-hiltneri:0.0574460244)94.2/100:0.0435326044,WP_208283396.1-Enterobacter-sichuanensis:0.0690623575)66.5/33:0.0162246223,WP_172688853.1-Enterobacterales:0.1169420941)52/6:0.0077462723)89.6/60:0.0244930069,(WP_046888360.1-Enterobacter-cloacae:0.0514832294,(((WP_095374511.1-Escherichia-coli:0.0094037262,WP_012906377.1-Citrobacter-rodentium:0.0764219718)98.9/100:0.0693114377,WP_096122580.1-Escherichia-coli:0.0571081902)47.4/98:0.0183753963,WP_162379936.1-Citrobacter-sp.-NCU1:0.1371354623)92.3/100:0.0392642355)89.9/67:0.0240184210)85.1/23:0.0187851319)94.5/70:0.0316990312,(((((WP_181680936.1-Citrobacter-sp.-RHB25-C09:0.0911512767,(WP_090049757.1-Citrobacter-amalonaticus:0.0237672731,WP_196013128.1-Citrobacter-farmeri:0.0286335228)99.7/100:0.0952435348)80/99:0.0381227578,WP_103775088.1-Citrobacter-amalonaticus:0.1335218194)99/100:0.1025428525,WP_121265736.1-Enterobacter-sp.-R1-2018:0.1623662948)87.9/98:0.0457237571,WP_259047621.1-Klebsiella-sp.-BIGb0407:0.1198622295)0/55:0.0116940631,WP_249031739.1-Tenebrionicola-larvae:0.1973185402)90.7/97:0.0345702728)85.6/48:0.0150324939)85.1/23:0.0126799155,((((WP_162648791.1-Citrobacter-freundii:0.0848769437,(WP_060683585.1-Citrobacter-portucalensis:0.1048378389,(WP_079798993.1-Salmonella-enterica:0.0790840691,WP_032255798.1-Enterobacteriaceae:0.0671065101)39.7/96:0.0070125521)95.6/96:0.0520650733)66/99:0.0320993758,(WP_195994783.1-Citrobacter-portucalensis:0.1438381102,WP_048218074.1-Citrobacter-freundii-complex:0.1309123422)93.1/97:0.0590055511)82.4/97:0.0238039510,(WP_079793206.1-Salmonella-enterica:0.0798966551,WP_001100807.1-Salmonella:0.0675448688)99.9/99:0.1137211014)87.6/52:0.0269022051,WP_015386517.1-Cronobacter-sakazakii:0.1451618471)68.8/14:0.0261311829)30.4/22:0.0072883721)89/50:0.0220371541,(WP_187216555.1-Escherichia-coli:0.1368049726,((((WP_049002539.1-Citrobacter-freundii:0.0969494123,(WP_249576357.1-Enterobacter-vonholyi:0.0468188301,WP_161617217.1-Yokenella-regensburgei:0.0344333691)96.4/100:0.0484423214)74.3/100:0.0181599435,(WP_079834758.1-Salmonella-enterica:0.0319101933,WP_058109100.1-Salmonella-enterica:0.0230796755)99.4/100:0.0764513505)96.8/88:0.0467689621,WP_032303308.1-Escherichia:0.0401470884)96.2/88:0.0474869641,WP_061892435.1-Escherichia-coli:0.0970008164)82.9/81:0.0128776990)87.3/84:0.0196666825)88/41:0.0180041032,(((((WP_109548670.1-Escherichia-coli:0.0340791437,WP_001100790.1-Escherichia:0.0223185728)93.1/100:0.0348390355,(WP_089706542.1-Escherichia-coli:0.0988950978,WP_097421351.1-Escherichia-coli:0.0649049119)85.8/95:0.0241977030)89.5/98:0.0365891967,WP_181237856.1-Escherichia-marmotae:0.0549713035)90.6/89:0.0292179564,WP_107179425.1-Escherichia:0.0659060796)44.8/37:0.0084774893,((WP_034458444.1-Buttiauxella-noackiae:0.1030284672,WP_253383865.1-unidentified-bacterial-endosymbiont:0.2046518220)89.8/98:0.0554665513,(WP_105591414.1-Pantoea-coffeiphila:0.0495463932,WP_123334936.1-Erwinia-sp.-JUb26:0.0292224591)99.9/100:0.1432630361)96/91:0.0595575432)27.2/37:0.0151361216)94.6/83:0.0276356214)75/8:0.0105596865,((WP_042093344.1-Escherichia-coli:0.1396366405,(WP_032659356.1-Enterobacter-hormaechei:0.2179734495,WP_029881813.1-Enterobacter:0.1823123930)92/78:0.0720641606)48/10:0.0193021007,WP_262938691.1-Escherichia-albertii:0.0769889476)93.6/10:0.0451978844)0/6:0.0019753571,((((WP_115348269.1-Hafnia-alvei:0.1110209559,WP_046458830.1-Obesumbacterium-proteus:0.2138384828)85.4/95:0.0457401942,WP_130961887.1-Hafnia-paralvei:0.1333378579)22.2/17:0.0277878567,WP_061059890.1-Hafnia-paralvei:0.1271276164)32.2/85:0.0301763609,(WP_175320495.1-Hafnia-paralvei:0.0590619159,WP_130996599.1-Hafnia-paralvei:0.1477447614)97.7/100:0.0896444604)99.4/93:0.0779549002)90.8/51:0.0240569870,(((WP_119938542.1-Enterobacter-huaxiensis:0.0447436335,WP_100029735.1-Escherichia-coli:0.1383195682)52.5/42:0.0509751811,WP_050871166.1-Enterobacter-cloacae-complex:0.0283370982)94.2/42:0.0526770559,(WP_228937168.1-Escherichia-marmotae:0.0280474081,WP_040079227.1-Escherichia-coli:0.0336746482)65.2/79:0.0042829408)97.6/53:0.0479496289)71.6/10:0.0058302084)74.1/22:0.0358379233)10.1/3:0.0180900067,(((WP_167863639.1-Erwinia-rhapontici:0.0999411062,WP_039385357.1-Pantoea-eucrina:0.1495364879)94.2/100:0.0574792523,WP_118663751.1-Erwinia-persicina:0.1075604163)66.7/99:0.0249170273,WP_062742814.1-Erwinia-persicina:0.1494013836)93.7/51:0.0713771068)0/16:0.0000025454)53.2/20:0.0176109684)80.9/47:0.0282802883,((((WP_238085128.1-Pseudescherichia-vulneris:0.0163209092,WP_032652659.1-Enterobacter-cloacae-complex:0.0820867429)71.3/80:0.0099514294,WP_129363899.1-Enterobacter-cloacae:0.0261425759)61.1/80:0.0144894433,(WP_257661844.1-Citrobacter-portucalensis:0.0993077273,WP_193134185.1-Citrobacter-freundii:0.0680865037)84.8/100:0.0268740253)93.1/94:0.0626285716,(WP_042284276.1-Citrobacter-sedlakii:0.1686646193,(WP_024193259.1-Escherichia-coli:0.0867774474,WP_099433281.1-Citrobacter-amalonaticus:0.0304980375)84.2/100:0.0632952587)97.2/100:0.0813275855)98.9/100:0.1024717488)85.6/83:0.0255915883,WP_171149131.1-Erwinia-sp.-JH02:0.1428840626)97.2/93:0.0567228114,((WP_241647798.1-Rosenbergiella-sp.-JB07:0.2084865614,WP_065370581.1-Kosakonia-sacchari:0.1450255368)67.6/46:0.0296997723,WP_181599127.1-Citrobacter-sp.-RHB36-C18:0.1629848504)80.9/46:0.0186223492)96.7/87:0.0518220654)79.4/67:0.0222830392,((((((((((((WP_182931197.1-Enterobacter-cloacae:0.1157353015,WP_211776196.1-Klebsiella-pneumoniae:0.0491712881)79.6/83:0.0204610966,(WP_047723323.1-Klebsiella-michiganensis:0.0370084778,WP_104011323.1-Klebsiella-michiganensis:0.0377633727)36.2/43:0.0091138915)71.5/64:0.0080309378,((WP_139968923.1-Klebsiella-pneumoniae:0.0771126222,WP_065799756.1-Klebsiella-pneumoniae:0.0055178055)0/100:0.0000025648,WP_095100910.1-Raoultella-sp.-10-1:0.0617168694)93/93:0.0301635516)87.9/45:0.0167745199,(WP_048284323.1-Pluralibacter-gergoviae:0.0998517179,WP_221694709.1-Klebsiella-aerogenes:0.0585655491)86.1/53:0.0243202655)0/26:0.0053406382,WP_234579544.1-Raoultella-terrigena:0.0332409706)76/30:0.0062011266,(WP_213760001.1-Klebsiella-aerogenes:0.0572466034,WP_223370324.1-Klebsiella-pneumoniae:0.0281712297)90/43:0.0168069422)73.3/32:0.0055667381,WP_149270444.1-Klebsiella-aerogenes:0.0343289954)57.9/44:0.0113079066,(((((WP_095100919.1-Raoultella-sp.-10-1:0.0474689190,WP_212709529.1-Tatumella-sp.-JGM130:0.0114930363)100/100:0.0970921544,WP_236948387.1-Klebsiella-sp.-2019SCSN059:0.0248697172)73.5/62:0.0084341499,((WP_114692711.1-Klebsiella-pneumoniae:0.0336550078,WP_259891680.1-Klebsiella-variicola:0.0280385677)0/47:0.0000020786,WP_123755035.1-Raoultella-sp.-BIGb0399:0.0333622516)85.8/65:0.0109689186)79.4/50:0.0094750460,WP_128320364.1-Raoultella-planticola:0.0470243417)68.7/37:0.0072204811,WP_046852082.1-Raoultella-planticola:0.0298113685)84.7/45:0.0100212276)77.2/48:0.0057582635,(WP_234537246.1-Raoultella-ornithinolytica:0.0399270763,WP_113990968.1-Raoultella:0.0171193408)97.3/99:0.0395815601)72.6/53:0.0083798518,WP_186601692.1-Klebsiella-quasipneumoniae:0.0196519434)92/69:0.0598809966,WP_236948353.1-Klebsiella-sp.-2019SCSN059:0.0617449664)96.3/100:0.0536820630,WP_123822891.1-Klebsiella:0.0264104651)98.9/100:0.0770305870)91.2/94:0.0432601011,WP_072663757.1-Enterobacteriaceae:0.0679890490)40.2/77:0.0310226602,WP_021241446.1-Enterobacter-roggenkampii:0.1460036521)96/100:0.0621070187)97.1/100:0.0533206641,XP_011503919.1-Ceratosolen-solmsi-marchali:0.0731857653);

**Nematode RML-3, with bacterial and eukaryotic homologs**

**Shown in Figures S2**

ML tree method: IQ-TREE

Sequences: 2787

Substitution model: JTT+I+G4

(WP_063501760.1-Escherichia-coli:0.0473272683,(WP_044173167.1-Phytobacter-massiliensis:0.0459977662,((((((((WP_203451401.1-Lelliottia-sp.-RWM.1:0.0769145143,WP_185654835.1-Citrobacter-braakii:0.1438697905)89.1/100:0.0985060868,WP_183170804.1-Atlantibacter-sp.-RC6:0.2872496398)95.1/100:0.1319958827,(WP_090085547.1-Escherichia-coli:0.0525372469,(WP_103283990.1-Citrobacter:0.2267663018,WP_234083964.1-Enterobacter-quasiroggenkampii:0.1728745889)55.3/100:0.0333087707)99.7/100:0.2549786991)0/78:0.0031595878,(WP_160868038.1-Pantoea-sp.-Taur:0.4110265923,WP_008499438.1-Enterobacter-cloacae-complex:0.2657544203)77/96:0.0681466337)91.5/90:0.0566428340,(((((((((((((((((((WP_249615874.1-Craterilacuibacter-sp.-RT1T:0.5831885125,(WP_163316497.1-Crenobacter-sedimenti:0.1827838480,WP_160797365.1-Craterilacuibacter-sinensis:0.1720985962)97.7/100:0.2548037836)65.8/98:0.0476262346,WP_014088374.1-Pseudogulbenkiania-sp.-NH8B:0.2464324135)85.6/96:0.0979843773,WP_008952446.1-Pseudogulbenkiania-ferrooxidans:0.3283153124)84.5/98:0.0811472236,(WP_174873379.1-Vogesella-oryzae:0.2383462038,WP_147289921.1-Crenobacter-cavernae:0.1709289598)77.2/96:0.0437294594)15.6/66:0.0409856417,((((((((WP_004123988.1-Rhizobium-freirei:0.1238523092,WP_095433322.1-Rhizobium-sp.-11515TR:0.0425744339)72.3/81:0.0296355083,WP_106999541.1-Rhizobium-sp.-JAB6:0.0639077355)0/74:0.0000026835,((WP_174074493.1-Agrobacterium-rhizogenes:0.1211993580,((WP_183765619.1-unclassified-Rhizobium:0.0480693076,WP_120667780.1-unclassified-Rhizobium:0.0642365709)66.4/100:0.0343928313,WP_037195771.1-Rhizobium-lusitanum:0.0320589755)66.8/99:0.0146064901)80/76:0.0163479672,((WP_199146672.1-Rhizobium-sp.-ASV8:0.2108746369,WP_184706966.1-Rhizobium-lusitanum:0.0580398075)92.1/93:0.0617968424,(((WP_145321829.1-unclassified-Rhizobium:0.1335443331,WP_112341687.1-Rhizobium-tropici:0.0452855841)11.1/66:0.0128806811,WP_168880117.1-Rhizobium-sp.-P28RR-XV:0.0523194019)87.6/75:0.0355358752,WP_092719658.1-Rhizobium-multihospitium:0.0450643054)76.5/68:0.0100455398)0/56:0.0000023940)92.2/90:0.0246092649)88.4/80:0.0169005120,WP_183712806.1-unclassified-Rhizobium:0.0399622478)80.1/96:0.0275930046,WP_015341698.1-Rhizobium:0.0180605313)99.5/100:0.1985344024,(WP_043579071.1-Chromobacterium:0.0696683203,WP_071113424.1-Chromobacterium-sphagni:0.1290956580)99.7/100:0.2464837024)92.4/100:0.1002385508,(((WP_189531325.1-Paludibacterium-paludis:0.6095757867,(WP_159673755.1-Andreprevotia-sp.-IGB-42:0.2450238049,WP_084093192.1-Andreprevotia-lacus:0.0817355116)98.7/100:0.2405884253)70.7/100:0.0629822902,(WP_043620211.1-Chromobacterium-piscinae:0.2868405962,(WP_103900886.1-Chromobacterium-alticapitis:0.2807377629,WP_045052286.1-Chromobacterium-violaceum:0.3094363315)27.6/100:0.0651660967)96.9/100:0.1766464975)77.1/100:0.0377506193,WP_020932001.1-Cycloclasticus-zancles:0.3999169596)29.1/97:0.0343735036)83.4/97:0.0640728374,WP_137008594.1-Aquitalea-aquatilis:0.5318302369)0/43:0.0103998608)81.2/91:0.0773251251,WP_211007190.1-Pectobacterium-brasiliense:0.6272181998)5.8/58:0.0232308608,(((WP_147247296.1-Brenneria-salicis:0.0464620400,WP_172289906.1-unclassified-Brenneria:0.1290989531)86.6/100:0.0495904546,WP_039356706.1-Pectobacterium-actinidiae:0.0974644232)92.7/100:0.1184906207,((WP_198997094.1-Pectobacterium-carotovorum:0.0232381968,(WP_165707548.1-Pectobacterium:0.0406663135,WP_174866005.1-Pectobacterium-polaris:0.0344509140)98.8/100:0.0800861560)58.4/99:0.0196397247,WP_145962313.1-Pectobacterium-peruviense:0.0700333242)98.4/99:0.1594444362)93.8/90:0.1071283694)86.5/83:0.0490308462,((((((((WP_236369955.1-Aeromonas-veronii:0.0646386890,(((WP_042019775.1-Aeromonas-sobria:0.0644446781,WP_139495680.1-Aeromonas-veronii:0.0306282674)96.9/100:0.0714871856,(WP_098969895.1-Aeromonas-sp.-CU5:0.0569232973,WP_159163188.1-Aeromonas-veronii:0.0607712430)89.4/100:0.0462587851)91.1/100:0.0445064693,WP_113736183.1-Aeromonas-caviae:0.0522737000)39.2/78:0.0240542649)0/84:0.0000029734,(((((WP_080988732.1-Aeromonas-caviae:0.0547811236,WP_108536035.1-Aeromonas-sp.-HMWF014:0.0454292516)92.5/100:0.0449309375,(WP_252615985.1-Aeromonas-hydrophila:0.0727844945,WP_041208446.1-Aeromonas-jandaei:0.2768066525)72.4/97:0.0167999943)88.3/95:0.0232541466,WP_214003767.1-Aeromonas-caviae:0.0680073018)0/93:0.0000020555,WP_143237783.1-Aeromonas-caviae:0.0784252202)29.5/85:0.0140474407,(WP_113739100.1-Aeromonas-veronii:0.0634223334,(WP_226032029.1-Aeromonas-hydrophila:0.0380440431,WP_139459904.1-Aeromonas-veronii:0.1595525299)85.2/93:0.0182806826)24.1/86:0.0076689524)99.7/100:0.1035496505)95.5/97:0.0660634923,WP_201035193.1-Aeromonas-media:0.1576649706)82.9/90:0.0506190532,(WP_082969903.1-Aeromonas-dhakensis:0.1522446090,WP_104454419.1-Aeromonas-sp.-ASNIH2:0.0748787608)77/79:0.0364993844)0/85:0.0020414488,(WP_220777726.1-Shewanella-hafniensis:0.0375505885,WP_055647506.1-Shewanella-sp.-Sh95:0.0400344363)97.1/100:0.0611779255)94/86:0.0588375053,(WP_263173612.1-Shewanella-sp.-SM73:0.1808966536,((WP_263191039.1-Shewanella-sp.-SM87:0.0883833225,WP_263255112.1-Shewanella-sp.-SM32:0.0542439493)81.8/100:0.0214408163,(WP_263178828.1-Shewanella-sp.-SM101:0.0505595149,WP_109248233.1-Shewanella-algae:0.0903281724)92.8/100:0.0487548517)86.7/100:0.0388948097)88.9/99:0.0552502566)86.6/89:0.0527300209,(((((WP_059168599.1-Aeromonas-salmonicida:0.1263064877,WP_033115905.1-Aeromonas-lacus:0.0543783816)87.8/95:0.0276449457,(WP_058394234.1-Aeromonas-salmonicida:0.0369956854,WP_257690725.1-Aeromonas-veronii:0.0612819651)81.2/100:0.0269845162)75.3/95:0.0154198639,WP_223920301.1-Aeromonas-caviae:0.0468887940)99.6/95:0.1061536281,WP_226058924.1-Shewanella-sp.-SR1:0.0470199044)96.1/94:0.1134311776,((WP_248067241.1-unclassified-Shewanella:0.1255632032,WP_148505453.1-Vibrio-cholerae:0.1240884665)84.2/100:0.0605016980,((WP_218558465.1-Shewanella-putrefaciens:0.1035046887,WP_011919582.1-Shewanella-putrefaciens:0.0305380840)99.7/100:0.1480969259,(((((WP_050718733.1-Aeromonas-tecta:0.1849223573,WP_100859741.1-Aeromonas-lusitana:0.0480089532)73.8/97:0.0111663719,WP_042654929.1-Aeromonas-encheleia:0.0714291874)96.5/100:0.0698625144,WP_077098334.1-Aeromonas-hydrophila:0.0736133506)79.6/95:0.0192925877,WP_252420256.1-Aeromonas-taiwanensis:0.1086978471)77.9/94:0.0160170224,WP_043167431.1-Aeromonas-hydrophila:0.1198640155)97.9/100:0.1578058899)62.1/98:0.0419542824)61.5/98:0.0729881393)87/94:0.0884940101)45.7/89:0.0671894920,(((((WP_073350970.1-Aeromonas-hydrophila:0.1522458887,WP_190284026.1-Aeromonas-hydrophila:0.0447667482)87.7/100:0.0477419349,WP_241344946.1-Aeromonas-sp.-MR16:0.0831220775)90.1/100:0.0959407406,WP_223952962.1-Aeromonas-caviae:0.2910519852)55.7/77:0.0616635186,WP_244081328.1-Aeromonas-caviae:0.2463379204)4.5/45:0.0352174950,((WP_208149206.1-Shewanella-algae:0.3969885301,(((WP_045500143.1-Vibrio-hyugaensis:0.2253345896,WP_045419977.1-Vibrio-campbellii:0.0978450684)29.8/98:0.0428841719,WP_025533544.1-Vibrio-parahaemolyticus:0.0860937416)93.1/100:0.0546616486,WP_011788809.1-Shewanella-sp.-W3-18-1:0.0903425977)60.5/99:0.0234623684)76.6/100:0.0321088930,WP_219142360.1-Aeromonas-sanarellii:0.1727131102)99.7/100:0.2131416435)44.6/67:0.0249113568)88.7/88:0.0603222025)77/82:0.0475923923,((WP_076573677.1-Aeromonas-sp.-RU39B:0.1682469386,WP_134696535.1-Aeromonas-taiwanensis:0.8331967735)91/99:0.1334250492,WP_108652490.1-Dongshaea-marina:0.2583988238)93/98:0.1102694594)66.9/77:0.0356609140,((((WP_234664799.1-Aeromonas-allosaccharophila:0.2004500286,(((((((WP_245131329.1-Aeromonas-veronii:0.2037882021,(WP_225159845.1-Aeromonas-caviae:0.1594643808,(WP_163135231.1-Aeromonas-rivipollensis:0.0397007750,WP_241279715.1-Aeromonas-sp.-MR19:0.0365021021)100/100:0.2916431545)48.4/99:0.0352016548)68.6/67:0.0170635845,((((WP_213320225.1-Aeromonas-caviae:0.0605667009,WP_203764723.1-Aeromonas-caviae:0.1173111705)94.8/100:0.1278494828,WP_069554789.1-Aeromonas-sp.-DNP9:0.1986300330)47.2/97:0.0681956485,(WP_257692500.1-Aeromonas-caviae:0.2298898828,WP_113719982.1-Aeromonas-salmonicida:0.1499220874)79.3/100:0.0718375447)88.8/99:0.0815332719,((WP_050666161.1-Aeromonas-schubertii:0.0714710024,WP_059168611.1-Aeromonas-salmonicida:0.1519140105)79.9/98:0.0373330237,WP_210600017.1-Aeromonas-sp.-MrichA-1:0.0560017998)79.9/98:0.0399002456)81/90:0.0379493255)89.3/57:0.0434435773,(((((WP_148612677.1-Aeromonas-sobria:0.0530219523,(WP_201355655.1-Aeromonas-hydrophila:0.0537071144,WP_201951905.1-Aeromonas-veronii:0.0400110415)42.2/100:0.0076859358)32.9/97:0.0138438435,WP_171268810.1-Aeromonas-media:0.0290139393)100/100:0.1793249705,(WP_042063659.1-Aeromonas-allosaccharophila:0.0421623673,WP_256684552.1-Aeromonas-salmonicida:0.0398906478)76.4/98:0.0208023777)64.7/99:0.0101096519,(((WP_212539243.1-Aeromonas-salmonicida:0.0413871375,((WP_187590672.1-Aeromonas-hydrophila:0.1466313119,WP_060586794.1-Aeromonas-schubertii:0.0626965377)2.3/67:0.0137445786,WP_017782281.1-Aeromonas-dhakensis:0.0436274261)99.3/100:0.1145270448)88.3/100:0.0307334231,WP_201904077.1-Aeromonas-veronii:0.0710884271)77.1/100:0.0235529475,WP_206874356.1-Aeromonas-veronii:0.0130548245)73.9/99:0.0103999208)93.3/99:0.0803310200,WP_042649704.1-Aeromonas-media:0.1528456166)90/85:0.0905699068)1.2/14:0.0090118717,((WP_101347551.1-Aeromonas-sobria:0.0825793016,WP_201955411.1-Aeromonas-veronii:0.0765583293)94.3/100:0.0801841214,(WP_136501435.1-Aeromonas-veronii:0.2017555011,(WP_268029297.1-Aeromonas-hydrophila:0.1338608995,WP_201942265.1-Aeromonas-veronii:0.1676156429)98/100:0.1522185493)87.3/98:0.0631779617)94.7/80:0.0693875869)80.8/43:0.0296788014,WP_139406182.1-Aeromonas-veronii:0.3764323460)68.4/73:0.0169209976,WP_253626250.1-Aeromonas-hydrophila:0.2287002567)65.1/38:0.0139558200,((((WP_268421127.1-Aeromonas-media:0.0837468250,WP_111910157.1-Aeromonas:0.0255248319)81.4/100:0.0208873561,WP_128343177.1-Aeromonas-caviae:0.0569845465)96/100:0.0662025044,WP_268037199.1-Aeromonas-dhakensis:0.1267906002)47.2/91:0.0143513556,(WP_139733908.1-Aeromonas-veronii:0.0188111912,((WP_234579942.1-Aeromonas-salmonicida:0.0902228666,(WP_234676856.1-Aeromonas-rivipollensis:0.0197000781,WP_269382839.1-Aeromonas-hydrophila:0.1013324244)72.2/100:0.0108841001)43.4/99:0.0113026402,WP_222134508.1-Vibrio-cholerae:0.1884492235)67.6/99:0.0123873282)99.4/100:0.1077617125)86.4/100:0.0339187533)71.7/38:0.0295580519)73.9/42:0.0235587395,((((WP_216990617.1-Aeromonas-sp.-FDAARGOS-1409:0.1535139092,WP_236563281.1-Aeromonas-veronii:0.1845603405)64.4/98:0.0410600322,(((((WP_223945983.1-Aeromonas-caviae:0.1431018098,((WP_040098646.1-Aeromonas:0.1188468950,((WP_263309349.1-Aeromonas-caviae:0.0481337702,WP_041217344.1-Aeromonas-hydrophila:0.0577319505)75.4/79:0.0089333744,(WP_257686696.1-Aeromonas-veronii:0.0436012588,WP_255912544.1-Aeromonas-sp.-JL9:0.0387148237)75.8/69:0.0098861122)82.1/91:0.0201686103)75.4/87:0.0143344354,((WP_139750984.1-Aeromonas-media:0.0381620345,WP_224430864.1-Aeromonas-rivuli:0.0317161923)77.1/89:0.0244592621,WP_041996113.1-Aeromonas-bivalvium:0.0676200518)99/100:0.0784104797)67.1/61:0.0049344993)79.4/63:0.0111395653,((WP_265458754.1-Aeromonas-salmonicida:0.0982757705,WP_125597700.1-Aeromonas-salmonicida:0.0280350426)70.5/100:0.0050702332,WP_163128050.1-Aeromonas-allosaccharophila:0.1090150566)0/65:0.0000027997)74.1/95:0.0079629895,(WP_180911080.1-Aeromonas-caviae:0.0490815699,WP_264818242.1-Aeromonas-caviae:0.0325115460)86.9/100:0.0164499815)87.6/96:0.0493743619,WP_265776268.1-Aeromonas-veronii:0.1536545490)79.2/75:0.0418804362,(((WP_201942221.1-Aeromonas-veronii:0.1980811181,(((WP_268854930.1-Aeromonas-enteropelogenes:0.1460522365,(WP_203131090.1-Aeromonas-jandaei:0.0740045440,((WP_042657830.1-Aeromonas:0.0506511878,WP_111901368.1-Aeromonas-veronii:0.0467107483)64.8/98:0.0261485804,WP_139692403.1-Aeromonas-sobria:0.0388481859)0/96:0.0000021694)76.1/98:0.0080406999)57.1/93:0.0139054926,WP_223380193.1-Aeromonas-enteropelogenes:0.0506919879)0/73:0.0000027820,WP_159143398.1-Aeromonas-veronii:0.0312902769)58.5/77:0.0426786428)77.2/94:0.0518765177,(WP_236321701.1-Aeromonas-veronii:0.0943403187,WP_257119864.1-Aeromonas-hydrophila:0.0242331097)93.6/99:0.0789233585)49.2/99:0.0302013501,(WP_257707542.1-Aeromonas-caviae:0.0686940575,WP_182951616.1-Aeromonas-sp.-WP2-W18-CRE-05:0.0735160494)97.6/100:0.1026363445)88.4/99:0.0876869737)40/74:0.0382358715)86.1/98:0.0477710484,WP_265468300.1-Aeromonas-salmonicida:0.1573403330)84.1/98:0.0462719890,(WP_100646402.1-Aeromonas-veronii:0.0531045614,((WP_252601295.1-Aeromonas-dhakensis:0.0287526964,WP_139432874.1-Aeromonas-dhakensis:0.1831596308)87.7/100:0.0379964783,WP_106553824.1-Aeromonas-hydrophila:0.0762925730)92.1/100:0.0511659915)99.9/100:0.2116675475)60.4/86:0.0342870775)81.1/79:0.0286127461,WP_210550341.1-Aeromonas-sp.-SrichE-2G:0.4594191057)25.9/38:0.0182694755,((WP_224211331.1-Aeromonas-enteropelogenes:0.2373731681,WP_082041448.1-Aeromonas-eucrenophila:0.3580589912)80.5/99:0.0926163152,(((WP_268010485.1-Aeromonas-dhakensis:0.0916297212,WP_098985072.1-Aeromonas-sp.-CA23:0.0526733783)97.5/100:0.1494397581,(WP_181268965.1-Aeromonas-veronii:0.0466828565,(WP_129146711.1-Aeromonas-caviae:0.0295812927,(WP_081843667.1-Aeromonas-taiwanensis:0.0242960057,WP_139744464.1-Aeromonas-media:0.0483440309)86.3/100:0.0290674570)79.7/100:0.0196464063)83.4/100:0.0908717300)95.4/100:0.1860546863,WP_200561028.1-unclassified-Erwinia:0.4325750497)92/99:0.1367544146)85.5/39:0.0576114456)74.6/86:0.0156499524)0/20:0.0000029410,(((((WP_250059255.1-Halomonas-sp.-ATCH28:0.3697457271,(WP_044615503.1-Gynuella-sunshinyii:0.6319068493,WP_179153381.1-Chromohalobacter-salexigens:0.2809415662)49.6/84:0.0415650584)83.1/92:0.0762635632,WP_250939176.1-unclassified-Larsenimonas:0.4108289454)96.2/91:0.2094736927,((((((((((((((((((((((((((WP_136867012.1-Guyparkeria-sp.-SB14A:0.3861890486,WP_058575749.1-unclassified-Guyparkeria:0.2950454650)94.8/100:0.1266860011,(WP_218634960.1-Marinobacter-arenosus:0.1592445097,WP_240716217.1-Halomonas-neptunia:0.1146069472)96.5/100:0.1367250709)79.6/100:0.0407350955,WP_106478747.1-Phytohalomonas-tamaricis:0.3443943712)95.6/100:0.1525268817,WP_153720048.1-Spiribacter-sp.-C176:0.4590042200)36.9/86:0.0133503229,(WP_177138191.1-Hydrogenophaga-aromaticivorans:0.1238724392,(WP_133157403.1-Hydrogenophaga-pseudoflava:0.2089188833,(WP_236964528.1-Hydrogenophaga-sp.-SL48:0.0897475761,(WP_226493830.1-Hydrogenophaga-taeniospiralis:0.0459612683,WP_210824367.1-Hydrogenophaga-aromaticivorans:0.0846032082)85.3/100:0.0642539826)98.3/100:0.1204503306)81.5/96:0.0410673421)86.9/94:0.0503236091)93.8/91:0.0910347919,((WP_138213821.1-Pseudomonas-nitroreducens:0.0451208152,WP_017517140.1-Pseudomonas-nitroreducens:0.0287033994)100/100:0.3853668329,WP_085721622.1-Pseudomonas:0.1687624173)70.2/90:0.0233862237)41.5/85:0.0608418388,WP_104727848.1-Pseudomonas-oleovorans:0.1617399021)90.4/90:0.0712395156,(WP_169965430.1-unclassified-Pseudomonas:0.0559886847,WP_047278503.1-Pseudomonas-lundensis:0.0329117770)99.4/100:0.1461424025)82.6/80:0.0342883483,((WP_065086507.1-Pseudomonas:0.3059295556,WP_158658301.1-Achromobacter-sp.-AONIH1:0.5759256053)74.5/78:0.0485935357,WP_205489922.1-Pseudomonas-lactucae:0.2165174306)77.1/68:0.0267298988)77.5/54:0.0263318743,(WP_042945119.1-Pseudomonas-fluorescens:0.0910096103,WP_181643258.1-Pseudomonas-sivasensis:0.0556937101)100/100:0.1782725013)74/78:0.0201473503,(((((((WP_225237063.1-Massilia-oculi:0.4127317759,WP_147937383.1-Massilia-arenae:0.1375581625)98.8/100:0.2625333376,WP_136222466.1-Massilia-sp.-Mn16-1_5:0.1090033197)3.1/91:0.0240622137,WP_192051557.1-unclassified-Massilia:0.1378646352)98.6/100:0.1603109266,WP_155457065.1-Massilia-eburnea:0.2324296737)86/95:0.0447441885,WP_214501190.1-Pseudomonas-sp.-RC3H12:0.3104079026)85.3/90:0.0337255122,((((WP_029241333.1-Pseudomonas:0.0796372413,((WP_054996663.1-Pseudomonas-ficuserectae:0.0526142213,(WP_007252556.1-Pseudomonas-syringae-group:0.0604665510,WP_025390602.1-Pseudomonas-syringae:0.0253098961)55.8/100:0.0268040151)93.5/100:0.0577945384,WP_125830676.1-Pseudomonas-sp.-v388:0.0897723088)92/100:0.0637132370)84.6/100:0.0816382021,((WP_016973326.1-Pseudomonas-tolaasii:0.1443475146,WP_099548838.1-Pseudomonas-sp.-2995-3:0.1179081996)79.4/100:0.0467411256,(WP_177015116.1-Pseudomonas-costantinii:0.0610262890,WP_141063556.1-Pseudomonas-fluorescens:0.1208124565)93/100:0.0858749386)95.8/100:0.1129905000)96.3/100:0.1262461334,(WP_033955816.1-Pseudomonas-aeruginosa:0.0353237348,WP_003153917.1-Pseudomonas-aeruginosa-group:0.0441763925)99.9/100:0.2699428454)93.9/95:0.1007327574,((WP_058488093.1-Pseudomonas:0.2004234436,WP_043256200.1-Pseudomonas-knackmussii:0.1667140851)69.4/95:0.0526339413,((WP_088420173.1-Pseudomonas-nitroreducens:0.1614780262,WP_254475170.1-unclassified-Pseudomonas:0.0823481566)88.1/100:0.0443029801,WP_205338011.1-Pseudomonas-sp.-PDNC002:0.0711221638)99.6/100:0.1954041342)81.2/94:0.0431697652)79.6/78:0.0490410488)0/51:0.0000024047,((WP_077733980.1-Paenalcaligenes-hominis:0.0561478723,WP_167661700.1-Paenalcaligenes-hominis:0.0361974537)100/100:0.3132441505,((((WP_191471912.1-Pseudomonas-asiatica:0.2413341397,(WP_043318452.1-Pseudomonas-sp.-PI1:0.0572299068,WP_043268312.1-Pseudomonas:0.0387118028)47.4/100:0.0305550115)93.5/100:0.0792545124,(WP_089390899.1-Pseudomonas-delhiensis:0.1563426771,WP_076426141.1-Pseudomonas-alcaligenes:0.1926388278)10.4/58:0.0211550720)96.8/100:0.0932144523,WP_233297479.1-Pseudomonas:0.2085536668)74.8/99:0.0185611927,(WP_184594840.1-Pseudomonas-nitritireducens:0.2209404733,WP_192381882.1-Pseudomonas-sp.-PDM20:0.1468548615)98.9/100:0.1827160801)90.2/85:0.0545216201)49.7/64:0.0262488795)82.8/54:0.0253021076)73.7/50:0.0148604271,(((((((WP_269022943.1-Methylomonas-sp.-MP1:0.1188725092,(WP_196437371.1-Methylomonas-sp.-LL1:0.1602573940,(WP_150047608.1-Methylomonas-rhizoryzae:0.3826309767,WP_066983474.1-Methylomonas-lenta:0.1043634302)87.6/100:0.0777487362)90.2/100:0.0811181763)88.4/100:0.0783078815,(WP_140911374.1-Methylomonas-koyamae:0.2780985752,(WP_071159776.1-Methylomonas-sp.-LWB:0.2109112462,((WP_020485671.1-Methylomonas-sp.-MK1:0.2408316808,(WP_256607223.1-Methylomonas-sp.-WSC-7:0.0897840645,(WP_205450646.1-Methylomonas-sp.-EFPC1:0.1045122433,WP_064036612.1-Methylomonas-methanica:0.0000023961)44.4/100:0.0098506184)90.2/100:0.0720711751)90.8/99:0.0885004270,WP_036273427.1-Methylomonas:0.1441055318)0/98:0.0000029613)83.1/100:0.0543022299)87.9/100:0.0695857832)86.3/100:0.0647933619,WP_064030819.1-Methylomonas-koyamae:0.2603647008)90.4/74:0.1083463138,(WP_019964198.1-Pannonibacter-phragmitetus:0.8274060240,WP_138632053.1-Pseudoalteromonas-spongiae:0.3354948840)86.5/74:0.1067799050)94.2/100:0.1152999179,((((WP_006216420.1-Achromobacter-piechaudii:0.1475134193,WP_219991400.1-Achromobacter-sp.-ES-001:0.1053475194)95.3/100:0.0673160034,WP_183017169.1-Achromobacter-sp.-UMC46:0.0571224340)88/92:0.0446677337,WP_100503206.1-Achromobacter-ruhlandii:0.1781354279)71.3/87:0.0214934790,WP_175192416.1-Achromobacter-deleyi:0.0926881508)100/100:0.3049797809)82.8/94:0.0561815737,(WP_209394989.1-Rhodoferax-sp.-AJA081-3:0.3756580921,WP_094844257.1-Bordetella-genomosp.-11:0.2422743574)92.8/91:0.1159308421)73.9/85:0.0549699124,(WP_076518421.1-Achromobacter-sp.-MFA1-R4:0.0770221383,(WP_104142565.1-Achromobacter-spanius:0.0335236736,WP_120157261.1-Achromobacter-sp.-B7:0.0641868321)93.8/100:0.0705171527)99.5/99:0.1515673219)84.2/38:0.0422128411)50/46:0.0455066418,(WP_209872004.1-Pseudomonas-sp.-BP8:0.2365615700,WP_066093302.1-Hydrogenophaga-crassostreae:0.2187337973)89.6/73:0.0828839046)84.4/71:0.0363040721,(WP_020564242.1-Methylosarcina-fibrata:0.2678906146,(WP_125865093.1-Stutzerimonas-stutzeri:0.2166166512,WP_200290630.1-Stutzerimonas-balearica:0.1523897009)83.3/98:0.0654795735)73.3/46:0.0292840482)73.9/42:0.0103671427,(WP_101152997.1-Pseudomonas-sp.-Choline-3u-10:0.1565956810,((WP_046617631.1-Stutzerimonas-stutzeri:0.0923118992,WP_135265705.1-Stutzerimonas-stutzeri:0.0594630833)91.6/100:0.0890446128,WP_003299494.1-Stutzerimonas-stutzeri:0.1116628695)86.1/100:0.0414945523)93.9/100:0.0641279389)93.6/86:0.0487685807,(WP_180152801.1-Parapusillimonas-granuli:0.1780696145,(WP_088146851.1-Achromobacter-denitrificans:0.1156650690,WP_105555808.1-Achromobacter-sp.-MYb9:0.1185578792)97.4/100:0.1441665942)87.5/92:0.0590089109)67/74:0.0120156885,WP_046857648.1-Pseudomonas-sp.-CCOS-191:0.2817617004)90.8/94:0.0375710090,WP_074973890.1-Pseudomonas-otitidis:0.2757227802)81.6/85:0.0200891592,((((WP_102075761.1-Pollutimonas-subterranea:0.2546866957,(WP_066098770.1-Halothiobacillus-sp.-LS2:0.1906405513,WP_070073009.1-Acidihalobacter-aeolianus:0.2574107624)93.2/98:0.1523028329)70.3/64:0.0193034863,WP_217963608.1-Candidimonas-humi:0.3430786050)82.2/67:0.0522427700,(((WP_057247240.1-Duganella-sp.-Root198D2:0.3496702054,WP_212592286.1-Janthinobacterium-sp.-TND4EL3:0.2244075260)90.7/96:0.1091070065,WP_102071501.1-Pollutimonas-nitritireducens:0.3327187817)36.9/61:0.0260224265,WP_047550623.1-Methylotenera-sp.-G11:0.2876203458)28.5/61:0.0580564580)80.8/87:0.0335334373,(WP_019141909.1-Noviherbaspirillum-massiliense:0.2195317163,WP_146330669.1-Noviherbaspirillum-sp.-UKPF54:0.3159940748)66.2/87:0.0721963480)86.5/92:0.0332911629)74.4/79:0.0113043631,((((WP_227390214.1-Halomonas-malpeensis:0.7825759270,WP_058422493.1-Pseudomonas:0.1877583749)85.1/100:0.1168571062,(WP_092579398.1-Achromobacter-sp.-NFACC18-2:0.5684619968,WP_223462047.1-Pseudomonas-sp.-A-R-26:0.2185556918)71.9/99:0.0418786948)93.2/99:0.1039247417,WP_011111299.1-Nitrosomonas-europaea:0.2360941039)31.2/88:0.0197523379,WP_068803504.1-Immundisolibacter-cernigliae:0.2834526989)87.1/85:0.0711911651)77.5/84:0.0220186427,((WP_230502293.1-Janthinobacterium-sp.-FW305-128:0.5062259761,WP_249918475.1-Pseudomonas-sp.-R3.Fl:0.2268627207)76.2/98:0.0564624207,WP_227683402.1-Pusillimonas-sp.-MFBS29:0.3975484558)48.8/66:0.0320737547)83.8/68:0.0290016845,((WP_139871681.1-Candidatus-Methylopumilus-universalis:0.2612486806,(WP_139929800.1-Candidatus-Methylopumilus-universalis:0.3604989433,WP_140005423.1-Candidatus-Methylopumilus-planktonicus:0.0979658018)89.5/100:0.1346993089)99.8/100:0.3111155790,(((WP_220354719.1-Rhizobium-cauense:0.2905946722,WP_016557527.1-Rhizobium-grahamii:0.1588158943)89.3/98:0.0640146272,((WP_184450570.1-Rhizobium-sp.-BK538:0.2665947742,(WP_072373837.1-Rhizobium-tibeticum:0.1137059990,WP_007537558.1-Rhizobium-mesoamericanum:0.0679186554)84.8/99:0.0501740837)28.4/96:0.0309271550,WP_018896487.1-Rhizobium-sp.-2MFCol3.1:0.2679419346)68.9/84:0.0238960973)41/86:0.0277231905,(WP_153273632.1-Rhizobium-grahamii:0.2922139509,WP_233136718.1-Agrobacterium-vaccinii:0.3053929346)73.2/98:0.0513407103)97.2/100:0.1440831234)90.1/96:0.1102470514)87.8/92:0.0577261421,(((((((((((WP_213263532.1-Burkholderia-pyrrocinia:0.1851382431,WP_060279477.1-Burkholderia-cepacia-complex:0.1450449214)97.6/100:0.1928981784,(((WP_208101977.1-Burkholderia-sp.-D-99:0.0406916227,WP_205714023.1-Burkholderia-cepacia:0.0356098538)73.4/100:0.0092165431,WP_174919663.1-Burkholderia-metallica:0.0268405464)87.7/100:0.0514066307,WP_077021900.1-Burkholderia-cenocepacia:0.1025972024)99/100:0.2781827776)73.9/100:0.0787547283,WP_083560927.1-Oceanococcus-atlanticus:0.6965979049)84.6/39:0.0859831078,((((((((((((WP_065305794.1-Janthinobacterium-psychrotolerans:0.1627463853,WP_035822645.1-Janthinobacterium-sp.-RA13:0.0759357208)0/81:0.0000029700,WP_034778721.1-Janthinobacterium-lividum:0.0453308801)44.9/90:0.0084912980,WP_139090581.1-Janthinobacterium-lividum:0.1041472800)72.8/90:0.0082761776,WP_152282105.1-Janthinobacterium-violaceinigrum:0.0246403851)75.8/91:0.0085038234,WP_183742413.1-unclassified-Janthinobacterium:0.0577176074)73.9/91:0.0079526882,((((WP_094442213.1-Janthinobacterium-sp.-PC23-8:0.0795571354,WP_230539136.1-Janthinobacterium-sp.-FW305-129:0.0640213937)24.2/100:0.0078909840,WP_215403374.1-Janthinobacterium-sp.-JC611:0.0291818453)72.7/100:0.0111367890,WP_102124214.1-Janthinobacterium-sp.-ROICE36:0.0413551227)77.2/99:0.0086593599,WP_096238059.1-Janthinobacterium-svalbardensis:0.0241793892)77.3/99:0.0081690677)99/99:0.1395493130,WP_023495308.1-Methyloglobulus-morosus:0.2187807798)83.5/43:0.0260648994,(WP_101482628.1-unclassified-Janthinobacterium:0.2504462505,(WP_099917421.1-Massilia-psychrophila:0.1729130529,(WP_205319084.1-Glaciimonas-sp.-PAMC28666:0.0877467157,WP_092396716.1-Collimonas-sp.-OK412:0.1521995334)69.6/96:0.0161288148)88.5/100:0.0386170613)52.5/42:0.0040625977)99/99:0.1189358367,((WP_162020603.1-Undibacterium-crateris:0.1910846424,WP_186884883.1-Undibacterium:0.0574299452)85/97:0.0355314972,WP_186861877.1-Undibacterium-griseum:0.0575826228)100/100:0.1855039641)68.6/88:0.0143768967,WP_217900122.1-Noviherbaspirillum-humi:0.2467495018)83.5/90:0.0466073970,((WP_195874647.1-Herminiimonas-contaminans:0.1627658275,WP_041297045.1-Janthinobacterium-sp.-Marseille:0.1139708558)78.4/100:0.0301720333,(WP_076590920.1-Herminiimonas-arsenitoxidans:0.2008140234,(WP_206604468.1-Herminiimonas-fonticola:0.0550656650,WP_188420499.1-Oxalicibacterium-solurbis:0.2216637021)73.4/89:0.0199265650)85/99:0.0488283939)98.5/99:0.1257564856)52.9/40:0.0608898397,(WP_247737070.1-Thauera-aromatica:0.0631362975,WP_039805139.1-Azotobacter-chroococcum:0.1942483749)97.9/100:0.1601564223)79.2/96:0.0675012808)80.3/71:0.0392303984,((((WP_239190817.1-Candidatus-Nitrotoga-sp.-HW29:0.0702006759,WP_239797721.1-Candidatus-Nitrotoga-arctica:0.0355493419)63.7/74:0.0094805195,WP_239287644.1-Candidatus-Nitrotoga-sp.-1052:0.0261516731)67.6/71:0.0064491991,WP_239235002.1-Candidatus-Nitrotoga-sp.-BS:0.0680609924)100/100:0.3475112909,((WP_237889684.1-Dissulfurimicrobium-hydrothermale:0.4104233824,WP_009207329.1-Sulfuricella-denitrificans:0.3965288545)81/33:0.0529925819,WP_237247350.1-Sideroxyarcus-emersonii:0.2379135710)85.6/69:0.1017150516)74.5/79:0.0211070933)61.9/31:0.0445549303,(WP_054689510.1-Desulfosarcina-cetonica:0.3568086265,((WP_104228621.1-Solimonas-fluminis:0.6411532726,WP_217345508.1-Noviherbaspirillum-sp.-L7-7A:0.2159397321)85.9/88:0.0809858798,WP_047549861.1-Methylotenera-sp.-G11:0.3158679999)63/85:0.0366110915)87.9/31:0.0446924961)84.7/56:0.0294749859,((((((((WP_072282603.1-Syntrophotalea-acetylenivorans:0.1647307238,(WP_092056970.1-Desulfuromonas-acetexigens:0.1847558215,((WP_028317018.1-Desulfobulbus-elongatus:0.4875466392,WP_043770486.1-Desulfobulbus-propionicus:0.2695190476)63.8/67:0.0690310792,WP_148894253.1-Geothermobacter-ehrlichii:0.4362917779)91.1/68:0.1024929666)84.5/67:0.0645887416)63.1/67:0.0193533898,WP_155876508.1-Desulfuromonas-sp.-AOP6:0.4592559656)98.1/100:0.1515952527,WP_053550565.1-Desulfuromonas-soudanensis:0.3430246595)81.7/95:0.0657014671,WP_027714505.1-Desulfuromonas-sp.-TF:0.3409479906)71.8/90:0.0283773424,WP_072282412.1-Syntrophotalea-acetylenivorans:0.3061237659)79.5/95:0.0320935696,((((((((((((((WP_089129802.1-Tolypothrix-sp.-NIES-4075:0.1006581459,WP_096563566.1-Scytonema-sp.-NIES-4073:0.1506820544)81.7/95:0.0550156803,WP_048866388.1-Scytonema-tolypothrichoides:0.1123118764)51.4/94:0.0325332735,WP_096563464.1-Scytonema-sp.-NIES-4073:0.1177709061)99.3/100:0.2157574300,WP_096557790.1-Nostoc-sp.-NIES-4103:0.1974455169)90.4/95:0.0527554896,WP_016866333.1-Fischerella:0.0916684795)75.4/55:0.0139805193,(((WP_015128325.1-Calothrix-sp.-PCC-7507:0.1555751825,WP_017653186.1-Fortiea-contorta:0.2246571298)18.9/100:0.0547101544,((WP_193199005.1-Nostoc-sp.-MG11:0.0605094894,WP_162396505.1-Nostoc-sp.-B-2019:0.0561090222)95.9/100:0.0859980862,WP_214439445.1-Atlanticothrix-silvestris:0.1730465160)64.5/100:0.0658979405)82.1/99:0.0374217924,(((WP_045870014.1-unclassified-Tolypothrix:0.0342887555,WP_190656031.1-Nostoc-linckia:0.0439628742)98.4/100:0.0928546951,(WP_190743070.1-Tolypothrix-sp.-FACHB-123:0.0604097395,WP_190545010.1-Calothrix:0.0404006584)97.9/100:0.0924972410)36.6/100:0.0425429871,WP_086758595.1-Nostoc-sp.-106C:0.0826956438)63.3/99:0.0417811516)86/75:0.0410369661)85.7/76:0.0379033575,((((WP_009344618.1-Cylindrospermopsis-raciborskii:0.0946959539,WP_240868223.1-Cylindrospermopsis-raciborskii:0.0250927256)99.9/100:0.1683437122,WP_096570274.1-Sphaerospermopsis:0.0666679167)12.2/100:0.0127388834,WP_220611028.1-Sphaerospermopsis-torques-reginae:0.0658956243)98.1/100:0.1083453900,((WP_237989437.1-Nostoc-sp.-UHCC-0870:0.1033774701,WP_190559626.1-Anabaena-sphaerica:0.1177036488)0/45:0.0268431376,WP_206263219.1-Nostoc-sp.-UHCC-0702:0.1077576840)90.6/100:0.0619587852)91.8/99:0.0514932626)97.5/100:0.1269179412,(WP_015193936.1-Stanieria-cyanosphaera:0.0492669037,WP_096384422.1-Stanieria-sp.-NIES-3757:0.0957309424)100/100:0.2469079366)89.2/99:0.0497862159,WP_015143663.1-Pleurocapsa-sp.-PCC-7327:0.1502518397)93.1/99:0.0729829667,WP_013324143.1-Gloeothece-verrucosa:0.1310004437)89.6/98:0.0575453144,(WP_106909014.1-Microcystis-sp.-MC19:0.3286310427,WP_124977990.1-Aphanothece-sacrum:0.2799012834)0/21:0.0153980566)65.5/56:0.0115649979,(((((WP_106232846.1-Pleurocapsa-sp.-CCALA-161:0.2116480472,WP_229639079.1-Waterburya-agarophytonicola:0.0382392015)81.7/100:0.0418410220,WP_096721927.1-Chondrocystis-sp.-NIES-4102:0.1085194337)97/100:0.1461632365,WP_006510035.1-Xenococcus-sp.-PCC-7305:0.2412069420)95.9/100:0.1386853176,WP_264322079.1-Zarconia-navalis:0.2188417914)86.7/52:0.0587881329,(WP_012595695.1-Rippkaea-orientalis:0.2586716735,(WP_007308862.1-Crocosphaera-watsonii:0.1027033788,((WP_008278255.1-Crocosphaera-chwakensis:0.0729560268,WP_107668138.1-Cyanothece-sp.-BG0011:0.0480658350)29.4/76:0.0085567662,WP_009545547.1-Crocosphaera-subtropica:0.0303585125)77.1/77:0.0228852163)97.3/100:0.1374285497)80.7/56:0.0425884516)85.2/91:0.0379322280)85.3/88:0.0304453549,((((((((((WP_102179481.1-Fischerella-thermalis:0.0264577962,((WP_096675774.1-Fischerella-sp.-NIES-4106:0.0614864046,WP_053458209.1-Hapalosiphon-sp.-MRB220:0.0399152918)68.5/100:0.0081832701,(WP_131119525.1-Westiellopsis-prolifica:0.0617299834,WP_016860528.1-Fischerella-muscicola:0.0324889086)84.9/100:0.0174950005)96.9/100:0.0951549077)100/100:0.2752240003,(WP_190825375.1-Iningainema-tapete:0.3066842512,(WP_015202791.1-Crinalium-epipsammum:0.2661376944,WP_035154029.1-Calothrix-sp.-336-3:0.4614309291)80/99:0.0628765725)58.1/84:0.0462751708)34.2/85:0.0114597801,((WP_099102643.1-Nostoc-sp.-Peltigera-malacea-cyanobiont-DB3992:0.1052611596,((WP_190950926.1-Nostoc-punctiforme:0.0575809626,(WP_221704991.1-Desmonostoc-muscorum:0.0581267823,WP_096537650.1-Nostoc-linckia:0.0218933756)72.4/95:0.0088703526)53.4/86:0.0299485915,WP_190959861.1-Nostoc:0.0239976236)97.9/97:0.0987657900)99.2/100:0.1366489285,((((((WP_193938462.1-Cuspidothrix-issatschenkoi:0.1802354629,WP_168537172.1-Anabaena-sp.-UHCC-0253:0.0494227672)89.8/80:0.0383823335,(((WP_190370804.1-Anabaena-sp.-FACHB-1237:0.1875504633,(WP_190392428.1-Dolichospermum-sp.-FACHB-1091:0.0158806473,(WP_028089231.1-Dolichospermum-circinale:0.0235885033,WP_137908967.1-Nostocales:0.0593024959)98.7/100:0.0777918349)98.6/100:0.1051376575)92.7/86:0.0463136019,((WP_096666215.1-Dolichospermum-compactum:0.0754911508,(WP_190387835.1-Aphanizomenon:0.0745216102,WP_168466156.1-Aphanizomenon:0.0230458796)43.2/98:0.0173093364)65.1/96:0.0191523394,WP_015080416.1-Anabaena-sp.-90:0.0409020520)93.9/96:0.0323236975)44.2/70:0.0075573939,WP_168502193.1-Anabaena-sp.-UHCC-0187:0.0075398064)73.2/70:0.0169122818)99.8/100:0.1511200245,(WP_010998622.1-Nostocaceae:0.0732501602,WP_011320078.1-Nostocaceae:0.0758089828)76.3/100:0.0229504217)94.7/95:0.0610053571,((WP_015138800.1-Nostoc-sp.-PCC-7524:0.0917252509,WP_066425233.1-Anabaena-sp.-4-3:0.1144020065)97.8/99:0.0868635970,(WP_199337214.1-Nostoc-sp.-FACHB-152:0.1442365284,(WP_239729951.1-Nodularia-sphaerocarpa:0.0817054223,WP_173403323.1-Nodularia-spumigena:0.0614110327)94.5/100:0.0569575517)86.1/88:0.0294099909)46.8/81:0.0094959879)84.4/91:0.0219411409,WP_199337786.1-Nostoc-sp.-FACHB-280:0.1627290837)95.9/100:0.0815349121,(WP_096624129.1-Calothrix-sp.-NIES-3974:0.0968633070,WP_095721999.1-Calothrix-elsteri:0.1158025671)98.7/100:0.1404157252)80/100:0.0277148187)96.5/100:0.0774049459)96.7/100:0.0847404684,(WP_194029353.1-Lusitaniella-coriacea:0.3013154195,WP_192216758.1-Microcoleus-sp.-FACHB-SPT15:0.2177029983)76.9/89:0.0308424642)92.6/85:0.0393263430,((((((WP_224410051.1-Oscillatoria-salina:0.2546470670,WP_250123015.1-Chroococcidiopsis-sp.-CCMEE-29:0.1955057920)91.1/97:0.0906366849,WP_190531554.1-Coleofasciculus-sp.-FACHB-542:0.2259126927)25.3/94:0.0147128457,WP_073549891.1-Chroogloeocystis-siderophila:0.2464025299)22.1/32:0.0280188186,((WP_233747769.1-Leptodesmis-sichuanensis:0.2863909746,WP_223049852.1-Leptothermofonsia-sichuanensis:0.1175452358)54.8/33:0.0112790594,WP_224345144.1-Kovacikia-minuta:0.1670622267)86.8/35:0.0511051739)78.9/86:0.0152503284,((WP_193873404.1-unclassified-Coleofasciculus:0.1757654994,WP_254010234.1-Symplocastrum-sp.-BBK-W-15:0.1573408676)81.7/92:0.0526463285,(WP_190666400.1-Coleofasciculus-sp.-FACHB-501:0.1404378426,WP_190525322.1-Coleofasciculus-sp.-FACHB-T130:0.0798894069)95.9/100:0.0858620679)34.5/61:0.0124723870)43.9/83:0.0117484891,WP_008178350.1-Moorena-producens:0.2970602874)77.7/86:0.0206566401)68.6/80:0.0138452117,WP_190493486.1-unclassified-Microcoleus:0.3498744605)73.2/79:0.0132824989,(WP_226578226.1-Microseira-wollei:0.2376451540,((WP_073595522.1-Phormidium-ambiguum:0.0852331316,WP_193982193.1-Phormidium-sp.-LEGE-05292:0.0912283941)99/100:0.1461778329,(WP_249064905.1-Argonema-antarcticum:0.0685647485,WP_190468950.1-unclassified-Planktothrix:0.1073825178)39.5/95:0.0135699825)86/90:0.0545731958)93.2/95:0.0658143195)77.8/69:0.0264700455,((((((((((WP_202221768.1-Okeania-sp.-KiyG1:0.0529763195,WP_094672483.1-Hydrocoleum-sp.-CS-953:0.0571050643)35.8/98:0.0167788141,WP_124146462.1-Okeania-hirsuta:0.0109759885)99.5/100:0.2049747359,WP_071516849.1-Geitlerinema-sp.-PCC-9228:0.2868491162)79/98:0.0892459464,WP_008052766.1-Limnospira-indica:0.3058962457)42/77:0.0462351535,(WP_009783380.1-Lyngbya:0.0927495917,WP_046277942.1-Limnoraphis-robusta:0.0612955571)95.4/100:0.0840519060)33.8/78:0.0487504437,(WP_083623900.1-Planktothrix-serta:0.0358088835,(WP_072717475.1-Planktothrix:0.0287851056,(WP_079676747.1-Planktothrix-sp.-PCC-11201:0.0286742388,(WP_026797127.1-Planktothrix:0.0318256201,WP_214559879.1-Planktothrix-agardhii:0.0509337820)58.5/94:0.0345564342)99.2/100:0.1116568490)25.1/66:0.0185865441)97.3/100:0.1032183313)88.8/98:0.0566378642,WP_190877489.1-unclassified-Planktothricoides:0.1916858509)87.1/70:0.0486919559,(((WP_261197618.1-unclassified-Laspinema:0.0540831443,(WP_207087219.1-Phormidium-pseudopriestleyi:0.0465472987,WP_261204505.1-Laspinema-sp.-D2d:0.0466206241)73.1/100:0.0214418995)88.4/100:0.0821699565,WP_006529211.1-Gloeocapsa-sp.-PCC-73106:0.4559812901)31.5/64:0.0425309320,WP_200665545.1-Oxynema-sp.-CENA135:0.2831132322)0/25:0.0404192419)70.3/62:0.0410617740,(WP_017715496.1-Oscillatoria-sp.-PCC-10802:0.1993029412,(WP_190665495.1-Microcoleus-sp.-FACHB-672:0.0747493391,WP_261220767.1-Ancylothrix-sp.-D3o:0.2393461366)95.2/100:0.0814485868)82/99:0.0537181259)43.4/56:0.0162452449,(WP_007355666.1-Kamptonema:0.1211393161,(((((WP_236507253.1-Tychonema-sp.-BBK16:0.0494815595,WP_096828456.1-Tychonema-bourrellyi:0.0822679648)93.6/100:0.0434591742,WP_242724542.1-Microcoleus-vaginatus:0.0401022364)83.8/100:0.0163257226,WP_015174083.1-Oscillatoria-nigro-viridis:0.0237668076)87.1/100:0.0256501462,(WP_194009709.1-unclassified-Tychonema:0.0451657238,WP_265235823.1-Lyngbya-sp.-CCAP-1446-10:0.0522437232)88.5/100:0.0247808182)0/25:0.0113400472,WP_193973976.1-Microcoleus-sp.-LEGE-07076:0.0506436853)99/100:0.1350609309)95.2/82:0.0724840717)39/81:0.0110184854)91.7/90:0.0657659128,WP_069965800.1-unclassified-Desertifilum:0.3461825231)79.4/84:0.0245840649,(((((WP_204137747.1-Halomicronema-sp.-CCY15110:0.0619671075,WP_039724568.1-Leptolyngbya-sp.-LK:0.0952918677)99.7/100:0.2021755707,WP_023076112.1-Leptolyngbya-sp.-Heron-Island-J:0.3283861897)66.5/99:0.1258881339,WP_017324474.1-Synechococcus-sp.-PCC-7336:0.3802561585)85.5/98:0.0560054168,WP_190772446.1-unclassified-Leptolyngbya:0.2349926680)83.9/96:0.0590558648,(WP_017661508.1-Baaleninema-simplex:0.4029776432,WP_036478304.1-Myxosarcina-sp.-GI1:0.2843123476)77.7/87:0.0308894188)82.8/87:0.0268001919)89.3/93:0.0383612299)99.2/100:0.1935625663,((((WP_205222281.1-Desulfogranum-marinum:0.3322736508,WP_205242362.1-Desulfobulbus-alkaliphilus:0.1720909710)90.8/91:0.0871923014,WP_267928740.1-Desulfolithobacter-dissulfuricans:0.1320894132)93.8/96:0.1106744597,WP_066347484.1-Geminocystis-sp.-NIES-3708:0.5920298276)56.4/75:0.0643658223,((WP_250680171.1-Geomonas-sp.-Red32:0.2456119395,WP_199381999.1-Geomesophilobacter-sediminis:0.2850890867)83/100:0.0624108648,WP_205218910.1-Desulforhopalus-vacuolatus:0.4304149291)86.9/99:0.0589564946)54.2/66:0.0203919288)90.7/88:0.0608854190)88.8/84:0.0588070610,(((WP_015405619.1-Desulfocapsa-sulfexigens:0.3258306991,(((WP_163339734.1-Desulfopila-sp.-IMCC35008:0.4055017705,WP_136805251.1-Desulfosediminicola-flagellatus:0.3204275190)73.6/99:0.0352301943,WP_073615262.1-Desulfopila-aestuarii:0.3121537523)94/100:0.1104508328,(WP_228856481.1-Desulfomarina-profundi:0.4234323528,WP_183349429.1-Desulfoprunum-benzoelyticum:0.2125825829)88.2/100:0.0776751302)68/77:0.0452259599)90.4/78:0.0485581218,(WP_028582690.1-Desulfogranum-japonicum:0.3836823264,WP_040200185.1-Geoalkalibacter-subterraneus:0.3755865966)81.7/36:0.0696780269)80.9/21:0.0304982074,WP_221248821.1-Desulfuromonas-versatilis:0.4968627675)0/12:0.0000020712)88.2/27:0.0440237693,(WP_137423090.1-Desulfoglaeba-alkanexedens:0.4306033819,WP_103114677.1-Geothermobacter-hydrogeniphilus:0.4249181445)81.9/32:0.0746068277)90.7/98:0.0691270975)82.9/29:0.0258585610,((((((((WP_212718705.1-unclassified-Synechocystis:0.0343595921,WP_228015966.1-Synechocystis-salina:0.1288323601)83.8/99:0.0428751136,WP_010873214.1-unclassified-Synechocystis:0.0441020856)100/100:0.5174985253,((WP_099797666.1-Thermostichus-lividus:0.5564717712,WP_196167318.1-Pseudomonas-monteilii:0.3771102604)88.7/86:0.1383804116,WP_048706771.1-Thauera-humireducens:0.3107434856)39.5/78:0.0409333878)88.7/91:0.0827025116,(WP_220807781.1-Noviherbaspirillum-aridicola:0.6449368802,WP_119020982.1-Halomonas-sp.-JS92-SW72:0.3266539547)95.4/89:0.2938272725)77/62:0.0146343353,((WP_084055900.1-Desulfacinum-hydrothermale:0.3830527690,WP_123289658.1-Desulfosoma-caldarium:0.3746185102)98.3/100:0.2498036207,(((WP_200222041.1-Achromobacter-xylosoxidans:0.1878528783,WP_259251071.1-Achromobacter-sp.-JUb104:0.2187880522)93.5/100:0.1004967854,(WP_088588627.1-Achromobacter-marplatensis:0.1908744626,WP_169273406.1-Achromobacter-sp.-Bel:0.1024358693)95/100:0.1083503498)85.4/94:0.0506799795,(((WP_111826915.1-Achromobacter-sp.-HZ01:0.1518381048,WP_102775120.1-Achromobacter:0.0240377619)99.7/100:0.1787188455,(WP_129525645.1-Achromobacter-agilis:0.0403911706,(WP_129240460.1-Achromobacter-veterisilvae:0.0540459157,WP_043544564.1-Achromobacter-sp.-RTa:0.0364230768)85.9/98:0.0218477938)71.1/97:0.0146044208)97.8/100:0.0990325552,((WP_175208175.1-Achromobacter-anxifer:0.0542433359,((WP_088158656.1-Achromobacter-xylosoxidans:0.1160622987,WP_175164188.1-Achromobacter-aegrifaciens:0.0455454368)70.4/56:0.0162369198,(((WP_251864322.1-Achromobacter-sp.-Marseille-Q4962:0.0899340195,WP_164793551.1-Achromobacter-insolitus:0.0404783157)97.8/89:0.0708493885,WP_207472676.1-Achromobacter-sp.-SD115:0.0141034062)89.6/90:0.0407046504,WP_125471167.1-Achromobacter:0.0237661569)93.5/98:0.0562312423)39.5/61:0.0283447771)98.9/100:0.1384767125,WP_174717299.1-Achromobacter-denitrificans:0.1058857634)0/68:0.0000022172)89.4/95:0.0451545781)98.6/100:0.2034346236)70.3/62:0.0207699421)94/94:0.0463531843,(((((((((((WP_238511579.1-Cupriavidus-gilardii:0.2826164712,(WP_226989104.1-Desulfuromonas-sp.-TF:0.5528167062,WP_227459030.1-Cupriavidus-pauculus:0.4507603935)93.1/100:0.1954542558)84.9/98:0.0906022728,WP_224042884.1-Cupriavidus-respiraculi:0.3805710856)44.2/93:0.0220363069,WP_213445339.1-Variovorax-sp.-PCZ-1:0.5080765394)79.8/60:0.0445147561,WP_116600700.1-Cupriavidus-alkaliphilus:0.2127954424)34.3/33:0.0090691301,(WP_018008101.1-Cupriavidus-neocaledonicus:0.0616197190,WP_183298446.1-Cupriavidus-alkaliphilus:0.0691130601)99.9/100:0.1947697316)92.6/85:0.0480232342,WP_250479419.1-unclassified-Caballeronia:0.5260205645)0/40:0.0000899052,WP_226921657.1-Kinneretia-sp.-XES5:0.4565900020)60/55:0.0185768961,((((((((((((WP_207000528.1-Trinickia-sp.-DHG64:0.1500840619,WP_124149903.1-Paraburkholderia-dinghuensis:0.1363962025)75.6/100:0.0559675772,(WP_266250364.1-Paraburkholderia-megapolitana:0.2012966707,WP_026121090.1-Paraburkholderia-kururiensis:0.1416652054)76.7/100:0.0758978698)97.5/100:0.1096263387,(((((((((((WP_059459219.1-Burkholderia-vietnamiensis:0.2940277883,WP_121276956.1-Trinickia-fusca:0.1420948537)63.6/92:0.0483183849,WP_134195781.1-Paraburkholderia-rhizosphaerae:0.3498391276)79.7/99:0.0290930910,WP_136892825.1-Trinickia-sp.-7GSK02:0.0507943987)78.5/91:0.0261856122,(WP_201650988.1-Paraburkholderia-sabiae:0.0515690473,WP_205970438.1-Paraburkholderia-sp.-Tr-20389:0.0474700005)19.3/88:0.0177434559)49.1/52:0.0188446567,WP_054928771.1-Paraburkholderia-caribensis:0.0413184670)78.6/74:0.0159495301,(WP_059889870.1-Burkholderia-stagnalis:0.2527469572,WP_206951398.1-Trinickia-sp.-4D114:0.1916503045)75.6/47:0.0246910333)82/56:0.0167803638,WP_209935850.1-Paraburkholderia-sp.-LEh10:0.1262490889)86.3/47:0.0254671872,((WP_168792792.1-Paraburkholderia-aromaticivorans:0.0983148892,(WP_004544228.1-Burkholderia:0.0615786443,WP_059583574.1-pseudomallei-group:0.0401829998)78.6/99:0.0256909540)85.3/99:0.0378983620,(((WP_175791145.1-Burkholderia-ambifaria:0.1208336190,WP_059842355.1-Burkholderia-multivorans:0.0445167733)77.2/99:0.0142732434,WP_105772332.1-Burkholderia-multivorans:0.0425496595)90.3/99:0.0291230036,(WP_006496666.1-Burkholderia-cenocepacia:0.0323751785,WP_059716874.1-Burkholderia-ubonensis:0.0484948147)91.7/94:0.0340346624)0/14:0.0000021744)82.3/21:0.0170054294)72.7/13:0.0112497524,(WP_060156927.1-Burkholderia-ubonensis:0.1337101715,(WP_061168975.1-Caballeronia-hypogeia:0.0736232782,WP_069272519.1-Burkholderia-cepacia:0.3020589268)88.5/93:0.0436175551)75.4/6:0.0083888104)43.6/29:0.0170801636,(WP_060092900.1-Burkholderia-vietnamiensis:0.0301381861,WP_226113708.1-Burkholderia-sp.-AU33545:0.0507000806)100/100:0.1942305184)95.6/98:0.0620879613,((WP_201138986.1-Paraburkholderia-domus:0.0552387508,WP_266262001.1-Paraburkholderia-madseniana:0.0287297717)99.1/100:0.1252825575,WP_009889585.1-Burkholderia-thailandensis:0.1166549743)79.5/99:0.0178574406)92.7/99:0.0605461208)71/96:0.0294080595,WP_205976236.1-Paraburkholderia-sp.-Se-20369:0.2967780051)87.8/98:0.0437137792,WP_179404315.1-Burkholderia-guangdongensis:0.2274117824)51.3/86:0.0205504433,WP_038711716.1-Burkholderia-sp.-lig30:0.3274295389)33.2/33:0.0677666872,((WP_063494936.1-Paraburkholderia-phytofirmans:0.0854377208,WP_254318368.1-Paraburkholderia-sp.-CNPSo-3272:0.0070566858)60.8/34:0.0443976998,WP_042262232.1-Paraburkholderia-heleia:0.0297974425)99.8/100:0.1777695553)56.5/98:0.0433723302,WP_056136218.1-Massilia-sp.-Root133:0.2822977887)95/86:0.0843129257,WP_135702508.1-Cupriavidus-oxalaticus:0.2043576189)80.8/81:0.0205102769,((((WP_108647113.1-Polynucleobacter-rarus:0.0161924256,WP_084281813.1-Polynucleobacter-kasalickyi:0.0633816658)100/100:0.7058287473,(((WP_215371371.1-Polynucleobacter-sp.-MWH-UH24A:0.2275354992,WP_173942040.1-Polynucleobacter-antarcticus:0.2244916535)97.8/100:0.1534315802,((WP_216254756.1-Polynucleobacter-sp.-AP-Melu-500A-A1:0.0674081956,WP_216218975.1-Polynucleobacter-sp.-AP-Latsch-80-C2:0.0470504125)99.1/100:0.1171399319,(((((WP_215319389.1-Polynucleobacter-paneuropaeus:0.1172871436,(WP_215352354.1-Polynucleobacter-sp.-MWH-P3-07-1:0.0533970192,WP_215380622.1-Polynucleobacter-paludilacus:0.0457578705)35.6/100:0.0410127714)96.2/100:0.0746807344,WP_062307935.1-Polynucleobacter-sinensis:0.0643165800)81.7/75:0.0209546507,WP_216197047.1-Polynucleobacter-sp.-UK-Mo-2m-Kol15:0.0843407460)75/69:0.0080961143,((((((WP_215376847.1-Polynucleobacter-asymbioticus:0.0306453246,WP_215328794.1-Polynucleobacter-sp.-AP-Sving-400A-A2:0.0636850909)97.9/100:0.0661432519,WP_173959551.1-Polynucleobacter-arcticus:0.0357060370)92.7/100:0.0297107354,WP_216225867.1-Polynucleobacter-sp.-MG-27-Goln-C1:0.1017720246)39/100:0.0076189167,(WP_216168716.1-Polynucleobacter-bastaniensis:0.0989791316,WP_216178894.1-Polynucleobacter-sp.-80A-SIGWE:0.0161566764)86.5/100:0.0203715017)88.5/100:0.0300870429,WP_215299358.1-Polynucleobacter-sp.-AP-Nino-20-G2:0.1072657210)87.3/84:0.0253238739,(WP_216268685.1-Polynucleobacter-sp.-Nonnen-W13:0.0589902118,(WP_215334312.1-Polynucleobacter-sp.-AP-Ainpum-60-G11:0.0549384818,WP_251374783.1-Polynucleobacter-corsicus:0.0425364677)43.9/64:0.0069338264)82.7/74:0.0137285131)66.6/71:0.0070795547)89.4/83:0.0341311872,WP_076024025.1-Polynucleobacter-sphagniphilus:0.1097184113)72.4/79:0.0111568081)95.2/90:0.0686661199)52.3/70:0.0341117298,(((WP_128113711.1-Polynucleobacter-necessarius:0.1204676693,WP_215385554.1-Polynucleobacter-sp.-AP-Kolm-20A-A1:0.0876032765)94.5/100:0.0905076973,WP_216177372.1-Polynucleobacter-sp.-AP-Feld-500C-C5:0.2064467743)81.4/94:0.0392665427,WP_114639791.1-Polynucleobacter-necessarius:0.1383144440)62.2/97:0.0491873510)97.4/99:0.3160392439)98.5/96:0.3982434794,(WP_115680924.1-Cupriavidus-taiwanensis:0.2867375057,(WP_071014919.1-Cupriavidus-sp.-USMAHM13:0.2554333092,WP_174426450.1-Cupriavidus-basilensis:0.4933531257)83.6/62:0.0777925108)76.6/59:0.0362009271)80.9/66:0.0306478101,WP_018313502.1-Cupriavidus-sp.-UYPR2.512:0.3570197862)0/50:0.0000028594)85.1/83:0.0454339187,WP_228215938.1-Aromatoleum-toluclasticum:0.3175237854)83.9/75:0.0352324089,WP_150984056.1-Cupriavidus-basilensis:0.5937958927)75.8/61:0.0219158134)92.9/91:0.0734983409,((WP_250024752.1-Cupriavidus-campinensis:0.2004909562,WP_094797980.1-Bordetella-genomosp.-5:0.2806330168)88.6/97:0.0887160715,WP_227181580.1-Leeia-sp.-TBRC-13508:0.4420794160)37.3/81:0.0600356699)33.2/68:0.0191494771,WP_136680209.1-Neptunomonas-sp.-XY-337:0.2689865513)90.4/83:0.0612302586,((((((((((((WP_209862330.1-Pseudomonas-sp.-BP8:0.6545605959,(WP_054891693.1-Pseudomonas:0.0981927330,WP_236238170.1-Pseudomonas-faucium:0.1917771824)6/68:0.0330379720)60.7/84:0.0194369914,(WP_158461286.1-Pseudomonas-fluorescens:0.3003196309,(WP_200141208.1-unclassified-Pseudomonas:0.2496443604,WP_029612831.1-Pseudomonas-parafulva:0.0231994026)94.2/100:0.0811429237)79.3/100:0.0521551877)90/59:0.0548809389,(((WP_217887155.1-Pseudomonas-xanthosomae:0.0418013654,WP_217840807.1-Pseudomonas-fakonensis:0.0506923209)68.2/100:0.0133746350,WP_186654015.1-Pseudomonas-xantholysinigenes:0.2805222430)80.5/98:0.0481222051,(WP_201176141.1-Pseudomonas-sp.-S31:0.0240589529,WP_261240966.1-unclassified-Pseudomonas:0.0673650777)99.7/100:0.1413008171)86.4/82:0.0221789177)70.4/48:0.0217549651,WP_225786622.1-Pseudomonas:0.1090567520)60.1/66:0.0396916536,WP_110605176.1-Pseudomonas-soli:0.3315649619)95.9/100:0.0823948660,((((WP_181290220.1-Pseudomonas-brassicacearum:0.2204556151,WP_249678359.1-Pseudomonas-sp.-PIA16:0.1956392366)71.8/64:0.0162586959,WP_043192328.1-Pseudomonas-rhizosphaerae:0.2943172363)84.5/64:0.0317793611,WP_268799018.1-Pseudomonas-sp.-BSw22131:0.2227864994)84.4/92:0.0446475690,((WP_201004194.1-Pseudomonas-cichorii:0.0796439642,WP_176569640.1-Pseudomonas-eucalypticola:0.1804549507)90.7/96:0.0672108099,(WP_219063768.1-Pseudomonas-sp.-UMAB-08:0.2558891348,(WP_222540780.1-Pseudomonas-sp.-DR208:0.1556743465,(((WP_065875813.1-Pseudomonas:0.0553010627,(WP_033898696.1-Pseudomonas-fluorescens-group:0.0432909408,(WP_186687394.1-Pseudomonas-sp.-SWRI18:0.0662534049,WP_169378974.1-Pseudomonas:0.0197202851)84.2/100:0.0321285757)22.7/100:0.0136155397)99/100:0.0816423050,(WP_253509974.1-Pseudomonas-rhodesiae:0.1464754126,WP_093202821.1-Pseudomonas-yamanorum:0.0368159320)87/100:0.0444791352)30.5/97:0.0165810020,WP_105697656.1-Pseudomonas-poae:0.0646167108)0/97:0.0000025529)64.3/100:0.0485072100)89.6/100:0.0520276326)68.6/95:0.0319892659)97.4/100:0.1227994162)95/98:0.0682396702,WP_252273535.1-Pseudomonas-sp.-YIM-B01952:0.1325788434)88.9/98:0.0389453953,WP_143488038.1-Pseudomonas-mangiferae:0.1703654213)83.7/98:0.0324700333,(((WP_058067682.1-Pseudomonas-sp.-TTU2014-080ASC:0.1803395482,(WP_192399936.1-Pseudomonas-sp.-PDM15:0.1852976769,WP_226282979.1-Pseudomonas-lalkuanensis:0.0950575405)88.5/100:0.0586515381)91.5/58:0.0513167782,(WP_003453405.1-Pseudomonas:0.0939201248,WP_025281494.1-Ectothiorhodospira-haloalkaliphila:0.5955986874)80.4/91:0.0681584223)75.4/51:0.0155366095,((((WP_077525673.1-Pseudomonas-resinovorans:0.1358162364,(WP_137821430.1-Pseudomonas-sp.-D-2018:0.0910480645,(WP_153584563.1-Caenimonas-koreensis:0.3398363201,(WP_007185580.1-Hydrocarboniphaga-effusa:0.0735083228,WP_007185565.1-Hydrocarboniphaga-effusa:0.1093309189)99.3/100:0.2297843803)99/100:0.2550554406)46.4/66:0.0206270457)96/83:0.0817617106,(WP_016495240.1-Pseudomonas-resinovorans:0.0791845785,WP_044873077.1-Pseudomonas-sp.-LFM046:0.0706056033)69.2/98:0.0325468704)79.1/17:0.0175620151,(WP_151138079.1-Pseudomonas-lalkuanensis:0.0663562946,WP_028626762.1-Pseudomonas-resinovorans:0.0431879119)80.1/16:0.0211594566)74.4/12:0.0089305546,WP_120654068.1-Pseudomonas-sp.-DY-1:0.1203177687)65.8/49:0.0079324106)91.8/93:0.0527955601)85.5/93:0.0342346099,((((((WP_090448460.1-Pseudomonas-benzenivorans:0.2383160132,(WP_108096457.1-Pseudomonas-sp.-GV071:0.1910219724,(WP_233685494.1-Pseudomonas-anguilliseptica:0.0684412115,WP_257778987.1-Pseudomonas-sp.-32.2.56:0.0065679376)99.3/100:0.1303609846)79.4/91:0.0306072485)77.2/91:0.0107881689,((WP_025164100.1-Pseudomonas-taeanensis:0.0747715627,WP_147170838.1-Pseudomonas-sp.-SJZ079:0.0040665912)93.3/100:0.0635749445,(WP_208707982.1-Pseudomonas-sp.-LPB0260:0.0314047583,WP_255840290.1-Pseudomonas-benzenivorans:0.0505403703)97.1/100:0.0809897706)80.1/95:0.0220936035)87.9/94:0.0296888017,WP_069518708.1-Pseudomonas:0.1661983887)84.4/87:0.0340460560,((WP_131198269.1-Pseudomonas-dryadis:0.1250633914,(WP_131178052.1-Pseudomonas-daroniae:0.0766418940,(WP_084307530.1-Pseudomonas-flavescens:0.0910637063,(WP_249909557.1-Pseudomonas-fulva:0.0573997141,(WP_074889242.1-Pseudomonas-argentinensis:0.0337481763,WP_027910132.1-Pseudomonas-sp.-URMO17WK12-I4:0.0549291156)38.9/92:0.0179299404)82.6/100:0.0342321438)83.3/93:0.0426481862)95/100:0.0883500094)97.1/100:0.0949377725,(((WP_119694090.1-Pseudomonas:0.0654205681,WP_106738315.1-Pseudomonas-mendocina:0.0569977009)89.6/100:0.0442921576,WP_150606948.1-Pseudomonas-oleovorans:0.0682833547)92.7/83:0.0531685200,(WP_159970704.1-Pseudomonas-sp.-8Z:0.1035679791,(WP_021444205.1-Pseudomonas-sp.-EGD-AK9:0.0619262618,((WP_024309956.1-Pseudomonas-sp.-P818:0.0665863756,(WP_143497094.1-Pseudomonas-sp.-ALS1131:0.0485281211,WP_090337209.1-Pseudomonas-chengduensis:0.0250588579)0/48:0.0000029559)84.3/66:0.0208649662,WP_074939507.1-Pseudomonas-composti:0.0257922022)87.3/49:0.0237183270)83.4/17:0.0222199424)72.9/24:0.0103831696)93/99:0.0561798978)39/91:0.0181773838)70.9/85:0.0134056770,((WP_047193087.1-Schlegelella-brevitalea:0.3979617111,WP_019561187.1-Caldimonas-manganoxidans:0.2286037041)96.3/99:0.2027394711,WP_074865046.1-Atopomonas-hussainii:0.1590097844)99.7/99:0.2261436264)37.2/48:0.0171361964,WP_238040454.1-Pseudomonas-sp.-MMS21-TM103:0.1705568969)77.2/59:0.0349403444)91.7/99:0.0585119177,(((WP_116346290.1-Cupriavidus-taiwanensis:0.1248622834,WP_124685047.1-Cupriavidus-pauculus:0.3224530469)100/100:0.6313652803,WP_220803742.1-Pseudomonas-sp.-NCCP-436:0.1042623479)82.3/84:0.0887699271,WP_090225506.1-Pseudomonas-kuykendallii:0.1471680838)46.6/79:0.0481703818)87.4/98:0.0527364498,((WP_183165579.1-Azomonas-macrocytogenes:0.2154449292,((((WP_102836461.1-Stutzerimonas-stutzeri:0.0341800018,(WP_256071125.1-Stutzerimonas-stutzeri:0.0425074651,WP_045423249.1-Pseudomonadaceae:0.0342492621)82.6/97:0.0167203691)96.9/98:0.0646957163,WP_255961233.1-Stutzerimonas-stutzeri:0.0346521073)65.1/86:0.0043044928,WP_025243649.1-Stutzerimonas-stutzeri:0.0580972529)95.5/99:0.0712746267,(WP_122077451.1-Pseudomonadaceae:0.0415565838,((WP_037024448.1-Pseudomonas-saudiphocaensis:0.0831864982,WP_181084609.1-Stutzerimonas-stutzeri:0.0085329850)91/100:0.0346681402,(WP_011911332.1-Stutzerimonas-stutzeri:0.0504670097,(((WP_230358311.1-Pseudomonas-sp.-NP21570:0.0204678875,(WP_193680295.1-Pseudomonas-lopnurensis:0.0567933655,WP_200589398.1-Stutzerimonas-stutzeri:0.0278035324)49.5/100:0.0416080099)89.7/91:0.0325706927,(WP_237255234.1-Pseudomonas-oligotrophica:0.0854825726,WP_043295157.1-Stutzerimonas-degradans:0.0234383817)94.4/100:0.0522831533)77/82:0.0087585280,WP_120997453.1-Pseudomonas:0.0419176401)88.7/91:0.0273685703)87.8/97:0.0274715113)77.4/74:0.0172956857)0/73:0.0000020692)81.2/98:0.0480098026)51.5/99:0.0486994003,WP_028238613.1-Stutzerimonas-azotifigens:0.0801704458)99.3/99:0.1521220697)58.1/66:0.0560428898)71/60:0.0128852842)82.8/56:0.0309173076,((WP_199264031.1-Desulfobulbus-oligotrophicus:0.3237840302,WP_226440716.1-Quatrionicoccus-australiensis:0.3646648895)28.6/71:0.0916365090,WP_091467897.1-Methylophilus-rhizosphaerae:0.4562268107)86.4/85:0.1022132679)73.9/84:0.0400390996,((((((WP_198111724.1-Inhella-proteolytica:0.5142452878,(WP_241396682.1-Pelomonas-sp.-CA6:0.2084974620,(WP_028996355.1-Azohydromonas-australica:0.0938186384,WP_066337818.1-Azohydromonas-lata:0.1150663228)98.9/100:0.1726295475)9.1/40:0.0805811497)63.4/67:0.0762571776,WP_037482909.1-Sphaerotilus-natans:0.1680630900)97.8/99:0.2388432759,WP_160809027.1-Nitrosomonas-sp.-HPC101:0.4841928793)88.9/69:0.0985567408,(WP_253536546.1-Ralstonia-sp.-21YRMH01-3:0.4662110678,WP_154206467.1-Ralstonia-pickettii:0.2246669450)76.3/68:0.0388719254)90/67:0.0665313333,(((KAI6244242.1-Aphelenchoides-fujianensis:0.6659952426,KAI6170547.1-Aphelenchoides-bicaudatus:0.9437799508)93.1/100:0.2667768289,WP_110520589.1-Herpetosiphon-llansteffanensis:0.7996534027)94.8/93:0.2913880184,(WP_234070476.1-Myxococcus-stipitatus:0.3287728312,WP_120555541.1-Corallococcus-aberystwythensis:0.3199066331)98.1/100:0.4278772724)93.2/92:0.2065412218)89.6/86:0.0927434166,((((WP_036252969.1-Methylobacter-sp.-BBA5.1:0.2881465703,((((WP_019918777.1-Methyloversatilis-discipulorum:0.0543189385,WP_018229738.1-Methyloversatilis-universalis:0.0730125383)95.2/100:0.1318653824,WP_183409942.1-Litorivivens-lipolytica:0.5965459132)37/89:0.0684812835,(WP_101941536.1-Uliginosibacterium-sp.-TH139:0.1196794219,WP_095523004.1-Candidatus-Dactylopiibacterium-carminicum:0.1313666324)97.4/100:0.1865762246)93/100:0.1238697598,WP_101249932.1-Telmatospirillum-siberiense:0.5943447134)89.6/99:0.0906248945)89.8/97:0.0558410239,WP_220807643.1-Noviherbaspirillum-aridicola:0.4875156003)76.3/91:0.0157885305,((((WP_109041087.1-Azospira-sp.-I13:0.1405148735,WP_014237023.1-Azospira-oryzae:0.1541459049)97.3/100:0.1715443268,WP_067106371.1-Methylobacillus-sp.-MM3:0.2836212159)37.4/95:0.0601631214,WP_201673630.1-Ramlibacter-monticola:0.2880864601)77.9/97:0.0391423796,WP_212785890.1-Ferrigenium-kumadai:0.2734464265)91.4/95:0.0741820523)72.1/81:0.0159247135,(((((((WP_074796726.1-Nitrosospira-briensis:0.0862973083,(WP_090882638.1-Nitrosovibrio-sp.-Nv6:0.0721392174,WP_097065525.1-Nitrosovibrio-sp.-Nv4:0.0366220161)88.5/100:0.0685421558)99.9/100:0.2940270086,(WP_106935816.1-unclassified-Variovorax:0.3176425237,WP_122244186.1-Vandammella-animalimorsus:0.2728301316)87.4/99:0.0633686911)91.1/96:0.0558057207,(((WP_223986959.1-Cupriavidus-pampae:0.0730722416,((WP_011517306.1-Cupriavidus-metallidurans:0.0624605731,WP_019451253.1-Cupriavidus-sp.-BIS7:0.0647828031)0/80:0.0000021535,WP_082079449.1-Cupriavidus-basilensis:0.1012077394)80.2/84:0.0317356002)99.5/100:0.2001265611,WP_169420802.1-Ramlibacter-agri:0.2739442659)30.3/64:0.0250711534,WP_137918285.1-Hydrogenophaga-sp.-2FB:0.1981795707)36.4/31:0.0235954037)75.3/44:0.0152812089,(WP_026436726.1-Acidovorax-sp.-JHL-9:0.2576307356,((((((((WP_169107474.1-Comamonas-suwonensis:0.0557433285,WP_219343559.1-Comamonas-sp.-Y33R10-2:0.1065201897)93.1/100:0.0630630390,(WP_003073420.1-Comamonas:0.0850591519,((WP_211457163.1-Comamonas-sp.-PE63:0.0608549435,WP_148729134.1-Comamonas-sp.-Z3:0.0526854266)50.3/94:0.0080790847,WP_034391098.1-Comamonas-thiooxydans:0.0198996318)74.4/94:0.0104686773)87.7/100:0.0332466218)64.9/100:0.0193355512,WP_218241177.1-Comamonas-fluminis:0.1399209386)99.1/100:0.2032978027,WP_066475960.1-Comamonas-terrae:0.2764658889)28.3/25:0.0477994506,WP_012203208.1-Delftia-acidovorans:0.2593552996)24.1/21:0.0211823745,WP_058879441.1-Comamonas-kerstersii:0.1456179262)99.7/100:0.2607433187,WP_166067558.1-Diaphorobacter-sp.-HDW4B:0.2160651340)83.5/99:0.0651170005,WP_086926522.1-Acidovorax-carolinensis:0.2164299053)35.7/91:0.0458176820)91.6/96:0.0576206656)80/81:0.0216848353,WP_174484170.1-methanotrophic-endosymbiont-of-Bathymodiolus-puteoserpentis:0.3026785433)0/48:0.0000025588,(((WP_219210863.1-Variovorax-boronicumulans:0.2310376610,((((WP_126747644.1-Variovorax-sp.-DXTD-1:0.1794789870,WP_068630051.1-Variovorax-sp.-PAMC-28711:0.3355426709)0/53:0.0000029647,(WP_093207264.1-Variovorax-sp.-YR750:0.1444950149,WP_206177103.1-Variovorax-sp.-RKNM96:0.1297674993)58.3/92:0.0506270506)90.6/75:0.0725923301,(WP_093441791.1-Variovorax-sp.-770b2:0.1572103391,WP_160553990.1-Xylophilus-rhododendri:0.2482828416)23.6/61:0.0585207993)57.3/62:0.0387640211,WP_192194832.1-Variovorax-sp.-38R:0.2741058644)86.7/99:0.0490725240)18/88:0.0409767149,WP_056898839.1-Pseudorhodoferax-sp.-Leaf274:0.3697920895)89.4/73:0.0447293533,WP_137890779.1-Ramlibacter-sp.-2FC:0.1522142637)63/25:0.0104658887)84.8/30:0.0225186818,WP_057673948.1-Curvibacter-sp.-PAE-UM:0.2553697872)87.8/86:0.0433474432)69.5/95:0.0271752876)97.6/96:0.0884211062)72/18:0.0126143451)82.4/26:0.0176269732,(((((((WP_015170385.1-Geitlerinema-sp.-PCC-7407:0.2935042897,WP_123210144.1-Stagnimonas-aquatica:1.0828829713)29.8/76:0.1162631593,WP_204103763.1-Spirulina-major:0.5704839137)13.4/67:0.0371560846,(WP_002810067.1-Nitrosococcus-oceani:0.0547826530,WP_013221233.1-Nitrosococcus-watsonii:0.0902288232)100/100:0.3087171344)73.5/69:0.0323176125,WP_258826205.1-Massilia-agri:0.4666670992)76.3/84:0.0310872506,(WP_040097704.1-Geoalkalibacter-ferrihydriticus:0.3354828973,WP_066727622.1-Desulfuromonas-sp.-DDH964:0.3327702690)88.8/67:0.0808997906)0/35:0.0499938232,WP_054772520.1-Methylogaea-oryzae:0.3118452594)63.1/35:0.0201785287,((((WP_014706296.1-Methylophaga-nitratireducenticrescens:0.0000023571,WP_136552334.1-Methylophaga-sp.-SB9B:0.0920176546)100/100:0.5611954689,((WP_148717329.1-Chitinolyticbacter-meiyuanensis:0.8566576879,(WP_046292890.1-Pandoraea-oxalativorans:0.1220242567,WP_150719538.1-Pandoraea-capi:0.1015243403)99.9/100:0.4507576789)82.8/97:0.0941568805,WP_216129682.1-Azospira-inquinata:0.4281697719)55.2/94:0.0642482720)24.9/75:0.0249046985,(WP_108623287.1-Orrella-marina:0.4910007502,WP_243478771.1-Orrella-sp.-f23:0.4652651726)92.7/100:0.1874511036)91.2/84:0.0700291668,(((WP_205202385.1-Azonexus-hydrophilus:0.3592290660,WP_173581083.1-Azonexus-fungiphilus:0.2757725356)92.4/98:0.1034514618,((((WP_136386324.1-Pseudothauera-rhizosphaerae:0.2612590576,WP_187718084.1-Thauera-sp.-CAU-1555:0.1701513995)73.2/85:0.0526948987,WP_114649035.1-Pseudothauera-hydrothermalis:0.0838841862)99.3/100:0.2242017281,((((((WP_148817865.1-Pigmentiphaga-aceris:0.4383698804,WP_004074545.1-Desulfobacter-postgatei:0.2922338486)85.2/99:0.0746246449,(WP_168987795.1-Azoarcus-taiwanensis:0.2982231481,WP_018865194.1-Thioalkalivibrio-sp.-ARh3:0.2384109785)82.1/93:0.0669290315)88.6/90:0.0503384899,(WP_256068944.1-Stutzerimonas-stutzeri:0.2578177350,(((((((((WP_060482506.1-Pseudomonas-sp.-NBRC-111119:0.2296675953,WP_159409308.1-Pseudomonas-putida:0.1091826484)91.7/93:0.0624036268,(((WP_182365203.1-Pseudomonas-juntendi:0.0872333281,WP_182365213.1-Pseudomonas:0.1260357861)78.7/99:0.0246815751,((WP_110607173.1-Pseudomonas-fulva:0.0160793357,WP_088523134.1-unclassified-Pseudomonas:0.0601338492)95.8/100:0.0467382230,(WP_167339957.1-Pseudomonas-fulva:0.0381272959,WP_186476580.1-Pseudomonas-promysalinigenes:0.0504321421)78.1/100:0.0245581640)77/98:0.0148715887)87.7/97:0.0199037382,(WP_201214098.1-Pseudomonas-sp.-S36:0.0250330308,WP_054890059.1-Pseudomonas:0.0750736205)96.8/100:0.0576289912)72.7/78:0.0068848898)76.2/67:0.0177208631,(WP_115272924.1-Pseudomonas-putida:0.1595795524,WP_267821885.1-Pseudomonas-putida:0.0627358151)80.8/86:0.0434042653)0/94:0.0000026050,(WP_167277242.1-Pantoea-sp.-Tr-811:0.0202476511,WP_103449615.1-Pseudomonas-putida:0.0688036532)97.8/100:0.0704003578)34.6/94:0.0092567805,((WP_196129978.1-Pseudomonas-fulva:0.1142467207,WP_201164743.1-Pseudomonas-sp.-S60:0.0352404884)98.8/100:0.1096079216,(((WP_174142546.1-Pseudomonas-putida:0.1699160288,WP_027907237.1-Pseudomonas:0.0461066745)71.9/100:0.0109662810,WP_075044077.1-Pseudomonas-putida:0.0571518575)77.1/100:0.0342400785,(WP_023379297.1-Pseudomonas-sp.-VLB120:0.1178200015,WP_202675963.1-Pseudomonas-sp.-SK2:0.0447150585)98.7/100:0.0899020687)91.1/100:0.0634730170)89.8/100:0.0396501056)94.5/99:0.0466369141,(WP_060394622.1-Pseudomonas-entomophila:0.1975471737,(WP_230064532.1-Pseudomonas-sp.-CIP-10:0.0262585652,WP_197863301.1-Pseudomonas-putida:0.0490488437)71.8/99:0.0465052597)0/75:0.0049811511)69/58:0.0099444979,WP_029885231.1-Pseudomonas:0.0362379235)86.1/76:0.0387598636,(WP_046489762.1-Pseudomonas-veronii:0.1250877578,(WP_059395284.1-Pseudomonas-putida:0.1262108855,(WP_009685980.1-Pseudomonas:0.0912212020,WP_024086460.1-Pseudomonas:0.0821483290)66/100:0.0421614609)89.3/100:0.0483146832)74.9/98:0.0504013698)98.3/100:0.1226189399,WP_183165449.1-Azomonas-macrocytogenes:0.1498285108)48.6/99:0.0336888713)44.4/39:0.0480015557)84.1/38:0.0328167041,((WP_238617884.1-Ectothiorhodospira-shaposhnikovii:0.1737894342,WP_025282248.1-Ectothiorhodospira-haloalkaliphila:0.1436030595)91.2/100:0.0818487005,WP_238622182.1-unclassified-Ectothiorhodospira:0.1838206028)82.3/99:0.0420503627)77.5/94:0.0955835510,WP_013164002.1-Desulfurivibrio-alkaliphilus:0.3643155639)19.4/20:0.0562066714,WP_026685886.1-Azovibrio-restrictus:0.1881656650)87.2/22:0.0788264716)92.7/89:0.1008789880,((((WP_167682126.1-Rhodocyclus-gracilis:0.3931903296,(WP_153115862.1-Rhodocyclus-tenuis:0.0557146758,WP_200385811.1-Rhodocyclus-tenuis:0.0271720932)92.7/100:0.0870280297)88.7/100:0.0752954806,WP_201212349.1-Rhodocyclus-purpureus:0.2140624896)96.1/100:0.1699562149,WP_153110537.1-Propionivibrio-limicola:0.3183943401)69.7/100:0.0366056326,WP_263768642.1-Propionivibrio-sp.-SG131:0.3527513281)79.6/88:0.0645401807)41.8/95:0.0200872664)38.5/88:0.0170533690,(((WP_121459656.1-Azonexus-fungiphilus:0.3452150259,WP_117174481.1-Rhodoferax-lacus:0.2845276096)76.1/97:0.0614620044,((WP_094482496.1-Rhodoferax-sp.-TH121:0.1022729046,WP_105262671.1-Rhodoferax-sp.-TS-BS-61-7:0.0505747204)90.7/100:0.0597129126,WP_214121631.1-Curvibacter-sp.-CHRR-16:0.3292031555)96.1/100:0.1429489538)73.1/97:0.0509416617,(WP_066884254.1-Dechloromonas:0.1303915366,WP_027458229.1-Dechloromonas-agitata:0.1422641579)96.8/100:0.1075866687)87.8/97:0.0847572544)91.1/94:0.0675329513)86.8/86:0.0301759567)87.1/59:0.0356047177)88.7/80:0.0436420941,(WP_176062241.1-Anaeromyxobacter-diazotrophicus:0.2280989552,WP_248361410.1-Anaeromyxobacter-oryzae:0.3204234154)99.9/100:0.3454409940)84.7/25:0.0346819207,((WP_044881819.1-Neochlamydia-sp.-EPS4:0.0753510674,WP_166155173.1-unclassified-Neochlamydia:0.0000021458)100/100:0.5773299731,((((WP_129151777.1-Achromobacter-aloeverae:0.1219013842,WP_088599355.1-unclassified-Achromobacter:0.0875445905)96.7/100:0.1124652946,WP_057654143.1-Bordetella-sp.-N:0.2380398163)76.2/100:0.0686278054,WP_086067427.1-Bordetella:0.2922572063)92.3/100:0.0942158678,(WP_066641528.1-Bordetella-sp.-H567:0.1253882957,(WP_086058993.1-Bordetella-genomosp.-9:0.2445916902,WP_094856450.1-Bordetella-genomosp.-10:0.1671223205)90.1/100:0.1075041181)84.1/100:0.0777021948)83.7/100:0.0816370569)86.1/96:0.0475336178)78.5/19:0.0381617032)91.5/73:0.1063278360,((WP_188630571.1-Oceanisphaera-marina:0.1640417162,WP_204795415.1-Oceanisphaera-litoralis:0.0622230554)7.7/77:0.0162328562,WP_104485062.1-Oceanisphaera-arctica:0.1275984590)100/100:0.2763603575)87.8/91:0.0661656976,(WP_189493202.1-Formosimonas-limnophila:0.3213117116,WP_133620003.1-Hydromonas-duriensis:0.3427491957)98.6/99:0.2538081733)46.5/85:0.0203147584,(WP_094838100.1-Bordetella-genomosp.-4:0.2433343415,WP_160351730.1-Bordetella-sp.-15P40C-2:0.2889073182)100/100:0.4382231400)76/84:0.0317840750)80.1/78:0.0303433004,((WP_151054911.1-Cellvibrio-sp.-KY-YJ-3:0.2618535330,WP_078044787.1-Cellvibrio-sp.-PSBB023:0.2971202947)78.9/97:0.0509546994,(WP_012489013.1-Cellvibrio-japonicus:0.2189538165,WP_087465811.1-Cellvibrio-sp.-PSBB006:0.2173105460)87.3/97:0.0560040566)99.6/100:0.2067559106)42.1/67:0.0181489889,(((WP_132257458.1-Paucimonas-lemoignei:0.0941470190,WP_119785517.1-Noviherbaspirillum-sedimenti:0.0746187728)100/100:0.4371290936,(WP_026470910.1-Alkanindiges-illinoisensis:0.3107122911,(WP_248147450.1-Moraxella-osloensis:0.4732036030,WP_092690500.1-Acinetobacter-kyonggiensis:0.1988695439)83.7/100:0.1449326515)98.9/100:0.3151445119)98.5/100:0.2408114483,(WP_208005626.1-Oceanisphaera-pacifica:0.1113712828,(WP_087034853.1-Oceanisphaera-profunda:0.0776220843,WP_087034873.1-Oceanisphaera-profunda:0.1503740661)84.5/100:0.0914773278)95.4/100:0.1672640720)74/78:0.0427190498)86.4/81:0.0474973883)89.5/83:0.0564554254,((WP_047682696.1-Xenorhabdus:0.0604421790,WP_099124588.1-Xenorhabdus-stockiae:0.1037413558)94.5/100:0.1469282413,(WP_074024273.1-Xenorhabdus-eapokensis:0.0909609677,WP_217470569.1-Photorhabdus-akhurstii:0.0871590528)93.6/100:0.1301888359)98.5/99:0.1622763880)79.9/99:0.0267045812,(((WP_035600513.1-Edwardsiella-tarda:0.3675316750,WP_125117341.1-Aeromonas-caviae:0.3737263033)12.5/72:0.0460399372,(WP_048221111.1-Citrobacter:0.1501890138,WP_263807164.1-Salmonella-enterica:0.2371583954)99.9/100:0.3255259539)77.1/100:0.0230997539,(WP_202782010.1-Enterobacter-asburiae:0.1120457566,WP_144817260.1-Pseudescherichia-sp.-L3:0.0349549180)99.9/100:0.2357377552)96.7/100:0.0894179153)87.1/98:0.0274329870,((WP_061279316.1-Kluyvera-cryocrescens:0.4351618941,WP_172410167.1-Escherichia-coli:0.4379639012)50.5/98:0.1096880105,(WP_105591414.1-Pantoea-coffeiphila:0.0949443697,WP_123334936.1-Erwinia-sp.-JUb26:0.0323416882)100/100:0.3090066179)85.7/96:0.0478624279)87/92:0.0249578519,(((((((((((((((((WP_085037002.1-Cronobacter-sakazakii:0.2637738561,WP_076770125.1-Kosakonia-cowanii:0.1817342513)95.2/100:0.2063122334,WP_216981205.1-Lelliottia-amnigena:0.3745925245)87.1/100:0.1045489728,((WP_227534785.1-Erwinia-rhapontici:0.1771808482,WP_017800228.1-Erwinia-toletana:0.2118491705)91.2/100:0.1182516310,WP_255851625.1-Erwinia-persicina:0.2514465230)53.7/100:0.0979912932)80.4/100:0.0546856329,(WP_089585421.1-Escherichia-coli:0.0671396101,WP_104705897.1-Escherichia-coli:0.0273642097)99.1/100:0.1616057981)86.1/100:0.0838181966,((WP_016236761.1-Enterobacteriaceae:0.1153580003,WP_024239504.1-Escherichia-coli:0.1692723101)72.5/99:0.0489299750,WP_046458830.1-Obesumbacterium-proteus:0.4295069873)68.6/100:0.0687249882)52.3/98:0.0720021663,(WP_001610415.1-Escherichia-coli:0.0487458383,WP_138183825.1-Escherichia-sp.-E4385:0.0330086374)99.8/100:0.2298364926)87.8/98:0.0623386887,WP_033870520.1-Enterobacteriaceae:0.2711074916)60.6/75:0.0366482480,WP_100272167.1-Citrobacter-portucalensis:0.3633629195)76.7/51:0.0224663249,(((WP_157959988.1-Citrobacter:0.6286348717,WP_267387874.1-Citrobacter-portucalensis:0.3339738818)65.4/71:0.0814203494,WP_138098269.1-Jejubacter-calystegiae:0.2540147888)0/50:0.0062459668,((WP_095844187.1-Escherichia-coli:0.0629473669,(WP_086118650.1-Escherichia-coli:0.0330156553,WP_089538038.1-Escherichia-coli:0.0422432626)86.3/100:0.0276879382)95/100:0.1309370053,WP_179154141.1-Citrobacter-freundii:0.4054604788)92.8/89:0.1186017599)64.7/52:0.0075590111)89.7/93:0.0402452887,((((WP_137650031.1-Escherichia-albertii:0.1592507419,WP_073446180.1-Escherichia-coli:0.2951847989)87.9/85:0.0679601294,(WP_001100786.1-Escherichia-coli:0.1571583119,(WP_248249449.1-Enterobacter-roggenkampii:0.1902783572,WP_032936727.1-Citrobacter-freundii:0.1557869769)99.7/100:0.2158145485)26.5/94:0.0103543777)73.6/83:0.0331056338,((WP_001100804.1-Enterobacteriaceae:0.1728199330,WP_001297344.1-Escherichia-coli:0.2635799847)28.2/69:0.0459627427,(WP_213789144.1-Enterobacter-hormaechei:0.0308311701,((WP_148242178.1-Enterobacter-asburiae:0.0629530123,WP_237994110.1-Enterobacter-sp.-DRP3:0.0232880946)71.3/100:0.0054617944,WP_237424876.1-Enterobacter-cloacae-complex-sp.-ECC445:0.0590427581)80.8/100:0.0355984078)100/100:0.3139234593)89.4/77:0.0845590478)67.4/68:0.0241309146,(WP_174633339.1-Escherichia-coli:0.1384348173,WP_072664248.1-Escherichia:0.1188811455)96.6/100:0.1045401095)85.8/76:0.0387259991)84.9/79:0.0451250326,WP_086539147.1-Citrobacter-freundii:0.3980256345)15.6/13:0.0127758916,WP_100029735.1-Escherichia-coli:0.1973227690)78.4/46:0.0270228888,(((WP_107192407.1-Escherichia-albertii:0.0201155344,WP_181673375.1-Escherichia-fergusonii:0.0594734128)98.6/100:0.1542617153,((WP_032214194.1-Escherichia-coli:0.0619336897,(WP_059217428.1-Escherichia-albertii:0.0391833626,(WP_073278747.1-Escherichia-coli:0.1047636649,WP_087893724.1-Enterobacteriaceae:0.0391729528)99.8/91:0.1946120987)79/64:0.0184710972)76.7/61:0.0312020681,WP_085459683.1-Escherichia-coli:0.0902999951)98.6/87:0.2111964589)80.1/64:0.0852111577,(WP_167816599.1-Enterobacter-cloacae:0.3722301722,(WP_262183194.1-Enterobacter-mori:0.1858163347,(WP_063412371.1-Enterobacter-cloacae:0.0554183826,WP_047027451.1-Enterobacter-kobei:0.0624353195)98.2/100:0.1350880954)93.4/100:0.1697097551)91.7/62:0.1360646348)42/60:0.0225838351)78.9/58:0.0322437185,((WP_119938542.1-Enterobacter-huaxiensis:0.1396041300,WP_050871166.1-Enterobacter-cloacae-complex:0.0400803236)95/100:0.0671751564,(WP_228937168.1-Escherichia-marmotae:0.0401161426,WP_040079227.1-Escherichia-coli:0.0469244876)81.7/95:0.0229596060)94.2/95:0.0659587556)84.5/59:0.0308125307,(((WP_168756429.1-Aeromonas-hydrophila:0.6983633547,(WP_021577651.1-Escherichia-coli:0.1735980467,(WP_021540003.1-Escherichia-coli:0.0144066984,WP_047657564.1-Enterobacteriaceae:0.0633047761)98.2/100:0.1303141581)0/61:0.0144615808)83.9/87:0.0405496501,(((((WP_170897747.1-Escherichia-coli:0.2077256632,WP_001100781.1-Escherichia-coli:0.2356954566)76.8/37:0.0436831924,WP_208464084.1-Escherichia-coli:0.1410787163)95.2/59:0.1003269756,(WP_220393175.1-Citrobacter-freundii:0.3380966723,WP_219093375.1-Escherichia-coli:0.2370505753)75.3/96:0.0464084932)44/26:0.0379753816,WP_001542972.1-Escherichia-coli:0.2137173233)17.4/50:0.0091676128,((WP_042093344.1-Escherichia-coli:0.2764841526,WP_089706542.1-Escherichia-coli:0.0880810581)80.4/98:0.0568419695,WP_262938691.1-Escherichia-albertii:0.1577257363)75.7/71:0.0274889711)90.5/25:0.0534348443)0/43:0.0000027537,(((((((((WP_127791218.1-Escherichia-coli:0.1556110871,(WP_025210617.1-Escherichia-coli:0.1735760159,WP_001527244.1-Escherichia-coli:0.0972191592)28.8/99:0.0440248901)77.6/93:0.0322586241,(WP_250341216.1-Pantoea-eucrina:0.3631010180,(((WP_049002539.1-Citrobacter-freundii:0.1496859810,(WP_249576357.1-Enterobacter-vonholyi:0.0728362492,WP_161617217.1-Yokenella-regensburgei:0.0527867682)95.9/100:0.0821429327)85.2/100:0.0352092645,((WP_079834758.1-Salmonella-enterica:0.0034246313,(WP_079793206.1-Salmonella-enterica:0.1638173770,WP_001100807.1-Salmonella:0.0991374393)99.6/100:0.1972807889)76.3/100:0.0477633923,WP_058109100.1-Salmonella-enterica:0.0376840013)99.6/100:0.1103171083)91/100:0.0465583236,WP_032303308.1-Escherichia:0.0922513371)75.4/100:0.0335502261)77.6/95:0.0402635538)68.1/86:0.0194088322,(WP_001100798.1-Enterobacteriaceae:0.1553141053,(WP_085454021.1-Escherichia-coli:0.1517302823,(WP_088541377.1-Escherichia-coli:0.0337748892,WP_222691870.1-Escherichia-marmotae:0.0769326109)82.2/100:0.0458367205)37.3/91:0.0417131930)51.4/76:0.0364763349)70/76:0.0421133021,((WP_059222105.1-Escherichia-albertii:0.1531269399,WP_187209312.1-Escherichia-coli:0.1063907849)82/91:0.0406297597,WP_218070502.1-Escherichia-coli:0.1180436287)20.1/57:0.0159776877)77.2/42:0.0271249150,(((((WP_096972986.1-Escherichia-coli:0.1395181666,(WP_157704420.1-Escherichia-coli:0.1094231778,WP_042200020.1-Enterobacteriaceae:0.0932015443)77.4/97:0.0241402975)73.8/70:0.0192946879,(WP_097421351.1-Escherichia-coli:0.1269854605,(WP_137653986.1-Escherichia-albertii:0.0391740282,WP_001100785.1-Escherichia-coli:0.0453456695)90.7/100:0.0637157614)79.5/99:0.0457973080)83.6/56:0.0349396881,WP_053290779.1-Escherichia-coli:0.1425309468)41.4/55:0.0159293141,WP_063610377.1-Escherichia-coli:0.1379356711)80.8/38:0.0272552087,((WP_146314246.1-Escherichia-coli:0.0477960212,WP_096967326.1-Escherichia-coli:0.0304205809)98.5/100:0.1104187350,WP_181465273.1-Escherichia-fergusonii:0.0831214002)85.2/85:0.0359325003)89.9/80:0.0502780288)61.7/40:0.0159756738,(((((WP_115348269.1-Hafnia-alvei:0.1502441982,WP_032659356.1-Enterobacter-hormaechei:0.4969029671)88.3/94:0.0898777608,(WP_130961887.1-Hafnia-paralvei:0.2328091082,WP_061059890.1-Hafnia-paralvei:0.2429824262)42.7/100:0.0246264044)82.7/87:0.0474495006,(WP_120162698.1-Rahnella-variigena:0.0701292016,WP_183847446.1-Rahnella-inusitata:0.0792176335)100/100:0.4238920164)88.5/75:0.0716218664,((WP_175320495.1-Hafnia-paralvei:0.1468230591,WP_130996599.1-Hafnia-paralvei:0.2075127465)98.3/99:0.1342670379,(((WP_171269898.1-Aeromonas-media:0.1463289953,WP_216983019.1-Aeromonas-sp.-FDAARGOS-1415:0.0880322103)97.2/100:0.1274311253,WP_168366848.1-Dickeya-zeae:0.2229645200)76.4/84:0.0620310053,(WP_064338398.1-Aeromonas-veronii:0.1797335777,((((WP_042864139.1-Aeromonas-caviae:0.0517287990,WP_225159881.1-Aeromonas-caviae:0.0497853901)95.1/100:0.1017863963,(WP_252995952.1-Aeromonas-encheleia:0.2960493004,(WP_244056308.1-Aeromonas-caviae:0.1445380979,WP_202046159.1-Aeromonas-media:0.1676529036)92.6/100:0.0952362720)86.7/54:0.0504118823)85/54:0.0502549896,(((WP_227095090.1-Aeromonas-veronii:0.0852127565,WP_152109808.1-Plesiomonas-shigelloides:0.1077682799)98.6/100:0.1662778897,WP_045790994.1-Aeromonas-hydrophila:0.1895282174)81.3/82:0.0503195570,(((((((WP_247163170.1-Escherichia-coli:0.0232195087,WP_038644959.1-Pantoea:0.0961535512)99.1/100:0.1166893006,(WP_252422582.1-Aeromonas-taiwanensis:0.0213106587,WP_263719954.1-Aeromonas-caviae:0.1074287180)79.7/96:0.0552041045)89.7/99:0.0482657602,WP_033131297.1-Aeromonas-aquatica:0.0281134677)95.9/95:0.0522169299,WP_241344952.1-Aeromonas-sp.-MR16:0.0157586463)88.1/96:0.0310612272,WP_160838676.1-Aeromonas-caviae:0.0925797448)72.3/89:0.0183339973,(WP_167569591.1-Aeromonas-veronii:0.1598193703,(WP_116547128.1-Plesiomonas-shigelloides:0.1137605327,(((WP_139496062.1-Escherichia-coli:0.0803988204,WP_201996661.1-Aeromonas-veronii:0.0381428618)32.2/99:0.0082767527,WP_215269322.1-Aeromonas-dhakensis:0.0548475618)67.3/99:0.0086205037,WP_235463836.1-Aeromonas-australiensis:0.0223806675)40.6/86:0.0212849227)60.7/98:0.0112373828)32.5/50:0.0454686123)42.7/56:0.0374828320,WP_049048056.1-Aeromonas-hydrophila:0.1870770380)97.2/100:0.1097698325)76.4/83:0.0224184682)81.6/56:0.0304007483,(WP_223932214.1-Aeromonas-caviae:0.0139008882,WP_161506954.1-Aeromonas-media:0.1580307388)100/100:0.1956058248)64/61:0.0319351220)83.2/64:0.0477324850)89.8/100:0.0615727701)74.8/61:0.0165251953)87.6/90:0.0550411267,(WP_034458444.1-Buttiauxella-noackiae:0.1978010151,WP_253383865.1-unidentified-bacterial-endosymbiont:0.4080471229)58.4/96:0.0726689524)94.1/87:0.1004229327)55.3/61:0.0191975849,(((WP_187216555.1-Escherichia-coli:0.2371896742,(WP_262931739.1-Escherichia-albertii:0.0535314757,WP_160188772.1-Escherichia-coli:0.0455123815)96.9/100:0.1061792968)88.4/92:0.0574897754,WP_015386517.1-Cronobacter-sakazakii:0.2531613248)80.6/60:0.0333656722,(((((WP_232904305.1-Buttiauxella-sp.-W03-F01:0.2248720655,WP_259388919.1-Enterobacter-huaxiensis:0.0643521313)78.2/100:0.0316305886,WP_230350752.1-Lelliottia-sp.-WAP21:0.1827666126)84.4/100:0.0593610655,WP_034915863.1-Erwinia-sp.-9145:0.2111118638)52.1/99:0.0491103581,WP_105963664.1-Escherichia-coli:0.0719094872)82.8/88:0.0319356044,WP_107179425.1-Escherichia:0.1594933628)83.1/61:0.0256008999)13.7/50:0.0072156404)87.6/64:0.0324432677,(((((((((((WP_193360946.1-Enterobacter-cloacae-complex-sp.-P39RS:0.0685855576,WP_182945676.1-Enterobacter-cloacae:0.0292583542)99.1/100:0.1184609447,WP_265185975.1-Enterobacter-mori:0.0952398530)82.8/100:0.0325457384,((WP_032636062.1-Enterobacter-kobei:0.0886917348,(WP_063137673.1-Enterobacter-cloacae-complex:0.0493812695,WP_165464597.1-Enterobacter-cloacae:0.0625851330)89.8/99:0.0577406390)47.9/14:0.0516310394,WP_138014015.1-Enterobacter-ludwigii:0.0679372320)85.4/99:0.0388588641)59.3/78:0.0476952695,WP_239609625.1-Lelliottia-amnigena:0.1511067330)0/7:0.0458760638,((WP_096205048.1-Enterobacter:0.0348210347,WP_100165931.1-Enterobacter-mori:0.0442802750)93.3/100:0.0713432314,WP_242172213.1-Enterobacter:0.1295965020)92.1/100:0.0795563089)70.4/62:0.0360003631,WP_033487312.1-Enterobacter-cloacae-complex:0.0656613707)87.2/57:0.0379308251,((((WP_042716137.1-Enterobacter-sp.-B509:0.0171854341,WP_241820523.1-Enterobacter-kobei:0.0908128434)97.7/100:0.1219477565,(WP_172688853.1-Enterobacterales:0.1427421084,WP_126546226.1-Enterobacter-quasimori:0.1137997217)85.4/100:0.0568780788)86.6/87:0.0495023407,((WP_045338355.1-Enterobacter-hormaechei:0.1942146361,((WP_182115563.1-Enterobacter-kobei:0.0551572472,WP_213381159.1-Enterobacter-cloacae:0.0740728804)96.6/100:0.1132981410,WP_063158410.1-Enterobacter-hormaechei:0.0839856026)38.4/58:0.0469157951)78.5/94:0.0265674072,WP_063869545.1-Enterobacter-hormaechei:0.0694547684)57.1/56:0.0392362458)75.2/51:0.0119853705,(WP_046888360.1-Enterobacter-cloacae:0.0868706229,WP_126315015.1-Enterobacter-hormaechei:0.1238928810)86.7/93:0.0343713824)27.5/53:0.0144658017)92.6/94:0.0525941901,(WP_191149682.1-Erwinia-aphidicola:0.2496918500,WP_208283396.1-Enterobacter-sichuanensis:0.1316110518)47.9/63:0.0257512143)76.7/65:0.0144395159,(((((WP_181680936.1-Citrobacter-sp.-RHB25-C09:0.1375611727,(WP_090049757.1-Citrobacter-amalonaticus:0.0349131938,WP_196013128.1-Citrobacter-farmeri:0.0411610332)99.1/100:0.1540599298)85.6/100:0.0617643785,WP_103775088.1-Citrobacter-amalonaticus:0.2182573895)98.2/100:0.1737669562,WP_121265736.1-Enterobacter-sp.-R1-2018:0.2924679294)88.5/92:0.0784080753,WP_259047621.1-Klebsiella-sp.-BIGb0407:0.2104607969)0/89:0.0052915912,WP_249031739.1-Tenebrionicola-larvae:0.3722659581)85.8/92:0.0583442116)79.1/46:0.0147277749,(((((WP_141109706.1-Escherichia-albertii:0.1092512114,WP_001572827.1-Escherichia-coli:0.0744068624)36/74:0.0280055542,WP_203360733.1-Citrobacter-sp.-R56:0.0699541191)88/79:0.0581449656,WP_258993080.1-Scandinavium-hiltneri:0.1072286997)95.7/100:0.1122787602,((WP_095374511.1-Escherichia-coli:0.0126994031,WP_012906377.1-Citrobacter-rodentium:0.1159112925)97.1/100:0.1188267239,(WP_096122580.1-Escherichia-coli:0.0680520364,WP_162379936.1-Citrobacter-sp.-NCU1:0.2319274699)82.6/95:0.0455109334)64.2/98:0.0537485994)35.3/77:0.0156107828,WP_259000881.1-Scandinavium-goeteborgense:0.2134301299)57.5/54:0.0321407550)74.4/39:0.0108325016,((((WP_075180894.1-Pantoea-sp.-1.19:0.3144122688,(WP_162648791.1-Citrobacter-freundii:0.1615214178,(WP_060683585.1-Citrobacter-portucalensis:0.1527694222,(WP_079798993.1-Salmonella-enterica:0.1171039720,WP_032255798.1-Enterobacteriaceae:0.0995942519)60.5/97:0.0076490496)94/97:0.0841601165)88.2/99:0.0659739081)85.6/44:0.0545431078,(WP_195994783.1-Citrobacter-portucalensis:0.3076129116,WP_048218074.1-Citrobacter-freundii-complex:0.1826132486)81.7/95:0.0469562539)75.7/66:0.0258228807,(WP_181624464.1-Citrobacter-freundii:0.2905521314,WP_001515521.1-Escherichia-coli:0.2283084017)91.3/100:0.1303514388)39.8/72:0.0492632850,((((WP_058662478.1-Enterobacteriaceae:0.0504892212,WP_061709645.1-Pseudenterobacter-timonensis:0.0607839865)99.7/100:0.2404259286,WP_112164393.1-Rahnella-inusitata:0.2003072380)64/99:0.0752800916,WP_157953065.1-Limnobaculum-parvum:0.2848873639)80.6/100:0.0734532160,WP_047609274.1-Rahnella-aquatilis:0.2379748862)84.7/81:0.0564746247)95/80:0.0615548589)75.8/64:0.0109154467)74.4/72:0.0156009313,(((WP_109548670.1-Escherichia-coli:0.0569614659,WP_001100790.1-Escherichia:0.0281873814)95.6/100:0.1207523810,WP_061892435.1-Escherichia-coli:0.2114071133)0/92:0.0046960678,WP_181237856.1-Escherichia-marmotae:0.0893319854)89.8/95:0.0578548429)76/89:0.0219554279)87.7/85:0.0248813364)77.1/66:0.0189225681,(((((((((((((((((WP_151711904.1-Enterobacter-ludwigii:0.0356751710,(WP_032941356.1-Citrobacter-freundii:0.2652209316,WP_058678686.1-Enterobacter:0.0932639446)92/100:0.0788969848)92.1/100:0.0912809716,WP_223560970.1-unclassified-Enterobacter:0.0000021096)98.6/100:0.3563580245,WP_044720521.1-Escherichia-coli:0.5377051936)99.5/100:0.4461926897,(WP_159241675.1-Proteus-terrae:0.2527024873,WP_195697765.1-Providencia-rettgeri:0.2345948544)4.3/81:0.0516563352)86.2/93:0.0569386221,WP_238563424.1-Providencia-rettgeri:0.1491318812)72.6/87:0.0217349983,WP_165880752.1-Providencia-rettgeri:0.2501382938)85.1/99:0.0417392778,WP_231068620.1-Providencia-huaxiensis:0.1923689601)87.9/93:0.0348494546,(((WP_187128479.1-Providencia-sp.-JUb39:0.2040707037,WP_193115284.1-Escherichia-coli:0.2050090819)83.5/100:0.0602205987,WP_154625743.1-Providencia-stuartii:0.2061570473)35.4/99:0.0321533257,WP_004924376.1-Providencia-stuartii:0.2730155067)0/89:0.0000026787)78.6/96:0.0283139511,(WP_169342312.1-Providencia-stuartii:0.2256185383,WP_164565644.1-Providencia-rettgeri:0.2713564795)81.1/98:0.0682867913)86.9/100:0.0679923178,WP_234505032.1-Kluyvera-intermedia:0.5497382041)74.3/98:0.0683961147,WP_257124601.1-Aeromonas-veronii:0.5098620709)78.4/66:0.0225411861,(WP_038258850.1-Xenorhabdus-bovienii:0.3352587836,WP_086113286.1-Xenorhabdus-beddingii:0.2675557780)45.6/59:0.1038483319)72/74:0.0262978488,(WP_038223662.1-Xenorhabdus-bovienii:0.2831243552,((WP_161748440.1-Proteus-sp.-G2658:0.2894381537,WP_049199273.1-Proteus-mirabilis:0.0709553985)59.1/100:0.0596914015,WP_157954980.1-Proteus-terrae:0.0821167136)98.1/100:0.1726843305)87.3/89:0.0665110235)79.3/89:0.0402050728,(WP_064374050.1-Pantoea:0.4458747385,((WP_036421240.1-Morganella-morganii:0.2084120573,(WP_264998074.1-Morganella-morganii:0.1030698738,WP_186849509.1-Morganella-morganii:0.1077804612)94.8/100:0.0855455454)92.3/100:0.0843460462,((((WP_230863267.1-Enterobacter-cloacae:0.6765455619,WP_164716793.1-Cedecea-lapagei:0.4664470369)19.7/70:0.1101379570,WP_034496584.1-Buttiauxella-agrestis:0.4912105501)65.8/71:0.0601505682,WP_050862535.1-Enterobacter:0.5428471757)77.8/71:0.0403751729,WP_029399168.1-Escherichia-coli:0.4011504178)99.1/99:0.1785722124)90/99:0.0744649422)85.1/85:0.0477389335)71.4/81:0.0157849172,(WP_039304623.1-Cedecea-neteri:0.5043881673,((WP_159282060.1-Rahnella-variigena:0.4690980324,WP_106902182.1-Citrobacter-freundii:0.2316059348)75.9/87:0.1043068846,(WP_126291239.1-Enterobacter-hormaechei:0.1977487974,((WP_062932819.1-Enterobacter-hormaechei:0.0447971955,WP_252042312.1-Enterobacter-chengduensis:0.0464350396)82.1/99:0.0268986844,((WP_049008289.1-Enterobacter-cloacae:0.0807056880,WP_048999124.1-Enterobacter:0.0472244609)68/26:0.0094861690,WP_063144508.1-Enterobacter-cloacae:0.0518643645)29.7/25:0.0098160059)82.3/96:0.0680575676)96.5/100:0.1441476526)36.4/58:0.0236186672)96/97:0.0911224877)91.8/66:0.0611927582,(((WP_108569539.1-unclassified-Pantoea:0.3643863864,WP_028018520.1-Enterobacter-cloacae-complex:0.8284026202)92.3/100:0.2171849797,WP_269535377.1-Citrobacter-sedlakii:0.2822019687)84.2/90:0.0819760803,WP_029881813.1-Enterobacter:0.4220197262)80.5/52:0.0435953926)67.1/41:0.0262504426,((((((WP_090123257.1-Kosakonia-arachidis:0.1022461758,WP_020454293.1-Kosakonia-sp.-SOY2:0.1273338888)89.8/100:0.0606976544,WP_017458471.1-Kosakonia-sacchari:0.1835613085)81.3/100:0.0549457563,(WP_200002836.1-Citrobacter-freundii:0.2148653933,WP_080329661.1-Enterobacter-cloacae-complex:0.1773229921)98.9/100:0.2319108646)85.8/100:0.1177215395,WP_090460354.1-Enterobacter-sp.-kpr-6:0.1025394548)98.3/100:0.1848470674,((WP_105274811.1-Escherichia:0.2202712218,WP_064512195.1-Buttiauxella-gaviniae:0.4107190126)95.6/63:0.1338156588,WP_140919905.1-Insectihabitans-xujianqingii:0.2945722094)39.7/19:0.0238255627)87.8/19:0.0714425875,((WP_194857598.1-Citrobacter-freundii:0.4529722699,((WP_141032926.1-Escherichia-coli:0.1860594226,WP_196390204.1-Enterobacter-mori:0.0543489455)51.6/97:0.0465478865,(WP_252032207.1-Enterobacter-asburiae:0.0621486544,WP_111966211.1-Enterobacter-cloacae-complex:0.0342525102)80.6/94:0.0315365314)95.7/99:0.1843187929)99.7/100:0.3281149492,WP_070133611.1-Candidatus-Erwinia-dacicola:0.3005628884)76.8/55:0.0386376717)82/41:0.0422970507)87.5/68:0.0414484032)94.3/84:0.0613145349,((((WP_103946175.1-Lelliottia-aquatilis:0.1198945831,WP_235147774.1-Enterobacter-cloacae:0.3660454928)80.3/99:0.0243753935,WP_123427447.1-Lelliottia:0.0761650389)96.9/100:0.1136822008,((((WP_214593796.1-Enterobacter-mori:0.0549663404,WP_086532312.1-Enterobacter-ludwigii:0.0943271194)95.6/100:0.1172938119,WP_045339085.1-Enterobacter-cloacae-complex:0.2267499153)44.3/100:0.0530931043,((WP_047742849.1-Enterobacter:0.1003386902,WP_040022520.1-Enterobacter-cloacae:0.0909085243)97.7/100:0.1554462067,(WP_129995405.1-Enterobacter-cloacae:0.0776457610,WP_042099316.1-Escherichia-coli:0.0950880354)63.5/95:0.0201646923)90.8/99:0.1299468970)87.4/100:0.0704386612,WP_153879942.1-Citrobacter-freundii:0.2321883173)85.3/85:0.0515829912)84.7/68:0.0411416526,(((((WP_097405671.1-Kosakonia:0.0573107047,WP_161518249.1-Kosakonia-sp.-MH5:0.0811592063)89.8/100:0.0726175144,WP_128482629.1-Kosakonia-cowanii:0.1443143765)98.7/100:0.1801198048,WP_206164314.1-Citrobacter:0.2856333254)69.2/99:0.0712355083,((((WP_238085128.1-Pseudescherichia-vulneris:0.0259480430,WP_032652659.1-Enterobacter-cloacae-complex:0.1203558008)65.7/89:0.0154730156,WP_129363899.1-Enterobacter-cloacae:0.0370067841)74/89:0.0214039295,(WP_257661844.1-Citrobacter-portucalensis:0.1493650485,WP_193134185.1-Citrobacter-freundii:0.1014573467)85.4/99:0.0352361528)94.9/98:0.1024010951,(WP_042284276.1-Citrobacter-sedlakii:0.2930554451,(WP_024193259.1-Escherichia-coli:0.1319653239,WP_099433281.1-Citrobacter-amalonaticus:0.0414352188)69/100:0.1034594964)93.7/100:0.1194302622)95/99:0.1883572440)84.3/93:0.0661713329,(((WP_167863639.1-Erwinia-rhapontici:0.1630652902,WP_039385357.1-Pantoea-eucrina:0.2603855061)88.6/100:0.1085524278,WP_118663751.1-Erwinia-persicina:0.1782095953)44.2/97:0.0716591054,WP_062742814.1-Erwinia-persicina:0.2548920660)88.5/96:0.0711354471)32.4/66:0.0210866494)89.5/77:0.0450443539)77.2/76:0.0189858489)86.2/86:0.0227449171,WP_202667543.1-Lelliottia-amnigena:0.2889821156)38.7/85:0.0109723040,((((((((((WP_172912938.1-Pantoea-stewartii:0.1573359094,WP_145394131.1-Pantoea-sp.-SJZ147:0.1730064016)99.5/100:0.3727370734,WP_061356323.1-Escherichia-coli:0.9732437787)88.4/86:0.1492433589,WP_153752706.1-unclassified-Citrobacter:0.6017304199)77/85:0.0566999879,WP_043158915.1-Aeromonas-hydrophila:0.5434543167)90.1/83:0.1139582559,WP_097364011.1-Edwardsiella-tarda:0.4443242471)4.6/72:0.0269601062,WP_086868392.1-Kosakonia-pseudosacchari:1.0287567321)80.6/38:0.0654427907,((((((((((((((((((((WP_167868984.1-Shewanella-sp.-WXL01:0.5421781570,(WP_222525605.1-Ferrimonas-balearica:0.2017743824,WP_013344594.1-Ferrimonas-balearica:0.1620743597)90.2/100:0.1405305545)89.9/99:0.1022976131,(WP_016959527.1-Enterovibrio-norvegicus:0.0914513778,WP_229491984.1-Enterovibrio-norvegicus:0.1417994002)66.8/100:0.0628874655)85.3/88:0.0504831331,(((((((WP_047049276.1-Vibrio-mexicanus:0.1674145459,WP_193630602.1-Shewanella-benthica:0.2599260536)85.5/93:0.0527742799,WP_047874860.1-Photobacterium-aphoticum:0.2721019256)73.6/90:0.0265654418,WP_020335761.1-Vibrio-natriegens:0.1442335769)85.1/88:0.0350007054,((((WP_269599241.1-Salinivibrio-kushneri:0.0877955825,WP_136483218.1-Vibrio-sp.-H11:0.0731350137)98.3/100:0.1206927788,(WP_012143346.1-Shewanella-sediminis:0.2556882258,WP_140076235.1-Vibrio-parahaemolyticus:0.1419889684)74.9/99:0.0174350536)48.3/92:0.0202383123,WP_205390674.1-Vibrio:0.1070005515)78.5/92:0.0216433629,((WP_167678613.1-Shewanella-aestuarii:0.2312034275,(WP_113818410.1-Psychromonas-sp.-B3M02:0.0630327146,(WP_025564776.1-Psychromonas-sp.-SP041:0.0591065482,WP_166372414.1-Psychromonas-sp.-SA13A:0.0183603013)83.2/100:0.0702157103)88.7/100:0.0602520900)61.4/53:0.0244804425,((((((((WP_188955407.1-Shewanella-ulleungensis:0.0632988635,WP_137222113.1-Shewanella-sp.-MEBiC00475:0.0798215581)92.7/97:0.0659605431,WP_155694061.1-Psychrosphaera-haliotis:0.1772390025)96.6/54:0.0824336072,WP_126005793.1-Pseudoalteromonas-sp.-Xi13:0.0402498628)46.3/45:0.0036656007,WP_256848444.1-Pseudoalteromonas-carrageenovora:0.0324341663)39.8/42:0.0227001091,WP_182693438.1-Pseudoalteromonas-sp.-SR45-5:0.0725365111)88.9/54:0.0379604146,WP_138577097.1-unclassified-Pseudoalteromonas:0.1236706490)42.8/53:0.0140991805,(WP_264390127.1-Pseudoalteromonas-sp.-A3:0.0316957713,(WP_096739047.1-Pseudoalteromonas-atlantica:0.0471444241,WP_248462211.1-Pseudoalteromonas-sp.-2CM37A:0.0232919702)90/100:0.0258832701)0/94:0.0040869339)97/96:0.1073692818,WP_022943097.1-Psychromonas-hadalis:0.1658111517)72/51:0.0345957032)88/92:0.0395695496)71.3/82:0.0260124533)50.8/80:0.0170820467,(WP_050648448.1-Vibrio-sp.-J2-4:0.1684019859,WP_264874461.1-Vibrio-agarivorans:0.1187196836)77.1/93:0.0496964644)81.3/97:0.0220256399,WP_164649367.1-Vibrio-astriarenae:0.1078393379)83.6/98:0.0344785420,(WP_084884043.1-Vibrio-sp.-qd031:0.1635779844,WP_159656123.1-Vibrio-atypicus:0.1344424777)86.7/96:0.0471714522)76.5/88:0.0199422531)0/58:0.0327613800,WP_093326971.1-Thalassotalea-agarivorans:0.1404852950)66/82:0.0800761614,(((WP_142941196.1-Aliikangiella-marina:0.4232320328,(WP_068546663.1-Thalassotalea-crassostreae:0.3019287316,(WP_081148784.1-Cognaticolwellia-beringensis:0.0579903437,WP_007634918.1-Paraglaciecola-psychrophila:0.0460655278)99.6/100:0.2317781094)0/59:0.0209984913)88.7/100:0.0648341009,((((((((((WP_160153838.1-Microbulbifer-sp.-ALW1:0.3235559940,((WP_263600466.1-Microbulbifer-agarilyticus:0.1747506330,WP_078085574.1-Microbulbifer-mangrovi:0.1007820862)59.7/98:0.0167944477,WP_029249715.1-Microbulbifer-agarilyticus:0.1267275811)92.1/98:0.1104742361)96.3/100:0.1339464018,(WP_138624780.1-Pseudoalteromonas-sp.-S4492:0.3482359522,WP_130050482.1-Pseudoalteromonas-sp.-MEBiC-03485:0.1240451213)97.7/93:0.2419007874)38.2/7:0.0167559193,(((WP_085356321.1-Neisseria-dumasiana:0.2732576748,((WP_101133177.1-Neisseria-meningitidis:0.0767978990,WP_005552781.1-Aggregatibacter-actinomycetemcomitans:0.2531452598)8.1/48:0.0165778823,(WP_127202078.1-Neisseria-meningitidis:0.1433385369,WP_237092367.1-Neisseria-sp.-ZJ104:0.1850010863)88.5/98:0.0681919751)49.1/67:0.0489470581)93.3/99:0.2292687872,WP_031579865.1-Ruminobacter-sp.-RM87:0.5897348527)94/99:0.1528646854,(WP_189610147.1-Saccharospirillum-salsuginis:0.4120464906,WP_111640782.1-Marinimicrobium-alkaliphilum:0.4114247097)81.6/88:0.1183953025)37/5:0.0569439209)64.8/84:0.0389060956,WP_118961222.1-Colwellia-sp.-RSH04:0.5068150249)59.9/96:0.0344361113,WP_186505417.1-Neptunicella-marina:0.2493833797)91.3/97:0.0854192680,(WP_102798355.1-Bowmanella-denitrificans:0.0861974781,WP_214344498.1-Bowmanella-yangjiangensis:0.0459621807)99.9/100:0.2446561453)50/86:0.0307049015,(WP_124026287.1-Alteromonas-sediminis:0.1950544421,WP_100655941.1-Alteromonas-flava:0.2906109785)99.8/100:0.2232738694)61.3/70:0.0253286944,(WP_216048042.1-Aliiglaciecola-lipolytica:0.2774992696,(WP_131095783.1-Alteromonas-sp.-KUL42:0.0964091440,((((((WP_100972994.1-Alteromonas-sp.-MB-3u-76:0.1578742392,(WP_071816068.1-Alteromonas-sp.-V450:0.1184990097,WP_198816918.1-Alteromonas-sp.-IB21:0.0254634535)70.8/100:0.0080405775)75.8/89:0.0107735123,(((WP_156078883.1-Alteromonas:0.0670785155,((((WP_063457834.1-Alteromonas-stellipolaris:0.0399928453,(WP_202940767.1-Alteromonas-macleodii:0.0264036698,WP_061095937.1-Alteromonas-macleodii:0.0484571394)84.2/79:0.0150116630)75.4/73:0.0072993278,((WP_044058190.1-Alteromonas-australica:0.0490726648,WP_159580202.1-Alteromonas-sp.-KUL106:0.0240830611)78.8/100:0.0153536064,(WP_014977514.1-Alteromonas-macleodii:0.0412211807,WP_239220085.1-Alteromonas-sp.-MmMcT2-2:0.0325693763)0/97:0.0000027410)44.9/72:0.0079613509)68.8/90:0.0064000018,WP_187447473.1-Alteromonas-sp.-BZK5:0.0256847360)73.5/84:0.0085945769,WP_202262575.1-unclassified-Alteromonas:0.0512064689)48.9/71:0.0105385560)92.2/99:0.0310867057,(WP_163106912.1-Alteromonas-genovensis:0.0631895080,WP_057795858.1-Alteromonas-stellipolaris:0.0459398750)0/64:0.0000022023)80/81:0.0173559453,WP_269520955.1-Alteromonas-sp.-BMJM2:0.0910524804)63.5/81:0.0066697358)94.9/97:0.0410256914,WP_163085646.1-Alteromonas-profundi:0.0407744323)71.8/78:0.0070299706,(WP_130638173.1-Alteromonas-sp.-KUL49:0.0754662180,WP_115592514.1-Alteromonas-aestuariivivens:0.0567677014)57.2/98:0.0217024770)88.8/97:0.0271828857,((((WP_018984642.1-Salinimonas-chungwhensis:0.1224933281,WP_265616768.1-Alteromonas-sp.-ASW11-7:0.0805271948)84.9/89:0.0528580122,WP_216028209.1-Alteromonas-sp.-C1M14:0.0609296588)79.4/56:0.0214635934,(WP_265607972.1-Alteromonas-sp.-ASW11-130:0.0834256255,WP_119499891.1-Alteromonas-sp.-RKMC-009:0.0726422836)89/57:0.0345862072)62.4/73:0.0111454356,(WP_216026947.1-Aestuariibacter-sp.-A3R04:0.0730157158,(WP_070176406.1-Alteromonas-lipolytica:0.0424894255,WP_073325432.1-Marisediminitalea-aggregata:0.0607906785)91.2/100:0.0531143298)65.9/90:0.0272531597)94.3/88:0.0546795914)65.5/88:0.0236925883,(WP_262993134.1-Alteromonas-sp.-ASW11-19:0.1042389334,(WP_218354448.1-Alteromonas-lipotrueiana:0.0891303445,WP_195811108.1-Salinimonas-marina:0.0813121505)80.6/96:0.0514535836)75.7/89:0.0178300291)91.6/97:0.0523236325)91.6/98:0.0526923940)85.7/62:0.0387309606)91.4/70:0.0491301221,(WP_057181266.1-Colwellia:0.2567089777,WP_009839692.1-Pseudoalteromonas-tunicata:0.7639735124)70.1/78:0.0218759139)95.3/93:0.0874960140,(WP_250459773.1-Microbulbifer-sp.-GX-H0434:0.1454955230,WP_152451766.1-Microbulbifer-sp.-THAF38:0.2116117538)97.2/100:0.1676238931)81.1/77:0.0368546365)89.8/64:0.1329411632,(WP_138986566.1-Neptunomonas-concharum:0.4780062761,((WP_255857717.1-Bacterioplanoides-sp.-SCSIO-12839:0.4128384509,WP_116686321.1-Pelagibaculum-spongiae:0.6694386513)91.2/85:0.1393296156,((WP_220103444.1-Neiella-holothuriorum:0.1058417353,WP_087505717.1-Neiella-marina:0.2209050293)93.7/100:0.0910100994,((WP_028862352.1-Psychromonas-aquimarina:0.2456084831,WP_187408568.1-Saccharophagus-sp.-K07:0.1881901930)68.1/100:0.0539023857,(WP_251261538.1-Echinimonas-agarilytica:0.1792604448,(WP_096084688.1-Agaribacterium-haliotis:0.5010289782,WP_188151970.1-Teredinibacter-waterburyi:0.2531941939)92.2/99:0.1743103145)28.9/94:0.0414152317)93.8/99:0.1342403086)88.4/95:0.0628948908)77/53:0.0363704962)49.8/52:0.0284806160)33.4/80:0.0672083049)76.1/96:0.0653262509,(WP_248714967.1-Agarivorans-sp.-TSD2052:0.0906989208,WP_152781274.1-Agarivorans-sp.-B2Z047:0.0963510659)99.4/100:0.1653956743)98/97:0.1691235967,WP_234495031.1-Vibrio-maritimus:0.2926486837)84.1/91:0.0630963701,(WP_020332409.1-Vibrio-fluvialis:0.0978361564,WP_096629284.1-Salinivibrio-sp.-YCSC6:0.2270157961)85.2/94:0.0694375988)81.6/86:0.0265785500,((((WP_067416136.1-Enterovibrio-coralii:0.1095050464,((WP_171342182.1-Vibrio-cyclitrophicus:0.2273830931,(WP_259522409.1-Shewanella-baltica:0.1078071130,WP_173243873.1-Shewanella-sp.-DC2-4:0.2242356907)96.3/100:0.1448092162)71.9/94:0.0403839285,WP_161972844.1-Enterovibrio-baiacu:0.0896756085)95/98:0.1260365248)90/93:0.0860582667,((WP_223418564.1-Shewanella-inventionis:0.1441185081,WP_011497090.1-Shewanella-denitrificans:0.3000840098)69.3/98:0.1035604526,((WP_197173106.1-Pseudoalteromonas-sp.-NZS11_1:0.2256099933,WP_182748079.1-Pseudoalteromonas-sp.-SR41-1:0.2000101602)90.8/98:0.0833003359,WP_149593442.1-Pseudoalteromonas-sp.-FUC4:0.1749974758)76.6/98:0.0367181485)76.2/97:0.0288150609)82.5/87:0.0364970216,(WP_102038478.1-Shewanella-sp.-GutCb:0.2394704301,WP_166378370.1-Pseudoalteromonas-sp.-Z9A4:0.2973826350)71.8/79:0.0259666092)75.1/68:0.0152133276,((WP_017031451.1-Vibrio-breoganii:0.0959935830,WP_248975945.1-Shewanella-kaireitica:0.2077418021)92.3/100:0.0778099022,(WP_193262143.1-Vibrio-parahaemolyticus:0.3228895510,WP_112353352.1-Shewanella-benthica:0.2582552954)73.3/98:0.0193267135)91.5/83:0.0654481314)90.2/88:0.0528320782)77.1/69:0.0381775803,((((WP_262601261.1-Endozoicomonas-euniceicola:0.1209360263,WP_066015073.1-Endozoicomonas-atrinae:0.1678585431)92/100:0.1081572558,WP_257287521.1-Endozoicomonas-sp.-SESOKO2:0.1492789366)84.9/100:0.1081374590,(WP_067585802.1-Endozoicomonas-ascidiicola:0.1600156585,(WP_020580625.1-Endozoicomonas-elysicola:0.1247467482,WP_257293945.1-Endozoicomonas-sp.-YOMI1:0.1472351190)93.6/100:0.1238522989)99.1/100:0.2985366462)98.5/100:0.3178274421,WP_011189464.1-Desulfotalea-psychrophila:0.5820214117)76.2/83:0.0946676658)0/65:0.0047345853,(WP_038175370.1-Vibrio-pacinii:0.1574238534,(((((WP_047883806.1-Photobacterium-ganghwense:0.1636269181,WP_146157089.1-Photobacterium-rosenbergii:0.2236903773)79.6/74:0.0483489575,(WP_187463805.1-Photobacterium-sp.-BZF1:0.0630935111,WP_039467548.1-Photobacterium-gaetbulicola:0.1868253731)89.3/74:0.0796417259)93.8/100:0.0934565666,WP_239411815.1-Photobacterium-sp.-OFAV2-7:0.1816534140)63.5/97:0.0510292949,WP_075764060.1-Photobacterium-proteolyticum:0.1953775699)90.1/95:0.0618244449,WP_171749982.1-Vibrio-sp.-RE88:0.2298157741)0/46:0.0000020790)87.7/81:0.0546834783)93.2/86:0.0506536385,(((WP_190336189.1-Vibrio-sp.-S9_S30:0.1315094133,WP_022612811.1-Vibrio-nigripulchritudo:0.1360663350)92.8/100:0.0883226303,(((WP_261835819.1-Vibrio-ishigakensis:0.1855326561,WP_017221048.1-Moritella-dasanensis:0.1741167778)72.2/91:0.0496517694,WP_193154481.1-Vibrio-navarrensis:0.2565059356)43.7/91:0.0457072548,WP_102446429.1-Vibrio-breoganii:0.2165442999)64.3/89:0.0228442166)82.3/95:0.0507625595,(WP_070968050.1-Vibrio-sonorensis:0.2856829794,WP_102329601.1-Vibrio-breoganii:0.2486399690)73.3/82:0.0361880188)81.8/83:0.0435444498)71.6/24:0.0134596773,((((((((WP_188026786.1-Tolumonas-osonensis:0.0367034553,WP_024870911.1-Tolumonas-lignilytica:0.1515740437)91.2/100:0.1269644335,WP_105076797.1-Pantoea-ananatis:0.5206771425)57.9/63:0.1129880281,((((WP_144135068.1-Kosakonia-radicincitans:0.4056604257,((WP_088208149.1-Enterobacter-cancerogenus:0.1352938612,WP_121932958.1-Enterobacter-hormaechei:0.1205558056)88.4/100:0.0634677581,WP_023330866.1-Enterobacter:0.0515166497)96.5/100:0.1396407466)90/92:0.1015085585,WP_120453320.1-Kalamiella-piersonii:0.1351820172)72.2/74:0.0271481335,(WP_269294270.1-Pantoea-sp.-LMR881:0.0708344549,WP_215846277.1-Pantoea-sp.-EnvD:0.0819237493)97.7/100:0.1027576633)42.4/68:0.0363634717,((WP_214216788.1-Rosenbergiella-epipactidis:0.0731965658,WP_241623482.1-Rosenbergiella-epipactidis:0.0285557215)99.8/100:0.1912027004,(WP_187508308.1-Pantoea-agglomerans:0.2481205500,(((WP_192411534.1-Pantoea-sp.-PNT02:0.1426690988,WP_097116658.1-Pantoea-sp.-GL120224-02:0.0000021088)94.8/65:0.0342509375,((WP_150019269.1-unclassified-Pantoea:0.0914424490,WP_069025179.1-Pantoea-agglomerans:0.0077090863)15.4/44:0.0077269008,WP_140917215.1-Pantoea-deleyi:0.0521075088)92/94:0.0383265728)78.7/61:0.0253703551,WP_125311258.1-Pantoea-dispersa:0.0712872756)97.7/99:0.1542365202)42.9/44:0.0217573647)86.4/76:0.0572177291)78.6/73:0.0356697702)15.7/46:0.0748188399,((WP_076542412.1-Shewanella-sp.-UCD-KL21:0.4687170193,WP_007466453.1-Photobacterium-marinum:0.2018931386)86.1/93:0.0892301149,WP_120356561.1-Alginatibacterium-sediminis:0.3693750546)91.4/73:0.1153891363)64.4/73:0.0466652241,((WP_075989065.1-Vibrio-fluvialis:0.0445493286,WP_044366945.1-Vibrio-fluvialis:0.0388947064)74.2/81:0.0432411609,(WP_076657081.1-unclassified-Vibrio:0.0807859654,WP_239929047.1-Vibrio-kanaloae:0.0440405170)97.4/100:0.1274622952)97.1/100:0.1565671842)85/100:0.0503701573,((WP_192015074.1-Photobacterium-arenosum:0.0934823294,WP_088876367.1-Vibrio-mediterranei:0.3361141927)88.5/100:0.0908913107,WP_022590465.1-Vibrio-nigripulchritudo:0.2851277584)79.6/98:0.0445557501)92.6/98:0.0530919374,(((WP_102941572.1-Vibrio-diazotrophicus:0.2017878651,WP_045977433.1-Vibrio-neptunius:0.3037750615)0/9:0.0103764410,(WP_123015137.1-Vibrio-zhugei:0.0617857508,WP_077313846.1-Vibrio-palustris:0.0209471531)99.9/100:0.2161406633)60.8/10:0.0241561212,(WP_102455158.1-Vibrio-breoganii:0.1559004107,(WP_188011373.1-Vibrio-vulnificus:0.2508886259,WP_102451772.1-Vibrio-breoganii:0.1821215469)95/99:0.1381930135)90.2/97:0.0796734941)85.9/73:0.0441088072)79.5/73:0.0277604553,WP_180567593.1-Oceanospirillales:0.2140409324)10.2/24:0.0137989451)92.6/31:0.0671015525,((((((((((((WP_011771697.1-Psychromonas-ingrahamii:0.1526667351,WP_182690147.1-Psychromonas-sp.-SR45-3:0.1581989455)99.1/99:0.1835650134,(WP_102986644.1-Vibrio-diazotrophicus:0.2053584702,WP_130066657.1-Aliivibrio-finisterrensis:0.1799790504)75.6/76:0.0323654496)70/60:0.0101067439,((WP_065600047.1-Aliivibrio-sp.-1S128:0.1361695329,(WP_017063118.1-Vibrio:0.1995951810,WP_203433235.1-Shewanella-sp.-LZH-2:0.3061463528)0/33:0.0000023188)78.7/45:0.0378315035,WP_102285079.1-Vibrio-lentus:0.1957199793)76.7/53:0.0301344385)87/78:0.0294270975,((WP_048661380.1-Vibrio-crassostreae:0.1015240605,WP_193846925.1-Vibrio-sp.-OPT10:0.1237528179)84.7/98:0.0429780840,(((WP_102481006.1-Vibrio-breoganii:0.1566101351,WP_258489575.1-Vibrio-alginolyticus:0.1664064519)74.9/89:0.0577108126,(WP_261874894.1-Vibrio-rarus:0.1136075860,WP_138522257.1-Pseudoalteromonas-galatheae:0.4166137040)86.4/91:0.0988629284)82.2/96:0.0452925651,WP_102533425.1-Vibrio-breoganii:0.2096123890)82/93:0.0287506850)91/91:0.0395126307)0/34:0.0042325268,((WP_063656223.1-Aliivibrio-fischeri:0.1134813611,WP_076646094.1-Vibrio-sp.-10N.222.47.A9:0.1926314702)43/82:0.0353449016,WP_239937663.1-Vibrio-chagasii:0.2068477559)92.1/96:0.0873806719)74.8/63:0.0219424253,WP_017020702.1-Aliivibrio-logei:0.2813651895)65.6/70:0.0274676303,(((((((WP_233983986.1-Vibrio-fluvialis:0.1112956126,((((((WP_155682326.1-Aliivibrio-fischeri:0.1650593813,WP_102482947.1-Vibrio-splendidus:0.1271115464)54.4/89:0.0306250445,WP_102394203.1-Vibrio-sp.-10N.222.54.B6:0.1342678999)74.9/91:0.0274394761,WP_016784063.1-Vibrio:0.1537394863)5/54:0.0372931663,WP_140342839.1-Vibrio-parahaemolyticus:0.2358611038)80.4/98:0.0352024311,(WP_086585081.1-Vibrio-parahaemolyticus:0.0297083374,WP_217557493.1-Vibrio-metschnikovii:0.0851701845)88.1/100:0.0598391729)76.4/98:0.0450221281,((WP_068331709.1-Photobacterium-jeanii:0.2139805293,WP_077679840.1-Vibrio-kanaloae:0.1164346472)13.3/90:0.0547826994,(WP_031414678.1-Vibrio-parahaemolyticus:0.1572337164,WP_248468589.1-Shewanella-sp.-1CM18E:0.1901803280)50.7/100:0.0864600278)80.3/96:0.0592455443)85.8/94:0.0368283327)75.1/56:0.0151805205,((((((((WP_155660100.1-Aliivibrio-fischeri:0.1102717840,WP_095457151.1-Vibrio:0.1231055753)89.8/100:0.0579192931,(WP_252020469.1-Vibrio-sp.-SCSIO-43136:0.1198947618,WP_243584106.1-Vibrio-splendidus:0.2437025230)81.1/100:0.0460904958)86.6/100:0.0688941181,WP_166696422.1-Providencia-rettgeri:0.1938244914)97/100:0.1412255128,WP_172561824.1-Vibrio-furnissii:0.1254932198)78.1/15:0.0354281346,((WP_165312521.1-Vibrio-ziniensis:0.1319055096,(WP_065296243.1-Vibrio-natriegens:0.1187243190,(WP_009843931.1-Vibrio-sp.-AND4:0.0822363088,WP_017094106.1-Vibrio-splendidus:0.1243657843)75.9/100:0.0400380855)89.3/100:0.0605719055)74.1/48:0.0333570748,((WP_217542775.1-Vibrio-metschnikovii:0.0670422788,WP_113593871.1-Vibrio:0.1116143746)97.6/100:0.1236187656,WP_238819535.1-Vibrio-furnissii:0.1356494749)86.1/94:0.0503493995)68.1/42:0.0167569270)48.2/14:0.0349622915,(WP_047110243.1-Vibrio-vulnificus:0.3138544743,WP_102313365.1-Vibrio-cyclitrophicus:0.1930949180)96.4/99:0.1357996809)0/28:0.0154913067,WP_143680610.1-Vibrio-furnissii:0.1605066789)75.3/28:0.0261478162,((((WP_158107962.1-Vibrio-furnissii:0.1458127483,WP_185890410.1-Vibrio-vulnificus:0.1876455672)83.4/99:0.0351888521,WP_172855536.1-Vibrio-coralliilyticus:0.3576141435)74.5/52:0.0261057333,WP_225486226.1-Vibrio-vulnificus:0.2005295327)12.6/51:0.0431844855,(WP_234235026.1-Shewanella-sp.-AS16:0.2533897846,WP_088905011.1-Shewanella-marisflavi:0.2765643370)85.3/40:0.1191850463)0/31:0.0070260984)16.4/31:0.0061316600)87.6/86:0.0353671228,(((WP_211915530.1-Vibrio-sp.-B1REV9:0.1409942656,WP_017420706.1-Vibrio-vulnificus:0.1310488681)61.9/100:0.0168423774,(WP_192928291.1-Vibrio-vulnificus:0.0504623970,WP_132698543.1-Vibrio-harveyi-group:0.1271076857)98.3/100:0.1236930207)93.4/100:0.0727902426,WP_206974914.1-Vibrio-parahaemolyticus:0.2330295928)0/83:0.0000022857)86.1/94:0.0204817557,(WP_069533349.1-Vibrio-parahaemolyticus:0.2061428163,WP_229630939.1-Vibrio-coralliilyticus:0.2794546077)67.5/98:0.0230484312)89.8/97:0.0341127377,(((((WP_102563398.1-Vibrio-sp.-10N.286.49.B3:0.1567048162,WP_120030433.1-Vibrio-sinensis:0.1365127348)0/32:0.0327679616,(WP_145570128.1-Vibrio-tasmaniensis:0.1842383842,WP_102475754.1-Shewanella-sp.-10N.286.52.C2:0.2951400393)87.2/88:0.0697413574)59.1/69:0.0410793734,(((WP_039841338.1-Vibrio-harveyi-group:0.1266149357,WP_193167845.1-Vibrio-navarrensis:0.4143051533)76.7/98:0.0992925912,WP_144217665.1-Shewanella-algae:0.3248123162)0/51:0.0238662585,(((((WP_033540599.1-Shewanella:0.1993279549,WP_031815865.1-Vibrio-parahaemolyticus:0.1618468348)86.3/98:0.0667120713,(((WP_064489977.1-Vibrio-parahaemolyticus:0.0597136781,WP_201976964.1-Aeromonas-enteropelogenes:0.1459583172)94/100:0.1176568033,WP_224657569.1-Vibrio-vulnificus:0.1852592164)75.5/98:0.0571958714,WP_193250619.1-Vibrio-parahaemolyticus:0.1966772057)10.5/98:0.0622522297)0/3:0.0000027589,WP_258516089.1-Vibrio-alginolyticus:0.1314024949)59.9/87:0.0344109249,WP_106651046.1-Shewanella-sp.-Pdp11:0.2851780747)21.8/13:0.0197707777,(((WP_103194250.1-Vibrio-vulnificus:0.0494227824,WP_158146950.1-Vibrio-fluvialis:0.1501880068)93/100:0.0811067668,WP_233927909.1-Vibrio-fluvialis:0.1790457153)88.5/90:0.0666066934,(WP_211689658.1-Moritella-sp.-28:0.3950394605,(WP_011716450.1-Shewanella-sp.-ANA-3:0.3573322195,WP_206625521.1-Vibrio-vulnificus:0.1216080251)52.5/99:0.0639104292)85/94:0.0814763410)67.9/65:0.0223519111)91.3/46:0.0562717986)0/12:0.0000020230)85.6/42:0.0333758425,WP_020328057.1-Vibrio-fluvialis:0.1857558453)82.6/85:0.0349228433,(WP_099031861.1-Pseudoalteromonas-sp.-GCY:0.1857266757,WP_072616325.1-Vibrio-vulnificus:0.2173118762)69.4/92:0.0273346659)87.1/93:0.0397633673)73.5/56:0.0099586645,((((WP_268674123.1-Vibrio-scophthalmi:0.2087507842,(((WP_162023628.1-Shewanella-sp.-Arc9-LZ:0.0897423776,WP_248997855.1-Shewanella-marinintestina:0.0627225477)90.9/100:0.1162321831,((WP_009847952.1-Vibrio:0.2204546685,WP_233894267.1-Vibrio-kanaloae:0.1794151244)83.7/100:0.0621904810,(WP_048661352.1-Vibrio-crassostreae:0.0794159643,WP_114634227.1-Vibrio-splendidus:0.0864883454)87/100:0.1081127926)92.1/100:0.1007641418)72.7/96:0.0314429251,WP_238130349.1-Vibrio-cincinnatiensis:0.1990225236)69.2/85:0.0283722581)71.7/74:0.0464848303,((((((((WP_171343915.1-Vibrio-cyclitrophicus:0.1031782712,(WP_102248986.1-Vibrio-tasmaniensis:0.2149683539,WP_135445135.1-Vibrio-tasmaniensis:0.1195984915)97.9/100:0.1205517529)6.7/90:0.0172960582,WP_161753058.1-Vibrio-toranzoniae:0.1923714239)0.9/92:0.0072524835,WP_220058598.1-Shewanella-sp.-FJAT-53555:0.4295994620)87.3/97:0.0498720406,(((WP_102276040.1-Vibrio-lentus:0.1571864478,WP_102457030.1-Vibrio-cyclitrophicus:0.1245461213)84.5/99:0.0454355676,(WP_102503160.1-Vibrio-splendidus:0.2191068601,WP_155677763.1-Aliivibrio-fischeri:0.1170827358)93.2/98:0.0842660752)29.9/61:0.0151244745,(WP_102420589.1-Vibrio-cyclitrophicus:0.1533493776,WP_132940416.1-Vibrio-crassostreae:0.1603766160)48.9/64:0.0323558102)91.4/97:0.0635740198)76.7/84:0.0157161184,(((((WP_172851426.1-Vibrio-coralliilyticus:0.2327995022,WP_182742106.1-Shewanella-sp.-SG44-6:0.2820226778)85.2/97:0.0738302640,WP_108186198.1-Vibrio-splendidus:0.1506130252)83.6/83:0.0549874318,WP_046209502.1-Vibrio:0.1972215089)80.5/70:0.0242956554,WP_017085924.1-Vibrio-splendidus:0.1916892277)78.7/67:0.0271594109,((WP_239850080.1-Shewanella-sp.-Isolate8:0.0974603902,WP_121852788.1-Parashewanella-spongiae:0.2940953716)0.7/20:0.0448082425,WP_069440067.1-Pseudoalteromonas-tetraodonis:0.3564308953)93.6/95:0.1145327316)77.1/86:0.0315604820)75.2/78:0.0126632997,(WP_139685731.1-Vibrio-tasmaniensis:0.1642376203,(((WP_061039794.1-Vibrio-coralliirubri:0.0960391703,(WP_044365158.1-Vibrio-fluvialis:0.3818961585,WP_025523961.1-Vibrio-parahaemolyticus:0.1917700991)87.7/100:0.0654255120)87.8/98:0.0808238465,WP_065194916.1-Photobacterium-phosphoreum:0.1948010507)84.1/100:0.0679158427,WP_248386338.1-Vibrio-sp.-ED004:0.2851482763)37.6/89:0.0263027881)89.8/86:0.0829410752)73.4/81:0.0164112429,WP_248368760.1-Vibrio-sp.-1CM8B:0.1879788527)87.4/82:0.0288666158,((((((((WP_155682312.1-Aliivibrio-fischeri:0.1245214121,WP_012532661.1-Aliivibrio-fischeri:0.1414618737)84.4/99:0.0653468859,WP_102428030.1-Vibrio-sp.-10N.261.54.E10:0.1684367466)80.5/98:0.0422663690,(WP_108162474.1-Vibrio-splendidus:0.0297250777,WP_004742905.1-Vibrio-tubiashii:0.0735887949)99.7/100:0.1501819305)79.2/37:0.0430978621,WP_017069285.1-Vibrio-crassostreae:0.0956575639)6/49:0.0191069018,WP_016768872.1-Vibrio-cyclitrophicus:0.1635072582)72.8/81:0.0621073585,(((WP_005597034.1-Vibrio-scophthalmi:0.1770692879,(WP_108176621.1-Vibrio-splendidus:0.0749765614,WP_239918548.1-Vibrio-sp.-Isolate22:0.0205152594)93.3/100:0.0953917144)81/96:0.0437294425,WP_135445143.1-Vibrio-tasmaniensis:0.2140141220)81.1/80:0.0570207435,(((WP_012551417.1-Aliivibrio-salmonicida:0.0968666037,WP_012276595.1-Shewanella-halifaxensis:0.0848373184)85.7/100:0.0821073664,WP_135385147.1-Vibrio-tasmaniensis:0.1807185884)90.5/94:0.0726356289,WP_265133188.1-Vibrio-splendidus:0.1271431864)51.8/60:0.0166110476)88.4/86:0.0631589346)33.1/32:0.0261475868,((WP_130049361.1-Aliivibrio-finisterrensis:0.0939046142,(WP_019826652.1-Vibrio-splendidus:0.1566674845,WP_102571993.1-Vibrio:0.1690419485)37.6/97:0.0420736476)93.9/91:0.0833114938,(WP_260256886.1-Vibrio-sp.-NC2:0.1197284748,(WP_102298256.1-Vibrio-cyclitrophicus:0.1292236251,WP_248388778.1-Vibrio-cyclitrophicus:0.2041130699)82.5/59:0.0347739048)0/37:0.0000029850)84.2/43:0.0451547547)41.9/61:0.0158949127,((WP_258428294.1-Vibrio-sp.-RM-69-4:0.1645113574,(WP_193834777.1-Vibrio-sp.-OPT20:0.0632245004,(WP_029865567.1-Vibrio-parahaemolyticus:0.0770513668,WP_213967907.1-Vibrio-alginolyticus:0.0352802729)99.3/100:0.1039656801)75.6/100:0.0232832025)90.2/99:0.0605825963,(((WP_004733311.1-Vibrio-splendidus:0.1237322221,WP_171315631.1-Vibrio-splendidus:0.1647886242)2.3/47:0.0277437995,WP_102353859.1-Vibrio-cyclitrophicus:0.1778524835)71.1/93:0.0490392634,WP_102385704.1-Vibrio-cyclitrophicus:0.1867506980)61.1/85:0.0498155269)86.2/77:0.0401840815)77.7/61:0.0156541460)0/28:0.0000027976)92.9/79:0.0305720130,(((WP_059021773.1-Vibrio-coralliirubri:0.0531183870,WP_135380937.1-Vibrio-tasmaniensis:0.1567192775)86.4/97:0.0718425490,WP_182796407.1-Vibrio-sp.-SG41-7:0.2221532887)66.9/79:0.0453143755,((((WP_161672404.1-Vibrio-toranzoniae:0.1121878662,WP_017040654.1-Vibrio-genomosp.-F10:0.2028294850)90.3/98:0.0927664826,(WP_137008959.1-Vibrio-kanaloae:0.1666785543,WP_135353160.1-Aliivibrio-fischeri:0.1050528006)79.4/100:0.0525336718)84.8/97:0.0470631332,WP_248638627.1-Vibrio-crassostreae:0.1650012245)78.4/88:0.0332103303,((WP_155675904.1-Aliivibrio-fischeri:0.1290422334,WP_017105169.1-Vibrio:0.1777692405)46/87:0.0285152802,((WP_102459739.1-Vibrio-lentus:0.1351876918,WP_102363333.1-Vibrio-cyclitrophicus:0.1607583819)48.7/63:0.0402790022,(WP_102554090.1-Vibrio-splendidus:0.1700231900,WP_130314193.1-Vibrio-vulnificus:0.2171409689)0/50:0.0000023966)88.1/82:0.0475503604)72.9/89:0.0343843697)84.7/89:0.0323353520)72.5/76:0.0122672774)83.5/92:0.0264380427,WP_058428281.1-Pseudoalteromonas:0.3225015633)63.9/66:0.0305132954)19.8/50:0.0088601399,(WP_208844179.1-Pseudoalteromonas-xiamenensis:0.1792650451,WP_233976434.1-Vibrio:0.2206106167)38.2/66:0.0430014871)91.8/96:0.0426832639)75.3/62:0.0107386546,((((WP_102457123.1-Vibrio-breoganii:0.1665996340,(WP_172380549.1-Vibrio-sp.-Vb339:0.1383446153,WP_102443327.1-Vibrio-breoganii:0.1318278467)90.2/95:0.1077690417)81.9/75:0.0481439553,WP_077647682.1-Salinivibrio-sp.-IB643:0.1408242606)21.5/71:0.0215397132,(WP_045031875.1-Photobacterium-phosphoreum:0.0280587277,WP_107213458.1-Photobacterium-phosphoreum:0.0461460382)100/100:0.2424563212)0/64:0.0000020228,(((((((WP_017027407.1-Vibrio-breoganii:0.1353424070,(WP_102532836.1-Vibrio-sp.-10N.286.48.B7:0.1506759140,WP_254101081.1-Alteromonas-sp.-LMIT007:0.1905401315)87.6/100:0.0705546456)83.2/99:0.0442760086,WP_108083222.1-Vibrio-sp.-ZF-223:0.1874962857)31.8/13:0.0097892987,(WP_108189654.1-Vibrio-splendidus:0.2130938723,(WP_102448490.1-Vibrio-splendidus:0.1360835419,WP_243579549.1-Vibrio-splendidus:0.2489957289)78.5/99:0.0361273806)74.9/13:0.0396790178)88.6/79:0.0398478716,((WP_138589904.1-Pseudoalteromonas-aurantia:0.1687823977,((WP_031827242.1-Vibrio-parahaemolyticus:0.3387463654,WP_194091044.1-Vibrio-hibernica:0.2135569640)16.8/75:0.0511625161,WP_110456309.1-Shewanella-algidipiscicola:0.2532575496)84.5/73:0.0622159214)79.5/70:0.0373586032,(WP_011638275.1-Shewanella-frigidimarina:0.2093318115,WP_012549058.1-Aliivibrio-salmonicida:0.2676070950)51.1/54:0.0248507274)83.1/45:0.0369518466)83.4/29:0.0180152390,((WP_248991952.1-Shewanella-profunda:0.1335289891,(WP_220774251.1-Shewanella-glacialipiscicola:0.0034251706,WP_182751750.1-Shewanella-sp.-SG44-2:0.2287889947)98.5/100:0.1887307567)59.7/77:0.0620919755,WP_059744187.1-Shewanella-frigidimarina:0.2908553329)85/76:0.0453442507)74.8/17:0.0123347972,(((WP_061023752.1-Vibrio-splendidus:0.0965404565,WP_220768829.1-Shewanella-sp.-MBTL60-007:0.1438954972)84.3/81:0.0404117513,(WP_063660043.1-Aliivibrio-fischeri:0.2061572834,WP_204999274.1-Parashewanella-hymeniacidonis:0.2581281695)16.3/37:0.0126098957)81.7/34:0.0414060897,WP_182754472.1-Shewanella-sp.-SG41-3:0.1996628821)65.1/23:0.0097496291)59.8/40:0.0238727343,(WP_238813939.1-Vibrio-vulnificus:0.1139644885,WP_222242368.1-Pseudoalteromonas-piscicida:0.5036653282)77.3/98:0.0409910649)95.8/80:0.0565837679)89.5/69:0.0327124568)79.6/81:0.0178096423,(WP_218529053.1-Shewanella-sp.-NIFS-20-20:0.1737335375,WP_225483056.1-Vibrio-vulnificus:0.0755688101)99.5/99:0.1913246296)77.2/52:0.0092069094,((((((((WP_193255073.1-Vibrio-navarrensis:0.0561872331,WP_258469755.1-Vibrio-alginolyticus:0.1229143660)98.3/100:0.1332196497,(WP_095430409.1-Vibrio-vulnificus:0.0827489844,WP_114994956.1-Grimontia-hollisae:0.2449313178)93.4/100:0.0945354727)82.8/91:0.0548519500,((((WP_268809473.1-Vibrio-parahaemolyticus:0.1568979921,WP_023404414.1-Vibrio-halioticoli:0.2972780968)79.9/96:0.0516890108,WP_028024862.1-Enterovibrio-calviensis:0.2300723706)32/68:0.0247638321,WP_017007824.1-Enterovibrio-norvegicus:0.2188845992)77.8/74:0.0290995003,(WP_211678393.1-Moritella-sp.-36:0.2933721869,(WP_138627230.1-Pseudoalteromonas-sp.-S4741:0.2338951563,WP_269377025.1-Vibrio-sinaloensis:0.2051704577)83.1/100:0.0711043059)88.4/100:0.0720452787)74.5/93:0.0382037296)87.9/85:0.0405076483,WP_068898547.1-Enterovibrio-pacificus:0.2144232497)71.3/82:0.0078079493,(WP_069212356.1-Vibrio-cholerae:0.0759230031,WP_202641945.1-Vibrio-fluvialis:0.1180244645)99.4/99:0.1641929495)82.2/75:0.0344984789,(((((((WP_041510674.1-Shewanella:0.0320129045,(WP_187575439.1-Shewanella-algae:0.0965565553,(WP_208158397.1-Shewanella-algae:0.1021379290,WP_208157589.1-Shewanella-algae:0.0442503581)98.3/100:0.0903210853)44/82:0.0158215005)94.4/98:0.0867833249,(WP_048899071.1-Photobacterium-swingsii:0.2197805643,(((WP_045569365.1-Vibrio:0.0906618255,WP_039437343.1-Vibrio-navarrensis:0.0571183477)98.2/100:0.1272766133,WP_176299354.1-Vibrio-sp.-05-20-BW147:0.1857201855)79.9/100:0.0650098266,(WP_140233673.1-Shewanella-polaris:0.2705652090,WP_192022372.1-Shewanella-sp.-WPAGA9:0.2677154124)69.2/83:0.0796098978)54.3/83:0.0685176907)60.8/70:0.0598751217)51.1/60:0.0207696318,(WP_101201417.1-Paraglaciecola-sp.-MB-3u-78:0.2660808179,WP_039484180.1-Vibrio-sinaloensis:0.1743526488)91.2/90:0.0793959361)27.7/79:0.0122224715,WP_036979211.1-Providencia-alcalifaciens:0.3714865896)70.8/39:0.0217991770,(WP_107294031.1-Photobacterium-phosphoreum:0.0440843377,WP_107276976.1-Photobacterium-sp.-NCIMB-13483:0.0827890527)99.9/100:0.1743257397)84/39:0.0351321058,((((((((((WP_065208126.1-Photobacterium-phosphoreum:0.1612648402,((WP_232585319.1-Photobacterium-iliopiscarium:0.1188748766,(WP_232582982.1-Photobacterium-phosphoreum:0.0680670630,((WP_256383785.1-Photobacterium-toruni:0.0566677706,WP_232578536.1-Photobacterium-carnosum:0.0668871259)0/95:0.0000020322,WP_107228404.1-Photobacterium-phosphoreum:0.1178678357)93.2/100:0.0501849869)94.2/100:0.0665796786)77.5/75:0.0251917021,WP_107228425.1-Photobacterium-phosphoreum:0.2517113982)66.5/91:0.0311570436)87.9/81:0.0630935521,((WP_232575020.1-Photobacterium-carnosum:0.1598337097,(WP_045045404.1-Photobacterium-kishitanii:0.0456938975,(WP_045030671.1-Photobacterium-phosphoreum:0.0615660612,WP_107264793.1-Photobacterium-phosphoreum:0.0332826186)80.3/60:0.0221480397)94.5/100:0.0790432600)88.5/40:0.0576857237,(WP_232600381.1-Photobacterium-carnosum:0.0858887656,(WP_107213932.1-Photobacterium-phosphoreum:0.1277115786,WP_065173166.1-Photobacterium-kishitanii:0.1946156262)90.8/99:0.0810877518)78.1/46:0.0305050312)4.9/18:0.0086875297)82.2/58:0.0297046016,WP_237041371.1-Photobacterium-damselae:0.1184251864)33.3/44:0.0262430130,((((((WP_151182432.1-Photobacterium-damselae:0.1708029952,WP_106261939.1-Photobacterium-damselae:0.1092771144)32.1/89:0.0267467925,WP_115426223.1-Photobacterium-damselae:0.0837835615)61.3/98:0.0330614173,WP_053987023.1-Photobacterium-leiognathi:0.1393368624)4.3/71:0.0208163490,WP_086367281.1-Photobacterium-damselae:0.0788567615)83.5/88:0.0399609795,((WP_238087945.1-Photobacterium-damselae:0.0800619158,(WP_065172005.1-Photobacterium-damselae:0.1072146434,WP_107185088.1-Photobacterium-leiognathi:0.0952576656)82.6/100:0.0291559522)52.6/94:0.0327727251,WP_151170663.1-Photobacterium-damselae:0.0616166709)11.6/62:0.0222695246)82/80:0.0326265742,WP_045063675.1-Photobacterium-leiognathi:0.0967639194)92.4/88:0.0605907153)0.9/16:0.0205253142,((WP_107293979.1-Photobacterium-phosphoreum:0.1709678132,(WP_045039948.1-Photobacterium-iliopiscarium:0.2306681022,((WP_065176337.1-Photobacterium-aquimaris:0.0832490669,WP_107199726.1-Photobacterium-phosphoreum:0.0974708368)54.6/100:0.0275759460,WP_023931546.1-Photobacterium-leiognathi:0.1697204231)57.2/94:0.0485309361)89.8/100:0.0713428365)45.9/88:0.0641904300,(WP_087853896.1-Photobacterium-andalusiense:0.0532315244,WP_107274737.1-Photobacterium-phosphoreum:0.1289277851)93.4/100:0.0843097913)80.5/67:0.0394076866)85.9/88:0.0359670195,(WP_237042700.1-Photobacterium-damselae:0.0931205934,(WP_013365656.1-Enterobacter:0.2370063239,(((WP_258988101.1-Scandinavium-hiltneri:0.0919313966,WP_181620050.1-Enterobacter-sp.-RHBSTW-00994:0.0674403651)93.9/100:0.0736662224,WP_193129600.1-Klebsiella-aerogenes:0.1027997088)94.7/100:0.0872337608,WP_214579582.1-Enterobacter-asburiae:0.1079609040)82.5/99:0.0650596656)99.9/100:0.2551856648)74.6/93:0.0243458284)87.5/86:0.0395321339,((WP_175242188.1-Vibrio-cholerae:0.0889506308,(WP_199351799.1-Vibrio-cholerae:0.1041059052,(WP_095464062.1-Vibrio-cholerae:0.1060068675,WP_149560528.1-Vibrio-cholerae:0.1288527243)11.1/51:0.0609374899)52.8/94:0.0334332043)80.4/94:0.0354391170,(WP_000864154.1-Vibrio-cholerae:0.0722957368,(WP_102786270.1-Vibrio-cholerae:0.0874855847,WP_095478449.1-Vibrio:0.1186595786)72.3/91:0.0344486845)87.2/95:0.0470857028)19.2/60:0.0198368121)85.1/83:0.0435700969,((WP_161119826.1-Photobacterium-lucens:0.1681015796,WP_017004373.1-Enterovibrio-norvegicus:0.2760712344)20.9/85:0.0322524001,(((WP_230432956.1-Plesiomonas-shigelloides:0.1356055166,WP_222763364.1-Vibrio-fluvialis:0.1616541533)73.9/100:0.0270254581,(WP_006646466.1-Photobacterium-sp.-SKA34:0.2732056533,((WP_107253996.1-Photobacterium-indicum:0.1742309608,WP_107295906.1-Photobacterium-phosphoreum:0.1994621638)89.9/100:0.0799024874,WP_251878087.1-Grimontia-sp.-020920N:0.1300298486)62.5/98:0.0130921532)86.8/86:0.0460659202)76.4/67:0.0250240242,(WP_171138527.1-Vibrio-plantisponsor:0.0973645429,WP_102966424.1-Vibrio:0.0956842760)99.4/100:0.1765801237)84.4/84:0.0392849619)66.4/70:0.0414911299)71.4/71:0.0147001426,(((WP_208891742.1-Vibrio-anguillarum:0.1941232132,WP_148533341.1-Vibrio-cholerae:0.0726805163)83.3/99:0.0483317182,(WP_161680902.1-Proteus:0.5154323296,WP_107294203.1-Photobacterium-sp.-GB-50:0.1463562560)85.7/78:0.0712629678)71.4/77:0.0362226757,(WP_000864156.1-Vibrio-cholerae:0.0839728073,WP_000864152.1-Vibrio-cholerae:0.1225472017)93.7/99:0.0837749821)78.7/76:0.0326994850)75.8/75:0.0120869106,(WP_094127815.1-Vibrio-sp.-V09_P4A23P171:0.1327826691,WP_199867560.1-Vibrio-anguillarum:0.1149629931)97.4/100:0.1070468875)78.1/51:0.0188144261)79.1/58:0.0154230176,(((((((((WP_239941760.1-Vibrio-mediterranei:0.1378387707,WP_062462511.1-Vibrio-mediterranei:0.1285781374)43/75:0.0336973560,WP_221070344.1-Vibrio-alfacsensis:0.2086250869)74.7/72:0.0215079734,(WP_171392669.1-Vibrio:0.1719047225,WP_101064017.1-Moritella-sp.-Urea-trap-13:0.3044481571)77.6/87:0.0246333939)75/73:0.0130696930,(WP_102532726.1-Vibrio-sp.-10N.286.48.B7:0.1047160239,WP_250617847.1-Vibrio-sp.-S4B1:0.1181990676)87.1/98:0.0457931147)73.6/73:0.0162542182,(WP_252012797.1-Vibrio-sp.-SCSIO-43140:0.1624795589,WP_252012765.1-Vibrio-sp.-SCSIO-43140:0.1553439817)85.8/97:0.0538980114)96/96:0.0914950261,(WP_268679810.1-Vibrio-barjaei:0.1391623464,(WP_261894433.1-Vibrio-porteresiae:0.2645468329,((WP_117235674.1-Vibrio-maerlii:0.2254762759,WP_265676699.1-Vibrio-sp.-ZSDZ65:0.1487097176)73/91:0.0223853610,WP_103882085.1-Vibrio-hangzhouensis:0.0594748550)78.2/98:0.0413166843)92.6/96:0.0810161293)0/74:0.0000026480)85.7/85:0.0493481764,((WP_261600599.1-Photobacterium-sp.-TY1-4:0.1175778452,(WP_114765998.1-Vibrio-rhodolitus:0.1452312061,WP_182108830.1-Vibrio-marinisediminis:0.1299979592)94.7/100:0.1220691900)75.1/95:0.0268993832,(((WP_170899585.1-Vibrio-sp.-3-2-1:0.1358603131,WP_017221010.1-Moritella-dasanensis:0.1592928071)86.8/100:0.0565220571,WP_107284330.1-Photobacterium-lipolyticum:0.1340899043)90/98:0.0645235833,(WP_159197271.1-Escherichia-coli:0.1420268714,WP_088133934.1-Vibrio-gazogenes:0.3262429041)0/53:0.0049816987)83.4/87:0.0258118055)85.9/84:0.0435593569)88.2/74:0.0442608440,((WP_020431327.1-Vibrio-fluvialis:0.1258401823,WP_055453311.1-Vibrio-fluvialis:0.0541026417)97.4/100:0.1277574668,(WP_045898906.1-Enterobacteriaceae:0.1790674208,(WP_182744595.1-Shewanella-sp.-SR44-4:0.2960657807,WP_152797327.1-Salinivibrio-sp.-VYel1:0.2422216644)91.8/95:0.1232943099)0/61:0.0064104035)71/70:0.0304911513)40.7/34:0.0108698229,((((((((WP_211647400.1-Photobacterium-sp.-GJ3:0.0565150922,WP_120511428.1-Photobacterium-salinisoli:0.0663885498)94.9/99:0.0844234761,((WP_050928752.1-Yersinia-enterocolitica:0.3004294891,WP_012883389.1-Dickeya-parazeae:0.2891571300)70.6/99:0.0697274499,((WP_035339894.1-Dickeya:0.0371380124,WP_125258312.1-Dickeya-lacustris:0.1017014064)99.8/100:0.2542077797,WP_042868619.1-Dickeya:0.0943718511)95.4/100:0.1250005391)95.4/98:0.1257555212)0/87:0.0000024172,WP_046218796.1-Photobacterium-halotolerans:0.2429200280)73.3/89:0.0388519913,((WP_015854950.1-Musicola-paradisiaca:0.0642238246,WP_192457620.1-Musicola-keenii:0.0882651426)100/100:0.3835595617,((WP_200225058.1-Rahnella-aceris:0.1485219947,WP_195814554.1-Rahnella:0.1391432435)87.8/100:0.0524851878,WP_169403689.1-Rouxiella-aceris:0.1032096626)95.3/100:0.1403745499)84.6/93:0.0464669657)87.2/90:0.0283519305,(WP_047463293.1-Vibrio-fluvialis:0.1510862269,WP_148560072.1-Vibrio-mimicus:0.1973083005)92.6/100:0.0944112841)78.4/88:0.0146269532,WP_124938567.1-Vibrio-viridaestus:0.2139409256)40.3/57:0.0141772074,(WP_112404745.1-Vibrio-diazotrophicus:0.0646483687,WP_202661182.1-Vibrio-fluvialis:0.0704260416)96.6/100:0.0876215601)15.3/45:0.0186506752,((((WP_263100201.1-Kosakonia-sp.-ML.JS2a:0.0387498029,WP_200550470.1-Kosakonia-sp.-LAM2021:0.1104568578)97.7/100:0.1456430608,(WP_094109047.1-Lonsdalea-iberica:0.1391813771,(WP_026742149.1-Lonsdalea-quercina:0.0369297222,WP_094117837.1-Lonsdalea-britannica:0.0474106785)94.8/100:0.1137358167)84.9/95:0.0537172171)86.6/93:0.0734946257,WP_172533652.1-Vibrio-aestuarianus:0.4312789059)60.7/68:0.0495524703,(WP_038178862.1-Vibrio-rhizosphaerae:0.1855251265,WP_107187026.1-Photobacterium-kishitanii:0.2168021087)46.2/76:0.0278518601)85.4/86:0.0558735479)88.8/91:0.0395452117)78.7/92:0.0105391643)77.3/55:0.0159983736)88.9/79:0.0382288133,(((((WP_239872404.1-Shewanella-insulae:0.1503269448,WP_101040106.1-Psychromonas-sp.-MB-3u-54:0.3124013996)61.2/52:0.0336028274,WP_238097025.1-Vibrio-cincinnatiensis:0.2028586512)77.1/52:0.0349423918,WP_118894008.1-Vibrio-vulnificus:0.2003763828)75.3/53:0.0476580210,WP_235060641.1-Vibrio-sp.-SS-MA-C1-2:0.3639110428)76.5/66:0.0388101966,(WP_045460498.1-Vibrio-hyugaensis:0.1922169868,WP_220051788.1-Shewanella-sp.-FJAT-51860:0.1920944686)76.7/90:0.0684395475)53.9/50:0.0199026948)73.9/20:0.0106694400,((WP_135455718.1-Vibrio-echinoideorum:0.1734283274,((WP_137027437.1-Vibrio-kanaloae:0.0292799957,WP_108117598.1-Vibrio-splendidus:0.0950393107)84.9/100:0.0494307654,WP_137025093.1-Vibrio-kanaloae:0.2077126614)12.7/56:0.0205356988)80.8/55:0.0443230093,WP_017449116.1-Vibrio-parahaemolyticus:0.1849142214)81.8/82:0.0434238131)85.1/90:0.0240269761)79.5/59:0.0137029942,(WP_213972370.1-Vibrio-alginolyticus:0.2322828911,WP_017024716.1-Vibrio-rumoiensis:0.2018493322)94.5/91:0.0815231925)75.9/90:0.0167608429,(WP_014230611.1-Vibrio-sp.-EJY3:0.1355308748,WP_243025268.1-Vibrio-cincinnatiensis:0.6173018658)54.4/51:0.0631289469)80.9/69:0.0223363020)75.9/23:0.0280537809,((((((WP_237041393.1-Photobacterium-damselae:0.1696321379,WP_080158029.1-Photobacterium-piscicola:0.1890039858)85.7/82:0.0695180679,(WP_047876260.1-Photobacterium-aphoticum:0.1483237414,(WP_188234612.1-Vibrio-sp.-Y2-5:0.0886367223,WP_057556016.1-Vibrio-cholerae:0.0136985359)99.9/100:0.2696948818)83.3/91:0.0864165731)82.7/66:0.0432570521,WP_102395415.1-Enterovibrio-norvegicus:0.1741071531)0/9:0.0225771729,WP_045069930.1-Photobacterium-leiognathi:0.2279516044)64.1/14:0.0282923627,(((WP_267741605.1-Vibrio-cholerae:0.1193253635,WP_148560094.1-Vibrio-mimicus:0.2401809521)58.5/63:0.0510983297,((WP_172567459.1-Vibrio-fluvialis:0.0720730770,WP_005395829.1-Vibrio-diabolicus:0.3800238108)79.8/94:0.0460135540,WP_199354891.1-Vibrio-cholerae:0.0551312085)95.4/94:0.1019653691)14.5/53:0.0302108151,WP_102137258.1-Vibrio-cholerae:0.1409836838)86.2/86:0.0564083092)79.2/27:0.0351755249,(WP_017008437.1-Enterovibrio-norvegicus:0.1424713277,(WP_232612305.1-Photobacterium-phosphoreum:0.4130548969,WP_136989485.1-Vibrio-sp.-F13:0.3326925548)97.3/99:0.2051506193)80.9/43:0.0405394257)78.6/32:0.0255505631)75.8/3:0.0138466148,((((WP_046223474.1-Vibrio-sp.-ECSMB14106:0.1232727553,WP_102562373.1-Vibrio-breoganii:0.1819094176)79.3/100:0.0448870523,WP_218382835.1-Photobacterium-sp.-WH24:0.2224145062)54/66:0.0488732020,((WP_102546929.1-Vibrio-breoganii:0.2043951223,WP_102461639.1-Vibrio-breoganii:0.1518663120)73.8/97:0.0318119162,(WP_135455667.1-Vibrio-echinoideorum:0.1631877573,(WP_141346804.1-Vibrio-inusitatus:0.2048232374,WP_102282524.1-Vibrio-cyclitrophicus:0.2054453951)84.2/99:0.0562282752)61.1/42:0.0132458733)83.4/66:0.0632738342)72.2/99:0.0342800965,WP_193290728.1-Vibrio-parahaemolyticus:0.3544308934)88.1/62:0.0699321229)84.3/71:0.0233786037,(((WP_232600401.1-Photobacterium-carnosum:0.1607338163,(WP_060997345.1-Photobacterium-aquimaris:0.1729090823,((WP_232603643.1-Photobacterium-carnosum:0.1185017802,WP_232615715.1-Photobacterium-carnosum:0.1292810920)76.1/93:0.0235059846,WP_237309237.1-Vibrio:0.5028946692)66/86:0.0138012351)44.7/79:0.0591701816)81.1/88:0.0308130755,WP_176284769.1-Photobacterium-damselae:0.2310002353)88.6/84:0.0513844620,((WP_188008910.1-Grimontia-hollisae:0.1500803297,WP_065173140.1-Photobacterium-kishitanii:0.1888571891)88.9/98:0.0949021640,(WP_108452928.1-Enterobacter-cloacae:0.4988713030,WP_133040538.1-Shewanella-fodinae:0.3358232157)89.2/78:0.1038800788)60.9/55:0.0240035108)77.9/59:0.0196988012)75/9:0.0136080020,((((((((WP_249572008.1-Vibrio-cholerae:0.1113522136,WP_088132129.1-Vibrio-cholerae:0.1533933764)52.3/95:0.0424070894,WP_255122387.1-Vibrio-cholerae:0.1527770868)76.1/94:0.0275623717,(WP_000864153.1-Vibrio-cholerae:0.0194036970,WP_251878143.1-Grimontia-sp.-020920N:0.2162367210)96.2/96:0.1239580745)76.3/74:0.0413719317,WP_045618469.1-Vibrio-vulnificus:0.1700910697)0/21:0.0000027091,(WP_152138288.1-Plesiomonas-shigelloides:0.1916367213,((WP_148519618.1-Vibrio-cholerae:0.0358245613,(WP_167570263.1-Aeromonas-veronii:0.0718517491,WP_000864158.1-Vibrio-metoecus:0.0457413542)93.4/100:0.0377798980)83/100:0.0229681817,WP_148520854.1-Vibrio-cholerae:0.0369473082)93.3/100:0.0941184201)80.6/95:0.0427395179)84.7/80:0.0370209861,(WP_103155729.1-Vibrio-vulnificus:0.1615391764,WP_129120576.1-Veronia-nyctiphanis:0.2362659184)51.2/79:0.0124758651)83.8/81:0.0243122008,(((((WP_243584123.1-Vibrio-splendidus:0.1306191202,WP_259574191.1-Shewanella-baltica:0.3196941263)55.3/65:0.0726817148,((WP_102462022.1-Vibrio-splendidus:0.0892684646,WP_187463684.1-Photobacterium-sp.-BZF1:0.0674610624)94.5/100:0.1246081629,WP_029788868.1-Vibrio:0.3660028850)20.3/42:0.0298589884)78.4/72:0.0463862156,(((((((WP_261298106.1-Shewanella-sp.-SP1S2-7:0.1249913763,WP_167678467.1-Shewanella-aestuarii:0.1182962794)100/100:0.3346133809,WP_248994245.1-Shewanella-gaetbuli:0.2883230266)92.2/100:0.1151098840,WP_212599444.1-Shewanella-sp.-M16:0.3421347938)87.7/99:0.0886658890,WP_261923504.1-Shewanella-sp.-NFH-SH190041:0.2578037847)84.2/99:0.0593903707,((WP_069589493.1-Salinivibrio-sp.-DV:0.1141292920,WP_077662143.1-Salinivibrio-proteolyticus:0.0180687914)74.5/68:0.0222783908,((WP_077659434.1-Salinivibrio-kushneri:0.1651277337,WP_077659452.1-Salinivibrio-kushneri:0.0401293033)28.5/68:0.0280771105,WP_269578450.1-Salinivibrio-kushneri:0.1033294380)84.1/96:0.0620423515)100/100:0.3434279576)84.2/80:0.0617819787,(WP_213886549.1-Vibrio-alginolyticus:0.2358929648,WP_158173598.1-Vibrio-alginolyticus:0.1219773433)94.3/100:0.1290092734)34.4/64:0.0353251395,((WP_041156219.1-Vibrio-mytili:0.1841606789,(WP_047480300.1-Vibrio-campbellii:0.0711659345,WP_045494469.1-Vibrio-hyugaensis:0.0576734233)98.3/100:0.1502356440)93.1/100:0.1153527787,WP_102523418.1-Vibrio-tapetis:0.2446169878)80.1/84:0.0669262822)86/65:0.0557854246)77/77:0.0243507613,WP_163929979.1-Paraferrimonas-sp.-SM1919:0.1708886206)32.6/78:0.0494026673,WP_087845049.1-Photobacterium-malacitanum:0.1919348869)85.5/98:0.0476185312)73.4/76:0.0119989683,(WP_161129723.1-Photobacterium-lucens:0.1734777478,(WP_084988554.1-Vibrio:0.2122939838,WP_206634697.1-Vibrio-vulnificus:0.1698620297)79.1/100:0.0562805498)90.3/94:0.0685383202)80.1/72:0.0309644803)81.4/91:0.0715699718,(WP_232581009.1-Photobacterium-phosphoreum:0.6690659364,(WP_073514120.1-Vibrio-parahaemolyticus:0.0822951464,WP_032481483.1-Vibrio:0.0138437000)99.6/100:0.7324493195)99.4/100:0.4184274484)93.8/98:0.1596097790,((WP_162556830.1-Proteus:0.1048690104,WP_161779122.1-Proteus-sp.-G2675:0.1023064944)99.5/100:0.2493136636,WP_236366479.1-Aeromonas-veronii:0.2736023659)0/2:0.0000020108)97.8/98:0.1593084478)31.9/29:0.0531183327,WP_225633057.1-Aeromonas-enteropelogenes:0.2904053306)90/35:0.0688493405,WP_102949229.1-Aeromonas-caviae:0.3813132863)64.9/81:0.0525731589,((WP_071788415.1-Franconibacter-pulveris:0.5309901629,(WP_020232007.1-Escherichia-coli:0.3620690509,(WP_228338545.1-Enterobacter-hormaechei:0.1598677426,(WP_109845273.1-Enterobacter:0.0616599612,WP_150146763.1-Enterobacter-roggenkampii:0.0520914482)88.3/100:0.1099564684)98.7/100:0.2339168550)91.3/100:0.1297020005)95.2/98:0.1636972880,((((((((((((((WP_051234278.1-Halomonas-halocynthiae:0.8535524595,WP_110648656.1-Salinicola-peritrichatus:0.3933305731)98.9/95:0.3502153580,(WP_081193716.1-Halomonas-sp.-BC1:0.3230432197,WP_035566812.1-Halomonas:0.2580975641)72.9/82:0.0491480205)72.6/69:0.0327306312,((WP_224850395.1-Marinobacter-nauticus:0.2046747751,WP_110630630.1-Salinicola-salarius:0.1040141203)77.4/100:0.0261800372,((WP_193085952.1-Halomonas-sp.-FME65:0.1602488666,(((WP_159342670.1-Halomonas-meridiana:0.4084122710,WP_253554573.1-Halomonas-sp.-DN3:0.2290418663)0/95:0.0109885868,WP_102589198.1-Halomonas-urumqiensis:0.1360626209)32.6/96:0.0223167630,(WP_193343673.1-Halomonas-elongata:0.0753682854,WP_064701650.1-Halomonas-caseinilytica:0.0185712631)99.8/100:0.1816384272)91.7/93:0.0356015085)0/87:0.0000020143,((((WP_079551663.1-Halomonas-subglaciescola:0.0808543285,WP_195687443.1-Halomonas-sp.-KAO:0.0650873241)87.1/98:0.0234268038,WP_027961801.1-Halomonas-halodenitrificans:0.0393227016)86.7/98:0.0171736709,WP_089675781.1-Halomonas-aquamarina:0.0532213943)30.5/75:0.0093153668,((WP_253557393.1-unclassified-Halomonas:0.0560446572,WP_089676106.1-Halomonas-aquamarina:0.0334488141)0/94:0.0000020359,WP_062362595.1-Halomonas-axialensis:0.0423285565)80.3/77:0.0148813384)94.2/97:0.0451384703)87.2/93:0.0241941038)96.4/99:0.0850742991)90.3/90:0.0456143988,((WP_248234415.1-Marinobacter-bryozoorum:0.1245640260,(WP_228223924.1-Marinobacter-nauticus:0.0720808667,WP_098418752.1-Marinobacter-sp.-LV10MA510-1:0.1562102574)89.5/100:0.0897735461)93.3/100:0.1037542419,(WP_021818206.1-Halomonas-huangheensis:0.1750265559,(WP_149282777.1-Halomonas-sp.-Y2R2:0.0939118044,WP_073436013.1-Halomonas-cupida:0.0930088384)92.4/100:0.1048592018)77.9/99:0.0497753970)87.4/98:0.0592221631)81.2/89:0.0270205083,(WP_127061154.1-Halomonas-nanhaiensis:0.2015131940,(WP_096281135.1-Halomonas-sp.-JB37:0.3068121158,(WP_027958907.1-unclassified-Halomonas:0.0387244863,WP_064236373.1-unclassified-Halomonas:0.0761141280)92.3/100:0.1059545338)84.1/100:0.0713477701)67.1/81:0.0353215314)86.7/89:0.0584820288,(((((WP_066801876.1-Moraxella-oblonga:0.1822430914,(WP_197128702.1-Psychrobacter-sp.-NZS113:0.2198494501,WP_201616712.1-Psychrobacter-immobilis:0.2017643787)98.8/100:0.2364053305)92.2/100:0.2247994650,WP_036548095.1-Nitrincola-lacisaponensis:0.5279395041)73.2/60:0.0776077686,((WP_041157945.1-Halomonas-sp.-KHS3:0.0810151449,(WP_120386672.1-Halomonas-alkaliphila:0.1666216479,(WP_141441391.1-Halomonas-titanicae:0.0890772547,WP_226251132.1-unclassified-Halomonas:0.0844368465)32.6/87:0.0135595903)80/87:0.0782903366)98.8/99:0.1433077668,WP_228219791.1-Halomonas-meridiana:0.1095818158)94.9/96:0.0979788320)88.5/57:0.0591785380,((((((WP_030071911.1-Halomonas-alkaliantarctica:0.0603078149,WP_095603285.1-Halomonas-humidisoli:0.0300379590)80.3/54:0.0233299742,WP_253485683.1-Halomonas-sp.-XH26:0.0363518336)54.8/54:0.0126938043,WP_038485954.1-Halomonas-campaniensis:0.0259627377)98.8/100:0.0998867093,WP_162218282.1-Halomonas-alkaliphila:0.0336503669)96.3/100:0.0575926018,WP_078087027.1-unclassified-Halomonas:0.0321096129)0/97:0.0006299379,WP_096655487.1-Halomonas-nigrificans:0.0557313301)98.8/100:0.1393459407)76.8/42:0.0336960924,(((((WP_158774340.1-Cobetia-sp.-L2A1:0.0910020925,WP_144727127.1-Cobetia-crustatorum:0.0364372504)15.2/96:0.0125016219,WP_088744253.1-Cobetia-sp.-QF-1:0.0745925498)89.9/99:0.0359659670,WP_158774127.1-Cobetia-sp.-L2A1:0.0264423126)82.1/100:0.0169862680,(WP_077373671.1-Cobetia:0.0563165619,((WP_213115235.1-Cobetia-sp.-MC34:0.0403097158,WP_200017180.1-Cobetia-sp.-cqz5-12:0.0778746495)94/100:0.0735092230,WP_225346840.1-Cobetia-amphilecti:0.0523032793)86.3/100:0.0433583847)92.3/100:0.0377783149)97.7/100:0.1374847015,(WP_017842145.1-Methylotuvimicrobium-buryatense:0.5297133408,(WP_251978361.1-unclassified-Salinicola:0.0872645964,WP_189443129.1-Salinicola-rhizosphaerae:0.1345776504)95.7/100:0.1360086259)87.7/62:0.0728711376)98/62:0.1323695849)73/96:0.0193692027)96.2/94:0.1539361366,WP_179929642.1-Halomonas-salicampi:0.4793633712)49.2/95:0.0493544766,WP_254581577.1-Marinobacterium-sedimentorum:0.4927219089)94.9/100:0.1204507756,(WP_015904383.1-Desulforapulum-autotrophicum:0.3571186160,WP_111960646.1-Desulfobacter-hydrogenophilus:0.4090137795)87.7/100:0.1649233617)89.9/98:0.1155004560,((((WP_151033206.1-Cellvibrio-sp.-KY-GH-1:0.0945477395,WP_049631514.1-Cellvibrio-sp.-pealriver:0.0988319025)84.5/100:0.1015704609,WP_062059887.1-Cellvibrio-sp.-OA-2007:0.0568193498)99.7/100:0.3447153869,WP_178968040.1-Neptunomonas-phycophila:0.4915541797)85.4/96:0.1834102113,WP_156872703.1-Ketobacter-sp.-MCCC-1A13808:0.5111865700)76.2/49:0.0705572432)76.2/47:0.0308243478,(((((((((((WP_018154793.1-Methanothermococcus-thermolithotrophicus:0.1858654080,WP_048201153.1-Methanocaldococcus-bathoardescens:0.0784657152)90.1/98:0.0753839655,((WP_052562942.1-Candidatus-Brocadia-sinica:0.4975143543,WP_173298646.1-Thermanaeromonas-sp.-C210:0.3837524998)87.8/95:0.1084024966,(((WP_011869065.1-Methanococcus-maripaludis:0.1169740735,WP_013998786.1-Methanococcus-maripaludis:0.0747421721)81.6/100:0.0476040519,WP_181493664.1-Methanococcus-maripaludis:0.0945022614)42/86:0.0304974732,WP_181504225.1-Methanococcus-maripaludis:0.0581719442)99.9/100:0.3293512082)14/86:0.0638557571)82.2/91:0.0721030605,((WP_119360783.1-Meiothermus-luteus:0.2348588222,WP_027877894.1-Meiothermus-cerbereus:0.0685923761)79.2/99:0.0834286099,WP_208013758.1-Meiothermus-sp.-CFH-77666:0.0830061970)100/100:0.9985199959)31.7/50:0.1166691871,((WP_010868297.1-Pyrococcus-abyssi:0.0290682318,((((WP_013798345.1-Methanotorris-igneus:0.0487473082,WP_013866394.1-Methanothermococcus-okinawensis:0.2442174287)62/96:0.0325292485,WP_007044577.1-Methanotorris-formicicus:0.0818366331)99.3/100:0.1109241844,((((((WP_148120035.1-Methanofervidicoccus-sp.-A16:0.2531375691,WP_013468354.1-Thermococcus-barophilus:0.0413595659)65.6/80:0.0256742486,WP_013906489.1-Pyrococcus-yayanosii:0.0308413811)81.1/87:0.0161475030,((WP_088855609.1-Thermococcus-siculi:0.0709783209,WP_087036111.1-Thermococcus-litoralis:0.0451872722)85.6/100:0.0338174562,(WP_167888701.1-Thermococcus-sp.-MV5:0.0480179671,WP_209475361.1-Thermococcus-stetteri:0.1239711529)26.2/14:0.0451852685)82.9/11:0.0273318959)0/10:0.0000021694,((((((((((WP_237702072.1-Thermococcus-sp.-AM4:0.0436057754,WP_223211916.1-Thermococcus-henrietii:0.0749635796)96.7/100:0.0702764986,WP_014733609.1-Pyrococcus-sp.-ST04:0.1579551620)87.7/99:0.0258147245,(WP_139679991.1-Thermococcus-indicus:0.0421765259,WP_148882943.1-Thermococcus-aciditolerans:0.0500034806)87.7/99:0.0268779642)25.4/65:0.0171038859,WP_058938040.1-Thermococcus-celericrescens:0.0378963366)73.1/68:0.0090125710,(WP_088865724.1-Thermococcus-barossii:0.0791804267,(WP_167711671.1-unclassified-Thermococcus:0.0321860441,WP_232460978.1-Thermococcus-radiotolerans:0.0651970153)75/78:0.0075609929)74.2/66:0.0094352550)82.2/80:0.0218158313,WP_258085050.1-Thermococcus-sp.-813A4:0.0377477293)84.4/98:0.0169766769,WP_088181368.1-Thermococcus-sp.-5-4:0.0758073550)91.7/99:0.0378056056,(WP_152879125.1-Thermococcus-sp.-101-C5:0.0591730179,WP_015850310.1-Thermococcus-sibiricus:0.0705434273)73.2/100:0.0090380620)90.8/99:0.0255098089,WP_253304977.1-Thermococcus-aggregans:0.0332399956)83.5/73:0.0181939928,WP_068664737.1-Thermococcus-piezophilus:0.0796906820)94.8/88:0.0617219944)36.6/63:0.0213198801,WP_048053121.1-Pyrococcus-horikoshii:0.0416011587)73.3/64:0.0146594716,(WP_167913028.1-Thermococcus-sp.-21S7:0.1006646166,WP_048150691.1-Palaeococcus-ferrophilus:0.0318878194)95.1/100:0.0588335962)81.7/77:0.0157846975)82.2/77:0.0240483767,WP_088862340.1-Thermococcus-celer:0.1235414015)72.6/89:0.0766770405)91.3/100:0.0809227982,WP_048103006.1-Aciduliprofundum-boonei:0.2996605431)100/100:0.3118564141)89/49:0.1110785429,((WP_245533116.1-Coprothermobacter-platensis:0.7714949960,WP_022856039.1-Thermodesulfobacterium:0.2730144387)35/56:0.1000849824,WP_150071182.1-Thermodesulfobacterium-sp.-TA1:0.3339666469)90.3/56:0.1419943118)96.1/94:0.1275064978,WP_028950938.1-Sulfurihydrogenibium-subterraneum:0.1538910496)0/11:0.0535876700,(WP_029522925.1-Persephonella-sp.-KM09-Lau-8:0.1475775217,(WP_200674641.1-Persephonella-atlantica:0.0908549764,WP_096999769.1-Persephonella-hydrogeniphila:0.1471978720)93.2/86:0.0793734328)80.5/86:0.0745629271)92.9/88:0.0933156616,WP_089322203.1-Desulfurobacterium-atlanticum:0.1911494917)86.6/88:0.0378107408,WP_132526207.1-Phorcysia-thermohydrogeniphila:0.1102370273)88.7/88:0.0631566091,WP_013638952.1-Desulfurobacterium-thermolithotrophum:0.0000026757)98.3/100:0.2428250801,WP_011996389.1-Coxiella-burnetii:0.7308772728)99/98:0.3663096916)89.7/95:0.0971103068,(((((WP_230291920.1-Croceicoccus-sp.-Ery5:0.5130394128,WP_164156863.1-Sandarakinorhabdus-rubra:0.3281561344)88.7/98:0.1194198612,((WP_116367876.1-Parahaliea-mediterranea:0.0892453451,WP_148063811.1-Parahaliea-aestuarii:0.1276159260)99.9/100:0.3843283689,WP_129645432.1-Steroidobacter-agariperforans:0.4840922239)87.8/94:0.1105045844)82.5/94:0.0847734202,(WP_070114691.1-Acidithiobacillus-caldus:0.3854816862,(((WP_009566950.1-Acidithiobacillus:0.0604990021,WP_215870853.1-unclassified-Acidithiobacillus:0.0886335624)95.9/100:0.0888595605,WP_163095688.1-Acidithiobacillus-ferrianus:0.0497222726)72.9/99:0.0386347395,WP_101536718.1-Acidithiobacillus-sp.-SH:0.1976747253)93.9/100:0.1344495255)97.3/100:0.2178919482)36.2/80:0.1227024703,WP_152145882.1-Alkalilimnicola-sp.-S0819:0.3612393245)95.5/97:0.1954530877,((((((((((WP_018882239.1-unclassified-Thioalkalivibrio:0.5307640553,WP_125198734.1-Guyparkeria-sp.-SCN-R1:0.2341104983)82.7/100:0.0683033138,WP_250072676.1-Guyparkeria-hydrothermalis:0.2615827300)17.8/92:0.0378360392,(((WP_026296529.1-Thioalkalivibrio-sp.-ALJ2:0.1635816038,WP_019569965.1-Thioalkalivibrio-sp.-ALE11:0.0952700681)87.5/100:0.0323601685,WP_028484075.1-Thioalkalivibrio-sp.-ALE17:0.0490113641)73.3/100:0.0104332200,WP_018169543.1-Thioalkalivibrio-sp.-ALMg9:0.0379213668)95.9/100:0.1198551249)99.9/100:0.3215684734,WP_092999033.1-Thiohalomonas-denitrificans:0.0797289485)36.1/97:0.0603242018,WP_019018778.1-Halomonas-lutea:0.3821008121)91.6/100:0.1736696753,(WP_125841544.1-Stutzerimonas-stutzeri:0.4765488374,((WP_262731117.1-Alcanivorax-marinus:0.2293686137,(WP_106232428.1-Halomonas-ventosae:0.0971797732,WP_228234339.1-Alcanivorax-marinus:0.2301666078)79.7/100:0.0507542836)83.1/100:0.0346258708,WP_175070852.1-Halomonas-taeanensis:0.1438906277)30.7/99:0.1044302607)85/100:0.0994535220)87.7/99:0.0656683457,((WP_146294265.1-Euhalothece-natronophila:0.3006613911,WP_159299499.1-Spiribacter-sp.-SSL99:0.2895778919)89.8/100:0.1032253652,WP_011630054.1-Alkalilimnicola-ehrlichii:0.2217099502)90.9/99:0.0820629997)85.6/98:0.0517236924,WP_127477393.1-Sulfurivermis-fontis:0.4262356643)80.9/95:0.0535265103,WP_028988655.1-Thermithiobacillus-tepidarius:0.3597363494)94.1/100:0.1420285459,WP_069857109.1-Desulfoplanes-formicivorans:0.3890666439)89.2/99:0.0905981478)83.5/86:0.0879727872)94.1/98:0.1201947989,((((((WP_132974196.1-Thiogranum-longum:0.2722298979,WP_201344172.1-Thiohalobacter-sp.-COW1:0.1803486207)96/100:0.2695770300,WP_198243301.1-methane-oxidizing-endosymbiont-of-Gigantopelta-aegis:0.4947346796)81.7/100:0.2001983432,WP_062153084.1-Beggiatoa-leptomitoformis:0.6806162821)80.1/75:0.1342769664,((((((((((((WP_036226274.1-Mesoaciditoga-lauensis:0.2489037825,((((WP_077198159.1-Thermosipho-affectus:0.0966288463,WP_012057290.1-Thermosipho-melanesiensis:0.2279545567)57.6/91:0.0453492681,(WP_127352844.1-Caldicellulosiruptor-changbaiensis:0.0770504064,WP_014043315.1-Caldicellulosiruptor-acetigenus:0.0329646618)100/100:0.4997184565)86.9/100:0.0738645720,((((((((WP_117708548.1-Fusobacterium-mortiferum:0.0006857915,WP_204716698.1-Fusobacterium-mortiferum:0.0909416676)74.2/70:0.0222212015,WP_115270465.1-Fusobacterium-necrogenes:0.0846543260)83.2/71:0.0425304502,WP_204716813.1-Fusobacterium-mortiferum:0.5384376304)73.3/97:0.0809049922,(((WP_130889691.1-Fusobacterium-ulcerans:0.0518592115,((WP_005950923.1-Fusobacterium-varium:0.0409053350,WP_070580525.1-Fusobacterium:0.0707093408)15.3/100:0.0143485275,WP_227372109.1-Fusobacterium-ulcerans:0.0434523671)84.7/100:0.0195819161)75.4/100:0.0132904187,(WP_241635364.1-Fusobacterium-gastrosuis:0.1442948669,(WP_074518138.1-Fusobacterium-necrophorum:0.0967199422,(WP_211495349.1-Fusobacterium-necrophorum:0.1648531224,WP_032890075.1-Fusobacterium-nucleatum:0.8366436430)88.7/98:0.0914315157)93.5/98:0.0891755203)98.9/98:0.0948071409)86.6/98:0.0355810196,WP_261793436.1-Fusobacterium-varium:0.3383236099)84.1/97:0.0511643299)96.6/100:0.1642305246,((WP_147004474.1-Leptotrichia-wadei:0.0332149274,WP_018499460.1-Leptotrichia-wadei:0.0395343626)90.6/100:0.0838545060,(WP_068155536.1-Leptotrichia-sp.-oral-taxon-847:0.0847714434,(WP_099992503.1-Fusobacterium-pseudoperiodonticum:0.1595151582,(WP_211617428.1-Leptotrichia-sp.-oral-taxon-218:0.0172895132,WP_094080762.1-Leptotrichia-sp.-oral-taxon-498:0.0560639884)96.2/100:0.0710979342)3.8/63:0.0436735845)91.8/100:0.0933288979)99.3/100:0.3159630081)88.9/96:0.0862251812,WP_013388736.1-Ilyobacter-polytropus:0.5952035697)67.6/79:0.0516267425,WP_012548442.1-Dictyoglomus-thermophilum:0.4337765088)78.3/78:0.0208944112,WP_173194267.1-Paenibacillus-frigoriresistens:0.4252856753)98.2/100:0.1108566952)87.7/88:0.0649632561,WP_047753768.1-Kosmotoga-pacifica:0.2029673319)83.8/88:0.0451400968)95.6/100:0.1254509353,(WP_099829233.1-Mesotoga-sp.-Brook.08.105.5.1:0.2041875471,WP_099830786.1-Mesotoga-sp.-Brook.08.105.5.1:0.2291325669)98.8/100:0.1798408956)96.1/99:0.1838776443,WP_038266426.1-Peptoclostridium-litorale:0.6403852423)56.5/83:0.0307031170,(((WP_009529486.1-Peptoanaerobacter-stomatis:0.2584906357,WP_068913445.1-Criibacterium-bergeronii:0.1982354725)99.6/100:0.3317954106,(WP_111570784.1-Halanaerobium-saccharolyticum:0.1001156193,(WP_089717478.1-Halanaerobium-congolense:0.0678233637,WP_089862761.1-Halanaerobium-salsuginis:0.0337052005)82.6/100:0.0466473611)99.3/100:0.2352971666)90.8/93:0.1045331644,((WP_148566871.1-Acetobacterium-paludosum:0.4280893877,WP_213236459.1-Fusibacter-paucivorans:0.7657197078)92.1/100:0.2086197154,(WP_138788932.1-Paenibacillus-thermoaerophilus:0.4387325703,WP_078711972.1-Anaerorhabdus-furcosa:0.8270606825)90.7/100:0.2377410362)85.8/98:0.0887887401)88.5/85:0.0577336092)77.7/82:0.0198640159,(((((((WP_068555806.1-Thermotalea-metallivorans:0.1611773917,WP_073236503.1-Desulforamulus-putei:0.1428544786)87.4/76:0.0433929757,WP_090442301.1-Natronincola-peptidivorans:0.2291254223)0/13:0.0202093268,WP_035164300.1-Caloranaerobacter-azorensis:0.1132485924)67.4/67:0.0402741257,WP_095133872.1-Anaeromicrobium-sediminis:0.2462005338)89.7/98:0.0618533825,(((((WP_266162911.1-Caproicibacterium-sp.-BJN0003:0.2102445757,WP_256135338.1-Bittarella-massiliensis:0.3972308658)97.2/98:0.2413042423,WP_220321633.1-Clostridium-tagluense:0.3873011397)44.5/86:0.0311466446,WP_160686059.1-Clostridium-sp.-C2-6-12:0.3491321614)88.7/85:0.0718426513,(WP_105204772.1-Neobittarella-massiliensis:0.2044465197,WP_022475313.1-Huintestinicola-butyrica:0.4278372181)97.9/91:0.2081075065)23.1/80:0.0308358725,((WP_069974893.1-Geosporobacter-ferrireducens:0.5592286765,(WP_223038819.1-Crassaminicella-profunda:0.1092964448,(WP_148809648.1-Crassaminicella-thermophila:0.0716524174,WP_218283043.1-Crassaminicella-sp.-143-21:0.3982002301)89.9/100:0.0550536712)87.8/100:0.0862131612)93.3/100:0.1382578793,WP_250858284.1-Oceanirhabdus-seepicola:0.2842413438)94.3/100:0.1357735295)93.3/85:0.0857295496)77.8/59:0.0227791366,(((((WP_252130283.1-Clostridium-sp.-SYSU_GA19001:0.4082684396,WP_202749105.1-Clostridium-sp.-YIM-B02515:0.1643121893)97.9/100:0.1787455037,(((WP_104410444.1-Clostridium-algidicarnis:0.0505088655,WP_185162668.1-Clostridium-algidicarnis:0.0906647692)100/100:0.2526536605,((WP_090038599.1-Clostridium-frigidicarnis:0.1934224813,WP_209459870.1-Youngiibacter-multivorans:0.2794411052)91.9/100:0.0984636089,(WP_164955553.1-Clostridium-estertheticum:0.2338959149,(WP_125002390.1-Clostridium-tagluense:0.2485703052,(WP_226136064.1-Clostridium-estertheticum:0.0336731710,WP_216146875.1-Clostridium-frigoris:0.0896252103)97.9/100:0.1362846510)83.2/100:0.0626927340)93.9/98:0.1100140538)30/96:0.0373101362)87.1/79:0.0591911024,WP_216462460.1-Clostridium-bornimense:0.2031126709)33.2/68:0.0207526175)84.6/59:0.0382053915,(WP_234123145.1-Clostridium-hydrogenum:0.2808205649,((WP_084114600.1-Clostridium-acidisoli:0.1274060249,(WP_015614219.1-Clostridium-pasteurianum:0.0347619186,(WP_010235680.1-Clostridium-arbusti:0.0159743337,WP_003443690.1-Clostridium-pasteurianum:0.0642589111)86.7/100:0.0287773630)94.8/100:0.0467001586)10.1/93:0.0110143975,WP_026884435.1-Clostridium-akagii:0.1272155607)90.8/100:0.1165589852)63.4/78:0.0246591073)93.4/84:0.0700613624,(((WP_165412483.1-Clostridium-estertheticum:0.0852767681,WP_216302578.1-unclassified-Clostridium:0.0132666480)100/100:0.2839792595,((WP_204598070.1-Clostridium-pascui:0.1918243929,WP_027632043.1-Clostridium-hydrogeniformans:0.1909879996)68.2/92:0.0143316638,WP_216457288.1-Clostridium-simiarum:0.1984748220)25.2/93:0.0201562381)83.4/98:0.0447084985,((WP_066398085.1-Cytobacillus-eiseniae:0.4213152303,((WP_076307499.1-Paenibacillus-odorifer:0.0782583135,((WP_042140168.1-Paenibacillus-sp.-FSL-P4-0081:0.0130020912,WP_036721118.1-Paenibacillus-sp.-FSL-R7-277:0.0940469563)91.9/100:0.0483566277,WP_074087222.1-Paenibacillus-sp.-P32E:0.0444402856)95.1/100:0.1148598564)97.7/100:0.1685772818,(((WP_056034132.1-Paenibacillus-sp.-Leaf72:0.1287709672,WP_197936429.1-Paenibacillus-sp.-GSMTC-2017:0.1958422005)89.9/100:0.1000247961,WP_090798944.1-Paenibacillus-sp.-GP183:0.3147039208)83/96:0.0543849165,WP_262684732.1-Paenibacillus-sp.-WQ-127069:0.2327985082)75.8/80:0.0609671890)31.2/69:0.0223058613)74.1/96:0.0684580453,WP_036935152.1-Pseudobacteroides-cellulosolvens:0.2712156268)95.1/98:0.1287851788)51.6/82:0.0372352021)88.4/96:0.0466810764,((((WP_091684654.1-Anaerocolumna-aminovalerica:0.2907434248,(WP_033168390.1-Clostridium-sp.-KNHs205:0.1208231139,((WP_073588199.1-Anaerocolumna-xylanovorans:0.0390495342,WP_185256524.1-Anaerocolumna-chitinilytica:0.1832522590)75.8/100:0.0622077878,WP_073279502.1-Anaerocolumna-jejuensis:0.1446944563)57.6/100:0.0612718846)99.1/100:0.2177266440)94.1/100:0.1290289783,((((WP_073273890.1-Anaerocolumna-jejuensis:0.1869522641,WP_033167869.1-Clostridium-sp.-KNHs205:0.1962898564)57.7/99:0.0595718034,((WP_010268185.1-Paenibacillus-senegalensis:0.2381354584,WP_155699987.1-Paenibacillus-psychroresistens:0.0781591946)43.1/96:0.0294426698,(WP_202751253.1-Clostridium-sp.-YIM-B02515:0.0746868349,WP_090040610.1-Clostridium-frigidicarnis:0.0385351755)21.8/82:0.0101971988)93.3/94:0.0929974017)99.5/100:0.2296729941,((WP_249305773.1-Jutongia-hominis:0.2877004259,(WP_055944671.1-unclassified-Clostridium:0.0456642539,WP_216564272.1-Eubacterium-sp.-MSJ-13:0.0374084857)81.9/100:0.0460316903)59/92:0.0309347766,WP_022463459.1-Jutongia-huaianensis:0.0886227233)99.8/100:0.3002085144)77.5/100:0.0539536771,WP_058352784.1-Acetivibrio-ethanolgignens:0.4117472104)95.2/99:0.1690745621)79.9/90:0.0395320583,(((WP_253721803.1-Treponema-sp.-OMZ-787:0.7419215854,WP_005539496.1-Johnsonella-ignava:0.1717234479)90.1/95:0.1105894476,(WP_060930762.1-Lachnoanaerobaculum-saburreum:0.2685714598,((((((((((((WP_024837362.1-Lacrimispora:0.1312847206,WP_077612318.1-Clostridium-sp.-Marseille-P2415:0.0447970742)72.8/98:0.0091969169,(WP_024293365.1-Lachnospiraceae:0.0488802439,WP_092242699.1-Lacrimispora:0.0344061659)95.1/100:0.0472156314)87.8/98:0.0230935333,WP_138313639.1-Hungatella-hathewayi:0.0476413866)23.4/58:0.0181177987,((WP_124065139.1-Clostridium-sp.-E02:0.1284122587,WP_117415169.1-Clostridium-indicum:0.0583582920)75.9/93:0.0124573061,WP_143321893.1-Clostridium-sp.-HBUAS56010:0.0413852827)43.5/86:0.0043345352)98.3/100:0.0659613432,((WP_231062061.1-Lacrimispora-sp.-NSJ-141:0.1224782108,WP_249304640.1-Qiania-dongpingensis:0.1008109903)94.3/99:0.0826419325,(WP_227052204.1-Anaerostipes-caccae:0.0362312555,WP_137330070.1-Anaerostipes-rhamnosivorans:0.1205134200)99.7/100:0.1468440513)71.7/69:0.0159880212)88.1/49:0.0215232133,((WP_186424215.1-Lacrimispora-celerecrescens:0.2176071708,WP_118541387.1-unclassified-Clostridium:0.0541375249)84.8/46:0.0406753176,(((WP_215718428.1-Anaerostipes-hadrus:0.0166430422,WP_075760385.1-Anaerostipes-sp.-494a:0.0742653293)99.9/100:0.1187965872,WP_215700061.1-Clostridium-sp.-MCC353:0.0872724700)0/37:0.0000029425,((((WP_045093538.1-Clostridium-sp.-FS41:0.0610720421,WP_025485467.1-Lachnospiraceae:0.0813790509)83.3/95:0.0268154888,((WP_154471253.1-Clostridium-porci:0.1779945532,WP_002577448.1-Enterocloster:0.0792401067)0/14:0.0180738826,WP_178313416.1-Lacrimispora-sp.-210928-DFI.3.58:0.0250755630)0/75:0.0000026762)93.8/85:0.0413616475,WP_066545911.1-Clostridium-sp.-AT4:0.0443776949)73.8/80:0.0068222181,WP_024736308.1-Enterocloster-asparagiformis:0.0523307275)92.7/85:0.0257726219)74.3/37:0.0083715893)78.9/47:0.0139000012)21.6/64:0.0247039310,WP_006781439.1-Hungatella-hathewayi:0.1313273963)87.7/95:0.0219052380,WP_118243934.1-Clostridium-sp.-AM58-1XD:0.0577090203)72.1/80:0.0089184070,((WP_087223935.1-Lachnoclostridium-sp.-An14:0.0575000194,WP_101876396.1-Lachnoclostridium-edouardi:0.0694190972)90.7/100:0.0362206205,(WP_008399284.1-unclassified-Clostridium:0.0414690713,WP_118742831.1-Clostridium-sp.-AM28-20LB:0.0410707107)71.8/88:0.0085742244)42.7/93:0.0165125042)96.6/96:0.0549719985,(WP_003497689.1-Clostridium:0.0597736221,WP_125142063.1-Clostridium-transplantifaecale:0.0580332155)57.7/100:0.0109406658)90.4/96:0.0471969562,WP_251497048.1-Otoolea-sp.-DSM-111138:0.1833458473)21.9/78:0.0397894145,WP_118555425.1-Lachnotalea-sp.-AF33-28:0.3055761060)82.9/94:0.0719328261)94.9/100:0.1142975833)79/92:0.0767564367,((((((((((((WP_035764434.1-Butyrivibrio-sp.-NC2002:0.1186806839,(((WP_026526812.1-Butyrivibrio-sp.-VCD2006:0.2072317664,WP_026491547.1-unclassified-Butyrivibrio:0.0198275439)63.2/100:0.0476203991,WP_044913869.1-Butyrivibrio-sp.-WCE2006:0.0315746544)91/100:0.0899164453,WP_091822189.1-Butyrivibrio-sp.-ob235:0.0895488664)79.7/100:0.0513671849)97/100:0.1499878777,((((((WP_022768176.1-unclassified-Butyrivibrio:0.0299951623,WP_022766545.1-Butyrivibrio-sp.-XPD2006:0.0717511034)99.4/100:0.1270024202,WP_034446055.1-Butyrivibrio-sp.-AE2032:0.0823648130)93/96:0.0655496373,WP_024867291.1-Butyrivibrio-sp.-FCS014:0.0390979809)86.9/67:0.0371930560,(WP_071175335.1-Butyrivibrio:0.0946207554,WP_026513668.1-Butyrivibrio-sp.-LB2008:0.0597073571)87.6/67:0.0530699638)70.8/98:0.0322705527,WP_026663187.1-Butyrivibrio-proteoclasticus:0.0906840377)71.9/84:0.0218033156,(WP_026652320.1-Butyrivibrio-proteoclasticus:0.0576085775,(WP_013279951.1-Butyrivibrio-proteoclasticus:0.0391236693,WP_092060841.1-Butyrivibrio-sp.-INlla14:0.0661991143)82/100:0.0218501986)0/95:0.0000023695)96.8/98:0.1095264265)96.2/100:0.1263139192,((WP_019160740.1-Blautia:0.0805890339,WP_173738571.1-Blautia-schinkii:0.0295619426)86.1/100:0.0357265241,(WP_158421413.1-Blautia:0.0498718492,(WP_148445023.1-Eubacteriales:0.0368384896,WP_187566223.1-Blautia-faecis:0.0666311105)93.5/100:0.0384232388)74.5/98:0.0222013067)96.8/100:0.1002457252)95/98:0.0730287039,WP_178047055.1-Fusicatenibacter-sp.-CLA-AA-H277:0.1355852401)69.9/89:0.0165814924,WP_109624131.1-Murimonas-intestini:0.1076560632)89.6/91:0.0404759637,(WP_230400100.1-Novisyntrophococcus-fermenticellae:0.2805976053,((WP_173751851.1-Blautia-caecimuris:0.0618328884,WP_117997894.1-Blautia-obeum:0.0165339782)99.6/100:0.1064843490,(((WP_158367551.1-Lachnospiraceae:0.0461711138,WP_235247881.1-Faecalicatena-contorta:0.1036044071)74.2/99:0.0530696137,(WP_251390531.1-Mediterraneibacter-sp.-DSM-111629:0.0568384808,WP_073962323.1-Ruminococcus:0.0362229051)86.3/100:0.0340822214)83.2/79:0.0229841999,WP_070089756.1-Merdimonas-faecis:0.0416607545)81.6/84:0.0184502313)75.7/66:0.0103352903)84.7/70:0.0290939736)81.4/93:0.0225844651,WP_132379639.1-Muricomes-intestini:0.1341560443)96.2/94:0.0604204281,WP_087156947.1-Lachnoclostridium-sp.-An181:0.0732838610)67.3/55:0.0222628087,(WP_073994331.1-Parasporobacterium-paucivorans:0.1894616017,(WP_099466994.1-Konateibacter-massiliensis:0.0959237527,WP_151145296.1-Candidatus-Galacturonibacter-soehngenii:0.0710394522)96.6/100:0.1166570682)73.8/92:0.0830237289)38.6/81:0.0272497070,WP_115483253.1-Anaerosacchariphilus-polymeriproducens:0.1427904411)36.3/92:0.0538891696,WP_158351687.1-Diplocloster-modestus:0.3217677888)67.5/99:0.0757182408,WP_158346166.1-Lachnospiraceae:0.3211406789)97.1/95:0.1670184214)86.5/92:0.0457234372)88/95:0.0723772562,WP_226914794.1-Intestinibacter-bartlettii:0.3901019197)97.1/100:0.1838650127)78.8/70:0.0219818759)90.8/96:0.0456261913,WP_119970050.1-Clostridium-fermenticellae:0.3759819081)76.5/86:0.0158249104)84.5/83:0.0300467487,(((((((WP_088538685.1-Methanobrevibacter-sp.-87.7:0.0686906057,WP_016359256.1-Methanobrevibacter-sp.-AbM4:0.1605295264)97.7/100:0.1328895174,WP_067148181.1-Methanobrevibacter-olleyae:0.1625292493)80.8/100:0.0401675561,WP_116670296.1-Methanobrevibacter-woesei:0.3243678673)99.8/95:0.1882295737,(WP_248612240.1-Methanobacterium-alcaliphilum:0.1821826061,((WP_071907175.1-Methanobacterium-congolense:0.0289850046,WP_209584978.1-Methanobacterium-aggregans:0.0567416559)99.6/100:0.1483125672,(WP_069583666.1-Methanobacterium:0.2357059931,WP_048190917.1-Methanobacterium-sp.-SMA-27:0.1720796181)89.6/95:0.0731742505)37.3/92:0.0272438754)80.5/82:0.0283651585)61.1/29:0.0182276605,(WP_011406138.1-Methanosphaera:0.2209672368,(WP_112123902.1-Methanosphaera-sp.-BMS:0.1409114901,(WP_069592847.1-Methanosphaera-sp.-WGK6:0.0382143594,WP_095608065.1-Methanosphaera-cuniculi:0.1394033687)96.8/100:0.1033296236)25.8/100:0.0474968859)97.6/100:0.1203223416)69/30:0.0154718095,(WP_013644504.1-Methanobacterium-lacus:0.2867952353,WP_054834988.1-Methanobrevibacter-arboriphilus:0.2322842960)89.2/87:0.0627215481)87.9/93:0.0549257820,(WP_112093115.1-Methanothermobacter-tenebrarum:0.3718290855,WP_013413362.1-Methanothermus-fervidus:0.2917390169)98.9/100:0.2237384080)89.5/93:0.0804612796)85.9/88:0.0498442782,(((((((((((((((((((((((((WP_054742706.1-Cellulosilyticum-ruminicola:0.0977148421,WP_069999280.1-Cellulosilyticum-sp.-I15G10I2:0.3622292687)69/83:0.0259727809,(WP_177671917.1-Zhenhengia-yiwuensis:0.0589729928,WP_053982981.1-Niameybacter-massiliensis:0.1401182123)97.9/100:0.1288648425)99.6/99:0.1900221646,WP_152890015.1-Clostridium-tarantellae:0.1510746615)92.5/82:0.0640976731,(WP_195959433.1-Clostridium-tertium:0.0821048758,WP_195429094.1-Clostridium-sp.-D46t1_190503_E9:0.0313651054)57.7/80:0.0039871418)81.7/80:0.0181881540,WP_047001949.1-Clostridium-sp.-C8:0.0742727073)76.8/95:0.0311021647,WP_066890539.1-Clostridium-nigeriense:0.0595854273)74.9/93:0.0085705745,WP_016207773.1-Clostridium-sartagoforme:0.0379492209)92.3/99:0.0310304516,WP_095406848.1-Romboutsia-maritimum:0.1578861298)77.2/47:0.0087549587,(WP_197096717.1-Paeniclostridium-sordellii:0.1187360537,WP_238903698.1-Clostridium-sp.-YIM-B02506:0.0740356879)62.2/35:0.0332194526)89.6/84:0.0257892526,((((WP_029452249.1-Clostridium-algidicarnis:0.1313665064,WP_216183125.1-Clostridium-gasigenes:0.0275971663)37.2/99:0.0349382689,WP_216313757.1-Clostridium-gasigenes:0.0976656997)96.9/98:0.0816198765,WP_066677671.1-Clostridium-septicum:0.0550925771)71.1/82:0.0238234520,((((WP_195972826.1-Clostridium-thermobutyricum:0.1080453743,(WP_209795868.1-Clostridium-moniliforme:0.0824510349,WP_194191032.1-Clostridium-chrysemydis:0.1417499445)97.1/100:0.0688128363)72.9/100:0.0114889812,(WP_073337580.1-Clostridium-grantii:0.1747243494,(WP_072892386.1-Clostridium-fallax:0.1723870549,WP_072832886.1-Clostridium-collagenovorans:0.0356668029)77.9/98:0.0299802959)88.8/92:0.0381171864)68.9/87:0.0078877252,WP_040194942.1-Clostridium-culturomicium:0.0854160528)90.9/91:0.0346025226,WP_055184115.1-Clostridium-paraputrificum:0.0837779599)86.7/92:0.0291605666)74.4/77:0.0144548480)89.4/82:0.0277100438,((((WP_094369216.1-Romboutsia-weinsteinii:0.2120254713,WP_021359979.1-Clostridioides-difficile:0.4347836790)0.8/82:0.0506517263,(WP_074918830.1-Terrisporobacter-glycolicus:0.0917344397,WP_195929149.1-Clostridium-sp.-1001270J_160509_D11:0.0434157991)82.6/100:0.0312757683)91.6/91:0.0449176878,(WP_258242885.1-Romboutsia-lituseburensis:0.1629799117,(WP_057560648.1-Paeniclostridium-sordellii:0.1107294031,((WP_057580750.1-Paeniclostridium-sordellii:0.1459197402,WP_057538715.1-Paeniclostridium-sordellii:0.0501511580)82.2/100:0.0197771385,WP_195252007.1-Romboutsia-sp.-1001713B170207_170306_H8:0.0701786171)82.5/100:0.0182665533)47.3/97:0.0171872876)87/90:0.0304516906)71.9/77:0.0125742814,((((WP_055265352.1-Clostridium:0.0445360311,(WP_186859336.1-Clostridium-sp.-NSJ-6:0.0746531369,(WP_055265304.1-Clostridium-disporicum:0.0756895001,(WP_154821366.1-Clostridium-saudiense:0.0103035862,WP_195245504.1-Clostridium-celatum:0.0845244643)79.1/100:0.0333700129)69.1/100:0.0405918308)79.4/100:0.0646425027)86.6/100:0.0891313987,WP_186834444.1-Eubacteriales:0.1555653089)96.6/100:0.0935054426,((WP_235068992.1-Turicibacter-sp.-TJ11:0.0765982140,((WP_161831114.1-Turicibacter-sp.-TS3:0.0693632450,WP_155223812.1-Turicibacter-sanguinis:0.0323675734)80.4/50:0.0207571459,WP_195933776.1-Turicibacter-sanguinis:0.0322430382)60.5/50:0.0083565885)84.5/50:0.0228743305,WP_212726030.1-Turicibacter-bilis:0.0246033521)99.2/100:0.1083823918)47.6/97:0.0216150262,(WP_066874757.1-Clostridium-mediterraneense:0.2503356392,WP_198610231.1-Clostridium-perfringens:0.0231328011)59.3/85:0.0615078317)97/98:0.0949078946)35.2/75:0.0180608694)86/88:0.0194529396,WP_163195303.1-Clostridium-thermarum:0.1600874659)90.4/93:0.0322176787,(((WP_092726456.1-Romboutsia-lituseburensis:0.2840549122,WP_055258173.1-Sarcina-ventriculi:0.1168375159)81.1/95:0.0283958945,((WP_055666468.1-Desnuesiella-massiliensis:0.1667339499,((((WP_099190526.1-Tepidibacter-mesophilus:0.2265105819,((((WP_227847393.1-Clostridioides-sp.-ZZV15-6598:0.1465013819,WP_227846117.1-Clostridioides-sp.-ES-S-0145-01:0.0411609672)95.9/100:0.0555225559,(WP_227842647.1-unclassified-Clostridioides:0.0409428180,WP_003420982.1-Clostridioides-difficile:0.1197233429)85.9/100:0.0281888380)82.6/100:0.0342844958,WP_038810653.1-Clostridioides-difficile:0.0438555003)100/100:0.3848263250,WP_226126924.1-Clostridium-algoriphilum:0.4151327446)74.9/98:0.0543075880)46.3/95:0.0657443617,((WP_142413383.1-Hathewaya-massiliensis:0.2592101909,WP_035130905.1-Clostridium-sulfidigenes:0.2835626508)46.5/99:0.0385387404,WP_072903743.1-Hathewaya-proteolytica:0.5022122203)96.7/98:0.1439408511)85.1/97:0.0378046334,WP_095404508.1-Romboutsia-maritimum:0.4130116633)0/86:0.0067200071,WP_072725426.1-Tepidibacter-thalassicus:0.2243274152)91/95:0.0493027045)73.7/98:0.0463125667,WP_072990516.1-Clostridium-cavendishii:0.3060282594)71/49:0.0150264212)87.7/51:0.0339134556,((WP_216288229.1-Clostridium-psychrophilum:0.1363537874,(WP_220321605.1-Clostridium-tagluense:0.1108227116,(WP_216137569.1-Clostridium-estertheticum:0.0539152640,WP_220704225.1-Clostridium-estertheticum:0.0502538124)65.3/90:0.0169388980)7.6/45:0.0453735661)99.1/100:0.1359141312,WP_133628539.1-Fonticella-tunisiensis:0.0503198011)92.5/71:0.0437507253)84.9/90:0.0230353445)90.7/88:0.0393200557,(WP_066825124.1-Clostridium-tepidiprofundi:0.1494892229,WP_072889493.1-Tepidibacter-formicigenes:0.1007332025)77.5/99:0.0151883337)89.4/98:0.0629012948,WP_055069574.1-Clostridium-massiliamazoniense:0.1669191303)87/87:0.0374529003,(WP_106059304.1-Clostridium-vincentii:0.1290677986,WP_143316579.1-Clostridium:0.0320184111)94.1/100:0.0551569035)21.1/80:0.0210235994,WP_261829490.1-Clostridium-sp.-TW13:0.2541430785)88.8/97:0.0363441348,WP_090010800.1-Clostridium-sp.-DSM-8431:0.1169120663)99.9/99:0.1471216998,(((WP_261829154.1-Clostridium-sp.-TW13:0.1129550694,(WP_022750218.1-Clostridium-saccharobutylicum:0.0479818396,WP_226126241.1-Clostridium-algoriphilum:0.1778362543)81.6/87:0.0276161272)82.7/58:0.0231575233,((WP_077855754.1-Clostridium-beijerinckii:0.0473761439,WP_041894754.1-Clostridium-beijerinckii:0.0363739315)99.2/100:0.1042961853,(WP_015391310.1-Clostridium-saccharoperbutylacetonicum:0.0666861854,(WP_077846315.1-Clostridium-puniceum:0.0650983273,WP_238915641.1-Clostridium-sp.-YIM-B02555:0.0779147338)85.3/100:0.0281233137)90/99:0.0312195672)78.1/82:0.0172936968)74.1/32:0.0081412418,(WP_185652834.1-Clostridium-sp.-DJ247:0.0261334289,WP_185652641.1-Clostridium-sp.-DJ247:0.0894748604)94.7/100:0.0441157628)86.7/52:0.0231947006)73.8/43:0.0105112663,(WP_077848125.1-Clostridium-puniceum:0.1708180722,WP_224033465.1-Clostridium-gelidum:0.1104065773)59.8/47:0.0145658016)76/64:0.0199964794,(WP_169105633.1-Clostridium-sp.-SM-530-WT-3G:0.0758542249,(WP_027636628.1-Clostridium-butyricum:0.1066183904,WP_077866663.1-Clostridium-saccharobutylicum:0.2806218154)58.7/95:0.0217004933)94.7/94:0.0489479266)75.2/77:0.0110104629,((WP_265859166.1-Clostridium-botulinum:0.1513870612,(WP_009169009.1-Clostridium-sp.-DL-VIII:0.0666233181,WP_077856313.1-Clostridium-beijerinckii:0.0198278516)93/67:0.0680081258)67.8/63:0.0417766547,WP_154531076.1-Inconstantimicrobium-porci:0.1577243853)81.5/85:0.0299077399)92.2/75:0.0470724962,((((WP_009169246.1-unclassified-Clostridium:0.1411855037,WP_077843022.1-Clostridium-beijerinckii:0.1876818787)83.4/100:0.0429284458,WP_041896941.1-Clostridium-beijerinckii:0.2345561908)89.1/100:0.0707450961,WP_069680896.1-Clostridium-taeniosporum:0.2815231779)62.3/99:0.0634890956,WP_160359894.1-Clostridium-chromiireducens:0.3510798316)62.6/85:0.0170105273)81/82:0.0560885747,WP_079421587.1-Clostridium-oryzae:0.5710483090)79.5/82:0.0822742512,WP_216390068.1-Clostridium-sp.-MSJ-8:0.2587171370)94.4/85:0.1182728845)93.9/86:0.1435115481,(((WP_072827923.1-Fibrobacter-sp.-UWB12:0.0388830894,WP_088630837.1-unclassified-Fibrobacter:0.0516599723)40/75:0.0248828784,WP_073304542.1-Fibrobacter:0.2540811994)94.5/100:0.1024903369,(WP_097034884.1-Fibrobacter-sp.-UWT3:0.0302342720,(WP_073056819.1-unclassified-Fibrobacter:0.0301603282,(WP_073155408.1-Fibrobacter:0.0573237284,WP_073188830.1-Fibrobacter-sp.-UWP2:0.0336713399)52.5/81:0.0410245075)45.4/70:0.0212963201)94.3/95:0.1269075278)99.4/100:0.2628422385)1.6/33:0.0261659200,(((WP_058965344.1-Fournierella-massiliensis:0.2568664787,WP_185752312.1-Butyrivibrio-sp.-AE2032:0.4833240674)79.4/84:0.0870043931,WP_155202396.1-Ruthenibacterium-lactatiformans:0.3026302705)93.2/99:0.1385042622,(WP_186886939.1-Anaerofilum-hominis:0.2203889322,WP_102049807.1-Pygmaiobacter-massiliensis:0.3234474590)87.9/87:0.1118023596)98.8/100:0.2216314132)96.3/99:0.2102344196,WP_236098952.1-Dethiosulfovibrio:0.8448444910)92.6/98:0.1730713384,(((((((((((((((WP_008983868.1-Alishewanella-agri:1.6010622680,WP_265845290.1-Plesiomonas-shigelloides:0.6816993782)97.7/100:0.5485644335,(WP_247177500.1-Escherichia-coli:0.5537092731,YP_009985672.1-Escherichia-phage-muut:0.3208326077)95.9/100:0.2869303913)88.1/99:0.0991354098,WP_096000273.1-Klebsiella-pneumoniae:0.2102355723)82.4/85:0.0527501407,(((WP_267333169.1-unclassified-Gilliamella:0.2154793249,WP_267361838.1-Gilliamella-sp.-B2772:0.1830458470)71.7/100:0.0140228775,WP_086361229.1-Gilliamella:0.1644051484)48.8/100:0.0346706203,(((((WP_086319877.1-Gilliamella-apis:0.0254087899,WP_198214961.1-Gilliamella-sp.-M0320:0.0711772201)91.7/100:0.0401386864,WP_039129858.1-Gilliamella:0.0314149036)76.9/100:0.0150034989,(WP_065737496.1-Gilliamella-apicola:0.0464243368,WP_267331332.1-unclassified-Gilliamella:0.0767938874)95.7/100:0.0680494729)89.3/99:0.0474332676,((WP_065579324.1-Gilliamella-apis:0.1023035791,WP_160432071.1-Gilliamella-sp.-Pas-s25:0.1486736268)85.4/93:0.0280416077,(WP_065587409.1-Gilliamella-apicola:0.0778195099,WP_065580031.1-Gilliamella-apicola:0.0823618619)92/100:0.0394315548)74.1/92:0.0172170441)88.9/99:0.0480681701,((WP_065574597.1-Gilliamella-apicola:0.0694622698,WP_065569248.1-Gilliamella-apicola:0.0481008911)39.2/100:0.0087148303,WP_065610260.1-Gilliamella-apicola:0.0532420183)97.7/100:0.0864866020)68.7/96:0.0164640715)99.3/100:0.2085317575)66.9/72:0.0446742250,((((((((((WP_171588903.1-Pelistega-europaea:0.4771401158,(WP_087653467.1-Klebsiella-Raoultella-group:0.1752260317,WP_028357569.1-Brackiella-oedipodis:0.3234605238)68.1/98:0.0518155764)80/97:0.0462976806,((WP_029991521.1-Tatumella-ptyseos:0.1111249947,WP_212707699.1-Tatumella-sp.-JGM130:0.0521972929)51.9/91:0.0446597283,WP_087487466.1-Tatumella-citrea:0.0510423906)96.7/100:0.0950352386)86/87:0.0385747394,((WP_024553491.1-Franconibacter-helveticus:0.3746779061,WP_154751639.1-unclassified-Erwinia:0.2779577188)59.1/94:0.0699450281,WP_140470594.1-Ewingella-americana:0.3270797754)86.6/94:0.0605469992)73.9/86:0.0126639394,((WP_040666911.1-Neisseria-wadsworthii:0.1306293059,WP_135034416.1-Neisseria:0.2089754389)95/98:0.0884264998,(((WP_214236207.1-Rosenbergiella-sp.-S61:0.0908021427,WP_241647140.1-Rosenbergiella-sp.-JB07:0.1630807690)45.9/100:0.0514764320,WP_214195533.1-Rosenbergiella-collisarenosi:0.1119239872)95.6/100:0.1143410386,WP_101506252.1-Erwinia-sp.-B116:0.1849875853)78/100:0.0592237018)81.3/95:0.0291077729)89.3/90:0.0417710195,(WP_142430937.1-Klebsiella-oxytoca:0.1788949577,((WP_239614184.1-Raoultella-ornithinolytica:0.1156629345,WP_186921353.1-Klebsiella-quasipneumoniae:0.2136085959)74.1/76:0.0172948966,((WP_032699180.1-Raoultella:0.0707952797,WP_228399925.1-Raoultella:0.0400220375)72/98:0.0175479414,WP_004138711.1-Klebsiella:0.0245736153)95.2/100:0.0570984060)69.3/76:0.0267998383)77.6/96:0.0323083168)67.4/75:0.0165787422,((((WP_228398973.1-Insectihabitans-xujianqingii:0.1084437359,WP_074823218.1-Pragia-fontium:0.1247232121)96.2/100:0.0836490146,WP_130971788.1-Hafnia-alvei:0.1593541333)89.8/100:0.0487520333,((WP_112167140.1-Klebsiella:0.1594807290,WP_200519240.1-Klebsiella-michiganensis:0.1317414546)92.5/99:0.0670461741,WP_137713144.1-Brenneria-rubrifaciens:0.1539988612)72.9/99:0.0197205172)82.3/34:0.0170234355,((WP_131063651.1-Raoultella-planticola:0.0320912465,WP_163980548.1-Raoultella-ornithinolytica:0.0457964909)99/100:0.0908241688,((WP_148250181.1-Raoultella-terrigena:0.1188381248,WP_032686784.1-Raoultella-planticola:0.0436514634)30.8/100:0.0402902259,(WP_032441370.1-Klebsiella-pneumoniae:0.1545855234,(WP_015583911.1-Raoultella-ornithinolytica:0.0941914840,WP_265776798.1-Klebsiella-oxytoca:0.1860085510)60/100:0.0268015565)98.2/99:0.1516408805)85.7/83:0.0488919897)72/71:0.0144044732)74.2/29:0.0121919880)51.1/33:0.0080933345,WP_165428828.1-Rouxiella-badensis:0.1632740654)76.9/79:0.0128758241,(WP_142514871.1-Klebsiella-huaxiensis:0.1861316866,(WP_024360036.1-Klebsiella:0.0661665309,WP_234284225.1-Klebsiella-oxytoca:0.1133932082)78.7/90:0.0541256944)93.8/99:0.0735947558)71.8/91:0.0089796732,(((((WP_136167279.1-Brenneria-sp.-CFCC-11842:0.0559487327,WP_264089117.1-unclassified-Brenneria:0.1304887408)64/99:0.0487672620,WP_109053612.1-Brenneria-roseae:0.1273563336)80.7/99:0.0327047035,((((((((((WP_180742451.1-Pectobacterium-sp.-CFBP8739:0.2763309303,WP_039517132.1-Pectobacterium-brasiliense:0.0196020365)81/95:0.0220207642,WP_039280054.1-Pectobacterium-brasiliense:0.0416384898)89.1/94:0.0265151077,WP_039346077.1-Pectobacterium-fontis:0.0382792138)72.1/87:0.0078956502,(((WP_047710158.1-Pectobacterium-atrosepticum:0.0705450833,(WP_198482318.1-Pectobacterium-parmentieri:0.0203912810,WP_107169883.1-Pectobacterium:0.0698339118)87.2/70:0.0274627062)44.9/71:0.0153153092,WP_033071760.1-Pectobacterium-parmentieri:0.0356562131)93.2/85:0.0516092321,WP_226305805.1-Pectobacterium-versatile:0.0290531112)81.2/74:0.0285370122)88/70:0.0255525462,(((((WP_180777316.1-Pectobacterium-aroidearum:0.0972964523,(((WP_233968669.1-Pectobacterium-polaris:0.0703815821,WP_048259384.1-Pectobacterium-peruviense:0.0172679312)92.5/100:0.0384189129,WP_138255615.1-Pectobacterium:0.0338320780)72.8/76:0.0093062413,WP_039504461.1-Pectobacterium:0.0386780919)41.8/76:0.0132240965)87.4/98:0.0285004399,WP_151242707.1-Pectobacterium-carotovorum:0.0453529516)0/62:0.0000027447,(WP_205946867.1-Pectobacterium:0.0763871405,WP_116186106.1-Pectobacterium-aquaticum:0.0364677817)71.2/82:0.0259242668)76.2/79:0.0166751005,WP_205558955.1-Pectobacterium-brasiliense:0.0719625025)73.3/77:0.0092239645,WP_103972204.1-Pectobacterium:0.0195214042)73.9/72:0.0095748931)94.7/93:0.0412233628,WP_233976643.1-Pectobacterium-versatile:0.0270152809)68.5/92:0.0089830250,WP_109411993.1-Pectobacterium-polaris:0.0704917270)75.3/93:0.0099413434,((WP_249680869.1-Pectobacterium-polaris:0.0482193122,WP_125232508.1-Pectobacterium-versatile:0.0654092131)81.3/97:0.0175084927,WP_005974084.1-Pectobacterium-wasabiae:0.0418086927)83.1/99:0.0198940303)71.5/99:0.0135344790,WP_119888241.1-Pectobacterium:0.0697515737)81.1/100:0.0405982362,(WP_132452860.1-Samsonia-erythrinae:0.1611070307,WP_130637964.1-Pectobacterium-versatile:0.0728105453)69.9/100:0.0279907900)86.6/100:0.0399907410)96.9/100:0.0989913752,((((((WP_168640580.1-Dickeya-sp.-CFBP-2040:0.1404907079,(WP_027712673.1-Dickeya-chrysanthemi:0.0552386455,WP_012770533.1-Dickeya:0.0427881102)75.9/94:0.0088490704)73.6/80:0.0086820543,(WP_012883965.1-Dickeya-parazeae:0.0861115596,WP_236306746.1-Dickeya-zeae:0.0422631106)77.4/98:0.0188463761)74.6/81:0.0081732047,((WP_038918362.1-Dickeya:0.0376685901,WP_123251184.1-Dickeya-undicola:0.0608920752)46.6/100:0.0092017130,WP_022632763.1-Dickeya-solani:0.0339007096)89.5/100:0.0283201968)98.8/100:0.1324669668,(WP_148877341.1-Serratia-marcescens:0.0859456373,WP_193158894.1-Serratia-marcescens:0.0643228360)95.6/100:0.0860926382)76.2/99:0.0430218031,WP_062791489.1-Serratia-plymuthica:0.0927746943)73.2/73:0.0283501399,((WP_121576612.1-Brenneria-alni:0.1175832556,WP_095845203.1-Gibbsiella-quercinecans:0.0980411111)3.8/37:0.0073888682,((WP_214648782.1-Pectobacterium-carotovorum:0.0449776996,((WP_261846379.1-Pectobacterium-carotovorum:0.0749080466,WP_121590317.1-Brenneria-goodwinii:0.0080589413)93.7/100:0.0372162707,(WP_182099415.1-Pectobacterium-aroidearum:0.0448329618,WP_039482612.1-Pectobacterium:0.0354482889)71.2/100:0.0092483661)22.1/99:0.0088076982)0/99:0.0000020831,WP_249228983.1-Brenneria-tiliae:0.0662175453)96.2/99:0.0580228799)84.4/56:0.0243444428)75.5/75:0.0185656768)93.3/93:0.0520598481,WP_248464278.1-Pectobacterium-versatile:0.2111550795)88.7/89:0.0406019545)74.4/78:0.0107647645,WP_193937755.1-Klebsiella-grimontii:0.2609897820)72.7/68:0.0219768135)93.3/88:0.0818929197,(WP_246757812.1-Erwinia-phyllosphaerae:0.2613469566,WP_232759612.1-unclassified-Pantoea:0.1534950383)94.7/98:0.1240747145)24.1/55:0.0137071512,((WP_146378871.1-Cronobacter-turicensis:0.0757950186,WP_032180402.1-Escherichia-coli:0.1231361542)99.8/100:0.2494515769,((WP_126816534.1-Enterobacter-mori:0.1134623065,WP_049841125.1-Klebsiella-sp.-RIT-PI-d:0.5190764564)54.5/74:0.0507981578,WP_256879438.1-Escherichia-albertii:0.1137730767)90/74:0.0756903534)91.5/85:0.0634883500)78.6/62:0.0583299704,WP_253460089.1-Erwinia-persicina:0.3887328488)72.7/72:0.0422821767,(((WP_168836617.1-Vibrio-agarilyticus:0.1603334907,(WP_076537303.1-Shewanella-sp.-UCD-KL21:0.0274820893,WP_102531614.1-Shewanella-sp.-10N.286.52.B9:0.0688219701)100/100:0.7532322610)89.8/98:0.1259830557,(WP_137671336.1-Agarivorans-sp.-Toyoura001:0.0784685927,WP_221077378.1-Agarivorans-albus:0.0494243505)93/100:0.1073501487)97.3/98:0.1481118767,WP_100256321.1-Reinekea-forsetii:0.8658361551)0/46:0.0000021739)82.5/99:0.0391028553,(((WP_145010177.1-Pseudomonas-psychrotolerans:0.0226629841,WP_077172422.1-Pseudomonas-psychrotolerans:0.0542110361)100/100:0.2935072802,WP_084334293.1-Pseudomonas-indica:0.1370634967)94.6/100:0.1276454119,(WP_261844981.1-Amphritea-ceti:0.4194807582,((((((WP_269124033.1-Marinomonas-sp.-15G1-11:0.0395002584,WP_191602238.1-Marinomonas-algicola:0.1085231118)99/100:0.1792048976,WP_048693477.1-Catenovulum-maritimum:0.2757963662)91.8/100:0.0948802322,WP_228191388.1-Marinomonas-communis:0.2032387897)0/13:0.0129609371,WP_067216963.1-Marinomonas-gallaica:0.0926901544)79.9/90:0.0324377446,WP_067204681.1-Marinomonas-aquimarina:0.0331742540)86.9/90:0.0329019611,WP_055461762.1-Marinomonas-fungiae:0.0734716717)81.3/96:0.0541429334)93.3/100:0.1222994855)74.2/88:0.0494928279)94.6/96:0.1198509435,(WP_199145406.1-Citrobacter-sp.-ASV29:0.4115350148,WP_148513713.1-Vibrio:0.2499757289)89.9/99:0.1261321032)64.8/82:0.0093552372,((((WP_138320019.1-Thalassotalea-litorea:0.2003587614,WP_136736185.1-Thalassotalea-mangrovi:0.1697414539)99.9/100:0.3918903266,WP_114327629.1-Candidatus-Colwellia-aromaticivorans:0.1524961335)85.9/99:0.0755498086,WP_101108574.1-Psychromonas-sp.-psych-6C06:0.3165958375)20/92:0.0409240882,((WP_016794198.1-Vibrio-cyclitrophicus:0.0534298324,WP_102424249.1-Vibrio-cyclitrophicus:0.0589993048)87.7/100:0.0456566252,(WP_261903470.1-Vibrio-fortis:0.1047569134,WP_240207223.1-Vibrio-sp.-CyArs1:0.0456535513)90/100:0.0466025773)98.4/100:0.1670075398)91.9/95:0.0962034451)83.6/96:0.0422552830,(((((WP_127021653.1-Rheinheimera-mangrovi:0.4334294604,WP_132584255.1-Rheinheimera-sp.-D18:0.4662662100)16.2/70:0.0401729614,(WP_131912015.1-Celerinatantimonas-diazotrophica:0.2702232685,(((WP_027875186.1-Spongiibacter-marinus:0.2030693799,WP_204995064.1-Spongiibacter-marinus:0.1177523606)75.3/100:0.0442690442,((((WP_184464382.1-Zhongshania-antarctica:0.0595710653,WP_198570926.1-Spongiibacter-nanhainus:0.1138242577)69.9/99:0.0065298314,WP_103683243.1-Marortus-luteolus:0.0396126693)98.9/99:0.1433048437,WP_062382560.1-Zhongshania-aliphaticivorans:0.0663353094)83.6/99:0.0577177778,WP_168451636.1-Spongiibacter-thalassae:0.0561831956)90.5/99:0.0620919619)74.8/99:0.0489988062,((WP_092385961.1-Halopseudomonas-salegens:0.2290709292,WP_036859756.1-Porticoccus-hydrocarbonoclasticus:0.3369222493)91.6/97:0.1434200083,((WP_076722556.1-Marinobacter-lutaoensis:0.2961661863,((WP_106695101.1-Marinobacter-shengliensis:0.0768279701,(WP_222473125.1-Marinobacter-daepoensis:0.0701828656,WP_092030923.1-Marinobacter-sp.-DSM-26671:0.1514373431)86.2/19:0.0429913733)0/15:0.0000020454,WP_148609564.1-Marinobacter-sp.-BW6:0.1365402421)96.4/99:0.1333765591)27.6/90:0.0488499863,WP_036130839.1-Marinobacter-nitratireducens:0.1465464106)70.5/91:0.0666291810)77.3/100:0.0713264548)87.6/97:0.0731399208)76/96:0.0562750986)97.2/99:0.1525480587,WP_067584343.1-Endozoicomonas-ascidiicola:0.2561723655)79.6/88:0.0613737305,WP_150896168.1-Vibrio-fortis:0.3158024995)49.4/86:0.0549244743,((WP_260491935.1-Pseudoalteromonas-flavipulchra:0.2701626518,(WP_039494249.1-Pseudoalteromonas-flavipulchra:0.1212469131,WP_055022295.1-Pseudoalteromonas-sp.-P1-26:0.0801049990)33.3/93:0.0435988295)81/100:0.0357472550,(WP_149602089.1-Pseudoalteromonas-distincta:0.0491822454,WP_248518672.1-Pseudoalteromonas-sp.-2CM32C:0.0336972243)97.9/100:0.1434117461)99.6/100:0.2591331409)32/42:0.0296262746)98.7/100:0.1749735622,(((((((((WP_094934028.1-Enterobacter-cloacae-complex:0.7514743725,WP_016795256.1-Vibrio-cyclitrophicus:0.2012740140)87.9/92:0.1169009284,WP_039473409.1-Vibrio-sinaloensis:0.3609232080)87.3/92:0.0839306587,(WP_010368546.1-Pseudoalteromonas-piscicida:0.5073364085,WP_086590801.1-Vibrio-parahaemolyticus:0.4859080144)81.9/98:0.0718153898)89.6/94:0.0938317755,(((WP_269408820.1-Pseudoalteromonas-shioyasakiensis:0.2505605629,WP_207379479.1-Shewanella-sedimentimangrovi:0.2496984729)95.5/100:0.0994066859,(WP_061013169.1-Vibrionaceae:0.3001299956,WP_155662549.1-Aliivibrio-fischeri:0.1754689700)85.6/74:0.0984177695)0/70:0.0000027003,WP_171328835.1-Vibrio-splendidus:0.3166428181)92.9/92:0.0755319336)79.2/88:0.0656560164,WP_016537220.1-Cedecea-davisae:0.2189828850)72.8/48:0.0205901468,(WP_195504469.1-Citrobacter-farmeri:0.3404522840,WP_064554809.1-Buttiauxella-noackiae:0.3639618519)99.2/100:0.2913414134)79/50:0.0462037451,(WP_240304790.1-Vibrio-campbellii:0.6187480560,WP_004741399.1-Vibrio:0.3040786615)90/90:0.1804242343)90.2/68:0.0705233615,(WP_057065012.1-Citrobacter:0.7702026017,WP_001100815.1-Escherichia-coli:0.3665810998)93.3/95:0.1663413274)91.6/100:0.1122116301,WP_216735790.1-Hahella-sp.-HN01:0.3513301214)79/98:0.0646836707)89.6/99:0.1513381044,(WP_239493201.1-Luteitalea-sp.-TBR-22:0.4439983210,WP_058867107.1-Chloracidobacterium-thermophilum:0.5068053628)98.3/99:0.3284015316)84.4/86:0.0903540420)85.4/23:0.0914203073,((((((((((((GJNO01008138.1-Laxus-oneistus-TSA:1.6781230482,GGJS01006545.1-Plectus-murrayi-TSA:0.3293360824)81.9/27:0.1873976562,(((((VDM95900.1-Thelazia-callipaeda:0.2780531066,(VDO27950.1-Onchocerca-flexuosa:0.1488647594,(VDO31788.1-Brugia-timori:0.1392891992,(((XP_003135693.1-Loa-loa:0.0973510974,VDK73328.1-Litomosoides-sigmodontis:0.0614411123)62.8/73:0.0128100405,CAG9536712.1-Cercopithifilaria-johnstoni:0.0367975541)78.7/73:0.0157980261,VBB33646.1-Acanthocheilonema-viteae:0.1094562639)84.3/97:0.0322223257)82.2/97:0.0387405646)97.3/100:0.1514705378)99.9/100:0.4038063525,(VDD97390.1-Enterobius-vermicularis:0.4595843339,KHN87800.1-Toxocara-canis:0.2229127322)84.2/79:0.0741977588)70.2/75:0.0672532396,VDN56932.1-Dracunculus-medinensis:0.4134643121)95.2/100:0.1731985230,(KAI1709308.1-Ditylenchus-destructor:0.1596238695,KAI1730870.1-Ditylenchus-destructor:0.0310645021)99.3/100:0.2252938593)90.7/15:0.0767035546,TKR72632.1-Steinernema-carpocapsae:0.2931325767)31.7/11:0.0110058091)93.7/82:0.1017962496,KAE9552274.1-Halicephalobus-sp.-NKZ332:0.3754310561)79.7/84:0.1310457190,((CAD5228290.1-Bursaphelenchus-okinawaensis:0.0424419312,((KAI6206096.1-Aphelenchoides-besseyi:0.1895026666,KAI6191010.1-Aphelenchoides-bicaudatus:0.1053631000)64/100:0.0518146723,KAI6234710.1-Aphelenchoides-fujianensis:0.1329233634)99.7/100:0.1894493487)77.6/94:0.0260423539,CAD5235627.1-Bursaphelenchus-xylophilus:0.0282901608)98.4/100:0.1610150085)81.9/59:0.0830693555,KAH7695359.1-Aphelenchus-avenae:0.3580878976)99.3/100:0.1950403938,(PAV65180.1-Diploscapter-pachys:0.2992758349,((KHJ99538.1-Oesophagostomum-dentatum:0.0678247567,EPB80542.1-Ancylostoma-ceylanicum:0.0529709274)96.9/100:0.1134309441,((VDO65958.1-Haemonchus-placei:0.1683052131,VDL70484.1-Nippostrongylus-brasiliensis:0.1103838997)87.8/100:0.0335931164,VDO95824.1-Heligmosomoides-polygyrus:0.0908879275)99.2/100:0.1599015634)98.7/100:0.1825552408)34/65:0.0432290836)79.4/66:0.0326894311,CAI4229259.1-Auanema-sp.-JU1783:0.5114961309)91/97:0.1119622664,(CAD6189320.1-Caenorhabditis-auriculariae:0.4976483814,(CAB3400093.1-Caenorhabditis-bovis:0.2310902278,(NP_509046.1-Caenorhabditis-elegans:0.0235737437,(XP_003117635.1-Caenorhabditis-remanei:0.0335100918,CAI5454992.1-Caenorhabditis-angaria:0.0868634628)0/5:0.0202665251)94.5/94:0.1057834751)90.9/100:0.1204881664)80.4/100:0.0895383632)99.9/100:0.8829142516,(WP_042281872.1-Candidatus-Protochlamydia-sp.-R18:0.1135800178,WP_068471358.1-Parachlamydia-sp.-C2:0.2575159169)100/100:1.2686094610)90.1/80:0.3150126284,(WP_161826415.1-Petrachloros-mirabilis:0.6474600377,WP_099324654.1-Candidatus-Kuenenia-stuttgartiensis:0.7145970365)82.5/63:0.1059650669)87.1/45:0.1134587212,((WP_255135177.1-Gracilimonas-sp.-CAU-1638:0.4103367911,WP_020401568.1-Gracilimonas-tropica:0.3497830174)99.9/100:0.6675179759,WP_008811140.1-Parasutterella-excrementihominis:0.7230794738)62.5/57:0.1469993379)92.7/50:0.1201932149,(((((((((WP_176021856.1-Stenotrophomonas-sp.-NA06056:0.0528099758,WP_049440708.1-Stenotrophomonas-maltophilia:0.0621237858)97.6/100:0.1908678843,WP_132999175.1-Luteimonas-arsenica:0.2908450480)95.1/98:0.1488056901,WP_078758981.1-Lysobacter-spongiicola:0.3113819208)87.9/97:0.0869360113,(((((WP_019467068.1-Dyella-japonica:0.2820835193,WP_204636097.1-Dyella-kyungheensis:0.1683062719)92/96:0.0525813780,WP_049621948.1-Frateuria-defendens:0.1711705669)76.5/57:0.0242498359,((((((WP_131151171.1-Dyella:0.2602555909,WP_130619635.1-Dyella-amyloliquefaciens:0.0770364730)81.7/78:0.0339268129,WP_199044928.1-Dyella-sp.-ASV24:0.0951827231)71.5/34:0.0091736964,(WP_266169930.1-Dyella-sp.-RXD159:0.1020654788,(WP_114846719.1-Dyella-tabacisoli:0.0602494496,WP_090451016.1-Dyella-sp.-OK004:0.0884807139)85.7/61:0.0324884773)80.6/61:0.0159509030)75.5/35:0.0099091098,WP_214570707.1-unclassified-Dyella:0.2031187698)79.2/56:0.0242931954,(WP_111981026.1-Dyella-jiangningensis:0.2001013512,WP_266183414.1-Dyella-sp.-RBB1W86:0.1011906277)85.8/56:0.0368439167)93.3/83:0.0677146660,((WP_233843762.1-Dyella-sp.-2HG41-7:0.0472950850,WP_188793403.1-Dyella-nitratireducens:0.0895652924)31.7/96:0.0452981383,WP_126673574.1-Dyella-dinghuensis:0.0708331148)99/100:0.1302800276)75.7/55:0.0142042674)87.1/35:0.0294647595,(((((WP_063674027.1-Dyella-thiooxydans:0.1827267534,WP_253565187.1-Dyella-sp.-Sa:0.0828069629)84.1/92:0.0340782998,(WP_209622196.1-Frateuria-flava:0.0947055800,WP_091339503.1-Frateuria-terrea:0.0557835990)97.1/100:0.0790624640)24.1/78:0.0192429227,WP_201315287.1-Dyella-sp.-EPa41:0.1770720096)78.3/92:0.0245583561,(WP_188800319.1-Dyella-caseinilytica:0.1945611640,WP_192554051.1-Dyella-acidiphila:0.1590516063)97.1/100:0.1214100221)81.1/74:0.0270403137,(WP_179481164.1-unclassified-Rhodanobacter:0.1708973724,((((((WP_214557828.1-unclassified-Rhodanobacter:0.0424393789,WP_068095172.1-unclassified-Rhodanobacter:0.0388664736)32.9/100:0.0258623709,(WP_008437080.1-Rhodanobacter-thiooxydans:0.0536696878,(WP_063107523.1-Rhodanobacter-thiooxydans:0.0390097851,WP_211005527.1-Rhodanobacter-sp.-B2A1Ga4:0.0472384426)44.8/100:0.0084246620)58.1/100:0.0236364367)96.5/100:0.0630700367,WP_072762453.1-Rhodanobacter-sp.-OK091:0.0147082670)71.7/91:0.0100958601,((WP_077439480.1-Rhodanobacter-sp.-C01:0.1356449741,WP_158752649.1-Dyella-sp.-S184:0.0313828980)55.4/96:0.0170152088,WP_077445946.1-Rhodanobacter-sp.-C05:0.0623783317)83.6/96:0.0254874502)68.3/90:0.0253828972,WP_184416890.1-Rhodanobacter-sp.-MP7CTX1:0.0677474086)90.4/100:0.0469424980,WP_015446789.1-Rhodanobacter-denitrificans:0.0566684997)78.4/100:0.0273945662)91.2/77:0.0458414136)34.1/50:0.0197136770)82.9/78:0.0362309071,((WP_168608661.1-Oleiagrimonas-citrea:0.0781913552,WP_113064358.1-Oleiagrimonas-sp.-MCCC-1A03011:0.0245200468)79.4/100:0.0491337944,WP_043099571.1-Oleiagrimonas-soli:0.1040384366)100/100:0.1802433769)95.1/99:0.1082708745)99.6/100:0.3052076430,(WP_256181981.1-Cloacibacillus-evryensis:0.2955827340,WP_087368268.1-Cloacibacillus-sp.-An23:0.1347117192)100/100:0.7067112239)82.8/65:0.0516205399,((WP_084578244.1-Sporomusa-malonica:0.5748235111,WP_131919244.1-Heliophilum-fasciatum:0.3588703037)66.5/95:0.0406958902,((WP_090940141.1-Pelosinus-propionicus:0.3324803879,WP_240986382.1-Acididesulfobacillus-acetoxydans:0.3415858676)84.2/100:0.0871177166,WP_092486361.1-Desulfoscipio-geothermicus:0.1806834990)84.9/100:0.0814983316)89.2/93:0.0918701996)80.4/53:0.0322137133,((WP_246588317.1-Pelorhabdus-rhamnosifermentans:0.2470529314,((WP_092074275.1-Dendrosporobacter-quercicolus:0.3402662074,(WP_145090815.1-Sporomusa-sp.-KB1:0.0563327783,WP_093793074.1-Sporomusa-acidovorans:0.0897621123)94.6/100:0.1047105384)55.4/99:0.0851657934,(((WP_007952170.1-Pelosinus-fermentans:0.1334720927,WP_007939097.1-Pelosinus:0.1362478908)93.9/100:0.0906896942,(((WP_132081660.1-Anaerospora-hongkongensis:0.1233007258,WP_132083235.1-Anaerospora-hongkongensis:0.2419450202)96.7/100:0.1463025276,WP_188399481.1-Sporomusa-sp.-GT1:0.3498683376)61.3/42:0.0228406556,((WP_094603851.1-Sporomusa-silvacetica:0.1559353593,WP_144352952.1-Sporomusa-termitida:0.1720565780)89.2/100:0.0886777842,WP_054258662.1-Propionispora-sp.-2-2-37:0.2903321960)73.8/47:0.0659547691)70.4/75:0.0468180892)78.2/78:0.0297429168,(WP_054258683.1-Propionispora-sp.-2-2-37:0.1867322964,((WP_149735423.1-Propionispora-hippei:0.1017865342,WP_091744506.1-Propionispora-vibrioides:0.1763667393)66.1/76:0.0287210347,WP_188400350.1-Sporomusa-sp.-GT1:0.3839610508)50.2/44:0.0319171525)95.6/95:0.1090178239)83.4/96:0.0496011581)0/58:0.0215581494)80.9/97:0.0553008944,WP_110954047.1-Massilibacillus-massiliensis:0.3513370562)95.2/100:0.1534005489)86/74:0.0674185192,(((((((WP_200343816.1-Halochromatium-glycolicum:0.2085398946,WP_201222000.1-Halochromatium-roseum:0.1593215519)96/100:0.1579170977,(WP_200237937.1-Thiohalocapsa-halophila:0.0864388632,WP_058558062.1-Thiohalocapsa-sp.-ML1:0.2983207949)96.5/100:0.1277082118)97.9/100:0.1752912640,(WP_182584206.1-Thiospirillum-jenense:0.5221001390,(WP_200286492.1-Rhabdochromatium-marinum:0.1654576072,WP_009150811.1-Thiorhodovibrio-sp.-970:0.0839279804)91.1/100:0.1233616510)53.8/99:0.1452455503)92.7/100:0.1000967561,WP_018175268.1-unclassified-Thioalkalivibrio:0.5612040005)77.7/92:0.0235233395,((((((WP_200207675.1-Chromatium-weissei:0.2628073976,WP_200154756.1-Chromatium-okenii:0.1253838488)98.7/92:0.1715199140,(WP_200158060.1-Allochromatium-vinosum:0.0716721349,WP_200376120.1-Thiocystis-violacea:0.1379440423)81.7/100:0.0505771908)68/81:0.0221443803,(((WP_043749566.1-Imhoffiella-purpurea:0.0933127398,WP_007039504.1-Thiorhodococcus-drewsii:0.0768641861)86/100:0.0400264799,((WP_164456073.1-Thiorhodococcus-minor:0.1158307729,WP_200330387.1-Thiocystis-violacea:0.1093957985)82.8/99:0.0281835736,WP_164652045.1-Thiorhodococcus-mannitoliphagus:0.0569790830)94.4/100:0.0762485328)89.5/99:0.0295769186,(WP_201092787.1-Thiocystis-minor:0.0904968148,WP_132978621.1-Thiobaca-trueperi:0.1249058330)83.6/92:0.0283251130)85.9/91:0.0311580608)86/91:0.0434367199,(WP_200388819.1-Thiocapsa-imhoffii:0.2179547522,((WP_007191850.1-Thiocapsa-marina:0.0515517171,WP_120796078.1-Thiocapsa-rosea:0.0442326935)74.2/100:0.0080425361,WP_232409912.1-Thiocapsa-bogorovii:0.0264946938)96.8/100:0.1038170116)94.2/100:0.1039611729)95.7/100:0.1189585792,(WP_100918325.1-Candidatus-Thiodictyon-syntrophicum:0.1047132046,WP_020502667.1-Lamprocystis-purpurea:0.1315790748)94.5/100:0.1105753732)75.9/99:0.0804581157,WP_089729519.1-Candidatus-Thiosymbion-oneisti:0.3987498396)88.2/95:0.0740605734)91.2/93:0.1076033721,(WP_141730858.1-Oligoflexus-tunisiensis:0.5379284745,WP_101758446.1-Oceanicoccus-sp.-KOV_DT_Chl:0.5294973327)0/51:0.1814790188)34/69:0.1082446374,WP_141198172.1-Persicimonas-caeni:0.5963197341)83.5/93:0.1144988123)92.3/96:0.1386681398,WP_207681674.1-Desulfonema-magnum:0.7631471149)49.3/47:0.0643867306)89.1/80:0.0684111748)84.3/27:0.0469655002)56.7/56:0.0324289874,WP_230840800.1-Gloeobacter-morelensis:0.6388374262)89.3/24:0.0782796346,((WP_110265962.1-Sinimarinibacterium-flocculans:0.5033575787,WP_077329105.1-Virgibacillus-siamensis:0.5712657268)90.7/96:0.2195381706,((((((((((((WP_160564586.1-Apibacter-sp.-B2912:0.1908954974,WP_055424523.1-Apibacter-mensalis:0.1200062065)0/4:0.0406414208,(WP_160568957.1-unclassified-Apibacter:0.0305589699,WP_166004639.1-Apibacter-sp.-B2966:0.0508932152)96.5/100:0.1044660166)96.4/100:0.1296953762,(WP_105245553.1-Apibacter-adventoris:0.2298679428,WP_146261858.1-Apibacter-muscae:0.2008994917)77.5/99:0.0897523269)100/100:0.3731927152,(WP_068596492.1-Vaginella-massiliensis:0.2129865072,((WP_230477342.1-Faecalibacter-bovis:0.1443315129,WP_125349156.1-Empedobacter-falsenii:0.1343644916)76.6/98:0.0442950620,(WP_125113420.1-Algoriella-xinjiangensis:0.1157778757,WP_092908829.1-Algoriella-xinjiangensis:0.0433894569)93.3/100:0.0667500712)96.9/98:0.1621351426)98.6/100:0.2248561255)74.8/75:0.0510386222,(WP_084015419.1-Moheibacter-sediminis:0.4972398947,WP_182042771.1-Moheibacter-lacus:0.3768810521)91.6/98:0.1643831653)93.1/99:0.1452343322,(WP_014783622.1-Aequorivita-sublithincola:0.4177333288,((((WP_230143398.1-unclassified-Pedobacter:0.1493252404,WP_215166022.1-Pedobacter-sp.-ISL-64:0.1113792927)93.8/100:0.1066838357,WP_133260761.1-Pedobacter-changchengzhani:0.1624329927)89.4/57:0.0807918384,((WP_056872627.1-Pedobacter-sp.-Leaf194:0.1253716489,(WP_123203876.1-Pedobacter-jejuensis:0.0989524373,(WP_188416210.1-Pedobacter-mendelii:0.0371132416,WP_056087060.1-Pedobacter-sp.-Leaf176:0.0674388843)24.8/77:0.0183772959)39.9/100:0.0225573041)95.9/97:0.0789131963,WP_109929147.1-Pedobacter-paludis:0.0521204401)81.5/91:0.0459940392)56.9/48:0.0360388105,WP_199156598.1-Pedobacter-sp.-ASV2:0.1276807074)99.4/96:0.3241019972)99/100:0.3318233278)87.5/100:0.1671300754,WP_111126400.1-Mesonia-sp.-K7:1.0813450272)0/83:0.0000024146,(((WP_107809536.1-Elizabethkingia-sp.-YR214:0.1969455912,WP_219971314.1-Chryseobacterium-sp.-LJ756:0.1903498747)90.7/78:0.0776575517,WP_076395614.1-Chryseobacterium:0.0584676474)69.4/78:0.0704120173,WP_088467981.1-Chryseobacterium-sp.-VAUSW3:0.1675768227)99.9/100:0.2996697693)95.1/96:0.1858508288,((((WP_205722928.1-Fulvivirga-lutea:0.7010413050,((((((((((WP_053470876.1-unclassified-Flavobacterium:0.2187626887,WP_056200692.1-Flavobacterium-sp.-Root186:0.0406668459)10.2/72:0.0087050722,WP_095952323.1-Flavobacterium-sp.-ACN6:0.0553632871)55/96:0.0158310554,WP_115889823.1-Flavobacterium-cutihirudinis:0.0634273708)76.5/99:0.0088324061,WP_198857739.1-Flavobacterium-sp.-IB48:0.0469837244)72.3/98:0.0082143987,((WP_269237486.1-Flavobacterium-sp.-IMCC34775:0.1073176771,WP_163393939.1-Flavobacterium-limi:0.0475470753)0/100:0.0000026734,WP_264205577.1-Flavobacterium-sp.-LS1R49:0.0471385829)32.7/74:0.0083213237)90.2/99:0.0300346310,((WP_173968237.1-Flavobacterium-collinsii:0.0928883396,WP_264525312.1-Flavobacterium-sp.-N502536:0.0645501106)94.2/99:0.0466101798,WP_070907856.1-Flavobacterium-tructae:0.1066738765)76.6/98:0.0102644588)73.8/98:0.0101414343,WP_091091738.1-Flavobacterium-gillisiae:0.1329605818)97.4/100:0.0866590512,WP_248907802.1-Flavobacterium-sp.-K5-23:0.1669719343)90/99:0.0569953441,(((WP_116761981.1-Flavobacterium-laiguense:0.0534410292,WP_143377960.1-Flavobacterium-sp.-ZT3R18:0.0659485555)96.8/100:0.0844716464,((WP_144071503.1-Flavobacterium-franklandianum:0.0848165874,WP_133277350.1-Flavobacterium-nackdongense:0.1145571240)47.3/99:0.0472075778,WP_103806521.1-Flavobacterium-alvei:0.1391309348)82.6/59:0.0274996449)52/58:0.0100267388,WP_194311471.1-Flavobacterium-soyangense:0.1743016510)84.7/94:0.0782406977)97.9/99:0.2019551545,(WP_140000461.1-Flavobacterium-microcysteis:0.3162479080,(((WP_194640005.1-Flavobacterium-sp.-HJJ:0.1259373398,WP_073246022.1-Flavobacterium-flevense:0.0851391309)65.9/100:0.0246334679,WP_100843428.1-Flavobacterium-sp.-5:0.1443180304)90.9/99:0.0764864438,WP_140487657.1-Flavobacterium-sp.-GSA192:0.0518220205)99.3/100:0.2400628425)93.7/100:0.1400088801)94.8/98:0.1575050388)88.2/95:0.1260481396,WP_057952567.1-Salinivirga-cyanobacteriivorans:0.8314871178)42.5/71:0.0581551394,(WP_116872776.1-Terrimonas-sp.:0.6410107473,(WP_004155558.1-Microscilla-marina:0.4768098288,WP_101358295.1-Raineya-orbicola:0.3463713324)98.2/100:0.4101785643)78.5/87:0.1521427381)93.8/96:0.1385677833,(((((WP_233378072.1-Maribellus-sp.-CM-23:0.0841452258,WP_247978154.1-Maribellus-sp.-YY47:0.0927929963)96.3/100:0.1513157050,(WP_167609329.1-Maribellus-sediminis:0.2213469360,(WP_045033348.1-Draconibacterium-sediminis:0.0326587157,WP_163345719.1-Draconibacterium-halophilum:0.1053993832)99.1/100:0.2086129220)93.1/100:0.1394988011)99.9/100:0.4007581668,(WP_106543043.1-Prolixibacter-denitrificans:0.1524061790,(WP_025865917.1-Prolixibacter-bellariivorans:0.0268524021,WP_153632016.1-Prolixibacter-sp.-SD074:0.0732456263)0/93:0.0000020569)99.7/100:0.3165057137)74.3/98:0.0982331640,((WP_159517929.1-Sunxiuqinia-indica:0.1479388238,WP_053182762.1-Sunxiuqinia-dokdonensis:0.1422183971)87.1/100:0.0717419533,WP_133466511.1-Sunxiuqinia-elliptica:0.1977314575)98.7/100:0.2236334990)95/99:0.1285187233,((WP_068701711.1-Paludibacter-jiangxiensis:0.5263045735,(WP_011500232.1-Methanococcoides-burtonii:0.2909267597,((((((WP_128503182.1-Methanosarcina-sp.-MSH10X1:0.2231276487,WP_011033115.1-Methanosarcina-mazei:0.0886181783)12.3/39:0.0413511284,WP_048134921.1-Methanosarcina-sp.-1.H.T.1A.1:0.0447478754)94.9/95:0.0873885603,WP_048167026.1-Methanosarcina:0.1229987940)41/81:0.0469901990,WP_048177163.1-Methanosarcina-sp.-MTP4:0.1046647518)85.7/93:0.0950263547,(WP_048137589.1-Methanosarcina-horonobensis:0.1889457695,((WP_048184631.1-Methanosarcina-siciliae:0.0748068989,WP_048173663.1-Methanosarcina-siciliae:0.0759039973)0.2/48:0.0147349484,WP_011023680.1-Methanosarcina-acetivorans:0.0458511760)96/100:0.1079307774)73.6/86:0.0849363411)95.4/97:0.1438034783,WP_135611779.1-Methanococcoides-sp.-AM1:0.2662047110)43.5/31:0.0275286121)97.8/97:0.2426233183)53.4/95:0.0840748949,((WP_101263396.1-Labilibaculum-filiforme:0.0506751031,((WP_152734748.1-Marinifilum-sp.-N1E240:0.1066771165,(WP_171595603.1-Marinifilum-caeruleilacunae:0.0841499018,(WP_268189532.1-Marinifilum-sp.-D737:0.0655592723,WP_054720444.1-Marinifilum-fragile:0.0422397993)88.4/100:0.0271778724)88.9/100:0.0333089380)83.3/100:0.0537815146,WP_127342456.1-Ancylomarina-longa:0.2083856404)98.7/85:0.1256328841)45.8/66:0.0292822320,WP_245870435.1-Labilibaculum-manganireducens:0.0414638280)99.9/100:0.3180248931)73.6/95:0.0566494952)78.2/92:0.1388554843)77.9/79:0.1102538359)55.4/96:0.1020136203,(((((WP_013633636.1-Pseudopedobacter-saltans:0.1848810900,WP_154287284.1-Pedobacter-puniceum:0.2196461414)91.5/100:0.0930567122,WP_160842504.1-Pedobacter-sp.-HMF7647:0.0931711173)91.6/100:0.1485837525,(WP_074492796.1-Mucilaginibacter-polytrichastri:0.2412909938,(WP_258137696.1-Mucilaginibacter-sp.-BJC16-A38:0.2436603399,(WP_259068220.1-Mucilaginibacter-sp.-X4EP1:0.0759659838,WP_144912900.1-Mucilaginibacter-frigoritolerans:0.1093234980)53.5/85:0.0894118225)86.8/100:0.1610731875)69/76:0.1623321884)99.8/100:0.4055213934,WP_130092062.1-Brumimicrobium-glaciale:0.6283118486)23.5/91:0.0757438647,WP_163286637.1-Cryomorpha-ignava:0.7762145016)98.4/100:0.2749884102)87.1/37:0.1494955392,((WP_040596643.1-Prevotella-stercorea:0.1170397704,WP_207153370.1-Prevotella-herbatica:0.1184705587)99.9/100:0.3683175789,(WP_223927321.1-Prevotella-lacticifex:0.3318045876,WP_009010825.1-Prevotella-sp.-C561:0.2026368483)99.8/100:0.5656945814)82.1/93:0.1701210560)83.5/22:0.1223553313,(WP_085659180.1-Prosthecochloris-sp.-HL-130-GSB:0.5925365820,WP_038563105.1-Draconibacterium-orientale:0.4573065565)88.2/100:0.1376373337)99.6/99:0.3661542951)22.4/10:0.0483246038)92.5/89:0.0947205516)92.1/97:0.0892848103,((WP_133132051.1-Legionella-sp.-W10-070:0.2758570362,((((((((((((WP_027223798.1-Legionella-pneumophila:0.0382604061,WP_265717942.1-Legionella-sp.-PATHC032:0.0475714410)92.2/99:0.0367414889,WP_106142999.1-Legionella-pneumophila:0.0457428050)69.7/92:0.0056127153,(WP_061513621.1-Legionella:0.0158550975,WP_027226296.1-Legionella-pneumophila:0.0701099783)99.8/93:0.0821648334)86.5/99:0.0169493295,WP_044499121.1-Legionella-pneumophila:0.0310145517)87.3/99:0.0201318923,WP_027220106.1-Legionella-pneumophila:0.0199985471)81.5/99:0.0227084853,WP_027266778.1-Legionella-pneumophila:0.0427109663)96.6/100:0.1007194870,(WP_058494182.1-Legionella-worsleiensis:0.1815144386,(WP_127106294.1-Legionella-sp.-km535:0.0512163004,WP_058474153.1-Legionella-quateirensis:0.0390366501)89.4/99:0.0497492012)94.8/100:0.0880184607)94.5/99:0.0897625093,WP_031567420.1-Legionella-wadsworthii:0.2488106256)64.2/90:0.0212732542,WP_045097199.1-Legionella-fallonii:0.1557000469)87.1/90:0.0358730367,(WP_232890831.1-unclassified-Legionella:0.0535215105,(WP_058459148.1-Legionella-bozemanae:0.0295369861,WP_058518697.1-Legionella-parisiensis:0.0592551956)85.1/100:0.0356196157)98.7/100:0.1168479645)55.2/82:0.0286626848,(WP_058463758.1-Legionella-cincinnatiensis:0.0265807686,WP_128130294.1-Legionella-sainthelensi:0.0673691683)93.3/100:0.0453660410)0/80:0.0000028815,WP_058513550.1-Legionella-santicrucis:0.1368901621)99.9/100:0.4754308331)98.2/100:0.2972151573,(WP_205000141.1-Parashewanella-hymeniacidonis:0.0654659989,WP_129946484.1-Shewanella-sp.-OPT22:0.0763045938)100/100:0.7905413884)89.4/100:0.1235236610)97/100:0.2020486057)85.2/84:0.0502909697)92.2/99:0.0630801045)78.4/99:0.0303723075,WP_171149131.1-Erwinia-sp.-JH02:0.2224344246)98.8/100:0.1260087049,((WP_241647798.1-Rosenbergiella-sp.-JB07:0.3845109619,WP_065370581.1-Kosakonia-sacchari:0.2373349152)87.9/100:0.0546815574,WP_181599127.1-Citrobacter-sp.-RHB36-C18:0.2976632192)77.6/99:0.0255378961)89.2/99:0.0668107845)81.6/93:0.0354311897,((((((((((((((WP_182931197.1-Enterobacter-cloacae:0.2156198695,WP_211776196.1-Klebsiella-pneumoniae:0.0826270274)60.5/96:0.0109862742,WP_047723323.1-Klebsiella-michiganensis:0.0550372492)76.8/83:0.0107612878,(WP_048284323.1-Pluralibacter-gergoviae:0.1808904799,((WP_139968923.1-Klebsiella-pneumoniae:0.1227346793,WP_065799756.1-Klebsiella-pneumoniae:0.0074672031)0/91:0.0000020865,WP_095100910.1-Raoultella-sp.-10-1:0.0863123173)69.6/90:0.0094097536)91.2/95:0.0370117706)84.9/74:0.0159702955,WP_104011323.1-Klebsiella-michiganensis:0.0476265143)88.5/87:0.0233184929,WP_234579544.1-Raoultella-terrigena:0.0314344723)53.2/54:0.0077391872,WP_186601692.1-Klebsiella-quasipneumoniae:0.0393597936)84.2/61:0.0159731756,WP_149270444.1-Klebsiella-aerogenes:0.0582792797)78.2/40:0.0093090289,(WP_213760001.1-Klebsiella-aerogenes:0.0858827958,WP_223370324.1-Klebsiella-pneumoniae:0.0393816946)45.1/80:0.0150388840)50.5/27:0.0037599624,WP_221694709.1-Klebsiella-aerogenes:0.1269502064)70.7/75:0.0203662398,WP_128320364.1-Raoultella-planticola:0.0644956338)61.8/70:0.0095182161,((WP_234537246.1-Raoultella-ornithinolytica:0.0632018837,WP_113990968.1-Raoultella:0.0187964385)99.1/99:0.0654892634,WP_046852082.1-Raoultella-planticola:0.0463603824)55.6/12:0.0114674818)0/7:0.0000024391,(WP_236948387.1-Klebsiella-sp.-2019SCSN059:0.0464453602,((WP_114692711.1-Klebsiella-pneumoniae:0.0471828079,WP_259891680.1-Klebsiella-variicola:0.0387554429)0/30:0.0000028737,WP_123755035.1-Raoultella-sp.-BIGb0399:0.0457106857)84.6/73:0.0141174252)77.9/52:0.0124446841)96.9/83:0.1038404454,((WP_095100919.1-Raoultella-sp.-10-1:0.0733230052,WP_212709529.1-Tatumella-sp.-JGM130:0.0177687216)98.3/100:0.1077530829,WP_236948353.1-Klebsiella-sp.-2019SCSN059:0.1094917083)76.3/84:0.0271937236)87.8/100:0.0624382899,WP_123822891.1-Klebsiella:0.0343320417)98.9/100:0.1285225370)87.7/99:0.0590310799,WP_072663757.1-Enterobacteriaceae:0.1146737436)31.4/87:0.0383061871,WP_021241446.1-Enterobacter-roggenkampii:0.2464989933)95.9/100:0.1003607402)95.3/100:0.0709943625,XP_011503919.1-Ceratosolen-solmsi-marchali:0.1118804155);

**Nematode RML-3, with bacterial and eukaryotic homologs**

**Shown in Figure S2**

ML tree method: IQ-TREE

Sequences: 2787

Substitution model: LG

(WP_063501760.1-Escherichia-coli:0.0336676231,(WP_044173167.1-Phytobacter-massiliensis:0.0260455916,((((((((WP_203451401.1-Lelliottia-sp.-RWM.1:0.0559140089,WP_185654835.1-Citrobacter-braakii:0.0976933602)93.2/100:0.0649611260,WP_183170804.1-Atlantibacter-sp.-RC6:0.1674963591)98.2/99:0.0829053402,(WP_160868038.1-Pantoea-sp.-Taur:0.2220360356,WP_008499438.1-Enterobacter-cloacae-complex:0.1551053172)80.9/95:0.0436196924)52.8/78:0.0147282570,((WP_090085547.1-Escherichia-coli:0.0503879528,WP_103283990.1-Citrobacter:0.1438107312)8.2/44:0.0135812663,WP_234083964.1-Enterobacter-quasiroggenkampii:0.1017960535)100/100:0.1371402706)88.4/98:0.0381818550,((((((((((((((((((((((((((((((((((((((((((((((((((((((((((WP_249615874.1-Craterilacuibacter-sp.-RT1T:0.2920150996,(WP_163316497.1-Crenobacter-sedimenti:0.1166831990,WP_160797365.1-Craterilacuibacter-sinensis:0.1027107583)96.7/100:0.1217883683)82.4/100:0.0424349384,WP_014088374.1-Pseudogulbenkiania-sp.-NH8B:0.1497854956)43.7/97:0.0435962529,WP_008952446.1-Pseudogulbenkiania-ferrooxidans:0.1816634795)0/84:0.0290029049,(((((WP_147247296.1-Brenneria-salicis:0.0329450553,WP_172289906.1-unclassified-Brenneria:0.0843713769)87.5/100:0.0319854749,WP_039356706.1-Pectobacterium-actinidiae:0.0703249402)69.9/61:0.0565837138,((WP_198997094.1-Pectobacterium-carotovorum:0.0156908573,(WP_165707548.1-Pectobacterium:0.0285023617,WP_174866005.1-Pectobacterium-polaris:0.0230108429)99.3/100:0.0543751931)77.6/60:0.0164919903,WP_145962313.1-Pectobacterium-peruviense:0.0448369722)99.4/99:0.1108171738)11.2/73:0.0375646495,(((WP_208149206.1-Shewanella-algae:0.2312246025,WP_011788809.1-Shewanella-sp.-W3-18-1:0.0518420525)49.6/66:0.0125338554,((WP_045500143.1-Vibrio-hyugaensis:0.1373408192,WP_045419977.1-Vibrio-campbellii:0.0678824963)58.4/84:0.0314471939,WP_025533544.1-Vibrio-parahaemolyticus:0.0535830754)93.8/89:0.0346548748)82.6/99:0.0446611070,WP_219142360.1-Aeromonas-sanarellii:0.0994732227)99.5/100:0.1271677447)87.1/99:0.0572861799,(WP_222134508.1-Vibrio-cholerae:0.1412152444,WP_137008594.1-Aquitalea-aquatilis:0.2455587798)9.1/64:0.0441500775)95.9/88:0.0725818907)0/17:0.0153681955,(WP_174873379.1-Vogesella-oryzae:0.1490691802,WP_147289921.1-Crenobacter-cavernae:0.0890989467)92.1/88:0.0535644037)97.1/91:0.0702392216,(((((((((WP_004123988.1-Rhizobium-freirei:0.0824439039,WP_095433322.1-Rhizobium-sp.-11515TR:0.0320420872)80.8/85:0.0165614101,((WP_183712806.1-unclassified-Rhizobium:0.0335367286,WP_015341698.1-Rhizobium:0.0256066186)45.2/75:0.0114855587,WP_106999541.1-Rhizobium-sp.-JAB6:0.0460249734)0/74:0.0000028242)71.2/86:0.0173513060,WP_184706966.1-Rhizobium-lusitanum:0.0785681218)20.1/75:0.0020346240,(WP_174074493.1-Agrobacterium-rhizogenes:0.0784013541,((WP_183765619.1-unclassified-Rhizobium:0.0332855672,WP_120667780.1-unclassified-Rhizobium:0.0448980147)89.1/100:0.0284847114,WP_037195771.1-Rhizobium-lusitanum:0.0218333180)40.2/93:0.0115695419)80.9/70:0.0149581291)76.1/79:0.0076806188,WP_092719658.1-Rhizobium-multihospitium:0.0313480347)90.6/98:0.0232379892,WP_168880117.1-Rhizobium-sp.-P28RR-XV:0.0354773005)75.2/74:0.0058396155,WP_112341687.1-Rhizobium-tropici:0.0345554405)89.3/95:0.0185892963,(WP_199146672.1-Rhizobium-sp.-ASV8:0.1414118624,WP_145321829.1-unclassified-Rhizobium:0.0758068155)16.7/70:0.0085429324)99.7/100:0.1121309606,(WP_043579071.1-Chromobacterium:0.0473252958,WP_071113424.1-Chromobacterium-sphagni:0.0839322640)99.9/100:0.1229195448)94.4/96:0.0529926083)67.8/80:0.0118283158,((WP_189531325.1-Paludibacterium-paludis:0.3015902390,(WP_043620211.1-Chromobacterium-piscinae:0.1410212490,(WP_103900886.1-Chromobacterium-alticapitis:0.1674372079,WP_045052286.1-Chromobacterium-violaceum:0.1627961900)94.5/100:0.0801586915)91.7/100:0.0611512749)82.3/85:0.0299903828,(WP_020932001.1-Cycloclasticus-zancles:0.2229931264,(((WP_213263532.1-Burkholderia-pyrrocinia:0.0912664507,WP_060279477.1-Burkholderia-cepacia-complex:0.1156639752)98.9/100:0.1126290078,(((WP_208101977.1-Burkholderia-sp.-D-99:0.0297456180,WP_205714023.1-Burkholderia-cepacia:0.0258563172)44.5/100:0.0072356531,WP_174919663.1-Burkholderia-metallica:0.0182438011)50.7/99:0.0302182169,WP_077021900.1-Burkholderia-cenocepacia:0.0740309698)100/100:0.1497348711)85.8/91:0.0549907268,(WP_159673755.1-Andreprevotia-sp.-IGB-42:0.1432223686,WP_084093192.1-Andreprevotia-lacus:0.0672718032)99.2/100:0.1108940388)43.2/77:0.0326322362)75.2/45:0.0427460021)40/42:0.0290013785)91/98:0.0561705239,(WP_188630571.1-Oceanisphaera-marina:0.0993822867,(WP_104485062.1-Oceanisphaera-arctica:0.0870519444,WP_204795415.1-Oceanisphaera-litoralis:0.0397164824)46.7/51:0.0170507553)100/100:0.1677728400)90.8/45:0.0522774353,((WP_136867012.1-Guyparkeria-sp.-SB14A:0.1961630920,(((WP_139871681.1-Candidatus-Methylopumilus-universalis:0.1275155799,(WP_139929800.1-Candidatus-Methylopumilus-universalis:0.2187354992,WP_140005423.1-Candidatus-Methylopumilus-planktonicus:0.0574099154)97.9/100:0.1070201672)89.2/100:0.0670442427,WP_019964198.1-Pannonibacter-phragmitetus:0.3918709004)96.2/95:0.0894843301,WP_058575749.1-unclassified-Guyparkeria:0.1483650130)89.1/83:0.0501023593)93.6/82:0.0686830102,((WP_110265962.1-Sinimarinibacterium-flocculans:0.2669551963,(WP_234070476.1-Myxococcus-stipitatus:0.2047612743,WP_120555541.1-Corallococcus-aberystwythensis:0.1899298359)99/100:0.1692538486)75.2/100:0.0662291804,(((((WP_169107474.1-Comamonas-suwonensis:0.0437857517,WP_219343559.1-Comamonas-sp.-Y33R10-2:0.0690912339)95.2/100:0.0443725516,(WP_003073420.1-Comamonas:0.0613152042,((WP_211457163.1-Comamonas-sp.-PE63:0.0436902520,WP_148729134.1-Comamonas-sp.-Z3:0.0382155266)39.2/93:0.0058400225,WP_034391098.1-Comamonas-thiooxydans:0.0142368904)74.4/93:0.0076665937)89/99:0.0254258970)8/89:0.0101040821,WP_218241177.1-Comamonas-fluminis:0.0951140723)99.8/100:0.1231697729,WP_012203208.1-Delftia-acidovorans:0.1611665987)85.1/95:0.0321902307,(WP_066475960.1-Comamonas-terrae:0.1327777193,WP_058879441.1-Comamonas-kerstersii:0.1092951539)62.7/93:0.0241350629)99.4/99:0.1523387144)93.3/99:0.0767662401)20.1/38:0.0448335292)77.5/24:0.0342259726,((WP_167682126.1-Rhodocyclus-gracilis:0.2249449144,(WP_153115862.1-Rhodocyclus-tenuis:0.0374556355,WP_200385811.1-Rhodocyclus-tenuis:0.0192794561)91.9/100:0.0711510479)83.2/100:0.0524239158,WP_201212349.1-Rhodocyclus-purpureus:0.1219113390)99.8/100:0.1292242589)89.4/26:0.0428570251,((((WP_094838100.1-Bordetella-genomosp.-4:0.1529518487,WP_160351730.1-Bordetella-sp.-15P40C-2:0.1829873121)99.6/100:0.1706955798,(WP_083560927.1-Oceanococcus-atlanticus:0.2080417359,WP_104228621.1-Solimonas-fluminis:0.2312442695)92.4/100:0.0927035172)73.5/96:0.0528222952,(WP_046292890.1-Pandoraea-oxalativorans:0.0763552288,WP_150719538.1-Pandoraea-capi:0.0798815723)100/100:0.2714713446)90.6/82:0.0752847726,(((WP_176062241.1-Anaeromyxobacter-diazotrophicus:0.1367383190,WP_248361410.1-Anaeromyxobacter-oryzae:0.1832560166)99.5/99:0.1402396952,(((WP_018882239.1-unclassified-Thioalkalivibrio:0.2757227636,WP_125198734.1-Guyparkeria-sp.-SCN-R1:0.1382873727)89.9/100:0.0649099742,(((WP_026296529.1-Thioalkalivibrio-sp.-ALJ2:0.1149500103,WP_019569965.1-Thioalkalivibrio-sp.-ALE11:0.0693668407)84.9/98:0.0207979322,WP_028484075.1-Thioalkalivibrio-sp.-ALE17:0.0360298445)80.2/97:0.0099506977,WP_018169543.1-Thioalkalivibrio-sp.-ALMg9:0.0260474974)99.1/99:0.0786595004)7.3/28:0.0246140407,WP_250072676.1-Guyparkeria-hydrothermalis:0.1569392176)98.5/100:0.1082821991)60.6/81:0.0414718429,WP_205202385.1-Azonexus-hydrophilus:0.1833624501)16.1/32:0.0208753547)80.1/28:0.0264393595)61.8/23:0.0338583069,(WP_173581083.1-Azonexus-fungiphilus:0.1792147847,WP_153110537.1-Propionivibrio-limicola:0.1883916576)93.8/49:0.0708835102)0/7:0.0000025566,(((((((WP_269022943.1-Methylomonas-sp.-MP1:0.0809736101,(WP_196437371.1-Methylomonas-sp.-LL1:0.0917200016,(WP_150047608.1-Methylomonas-rhizoryzae:0.2162468424,WP_066983474.1-Methylomonas-lenta:0.0692231920)87.2/99:0.0443566401)92.5/99:0.0539575459)91.3/99:0.0537620869,(WP_140911374.1-Methylomonas-koyamae:0.1578106497,((WP_071159776.1-Methylomonas-sp.-LWB:0.1288098813,(WP_020485671.1-Methylomonas-sp.-MK1:0.1405832605,(WP_256607223.1-Methylomonas-sp.-WSC-7:0.0597623337,(WP_205450646.1-Methylomonas-sp.-EFPC1:0.0696618705,WP_064036612.1-Methylomonas-methanica:0.0000025932)69.7/100:0.0074621131)93.3/100:0.0496296509)90.2/98:0.0507706072)30.4/87:0.0049927777,WP_036273427.1-Methylomonas:0.0907636314)87.2/99:0.0360955701)51.9/98:0.0397921869)86.9/98:0.0463136535,(WP_148817865.1-Pigmentiphaga-aceris:0.2272344689,WP_064030819.1-Methylomonas-koyamae:0.1339674883)58/67:0.0396522093)96.3/62:0.0801093456,((((WP_138632053.1-Pseudoalteromonas-spongiae:0.2488970363,WP_168987795.1-Azoarcus-taiwanensis:0.1583126082)0.7/19:0.0390840554,(WP_018865194.1-Thioalkalivibrio-sp.-ARh3:0.1148868902,WP_048706771.1-Thauera-humireducens:0.2484128553)81.9/96:0.0361699219)93.8/45:0.0553181523,((((WP_238617884.1-Ectothiorhodospira-shaposhnikovii:0.1099404402,WP_025282248.1-Ectothiorhodospira-haloalkaliphila:0.0977013158)92.9/100:0.0607516991,WP_238622182.1-unclassified-Ectothiorhodospira:0.0990957350)87.9/100:0.0519927182,(((((((((WP_060482506.1-Pseudomonas-sp.-NBRC-111119:0.1318708715,WP_159409308.1-Pseudomonas-putida:0.0824856968)83.8/54:0.0486108735,((WP_196129978.1-Pseudomonas-fulva:0.0744997893,WP_201164743.1-Pseudomonas-sp.-S60:0.0333741027)98.8/99:0.0667278282,(((WP_174142546.1-Pseudomonas-putida:0.1127674565,WP_027907237.1-Pseudomonas:0.0331533576)44.5/100:0.0131416979,WP_075044077.1-Pseudomonas-putida:0.0387206324)24.4/100:0.0220591445,(WP_023379297.1-Pseudomonas-sp.-VLB120:0.0815265061,WP_202675963.1-Pseudomonas-sp.-SK2:0.0304451750)98.3/100:0.0594554304)94/99:0.0398962524)90.1/99:0.0257329422)77.1/26:0.0096929684,((WP_182365203.1-Pseudomonas-juntendi:0.0605500863,(WP_182365213.1-Pseudomonas:0.0914989676,(((WP_110607173.1-Pseudomonas-fulva:0.0121645642,WP_088523134.1-unclassified-Pseudomonas:0.0417562514)86.8/100:0.0207706790,(WP_201214098.1-Pseudomonas-sp.-S36:0.0123531777,WP_054890059.1-Pseudomonas:0.0593997927)98/100:0.0487478156)85.5/42:0.0158264066,(WP_167339957.1-Pseudomonas-fulva:0.0263614035,WP_186476580.1-Pseudomonas-promysalinigenes:0.0384623021)88.5/98:0.0179268370)66.2/42:0.0119605343)15.8/37:0.0117026657)89.5/63:0.0177494023,(WP_115272924.1-Pseudomonas-putida:0.0987670609,WP_267821885.1-Pseudomonas-putida:0.0396549470)84.7/65:0.0266053329)0/26:0.0000028680)30.5/32:0.0115779169,(WP_167277242.1-Pantoea-sp.-Tr-811:0.0174603174,WP_103449615.1-Pseudomonas-putida:0.0468448275)98.2/100:0.0417098090)90.6/80:0.0217180943,(WP_046489762.1-Pseudomonas-veronii:0.0912488101,(WP_059395284.1-Pseudomonas-putida:0.0680385312,(WP_009685980.1-Pseudomonas:0.0623753969,WP_024086460.1-Pseudomonas:0.0542052580)91.1/100:0.0401187457)86.3/100:0.0357977267)92.5/96:0.0471554846)69/71:0.0119770447,WP_029885231.1-Pseudomonas:0.0259155068)21.1/56:0.0093234398,(WP_230064532.1-Pseudomonas-sp.-CIP-10:0.0178714790,WP_197863301.1-Pseudomonas-putida:0.0353847141)92.1/99:0.0297523837)95.8/72:0.0561220354,WP_060394622.1-Pseudomonas-entomophila:0.0818306426)98.2/82:0.0690791909,WP_183165449.1-Azomonas-macrocytogenes:0.0977392217)78.3/100:0.0285912136)85.6/79:0.0289912873,(WP_256068944.1-Stutzerimonas-stutzeri:0.1134454829,WP_026685886.1-Azovibrio-restrictus:0.1322952068)94.6/96:0.0566542851)81.2/73:0.0195249824)70.8/38:0.0150545850,WP_004074545.1-Desulfobacter-postgatei:0.2253650894)88/47:0.0334127238)90.8/91:0.0398101887,(WP_136386324.1-Pseudothauera-rhizosphaerae:0.1232938466,(WP_114649035.1-Pseudothauera-hydrothermalis:0.0769276362,WP_187718084.1-Thauera-sp.-CAU-1555:0.1135367014)78.1/98:0.0615044853)99.4/100:0.1116956447)88.1/84:0.0291584077,((WP_013164002.1-Desulfurivibrio-alkaliphilus:0.1880613016,WP_263768642.1-Propionivibrio-sp.-SG131:0.1896091869)51.1/69:0.0650939280,(WP_033955816.1-Pseudomonas-aeruginosa:0.0228625422,WP_003153917.1-Pseudomonas-aeruginosa-group:0.0318226583)99.9/100:0.1599114096)85.5/63:0.0254643214)93.8/95:0.0453890196,(((WP_121459656.1-Azonexus-fungiphilus:0.1879249168,((WP_094482496.1-Rhodoferax-sp.-TH121:0.0594194402,WP_105262671.1-Rhodoferax-sp.-TS-BS-61-7:0.0423384631)86.5/100:0.0334848717,WP_214121631.1-Curvibacter-sp.-CHRR-16:0.2024679438)98/100:0.0971063290)14.8/39:0.0127787106,WP_117174481.1-Rhodoferax-lacus:0.1737072061)88.4/70:0.0287460545,(WP_066884254.1-Dechloromonas:0.0873066571,WP_027458229.1-Dechloromonas-agitata:0.0817483559)96.6/100:0.0638409906)98.2/72:0.0741665096)36.8/20:0.0280742523)95.1/41:0.0556585840,(((((((((((WP_250059255.1-Halomonas-sp.-ATCH28:0.2563879940,WP_179153381.1-Chromohalobacter-salexigens:0.1345766705)94.7/99:0.0702539096,WP_250939176.1-unclassified-Larsenimonas:0.1868075042)84.1/98:0.0459080813,(WP_218634960.1-Marinobacter-arenosus:0.0758741638,WP_240716217.1-Halomonas-neptunia:0.0886749080)99.8/100:0.1278402509)81.7/62:0.0255395827,(WP_106478747.1-Phytohalomonas-tamaricis:0.1588981999,(WP_138213821.1-Pseudomonas-nitroreducens:0.0288730167,WP_017517140.1-Pseudomonas-nitroreducens:0.0241415552)100/100:0.1918998198)91.2/95:0.0628959243)96.6/71:0.0743942510,((WP_029241333.1-Pseudomonas:0.0531944658,((WP_054996663.1-Pseudomonas-ficuserectae:0.0338582834,(WP_007252556.1-Pseudomonas-syringae-group:0.0404045790,WP_025390602.1-Pseudomonas-syringae:0.0237260848)37.7/100:0.0188250886)93.6/100:0.0398440378,WP_125830676.1-Pseudomonas-sp.-v388:0.0635865360)66/98:0.0432482097)97/100:0.0743077387,((WP_016973326.1-Pseudomonas-tolaasii:0.0899162499,WP_099548838.1-Pseudomonas-sp.-2995-3:0.0933571037)33.8/98:0.0342225687,(WP_177015116.1-Pseudomonas-costantinii:0.0459902843,WP_141063556.1-Pseudomonas-fluorescens:0.0851640925)93.9/100:0.0519218183)92.8/99:0.0567846957)98.3/100:0.0913645046)92.5/29:0.0482363545,(((WP_099797666.1-Thermostichus-lividus:0.4206374667,WP_085721622.1-Pseudomonas:0.1371479096)53/29:0.0318530099,(((WP_169965430.1-unclassified-Pseudomonas:0.0368131171,WP_047278503.1-Pseudomonas-lundensis:0.0230566363)96.3/100:0.0735649889,WP_205489922.1-Pseudomonas-lactucae:0.1412518047)94.2/85:0.0696556917,(WP_104727848.1-Pseudomonas-oleovorans:0.0724802576,WP_196167318.1-Pseudomonas-monteilii:0.2506624075)73.4/31:0.0153896164)82.7/26:0.0242391753)85.2/34:0.0255090655,(WP_042945119.1-Pseudomonas-fluorescens:0.0603601756,WP_181643258.1-Pseudomonas-sivasensis:0.0426184730)99.7/100:0.1021453094)15.2/48:0.0112285170)87.5/28:0.0173552866,((((((((WP_044615503.1-Gynuella-sunshinyii:0.2999869953,WP_227683402.1-Pusillimonas-sp.-MFBS29:0.1249640144)57.2/98:0.0709417251,WP_092579398.1-Achromobacter-sp.-NFACC18-2:0.3080306242)37.8/93:0.0314827674,((WP_230502293.1-Janthinobacterium-sp.-FW305-128:0.2959639215,WP_249918475.1-Pseudomonas-sp.-R3.Fl:0.1186170934)69.5/97:0.0277941258,((((WP_220354719.1-Rhizobium-cauense:0.2013848458,WP_072373837.1-Rhizobium-tibeticum:0.0611336854)0/24:0.0241536238,WP_007537558.1-Rhizobium-mesoamericanum:0.0562592595)92/56:0.0508894368,WP_016557527.1-Rhizobium-grahamii:0.1312331125)0/55:0.0132910273,(((WP_184450570.1-Rhizobium-sp.-BK538:0.1383530666,WP_153273632.1-Rhizobium-grahamii:0.1762175195)36.7/85:0.0224257918,WP_233136718.1-Agrobacterium-vaccinii:0.1991227915)23.9/85:0.0200027970,WP_018896487.1-Rhizobium-sp.-2MFCol3.1:0.1641939367)58.6/66:0.0308809340)99.9/100:0.1582913491)3.5/25:0.0197038549)79.9/71:0.0375790838,WP_068803504.1-Immundisolibacter-cernigliae:0.1680513932)61.7/54:0.0353612632,(WP_019141909.1-Noviherbaspirillum-massiliense:0.1039575150,WP_146330669.1-Noviherbaspirillum-sp.-UKPF54:0.1758718487)62.6/93:0.0530031006)91.7/94:0.0488913262,(((WP_227390214.1-Halomonas-malpeensis:0.4179647709,WP_058422493.1-Pseudomonas:0.0936201883)78.8/99:0.0480379382,WP_223462047.1-Pseudomonas-sp.-A-R-26:0.1546736494)84.4/99:0.0436714966,(((WP_225237063.1-Massilia-oculi:0.2266152291,WP_147937383.1-Massilia-arenae:0.1155378652)96.5/100:0.1085553615,(WP_192051557.1-unclassified-Massilia:0.0930950146,WP_136222466.1-Massilia-sp.-Mn16-1_5:0.0751434454)84.3/100:0.0567744101)94.6/100:0.0703066265,(WP_057247240.1-Duganella-sp.-Root198D2:0.1621414404,WP_155457065.1-Massilia-eburnea:0.0925680421)95/100:0.0576642307)95/95:0.0554045768)22.1/80:0.0179053176)88.2/66:0.0346671952,((((WP_102075761.1-Pollutimonas-subterranea:0.1186020377,WP_102071501.1-Pollutimonas-nitritireducens:0.1835031891)66.9/81:0.0427686547,(WP_212592286.1-Janthinobacterium-sp.-TND4EL3:0.1674952605,WP_217963608.1-Candidimonas-humi:0.1661849540)94.1/77:0.0575010409)72.8/70:0.0226782383,WP_047550623.1-Methylotenera-sp.-G11:0.1929278199)89.2/73:0.0369987863,(WP_066098770.1-Halothiobacillus-sp.-LS2:0.0938088103,WP_070073009.1-Acidihalobacter-aeolianus:0.1779564462)96.5/99:0.0856462828)90/73:0.0288608096)94.1/84:0.0348015029,(((((((WP_191471912.1-Pseudomonas-asiatica:0.1639238599,(WP_043318452.1-Pseudomonas-sp.-PI1:0.0432007129,WP_043268312.1-Pseudomonas:0.0264555539)83.4/100:0.0208993040)96.4/100:0.0696473748,WP_089390899.1-Pseudomonas-delhiensis:0.0826669805)76.6/98:0.0354396814,WP_076426141.1-Pseudomonas-alcaligenes:0.0948373151)92/99:0.0486444622,WP_214501190.1-Pseudomonas-sp.-RC3H12:0.2111833465)0/22:0.0165911373,WP_233297479.1-Pseudomonas:0.1317348745)95.8/96:0.0581476216,((((WP_020564242.1-Methylosarcina-fibrata:0.1371960500,WP_074973890.1-Pseudomonas-otitidis:0.1597510154)91.6/76:0.0533581204,(WP_101152997.1-Pseudomonas-sp.-Choline-3u-10:0.0990777155,((WP_046617631.1-Stutzerimonas-stutzeri:0.0542823370,WP_135265705.1-Stutzerimonas-stutzeri:0.0493494717)92.5/100:0.0504154157,WP_003299494.1-Stutzerimonas-stutzeri:0.0792294406)73.9/100:0.0253222558)91.7/96:0.0427608942)42.7/60:0.0266547418,((WP_180152801.1-Parapusillimonas-granuli:0.1215373720,(WP_088146851.1-Achromobacter-denitrificans:0.0819540959,WP_105555808.1-Achromobacter-sp.-MYb9:0.0714863294)98.4/100:0.0646221119)29.9/91:0.0245273323,(WP_011111299.1-Nitrosomonas-europaea:0.1320640347,WP_046857648.1-Pseudomonas-sp.-CCOS-191:0.1408054162)67.9/91:0.0313118861)95.5/89:0.0572058755)53.6/62:0.0191963526,(WP_125865093.1-Stutzerimonas-stutzeri:0.1199181786,WP_200290630.1-Stutzerimonas-balearica:0.1037282110)92.6/93:0.0631773900)72.9/40:0.0147431604)74.7/58:0.0065182972,((((WP_209872004.1-Pseudomonas-sp.-BP8:0.1272502792,WP_066093302.1-Hydrogenophaga-crassostreae:0.1427673960)77.5/84:0.0302960716,(WP_094844257.1-Bordetella-genomosp.-11:0.1513374601,WP_166067558.1-Diaphorobacter-sp.-HDW4B:0.1347186639)95/97:0.0644676576)76.1/75:0.0238943577,((WP_177138191.1-Hydrogenophaga-aromaticivorans:0.1046215575,WP_133157403.1-Hydrogenophaga-pseudoflava:0.1249394899)90.3/97:0.0428373096,((WP_236964528.1-Hydrogenophaga-sp.-SL48:0.0920402066,WP_226493830.1-Hydrogenophaga-taeniospiralis:0.0410280351)17.1/80:0.0161872328,WP_210824367.1-Hydrogenophaga-aromaticivorans:0.0401556916)95.8/100:0.0565105110)88.4/97:0.0354686272)83.6/88:0.0356674848,(WP_065086507.1-Pseudomonas:0.1729941789,(WP_184594840.1-Pseudomonas-nitritireducens:0.1256025189,WP_192381882.1-Pseudomonas-sp.-PDM20:0.1079828276)97.8/100:0.0957795981)89.8/96:0.0458809624)89.6/83:0.0310184374)88/84:0.0250284675)0/38:0.0000020368)91.8/35:0.0301164421,(WP_076518421.1-Achromobacter-sp.-MFA1-R4:0.0569111107,(WP_104142565.1-Achromobacter-spanius:0.0200999420,(WP_120157261.1-Achromobacter-sp.-B7:0.0175230685,WP_158658301.1-Achromobacter-sp.-AONIH1:0.2891691315)82.6/44:0.0308030239)96.1/44:0.0502537295)99.8/98:0.1102652615)77.9/29:0.0121126337,((WP_058488093.1-Pseudomonas:0.1183039693,WP_043256200.1-Pseudomonas-knackmussii:0.1021087599)91.7/100:0.0466235408,((WP_088420173.1-Pseudomonas-nitroreducens:0.1028300574,WP_254475170.1-unclassified-Pseudomonas:0.0599702819)31.9/100:0.0207188735,WP_205338011.1-Pseudomonas-sp.-PDNC002:0.0595128154)99.4/100:0.1042888691)93.1/98:0.0476930199)85.9/30:0.0241146638,(WP_077733980.1-Paenalcaligenes-hominis:0.0380874961,WP_167661700.1-Paenalcaligenes-hominis:0.0266629754)100/100:0.2090727883)88.1/29:0.0240098772,((((WP_014706296.1-Methylophaga-nitratireducenticrescens:0.0006869231,WP_136552334.1-Methylophaga-sp.-SB9B:0.0650503141)100/100:0.2943334596,((WP_148717329.1-Chitinolyticbacter-meiyuanensis:0.3216749515,WP_123210144.1-Stagnimonas-aquatica:0.3653895645)95.7/100:0.1370955298,WP_204103763.1-Spirulina-major:0.2262912789)88.1/89:0.0559408549)36.6/53:0.0185781265,WP_216129682.1-Azospira-inquinata:0.2290299325)63.5/71:0.0580398592,WP_258826205.1-Massilia-agri:0.2563546153)86.8/73:0.0379803071)94.3/35:0.0393411657)83.5/79:0.0185564670,((WP_212718705.1-unclassified-Synechocystis:0.0000022910,(WP_228015966.1-Synechocystis-salina:0.0875495395,WP_010873214.1-unclassified-Synechocystis:0.0605634191)86.6/99:0.0251728541)100/100:0.3482828116,((((((WP_019918777.1-Methyloversatilis-discipulorum:0.0428161249,WP_018229738.1-Methyloversatilis-universalis:0.0447161061)99.5/100:0.0853036314,((WP_101941536.1-Uliginosibacterium-sp.-TH139:0.0877360460,WP_095523004.1-Candidatus-Dactylopiibacterium-carminicum:0.0812404838)97.7/100:0.1150704956,WP_183409942.1-Litorivivens-lipolytica:0.3137106262)51.7/98:0.0354670093)97.2/100:0.0823906498,((WP_047193087.1-Schlegelella-brevitalea:0.2081469211,WP_019561187.1-Caldimonas-manganoxidans:0.1568959769)93.1/100:0.0898974782,WP_074865046.1-Atopomonas-hussainii:0.1502631548)95.7/100:0.0776915627)95.8/95:0.0528211229,WP_201673630.1-Ramlibacter-monticola:0.1950677312)7.4/76:0.0124257400,(WP_109041087.1-Azospira-sp.-I13:0.0970187514,WP_014237023.1-Azospira-oryzae:0.1068055806)97.8/100:0.0774230485)95/80:0.0482809902,(((((WP_129151777.1-Achromobacter-aloeverae:0.0821974961,WP_088599355.1-unclassified-Achromobacter:0.0609566161)98.9/100:0.1019285461,WP_057654143.1-Bordetella-sp.-N:0.1341115525)83.7/99:0.0518526816,WP_086067427.1-Bordetella:0.1666977321)91/100:0.0484970145,(WP_066641528.1-Bordetella-sp.-H567:0.0632440157,(WP_086058993.1-Bordetella-genomosp.-9:0.1453421890,WP_094856450.1-Bordetella-genomosp.-10:0.1272889120)96.5/100:0.0821343818)63/100:0.0585424498)98.6/100:0.0912018360,(WP_091467897.1-Methylophilus-rhizosphaerae:0.2238331642,WP_067106371.1-Methylobacillus-sp.-MM3:0.1604338840)63.8/90:0.0366358028)76.7/95:0.0111423315)81.2/89:0.0213830421)93.1/68:0.0435869155)89.9/59:0.0252455628,WP_054772520.1-Methylogaea-oryzae:0.1690404276)76/23:0.0105659262,((((((((WP_072282603.1-Syntrophotalea-acetylenivorans:0.1008445313,(WP_092056970.1-Desulfuromonas-acetexigens:0.1128379152,(WP_043770486.1-Desulfobulbus-propionicus:0.1660574107,WP_148894253.1-Geothermobacter-ehrlichii:0.2567647922)47.8/74:0.0566252466)94.5/74:0.0600273020)75.2/74:0.0261743374,(WP_028317018.1-Desulfobulbus-elongatus:0.2316773691,WP_155876508.1-Desulfuromonas-sp.-AOP6:0.1700248786)91.4/98:0.0767159869)95.4/100:0.0806329440,WP_072282412.1-Syntrophotalea-acetylenivorans:0.1495412615)90.6/99:0.0575564917,(((((((WP_089129802.1-Tolypothrix-sp.-NIES-4075:0.0668309310,WP_096563566.1-Scytonema-sp.-NIES-4073:0.0981603640)63.4/99:0.0319921391,(WP_048866388.1-Scytonema-tolypothrichoides:0.0771307915,WP_096563464.1-Scytonema-sp.-NIES-4073:0.0969222167)0/5:0.0149914739)99.9/100:0.1397210118,(((((((((WP_009344618.1-Cylindrospermopsis-raciborskii:0.0619321646,WP_240868223.1-Cylindrospermopsis-raciborskii:0.0225677883)99.4/100:0.0895242338,(WP_096570274.1-Sphaerospermopsis:0.0386038671,WP_220611028.1-Sphaerospermopsis-torques-reginae:0.0531189819)47.5/100:0.0274342531)98.7/100:0.0685952165,WP_190559626.1-Anabaena-sphaerica:0.0625957392)81.3/87:0.0308074029,WP_237989437.1-Nostoc-sp.-UHCC-0870:0.0679893642)88.6/87:0.0250610803,WP_206263219.1-Nostoc-sp.-UHCC-0702:0.0611421353)62.6/98:0.0291845146,WP_096557790.1-Nostoc-sp.-NIES-4103:0.1312491932)78/99:0.0378441278,WP_016866333.1-Fischerella:0.0645826490)89.4/99:0.0304100651,((WP_015128325.1-Calothrix-sp.-PCC-7507:0.1047116727,WP_017653186.1-Fortiea-contorta:0.1354630201)63.1/100:0.0379846718,((WP_193199005.1-Nostoc-sp.-MG11:0.0425924603,WP_162396505.1-Nostoc-sp.-B-2019:0.0390383494)91.5/100:0.0495020571,WP_214439445.1-Atlanticothrix-silvestris:0.1213508560)90.7/99:0.0629558686)75.5/97:0.0235564073)39.5/83:0.0129945527,(((WP_045870014.1-unclassified-Tolypothrix:0.0243875672,WP_190656031.1-Nostoc-linckia:0.0318207548)99.2/100:0.0661106448,(WP_190743070.1-Tolypothrix-sp.-FACHB-123:0.0417862914,WP_190545010.1-Calothrix:0.0313569281)98.2/100:0.0607029034)70.9/100:0.0313060592,WP_086758595.1-Nostoc-sp.-106C:0.0557419759)19.7/84:0.0325793511)90.2/97:0.0462681999)97.5/100:0.0859316323,(WP_054689510.1-Desulfosarcina-cetonica:0.1995998279,WP_009207329.1-Sulfuricella-denitrificans:0.1935108041)51.2/14:0.0446247418)0/6:0.0140686512,((((WP_239190817.1-Candidatus-Nitrotoga-sp.-HW29:0.0524545368,WP_239287644.1-Candidatus-Nitrotoga-sp.-1052:0.0136525209)85.9/100:0.0146873110,WP_239235002.1-Candidatus-Nitrotoga-sp.-BS:0.0460688603)59.8/98:0.0041234281,WP_239797721.1-Candidatus-Nitrotoga-arctica:0.0186907854)99.9/100:0.1641286843,(WP_044881819.1-Neochlamydia-sp.-EPS4:0.0386658223,WP_166155173.1-unclassified-Neochlamydia:0.0136435793)100/100:0.2687582841)92.1/15:0.0478975642)88.6/80:0.0359586982,((((((((((((((WP_102179481.1-Fischerella-thermalis:0.0238353240,((WP_096675774.1-Fischerella-sp.-NIES-4106:0.0424101878,WP_053458209.1-Hapalosiphon-sp.-MRB220:0.0285754325)71.3/90:0.0053198741,(WP_131119525.1-Westiellopsis-prolifica:0.0412507073,WP_016860528.1-Fischerella-muscicola:0.0223022758)85.2/100:0.0119032384)98.4/96:0.0629399800)100/100:0.1501426591,(WP_190825375.1-Iningainema-tapete:0.1783496691,(WP_015202791.1-Crinalium-epipsammum:0.1682558135,WP_035154029.1-Calothrix-sp.-336/3:0.2537360747)89.5/97:0.0575578675)50.2/88:0.0314349838)83.4/92:0.0222062873,((WP_099102643.1-Nostoc-sp.-Peltigera-malacea-cyanobiont-DB3992:0.0639703481,((WP_190950926.1-Nostoc-punctiforme:0.0409982306,(WP_221704991.1-Desmonostoc-muscorum:0.0411989757,WP_096537650.1-Nostoc-linckia:0.0151859113)73.7/99:0.0067043870)2/62:0.0203727922,WP_190959861.1-Nostoc:0.0183604064)97.8/95:0.0764071174)99.6/100:0.1013159360,((((((WP_193938462.1-Cuspidothrix-issatschenkoi:0.1332720258,((WP_096666215.1-Dolichospermum-compactum:0.0509557410,(WP_190387835.1-Aphanizomenon:0.0535381174,WP_168466156.1-Aphanizomenon:0.0164265624)51.6/98:0.0163384157)83/90:0.0136331050,WP_015080416.1-Anabaena-sp.-90:0.0258897134)86.4/94:0.0121185611)86/85:0.0183304069,(((WP_190370804.1-Anabaena-sp.-FACHB-1237:0.1136741948,(WP_190392428.1-Dolichospermum-sp.-FACHB-1091:0.0136498241,(WP_028089231.1-Dolichospermum-circinale:0.0170289770,WP_137908967.1-Nostocales:0.0408003555)98.2/100:0.0484112418)98.5/100:0.0756661677)28.5/82:0.0237308926,WP_168537172.1-Anabaena-sp.-UHCC-0253:0.0601247930)84.4/65:0.0140472886,WP_168502193.1-Anabaena-sp.-UHCC-0187:0.0065049808)70.7/68:0.0046338005)99.9/100:0.1092153443,(WP_010998622.1-Nostocaceae:0.0516032724,WP_011320078.1-Nostocaceae:0.0501902904)83.7/100:0.0193055332)94.1/94:0.0409849123,((WP_015138800.1-Nostoc-sp.-PCC-7524:0.0642303217,WP_066425233.1-Anabaena-sp.-4-3:0.0770798979)97.1/100:0.0572799371,(WP_199337214.1-Nostoc-sp.-FACHB-152:0.0969079601,(WP_239729951.1-Nodularia-sphaerocarpa:0.0578808570,WP_173403323.1-Nodularia-spumigena:0.0418050038)95/99:0.0405453485)89.4/95:0.0200323313)73.1/87:0.0067856733)81.4/89:0.0151522772,WP_199337786.1-Nostoc-sp.-FACHB-280:0.1073776941)96.7/100:0.0576920774,(WP_096624129.1-Calothrix-sp.-NIES-3974:0.0640586689,WP_095721999.1-Calothrix-elsteri:0.0798783613)99.2/100:0.0775344754)65.3/100:0.0228645089)94.3/100:0.0474630584)90.8/100:0.0357982761,(WP_224410051.1-Oscillatoria-salina:0.1516635899,WP_250123015.1-Chroococcidiopsis-sp.-CCMEE-29:0.1250731722)51/95:0.0468502414)66.4/92:0.0282555781,WP_073549891.1-Chroogloeocystis-siderophila:0.1447528583)7/29:0.0290945581,(((WP_233747769.1-Leptodesmis-sichuanensis:0.1681988085,WP_223049852.1-Leptothermofonsia-sichuanensis:0.0844746658)78.1/43:0.0152647444,WP_224345144.1-Kovacikia-minuta:0.1007340231)37.7/41:0.0161336783,WP_192216758.1-Microcoleus-sp.-FACHB-SPT15:0.1347715020)91.3/73:0.0252795918)41.1/71:0.0145381839,((WP_008178350.1-Moorena-producens:0.1740501776,WP_193873404.1-unclassified-Coleofasciculus:0.1064851450)73/76:0.0121666247,((WP_190666400.1-Coleofasciculus-sp.-FACHB-501:0.1029910216,WP_190525322.1-Coleofasciculus-sp.-FACHB-T130:0.0475628064)96.8/100:0.0561582364,WP_254010234.1-Symplocastrum-sp.-BBK-W-15:0.1219916056)79.4/87:0.0219394904)25.5/54:0.0142321814)91.8/89:0.0293388249,((((WP_015170385.1-Geitlerinema-sp.-PCC-7407:0.1676561685,(WP_194029353.1-Lusitaniella-coriacea:0.1450774599,WP_017324474.1-Synechococcus-sp.-PCC-7336:0.2281131537)93.6/99:0.0565659192)94.4/98:0.0549766617,((WP_069965800.1-unclassified-Desertifilum:0.1921061860,(WP_073595522.1-Phormidium-ambiguum:0.0650871876,WP_193982193.1-Phormidium-sp.-LEGE-05292:0.0526628388)95.3/100:0.0551669097)89.6/92:0.0435393910,((WP_226578226.1-Microseira-wollei:0.1566759004,WP_249064905.1-Argonema-antarcticum:0.0484937034)50.7/82:0.0170696729,WP_190468950.1-unclassified-Planktothrix:0.0751919300)83.3/82:0.0165500228)72.9/92:0.0247897426)88/75:0.0266162558,WP_007355666.1-Kamptonema:0.1146949081)69.8/66:0.0106212176,WP_190493486.1-unclassified-Microcoleus:0.1945287914)86.1/75:0.0211195535)85.2/69:0.0239025178,(((WP_261197618.1-unclassified-Laspinema:0.0429641663,WP_207087219.1-Phormidium-pseudopriestleyi:0.0334285672)0/16:0.0137955226,WP_261204505.1-Laspinema-sp.-D2d:0.0272712890)100/100:0.0996961471,(WP_017715496.1-Oscillatoria-sp.-PCC-10802:0.1399724674,(WP_190665495.1-Microcoleus-sp.-FACHB-672:0.0619931452,WP_261220767.1-Ancylothrix-sp.-D3o:0.1428909470)92.3/100:0.0471029251)66.7/96:0.0389188784)27.3/77:0.0285298432)81/56:0.0167754707,(((((((WP_204137747.1-Halomicronema-sp.-CCY15110:0.0405799410,WP_039724568.1-Leptolyngbya-sp.-LK:0.0659452910)99.3/100:0.1352060315,WP_023076112.1-Leptolyngbya-sp.-Heron-Island-J:0.1660133928)86.5/100:0.1070860739,WP_190772446.1-unclassified-Leptolyngbya:0.1425370651)94.3/92:0.0775985267,((((WP_017661508.1-Baaleninema-simplex:0.2027574406,WP_008052766.1-Limnospira-indica:0.1545198080)91.4/86:0.0557600500,(((WP_202221768.1-Okeania-sp.-KiyG1:0.0358000054,WP_094672483.1-Hydrocoleum-sp.-CS-953:0.0372950011)33.9/90:0.0129815637,WP_124146462.1-Okeania-hirsuta:0.0086745923)99.9/100:0.1332662484,WP_071516849.1-Geitlerinema-sp.-PCC-9228:0.1638971572)82.8/93:0.0447142747)88/90:0.0441112590,(WP_009783380.1-Lyngbya:0.0723126693,WP_046277942.1-Limnoraphis-robusta:0.0406711162)92/100:0.0418457031)93.8/90:0.0537782156,((WP_083623900.1-Planktothrix-serta:0.0329315540,WP_072717475.1-Planktothrix:0.0247133849)87.8/100:0.0175800369,(WP_079676747.1-Planktothrix-sp.-PCC-11201:0.0215963309,(WP_026797127.1-Planktothrix:0.0230891923,WP_214559879.1-Planktothrix-agardhii:0.0351783292)77.9/100:0.0238953683)98.2/100:0.0580718411)96.2/100:0.0550948150)29.2/75:0.0266700064)57.5/49:0.0060442365,WP_200665545.1-Oxynema-sp.-CENA135:0.1750227516)69.2/50:0.0172350683,WP_190877489.1-unclassified-Planktothricoides:0.1425822789)88.8/85:0.0288592420,(WP_190531554.1-Coleofasciculus-sp.-FACHB-542:0.1527452445,((((WP_236507253.1-Tychonema-sp.-BBK16:0.0351634423,WP_096828456.1-Tychonema-bourrellyi:0.0569357092)93.5/100:0.0306579716,WP_242724542.1-Microcoleus-vaginatus:0.0293876512)83.2/92:0.0109194458,WP_015174083.1-Oscillatoria-nigro-viridis:0.0170648930)72.5/92:0.0047779068,((WP_194009709.1-unclassified-Tychonema:0.0322516433,WP_265235823.1-Lyngbya-sp.-CCAP-1446/10:0.0366360852)88.8/100:0.0173120660,WP_193973976.1-Microcoleus-sp.-LEGE-07076:0.0433919632)79/92:0.0155969993)100/100:0.0898198246)19.1/29:0.0279663227)0/8:0.0000021740)93/95:0.0423359951,(WP_015193936.1-Stanieria-cyanosphaera:0.0241152159,WP_096384422.1-Stanieria-sp.-NIES-3757:0.0725758467)100/100:0.1552445057)86.2/89:0.0351010035,(((((((WP_106232846.1-Pleurocapsa-sp.-CCALA-161:0.1316282092,WP_229639079.1-Waterburya-agarophytonicola:0.0267393555)75.7/100:0.0313612738,WP_096721927.1-Chondrocystis-sp.-NIES-4102:0.0692901728)99.6/100:0.1117861731,WP_006510035.1-Xenococcus-sp.-PCC-7305:0.1177828317)93.7/100:0.0616970298,WP_036478304.1-Myxosarcina-sp.-GI1:0.1291676782)88.8/95:0.0365725838,WP_012595695.1-Rippkaea-orientalis:0.1360214999)76.9/53:0.0265578104,(WP_264322079.1-Zarconia-navalis:0.1733253763,(((WP_007308862.1-Crocosphaera-watsonii:0.0710843393,WP_107668138.1-Cyanothece-sp.-BG0011:0.0284590153)86.6/91:0.0117441289,WP_009545547.1-Crocosphaera-subtropica:0.0175476002)84.3/89:0.0112937982,WP_008278255.1-Crocosphaera-chwakensis:0.0400869282)99.8/100:0.0928746186)25.7/21:0.0264224511)84.3/91:0.0273063073,((WP_006529211.1-Gloeocapsa-sp.-PCC-73106:0.2180220334,WP_124977990.1-Aphanothece-sacrum:0.1137891584)94.3/79:0.0592313159,(WP_106909014.1-Microcystis-sp.-MC19:0.1939878610,WP_015143663.1-Pleurocapsa-sp.-PCC-7327:0.1444451868)26.3/75:0.0214614387)77.8/72:0.0136269315)85/84:0.0247658477)81.9/98:0.0314539276,WP_013324143.1-Gloeothece-verrucosa:0.0763378853)86.7/100:0.0304310296,WP_066347484.1-Geminocystis-sp.-NIES-3708:0.3357862886)32.9/13:0.0277381793)54.2/9:0.0105570713,(WP_237889684.1-Dissulfurimicrobium-hydrothermale:0.2294944284,(WP_250680171.1-Geomonas-sp.-Red32:0.1416711760,WP_199381999.1-Geomesophilobacter-sediminis:0.1592240846)51.3/95:0.0528146032)48/11:0.0392115634)90.7/67:0.0408275770)94.7/66:0.0505309742,((((WP_205222281.1-Desulfogranum-marinum:0.1593084937,(WP_205242362.1-Desulfobulbus-alkaliphilus:0.1204624530,WP_267928740.1-Desulfolithobacter-dissulfuricans:0.1323718107)86.4/100:0.0483253379)69.5/98:0.0401800340,WP_028582690.1-Desulfogranum-japonicum:0.2032916263)84.9/98:0.0418108775,WP_205218910.1-Desulforhopalus-vacuolatus:0.2540117133)90/43:0.0482859204,((WP_137423090.1-Desulfoglaeba-alkanexedens:0.2558491923,WP_027714505.1-Desulfuromonas-sp.-TF:0.1789248744)68.1/47:0.0536142578,((((WP_163339734.1-Desulfopila-sp.-IMCC35008:0.2149635782,WP_136805251.1-Desulfosediminicola-flagellatus:0.1785730466)79.8/91:0.0236584354,WP_073615262.1-Desulfopila-aestuarii:0.1759692840)92.3/100:0.0491554469,WP_103114677.1-Geothermobacter-hydrogeniphilus:0.2855132737)70.4/26:0.0417297940,(WP_228856481.1-Desulfomarina-profundi:0.2457103462,WP_183349429.1-Desulfoprunum-benzoelyticum:0.1366780997)82.1/99:0.0440209376)60.3/27:0.0387583630)87.1/43:0.0293852504)82.9/24:0.0236018722)65.5/23:0.0315271949,(WP_015405619.1-Desulfocapsa-sulfexigens:0.2083095561,(WP_040200185.1-Geoalkalibacter-subterraneus:0.1655334122,WP_053550565.1-Desulfuromonas-soudanensis:0.1711134935)55.2/76:0.0501423181)77.8/18:0.0197070874)64.8/70:0.0182900449,WP_221248821.1-Desulfuromonas-versatilis:0.2452809946)92.4/94:0.0379663098,(((((((WP_238511579.1-Cupriavidus-gilardii:0.2080381787,WP_224042884.1-Cupriavidus-respiraculi:0.2113439278)65.2/89:0.0405845029,(WP_213445339.1-Variovorax-sp.-PCZ-1:0.2778999105,(WP_116600700.1-Cupriavidus-alkaliphilus:0.1229997667,(WP_018008101.1-Cupriavidus-neocaledonicus:0.0412745684,WP_183298446.1-Cupriavidus-alkaliphilus:0.0453264096)99.5/100:0.1115481794)9.3/91:0.0190733267)34.2/75:0.0433031122)95.2/91:0.0517453902,((((WP_183165579.1-Azomonas-macrocytogenes:0.1357558833,((((WP_102836461.1-Stutzerimonas-stutzeri:0.0234230620,(WP_256071125.1-Stutzerimonas-stutzeri:0.0296075316,WP_045423249.1-Pseudomonadaceae:0.0237155517)86.2/100:0.0117784990)97.2/100:0.0421698002,(WP_255961233.1-Stutzerimonas-stutzeri:0.0237357408,WP_025243649.1-Stutzerimonas-stutzeri:0.0413280282)0/29:0.0012937478)96/100:0.0447028820,((WP_037024448.1-Pseudomonas-saudiphocaensis:0.0595379116,WP_181084609.1-Stutzerimonas-stutzeri:0.0058450433)94.3/100:0.0259443124,(WP_011911332.1-Stutzerimonas-stutzeri:0.0338296683,(WP_230358311.1-Pseudomonas-sp.-NP21570:0.0210896435,((WP_193680295.1-Pseudomonas-lopnurensis:0.0312262249,WP_200589398.1-Stutzerimonas-stutzeri:0.0277162554)89.3/98:0.0208433932,((WP_237255234.1-Pseudomonas-oligotrophica:0.0590099134,WP_043295157.1-Stutzerimonas-degradans:0.0146812181)97.4/100:0.0352571789,WP_120997453.1-_Pseudomonas:0.0280052302)86.1/20:0.0161157895)74.4/19:0.0095406388)91.1/31:0.0240718610)85.4/92:0.0174028237)74.8/94:0.0121031974)0/16:0.0000029185,WP_122077451.1-Pseudomonadaceae:0.0314399351)86.3/90:0.0250945042)56.5/69:0.0286539002,WP_028238613.1-Stutzerimonas-azotifigens:0.0745226594)98.8/99:0.0667453133,(WP_228215938.1-Aromatoleum-toluclasticum:0.1613486052,WP_226921657.1-Kinneretia-sp.-XES5:0.2096800795)88.1/98:0.0343857663)47.9/60:0.0131712160,((WP_115680924.1-Cupriavidus-taiwanensis:0.1769278495,WP_018313502.1-Cupriavidus-sp.-UYPR2.512:0.1965835739)68.1/95:0.0276873765,WP_135702508.1-Cupriavidus-oxalaticus:0.1162256606)95.4/83:0.0492467865)93.6/80:0.0385309746)79.8/23:0.0337389767,((((((((((((((((WP_209862330.1-Pseudomonas-sp.-BP8:0.3059038136,WP_225786622.1-Pseudomonas:0.0246690480)94.3/99:0.0541479550,(((((WP_054891693.1-Pseudomonas:0.0619749602,WP_236238170.1-Pseudomonas-faucium:0.1138047522)44.3/96:0.0257407605,WP_158461286.1-Pseudomonas-fluorescens:0.1731315166)23.4/94:0.0282291547,(WP_200141208.1-unclassified-Pseudomonas:0.1494946180,WP_029612831.1-Pseudomonas-parafulva:0.0221143699)94.1/100:0.0552339119)83.6/99:0.0415463771,WP_110605176.1-Pseudomonas-soli:0.1887613390)0/15:0.0251351980,(((WP_217887155.1-Pseudomonas-xanthosomae:0.0356381144,WP_217840807.1-Pseudomonas-fakonensis:0.0316318161)87.2/100:0.0290358022,WP_186654015.1-Pseudomonas-xantholysinigenes:0.1558431990)69.9/93:0.0277155667,(WP_201176141.1-Pseudomonas-sp.-S31:0.0343589303,WP_261240966.1-unclassified-Pseudomonas:0.0320172069)99.4/100:0.0730440482)86.5/85:0.0254357732)76.8/31:0.0151337637)97.7/100:0.0712707417,((WP_181290220.1-Pseudomonas-brassicacearum:0.1236573752,((((WP_201004194.1-Pseudomonas-cichorii:0.0704751399,WP_176569640.1-Pseudomonas-eucalypticola:0.1039578374)88.9/63:0.0395227389,(WP_219063768.1-Pseudomonas-sp.-UMAB-08:0.1602623320,((WP_222540780.1-Pseudomonas-sp.-DR208:0.0966702265,(WP_253509974.1-Pseudomonas-rhodesiae:0.0892383910,WP_093202821.1-Pseudomonas-yamanorum:0.0273658827)91.3/100:0.0339043150)23/76:0.0143626545,((WP_065875813.1-Pseudomonas:0.0368919990,(WP_033898696.1-Pseudomonas-fluorescens-group:0.0296559202,(WP_186687394.1-Pseudomonas-sp.-SWRI18:0.0465786572,WP_169378974.1-Pseudomonas:0.0139325052)77.9/100:0.0223100450)81.6/100:0.0115057906)99.5/100:0.0630295761,WP_105697656.1-Pseudomonas-poae:0.0355179649)74.8/76:0.0093305622)84.1/100:0.0369739143)85.6/98:0.0385127551)87.7/63:0.0250928938,(WP_249678359.1-Pseudomonas-sp.-PIA16:0.1160246804,WP_043192328.1-Pseudomonas-rhizosphaerae:0.1627781837)79.9/48:0.0389852510)71.7/40:0.0106141758,WP_268799018.1-Pseudomonas-sp.-BSw22131:0.1385454934)90.5/90:0.0396592327)91.6/98:0.0413930060,WP_108096457.1-Pseudomonas-sp.-GV071:0.0864977637)76.3/100:0.0419892756)89.8/99:0.0384163795,WP_252273535.1-Pseudomonas-sp.-YIM-B01952:0.0961989232)87.4/99:0.0209121597,WP_143488038.1-Pseudomonas-mangiferae:0.1166975426)53.4/98:0.0146297059,((((WP_090448460.1-Pseudomonas-benzenivorans:0.1546545885,((WP_025164100.1-Pseudomonas-taeanensis:0.0485265497,WP_147170838.1-Pseudomonas-sp.-SJZ079:0.0057246115)98.2/100:0.0456634628,((WP_208707982.1-Pseudomonas-sp.-LPB0260:0.0264455212,WP_255840290.1-Pseudomonas-benzenivorans:0.0312928299)93.3/100:0.0369022749,(WP_233685494.1-Pseudomonas-anguilliseptica:0.0532096215,WP_257778987.1-Pseudomonas-sp.-32.2.56:0.0000026624)100/100:0.1058374610)64.1/43:0.0191055373)74.2/41:0.0067186412)87.9/83:0.0219280605,(WP_238040454.1-Pseudomonas-sp.-MMS21-TM103:0.1034164666,WP_069518708.1-Pseudomonas:0.0917703667)85.4/95:0.0233566255)77.1/79:0.0171116855,(WP_131198269.1-Pseudomonas-dryadis:0.0848662961,(WP_131178052.1-Pseudomonas-daroniae:0.0515048158,(WP_084307530.1-Pseudomonas-flavescens:0.0578629056,(WP_249909557.1-Pseudomonas-fulva:0.0377584580,(WP_074889242.1-Pseudomonas-argentinensis:0.0242549551,WP_027910132.1-Pseudomonas-sp.-URMO17WK12_I4:0.0387521767)66.2/99:0.0126056896)85.3/100:0.0282506468)80.7/98:0.0227044306)95.8/100:0.0545077680)97.5/98:0.0682670161)6.5/49:0.0232154216,((((((((WP_119694090.1-Pseudomonas:0.0417742171,WP_106738315.1-Pseudomonas-mendocina:0.0446485038)91/100:0.0296026487,WP_150606948.1-Pseudomonas-oleovorans:0.0486763753)95.5/98:0.0339230569,WP_024309956.1-Pseudomonas-sp.-P818:0.0333541027)83.2/90:0.0134134125,WP_090337209.1-Pseudomonas-chengduensis:0.0195867434)68.7/76:0.0053564675,WP_074939507.1-Pseudomonas-composti:0.0283542316)28.6/76:0.0131567471,WP_143497094.1-Pseudomonas-sp.-ALS1131:0.0242945305)86.2/100:0.0160309413,WP_021444205.1-Pseudomonas-sp.-EGD-AK9:0.0460206741)85.7/100:0.0195310077,WP_159970704.1-Pseudomonas-sp.-8Z:0.0630862342)91.7/100:0.0445343232)91.6/99:0.0395823773)90.5/85:0.0419448177,(WP_192399936.1-Pseudomonas-sp.-PDM15:0.1226587113,WP_226282979.1-Pseudomonas-lalkuanensis:0.0522499714)97.7/100:0.0715338807)90.5/70:0.0268093896,WP_028626762.1-Pseudomonas-resinovorans:0.0393540834)41/55:0.0059761524,(WP_120654068.1-Pseudomonas-sp.-DY-1:0.0740935973,WP_151138079.1-Pseudomonas-lalkuanensis:0.0366408778)80/87:0.0135021921)81.5/60:0.0182398336,((WP_058067682.1-Pseudomonas-sp.-TTU2014-080ASC:0.1279496913,WP_137821430.1-Pseudomonas-sp.-D-2018:0.0654125151)0/18:0.0207854387,WP_077525673.1-Pseudomonas-resinovorans:0.0997734141)95.5/66:0.0542754924)76.7/50:0.0078193410,(WP_016495240.1-Pseudomonas-resinovorans:0.0444798538,WP_044873077.1-Pseudomonas-sp.-LFM046:0.0562810781)75.4/81:0.0206261687)90.7/79:0.0347236495,WP_003453405.1-Pseudomonas:0.0769843946)84.6/86:0.0341999565,WP_025281494.1-Ectothiorhodospira-haloalkaliphila:0.3041408485)89.4/93:0.0333451372,(WP_220803742.1-Pseudomonas-sp.-NCCP-436:0.1274284413,WP_090225506.1-Pseudomonas-kuykendallii:0.0744994818)89.6/99:0.0349705953)74.6/89:0.0305117895,((WP_002810067.1-Nitrosococcus-oceani:0.0551410905,WP_013221233.1-Nitrosococcus-watsonii:0.0448336577)100/100:0.1825364095,(WP_040097704.1-Geoalkalibacter-ferrihydriticus:0.1905631269,WP_066727622.1-Desulfuromonas-sp.-DDH964:0.1917211981)88.6/97:0.0416452431)89.7/95:0.0413093170)42.8/50:0.0171595554,(((WP_116346290.1-Cupriavidus-taiwanensis:0.0864688850,WP_124685047.1-Cupriavidus-pauculus:0.1905953866)100/100:0.2514040545,WP_226440716.1-Quatrionicoccus-australiensis:0.1992484915)0/71:0.0181170309,((WP_220807781.1-Noviherbaspirillum-aridicola:0.2651426174,WP_119020982.1-Halomonas-sp.-JS92-SW72:0.2710270805)90.1/97:0.0724435650,((WP_226989104.1-Desulfuromonas-sp.-TF:0.3195047936,WP_227459030.1-Cupriavidus-pauculus:0.1858490000)89.1/100:0.0970194017,WP_150984056.1-Cupriavidus-basilensis:0.2677832288)87.4/97:0.0497432732)93/91:0.0685954881)94.2/65:0.0624181447)85.8/34:0.0293195597)77/32:0.0122190642,((WP_084055900.1-Desulfacinum-hydrothermale:0.2141862007,WP_123289658.1-Desulfosoma-caldarium:0.2072019378)99.4/98:0.1788556587,WP_136680209.1-Neptunomonas-sp.-XY-337:0.1678374459)82.5/64:0.0248768174)78.2/42:0.0133084104,((((WP_200222041.1-Achromobacter-xylosoxidans:0.1211366575,WP_259251071.1-Achromobacter-sp.-JUb104:0.1333928278)95.2/100:0.0663580975,(WP_088588627.1-Achromobacter-marplatensis:0.1273064478,WP_169273406.1-Achromobacter-sp.-Bel:0.0764795709)93.2/100:0.0644244550)0/55:0.0262900382,((((((WP_111826915.1-Achromobacter-sp.-HZ01:0.0851218336,WP_102775120.1-Achromobacter:0.0206355396)99.9/100:0.1224832522,WP_129525645.1-Achromobacter-agilis:0.0224234327)0.8/36:0.0190388658,WP_043544564.1-Achromobacter-sp.-RTa:0.0259924708)21.9/66:0.0194825699,WP_129240460.1-Achromobacter-veterisilvae:0.0255784674)98.3/99:0.0633699988,(((WP_175208175.1-Achromobacter-anxifer:0.0459655434,(((WP_251864322.1-Achromobacter-sp.-Marseille-Q4962:0.0640130904,WP_164793551.1-Achromobacter-insolitus:0.0283117961)97.5/98:0.0482424280,WP_207472676.1-Achromobacter-sp.-SD115:0.0086635106)92.9/98:0.0293636749,WP_125471167.1-Achromobacter:0.0178100935)94.1/98:0.0368687632)86.2/69:0.0127528326,WP_175164188.1-Achromobacter-aegrifaciens:0.0397045188)86.7/68:0.0266870508,WP_088158656.1-Achromobacter-xylosoxidans:0.0556777474)99.3/100:0.0826071377)76.4/95:0.0131340216,WP_174717299.1-Achromobacter-denitrificans:0.0742154538)84.5/98:0.0254896105)99.8/100:0.1180675316,(WP_006216420.1-Achromobacter-piechaudii:0.0796252349,(((WP_175192416.1-Achromobacter-deleyi:0.0643395570,WP_100503206.1-Achromobacter-ruhlandii:0.1231541031)74.1/90:0.0313866044,WP_183017169.1-Achromobacter-sp.-UMC46:0.0518382231)90.9/90:0.0384666699,WP_219991400.1-Achromobacter-sp.-ES-001:0.0652051202)48.6/89:0.0348005147)100/100:0.1557064276)93.5/95:0.0533472612)83.1/28:0.0245188842,(((((((((((((((((WP_065305794.1-Janthinobacterium-psychrotolerans:0.1175988889,(WP_139090581.1-Janthinobacterium-lividum:0.0581874956,WP_230539136.1-Janthinobacterium-sp.-FW305-129:0.0332824572)59.8/99:0.0098118788)74.6/62:0.0055029099,WP_215403374.1-Janthinobacterium-sp.-JC611:0.0116386366)87.6/62:0.0116630813,WP_094442213.1-Janthinobacterium-sp.-PC23-8:0.0597249888)74.3/62:0.0056986655,WP_102124214.1-Janthinobacterium-sp.-ROICE36:0.0292655078)77.7/75:0.0057811454,WP_096238059.1-Janthinobacterium-svalbardensis:0.0172751402)74.9/74:0.0057196295,(((WP_035822645.1-Janthinobacterium-sp.-RA13:0.0548316108,WP_034778721.1-Janthinobacterium-lividum:0.0358131278)73.4/99:0.0053664356,WP_152282105.1-Janthinobacterium-violaceinigrum:0.0176290034)75.5/99:0.0056931976,WP_183742413.1-unclassified-Janthinobacterium:0.0409782001)70.3/98:0.0057585836)99.3/99:0.0772250950,WP_101482628.1-unclassified-Janthinobacterium:0.1481534251)25.8/93:0.0215297425,(WP_099917421.1-Massilia-psychrophila:0.1106634385,(WP_205319084.1-Glaciimonas-sp.-PAMC28666:0.0569015631,WP_092396716.1-Collimonas-sp.-OK412:0.1016071858)27.9/89:0.0131452013)89.4/95:0.0302717180)73.7/74:0.0077139464,WP_023495308.1-Methyloglobulus-morosus:0.1385043069)97.7/95:0.0633829263,WP_217900122.1-Noviherbaspirillum-humi:0.1676805600)76/42:0.0167069211,(((WP_162020603.1-Undibacterium-crateris:0.1185556003,WP_186861877.1-Undibacterium-griseum:0.0582221713)14.1/57:0.0124664707,WP_186884883.1-Undibacterium:0.0359126375)99/100:0.1128424684,(((((((((WP_207000528.1-Trinickia-sp.-DHG64:0.0942466713,WP_124149903.1-Paraburkholderia-dinghuensis:0.0912849280)85.4/100:0.0411449471,(WP_266250364.1-Paraburkholderia-megapolitana:0.1364474242,WP_026121090.1-Paraburkholderia-kururiensis:0.0835850542)79.7/100:0.0493003073)92.8/100:0.0536479158,((WP_179404315.1-Burkholderia-guangdongensis:0.1320137481,(WP_063494936.1-Paraburkholderia-phytofirmans:0.0328418339,(WP_254318368.1-Paraburkholderia-sp.-CNPSo-3272:0.0000022888,WP_042262232.1-Paraburkholderia-heleia:0.0593653742)91.4/100:0.0328684163)99.6/100:0.1042033729)9.4/35:0.0308281856,WP_038711716.1-Burkholderia-sp.-lig30:0.1729364060)91/82:0.0523221536)88/61:0.0691165359,((WP_201138986.1-Paraburkholderia-domus:0.0417763760,WP_266262001.1-Paraburkholderia-madseniana:0.0199866867)99.3/100:0.0897914167,WP_009889585.1-Burkholderia-thailandensis:0.0712462873)82.9/99:0.0269208262)81.3/35:0.0251242758,(((((((((WP_059459219.1-Burkholderia-vietnamiensis:0.1821869493,WP_121276956.1-Trinickia-fusca:0.0914738046)34.9/100:0.0325833085,WP_134195781.1-Paraburkholderia-rhizosphaerae:0.1998690378)84.4/51:0.0246887056,WP_136892825.1-Trinickia-sp.-7GSK02:0.0393212621)81/51:0.0131362703,WP_201650988.1-Paraburkholderia-sabiae:0.0354072418)43.5/41:0.0125547731,WP_205970438.1-Paraburkholderia-sp.-Tr-20389:0.0375432049)54.6/41:0.0132337778,WP_054928771.1-Paraburkholderia-caribensis:0.0302728719)86.3/51:0.0208980919,(WP_205976236.1-Paraburkholderia-sp.-Se-20369:0.1840343455,WP_206951398.1-Trinickia-sp.-4D114:0.0617504281)98.5/92:0.0822174165)82.6/49:0.0149131436,WP_209935850.1-Paraburkholderia-sp.-LEh10:0.0771421706)89.1/51:0.0201790361,((((WP_060156927.1-Burkholderia-ubonensis:0.0725328029,WP_059889870.1-Burkholderia-stagnalis:0.1554246510)89.7/98:0.0275103970,(WP_006496666.1-Burkholderia-cenocepacia:0.0180273954,WP_059716874.1-Burkholderia-ubonensis:0.0398213578)89.2/100:0.0170055058)74.3/85:0.0095083752,(WP_061168975.1-Caballeronia-hypogeia:0.0582389094,WP_069272519.1-Burkholderia-cepacia:0.1739017770)86.2/82:0.0212207737)38.5/88:0.0120313477,((WP_175791145.1-Burkholderia-ambifaria:0.0835983894,WP_059842355.1-Burkholderia-multivorans:0.0298858229)83.8/91:0.0114031742,WP_105772332.1-Burkholderia-multivorans:0.0292647593)88.6/94:0.0229270676)64.3/81:0.0069439770)73.2/48:0.0068105325)86/35:0.0159454419,(WP_060092900.1-Burkholderia-vietnamiensis:0.0203361221,WP_226113708.1-Burkholderia-sp.-AU33545:0.0400026360)100/100:0.1268593508)72.1/38:0.0084435231,WP_168792792.1-Paraburkholderia-aromaticivorans:0.0573778760)72.6/37:0.0113082186,(WP_004544228.1-Burkholderia:0.0455064228,WP_059583574.1-pseudomallei-group:0.0297519022)86.1/99:0.0127436299)97.8/85:0.0896579773,(WP_056136218.1-Massilia-sp.-Root133:0.0839066898,WP_108623287.1-Orrella-marina:0.4139176356)73.9/98:0.0865044755)95.9/99:0.0795963751)78.2/82:0.0456164162)79.8/47:0.0174473040,((WP_247737070.1-Thauera-aromatica:0.0556097673,WP_039805139.1-Azotobacter-chroococcum:0.1142382187)98.9/100:0.0894080364,((WP_195874647.1-Herminiimonas-contaminans:0.1026542702,WP_041297045.1-Janthinobacterium-sp.-Marseille:0.0756659248)92.9/100:0.0398970490,((WP_076590920.1-Herminiimonas-arsenitoxidans:0.1413753672,WP_206604468.1-Herminiimonas-fonticola:0.0338715301)79.5/45:0.0178809475,WP_188420499.1-Oxalicibacterium-solurbis:0.1281939382)73.8/92:0.0256933550)98.6/99:0.0853143614)50.7/38:0.0207216308)79.5/84:0.0358403025,(WP_199264031.1-Desulfobulbus-oligotrophicus:0.1945853967,WP_153720048.1-Spiribacter-sp.-C176:0.2140918608)92.3/98:0.0662365015)54.4/48:0.0278723170,(WP_250024752.1-Cupriavidus-campinensis:0.1309922643,WP_094797980.1-Bordetella-genomosp.-5:0.1381050922)98.4/98:0.0772260648)23.3/12:0.0117161594,WP_250479419.1-unclassified-Caballeronia:0.2382323557)38.2/30:0.0287749606,((WP_243478771.1-Orrella-sp.-f23:0.3106303458,WP_227181580.1-Leeia-sp.-TBRC-13508:0.1741127643)87.6/100:0.0557181869,(WP_071014919.1-Cupriavidus-sp.-USMAHM13:0.1546667829,WP_174426450.1-Cupriavidus-basilensis:0.2555729949)90.1/97:0.0722447690)89.9/58:0.0737139454)40.1/74:0.0539217102,((WP_217345508.1-Noviherbaspirillum-sp.-L7-7A:0.1396342756,WP_237247350.1-Sideroxyarcus-emersonii:0.2062208688)78.4/78:0.0401560651,WP_047549861.1-Methylotenera-sp.-G11:0.1678379896)50.4/94:0.0381027256)31.9/39:0.0430718897)89.4/66:0.0225248367)90.1/60:0.0242048542)80.4/23:0.0335264015,((WP_153584563.1-Caenimonas-koreensis:0.2109270724,(WP_007185580.1-Hydrocarboniphaga-effusa:0.0601181353,WP_007185565.1-Hydrocarboniphaga-effusa:0.0651096208)99.5/100:0.1406130171)93.2/65:0.0959434487,WP_209394989.1-Rhodoferax-sp.-AJA081-3:0.2433503921)41.1/39:0.0478293129)48.2/57:0.0283871525,((((((((WP_198111724.1-Inhella-proteolytica:0.2478214775,((WP_037482909.1-Sphaerotilus-natans:0.1435521574,(WP_028996355.1-Azohydromonas-australica:0.0617330471,WP_066337818.1-Azohydromonas-lata:0.0799470158)96.4/100:0.0790863764)92.5/100:0.0645384500,WP_241396682.1-Pelomonas-sp.-CA6:0.1280164878)76/100:0.0758426184)98.1/100:0.1378586134,WP_220807643.1-Noviherbaspirillum-aridicola:0.1972982728)84/96:0.0714013217,WP_122244186.1-Vandammella-animalimorsus:0.1560961532)93.9/71:0.0679041221,((WP_026436726.1-Acidovorax-sp.-JHL-9:0.1352618330,WP_086926522.1-Acidovorax-carolinensis:0.1364344636)85.1/96:0.0330087931,WP_174484170.1-methanotrophic-endosymbiont-of-Bathymodiolus-puteoserpentis:0.1521299445)79.8/92:0.0212529389)90.5/72:0.0387538079,(((((WP_223986959.1-Cupriavidus-pampae:0.0701107645,WP_019451253.1-Cupriavidus-sp.-BIS7:0.0413976350)12.2/73:0.0055009165,WP_082079449.1-Cupriavidus-basilensis:0.0728735822)78.5/84:0.0170222411,WP_011517306.1-Cupriavidus-metallidurans:0.0297205205)99.8/100:0.1207523443,WP_137918285.1-Hydrogenophaga-sp.-2FB:0.1155140724)68.2/71:0.0316404245,WP_137890779.1-Ramlibacter-sp.-2FC:0.0889207889)37.6/70:0.0211106206)83.5/80:0.0192168082,((((WP_101249932.1-Telmatospirillum-siberiense:0.3365609994,WP_056898839.1-Pseudorhodoferax-sp.-Leaf274:0.1717214819)79.8/87:0.0386573088,((WP_106935816.1-unclassified-Variovorax:0.1901184868,(((WP_126747644.1-Variovorax-sp.-DXTD-1:0.1286741992,WP_206177103.1-Variovorax-sp.-RKNM96:0.0808227694)58.4/51:0.0244193931,WP_068630051.1-Variovorax-sp.-PAMC-28711:0.1857834353)23.7/8:0.0134076228,(WP_093207264.1-Variovorax-sp.-YR750:0.0800117825,((WP_192194832.1-Variovorax-sp.-38R:0.1785541479,WP_093441791.1-Variovorax-sp.-770b2:0.1099609268)13/8:0.0280938300,WP_160553990.1-Xylophilus-rhododendri:0.1252023663)96.5/98:0.0656832023)22.8/6:0.0242092940)92/100:0.0386402785)0/42:0.0229559805,WP_219210863.1-Variovorax-boronicumulans:0.1487044484)87/72:0.0266252933)91.8/96:0.0400417132,(WP_057673948.1-Curvibacter-sp.-PAE-UM:0.1304348715,WP_169420802.1-Ramlibacter-agri:0.1624143787)25.6/93:0.0273837230)77.7/89:0.0136823425,(WP_074796726.1-Nitrosospira-briensis:0.0651128171,(WP_090882638.1-Nitrosovibrio-sp.-Nv6:0.0507640753,WP_097065525.1-Nitrosovibrio-sp.-Nv4:0.0337421170)89.7/100:0.0425910635)100/100:0.1912002538)57/73:0.0054094700)88.7/98:0.0340231611,(WP_036252969.1-Methylobacter-sp.-BBA5.1:0.2073744449,WP_212785890.1-Ferrigenium-kumadai:0.1717214998)83.4/83:0.0391538302)89.2/83:0.0303074096,((WP_160809027.1-Nitrosomonas-sp.-HPC101:0.3097369956,WP_154206467.1-Ralstonia-pickettii:0.1278417628)49.6/99:0.0513525502,WP_253536546.1-Ralstonia-sp.-21YRMH01-3:0.2636493479)92.3/99:0.0492410561)83.2/84:0.0287692934)81.3/71:0.0343123056,((WP_108647113.1-Polynucleobacter-rarus:0.0263337504,WP_084281813.1-Polynucleobacter-kasalickyi:0.0309805574)100/100:0.3517510553,((WP_215371371.1-Polynucleobacter-sp.-MWH-UH24A:0.1632566702,WP_173942040.1-Polynucleobacter-antarcticus:0.1394372791)93.1/100:0.0803941365,((((WP_128113711.1-Polynucleobacter-necessarius:0.0991048335,WP_215385554.1-Polynucleobacter-sp.-AP-Kolm-20A-A1:0.0617688287)94.3/100:0.0661713647,WP_216177372.1-Polynucleobacter-sp.-AP-Feld-500C-C5:0.1451190645)84/100:0.0357333093,WP_114639791.1-Polynucleobacter-necessarius:0.0999146670)89.4/99:0.0405547550,((WP_216254756.1-Polynucleobacter-sp.-AP-Melu-500A-A1:0.0458960839,WP_216218975.1-Polynucleobacter-sp.-AP-Latsch-80-C2:0.0424089244)99.3/100:0.0791916622,(((((((WP_215319389.1-Polynucleobacter-paneuropaeus:0.0830298282,(WP_215352354.1-Polynucleobacter-sp.-MWH-P3-07-1:0.0410254578,WP_215380622.1-Polynucleobacter-paludilacus:0.0334462826)87.9/100:0.0366378653)97.5/99:0.0552334044,WP_062307935.1-Polynucleobacter-sinensis:0.0474762515)77.6/84:0.0112648451,WP_216197047.1-Polynucleobacter-sp.-UK-Mo-2m-Kol15:0.0625590271)74.9/66:0.0057980895,(((((WP_215376847.1-Polynucleobacter-asymbioticus:0.0248553013,WP_215328794.1-Polynucleobacter-sp.-AP-Sving-400A-A2:0.0496812747)98.2/100:0.0487083415,WP_173959551.1-Polynucleobacter-arcticus:0.0278901011)91.3/77:0.0217976225,WP_216225867.1-Polynucleobacter-sp.-MG-27-Goln-C1:0.0810069312)59.8/77:0.0062600866,(WP_216168716.1-Polynucleobacter-bastaniensis:0.0735033003,WP_216178894.1-Polynucleobacter-sp.-80A-SIGWE:0.0135567844)83.9/100:0.0153843895)88.2/77:0.0207346027,WP_215299358.1-Polynucleobacter-sp.-AP-Nino-20-G2:0.0830413882)88.3/47:0.0199012285)72.8/40:0.0057420219,WP_215334312.1-Polynucleobacter-sp.-AP-Ainpum-60-G11:0.0391889184)77.9/39:0.0111923589,(WP_216268685.1-Polynucleobacter-sp.-Nonnen-W13:0.0400010914,WP_251374783.1-Polynucleobacter-corsicus:0.0335498795)73.6/92:0.0056354379)92.7/93:0.0281321436,WP_076024025.1-Polynucleobacter-sphagniphilus:0.0788863940)67.8/54:0.0154686648)96.7/93:0.0564086377)72.3/93:0.0338765594)98.2/100:0.1447086373)97.9/100:0.1367624207)95.4/90:0.0617711496,(WP_070114691.1-Acidithiobacillus-caldus:0.2233451315,(((WP_009566950.1-Acidithiobacillus:0.0367314800,WP_215870853.1-unclassified-Acidithiobacillus:0.0664821367)96.8/100:0.0544901237,WP_163095688.1-Acidithiobacillus-ferrianus:0.0411580714)46.2/84:0.0395420626,WP_101536718.1-Acidithiobacillus-sp.-SH:0.1137810484)96.1/100:0.0730951940)99/100:0.1163190512)83.6/75:0.0310577603,((WP_230291920.1-Croceicoccus-sp.-Ery5:0.2935730541,WP_164156863.1-Sandarakinorhabdus-rubra:0.1769526569)56.8/100:0.0580237660,((WP_116367876.1-Parahaliea-mediterranea:0.0646108606,WP_148063811.1-Parahaliea-aestuarii:0.0872586074)99.9/100:0.1884141901,WP_129645432.1-Steroidobacter-agariperforans:0.2972286420)91/80:0.0778644410)87.6/64:0.0552993195)82.6/61:0.0337100666,((WP_132974196.1-Thiogranum-longum:0.1325044103,WP_201344172.1-Thiohalobacter-sp.-COW1:0.1409565224)98.8/100:0.1478698867,(((((((((((WP_036226274.1-Mesoaciditoga-lauensis:0.1572464335,WP_047753768.1-Kosmotoga-pacifica:0.1302011466)0/38:0.0249758857,(((WP_077198159.1-Thermosipho-affectus:0.0701243595,WP_012057290.1-Thermosipho-melanesiensis:0.1424459283)79/100:0.0357618992,(WP_127352844.1-Caldicellulosiruptor-changbaiensis:0.0516028277,WP_014043315.1-Caldicellulosiruptor-acetigenus:0.0274050236)100/100:0.2807970205)89.2/100:0.0461513267,((((((((WP_117708548.1-Fusobacterium-mortiferum:0.0090408228,(WP_204716698.1-Fusobacterium-mortiferum:0.0160654963,WP_204716813.1-Fusobacterium-mortiferum:0.3160818382)87.3/93:0.0412736310)84.3/93:0.0213083791,WP_115270465.1-Fusobacterium-necrogenes:0.0499520163)98.7/98:0.0809647860,(((WP_130889691.1-Fusobacterium-ulcerans:0.0388492109,(WP_005950923.1-Fusobacterium-varium:0.0378234172,(WP_070580525.1-Fusobacterium:0.0496267199,WP_227372109.1-Fusobacterium-ulcerans:0.0221312826)8.6/99:0.0128503078)85.9/100:0.0135173044)47.8/99:0.0115955998,(WP_241635364.1-Fusobacterium-gastrosuis:0.0996622537,(WP_074518138.1-Fusobacterium-necrophorum:0.0685846112,(WP_211495349.1-Fusobacterium-necrophorum:0.1268219523,WP_032890075.1-Fusobacterium-nucleatum:0.4280317345)82.3/37:0.0496891279)96.1/37:0.0623193850)97.6/37:0.0655897184)88.6/37:0.0329739506,WP_261793436.1-Fusobacterium-varium:0.2256474621)92.4/38:0.0442582281)90.1/40:0.0604993907,(((((WP_013633636.1-Pseudopedobacter-saltans:0.1158296663,WP_154287284.1-Pedobacter-puniceum:0.1271039117)81.4/39:0.0567497663,WP_160842504.1-Pedobacter-sp.-HMF7647:0.0641594000)46.4/39:0.0915301590,(WP_074492796.1-Mucilaginibacter-polytrichastri:0.1370145859,(WP_258137696.1-Mucilaginibacter-sp.-BJC16-A38:0.1432433825,(WP_259068220.1-Mucilaginibacter-sp.-X4EP1:0.0552114094,WP_144912900.1-Mucilaginibacter-frigoritolerans:0.0686136120)83.3/100:0.0709492809)75.1/100:0.0941913822)90.4/100:0.0870863386)95.1/93:0.1713942309,WP_163286637.1-Cryomorpha-ignava:0.3910564528)78.9/99:0.1167377351,WP_130092062.1-Brumimicrobium-glaciale:0.2292429092)99.9/100:0.2775886540)89.4/70:0.0560556067,((((WP_147004474.1-Leptotrichia-wadei:0.0215613348,WP_018499460.1-Leptotrichia-wadei:0.0332658003)99.8/100:0.1165151683,WP_068155536.1-Leptotrichia-sp.-oral-taxon-847:0.0614205502)20/35:0.0273617933,(WP_211617428.1-Leptotrichia-sp.-oral-taxon-218:0.0129029779,WP_094080762.1-Leptotrichia-sp.-oral-taxon-498:0.0442124770)96/100:0.0561854761)60.3/35:0.0266448377,WP_099992503.1-Fusobacterium-pseudoperiodonticum:0.0820597604)100/100:0.1936597875)92.3/98:0.0552689383,WP_013388736.1-Ilyobacter-polytropus:0.3108703529)86.6/78:0.0325722282,WP_012548442.1-Dictyoglomus-thermophilum:0.2684467496)76.2/72:0.0237812208,WP_173194267.1-Paenibacillus-frigoriresistens:0.2330716075)97.6/100:0.0904454150)89.4/100:0.0487528288)98.5/100:0.0907543146,(WP_099829233.1-Mesotoga-sp.-Brook.08.105.5.1:0.1414587318,WP_099830786.1-Mesotoga-sp.-Brook.08.105.5.1:0.1409863797)98/100:0.0995470467)91.9/100:0.0766903929,(((((((((((((((((((((((WP_054742706.1-Cellulosilyticum-ruminicola:0.0765793244,(WP_177671917.1-Zhenhengia-yiwuensis:0.0379363384,WP_053982981.1-Niameybacter-massiliensis:0.0956166834)99.6/100:0.0896578742)94.6/100:0.0502300293,WP_069999280.1-Cellulosilyticum-sp.-I15G10I2:0.1791889277)99/99:0.0976650662,WP_072892386.1-Clostridium-fallax:0.0828209200)89.1/84:0.0343866195,(WP_073337580.1-Clostridium-grantii:0.1168835989,WP_072832886.1-Clostridium-collagenovorans:0.0313601593)46.4/90:0.0105227785)89.1/74:0.0182004891,(((WP_152890015.1-Clostridium-tarantellae:0.1248144871,WP_195972826.1-Clostridium-thermobutyricum:0.0620516951)78.1/97:0.0345838951,(WP_209795868.1-Clostridium-moniliforme:0.0593677951,WP_194191032.1-Clostridium-chrysemydis:0.0904766738)91.5/99:0.0446275879)73.6/94:0.0103330887,WP_040194942.1-Clostridium-culturomicium:0.0639897976)86.1/89:0.0144160569)83.8/85:0.0122609982,((((((WP_094369216.1-Romboutsia-weinsteinii:0.1201371964,(WP_079421587.1-Clostridium-oryzae:0.3032477652,WP_021359979.1-Clostridioides-difficile:0.1817472470)77.1/99:0.0779213326)90/96:0.0541090460,(WP_074918830.1-Terrisporobacter-glycolicus:0.0568048145,WP_195929149.1-Clostridium-sp.-1001270J_160509_D11:0.0348882851)81.1/100:0.0197534791)76.7/95:0.0305450321,(((((WP_055265352.1-Clostridium:0.0351792260,(WP_186859336.1-Clostridium-sp.-NSJ-6:0.0545773798,(WP_055265304.1-Clostridium-disporicum:0.0546539660,(WP_154821366.1-Clostridium-saudiense:0.0088569640,WP_195245504.1-Clostridium-celatum:0.0609959094)73.2/99:0.0266165469)51.1/94:0.0253384138)75.3/94:0.0467341551)75.1/100:0.0526765709,WP_186834444.1-Eubacteriales:0.1062768357)93.6/100:0.0503881709,((((WP_235068992.1-Turicibacter-sp.-TJ11:0.0527033653,WP_212726030.1-Turicibacter-bilis:0.0349578349)50.2/54:0.0051769897,WP_195933776.1-Turicibacter-sanguinis:0.0231027042)87.7/54:0.0154411073,WP_155223812.1-Turicibacter-sanguinis:0.0224077418)86.9/62:0.0187668373,WP_161831114.1-Turicibacter-sp.-TS3:0.0312668487)99.2/99:0.0849185065)81/92:0.0213962494,WP_055258173.1-Sarcina-ventriculi:0.0857278455)81.2/53:0.0281608603,(WP_066874757.1-Clostridium-mediterraneense:0.1530005160,WP_198610231.1-Clostridium-perfringens:0.0303847371)79.2/87:0.0296250451)95.5/99:0.0473359486)52.4/50:0.0138710020,((((WP_092726456.1-Romboutsia-lituseburensis:0.1532427514,WP_258242885.1-Romboutsia-lituseburensis:0.0317492861)98.8/99:0.0754902364,WP_057560648.1-Paeniclostridium-sordellii:0.0547509320)89.7/97:0.0275292674,(WP_057580750.1-Paeniclostridium-sordellii:0.0987576379,WP_057538715.1-Paeniclostridium-sordellii:0.0371714923)81.7/100:0.0156698174)25.1/52:0.0127577131,WP_195252007.1-Romboutsia-sp.-1001713B170207_170306_H8:0.0412154314)94/97:0.0389928143)85.8/69:0.0176922960,(WP_095406848.1-Romboutsia-maritimum:0.1092141570,(((WP_195959433.1-Clostridium-tertium:0.0620034391,WP_195429094.1-Clostridium-sp.-D46t1_190503_E9:0.0164346518)83.9/99:0.0167598750,WP_047001949.1-Clostridium-sp.-C8:0.0527868044)81.8/81:0.0152859199,(WP_066890539.1-Clostridium-nigeriense:0.0429160147,WP_016207773.1-Clostridium-sartagoforme:0.0320981083)0/40:0.0033377716)75/96:0.0286963235)0/32:0.0148303849)77.5/60:0.0108099571,((((WP_029452249.1-Clostridium-algidicarnis:0.0898686286,WP_216183125.1-Clostridium-gasigenes:0.0202690257)59.3/99:0.0308788211,WP_216313757.1-Clostridium-gasigenes:0.0614337001)97.6/100:0.0537232178,(WP_197096717.1-Paeniclostridium-sordellii:0.0710632272,WP_238903698.1-Clostridium-sp.-YIM-B02506:0.0566574976)83/76:0.0166791894)72.6/33:0.0061029926,(WP_163195303.1-Clostridium-thermarum:0.1211846708,WP_066677671.1-Clostridium-septicum:0.0264238944)73/99:0.0149729443)89.9/91:0.0171480184)94.4/82:0.0274919611)75.2/77:0.0083721503,WP_261829490.1-Clostridium-sp.-TW13:0.1646603789)86/29:0.0186591008,(WP_072990516.1-Clostridium-cavendishii:0.2041997659,((((WP_216288229.1-Clostridium-psychrophilum:0.1208450805,WP_216137569.1-Clostridium-estertheticum:0.0284659664)71/100:0.0109834096,WP_220704225.1-Clostridium-estertheticum:0.0345252089)91.3/100:0.0393698675,WP_220321605.1-Clostridium-tagluense:0.0529785541)99.4/100:0.0771076520,WP_133628539.1-Fonticella-tunisiensis:0.0631774411)75.2/93:0.0104121506)89.7/91:0.0304190708)80.9/28:0.0109043737,(((((((((WP_169105633.1-Clostridium-sp.-SM-530-WT-3G:0.0543054602,WP_027636628.1-Clostridium-butyricum:0.0818686195)48.2/54:0.0226993471,WP_077866663.1-Clostridium-saccharobutylicum:0.1748075420)89.5/71:0.0290750694,WP_261829154.1-Clostridium-sp.-TW13:0.0639590450)86.7/13:0.0183338285,(((((WP_185652834.1-Clostridium-sp.-DJ247:0.0194965102,WP_185652641.1-Clostridium-sp.-DJ247:0.0625227872)90.9/93:0.0248734080,((WP_022750218.1-Clostridium-saccharobutylicum:0.0361768896,(WP_015391310.1-Clostridium-saccharoperbutylacetonicum:0.0570277240,(WP_077846315.1-Clostridium-puniceum:0.0441696998,WP_238915641.1-Clostridium-sp.-YIM-B02555:0.0535228414)85.1/98:0.0146994036)88/98:0.0219112530)24.8/92:0.0198974483,WP_226126241.1-Clostridium-algoriphilum:0.1175498099)31.7/33:0.0118656640)38.6/36:0.0033916302,((((WP_009169246.1-unclassified-Clostridium:0.0955577802,WP_077843022.1-Clostridium-beijerinckii:0.1232833109)83.9/100:0.0281235664,WP_041896941.1-Clostridium-beijerinckii:0.1562118364)97.5/100:0.0710567782,(WP_069680896.1-Clostridium-taeniosporum:0.1683522014,WP_160359894.1-Clostridium-chromiireducens:0.1998087278)12.5/52:0.0318995026)93.5/65:0.0563442879,(WP_265859166.1-Clostridium-botulinum:0.0962379682,((WP_154531076.1-Inconstantimicrobium-porci:0.1257561608,WP_009169009.1-Clostridium-sp.-DL-VIII:0.0167615510)69.1/34:0.0459597216,WP_077856313.1-Clostridium-beijerinckii:0.0000029991)96.2/34:0.0566251574)41.1/37:0.0266562858)88.8/35:0.0229254769)72.3/29:0.0152130541,WP_224033465.1-Clostridium-gelidum:0.0946474225)73.5/7:0.0057307725,(WP_077848125.1-Clostridium-puniceum:0.1202995126,(WP_077855754.1-Clostridium-beijerinckii:0.0367305146,WP_041894754.1-Clostridium-beijerinckii:0.0247463979)99.5/100:0.0735412136)68.7/7:0.0065250517)0/7:0.0000022849)99.1/99:0.0804315356,WP_216390068.1-Clostridium-sp.-MSJ-8:0.1928538793)80.1/98:0.0205737139,WP_090010800.1-Clostridium-sp.-DSM-8431:0.0932254308)74.5/85:0.0414311808,(WP_106059304.1-Clostridium-vincentii:0.0815673047,WP_143316579.1-Clostridium:0.0301992457)73.2/100:0.0192678311)80.5/82:0.0172775887,(WP_055069574.1-Clostridium-massiliamazoniense:0.1074029460,WP_148566871.1-Acetobacterium-paludosum:0.2855622534)35/51:0.0244470072)57.5/61:0.0180824147,WP_055184115.1-Clostridium-paraputrificum:0.0561833375)91.8/60:0.0240269507)90.3/90:0.0272031972,WP_066825124.1-Clostridium-tepidiprofundi:0.0861660508)93.8/93:0.0386691931,WP_072889493.1-Tepidibacter-formicigenes:0.0376230702)96.7/98:0.0680034901,WP_072725426.1-Tepidibacter-thalassicus:0.1285921918)7.3/67:0.0240704378,WP_055666468.1-Desnuesiella-massiliensis:0.1446826566)18.6/77:0.0286586565,WP_095404508.1-Romboutsia-maritimum:0.2335693365)87.1/100:0.0297129748,(WP_142413383.1-Hathewaya-massiliensis:0.1670221215,WP_035130905.1-Clostridium-sulfidigenes:0.1707624793)99/100:0.1038174347)91.3/95:0.0449475659,((((WP_227847393.1-Clostridioides-sp.-ZZV15-6598:0.0916535406,WP_227846117.1-Clostridioides-sp.-ES-S-0145-01:0.0279723613)96.1/100:0.0406229291,(WP_227842647.1-unclassified-Clostridioides:0.0290940466,WP_003420982.1-Clostridioides-difficile:0.0797743143)85.1/95:0.0202460461)52.8/95:0.0190248970,WP_038810653.1-Clostridioides-difficile:0.0354175258)100/100:0.2159202006,WP_226126924.1-Clostridium-algoriphilum:0.2359924416)87.8/99:0.0457219353)91.3/94:0.0375062947,WP_099190526.1-Tepidibacter-mesophilus:0.1019925833)95.4/94:0.0532723526,(WP_111570784.1-Halanaerobium-saccharolyticum:0.0678346887,(WP_089717478.1-Halanaerobium-congolense:0.0482594234,WP_089862761.1-Halanaerobium-salsuginis:0.0263183634)90.3/100:0.0366286477)100/100:0.2071867214)74.3/70:0.0153173459,WP_072903743.1-Hathewaya-proteolytica:0.2849506772)86.3/68:0.0330913360,((WP_009529486.1-Peptoanaerobacter-stomatis:0.1345177655,WP_068913445.1-Criibacterium-bergeronii:0.1424527225)100/100:0.1849724613,((WP_069974893.1-Geosporobacter-ferrireducens:0.2765713786,((WP_223038819.1-Crassaminicella-profunda:0.0939027936,WP_148809648.1-Crassaminicella-thermophila:0.0681548681)53.1/100:0.0306709285,WP_218283043.1-Crassaminicella-sp.-143-21:0.2181700766)61.4/100:0.0607592595)88.1/100:0.0697729041,WP_250858284.1-Oceanirhabdus-seepicola:0.1637785593)96.2/100:0.0798787412)7.1/58:0.0101984813)97.8/70:0.0681459425,((((((WP_068555806.1-Thermotalea-metallivorans:0.0957539592,WP_073236503.1-Desulforamulus-putei:0.0949849702)84/99:0.0212486317,WP_090442301.1-Natronincola-peptidivorans:0.1444969475)62.9/63:0.0192982032,((((WP_252130283.1-Clostridium-sp.-SYSU_GA19001:0.2402688707,WP_202749105.1-Clostridium-sp.-YIM-B02515:0.0900303224)96.5/99:0.1042445112,((((WP_066398085.1-Cytobacillus-eiseniae:0.2183039580,(WP_056034132.1-Paenibacillus-sp.-Leaf72:0.0845594298,WP_197936429.1-Paenibacillus-sp.-GSMTC-2017:0.1195418438)93.6/100:0.0634879700)84.4/90:0.0290610800,(WP_076307499.1-Paenibacillus-odorifer:0.0514435032,((WP_042140168.1-Paenibacillus-sp.-FSL-P4-0081:0.0110568533,WP_036721118.1-Paenibacillus-sp.-FSL-R7-277:0.0631768300)90.8/100:0.0303389793,WP_074087222.1-Paenibacillus-sp.-P32E:0.0303123082)93.1/100:0.0727277958)98.8/100:0.0848927668)92.1/90:0.0449879309,(WP_262684732.1-Paenibacillus-sp.-WQ-127069:0.1349284605,WP_090798944.1-Paenibacillus-sp.-GP183:0.1972764225)80.5/90:0.0339471613)87.5/100:0.0438200033,WP_036935152.1-Pseudobacteroides-cellulosolvens:0.1563866968)91.4/99:0.0632800450)80.6/80:0.0259227709,((WP_165412483.1-Clostridium-estertheticum:0.0598315085,WP_216302578.1-unclassified-Clostridium:0.0075143388)100/100:0.1764402564,((WP_204598070.1-Clostridium-pascui:0.1152759122,WP_216457288.1-Clostridium-simiarum:0.1239253591)8.6/48:0.0278218242,WP_027632043.1-Clostridium-hydrogeniformans:0.1166721226)55.5/83:0.0248692267)85.1/96:0.0288389547)90.5/87:0.0307792066,((((WP_104410444.1-Clostridium-algidicarnis:0.0563477208,WP_185162668.1-Clostridium-algidicarnis:0.0434407097)99.8/100:0.1323743757,WP_234123145.1-Clostridium-hydrogenum:0.1566060305)83.6/79:0.0336589316,(WP_084114600.1-Clostridium-acidisoli:0.0818065775,(WP_026884435.1-Clostridium-akagii:0.0937817576,(WP_015614219.1-Clostridium-pasteurianum:0.0260075424,(WP_010235680.1-Clostridium-arbusti:0.0114123832,WP_003443690.1-Clostridium-pasteurianum:0.0467851030)88.1/100:0.0201635498)93/100:0.0318976379)29.2/90:0.0104429593)97.2/100:0.0595884147)81.6/80:0.0164129684,(((WP_090038599.1-Clostridium-frigidicarnis:0.1229341961,WP_209459870.1-Youngiibacter-multivorans:0.1706806016)93.7/100:0.0629717856,((WP_164955553.1-Clostridium-estertheticum:0.1608259349,WP_125002390.1-Clostridium-tagluense:0.1267042594)72.8/97:0.0150574892,(WP_226136064.1-Clostridium-estertheticum:0.0220281825,WP_216146875.1-Clostridium-frigoris:0.0648701327)98.8/100:0.0882435571)98.8/100:0.0920622336)90.1/98:0.0349815437,WP_216462460.1-Clostridium-bornimense:0.1363248390)77.7/56:0.0211971725)96.2/95:0.0437251035)93.1/99:0.0387427806)90.6/54:0.0456966473,WP_035164300.1-Caloranaerobacter-azorensis:0.0638252357)71.5/52:0.0137700840,WP_095133872.1-Anaeromicrobium-sediminis:0.1488282911)95.5/87:0.0524492333,(((((((WP_088538685.1-Methanobrevibacter-sp.-87.7:0.0575598970,WP_016359256.1-Methanobrevibacter-sp.-AbM4:0.1033366543)98.3/100:0.0846372697,WP_067148181.1-Methanobrevibacter-olleyae:0.1056886087)84/100:0.0332565008,WP_116670296.1-Methanobrevibacter-woesei:0.1965948214)99.4/97:0.1101814813,(WP_248612240.1-Methanobacterium-alcaliphilum:0.1193199072,((WP_071907175.1-Methanobacterium-congolense:0.0208378362,WP_209584978.1-Methanobacterium-aggregans:0.0390784959)99.5/100:0.0967421397,(WP_069583666.1-Methanobacterium:0.1463919923,WP_048190917.1-Methanobacterium-sp.-SMA-27:0.1140629602)92.8/100:0.0577497043)74.5/96:0.0153503473)89.3/98:0.0254634078)47.1/87:0.0128156298,(WP_011406138.1-Methanosphaera:0.1273894136,(WP_112123902.1-Methanosphaera-sp.-BMS:0.0861241061,(WP_069592847.1-Methanosphaera-sp.-WGK6:0.0321081228,WP_095608065.1-Methanosphaera-cuniculi:0.0925283383)97.8/100:0.0724267418)44.1/99:0.0394991856)95.9/100:0.0635450199)88.9/95:0.0279533419,(WP_013644504.1-Methanobacterium-lacus:0.1756327186,WP_054834988.1-Methanobrevibacter-arboriphilus:0.1475703408)85.5/84:0.0333240958)97.1/81:0.0608411609,((WP_112093115.1-Methanothermobacter-tenebrarum:0.2200889009,WP_013413362.1-Methanothermus-fervidus:0.1612623520)94/100:0.1023415083,WP_213236459.1-Fusibacter-paucivorans:0.4686582792)85.2/70:0.0356819667)91/80:0.0285535823)17.5/42:0.0077001297)80.7/69:0.0234312335,WP_038266426.1-Peptoclostridium-litorale:0.3432530780)55.7/83:0.0164329458,((((((((WP_058965344.1-Fournierella-massiliensis:0.1480378244,WP_185752312.1-Butyrivibrio-sp.-AE2032:0.2551560362)41.6/98:0.0540090708,WP_155202396.1-Ruthenibacterium-lactatiformans:0.1557516295)82.4/100:0.0741508745,(WP_186886939.1-Anaerofilum-hominis:0.1521898973,WP_102049807.1-Pygmaiobacter-massiliensis:0.1869087698)67.6/99:0.0635493523)99.8/100:0.1825913537,((WP_266162911.1-Caproicibacterium-sp.-BJN0003:0.1243121468,WP_256135338.1-Bittarella-massiliensis:0.2261817240)66/99:0.0588778930,WP_220321633.1-Clostridium-tagluense:0.2378852043)54.6/91:0.0637532218)47.3/82:0.0310148546,((((((((WP_072827923.1-Fibrobacter-sp.-UWB12:0.0228834710,WP_073304542.1-Fibrobacter:0.1702457979)47.1/92:0.0218556910,WP_088630837.1-unclassified-Fibrobacter:0.0154816541)100/100:0.1642747974,WP_073188830.1-Fibrobacter-sp.-UWP2:0.0000022395)71.4/90:0.0291587036,WP_073155408.1-Fibrobacter:0.0363959275)54.2/41:0.0296069513,WP_097034884.1-Fibrobacter-sp.-UWT3:0.0371492921)48.6/44:0.0125724030,WP_073056819.1-unclassified-Fibrobacter:0.0095051122)100/99:0.2187739140,WP_022475313.1-Huintestinicola-butyrica:0.1929346238)71.8/99:0.0533727174,WP_105204772.1-Neobittarella-massiliensis:0.1272462410)96.9/98:0.0745350705)80.9/87:0.0258152706,((((WP_091684654.1-Anaerocolumna-aminovalerica:0.1589632809,(WP_033168390.1-Clostridium-sp.-KNHs205:0.0820613369,((WP_073588199.1-Anaerocolumna-xylanovorans:0.0337756277,WP_185256524.1-Anaerocolumna-chitinilytica:0.1148014568)46.4/100:0.0367808415,WP_073279502.1-Anaerocolumna-jejuensis:0.0967685378)62.5/100:0.0354807879)99.8/100:0.1339701805)97.4/100:0.0897413712,((((WP_060930762.1-Lachnoanaerobaculum-saburreum:0.1466957907,((((((((((((((WP_024837362.1-Lacrimispora:0.0814429895,WP_124065139.1-Clostridium-sp.-E02:0.0566938279)87.4/58:0.0207203386,WP_117415169.1-Clostridium-indicum:0.0428280752)75.9/31:0.0095580486,WP_143321893.1-Clostridium-sp.-HBUAS56010:0.0297379384)59.5/29:0.0112650455,((WP_024293365.1-Lachnospiraceae:0.0329744189,WP_092242699.1-Lacrimispora:0.0244016927)94.5/99:0.0342009676,WP_077612318.1-Clostridium-sp.-Marseille-P2415:0.0367283355)86.8/41:0.0188563336)25/67:0.0078998667,WP_138313639.1-Hungatella-hathewayi:0.0276073869)73.6/93:0.0151455107,WP_186424215.1-Lacrimispora-celerecrescens:0.1374582665)95/62:0.0449618318,((((WP_231062061.1-Lacrimispora-sp.-NSJ-141:0.0872054470,WP_249304640.1-Qiania-dongpingensis:0.0557982596)96/94:0.0505794404,WP_118541387.1-unclassified-Clostridium:0.0632686401)73.8/30:0.0070912761,((WP_215718428.1-Anaerostipes-hadrus:0.0112518488,WP_075760385.1-Anaerostipes-sp.-494a:0.0472584789)99/100:0.0501753200,((WP_227052204.1-Anaerostipes-caccae:0.0300823961,WP_137330070.1-Anaerostipes-rhamnosivorans:0.0692899902)99.2/100:0.0792275886,WP_006781439.1-Hungatella-hathewayi:0.0810349311)84.1/78:0.0219435171)91.7/65:0.0228279218)84.8/25:0.0109928869,(WP_215700061.1-Clostridium-sp.-MCC353:0.0592347548,(((((WP_251497048.1-Otoolea-sp.-DSM-111138:0.1366929342,(WP_003497689.1-_Clostridium:0.0367252591,WP_125142063.1-Clostridium-transplantifaecale:0.0469256834)81.2/90:0.0208234326)92.3/85:0.0315837845,WP_118742831.1-Clostridium-sp.-AM28-20LB:0.0130004615)30.5/68:0.0164433710,WP_008399284.1-unclassified-Clostridium:0.0291642080)0/65:0.0000025365,(WP_087223935.1-Lachnoclostridium-sp.-An14:0.0384383388,WP_101876396.1-Lachnoclostridium-edouardi:0.0456905997)95.7/100:0.0328444928)89.4/86:0.0175745374,WP_118243934.1-Clostridium-sp.-AM58-1XD:0.0463621338)96.1/98:0.0290187692)72.7/30:0.0056754178)83.9/58:0.0113911569)74.9/43:0.0064164033,WP_024736308.1-Enterocloster-asparagiformis:0.0410055428)74.7/38:0.0057541517,WP_066545911.1-Clostridium-sp.-AT4:0.0234617151)85.9/51:0.0281605791,WP_002577448.1-Enterocloster:0.0577932442)74.3/30:0.0064847083,(WP_045093538.1-Clostridium-sp.-FS41:0.0414230022,WP_025485467.1-Lachnospiraceae:0.0541821606)89/99:0.0174825681)87.2/30:0.0114561430,WP_178313416.1-Lacrimispora-sp.-210928-DFI.3.58:0.0057038232)82.1/31:0.0157264185,WP_154471253.1-Clostridium-porci:0.0982559436)93.6/98:0.0525513683,WP_118555425.1-Lachnotalea-sp.-AF33-28:0.1617576927)80.2/100:0.0491207053)94.8/100:0.0783099543,WP_005539496.1-Johnsonella-ignava:0.1714767711)88.7/100:0.0494139558,((((((((((((((((WP_035764434.1-Butyrivibrio-sp.-NC2002:0.0923047109,((WP_026526812.1-Butyrivibrio-sp.-VCD2006:0.1359107481,WP_026491547.1-unclassified-Butyrivibrio:0.0162948499)40.7/99:0.0347734812,WP_044913869.1-Butyrivibrio-sp.-WCE2006:0.0200061721)93.6/100:0.0547268301)36.4/97:0.0366770630,WP_091822189.1-Butyrivibrio-sp.-ob235:0.0450774834)99.3/100:0.1071158175,(((((((WP_022768176.1-unclassified-Butyrivibrio:0.0199840724,WP_022766545.1-Butyrivibrio-sp.-XPD2006:0.0480833978)99/100:0.0803151193,WP_034446055.1-Butyrivibrio-sp.-AE2032:0.0563869318)88.6/99:0.0411549832,WP_024867291.1-Butyrivibrio-sp.-FCS014:0.0236145272)87.1/33:0.0245697379,(WP_071175335.1-Butyrivibrio:0.0569575111,WP_026513668.1-Butyrivibrio-sp.-LB2008:0.0411787430)80/33:0.0325315743)85.3/98:0.0235040940,WP_026663187.1-Butyrivibrio-proteoclasticus:0.0600765738)71.1/45:0.0146128642,(WP_013279951.1-Butyrivibrio-proteoclasticus:0.0278230253,WP_092060841.1-Butyrivibrio-sp.-INlla14:0.0428278429)87.7/47:0.0142126704)0/23:0.0000028016,WP_026652320.1-Butyrivibrio-proteoclasticus:0.0373506377)96.3/97:0.0693046582)98/100:0.0880528118,(((WP_019160740.1-Blautia:0.0536321118,WP_173738571.1-Blautia-schinkii:0.0236169855)95.5/100:0.0382775794,WP_158421413.1-Blautia:0.0289165235)49.5/58:0.0202939562,(WP_148445023.1-Eubacteriales:0.0272423374,WP_187566223.1-Blautia-faecis:0.0428539854)77/100:0.0112499906)98.2/100:0.0754420400)81.6/98:0.0457706647,WP_109624131.1-Murimonas-intestini:0.0588271188)40.9/54:0.0190754856,WP_178047055.1-Fusicatenibacter-sp.-CLA-AA-H277:0.0934929699)66.3/94:0.0231106673,(WP_173751851.1-Blautia-caecimuris:0.0443286813,WP_117997894.1-Blautia-obeum:0.0131375271)96.4/100:0.0544742695)68.7/79:0.0288761599,(((WP_158367551.1-Lachnospiraceae:0.0249710296,WP_235247881.1-Faecalicatena-contorta:0.0725199167)84.3/100:0.0365857995,(WP_251390531.1-Mediterraneibacter-sp.-DSM-111629:0.0307299073,WP_073962323.1-_Ruminococcus:0.0296462451)89.2/100:0.0227897729)82.2/89:0.0145697840,WP_070089756.1-Merdimonas-faecis:0.0299640938)77.8/83:0.0179148788)0/64:0.0017577844,WP_230400100.1-Novisyntrophococcus-fermenticellae:0.1487510780)87.9/94:0.0284979511,WP_132379639.1-Muricomes-intestini:0.0909547386)95.5/99:0.0438515807,WP_087156947.1-Lachnoclostridium-sp.-An181:0.0417624614)84.3/92:0.0259818874,WP_115483253.1-Anaerosacchariphilus-polymeriproducens:0.1062074808)80.2/83:0.0244507672,WP_073994331.1-Parasporobacterium-paucivorans:0.1350454069)40.9/81:0.0181035530,(WP_099466994.1-Konateibacter-massiliensis:0.0697888738,WP_151145296.1-Candidatus-Galacturonibacter-soehngenii:0.0443279874)98.8/100:0.0873925704)88.4/93:0.0359135440,WP_158351687.1-Diplocloster-modestus:0.1782389892)88.2/97:0.0350058864,WP_158346166.1-Lachnospiraceae:0.1935714623)98.4/100:0.0913528624)63.7/99:0.0246736951,WP_226914794.1-Intestinibacter-bartlettii:0.2499586819)88.8/99:0.0338943391)81.2/95:0.0309843540,(((((WP_073273890.1-Anaerocolumna-jejuensis:0.1249321965,WP_033167869.1-Clostridium-sp.-KNHs205:0.1429527178)94/83:0.0766998380,((WP_155699987.1-Paenibacillus-psychroresistens:0.0805238640,WP_090040610.1-Clostridium-frigidicarnis:0.0223786414)87.2/78:0.0207745408,WP_202751253.1-Clostridium-sp.-YIM-B02515:0.0437550373)81.6/78:0.0296095538)5.8/69:0.0381812056,WP_010268185.1-Paenibacillus-senegalensis:0.1498085344)98.8/100:0.1333564811,(WP_249305773.1-Jutongia-hominis:0.1514615598,(WP_022463459.1-Jutongia-huaianensis:0.0671409874,(WP_055944671.1-unclassified-Clostridium:0.0384273914,WP_216564272.1-Eubacterium-sp.-MSJ-13:0.0249888166)94.5/100:0.0467210652)44.7/100:0.0378905706)99.8/100:0.1394890808)88.1/100:0.0409480213,(WP_253721803.1-Treponema-sp.-OMZ-787:0.3070299405,WP_058352784.1-Acetivibrio-ethanolgignens:0.1702795295)86.2/93:0.0734305215)97.4/91:0.0657452846)92.7/92:0.0532905890,(WP_138788932.1-Paenibacillus-thermoaerophilus:0.2828974826,WP_078711972.1-Anaerorhabdus-furcosa:0.3846532495)97/89:0.1611153719)80.1/83:0.0646518012)95.7/91:0.0720234631,WP_119970050.1-Clostridium-fermenticellae:0.2135340879)0/21:0.0224218982,WP_160686059.1-Clostridium-sp.-C2-6-12:0.1977071674)81.7/81:0.0346556028)93.8/99:0.0693867131)97/99:0.0972512306,((((WP_018154793.1-Methanothermococcus-thermolithotrophicus:0.1055689707,WP_048201153.1-Methanocaldococcus-bathoardescens:0.0644128276)91.8/100:0.0447653148,((WP_052562942.1-Candidatus-Brocadia-sinica:0.2832583921,WP_173298646.1-Thermanaeromonas-sp.-C210:0.2346222116)64.9/99:0.0586732090,(((WP_011869065.1-Methanococcus-maripaludis:0.0795347479,WP_013998786.1-Methanococcus-maripaludis:0.0478391592)84.3/100:0.0332923348,WP_181493664.1-Methanococcus-maripaludis:0.0640863418)81.1/100:0.0258479866,WP_181504225.1-Methanococcus-maripaludis:0.0357595724)100/100:0.1815193719)87.5/99:0.0532163978)97.3/99:0.0861190479,(((WP_245533116.1-Coprothermobacter-platensis:0.2954811498,(WP_119360783.1-Meiothermus-luteus:0.0835978513,(WP_208013758.1-Meiothermus-sp.-CFH-77666:0.1066772997,WP_027877894.1-Meiothermus-cerbereus:0.0531254810)87/99:0.0765292581)100/100:0.4010013411)94.6/80:0.1281285090,(WP_150071182.1-Thermodesulfobacterium-sp.-TA1:0.1739507075,WP_022856039.1-Thermodesulfobacterium:0.1906756330)33.8/80:0.0433186662)91/98:0.0790990702,(((((WP_011996389.1-Coxiella-burnetii:0.4984622693,WP_013638952.1-Desulfurobacterium-thermolithotrophum:0.0325720340)11.1/81:0.0121684477,WP_132526207.1-Phorcysia-thermohydrogeniphila:0.0708837384)86.5/100:0.0254266548,WP_089322203.1-Desulfurobacterium-atlanticum:0.1307348444)93.1/100:0.0571823183,(WP_029522925.1-Persephonella-sp.-KM09-Lau-8:0.0957235369,(WP_200674641.1-Persephonella-atlantica:0.0693493271,WP_096999769.1-Persephonella-hydrogeniphila:0.0934795851)93/98:0.0538525128)79.9/87:0.0569393254)9.4/61:0.0302930910,WP_028950938.1-Sulfurihydrogenibium-subterraneum:0.1045029689)97.2/100:0.0912154442)94.6/98:0.0820473444)57.9/91:0.0579475272,((WP_010868297.1-Pyrococcus-abyssi:0.0340576881,(((((WP_013798345.1-Methanotorris-igneus:0.0315800209,WP_013866394.1-Methanothermococcus-okinawensis:0.1580243586)80.2/96:0.0264457142,WP_007044577.1-Methanotorris-formicicus:0.0569717401)98.4/97:0.0637428936,(((((WP_237702072.1-Thermococcus-sp.-AM4:0.0391420713,WP_223211916.1-Thermococcus-henrietii:0.0475196359)98.1/100:0.0554337215,WP_014733609.1-Pyrococcus-sp.-ST04:0.1060701799)50.1/67:0.0120686517,((((WP_088865724.1-Thermococcus-barossii:0.0577327915,((WP_167711671.1-unclassified-Thermococcus:0.0189675298,WP_088181368.1-Thermococcus-sp.-5-4:0.0645458005)74/76:0.0058343274,WP_232460978.1-Thermococcus-radiotolerans:0.0439074206)0/65:0.0000021432)78.8/67:0.0115602108,WP_258085050.1-Thermococcus-sp.-813A4:0.0312660325)65.4/47:0.0136699778,(WP_139679991.1-Thermococcus-indicus:0.0291081949,WP_148882943.1-Thermococcus-aciditolerans:0.0335120736)89/73:0.0214003360)74.4/66:0.0058898663,WP_058938040.1-Thermococcus-celericrescens:0.0286101234)84.7/79:0.0151470051)87.7/96:0.0295250658,((WP_152879125.1-Thermococcus-sp.-101-C5:0.0450224593,WP_015850310.1-Thermococcus-sibiricus:0.0441938080)41.3/100:0.0116210278,(WP_209475361.1-Thermococcus-stetteri:0.0733826024,WP_253304977.1-Thermococcus-aggregans:0.0173701718)83.2/59:0.0136898125)85.6/52:0.0168180697)73.7/63:0.0093067930,WP_068664737.1-Thermococcus-piezophilus:0.0512670639)94.4/96:0.0325951506)80/53:0.0169112420,((WP_167913028.1-Thermococcus-sp.-21S7:0.0670589577,WP_048150691.1-Palaeococcus-ferrophilus:0.0234171056)97.5/98:0.0422844988,((((WP_088855609.1-Thermococcus-siculi:0.0479995972,WP_087036111.1-Thermococcus-litoralis:0.0308167507)87.4/100:0.0180411199,WP_167888701.1-Thermococcus-sp.-MV5:0.0643120155)88.9/92:0.0187751477,(WP_013468354.1-Thermococcus-barophilus:0.0461413103,WP_013906489.1-Pyrococcus-yayanosii:0.0187754376)86.8/94:0.0117914020)86.6/60:0.0123015656,WP_048053121.1-Pyrococcus-horikoshii:0.0246912178)80.7/53:0.0102134240)86.2/55:0.0136097820)89.9/81:0.0214935955,(WP_148120035.1-Methanofervidicoccus-sp.-A16:0.1480487055,WP_088862340.1-Thermococcus-celer:0.0779040746)0/51:0.0000022165)54.7/76:0.0448931812)96.2/95:0.0539770825,WP_048103006.1-Aciduliprofundum-boonei:0.1644820013)99.1/100:0.1353729213)99.9/100:0.1611918214)65.9/99:0.0629365429,WP_236098952.1-Dethiosulfovibrio:0.3652451052)97.3/100:0.1494234346,((((KAI6244242.1-Aphelenchoides-fujianensis:0.3744342901,KAI6170547.1-Aphelenchoides-bicaudatus:0.4519349066)82.5/100:0.0967746809,WP_110520589.1-Herpetosiphon-llansteffanensis:0.4273859353)79.9/100:0.0743573145,WP_077329105.1-Virgibacillus-siamensis:0.3573473673)41.1/94:0.0763541640,(((((((((WP_160564586.1-Apibacter-sp.-B2912:0.0982493050,(WP_055424523.1-Apibacter-mensalis:0.0799827097,(WP_160568957.1-unclassified-Apibacter:0.0142571100,WP_166004639.1-Apibacter-sp.-B2966:0.0423848129)98.9/100:0.0796001220)77.9/100:0.0481783207)97.3/100:0.0913816887,(WP_105245553.1-Apibacter-adventoris:0.1678972141,WP_146261858.1-Apibacter-muscae:0.0976156463)83.8/100:0.0444398448)100/100:0.1721831261,(WP_084015419.1-Moheibacter-sediminis:0.2772294338,WP_182042771.1-Moheibacter-lacus:0.2027297821)94.7/100:0.0858075944)84.5/100:0.0508232663,(WP_068596492.1-Vaginella-massiliensis:0.1311613710,((WP_230477342.1-Faecalibacter-bovis:0.0956146715,WP_125349156.1-Empedobacter-falsenii:0.0902560705)84.2/100:0.0321280307,(WP_125113420.1-Algoriella-xinjiangensis:0.0734567751,WP_092908829.1-Algoriella-xinjiangensis:0.0352478908)94.1/100:0.0463908332)97.8/100:0.0986216006)97.4/100:0.1040635803)96.3/100:0.0938775573,(WP_111126400.1-Mesonia-sp.-K7:0.4741650445,(WP_014783622.1-Aequorivita-sublithincola:0.2160091937,((((WP_230143398.1-unclassified-Pedobacter:0.0966777665,WP_215166022.1-Pedobacter-sp.-ISL-64:0.0776522150)96.2/100:0.0720438429,WP_133260761.1-Pedobacter-changchengzhani:0.1029990400)90.2/100:0.0495009337,((WP_056872627.1-Pedobacter-sp.-Leaf194:0.0826667714,(WP_123203876.1-Pedobacter-jejuensis:0.0648088278,(WP_188416210.1-Pedobacter-mendelii:0.0267072101,WP_056087060.1-Pedobacter-sp.-Leaf176:0.0455856235)62.1/97:0.0171909792)16.4/99:0.0148185239)97.1/85:0.0534140262,WP_109929147.1-Pedobacter-paludis:0.0305981641)71/35:0.0292470531)79.7/35:0.0434988970,WP_199156598.1-Pedobacter-sp.-ASV2:0.0729952306)99.3/85:0.1672900726)84.9/100:0.1183121553)62.2/63:0.0900790126)93.1/97:0.0769991424,((WP_107809536.1-Elizabethkingia-sp.-YR214:0.1289538240,WP_219971314.1-Chryseobacterium-sp.-LJ756:0.1149667518)71.2/100:0.0178706265,(WP_076395614.1-Chryseobacterium:0.0416371254,WP_088467981.1-Chryseobacterium-sp.-VAUSW3:0.1430497195)63.8/75:0.0399739376)99.9/100:0.1806795502)99/96:0.1054062533,((((WP_233378072.1-Maribellus-sp.-CM-23:0.0567246453,WP_247978154.1-Maribellus-sp.-YY47:0.0674746669)98.8/100:0.1113159000,(WP_167609329.1-Maribellus-sediminis:0.1440208822,(WP_045033348.1-Draconibacterium-sediminis:0.0280615854,WP_163345719.1-Draconibacterium-halophilum:0.0694458894)99.6/100:0.1179079895)94.3/100:0.0727105172)99.8/100:0.2093211589,(((WP_106543043.1-Prolixibacter-denitrificans:0.1046640709,WP_025865917.1-Prolixibacter-bellariivorans:0.0197535561)66.3/76:0.0047004196,WP_153632016.1-Prolixibacter-sp.-SD074:0.0489956405)100/100:0.1937352842,((WP_159517929.1-Sunxiuqinia-indica:0.0993682428,WP_053182762.1-Sunxiuqinia-dokdonensis:0.0835634345)81.3/100:0.0577839267,WP_133466511.1-Sunxiuqinia-elliptica:0.1241983048)98.3/100:0.1216866955)85.3/100:0.0777490475)68.1/96:0.0467233950,((WP_068701711.1-Paludibacter-jiangxiensis:0.2772895917,((WP_011500232.1-Methanococcoides-burtonii:0.1700778248,WP_135611779.1-Methanococcoides-sp.-AM1:0.1530096490)81.6/100:0.0630712498,((((WP_128503182.1-Methanosarcina-sp.-MSH10X1:0.1419423245,(WP_011033115.1-Methanosarcina-mazei:0.0755942576,WP_048134921.1-Methanosarcina-sp.-1.H.T.1A.1:0.0387630552)17.9/78:0.0342419676)42.3/96:0.0464236408,WP_048167026.1-Methanosarcina:0.0745287832)77.5/70:0.0374050874,WP_048177163.1-Methanosarcina-sp.-MTP4:0.0794585674)7.9/66:0.0411570554,(WP_048137589.1-Methanosarcina-horonobensis:0.1325749886,((WP_048184631.1-Methanosarcina-siciliae:0.0530163853,WP_048173663.1-Methanosarcina-siciliae:0.0530186590)55.9/100:0.0109845486,WP_011023680.1-Methanosarcina-acetivorans:0.0320526432)95.8/100:0.0644518844)89/100:0.0638933147)75.2/82:0.0787400213)98.8/100:0.1403606476)78/99:0.0684715652,(((WP_101263396.1-Labilibaculum-filiforme:0.0372050401,WP_245870435.1-Labilibaculum-manganireducens:0.0417256994)98.8/100:0.0826780602,(WP_152734748.1-Marinifilum-sp.-N1E240:0.0729226530,(WP_171595603.1-Marinifilum-caeruleilacunae:0.0587359398,(WP_268189532.1-Marinifilum-sp.-D737:0.0483700731,WP_054720444.1-Marinifilum-fragile:0.0321406666)84.1/100:0.0184997025)86.3/100:0.0223155291)94/100:0.0476911777)54.1/99:0.0376048161,WP_127342456.1-Ancylomarina-longa:0.1021640645)99.8/100:0.1510208526)86.6/97:0.0426278478)90.9/95:0.0665626335)82.2/52:0.0298627401,(((WP_205722928.1-Fulvivirga-lutea:0.3218742703,(((((((((((WP_053470876.1-unclassified-Flavobacterium:0.1387579926,WP_095952323.1-Flavobacterium-sp.-ACN6:0.0362432310)65.9/99:0.0065398516,WP_056200692.1-Flavobacterium-sp.-Root186:0.0310767450)86.8/99:0.0116702056,WP_115889823.1-Flavobacterium-cutihirudinis:0.0479158682)37.1/100:0.0053955518,WP_198857739.1-Flavobacterium-sp.-IB48:0.0361739509)79.8/100:0.0083822697,(WP_269237486.1-Flavobacterium-sp.-IMCC34775:0.0743834964,WP_163393939.1-Flavobacterium-limi:0.0335607524)0/98:0.0000023084)74.6/98:0.0108802801,WP_264205577.1-Flavobacterium-sp.-LS1R49:0.0226666277)90/99:0.0179343080,((WP_173968237.1-Flavobacterium-collinsii:0.0669229734,WP_264525312.1-Flavobacterium-sp.-N502536:0.0420178661)92.8/99:0.0287145359,WP_070907856.1-Flavobacterium-tructae:0.0737822692)72.8/88:0.0064250798)77.2/87:0.0087249314,WP_091091738.1-Flavobacterium-gillisiae:0.0886267046)98.6/96:0.0579330381,WP_248907802.1-Flavobacterium-sp.-K5-23:0.1078845944)53/95:0.0297358296,(((WP_116761981.1-Flavobacterium-laiguense:0.0420164604,WP_143377960.1-Flavobacterium-sp.-ZT3R18:0.0391605487)98.6/100:0.0650967361,((WP_144071503.1-Flavobacterium-franklandianum:0.0588009776,WP_133277350.1-Flavobacterium-nackdongense:0.0813872484)33/98:0.0246227820,WP_194311471.1-Flavobacterium-soyangense:0.1067838950)84.8/97:0.0292618050)81.9/95:0.0264596232,WP_103806521.1-Flavobacterium-alvei:0.0795699098)84.7/97:0.0352122860)99.8/100:0.1358684372,(WP_140000461.1-Flavobacterium-microcysteis:0.1736203747,(((WP_194640005.1-Flavobacterium-sp.-HJJ:0.0801178141,WP_073246022.1-Flavobacterium-flevense:0.0586355398)76.7/100:0.0226154144,WP_100843428.1-Flavobacterium-sp.-5:0.0854704161)74.8/100:0.0427470419,WP_140487657.1-Flavobacterium-sp.-GSA192:0.0445338643)98.9/100:0.1403280381)92.9/100:0.0879529562)97/100:0.1069853810)84.4/70:0.0928068044,(WP_116872776.1-Terrimonas-sp.:0.2751176087,(WP_004155558.1-Microscilla-marina:0.2508619420,WP_101358295.1-Raineya-orbicola:0.1671207942)100/100:0.2422215294)51.2/70:0.0689713034)87.3/95:0.0664091739,WP_057952567.1-Salinivirga-cyanobacteriivorans:0.3973645660)28.3/54:0.0608308412)96.2/99:0.1108067756,((WP_085659180.1-Prosthecochloris-sp.-HL-130-GSB:0.2996354286,WP_038563105.1-Draconibacterium-orientale:0.2383107338)95.1/100:0.1141558453,((WP_040596643.1-Prevotella-stercorea:0.0812820149,WP_207153370.1-Prevotella-herbatica:0.0750114585)98.5/100:0.1895338920,(WP_223927321.1-Prevotella-lacticifex:0.1729951576,WP_009010825.1-Prevotella-sp.-C561:0.1358216878)100/100:0.2580233079)77.1/96:0.0774253799)95/100:0.1219651013)98.5/100:0.1470734648)69.2/92:0.0407964519)95.9/97:0.0928609990,(((((((((((WP_176021856.1-Stenotrophomonas-sp.-NA06056:0.0428211771,WP_049440708.1-Stenotrophomonas-maltophilia:0.0461155764)99.2/100:0.1542387993,WP_132999175.1-Luteimonas-arsenica:0.1572587663)56.6/100:0.0896083485,WP_078758981.1-Lysobacter-spongiicola:0.2355186402)94.3/99:0.1083945361,((WP_168608661.1-Oleiagrimonas-citrea:0.0606373526,WP_113064358.1-Oleiagrimonas-sp.-MCCC-1A03011:0.0151226381)94/100:0.0511034098,WP_043099571.1-Oleiagrimonas-soli:0.0595737797)98.2/100:0.0973974711)32.6/58:0.0382029333,((WP_233843762.1-Dyella-sp.-2HG41-7:0.0340944954,WP_188793403.1-Dyella-nitratireducens:0.0674142974)92.6/100:0.0383776525,WP_126673574.1-Dyella-dinghuensis:0.0457722587)96.8/100:0.0683196418)85.9/72:0.0321934081,(((((WP_063674027.1-Dyella-thiooxydans:0.1219939619,WP_253565187.1-Dyella-sp.-Sa:0.0632392191)84.2/98:0.0250038560,(WP_209622196.1-Frateuria-flava:0.0713345526,WP_091339503.1-Frateuria-terrea:0.0407888665)97.9/100:0.0590314224)72.7/97:0.0235066494,WP_201315287.1-Dyella-sp.-EPa41:0.1132033647)34.2/76:0.0106306831,(WP_188800319.1-Dyella-caseinilytica:0.1355712866,WP_192554051.1-Dyella-acidiphila:0.1109328239)97.6/100:0.0847600917)85.7/84:0.0281084003,((WP_049621948.1-Frateuria-defendens:0.1223652445,WP_179481164.1-unclassified-Rhodanobacter:0.0779328657)78.9/95:0.0361466233,((((((WP_214557828.1-unclassified-Rhodanobacter:0.0327088172,WP_068095172.1-unclassified-Rhodanobacter:0.0262584370)62.9/100:0.0178270837,((WP_008437080.1-Rhodanobacter-thiooxydans:0.0380893894,WP_063107523.1-Rhodanobacter-thiooxydans:0.0264283112)37/93:0.0102176676,WP_211005527.1-Rhodanobacter-sp.-B2A1Ga4:0.0332768799)86.2/93:0.0167397356)97.4/100:0.0464891677,WP_072762453.1-Rhodanobacter-sp.-OK091:0.0111299290)82.3/62:0.0126052407,((WP_077439480.1-Rhodanobacter-sp.-C01:0.0904854818,WP_077445946.1-Rhodanobacter-sp.-C05:0.0408853809)22.7/88:0.0120762747,WP_158752649.1-Dyella-sp.-S184:0.0278355517)85.5/50:0.0194492651)83.6/50:0.0186083318,WP_184416890.1-Rhodanobacter-sp.-MP7CTX1:0.0419551931)92.7/98:0.0300514997,WP_015446789.1-Rhodanobacter-denitrificans:0.0521371979)64.1/99:0.0254924439)76.6/79:0.0333879690)0/54:0.0134709883)52.5/51:0.0309968129,WP_204636097.1-Dyella-kyungheensis:0.1337069461)0/24:0.0188132003,(((WP_131151171.1-Dyella:0.1688067643,WP_214570707.1-unclassified-Dyella:0.1211806482)80.6/64:0.0165868899,((WP_130619635.1-Dyella-amyloliquefaciens:0.0695002136,WP_199044928.1-Dyella-sp.-ASV24:0.0623382925)80.8/59:0.0116850122,(WP_266169930.1-Dyella-sp.-RXD159:0.0700689232,(WP_114846719.1-Dyella-tabacisoli:0.0434126589,WP_090451016.1-Dyella-sp.-OK004:0.0634482297)84.8/99:0.0232988439)85/89:0.0119954179)73.6/64:0.0059757553)87.5/83:0.0207121310,(WP_111981026.1-Dyella-jiangningensis:0.1366522212,WP_266183414.1-Dyella-sp.-RBB1W86:0.0699685481)62.1/95:0.0296198050)17.1/89:0.0211876749)93.6/72:0.0688169715,WP_019467068.1-Dyella-japonica:0.1641840221)99.8/100:0.1859233604,(((((WP_256181981.1-Cloacibacillus-evryensis:0.1526020109,WP_087368268.1-Cloacibacillus-sp.-An23:0.1183587291)99.9/100:0.3317490031,WP_141198172.1-Persicimonas-caeni:0.3240258895)91.1/49:0.1178044642,(((WP_084578244.1-Sporomusa-malonica:0.3276217545,(WP_090940141.1-Pelosinus-propionicus:0.1856588654,WP_240986382.1-Acididesulfobacillus-acetoxydans:0.2008900396)72.6/98:0.0404397634)1.6/9:0.0410801270,WP_092486361.1-Desulfoscipio-geothermicus:0.1303096734)89/14:0.0521993941,WP_131919244.1-Heliophilum-fasciatum:0.1736273909)88.5/79:0.0533061456)71.9/28:0.0222270489,((((((WP_200343816.1-Halochromatium-glycolicum:0.1429929839,WP_201222000.1-Halochromatium-roseum:0.1070064841)99.1/100:0.1190739161,(WP_200237937.1-Thiohalocapsa-halophila:0.0543233587,WP_058558062.1-Thiohalocapsa-sp.-ML1:0.2013234053)94.6/100:0.0865547938)97.1/100:0.1128055698,(WP_182584206.1-Thiospirillum-jenense:0.2888360428,(WP_200286492.1-Rhabdochromatium-marinum:0.1139409383,WP_009150811.1-Thiorhodovibrio-sp.-970:0.0685489300)95.9/100:0.1016635311)94.6/100:0.0969206374)90.8/100:0.0677003327,((((((WP_200207675.1-Chromatium-weissei:0.1735336875,WP_200154756.1-Chromatium-okenii:0.0751651538)99/99:0.1094165482,(WP_200158060.1-Allochromatium-vinosum:0.0521226146,WP_200376120.1-Thiocystis-violacea:0.0992095976)72.5/100:0.0366834601)39/68:0.0207276590,(((WP_043749566.1-Imhoffiella-purpurea:0.0640720721,WP_007039504.1-Thiorhodococcus-drewsii:0.0533419962)88.5/100:0.0301092456,((WP_164456073.1-Thiorhodococcus-minor:0.0912087684,WP_164652045.1-Thiorhodococcus-mannitoliphagus:0.0356427911)0/25:0.0148853086,WP_200330387.1-Thiocystis-violacea:0.0743864456)96/100:0.0516150910)87.8/100:0.0202148213,(WP_201092787.1-Thiocystis-minor:0.0675832017,WP_132978621.1-Thiobaca-trueperi:0.0917628989)81.1/72:0.0158071284)85.6/72:0.0220768225)86.7/72:0.0361117425,(WP_200388819.1-Thiocapsa-imhoffii:0.1381842409,((WP_007191850.1-Thiocapsa-marina:0.0390003326,WP_232409912.1-Thiocapsa-bogorovii:0.0228754571)48.4/95:0.0123540564,WP_120796078.1-Thiocapsa-rosea:0.0199979038)98/100:0.0806052979)94.3/100:0.0695944585)94.7/100:0.0848116732,(WP_100918325.1-Candidatus-Thiodictyon-syntrophicum:0.0799853717,WP_020502667.1-Lamprocystis-purpurea:0.1009712222)94.8/100:0.0722297488)85.1/100:0.0535344057,WP_089729519.1-Candidatus-Thiosymbion-oneisti:0.2461691979)92.9/95:0.0590329135)56.9/90:0.0386147181,(WP_018175268.1-unclassified-Thioalkalivibrio:0.2775770646,WP_101758446.1-Oceanicoccus-sp.-KOV_DT_Chl:0.3526538124)18.7/31:0.0660388461)93/35:0.0847238423,WP_141730858.1-Oligoflexus-tunisiensis:0.3412692870)88.3/74:0.0513232375)88.4/47:0.0616138809,((WP_246588317.1-Pelorhabdus-rhamnosifermentans:0.1682527278,(((WP_092074275.1-Dendrosporobacter-quercicolus:0.1945965647,(WP_145090815.1-Sporomusa-sp.-KB1:0.0497515915,WP_093793074.1-Sporomusa-acidovorans:0.0593915576)98/100:0.0838898968)22.9/86:0.0392846413,((WP_054258683.1-Propionispora-sp.-2/2-37:0.1271033489,(WP_149735423.1-Propionispora-hippei:0.0653855846,WP_091744506.1-Propionispora-vibrioides:0.1258442734)68.1/98:0.0269742342)50.8/98:0.0320411965,WP_188400350.1-Sporomusa-sp.-GT1:0.2174504685)70.6/78:0.0562361893)54.9/72:0.0349630441,((WP_007952170.1-Pelosinus-fermentans:0.0927225216,WP_007939097.1-Pelosinus:0.0949700659)97.4/99:0.0639941831,(((WP_132081660.1-Anaerospora-hongkongensis:0.1100826397,WP_132083235.1-Anaerospora-hongkongensis:0.1443962520)98.5/100:0.0992172196,(WP_188399481.1-Sporomusa-sp.-GT1:0.2112983453,(WP_094603851.1-Sporomusa-silvacetica:0.1003841598,WP_144352952.1-Sporomusa-termitida:0.1206553338)95/100:0.0586883838)53.5/92:0.0287966894)53.9/44:0.0215292187,WP_054258662.1-Propionispora-sp.-2/2-37:0.1966046693)49.5/19:0.0285027425)70.5/32:0.0164497826)91.1/94:0.0505196731)92/99:0.0582842758,WP_110954047.1-Massilibacillus-massiliensis:0.1916622778)98.2/100:0.1183160539)82.9/95:0.0406761016)93.2/99:0.0756535907,((WP_207681674.1-Desulfonema-magnum:0.3001072679,WP_099324654.1-Candidatus-Kuenenia-stuttgartiensis:0.3655210363)94.6/99:0.1079272608,WP_230840800.1-Gloeobacter-morelensis:0.3008390736)94.1/95:0.1081010678)19.3/84:0.0623623409)35.5/71:0.0299188447,(((((((GJNO01008138.1-Laxus-oneistus-TSA:0.4422622404,(((((VDM95900.1-Thelazia-callipaeda:0.1586517767,(((((VDO27950.1-Onchocerca-flexuosa:0.1358672540,VDO31788.1-Brugia-timori:0.0989113074)81.6/48:0.0244860823,XP_003135693.1-Loa-loa:0.0658306307)91.6/47:0.0256524788,CAG9536712.1-Cercopithifilaria-johnstoni:0.0275299841)84.8/47:0.0149198463,VBB33646.1-Acanthocheilonema-viteae:0.0854404042)66.4/47:0.0085014223,VDK73328.1-Litomosoides-sigmodontis:0.0342236446)99.8/100:0.1461061806)100/100:0.2376374446,(VDD97390.1-Enterobius-vermicularis:0.3003932125,KHN87800.1-Toxocara-canis:0.1544634363)76.7/80:0.0509372192)41.3/77:0.0567590978,VDN56932.1-Dracunculus-medinensis:0.2437879959)94.9/99:0.0969859793,(((((KAH7695359.1-Aphelenchus-avenae:0.2653861603,((CAD5228290.1-Bursaphelenchus-okinawaensis:0.0462430390,CAD5235627.1-Bursaphelenchus-xylophilus:0.0292178519)75.4/100:0.0521237345,((KAI6206096.1-Aphelenchoides-besseyi:0.1315500506,KAI6191010.1-Aphelenchoides-bicaudatus:0.0783928472)80.8/100:0.0563954759,KAI6234710.1-Aphelenchoides-fujianensis:0.0911166910)96.6/100:0.0875392280)92.2/100:0.0930743049)92.7/99:0.0936570741,KAE9552274.1-Halicephalobus-sp.-NKZ332:0.2217754100)89.4/30:0.0735671650,TKR72632.1-Steinernema-carpocapsae:0.1725881434)37.2/11:0.0268385187,GGJS01006545.1-Plectus-murrayi-TSA:0.3079868722)11.3/11:0.0328164094,(KAI1709308.1-Ditylenchus-destructor:0.1236674542,KAI1730870.1-Ditylenchus-destructor:0.0216660858)99.9/100:0.1713925746)78.2/32:0.0617132336)92.2/95:0.0834381039,((CAI4229259.1-Auanema-sp.-JU1783:0.3144998760,(PAV65180.1-Diploscapter-pachys:0.2039033883,(CAD6189320.1-Caenorhabditis-auriculariae:0.2873855796,(CAB3400093.1-Caenorhabditis-bovis:0.1832644302,((NP_509046.1-Caenorhabditis-elegans:0.0331197857,XP_003117635.1-Caenorhabditis-remanei:0.0229126708)71.4/100:0.0350842539,CAI5454992.1-Caenorhabditis-angaria:0.0350165640)83.2/100:0.0306890743)91.9/100:0.0759758809)97.9/100:0.1224402587)34.2/68:0.0235860740)69.8/60:0.0387888650,((KHJ99538.1-Oesophagostomum-dentatum:0.0488254558,EPB80542.1-Ancylostoma-ceylanicum:0.0467269816)97/100:0.0846134705,((VDO65958.1-Haemonchus-placei:0.1234553586,VDL70484.1-Nippostrongylus-brasiliensis:0.0801337266)69.9/100:0.0209475462,VDO95824.1-Heligmosomoides-polygyrus:0.0690665909)99.5/100:0.1063798482)98.9/99:0.1135931969)38.4/39:0.0574222061)89.9/95:0.2045397117)99.9/100:0.3662135043,(WP_042281872.1-Candidatus-Protochlamydia-sp.-R18:0.1367604329,WP_068471358.1-Parachlamydia-sp.-C2:0.1008602854)100/100:0.3632733813)87.1/93:0.1363999284,(WP_008983868.1-Alishewanella-agri:0.4709085135,WP_265845290.1-Plesiomonas-shigelloides:0.4902536470)82.7/94:0.0810717428)0/24:0.0552110122,WP_161826415.1-Petrachloros-mirabilis:0.3304728309)87.9/99:0.0916002735,(WP_239493201.1-Luteitalea-sp.-TBR-22:0.2405489680,WP_058867107.1-Chloracidobacterium-thermophilum:0.2984561926)97.4/100:0.1582599050)90.9/77:0.0951238174,(((((((WP_094934028.1-Enterobacter-cloacae-complex:0.3158363904,WP_086590801.1-Vibrio-parahaemolyticus:0.1990520773)91.2/85:0.1155023904,(WP_010368546.1-Pseudoalteromonas-piscicida:0.2845681145,(WP_039473409.1-Vibrio-sinaloensis:0.2576124707,WP_016795256.1-Vibrio-cyclitrophicus:0.1545240096)81.1/93:0.0471964843)58.2/94:0.0424287811)81.6/89:0.0295089320,(((WP_195504469.1-Citrobacter-farmeri:0.3413006721,(WP_269408820.1-Pseudoalteromonas-shioyasakiensis:0.1691156022,WP_207379479.1-Shewanella-sedimentimangrovi:0.1513928722)35.2/95:0.0383570685)25.9/77:0.0126779707,(WP_061013169.1-Vibrionaceae:0.1761971312,WP_155662549.1-Aliivibrio-fischeri:0.1060763015)91.7/100:0.0556599361)80.1/83:0.0280237810,WP_171328835.1-Vibrio-splendidus:0.1902186565)90.9/93:0.0553517214)89.6/93:0.0594231331,WP_016537220.1-Cedecea-davisae:0.1230787637)91/92:0.0495902913,((WP_001100815.1-Escherichia-coli:0.2927313892,(WP_240304790.1-Vibrio-campbellii:0.2884014421,WP_004741399.1-Vibrio:0.1882913182)49.4/92:0.0660449471)87.8/88:0.0433821209,WP_064554809.1-Buttiauxella-noackiae:0.2922315762)76.7/85:0.0285461996)98.4/94:0.1168154150,WP_216735790.1-Hahella-sp.-HN01:0.1482266056)68.6/86:0.0862828491,WP_057065012.1-Citrobacter:0.3693716918)95.1/91:0.0987926533)10.3/26:0.0231631254,WP_062153084.1-Beggiatoa-leptomitoformis:0.3308837604)82.1/74:0.0529879124)69.6/76:0.0575238160,(((WP_255135177.1-Gracilimonas-sp.-CAU-1638:0.1936890406,WP_020401568.1-Gracilimonas-tropica:0.2045391533)100/100:0.3404414684,((((((((WP_171588903.1-Pelistega-europaea:0.2374696053,(WP_267333169.1-unclassified-Gilliamella:0.1256907285,((((WP_086361229.1-Gilliamella:0.0720547703,(WP_247177500.1-Escherichia-coli:0.3218828503,YP_009985672.1-Escherichia-phage-muut:0.1900986496)99.8/100:0.2168343887)69.4/38:0.0409357105,WP_267361838.1-Gilliamella-sp.-B2772:0.1222352859)60.2/26:0.0300938390,((((WP_086319877.1-Gilliamella-apis:0.0163606060,WP_198214961.1-Gilliamella-sp.-M0320:0.0500054198)93.7/99:0.0276699203,WP_039129858.1-Gilliamella:0.0202247013)74/99:0.0083133909,(WP_065737496.1-Gilliamella-apicola:0.0305257091,WP_267331332.1-unclassified-Gilliamella:0.0517138308)96.9/100:0.0473812403)92.5/100:0.0329213919,((WP_065579324.1-Gilliamella-apis:0.0673297389,WP_160432071.1-Gilliamella-sp.-Pas-s25:0.0925247724)66.7/82:0.0167966227,(WP_065587409.1-Gilliamella-apicola:0.0502753016,WP_065580031.1-Gilliamella-apicola:0.0567628119)90.4/100:0.0261466098)33.3/30:0.0105645067)90.2/94:0.0298819316)83.1/26:0.0298567039,((WP_065574597.1-Gilliamella-apicola:0.0466514974,WP_065569248.1-Gilliamella-apicola:0.0341089242)76.1/100:0.0098406638,WP_065610260.1-Gilliamella-apicola:0.0292476357)96.6/100:0.0435777426)0/26:0.0046664018)99.5/100:0.1071482394)63.1/62:0.0067199379,(WP_024553491.1-Franconibacter-helveticus:0.2391296868,(WP_040666911.1-Neisseria-wadsworthii:0.0673556600,WP_135034416.1-Neisseria:0.1414970094)81.3/97:0.0413316189)57/13:0.0057912170)96.9/46:0.0383444541,((((WP_214236207.1-Rosenbergiella-sp.-S61:0.0525465459,(WP_241647140.1-Rosenbergiella-sp.-JB07:0.0926477479,WP_214195533.1-Rosenbergiella-collisarenosi:0.0819831610)43.9/75:0.0207411498)98.4/100:0.0890262877,(((WP_032441370.1-Klebsiella-pneumoniae:0.1031872998,(WP_015583911.1-Raoultella-ornithinolytica:0.0520512403,WP_265776798.1-Klebsiella-oxytoca:0.1231223054)88.2/100:0.0435129551)95.1/99:0.0589842607,(WP_029991521.1-Tatumella-ptyseos:0.0633912826,WP_212707699.1-Tatumella-sp.-JGM130:0.0360925838)20.6/51:0.0122341293)67.1/30:0.0095962140,WP_087487466.1-Tatumella-citrea:0.0443530499)69.8/94:0.0234360627)76.2/76:0.0357142935,WP_087653467.1-Klebsiella/Raoultella-group:0.1607244230)61.4/13:0.0274082526,(WP_154751639.1-unclassified-Erwinia:0.1758226182,WP_140470594.1-Ewingella-americana:0.1863532242)74.4/34:0.0305457485)75.4/25:0.0153812868)77.2/36:0.0305112947,((((((WP_228398973.1-Insectihabitans-xujianqingii:0.0765731016,WP_074823218.1-Pragia-fontium:0.0752997171)97/100:0.0608127760,WP_130971788.1-Hafnia-alvei:0.1057695330)69.7/98:0.0250101191,WP_165428828.1-Rouxiella-badensis:0.0893835408)73.9/94:0.0063548820,WP_137713144.1-Brenneria-rubrifaciens:0.0962565019)82.8/75:0.0112175623,((WP_193937755.1-Klebsiella-grimontii:0.1444792087,(WP_142514871.1-Klebsiella-huaxiensis:0.1226989619,(WP_024360036.1-Klebsiella:0.0413064775,WP_234284225.1-Klebsiella-oxytoca:0.0681188389)82.1/100:0.0329544768)84.2/98:0.0230353665)83.1/98:0.0131554421,(((((((((WP_136167279.1-Brenneria-sp.-CFCC-11842:0.0355007981,WP_264089117.1-unclassified-Brenneria:0.0808378862)48.3/99:0.0312538709,WP_109053612.1-Brenneria-roseae:0.0742753777)87/100:0.0230688385,(((((((((WP_180742451.1-Pectobacterium-sp.-CFBP8739:0.1491174866,WP_180777316.1-Pectobacterium-aroidearum:0.0067321400)96.5/95:0.0385252671,WP_107169883.1-Pectobacterium:0.0364017429)76.1/94:0.0069865214,WP_039504461.1-Pectobacterium:0.0290371822)88/85:0.0160325893,(((((((WP_047710158.1-Pectobacterium-atrosepticum:0.0489855908,(WP_033071760.1-Pectobacterium-parmentieri:0.0259863964,WP_198482318.1-Pectobacterium-parmentieri:0.0342138185)0/42:0.0057469912)94.4/99:0.0328510994,WP_226305805.1-Pectobacterium-versatile:0.0185721247)49.7/98:0.0202313788,((WP_039280054.1-Pectobacterium-brasiliense:0.0278621591,WP_039517132.1-Pectobacterium-brasiliense:0.0281871180)88/92:0.0185031222,WP_039346077.1-Pectobacterium-fontis:0.0249152079)73.8/85:0.0071332369)90/93:0.0213199331,WP_103972204.1-Pectobacterium:0.0123759168)75.4/86:0.0060255228,WP_205558955.1-Pectobacterium-brasiliense:0.0486780555)74.6/84:0.0061751673,WP_116186106.1-Pectobacterium-aquaticum:0.0363997371)75.7/77:0.0055597383,WP_151242707.1-Pectobacterium-carotovorum:0.0302950979)86.9/95:0.0128011058)79.7/85:0.0119963542,((WP_109411993.1-Pectobacterium-polaris:0.0385933496,(WP_233968669.1-Pectobacterium-polaris:0.0477448118,WP_048259384.1-Pectobacterium-peruviense:0.0125295951)88.2/77:0.0211203554)87.7/76:0.0184382805,WP_138255615.1-Pectobacterium:0.0157410658)76.3/57:0.0088358790)0/64:0.0000022260,(WP_205946867.1-Pectobacterium:0.0489016794,WP_233976643.1-Pectobacterium-versatile:0.0182516228)82.4/96:0.0120172507)29.2/61:0.0107001188,((WP_249680869.1-Pectobacterium-polaris:0.0306676558,WP_125232508.1-Pectobacterium-versatile:0.0435258615)71.6/100:0.0063852474,WP_005974084.1-Pectobacterium-wasabiae:0.0299081259)87.2/98:0.0139389737)76.7/77:0.0118937110,WP_119888241.1-Pectobacterium:0.0479089032)65/99:0.0241158455,(WP_132452860.1-Samsonia-erythrinae:0.0988200537,WP_130637964.1-Pectobacterium-versatile:0.0429599013)0.2/64:0.0138155032)90.1/100:0.0333808164)97.1/100:0.0627697824,((WP_121576612.1-Brenneria-alni:0.0790542300,((WP_214648782.1-Pectobacterium-carotovorum:0.0306457864,((WP_261846379.1-Pectobacterium-carotovorum:0.0485852362,WP_121590317.1-Brenneria-goodwinii:0.0062063034)94.6/97:0.0237857133,(WP_182099415.1-Pectobacterium-aroidearum:0.0298837376,WP_039482612.1-Pectobacterium:0.0243380926)74/93:0.0064194113)14.8/86:0.0058689764)73.6/85:0.0060156389,WP_249228983.1-Brenneria-tiliae:0.0376348629)91.7/96:0.0272065138)71.2/78:0.0097325696,WP_062791489.1-Serratia-plymuthica:0.0743091140)79.7/65:0.0162094747)82.6/71:0.0208805907,((WP_168640580.1-Dickeya-sp.-CFBP-2040:0.0818916184,((((((WP_038918362.1-Dickeya:0.0269592697,WP_123251184.1-Dickeya-undicola:0.0397527532)63.1/100:0.0058531835,WP_022632763.1-Dickeya-solani:0.0233979810)93.4/100:0.0236094429,WP_027712673.1-Dickeya-chrysanthemi:0.0363172713)74.3/88:0.0073804265,WP_012883965.1-Dickeya-parazeae:0.0622977573)74/35:0.0065741646,WP_012770533.1-Dickeya:0.0306055904)76.6/35:0.0059233693,WP_236306746.1-Dickeya-zeae:0.0360364603)60.7/49:0.0033547299)99/100:0.0731685972,(WP_148877341.1-Serratia-marcescens:0.0469403014,WP_193158894.1-Serratia-marcescens:0.0438895788)95.4/100:0.0563562260)87.3/98:0.0320646289)13.1/36:0.0085383233,WP_095845203.1-Gibbsiella-quercinecans:0.0464682191)63.5/80:0.0156007239,WP_096000273.1-Klebsiella-pneumoniae:0.1337037756)74.8/87:0.0139117860,WP_248464278.1-Pectobacterium-versatile:0.1252562163)83.1/99:0.0257838057,(WP_112167140.1-Klebsiella:0.1075248269,WP_200519240.1-Klebsiella-michiganensis:0.0715287201)92.2/100:0.0497935033)30.1/77:0.0139621103)90.7/76:0.0200921639)74.7/27:0.0060563865,(((WP_131063651.1-Raoultella-planticola:0.0228597660,WP_163980548.1-Raoultella-ornithinolytica:0.0316212260)99.6/100:0.0653644170,WP_142430937.1-Klebsiella-oxytoca:0.0966565563)83.4/36:0.0123640924,((WP_148250181.1-Raoultella-terrigena:0.0719075146,WP_032686784.1-Raoultella-planticola:0.0354033787)93.6/99:0.0436087506,((WP_239614184.1-Raoultella-ornithinolytica:0.0731148613,WP_186921353.1-Klebsiella-quasipneumoniae:0.1228190522)82.2/96:0.0159432257,((WP_032699180.1-Raoultella:0.0455068660,WP_228399925.1-Raoultella:0.0270570058)75/100:0.0155146519,WP_004138711.1-Klebsiella:0.0134654269)94.9/100:0.0349528231)87.5/96:0.0224092202)15.7/31:0.0074547468)86.6/80:0.0180093962)85.8/82:0.0182704314)56.6/16:0.0258260712,(WP_028357569.1-Brackiella-oedipodis:0.1884489940,WP_101506252.1-Erwinia-sp.-B116:0.1073355974)83.5/27:0.0289994968)94/33:0.0527598221,(((WP_253460089.1-Erwinia-persicina:0.1904315703,(WP_246757812.1-Erwinia-phyllosphaerae:0.1341548093,WP_232759612.1-unclassified-Pantoea:0.1136443042)75.2/100:0.0463203410)58.7/68:0.0297670561,(((WP_168836617.1-Vibrio-agarilyticus:0.0960963443,(WP_076537303.1-Shewanella-sp.-UCD-KL21:0.0223633856,WP_102531614.1-Shewanella-sp.-10N.286.52.B9:0.0503161011)100/100:0.3609077026)92.9/100:0.0707068461,(WP_137671336.1-Agarivorans-sp.-Toyoura001:0.0401496845,WP_221077378.1-Agarivorans-albus:0.0457352168)95.8/100:0.0822969861)97.4/100:0.0845129360,((((((((((((WP_127021653.1-Rheinheimera-mangrovi:0.2191083556,(WP_138320019.1-Thalassotalea-litorea:0.1162752933,WP_136736185.1-Thalassotalea-mangrovi:0.1092093979)99.9/100:0.1719000469)64.6/100:0.0399498718,WP_114327629.1-Candidatus-Colwellia-aromaticivorans:0.0976446019)88.1/100:0.0399019232,WP_101108574.1-Psychromonas-sp.-psych-6C06:0.1640940220)85.1/100:0.0520456132,((WP_016794198.1-Vibrio-cyclitrophicus:0.0327694640,WP_102424249.1-Vibrio-cyclitrophicus:0.0385524920)89.1/100:0.0326072776,(WP_261903470.1-Vibrio-fortis:0.0707708962,WP_240207223.1-Vibrio-sp.-CyArs1:0.0314303523)88.3/100:0.0276685403)95.6/100:0.0766858821)84.9/89:0.0521448837,WP_150896168.1-Vibrio-fortis:0.1507366276)80.3/88:0.0584323929,(((WP_260491935.1-Pseudoalteromonas-flavipulchra:0.1593630507,WP_039494249.1-Pseudoalteromonas-flavipulchra:0.0583718155)45.5/98:0.0285561974,WP_055022295.1-Pseudoalteromonas-sp.-P1-26:0.0507799341)87.8/99:0.0381395687,(WP_149602089.1-Pseudoalteromonas-distincta:0.0377302137,WP_248518672.1-Pseudoalteromonas-sp.-2CM32C:0.0175735928)98.7/100:0.0891627924)98.9/100:0.1256455918)42.8/79:0.0458847734,(WP_199145406.1-Citrobacter-sp.-ASV29:0.2014993925,WP_148513713.1-Vibrio:0.1197328638)88.9/96:0.0621426477)85.1/93:0.0395421068,WP_067584343.1-Endozoicomonas-ascidiicola:0.1944435816)84.6/97:0.0419234614,(WP_132584255.1-Rheinheimera-sp.-D18:0.2344689651,(((((WP_131912015.1-Celerinatantimonas-diazotrophica:0.1674534511,((WP_092385961.1-Halopseudomonas-salegens:0.1496830195,WP_036859756.1-Porticoccus-hydrocarbonoclasticus:0.1893464944)91/99:0.0747511970,((WP_076722556.1-Marinobacter-lutaoensis:0.1717925914,WP_036130839.1-Marinobacter-nitratireducens:0.0955749821)24.7/82:0.0493057561,((WP_106695101.1-Marinobacter-shengliensis:0.0547017054,(WP_092030923.1-Marinobacter-sp.-DSM-26671:0.0929069040,WP_148609564.1-Marinobacter-sp.-BW6:0.0755897059)85.3/79:0.0297469051)74.2/78:0.0120164228,WP_222473125.1-Marinobacter-daepoensis:0.0499695614)96.1/99:0.0830391095)24.8/87:0.0375367029)87.4/100:0.0441446049)88.4/99:0.0358869279,(WP_027875186.1-Spongiibacter-marinus:0.1257812563,WP_204995064.1-Spongiibacter-marinus:0.0844613368)92.2/100:0.0448033567)91.4/99:0.0330289803,WP_168451636.1-Spongiibacter-thalassae:0.0413147159)75.2/99:0.0371707808,WP_062382560.1-Zhongshania-aliphaticivorans:0.0577105234)88.3/100:0.0545337255,((WP_184464382.1-Zhongshania-antarctica:0.0434238539,WP_103683243.1-Marortus-luteolus:0.0311333059)72.2/100:0.0136854449,WP_198570926.1-Spongiibacter-nanhainus:0.0636159774)32.8/68:0.0374491876)90.2/100:0.0571715195)91.2/100:0.0679572092)92.5/86:0.0501720455,((WP_145010177.1-Pseudomonas-psychrotolerans:0.0172435346,WP_077172422.1-Pseudomonas-psychrotolerans:0.0361841998)99.9/100:0.1596027848,WP_084334293.1-Pseudomonas-indica:0.0986315115)51.4/100:0.0353523674)90/86:0.0449326679,(WP_261844981.1-Amphritea-ceti:0.2151082212,((((((WP_269124033.1-Marinomonas-sp.-15G1-11:0.0257399587,WP_191602238.1-Marinomonas-algicola:0.0660245300)98.7/100:0.0997719031,WP_048693477.1-Catenovulum-maritimum:0.1477435379)90.2/100:0.0544546801,WP_228191388.1-Marinomonas-communis:0.1064716768)68.9/100:0.0330695792,WP_067216963.1-Marinomonas-gallaica:0.0528693852)17/90:0.0225648012,WP_067204681.1-Marinomonas-aquimarina:0.0300548726)88.7/95:0.0265870145,WP_055461762.1-Marinomonas-fungiae:0.0455259943)89.4/99:0.0443938060)91.7/99:0.0538271832)79/99:0.0318294577,WP_100256321.1-Reinekea-forsetii:0.3793316696)75.8/88:0.0106484225)94.9/94:0.0511213565)24.9/79:0.0290346471,(WP_146378871.1-Cronobacter-turicensis:0.0650113597,WP_032180402.1-Escherichia-coli:0.0615214873)100/100:0.1219681359)17/42:0.0291982739)65.6/20:0.0353805490,(WP_126816534.1-Enterobacter-mori:0.0987679046,WP_256879438.1-Escherichia-albertii:0.0653222267)94.9/100:0.0711417360)82.7/34:0.0397048196,WP_049841125.1-Klebsiella-sp.-RIT-PI-d:0.2055676011)99.3/100:0.1693652057)61.4/12:0.0642392201,(WP_008811140.1-Parasutterella-excrementihominis:0.3292896345,((WP_133132051.1-Legionella-sp.-W10-070:0.1978074687,((((((((((((WP_027223798.1-Legionella-pneumophila:0.0297318236,WP_265717942.1-Legionella-sp.-PATHC032:0.0338438036)94/97:0.0279641273,WP_106142999.1-Legionella-pneumophila:0.0339980684)70.2/65:0.0043819751,(WP_061513621.1-Legionella:0.0138547252,WP_027226296.1-Legionella-pneumophila:0.0500977403)98.8/91:0.0594104847)87.7/76:0.0123677007,WP_044499121.1-Legionella-pneumophila:0.0236938202)84.1/76:0.0132530868,WP_027220106.1-Legionella-pneumophila:0.0155540501)75.6/73:0.0168141733,WP_027266778.1-Legionella-pneumophila:0.0309142532)98/100:0.0749308521,(WP_058494182.1-Legionella-worsleiensis:0.1185233965,(WP_127106294.1-Legionella-sp.-km535:0.0350249916,WP_058474153.1-Legionella-quateirensis:0.0284528467)93.4/93:0.0398677118)89.7/96:0.0509030320)98.8/92:0.0728763901,WP_045097199.1-Legionella-fallonii:0.1061589046)0/37:0.0056342425,(WP_031567420.1-Legionella-wadsworthii:0.1662667754,(WP_232890831.1-unclassified-Legionella:0.0379979420,(WP_058459148.1-Legionella-bozemanae:0.0228798966,WP_058518697.1-Legionella-parisiensis:0.0446942747)89.5/100:0.0273551406)98.7/100:0.0688882702)2.2/40:0.0323500095)83.3/83:0.0330813144,WP_058513550.1-Legionella-santicrucis:0.1017584102)90.5/84:0.0296704103,WP_058463758.1-Legionella-cincinnatiensis:0.0231856856)59/77:0.0320355237,WP_128130294.1-Legionella-sainthelensi:0.0153793789)99.9/99:0.2384164570)95.5/100:0.1214740281,(WP_205000141.1-Parashewanella-hymeniacidonis:0.0609877458,WP_129946484.1-Shewanella-sp.-OPT22:0.0333166184)100/100:0.3683950214)95.7/98:0.1085234682)33/5:0.0549853548)51.4/21:0.0472879099)90.2/29:0.0729582007,WP_198243301.1-methane-oxidizing-endosymbiont-of-Gigantopelta-aegis:0.2997726665)75/66:0.1180924961)99.2/100:0.1475548808)56.8/69:0.0491889778,WP_152145882.1-Alkalilimnicola-sp.-S0819:0.2435214257)73.6/68:0.0439277678,((((((WP_125841544.1-Stutzerimonas-stutzeri:0.2432467673,((WP_262731117.1-Alcanivorax-marinus:0.1341306085,(WP_106232428.1-Halomonas-ventosae:0.0691674336,WP_228234339.1-Alcanivorax-marinus:0.1535357049)78/100:0.0360850746)79.9/99:0.0199200826,WP_175070852.1-Halomonas-taeanensis:0.0928024503)39.7/98:0.0755999734)91/100:0.0629364253,(WP_019018778.1-Halomonas-lutea:0.2108819713,WP_092999033.1-Thiohalomonas-denitrificans:0.0951224659)90.8/100:0.0961688369)87.2/100:0.0377030478,((WP_146294265.1-Euhalothece-natronophila:0.1709785554,WP_159299499.1-Spiribacter-sp.-SSL99:0.1688749224)93.6/100:0.0611294312,WP_011630054.1-Alkalilimnicola-ehrlichii:0.1539363160)91.7/100:0.0535258386)93.2/100:0.0572573912,WP_127477393.1-Sulfurivermis-fontis:0.2107110674)55/66:0.0345973329,WP_028988655.1-Thermithiobacillus-tepidarius:0.1866015972)94.2/99:0.0612745526,WP_069857109.1-Desulfoplanes-formicivorans:0.2671237804)81.5/96:0.0440818085)84.2/93:0.0693010872,(((WP_151033206.1-Cellvibrio-sp.-KY-GH-1:0.0751191151,WP_049631514.1-Cellvibrio-sp.-pealriver:0.0600868790)88.2/100:0.0630137923,WP_062059887.1-Cellvibrio-sp.-OA-2007:0.0398431877)100/100:0.2472029934,WP_156872703.1-Ketobacter-sp.-MCCC-1A13808:0.2534456908)90.4/72:0.0755735289)84.5/74:0.0519722325,(WP_015904383.1-Desulforapulum-autotrophicum:0.2166859492,WP_111960646.1-Desulfobacter-hydrogenophilus:0.2116312279)98.6/100:0.1266401374)78.6/70:0.0410206446,WP_178968040.1-Neptunomonas-phycophila:0.2802799170)18.3/75:0.0258053224,(((((((((((WP_051234278.1-Halomonas-halocynthiae:0.3599403266,WP_110648656.1-Salinicola-peritrichatus:0.2361194383)89/100:0.0996109022,WP_254581577.1-Marinobacterium-sedimentorum:0.2290648377)71.4/100:0.0569053676,WP_081193716.1-Halomonas-sp.-BC1:0.1497407026)78.4/99:0.0652622255,WP_035566812.1-Halomonas:0.1646563851)81.5/97:0.0532554055,((((((((WP_224850395.1-Marinobacter-nauticus:0.1163103364,WP_110630630.1-Salinicola-salarius:0.0678784195)73.6/100:0.0345039184,(WP_193085952.1-Halomonas-sp.-FME65:0.0853121380,(((WP_159342670.1-Halomonas-meridiana:0.2092304191,WP_253554573.1-Halomonas-sp.-DN3:0.1365574758)84.9/100:0.0320444633,WP_102589198.1-Halomonas-urumqiensis:0.0972763789)57.4/96:0.0165687085,(WP_193343673.1-Halomonas-elongata:0.0530759647,WP_064701650.1-Halomonas-caseinilytica:0.0135996931)100/100:0.1135298708)90.8/91:0.0215735726)80.5/73:0.0159954139)93.7/58:0.0283260170,WP_089675781.1-Halomonas-aquamarina:0.0303339175)78.5/57:0.0122077558,((WP_253557393.1-unclassified-Halomonas:0.0375912198,WP_089676106.1-Halomonas-aquamarina:0.0239222631)0/94:0.0000029035,WP_062362595.1-Halomonas-axialensis:0.0302229549)77.6/95:0.0155962273)74.1/58:0.0124221820,WP_027961801.1-Halomonas-halodenitrificans:0.0273309235)85.8/58:0.0162176352,WP_195687443.1-Halomonas-sp.-KAO:0.0465545673)43.9/60:0.0187288704,WP_079551663.1-Halomonas-subglaciescola:0.0431588054)96.7/99:0.0547079193,(WP_248234415.1-Marinobacter-bryozoorum:0.0804633650,(WP_228223924.1-Marinobacter-nauticus:0.0368384241,WP_098418752.1-Marinobacter-sp.-LV10MA510-1:0.1051810619)88.4/100:0.0536996061)96.5/100:0.0585465520)79.3/48:0.0387413214)78.9/48:0.0276612399,(WP_021818206.1-Halomonas-huangheensis:0.1093358884,(WP_017842145.1-Methylotuvimicrobium-buryatense:0.2567134172,(WP_149282777.1-Halomonas-sp.-Y2R2:0.0691383055,WP_073436013.1-Halomonas-cupida:0.0641276986)0/69:0.0229061177)76.4/98:0.0447752193)95.2/98:0.0582942484)94.7/96:0.0439091196,WP_127061154.1-Halomonas-nanhaiensis:0.1240453484)79.5/82:0.0153034110,(WP_096281135.1-Halomonas-sp.-JB37:0.1582047073,(WP_027958907.1-unclassified-Halomonas:0.0382665262,WP_064236373.1-unclassified-Halomonas:0.0401148509)96.3/100:0.0701391524)89.1/100:0.0396151723)55.7/81:0.0248955742,(((((((WP_158774340.1-Cobetia-sp.-L2A1:0.0578664906,(WP_144727127.1-Cobetia-crustatorum:0.0281315615,WP_088744253.1-Cobetia-sp.-QF-1:0.0532795228)43.5/100:0.0056676309)93.2/100:0.0267067069,WP_158774127.1-Cobetia-sp.-L2A1:0.0233290617)81.2/99:0.0120337035,(WP_077373671.1-Cobetia:0.0458822118,((WP_213115235.1-Cobetia-sp.-MC34:0.0348879326,WP_200017180.1-Cobetia-sp.-cqz5-12:0.0477185219)94.8/100:0.0437409059,WP_225346840.1-Cobetia-amphilecti:0.0324631150)89/100:0.0270549960)93.2/100:0.0258007845)98.7/100:0.1026276594,(WP_036548095.1-Nitrincola-lacisaponensis:0.2628230982,(WP_251978361.1-unclassified-Salinicola:0.0733019994,WP_189443129.1-Salinicola-rhizosphaerae:0.0746018565)98.9/100:0.1185963354)11.9/89:0.0284899406)88.2/82:0.0474317768,(((WP_041157945.1-Halomonas-sp.-KHS3:0.0858942126,WP_141441391.1-Halomonas-titanicae:0.0562450638)78.7/96:0.0209257919,WP_226251132.1-unclassified-Halomonas:0.0607049849)90.9/97:0.0395544611,WP_120386672.1-Halomonas-alkaliphila:0.0880952958)94.4/100:0.0604042996)64.3/81:0.0495745800,WP_228219791.1-Halomonas-meridiana:0.0952088856)97.9/96:0.0801711860,((((((WP_030071911.1-Halomonas-alkaliantarctica:0.0407880879,WP_095603285.1-Halomonas-humidisoli:0.0230844197)79.2/70:0.0172806948,WP_253485683.1-Halomonas-sp.-XH26:0.0242403432)73.5/70:0.0108891119,WP_038485954.1-Halomonas-campaniensis:0.0187472284)98.5/100:0.0615427518,WP_162218282.1-Halomonas-alkaliphila:0.0271872074)96.2/100:0.0390763779,WP_078087027.1-unclassified-Halomonas:0.0226997145)77.9/94:0.0124028547,WP_096655487.1-Halomonas-nigrificans:0.0322252576)96.6/100:0.0582183795)69.1/80:0.0369105407)94.4/88:0.0676443461,WP_179929642.1-Halomonas-salicampi:0.2544235898)90.2/87:0.0541243534,(WP_138986566.1-Neptunomonas-concharum:0.1987882672,(WP_066801876.1-Moraxella-oblonga:0.0753057764,(WP_197128702.1-Psychrobacter-sp.-NZS113:0.1539663769,WP_201616712.1-Psychrobacter-immobilis:0.1036925374)98.9/100:0.1408962431)99.2/100:0.1229570669)57.5/71:0.0525460948)93.3/83:0.0544984459)98.8/83:0.1001955441,(((WP_255857717.1-Bacterioplanoides-sp.-SCSIO-12839:0.2327384990,((((WP_262601261.1-Endozoicomonas-euniceicola:0.0831799054,WP_066015073.1-Endozoicomonas-atrinae:0.1020523514)96.6/100:0.0816874850,WP_257287521.1-Endozoicomonas-sp.-SESOKO2:0.0745881591)81.7/100:0.0559458512,(WP_067585802.1-Endozoicomonas-ascidiicola:0.1206217949,(WP_020580625.1-Endozoicomonas-elysicola:0.0910232192,WP_257293945.1-Endozoicomonas-sp.-YOMI1:0.0989899873)76.1/100:0.0571728294)99.9/100:0.1770087482)92.2/100:0.1013000650,WP_116686321.1-Pelagibaculum-spongiae:0.2278270663)93/100:0.1012076453)97.4/64:0.0986348138,((WP_220103444.1-Neiella-holothuriorum:0.0835209206,WP_087505717.1-Neiella-marina:0.1216299385)94.3/100:0.0525082518,((WP_028862352.1-Psychromonas-aquimarina:0.1481886260,WP_187408568.1-Saccharophagus-sp.-K07:0.1128460719)87.9/100:0.0446115397,WP_251261538.1-Echinimonas-agarilytica:0.1178549583)98.3/90:0.0751207508)40.6/32:0.0263717159)68.5/26:0.0432726070,(WP_248714967.1-Agarivorans-sp.-TSD2052:0.0587170965,WP_152781274.1-Agarivorans-sp.-B2Z047:0.0602456870)99.4/100:0.1083953468)90.4/26:0.0419019023)14.5/12:0.0288496450,(WP_096084688.1-Agaribacterium-haliotis:0.1746833605,WP_188151970.1-Teredinibacter-waterburyi:0.1965611759)98.9/75:0.1179776526)79.3/73:0.0228770254,((((((((((WP_167868984.1-Shewanella-sp.-WXL01:0.2936044033,WP_047049276.1-Vibrio-mexicanus:0.0992811234)90/99:0.0459325498,WP_264874461.1-Vibrio-agarivorans:0.0757750306)83.2/97:0.0150361094,WP_020335761.1-Vibrio-natriegens:0.0888521451)75.2/95:0.0169794560,((WP_022943097.1-Psychromonas-hadalis:0.1007322011,(WP_113818410.1-Psychromonas-sp.-B3M02:0.0378057333,(WP_025564776.1-Psychromonas-sp.-SP041:0.0427055537,WP_166372414.1-Psychromonas-sp.-SA13A:0.0115420730)96.6/100:0.0537968131)94.6/100:0.0436322913)88.5/90:0.0311388076,WP_047874860.1-Photobacterium-aphoticum:0.1756852349)3.8/11:0.0173288038)68.9/52:0.0221685310,(WP_050648448.1-Vibrio-sp.-J2-4:0.1085037390,(WP_140076235.1-Vibrio-parahaemolyticus:0.1065924196,WP_205390674.1-Vibrio:0.0658092705)77.8/97:0.0136208846)83.1/95:0.0186057692)83.6/85:0.0140307432,WP_164649367.1-Vibrio-astriarenae:0.0684202204)80.1/37:0.0106229349,(((WP_193630602.1-Shewanella-benthica:0.1473684177,WP_159656123.1-Vibrio-atypicus:0.0496914811)90.5/99:0.0483542725,(WP_269599241.1-Salinivibrio-kushneri:0.0559382494,WP_136483218.1-Vibrio-sp.-H11:0.0594693265)94.9/100:0.0584597791)43.6/85:0.0261002784,(WP_084884043.1-Vibrio-sp.-qd031:0.0828063968,WP_012143346.1-Shewanella-sediminis:0.1668951532)86.8/96:0.0235137642)92.6/74:0.0329962034)48.7/25:0.0111982562,((WP_222525605.1-Ferrimonas-balearica:0.1221568760,WP_013344594.1-Ferrimonas-balearica:0.0967387773)99.5/100:0.1307354795,(WP_016959527.1-Enterovibrio-norvegicus:0.0480345961,WP_229491984.1-Enterovibrio-norvegicus:0.1053892300)88.2/99:0.0514928642)91/82:0.0419552178)87.8/38:0.0208322689,WP_093326971.1-Thalassotalea-agarivorans:0.0972331733)0/22:0.0000023121,((((((WP_167678613.1-Shewanella-aestuarii:0.1129181271,WP_137222113.1-Shewanella-sp.-MEBiC00475:0.0294017432)90.4/100:0.0409883837,WP_188955407.1-Shewanella-ulleungensis:0.0446828219)98.9/100:0.0817559523,((WP_182693438.1-Pseudoalteromonas-sp.-SR45-5:0.0561947670,WP_256848444.1-Pseudoalteromonas-carrageenovora:0.0158110979)78.2/45:0.0115729266,WP_126005793.1-Pseudoalteromonas-sp.-Xi13:0.0282954097)71.2/41:0.0085485611)74.1/28:0.0283253794,WP_138577097.1-unclassified-Pseudoalteromonas:0.0844977485)61.6/26:0.0085569899,(WP_264390127.1-Pseudoalteromonas-sp.-A3:0.0216527602,(WP_096739047.1-Pseudoalteromonas-atlantica:0.0346986075,WP_248462211.1-Pseudoalteromonas-sp.-2CM37A:0.0166539327)92.9/100:0.0225781191)80.9/98:0.0167576620)97.5/100:0.0649232307,WP_155694061.1-Psychrosphaera-haliotis:0.0962321034)92.9/72:0.0342000156)97.3/91:0.0692653128)28.3/53:0.0426271956,((WP_142941196.1-Aliikangiella-marina:0.2250985779,(WP_081148784.1-Cognaticolwellia-beringensis:0.0426627336,WP_007634918.1-Paraglaciecola-psychrophila:0.0292074782)99.9/100:0.1290690324)74.3/94:0.0401841197,WP_068546663.1-Thalassotalea-crassostreae:0.1651347782)80.1/80:0.0382673323)78/86:0.0183913264,(WP_250459773.1-Microbulbifer-sp.-GX-H0434:0.1154520756,WP_152451766.1-Microbulbifer-sp.-THAF38:0.1113422513)99.2/100:0.1060830623)96.8/95:0.0702368681,((WP_216048042.1-Aliiglaciecola-lipolytica:0.1468863244,WP_057181266.1-Colwellia:0.1594496543)23/41:0.0365492209,(WP_131095783.1-Alteromonas-sp.-KUL42:0.0646436423,(((((((WP_100972994.1-Alteromonas-sp.-MB-3u-76:0.0836253968,WP_269520955.1-Alteromonas-sp.-BMJM2:0.0413863926)88.7/100:0.0236543257,(((((((WP_156078883.1-Alteromonas:0.0467014979,WP_202262575.1-unclassified-Alteromonas:0.0289799564)83.9/90:0.0138640916,WP_187447473.1-Alteromonas-sp.-BZK5:0.0174446207)75.2/58:0.0054033328,((WP_044058190.1-Alteromonas-australica:0.0348243982,WP_159580202.1-Alteromonas-sp.-KUL106:0.0173145150)87.2/98:0.0113571037,(WP_014977514.1-Alteromonas-macleodii:0.0290176945,WP_239220085.1-Alteromonas-sp.-MmMcT2-2:0.0229238831)0/85:0.0000020392)76.2/75:0.0057110769)0/34:0.0000024244,WP_202940767.1-Alteromonas-macleodii:0.0345726678)71.6/28:0.0057710687,(WP_063457834.1-Alteromonas-stellipolaris:0.0280126250,WP_057795858.1-Alteromonas-stellipolaris:0.0407906906)68.6/86:0.0057089662)87.9/87:0.0118883185,WP_061095937.1-Alteromonas-macleodii:0.0347574050)76.7/73:0.0107104119,WP_163106912.1-Alteromonas-genovensis:0.0413376819)56.3/46:0.0118986211)61.3/42:0.0049983089,(WP_071816068.1-Alteromonas-sp.-V450:0.0777228445,WP_198816918.1-Alteromonas-sp.-IB21:0.0181239867)86.9/100:0.0206406970)91.8/97:0.0252853053,WP_163085646.1-Alteromonas-profundi:0.0346210679)72.7/46:0.0058372426,(WP_130638173.1-Alteromonas-sp.-KUL49:0.0473108431,WP_115592514.1-Alteromonas-aestuariivivens:0.0409805725)81.1/65:0.0121341583)91.8/97:0.0235117656,((((WP_018984642.1-Salinimonas-chungwhensis:0.0840252681,WP_265616768.1-Alteromonas-sp.-ASW11-7:0.0544387876)85.1/94:0.0385551982,WP_216028209.1-Alteromonas-sp.-C1M14:0.0421848688)42.6/65:0.0168740951,(WP_265607972.1-Alteromonas-sp.-ASW11-130:0.0587682906,WP_119499891.1-Alteromonas-sp.-RKMC-009:0.0475661528)89.1/52:0.0234307693)76.5/62:0.0093317109,(WP_216026947.1-Aestuariibacter-sp.-A3R04:0.0508706694,(WP_070176406.1-Alteromonas-lipolytica:0.0308297409,WP_073325432.1-Marisediminitalea-aggregata:0.0429736424)91.3/100:0.0358204942)26/65:0.0175644200)91.2/98:0.0273698488)77.7/93:0.0168842548,(WP_262993134.1-Alteromonas-sp.-ASW11-19:0.0611450622,(WP_218354448.1-Alteromonas-lipotrueiana:0.0687720274,WP_195811108.1-Salinimonas-marina:0.0495623352)74.1/96:0.0362692604)77.9/97:0.0177115665)91.8/99:0.0344131172)96.2/99:0.0515745823)55/41:0.0114098782)85.9/53:0.0278858911,(WP_009839692.1-Pseudoalteromonas-tunicata:0.3761130886,(WP_124026287.1-Alteromonas-sediminis:0.1071333641,WP_100655941.1-Alteromonas-flava:0.1822888452)94.7/95:0.0850827654)86.9/58:0.0383352062)23.4/75:0.0180771535,(WP_102798355.1-Bowmanella-denitrificans:0.0506017181,WP_214344498.1-Bowmanella-yangjiangensis:0.0397285461)99.9/100:0.1343032697)94.4/99:0.0648674449,WP_186505417.1-Neptunicella-marina:0.1545974478)83.6/89:0.0289646409,(WP_160153838.1-Microbulbifer-sp.-ALW1:0.1965262155,((WP_263600466.1-Microbulbifer-agarilyticus:0.1092447545,WP_078085574.1-Microbulbifer-mangrovi:0.0738077356)34.3/86:0.0103128500,WP_029249715.1-Microbulbifer-agarilyticus:0.0909322245)91.3/87:0.0603197155)97.1/100:0.0814096070)79.8/89:0.0277757935,WP_031579865.1-Ruminobacter-sp.-RM87:0.3326514380)92.2/33:0.0675881420,(WP_118961222.1-Colwellia-sp.-RSH04:0.2214366714,(WP_189610147.1-Saccharospirillum-salsuginis:0.1932004233,WP_111640782.1-Marinimicrobium-alkaliphilum:0.2416622249)90.8/99:0.0533018574)81.1/65:0.0310161587)82.7/31:0.0365441788,(((WP_085356321.1-Neisseria-dumasiana:0.1598521231,(WP_127202078.1-Neisseria-meningitidis:0.1009531245,WP_237092367.1-Neisseria-sp.-ZJ104:0.1142139327)18.8/98:0.0366896531)39.1/64:0.0160515861,WP_101133177.1-Neisseria-meningitidis:0.0607639262)85.9/70:0.0329650794,WP_005552781.1-Aggregatibacter-actinomycetemcomitans:0.1309469556)99.8/100:0.1349793254)90.4/65:0.0509023594,(WP_138624780.1-Pseudoalteromonas-sp.-S4492:0.1080758500,WP_130050482.1-Pseudoalteromonas-sp.-MEBiC-03485:0.1586744809)86.2/94:0.0567816137)99.4/96:0.1159562616,((((((((((((((((WP_188026786.1-Tolumonas-osonensis:0.0204380799,WP_024870911.1-Tolumonas-lignilytica:0.1016595965)99.3/100:0.0903898418,(((WP_047883806.1-Photobacterium-ganghwense:0.0850772678,(WP_105076797.1-Pantoea-ananatis:0.2873620761,WP_146157089.1-Photobacterium-rosenbergii:0.1265186241)79.7/61:0.0297828809)80.8/63:0.0224749317,(WP_075764060.1-Photobacterium-proteolyticum:0.1311409454,WP_239411815.1-Photobacterium-sp.-OFAV2-7:0.0830133975)93.6/98:0.0758945868)75.1/61:0.0409286829,WP_187463805.1-Photobacterium-sp.-BZF1:0.0643038569)85.2/41:0.0401437074)95.5/73:0.0550055390,((((WP_144135068.1-Kosakonia-radicincitans:0.2099946086,((WP_088208149.1-Enterobacter-cancerogenus:0.0807247376,WP_121932958.1-Enterobacter-hormaechei:0.0816765090)15.4/79:0.0337978791,WP_023330866.1-Enterobacter:0.0465951358)94.4/100:0.0733758687)92.2/99:0.0733507420,WP_120453320.1-Kalamiella-piersonii:0.0817934961)31/60:0.0228658357,(((WP_214216788.1-Rosenbergiella-epipactidis:0.0483210159,WP_241623482.1-Rosenbergiella-epipactidis:0.0220087644)99.8/100:0.1054804311,(WP_192411534.1-Pantoea-sp.-PNT02:0.0621017566,(WP_125311258.1-Pantoea-dispersa:0.0812468195,((WP_150019269.1-unclassified-Pantoea:0.0504266209,(WP_069025179.1-Pantoea-agglomerans:0.0057544649,WP_140917215.1-Pantoea-deleyi:0.0451334002)75.7/79:0.0180689836)95.3/72:0.0278983404,WP_097116658.1-Pantoea-sp.-GL120224-02:0.0113668850)68.3/48:0.0057046528)88.6/90:0.0377670902)96.1/99:0.0601503309)90.3/100:0.0398205250,WP_187508308.1-Pantoea-agglomerans:0.1250788565)16.1/65:0.0279717907)71.9/70:0.0213808649,(WP_269294270.1-Pantoea-sp.-LMR881:0.0374286132,WP_215846277.1-Pantoea-sp.-EnvD:0.0686005170)96.7/100:0.0577864334)96.3/100:0.0654204416)89.1/53:0.0311234039,(WP_020431327.1-Vibrio-fluvialis:0.0620770809,WP_055453311.1-Vibrio-fluvialis:0.0534421254)93.9/88:0.0395905494)88.5/49:0.0289868610,(WP_117235674.1-Vibrio-maerlii:0.1130756315,((WP_103882085.1-Vibrio-hangzhouensis:0.0461347004,(WP_120356561.1-Alginatibacterium-sediminis:0.2205637990,WP_039467548.1-Photobacterium-gaetbulicola:0.0713223185)95.2/52:0.0637234985)30.8/49:0.0104399646,WP_265676699.1-Vibrio-sp.-ZSDZ65:0.1082901828)82/52:0.0177130166)89.7/52:0.0272144040)68.2/33:0.0237389641,(WP_022590465.1-Vibrio-nigripulchritudo:0.1595510585,WP_045977433.1-Vibrio-neptunius:0.1268396337)94.5/97:0.0555147928)63.8/43:0.0221109070,(((WP_200225058.1-Rahnella-aceris:0.0940712833,WP_195814554.1-Rahnella:0.0923076752)91.8/100:0.0481987324,WP_169403689.1-Rouxiella-aceris:0.0511520647)98.2/100:0.0952557643,(WP_172533652.1-Vibrio-aestuarianus:0.2244700553,WP_101064017.1-Moritella-sp.-Urea-trap-13:0.1273254808)74.4/37:0.0589694686)80.9/36:0.0475626817)88.7/20:0.0349560014,((WP_261600599.1-Photobacterium-sp.-TY1-4:0.0650980209,(WP_114765998.1-Vibrio-rhodolitus:0.0925538356,WP_182108830.1-Vibrio-marinisediminis:0.0752118143)96.8/100:0.0784689717)80/48:0.0281681398,((WP_170899585.1-Vibrio-sp.-3-2-1:0.0843152290,WP_017221010.1-Moritella-dasanensis:0.0964497579)92.4/100:0.0390747937,WP_107284330.1-Photobacterium-lipolyticum:0.0753016910)89.5/100:0.0439438810)85.4/80:0.0265861404)0/18:0.0196421862,((((WP_211647400.1-Photobacterium-sp.-GJ3:0.0472273181,WP_120511428.1-Photobacterium-salinisoli:0.0309069407)94.1/100:0.0430507820,((WP_050928752.1-Yersinia-enterocolitica:0.1755145792,WP_012883389.1-Dickeya-parazeae:0.1721406972)94.6/100:0.0631993206,((WP_035339894.1-Dickeya:0.0253482354,WP_125258312.1-Dickeya-lacustris:0.0652259063)100/100:0.1641036308,WP_042868619.1-Dickeya:0.0548676802)95.2/100:0.0661718923)94.9/100:0.0700524873)58.7/93:0.0166780682,((WP_038178862.1-Vibrio-rhizosphaerae:0.0900751281,WP_107187026.1-Photobacterium-kishitanii:0.1637929547)84.4/87:0.0481314130,(((WP_070968050.1-Vibrio-sonorensis:0.1452547792,WP_269377025.1-Vibrio-sinaloensis:0.1101848836)93.5/99:0.0499689394,WP_046218796.1-Photobacterium-halotolerans:0.1109339181)54.2/98:0.0276812997,(WP_123015137.1-Vibrio-zhugei:0.0325513345,WP_077313846.1-Vibrio-palustris:0.0267162402)99.9/100:0.1189488942)79.8/99:0.0277474727)91.4/86:0.0406464216)96.5/75:0.0518759583,(WP_124938567.1-Vibrio-viridaestus:0.1206446994,WP_088133934.1-Vibrio-gazogenes:0.1752508891)55.4/96:0.0059748729)94/69:0.0363123050)84.8/36:0.0175450063,(((((((((WP_067416136.1-Enterovibrio-coralii:0.1074503524,WP_102395415.1-Enterovibrio-norvegicus:0.0771204555)85.2/59:0.0406792020,(((WP_222763364.1-Vibrio-fluvialis:0.0842595753,(WP_102137258.1-Vibrio-cholerae:0.0768565578,(WP_172567459.1-Vibrio-fluvialis:0.0748325632,WP_199354891.1-Vibrio-cholerae:0.0369189590)96.2/100:0.0624227948)91.3/94:0.0607071566)86.5/33:0.0384304143,(WP_148560094.1-Vibrio-mimicus:0.1192266756,WP_103155729.1-Vibrio-vulnificus:0.0917260463)90.1/90:0.0378019801)86.5/19:0.0223898060,(WP_152138288.1-Plesiomonas-shigelloides:0.1069227382,((WP_148519618.1-Vibrio-cholerae:0.0231397173,(WP_167570263.1-Aeromonas-veronii:0.0524134639,WP_000864158.1-Vibrio-metoecus:0.0293420196)93.2/100:0.0251836552)74.2/100:0.0078271567,WP_148520854.1-Vibrio-cholerae:0.0353379139)92.7/100:0.0389480036)87.7/51:0.0237230382)77/19:0.0168588425)81.7/25:0.0231541162,(((WP_000864153.1-Vibrio-cholerae:0.0208232923,WP_251878143.1-Grimontia-sp.-020920N:0.1338007885)89.2/96:0.0499536028,((WP_161129723.1-Photobacterium-lucens:0.1000651525,WP_206634697.1-Vibrio-vulnificus:0.1291105515)58.3/72:0.0208715133,WP_084988554.1-Vibrio:0.1118241916)93.1/99:0.0490928367)50.2/68:0.0325730607,WP_047876260.1-Photobacterium-aphoticum:0.1529632151)23.3/30:0.0110691257)91.8/40:0.0434932902,WP_267741605.1-Vibrio-cholerae:0.0951394852)65.9/23:0.0100925415,(WP_237042700.1-Photobacterium-damselae:0.0861364656,(((WP_249572008.1-Vibrio-cholerae:0.0851404697,WP_088132129.1-Vibrio-cholerae:0.0850410684)66.4/35:0.0367302430,(WP_255122387.1-Vibrio-cholerae:0.0585632633,(WP_217542775.1-Vibrio-metschnikovii:0.0833981897,WP_113593871.1-Vibrio:0.0332134772)97.6/100:0.0647927900)34.1/77:0.0282765075)43.8/28:0.0242823546,(WP_000864156.1-Vibrio-cholerae:0.0492996248,WP_000864152.1-Vibrio-cholerae:0.0740721575)92.1/99:0.0411589732)33.3/22:0.0298514180)86.9/27:0.0224687172)91.3/86:0.0238725568,(((WP_175242188.1-Vibrio-cholerae:0.0762341343,WP_095464062.1-Vibrio-cholerae:0.0532934790)85.5/58:0.0239377338,((WP_017004373.1-Enterovibrio-norvegicus:0.1084389775,WP_011189464.1-Desulfotalea-psychrophila:0.3317796440)35.1/57:0.0305601020,WP_107253996.1-Photobacterium-indicum:0.0964322687)97.4/99:0.0637270889)44.8/56:0.0077263268,(((WP_199351799.1-Vibrio-cholerae:0.0616175132,WP_065173166.1-Photobacterium-kishitanii:0.1332159187)20.1/7:0.0241586926,WP_149560528.1-Vibrio-cholerae:0.0677805871)91.6/82:0.0537560719,((WP_000864154.1-Vibrio-cholerae:0.0679813328,WP_102786270.1-Vibrio-cholerae:0.0572051039)71.4/10:0.0139804184,WP_095478449.1-Vibrio:0.0707480657)77.1/90:0.0214216982)65.8/9:0.0143895181)85.9/72:0.0210052970)81.5/82:0.0143026132,(WP_161119826.1-Photobacterium-lucens:0.0979234284,(WP_188234612.1-Vibrio-sp.-Y2-5:0.0544005989,WP_057556016.1-Vibrio-cholerae:0.0163287452)100/98:0.1486068113)90.2/94:0.0414695204)0/35:0.0145595081,(WP_065208126.1-Photobacterium-phosphoreum:0.1062349360,((((((WP_232600401.1-Photobacterium-carnosum:0.0748235422,WP_176284769.1-Photobacterium-damselae:0.1233911633)59.7/76:0.0506616805,WP_060997345.1-Photobacterium-aquimaris:0.0945539614)9.5/52:0.0275299650,(WP_232582982.1-Photobacterium-phosphoreum:0.0409513613,((WP_256383785.1-Photobacterium-toruni:0.0393115636,WP_232578536.1-Photobacterium-carnosum:0.0457776377)0/83:0.0020821640,WP_107228404.1-Photobacterium-phosphoreum:0.0714589742)93.6/100:0.0344258305)89.1/100:0.0383806481)75.2/38:0.0211535650,(WP_232575020.1-Photobacterium-carnosum:0.0976186507,WP_232585319.1-Photobacterium-iliopiscarium:0.0730732324)53.6/95:0.0201439232)59.8/58:0.0337113313,(WP_232600381.1-Photobacterium-carnosum:0.0501103058,WP_107213932.1-Photobacterium-phosphoreum:0.1163388180)0/58:0.0024799070)87.8/80:0.0265340465,(WP_232603643.1-Photobacterium-carnosum:0.0966464648,WP_232615715.1-Photobacterium-carnosum:0.0621439345)92.8/93:0.0405383384)38.3/49:0.0136099445)96.5/87:0.0521482966)0/46:0.0092024765,((((WP_048899071.1-Photobacterium-swingsii:0.1092381966,(((WP_107293979.1-Photobacterium-phosphoreum:0.0856063911,WP_107228425.1-Photobacterium-phosphoreum:0.1044006501)76.3/83:0.0418010703,WP_107295906.1-Photobacterium-phosphoreum:0.1052018424)87.9/59:0.0366161563,((WP_045039948.1-Photobacterium-iliopiscarium:0.1010451833,WP_193290728.1-Vibrio-parahaemolyticus:0.1842278670)51.6/65:0.0436309869,((WP_065176337.1-Photobacterium-aquimaris:0.0592688389,WP_107199726.1-Photobacterium-phosphoreum:0.0676736884)27/96:0.0172442971,WP_023931546.1-Photobacterium-leiognathi:0.1116908055)85.9/99:0.0501198008)80.6/33:0.0283895912)28.1/30:0.0274328049)81.6/27:0.0274320853,(((WP_045045404.1-Photobacterium-kishitanii:0.0453098108,WP_107264793.1-Photobacterium-phosphoreum:0.0235527474)89.5/97:0.0246375715,WP_045030671.1-Photobacterium-phosphoreum:0.0171919405)95.3/99:0.0452552833,(WP_087853896.1-Photobacterium-andalusiense:0.0307502425,WP_107274737.1-Photobacterium-phosphoreum:0.0877867992)98.7/100:0.0603802062)86.1/66:0.0245122389)75.6/26:0.0130947176,WP_065194916.1-Photobacterium-phosphoreum:0.1308203193)87.9/29:0.0250433674,((((WP_237041393.1-Photobacterium-damselae:0.0780326255,WP_080158029.1-Photobacterium-piscicola:0.1422564174)96.6/98:0.0731976643,(((WP_238087945.1-Photobacterium-damselae:0.0501386101,(WP_065172005.1-Photobacterium-damselae:0.0666256943,WP_107185088.1-Photobacterium-leiognathi:0.0607231400)73.8/96:0.0201047066)78.6/96:0.0221374961,WP_151170663.1-Photobacterium-damselae:0.0323616543)74.2/94:0.0250493998,WP_045063675.1-Photobacterium-leiognathi:0.0661343908)81.8/92:0.0200506938)74.6/83:0.0101454720,((WP_151182432.1-Photobacterium-damselae:0.1002561335,(WP_106261939.1-Photobacterium-damselae:0.0695644007,WP_115426223.1-Photobacterium-damselae:0.0538148202)42/88:0.0149021616)41.1/86:0.0189134037,(WP_053987023.1-Photobacterium-leiognathi:0.0872634019,WP_086367281.1-Photobacterium-damselae:0.0473893140)82.8/79:0.0171687872)90.6/86:0.0277477958)84.6/88:0.0237693625,WP_237041371.1-Photobacterium-damselae:0.0697256057)52.4/89:0.0237544482)77.4/27:0.0304089963)91/34:0.0303639819)79.4/18:0.0106544766,((((((WP_190336189.1-Vibrio-sp.-S9_S30:0.0722091791,WP_022612811.1-Vibrio-nigripulchritudo:0.0775859888)95.5/100:0.0402735648,((WP_075989065.1-Vibrio-fluvialis:0.0376669152,WP_044366945.1-Vibrio-fluvialis:0.0233718738)88.2/100:0.0381664965,(WP_076657081.1-unclassified-Vibrio:0.0546175329,WP_239929047.1-Vibrio-kanaloae:0.0319335820)98/100:0.0707478093)99.5/100:0.0922819596)85/99:0.0281232757,WP_006646466.1-Photobacterium-sp.-SKA34:0.1382713006)82.3/33:0.0195174667,(WP_224657569.1-Vibrio-vulnificus:0.1201961507,((((WP_261298106.1-Shewanella-sp.-SP1S2-7:0.0802653113,WP_167678467.1-Shewanella-aestuarii:0.0735563794)99.8/100:0.1541636201,WP_248994245.1-Shewanella-gaetbuli:0.1960027633)99.4/99:0.1510837149,(((WP_069589493.1-Salinivibrio-sp.-DV:0.0832963757,WP_077662143.1-Salinivibrio-proteolyticus:0.0139331169)37.9/76:0.0235219696,((WP_077659434.1-Salinivibrio-kushneri:0.1019944121,WP_077659452.1-Salinivibrio-kushneri:0.0264219604)83/100:0.0225111810,WP_269578450.1-Salinivibrio-kushneri:0.0708645784)88.5/100:0.0359653235)99.8/100:0.1644634608,WP_136989485.1-Vibrio-sp.-F13:0.2343274072)5.4/43:0.0413352895)56.3/27:0.0323179564,(WP_212599444.1-Shewanella-sp.-M16:0.1775530153,WP_220058598.1-Shewanella-sp.-FJAT-53555:0.1723743952)92.7/57:0.0625986925)77.5/42:0.0455054324)83.3/36:0.0254314009)75.1/16:0.0128088303,(((WP_192015074.1-Photobacterium-arenosum:0.0675528881,WP_088876367.1-Vibrio-mediterranei:0.1624845145)88.8/100:0.0361671943,(WP_028024862.1-Enterovibrio-calviensis:0.1417496027,WP_017007824.1-Enterovibrio-norvegicus:0.1389559254)83.9/99:0.0307823956)92.2/99:0.0587912936,((WP_069212356.1-Vibrio-cholerae:0.0597070841,WP_202641945.1-Vibrio-fluvialis:0.0762511718)97.1/100:0.0890487005,WP_068898547.1-Enterovibrio-pacificus:0.1311524069)68.4/29:0.0118112893)87.9/23:0.0350719344)0/18:0.0110655368,(((((WP_102986644.1-Vibrio-diazotrophicus:0.1160387950,WP_045460498.1-Vibrio-hyugaensis:0.1144546390)90.5/50:0.0371944096,(WP_268679810.1-Vibrio-barjaei:0.0730282153,(WP_013365656.1-_Enterobacter:0.1270406104,(((WP_258988101.1-Scandinavium-hiltneri:0.0623496529,WP_181620050.1-Enterobacter-sp.-RHBSTW-00994:0.0516803786)95.6/100:0.0550460344,WP_193129600.1-Klebsiella-aerogenes:0.0686942048)95.4/100:0.0560569448,WP_214579582.1-Enterobacter-asburiae:0.0701580904)93.1/100:0.0536271120)100/100:0.1344360653)87/85:0.0342191127)73/35:0.0280025559,(((WP_094127815.1-Vibrio-sp.-V09_P4A23P171:0.0969785000,WP_199867560.1-Vibrio-anguillarum:0.0523837024)97.9/99:0.0711206791,WP_045898906.1-Enterobacteriaceae:0.0961443799)82.9/95:0.0211129532,((WP_182744595.1-Shewanella-sp.-SR44-4:0.1580062530,WP_152797327.1-Salinivibrio-sp.-VYel1:0.1339574503)90.8/100:0.0356921125,WP_039484180.1-Vibrio-sinaloensis:0.1213002482)94.5/100:0.0534650851)86.5/64:0.0219457212)69.3/36:0.0134552369,(((((WP_208891742.1-Vibrio-anguillarum:0.1015502010,WP_230432956.1-Plesiomonas-shigelloides:0.0921655440)16.6/56:0.0287718886,WP_148533341.1-Vibrio-cholerae:0.0614207421)70.4/94:0.0171987525,WP_107294203.1-Photobacterium-sp.-GB-50:0.1350075635)91.2/47:0.0374414080,(((WP_112404745.1-Vibrio-diazotrophicus:0.0518406413,WP_202661182.1-Vibrio-fluvialis:0.0374127060)94.6/100:0.0489157152,WP_238097025.1-Vibrio-cincinnatiensis:0.1480854402)0/71:0.0000027387,(((WP_263100201.1-Kosakonia-sp.-ML.JS2a:0.0254244378,WP_200550470.1-Kosakonia-sp.-LAM2021:0.0774775097)99.6/100:0.1045273071,WP_094109047.1-Lonsdalea-iberica:0.1016008538)17.1/98:0.0163549413,(WP_026742149.1-Lonsdalea-quercina:0.0170479326,WP_094117837.1-Lonsdalea-britannica:0.0416377604)99/100:0.0654602827)98.7/100:0.0759055025)97/95:0.0495483179)44.1/44:0.0069472856,(WP_107294031.1-Photobacterium-phosphoreum:0.0390081911,WP_107276976.1-Photobacterium-sp.-NCIMB-13483:0.0477927808)99.7/100:0.0993202302)89.8/21:0.0228328794)13.3/25:0.0064452539,WP_211689658.1-Moritella-sp.-28:0.2404384588)56.6/50:0.0380250028)86.3/23:0.0154337439)86.8/35:0.0123369546,((((((WP_011771697.1-Psychromonas-ingrahamii:0.0939968539,WP_182690147.1-Psychromonas-sp.-SR45-3:0.0963399176)91.3/92:0.0691268604,WP_161680902.1-Proteus:0.2369530632)71.3/74:0.0449926772,WP_014230611.1-Vibrio-sp.-EJY3:0.1100360824)59/30:0.0068281309,(((((WP_193255073.1-Vibrio-navarrensis:0.0344145452,WP_258469755.1-Vibrio-alginolyticus:0.0839606442)98.2/96:0.0518654725,((((WP_172851426.1-Vibrio-coralliilyticus:0.1202546107,WP_025523961.1-Vibrio-parahaemolyticus:0.1343994241)51.9/66:0.0306172029,(WP_213886549.1-Vibrio-alginolyticus:0.1262172790,WP_158173598.1-Vibrio-alginolyticus:0.0879619480)81.8/100:0.0344280415)89.9/66:0.0499720563,(WP_044365158.1-Vibrio-fluvialis:0.2079259887,WP_138522257.1-Pseudoalteromonas-galatheae:0.1649413219)22.3/53:0.0400467215)89.2/99:0.0375025729,(WP_220051788.1-Shewanella-sp.-FJAT-51860:0.1102696343,WP_029788868.1-Vibrio:0.1554880945)89.6/97:0.0378523310)70.8/56:0.0259497961)45/46:0.0218640719,WP_261874894.1-Vibrio-rarus:0.1310649473)87.8/68:0.0295246983,(((((WP_155660100.1-Aliivibrio-fischeri:0.0608783468,WP_095457151.1-Vibrio:0.0818274341)88.5/100:0.0360092274,(WP_252020469.1-Vibrio-sp.-SCSIO-43136:0.0796305872,WP_243584106.1-Vibrio-splendidus:0.1397056870)36.3/95:0.0396783776)95.8/100:0.0553861211,(WP_166696422.1-Providencia-rettgeri:0.1000023365,WP_088905011.1-Shewanella-marisflavi:0.1490619437)69.3/99:0.0350227586)91.9/99:0.0386798892,(WP_095430409.1-Vibrio-vulnificus:0.0484836964,WP_114994956.1-Grimontia-hollisae:0.1612044202)97.6/100:0.0799616828)66/82:0.0258334806,(WP_045031875.1-Photobacterium-phosphoreum:0.0222297867,WP_107213458.1-Photobacterium-phosphoreum:0.0301399008)99.9/100:0.0935939839)81.1/68:0.0196247943)89.5/99:0.0286962756,((((WP_239941760.1-Vibrio-mediterranei:0.0841441417,(((WP_252012797.1-Vibrio-sp.-SCSIO-43140:0.0684232666,WP_234495031.1-Vibrio-maritimus:0.1530155997)76.2/97:0.0324529018,(WP_102532726.1-Vibrio-sp.-10N.286.48.B7:0.0789602636,WP_250617847.1-Vibrio-sp.-S4B1:0.0432036061)94.2/94:0.0469926106)55.1/82:0.0315822298,WP_252012765.1-Vibrio-sp.-SCSIO-43140:0.1052526867)83.5/83:0.0205375851)92.1/64:0.0427634000,(WP_062462511.1-Vibrio-mediterranei:0.0633372139,WP_171392669.1-Vibrio:0.1032210144)32.3/46:0.0182001951)82/69:0.0227870912,WP_221070344.1-Vibrio-alfacsensis:0.1051063056)58.4/75:0.0282808062,WP_102533425.1-Vibrio-breoganii:0.1054595559)92.3/95:0.0384980240)89.2/89:0.0223816226)40.3/26:0.0069035835,((((WP_102457123.1-Vibrio-breoganii:0.0915761786,(((WP_238813939.1-Vibrio-vulnificus:0.0636376830,WP_182742106.1-Shewanella-sp.-SG44-6:0.2111138916)32.5/69:0.0349021403,WP_102329601.1-Vibrio-breoganii:0.1715301243)27.4/36:0.0154224027,(WP_172380549.1-Vibrio-sp.-Vb339:0.0867539188,WP_102443327.1-Vibrio-breoganii:0.0729437302)96.3/98:0.0596040148)73.8/30:0.0090620351)71/45:0.0188758998,(((((WP_045569365.1-Vibrio:0.0670699750,WP_039437343.1-Vibrio-navarrensis:0.0385543178)98.4/100:0.0752356398,WP_176299354.1-Vibrio-sp.-05-20-BW147:0.1119820843)88/100:0.0363506051,WP_140233673.1-Shewanella-polaris:0.1743078544)97.8/95:0.0721655427,WP_108186198.1-Vibrio-splendidus:0.1394183169)34.9/34:0.0190456418,((WP_102941572.1-Vibrio-diazotrophicus:0.1118384028,(WP_102455158.1-Vibrio-breoganii:0.1187367750,WP_166378370.1-Pseudoalteromonas-sp.-Z9A4:0.1270972098)82.9/100:0.0404605532)83.9/100:0.0308085291,WP_005395829.1-Vibrio-diabolicus:0.1541153681)77.5/92:0.0322852904)64.2/62:0.0231844770)38.7/74:0.0227975439,WP_077647682.1-Salinivibrio-sp.-IB643:0.0955893102)85/70:0.0276476287,((((WP_017027407.1-Vibrio-breoganii:0.0691804881,WP_102532836.1-Vibrio-sp.-10N.286.48.B7:0.1149509369)45.2/21:0.0271099416,((WP_108083222.1-Vibrio-sp.-ZF-223:0.1076031539,WP_254101081.1-Alteromonas-sp.-LMIT007:0.1426763030)60.5/15:0.0151293523,((WP_108189654.1-Vibrio-splendidus:0.1044742273,(((WP_239850080.1-Shewanella-sp.-Isolate8:0.0615392409,((WP_011716450.1-Shewanella-sp.-ANA-3:0.1867672927,WP_206625521.1-Vibrio-vulnificus:0.0781265956)72.3/88:0.0537828386,WP_211678393.1-Moritella-sp.-36:0.2067054174)83.6/31:0.0318968113)73.3/25:0.0119736347,WP_121852788.1-Parashewanella-spongiae:0.1605060695)72.7/26:0.0185666855,WP_069440067.1-Pseudoalteromonas-tetraodonis:0.2132378602)94.7/85:0.0499261559)81.4/41:0.0246415070,(WP_102448490.1-Vibrio-splendidus:0.0831425555,WP_239872404.1-Shewanella-insulae:0.0908583571)54.7/68:0.0282578841)89.3/59:0.0292436486)82.2/15:0.0216667932)87.6/66:0.0240442703,((((WP_061023752.1-Vibrio-splendidus:0.0451356516,(WP_220768829.1-Shewanella-sp.-MBTL60-007:0.0811489480,(WP_110456309.1-Shewanella-algidipiscicola:0.1441207110,WP_204999274.1-Parashewanella-hymeniacidonis:0.1029748464)91.5/96:0.0643079025)29.8/61:0.0221623866)92.4/61:0.0388898021,(WP_182754472.1-Shewanella-sp.-SG41-3:0.1031233494,WP_192022372.1-Shewanella-sp.-WPAGA9:0.1731913826)27/46:0.0253608871)54.2/45:0.0292291622,(WP_248991952.1-Shewanella-profunda:0.1031025088,WP_059744187.1-Shewanella-frigidimarina:0.1398146037)76/45:0.0320158304)89/72:0.0301659740,(WP_011638275.1-Shewanella-frigidimarina:0.1074801816,((WP_197173106.1-Pseudoalteromonas-sp.-NZS11_1:0.1131684897,WP_182748079.1-Pseudoalteromonas-sp.-SR41-1:0.1413693426)94.3/100:0.0604059817,(WP_220774251.1-Shewanella-glacialipiscicola:0.0500958247,WP_182751750.1-Shewanella-sp.-SG44-2:0.0969868754)94.5/100:0.0618922884)50.5/94:0.0223039325)90.6/98:0.0284871768)77.6/62:0.0084202222)39.8/33:0.0111579741,(WP_193154481.1-Vibrio-navarrensis:0.1272304627,WP_130314193.1-Vibrio-vulnificus:0.1201022091)97.9/96:0.0786678202)76.5/72:0.0103238426)87.9/85:0.0193337381)91.9/86:0.0267320593,(((((((((WP_046223474.1-Vibrio-sp.-ECSMB14106:0.0583035046,WP_102562373.1-Vibrio-breoganii:0.1143670939)87.6/68:0.0381217577,((WP_223418564.1-Shewanella-inventionis:0.0731540665,WP_011497090.1-Shewanella-denitrificans:0.1667163426)95.2/85:0.0615279815,(WP_172855536.1-Vibrio-coralliilyticus:0.1622656516,WP_234235026.1-Shewanella-sp.-AS16:0.1454654848)84.8/88:0.0355799587)80.6/80:0.0326249868)89.1/64:0.0327452161,((WP_102546929.1-Vibrio-breoganii:0.1143261764,(WP_102461639.1-Vibrio-breoganii:0.0727922376,WP_102446429.1-Vibrio-breoganii:0.1341286650)89.1/96:0.0484166757)37.1/78:0.0308290796,WP_102282524.1-Vibrio-cyclitrophicus:0.1341644635)79.7/67:0.0188205630)88.1/66:0.0341332555,((WP_135455667.1-Vibrio-echinoideorum:0.0565086314,WP_218382835.1-Photobacterium-sp.-WH24:0.1470966835)82.3/87:0.0359389067,(WP_261835819.1-Vibrio-ishigakensis:0.1129102628,(WP_141346804.1-Vibrio-inusitatus:0.1228497768,WP_017221048.1-Moritella-dasanensis:0.1115935535)0/7:0.0285420021)80.4/92:0.0489286437)87.2/79:0.0378619325)71.5/90:0.0124521132,WP_258489575.1-Vibrio-alginolyticus:0.1068510532)51.6/28:0.0307731777,(WP_102481006.1-Vibrio-breoganii:0.0826161041,(WP_017031451.1-Vibrio-breoganii:0.0404833199,WP_248975945.1-Shewanella-kaireitica:0.1369373387)97.5/100:0.0695924428)82.3/48:0.0167352808)84.7/33:0.0272217315,(WP_045069930.1-Photobacterium-leiognathi:0.1296584987,(((WP_261923504.1-Shewanella-sp.-NFH-SH190041:0.1593675244,WP_149593442.1-Pseudoalteromonas-sp.-FUC4:0.1272267702)84.4/92:0.0357136579,(WP_041156219.1-Vibrio-mytili:0.0981470299,(WP_047480300.1-Vibrio-campbellii:0.0457204882,WP_045494469.1-Vibrio-hyugaensis:0.0380088736)99.7/100:0.1156705904)98.4/100:0.0938638743)27.7/90:0.0262691537,WP_102523418.1-Vibrio-tapetis:0.1322603916)91.7/97:0.0403796865)35.5/27:0.0146979975)72.3/95:0.0325887527,WP_129120576.1-Veronia-nyctiphanis:0.1419651189)29/89:0.0143366583,(((WP_017449116.1-Vibrio-parahaemolyticus:0.0975569034,WP_203433235.1-Shewanella-sp.-LZH-2:0.1687273099)79.9/95:0.0229470089,WP_238819535.1-Vibrio-furnissii:0.0898623562)74/71:0.0079667478,(((((WP_017020702.1-Aliivibrio-logei:0.1113239824,WP_194091044.1-Vibrio-hibernica:0.1591377156)80.8/100:0.0475358880,WP_016784063.1-Vibrio:0.0769634858)48.3/98:0.0280876869,WP_140342839.1-Vibrio-parahaemolyticus:0.1523162866)81/93:0.0186974462,((((WP_243584123.1-Vibrio-splendidus:0.0692660169,WP_259574191.1-Shewanella-baltica:0.1914968325)85.6/98:0.0352523361,(WP_163929979.1-Paraferrimonas-sp.-SM1919:0.0787041684,WP_087845049.1-Photobacterium-malacitanum:0.1150013791)92.1/100:0.0486371291)16/68:0.0213980246,(WP_102462022.1-Vibrio-splendidus:0.0563135704,WP_187463684.1-Photobacterium-sp.-BZF1:0.0397597709)98/100:0.0751551722)94.1/72:0.0545818462,((WP_218529053.1-Shewanella-sp.-NIFS-20-20:0.1107577411,WP_225483056.1-Vibrio-vulnificus:0.0548079538)95.4/100:0.0641828348,WP_012549058.1-Aliivibrio-salmonicida:0.1558500602)75.8/97:0.0168464866)6.8/10:0.0099794543)39.3/10:0.0115636158,(WP_165312521.1-Vibrio-ziniensis:0.0906182174,(WP_206974914.1-Vibrio-parahaemolyticus:0.0736420295,WP_058428281.1-Pseudoalteromonas:0.1613017312)86/98:0.0397060937)89.2/65:0.0460167363)89.1/91:0.0281602557)89.4/89:0.0266633018)95.2/88:0.0383950501)90.6/68:0.0251261624)73.9/13:0.0054534802,(((((WP_038175370.1-Vibrio-pacinii:0.1023742920,WP_180567593.1-Oceanospirillales:0.1230534924)74.2/82:0.0124257304,WP_159197271.1-Escherichia-coli:0.0746240804)84.7/83:0.0281332531,WP_261894433.1-Vibrio-porteresiae:0.1502991448)79/74:0.0210457589,((WP_020332409.1-Vibrio-fluvialis:0.0471300949,WP_096629284.1-Salinivibrio-sp.-YCSC6:0.1275576556)55.8/99:0.0398238390,WP_171749982.1-Vibrio-sp.-RE88:0.1314245155)78/75:0.0270244554)60.4/95:0.0186999155,((WP_017008437.1-Enterovibrio-norvegicus:0.0971566285,WP_251878087.1-Grimontia-sp.-020920N:0.0994644785)43.7/99:0.0202392568,(WP_188008910.1-Grimontia-hollisae:0.0978529289,(WP_138589904.1-Pseudoalteromonas-aurantia:0.1221612245,WP_065173140.1-Photobacterium-kishitanii:0.1024000746)67.4/95:0.0210087134)96.8/100:0.0688632735)89.9/98:0.0314051492)92.2/85:0.0328165981)36.2/10:0.0060244404,(((((((WP_268674123.1-Vibrio-scophthalmi:0.0705380781,WP_102475754.1-Shewanella-sp.-10N.286.52.C2:0.1368996451)96.6/98:0.0572741779,((WP_076646094.1-Vibrio-sp.-10N.222.47.A9:0.1095647414,(WP_102385704.1-Vibrio-cyclitrophicus:0.0852947193,WP_239937663.1-Vibrio-chagasii:0.1130898911)86.4/99:0.0611532570)34.3/32:0.0181168106,WP_135445143.1-Vibrio-tasmaniensis:0.1345683297)22.4/33:0.0175084347)72/45:0.0164736178,(WP_137027437.1-Vibrio-kanaloae:0.0281812838,WP_108117598.1-Vibrio-splendidus:0.0546398644)95.7/100:0.0440573153)80.1/42:0.0169089641,WP_135455718.1-Vibrio-echinoideorum:0.0901654218)89.8/33:0.0357201940,(((WP_161672404.1-Vibrio-toranzoniae:0.0637812215,WP_017040654.1-Vibrio-genomosp.-F10:0.1269178732)94.5/97:0.0400087191,((WP_065600047.1-Aliivibrio-sp.-1S128:0.0715204694,WP_155682326.1-Aliivibrio-fischeri:0.0725977291)46.1/90:0.0207669335,(WP_102482947.1-Vibrio-splendidus:0.0564358111,WP_102394203.1-Vibrio-sp.-10N.222.54.B6:0.1023899410)84.6/74:0.0381981951)91.6/48:0.0315562484)85.1/32:0.0166728630,((WP_161753058.1-Vibrio-toranzoniae:0.0983054612,WP_102554090.1-Vibrio-splendidus:0.0943090277)56.7/99:0.0347916873,(WP_102298256.1-Vibrio-cyclitrophicus:0.0682496339,WP_017085924.1-Vibrio-splendidus:0.1080707779)75.3/53:0.0287490043)89/52:0.0238609027)55.2/22:0.0244044606)0/23:0.0000020772,((((((WP_059021773.1-Vibrio-coralliirubri:0.0433803243,WP_135380937.1-Vibrio-tasmaniensis:0.0755884754)92.1/100:0.0315703880,(WP_238130349.1-Vibrio-cincinnatiensis:0.1305852521,WP_102276040.1-Vibrio-lentus:0.0672476251)85.4/76:0.0348277138)74.1/40:0.0194970771,((WP_005597034.1-Vibrio-scophthalmi:0.1087290971,(WP_108176621.1-Vibrio-splendidus:0.0435606130,WP_239918548.1-Vibrio-sp.-Isolate22:0.0264595939)97.1/100:0.0544054205)84.8/99:0.0294764661,((WP_102459739.1-Vibrio-lentus:0.0630904931,WP_102363333.1-Vibrio-cyclitrophicus:0.1206759360)73.1/98:0.0353902740,WP_248388778.1-Vibrio-cyclitrophicus:0.1016006175)72.5/48:0.0276793921)84.6/87:0.0276747407)83.9/82:0.0282075236,((((((WP_155682312.1-Aliivibrio-fischeri:0.0592864942,(WP_012532661.1-Aliivibrio-fischeri:0.0962183756,WP_155677763.1-Aliivibrio-fischeri:0.1062470623)46.5/85:0.0271404252)44/86:0.0236116291,(((((((WP_139685731.1-Vibrio-tasmaniensis:0.0928081155,WP_248386338.1-Vibrio-sp.-ED004:0.1485164966)83.7/96:0.0339443234,(WP_248368760.1-Vibrio-sp.-1CM8B:0.0943685984,(WP_046209502.1-Vibrio:0.0880962440,WP_061039794.1-Vibrio-coralliirubri:0.1009895012)91.1/100:0.0523826488)83.8/95:0.0327134116)39.2/29:0.0110023217,WP_102457030.1-Vibrio-cyclitrophicus:0.1020173248)92/95:0.0383028846,(WP_108162474.1-Vibrio-splendidus:0.0272144495,WP_004742905.1-Vibrio-tubiashii:0.0465857249)98.7/100:0.0722267837)81.5/28:0.0183603398,(WP_016768872.1-Vibrio-cyclitrophicus:0.0906801184,WP_017069285.1-Vibrio-crassostreae:0.0711869654)90/19:0.0322510119)23.5/16:0.0210271599,((WP_102503160.1-Vibrio-splendidus:0.1300382906,WP_132940416.1-Vibrio-crassostreae:0.0963591721)28.5/18:0.0364165187,(WP_102420589.1-Vibrio-cyclitrophicus:0.0717950975,WP_101201417.1-Paraglaciecola-sp.-MB-3u-78:0.1712339109)36.7/47:0.0171048868)89.9/23:0.0450272924)90.6/68:0.0415727727,(WP_258428294.1-Vibrio-sp.-RM-69-4:0.0773575294,WP_102428030.1-Vibrio-sp.-10N.261.54.E10:0.1164460195)84.4/58:0.0329031301)65/72:0.0229272594)88.8/73:0.0298497238,((WP_130049361.1-Aliivibrio-finisterrensis:0.0553437971,WP_019826652.1-Vibrio-splendidus:0.1106577785)69/97:0.0238758265,WP_260256886.1-Vibrio-sp.-NC2:0.0899671043)81.6/97:0.0179984636)15.7/56:0.0139393908,(WP_193834777.1-Vibrio-sp.-OPT20:0.0396433514,(WP_029865567.1-Vibrio-parahaemolyticus:0.0544383187,WP_213967907.1-Vibrio-alginolyticus:0.0318109481)98.1/100:0.0742201776)97.7/100:0.0705044991)87.2/93:0.0330163812,(((((((((WP_162023628.1-Shewanella-sp.-Arc9-LZ:0.0568004168,WP_248997855.1-Shewanella-marinintestina:0.0468594603)95/100:0.0503295912,((WP_009847952.1-Vibrio:0.1405075199,WP_233894267.1-Vibrio-kanaloae:0.1037205250)42.6/99:0.0359525454,(WP_048661352.1-Vibrio-crassostreae:0.0621979985,WP_114634227.1-Vibrio-splendidus:0.0553819819)95/100:0.0674029619)90.3/100:0.0565854211)86.1/99:0.0365413162,WP_017094106.1-Vibrio-splendidus:0.0619421518)89.3/99:0.0259582539,WP_065296243.1-Vibrio-natriegens:0.0972730165)84/83:0.0225953904,(WP_009843931.1-Vibrio-sp.-AND4:0.0650514882,(WP_102353859.1-Vibrio-cyclitrophicus:0.0970249801,WP_171315631.1-Vibrio-splendidus:0.0909853213)91.4/100:0.0488694187)23.4/51:0.0076050762)34.4/75:0.0214620902,WP_102285079.1-Vibrio-lentus:0.1113607763)80.3/91:0.0229787408,WP_004733311.1-Vibrio-splendidus:0.0791445781)79.7/96:0.0190264685,WP_017063118.1-Vibrio:0.0920617473)76.4/94:0.0127625363,WP_265133188.1-Vibrio-splendidus:0.0887794145)83.6/98:0.0185905500)68.3/86:0.0108230883,WP_182796407.1-Vibrio-sp.-SG41-7:0.1461273484)79.4/47:0.0146541164)85.1/42:0.0245539224,(((WP_063656223.1-Aliivibrio-fischeri:0.0754590098,WP_155675904.1-Aliivibrio-fischeri:0.0863821085)84.8/90:0.0268686151,(WP_137008959.1-Vibrio-kanaloae:0.0970115945,WP_135353160.1-Aliivibrio-fischeri:0.0613694719)92.7/100:0.0558271942)85.8/35:0.0224212682,(WP_017105169.1-Vibrio:0.1183930526,(WP_137025093.1-Vibrio-kanaloae:0.1044686132,WP_077679840.1-Vibrio-kanaloae:0.0527297994)77.2/100:0.0425499410)58.3/33:0.0203999086)74.3/46:0.0232877167)60.6/31:0.0193043933,(WP_063660043.1-Aliivibrio-fischeri:0.0676143232,WP_243579549.1-Vibrio-splendidus:0.1467070738)96.4/96:0.0663570312)88.4/37:0.0334330961)76.3/42:0.0066370062,((WP_171343915.1-Vibrio-cyclitrophicus:0.0389057529,(WP_248638627.1-Vibrio-crassostreae:0.1049816059,(WP_102248986.1-Vibrio-tasmaniensis:0.1361764204,WP_135445135.1-Vibrio-tasmaniensis:0.0867822084)98.4/100:0.0831074680)88.2/54:0.0282135081)87.2/52:0.0361259082,(WP_048661380.1-Vibrio-crassostreae:0.0604200657,WP_193846925.1-Vibrio-sp.-OPT10:0.0710765609)92.3/97:0.0373910774)89.7/44:0.0321422031)85.9/67:0.0220897180)92.6/34:0.0375875720,(((WP_120030433.1-Vibrio-sinensis:0.0756308350,WP_235060641.1-Vibrio-sp.-SS-MA-C1-2:0.1761073243)73/82:0.0100222278,WP_213972370.1-Vibrio-alginolyticus:0.1462266257)0/48:0.0059388038,(((((WP_012551417.1-Aliivibrio-salmonicida:0.0707289664,WP_012276595.1-Shewanella-halifaxensis:0.0527783299)90.4/96:0.0482909270,WP_135385147.1-Vibrio-tasmaniensis:0.1111299413)94.8/98:0.0540747656,WP_130066657.1-Aliivibrio-finisterrensis:0.0949840161)63.4/74:0.0251090627,WP_102571993.1-Vibrio:0.0887670390)92.7/69:0.0402277809,(((WP_171342182.1-Vibrio-cyclitrophicus:0.1111248482,WP_161972844.1-Enterovibrio-baiacu:0.0921700968)78.2/88:0.0277226255,(WP_193262143.1-Vibrio-parahaemolyticus:0.1839402777,(WP_102038478.1-Shewanella-sp.-GutCb:0.1698310634,WP_112353352.1-Shewanella-benthica:0.1148911867)81.4/99:0.0452814461)92.1/98:0.0518866584)84.9/76:0.0439451481,(WP_259522409.1-Shewanella-baltica:0.0861849126,WP_173243873.1-Shewanella-sp.-DC2-4:0.1042674613)93.8/97:0.0692385919)87.9/79:0.0430547316)83.9/67:0.0222616213)90.3/78:0.0361891810)5.1/10:0.0188153906,(((((((((WP_233983986.1-Vibrio-fluvialis:0.0702371046,WP_143680610.1-Vibrio-furnissii:0.0973383589)76.9/50:0.0163305606,((WP_172561824.1-Vibrio-furnissii:0.0757903871,(((WP_243025268.1-Vibrio-cincinnatiensis:0.2944074432,WP_222242368.1-Pseudoalteromonas-piscicida:0.2075228582)38.6/75:0.0639365516,WP_031827242.1-Vibrio-parahaemolyticus:0.1909688750)87.1/75:0.0592158018,(WP_047110243.1-Vibrio-vulnificus:0.1855010522,WP_102313365.1-Vibrio-cyclitrophicus:0.1234642390)90.7/78:0.0688650803)61.8/94:0.0392016702)74.6/75:0.0241975367,(WP_086585081.1-Vibrio-parahaemolyticus:0.0245112992,WP_217557493.1-Vibrio-metschnikovii:0.0462371223)96.9/100:0.0606617499)81.2/72:0.0316406896)41.6/21:0.0097022462,WP_233976434.1-Vibrio:0.1404129943)75.8/20:0.0067006990,((WP_158107962.1-Vibrio-furnissii:0.0928751801,WP_106651046.1-Shewanella-sp.-Pdp11:0.1159760298)93.4/39:0.0448293789,(((((WP_185890410.1-Vibrio-vulnificus:0.1169930100,WP_225486226.1-Vibrio-vulnificus:0.1056307942)66.8/57:0.0197356716,(WP_099031861.1-Pseudoalteromonas-sp.-GCY:0.0841543730,(WP_188011373.1-Vibrio-vulnificus:0.1196950424,WP_102451772.1-Vibrio-breoganii:0.1442215895)94.9/100:0.0568873742)21.3/36:0.0172846215)16.6/24:0.0108462380,WP_144217665.1-Shewanella-algae:0.1734198879)56.6/77:0.0325615353,(WP_031414678.1-Vibrio-parahaemolyticus:0.0970804994,WP_248468589.1-Shewanella-sp.-1CM18E:0.1296319170)94/99:0.0619643734)21.8/33:0.0080910717,WP_208844179.1-Pseudoalteromonas-xiamenensis:0.1295813004)85.9/35:0.0352211786)84.6/27:0.0246576353)78.7/59:0.0152254769,WP_229630939.1-Vibrio-coralliilyticus:0.1959613331)89.6/88:0.0285537738,(((((WP_102563398.1-Vibrio-sp.-10N.286.49.B3:0.0680155287,(WP_064489977.1-Vibrio-parahaemolyticus:0.0578450416,WP_201976964.1-Aeromonas-enteropelogenes:0.0818116730)95.3/100:0.0506717905)93.6/100:0.0431825755,WP_020328057.1-Vibrio-fluvialis:0.1102168381)0/24:0.0029690423,WP_039841338.1-Vibrio-harveyi-group:0.1228981233)94.1/56:0.0420663034,(((((((WP_041510674.1-Shewanella:0.0305196786,WP_187575439.1-Shewanella-algae:0.0606217980)85.8/97:0.0349075793,(WP_208158397.1-Shewanella-algae:0.0608165003,WP_208157589.1-Shewanella-algae:0.0370193311)89.1/100:0.0352305137)92.3/100:0.0491726464,((WP_171138527.1-Vibrio-plantisponsor:0.0436706194,WP_102966424.1-Vibrio:0.0820069983)98.4/100:0.0875752021,WP_072616325.1-Vibrio-vulnificus:0.1380921046)90.6/95:0.0453123534)26.3/72:0.0167051302,((WP_033540599.1-Shewanella:0.1009632016,WP_031815865.1-Vibrio-parahaemolyticus:0.1149725672)86.1/91:0.0369321074,(WP_193250619.1-Vibrio-parahaemolyticus:0.1316181801,WP_258516089.1-Vibrio-alginolyticus:0.0740572092)84.3/75:0.0185741114)89/85:0.0468970893)25.2/15:0.0113801223,WP_145570128.1-Vibrio-tasmaniensis:0.1260693576)59/25:0.0241174420,(WP_118894008.1-Vibrio-vulnificus:0.0950421247,WP_023404414.1-Vibrio-halioticoli:0.1664501141)53.4/49:0.0260264842)77.1/40:0.0176643580,(((WP_076542412.1-Shewanella-sp.-UCD-KL21:0.2207139711,WP_007466453.1-Photobacterium-marinum:0.1478801488)95.3/99:0.0816749649,(WP_036979211.1-Providencia-alcalifaciens:0.1270822465,WP_101040106.1-Psychromonas-sp.-MB-3u-54:0.1794372679)8.7/12:0.0261421881)22.6/11:0.0389960481,(WP_015854950.1-Musicola-paradisiaca:0.0355838831,WP_192457620.1-Musicola-keenii:0.0772062425)100/100:0.1638652854)92.4/96:0.0495186319)69.8/37:0.0124853245)87.1/51:0.0243567493,(((WP_047463293.1-Vibrio-fluvialis:0.0565035573,WP_193167845.1-Vibrio-navarrensis:0.2435949413)69.8/83:0.0404677351,WP_148560072.1-Vibrio-mimicus:0.1241963236)95.2/84:0.0425539675,WP_068331709.1-Photobacterium-jeanii:0.1591837327)85.4/49:0.0206892077)86.3/49:0.0195582320)80.4/47:0.0062887840,((WP_211915530.1-Vibrio-sp.-B1REV9:0.0782118925,((WP_045618469.1-Vibrio-vulnificus:0.1207064068,(WP_192928291.1-Vibrio-vulnificus:0.0353114006,WP_132698543.1-Vibrio-harveyi-group:0.0840847991)98.5/100:0.0785994626)43/61:0.0131707382,WP_017420706.1-Vibrio-vulnificus:0.0806164907)84.6/60:0.0192947051)0.7/60:0.0033356494,WP_017024716.1-Vibrio-rumoiensis:0.1451602203)97/61:0.0466958794)68.7/31:0.0037824188,(WP_069533349.1-Vibrio-parahaemolyticus:0.1040787156,((WP_103194250.1-Vibrio-vulnificus:0.0363468384,WP_158146950.1-Vibrio-fluvialis:0.0993393979)94.9/100:0.0590829650,WP_233927909.1-Vibrio-fluvialis:0.1025731337)89/100:0.0626464714)88/96:0.0330727636)76.2/43:0.0128167900,WP_268809473.1-Vibrio-parahaemolyticus:0.1220004221)85.5/37:0.0232280195)85/53:0.0352517753)25/12:0.0280049500,(((WP_108452928.1-Enterobacter-cloacae:0.2050822245,(WP_133040538.1-Shewanella-fodinae:0.2177117968,WP_237309237.1-Vibrio:0.1644276180)93/95:0.0573468176)87.1/64:0.0319983194,WP_138627230.1-Pseudoalteromonas-sp.-S4741:0.1473001439)29.3/36:0.0192853493,WP_232612305.1-Photobacterium-phosphoreum:0.2655495466)87.8/64:0.0325745254)99.8/98:0.1239085828,((WP_162556830.1-Proteus:0.0630300395,WP_161779122.1-Proteus-sp.-G2675:0.0804902693)99.9/100:0.1612533020,WP_236366479.1-Aeromonas-veronii:0.1483152627)18.8/52:0.0184470775)78.6/52:0.0385530490,((WP_071788415.1-Franconibacter-pulveris:0.3249958126,WP_020232007.1-Escherichia-coli:0.1430387327)7.2/69:0.0533526488,((WP_232581009.1-Photobacterium-phosphoreum:0.3286166927,(WP_073514120.1-Vibrio-parahaemolyticus:0.0538830681,WP_032481483.1-Vibrio:0.0164446860)100/100:0.3506218321)99/99:0.2074429916,(WP_228338545.1-Enterobacter-hormaechei:0.1238691429,(WP_109845273.1-Enterobacter:0.0462900051,WP_150146763.1-Enterobacter-roggenkampii:0.0360711281)77.1/100:0.0599743399)48.3/99:0.0790574585)73.1/59:0.0879042452)97.9/94:0.1016787414)95.5/97:0.0760254509,WP_097364011.1-Edwardsiella-tarda:0.2400821594)36.8/41:0.0242504951,((WP_172912938.1-Pantoea-stewartii:0.1183663290,WP_145394131.1-Pantoea-sp.-SJZ147:0.0883164056)99.8/100:0.1965021887,(WP_086868392.1-Kosakonia-pseudosacchari:0.3249497831,WP_153752706.1-unclassified-Citrobacter:0.2901625135)8/43:0.0954065285)96.3/92:0.1036505381)81.9/90:0.0376826565,WP_225633057.1-Aeromonas-enteropelogenes:0.1494235501)86/91:0.0278548752,WP_102949229.1-Aeromonas-caviae:0.2312284606)89.1/98:0.0352483778,(((WP_189493202.1-Formosimonas-limnophila:0.1716098321,WP_133620003.1-Hydromonas-duriensis:0.1862636245)98.7/100:0.1339820232,WP_061279316.1-Kluyvera-cryocrescens:0.2559784131)32.9/34:0.0379636202,(WP_168756429.1-Aeromonas-hydrophila:0.2804327516,WP_043158915.1-Aeromonas-hydrophila:0.2618608695)77.5/46:0.0694592576)87.9/96:0.0398045546)92.7/98:0.0453785882,(((((((((((((((((((((WP_151711904.1-Enterobacter-ludwigii:0.0318600213,(WP_032941356.1-Citrobacter-freundii:0.1645578753,WP_058678686.1-Enterobacter:0.0696678000)90.7/100:0.0520263682)54.6/99:0.0393508099,WP_223560970.1-unclassified-Enterobacter:0.0225801150)99.9/100:0.1836171456,WP_044720521.1-Escherichia-coli:0.2365377354)97.9/100:0.1519077821,WP_230863267.1-Enterobacter-cloacae:0.2530239381)81.7/85:0.0983021716,((WP_029399168.1-Escherichia-coli:0.1848035273,WP_050862535.1-Enterobacter:0.2510515642)37.3/94:0.0489982707,(WP_034496584.1-Buttiauxella-agrestis:0.2475823539,WP_164716793.1-Cedecea-lapagei:0.2363434135)92.4/100:0.0817825224)88.8/99:0.0774765684)84.2/81:0.0628146975,(WP_036421240.1-Morganella-morganii:0.1214858834,(WP_264998074.1-Morganella-morganii:0.0694079822,WP_186849509.1-Morganella-morganii:0.0665996118)95.4/100:0.0698470233)93.2/100:0.0607840136)79.1/81:0.0441331888,WP_064374050.1-Pantoea:0.2422440954)0/28:0.0251432850,(WP_039304623.1-Cedecea-neteri:0.2692112463,(WP_106902182.1-Citrobacter-freundii:0.1726015027,(WP_126291239.1-Enterobacter-hormaechei:0.1200176117,((((WP_062932819.1-Enterobacter-hormaechei:0.0267995076,WP_252042312.1-Enterobacter-chengduensis:0.0365091378)84.1/100:0.0226067705,WP_048999124.1-Enterobacter:0.0333837874)75.9/80:0.0074048423,WP_063144508.1-Enterobacter-cloacae:0.0310200714)84.6/80:0.0183638835,WP_049008289.1-Enterobacter-cloacae:0.0405474316)92.5/95:0.0484831857)94.7/100:0.0708623135)86.4/100:0.0487408532)89.3/60:0.0573745943)92.7/84:0.0370754575,(WP_038223662.1-Xenorhabdus-bovienii:0.1416736317,(WP_161748440.1-Proteus-sp.-G2658:0.1411894295,(WP_049199273.1-Proteus-mirabilis:0.0611071663,WP_157954980.1-Proteus-terrae:0.0790414100)65.4/98:0.0566685556)94/100:0.0714147128)93.3/93:0.0568970742)89.5/58:0.0357549562,(((WP_234505032.1-Kluyvera-intermedia:0.2892853747,(WP_257124601.1-Aeromonas-veronii:0.2460364171,WP_211007190.1-Pectobacterium-brasiliense:0.2791022678)28.8/91:0.0278989621)90.5/91:0.0494030510,(((((((WP_159241675.1-Proteus-terrae:0.1494324930,WP_195697765.1-Providencia-rettgeri:0.1290041434)95.3/93:0.0664874921,WP_165880752.1-Providencia-rettgeri:0.1245614627)29.5/81:0.0173121329,WP_231068620.1-Providencia-huaxiensis:0.1372831036)36.1/66:0.0172296552,WP_238563424.1-Providencia-rettgeri:0.0998581844)87.7/84:0.0285016405,((WP_187128479.1-Providencia-sp.-JUb39:0.1176983725,WP_193115284.1-Escherichia-coli:0.1036455928)90.5/100:0.0442539600,WP_164565644.1-Providencia-rettgeri:0.1792597053)32.6/65:0.0330374283)81.8/72:0.0349232322,(WP_154625743.1-Providencia-stuartii:0.1031298641,WP_004924376.1-Providencia-stuartii:0.1150972336)80.6/84:0.0377315406)89.1/74:0.0374733005,WP_169342312.1-Providencia-stuartii:0.1324482589)91.4/100:0.0375370559)94.7/92:0.0593308771,(WP_038258850.1-Xenorhabdus-bovienii:0.1950914765,WP_086113286.1-Xenorhabdus-beddingii:0.1461438965)85.9/80:0.0805777510)75.4/84:0.0256006809)67.9/70:0.0197027952,(((WP_115348269.1-Hafnia-alvei:0.1179013300,WP_046458830.1-Obesumbacterium-proteus:0.2030300931)42.8/80:0.0345686611,(WP_130961887.1-Hafnia-paralvei:0.1452123981,WP_061059890.1-Hafnia-paralvei:0.1308414398)84.5/80:0.0316881121)75.3/92:0.0358149607,(WP_120162698.1-Rahnella-variigena:0.0452439837,WP_183847446.1-Rahnella-inusitata:0.0537966901)100/100:0.2044189600)86.7/99:0.0441568176)92.4/82:0.0396781665,(((((WP_090123257.1-Kosakonia-arachidis:0.0775567192,WP_020454293.1-Kosakonia-sp.-SOY2:0.0798912357)85.8/100:0.0380567666,(WP_200002836.1-Citrobacter-freundii:0.1366668641,WP_080329661.1-Enterobacter-cloacae-complex:0.1088098705)99.9/100:0.1530891357)43.4/43:0.0170979303,WP_017458471.1-Kosakonia-sacchari:0.1218597925)97.1/97:0.0693213449,WP_090460354.1-Enterobacter-sp.-kpr-6:0.0872199860)99.3/100:0.1108986946,(WP_070133611.1-Candidatus-Erwinia-dacicola:0.1538260812,WP_140919905.1-Insectihabitans-xujianqingii:0.1620629855)35.9/43:0.0336460791)93.4/62:0.0544460876)35.7/32:0.0157911870,(WP_194857598.1-Citrobacter-freundii:0.2469654954,((WP_141032926.1-Escherichia-coli:0.1173703763,WP_196390204.1-Enterobacter-mori:0.0362282734)15/96:0.0259837432,(WP_252032207.1-Enterobacter-asburiae:0.0482331828,WP_111966211.1-Enterobacter-cloacae-complex:0.0215513402)44.4/54:0.0279696513)96.9/99:0.1125715061)100/100:0.1727405803)75.7/27:0.0150434006,(((WP_108569539.1-unclassified-Pantoea:0.1905188494,WP_028018520.1-Enterobacter-cloacae-complex:0.3489286252)98.3/100:0.1337844568,WP_269535377.1-Citrobacter-sedlakii:0.1493478658)92/98:0.0582085325,((WP_100272167.1-Citrobacter-portucalensis:0.1506219639,((WP_181624464.1-Citrobacter-freundii:0.1516494923,WP_001515521.1-Escherichia-coli:0.1379502237)95.1/100:0.0663738822,(((WP_058662478.1-Enterobacteriaceae:0.0431996045,WP_061709645.1-Pseudenterobacter-timonensis:0.0317146779)99.6/100:0.1224303915,WP_112164393.1-Rahnella-inusitata:0.1312459663)68.5/100:0.0614416408,WP_157953065.1-Limnobaculum-parvum:0.1609633120)82/89:0.0294646188)0/5:0.0182442343)87.3/98:0.0268925049,(WP_035600513.1-Edwardsiella-tarda:0.2021073348,((WP_202782010.1-Enterobacter-asburiae:0.0650584654,WP_144817260.1-Pseudescherichia-sp.-L3:0.0378773358)98.9/100:0.1261853119,WP_125117341.1-Aeromonas-caviae:0.2275118207)32.4/99:0.0468827877)92.3/93:0.0563161538)81.5/60:0.0277393754)83.6/51:0.0152324283)90.9/27:0.0291888591,((WP_021540003.1-Escherichia-coli:0.0000021047,WP_047657564.1-Enterobacteriaceae:0.0527965805)95/100:0.0662349299,WP_159282060.1-Rahnella-variigena:0.2835957712)28.8/26:0.0326147159)13.8/23:0.0195763146,((((((((WP_016236761.1-Enterobacteriaceae:0.0811940730,WP_024239504.1-Escherichia-coli:0.0980579244)91.7/100:0.0509618884,((WP_001610415.1-Escherichia-coli:0.0312227376,WP_138183825.1-Escherichia-sp.-E4385:0.0213454837)99/100:0.1033863626,(WP_089585421.1-Escherichia-coli:0.0400727350,WP_104705897.1-Escherichia-coli:0.0253001870)99.9/100:0.1382424902)89.7/100:0.0710314465)75.2/88:0.0488690936,WP_033870520.1-Enterobacteriaceae:0.1466517035)90.3/94:0.0357190794,(WP_179154141.1-Citrobacter-freundii:0.2061180973,WP_086539147.1-Citrobacter-freundii:0.1552367183)91.3/97:0.0600204772)87.1/80:0.0308631844,((WP_107192407.1-Escherichia-albertii:0.0190419785,WP_181673375.1-Escherichia-fergusonii:0.0350612532)99.5/100:0.1056959702,(((((WP_032214194.1-Escherichia-coli:0.0353232266,WP_085459683.1-Escherichia-coli:0.0857438757)10.3/65:0.0134082640,WP_059217428.1-Escherichia-albertii:0.0320830311)93.7/89:0.0434840802,(WP_073278747.1-Escherichia-coli:0.0554617548,WP_087893724.1-Enterobacteriaceae:0.0425866544)96.6/100:0.0712203413)91.5/99:0.0649309483,WP_167816599.1-Enterobacter-cloacae:0.2259409450)39.5/32:0.0643309556,(WP_262183194.1-Enterobacter-mori:0.1315382912,(WP_063412371.1-Enterobacter-cloacae:0.0397389105,WP_047027451.1-Enterobacter-kobei:0.0387268151)97.3/100:0.0702238221)99.4/100:0.1116493491)39.7/40:0.0245449326)87.2/61:0.0300414739)85.4/82:0.0187607728,WP_100029735.1-Escherichia-coli:0.1063990511)80.7/85:0.0190095138,WP_267387874.1-Citrobacter-portucalensis:0.2224916760)90.4/30:0.0301076680,(WP_001297344.1-Escherichia-coli:0.1246425559,WP_138098269.1-Jejubacter-calystegiae:0.1197308059)92.6/95:0.0585861521)71.2/28:0.0065530945)89.4/25:0.0296991019,((WP_119938542.1-Enterobacter-huaxiensis:0.0799045313,WP_050871166.1-Enterobacter-cloacae-complex:0.0319908604)92.6/100:0.0506426867,(WP_228937168.1-Escherichia-marmotae:0.0291964448,WP_040079227.1-Escherichia-coli:0.0316062678)79.8/80:0.0095880755)96/100:0.0437156455)80.6/27:0.0131286673,((((((((((WP_127791218.1-Escherichia-coli:0.1279583902,(((WP_021577651.1-Escherichia-coli:0.1075673524,WP_107179425.1-Escherichia:0.0653848252)64.2/94:0.0257423821,WP_059222105.1-Escherichia-albertii:0.0874163969)72.1/29:0.0107028906,WP_187209312.1-Escherichia-coli:0.0665514963)62.9/27:0.0108875213)53.3/33:0.0236415552,(WP_042093344.1-Escherichia-coli:0.1922875644,WP_181465273.1-Escherichia-fergusonii:0.0346700418)80.2/26:0.0374499229)79.8/9:0.0158704759,((((((((WP_170897747.1-Escherichia-coli:0.1547550501,WP_208464084.1-Escherichia-coli:0.0804370793)90.9/96:0.0416508475,WP_162379936.1-Citrobacter-sp.-NCU1:0.1343450521)60.7/85:0.0144619293,WP_096122580.1-Escherichia-coli:0.0556061324)82.3/89:0.0127134818,(WP_095374511.1-Escherichia-coli:0.0081150490,WP_012906377.1-Citrobacter-rodentium:0.0781759733)99.6/100:0.0722574799)90.4/95:0.0304310693,((WP_230350752.1-Lelliottia-sp.-WAP21:0.1077238427,WP_265185975.1-Enterobacter-mori:0.0759919517)83.2/88:0.0387730806,(((WP_141109706.1-Escherichia-albertii:0.0687291645,(WP_203360733.1-Citrobacter-sp.-R56:0.0552664543,WP_001572827.1-Escherichia-coli:0.0501060340)0.5/53:0.0203831677)87.2/100:0.0422368586,WP_258993080.1-Scandinavium-hiltneri:0.0611051338)96/100:0.0492922116,WP_046888360.1-Enterobacter-cloacae:0.0508711399)9.2/84:0.0040795741)77.7/88:0.0327856486)77.8/70:0.0106470745,((WP_250341216.1-Pantoea-eucrina:0.1853949396,WP_047609274.1-Rahnella-aquatilis:0.1549310546)85.8/72:0.0370105131,(WP_085454021.1-Escherichia-coli:0.0968816479,WP_105963664.1-Escherichia-coli:0.0455774257)75.4/84:0.0281594988)70.3/50:0.0048721169)95.9/66:0.0292770291,(((WP_063610377.1-Escherichia-coli:0.0489816245,WP_001100798.1-Enterobacteriaceae:0.0995714370)84/85:0.0227000259,(WP_088541377.1-Escherichia-coli:0.0278662901,WP_222691870.1-Escherichia-marmotae:0.0461859469)92.8/100:0.0301643512)31.1/70:0.0124050469,((WP_218070502.1-Escherichia-coli:0.0631007004,WP_015386517.1-Cronobacter-sakazakii:0.1619243104)64.5/61:0.0150608958,(WP_146314246.1-Escherichia-coli:0.0307648397,WP_096967326.1-Escherichia-coli:0.0218399387)97.8/100:0.0568433588)90.4/66:0.0358837716)84.8/65:0.0234236444)84.6/57:0.0172828824,((WP_001100781.1-Escherichia-coli:0.1661502056,WP_053290779.1-Escherichia-coli:0.0668846891)82.7/38:0.0310971307,((WP_105274811.1-Escherichia:0.1554634750,WP_064512195.1-Buttiauxella-gaviniae:0.1945186669)92.8/99:0.0706615936,WP_172410167.1-Escherichia-coli:0.1615119075)98.2/97:0.0830558484)75.9/15:0.0135731303)77.1/11:0.0102680505)69.7/66:0.0231747728,(WP_137653986.1-Escherichia-albertii:0.0276858251,WP_001100785.1-Escherichia-coli:0.0325358267)99.2/100:0.0701822657)75.8/49:0.0096888786,((((((((WP_175320495.1-Hafnia-paralvei:0.0947194674,WP_130996599.1-Hafnia-paralvei:0.0977290775)94.8/97:0.0885382438,(WP_171269898.1-Aeromonas-media:0.1004585134,WP_216983019.1-Aeromonas-sp.-FDAARGOS-1415:0.0571979501)98.2/100:0.0864538856)90.8/73:0.0486943362,WP_168366848.1-Dickeya-zeae:0.0905734014)70.7/73:0.0399911596,WP_252995952.1-Aeromonas-encheleia:0.1939351123)40/68:0.0235433085,(WP_227095090.1-Aeromonas-veronii:0.0809641279,WP_152109808.1-Plesiomonas-shigelloides:0.0557183324)99.7/100:0.0987885592)87/97:0.0345124304,(WP_223932214.1-Aeromonas-caviae:0.0232891423,WP_161506954.1-Aeromonas-media:0.0902882258)99.3/100:0.0930035831)96.8/76:0.0670035871,(WP_034458444.1-Buttiauxella-noackiae:0.1188895989,WP_253383865.1-unidentified-bacterial-endosymbiont:0.1987624622)13.8/67:0.0246089914)76.7/96:0.0193693779,(WP_105591414.1-Pantoea-coffeiphila:0.0500143478,WP_123334936.1-Erwinia-sp.-JUb26:0.0316472441)99.9/100:0.1534707854)94/96:0.0314326392)74.3/67:0.0085692498,(WP_001527244.1-Escherichia-coli:0.0913440495,(WP_157704420.1-Escherichia-coli:0.0799010104,WP_042200020.1-Enterobacteriaceae:0.0442072535)70.2/100:0.0211747767)74.5/85:0.0125675049)69.7/50:0.0167899756,WP_220393175.1-Citrobacter-freundii:0.2571180323)84.8/39:0.0233932271,(((((((((WP_193360946.1-Enterobacter-cloacae-complex-sp.-P39RS:0.0400654712,WP_182945676.1-Enterobacter-cloacae:0.0252828614)97.3/100:0.0533977027,WP_259388919.1-Enterobacter-huaxiensis:0.0725617209)86.5/69:0.0197206602,(((((WP_096205048.1-Enterobacter:0.0237766241,WP_100165931.1-Enterobacter-mori:0.0302363457)97.3/100:0.0653025227,WP_242172213.1-Enterobacter:0.0477742155)68.7/96:0.0277771232,WP_239609625.1-Lelliottia-amnigena:0.0816944331)87.3/97:0.0277456089,WP_063158410.1-Enterobacter-hormaechei:0.0678267015)82.1/93:0.0184427776,((WP_032636062.1-Enterobacter-kobei:0.0478337803,WP_138014015.1-Enterobacter-ludwigii:0.0446777288)89.6/100:0.0443307686,(WP_063137673.1-Enterobacter-cloacae-complex:0.0289091891,WP_165464597.1-Enterobacter-cloacae:0.0451401984)80/100:0.0169792482)91.7/100:0.0371821376)87.3/95:0.0211075770)86.9/68:0.0198394887,WP_033487312.1-Enterobacter-cloacae-complex:0.0498773275)92.7/93:0.0309223734,(((WP_042716137.1-Enterobacter-sp.-B509:0.0159378936,WP_241820523.1-Enterobacter-kobei:0.0593509537)99.2/100:0.0680554248,WP_063869545.1-Enterobacter-hormaechei:0.0546531440)67/87:0.0079092318,(WP_172688853.1-Enterobacterales:0.0733913156,WP_126546226.1-Enterobacter-quasimori:0.0763104847)93.9/99:0.0460942496)87.5/88:0.0279648021)78.2/75:0.0090309816,((((WP_214593796.1-Enterobacter-mori:0.0369516511,WP_086532312.1-Enterobacter-ludwigii:0.0634689404)98.1/100:0.0774538510,(((WP_047742849.1-Enterobacter:0.0583388192,WP_040022520.1-Enterobacter-cloacae:0.0683888130)98.5/100:0.0751144521,(WP_129995405.1-Enterobacter-cloacae:0.0567904073,WP_042099316.1-Escherichia-coli:0.0557379397)85.6/100:0.0255853957)92.9/100:0.0527189702,WP_045339085.1-Enterobacter-cloacae-complex:0.1161876900)38.9/97:0.0237046974)92.9/95:0.0557359791,WP_126315015.1-Enterobacter-hormaechei:0.0622407847)55.7/78:0.0240695725,WP_045338355.1-Enterobacter-hormaechei:0.1028735213)91.6/89:0.0359530980)84.1/80:0.0145912943,(WP_208283396.1-Enterobacter-sichuanensis:0.0640779334,(WP_182115563.1-Enterobacter-kobei:0.0553312567,WP_213381159.1-Enterobacter-cloacae:0.0332725468)97.8/100:0.0627607973)90.2/81:0.0245948251)87.9/89:0.0233885817,(((((WP_181680936.1-Citrobacter-sp.-RHB25-C09:0.0922839577,(WP_090049757.1-Citrobacter-amalonaticus:0.0286719819,WP_196013128.1-Citrobacter-farmeri:0.0238193605)99.5/100:0.0972391074)78.6/51:0.0344225935,WP_103775088.1-Citrobacter-amalonaticus:0.1376163920)99.4/100:0.1083740872,WP_121265736.1-Enterobacter-sp.-R1-2018:0.1645726575)81.5/51:0.0448367494,WP_259047621.1-Klebsiella-sp.-BIGb0407:0.1226355262)22.6/42:0.0120143507,WP_249031739.1-Tenebrionicola-larvae:0.2026176050)86.8/42:0.0248598277)42.5/36:0.0072627690,WP_259000881.1-Scandinavium-goeteborgense:0.1414281540)89.5/55:0.0224393782)78.9/31:0.0108687628,((((WP_001100804.1-Enterobacteriaceae:0.0804310342,(WP_213789144.1-Enterobacter-hormaechei:0.0048777916,((WP_148242178.1-Enterobacter-asburiae:0.0452114570,WP_237994110.1-Enterobacter-sp.-DRP3:0.0171425242)74/100:0.0050403657,WP_237424876.1-Enterobacter-cloacae-complex-sp.-ECC445:0.0402042091)91.6/100:0.0416431293)100/100:0.1950119650)91.5/100:0.0679261548,WP_075180894.1-Pantoea-sp.-1.19:0.1479750733)93.3/95:0.0596379800,(WP_162648791.1-Citrobacter-freundii:0.0747363646,((WP_060683585.1-Citrobacter-portucalensis:0.0998759215,(WP_079798993.1-Salmonella-enterica:0.0818211845,WP_032255798.1-Enterobacteriaceae:0.0673080733)73/87:0.0099054169)93.5/99:0.0519500653,(WP_195994783.1-Citrobacter-portucalensis:0.1617711365,WP_048218074.1-Citrobacter-freundii-complex:0.1283549268)74.2/99:0.0627285948)20.7/32:0.0321408874)93.3/86:0.0466440343)86.7/77:0.0242081232,(WP_174633339.1-Escherichia-coli:0.0874364067,WP_025210617.1-Escherichia-coli:0.0680737615)86.6/75:0.0339978661)93.2/56:0.0314205231)74.1/30:0.0065750975,(((((WP_187216555.1-Escherichia-coli:0.1085118832,((((WP_049002539.1-Citrobacter-freundii:0.0886166357,(WP_249576357.1-Enterobacter-vonholyi:0.0557616921,WP_161617217.1-Yokenella-regensburgei:0.0317952027)98.1/100:0.0586134427)79.9/100:0.0219115637,((WP_079834758.1-Salmonella-enterica:0.0309558214,WP_058109100.1-Salmonella-enterica:0.0303026870)40.6/97:0.0278832757,(WP_079793206.1-Salmonella-enterica:0.0815875040,WP_001100807.1-Salmonella:0.0704739454)98.5/100:0.0845226202)98.4/100:0.0634626872)89.9/100:0.0355192891,WP_032303308.1-Escherichia:0.0513961787)95.9/100:0.0532293434,WP_061892435.1-Escherichia-coli:0.1023942532)86.6/99:0.0163809868)84.5/99:0.0135146647,(WP_232904305.1-Buttiauxella-sp.-W03-F01:0.1217898981,WP_034915863.1-Erwinia-sp.-9145:0.1107909289)95.9/98:0.0507639410)90.5/73:0.0271297878,((((WP_262931739.1-Escherichia-albertii:0.0309775719,WP_160188772.1-Escherichia-coli:0.0355411186)93.5/100:0.0570738625,WP_072664248.1-Escherichia:0.0922202147)84.1/99:0.0328768790,(WP_089706542.1-Escherichia-coli:0.0801427754,WP_096972986.1-Escherichia-coli:0.0778846432)80.7/42:0.0326818054)69.9/40:0.0232533696,WP_097421351.1-Escherichia-coli:0.0744970940)94.6/40:0.0364546020)0/25:0.0000027503,((WP_109548670.1-Escherichia-coli:0.0421651892,WP_001100790.1-Escherichia:0.0181782411)98.6/100:0.0651997461,WP_181237856.1-Escherichia-marmotae:0.0547598896)74.8/60:0.0102815452)81.1/41:0.0159542564,WP_191149682.1-Erwinia-aphidicola:0.1272567392)92.3/42:0.0374838493)92.2/85:0.0241692379)87/27:0.0162710956,((((((WP_103946175.1-Lelliottia-aquatilis:0.0897080993,WP_123427447.1-Lelliottia:0.0498953332)76/99:0.0291028766,(WP_235147774.1-Enterobacter-cloacae:0.1204945293,WP_061356323.1-Escherichia-coli:0.4399498221)88.9/56:0.0805502803)96.6/52:0.0697258401,(WP_001100786.1-Escherichia-coli:0.0903687459,(WP_248249449.1-Enterobacter-roggenkampii:0.1193065623,WP_032936727.1-Citrobacter-freundii:0.0950477085)99.9/100:0.1242727293)89.6/51:0.0319807886)74.8/67:0.0096152367,WP_137650031.1-Escherichia-albertii:0.0995214609)89.7/71:0.0242177959,WP_262938691.1-Escherichia-albertii:0.1076463538)50.4/78:0.0091730078,(((WP_095844187.1-Escherichia-coli:0.0521683214,(WP_086118650.1-Escherichia-coli:0.0231146966,WP_089538038.1-Escherichia-coli:0.0300142939)70.2/99:0.0074852212)99.6/100:0.1102076537,WP_219093375.1-Escherichia-coli:0.1619583345)56.1/22:0.0403136173,(WP_001542972.1-Escherichia-coli:0.1112969054,WP_153879942.1-Citrobacter-freundii:0.1202638443)80.4/92:0.0224844525)87/27:0.0298549860)89.2/28:0.0267139739)52.9/63:0.0257281656,((WP_157959988.1-Citrobacter:0.2789252143,WP_073446180.1-Escherichia-coli:0.1208887310)95.3/98:0.0628297403,((((WP_238085128.1-Pseudescherichia-vulneris:0.0253552869,WP_129363899.1-Enterobacter-cloacae:0.0299703368)0/24:0.0063700165,WP_032652659.1-Enterobacter-cloacae-complex:0.0834101318)56.8/73:0.0150827809,(WP_257661844.1-Citrobacter-portucalensis:0.1005071325,WP_193134185.1-Citrobacter-freundii:0.0710606258)80.7/100:0.0222523516)96.9/96:0.0749856640,(WP_042284276.1-Citrobacter-sedlakii:0.1713106869,(WP_024193259.1-Escherichia-coli:0.0879418586,WP_099433281.1-Citrobacter-amalonaticus:0.0303537403)89.6/97:0.0651697276)95.2/100:0.0651957402)99.2/94:0.1051991142)31.8/41:0.0319765257)83.8/56:0.0214277155,(WP_032659356.1-Enterobacter-hormaechei:0.2346666299,WP_029881813.1-Enterobacter:0.1716270955)95.1/97:0.0849420196)87/55:0.0311335235)74.7/8:0.0090507547,((((WP_085037002.1-Cronobacter-sakazakii:0.1560972338,WP_076770125.1-Kosakonia-cowanii:0.1091000904)98.4/100:0.1267833931,WP_216981205.1-Lelliottia-amnigena:0.2030552213)91.7/98:0.0696892574,((WP_227534785.1-Erwinia-rhapontici:0.1117124224,WP_017800228.1-Erwinia-toletana:0.1167145751)93.4/100:0.0704224588,WP_255851625.1-Erwinia-persicina:0.1306311579)74/94:0.0350452870)95.6/98:0.0895905331,(((WP_167863639.1-Erwinia-rhapontici:0.1038388869,WP_039385357.1-Pantoea-eucrina:0.1571398011)92.2/100:0.0482431180,WP_118663751.1-Erwinia-persicina:0.1181746663)78.8/97:0.0302865199,WP_062742814.1-Erwinia-persicina:0.1416758927)58.7/99:0.0426554028)88.6/68:0.0366609898)76.7/21:0.0108719199,(((((WP_132257458.1-Paucimonas-lemoignei:0.0600199220,WP_119785517.1-Noviherbaspirillum-sedimenti:0.0591778308)100/100:0.2002429073,((WP_026470910.1-Alkanindiges-illinoisensis:0.2078086540,WP_248147450.1-Moraxella-osloensis:0.2342691967)23.7/90:0.0775592803,WP_092690500.1-Acinetobacter-kyonggiensis:0.1103289357)99.4/100:0.2172878801)99.1/98:0.1508767855,((WP_151054911.1-Cellvibrio-sp.-KY-YJ-3:0.1359294914,WP_078044787.1-Cellvibrio-sp.-PSBB023:0.1652725688)56.9/76:0.0320747477,(WP_012489013.1-Cellvibrio-japonicus:0.1093421985,WP_087465811.1-Cellvibrio-sp.-PSBB006:0.1498750044)29.2/75:0.0238487097)89.7/99:0.0531771693)85.2/96:0.0402925684,((((WP_097405671.1-Kosakonia:0.0389334101,WP_161518249.1-Kosakonia-sp.-MH5:0.0584339893)96.4/100:0.0603057029,WP_128482629.1-Kosakonia-cowanii:0.0880857050)97/100:0.0835458253,WP_206164314.1-Citrobacter:0.1888579224)69.4/93:0.0515882178,(WP_048221111.1-Citrobacter:0.1105049328,WP_263807164.1-Salmonella-enterica:0.1376729701)99.2/100:0.1265940123)87.6/94:0.0373873806)87.6/78:0.0316279750,(((WP_047682696.1-Xenorhabdus:0.0509103814,WP_099124588.1-Xenorhabdus-stockiae:0.0622075108)99.5/100:0.1051707590,(WP_074024273.1-Xenorhabdus-eapokensis:0.0586500871,WP_217470569.1-Photorhabdus-akhurstii:0.0674723595)92.2/100:0.0641781416)98/100:0.0752483825,(((((((((((((((((((WP_236369955.1-Aeromonas-veronii:0.0402953236,WP_113736183.1-Aeromonas-caviae:0.0319516145)73.1/100:0.0125418047,((WP_042019775.1-Aeromonas-sobria:0.0443860074,WP_139495680.1-Aeromonas-veronii:0.0209023795)98.5/100:0.0455625350,(WP_098969895.1-Aeromonas-sp.-CU5:0.0397781114,WP_159163188.1-Aeromonas-veronii:0.0404348537)93.1/100:0.0323934170)91.5/100:0.0251635977)87.7/54:0.0249924100,((((WP_263173612.1-Shewanella-sp.-SM73:0.1153022190,((WP_263191039.1-Shewanella-sp.-SM87:0.0576999271,WP_263255112.1-Shewanella-sp.-SM32:0.0409119665)66/89:0.0151709434,(WP_263178828.1-Shewanella-sp.-SM101:0.0349378027,WP_109248233.1-Shewanella-algae:0.0602129004)93.5/100:0.0299223999)89.8/98:0.0332380615)94.2/100:0.0497424416,(WP_220777726.1-Shewanella-hafniensis:0.0232525806,WP_055647506.1-Shewanella-sp.-Sh95:0.0287687910)87/100:0.0379298877)87.7/100:0.0337704979,(WP_082969903.1-Aeromonas-dhakensis:0.0918025644,WP_104454419.1-Aeromonas-sp.-ASNIH2:0.0460335313)73.2/56:0.0092422385)65.1/54:0.0318517584,WP_201035193.1-Aeromonas-media:0.0948921875)93.5/95:0.0449124590)97.6/100:0.0492006133,WP_143237783.1-Aeromonas-caviae:0.0402873462)72.5/56:0.0128540850,(WP_113739100.1-Aeromonas-veronii:0.0478615184,WP_226032029.1-Aeromonas-hydrophila:0.0351682931)84.8/86:0.0110533224)0/22:0.0000020310,(WP_214003767.1-Aeromonas-caviae:0.0314085247,(WP_139459904.1-Aeromonas-veronii:0.0469064216,WP_041208446.1-Aeromonas-jandaei:0.1122153932)99.4/96:0.0716130107)82.2/87:0.0155048028)85.9/31:0.0164055377,WP_252615985.1-Aeromonas-hydrophila:0.0629395118)73.7/34:0.0059991871,(WP_080988732.1-Aeromonas-caviae:0.0377944941,WP_108536035.1-Aeromonas-sp.-HMWF014:0.0299201477)91/100:0.0248952345)96.1/99:0.0530488027,(WP_100646402.1-Aeromonas-veronii:0.0505602971,((WP_252601295.1-Aeromonas-dhakensis:0.0196558593,WP_139432874.1-Aeromonas-dhakensis:0.1185649681)91.4/100:0.0276015558,WP_106553824.1-Aeromonas-hydrophila:0.0521190546)73.6/100:0.0195207013)99.9/100:0.0983256048)73.2/40:0.0105164828,((WP_268010485.1-Aeromonas-dhakensis:0.0609463203,WP_098985072.1-Aeromonas-sp.-CA23:0.0364289335)98.9/100:0.0997315311,(WP_181268965.1-Aeromonas-veronii:0.0271898789,(WP_129146711.1-Aeromonas-caviae:0.0215197455,(WP_081843667.1-Aeromonas-taiwanensis:0.0169318447,WP_139744464.1-Aeromonas-media:0.0379980963)83.6/100:0.0197016680)79.4/98:0.0202978113)89.5/98:0.0540060766)99.6/100:0.1182156616)94.7/40:0.0495516282,(WP_059168599.1-Aeromonas-salmonicida:0.0589617082,(WP_033115905.1-Aeromonas-lacus:0.0473543251,((WP_223920301.1-Aeromonas-caviae:0.0294691978,WP_226058924.1-Shewanella-sp.-SR1:0.1011686163)87/98:0.0199093473,(WP_058394234.1-Aeromonas-salmonicida:0.0265513926,WP_257690725.1-Aeromonas-veronii:0.0393237864)79.3/100:0.0143096484)39/98:0.0105329595)92.4/98:0.0362100851)79.1/97:0.0288969875)87.7/38:0.0350730892,(((WP_223945983.1-Aeromonas-caviae:0.0929134211,((WP_265458754.1-Aeromonas-salmonicida:0.0684160519,WP_125597700.1-Aeromonas-salmonicida:0.0208602683)0/98:0.0000024554,(WP_180911080.1-Aeromonas-caviae:0.0346948125,WP_264818242.1-Aeromonas-caviae:0.0229939509)90.4/100:0.0175570230)80.2/96:0.0097573272)0/44:0.0000026995,((WP_040098646.1-Aeromonas:0.0835340216,WP_163128050.1-Aeromonas-allosaccharophila:0.0725252605)0/46:0.0000905951,((WP_263309349.1-Aeromonas-caviae:0.0344537832,WP_041217344.1-Aeromonas-hydrophila:0.0394510009)62.4/75:0.0041652231,(WP_257686696.1-Aeromonas-veronii:0.0335028501,(WP_255912544.1-Aeromonas-sp.-JL9:0.0251209656,((WP_248067241.1-unclassified-Shewanella:0.0883623656,WP_148505453.1-Vibrio-cholerae:0.0723979445)0/99:0.0010697787,(WP_218558465.1-Shewanella-putrefaciens:0.0657147328,WP_011919582.1-Shewanella-putrefaciens:0.0293654079)100/100:0.1268216814)99.8/98:0.0927670608)0/91:0.0000020549)79.8/74:0.0091671550)92.7/96:0.0174678064)80.2/46:0.0058536889)77.7/82:0.0153169809,(((WP_139750984.1-Aeromonas-media:0.0265515523,WP_224430864.1-Aeromonas-rivuli:0.0247539373)79.1/100:0.0222196979,WP_041996113.1-Aeromonas-bivalvium:0.0442222222)87.2/100:0.0278546947,WP_265776268.1-Aeromonas-veronii:0.1072905021)77.5/65:0.0189225087)79.2/62:0.0209591008)97.6/45:0.0689899597,(WP_216990617.1-Aeromonas-sp.-FDAARGOS-1409:0.1038723274,(((((((((WP_201942221.1-Aeromonas-veronii:0.1387362644,WP_159143398.1-Aeromonas-veronii:0.0066292497)80/96:0.0133732664,WP_203131090.1-Aeromonas-jandaei:0.0471518703)73.9/52:0.0060164516,(WP_139692403.1-Aeromonas-sobria:0.0320813043,WP_111901368.1-Aeromonas-veronii:0.0312430418)18.3/49:0.0065055809)84/50:0.0142449423,WP_042657830.1-Aeromonas:0.0369693854)84.4/88:0.0119560585,WP_268854930.1-Aeromonas-enteropelogenes:0.0913538474)27.8/80:0.0064356270,WP_223380193.1-Aeromonas-enteropelogenes:0.0240972766)98.5/100:0.0698869455,((WP_236321701.1-Aeromonas-veronii:0.0577400598,(((WP_050718733.1-Aeromonas-tecta:0.1253830332,WP_100859741.1-Aeromonas-lusitana:0.0313080231)66.1/97:0.0105131975,((WP_077098334.1-Aeromonas-hydrophila:0.0503206120,WP_252420256.1-Aeromonas-taiwanensis:0.0751705893)11.5/94:0.0052249052,WP_043167431.1-Aeromonas-hydrophila:0.0918439046)94.1/97:0.0416367057)53/97:0.0312304850,WP_042654929.1-Aeromonas-encheleia:0.0262008674)97.9/100:0.0604834696)83.4/84:0.0153917165,WP_257119864.1-Aeromonas-hydrophila:0.0092979305)91.8/84:0.0331196117)28.3/57:0.0203827842,(WP_257707542.1-Aeromonas-caviae:0.0459631245,WP_182951616.1-Aeromonas-sp.-WP2-W18-CRE-05:0.0549571264)99.4/100:0.0788904284)74.7/86:0.0157803286,WP_045790994.1-Aeromonas-hydrophila:0.1685254077)63.8/89:0.0401685047)5.9/26:0.0103449393)76.2/73:0.0210679443,WP_236563281.1-Aeromonas-veronii:0.1200815093)79.2/39:0.0199909823,((WP_213320225.1-Aeromonas-caviae:0.0530164450,WP_203764723.1-Aeromonas-caviae:0.0650350096)96.7/100:0.0904690234,WP_069554789.1-Aeromonas-sp.-DNP9:0.1070455057)72.5/59:0.0405299635)30.4/56:0.0191635349,(((WP_064338398.1-Aeromonas-veronii:0.1099402411,((((((((WP_247163170.1-Escherichia-coli:0.0174780112,WP_038644959.1-Pantoea:0.0695550915)99.2/100:0.0760915436,(WP_252422582.1-Aeromonas-taiwanensis:0.0153290747,WP_263719954.1-Aeromonas-caviae:0.0780523539)70/88:0.0373577314)91.4/100:0.0332235188,WP_033131297.1-Aeromonas-aquatica:0.0220032984)93.1/91:0.0388820894,WP_241344952.1-Aeromonas-sp.-MR16:0.0128005619)85.7/91:0.0181322332,WP_160838676.1-Aeromonas-caviae:0.0709419792)88.8/100:0.0339214343,(((WP_116547128.1-Plesiomonas-shigelloides:0.0784116775,WP_201996661.1-Aeromonas-veronii:0.0286529109)81.4/50:0.0112386019,WP_235463836.1-Aeromonas-australiensis:0.0172329483)75.5/46:0.0052171388,(WP_139496062.1-Escherichia-coli:0.0529149553,WP_215269322.1-Aeromonas-dhakensis:0.0404752088)70.2/60:0.0058802537)87.1/98:0.0269021965)49.1/95:0.0355901723,WP_049048056.1-Aeromonas-hydrophila:0.0985332255)87.2/94:0.0386409434,WP_167569591.1-Aeromonas-veronii:0.0665708305)92.6/100:0.0547430204)37.9/76:0.0255430860,((WP_042864139.1-Aeromonas-caviae:0.0357442856,WP_225159881.1-Aeromonas-caviae:0.0350849765)94.5/100:0.0486003517,(WP_244056308.1-Aeromonas-caviae:0.1183326984,WP_202046159.1-Aeromonas-media:0.0778087092)96.9/100:0.0765061857)96/100:0.0656407476)89.9/86:0.0244271870,((((WP_148612677.1-Aeromonas-sobria:0.0377362840,WP_201355655.1-Aeromonas-hydrophila:0.0438173188)62.4/69:0.0058355008,WP_171268810.1-Aeromonas-media:0.0267013681)69.2/63:0.0051150741,WP_201951905.1-Aeromonas-veronii:0.0301805057)98.9/100:0.0756281168,(((((WP_212539243.1-Aeromonas-salmonicida:0.0283254557,((WP_187590672.1-Aeromonas-hydrophila:0.1031959558,WP_017782281.1-Aeromonas-dhakensis:0.0267032866)43.4/100:0.0140137652,WP_060586794.1-Aeromonas-schubertii:0.0450825560)99.5/100:0.0785858292)91.8/98:0.0226549872,WP_201904077.1-Aeromonas-veronii:0.0557050350)75.5/98:0.0118219409,WP_256684552.1-Aeromonas-salmonicida:0.0397433792)66.1/49:0.0083195374,WP_206874356.1-Aeromonas-veronii:0.0113094767)72.9/51:0.0128654010,WP_042063659.1-Aeromonas-allosaccharophila:0.0454783878)94.6/100:0.0527812620)90.2/96:0.0384530133)89.5/87:0.0286943424)76.2/68:0.0101573243,WP_265468300.1-Aeromonas-salmonicida:0.1183749875)83.9/96:0.0236003551,((((((((WP_234664799.1-Aeromonas-allosaccharophila:0.0988628949,(WP_257692500.1-Aeromonas-caviae:0.1189152593,WP_113719982.1-Aeromonas-salmonicida:0.0901813591)79.4/99:0.0450427702)94.7/97:0.0538663573,(WP_225159845.1-Aeromonas-caviae:0.0841145478,WP_210550341.1-Aeromonas-sp.-SrichE-2G:0.2402287146)80.3/65:0.0219389566)72.4/61:0.0250169651,WP_245131329.1-Aeromonas-veronii:0.1041642655)33.3/81:0.0299811397,((((WP_101347551.1-Aeromonas-sobria:0.0514239537,WP_201955411.1-Aeromonas-veronii:0.0512967038)98.4/100:0.0734582852,WP_136501435.1-Aeromonas-veronii:0.1228641432)92.5/73:0.0435457878,(((WP_050666161.1-Aeromonas-schubertii:0.0616944354,WP_210600017.1-Aeromonas-sp.-MrichA-1:0.0385435205)85.7/98:0.0188478282,(WP_268029297.1-Aeromonas-hydrophila:0.0885891481,WP_201942265.1-Aeromonas-veronii:0.1110657046)99.7/100:0.1134123169)78.1/67:0.0159754384,WP_059168611.1-Aeromonas-salmonicida:0.0880367554)88.3/75:0.0252096869)82.1/73:0.0153376752,(WP_163135231.1-Aeromonas-rivipollensis:0.0281679523,WP_241279715.1-Aeromonas-sp.-MR19:0.0278769070)100/100:0.1713773533)78.3/69:0.0083191181)85.8/93:0.0282960020,(WP_042649704.1-Aeromonas-media:0.0702305952,WP_139406182.1-Aeromonas-veronii:0.1882297740)89.6/94:0.0437062173)89.2/95:0.0241744277,((WP_268421127.1-Aeromonas-media:0.0443620032,(WP_111910157.1-Aeromonas:0.0247535271,WP_128343177.1-Aeromonas-caviae:0.0487731104)31.3/61:0.0143071336)95.5/100:0.0388640639,(WP_268037199.1-Aeromonas-dhakensis:0.0803842009,((WP_139733908.1-Aeromonas-veronii:0.0235355482,(WP_234676856.1-Aeromonas-rivipollensis:0.0155676389,WP_269382839.1-Aeromonas-hydrophila:0.0696113131)75.1/100:0.0104666540)70.3/98:0.0256392662,WP_234579942.1-Aeromonas-salmonicida:0.0364667990)95.6/100:0.0587391278)70.7/98:0.0165234189)69.4/99:0.0275471991)82.8/93:0.0223881848,WP_224211331.1-Aeromonas-enteropelogenes:0.1934171274)78.8/82:0.0183796269,(WP_253626250.1-Aeromonas-hydrophila:0.0649919375,((WP_073350970.1-Aeromonas-hydrophila:0.0643054062,((WP_241344946.1-Aeromonas-sp.-MR16:0.0591208550,WP_190284026.1-Aeromonas-hydrophila:0.0613215744)39.4/58:0.0193561912,(WP_223952962.1-Aeromonas-caviae:0.1273718006,WP_244081328.1-Aeromonas-caviae:0.1412375179)94.8/100:0.0655108434)82.2/93:0.0535299005)65.4/98:0.0083490343,WP_082041448.1-Aeromonas-eucrenophila:0.1717584157)98.4/98:0.0774422685)95.3/96:0.0449064346)85.6/91:0.0261591086)90.4/94:0.0400839775,(((WP_208005626.1-Oceanisphaera-pacifica:0.0612494621,(WP_087034853.1-Oceanisphaera-profunda:0.0637887605,WP_087034873.1-Oceanisphaera-profunda:0.0864944123)91.1/100:0.0583422007)96.3/100:0.0758967880,WP_200561028.1-unclassified-Erwinia:0.2037119145)88.5/95:0.0582803506,((WP_076573677.1-Aeromonas-sp.-RU39B:0.1037634563,WP_134696535.1-Aeromonas-taiwanensis:0.3984507163)97.3/100:0.0913555378,WP_108652490.1-Dongshaea-marina:0.1236438579)91.7/95:0.0558360191)55.2/97:0.0382361039)67.5/77:0.0091637796)85.8/84:0.0323710155)93/89:0.0389360526)68.4/79:0.0190725470,WP_202667543.1-Lelliottia-amnigena:0.1558445512)98.9/98:0.0862911061,((WP_241647798.1-Rosenbergiella-sp.-JB07:0.2225980210,WP_171149131.1-Erwinia-sp.-JH02:0.1812584207)0/11:0.0122368600,WP_181599127.1-Citrobacter-sp.-RHB36-C18:0.1558945754)70.6/11:0.0208724652)41/82:0.0444183141,WP_065370581.1-Kosakonia-sacchari:0.1472434744)93/99:0.0484086021)21.7/35:0.0102230733,((((((((((((((WP_182931197.1-Enterobacter-cloacae:0.1191922328,WP_048284323.1-Pluralibacter-gergoviae:0.1128754762)2.9/29:0.0139796360,WP_211776196.1-Klebsiella-pneumoniae:0.0513449049)73.1/32:0.0162694204,(WP_047723323.1-Klebsiella-michiganensis:0.0364801208,WP_104011323.1-Klebsiella-michiganensis:0.0394340849)7.8/46:0.0094244774)88.6/39:0.0202819896,((WP_139968923.1-Klebsiella-pneumoniae:0.0778195024,WP_065799756.1-Klebsiella-pneumoniae:0.0055326314)0/62:0.0000022409,WP_095100910.1-Raoultella-sp.-10-1:0.0620108123)88.1/70:0.0178534186)86.5/68:0.0182765928,WP_149270444.1-Klebsiella-aerogenes:0.0246867304)66.4/48:0.0223949736,WP_046852082.1-Raoultella-planticola:0.0267219347)84.9/50:0.0129311730,WP_123755035.1-Raoultella-sp.-BIGb0399:0.0279548446)72.9/41:0.0112444414,WP_114692711.1-Klebsiella-pneumoniae:0.0446253751)0/33:0.0000025937,(((WP_221694709.1-Klebsiella-aerogenes:0.0733792697,WP_213760001.1-Klebsiella-aerogenes:0.0546793682)62.2/55:0.0111573324,WP_223370324.1-Klebsiella-pneumoniae:0.0310706679)88.4/55:0.0198879199,WP_128320364.1-Raoultella-planticola:0.0407264793)87.4/40:0.0113451364)75/66:0.0055977726,WP_236948387.1-Klebsiella-sp.-2019SCSN059:0.0279786352)85.8/71:0.0110616064,WP_259891680.1-Klebsiella-variicola:0.0221641078)69/92:0.0056336299,((WP_234537246.1-Raoultella-ornithinolytica:0.0462408218,WP_113990968.1-Raoultella:0.0112254746)90.7/100:0.0336834333,(WP_234579544.1-Raoultella-terrigena:0.0398356261,WP_186601692.1-Klebsiella-quasipneumoniae:0.0168090183)92.5/96:0.0257526488)62.1/90:0.0030308716)97.8/93:0.0558201219,((WP_095100919.1-Raoultella-sp.-10-1:0.0414893085,WP_212709529.1-Tatumella-sp.-JGM130:0.0173913689)99/100:0.0586577440,WP_236948353.1-Klebsiella-sp.-2019SCSN059:0.0724130164)53.8/91:0.0172442492)91/98:0.0404789413,WP_123822891.1-Klebsiella:0.0343678962)98.1/99:0.0683204168)90.8/90:0.0406295084,WP_072663757.1-Enterobacteriaceae:0.0702660627)12.5/54:0.0295010295,WP_021241446.1-Enterobacter-roggenkampii:0.1392294919)96.5/100:0.0690649168)95.9/100:0.0551235411,XP_011503919.1-Ceratosolen-solmsi-marchali:0.0737941004);

**Nematode RML-3, with bacterial and eukaryotic homologs**

**Shown in Figure S2**

ML tree method: IQ-TREE

Sequences: 2787

Substitution model: LG+I+G4

(WP_063501760.1-Escherichia-coli:0.0480676898,(WP_044173167.1-Phytobacter-massiliensis:0.0499699057,(((((((WP_203451401.1-Lelliottia-sp.-RWM.1:0.0839676582,WP_185654835.1-Citrobacter-braakii:0.1421728469)91.6/100:0.1020786194,WP_183170804.1-Atlantibacter-sp.-RC6:0.2834731437)96.3/100:0.1285348782,((((((((((((((((((((((((((((WP_249615874.1-Craterilacuibacter-sp.-RT1T:0.7714470834,(WP_092579398.1-Achromobacter-sp.-NFACC18-2:0.5536715647,WP_223462047.1-Pseudomonas-sp.-A-R-26:0.2395764040)21.6/88:0.0553307291)0/75:0.0160549812,WP_058422493.1-Pseudomonas:0.2511463998)87.7/95:0.0600590319,WP_019141909.1-Noviherbaspirillum-massiliense:0.2688940236)92.9/43:0.0877783785,((((((((WP_250059255.1-Halomonas-sp.-ATCH28:0.4424888309,WP_179153381.1-Chromohalobacter-salexigens:0.2615015167)91.4/100:0.1238918808,WP_250939176.1-unclassified-Larsenimonas:0.3262746027)85.3/100:0.0732256539,WP_106478747.1-Phytohalomonas-tamaricis:0.3548539269)81.8/97:0.0637023134,(WP_058575749.1-unclassified-Guyparkeria:0.3910404695,(WP_218634960.1-Marinobacter-arenosus:0.1424419216,WP_240716217.1-Halomonas-neptunia:0.1261312132)95.7/100:0.1448619037)77.7/97:0.0506131688)96.6/100:0.1164965072,((((((((WP_269022943.1-Methylomonas-sp.-MP1:0.1287540083,(WP_140911374.1-Methylomonas-koyamae:0.2764965919,(WP_071159776.1-Methylomonas-sp.-LWB:0.2071711230,((WP_020485671.1-Methylomonas-sp.-MK1:0.2280849407,(WP_256607223.1-Methylomonas-sp.-WSC-7:0.0910120915,(WP_205450646.1-Methylomonas-sp.-EFPC1:0.1010680500,WP_064036612.1-Methylomonas-methanica:0.0000029332)0/95:0.0051792114)90.3/100:0.0836233736)91.3/96:0.0824649685,WP_036273427.1-Methylomonas:0.1520490379)0/73:0.0000020887)39.9/100:0.0384946563)91.8/100:0.1148540113)23/86:0.0630982149,(WP_196437371.1-Methylomonas-sp.-LL1:0.1555379651,(WP_150047608.1-Methylomonas-rhizoryzae:0.3895314659,WP_066983474.1-Methylomonas-lenta:0.1073063386)86.4/82:0.0723205152)14.5/67:0.0340872286)93.3/100:0.1406252283,WP_013164002.1-Desulfurivibrio-alkaliphilus:0.4039740552)49.5/22:0.0575934889,WP_138632053.1-Pseudoalteromonas-spongiae:0.4848963446)68.2/17:0.0541257565,(((((WP_136386324.1-Pseudothauera-rhizosphaerae:0.2157516293,(WP_114649035.1-Pseudothauera-hydrothermalis:0.1356673018,WP_187718084.1-Thauera-sp.-CAU-1555:0.1770250160)78.7/91:0.0640455753)98.8/100:0.2453467327,WP_026685886.1-Azovibrio-restrictus:0.2138073069)78.1/80:0.0494102202,(((WP_148817865.1-Pigmentiphaga-aceris:0.4797046065,(WP_238617884.1-Ectothiorhodospira-shaposhnikovii:0.1814820286,WP_025282248.1-Ectothiorhodospira-haloalkaliphila:0.1541073508)85.2/100:0.0432313597)82.7/100:0.0431353226,WP_238622182.1-unclassified-Ectothiorhodospira:0.1723573141)93.8/100:0.0780744371,((WP_256068944.1-Stutzerimonas-stutzeri:0.2583449839,((((((WP_060482506.1-Pseudomonas-sp.-NBRC-111119:0.2548801099,((WP_196129978.1-Pseudomonas-fulva:0.0983583785,WP_201164743.1-Pseudomonas-sp.-S60:0.0567057104)97.7/100:0.0965002183,(((WP_174142546.1-Pseudomonas-putida:0.1682540420,WP_027907237.1-Pseudomonas:0.0497665974)63.9/100:0.0169058506,WP_075044077.1-Pseudomonas-putida:0.0520911685)73.8/100:0.0389757993,(WP_023379297.1-Pseudomonas-sp.-VLB120:0.1175837062,WP_202675963.1-Pseudomonas-sp.-SK2:0.0437990230)97/100:0.0806736147)89.3/100:0.0488774895)85.2/100:0.0277838819)78/64:0.0263120348,(((WP_159409308.1-Pseudomonas-putida:0.1763200880,(((WP_182365203.1-Pseudomonas-juntendi:0.0875134349,WP_182365213.1-Pseudomonas:0.1317992918)76.9/99:0.0197930206,((WP_110607173.1-Pseudomonas-fulva:0.0164926841,WP_088523134.1-unclassified-Pseudomonas:0.0618094272)95.2/100:0.0476726024,(WP_167339957.1-Pseudomonas-fulva:0.0378941200,WP_186476580.1-Pseudomonas-promysalinigenes:0.0526525152)87.6/100:0.0255022641)80.9/45:0.0195906768)84.1/45:0.0196703224,(WP_201214098.1-Pseudomonas-sp.-S36:0.0253589221,WP_054890059.1-Pseudomonas:0.0764779082)96.7/100:0.0611429547)64.1/88:0.0062261514)48/26:0.0120482443,(WP_167277242.1-Pantoea-sp.-Tr-811:0.0237347324,WP_103449615.1-Pseudomonas-putida:0.0670251178)97.4/100:0.0709974428)0/26:0.0000023982,(WP_115272924.1-Pseudomonas-putida:0.1676941285,WP_267821885.1-Pseudomonas-putida:0.0629954118)89.3/91:0.0435721464)69.4/63:0.0166386113)87.7/98:0.0293735160,(WP_046489762.1-Pseudomonas-veronii:0.1344594891,(WP_059395284.1-Pseudomonas-putida:0.1108067296,(WP_009685980.1-Pseudomonas:0.0934085766,WP_024086460.1-Pseudomonas:0.0784285748)85.6/100:0.0530560838)85.9/100:0.0526537416)72.3/98:0.0695833875)76.4/70:0.0155337858,(WP_029885231.1-Pseudomonas:0.0274767882,(WP_230064532.1-Pseudomonas-sp.-CIP-10:0.0256485874,WP_197863301.1-Pseudomonas-putida:0.0507263000)96.3/100:0.0578378661)73.5/70:0.0069025766)95.2/69:0.0744108942,WP_060394622.1-Pseudomonas-entomophila:0.1467479975)97.7/100:0.1033003523,WP_183165449.1-Azomonas-macrocytogenes:0.1516898989)82.7/99:0.0340608247)42.8/79:0.0415616341,WP_004074545.1-Desulfobacter-postgatei:0.4029481557)52/34:0.0119854705)87.7/50:0.0652816422)82.1/63:0.0319941915,WP_168987795.1-Azoarcus-taiwanensis:0.3458964331)77.2/24:0.0219875391,WP_018865194.1-Thioalkalivibrio-sp.-ARh3:0.2671651054)87.2/16:0.0370998097)74.1/28:0.0163464063,WP_048706771.1-Thauera-humireducens:0.4473118434)91.4/66:0.0608489872,(((((WP_167682126.1-Rhodocyclus-gracilis:0.4288778380,(WP_153115862.1-Rhodocyclus-tenuis:0.0640799053,WP_200385811.1-Rhodocyclus-tenuis:0.0219697170)93/100:0.1138913049)95.6/100:0.1624629710,WP_201212349.1-Rhodocyclus-purpureus:0.1156915872)96.7/100:0.1617093347,WP_153110537.1-Propionivibrio-limicola:0.3487315876)69.5/67:0.0827787139,WP_153720048.1-Spiribacter-sp.-C176:0.3855232857)91.1/100:0.1134296047,(WP_263768642.1-Propionivibrio-sp.-SG131:0.3708957345,(WP_033955816.1-Pseudomonas-aeruginosa:0.0030405124,WP_003153917.1-Pseudomonas-aeruginosa-group:0.0782888766)99.5/100:0.2455637749)40.4/72:0.0649762875)90.2/66:0.0770395734)79.9/84:0.0374812217,(WP_209394989.1-Rhodoferax-sp.-AJA081-3:0.3925175849,WP_094844257.1-Bordetella-genomosp.-11:0.2221861380)92.5/86:0.1143308293)90.2/100:0.0549696589)89.6/99:0.0458964855,((((((((WP_225237063.1-Massilia-oculi:0.4257634708,WP_147937383.1-Massilia-arenae:0.1467497013)98.7/100:0.2420697414,WP_192051557.1-unclassified-Massilia:0.1210839884)14.6/72:0.0400653676,WP_136222466.1-Massilia-sp.-Mn16-1_5:0.1179370519)98.6/99:0.2094511706,WP_214501190.1-Pseudomonas-sp.-RC3H12:0.3201260799)76.6/80:0.0395029116,((WP_077733980.1-Paenalcaligenes-hominis:0.0706908098,WP_167661700.1-Paenalcaligenes-hominis:0.0207123410)100/100:0.3306442443,((((((WP_191471912.1-Pseudomonas-asiatica:0.2541401513,(WP_043318452.1-Pseudomonas-sp.-PI1:0.0501773726,WP_043268312.1-Pseudomonas:0.0436150728)34.6/100:0.0246640867)93/100:0.0819368516,(WP_089390899.1-Pseudomonas-delhiensis:0.1451487285,WP_076426141.1-Pseudomonas-alcaligenes:0.2009577284)51.9/100:0.0219810289)96.1/100:0.0897044831,WP_233297479.1-Pseudomonas:0.2181538558)62.7/97:0.0240633820,(WP_184594840.1-Pseudomonas-nitritireducens:0.2238053971,WP_192381882.1-Pseudomonas-sp.-PDM20:0.1470153979)99.3/97:0.1883235400)77.3/93:0.0339978628,(WP_058488093.1-Pseudomonas:0.1897247243,(WP_043256200.1-Pseudomonas-knackmussii:0.1711271884,((WP_088420173.1-Pseudomonas-nitroreducens:0.1559242240,WP_254475170.1-unclassified-Pseudomonas:0.0898530610)12/100:0.0308387090,WP_205338011.1-Pseudomonas-sp.-PDNC002:0.0850370822)99.5/100:0.1856474688)17.7/64:0.0483911171)95.5/94:0.1029765125)76.9/88:0.0101221336,WP_249918475.1-Pseudomonas-sp.-R3.Fl:0.3059535153)70.1/83:0.0067594741)70.4/70:0.0100526996)90.9/94:0.0277474427,(((WP_065086507.1-Pseudomonas:0.3329099204,WP_205489922.1-Pseudomonas-lactucae:0.2065678316)82.8/68:0.0530744556,((WP_169965430.1-unclassified-Pseudomonas:0.0578921682,WP_047278503.1-Pseudomonas-lundensis:0.0321045943)98.8/100:0.1493101251,(WP_104727848.1-Pseudomonas-oleovorans:0.1551938235,WP_085721622.1-Pseudomonas:0.2536752104)90.7/94:0.0765722541)74/82:0.0218876418)6.9/51:0.0275348546,(WP_042945119.1-Pseudomonas-fluorescens:0.0916748406,WP_181643258.1-Pseudomonas-sivasensis:0.0646972435)99.5/100:0.1835898156)77.8/92:0.0237194642)83.6/85:0.0300589359,((WP_076518421.1-Achromobacter-sp.-MFA1-R4:0.1069826477,(WP_104142565.1-Achromobacter-spanius:0.0280781448,WP_120157261.1-Achromobacter-sp.-B7:0.0667789452)85.9/53:0.0511578952)64/53:0.0616242308,WP_158658301.1-Achromobacter-sp.-AONIH1:0.5022065302)96.3/96:0.1405956708)54.8/38:0.0116334611,((WP_209872004.1-Pseudomonas-sp.-BP8:0.2232022769,WP_066093302.1-Hydrogenophaga-crassostreae:0.2150922192)89.8/98:0.0871174587,((WP_177138191.1-Hydrogenophaga-aromaticivorans:0.1682723411,WP_133157403.1-Hydrogenophaga-pseudoflava:0.2060784572)77.7/97:0.0435360906,(WP_236964528.1-Hydrogenophaga-sp.-SL48:0.1015538536,(WP_226493830.1-Hydrogenophaga-taeniospiralis:0.0509613098,WP_210824367.1-Hydrogenophaga-aromaticivorans:0.0800730552)51.7/100:0.0511529518)94.1/100:0.0880870510)89.8/99:0.0763654430)74.2/23:0.0646085085)76.1/23:0.0109881144)87.1/84:0.0242477349,((((WP_020564242.1-Methylosarcina-fibrata:0.2497223002,(WP_074973890.1-Pseudomonas-otitidis:0.1959338463,WP_068803504.1-Immundisolibacter-cernigliae:0.2857145873)83/94:0.0739611846)0/74:0.0705192333,(((WP_180152801.1-Parapusillimonas-granuli:0.1927163887,(WP_088146851.1-Achromobacter-denitrificans:0.1322242390,WP_105555808.1-Achromobacter-sp.-MYb9:0.1021527226)98/100:0.1115459760)77.6/100:0.0314041329,WP_011111299.1-Nitrosomonas-europaea:0.2527969766)44.3/98:0.0373296851,WP_046857648.1-Pseudomonas-sp.-CCOS-191:0.2710089339)77.7/96:0.0421003630)75.9/19:0.0304927868,(WP_101152997.1-Pseudomonas-sp.-Choline-3u-10:0.1589316886,((WP_046617631.1-Stutzerimonas-stutzeri:0.0879765499,WP_135265705.1-Stutzerimonas-stutzeri:0.0655828825)87.8/100:0.0802304553,WP_003299494.1-Stutzerimonas-stutzeri:0.1159884552)88.3/100:0.0428921670)94.2/100:0.0752414027)49.7/17:0.0246931315,(WP_125865093.1-Stutzerimonas-stutzeri:0.1983568786,WP_200290630.1-Stutzerimonas-balearica:0.1742825318)94.7/98:0.1079756832)76.6/21:0.0245534561)89.8/23:0.0376694766,((((((((WP_044615503.1-Gynuella-sunshinyii:0.4839638679,WP_227683402.1-Pusillimonas-sp.-MFBS29:0.2853105766)23.3/69:0.1039475730,WP_136867012.1-Guyparkeria-sp.-SB14A:0.4456261506)78.8/70:0.0550639618,((WP_019964198.1-Pannonibacter-phragmitetus:0.8918440922,WP_064030819.1-Methylomonas-koyamae:0.2038172477)83.2/100:0.1011043932,(WP_138213821.1-Pseudomonas-nitroreducens:0.0398270420,WP_017517140.1-Pseudomonas-nitroreducens:0.0340102763)100/100:0.3631779788)87.1/100:0.0650787869)75.5/70:0.0221222277,WP_102075761.1-Pollutimonas-subterranea:0.2010604444)87.5/100:0.0596996714,WP_102071501.1-Pollutimonas-nitritireducens:0.3771521568)80.3/96:0.0417528949,(WP_066098770.1-Halothiobacillus-sp.-LS2:0.1583716362,WP_070073009.1-Acidihalobacter-aeolianus:0.2947421703)92.6/97:0.1396089572)76.4/98:0.0185204601,((((WP_057247240.1-Duganella-sp.-Root198D2:0.2471195415,WP_212592286.1-Janthinobacterium-sp.-TND4EL3:0.3421016847)71.7/72:0.0696958503,WP_155457065.1-Massilia-eburnea:0.1754560071)88.6/71:0.0840550282,WP_217963608.1-Candidimonas-humi:0.3158813251)30.4/42:0.0583731403,WP_047550623.1-Methylotenera-sp.-G11:0.3067755866)84.4/42:0.0460443681)33/41:0.0252690429,(((WP_029241333.1-Pseudomonas:0.0781505861,((WP_016973326.1-Pseudomonas-tolaasii:0.1508763016,WP_099548838.1-Pseudomonas-sp.-2995-3:0.1197964795)28.7/100:0.0434502715,(WP_177015116.1-Pseudomonas-costantinii:0.0685104669,WP_141063556.1-Pseudomonas-fluorescens:0.1186858099)93.8/100:0.0816824424)99.1/100:0.1646808478)44.2/77:0.0512692569,((WP_054996663.1-Pseudomonas-ficuserectae:0.0510647953,(WP_007252556.1-Pseudomonas-syringae-group:0.0580924770,WP_025390602.1-Pseudomonas-syringae:0.0329191265)0/74:0.0218138620)94.1/97:0.0634543166,WP_125830676.1-Pseudomonas-sp.-v388:0.0824047126)61.3/31:0.0261315331)99.5/98:0.2005188211,WP_146330669.1-Noviherbaspirillum-sp.-UKPF54:0.3183496928)89.9/40:0.0636856719)81.7/33:0.0244098199)57.5/15:0.0114982092)88.7/79:0.0410069432,((((WP_139871681.1-Candidatus-Methylopumilus-universalis:0.3083255184,(WP_139929800.1-Candidatus-Methylopumilus-universalis:0.3583176780,WP_140005423.1-Candidatus-Methylopumilus-planktonicus:0.0985107908)82.4/100:0.0936365161)99.8/100:0.3556535638,(((WP_220354719.1-Rhizobium-cauense:0.3118969871,WP_016557527.1-Rhizobium-grahamii:0.1668145210)84.1/84:0.0634413252,((WP_184450570.1-Rhizobium-sp.-BK538:0.2608885211,(WP_072373837.1-Rhizobium-tibeticum:0.1218591811,WP_007537558.1-Rhizobium-mesoamericanum:0.0667249084)86.1/99:0.0622154161)55/97:0.0318259324,WP_018896487.1-Rhizobium-sp.-2MFCol3.1:0.2698984581)76.7/91:0.0259961285)76.6/79:0.0317558750,(WP_153273632.1-Rhizobium-grahamii:0.2836744181,WP_233136718.1-Agrobacterium-vaccinii:0.3250550438)68.1/88:0.0557954515)92.1/99:0.1417247873)88.3/98:0.1233624674,WP_230502293.1-Janthinobacterium-sp.-FW305-128:0.6372421678)74.8/20:0.0418997320,WP_216129682.1-Azospira-inquinata:0.5297503365)73.5/21:0.0144328991)90.5/40:0.0425997182,(((WP_046292890.1-Pandoraea-oxalativorans:0.1050147314,WP_150719538.1-Pandoraea-capi:0.1209481890)99.8/100:0.4197536926,WP_101249932.1-Telmatospirillum-siberiense:0.6247787728)89.1/43:0.1813498612,((((WP_129151777.1-Achromobacter-aloeverae:0.1198629181,WP_088599355.1-unclassified-Achromobacter:0.0849651547)96.4/100:0.1141210353,WP_057654143.1-Bordetella-sp.-N:0.2436888676)88.4/100:0.0751882070,WP_086067427.1-Bordetella:0.2762233203)91.1/100:0.0709720741,(WP_066641528.1-Bordetella-sp.-H567:0.1308146307,(WP_086058993.1-Bordetella-genomosp.-9:0.2532593671,WP_094856450.1-Bordetella-genomosp.-10:0.1755975660)81.2/100:0.1006340722)87.3/100:0.1016393452)94.5/99:0.1366464407)77.4/41:0.0346407699)76.1/75:0.0183780734,((((((((((WP_072282603.1-Syntrophotalea-acetylenivorans:0.1747812189,(WP_092056970.1-Desulfuromonas-acetexigens:0.2063786032,(((WP_028317018.1-Desulfobulbus-elongatus:0.4600472160,WP_155876508.1-Desulfuromonas-sp.-AOP6:0.3163553221)83.6/96:0.0851035901,WP_043770486.1-Desulfobulbus-propionicus:0.3063265001)72.2/88:0.0680250481,WP_148894253.1-Geothermobacter-ehrlichii:0.4941786148)73.5/89:0.0801120953)78.4/87:0.0709995863)98.1/100:0.1786547763,WP_053550565.1-Desulfuromonas-soudanensis:0.3175297240)64.8/69:0.0443089039,WP_027714505.1-Desulfuromonas-sp.-TF:0.3752289909)67.2/68:0.0269822254,WP_072282412.1-Syntrophotalea-acetylenivorans:0.3202697776)79.3/90:0.0284100213,((WP_250680171.1-Geomonas-sp.-Red32:0.2524235625,WP_199381999.1-Geomesophilobacter-sediminis:0.2724855782)87.9/87:0.0928854570,WP_205218910.1-Desulforhopalus-vacuolatus:0.4264899381)91/65:0.0955546723)77.1/25:0.0374332679,(WP_103114677.1-Geothermobacter-hydrogeniphilus:0.4266490977,WP_221248821.1-Desulfuromonas-versatilis:0.3507431509)91.4/95:0.1445922989)0.5/11:0.0271330030,((WP_015405619.1-Desulfocapsa-sulfexigens:0.2988609912,(((WP_163339734.1-Desulfopila-sp.-IMCC35008:0.3965645930,WP_136805251.1-Desulfosediminicola-flagellatus:0.3463626080)0/74:0.0000021073,WP_073615262.1-Desulfopila-aestuarii:0.3238487340)96.8/100:0.1523611292,(WP_228856481.1-Desulfomarina-profundi:0.4206667892,WP_183349429.1-Desulfoprunum-benzoelyticum:0.2397820322)79.4/100:0.0502337934)85.5/97:0.0696504201)85.3/96:0.0368413218,((WP_205222281.1-Desulfogranum-marinum:0.2852959535,(WP_205242362.1-Desulfobulbus-alkaliphilus:0.1987136682,WP_267928740.1-Desulfolithobacter-dissulfuricans:0.2160374360)53.7/97:0.0603140068)63.9/100:0.0787934472,WP_028582690.1-Desulfogranum-japonicum:0.3535888093)76.9/99:0.0823822433)90/93:0.0799237641)61.4/11:0.0170531326,(WP_137423090.1-Desulfoglaeba-alkanexedens:0.4565668520,WP_040200185.1-Geoalkalibacter-subterraneus:0.3270993539)70/82:0.1006654355)90.7/30:0.0473009287,((((((((((((((WP_089129802.1-Tolypothrix-sp.-NIES-4075:0.1088184505,WP_096563566.1-Scytonema-sp.-NIES-4073:0.1475618501)83.9/99:0.0469471758,WP_048866388.1-Scytonema-tolypothrichoides:0.1183288975)5.5/95:0.0205102258,WP_096563464.1-Scytonema-sp.-NIES-4073:0.1299575104)100/100:0.2516308856,((WP_015128325.1-Calothrix-sp.-PCC-7507:0.1329262720,WP_017653186.1-Fortiea-contorta:0.2307652543)65.5/100:0.0625199486,((WP_193199005.1-Nostoc-sp.-MG11:0.0489879814,WP_162396505.1-Nostoc-sp.-B-2019:0.0630781902)94/100:0.0750760321,WP_214439445.1-Atlanticothrix-silvestris:0.1702893318)75.8/100:0.0877615538)49.9/100:0.0252015610)27.2/25:0.0126249905,(((WP_045870014.1-unclassified-Tolypothrix:0.0333218789,WP_190656031.1-Nostoc-linckia:0.0435543345)98.2/100:0.0971509668,(WP_190743070.1-Tolypothrix-sp.-FACHB-123:0.0580030028,WP_190545010.1-Calothrix:0.0416963578)96/100:0.0893650112)73.6/100:0.0384020819,WP_086758595.1-Nostoc-sp.-106C:0.0793540865)87.3/96:0.0479896953)0/25:0.0042194548,((((((WP_009344618.1-Cylindrospermopsis-raciborskii:0.0912314510,WP_240868223.1-Cylindrospermopsis-raciborskii:0.0284585240)99.5/100:0.1717079734,WP_096570274.1-Sphaerospermopsis:0.0678001870)4.8/96:0.0118207314,WP_220611028.1-Sphaerospermopsis-torques-reginae:0.0653190057)98.9/100:0.1219378895,((WP_237989437.1-Nostoc-sp.-UHCC-0870:0.0880991188,WP_190559626.1-Anabaena-sphaerica:0.1197968506)54.2/98:0.0446944309,WP_206263219.1-Nostoc-sp.-UHCC-0702:0.0987307705)43.5/99:0.0342011791)69.2/100:0.0309526695,WP_096557790.1-Nostoc-sp.-NIES-4103:0.2062039588)36.8/25:0.0467900216,WP_016866333.1-Fischerella:0.1058609814)66.6/24:0.0489768550)79/89:0.0749161652,(WP_044881819.1-Neochlamydia-sp.-EPS4:0.0742588421,WP_166155173.1-unclassified-Neochlamydia:0.0000024347)100/100:0.5964603777)93/59:0.0950487229,WP_015143663.1-Pleurocapsa-sp.-PCC-7327:0.1868902159)67.8/45:0.0483767051,((((((((((WP_102179481.1-Fischerella-thermalis:0.0345637816,((WP_096675774.1-Fischerella-sp.-NIES-4106:0.0613213941,WP_053458209.1-Hapalosiphon-sp.-MRB220:0.0398685261)73.6/100:0.0079487833,(WP_131119525.1-Westiellopsis-prolifica:0.0595509813,WP_016860528.1-Fischerella-muscicola:0.0316059731)84.9/100:0.0170527193)95.5/100:0.0901661151)99.9/100:0.2433546631,(WP_190825375.1-Iningainema-tapete:0.3009954376,(WP_015202791.1-Crinalium-epipsammum:0.2794642448,WP_035154029.1-Calothrix-sp.-336/3:0.4533772133)80.1/93:0.0803489880)15.3/66:0.0263374120)85.7/66:0.0370702669,((WP_099102643.1-Nostoc-sp.-Peltigera-malacea-cyanobiont-DB3992:0.0961133416,((WP_190950926.1-Nostoc-punctiforme:0.0569511248,(WP_221704991.1-Desmonostoc-muscorum:0.0569483382,WP_096537650.1-Nostoc-linckia:0.0204506070)75.9/99:0.0082411479)48/96:0.0307275107,WP_190959861.1-Nostoc:0.0223176139)98.4/99:0.1047316025)99.2/100:0.1475087397,((((((WP_193938462.1-Cuspidothrix-issatschenkoi:0.1828534615,WP_168537172.1-Anabaena-sp.-UHCC-0253:0.0461485288)87.3/67:0.0334795133,(((WP_190370804.1-Anabaena-sp.-FACHB-1237:0.1908192458,(WP_190392428.1-Dolichospermum-sp.-FACHB-1091:0.0138101955,(WP_028089231.1-Dolichospermum-circinale:0.0232770757,WP_137908967.1-Nostocales:0.0599857355)99.1/100:0.0760692888)98.2/100:0.1040161491)90.4/87:0.0478234433,((WP_096666215.1-Dolichospermum-compactum:0.0757187726,(WP_190387835.1-Aphanizomenon:0.0748435940,WP_168466156.1-Aphanizomenon:0.0233912000)41.2/100:0.0168261191)65.2/98:0.0190840095,WP_015080416.1-Anabaena-sp.-90:0.0418117485)92.5/99:0.0325559779)52/68:0.0078485500,WP_168502193.1-Anabaena-sp.-UHCC-0187:0.0073751450)78.9/67:0.0229549966)99.9/100:0.1462796168,(WP_010998622.1-Nostocaceae:0.0742111191,WP_011320078.1-Nostocaceae:0.0713162251)77.2/100:0.0278580846)91.7/94:0.0547297701,((WP_015138800.1-Nostoc-sp.-PCC-7524:0.0895382440,WP_066425233.1-Anabaena-sp.-4-3:0.1120595676)98/100:0.0850709403,(WP_199337214.1-Nostoc-sp.-FACHB-152:0.1412310217,(WP_239729951.1-Nodularia-sphaerocarpa:0.0831670039,WP_173403323.1-Nodularia-spumigena:0.0571764783)94.5/100:0.0552181908)78/81:0.0296736888)74.9/78:0.0095460402)81.2/94:0.0196226508,WP_199337786.1-Nostoc-sp.-FACHB-280:0.1670613441)95.8/100:0.0808256026,(WP_096624129.1-Calothrix-sp.-NIES-3974:0.0920742850,WP_095721999.1-Calothrix-elsteri:0.1201294594)98.6/99:0.1358484924)78/99:0.0267728086)95.2/99:0.0655233910)93.2/97:0.0461262198,((WP_194029353.1-Lusitaniella-coriacea:0.2482231796,WP_204103763.1-Spirulina-major:0.6391598529)90.4/98:0.0790586136,(WP_224410051.1-Oscillatoria-salina:0.2611596200,WP_250123015.1-Chroococcidiopsis-sp.-CCMEE-29:0.1889694692)88/96:0.0546050713)89.3/86:0.0333111468)84.5/86:0.0279398098,((((WP_233747769.1-Leptodesmis-sichuanensis:0.2729936095,WP_223049852.1-Leptothermofonsia-sichuanensis:0.1235595210)81.6/84:0.0228544634,WP_224345144.1-Kovacikia-minuta:0.1487123532)6.8/54:0.0159662038,WP_192216758.1-Microcoleus-sp.-FACHB-SPT15:0.2113553299)85.8/79:0.0308829990,WP_073549891.1-Chroogloeocystis-siderophila:0.2316757779)85.5/84:0.0440174808)74.1/48:0.0120371047,((WP_193873404.1-unclassified-Coleofasciculus:0.1576639517,WP_254010234.1-Symplocastrum-sp.-BBK-W-15:0.1575359889)90.2/88:0.0556023971,(WP_190666400.1-Coleofasciculus-sp.-FACHB-501:0.1528755483,WP_190525322.1-Coleofasciculus-sp.-FACHB-T130:0.0710886979)95.6/100:0.0805147828)82.6/36:0.0190114846)81/30:0.0225955984,(WP_008178350.1-Moorena-producens:0.2573790156,WP_190493486.1-unclassified-Microcoleus:0.3123486495)84.8/36:0.0600635239)77.3/32:0.0196856189,(WP_226578226.1-Microseira-wollei:0.2293386039,((WP_073595522.1-Phormidium-ambiguum:0.0871847035,WP_193982193.1-Phormidium-sp.-LEGE-05292:0.0876388854)99.5/100:0.1363660160,(WP_249064905.1-Argonema-antarcticum:0.0711062384,WP_190468950.1-unclassified-Planktothrix:0.1042000914)36.1/97:0.0205929853)89.9/100:0.0467757168)94/99:0.0674996453)87.7/22:0.0273676611,(((((((((WP_202221768.1-Okeania-sp.-KiyG1:0.0565527313,WP_094672483.1-Hydrocoleum-sp.-CS-953:0.0538359929)75.3/100:0.0222715342,WP_124146462.1-Okeania-hirsuta:0.0051489186)99.5/100:0.2172180200,WP_071516849.1-Geitlerinema-sp.-PCC-9228:0.2836765751)92.7/99:0.1011117082,((WP_008052766.1-Limnospira-indica:0.3196340073,(WP_083623900.1-Planktothrix-serta:0.0397862964,(WP_072717475.1-Planktothrix:0.0222527316,(WP_079676747.1-Planktothrix-sp.-PCC-11201:0.0285065627,(WP_026797127.1-Planktothrix:0.0314137521,WP_214559879.1-Planktothrix-agardhii:0.0504276653)69.6/100:0.0343730057)98.8/100:0.1152669613)35.5/71:0.0194152448)97.6/100:0.1110783790)27.2/52:0.0205631096,(WP_009783380.1-Lyngbya:0.0980025035,WP_046277942.1-Limnoraphis-robusta:0.0640505797)97.1/100:0.0980994844)10.8/24:0.0336456379)85.9/100:0.0418705255,WP_190877489.1-unclassified-Planktothricoides:0.1980497067)88.2/99:0.0447112114,(((((WP_236507253.1-Tychonema-sp.-BBK16:0.0483002755,WP_096828456.1-Tychonema-bourrellyi:0.0807483341)93.5/100:0.0418598040,WP_242724542.1-Microcoleus-vaginatus:0.0408788961)80.3/100:0.0145199067,WP_015174083.1-Oscillatoria-nigro-viridis:0.0231050604)88.3/100:0.0232129528,(WP_194009709.1-unclassified-Tychonema:0.0447317920,WP_265235823.1-Lyngbya-sp.-CCAP-1446/10:0.0517434524)89.7/100:0.0262468078)47.1/88:0.0215470307,WP_193973976.1-Microcoleus-sp.-LEGE-07076:0.0409101075)99.4/100:0.1208305037)87/49:0.0441848687,((WP_261197618.1-unclassified-Laspinema:0.0514877958,(WP_207087219.1-Phormidium-pseudopriestleyi:0.0409755383,WP_261204505.1-Laspinema-sp.-D2d:0.0498822017)76.8/100:0.0267832773)98.6/100:0.1323389858,WP_200665545.1-Oxynema-sp.-CENA135:0.2753455147)87.8/99:0.0403527442)78.6/49:0.0141089604,((WP_017715496.1-Oscillatoria-sp.-PCC-10802:0.2075884426,(WP_190665495.1-Microcoleus-sp.-FACHB-672:0.0864445453,WP_261220767.1-Ancylothrix-sp.-D3o:0.2265501386)91.2/100:0.0790502494)90.1/100:0.0587416236,WP_007355666.1-Kamptonema:0.1939115941)72.2/16:0.0147230509)89.8/56:0.0386042336,(WP_190531554.1-Coleofasciculus-sp.-FACHB-542:0.2012109774,WP_069965800.1-unclassified-Desertifilum:0.3424499295)85.8/82:0.0410428718)0/33:0.0000025303)97.2/96:0.0694287059,(WP_015193936.1-Stanieria-cyanosphaera:0.0308238366,WP_096384422.1-Stanieria-sp.-NIES-3757:0.1102420424)100/100:0.2328785826)64.1/61:0.0339465863)79.5/73:0.0321154776,WP_013324143.1-Gloeothece-verrucosa:0.1251509568)83.8/77:0.0690939419,WP_124977990.1-Aphanothece-sacrum:0.2906788156)69.6/66:0.0094054028,(((WP_017661508.1-Baaleninema-simplex:0.4014737578,WP_264322079.1-Zarconia-navalis:0.1754174812)93.7/91:0.0972945303,(((((WP_106232846.1-Pleurocapsa-sp.-CCALA-161:0.2108976999,WP_229639079.1-Waterburya-agarophytonicola:0.0331867852)83.1/100:0.0395881832,WP_096721927.1-Chondrocystis-sp.-NIES-4102:0.1153587077)98.7/100:0.1819689359,WP_006510035.1-Xenococcus-sp.-PCC-7305:0.2205737850)91.7/98:0.0925764573,WP_036478304.1-Myxosarcina-sp.-GI1:0.2212367134)16/84:0.0432507112,WP_012595695.1-Rippkaea-orientalis:0.2667407123)58.6/23:0.0409497100)75.1/20:0.0154057073,((WP_007308862.1-Crocosphaera-watsonii:0.1069694054,WP_107668138.1-Cyanothece-sp.-BG0011:0.0413395887)9/94:0.0087919212,(WP_008278255.1-Crocosphaera-chwakensis:0.0748195107,WP_009545547.1-Crocosphaera-subtropica:0.0258779146)0/26:0.0102344862)99.9/100:0.2041763902)76.4/70:0.0226838563)85.2/53:0.0393677947,WP_106909014.1-Microcystis-sp.-MC19:0.3348908291)91.5/68:0.0767772396,(((((WP_204137747.1-Halomicronema-sp.-CCY15110:0.0591366045,WP_039724568.1-Leptolyngbya-sp.-LK:0.1006282200)99.8/100:0.2101559681,WP_023076112.1-Leptolyngbya-sp.-Heron-Island-J:0.3025889762)82.4/100:0.1109109307,WP_017324474.1-Synechococcus-sp.-PCC-7336:0.4051906558)86.9/100:0.0659736678,WP_190772446.1-unclassified-Leptolyngbya:0.2161433086)48.7/99:0.0372551272,WP_066347484.1-Geminocystis-sp.-NIES-3708:0.6734991443)81.4/25:0.0282187017)97.2/60:0.1144735238)36.9/81:0.0241219764,(((((((((WP_212718705.1-unclassified-Synechocystis:0.0317548771,WP_228015966.1-Synechocystis-salina:0.1379096859)24.9/100:0.0362261475,WP_010873214.1-unclassified-Synechocystis:0.0611532158)100/100:0.5047579323,(WP_099797666.1-Thermostichus-lividus:0.5298362793,WP_196167318.1-Pseudomonas-monteilii:0.3934599883)60/98:0.1324715082)19.7/44:0.0673059611,WP_015170385.1-Geitlerinema-sp.-PCC-7407:0.3923641371)90.9/91:0.0774729269,(WP_014706296.1-Methylophaga-nitratireducenticrescens:0.0000022203,WP_136552334.1-Methylophaga-sp.-SB9B:0.0917747922)100/100:0.5447084946)48.2/57:0.0264728147,WP_258826205.1-Massilia-agri:0.4868828807)83.9/81:0.0442802494,(WP_153584563.1-Caenimonas-koreensis:0.3580042502,(WP_007185580.1-Hydrocarboniphaga-effusa:0.0897415749,WP_007185565.1-Hydrocarboniphaga-effusa:0.1024817740)98.9/100:0.2624996017)98.2/100:0.2498925179)20.5/25:0.0049721499,((((((((((((((WP_238511579.1-Cupriavidus-gilardii:0.3490224656,((WP_116346290.1-Cupriavidus-taiwanensis:0.1826488188,WP_124685047.1-Cupriavidus-pauculus:0.2837942303)99.5/100:0.5029255500,WP_150984056.1-Cupriavidus-basilensis:0.4662167400)80.2/26:0.1309043312)78.6/19:0.0612825044,WP_224042884.1-Cupriavidus-respiraculi:0.3908100226)68.5/19:0.0483862552,(WP_116600700.1-Cupriavidus-alkaliphilus:0.2377999804,(WP_018008101.1-Cupriavidus-neocaledonicus:0.0506309582,WP_183298446.1-Cupriavidus-alkaliphilus:0.0778676295)97.4/100:0.1601987974)17.3/18:0.0383480495)94.7/78:0.0970100516,(((((((((WP_207000528.1-Trinickia-sp.-DHG64:0.1552156141,WP_124149903.1-Paraburkholderia-dinghuensis:0.1390061277)78.9/100:0.0601152946,(WP_266250364.1-Paraburkholderia-megapolitana:0.2107614653,WP_026121090.1-Paraburkholderia-kururiensis:0.1432101119)71.9/100:0.0718255339)94.1/100:0.0945059343,(((((((((((((WP_059459219.1-Burkholderia-vietnamiensis:0.2931489821,WP_121276956.1-Trinickia-fusca:0.1561509172)91.4/99:0.0690845363,WP_136892825.1-Trinickia-sp.-7GSK02:0.0435754099)82.4/99:0.0327156872,WP_201650988.1-Paraburkholderia-sabiae:0.0464689300)31.1/63:0.0169293556,WP_205970438.1-Paraburkholderia-sp.-Tr-20389:0.0478818798)84.4/68:0.0223949883,WP_054928771.1-Paraburkholderia-caribensis:0.0424860220)85.2/88:0.0256171589,(WP_059889870.1-Burkholderia-stagnalis:0.2699297632,WP_206951398.1-Trinickia-sp.-4D114:0.1957430573)23.1/85:0.0161964795)74.5/71:0.0150292454,WP_209935850.1-Paraburkholderia-sp.-LEh10:0.1263687699)72.7/87:0.0117711813,(WP_060092900.1-Burkholderia-vietnamiensis:0.0245896844,WP_226113708.1-Burkholderia-sp.-AU33545:0.0606432777)100/100:0.2090324746)75.2/35:0.0090125820,((WP_168792792.1-Paraburkholderia-aromaticivorans:0.0971945559,((WP_004544228.1-Burkholderia:0.0402274178,WP_069272519.1-Burkholderia-cepacia:0.2875695470)83.1/99:0.0284954058,WP_059583574.1-pseudomallei-group:0.0378084785)77.5/99:0.0277001768)86.2/96:0.0366966324,(((WP_175791145.1-Burkholderia-ambifaria:0.1370177842,WP_105772332.1-Burkholderia-multivorans:0.0394444793)45.4/51:0.0061232846,(WP_061168975.1-Caballeronia-hypogeia:0.1366058296,WP_059842355.1-Burkholderia-multivorans:0.0502936659)65.5/51:0.0096714496)90.6/91:0.0254542722,(WP_006496666.1-Burkholderia-cenocepacia:0.0373634765,WP_059716874.1-Burkholderia-ubonensis:0.0446010844)93.9/100:0.0350314050)73.9/78:0.0100102834)38.7/43:0.0212066185)81.3/53:0.0155195358,WP_060156927.1-Burkholderia-ubonensis:0.1332807634)91.3/97:0.0431693959,((WP_201138986.1-Paraburkholderia-domus:0.0567056951,WP_266262001.1-Paraburkholderia-madseniana:0.0317140928)99.5/100:0.1296015719,WP_009889585.1-Burkholderia-thailandensis:0.1219259785)83.7/100:0.0314823967)27.5/81:0.0267471681,WP_205976236.1-Paraburkholderia-sp.-Se-20369:0.3262217223)84.6/91:0.0377842891,WP_134195781.1-Paraburkholderia-rhizosphaerae:0.3480517033)87.8/98:0.0432059633)91.8/92:0.0683594560,WP_179404315.1-Burkholderia-guangdongensis:0.2320851042)72/42:0.0210714880,WP_038711716.1-Burkholderia-sp.-lig30:0.3024038173)69.7/44:0.0953556679,((WP_063494936.1-Paraburkholderia-phytofirmans:0.0887118903,WP_254318368.1-Paraburkholderia-sp.-CNPSo-3272:0.0075856593)76.8/45:0.0468529971,WP_042262232.1-Paraburkholderia-heleia:0.0297171907)99.3/100:0.1845954953)54.2/98:0.0518929335,WP_056136218.1-Massilia-sp.-Root133:0.2920768790)86.4/99:0.0801523369,WP_135702508.1-Cupriavidus-oxalaticus:0.2040319299)48.7/48:0.0093642453,((WP_115680924.1-Cupriavidus-taiwanensis:0.2815874271,WP_071014919.1-Cupriavidus-sp.-USMAHM13:0.3422474319)45.7/98:0.0594492569,WP_018313502.1-Cupriavidus-sp.-UYPR2.512:0.3554597578)68.1/93:0.0163483007)92.9/98:0.0502728075)74.1/77:0.0213561739,WP_228215938.1-Aromatoleum-toluclasticum:0.2917360051)83.3/74:0.0538441035,WP_199264031.1-Desulfobulbus-oligotrophicus:0.4369042899)80.3/71:0.0254355285,((((WP_183165579.1-Azomonas-macrocytogenes:0.2354919828,((WP_102836461.1-Stutzerimonas-stutzeri:0.0345187884,(WP_256071125.1-Stutzerimonas-stutzeri:0.0424760320,WP_045423249.1-Pseudomonadaceae:0.0344190004)82.9/100:0.0160940919)95.8/100:0.0642241197,(WP_255961233.1-Stutzerimonas-stutzeri:0.0347803969,WP_025243649.1-Stutzerimonas-stutzeri:0.0575656313)0/45:0.0018742248)94/97:0.0538775117)49.3/20:0.0099581568,(WP_011911332.1-Stutzerimonas-stutzeri:0.0596470873,(WP_230358311.1-Pseudomonas-sp.-NP21570:0.0295557642,((WP_193680295.1-Pseudomonas-lopnurensis:0.0439121884,WP_200589398.1-Stutzerimonas-stutzeri:0.0399329220)86.5/100:0.0284269228,((WP_237255234.1-Pseudomonas-oligotrophica:0.0924911143,WP_043295157.1-Stutzerimonas-degradans:0.0182417824)95.7/99:0.0527870710,WP_120997453.1-_Pseudomonas:0.0404632010)84.5/42:0.0232252577)76.3/42:0.0158581009)86.9/87:0.0320823536)86.4/99:0.0287344285)77.2/18:0.0096442239,(WP_122077451.1-Pseudomonadaceae:0.0434186895,(WP_037024448.1-Pseudomonas-saudiphocaensis:0.0849948726,WP_181084609.1-Stutzerimonas-stutzeri:0.0080925730)96.4/100:0.0523015968)0/27:0.0000021782)96.8/97:0.1098677765,WP_028238613.1-Stutzerimonas-azotifigens:0.0720416171)99.6/100:0.1456100579)83.5/70:0.0312559070,((WP_213445339.1-Variovorax-sp.-PCZ-1:0.4949324143,WP_226921657.1-Kinneretia-sp.-XES5:0.2166489294)91.3/96:0.1849781354,(WP_250024752.1-Cupriavidus-campinensis:0.1857133912,WP_094797980.1-Bordetella-genomosp.-5:0.2937367482)62.7/98:0.1222621043)71.4/47:0.0535158524)91.9/52:0.0641858062,((((((((((((((((((WP_209862330.1-Pseudomonas-sp.-BP8:0.6174081609,WP_225786622.1-Pseudomonas:0.0311875219)46.3/78:0.0832360682,(WP_201176141.1-Pseudomonas-sp.-S31:0.0344502750,WP_261240966.1-unclassified-Pseudomonas:0.0599443274)98.9/100:0.1307563976)82.2/55:0.0339357296,(((WP_054891693.1-Pseudomonas:0.0986537466,WP_236238170.1-Pseudomonas-faucium:0.1721236822)80.4/92:0.0357134226,WP_158461286.1-Pseudomonas-fluorescens:0.2860497481)6.3/67:0.0350330527,(WP_200141208.1-unclassified-Pseudomonas:0.2653322470,WP_029612831.1-Pseudomonas-parafulva:0.0080167024)96.9/99:0.1102562462)91.3/91:0.0685663420)17.1/24:0.0201148150,((WP_217887155.1-Pseudomonas-xanthosomae:0.0383290458,WP_217840807.1-Pseudomonas-fakonensis:0.0489304937)68.9/100:0.0142062605,WP_186654015.1-Pseudomonas-xantholysinigenes:0.2905187566)91.3/99:0.0646477770)80.3/93:0.0370892801,WP_110605176.1-Pseudomonas-soli:0.3071475019)96.6/100:0.0924702729,(((WP_181290220.1-Pseudomonas-brassicacearum:0.2145199759,(WP_249678359.1-Pseudomonas-sp.-PIA16:0.1966801311,WP_043192328.1-Pseudomonas-rhizosphaerae:0.2808163445)0/24:0.0254045031)85.9/54:0.0303261811,WP_268799018.1-Pseudomonas-sp.-BSw22131:0.2080195029)76.7/64:0.0380994705,((WP_201004194.1-Pseudomonas-cichorii:0.0870425530,WP_176569640.1-Pseudomonas-eucalypticola:0.1750831412)90.8/96:0.0688497721,(WP_219063768.1-Pseudomonas-sp.-UMAB-08:0.2779038961,((WP_222540780.1-Pseudomonas-sp.-DR208:0.1432392387,(WP_253509974.1-Pseudomonas-rhodesiae:0.1389471314,WP_093202821.1-Pseudomonas-yamanorum:0.0331196282)90.7/100:0.0521534680)74.7/76:0.0169756378,((WP_065875813.1-Pseudomonas:0.0551367215,(WP_033898696.1-Pseudomonas-fluorescens-group:0.0424060891,(WP_186687394.1-Pseudomonas-sp.-SWRI18:0.0668979554,WP_169378974.1-Pseudomonas:0.0220214548)79.2/100:0.0312637723)71.2/100:0.0148130782)99.5/100:0.0907946576,WP_105697656.1-Pseudomonas-poae:0.0516641533)73/76:0.0114302445)15/99:0.0397209566)86.5/100:0.0495839103)86.7/96:0.0520727371)93/99:0.0882135422)28.7/80:0.0392182497,WP_108096457.1-Pseudomonas-sp.-GV071:0.1823433644)84.5/96:0.0453192797,WP_252273535.1-Pseudomonas-sp.-YIM-B01952:0.1440826591)85.9/96:0.0280143455,WP_143488038.1-Pseudomonas-mangiferae:0.1860003041)47.7/83:0.0212430171,((((WP_090448460.1-Pseudomonas-benzenivorans:0.2568111581,(WP_025164100.1-Pseudomonas-taeanensis:0.0726818109,WP_147170838.1-Pseudomonas-sp.-SJZ079:0.0058314376)96.1/100:0.0712205945)49/53:0.0071728914,((WP_208707982.1-Pseudomonas-sp.-LPB0260:0.0357634281,WP_255840290.1-Pseudomonas-benzenivorans:0.0475619334)95.6/100:0.0649868560,((WP_131198269.1-Pseudomonas-dryadis:0.1596934362,(WP_131178052.1-Pseudomonas-daroniae:0.0779921555,(WP_084307530.1-Pseudomonas-flavescens:0.0879793754,(WP_249909557.1-Pseudomonas-fulva:0.0571067627,(WP_074889242.1-Pseudomonas-argentinensis:0.0332685796,WP_027910132.1-Pseudomonas-sp.-URMO17WK12_I4:0.0572522699)40.6/98:0.0171226348)83/98:0.0422398782)71.6/97:0.0329783587)85.8/100:0.0461076226)94.7/99:0.0927006083,(WP_233685494.1-Pseudomonas-anguilliseptica:0.0738809129,WP_257778987.1-Pseudomonas-sp.-32.2.56:0.0000026147)99.5/100:0.1342104028)53.1/87:0.0275675258)53.2/61:0.0309729791)72.3/77:0.0325661423,(WP_238040454.1-Pseudomonas-sp.-MMS21-TM103:0.1537302173,WP_069518708.1-Pseudomonas:0.1525102002)74.1/96:0.0339523873)86.7/92:0.0409488636,((((((WP_119694090.1-Pseudomonas:0.0645196390,WP_106738315.1-Pseudomonas-mendocina:0.0599325060)92.2/100:0.0538565454,WP_150606948.1-Pseudomonas-oleovorans:0.0578764964)94.2/97:0.0487721068,((WP_024309956.1-Pseudomonas-sp.-P818:0.0645773850,WP_090337209.1-Pseudomonas-chengduensis:0.0277067724)66.1/52:0.0056550478,WP_074939507.1-Pseudomonas-composti:0.0388576555)77/49:0.0074762135)75.3/94:0.0108849701,WP_143497094.1-Pseudomonas-sp.-ALS1131:0.0349494994)85.3/100:0.0231183552,WP_021444205.1-Pseudomonas-sp.-EGD-AK9:0.0633746890)82.7/100:0.0305457737,WP_159970704.1-Pseudomonas-sp.-8Z:0.0897300539)95.6/100:0.0722247206)93.5/93:0.0575674751)90.9/99:0.0549381470,(WP_058067682.1-Pseudomonas-sp.-TTU2014-080ASC:0.1824349884,(WP_192399936.1-Pseudomonas-sp.-PDM15:0.1698376315,WP_226282979.1-Pseudomonas-lalkuanensis:0.1154430161)78.8/100:0.0386867003)95.6/98:0.0651364946)88.2/63:0.0490938643,WP_028626762.1-Pseudomonas-resinovorans:0.0505279703)73/61:0.0082832596,(WP_120654068.1-Pseudomonas-sp.-DY-1:0.1068945988,WP_151138079.1-Pseudomonas-lalkuanensis:0.0516977197)87/97:0.0240658753)76.6/68:0.0130331862,(WP_016495240.1-Pseudomonas-resinovorans:0.0635399771,WP_044873077.1-Pseudomonas-sp.-LFM046:0.0837094094)39.8/95:0.0420110730)41.7/64:0.0168554850,(WP_077525673.1-Pseudomonas-resinovorans:0.1341201675,WP_137821430.1-Pseudomonas-sp.-D-2018:0.1135111496)91.7/98:0.0802111613)90.9/92:0.0467089695,(WP_003453405.1-Pseudomonas:0.1213817280,WP_025281494.1-Ectothiorhodospira-haloalkaliphila:0.5832009991)66.6/66:0.0210680112)84.5/99:0.0734813476,(WP_220803742.1-Pseudomonas-sp.-NCCP-436:0.2045196457,WP_090225506.1-Pseudomonas-kuykendallii:0.1254648407)80.8/98:0.0583797213)88.8/89:0.0731359247,WP_226440716.1-Quatrionicoccus-australiensis:0.5002946243)66.2/20:0.0267253088)20.6/23:0.0138831850,((WP_084055900.1-Desulfacinum-hydrothermale:0.3839755754,WP_123289658.1-Desulfosoma-caldarium:0.3871052027)99.7/99:0.3503844141,WP_136680209.1-Neptunomonas-sp.-XY-337:0.3055466627)0/18:0.0000026435)0/20:0.0000023718,((((WP_200222041.1-Achromobacter-xylosoxidans:0.1875371739,WP_259251071.1-Achromobacter-sp.-JUb104:0.2197132795)95.9/100:0.1117324847,(WP_088588627.1-Achromobacter-marplatensis:0.2080677105,WP_169273406.1-Achromobacter-sp.-Bel:0.1192974818)90.6/100:0.1046479751)52.9/87:0.0331191796,((((WP_111826915.1-Achromobacter-sp.-HZ01:0.1483764752,WP_102775120.1-Achromobacter:0.0254115252)99.9/100:0.1858123116,(WP_129525645.1-Achromobacter-agilis:0.0380691722,(WP_129240460.1-Achromobacter-veterisilvae:0.0559401643,WP_043544564.1-Achromobacter-sp.-RTa:0.0365378996)84.3/100:0.0247633093)64/100:0.0224700012)90.8/100:0.0827095682,(((WP_175208175.1-Achromobacter-anxifer:0.0726831936,(((WP_251864322.1-Achromobacter-sp.-Marseille-Q4962:0.0922874807,WP_164793551.1-Achromobacter-insolitus:0.0409692264)97.5/95:0.0706771357,WP_207472676.1-Achromobacter-sp.-SD115:0.0123989104)92.9/97:0.0451779585,WP_125471167.1-Achromobacter:0.0228182706)96.2/97:0.0527823106)80.3/85:0.0163794460,WP_175164188.1-Achromobacter-aegrifaciens:0.0544853298)85.7/86:0.0408147657,WP_088158656.1-Achromobacter-xylosoxidans:0.0821983021)99.5/100:0.1353415550)68.6/87:0.0126504730,WP_174717299.1-Achromobacter-denitrificans:0.1074479253)85.7/93:0.0377718312)99.5/94:0.2052917118,((WP_006216420.1-Achromobacter-piechaudii:0.1587118057,WP_219991400.1-Achromobacter-sp.-ES-001:0.0981915048)14.7/85:0.0364119614,((WP_175192416.1-Achromobacter-deleyi:0.0964802542,WP_100503206.1-Achromobacter-ruhlandii:0.1771838633)69.2/62:0.0458374741,WP_183017169.1-Achromobacter-sp.-UMC46:0.0729176603)48.1/62:0.0281206925)100/100:0.3185030020)89.1/57:0.0747225500)96.7/40:0.0663986927,((((((((((((WP_132974196.1-Thiogranum-longum:0.2917542746,WP_201344172.1-Thiohalobacter-sp.-COW1:0.2046313815)93.1/100:0.2230556928,WP_198243301.1-methane-oxidizing-endosymbiont-of-Gigantopelta-aegis:0.5656895179)86.2/60:0.2973602097,(((((((GJNO01008138.1-Laxus-oneistus-TSA:1.4540292929,(((((((((((VDM95900.1-Thelazia-callipaeda:0.2498752579,(VDO27950.1-Onchocerca-flexuosa:0.1569520813,(VDO31788.1-Brugia-timori:0.1285408212,(XP_003135693.1-Loa-loa:0.0828029886,((VBB33646.1-Acanthocheilonema-viteae:0.1071126003,VDK73328.1-Litomosoides-sigmodontis:0.0591968808)71.2/67:0.0152488854,CAG9536712.1-Cercopithifilaria-johnstoni:0.0331502243)85.5/67:0.0264708004)50/58:0.0257965019)76.6/58:0.0313344384)98/99:0.1759660773)100/100:0.4235740821,(VDD97390.1-Enterobius-vermicularis:0.4689197736,KHN87800.1-Toxocara-canis:0.2398061690)69/59:0.0746834073)80.9/57:0.0686316886,VDN56932.1-Dracunculus-medinensis:0.4064649573)95/100:0.1694593266,(KAI1709308.1-Ditylenchus-destructor:0.1563289246,KAI1730870.1-Ditylenchus-destructor:0.0294566820)99.6/100:0.2040866425)85.6/70:0.0765122244,GGJS01006545.1-Plectus-murrayi-TSA:0.5107788546)70.4/33:0.0477441698,TKR72632.1-Steinernema-carpocapsae:0.2635627303)88.1/67:0.0873565884,KAE9552274.1-Halicephalobus-sp.-NKZ332:0.3608773111)86.6/95:0.1267413900,((CAD5228290.1-Bursaphelenchus-okinawaensis:0.0539544187,CAD5235627.1-Bursaphelenchus-xylophilus:0.0417642229)0/68:0.0657215278,((KAI6206096.1-Aphelenchoides-besseyi:0.1828650284,KAI6191010.1-Aphelenchoides-bicaudatus:0.1041485285)77.9/100:0.0660584578,KAI6234710.1-Aphelenchoides-fujianensis:0.1226122808)94.1/100:0.1245568813)97.9/100:0.1388693086)55.3/73:0.0622167003,KAH7695359.1-Aphelenchus-avenae:0.3611220404)98.9/99:0.1945315920,((CAI4229259.1-Auanema-sp.-JU1783:0.5323442810,(CAD6189320.1-Caenorhabditis-auriculariae:0.4496761251,(CAB3400093.1-Caenorhabditis-bovis:0.2645863255,((NP_509046.1-Caenorhabditis-elegans:0.0369438740,XP_003117635.1-Caenorhabditis-remanei:0.0359004845)50.1/100:0.0510657922,CAI5454992.1-Caenorhabditis-angaria:0.0369774710)82/100:0.0523511354)92/100:0.1218726362)98/100:0.2048423037)70.4/94:0.0363687788,PAV65180.1-Diploscapter-pachys:0.3058724461)87.2/99:0.0478642518)73.5/99:0.0308933923,((KHJ99538.1-Oesophagostomum-dentatum:0.0668555847,EPB80542.1-Ancylostoma-ceylanicum:0.0528141132)96.2/100:0.1049436606,((VDO65958.1-Haemonchus-placei:0.1613583769,VDL70484.1-Nippostrongylus-brasiliensis:0.1010932389)85.3/100:0.0296811299,VDO95824.1-Heligmosomoides-polygyrus:0.0909655222)98.8/100:0.1540881793)93.1/100:0.1450768422)95.6/99:0.4017044812)99.5/100:0.9814769068,(WP_042281872.1-Candidatus-Protochlamydia-sp.-R18:0.1567083470,WP_068471358.1-Parachlamydia-sp.-C2:0.1961101403)99.7/100:0.8844556813)86.9/97:0.3266337302,(WP_008983868.1-Alishewanella-agri:1.2689331895,WP_265845290.1-Plesiomonas-shigelloides:1.1310894280)74.4/97:0.1645337853)86.1/100:0.1608631622,(WP_161826415.1-Petrachloros-mirabilis:0.5958253276,WP_099324654.1-Candidatus-Kuenenia-stuttgartiensis:0.7992502642)83.2/97:0.1261327217)79.5/92:0.0771204485,(WP_110520589.1-Herpetosiphon-llansteffanensis:1.1779252081,(WP_239493201.1-Luteitalea-sp.-TBR-22:0.4902861315,WP_058867107.1-Chloracidobacterium-thermophilum:0.5479064296)49.4/92:0.1187784261)69.2/89:0.2028764741)72.4/73:0.0774637714,((WP_255135177.1-Gracilimonas-sp.-CAU-1638:0.4277864099,WP_020401568.1-Gracilimonas-tropica:0.3552985447)99.9/100:0.9396134373,(((((WP_094934028.1-Enterobacter-cloacae-complex:0.7597557959,WP_016795256.1-Vibrio-cyclitrophicus:0.2272654272)51.3/42:0.0834609027,WP_039473409.1-Vibrio-sinaloensis:0.4178823404)89.1/39:0.0919849953,((((WP_195504469.1-Citrobacter-farmeri:0.3736265744,WP_064554809.1-Buttiauxella-noackiae:0.3542957919)89.2/100:0.2161418594,(WP_001100815.1-Escherichia-coli:0.4343402176,WP_240304790.1-Vibrio-campbellii:0.5627850427)91.4/98:0.2033723568)90/26:0.1109633519,((WP_269408820.1-Pseudoalteromonas-shioyasakiensis:0.2500374338,WP_207379479.1-Shewanella-sedimentimangrovi:0.2400621214)95.1/100:0.0806551099,((WP_061013169.1-Vibrionaceae:0.2993701272,WP_155662549.1-Aliivibrio-fischeri:0.1519030651)83/96:0.1000177049,WP_171328835.1-Vibrio-splendidus:0.3368484483)0/20:0.0000023984)88.6/96:0.0655321111)81.9/22:0.0442934493,(((WP_010368546.1-Pseudoalteromonas-piscicida:0.5137317312,WP_086590801.1-Vibrio-parahaemolyticus:0.4502978775)77.2/97:0.1224184712,WP_004741399.1-Vibrio:0.4431233767)53.5/19:0.0535712582,WP_016537220.1-Cedecea-davisae:0.1895040084)85.9/23:0.0530165719)91/40:0.0852868912)64.8/41:0.0321999141,WP_057065012.1-Citrobacter:1.0293001689)92/99:0.1704870908,WP_216735790.1-Hahella-sp.-HN01:0.3271664933)76/100:0.1522663075)82.8/75:0.1565400684)29.3/65:0.0653877575,WP_062153084.1-Beggiatoa-leptomitoformis:0.6443408144)42.1/40:0.0809021705)77.9/67:0.0675792962,WP_230840800.1-Gloeobacter-morelensis:0.6934752851)88.1/91:0.1178935509,(WP_110265962.1-Sinimarinibacterium-flocculans:0.3085727512,WP_077329105.1-Virgibacillus-siamensis:0.7659342846)95.7/86:0.1747920940)83.9/65:0.0936523402,(WP_234070476.1-Myxococcus-stipitatus:0.3767591862,WP_120555541.1-Corallococcus-aberystwythensis:0.3149893968)99.8/100:0.5244455178)79.9/37:0.0784299412,WP_152145882.1-Alkalilimnicola-sp.-S0819:0.6399701223)20.1/20:0.0179318645,((WP_220807781.1-Noviherbaspirillum-aridicola:0.5118392235,(WP_227390214.1-Halomonas-malpeensis:0.7805007625,WP_119020982.1-Halomonas-sp.-JS92-SW72:0.3171478088)65.6/98:0.2356757156)87.2/88:0.1391091501,((((((WP_018154793.1-Methanothermococcus-thermolithotrophicus:0.2006613483,WP_048201153.1-Methanocaldococcus-bathoardescens:0.0677439930)94.6/100:0.1171219943,((((((WP_010868297.1-Pyrococcus-abyssi:0.0434228538,WP_048103006.1-Aciduliprofundum-boonei:0.3944990505)23.8/17:0.0656794137,((WP_167913028.1-Thermococcus-sp.-21S7:0.1007650207,WP_048150691.1-Palaeococcus-ferrophilus:0.0310950093)96.8/100:0.0697014901,((((WP_088855609.1-Thermococcus-siculi:0.0694816124,WP_087036111.1-Thermococcus-litoralis:0.0456797452)82/100:0.0290110752,(WP_167888701.1-Thermococcus-sp.-MV5:0.0467617720,WP_209475361.1-Thermococcus-stetteri:0.1199657677)53.8/100:0.0528449105)82/96:0.0269575408,(WP_013468354.1-Thermococcus-barophilus:0.0647298831,WP_013906489.1-Pyrococcus-yayanosii:0.0296407644)85.3/100:0.0166506812)88.5/94:0.0184132360,WP_048053121.1-Pyrococcus-horikoshii:0.0380180476)60.5/87:0.0066427422)46.4/91:0.0258121117)82.1/17:0.0301817525,(((((((((((((WP_237702072.1-Thermococcus-sp.-AM4:0.0450144697,WP_223211916.1-Thermococcus-henrietii:0.0708461391)97.3/100:0.0723714785,WP_014733609.1-Pyrococcus-sp.-ST04:0.1626826621)77.1/100:0.0230195480,WP_058938040.1-Thermococcus-celericrescens:0.0362120064)59.4/53:0.0184625553,(WP_139679991.1-Thermococcus-indicus:0.0457030619,WP_148882943.1-Thermococcus-aciditolerans:0.0444867951)76.8/99:0.0335618791)75.8/91:0.0161134732,WP_088865724.1-Thermococcus-barossii:0.0701403500)82.5/40:0.0160600173,WP_232460978.1-Thermococcus-radiotolerans:0.0681858945)0/34:0.0015214221,WP_167711671.1-unclassified-Thermococcus:0.0306227658)60.5/93:0.0224901967,WP_258085050.1-Thermococcus-sp.-813A4:0.0406703783)80.2/95:0.0243632932,WP_088181368.1-Thermococcus-sp.-5-4:0.0677148600)92.6/94:0.0403599822,(WP_152879125.1-Thermococcus-sp.-101-C5:0.0682980442,WP_015850310.1-Thermococcus-sibiricus:0.0620921264)74.9/95:0.0092853879)81.4/92:0.0175144797,WP_253304977.1-Thermococcus-aggregans:0.0509058186)51.2/92:0.0249285143,WP_068664737.1-Thermococcus-piezophilus:0.0733826521)41.5/98:0.0251649423,WP_088862340.1-Thermococcus-celer:0.1163555658)79/51:0.0197502079)53.7/17:0.0223818997,((WP_013798345.1-Methanotorris-igneus:0.0508189037,WP_013866394.1-Methanothermococcus-okinawensis:0.2599715064)41.5/96:0.0285776511,WP_007044577.1-Methanotorris-formicicus:0.0858321488)94.7/96:0.0827793788)88.2/13:0.0375607383,WP_148120035.1-Methanofervidicoccus-sp.-A16:0.2037393252)100/100:0.4167611712,(((WP_245533116.1-Coprothermobacter-platensis:0.7656383086,WP_022856039.1-Thermodesulfobacterium:0.2406579169)63.8/100:0.1130130702,WP_150071182.1-Thermodesulfobacterium-sp.-TA1:0.2921563984)91.7/100:0.1629668798,((((WP_011996389.1-Coxiella-burnetii:1.0441251232,WP_013638952.1-Desulfurobacterium-thermolithotrophum:0.0025675509)84.9/99:0.0566163890,WP_132526207.1-Phorcysia-thermohydrogeniphila:0.1186412171)85.7/99:0.0360430299,WP_089322203.1-Desulfurobacterium-atlanticum:0.2087402399)90.6/99:0.0839286272,((WP_029522925.1-Persephonella-sp.-KM09-Lau-8:0.1761339685,(WP_200674641.1-Persephonella-atlantica:0.0914101197,WP_096999769.1-Persephonella-hydrogeniphila:0.1561064422)75/93:0.0550828806)91.1/93:0.0657646810,WP_028950938.1-Sulfurihydrogenibium-subterraneum:0.2066608996)68.6/93:0.0327582089)96.3/100:0.1394478526)45.7/99:0.0561349709)98.5/100:0.2367511133)61.5/70:0.0193944015,((WP_011869065.1-Methanococcus-maripaludis:0.1130033295,WP_013998786.1-Methanococcus-maripaludis:0.0750243249)59/100:0.0452134512,(WP_181504225.1-Methanococcus-maripaludis:0.0793150001,WP_181493664.1-Methanococcus-maripaludis:0.0957715676)9.4/95:0.0113975214)100/100:0.3580947329)90.5/70:0.1077102388,(WP_052562942.1-Candidatus-Brocadia-sinica:0.5639962983,WP_173298646.1-Thermanaeromonas-sp.-C210:0.4025898769)10.8/46:0.0449011871)89.2/90:0.2136786797,((WP_119360783.1-Meiothermus-luteus:0.2552523868,WP_027877894.1-Meiothermus-cerbereus:0.0693072858)84.4/100:0.0814294789,WP_208013758.1-Meiothermus-sp.-CFH-77666:0.0948442492)100/100:0.7828936897)98.2/100:0.3949431669,(KAI6244242.1-Aphelenchoides-fujianensis:0.6262096633,KAI6170547.1-Aphelenchoides-bicaudatus:1.0856019335)96.8/100:0.3513986018)35.6/51:0.0989827101)67.8/57:0.1123297930)95/94:0.1367258934,(((WP_018882239.1-unclassified-Thioalkalivibrio:0.5523400398,WP_125198734.1-Guyparkeria-sp.-SCN-R1:0.2375300522)82.9/100:0.0757609126,(((WP_026296529.1-Thioalkalivibrio-sp.-ALJ2:0.1642176221,WP_019569965.1-Thioalkalivibrio-sp.-ALE11:0.1021256482)88/100:0.0314267922,WP_028484075.1-Thioalkalivibrio-sp.-ALE17:0.0524678254)75.2/100:0.0106179007,WP_018169543.1-Thioalkalivibrio-sp.-ALMg9:0.0405366157)98.9/100:0.1502364937)24.4/90:0.0333446620,WP_250072676.1-Guyparkeria-hydrothermalis:0.2629476755)99/100:0.2033136004)82.1/61:0.0410064646,(WP_176062241.1-Anaeromyxobacter-diazotrophicus:0.2686483924,WP_248361410.1-Anaeromyxobacter-oryzae:0.2831527421)99.1/100:0.2561713734)84.4/21:0.0703197857,((WP_205202385.1-Azonexus-hydrophilus:0.2719668183,WP_173581083.1-Azonexus-fungiphilus:0.3540961486)25.4/81:0.0505145632,WP_121459656.1-Azonexus-fungiphilus:0.3752898012)86.6/95:0.0974745686)95.3/38:0.1135270215,((WP_002810067.1-Nitrosococcus-oceani:0.0737511867,WP_013221233.1-Nitrosococcus-watsonii:0.0753181390)100/100:0.3716353821,(WP_040097704.1-Geoalkalibacter-ferrihydriticus:0.3462883292,WP_066727622.1-Desulfuromonas-sp.-DDH964:0.3197898910)89.4/100:0.0668069549)55.4/96:0.0231915713)76.7/34:0.0216776428)87.7/35:0.0311908713,((((WP_070114691.1-Acidithiobacillus-caldus:0.4409148390,(((WP_009566950.1-Acidithiobacillus:0.0562941765,WP_215870853.1-unclassified-Acidithiobacillus:0.0960554715)96.6/100:0.0916916851,WP_163095688.1-Acidithiobacillus-ferrianus:0.0488327706)90.5/100:0.0673881035,WP_101536718.1-Acidithiobacillus-sp.-SH:0.1670624533)80.5/100:0.0847568912)100/100:0.3596576874,((WP_160809027.1-Nitrosomonas-sp.-HPC101:0.6013218170,WP_154206467.1-Ralstonia-pickettii:0.1901937957)67/100:0.0820349998,WP_253536546.1-Ralstonia-sp.-21YRMH01-3:0.5042415211)73.1/98:0.0507993144)76.9/94:0.0243759886,((((WP_036252969.1-Methylobacter-sp.-BBA5.1:0.3390854139,(((WP_019918777.1-Methyloversatilis-discipulorum:0.0611346298,WP_018229738.1-Methyloversatilis-universalis:0.0687926011)67.6/100:0.1023004346,WP_183409942.1-Litorivivens-lipolytica:0.6568285600)37.5/99:0.0954467848,(WP_101941536.1-Uliginosibacterium-sp.-TH139:0.1364343746,WP_095523004.1-Candidatus-Dactylopiibacterium-carminicum:0.1197531106)96.5/100:0.1893015483)95.1/100:0.1444302024)1.6/92:0.0291786020,((WP_047193087.1-Schlegelella-brevitalea:0.3831362487,WP_019561187.1-Caldimonas-manganoxidans:0.2900034894)47.2/95:0.1382983998,WP_074865046.1-Atopomonas-hussainii:0.2165211180)97/87:0.1773699120)89/62:0.0573820178,((((WP_109041087.1-Azospira-sp.-I13:0.1488583538,WP_014237023.1-Azospira-oryzae:0.1626420456)97.1/100:0.1732993659,WP_067106371.1-Methylobacillus-sp.-MM3:0.2660707449)40.9/88:0.0505073051,WP_201673630.1-Ramlibacter-monticola:0.2914541847)80.8/93:0.0355407724,WP_212785890.1-Ferrigenium-kumadai:0.2890987127)80.6/95:0.0599229176)73.2/29:0.0152452281,(((((WP_074796726.1-Nitrosospira-briensis:0.0878618882,(WP_090882638.1-Nitrosovibrio-sp.-Nv6:0.0716376875,WP_097065525.1-Nitrosovibrio-sp.-Nv4:0.0417593890)88.5/100:0.0704703670)99.8/100:0.2868205276,WP_122244186.1-Vandammella-animalimorsus:0.3301270055)28/59:0.0583060563,(WP_223986959.1-Cupriavidus-pampae:0.0709651358,((WP_011517306.1-Cupriavidus-metallidurans:0.0602082093,WP_019451253.1-Cupriavidus-sp.-BIS7:0.0638996468)0/90:0.0014735905,WP_082079449.1-Cupriavidus-basilensis:0.1009138308)78.4/93:0.0349201616)100/100:0.2431526563)67.8/58:0.0229758782,((WP_057673948.1-Curvibacter-sp.-PAE-UM:0.1950731719,WP_169420802.1-Ramlibacter-agri:0.2858029542)74.8/92:0.0565222913,WP_056898839.1-Pseudorhodoferax-sp.-Leaf274:0.3731039106)66.2/73:0.0121689544)25.6/35:0.0107589562,((((WP_106935816.1-unclassified-Variovorax:0.4040250797,WP_137918285.1-Hydrogenophaga-sp.-2FB:0.2175051358)75.2/39:0.0340186711,((WP_219210863.1-Variovorax-boronicumulans:0.2371790779,(((((WP_126747644.1-Variovorax-sp.-DXTD-1:0.1982710097,WP_206177103.1-Variovorax-sp.-RKNM96:0.1384006313)19.8/86:0.0202884748,WP_068630051.1-Variovorax-sp.-PAMC-28711:0.3515662479)22.9/81:0.0348600415,WP_093207264.1-Variovorax-sp.-YR750:0.1507605020)67.7/93:0.0575148261,(WP_093441791.1-Variovorax-sp.-770b2:0.1759549237,WP_160553990.1-Xylophilus-rhododendri:0.2261768707)36.5/44:0.0726415971)49.6/40:0.0391149207,WP_192194832.1-Variovorax-sp.-38R:0.2745510917)47.6/88:0.0444530821)93.6/85:0.0764669009,WP_137890779.1-Ramlibacter-sp.-2FC:0.1259758910)71.3/37:0.0160182472)0/73:0.0066043063,WP_174484170.1-methanotrophic-endosymbiont-of-Bathymodiolus-puteoserpentis:0.2987841031)37/68:0.0111730355,((WP_026436726.1-Acidovorax-sp.-JHL-9:0.2599241904,WP_086926522.1-Acidovorax-carolinensis:0.2227065914)24.2/99:0.0441123262,WP_166067558.1-Diaphorobacter-sp.-HDW4B:0.3021965386)90.2/99:0.0499589762)69/69:0.0094663581)94.7/38:0.0498381757)90.3/78:0.0383540468)91.1/89:0.0434194795,((((WP_108647113.1-Polynucleobacter-rarus:0.0319304410,WP_084281813.1-Polynucleobacter-kasalickyi:0.0484166215)100/100:0.6763017956,(((WP_215371371.1-Polynucleobacter-sp.-MWH-UH24A:0.2372827302,WP_173942040.1-Polynucleobacter-antarcticus:0.2194864642)96.7/100:0.1456216123,((WP_216254756.1-Polynucleobacter-sp.-AP-Melu-500A-A1:0.0674470353,WP_216218975.1-Polynucleobacter-sp.-AP-Latsch-80-C2:0.0480555270)99.4/100:0.1172567577,(((((WP_215319389.1-Polynucleobacter-paneuropaeus:0.1218470079,(WP_215352354.1-Polynucleobacter-sp.-MWH-P3-07-1:0.0560536810,WP_215380622.1-Polynucleobacter-paludilacus:0.0448312728)23.1/100:0.0419037678)95.7/99:0.0740445157,WP_062307935.1-Polynucleobacter-sinensis:0.0663521501)79.5/60:0.0205032341,WP_216197047.1-Polynucleobacter-sp.-UK-Mo-2m-Kol15:0.0858745632)30.1/57:0.0087045917,((((((WP_215376847.1-Polynucleobacter-asymbioticus:0.0324317084,WP_215328794.1-Polynucleobacter-sp.-AP-Sving-400A-A2:0.0652549997)97.8/100:0.0642672434,WP_173959551.1-Polynucleobacter-arcticus:0.0362377081)94.3/100:0.0294684449,WP_216225867.1-Polynucleobacter-sp.-MG-27-Goln-C1:0.1031738705)42.1/99:0.0076704804,(WP_216168716.1-Polynucleobacter-bastaniensis:0.1009483507,WP_216178894.1-Polynucleobacter-sp.-80A-SIGWE:0.0170736912)86.8/100:0.0198754542)87.5/99:0.0266691042,WP_215299358.1-Polynucleobacter-sp.-AP-Nino-20-G2:0.1133998311)87.9/49:0.0263382515,(WP_216268685.1-Polynucleobacter-sp.-Nonnen-W13:0.0604810957,(WP_215334312.1-Polynucleobacter-sp.-AP-Ainpum-60-G11:0.0554546044,WP_251374783.1-Polynucleobacter-corsicus:0.0408165268)71.8/34:0.0069469794)89.4/71:0.0163357560)49.4/30:0.0073068252)90.5/98:0.0337122142,WP_076024025.1-Polynucleobacter-sphagniphilus:0.1098202494)71.4/51:0.0123592844)95.2/100:0.0661633453)52.2/72:0.0184378535,(((WP_128113711.1-Polynucleobacter-necessarius:0.1247051327,WP_215385554.1-Polynucleobacter-sp.-AP-Kolm-20A-A1:0.0902303854)93.9/100:0.0828173980,WP_216177372.1-Polynucleobacter-sp.-AP-Feld-500C-C5:0.2058211261)72.3/98:0.0401676924,WP_114639791.1-Polynucleobacter-necessarius:0.1477298125)43.9/78:0.0685766160)99.4/98:0.2929703700)94.2/100:0.1865462921,(((WP_226989104.1-Desulfuromonas-sp.-TF:0.5745969481,WP_227459030.1-Cupriavidus-pauculus:0.3494589971)99/100:0.2675480854,WP_227181580.1-Leeia-sp.-TBRC-13508:0.2710639663)86/97:0.0917193885,(WP_108623287.1-Orrella-marina:0.5561897427,WP_243478771.1-Orrella-sp.-f23:0.4371957119)88.6/100:0.1486750281)90.4/91:0.1146953856)25.6/71:0.0476597701,WP_123210144.1-Stagnimonas-aquatica:1.2276541847)55.9/27:0.0667810471)88.5/32:0.0366303640)0/37:0.0000022311,(WP_054772520.1-Methylogaea-oryzae:0.2697435968,(WP_091467897.1-Methylophilus-rhizosphaerae:0.4378991498,((WP_117174481.1-Rhodoferax-lacus:0.3010189256,((WP_094482496.1-Rhodoferax-sp.-TH121:0.1216721763,WP_105262671.1-Rhodoferax-sp.-TS-BS-61-7:0.0418360057)87/100:0.0591783077,WP_214121631.1-Curvibacter-sp.-CHRR-16:0.3151710272)80.8/100:0.1310813907)89.3/94:0.0975835605,(WP_066884254.1-Dechloromonas:0.1131755512,WP_027458229.1-Dechloromonas-agitata:0.1604132132)87.7/100:0.0939726797)88.9/93:0.1047499652)87.9/88:0.0990717452)76.5/54:0.0335185446)86.2/28:0.0503128205)91.3/40:0.0500425865,((((((WP_239190817.1-Candidatus-Nitrotoga-sp.-HW29:0.0716329542,WP_239797721.1-Candidatus-Nitrotoga-arctica:0.0327477132)33/43:0.0090265454,WP_239235002.1-Candidatus-Nitrotoga-sp.-BS:0.0712884717)0/26:0.0000022997,WP_239287644.1-Candidatus-Nitrotoga-sp.-1052:0.0273478040)99.9/100:0.2998206769,WP_009207329.1-Sulfuricella-denitrificans:0.4195633698)90.1/75:0.0547565552,WP_054689510.1-Desulfosarcina-cetonica:0.3049044468)84.7/73:0.0497009073,(((((((((((((WP_065305794.1-Janthinobacterium-psychrotolerans:0.1642765429,WP_034778721.1-Janthinobacterium-lividum:0.0454643975)11.9/91:0.0030488592,WP_035822645.1-Janthinobacterium-sp.-RA13:0.0810509948)78.6/91:0.0102992136,WP_152282105.1-Janthinobacterium-violaceinigrum:0.0250201671)72.7/91:0.0081256251,WP_183742413.1-unclassified-Janthinobacterium:0.0588218015)75.9/90:0.0079068289,(((((WP_139090581.1-Janthinobacterium-lividum:0.0831474516,WP_230539136.1-Janthinobacterium-sp.-FW305-129:0.0505598965)85.6/99:0.0181211021,WP_094442213.1-Janthinobacterium-sp.-PC23-8:0.0791091268)29.6/99:0.0081250069,WP_215403374.1-Janthinobacterium-sp.-JC611:0.0311129723)77.7/100:0.0101255920,WP_102124214.1-Janthinobacterium-sp.-ROICE36:0.0418628193)74/92:0.0081643045,WP_096238059.1-Janthinobacterium-svalbardensis:0.0243109123)76.9/92:0.0082833949)98.8/98:0.1327538932,WP_023495308.1-Methyloglobulus-morosus:0.2287636666)79.3/73:0.0242959798,(WP_101482628.1-unclassified-Janthinobacterium:0.2474139022,((WP_099917421.1-Massilia-psychrophila:0.1125260943,WP_250479419.1-unclassified-Caballeronia:0.5042458603)53.5/76:0.0558019125,(WP_205319084.1-Glaciimonas-sp.-PAMC28666:0.0867883881,WP_092396716.1-Collimonas-sp.-OK412:0.1528390702)78.2/98:0.0219145542)85.3/74:0.0354794953)67.3/46:0.0061491388)96/76:0.1133358043,((WP_162020603.1-Undibacterium-crateris:0.1761269397,WP_186884883.1-Undibacterium:0.0616993064)80.3/79:0.0344530264,WP_186861877.1-Undibacterium-griseum:0.0569314719)100/100:0.1899280104)78/72:0.0339537347,((WP_247737070.1-Thauera-aromatica:0.0620045052,WP_039805139.1-Azotobacter-chroococcum:0.2065555221)99.4/100:0.2009540737,((WP_195874647.1-Herminiimonas-contaminans:0.1688174891,WP_041297045.1-Janthinobacterium-sp.-Marseille:0.1100288516)78.6/100:0.0323573873,(WP_076590920.1-Herminiimonas-arsenitoxidans:0.2077999377,(WP_206604468.1-Herminiimonas-fonticola:0.0508098151,WP_188420499.1-Oxalicibacterium-solurbis:0.2290429830)73.3/99:0.0191973014)87.1/100:0.0519204857)98.3/100:0.1258704454)45.7/62:0.0192696423)46.4/60:0.0510379671,WP_217900122.1-Noviherbaspirillum-humi:0.2395437933)89.3/85:0.0934125758,(WP_237889684.1-Dissulfurimicrobium-hydrothermale:0.4400769483,WP_237247350.1-Sideroxyarcus-emersonii:0.2339736281)78.9/82:0.1041208012)89.8/69:0.0578909861,WP_217345508.1-Noviherbaspirillum-sp.-L7-7A:0.3451775314)0/43:0.0000020366,((WP_006529211.1-Gloeocapsa-sp.-PCC-73106:0.3751822858,(WP_083560927.1-Oceanococcus-atlanticus:0.4713870711,WP_104228621.1-Solimonas-fluminis:0.3034643601)97.6/99:0.2725596396)79.6/88:0.0444882499,WP_047549861.1-Methylotenera-sp.-G11:0.2661858406)79.1/89:0.0316344866)93.2/68:0.0672953204)92.4/93:0.0475235751)87.4/41:0.0371739831)92.6/81:0.0554500154)93.7/91:0.0743140259,((WP_188630571.1-Oceanisphaera-marina:0.1784407698,WP_104485062.1-Oceanisphaera-arctica:0.1352949483)6/67:0.0149254216,WP_204795415.1-Oceanisphaera-litoralis:0.0541430096)100/100:0.2715289049)90/75:0.0608771416,((((WP_213263532.1-Burkholderia-pyrrocinia:0.1855831377,WP_060279477.1-Burkholderia-cepacia-complex:0.1668501651)95.2/100:0.1703443979,(((WP_208101977.1-Burkholderia-sp.-D-99:0.0406376598,WP_205714023.1-Burkholderia-cepacia:0.0355251568)72.4/100:0.0092822688,WP_174919663.1-Burkholderia-metallica:0.0266665073)84.2/100:0.0501138320,WP_077021900.1-Burkholderia-cenocepacia:0.1040127552)100/100:0.3329255487)88/100:0.0656094504,WP_174426450.1-Cupriavidus-basilensis:0.5590165531)85/100:0.0572999875,((WP_094838100.1-Bordetella-genomosp.-4:0.2445249712,WP_160351730.1-Bordetella-sp.-15P40C-2:0.2993618259)97.1/100:0.2864328916,WP_148717329.1-Chitinolyticbacter-meiyuanensis:0.9742622048)87.2/73:0.1055334805)63.1/61:0.0713147076)89.3/54:0.0703279154,((WP_189493202.1-Formosimonas-limnophila:0.2956620337,WP_133620003.1-Hydromonas-duriensis:0.3723057682)98.2/100:0.2502634369,(((WP_132257458.1-Paucimonas-lemoignei:0.0806073064,WP_119785517.1-Noviherbaspirillum-sedimenti:0.0942726864)99.9/100:0.4731136983,(WP_026470910.1-Alkanindiges-illinoisensis:0.2848857503,(WP_248147450.1-Moraxella-osloensis:0.4492849649,WP_092690500.1-Acinetobacter-kyonggiensis:0.2085055552)87.2/100:0.1437853249)99.3/100:0.3144044930)96.4/99:0.1683200531,(((((WP_198111724.1-Inhella-proteolytica:0.5769112138,WP_241396682.1-Pelomonas-sp.-CA6:0.1684210441)90/100:0.1142594726,(WP_028996355.1-Azohydromonas-australica:0.0937023843,WP_066337818.1-Azohydromonas-lata:0.1246028810)94.6/100:0.1308858918)52.3/94:0.0982813173,WP_037482909.1-Sphaerotilus-natans:0.1870730879)95.2/100:0.2175454530,WP_220807643.1-Noviherbaspirillum-aridicola:0.5069197602)68.6/59:0.1081818142,((((((WP_169107474.1-Comamonas-suwonensis:0.0591795433,WP_219343559.1-Comamonas-sp.-Y33R10-2:0.1045997129)94.8/100:0.0665094276,(WP_003073420.1-Comamonas:0.0879929687,((WP_211457163.1-Comamonas-sp.-PE63:0.0610912245,WP_148729134.1-Comamonas-sp.-Z3:0.0545008242)66.6/71:0.0082198326,WP_034391098.1-Comamonas-thiooxydans:0.0194244227)75.2/71:0.0100425796)84.3/85:0.0322136522)76.9/85:0.0186969562,WP_218241177.1-Comamonas-fluminis:0.1430240701)99.1/99:0.1906166037,WP_012203208.1-Delftia-acidovorans:0.2895750021)89.2/100:0.0920165278,WP_066475960.1-Comamonas-terrae:0.2315580424)72.3/65:0.0300621336,WP_058879441.1-Comamonas-kerstersii:0.1310302251)99.9/100:0.3181936692)75.1/76:0.0294147920)89.5/84:0.0814018532)90.1/63:0.0562234441)83.3/40:0.0332756663,(WP_208005626.1-Oceanisphaera-pacifica:0.1006309770,(WP_087034853.1-Oceanisphaera-profunda:0.0809414749,WP_087034873.1-Oceanisphaera-profunda:0.1563621326)89/100:0.0925420880)97.9/100:0.1402937243)90.8/38:0.0607953240,((((((((((((WP_004123988.1-Rhizobium-freirei:0.1246926730,WP_095433322.1-Rhizobium-sp.-11515TR:0.0457200753)76.3/94:0.0294177985,WP_106999541.1-Rhizobium-sp.-JAB6:0.0639489290)0/86:0.0000024982,((WP_174074493.1-Agrobacterium-rhizogenes:0.1181419230,((WP_183765619.1-unclassified-Rhizobium:0.0493237747,WP_120667780.1-unclassified-Rhizobium:0.0670558129)75.1/100:0.0375308610,WP_037195771.1-Rhizobium-lusitanum:0.0327280533)48.4/100:0.0164089070)73.7/85:0.0130781108,((((WP_199146672.1-Rhizobium-sp.-ASV8:0.2089302110,WP_184706966.1-Rhizobium-lusitanum:0.0582672513)85.2/96:0.0529675299,WP_145321829.1-unclassified-Rhizobium:0.1291811853)77/68:0.0158395229,(WP_168880117.1-Rhizobium-sp.-P28RR-XV:0.0516291161,WP_112341687.1-Rhizobium-tropici:0.0572385586)87/100:0.0173464709)78.4/67:0.0166301993,WP_092719658.1-Rhizobium-multihospitium:0.0465135275)74.4/72:0.0130725059)89.2/93:0.0245124664)88.1/94:0.0170996751,WP_183712806.1-unclassified-Rhizobium:0.0377627837)82.1/96:0.0223597960,WP_015341698.1-Rhizobium:0.0286861956)99.9/100:0.1991395996,(WP_043579071.1-Chromobacterium:0.0720918540,WP_071113424.1-Chromobacterium-sphagni:0.1326072361)99.7/100:0.2256814870)92.3/100:0.0829571266,(((WP_189531325.1-Paludibacterium-paludis:0.5735856749,(WP_159673755.1-Andreprevotia-sp.-IGB-42:0.2475420484,WP_084093192.1-Andreprevotia-lacus:0.0852151053)98.2/100:0.2397089386)39.7/100:0.0674920630,(WP_043620211.1-Chromobacterium-piscinae:0.2899051735,(WP_103900886.1-Chromobacterium-alticapitis:0.2778374194,WP_045052286.1-Chromobacterium-violaceum:0.3020290272)62.2/100:0.0875811481)95/100:0.1480419685)58.9/100:0.0322930621,WP_020932001.1-Cycloclasticus-zancles:0.4113875046)78.1/100:0.0367957455)87.8/98:0.0754204067,(((WP_174873379.1-Vogesella-oryzae:0.2415339217,WP_147289921.1-Crenobacter-cavernae:0.1684816890)76.6/98:0.0273647956,(WP_008952446.1-Pseudogulbenkiania-ferrooxidans:0.3048622349,WP_014088374.1-Pseudogulbenkiania-sp.-NH8B:0.2907258179)92/100:0.1262416159)78.8/90:0.0274382356,WP_137008594.1-Aquitalea-aquatilis:0.5238427769)21.7/61:0.0230122684)95.7/92:0.1055631673,((((WP_147247296.1-Brenneria-salicis:0.0467009195,WP_172289906.1-unclassified-Brenneria:0.1260557973)89.5/100:0.0531491787,WP_039356706.1-Pectobacterium-actinidiae:0.0945187911)94.1/99:0.1258336518,((WP_198997094.1-Pectobacterium-carotovorum:0.0221203580,(WP_165707548.1-Pectobacterium:0.0409924547,WP_174866005.1-Pectobacterium-polaris:0.0300965528)99/100:0.0787745013)63.3/99:0.0175012365,WP_145962313.1-Pectobacterium-peruviense:0.0680297613)97/99:0.1246145690)66.7/99:0.0624807934,WP_211007190.1-Pectobacterium-brasiliense:0.5989193393)67.8/58:0.0533346540)95.1/89:0.0749765993,(((((((((WP_236369955.1-Aeromonas-veronii:0.0599693347,WP_113736183.1-Aeromonas-caviae:0.0405659706)75.6/61:0.0079146063,((((((WP_080988732.1-Aeromonas-caviae:0.0556648614,WP_108536035.1-Aeromonas-sp.-HMWF014:0.0445605765)92.5/100:0.0430216304,(WP_252615985.1-Aeromonas-hydrophila:0.0690866122,WP_041208446.1-Aeromonas-jandaei:0.2726004536)75.7/99:0.0196001915)88.2/99:0.0235014298,WP_214003767.1-Aeromonas-caviae:0.0664802466)0/95:0.0000022036,(WP_113739100.1-Aeromonas-veronii:0.0703226714,(WP_226032029.1-Aeromonas-hydrophila:0.0300839762,WP_139459904.1-Aeromonas-veronii:0.1683842954)81.3/88:0.0197253474)80.4/78:0.0159128071)43.1/82:0.0177642852,WP_143237783.1-Aeromonas-caviae:0.0609013740)99.3/99:0.1141535269,((WP_042019775.1-Aeromonas-sobria:0.0630948287,WP_139495680.1-Aeromonas-veronii:0.0291969763)95.8/100:0.0662809550,(WP_098969895.1-Aeromonas-sp.-CU5:0.0586865703,WP_159163188.1-Aeromonas-veronii:0.0586117279)89.4/100:0.0450654249)87.5/100:0.0338343700)75.2/23:0.0101819601)99.8/98:0.0954009681,(WP_220777726.1-Shewanella-hafniensis:0.0392729170,WP_055647506.1-Shewanella-sp.-Sh95:0.0313380509)92.6/100:0.0258215335)44.1/18:0.0204454239,(WP_263173612.1-Shewanella-sp.-SM73:0.1729682731,((WP_263191039.1-Shewanella-sp.-SM87:0.0810066139,WP_263255112.1-Shewanella-sp.-SM32:0.0530924890)77.2/95:0.0193727997,(WP_263178828.1-Shewanella-sp.-SM101:0.0483775021,WP_109248233.1-Shewanella-algae:0.0800224345)93.8/100:0.0495866575)86.5/100:0.0505643182)94.5/100:0.0853996239)92.8/22:0.0615153052,WP_201035193.1-Aeromonas-media:0.1809009153)17.9/55:0.0108600948,WP_082969903.1-Aeromonas-dhakensis:0.1410490179)88.3/61:0.0445663701,WP_104454419.1-Aeromonas-sp.-ASNIH2:0.0210000274)95.7/100:0.1106134504,WP_223952962.1-Aeromonas-caviae:0.2771525072)38.6/48:0.0297152189,(((WP_073350970.1-Aeromonas-hydrophila:0.1554870668,WP_190284026.1-Aeromonas-hydrophila:0.0455324319)82.6/100:0.0398086042,WP_241344946.1-Aeromonas-sp.-MR16:0.0866107058)93.7/100:0.1076005823,(((WP_208149206.1-Shewanella-algae:0.3827837756,(((WP_045500143.1-Vibrio-hyugaensis:0.2226816883,WP_045419977.1-Vibrio-campbellii:0.1028380216)39.6/94:0.0420990327,WP_025533544.1-Vibrio-parahaemolyticus:0.0865859153)94.2/100:0.0561037600,WP_011788809.1-Shewanella-sp.-W3-18-1:0.0878970750)68.1/100:0.0241399387)81.1/100:0.0470332667,WP_219142360.1-Aeromonas-sanarellii:0.1488023377)98.3/99:0.1951722786,WP_244081328.1-Aeromonas-caviae:0.2475114554)73.1/87:0.0583449121)85.4/51:0.0657327965)89.7/90:0.0311747644)0/23:0.0000020830,((((WP_248067241.1-unclassified-Shewanella:0.1346756623,WP_148505453.1-Vibrio-cholerae:0.1113255933)91.8/100:0.1047819154,(WP_218558465.1-Shewanella-putrefaciens:0.0957161189,WP_011919582.1-Shewanella-putrefaciens:0.0339354878)98.5/100:0.1364037640)93/93:0.0883244607,(((((WP_050718733.1-Aeromonas-tecta:0.1768258829,WP_100859741.1-Aeromonas-lusitana:0.0454573341)76.2/99:0.0114886180,WP_042654929.1-Aeromonas-encheleia:0.0665365899)96.8/100:0.0699405763,WP_077098334.1-Aeromonas-hydrophila:0.0665511987)79.1/100:0.0187289781,WP_043167431.1-Aeromonas-hydrophila:0.1283549621)70.4/64:0.0128543020,WP_252420256.1-Aeromonas-taiwanensis:0.0920143736)89.3/100:0.0661502059)76.3/58:0.0519983407,(WP_163316497.1-Crenobacter-sedimenti:0.1639583384,WP_160797365.1-Craterilacuibacter-sinensis:0.1883351401)99.7/100:0.2902538510)93.3/52:0.0855411380)87.3/50:0.0318730411,WP_108652490.1-Dongshaea-marina:0.3476971877)75.4/17:0.0124295121)67.9/16:0.0106379261,(WP_076573677.1-Aeromonas-sp.-RU39B:0.1973394147,WP_134696535.1-Aeromonas-taiwanensis:0.8152381760)95.6/32:0.1998519205)91.3/51:0.0616479978,(((((((((((((WP_071788415.1-Franconibacter-pulveris:0.5939059722,(WP_232581009.1-Photobacterium-phosphoreum:0.7015275001,(WP_073514120.1-Vibrio-parahaemolyticus:0.0791065291,WP_032481483.1-Vibrio:0.0117101616)99.7/100:0.7854723435)96.2/99:0.4350297269)54.2/59:0.0791260167,(WP_020232007.1-Escherichia-coli:0.3458274543,(WP_228338545.1-Enterobacter-hormaechei:0.1588882898,(WP_109845273.1-Enterobacter:0.0678503015,WP_150146763.1-Enterobacter-roggenkampii:0.0465966832)89.4/100:0.1082107262)97.7/97:0.2255635959)80.2/62:0.0913100796)97.4/98:0.1590735777,(((((((((((((((((((((((((((((((((((((((WP_051234278.1-Halomonas-halocynthiae:0.8497124538,WP_110648656.1-Salinicola-peritrichatus:0.4244847816)93.1/99:0.2620471228,WP_254581577.1-Marinobacterium-sedimentorum:0.3489106109)68.8/93:0.1201472882,(WP_179929642.1-Halomonas-salicampi:0.5645044932,(((((((WP_066801876.1-Moraxella-oblonga:0.2199604709,(WP_197128702.1-Psychrobacter-sp.-NZS113:0.2466728528,WP_201616712.1-Psychrobacter-immobilis:0.1798844274)96.1/100:0.1921545817)99.6/100:0.3056025099,((((WP_041157945.1-Halomonas-sp.-KHS3:0.1255933724,WP_141441391.1-Halomonas-titanicae:0.0757531481)76.1/98:0.0334172383,WP_226251132.1-unclassified-Halomonas:0.0796623542)86.6/98:0.0540085416,WP_120386672.1-Halomonas-alkaliphila:0.1288493222)83/100:0.0896168337,WP_228219791.1-Halomonas-meridiana:0.1694227386)88.1/44:0.0829540163)82.4/26:0.0536487642,((((WP_158774340.1-Cobetia-sp.-L2A1:0.0863660915,(WP_144727127.1-Cobetia-crustatorum:0.0408401023,WP_088744253.1-Cobetia-sp.-QF-1:0.0758060884)27.2/100:0.0071214641)88.3/100:0.0387602813,WP_158774127.1-Cobetia-sp.-L2A1:0.0295134673)84.8/100:0.0186617593,(WP_077373671.1-Cobetia:0.0593105610,((WP_213115235.1-Cobetia-sp.-MC34:0.0445287948,WP_200017180.1-Cobetia-sp.-cqz5-12:0.0732349625)94.3/100:0.0706010317,WP_225346840.1-Cobetia-amphilecti:0.0535248267)81/100:0.0364961336)92.6/100:0.0366973332)99.3/100:0.1436006184,((WP_036548095.1-Nitrincola-lacisaponensis:0.4687143637,WP_017842145.1-Methylotuvimicrobium-buryatense:0.3808174563)78.4/75:0.1521039114,(WP_251978361.1-unclassified-Salinicola:0.0907083407,WP_189443129.1-Salinicola-rhizosphaerae:0.1413435235)97.5/97:0.1770348727)63.3/70:0.0528770060)34.3/27:0.0241220445)94.6/46:0.0858564697,(((((((((WP_224850395.1-Marinobacter-nauticus:0.2143525627,WP_110630630.1-Salinicola-salarius:0.0970941071)90.9/98:0.0662782224,((WP_193085952.1-Halomonas-sp.-FME65:0.1886734877,WP_089675781.1-Halomonas-aquamarina:0.0288451390)80.6/56:0.0207092003,((WP_253557393.1-unclassified-Halomonas:0.0606456347,WP_089676106.1-Halomonas-aquamarina:0.0343781488)0/91:0.0000024363,WP_062362595.1-Halomonas-axialensis:0.0436082423)38.8/89:0.0322810794)50.2/34:0.0151198256)15/67:0.0189163042,WP_027961801.1-Halomonas-halodenitrificans:0.0491700257)0/90:0.0000022959,WP_079551663.1-Halomonas-subglaciescola:0.0900657768)91.7/92:0.0309171532,WP_195687443.1-Halomonas-sp.-KAO:0.0628306906)86.2/97:0.0396830548,((WP_159342670.1-Halomonas-meridiana:0.3799547422,WP_253554573.1-Halomonas-sp.-DN3:0.2183634093)73.5/100:0.0469748861,WP_102589198.1-Halomonas-urumqiensis:0.1422166770)92.1/100:0.0618116584)74.6/99:0.0164340278,(WP_193343673.1-Halomonas-elongata:0.0736566047,WP_064701650.1-Halomonas-caseinilytica:0.0197729295)99.3/100:0.1270465200)96.2/99:0.0847380625,WP_035566812.1-Halomonas:0.2964906837)86.4/100:0.0411080389,(WP_149282777.1-Halomonas-sp.-Y2R2:0.0729586242,WP_073436013.1-Halomonas-cupida:0.1200266113)96.5/99:0.1077152344)75.6/82:0.0361126940)91.7/42:0.0423475986,((WP_248234415.1-Marinobacter-bryozoorum:0.1187680540,(WP_228223924.1-Marinobacter-nauticus:0.0727611772,WP_098418752.1-Marinobacter-sp.-LV10MA510-1:0.1634054359)89.8/100:0.0852262821)95.5/100:0.1002506570,WP_021818206.1-Halomonas-huangheensis:0.2048982691)85.7/85:0.0526208276)83.7/51:0.0318898792,((WP_127061154.1-Halomonas-nanhaiensis:0.2085054237,((((((WP_030071911.1-Halomonas-alkaliantarctica:0.0571646091,WP_095603285.1-Halomonas-humidisoli:0.0316885885)73.5/63:0.0229678739,WP_253485683.1-Halomonas-sp.-XH26:0.0347499836)43.6/63:0.0135206498,WP_038485954.1-Halomonas-campaniensis:0.0267897416)98.4/100:0.0915136175,WP_162218282.1-Halomonas-alkaliphila:0.0369284737)95/100:0.0543746028,WP_078087027.1-unclassified-Halomonas:0.0295512454)78.9/100:0.0165634955,WP_096655487.1-Halomonas-nigrificans:0.0424886078)99.2/100:0.1519597388)41.6/65:0.0143636267,(WP_096281135.1-Halomonas-sp.-JB37:0.2705518108,(WP_027958907.1-unclassified-Halomonas:0.0484042678,WP_064236373.1-unclassified-Halomonas:0.0639762965)95.6/100:0.1223264156)66.6/97:0.0684414916)81.2/69:0.0277411809)81.9/64:0.0914941036,WP_081193716.1-Halomonas-sp.-BC1:0.3467619350)76.8/71:0.0502213499)70.8/65:0.0404308019)93.4/100:0.1439954010,(WP_015904383.1-Desulforapulum-autotrophicum:0.3370353521,WP_111960646.1-Desulfobacter-hydrogenophilus:0.4213409567)93.4/97:0.2457268421)80.7/89:0.0554822862,(((((WP_151033206.1-Cellvibrio-sp.-KY-GH-1:0.0975103596,WP_049631514.1-Cellvibrio-sp.-pealriver:0.0936539368)70.4/100:0.1056286181,WP_062059887.1-Cellvibrio-sp.-OA-2007:0.0486809393)99.7/100:0.4167426134,WP_178968040.1-Neptunomonas-phycophila:0.4224864596)70.4/93:0.1832463616,WP_156872703.1-Ketobacter-sp.-MCCC-1A13808:0.6023766420)38.5/84:0.0950037365,((((((WP_125841544.1-Stutzerimonas-stutzeri:0.4598109149,((WP_262731117.1-Alcanivorax-marinus:0.2434324289,(WP_106232428.1-Halomonas-ventosae:0.0969323091,WP_228234339.1-Alcanivorax-marinus:0.2567804302)87.3/100:0.0516663307)83.5/100:0.0311043182,WP_175070852.1-Halomonas-taeanensis:0.1433324195)18.3/99:0.0977698424)87.1/100:0.1032376094,(WP_019018778.1-Halomonas-lutea:0.3712197808,WP_092999033.1-Thiohalomonas-denitrificans:0.1492796187)92/100:0.1770006034)87.3/100:0.0669271077,((WP_146294265.1-Euhalothece-natronophila:0.3077214204,WP_159299499.1-Spiribacter-sp.-SSL99:0.2940478584)91.7/100:0.1263768898,WP_011630054.1-Alkalilimnicola-ehrlichii:0.2255458564)88.6/100:0.0815918138)85.9/100:0.0691055217,WP_127477393.1-Sulfurivermis-fontis:0.3894887143)87.3/100:0.0636077270,WP_028988655.1-Thermithiobacillus-tepidarius:0.3286851464)90.3/100:0.1006503370,WP_069857109.1-Desulfoplanes-formicivorans:0.5000076738)95.7/100:0.1635738408)81.3/81:0.0532252645)95.9/92:0.1202357570,(WP_255857717.1-Bacterioplanoides-sp.-SCSIO-12839:0.3677919760,(((WP_262601261.1-Endozoicomonas-euniceicola:0.1250288505,WP_066015073.1-Endozoicomonas-atrinae:0.1743425764)92/100:0.1333414536,WP_257287521.1-Endozoicomonas-sp.-SESOKO2:0.1397311894)94.1/100:0.1578667873,(WP_067585802.1-Endozoicomonas-ascidiicola:0.1641517076,(WP_020580625.1-Endozoicomonas-elysicola:0.1228874737,WP_257293945.1-Endozoicomonas-sp.-YOMI1:0.1686984412)91.1/100:0.1222178083)96.4/100:0.2744760455)99.7/99:0.3600475105)76.2/23:0.1281499267)86.9/23:0.0568798865,(((WP_028862352.1-Psychromonas-aquimarina:0.2243055526,WP_187408568.1-Saccharophagus-sp.-K07:0.1933658322)91.6/100:0.0786546515,WP_251261538.1-Echinimonas-agarilytica:0.1841930279)82.3/95:0.0494907120,(WP_096084688.1-Agaribacterium-haliotis:0.4286321922,WP_188151970.1-Teredinibacter-waterburyi:0.2839709103)29.9/80:0.1183088536)79.2/98:0.1478797434)80.1/20:0.0263790510,(((WP_248714967.1-Agarivorans-sp.-TSD2052:0.0855763978,WP_152781274.1-Agarivorans-sp.-B2Z047:0.0960384263)98.9/100:0.1692916533,WP_116686321.1-Pelagibaculum-spongiae:0.7726364241)58.7/23:0.0215042281,(WP_220103444.1-Neiella-holothuriorum:0.1262485915,WP_087505717.1-Neiella-marina:0.2001225567)98.4/100:0.1430894731)84/23:0.0269824929)85.4/44:0.0740017330,(((WP_142941196.1-Aliikangiella-marina:0.4043119392,(WP_081148784.1-Cognaticolwellia-beringensis:0.0600478402,WP_007634918.1-Paraglaciecola-psychrophila:0.0419535867)99.3/100:0.2344511992)18/92:0.0449859252,WP_068546663.1-Thalassotalea-crassostreae:0.2743084208)83.1/98:0.0564918032,(((((((((WP_160153838.1-Microbulbifer-sp.-ALW1:0.3574828829,((WP_263600466.1-Microbulbifer-agarilyticus:0.1629510057,WP_078085574.1-Microbulbifer-mangrovi:0.1091022183)57.5/97:0.0153943030,WP_029249715.1-Microbulbifer-agarilyticus:0.1228642652)87.4/97:0.0757518061)97.2/100:0.1584690899,(((WP_085356321.1-Neisseria-dumasiana:0.2393965530,((WP_101133177.1-Neisseria-meningitidis:0.0704904898,WP_005552781.1-Aggregatibacter-actinomycetemcomitans:0.2562049230)8.5/73:0.0202522448,(WP_127202078.1-Neisseria-meningitidis:0.1491992159,WP_237092367.1-Neisseria-sp.-ZJ104:0.1842801325)92/100:0.0712729371)75.3/75:0.0578531910)92.8/100:0.2011892120,WP_031579865.1-Ruminobacter-sp.-RM87:0.5654560684)92.6/97:0.1609094654,((WP_138624780.1-Pseudoalteromonas-sp.-S4492:0.2595746114,WP_130050482.1-Pseudoalteromonas-sp.-MEBiC-03485:0.1977652283)89.2/100:0.1022750579,(WP_189610147.1-Saccharospirillum-salsuginis:0.3494999273,WP_111640782.1-Marinimicrobium-alkaliphilum:0.4450966361)75.8/94:0.0730781045)82.2/92:0.1284738409)49.5/82:0.0357319087)51.7/87:0.0325580674,WP_118961222.1-Colwellia-sp.-RSH04:0.5052010955)57.4/99:0.0291778922,WP_186505417.1-Neptunicella-marina:0.2499223801)90.5/99:0.0814183301,(WP_102798355.1-Bowmanella-denitrificans:0.0903564020,WP_214344498.1-Bowmanella-yangjiangensis:0.0417752308)99.8/100:0.2351482812)32.5/81:0.0240784353,(WP_124026287.1-Alteromonas-sediminis:0.1786286021,WP_100655941.1-Alteromonas-flava:0.3012183233)99.7/99:0.2125510252)75/77:0.0218619941,(WP_216048042.1-Aliiglaciecola-lipolytica:0.2727579532,(WP_131095783.1-Alteromonas-sp.-KUL42:0.0868949303,((((((WP_100972994.1-Alteromonas-sp.-MB-3u-76:0.1605349466,(WP_071816068.1-Alteromonas-sp.-V450:0.1113717752,WP_198816918.1-Alteromonas-sp.-IB21:0.0294376094)67.7/100:0.0063948235)77.8/23:0.0092803436,(((WP_156078883.1-Alteromonas:0.0683301898,((((WP_063457834.1-Alteromonas-stellipolaris:0.0391272079,(WP_202940767.1-Alteromonas-macleodii:0.0266317873,WP_061095937.1-Alteromonas-macleodii:0.0487997043)84.5/65:0.0146768493)0/64:0.0000020770,((WP_044058190.1-Alteromonas-australica:0.0481422468,WP_159580202.1-Alteromonas-sp.-KUL106:0.0236480860)74.8/97:0.0081285850,(WP_014977514.1-Alteromonas-macleodii:0.0417193873,WP_239220085.1-Alteromonas-sp.-MmMcT2-2:0.0323295565)73.9/46:0.0068507261)50/23:0.0078449560)80.9/20:0.0151604680,WP_187447473.1-Alteromonas-sp.-BZK5:0.0237123372)75.3/18:0.0085392233,WP_202262575.1-unclassified-Alteromonas:0.0493349683)64.2/19:0.0130831366)89.9/96:0.0286700596,(WP_163106912.1-Alteromonas-genovensis:0.0606527961,WP_057795858.1-Alteromonas-stellipolaris:0.0403590326)54.4/31:0.0034318375)42/70:0.0168876113,WP_269520955.1-Alteromonas-sp.-BMJM2:0.0910241511)75.8/22:0.0092733656)88.6/85:0.0307017730,WP_130638173.1-Alteromonas-sp.-KUL49:0.0889391726)0/48:0.0154618356,WP_115592514.1-Alteromonas-aestuariivivens:0.0731343319)71.9/56:0.0073527177,WP_163085646.1-Alteromonas-profundi:0.0344105868)90.2/99:0.0389188516,((((WP_262993134.1-Alteromonas-sp.-ASW11-19:0.1058260092,WP_195811108.1-Salinimonas-marina:0.0709957594)88.2/66:0.0376179394,WP_018984642.1-Salinimonas-chungwhensis:0.1252382289)73/34:0.0136734108,WP_218354448.1-Alteromonas-lipotrueiana:0.0926056849)88.7/64:0.0428399394,((((WP_265616768.1-Alteromonas-sp.-ASW11-7:0.1210917335,WP_119499891.1-Alteromonas-sp.-RKMC-009:0.0657384097)10.6/38:0.0279452964,WP_265607972.1-Alteromonas-sp.-ASW11-130:0.1026547696)50.3/28:0.0095758936,(WP_216026947.1-Aestuariibacter-sp.-A3R04:0.0747153707,(WP_070176406.1-Alteromonas-lipolytica:0.0442244129,WP_073325432.1-Marisediminitalea-aggregata:0.0601871293)92.7/100:0.0655450668)71.1/84:0.0261076063)63.4/28:0.0089464427,WP_216028209.1-Alteromonas-sp.-C1M14:0.0790406359)88.7/34:0.0352857990)71.7/58:0.0387811633)91.8/98:0.0543862003)92.1/99:0.0585857366)88.6/76:0.0474521743)94/76:0.0415241865,(WP_057181266.1-Colwellia:0.2492459876,WP_009839692.1-Pseudoalteromonas-tunicata:0.7481478956)74.9/91:0.0192032252)93.1/96:0.0787786470,(WP_250459773.1-Microbulbifer-sp.-GX-H0434:0.1484604001,WP_152451766.1-Microbulbifer-sp.-THAF38:0.2140270777)97.9/100:0.1676979636)79.4/96:0.0341319302)90.2/46:0.0800725535)90.7/67:0.0603282364,WP_138986566.1-Neptunomonas-concharum:0.4362548821)80.1/39:0.0280760971,((WP_167868984.1-Shewanella-sp.-WXL01:0.5361610060,(WP_016959527.1-Enterovibrio-norvegicus:0.0943262275,WP_229491984.1-Enterovibrio-norvegicus:0.1553309408)85.2/99:0.0544399575)44.6/32:0.0146559653,(((((WP_222525605.1-Ferrimonas-balearica:0.1834270433,WP_013344594.1-Ferrimonas-balearica:0.1759997989)99.6/100:0.2587138294,WP_084884043.1-Vibrio-sp.-qd031:0.1495862331)78.2/51:0.0269960211,((WP_193630602.1-Shewanella-benthica:0.2549908305,WP_159656123.1-Vibrio-atypicus:0.0578583744)82.5/94:0.0779703510,(WP_269599241.1-Salinivibrio-kushneri:0.0775360116,WP_136483218.1-Vibrio-sp.-H11:0.0873107073)92.7/100:0.0868156058)79/78:0.0436908836)75.9/47:0.0221310368,((((WP_047049276.1-Vibrio-mexicanus:0.2138623896,WP_047874860.1-Photobacterium-aphoticum:0.2787428975)59.9/69:0.0199644544,WP_020335761.1-Vibrio-natriegens:0.1378806828)83/46:0.0335112176,((((WP_050648448.1-Vibrio-sp.-J2-4:0.1823884657,WP_205390674.1-Vibrio:0.1075664479)14.1/29:0.0156817655,WP_140076235.1-Vibrio-parahaemolyticus:0.1608648776)79.4/19:0.0200147006,(WP_012143346.1-Shewanella-sediminis:0.2995182927,(((((WP_167678613.1-Shewanella-aestuarii:0.1785684942,WP_137222113.1-Shewanella-sp.-MEBiC00475:0.0382915130)71.1/98:0.0469316535,WP_188955407.1-Shewanella-ulleungensis:0.0760552596)92.9/99:0.0839260467,(WP_155694061.1-Psychrosphaera-haliotis:0.1831967721,((((WP_138577097.1-unclassified-Pseudoalteromonas:0.1243539693,(WP_264390127.1-Pseudoalteromonas-sp.-A3:0.0304311400,(WP_096739047.1-Pseudoalteromonas-atlantica:0.0479251368,WP_248462211.1-Pseudoalteromonas-sp.-2CM37A:0.0224211546)90.7/99:0.0270144638)77.3/85:0.0177291673)95.3/75:0.0522422828,WP_126005793.1-Pseudoalteromonas-sp.-Xi13:0.0397444278)66.6/12:0.0073089451,WP_256848444.1-Pseudoalteromonas-carrageenovora:0.0291448687)0/11:0.0168153430,WP_182693438.1-Pseudoalteromonas-sp.-SR45-5:0.0657552211)88.8/79:0.0503832004)9.9/29:0.0339906797)90.8/86:0.0678300434,(WP_113818410.1-Psychromonas-sp.-B3M02:0.0724918450,(WP_025564776.1-Psychromonas-sp.-SP041:0.0584895778,WP_166372414.1-Psychromonas-sp.-SA13A:0.0165173742)71.4/100:0.0609284613)88.1/99:0.0539397668)65.6/27:0.0129016036,WP_022943097.1-Psychromonas-hadalis:0.1990738968)82.5/85:0.0306223767)12.6/18:0.0160352198)43.8/21:0.0186078696,WP_264874461.1-Vibrio-agarivorans:0.1599468857)73.8/39:0.0096273505)86.6/41:0.0191556730,WP_164649367.1-Vibrio-astriarenae:0.1109684192)80.8/94:0.0235870820)75.1/59:0.0146072272,WP_093326971.1-Thalassotalea-agarivorans:0.1660568526)57/33:0.0666116410)92.9/96:0.0675457194)98.9/92:0.1858823289,WP_234495031.1-Vibrio-maritimus:0.1517730628)93.3/79:0.0978665686,(WP_062462511.1-Vibrio-mediterranei:0.1162694183,WP_171392669.1-Vibrio:0.1363159870)72.3/90:0.0563298035)88.3/79:0.0722815355,((((((((WP_230291920.1-Croceicoccus-sp.-Ery5:0.4665857522,WP_164156863.1-Sandarakinorhabdus-rubra:0.4397112793)88.4/100:0.1873444108,(WP_116367876.1-Parahaliea-mediterranea:0.1103237242,WP_148063811.1-Parahaliea-aestuarii:0.1250922156)99.5/100:0.3911224250)78.3/89:0.1178906072,WP_129645432.1-Steroidobacter-agariperforans:0.4318568580)98.8/100:0.2979102030,((((((((WP_036226274.1-Mesoaciditoga-lauensis:0.2866329278,((WP_077198159.1-Thermosipho-affectus:0.0881018407,(WP_012057290.1-Thermosipho-melanesiensis:0.2118406953,(WP_127352844.1-Caldicellulosiruptor-changbaiensis:0.0806687188,WP_014043315.1-Caldicellulosiruptor-acetigenus:0.0268644877)100/100:0.5335646444)49.4/91:0.0530349613)97.3/100:0.1202777226,WP_047753768.1-Kosmotoga-pacifica:0.2119904070)78.9/95:0.0229217001)64/95:0.0520434343,((((((((WP_117708548.1-Fusobacterium-mortiferum:0.0036502952,WP_204716698.1-Fusobacterium-mortiferum:0.0835489866)78.3/100:0.0242626098,WP_115270465.1-Fusobacterium-necrogenes:0.0795767659)60.2/100:0.0243364638,WP_204716813.1-Fusobacterium-mortiferum:0.5343969547)86.1/95:0.0775936749,(((WP_130889691.1-Fusobacterium-ulcerans:0.0494281119,((WP_005950923.1-Fusobacterium-varium:0.0391000705,WP_070580525.1-Fusobacterium:0.0661957927)39.8/100:0.0138829827,WP_227372109.1-Fusobacterium-ulcerans:0.0401426600)86.2/100:0.0189004861)73.4/100:0.0123815408,(WP_241635364.1-Fusobacterium-gastrosuis:0.1377619041,(WP_074518138.1-Fusobacterium-necrophorum:0.0967316399,(WP_211495349.1-Fusobacterium-necrophorum:0.1614448876,WP_032890075.1-Fusobacterium-nucleatum:0.8340414934)84.8/99:0.0862384025)93.1/99:0.0888159834)98.6/99:0.0859091469)84.5/99:0.0332432159,WP_261793436.1-Fusobacterium-varium:0.3227652467)85.4/94:0.0596107826)97.1/100:0.1712796607,((WP_147004474.1-Leptotrichia-wadei:0.0315481936,WP_018499460.1-Leptotrichia-wadei:0.0384129567)95.5/100:0.0993635650,(WP_068155536.1-Leptotrichia-sp.-oral-taxon-847:0.0796312793,(WP_099992503.1-Fusobacterium-pseudoperiodonticum:0.1466209911,(WP_211617428.1-Leptotrichia-sp.-oral-taxon-218:0.0179364013,WP_094080762.1-Leptotrichia-sp.-oral-taxon-498:0.0550407141)96.1/100:0.0640756792)59.2/100:0.0447803706)88.6/100:0.0740837663)99.5/100:0.2794450617)73.1/91:0.0591958199,WP_013388736.1-Ilyobacter-polytropus:0.6223170011)21.1/87:0.0426899779,WP_173194267.1-Paenibacillus-frigoriresistens:0.3966383413)36.5/75:0.0164473351,WP_012548442.1-Dictyoglomus-thermophilum:0.5026365404)97.1/100:0.1304106058)97.7/100:0.1304731274,(WP_099829233.1-Mesotoga-sp.-Brook.08.105.5.1:0.2028969970,WP_099830786.1-Mesotoga-sp.-Brook.08.105.5.1:0.2270354198)98.9/100:0.1800460365)91.8/100:0.1142173688,(((((((((((((((((WP_054742706.1-Cellulosilyticum-ruminicola:0.1081314192,(WP_177671917.1-Zhenhengia-yiwuensis:0.0562498079,WP_053982981.1-Niameybacter-massiliensis:0.1332368553)98.7/100:0.1344132308)93.7/100:0.0837827094,WP_069999280.1-Cellulosilyticum-sp.-I15G10I2:0.2735668394)98.9/100:0.1911110926,((((((((((((((WP_169105633.1-Clostridium-sp.-SM-530-WT-3G:0.0586580540,WP_077866663.1-Clostridium-saccharobutylicum:0.2831134918)29.1/36:0.0216754025,WP_027636628.1-Clostridium-butyricum:0.1150168369)90/36:0.0394415627,(((((WP_009169246.1-unclassified-Clostridium:0.1509421774,WP_160359894.1-Clostridium-chromiireducens:0.3979727315)0/15:0.0041649314,WP_077843022.1-Clostridium-beijerinckii:0.1805985914)75/15:0.0286600806,WP_041896941.1-Clostridium-beijerinckii:0.2546134717)84.9/16:0.0743045809,WP_069680896.1-Clostridium-taeniosporum:0.2640852320)93.9/92:0.0894455006,WP_265859166.1-Clostridium-botulinum:0.1487382420)81.3/46:0.0241566402)76.5/24:0.0133401728,(WP_022750218.1-Clostridium-saccharobutylicum:0.0541777464,(WP_226126241.1-Clostridium-algoriphilum:0.1565801955,((WP_154531076.1-Inconstantimicrobium-porci:0.1794600572,WP_009169009.1-Clostridium-sp.-DL-VIII:0.0163740573)83.7/100:0.0601519029,WP_077856313.1-Clostridium-beijerinckii:0.0058910505)96.8/100:0.1175455379)0/47:0.0024388855)87.3/45:0.0182334309)89.9/38:0.0231848236,(WP_185652834.1-Clostridium-sp.-DJ247:0.0264285019,WP_185652641.1-Clostridium-sp.-DJ247:0.0857480070)91.1/100:0.0281689543)83.8/38:0.0156146752,WP_077848125.1-Clostridium-puniceum:0.1867144462)73.2/20:0.0069832873,(WP_015391310.1-Clostridium-saccharoperbutylacetonicum:0.0697238229,(WP_077846315.1-Clostridium-puniceum:0.0622116368,WP_238915641.1-Clostridium-sp.-YIM-B02555:0.0764673804)87.7/100:0.0255792216)95.7/100:0.0433591962)63.9/20:0.0043806677,((WP_261829154.1-Clostridium-sp.-TW13:0.1015847539,WP_224033465.1-Clostridium-gelidum:0.1330892960)0/37:0.0000023266,(WP_077855754.1-Clostridium-beijerinckii:0.0512414062,WP_041894754.1-Clostridium-beijerinckii:0.0325063090)99.4/100:0.0937517624)85.9/38:0.0187033831)98.6/99:0.1245983107,WP_216390068.1-Clostridium-sp.-MSJ-8:0.2976866227)60.7/98:0.0223003174,WP_090010800.1-Clostridium-sp.-DSM-8431:0.1252072519)81.3/100:0.0545749900,(WP_106059304.1-Clostridium-vincentii:0.1286894001,WP_143316579.1-Clostridium:0.0281618475)81.9/100:0.0214000484)92.4/100:0.0527668748,WP_055184115.1-Clostridium-paraputrificum:0.0715504678)93.6/100:0.0472252730,(WP_261829490.1-Clostridium-sp.-TW13:0.2384823793,WP_148566871.1-Acetobacterium-paludosum:0.5152038259)0/97:0.0000020806)95.1/100:0.0383038666,(WP_066825124.1-Clostridium-tepidiprofundi:0.1314453447,WP_072889493.1-Tepidibacter-formicigenes:0.1094628752)76/88:0.0184274731)64.2/84:0.0110382293)30.5/43:0.0209121142,(((((((WP_094369216.1-Romboutsia-weinsteinii:0.2049324168,WP_021359979.1-Clostridioides-difficile:0.4157897841)68.8/41:0.0519312103,(WP_074918830.1-Terrisporobacter-glycolicus:0.0833047459,WP_195929149.1-Clostridium-sp.-1001270J_160509_D11:0.0478083044)84.4/100:0.0280314696)92.5/41:0.0519266686,(((WP_092726456.1-Romboutsia-lituseburensis:0.2155452145,WP_258242885.1-Romboutsia-lituseburensis:0.0384922499)99.3/100:0.1185776302,WP_057560648.1-Paeniclostridium-sordellii:0.0831962404)31.8/28:0.0299373399,((WP_057580750.1-Paeniclostridium-sordellii:0.1419820225,WP_057538715.1-Paeniclostridium-sordellii:0.0541530470)78.9/88:0.0168110586,WP_195252007.1-Romboutsia-sp.-1001713B170207_170306_H8:0.0637722425)52.8/85:0.0185486072)91.9/99:0.0443948209)87.2/21:0.0396781781,(((WP_029452249.1-Clostridium-algidicarnis:0.1104637943,(WP_216183125.1-Clostridium-gasigenes:0.0297408396,WP_216313757.1-Clostridium-gasigenes:0.1097710982)10/90:0.0374420152)96/100:0.0698643208,WP_066677671.1-Clostridium-septicum:0.0501538744)86.2/87:0.0258225131,((((((((WP_152890015.1-Clostridium-tarantellae:0.1783173909,WP_047001949.1-Clostridium-sp.-C8:0.0498360898)85.4/100:0.0305517918,WP_195959433.1-Clostridium-tertium:0.0836577102)28.8/100:0.0086958083,WP_195429094.1-Clostridium-sp.-D46t1_190503_E9:0.0277913563)91.7/100:0.0365301903,WP_066890539.1-Clostridium-nigeriense:0.0622000910)72.5/84:0.0081481537,WP_016207773.1-Clostridium-sartagoforme:0.0386018255)90.7/99:0.0286366548,WP_197096717.1-Paeniclostridium-sordellii:0.1254482595)52.3/75:0.0178356073,WP_238903698.1-Clostridium-sp.-YIM-B02506:0.0929475766)0/69:0.0023904992,WP_095406848.1-Romboutsia-maritimum:0.1540709983)86/80:0.0276924763)73/20:0.0105370750)75.1/12:0.0120428233,(((WP_195972826.1-Clostridium-thermobutyricum:0.1112911299,(WP_209795868.1-Clostridium-moniliforme:0.0856320423,WP_194191032.1-Clostridium-chrysemydis:0.1343459712)95.2/100:0.0624005587)76.1/100:0.0145256262,WP_040194942.1-Clostridium-culturomicium:0.0913242400)82.4/95:0.0304295059,(WP_073337580.1-Clostridium-grantii:0.1763109359,(WP_072892386.1-Clostridium-fallax:0.1606512997,WP_072832886.1-Clostridium-collagenovorans:0.0361832699)73.7/99:0.0204250698)35.5/99:0.0129128424)94.3/99:0.0490824881)97.3/23:0.0504526071,((((((WP_055265352.1-Clostridium:0.0351195252,(WP_186859336.1-Clostridium-sp.-NSJ-6:0.0703820036,(WP_055265304.1-Clostridium-disporicum:0.0759590855,(WP_154821366.1-Clostridium-saudiense:0.0121333034,WP_195245504.1-Clostridium-celatum:0.0806900247)74.4/100:0.0311437676)77.7/100:0.0392971459)87.1/100:0.0658742271)92.1/100:0.0895466949,WP_186834444.1-Eubacteriales:0.1508726486)94.7/100:0.0736900766,((WP_066874757.1-Clostridium-mediterraneense:0.2379273195,WP_198610231.1-Clostridium-perfringens:0.0251330621)88.2/18:0.0470006155,((WP_235068992.1-Turicibacter-sp.-TJ11:0.0742128454,((WP_161831114.1-Turicibacter-sp.-TS3:0.0682821613,WP_155223812.1-Turicibacter-sanguinis:0.0289764036)78.7/86:0.0213200555,WP_195933776.1-Turicibacter-sanguinis:0.0312509041)36.4/86:0.0082339373)87.6/86:0.0320671190,WP_212726030.1-Turicibacter-bilis:0.0151418748)99.9/100:0.1181128262)29.3/14:0.0247691763)93.9/22:0.0416478835,WP_055258173.1-Sarcina-ventriculi:0.0972374274)78/20:0.0176653997,WP_055069574.1-Clostridium-massiliamazoniense:0.1800684357)91.5/22:0.0384837497,WP_133628539.1-Fonticella-tunisiensis:0.0901967442)0/15:0.0000021570)3.8/11:0.0181165465,WP_163195303.1-Clostridium-thermarum:0.1695297625)83.6/69:0.0202659399)76.6/46:0.0406342216,(((WP_216288229.1-Clostridium-psychrophilum:0.1732357947,WP_216137569.1-Clostridium-estertheticum:0.0368311574)46/95:0.0103234390,WP_220704225.1-Clostridium-estertheticum:0.0477684899)93.2/100:0.0602504685,WP_220321605.1-Clostridium-tagluense:0.0760003389)98.4/100:0.1044528614)86/62:0.0453994946,(((WP_055666468.1-Desnuesiella-massiliensis:0.1290502601,WP_095404508.1-Romboutsia-maritimum:0.4138656423)45.7/63:0.0617714715,WP_072725426.1-Tepidibacter-thalassicus:0.2700742758)0/47:0.0000027245,WP_072990516.1-Clostridium-cavendishii:0.3239262779)28.7/16:0.0215030285)87/94:0.0466559866,(WP_142413383.1-Hathewaya-massiliensis:0.2490622124,WP_035130905.1-Clostridium-sulfidigenes:0.2992424499)98.4/99:0.1795418270)79.5/65:0.0296924903,WP_226126924.1-Clostridium-algoriphilum:0.4703125226)79.8/61:0.0287445661,(((WP_227847393.1-Clostridioides-sp.-ZZV15-6598:0.1335506017,WP_227846117.1-Clostridioides-sp.-ES-S-0145-01:0.0355475275)96.7/100:0.0540426599,(WP_227842647.1-unclassified-Clostridioides:0.0366980735,WP_003420982.1-Clostridioides-difficile:0.1096741942)84.7/99:0.0260687513)85.4/99:0.0369283840,WP_038810653.1-Clostridioides-difficile:0.0341573649)100/100:0.3651077396)82.3/63:0.0402698511,WP_079421587.1-Clostridium-oryzae:0.5282026132)2.9/58:0.0244873361,WP_099190526.1-Tepidibacter-mesophilus:0.2279103208)95.7/70:0.0884535463,WP_072903743.1-Hathewaya-proteolytica:0.5530072309)56/43:0.0233908739,(((((((WP_088538685.1-Methanobrevibacter-sp.-87.7:0.0797185858,WP_016359256.1-Methanobrevibacter-sp.-AbM4:0.1497631912)98.6/100:0.1290399283,WP_067148181.1-Methanobrevibacter-olleyae:0.1581431445)74.8/100:0.0398687289,WP_116670296.1-Methanobrevibacter-woesei:0.3299923751)99.9/93:0.1926014423,(WP_248612240.1-Methanobacterium-alcaliphilum:0.1676369172,((WP_071907175.1-Methanobacterium-congolense:0.0299392434,WP_209584978.1-Methanobacterium-aggregans:0.0526646502)99.7/100:0.1485321545,(WP_069583666.1-Methanobacterium:0.2419753548,WP_048190917.1-Methanobacterium-sp.-SMA-27:0.1746264493)88.4/97:0.0787886886)77.2/96:0.0296727664)82.8/94:0.0247764405)73.4/79:0.0170246051,(WP_011406138.1-Methanosphaera:0.1995414358,(WP_112123902.1-Methanosphaera-sp.-BMS:0.1292474520,(WP_069592847.1-Methanosphaera-sp.-WGK6:0.0388643909,WP_095608065.1-Methanosphaera-cuniculi:0.1407187982)98/100:0.1137988578)22.6/100:0.0431624504)97.7/100:0.1107012481)0/79:0.0006275380,(WP_013644504.1-Methanobacterium-lacus:0.2737410380,WP_054834988.1-Methanobrevibacter-arboriphilus:0.2463126435)89.5/95:0.0650932929)96/93:0.0792296615,(WP_112093115.1-Methanothermobacter-tenebrarum:0.3722493533,WP_013413362.1-Methanothermus-fervidus:0.2731799864)98.7/92:0.2099935441)51.9/16:0.0489882972)77.1/89:0.0250252444,((WP_009529486.1-Peptoanaerobacter-stomatis:0.2171226899,WP_068913445.1-Criibacterium-bergeronii:0.2266044743)99.1/100:0.2676401681,(WP_111570784.1-Halanaerobium-saccharolyticum:0.1047432602,(WP_089717478.1-Halanaerobium-congolense:0.0655676034,WP_089862761.1-Halanaerobium-salsuginis:0.0328568719)76/100:0.0333748298)99.7/100:0.3252618752)85.9/95:0.1057730162)76.4/39:0.0187223430,((((((WP_068555806.1-Thermotalea-metallivorans:0.1758579955,WP_073236503.1-Desulforamulus-putei:0.1351052721)72/100:0.0367279788,WP_095133872.1-Anaeromicrobium-sediminis:0.2468015405)87.1/81:0.0379969706,WP_035164300.1-Caloranaerobacter-azorensis:0.0941430135)79.8/49:0.0325466072,WP_090442301.1-Natronincola-peptidivorans:0.2083434515)88.9/97:0.0767535591,(((((WP_252130283.1-Clostridium-sp.-SYSU_GA19001:0.4109809082,WP_202749105.1-Clostridium-sp.-YIM-B02515:0.1468418060)97.6/99:0.1617197529,(((((WP_104410444.1-Clostridium-algidicarnis:0.0528544757,WP_185162668.1-Clostridium-algidicarnis:0.0846557619)99.9/100:0.2484587465,(WP_090038599.1-Clostridium-frigidicarnis:0.1888949665,WP_209459870.1-Youngiibacter-multivorans:0.2782464049)90/100:0.0793801867)63.7/51:0.0326306241,(WP_164955553.1-Clostridium-estertheticum:0.2405349071,(WP_125002390.1-Clostridium-tagluense:0.2198990322,(WP_226136064.1-Clostridium-estertheticum:0.0292779687,WP_216146875.1-Clostridium-frigoris:0.0912519948)98.3/100:0.1308515701)76.5/100:0.0421516592)75.8/100:0.1219862994)80.3/96:0.0303344208,((WP_084114600.1-Clostridium-acidisoli:0.1262872374,(WP_015614219.1-Clostridium-pasteurianum:0.0268165101,(WP_010235680.1-Clostridium-arbusti:0.0145864834,WP_003443690.1-Clostridium-pasteurianum:0.0636988490)91.4/100:0.0353351504)93.8/100:0.0450025382)39.3/99:0.0077343920,WP_026884435.1-Clostridium-akagii:0.1279343364)98.9/100:0.1226077969)72.9/36:0.0437032165,WP_216462460.1-Clostridium-bornimense:0.1939558126)87.8/39:0.0434378048)52.9/19:0.0451929240,((((WP_165412483.1-Clostridium-estertheticum:0.0818122949,WP_216302578.1-unclassified-Clostridium:0.0135974596)100/100:0.2699672297,WP_216457288.1-Clostridium-simiarum:0.1718518357)44.7/67:0.0350833063,(WP_204598070.1-Clostridium-pascui:0.1847410838,WP_027632043.1-Clostridium-hydrogeniformans:0.1773367651)72/95:0.0120711278)65.1/95:0.0564547661,(((WP_066398085.1-Cytobacillus-eiseniae:0.3806826064,(WP_056034132.1-Paenibacillus-sp.-Leaf72:0.1201726770,WP_197936429.1-Paenibacillus-sp.-GSMTC-2017:0.1948577678)92.6/100:0.0965738531)0.3/26:0.0287041976,(WP_076307499.1-Paenibacillus-odorifer:0.0776500128,((WP_042140168.1-Paenibacillus-sp.-FSL-P4-0081:0.0151501194,WP_036721118.1-Paenibacillus-sp.-FSL-R7-277:0.0906405529)91.4/100:0.0443760872,WP_074087222.1-Paenibacillus-sp.-P32E:0.0428230298)88/100:0.1091162269)98.5/100:0.1482884022)87.2/31:0.0541888308,(WP_262684732.1-Paenibacillus-sp.-WQ-127069:0.2354137330,WP_090798944.1-Paenibacillus-sp.-GP183:0.3328686896)69.7/31:0.0313778475)98.5/100:0.1774980582)89.4/20:0.0587687454)56.4/17:0.0276786240,WP_036935152.1-Pseudobacteroides-cellulosolvens:0.3009804905)73.1/34:0.0129884598,WP_234123145.1-Clostridium-hydrogenum:0.3244726302)87/72:0.0483460689)0/37:0.0000022623,((((WP_266162911.1-Caproicibacterium-sp.-BJN0003:0.1944022134,WP_256135338.1-Bittarella-massiliensis:0.4004708785)93/100:0.1804804034,WP_220321633.1-Clostridium-tagluense:0.3914333042)84.2/94:0.1103137398,((((WP_091684654.1-Anaerocolumna-aminovalerica:0.2546562222,(WP_033168390.1-Clostridium-sp.-KNHs205:0.1198543067,((WP_073588199.1-Anaerocolumna-xylanovorans:0.0406489052,WP_185256524.1-Anaerocolumna-chitinilytica:0.1894931418)74.2/100:0.0601726197,WP_073279502.1-Anaerocolumna-jejuensis:0.1537615449)26.1/99:0.0530504842)99.5/100:0.2336449722)95.6/100:0.1305090854,((((WP_253721803.1-Treponema-sp.-OMZ-787:0.7526367574,WP_005539496.1-Johnsonella-ignava:0.1799172166)87.9/99:0.0936558014,(WP_060930762.1-Lachnoanaerobaculum-saburreum:0.2516686986,(((((((((((((WP_024837362.1-Lacrimispora:0.1283845719,WP_077612318.1-Clostridium-sp.-Marseille-P2415:0.0437957755)0/98:0.0023453110,(WP_024293365.1-Lachnospiraceae:0.0472951130,WP_092242699.1-Lacrimispora:0.0324999941)96/100:0.0473042998)83.8/100:0.0206067040,((WP_124065139.1-Clostridium-sp.-E02:0.1209054481,WP_117415169.1-Clostridium-indicum:0.0566176358)78.6/95:0.0169092180,WP_143321893.1-Clostridium-sp.-HBUAS56010:0.0369885712)85.6/96:0.0248191204)14/44:0.0105467330,WP_138313639.1-Hungatella-hathewayi:0.0365989695)86.4/99:0.0332948023,((WP_227052204.1-Anaerostipes-caccae:0.0317928140,WP_137330070.1-Anaerostipes-rhamnosivorans:0.1207983435)98.9/100:0.1171701706,WP_006781439.1-Hungatella-hathewayi:0.1132451655)90.1/100:0.0567216876)13.3/70:0.0173152330,WP_186424215.1-Lacrimispora-celerecrescens:0.2123458964)83/57:0.0246140866,(WP_215718428.1-Anaerostipes-hadrus:0.0501919564,WP_075760385.1-Anaerostipes-sp.-494a:0.0348231607)99.4/100:0.0930971801)69.1/24:0.0054392323,((((WP_231062061.1-Lacrimispora-sp.-NSJ-141:0.1434769086,WP_249304640.1-Qiania-dongpingensis:0.0701433445)98/100:0.0882490987,WP_118541387.1-unclassified-Clostridium:0.0808101298)72.7/46:0.0118724063,WP_215700061.1-Clostridium-sp.-MCC353:0.0787467129)86.9/34:0.0271242521,(WP_024736308.1-Enterocloster-asparagiformis:0.0493809876,WP_066545911.1-Clostridium-sp.-AT4:0.0475333825)78.6/52:0.0156528685)86.3/23:0.0234095661)76.8/48:0.0133831715,WP_154471253.1-Clostridium-porci:0.1692427084)76.7/41:0.0117762405,WP_002577448.1-Enterocloster:0.0831406338)0/20:0.0000027966,(((WP_045093538.1-Clostridium-sp.-FS41:0.0622955082,WP_025485467.1-Lachnospiraceae:0.0792914082)44/99:0.0194389172,((WP_003497689.1-_Clostridium:0.0695337581,WP_125142063.1-Clostridium-transplantifaecale:0.0490386862)95.9/97:0.0640014683,((((WP_087223935.1-Lachnoclostridium-sp.-An14:0.0528877063,WP_101876396.1-Lachnoclostridium-edouardi:0.0666033631)90/98:0.0372135237,WP_118243934.1-Clostridium-sp.-AM58-1XD:0.0806165414)76.2/77:0.0108054418,WP_118742831.1-Clostridium-sp.-AM28-20LB:0.0395962237)85.8/76:0.0162790203,WP_008399284.1-unclassified-Clostridium:0.0245355994)75.8/77:0.0148253331)85.4/43:0.0408179349)79.4/22:0.0151370382,WP_178313416.1-Lacrimispora-sp.-210928-DFI.3.58:0.0236240120)49.1/21:0.0079124297)86/93:0.0475193573,WP_251497048.1-Otoolea-sp.-DSM-111138:0.2098544740)22.6/88:0.0594045726,WP_118555425.1-Lachnotalea-sp.-AF33-28:0.2674462839)79.5/99:0.0838987250)95.9/100:0.1182581788)83/98:0.0779073483,(((((((((((WP_035764434.1-Butyrivibrio-sp.-NC2002:0.1237446747,(((WP_026526812.1-Butyrivibrio-sp.-VCD2006:0.2024574896,WP_026491547.1-unclassified-Butyrivibrio:0.0210715911)66.9/100:0.0484254540,WP_044913869.1-Butyrivibrio-sp.-WCE2006:0.0292288117)93.2/100:0.0883842167,WP_091822189.1-Butyrivibrio-sp.-ob235:0.0905980348)62.8/100:0.0484370332)98.5/100:0.1551869547,(((((((((WP_022768176.1-unclassified-Butyrivibrio:0.0335666407,WP_022766545.1-Butyrivibrio-sp.-XPD2006:0.0678553703)99/100:0.1156322398,WP_034446055.1-Butyrivibrio-sp.-AE2032:0.1015271238)55.4/100:0.0242917738,WP_071175335.1-Butyrivibrio:0.0903962504)90.4/95:0.0646124768,WP_024867291.1-Butyrivibrio-sp.-FCS014:0.0346970384)77.6/95:0.0521235774,WP_026513668.1-Butyrivibrio-sp.-LB2008:0.0775009199)90.6/100:0.0418647125,WP_013279951.1-Butyrivibrio-proteoclasticus:0.0416564672)0/51:0.0000020186,WP_026652320.1-Butyrivibrio-proteoclasticus:0.0638348548)85.6/52:0.0189350721,WP_026663187.1-Butyrivibrio-proteoclasticus:0.1031041668)70.7/96:0.0111388916,WP_092060841.1-Butyrivibrio-sp.-INlla14:0.0734483053)94.8/100:0.0691590374)97.5/99:0.1283184580,((WP_019160740.1-Blautia:0.0809583685,WP_173738571.1-Blautia-schinkii:0.0286609697)87.8/100:0.0394669977,(WP_158421413.1-Blautia:0.0453873901,(WP_148445023.1-Eubacteriales:0.0353135678,WP_187566223.1-Blautia-faecis:0.0643429951)93.7/100:0.0407474961)46.8/83:0.0179016641)97.7/98:0.1133703109)95.9/94:0.0792404223,WP_132379639.1-Muricomes-intestini:0.1428126231)48.1/20:0.0120184162,(WP_109624131.1-Murimonas-intestini:0.0942943968,WP_178047055.1-Fusicatenibacter-sp.-CLA-AA-H277:0.1402820198)87.7/20:0.0404258854)68.3/19:0.0098325441,(WP_230400100.1-Novisyntrophococcus-fermenticellae:0.2751119189,((WP_173751851.1-Blautia-caecimuris:0.0609926870,WP_117997894.1-Blautia-obeum:0.0172037945)99.7/100:0.1037999863,(((WP_158367551.1-Lachnospiraceae:0.0457825405,WP_235247881.1-Faecalicatena-contorta:0.1021415203)74.2/94:0.0519674867,(WP_251390531.1-Mediterraneibacter-sp.-DSM-111629:0.0563903129,WP_073962323.1-_Ruminococcus:0.0343691475)88.1/100:0.0332730412)84/70:0.0234814440,WP_070089756.1-Merdimonas-faecis:0.0438931857)81.4/68:0.0176628999)72.1/66:0.0109664729)84.1/69:0.0277086034)97.2/98:0.0712070639,WP_087156947.1-Lachnoclostridium-sp.-An181:0.0741911346)70.6/76:0.0147557161,(WP_073994331.1-Parasporobacterium-paucivorans:0.1902288775,(WP_099466994.1-Konateibacter-massiliensis:0.0979718458,WP_151145296.1-Candidatus-Galacturonibacter-soehngenii:0.0659708762)96.3/100:0.1114099255)71/85:0.0866055680)57.8/80:0.0282422753,WP_115483253.1-Anaerosacchariphilus-polymeriproducens:0.1346049972)37.2/90:0.0451264947,WP_158351687.1-Diplocloster-modestus:0.3209239753)64/100:0.0670193785,WP_158346166.1-Lachnospiraceae:0.3342578946)97.5/93:0.1568878996)73/92:0.0237654620,WP_226914794.1-Intestinibacter-bartlettii:0.3970991433)89.7/66:0.0766456625)77.4/58:0.0340071722,((((WP_073273890.1-Anaerocolumna-jejuensis:0.1844565688,WP_033167869.1-Clostridium-sp.-KNHs205:0.2054373415)61.8/100:0.0751609724,(WP_010268185.1-Paenibacillus-senegalensis:0.2244934810,(WP_155699987.1-Paenibacillus-psychroresistens:0.0934389789,(WP_202751253.1-Clostridium-sp.-YIM-B02515:0.0802513212,WP_090040610.1-Clostridium-frigidicarnis:0.0330540191)32.9/88:0.0190554966)0/15:0.0384480581)60.3/92:0.0752864648)98.7/100:0.2102948702,((WP_249305773.1-Jutongia-hominis:0.2991412488,(WP_055944671.1-unclassified-Clostridium:0.0499285655,WP_216564272.1-Eubacterium-sp.-MSJ-13:0.0348116712)78.6/100:0.0440945374)26.9/91:0.0287769401,WP_022463459.1-Jutongia-huaianensis:0.0909123636)99.8/100:0.3049880047)76.5/100:0.0509112037,WP_058352784.1-Acetivibrio-ethanolgignens:0.3972660403)94/97:0.1192966448)95.8/99:0.1432647195,(WP_105204772.1-Neobittarella-massiliensis:0.1661460118,WP_022475313.1-Huintestinicola-butyrica:0.4220309765)97.9/100:0.1722553888)33.1/56:0.0230959767)91.1/72:0.1130491450,((WP_069974893.1-Geosporobacter-ferrireducens:0.5288601915,(WP_223038819.1-Crassaminicella-profunda:0.1080835398,(WP_148809648.1-Crassaminicella-thermophila:0.0693619518,WP_218283043.1-Crassaminicella-sp.-143-21:0.3946641309)84.4/99:0.0496907595)87.2/99:0.0798157077)94.7/100:0.1480381171,WP_250858284.1-Oceanirhabdus-seepicola:0.2430029086)94.6/100:0.1287341040)88.4/69:0.0774031588)87.7/85:0.0499449843)73.2/49:0.0239361069,(WP_119970050.1-Clostridium-fermenticellae:0.3320556603,((WP_213236459.1-Fusibacter-paucivorans:0.8035418927,((((((((((WP_160564586.1-Apibacter-sp.-B2912:0.1418724690,(WP_055424523.1-Apibacter-mensalis:0.1164957479,(WP_160568957.1-unclassified-Apibacter:0.0153766196,WP_166004639.1-Apibacter-sp.-B2966:0.0626753996)98.7/100:0.1160324287)79.1/100:0.0743561804)96.9/100:0.1347920306,(WP_105245553.1-Apibacter-adventoris:0.2479031642,WP_146261858.1-Apibacter-muscae:0.1531241672)83.1/100:0.0730376725)100/100:0.3292428205,(WP_084015419.1-Moheibacter-sediminis:0.4719988327,WP_182042771.1-Moheibacter-lacus:0.3640882558)96.3/100:0.1809540463)0/30:0.0208386580,(WP_068596492.1-Vaginella-massiliensis:0.1996932609,((WP_230477342.1-Faecalibacter-bovis:0.1390903179,WP_125349156.1-Empedobacter-falsenii:0.1281620258)85.1/93:0.0454897359,(WP_125113420.1-Algoriella-xinjiangensis:0.1071546494,WP_092908829.1-Algoriella-xinjiangensis:0.0445890184)93.3/100:0.0647581195)97.6/93:0.1617463887)97.6/100:0.2363635191)94.8/99:0.2197083342,(WP_014783622.1-Aequorivita-sublithincola:0.4962741163,(((WP_230143398.1-unclassified-Pedobacter:0.1352738900,WP_215166022.1-Pedobacter-sp.-ISL-64:0.1250765892)96.9/100:0.1078736589,(WP_199156598.1-Pedobacter-sp.-ASV2:0.1591851032,((WP_056872627.1-Pedobacter-sp.-Leaf194:0.1165811955,((WP_123203876.1-Pedobacter-jejuensis:0.0978887428,WP_188416210.1-Pedobacter-mendelii:0.0408169032)8.5/97:0.0191306581,WP_056087060.1-Pedobacter-sp.-Leaf176:0.0694416419)73.9/99:0.0188569011)97.2/99:0.0757710068,WP_109929147.1-Pedobacter-paludis:0.0537073642)64.1/76:0.0352920325)85/57:0.0613721450)66.4/57:0.0230167029,WP_133260761.1-Pedobacter-changchengzhani:0.1622924529)97.3/96:0.2754047293)98.7/100:0.3128603029)94.2/100:0.1726249047,(((WP_107809536.1-Elizabethkingia-sp.-YR214:0.1928272371,WP_219971314.1-Chryseobacterium-sp.-LJ756:0.1810041613)87.9/80:0.0781945769,WP_076395614.1-Chryseobacterium:0.0556503671)72.3/79:0.0848806079,WP_088467981.1-Chryseobacterium-sp.-VAUSW3:0.1550481781)99.5/100:0.2939150785)0/22:0.0040101114,WP_111126400.1-Mesonia-sp.-K7:1.0540610441)98.2/96:0.1942574841,((((WP_205722928.1-Fulvivirga-lutea:0.6293795695,(((((((((((((WP_053470876.1-unclassified-Flavobacterium:0.2072746707,WP_056200692.1-Flavobacterium-sp.-Root186:0.0466432867)0/50:0.0000022364,(WP_115889823.1-Flavobacterium-cutihirudinis:0.0558316237,WP_095952323.1-Flavobacterium-sp.-ACN6:0.0470292392)32/40:0.0222169203)23.5/100:0.0102842178,WP_198857739.1-Flavobacterium-sp.-IB48:0.0586019423)65.9/97:0.0078860122,(WP_269237486.1-Flavobacterium-sp.-IMCC34775:0.1019997331,WP_163393939.1-Flavobacterium-limi:0.0414909848)71.8/100:0.0108636096)74.5/86:0.0086911061,WP_264205577.1-Flavobacterium-sp.-LS1R49:0.0306968376)88.7/98:0.0189677838,((WP_173968237.1-Flavobacterium-collinsii:0.0945275918,WP_264525312.1-Flavobacterium-sp.-N502536:0.0549810432)90.7/100:0.0418636291,WP_070907856.1-Flavobacterium-tructae:0.1110254588)75.8/63:0.0100721637)66.8/62:0.0158389553,WP_091091738.1-Flavobacterium-gillisiae:0.1275368687)98.1/100:0.1049155735,WP_248907802.1-Flavobacterium-sp.-K5-23:0.1463970403)95.4/99:0.0674631097,WP_103806521.1-Flavobacterium-alvei:0.1094112817)74.1/28:0.0496257181,(WP_144071503.1-Flavobacterium-franklandianum:0.0875935759,WP_133277350.1-Flavobacterium-nackdongense:0.1081238092)87.8/97:0.0492211124)46.7/27:0.0094985691,(WP_116761981.1-Flavobacterium-laiguense:0.0576464516,WP_143377960.1-Flavobacterium-sp.-ZT3R18:0.0543798829)98.3/100:0.0822132990)83.6/29:0.0321129697,WP_194311471.1-Flavobacterium-soyangense:0.1426134558)98/99:0.2248030675,(WP_140000461.1-Flavobacterium-microcysteis:0.3056670297,(((WP_194640005.1-Flavobacterium-sp.-HJJ:0.1271426552,WP_073246022.1-Flavobacterium-flevense:0.0849210932)77.3/100:0.0268930121,WP_100843428.1-Flavobacterium-sp.-5:0.1396339683)91.3/94:0.0772688771,WP_140487657.1-Flavobacterium-sp.-GSA192:0.0539292310)99.4/100:0.2448422941)95.9/100:0.1668398137)92.5/98:0.1867375739)73.8/95:0.1663165704,(WP_116872776.1-Terrimonas-sp.:0.5979113365,(WP_004155558.1-Microscilla-marina:0.4982558849,WP_101358295.1-Raineya-orbicola:0.3082791037)98.6/100:0.4206146219)57.4/94:0.1097047778)60.7/29:0.0787295498,WP_057952567.1-Salinivirga-cyanobacteriivorans:0.8193035202)86.7/95:0.1295219239,((((((WP_233378072.1-Maribellus-sp.-CM-23:0.0867123106,WP_247978154.1-Maribellus-sp.-YY47:0.0861586643)95.4/100:0.1530713856,(WP_167609329.1-Maribellus-sediminis:0.2250862727,(WP_045033348.1-Draconibacterium-sediminis:0.0332701729,WP_163345719.1-Draconibacterium-halophilum:0.1057954523)98.7/100:0.1944745093)93.4/100:0.1469573049)99.9/100:0.4185962796,(WP_106543043.1-Prolixibacter-denitrificans:0.1492999552,(WP_025865917.1-Prolixibacter-bellariivorans:0.0281738350,WP_153632016.1-Prolixibacter-sp.-SD074:0.0741624530)0/97:0.0000029132)99.4/100:0.3088170536)46.8/100:0.0940955941,((WP_159517929.1-Sunxiuqinia-indica:0.1397056146,WP_053182762.1-Sunxiuqinia-dokdonensis:0.1398775849)78.1/100:0.0674875853,WP_133466511.1-Sunxiuqinia-elliptica:0.2153860871)99/100:0.2293795746)96.2/99:0.1522867205,(WP_068701711.1-Paludibacter-jiangxiensis:0.5571020882,(((WP_101263396.1-Labilibaculum-filiforme:0.0524617179,WP_245870435.1-Labilibaculum-manganireducens:0.0591333757)96.8/99:0.1118161635,(WP_152734748.1-Marinifilum-sp.-N1E240:0.1017112944,(WP_171595603.1-Marinifilum-caeruleilacunae:0.0836183510,(WP_268189532.1-Marinifilum-sp.-D737:0.0674624411,WP_054720444.1-Marinifilum-fragile:0.0427557731)88.6/100:0.0274229305)84/100:0.0307607071)93.7/100:0.0764986976)3.8/89:0.0457344117,WP_127342456.1-Ancylomarina-longa:0.1698711706)99.4/100:0.2851688541)77.1/52:0.0368088424)81.8/52:0.0652833990,(((((WP_011500232.1-Methanococcoides-burtonii:0.3037033778,WP_135611779.1-Methanococcoides-sp.-AM1:0.2546393597)96.2/97:0.1661106104,(WP_048137589.1-Methanosarcina-horonobensis:0.1972703245,((WP_048184631.1-Methanosarcina-siciliae:0.0763839095,WP_048173663.1-Methanosarcina-siciliae:0.0742927973)31.2/99:0.0156224048,WP_011023680.1-Methanosarcina-acetivorans:0.0452193626)96/99:0.1106016651)81.3/99:0.0772808539)90.2/27:0.0855166568,WP_048177163.1-Methanosarcina-sp.-MTP4:0.1211776778)45.2/27:0.0431593439,((WP_128503182.1-Methanosarcina-sp.-MSH10X1:0.2242911579,WP_011033115.1-Methanosarcina-mazei:0.0834433747)49.3/82:0.0430863698,WP_048134921.1-Methanosarcina-sp.-1.H.T.1A.1:0.0469760983)93.9/99:0.0834989110)83.7/27:0.0384847820,WP_048167026.1-Methanosarcina:0.0871734740)99.7/99:0.3388104457)88.7/97:0.1129654084)48/92:0.0881937867)93/99:0.1867578371,(((WP_085659180.1-Prosthecochloris-sp.-HL-130-GSB:0.6349479380,WP_038563105.1-Draconibacterium-orientale:0.3807954327)94.4/100:0.2415703920,(WP_040596643.1-Prevotella-stercorea:0.1306069261,WP_207153370.1-Prevotella-herbatica:0.1133671576)98.9/100:0.3700768017)70.7/96:0.0987238460,(WP_223927321.1-Prevotella-lacticifex:0.2826411623,WP_009010825.1-Prevotella-sp.-C561:0.2295075755)99.9/100:0.5790066874)79.6/99:0.2317541183)76.2/100:0.1786879293,((((((WP_013633636.1-Pseudopedobacter-saltans:0.1782939212,WP_154287284.1-Pedobacter-puniceum:0.2105849878)91.8/98:0.1134928942,WP_160842504.1-Pedobacter-sp.-HMF7647:0.0748950354)99.4/98:0.2906825217,(WP_258137696.1-Mucilaginibacter-sp.-BJC16-A38:0.2583362451,(WP_259068220.1-Mucilaginibacter-sp.-X4EP1:0.0816049774,WP_144912900.1-Mucilaginibacter-frigoritolerans:0.1016875520)90.8/100:0.0974131846)95.9/100:0.1621044680)26.1/97:0.0166070957,WP_074492796.1-Mucilaginibacter-polytrichastri:0.2563370328)98.3/100:0.3468452320,WP_163286637.1-Cryomorpha-ignava:0.7826700971)60/100:0.1796560081,WP_130092062.1-Brumimicrobium-glaciale:0.6355929924)93/100:0.1375632441)88.3/100:0.1745339586)60.1/57:0.0734059037,(WP_138788932.1-Paenibacillus-thermoaerophilus:0.5435873137,WP_078711972.1-Anaerorhabdus-furcosa:0.7900540575)81.3/57:0.1833208905)96.7/84:0.2165767229)83/24:0.0464020180)89.3/22:0.0564122667,(WP_038266426.1-Peptoclostridium-litorale:0.6163704749,WP_160686059.1-Clostridium-sp.-C2-6-12:0.3531869265)65.6/98:0.0318046985)89.2/26:0.0607985261)94.3/97:0.1459707114,((((WP_058965344.1-Fournierella-massiliensis:0.2440863892,WP_185752312.1-Butyrivibrio-sp.-AE2032:0.4899860083)86/97:0.0885671439,WP_155202396.1-Ruthenibacterium-lactatiformans:0.2759638930)95/98:0.1362478050,(WP_186886939.1-Anaerofilum-hominis:0.2050580468,WP_102049807.1-Pygmaiobacter-massiliensis:0.3585550934)88.8/99:0.1049453505)97.1/100:0.2221956125,((WP_072827923.1-Fibrobacter-sp.-UWB12:0.0226323156,(WP_088630837.1-unclassified-Fibrobacter:0.0447938694,WP_073304542.1-Fibrobacter:0.2810766614)29.9/93:0.0189406411)93.2/100:0.0981963232,((WP_097034884.1-Fibrobacter-sp.-UWT3:0.0361767828,(WP_073056819.1-unclassified-Fibrobacter:0.0212996288,WP_073155408.1-Fibrobacter:0.0760201767)41.3/56:0.0321228339)63/62:0.0198483194,WP_073188830.1-Fibrobacter-sp.-UWP2:0.0384935759)96.5/100:0.1310983555)98.8/100:0.2583230789)89.4/66:0.1194488195)92.3/100:0.1633109005,WP_236098952.1-Dethiosulfovibrio:0.7894980114)95.3/99:0.2247003385,((((((((((((((((((((WP_176021856.1-Stenotrophomonas-sp.-NA06056:0.0342328613,WP_049440708.1-Stenotrophomonas-maltophilia:0.0853164077)99.6/100:0.2698482937,WP_132999175.1-Luteimonas-arsenica:0.2462515991)99/100:0.2206371565,((WP_168608661.1-Oleiagrimonas-citrea:0.0820903460,WP_113064358.1-Oleiagrimonas-sp.-MCCC-1A03011:0.0258506436)74.6/100:0.0202382724,WP_043099571.1-Oleiagrimonas-soli:0.1359721496)97.4/100:0.1279263573)94.2/98:0.0917035833,(WP_063674027.1-Dyella-thiooxydans:0.1777120341,(WP_188800319.1-Dyella-caseinilytica:0.2297608730,WP_192554051.1-Dyella-acidiphila:0.1486950403)93.1/100:0.0944549124)38.1/89:0.0455444968)84/90:0.0284852681,WP_201315287.1-Dyella-sp.-EPa41:0.1761110687)76.2/34:0.0170868536,(WP_209622196.1-Frateuria-flava:0.1006971350,WP_091339503.1-Frateuria-terrea:0.0585252115)96.6/100:0.0703185468)75.9/31:0.0134065198,(WP_253565187.1-Dyella-sp.-Sa:0.0874788028,WP_131151171.1-Dyella:0.2679061950)83.9/37:0.0352825014)97.7/92:0.0728889734,((WP_019467068.1-Dyella-japonica:0.3326117513,WP_204636097.1-Dyella-kyungheensis:0.1503277278)91.5/92:0.0604002979,((WP_049621948.1-Frateuria-defendens:0.1835133980,WP_179481164.1-unclassified-Rhodanobacter:0.1141728052)79.6/95:0.0512354884,((((((WP_214557828.1-unclassified-Rhodanobacter:0.0458887182,WP_068095172.1-unclassified-Rhodanobacter:0.0378721374)55.9/100:0.0260279975,(WP_008437080.1-Rhodanobacter-thiooxydans:0.0567106440,(WP_063107523.1-Rhodanobacter-thiooxydans:0.0416319051,WP_211005527.1-Rhodanobacter-sp.-B2A1Ga4:0.0465858076)72.7/100:0.0086802555)49.5/100:0.0227177011)95.7/100:0.0676191551,WP_072762453.1-Rhodanobacter-sp.-OK091:0.0139756576)77.3/99:0.0165535674,((WP_077439480.1-Rhodanobacter-sp.-C01:0.1407923123,WP_158752649.1-Dyella-sp.-S184:0.0311926965)32.3/97:0.0137453843,WP_077445946.1-Rhodanobacter-sp.-C05:0.0683656776)76.3/97:0.0260577652)73.4/97:0.0276426769,WP_184416890.1-Rhodanobacter-sp.-MP7CTX1:0.0577460796)87.8/99:0.0414804285,WP_015446789.1-Rhodanobacter-denitrificans:0.0718894553)71.3/100:0.0321629499)83.6/88:0.0379348434)77.4/61:0.0195012237)38.4/62:0.0065937374,((WP_233843762.1-Dyella-sp.-2HG41-7:0.0404310804,WP_188793403.1-Dyella-nitratireducens:0.0998993946)74.9/96:0.0622839268,WP_126673574.1-Dyella-dinghuensis:0.0571758058)99.7/100:0.1386277218)90.7/93:0.0572363658,(WP_111981026.1-Dyella-jiangningensis:0.2123327728,WP_266183414.1-Dyella-sp.-RBB1W86:0.0968818456)82.9/94:0.0320282495)85.8/96:0.0354327266,((WP_130619635.1-Dyella-amyloliquefaciens:0.1025453446,WP_199044928.1-Dyella-sp.-ASV24:0.0871124059)73.6/72:0.0147451078,(WP_266169930.1-Dyella-sp.-RXD159:0.1035110979,(WP_114846719.1-Dyella-tabacisoli:0.0595470023,WP_090451016.1-Dyella-sp.-OK004:0.0904590465)82.8/87:0.0338108514)79.3/82:0.0149915596)76.9/87:0.0107146793)89/98:0.0670249513,WP_214570707.1-unclassified-Dyella:0.1379010867)97.9/100:0.1725845206,WP_078758981.1-Lysobacter-spongiicola:0.3916334643)99.1/100:0.2975853194,((((((((WP_200343816.1-Halochromatium-glycolicum:0.2240737106,WP_201222000.1-Halochromatium-roseum:0.1587219722)97.7/100:0.2052661485,(WP_200237937.1-Thiohalocapsa-halophila:0.0690791011,WP_058558062.1-Thiohalocapsa-sp.-ML1:0.3360554957)96.1/100:0.1399487366)96.4/100:0.1906209708,(WP_182584206.1-Thiospirillum-jenense:0.5487378255,(WP_200286492.1-Rhabdochromatium-marinum:0.1720844969,WP_009150811.1-Thiorhodovibrio-sp.-970:0.0956422872)94.3/100:0.1415215814)63/100:0.1716548557)93.2/100:0.1181521797,((((((WP_200207675.1-Chromatium-weissei:0.2788460271,WP_200154756.1-Chromatium-okenii:0.1220810406)98.8/100:0.1920962380,(WP_200158060.1-Allochromatium-vinosum:0.0788815207,WP_200376120.1-Thiocystis-violacea:0.1506475968)82.3/100:0.0556164882)69.9/84:0.0243254152,(((WP_043749566.1-Imhoffiella-purpurea:0.1016360962,WP_007039504.1-Thiorhodococcus-drewsii:0.0814601295)83.4/100:0.0412386020,((WP_164456073.1-Thiorhodococcus-minor:0.1229136012,WP_200330387.1-Thiocystis-violacea:0.1110906026)83.1/100:0.0274552258,WP_164652045.1-Thiorhodococcus-mannitoliphagus:0.0609299669)95.7/100:0.0815773818)89.8/100:0.0311258462,(WP_201092787.1-Thiocystis-minor:0.0989473250,WP_132978621.1-Thiobaca-trueperi:0.1358553055)81.6/100:0.0257737793)85.5/100:0.0297978034)90.7/93:0.0480499674,(WP_200388819.1-Thiocapsa-imhoffii:0.2153308078,((WP_007191850.1-Thiocapsa-marina:0.0538714114,WP_120796078.1-Thiocapsa-rosea:0.0459753558)64.9/100:0.0081286289,WP_232409912.1-Thiocapsa-bogorovii:0.0280114309)96.7/100:0.1219573577)95.2/100:0.1124057520)96.7/100:0.1191471765,(WP_100918325.1-Candidatus-Thiodictyon-syntrophicum:0.1085371768,WP_020502667.1-Lamprocystis-purpurea:0.1438874915)95.6/100:0.1225137403)61.5/86:0.0739470160,WP_089729519.1-Candidatus-Thiosymbion-oneisti:0.4258904294)89/98:0.0887795195)51.4/71:0.0280887493,WP_018175268.1-unclassified-Thioalkalivibrio:0.5686398337)84.8/70:0.0928516170,WP_101758446.1-Oceanicoccus-sp.-KOV_DT_Chl:0.6932226605)12.3/39:0.0645811435,WP_141730858.1-Oligoflexus-tunisiensis:0.6670697835)6.6/92:0.0389233305,WP_141198172.1-Persicimonas-caeni:0.7703704010)81.7/20:0.0616619695)84.7/20:0.0723533468,((WP_256181981.1-Cloacibacillus-evryensis:0.2835645160,WP_087368268.1-Cloacibacillus-sp.-An23:0.1508515531)100/100:0.8463149526,((WP_084578244.1-Sporomusa-malonica:0.5900268701,WP_131919244.1-Heliophilum-fasciatum:0.3541128897)73.1/79:0.0497481370,((WP_090940141.1-Pelosinus-propionicus:0.3478516631,WP_240986382.1-Acididesulfobacillus-acetoxydans:0.3616322558)85/99:0.0804467371,WP_092486361.1-Desulfoscipio-geothermicus:0.2010850752)85.1/79:0.0537221453)82.4/94:0.0573703985)59.6/21:0.0282211113)90.2/48:0.0989083976,((WP_246588317.1-Pelorhabdus-rhamnosifermentans:0.2475821573,((((WP_092074275.1-Dendrosporobacter-quercicolus:0.3336438639,WP_188400350.1-Sporomusa-sp.-GT1:0.3804407775)52.2/33:0.0487754363,(WP_054258683.1-Propionispora-sp.-2/2-37:0.1887553386,(WP_149735423.1-Propionispora-hippei:0.1136787369,WP_091744506.1-Propionispora-vibrioides:0.1726903950)51/97:0.0489663088)88.6/75:0.0512190177)87.1/33:0.0610720839,(((WP_007952170.1-Pelosinus-fermentans:0.1505023665,WP_007939097.1-Pelosinus:0.1352277642)86.8/100:0.0833723074,(WP_132081660.1-Anaerospora-hongkongensis:0.1155124799,WP_132083235.1-Anaerospora-hongkongensis:0.2594358078)97/100:0.1482998140)4.3/21:0.0527794230,(WP_188399481.1-Sporomusa-sp.-GT1:0.3507388688,((WP_094603851.1-Sporomusa-silvacetica:0.1606242125,WP_144352952.1-Sporomusa-termitida:0.1793444649)75.5/100:0.0659015217,WP_054258662.1-Propionispora-sp.-2/2-37:0.3251151040)85.9/93:0.0513562550)23.3/20:0.0623856878)59.8/64:0.0311020349)53.1/33:0.0370345352,(WP_145090815.1-Sporomusa-sp.-KB1:0.0559225113,WP_093793074.1-Sporomusa-acidovorans:0.0920750852)99/100:0.1516776282)85.3/99:0.0633669110)80/99:0.0556873194,WP_110954047.1-Massilibacillus-massiliensis:0.3633180838)90.9/100:0.1063138744)93.7/99:0.1625163118,WP_207681674.1-Desulfonema-magnum:0.7975844963)28.9/35:0.0887488776,WP_008811140.1-Parasutterella-excrementihominis:0.7545558776)92.1/99:0.1235305100,(WP_133132051.1-Legionella-sp.-W10-070:0.4356635182,((((((((((((WP_027223798.1-Legionella-pneumophila:0.0385306943,WP_265717942.1-Legionella-sp.-PATHC032:0.0460401028)93.6/87:0.0371016209,WP_106142999.1-Legionella-pneumophila:0.0448552806)69.7/85:0.0059048411,(WP_061513621.1-Legionella:0.0173626527,WP_027226296.1-Legionella-pneumophila:0.0686256094)99.7/98:0.0814245695)86.9/100:0.0167045014,WP_044499121.1-Legionella-pneumophila:0.0316273332)86.1/100:0.0192349514,WP_027220106.1-Legionella-pneumophila:0.0200755894)77.3/100:0.0229260156,WP_027266778.1-Legionella-pneumophila:0.0422901601)94.2/100:0.0913431836,(WP_058494182.1-Legionella-worsleiensis:0.1694490724,(WP_127106294.1-Legionella-sp.-km535:0.0496431422,WP_058474153.1-Legionella-quateirensis:0.0372009550)90.3/100:0.0515581271)96.2/99:0.0897363477)95.3/99:0.0903224008,WP_031567420.1-Legionella-wadsworthii:0.2446048988)56.9/95:0.0275079852,WP_045097199.1-Legionella-fallonii:0.1582036069)85.3/97:0.0316094720,(WP_232890831.1-unclassified-Legionella:0.0538778344,(WP_058459148.1-Legionella-bozemanae:0.0309044287,WP_058518697.1-Legionella-parisiensis:0.0599795271)83.1/100:0.0329371542)99.2/100:0.1163815621)80.4/91:0.0303361955,WP_058513550.1-Legionella-santicrucis:0.1398661756)0/68:0.0000025892,(WP_058463758.1-Legionella-cincinnatiensis:0.0265384391,WP_128130294.1-Legionella-sainthelensi:0.0677037978)72.7/100:0.0451809774)97.9/98:0.3517359743)99.7/99:0.3566577631)79.1/61:0.0687108740,((((((((((((WP_171588903.1-Pelistega-europaea:0.4853215724,(WP_040666911.1-Neisseria-wadsworthii:0.0917483708,WP_135034416.1-Neisseria:0.2557511192)75/96:0.0322605055)31.2/54:0.0460669690,(((WP_214236207.1-Rosenbergiella-sp.-S61:0.0881328309,WP_241647140.1-Rosenbergiella-sp.-JB07:0.1632917135)71.1/75:0.0440760044,WP_214195533.1-Rosenbergiella-collisarenosi:0.1254253002)90.5/100:0.0934732933,WP_101506252.1-Erwinia-sp.-B116:0.1933813463)89.3/71:0.0578443661)78.2/24:0.0255106926,WP_140470594.1-Ewingella-americana:0.3596547179)73.6/16:0.0094539200,(((WP_032441370.1-Klebsiella-pneumoniae:0.1494543884,(WP_015583911.1-Raoultella-ornithinolytica:0.0722412121,WP_265776798.1-Klebsiella-oxytoca:0.1894216249)78.7/100:0.0616468834)91.9/100:0.0788660052,(WP_029991521.1-Tatumella-ptyseos:0.0964039201,(WP_212707699.1-Tatumella-sp.-JGM130:0.0567815964,WP_087487466.1-Tatumella-citrea:0.0739978570)63.1/80:0.0178213741)76.7/96:0.0276037561)93.2/99:0.1016833096,WP_154751639.1-unclassified-Erwinia:0.3482929614)66.5/15:0.0123717990)74.3/22:0.0141166680,(WP_024553491.1-Franconibacter-helveticus:0.4263947516,WP_087653467.1-Klebsiella/Raoultella-group:0.2356884121)84.2/70:0.0494566511)95.3/94:0.0615842945,((((WP_228398973.1-Insectihabitans-xujianqingii:0.1196368082,WP_074823218.1-Pragia-fontium:0.1132974213)97.1/100:0.0919159524,WP_130971788.1-Hafnia-alvei:0.1622985477)85.1/100:0.0380763809,WP_165428828.1-Rouxiella-badensis:0.1369606615)73.4/100:0.0089841874,((WP_112167140.1-Klebsiella:0.1628359036,WP_200519240.1-Klebsiella-michiganensis:0.1248575024)90.1/65:0.0702401543,WP_137713144.1-Brenneria-rubrifaciens:0.1426730259)73.9/44:0.0202093888)70.4/43:0.0178628080)71.8/21:0.0104131633,(((WP_131063651.1-Raoultella-planticola:0.0275630135,WP_163980548.1-Raoultella-ornithinolytica:0.0488091557)98.9/100:0.0966640616,(WP_148250181.1-Raoultella-terrigena:0.1085434975,WP_032686784.1-Raoultella-planticola:0.0482230138)95.5/97:0.0651128231)14.5/63:0.0139822391,(WP_142430937.1-Klebsiella-oxytoca:0.1594289791,((WP_239614184.1-Raoultella-ornithinolytica:0.1173889030,WP_186921353.1-Klebsiella-quasipneumoniae:0.2181549847)69.7/90:0.0162531113,((WP_032699180.1-Raoultella:0.0669808162,WP_228399925.1-Raoultella:0.0399322475)77.8/98:0.0207413219,WP_004138711.1-Klebsiella:0.0204093910)94.5/100:0.0527619758)82.7/80:0.0307414395)86.7/75:0.0335856924)76.6/47:0.0136541372)82.2/24:0.0129423447,((((((WP_136167279.1-Brenneria-sp.-CFCC-11842:0.0543634693,WP_264089117.1-unclassified-Brenneria:0.1287915892)67.6/100:0.0481594475,WP_109053612.1-Brenneria-roseae:0.1200026191)82.6/100:0.0319325482,((((((WP_180742451.1-Pectobacterium-sp.-CFBP8739:0.2974001056,((((((((((WP_047710158.1-Pectobacterium-atrosepticum:0.0716016394,WP_198482318.1-Pectobacterium-parmentieri:0.0457348092)20/97:0.0111182385,WP_033071760.1-Pectobacterium-parmentieri:0.0366571779)92.3/100:0.0493606031,WP_226305805.1-Pectobacterium-versatile:0.0295150218)84/99:0.0263440712,WP_039517132.1-Pectobacterium-brasiliense:0.0516058597)74.5/91:0.0091646816,WP_039346077.1-Pectobacterium-fontis:0.0398865650)46.6/87:0.0084875258,WP_039280054.1-Pectobacterium-brasiliense:0.0612357518)92.7/87:0.0354994778,WP_103972204.1-Pectobacterium:0.0199303826)42.9/72:0.0088438118,WP_205558955.1-Pectobacterium-brasiliense:0.0715729225)74/80:0.0092552417,WP_116186106.1-Pectobacterium-aquaticum:0.0540375761)72.5/58:0.0076280953,WP_151242707.1-Pectobacterium-carotovorum:0.0451713483)91.3/86:0.0346303045)0/35:0.0000023723,(((WP_180777316.1-Pectobacterium-aroidearum:0.0725475737,WP_107169883.1-Pectobacterium:0.0544143785)85.4/85:0.0207826826,WP_039504461.1-Pectobacterium:0.0278568776)47.4/67:0.0172365315,((WP_233968669.1-Pectobacterium-polaris:0.0683386849,WP_048259384.1-Pectobacterium-peruviense:0.0164839440)93.9/81:0.0363237863,WP_138255615.1-Pectobacterium:0.0352134084)0/44:0.0000021644)75.6/52:0.0175766970)73.5/56:0.0104331719,(WP_205946867.1-Pectobacterium:0.0728105102,WP_233976643.1-Pectobacterium-versatile:0.0270879865)75.9/73:0.0141992910)63/56:0.0166004590,(WP_109411993.1-Pectobacterium-polaris:0.0722621315,((WP_249680869.1-Pectobacterium-polaris:0.0499983511,WP_125232508.1-Pectobacterium-versatile:0.0651317692)75.8/99:0.0120269167,WP_005974084.1-Pectobacterium-wasabiae:0.0436027621)62.7/100:0.0195316414)0/75:0.0000023599)83.3/73:0.0175387124,WP_119888241.1-Pectobacterium:0.0708136127)80.5/87:0.0362756065,(WP_132452860.1-Samsonia-erythrinae:0.1450061072,WP_130637964.1-Pectobacterium-versatile:0.0707742318)42.2/93:0.0228163329)88/96:0.0441853358)97.2/100:0.0862784178,((((((WP_168640580.1-Dickeya-sp.-CFBP-2040:0.1408670749,(WP_027712673.1-Dickeya-chrysanthemi:0.0543478632,WP_012770533.1-Dickeya:0.0436767992)73.1/96:0.0088777984)76.2/86:0.0086611918,(WP_012883965.1-Dickeya-parazeae:0.0871158514,WP_236306746.1-Dickeya-zeae:0.0431236992)85.1/94:0.0177024296)75.6/92:0.0077455857,((WP_038918362.1-Dickeya:0.0389325371,WP_123251184.1-Dickeya-undicola:0.0582676485)10.6/79:0.0075128473,WP_022632763.1-Dickeya-solani:0.0337163304)90.4/100:0.0280170351)97.7/100:0.1183382489,(WP_148877341.1-Serratia-marcescens:0.0826816158,WP_193158894.1-Serratia-marcescens:0.0659484413)95.1/100:0.0937034698)80.5/100:0.0390223056,WP_062791489.1-Serratia-plymuthica:0.0939209153)78.3/84:0.0272095059,(WP_121576612.1-Brenneria-alni:0.1204704865,((WP_214648782.1-Pectobacterium-carotovorum:0.0451212440,((WP_261846379.1-Pectobacterium-carotovorum:0.0749851311,WP_121590317.1-Brenneria-goodwinii:0.0084625898)94.5/100:0.0366569858,(WP_182099415.1-Pectobacterium-aroidearum:0.0446994079,WP_039482612.1-Pectobacterium:0.0359158198)72.2/97:0.0091248123)17.1/96:0.0088057127)64.7/99:0.0047481336,WP_249228983.1-Brenneria-tiliae:0.0609025321)92.1/100:0.0426380839)82.6/73:0.0285279740)76.2/66:0.0186060713)56.2/78:0.0114861674,(WP_096000273.1-Klebsiella-pneumoniae:0.2210439574,WP_095845203.1-Gibbsiella-quercinecans:0.0590917478)39.3/23:0.0342558161)82.5/72:0.0362173939,WP_248464278.1-Pectobacterium-versatile:0.2235648887)88.1/70:0.0384759315)0/14:0.0000020271,(WP_193937755.1-Klebsiella-grimontii:0.2411750602,(WP_142514871.1-Klebsiella-huaxiensis:0.1917907352,(WP_024360036.1-Klebsiella:0.0650363257,WP_234284225.1-Klebsiella-oxytoca:0.1011219984)62.1/98:0.0475669360)85.2/76:0.0481429402)29.8/68:0.0257163347)89.1/90:0.0471243318,((WP_253460089.1-Erwinia-persicina:0.3811332708,(WP_246757812.1-Erwinia-phyllosphaerae:0.2382798369,WP_232759612.1-unclassified-Pantoea:0.1900990274)67.4/99:0.0683464270)81.2/27:0.0522243568,(((WP_146378871.1-Cronobacter-turicensis:0.0941432273,WP_032180402.1-Escherichia-coli:0.0999678313)100/100:0.2488115749,((WP_126816534.1-Enterobacter-mori:0.1438706487,WP_256879438.1-Escherichia-albertii:0.1260450322)27.4/85:0.0627946788,WP_049841125.1-Klebsiella-sp.-RIT-PI-d:0.4589355846)23.3/90:0.0560583784)81.6/81:0.0408021028,((((WP_168836617.1-Vibrio-agarilyticus:0.1459915406,(WP_076537303.1-Shewanella-sp.-UCD-KL21:0.0282110040,WP_102531614.1-Shewanella-sp.-10N.286.52.B9:0.0641686403)100/100:0.7149154749)91/100:0.1205527176,(WP_137671336.1-Agarivorans-sp.-Toyoura001:0.0552542735,WP_221077378.1-Agarivorans-albus:0.0724019846)90.6/100:0.1129567612)95.9/100:0.1168767440,((((((((WP_127021653.1-Rheinheimera-mangrovi:0.5084315788,((WP_138320019.1-Thalassotalea-litorea:0.1936624622,WP_136736185.1-Thalassotalea-mangrovi:0.1624879525)99.9/100:0.3687268554,WP_114327629.1-Candidatus-Colwellia-aromaticivorans:0.1477130726)63.7/100:0.0218047247)80.3/98:0.0414344423,WP_101108574.1-Psychromonas-sp.-psych-6C06:0.2927857502)39.6/98:0.0499203314,((WP_016794198.1-Vibrio-cyclitrophicus:0.0560302532,WP_102424249.1-Vibrio-cyclitrophicus:0.0539073880)83.8/100:0.0370625551,(WP_261903470.1-Vibrio-fortis:0.1062295719,WP_240207223.1-Vibrio-sp.-CyArs1:0.0538112920)91.4/99:0.0518872570)99.3/100:0.1882920269)52.1/91:0.0773427730,(WP_199145406.1-Citrobacter-sp.-ASV29:0.4146525290,WP_148513713.1-Vibrio:0.2217830664)92.7/68:0.1346004153)35.8/44:0.0233352837,((WP_150896168.1-Vibrio-fortis:0.2664487052,WP_067584343.1-Endozoicomonas-ascidiicola:0.3448468741)66.6/66:0.0505329962,(((WP_260491935.1-Pseudoalteromonas-flavipulchra:0.2820911140,WP_039494249.1-Pseudoalteromonas-flavipulchra:0.1016684302)17.7/60:0.0323389876,WP_055022295.1-Pseudoalteromonas-sp.-P1-26:0.0880453623)80.8/98:0.0522844007,(WP_149602089.1-Pseudoalteromonas-distincta:0.0506276910,WP_248518672.1-Pseudoalteromonas-sp.-2CM32C:0.0323591280)97/100:0.1505189899)98.9/100:0.2824370946)87/47:0.0820025767)86.4/98:0.0742843952,(WP_132584255.1-Rheinheimera-sp.-D18:0.4530462983,(((((WP_131912015.1-Celerinatantimonas-diazotrophica:0.2959438582,((WP_092385961.1-Halopseudomonas-salegens:0.2635388794,WP_036859756.1-Porticoccus-hydrocarbonoclasticus:0.3382606401)84.7/94:0.1212146706,((WP_076722556.1-Marinobacter-lutaoensis:0.2896426439,((WP_106695101.1-Marinobacter-shengliensis:0.0837148871,WP_148609564.1-Marinobacter-sp.-BW6:0.1446754202)49.2/40:0.0074133001,(WP_222473125.1-Marinobacter-daepoensis:0.0601611710,WP_092030923.1-Marinobacter-sp.-DSM-26671:0.1660887434)33.9/38:0.0376725774)95.8/97:0.1173803784)37.9/94:0.0432971494,WP_036130839.1-Marinobacter-nitratireducens:0.1784688351)55.9/94:0.0680884902)87.5/100:0.0606248391)81.8/98:0.0550655911,(WP_027875186.1-Spongiibacter-marinus:0.1972813630,WP_204995064.1-Spongiibacter-marinus:0.1235501934)87.3/100:0.0560293776)87/98:0.0443240419,WP_168451636.1-Spongiibacter-thalassae:0.0568761630)21.9/97:0.0504467078,WP_062382560.1-Zhongshania-aliphaticivorans:0.0815728485)64.8/99:0.0932655281,((WP_184464382.1-Zhongshania-antarctica:0.0586287988,WP_198570926.1-Spongiibacter-nanhainus:0.1050225488)59.9/26:0.0089038064,WP_103683243.1-Marortus-luteolus:0.0388751809)52.5/25:0.0496027747)82.6/83:0.0696167800)96.3/96:0.1559317095)91.9/94:0.0803097896,((WP_145010177.1-Pseudomonas-psychrotolerans:0.0223903313,WP_077172422.1-Pseudomonas-psychrotolerans:0.0541988343)99.8/100:0.3075811555,WP_084334293.1-Pseudomonas-indica:0.1439882682)84.8/98:0.0882552371)84.2/92:0.0465464859,(WP_261844981.1-Amphritea-ceti:0.4454608452,((((((WP_269124033.1-Marinomonas-sp.-15G1-11:0.0311899687,WP_191602238.1-Marinomonas-algicola:0.1116009756)99.5/100:0.1887705777,WP_048693477.1-Catenovulum-maritimum:0.2665033487)88.8/100:0.0872926367,WP_228191388.1-Marinomonas-communis:0.1977736540)37.6/98:0.0300834215,WP_067216963.1-Marinomonas-gallaica:0.0928614607)79.9/93:0.0336039625,WP_067204681.1-Marinomonas-aquimarina:0.0329485628)88.1/93:0.0340745436,WP_055461762.1-Marinomonas-fungiae:0.0705068024)85.3/93:0.0542828632)94.7/98:0.1247383261)93.3/100:0.0609871189)0/82:0.0054075282,WP_100256321.1-Reinekea-forsetii:0.8110326186)91.9/90:0.0816130423)45.1/25:0.0147208813)43/71:0.0522812023)85.4/71:0.0588155222,((((WP_267333169.1-unclassified-Gilliamella:0.2088556894,WP_267361838.1-Gilliamella-sp.-B2772:0.1827924461)19.5/60:0.0113490819,WP_086361229.1-Gilliamella:0.1556058273)62.4/95:0.0321331067,(((((WP_086319877.1-Gilliamella-apis:0.0233799539,WP_198214961.1-Gilliamella-sp.-M0320:0.0691098162)92.7/100:0.0396145595,WP_039129858.1-Gilliamella:0.0286342373)73/100:0.0127124891,(WP_065737496.1-Gilliamella-apicola:0.0412125771,WP_267331332.1-unclassified-Gilliamella:0.0763797769)96.2/100:0.0701544203)88.6/99:0.0459115250,((WP_065579324.1-Gilliamella-apis:0.0986296123,WP_160432071.1-Gilliamella-sp.-Pas-s25:0.1384442850)82.5/80:0.0251042980,(WP_065587409.1-Gilliamella-apicola:0.0751936799,WP_065580031.1-Gilliamella-apicola:0.0807604495)89.3/100:0.0370428742)75/72:0.0145039879)91.5/99:0.0498910417,((WP_065574597.1-Gilliamella-apicola:0.0686325276,WP_065569248.1-Gilliamella-apicola:0.0453100599)50.9/100:0.0093608019,WP_065610260.1-Gilliamella-apicola:0.0502617935)97.5/100:0.0801972731)19.2/57:0.0132305213)96.4/96:0.1653068566,(WP_247177500.1-Escherichia-coli:0.6879296357,YP_009985672.1-Escherichia-phage-muut:0.2492971414)99.5/97:0.4634307860)0/41:0.0044825301)75.8/72:0.1223498702,WP_028357569.1-Brackiella-oedipodis:0.2613312874)99/100:0.2826973820)83.6/53:0.0626433873)57.4/64:0.0804330303,(WP_205000141.1-Parashewanella-hymeniacidonis:0.0739248363,WP_129946484.1-Shewanella-sp.-OPT22:0.0642817574)100/100:0.8851955523)89.7/91:0.1980827226)91/95:0.2097306025,((WP_261298106.1-Shewanella-sp.-SP1S2-7:0.1147734219,WP_167678467.1-Shewanella-aestuarii:0.1155586211)99.8/100:0.2907941967,WP_248994245.1-Shewanella-gaetbuli:0.3258565001)42.2/95:0.1045674614)75/47:0.0855515347,WP_212599444.1-Shewanella-sp.-M16:0.3502333004)86.1/46:0.0903376335,WP_261923504.1-Shewanella-sp.-NFH-SH190041:0.2416273139)52.1/45:0.0701822718,WP_102523418.1-Vibrio-tapetis:0.2154948901)82.3/25:0.0424071022)56.4/34:0.0562322796,(WP_041156219.1-Vibrio-mytili:0.1516778848,(WP_047480300.1-Vibrio-campbellii:0.0737625680,WP_045494469.1-Vibrio-hyugaensis:0.0499289099)98.2/100:0.1897409566)96.9/76:0.1811318411)75.4/44:0.0541861285,((WP_069589493.1-Salinivibrio-sp.-DV:0.1110257694,WP_077662143.1-Salinivibrio-proteolyticus:0.0184281769)74.8/100:0.0277924795,((WP_077659434.1-Salinivibrio-kushneri:0.1637433300,WP_077659452.1-Salinivibrio-kushneri:0.0360300243)73.5/100:0.0304778338,WP_269578450.1-Salinivibrio-kushneri:0.1049059824)85.9/100:0.0507625779)100/100:0.3680412082)25.9/15:0.0236597766,((WP_011638275.1-Shewanella-frigidimarina:0.1706866117,(WP_220774251.1-Shewanella-glacialipiscicola:0.0673094834,WP_182751750.1-Shewanella-sp.-SG44-2:0.1636393005)94/100:0.1023608145)85/100:0.0615169136,((WP_197173106.1-Pseudoalteromonas-sp.-NZS11_1:0.1546598434,WP_182748079.1-Pseudoalteromonas-sp.-SR41-1:0.2426531037)74.1/100:0.0539181613,WP_166378370.1-Pseudoalteromonas-sp.-Z9A4:0.2232815275)56.8/98:0.0453863876)87.1/96:0.0504584260)82.1/22:0.0412924567,(WP_014230611.1-Vibrio-sp.-EJY3:0.1503517182,WP_005395829.1-Vibrio-diabolicus:0.2735246950)84.5/22:0.0671868409)35.3/14:0.0171060151,WP_076646094.1-Vibrio-sp.-10N.222.47.A9:0.2031778577)0/9:0.0000020489,(((WP_102457123.1-Vibrio-breoganii:0.1426216625,(WP_031827242.1-Vibrio-parahaemolyticus:0.3943364912,(WP_047110243.1-Vibrio-vulnificus:0.3302799367,WP_102313365.1-Vibrio-cyclitrophicus:0.1777079768)90.2/98:0.1116461718)84.2/81:0.0691558522)34.5/28:0.0441661247,((((WP_061023752.1-Vibrio-splendidus:0.0878842187,WP_220768829.1-Shewanella-sp.-MBTL60-007:0.1419102130)86.4/94:0.0567589648,WP_182754472.1-Shewanella-sp.-SG41-3:0.1835354467)73.6/28:0.0336857649,((WP_248991952.1-Shewanella-profunda:0.1239570676,WP_059744187.1-Shewanella-frigidimarina:0.2655229574)82.7/42:0.0418641757,((WP_110456309.1-Shewanella-algidipiscicola:0.1974711934,WP_204999274.1-Parashewanella-hymeniacidonis:0.1790941138)8/50:0.0357727832,WP_194091044.1-Vibrio-hibernica:0.2813560746)87.3/73:0.0810116633)79.9/29:0.0315076193)53.7/72:0.0188265825,WP_222242368.1-Pseudoalteromonas-piscicida:0.4931964802)87.1/83:0.0500241803)75.2/19:0.0263472620,((WP_045031875.1-Photobacterium-phosphoreum:0.0304575581,WP_107213458.1-Photobacterium-phosphoreum:0.0417346348)98.6/100:0.1668314717,(WP_172380549.1-Vibrio-sp.-Vb339:0.1313511248,WP_102443327.1-Vibrio-breoganii:0.1403618446)56.4/95:0.0715465552)87.3/85:0.0596333741)92/18:0.0736592500)78.8/31:0.0208224333,((((WP_017027407.1-Vibrio-breoganii:0.1507838059,(WP_102532836.1-Vibrio-sp.-10N.286.48.B7:0.1606368720,WP_254101081.1-Alteromonas-sp.-LMIT007:0.1869572487)48.6/100:0.0409075358)73/86:0.0389269405,(((((WP_138589904.1-Pseudoalteromonas-aurantia:0.2099205997,(WP_218529053.1-Shewanella-sp.-NIFS-20-20:0.1414921066,WP_225483056.1-Vibrio-vulnificus:0.1023170512)90.3/100:0.0743523772)75.6/93:0.0558462121,WP_012549058.1-Aliivibrio-salmonicida:0.1870879587)79.4/90:0.0351158215,WP_193262143.1-Vibrio-parahaemolyticus:0.3330756795)74/70:0.0151692724,(WP_152797327.1-Salinivibrio-sp.-VYel1:0.3182169073,WP_129120576.1-Veronia-nyctiphanis:0.1892652342)86.7/95:0.0868164386)0/68:0.0000026124,(WP_017449116.1-Vibrio-parahaemolyticus:0.1647232808,WP_203433235.1-Shewanella-sp.-LZH-2:0.2521954835)83/67:0.0462136564)89/90:0.0361058692)76.1/34:0.0262105431,WP_108083222.1-Vibrio-sp.-ZF-223:0.1988284829)88.2/23:0.0421364993,((WP_108189654.1-Vibrio-splendidus:0.1962426622,WP_243579549.1-Vibrio-splendidus:0.2504565290)0/23:0.0067517631,WP_102448490.1-Vibrio-splendidus:0.1642435650)70.3/23:0.0166317809)89.8/42:0.0374913084)0/17:0.0000022156,((((WP_065600047.1-Aliivibrio-sp.-1S128:0.1248674355,WP_155682326.1-Aliivibrio-fischeri:0.1132651937)81.5/100:0.0363007113,WP_102482947.1-Vibrio-splendidus:0.1071125811)41/74:0.0362610139,WP_102394203.1-Vibrio-sp.-10N.222.54.B6:0.1791006603)42.1/44:0.0281975346,(WP_063660043.1-Aliivibrio-fischeri:0.1548198749,(WP_102385704.1-Vibrio-cyclitrophicus:0.1414928579,WP_239937663.1-Vibrio-chagasii:0.1694299605)80.5/89:0.0506777416)88.6/66:0.0570407991)42/74:0.0141586041)93.9/24:0.0264612209,((((WP_135455718.1-Vibrio-echinoideorum:0.1425154022,WP_048899071.1-Photobacterium-swingsii:0.1835469866)78.2/100:0.0254885163,(WP_161672404.1-Vibrio-toranzoniae:0.0720609695,WP_017040654.1-Vibrio-genomosp.-F10:0.2460259440)87.1/89:0.0582572001)78.1/74:0.0211279076,((WP_137027437.1-Vibrio-kanaloae:0.0291848276,WP_108117598.1-Vibrio-splendidus:0.0907852052)85/100:0.0433444831,WP_137025093.1-Vibrio-kanaloae:0.1941475649)83.5/47:0.0541604123)89.4/51:0.0392318342,((((((WP_017008437.1-Enterovibrio-norvegicus:0.1256439087,WP_188008910.1-Grimontia-hollisae:0.1914688379)93.2/100:0.0869534265,(((WP_103194250.1-Vibrio-vulnificus:0.0513212699,WP_158146950.1-Vibrio-fluvialis:0.1453769037)65.8/100:0.0178933639,WP_233927909.1-Vibrio-fluvialis:0.2196398497)98.3/100:0.1472512505,(WP_171342182.1-Vibrio-cyclitrophicus:0.1754772267,WP_161972844.1-Enterovibrio-baiacu:0.1574983758)62.1/100:0.0416924007)96.3/100:0.1308908037)0/42:0.0000025425,WP_243025268.1-Vibrio-cincinnatiensis:0.7245799488)11.4/43:0.0125181765,WP_017020702.1-Aliivibrio-logei:0.2292396599)51.4/43:0.0113245620,WP_016784063.1-Vibrio:0.1069619499)42.9/43:0.0228792487,(((WP_005597034.1-Vibrio-scophthalmi:0.1741146245,WP_135385147.1-Vibrio-tasmaniensis:0.2072816407)58.2/99:0.0529040472,(WP_108176621.1-Vibrio-splendidus:0.0604558453,WP_239918548.1-Vibrio-sp.-Isolate22:0.0329113392)76.9/100:0.0345228939)87.7/100:0.0627343998,WP_140342839.1-Vibrio-parahaemolyticus:0.2286912899)84/88:0.0485731999)91.9/41:0.0640051987)0/13:0.0000022933)92/52:0.0354030684,((((WP_171343915.1-Vibrio-cyclitrophicus:0.0805988968,((WP_048661380.1-Vibrio-crassostreae:0.1041785373,WP_193846925.1-Vibrio-sp.-OPT10:0.1049932686)76.1/100:0.0428840476,WP_238130349.1-Vibrio-cincinnatiensis:0.2313542285)45/73:0.0427451212)85.2/67:0.0413784584,(WP_161753058.1-Vibrio-toranzoniae:0.1725793337,(WP_102248986.1-Vibrio-tasmaniensis:0.2310484461,WP_135445135.1-Vibrio-tasmaniensis:0.0976492300)96.8/100:0.1138868706)85.8/80:0.0441349013)75.2/56:0.0112845583,(((WP_139685731.1-Vibrio-tasmaniensis:0.1533657676,WP_145570128.1-Vibrio-tasmaniensis:0.1870068917)82.2/97:0.0421168848,((WP_061039794.1-Vibrio-coralliirubri:0.0985605214,(WP_044365158.1-Vibrio-fluvialis:0.3607574524,WP_025523961.1-Vibrio-parahaemolyticus:0.1808503825)89.6/100:0.0640683385)89.7/100:0.0824675145,WP_065194916.1-Photobacterium-phosphoreum:0.1852438912)88.6/95:0.0957192639)61/16:0.0313359435,(WP_248368760.1-Vibrio-sp.-1CM8B:0.1757360002,((WP_046209502.1-Vibrio:0.1713378361,WP_108186198.1-Vibrio-splendidus:0.2056892878)33.3/77:0.0151778690,WP_248386338.1-Vibrio-sp.-ED004:0.2591138077)89.1/94:0.0643482520)64.7/15:0.0199924062)76.5/92:0.0120147062)91/80:0.0472429676,((((WP_063656223.1-Aliivibrio-fischeri:0.1431020518,WP_155675904.1-Aliivibrio-fischeri:0.1033936933)78.7/43:0.0293574223,WP_017105169.1-Vibrio:0.1874173016)87.2/40:0.0427625262,((WP_102459739.1-Vibrio-lentus:0.1345131697,WP_102363333.1-Vibrio-cyclitrophicus:0.1664358132)56.3/94:0.0507901240,(WP_102554090.1-Vibrio-splendidus:0.1702369553,WP_130314193.1-Vibrio-vulnificus:0.2039418475)56.1/93:0.0229578792)71.8/99:0.0192255330)84.5/79:0.0340469643,((WP_137008959.1-Vibrio-kanaloae:0.1588758836,WP_135353160.1-Aliivibrio-fischeri:0.0980235774)91.5/100:0.0765505115,WP_248638627.1-Vibrio-crassostreae:0.1679141553)79.4/95:0.0319964579)74.6/79:0.0127623733)83.9/18:0.0265572058)68/19:0.0260104812,WP_077679840.1-Vibrio-kanaloae:0.1641012782)43.3/35:0.0164078475,(((((((WP_268674123.1-Vibrio-scophthalmi:0.1605484541,WP_017085924.1-Vibrio-splendidus:0.1368437595)22.5/92:0.0579484409,WP_102420589.1-Vibrio-cyclitrophicus:0.1102478061)88/97:0.0763933647,(WP_102457030.1-Vibrio-cyclitrophicus:0.1412524535,WP_102298256.1-Vibrio-cyclitrophicus:0.1047465902)42.5/84:0.0413070299)84.5/77:0.0379773569,((WP_059021773.1-Vibrio-coralliirubri:0.0670004766,WP_135380937.1-Vibrio-tasmaniensis:0.1203001495)75.2/100:0.0217531249,WP_102276040.1-Vibrio-lentus:0.1526831809)74.7/94:0.0166127686)85.6/77:0.0512991816,WP_265133188.1-Vibrio-splendidus:0.1493946600)78.2/82:0.0318966691,(((WP_017063118.1-Vibrio:0.1510787275,WP_132940416.1-Vibrio-crassostreae:0.1331102767)86.2/97:0.0579491190,WP_206974914.1-Vibrio-parahaemolyticus:0.2176178462)24.3/21:0.0173342345,WP_248388778.1-Vibrio-cyclitrophicus:0.1676759208)77.5/21:0.0288838322)85.5/21:0.0384288460,((((WP_155682312.1-Aliivibrio-fischeri:0.0962490449,WP_012532661.1-Aliivibrio-fischeri:0.1601888052)29.1/30:0.0343145294,((WP_108162474.1-Vibrio-splendidus:0.0364403386,WP_004742905.1-Vibrio-tubiashii:0.0642086535)99.7/100:0.1502146602,(((WP_102503160.1-Vibrio-splendidus:0.1675060531,WP_155677763.1-Aliivibrio-fischeri:0.1591438690)92/42:0.1068300267,(WP_016768872.1-Vibrio-cyclitrophicus:0.1272434351,WP_102428030.1-Vibrio-sp.-10N.261.54.E10:0.2169293996)71.5/52:0.0290717012)45.2/16:0.0123958159,WP_017069285.1-Vibrio-crassostreae:0.1093312723)25/16:0.0371175469)88.6/59:0.0705501862)91.5/86:0.0650009339,((((WP_130049361.1-Aliivibrio-finisterrensis:0.0790342940,WP_135445143.1-Vibrio-tasmaniensis:0.2621625645)76/97:0.0222871785,(WP_019826652.1-Vibrio-splendidus:0.1389721732,WP_182796407.1-Vibrio-sp.-SG41-7:0.2411015818)84.2/99:0.0532347175)81.6/96:0.0379403364,WP_260256886.1-Vibrio-sp.-NC2:0.1366868099)76.2/91:0.0175594868,(WP_258428294.1-Vibrio-sp.-RM-69-4:0.1614438776,(WP_193834777.1-Vibrio-sp.-OPT20:0.0548354805,(WP_029865567.1-Vibrio-parahaemolyticus:0.0756074771,WP_213967907.1-Vibrio-alginolyticus:0.0400845820)98.5/100:0.1084029365)75.3/100:0.0273012933)92.7/86:0.0815091407)36/12:0.0107761803)85.7/89:0.0414655658,(((WP_162023628.1-Shewanella-sp.-Arc9-LZ:0.0951587153,WP_248997855.1-Shewanella-marinintestina:0.0481457840)97.7/100:0.1094581870,WP_017094106.1-Vibrio-splendidus:0.1109632536)82.2/95:0.0475107142,((((((WP_243584123.1-Vibrio-splendidus:0.1483835823,(WP_163929979.1-Paraferrimonas-sp.-SM1919:0.1108859530,WP_087845049.1-Photobacterium-malacitanum:0.1940999102)92.1/93:0.0863376211)74.2/35:0.0244058213,((WP_102462022.1-Vibrio-splendidus:0.0686724097,WP_187463684.1-Photobacterium-sp.-BZF1:0.0863517330)96.6/100:0.1345942092,WP_259574191.1-Shewanella-baltica:0.2904939127)12.9/42:0.0381913665)75.7/38:0.0851473426,(WP_213886549.1-Vibrio-alginolyticus:0.2677463898,WP_158173598.1-Vibrio-alginolyticus:0.0578585400)98.7/100:0.1204738048)80.2/35:0.0311883104,WP_009843931.1-Vibrio-sp.-AND4:0.0731902056)56.5/32:0.0115509001,WP_065296243.1-Vibrio-natriegens:0.1442707529)77.2/82:0.0267150344,((WP_004733311.1-Vibrio-splendidus:0.1303266593,WP_171315631.1-Vibrio-splendidus:0.1588207235)10.2/83:0.0239968943,WP_102353859.1-Vibrio-cyclitrophicus:0.1576364602)92.4/97:0.0688191887)76/88:0.0146469893)89.6/78:0.0594025435)36/19:0.0127354684)78.3/68:0.0189983205)93.5/69:0.0331775736,((((((((((((((((((WP_188026786.1-Tolumonas-osonensis:0.0391186927,WP_024870911.1-Tolumonas-lignilytica:0.1461327149)88.1/100:0.1002432263,WP_105076797.1-Pantoea-ananatis:0.5412257461)0/18:0.0847273339,((((WP_088208149.1-Enterobacter-cancerogenus:0.1418145318,WP_121932958.1-Enterobacter-hormaechei:0.1105339383)87.3/100:0.0560361165,WP_023330866.1-Enterobacter:0.0498703083)99.4/100:0.2061856272,(WP_120453320.1-Kalamiella-piersonii:0.0937406818,((WP_214216788.1-Rosenbergiella-epipactidis:0.0751452598,WP_241623482.1-Rosenbergiella-epipactidis:0.0237009848)99.2/100:0.1821443258,(WP_192411534.1-Pantoea-sp.-PNT02:0.0746645836,((WP_125311258.1-Pantoea-dispersa:0.0968065048,((WP_150019269.1-unclassified-Pantoea:0.0842453938,WP_140917215.1-Pantoea-deleyi:0.0505027380)49.5/89:0.0077321630,WP_069025179.1-Pantoea-agglomerans:0.0081953701)95.7/92:0.0438881848)87.4/91:0.0302998298,WP_097116658.1-Pantoea-sp.-GL120224-02:0.0000025783)91.9/96:0.0714437070)95.9/100:0.1111251655)91.4/96:0.0980274914)85.9/48:0.0410188271)64.7/45:0.0163388393,((WP_269294270.1-Pantoea-sp.-LMR881:0.0536217629,WP_215846277.1-Pantoea-sp.-EnvD:0.0985625247)94/100:0.0817408566,WP_187508308.1-Pantoea-agglomerans:0.2499165641)81.8/52:0.0343307830)93.9/95:0.0768468664)94.2/94:0.0674982357,((WP_120356561.1-Alginatibacterium-sediminis:0.4733099398,((WP_075989065.1-Vibrio-fluvialis:0.0485512706,WP_044366945.1-Vibrio-fluvialis:0.0407199592)39.5/94:0.0262442904,(WP_076657081.1-unclassified-Vibrio:0.0812622049,WP_239929047.1-Vibrio-kanaloae:0.0417480667)98.8/100:0.1332125995)88.4/100:0.0736827216)3.4/32:0.0709948818,(WP_182744595.1-Shewanella-sp.-SR44-4:0.2925300135,WP_144135068.1-Kosakonia-radicincitans:0.3861684700)86.2/95:0.0766666124)80.1/93:0.0317873436)82/100:0.0261731960,(WP_020431327.1-Vibrio-fluvialis:0.1287066541,WP_055453311.1-Vibrio-fluvialis:0.0654356330)85.4/100:0.0597998529)95.1/99:0.0595660449,((((WP_261600599.1-Photobacterium-sp.-TY1-4:0.0919713935,(WP_114765998.1-Vibrio-rhodolitus:0.1589977559,WP_182108830.1-Vibrio-marinisediminis:0.1077152520)95.3/100:0.1387980329)79.4/95:0.0497500074,(((WP_211647400.1-Photobacterium-sp.-GJ3:0.0632239638,WP_120511428.1-Photobacterium-salinisoli:0.0525596113)96.3/100:0.1154595167,((WP_200225058.1-Rahnella-aceris:0.1396084958,WP_195814554.1-Rahnella:0.1340478472)88.2/100:0.0528256726,WP_169403689.1-Rouxiella-aceris:0.0871580000)99.4/100:0.1961427305)32.9/23:0.0383136916,WP_101064017.1-Moritella-sp.-Urea-trap-13:0.2989474543)54.5/20:0.0199515984)31.1/23:0.0175282492,(((WP_170899585.1-Vibrio-sp.-3-2-1:0.1316224698,WP_017221010.1-Moritella-dasanensis:0.1696374057)85.2/100:0.0463634724,WP_107284330.1-Photobacterium-lipolyticum:0.1286682077)90.7/100:0.0550291674,WP_159197271.1-Escherichia-coli:0.1531421980)84.1/99:0.0414829122)87.8/23:0.0231719379,((WP_112404745.1-Vibrio-diazotrophicus:0.0607030449,WP_202661182.1-Vibrio-fluvialis:0.0699305575)94.5/99:0.0910025955,((((((WP_263100201.1-Kosakonia-sp.-ML.JS2a:0.0374535535,WP_200550470.1-Kosakonia-sp.-LAM2021:0.1063192100)98.3/100:0.1372819078,((WP_015854950.1-Musicola-paradisiaca:0.0479097900,WP_192457620.1-Musicola-keenii:0.1017891236)100/100:0.2832724751,(WP_094109047.1-Lonsdalea-iberica:0.1242042587,(WP_026742149.1-Lonsdalea-quercina:0.0450161898,WP_094117837.1-Lonsdalea-britannica:0.0386293369)96.8/100:0.0904445217)83.6/95:0.0389281060)71.9/64:0.0320653621)84.9/80:0.0786229058,(WP_232612305.1-Photobacterium-phosphoreum:0.3789077004,WP_136989485.1-Vibrio-sp.-F13:0.3088154356)92.5/99:0.1176570244)64/93:0.0301945811,WP_006646466.1-Photobacterium-sp.-SKA34:0.2322001614)0/31:0.0351636255,WP_172533652.1-Vibrio-aestuarianus:0.4231219092)59.3/68:0.0268038360,((WP_038178862.1-Vibrio-rhizosphaerae:0.1442501952,WP_046218796.1-Photobacterium-halotolerans:0.2440304552)20.3/21:0.0315146131,WP_107187026.1-Photobacterium-kishitanii:0.2392681576)85.3/21:0.0605477557)76/17:0.0373799935)83.7/22:0.0423618314)80.2/89:0.0193906912)78.2/81:0.0167074329,(((((WP_239941760.1-Vibrio-mediterranei:0.1382578052,WP_252012765.1-Vibrio-sp.-SCSIO-43140:0.1640675046)69.9/94:0.0432931378,WP_252012797.1-Vibrio-sp.-SCSIO-43140:0.1274992763)71.7/94:0.0452031223,(WP_102532726.1-Vibrio-sp.-10N.286.48.B7:0.1176758786,WP_250617847.1-Vibrio-sp.-S4B1:0.0819518264)86.4/100:0.0576332350)88.8/95:0.0690593052,WP_268679810.1-Vibrio-barjaei:0.1493540908)87.4/100:0.0504484480,WP_045898906.1-Enterobacteriaceae:0.1910555994)84.5/85:0.0397388366)89.2/93:0.0241190251,((((WP_041510674.1-Shewanella:0.0588084866,WP_187575439.1-Shewanella-algae:0.0840673278)83.5/99:0.0478672890,(WP_208158397.1-Shewanella-algae:0.0961843516,WP_208157589.1-Shewanella-algae:0.0477491467)92.2/100:0.0618811665)86/98:0.0695801290,(WP_045460498.1-Vibrio-hyugaensis:0.1791234489,WP_220051788.1-Shewanella-sp.-FJAT-51860:0.1925589721)84.5/99:0.0466584539)90.7/93:0.0491717794,((WP_094127815.1-Vibrio-sp.-V09_P4A23P171:0.1639952964,WP_268809473.1-Vibrio-parahaemolyticus:0.1699631374)90.9/98:0.0783978869,(WP_107294031.1-Photobacterium-phosphoreum:0.0493378114,WP_107276976.1-Photobacterium-sp.-NCIMB-13483:0.0700493719)99.1/100:0.1098505038)79/80:0.0204824248)74.2/19:0.0077669701)71.9/22:0.0071797886,((((((((((((WP_011771697.1-Psychromonas-ingrahamii:0.1222101553,WP_182690147.1-Psychromonas-sp.-SR45-3:0.1888003863)96.3/99:0.1385289638,WP_161119826.1-Photobacterium-lucens:0.1592173247)89.9/96:0.0763461596,((((WP_151182432.1-Photobacterium-damselae:0.1603981868,WP_086367281.1-Photobacterium-damselae:0.0687519304)51.5/46:0.0177897290,(WP_045045404.1-Photobacterium-kishitanii:0.0360075110,(WP_045030671.1-Photobacterium-phosphoreum:0.0595259638,WP_107264793.1-Photobacterium-phosphoreum:0.0321897123)88/97:0.0302287693)97.7/100:0.0969369650)6.8/44:0.0246587342,WP_053987023.1-Photobacterium-leiognathi:0.1158850912)75.3/41:0.0235365265,WP_115426223.1-Photobacterium-damselae:0.0963898384)82.8/70:0.0304502765)80.5/35:0.0398420985,(((((WP_065208126.1-Photobacterium-phosphoreum:0.1599422277,(((((WP_232582982.1-Photobacterium-phosphoreum:0.1102627568,WP_232578536.1-Photobacterium-carnosum:0.0627064691)73.3/61:0.0095709735,WP_107228404.1-Photobacterium-phosphoreum:0.1087594358)75.9/61:0.0128528190,WP_256383785.1-Photobacterium-toruni:0.0435040481)93.8/100:0.0747444703,(WP_107293979.1-Photobacterium-phosphoreum:0.1183700257,WP_107228425.1-Photobacterium-phosphoreum:0.2253173185)90.1/66:0.0798703278)64.2/55:0.0161970086,((WP_065176337.1-Photobacterium-aquimaris:0.0768660179,WP_107199726.1-Photobacterium-phosphoreum:0.1005335035)44.9/100:0.0333188334,WP_023931546.1-Photobacterium-leiognathi:0.1627815628)97.7/100:0.1264589348)80.4/63:0.0277720696)77.4/64:0.0117986718,WP_060997345.1-Photobacterium-aquimaris:0.1610631191)86.3/63:0.0382108803,((((WP_232575020.1-Photobacterium-carnosum:0.1770599524,WP_232603643.1-Photobacterium-carnosum:0.1661223222)43.4/33:0.0237903747,WP_232585319.1-Photobacterium-iliopiscarium:0.1559671142)37.7/33:0.0303689955,WP_232615715.1-Photobacterium-carnosum:0.1504394748)73.8/83:0.0126806803,WP_232600381.1-Photobacterium-carnosum:0.1023372060)89.2/90:0.0234358294)0/56:0.0000022625,WP_161680902.1-Proteus:0.5395454743)91.3/97:0.0423963642,WP_237041371.1-Photobacterium-damselae:0.0987651490)12.3/79:0.0068399067)59.7/37:0.0262367380,((((WP_238087945.1-Photobacterium-damselae:0.0697415954,(WP_065172005.1-Photobacterium-damselae:0.1042878673,WP_107185088.1-Photobacterium-leiognathi:0.0993574101)82.1/100:0.0280273561)70.8/49:0.0168439525,(WP_106261939.1-Photobacterium-damselae:0.1077937183,WP_017004373.1-Enterovibrio-norvegicus:0.2523688468)10.3/42:0.0613410533)84.3/44:0.0432052071,WP_151170663.1-Photobacterium-damselae:0.0395407985)74.7/43:0.0276703270,WP_045063675.1-Photobacterium-leiognathi:0.1085266317)85/45:0.0508312404)86.1/60:0.0403619028,WP_237042700.1-Photobacterium-damselae:0.1208658251)87.9/92:0.0362614936,((((WP_230432956.1-Plesiomonas-shigelloides:0.1183805459,WP_222763364.1-Vibrio-fluvialis:0.1529423815)58.9/100:0.0246335043,(WP_117235674.1-Vibrio-maerlii:0.1526297074,(WP_103882085.1-Vibrio-hangzhouensis:0.0705026242,WP_265676699.1-Vibrio-sp.-ZSDZ65:0.1583235348)91.2/100:0.0856702841)90.3/98:0.0850455597)26.5/65:0.0110871067,((WP_013365656.1-_Enterobacter:0.2916584596,WP_214579582.1-Enterobacter-asburiae:0.1022559547)79.3/63:0.0489123761,((WP_258988101.1-Scandinavium-hiltneri:0.0818040683,WP_181620050.1-Enterobacter-sp.-RHBSTW-00994:0.0758741866)95.6/100:0.0792273222,WP_193129600.1-Klebsiella-aerogenes:0.0836933292)47.4/99:0.0412040100)100/99:0.2571603177)75.8/36:0.0201773586,(WP_251878087.1-Grimontia-sp.-020920N:0.1726013497,(WP_171138527.1-Vibrio-plantisponsor:0.0878089263,WP_102966424.1-Vibrio:0.1078816041)97/99:0.1380605509)72.3/68:0.0145548457)96.4/98:0.0800326211)89.6/59:0.0412919635,((((WP_069212356.1-Vibrio-cholerae:0.0366615256,WP_202641945.1-Vibrio-fluvialis:0.1514506709)91.5/100:0.1060283950,(WP_107213932.1-Photobacterium-phosphoreum:0.1590789564,(WP_065173166.1-Photobacterium-kishitanii:0.1609391483,WP_047876260.1-Photobacterium-aphoticum:0.2197228959)70/99:0.0290546839)88.1/99:0.0663404642)79.4/92:0.0525756755,WP_199351799.1-Vibrio-cholerae:0.1100707314)45.1/70:0.0335467801,WP_175242188.1-Vibrio-cholerae:0.1165888907)83.9/47:0.0358364908)0/11:0.0000023873,(WP_000864154.1-Vibrio-cholerae:0.0998728657,WP_102786270.1-Vibrio-cholerae:0.1026402293)69.3/76:0.0231116710)74.4/13:0.0238787238,((WP_095464062.1-Vibrio-cholerae:0.1082053121,WP_149560528.1-Vibrio-cholerae:0.1105728995)86.8/98:0.0365195594,WP_095478449.1-Vibrio:0.1082010353)52.3/90:0.0405784737)67.2/55:0.0147644382,(WP_208891742.1-Vibrio-anguillarum:0.1975683622,WP_148533341.1-Vibrio-cholerae:0.0622469940)92.9/77:0.0780977828)56.5/17:0.0256607165,((WP_000864156.1-Vibrio-cholerae:0.0783233864,WP_000864152.1-Vibrio-cholerae:0.1133492590)91.6/100:0.0667679126,(WP_199867560.1-Vibrio-anguillarum:0.2114793309,(WP_107294203.1-Photobacterium-sp.-GB-50:0.1966863593,(WP_188234612.1-Vibrio-sp.-Y2-5:0.0882086114,WP_057556016.1-Vibrio-cholerae:0.0152008757)100/100:0.2741450067)52.4/81:0.0306696167)0/67:0.0000024789)91.8/65:0.0476298508)93.8/33:0.0467703270)87.9/55:0.0206824341,(((((((WP_192015074.1-Photobacterium-arenosum:0.0756202960,(WP_088876367.1-Vibrio-mediterranei:0.3129511938,WP_022590465.1-Vibrio-nigripulchritudo:0.3252544537)23.2/86:0.0309664871)90/94:0.0611353081,(WP_172851426.1-Vibrio-coralliilyticus:0.2354066254,(((WP_108452928.1-Enterobacter-cloacae:0.3340187163,WP_237309237.1-Vibrio:0.3666532740)83.6/100:0.0830430014,WP_182742106.1-Shewanella-sp.-SG44-6:0.3251523443)84.7/89:0.0659793137,WP_138627230.1-Pseudoalteromonas-sp.-S4741:0.2479853743)73/84:0.0422782377)84.4/97:0.0399478762)75.8/88:0.0533065096,((WP_095430409.1-Vibrio-vulnificus:0.0620539294,WP_114994956.1-Grimontia-hollisae:0.2654324098)95.2/100:0.1284708515,((WP_211678393.1-Moritella-sp.-36:0.2952627537,WP_269377025.1-Vibrio-sinaloensis:0.2476448365)85.7/96:0.0704856303,(WP_028024862.1-Enterovibrio-calviensis:0.2325723015,WP_017007824.1-Enterovibrio-norvegicus:0.2113904643)78.8/75:0.0346969768)77.6/77:0.0321302510)53.1/87:0.0198245809)89.8/91:0.0412133733,(WP_101201417.1-Paraglaciecola-sp.-MB-3u-78:0.2560041771,WP_039484180.1-Vibrio-sinaloensis:0.2140656343)82.4/90:0.0517188307)75.2/45:0.0116504239,(WP_068898547.1-Enterovibrio-pacificus:0.2063640073,WP_220058598.1-Shewanella-sp.-FJAT-53555:0.3942234354)73.6/86:0.0248989132)44.1/54:0.0192908436,WP_036979211.1-Providencia-alcalifaciens:0.2993595396)75.8/94:0.0199502803,(WP_087853896.1-Photobacterium-andalusiense:0.0382283516,WP_107274737.1-Photobacterium-phosphoreum:0.1385235269)98.8/100:0.1262970542)76.2/50:0.0218878605)74.7/72:0.0131548494,(((WP_064489977.1-Vibrio-parahaemolyticus:0.0565300742,WP_201976964.1-Aeromonas-enteropelogenes:0.1528363724)96.2/100:0.1219098068,WP_224657569.1-Vibrio-vulnificus:0.1609797047)67.5/67:0.0343022627,WP_102475754.1-Shewanella-sp.-10N.286.52.C2:0.3024256153)82.5/62:0.0379283828)85.6/59:0.0294065001,((((WP_033540599.1-Shewanella:0.1978414181,WP_031815865.1-Vibrio-parahaemolyticus:0.1489395930)77.9/100:0.0574595824,WP_258516089.1-Vibrio-alginolyticus:0.1291007206)28.5/100:0.0443994585,WP_193250619.1-Vibrio-parahaemolyticus:0.2167076938)91.5/99:0.0826875667,WP_023404414.1-Vibrio-halioticoli:0.3188532593)72.1/21:0.0242677932)78.7/21:0.0205451564,((WP_102986644.1-Vibrio-diazotrophicus:0.1975928178,(WP_192928291.1-Vibrio-vulnificus:0.0143718076,WP_132698543.1-Vibrio-harveyi-group:0.1577363072)98/100:0.1648050106)78.5/44:0.0411759358,WP_045039948.1-Photobacterium-iliopiscarium:0.2138787803)42.8/18:0.0049990191)83.2/52:0.0213657710,(WP_102563398.1-Vibrio-sp.-10N.286.49.B3:0.1568654660,WP_020328057.1-Vibrio-fluvialis:0.2117424424)33.2/45:0.0242317854)86.1/70:0.0319690772,((((((((((WP_193255073.1-Vibrio-navarrensis:0.0315556165,WP_258469755.1-Vibrio-alginolyticus:0.1493057417)96.4/100:0.1058667475,(WP_239872404.1-Shewanella-insulae:0.1345385581,(WP_239850080.1-Shewanella-sp.-Isolate8:0.0848524865,WP_069440067.1-Pseudoalteromonas-tetraodonis:0.3897245734)87.6/99:0.0902315493)88.8/77:0.0471295190)49.3/58:0.0249288009,(WP_211689658.1-Moritella-sp.-28:0.4049000546,WP_238097025.1-Vibrio-cincinnatiensis:0.2379856655)73.4/45:0.0311844103)81.4/40:0.0334796654,(((WP_011716450.1-Shewanella-sp.-ANA-3:0.3801337675,WP_101040106.1-Psychromonas-sp.-MB-3u-54:0.2476007229)80.9/78:0.0492574965,(WP_009847952.1-Vibrio:0.2436050248,WP_233894267.1-Vibrio-kanaloae:0.1491586168)86.2/100:0.0579964100)53.4/53:0.0189259870,(WP_048661352.1-Vibrio-crassostreae:0.0842527985,WP_114634227.1-Vibrio-splendidus:0.0783039889)95.5/100:0.1208904714)94.4/87:0.0802066001)74.6/55:0.0153366294,WP_102285079.1-Vibrio-lentus:0.1926231343)75.3/44:0.0309865115,((((WP_077647682.1-Salinivibrio-sp.-IB643:0.1534852720,WP_072616325.1-Vibrio-vulnificus:0.2103069398)0/75:0.0000020993,WP_225486226.1-Vibrio-vulnificus:0.2354980782)71.9/73:0.0529408311,WP_099031861.1-Pseudoalteromonas-sp.-GCY:0.1727207465)82.2/88:0.0541857138,(WP_208844179.1-Pseudoalteromonas-xiamenensis:0.1712001643,WP_233976434.1-Vibrio:0.2104770144)42/41:0.0431548316)67.5/59:0.0309830392)42.1/25:0.0282228736,WP_233983986.1-Vibrio-fluvialis:0.1047793610)74.5/25:0.0182960026,(WP_069533349.1-Vibrio-parahaemolyticus:0.1781760623,WP_229630939.1-Vibrio-coralliilyticus:0.2773861139)78.5/98:0.0331940130)44.3/24:0.0121072955,(((WP_068331709.1-Photobacterium-jeanii:0.1920592333,WP_031414678.1-Vibrio-parahaemolyticus:0.2145757721)45.5/96:0.0431175053,(WP_106651046.1-Shewanella-sp.-Pdp11:0.1823224974,WP_206625521.1-Vibrio-vulnificus:0.1907951929)35.5/47:0.0690266927)89.7/42:0.0646011447,((((WP_012551417.1-Aliivibrio-salmonicida:0.1071147589,WP_012276595.1-Shewanella-halifaxensis:0.0731338727)85.9/100:0.0886662777,WP_144217665.1-Shewanella-algae:0.2523914206)82.9/94:0.0574843741,(WP_130066657.1-Aliivibrio-finisterrensis:0.1892844585,WP_248468589.1-Shewanella-sp.-1CM18E:0.1767687874)79/86:0.0481563882)18.3/38:0.0283078052,WP_102571993.1-Vibrio:0.1588754376)89.9/97:0.0659831405)78.7/34:0.0291211508)42/71:0.0159609197,((WP_120030433.1-Vibrio-sinensis:0.1344540705,WP_235060641.1-Vibrio-sp.-SS-MA-C1-2:0.2621091643)71.2/96:0.0215469762,WP_213972370.1-Vibrio-alginolyticus:0.2651810936)79.4/90:0.0206555521)77.5/52:0.0180686407)77.3/78:0.0109320107,(WP_211915530.1-Vibrio-sp.-B1REV9:0.1238965756,(WP_017420706.1-Vibrio-vulnificus:0.1519225183,WP_017024716.1-Vibrio-rumoiensis:0.2297560179)0/37:0.0053336937)86.7/96:0.0442358908)80.8/70:0.0219326726,WP_039841338.1-Vibrio-harveyi-group:0.2718954275)74.6/28:0.0153256992,(((((WP_238813939.1-Vibrio-vulnificus:0.1293712841,(WP_118894008.1-Vibrio-vulnificus:0.1812352097,((WP_259522409.1-Shewanella-baltica:0.1464983046,WP_173243873.1-Shewanella-sp.-DC2-4:0.1745540590)96.6/99:0.1360492230,WP_121852788.1-Parashewanella-spongiae:0.2956234736)58.5/93:0.0423061000)74.6/75:0.0462609520)47.6/80:0.0516755633,(WP_158107962.1-Vibrio-furnissii:0.1066309279,WP_185890410.1-Vibrio-vulnificus:0.2315547878)88.1/93:0.0768416853)78.6/70:0.0285881998,WP_172561824.1-Vibrio-furnissii:0.1390170878)84.6/69:0.0447160471,((((((((WP_155660100.1-Aliivibrio-fischeri:0.1018778790,WP_095457151.1-Vibrio:0.1287363747)90.1/100:0.0544699030,(WP_252020469.1-Vibrio-sp.-SCSIO-43136:0.1097259283,WP_243584106.1-Vibrio-splendidus:0.2244441258)81.1/100:0.0429484862)91.2/100:0.0810264429,WP_166696422.1-Providencia-rettgeri:0.1779893559)88.6/100:0.0899662849,((WP_172855536.1-Vibrio-coralliilyticus:0.2804208437,WP_234235026.1-Shewanella-sp.-AS16:0.2456867093)87.8/100:0.0838204956,WP_088905011.1-Shewanella-marisflavi:0.1793946758)78.5/99:0.0374899417)86.9/99:0.0504703044,(WP_102038478.1-Shewanella-sp.-GutCb:0.2502531731,WP_112353352.1-Shewanella-benthica:0.2632105866)36.8/100:0.0898015538)63.9/99:0.0238771228,WP_029788868.1-Vibrio:0.2788429076)74.8/89:0.0214869686,WP_143680610.1-Vibrio-furnissii:0.1607969147)58.4/65:0.0263653889,WP_165312521.1-Vibrio-ziniensis:0.1376461773)66/63:0.0181119867)82.6/51:0.0395610856,WP_238819535.1-Vibrio-furnissii:0.1289085612)75.1/56:0.0237087557)75.9/68:0.0079832124)77.6/87:0.0075989727,(((WP_067416136.1-Enterovibrio-coralii:0.1112950228,WP_149593442.1-Pseudoalteromonas-sp.-FUC4:0.2331485515)95.2/92:0.0840047211,((((((((((WP_232600401.1-Photobacterium-carnosum:0.1425555236,WP_065173140.1-Photobacterium-kishitanii:0.2149594116)84.9/91:0.0547178908,(WP_176284769.1-Photobacterium-damselae:0.2402082101,WP_088132129.1-Vibrio-cholerae:0.1304202739)7.7/38:0.0306822794)87.8/91:0.0489775132,((WP_084988554.1-Vibrio:0.1661368185,WP_206634697.1-Vibrio-vulnificus:0.2302225462)90.4/90:0.0910746981,WP_152138288.1-Plesiomonas-shigelloides:0.2165146705)0/24:0.0042528225)76.2/25:0.0260766171,(WP_000864153.1-Vibrio-cholerae:0.0170309343,WP_251878143.1-Grimontia-sp.-020920N:0.2128853195)93.1/100:0.1198261000)33.5/52:0.0410626302,WP_045618469.1-Vibrio-vulnificus:0.1852692726)57/71:0.0248890989,(WP_249572008.1-Vibrio-cholerae:0.1272579388,WP_103155729.1-Vibrio-vulnificus:0.1531544975)30.3/53:0.0129103975)75.4/87:0.0179680531,WP_255122387.1-Vibrio-cholerae:0.1515525221)80.5/95:0.0188996148,((WP_237041393.1-Photobacterium-damselae:0.1203470223,WP_080158029.1-Photobacterium-piscicola:0.2481216046)93.1/90:0.1019079625,(((WP_267741605.1-Vibrio-cholerae:0.1568319900,(WP_172567459.1-Vibrio-fluvialis:0.1231121086,WP_199354891.1-Vibrio-cholerae:0.0405036678)94.3/100:0.1100553094)6.8/75:0.0396623817,WP_102137258.1-Vibrio-cholerae:0.1137952834)89.5/87:0.0697814661,(WP_161129723.1-Photobacterium-lucens:0.1799590861,WP_148560094.1-Vibrio-mimicus:0.2347434841)17.1/51:0.0120954573)59.2/61:0.0343658037)84/91:0.0506563537)71.9/75:0.0188565644,((WP_148519618.1-Vibrio-cholerae:0.0304481965,(WP_167570263.1-Aeromonas-veronii:0.0754807057,WP_000864158.1-Vibrio-metoecus:0.0394870650)92.2/100:0.0361187885)73.9/100:0.0157089709,WP_148520854.1-Vibrio-cholerae:0.0449604341)99/98:0.1102501791)84.2/65:0.0580845554,((((((WP_045569365.1-Vibrio:0.0976092554,WP_039437343.1-Vibrio-navarrensis:0.0538901820)95.8/100:0.1008864090,WP_176299354.1-Vibrio-sp.-05-20-BW147:0.2004709054)85.7/100:0.0626751103,WP_140233673.1-Shewanella-polaris:0.3098464839)91.9/99:0.1114257996,((WP_017031451.1-Vibrio-breoganii:0.1175000045,(WP_192022372.1-Shewanella-sp.-WPAGA9:0.2577280613,WP_248975945.1-Shewanella-kaireitica:0.1632578458)86.4/26:0.0535251975)80.3/26:0.0400327756,(WP_223418564.1-Shewanella-inventionis:0.1403502459,WP_011497090.1-Shewanella-denitrificans:0.3152553775)94.5/100:0.1612968805)59.4/25:0.0539565963)89.2/99:0.0821539582,(WP_217542775.1-Vibrio-metschnikovii:0.1228482248,WP_113593871.1-Vibrio:0.0564870187)97.9/100:0.1203102563)56.6/94:0.0214882546,(WP_086585081.1-Vibrio-parahaemolyticus:0.0661218706,WP_217557493.1-Vibrio-metschnikovii:0.0427704039)95.4/100:0.0909482618)88.9/96:0.0475994209)0/57:0.0000020464)93.6/46:0.0422182986,(WP_047463293.1-Vibrio-fluvialis:0.1002292574,WP_193167845.1-Vibrio-navarrensis:0.3737371049)96.3/88:0.1208693033)0/38:0.0000028879)89.8/41:0.0252942059,(((WP_107253996.1-Photobacterium-indicum:0.1828796503,WP_107295906.1-Photobacterium-phosphoreum:0.2118566889)76.3/100:0.0531555499,WP_102395415.1-Enterovibrio-norvegicus:0.1463097150)87.6/82:0.0587841224,(WP_045069930.1-Photobacterium-leiognathi:0.1764513054,WP_058428281.1-Pseudoalteromonas:0.2970481152)75.7/98:0.0386117434)83.3/93:0.0455772096)75.8/53:0.0090580431,(((WP_190336189.1-Vibrio-sp.-S9_S30:0.1184876994,WP_022612811.1-Vibrio-nigripulchritudo:0.1428657760)90.8/100:0.1056064482,(WP_193154481.1-Vibrio-navarrensis:0.2115853600,WP_148560072.1-Vibrio-mimicus:0.2739832611)10.7/34:0.0613256311)15.1/49:0.0225570463,WP_180567593.1-Oceanospirillales:0.2097524100)86.4/50:0.0509284034)84.6/72:0.0203247063,((((((WP_096629284.1-Salinivibrio-sp.-YCSC6:0.2636070629,WP_141346804.1-Vibrio-inusitatus:0.1856341439)45.2/74:0.0233053910,WP_102282524.1-Vibrio-cyclitrophicus:0.2173731560)85.9/59:0.0519500570,((WP_102546929.1-Vibrio-breoganii:0.2117390392,WP_102461639.1-Vibrio-breoganii:0.1701070675)81.4/98:0.0574161444,WP_135455667.1-Vibrio-echinoideorum:0.1604525097)0/35:0.0000026059)93.3/41:0.0793024679,(((WP_102455158.1-Vibrio-breoganii:0.1785337919,(WP_188011373.1-Vibrio-vulnificus:0.2702602753,WP_102451772.1-Vibrio-breoganii:0.1529677397)90.5/99:0.1180714727)38.4/25:0.0756692478,WP_102446429.1-Vibrio-breoganii:0.1817533194)65.7/21:0.0361825254,((WP_070968050.1-Vibrio-sonorensis:0.2732928921,(WP_261835819.1-Vibrio-ishigakensis:0.1674468140,WP_017221048.1-Moritella-dasanensis:0.1578493746)69.3/99:0.0670269959)3.1/64:0.0094584122,WP_102329601.1-Vibrio-breoganii:0.2764515641)71.7/18:0.0190065763)84.3/33:0.0496675881)77.9/47:0.0181420111,(WP_045977433.1-Vibrio-neptunius:0.2845600676,(WP_123015137.1-Vibrio-zhugei:0.0543805298,WP_077313846.1-Vibrio-palustris:0.0258117084)97.1/100:0.1627849695)85.8/87:0.0631427424)77.6/49:0.0186700737,WP_218382835.1-Photobacterium-sp.-WH24:0.2601216682)63.5/45:0.0112770092)83.4/52:0.0237690582,(((WP_046223474.1-Vibrio-sp.-ECSMB14106:0.1136834394,WP_102562373.1-Vibrio-breoganii:0.1663079851)38.6/98:0.0371727641,(WP_102481006.1-Vibrio-breoganii:0.1519381669,WP_138522257.1-Pseudoalteromonas-galatheae:0.3860321646)74.5/77:0.0607824652)29.3/74:0.0237713483,(WP_258489575.1-Vibrio-alginolyticus:0.2130524059,WP_261874894.1-Vibrio-rarus:0.1928734042)70.6/73:0.0269413511)93.1/95:0.0566721826)72.9/53:0.0121200715,(((WP_038175370.1-Vibrio-pacinii:0.1788454228,((((WP_047883806.1-Photobacterium-ganghwense:0.1605827879,WP_146157089.1-Photobacterium-rosenbergii:0.2171573312)82.8/53:0.0493472264,(WP_187463805.1-Photobacterium-sp.-BZF1:0.0592141057,WP_039467548.1-Photobacterium-gaetbulicola:0.1944983622)91.4/54:0.0808395405)87.8/100:0.0708704061,WP_239411815.1-Photobacterium-sp.-OFAV2-7:0.1991372221)29.7/100:0.0577462580,WP_075764060.1-Photobacterium-proteolyticum:0.1728716896)90.1/100:0.0529773663)0/94:0.0000020754,WP_171749982.1-Vibrio-sp.-RE88:0.1927708294)81.8/99:0.0637565745,WP_020332409.1-Vibrio-fluvialis:0.1194555925)93.2/78:0.0795357237)86.3/89:0.0336988564,((((WP_221070344.1-Vibrio-alfacsensis:0.1270716460,WP_011189464.1-Desulfotalea-psychrophila:0.5344820157)79.6/93:0.0766612770,WP_102533425.1-Vibrio-breoganii:0.1773111284)81.4/90:0.0426226852,(WP_124938567.1-Vibrio-viridaestus:0.1398200339,WP_088133934.1-Vibrio-gazogenes:0.2740457875)88.5/100:0.0559658508)78.2/87:0.0245893228,(WP_102941572.1-Vibrio-diazotrophicus:0.1699099324,WP_261894433.1-Vibrio-porteresiae:0.2453464207)83/90:0.0444081327)59.3/77:0.0066594838)86.7/94:0.0454852649,WP_193290728.1-Vibrio-parahaemolyticus:0.3216416151)84.9/98:0.0613475465,(((WP_007466453.1-Photobacterium-marinum:0.3833697115,WP_042868619.1-Dickeya:0.0963061004)47.8/89:0.0307383629,(WP_035339894.1-Dickeya:0.0351206274,WP_125258312.1-Dickeya-lacustris:0.0972564384)100/100:0.2598554208)83/93:0.0623652242,(WP_050928752.1-Yersinia-enterocolitica:0.3326222613,WP_012883389.1-Dickeya-parazeae:0.2736671259)87.7/96:0.0951335789)83.3/94:0.0485304574)81.1/97:0.0479037326,WP_133040538.1-Shewanella-fodinae:0.4091896557)54/99:0.0424224750,WP_076542412.1-Shewanella-sp.-UCD-KL21:0.5129470603)93.4/92:0.0873877638,((WP_162556830.1-Proteus:0.0984274864,WP_161779122.1-Proteus-sp.-G2675:0.1065778733)99.3/100:0.2434097139,WP_236366479.1-Aeromonas-veronii:0.2917243010)75.8/97:0.0508276569)83.1/88:0.0426548467)93.3/99:0.0997084099,(WP_097364011.1-Edwardsiella-tarda:0.3953963988,(WP_086868392.1-Kosakonia-pseudosacchari:0.7843916649,WP_153752706.1-unclassified-Citrobacter:0.5849147549)92.9/93:0.2342793876)77.5/82:0.0734073995)80.4/75:0.0432001723,WP_102949229.1-Aeromonas-caviae:0.4318738383)0/8:0.0000025190,WP_225633057.1-Aeromonas-enteropelogenes:0.2534117416)88.8/98:0.0765931576,WP_043158915.1-Aeromonas-hydrophila:0.6631907917)10/45:0.0142484231,(WP_168756429.1-Aeromonas-hydrophila:0.6012827575,WP_061279316.1-Kluyvera-cryocrescens:0.3902985963)90.2/98:0.1229269348)90/94:0.0829799947,(WP_100646402.1-Aeromonas-veronii:0.0773236100,((WP_252601295.1-Aeromonas-dhakensis:0.0258720018,WP_139432874.1-Aeromonas-dhakensis:0.1761322214)87.5/100:0.0418869047,WP_106553824.1-Aeromonas-hydrophila:0.0733006012)78.5/100:0.0272371250)99.9/100:0.2412011031)69.7/23:0.0540612841,WP_224211331.1-Aeromonas-enteropelogenes:0.3409918440)66.2/14:0.0137542870,(WP_253626250.1-Aeromonas-hydrophila:0.1210270799,WP_082041448.1-Aeromonas-eucrenophila:0.4124826445)55.6/14:0.0731739395)74.6/14:0.0156453664,(((((((((WP_234664799.1-Aeromonas-allosaccharophila:0.1846230290,(WP_257692500.1-Aeromonas-caviae:0.2011961004,WP_113719982.1-Aeromonas-salmonicida:0.1919164398)6.2/90:0.0479108861)14.7/87:0.0369207385,WP_069554789.1-Aeromonas-sp.-DNP9:0.1925239172)29.9/97:0.0665821969,(WP_213320225.1-Aeromonas-caviae:0.0665022619,WP_203764723.1-Aeromonas-caviae:0.1190753477)97.3/100:0.1800390708)81.4/75:0.0537567216,((WP_050666161.1-Aeromonas-schubertii:0.0733607346,WP_059168611.1-Aeromonas-salmonicida:0.1522913114)50.1/78:0.0262868691,WP_210600017.1-Aeromonas-sp.-MrichA-1:0.0655872950)84.4/89:0.0453877314)81.2/56:0.0263182339,((WP_245131329.1-Aeromonas-veronii:0.1943280362,WP_225159845.1-Aeromonas-caviae:0.1670426532)3.9/18:0.0323524655,(WP_163135231.1-Aeromonas-rivipollensis:0.0373031934,WP_241279715.1-Aeromonas-sp.-MR19:0.0370472441)100/100:0.3002679048)77.2/51:0.0293935116)54.6/42:0.0462277750,((WP_101347551.1-Aeromonas-sobria:0.0786592693,WP_201955411.1-Aeromonas-veronii:0.0782323935)95.2/100:0.0945953441,(WP_136501435.1-Aeromonas-veronii:0.2102460073,(WP_268029297.1-Aeromonas-hydrophila:0.1385103160,WP_201942265.1-Aeromonas-veronii:0.1754467459)97.4/100:0.1469902699)80.7/98:0.0553312454)89/83:0.0620304683)68.5/85:0.0220102936,((((WP_148612677.1-Aeromonas-sobria:0.0462431511,((WP_201355655.1-Aeromonas-hydrophila:0.0562183428,WP_201951905.1-Aeromonas-veronii:0.0397848407)76.2/81:0.0077143027,WP_171268810.1-Aeromonas-media:0.0332207204)78.1/57:0.0184329765)99.9/100:0.1922346015,WP_042063659.1-Aeromonas-allosaccharophila:0.0367582785)0/44:0.0099547638,WP_256684552.1-Aeromonas-salmonicida:0.0408782975)76.5/76:0.0214378187,(((WP_212539243.1-Aeromonas-salmonicida:0.0394458819,((WP_187590672.1-Aeromonas-hydrophila:0.1564037987,WP_017782281.1-Aeromonas-dhakensis:0.0345868971)60.3/100:0.0186522710,WP_060586794.1-Aeromonas-schubertii:0.0619278319)99.4/100:0.1127205781)90.1/100:0.0358976945,WP_201904077.1-Aeromonas-veronii:0.0725713884)79.5/100:0.0241569647,WP_206874356.1-Aeromonas-veronii:0.0128472989)79.3/91:0.0182433650)99.4/100:0.1400125913)0/14:0.0284625909,(WP_042649704.1-Aeromonas-media:0.1312362375,WP_139406182.1-Aeromonas-veronii:0.3373826977)82.1/87:0.0572217538)76.3/53:0.0246834492,(((WP_268421127.1-Aeromonas-media:0.0850637928,WP_111910157.1-Aeromonas:0.0258263699)77/98:0.0173589419,WP_128343177.1-Aeromonas-caviae:0.0592839711)94.2/100:0.0680948892,(WP_268037199.1-Aeromonas-dhakensis:0.1306107113,(WP_139733908.1-Aeromonas-veronii:0.0208883064,((WP_234579942.1-Aeromonas-salmonicida:0.0882109112,(WP_234676856.1-Aeromonas-rivipollensis:0.0188082091,WP_269382839.1-Aeromonas-hydrophila:0.1002020454)44/100:0.0095846869)65.9/92:0.0118931620,WP_222134508.1-Vibrio-cholerae:0.1892238190)56.2/91:0.0074608697)97.4/98:0.0911148620)49.1/92:0.0281535376)86.7/98:0.0549791304)74.5/34:0.0247506928)38.8/21:0.0282091936,((WP_216990617.1-Aeromonas-sp.-FDAARGOS-1409:0.1582365811,((((((WP_059168599.1-Aeromonas-salmonicida:0.0700845882,((WP_033115905.1-Aeromonas-lacus:0.0548626565,WP_200561028.1-unclassified-Erwinia:0.5389128103)0/36:0.0000028228,((WP_223920301.1-Aeromonas-caviae:0.0447296525,WP_226058924.1-Shewanella-sp.-SR1:0.1526827554)77.8/98:0.0215987285,(WP_058394234.1-Aeromonas-salmonicida:0.0379994611,WP_257690725.1-Aeromonas-veronii:0.0602487242)76.6/100:0.0235194728)88.2/97:0.0253624671)89.7/41:0.0553532730)62.8/39:0.0527873640,((WP_268010485.1-Aeromonas-dhakensis:0.0959404584,WP_098985072.1-Aeromonas-sp.-CA23:0.0514434525)97.9/100:0.1606196472,(WP_181268965.1-Aeromonas-veronii:0.0391600835,(WP_129146711.1-Aeromonas-caviae:0.0285968979,(WP_081843667.1-Aeromonas-taiwanensis:0.0226882003,WP_139744464.1-Aeromonas-media:0.0503319875)87.2/100:0.0255693637)86/100:0.0244367526)70/100:0.0773015850)100/100:0.2512597490)89/100:0.0618646820,((WP_223945983.1-Aeromonas-caviae:0.1474531475,(((WP_139750984.1-Aeromonas-media:0.0341355185,(WP_224430864.1-Aeromonas-rivuli:0.0283701863,WP_041996113.1-Aeromonas-bivalvium:0.0911293831)41.5/26:0.0079168465)98.2/100:0.0816534953,WP_163128050.1-Aeromonas-allosaccharophila:0.0822598363)41.3/27:0.0292317489,((WP_265458754.1-Aeromonas-salmonicida:0.0978092367,WP_125597700.1-Aeromonas-salmonicida:0.0278653975)33.4/100:0.0025241086,(WP_180911080.1-Aeromonas-caviae:0.0479559573,WP_264818242.1-Aeromonas-caviae:0.0318819942)91.1/100:0.0243582554)0/95:0.0000022898)80.1/26:0.0129821381)69.3/19:0.0065936258,(WP_040098646.1-Aeromonas:0.1152762729,((WP_263309349.1-Aeromonas-caviae:0.0494914429,(WP_041217344.1-Aeromonas-hydrophila:0.0500797265,WP_255912544.1-Aeromonas-sp.-JL9:0.0403844026)75.8/95:0.0083620657)75/95:0.0074642541,WP_257686696.1-Aeromonas-veronii:0.0514162009)71.6/100:0.0131193915)75.3/99:0.0163718436)83.2/59:0.0336125033)77.3/87:0.0272148964,WP_265776268.1-Aeromonas-veronii:0.1479060836)85.3/86:0.0633075737,(((WP_201942221.1-Aeromonas-veronii:0.2034714869,(((WP_268854930.1-Aeromonas-enteropelogenes:0.1492810085,(WP_203131090.1-Aeromonas-jandaei:0.0735702179,((WP_042657830.1-Aeromonas:0.0546290341,WP_111901368.1-Aeromonas-veronii:0.0471068500)66/79:0.0215469636,WP_139692403.1-Aeromonas-sobria:0.0382051364)0/75:0.0000021431)75.9/77:0.0078809697)0/18:0.0131435747,WP_223380193.1-Aeromonas-enteropelogenes:0.0509020277)0/27:0.0000028043,WP_159143398.1-Aeromonas-veronii:0.0307545863)21.4/28:0.0321620548)89.9/98:0.0763389147,(WP_236321701.1-Aeromonas-veronii:0.0920978663,WP_257119864.1-Aeromonas-hydrophila:0.0210172548)90.2/63:0.0651985819)7.5/76:0.0230942449,(WP_257707542.1-Aeromonas-caviae:0.0687398220,WP_182951616.1-Aeromonas-sp.-WP2-W18-CRE-05:0.0751893571)98.9/100:0.1295030762)85/62:0.0676381230)0/38:0.0053081026,WP_236563281.1-Aeromonas-veronii:0.2012455584)16.7/34:0.0253309732)90.7/58:0.0640547378,WP_265468300.1-Aeromonas-salmonicida:0.1657854929)87.6/32:0.0554927339)88.1/65:0.0368752339)76.1/51:0.0271589184,(((WP_047682696.1-Xenorhabdus:0.0688527351,WP_099124588.1-Xenorhabdus-stockiae:0.0992863724)98.4/100:0.1767588995,(WP_074024273.1-Xenorhabdus-eapokensis:0.0828662649,WP_217470569.1-Photorhabdus-akhurstii:0.0955129698)88.7/100:0.0885763438)81.6/100:0.0549103966,(WP_048221111.1-Citrobacter:0.1715460441,WP_263807164.1-Salmonella-enterica:0.2250142067)99.8/100:0.2981147851)61.6/34:0.0847954118)22.7/22:0.0427223265,((WP_151054911.1-Cellvibrio-sp.-KY-YJ-3:0.2583628068,WP_078044787.1-Cellvibrio-sp.-PSBB023:0.2670321675)75.5/86:0.0316031506,(WP_012489013.1-Cellvibrio-japonicus:0.1842847318,WP_087465811.1-Cellvibrio-sp.-PSBB006:0.2483966986)86.9/86:0.0510475251)98.9/100:0.1892021546)92.6/92:0.0733254486,((WP_103946175.1-Lelliottia-aquatilis:0.1165031665,WP_123427447.1-Lelliottia:0.0911434664)33.8/87:0.0150601633,WP_235147774.1-Enterobacter-cloacae:0.3433764199)96.2/97:0.1046722396)92.9/86:0.0668720422,((WP_001100804.1-Enterobacteriaceae:0.1507506060,(WP_213789144.1-Enterobacter-hormaechei:0.0216060744,((WP_148242178.1-Enterobacter-asburiae:0.0610028165,WP_237994110.1-Enterobacter-sp.-DRP3:0.0228892169)56.7/95:0.0061478958,WP_237424876.1-Enterobacter-cloacae-complex-sp.-ECC445:0.0560353764)84.7/95:0.0434532530)100/100:0.3447905815)91.6/66:0.1015315633,WP_001100786.1-Escherichia-coli:0.1663293987)83.2/26:0.0425932922)0/16:0.0106893219,((WP_137650031.1-Escherichia-albertii:0.1436191781,WP_073446180.1-Escherichia-coli:0.2929842204)90.2/62:0.0550452615,((((WP_214593796.1-Enterobacter-mori:0.0572726069,WP_086532312.1-Enterobacter-ludwigii:0.0857558387)95.3/100:0.1317258884,((WP_047742849.1-Enterobacter:0.0998242723,WP_040022520.1-Enterobacter-cloacae:0.1071183030)97.3/100:0.1465159042,(WP_129995405.1-Enterobacter-cloacae:0.0728343381,WP_042099316.1-Escherichia-coli:0.0989924697)78.3/100:0.0235777194)93.7/100:0.1288196256)25.9/96:0.0517562113,WP_045339085.1-Enterobacter-cloacae-complex:0.2073607257)88.6/98:0.0705217368,WP_153879942.1-Citrobacter-freundii:0.2355525687)77.8/84:0.0326756747)68.3/15:0.0093039829)88.8/75:0.0387263537,WP_202667543.1-Lelliottia-amnigena:0.3068702347)24.7/41:0.0089702374,((((WP_187216555.1-Escherichia-coli:0.2214491439,WP_059222105.1-Escherichia-albertii:0.1547674268)82.8/99:0.0499521458,WP_262938691.1-Escherichia-albertii:0.1406318126)0/13:0.0000026337,(WP_137653986.1-Escherichia-albertii:0.0228558009,WP_001100785.1-Escherichia-coli:0.0694634266)98.6/100:0.1095802341)55.7/25:0.0242364951,WP_021577651.1-Escherichia-coli:0.2015219056)83.1/25:0.0316613567)22.5/15:0.0134285037,(((WP_257124601.1-Aeromonas-veronii:0.6483854127,WP_169342312.1-Providencia-stuartii:0.2200222616)70.5/50:0.0312451105,(WP_154625743.1-Providencia-stuartii:0.1768758858,WP_004924376.1-Providencia-stuartii:0.1988012667)69.1/62:0.0212857009)85.4/49:0.0574954942,((((((WP_159241675.1-Proteus-terrae:0.2200657023,WP_061356323.1-Escherichia-coli:0.9569894296)64.1/46:0.0472278954,WP_195697765.1-Providencia-rettgeri:0.2587302634)90.6/46:0.0820505560,WP_238563424.1-Providencia-rettgeri:0.1580177205)72.6/44:0.0165813720,WP_165880752.1-Providencia-rettgeri:0.2390382911)85.9/44:0.0414541262,WP_231068620.1-Providencia-huaxiensis:0.1895340247)85.3/56:0.0351669085,((WP_187128479.1-Providencia-sp.-JUb39:0.1996149238,WP_193115284.1-Escherichia-coli:0.1932931232)91.1/100:0.0850808962,WP_164565644.1-Providencia-rettgeri:0.3472930242)68.7/64:0.0201377460)71.4/47:0.0190153273)99.9/93:0.2036808495)87.5/24:0.0281987219,((((((((((((((WP_085037002.1-Cronobacter-sakazakii:0.2683659550,WP_076770125.1-Kosakonia-cowanii:0.1885743896)97/100:0.2143610554,WP_216981205.1-Lelliottia-amnigena:0.3748997918)78.6/100:0.0800797784,((WP_227534785.1-Erwinia-rhapontici:0.1767690366,WP_017800228.1-Erwinia-toletana:0.2005047786)88.5/100:0.1041544714,WP_255851625.1-Erwinia-persicina:0.2554770546)72.5/100:0.1066001879)86.3/100:0.0642595927,(WP_089585421.1-Escherichia-coli:0.0611885867,WP_104705897.1-Escherichia-coli:0.0292059021)97.6/100:0.1499781686)83.4/99:0.0874698608,(WP_001610415.1-Escherichia-coli:0.0522490956,WP_138183825.1-Escherichia-sp.-E4385:0.0266926516)99.7/100:0.2480698458)88/98:0.0799959800,(WP_016236761.1-Enterobacteriaceae:0.1194536726,WP_024239504.1-Escherichia-coli:0.1576779713)92.2/100:0.1102430020)39.8/73:0.0665512706,((WP_107192407.1-Escherichia-albertii:0.0227762365,WP_181673375.1-Escherichia-fergusonii:0.0557970759)99/100:0.1869431128,((WP_032214194.1-Escherichia-coli:0.0592570431,(WP_059217428.1-Escherichia-albertii:0.0428733591,(WP_073278747.1-Escherichia-coli:0.1098308996,WP_087893724.1-Enterobacteriaceae:0.0309756880)99.9/100:0.1864390602)75/95:0.0147667492)79.7/94:0.0346259058,WP_085459683.1-Escherichia-coli:0.0871345654)96.6/97:0.1537078149)92.4/95:0.1089421425)51.6/23:0.0118580103,(WP_138098269.1-Jejubacter-calystegiae:0.2361798125,((WP_179154141.1-Citrobacter-freundii:0.3629959608,WP_086539147.1-Citrobacter-freundii:0.2791650546)51.3/85:0.0812639580,WP_033870520.1-Enterobacteriaceae:0.2963431101)77.8/59:0.0303991621)80.6/56:0.0237667273)44.6/32:0.0265922738,((WP_157959988.1-Citrobacter:0.5866775476,WP_267387874.1-Citrobacter-portucalensis:0.3275410517)85.8/87:0.0709043553,(WP_095844187.1-Escherichia-coli:0.0794328683,(WP_086118650.1-Escherichia-coli:0.0328143447,WP_089538038.1-Escherichia-coli:0.0427409640)64.3/98:0.0086740279)100/100:0.2404636736)0/54:0.0000025987)90.1/93:0.0418365266,(WP_248249449.1-Enterobacter-roggenkampii:0.2358727864,WP_032936727.1-Citrobacter-freundii:0.1363706412)99.4/100:0.2671436110)85.7/86:0.0372736956,((WP_100272167.1-Citrobacter-portucalensis:0.2975237438,(WP_181624464.1-Citrobacter-freundii:0.2889809812,WP_001515521.1-Escherichia-coli:0.2150079548)82.3/100:0.0582916056)90.2/97:0.0653754455,((((WP_058662478.1-Enterobacteriaceae:0.0547195979,WP_061709645.1-Pseudenterobacter-timonensis:0.0578237675)99.9/100:0.2454516148,WP_112164393.1-Rahnella-inusitata:0.1889567846)84.9/100:0.0701471389,WP_047609274.1-Rahnella-aquatilis:0.2391869153)89/97:0.0675041641,WP_157953065.1-Limnobaculum-parvum:0.2624239477)82.9/94:0.0404895755)94.4/94:0.0790221902)80.8/57:0.0248636370,(((WP_119938542.1-Enterobacter-huaxiensis:0.1275752403,WP_050871166.1-Enterobacter-cloacae-complex:0.0490246768)83.3/100:0.0365961549,(WP_167816599.1-Enterobacter-cloacae:0.3392782107,(WP_262183194.1-Enterobacter-mori:0.1751187851,(WP_063412371.1-Enterobacter-cloacae:0.0487889852,WP_047027451.1-Enterobacter-kobei:0.0644482383)97.1/100:0.1475011984)96.8/100:0.1947746112)97.2/94:0.1490397468)86.5/89:0.0298733909,(WP_228937168.1-Escherichia-marmotae:0.0414010299,WP_040079227.1-Escherichia-coli:0.0464550802)82.6/100:0.0183196246)94.4/89:0.0589497770)68.9/39:0.0124314757,((WP_001297344.1-Escherichia-coli:0.2435122185,WP_072664248.1-Escherichia:0.1690841423)87.5/84:0.0739804330,(WP_100029735.1-Escherichia-coli:0.2367255268,(((WP_021540003.1-Escherichia-coli:0.0000022314,WP_047657564.1-Enterobacteriaceae:0.0768703019)98.9/100:0.1176381203,((((WP_035600513.1-Edwardsiella-tarda:0.3774131904,(WP_106902182.1-Citrobacter-freundii:0.2741637962,(WP_126291239.1-Enterobacter-hormaechei:0.2022640546,((((WP_062932819.1-Enterobacter-hormaechei:0.0388269440,WP_252042312.1-Enterobacter-chengduensis:0.0510112040)78.1/100:0.0318836161,WP_048999124.1-Enterobacter:0.0486956771)73.7/99:0.0100058946,WP_063144508.1-Enterobacter-cloacae:0.0451440062)85.2/99:0.0298259748,WP_049008289.1-Enterobacter-cloacae:0.0557090002)80.9/100:0.0599801365)92.6/100:0.1065831726)92.9/100:0.1015030440)83.3/70:0.0618875999,(WP_202782010.1-Enterobacter-asburiae:0.0878521639,WP_144817260.1-Pseudescherichia-sp.-L3:0.0576482943)100/100:0.2849439331)75.9/22:0.0551024638,WP_269535377.1-Citrobacter-sedlakii:0.3014952399)62.8/15:0.0422664971,((WP_105274811.1-Escherichia:0.2747969984,WP_064512195.1-Buttiauxella-gaviniae:0.3726956481)86.6/98:0.0902368206,WP_172410167.1-Escherichia-coli:0.3924808327)90.2/98:0.1081447727)75.5/13:0.0135987779)75.8/13:0.0162376235,WP_159282060.1-Rahnella-variigena:0.5401421068)46.5/17:0.0160592167)0/12:0.0144139024)80.7/16:0.0236076564)72.7/16:0.0099843824,((((((((WP_172912938.1-Pantoea-stewartii:0.1531190535,WP_145394131.1-Pantoea-sp.-SJZ147:0.1644333305)99.9/100:0.4036760150,WP_039304623.1-Cedecea-neteri:0.3788770414)91.6/81:0.1550765178,((WP_038258850.1-Xenorhabdus-bovienii:0.3401340880,(WP_038223662.1-Xenorhabdus-bovienii:0.2451000505,(WP_161748440.1-Proteus-sp.-G2658:0.2440412777,(WP_049199273.1-Proteus-mirabilis:0.0810117908,WP_157954980.1-Proteus-terrae:0.1269039693)37.6/82:0.0896155760)95.4/100:0.1349793881)72.9/88:0.0718954820)64.2/75:0.0499780593,WP_086113286.1-Xenorhabdus-beddingii:0.3293901284)75.8/76:0.0320139524)86.2/75:0.0546339309,(((((WP_151711904.1-Enterobacter-ludwigii:0.0414560539,(WP_032941356.1-Citrobacter-freundii:0.2618606534,WP_058678686.1-Enterobacter:0.0982932310)89.7/98:0.0763971946)83.6/98:0.0838486415,WP_223560970.1-unclassified-Enterobacter:0.0000026048)98.3/99:0.3599733238,WP_044720521.1-Escherichia-coli:0.4905769183)98.4/100:0.3622606842,WP_064374050.1-Pantoea:0.3209037312)90.1/79:0.1259084887,(((WP_234505032.1-Kluyvera-intermedia:0.5386477564,WP_230863267.1-Enterobacter-cloacae:0.5660274342)88.1/95:0.1217911549,(WP_029399168.1-Escherichia-coli:0.4316054190,((WP_034496584.1-Buttiauxella-agrestis:0.4486376745,WP_164716793.1-Cedecea-lapagei:0.4772894572)86.2/100:0.1258679798,WP_050862535.1-Enterobacter:0.5576417864)68.1/99:0.0299513019)86/99:0.1219285300)88.9/96:0.1001415587,(WP_036421240.1-Morganella-morganii:0.1811265255,(WP_264998074.1-Morganella-morganii:0.1057071279,WP_186849509.1-Morganella-morganii:0.1058611415)94.8/100:0.1195747952)91.9/100:0.1050968370)61.1/40:0.0575934187)83/25:0.0414822107)86.6/22:0.0473107630,((((WP_090123257.1-Kosakonia-arachidis:0.1042132768,WP_020454293.1-Kosakonia-sp.-SOY2:0.1282822273)88.3/100:0.0533183347,WP_017458471.1-Kosakonia-sacchari:0.1916736279)80.9/100:0.0556445446,(WP_200002836.1-Citrobacter-freundii:0.2220600307,WP_080329661.1-Enterobacter-cloacae-complex:0.1805208172)99/100:0.2270532687)86.4/100:0.1126270239,WP_090460354.1-Enterobacter-sp.-kpr-6:0.0932823754)99.6/100:0.2767583877)44.1/49:0.0144076559,(WP_194857598.1-Citrobacter-freundii:0.4406973844,((WP_141032926.1-Escherichia-coli:0.1762817508,WP_196390204.1-Enterobacter-mori:0.0544522136)78.9/96:0.0436642028,(WP_252032207.1-Enterobacter-asburiae:0.0632145116,WP_111966211.1-Enterobacter-cloacae-complex:0.0331599041)71.6/96:0.0305446424)94.2/96:0.1832547145)99.3/100:0.3197039120)75.6/79:0.0330568839,(((WP_115348269.1-Hafnia-alvei:0.1710498730,(WP_046458830.1-Obesumbacterium-proteus:0.3536527091,WP_125117341.1-Aeromonas-caviae:0.4414976029)30.5/33:0.0543917052)32.1/30:0.0471870715,(WP_130961887.1-Hafnia-paralvei:0.2466306965,WP_061059890.1-Hafnia-paralvei:0.2469171070)79.7/82:0.0413557115)32.9/29:0.0427534208,(WP_120162698.1-Rahnella-variigena:0.0778970979,WP_183847446.1-Rahnella-inusitata:0.0798168252)100/100:0.4369246623)93.4/58:0.1145170958)90.6/18:0.0608963825,((((((WP_097405671.1-Kosakonia:0.0541149197,WP_161518249.1-Kosakonia-sp.-MH5:0.0846192041)90.4/100:0.0747139918,WP_128482629.1-Kosakonia-cowanii:0.1441469705)97.7/100:0.1915185038,WP_206164314.1-Citrobacter:0.2399974536)92.3/100:0.0997371603,WP_171149131.1-Erwinia-sp.-JH02:0.2500763297)73.2/78:0.0586846451,((((WP_238085128.1-Pseudescherichia-vulneris:0.0290354585,WP_032652659.1-Enterobacter-cloacae-complex:0.1208436837)9.7/72:0.0116944367,WP_129363899.1-Enterobacter-cloacae:0.0399152840)64.9/78:0.0203386275,(WP_257661844.1-Citrobacter-portucalensis:0.1551420129,WP_193134185.1-Citrobacter-freundii:0.1030435349)85.7/100:0.0362698270)93.3/89:0.1213389281,(WP_042284276.1-Citrobacter-sedlakii:0.2898742199,(WP_024193259.1-Escherichia-coli:0.1320954386,WP_099433281.1-Citrobacter-amalonaticus:0.0380060083)82.5/100:0.1085399366)94/100:0.1115312512)97.3/98:0.1957090154)86.3/51:0.0766749827,(((((WP_167863639.1-Erwinia-rhapontici:0.1868831424,WP_039385357.1-Pantoea-eucrina:0.2454358992)89.1/99:0.1080345789,WP_118663751.1-Erwinia-persicina:0.1732399196)76.1/99:0.0823773354,WP_062742814.1-Erwinia-persicina:0.2067164646)89/97:0.0850397916,WP_191149682.1-Erwinia-aphidicola:0.1667730067)79.7/92:0.0421046758,((WP_070133611.1-Candidatus-Erwinia-dacicola:0.2648424467,WP_140919905.1-Insectihabitans-xujianqingii:0.2761516004)61/80:0.0201693788,WP_029881813.1-Enterobacter:0.4154521476)84.3/100:0.0518415653)10.5/67:0.0208115287)52.8/84:0.0329699866)84.1/18:0.0232674643)72.9/24:0.0085195264)83.4/18:0.0166807761,(((((((((((WP_108569539.1-unclassified-Pantoea:0.2262603780,WP_028018520.1-Enterobacter-cloacae-complex:0.9266018250)97.3/98:0.3074300312,WP_250341216.1-Pantoea-eucrina:0.0980918367)98.3/96:0.2224478854,(((WP_049002539.1-Citrobacter-freundii:0.1621517542,(WP_249576357.1-Enterobacter-vonholyi:0.0779528609,WP_161617217.1-Yokenella-regensburgei:0.0542735422)95.1/100:0.0720852750)83.6/100:0.0336263279,((WP_079834758.1-Salmonella-enterica:0.0000026656,(WP_079793206.1-Salmonella-enterica:0.1583993736,WP_001100807.1-Salmonella:0.0948182527)99.4/100:0.1989289086)77.4/99:0.0517838640,WP_058109100.1-Salmonella-enterica:0.0373699874)99.2/99:0.1170954785)89.8/99:0.0509804422,WP_032303308.1-Escherichia:0.0873926002)87.9/99:0.0598547796)75/90:0.0323306598,(WP_127791218.1-Escherichia-coli:0.1698541707,(((WP_174633339.1-Escherichia-coli:0.1447975563,WP_025210617.1-Escherichia-coli:0.1114268081)76.4/90:0.0661771778,WP_001527244.1-Escherichia-coli:0.0971273382)52.8/28:0.0192464431,(WP_085454021.1-Escherichia-coli:0.1417663909,(WP_088541377.1-Escherichia-coli:0.0434581368,WP_222691870.1-Escherichia-marmotae:0.0683152822)92.1/98:0.0586489746)83.8/83:0.0427819466)34.9/24:0.0343193011)76/89:0.0291963871)88.8/67:0.0381232524,((((((WP_171269898.1-Aeromonas-media:0.1221926015,WP_216983019.1-Aeromonas-sp.-FDAARGOS-1415:0.1156063502)98.6/100:0.1688593874,(((((((WP_064338398.1-Aeromonas-veronii:0.1728466562,WP_252995952.1-Aeromonas-encheleia:0.3113608312)84/84:0.0586516278,(((((WP_247163170.1-Escherichia-coli:0.0178557455,WP_038644959.1-Pantoea:0.1042188181)99.3/100:0.1241723198,((WP_252422582.1-Aeromonas-taiwanensis:0.0098769532,WP_263719954.1-Aeromonas-caviae:0.1170560379)74.5/96:0.0093225278,(WP_244056308.1-Aeromonas-caviae:0.0712851324,WP_202046159.1-Aeromonas-media:0.2311849151)98.3/96:0.1525950679)91.3/86:0.0500097867)89.8/88:0.0474662111,WP_033131297.1-Aeromonas-aquatica:0.0263010516)97.8/87:0.0556596813,WP_241344952.1-Aeromonas-sp.-MR16:0.0166517148)80.4/87:0.0159686576,(WP_160838676.1-Aeromonas-caviae:0.0987492112,((WP_049048056.1-Aeromonas-hydrophila:0.1844388935,WP_167569591.1-Aeromonas-veronii:0.1374624319)18.8/50:0.0458643397,(((WP_116547128.1-Plesiomonas-shigelloides:0.1221840625,WP_201996661.1-Aeromonas-veronii:0.0334986282)35.6/83:0.0136403953,WP_235463836.1-Aeromonas-australiensis:0.0303761862)0/27:0.0000026421,(WP_139496062.1-Escherichia-coli:0.0818064876,WP_215269322.1-Aeromonas-dhakensis:0.0491825773)73.7/62:0.0081151271)83.2/40:0.0279948136)94.5/76:0.0642447096)75.2/53:0.0114972404)98.4/88:0.1415388095)77.7/37:0.0431395549,((WP_042864139.1-Aeromonas-caviae:0.0443745120,WP_225159881.1-Aeromonas-caviae:0.0542467668)94.3/100:0.1035822334,WP_045790994.1-Aeromonas-hydrophila:0.2121370783)69.6/84:0.0537749091)79.6/87:0.0424285297,(WP_223932214.1-Aeromonas-caviae:0.0126935348,WP_161506954.1-Aeromonas-media:0.1574727518)99.8/100:0.1793061813)66.9/94:0.0556446412,WP_210550341.1-Aeromonas-sp.-SrichE-2G:0.3872137690)42.1/47:0.0356283612,(WP_227095090.1-Aeromonas-veronii:0.1202456503,WP_152109808.1-Plesiomonas-shigelloides:0.0732857682)92.2/100:0.0953553892)91.1/93:0.0713612599,WP_168366848.1-Dickeya-zeae:0.1850854957)88.6/93:0.0641587882)87/100:0.0533691307,WP_075180894.1-Pantoea-sp.-1.19:0.2280284918)68.8/98:0.0654616460,(WP_034458444.1-Buttiauxella-noackiae:0.2145447893,WP_253383865.1-unidentified-bacterial-endosymbiont:0.3636216327)0/18:0.0542033269)74.3/98:0.0284232213,(WP_105591414.1-Pantoea-coffeiphila:0.0814447723,WP_123334936.1-Erwinia-sp.-JUb26:0.0491801431)100/100:0.2565408359)94.4/99:0.0783179824,((((WP_259047621.1-Klebsiella-sp.-BIGb0407:0.2324405363,WP_105963664.1-Escherichia-coli:0.0538574398)83.4/97:0.0449301906,WP_063610377.1-Escherichia-coli:0.1063308381)38.8/26:0.0102225809,WP_053290779.1-Escherichia-coli:0.1684999013)83.2/34:0.0405657374,(WP_218070502.1-Escherichia-coli:0.0969255096,(WP_187209312.1-Escherichia-coli:0.1147355279,WP_015386517.1-Cronobacter-sakazakii:0.2608884939)70.9/33:0.0226397473)78.1/31:0.0336671499)90.5/44:0.0656442804)47.5/29:0.0202210726)75.8/23:0.0107146013,((((WP_262931739.1-Escherichia-albertii:0.0285665725,WP_160188772.1-Escherichia-coli:0.0726948137)99.1/100:0.1409474132,WP_096972986.1-Escherichia-coli:0.1456501819)74.6/83:0.0174531082,WP_097421351.1-Escherichia-coli:0.1305682636)77/82:0.0184203668,(WP_157704420.1-Escherichia-coli:0.1298795714,WP_042200020.1-Enterobacteriaceae:0.0752450125)74.2/100:0.0544463637)77.1/18:0.0200044667)78.2/13:0.0095197472,((WP_146314246.1-Escherichia-coli:0.0489148830,WP_096967326.1-Escherichia-coli:0.0252701320)98.1/100:0.1150964260,WP_181465273.1-Escherichia-fergusonii:0.0808925695)94.1/98:0.0856911485)71.8/21:0.0110857381,(((((((WP_193360946.1-Enterobacter-cloacae-complex-sp.-P39RS:0.0622590337,WP_182945676.1-Enterobacter-cloacae:0.0401444756)98.3/100:0.1259913992,((((WP_232904305.1-Buttiauxella-sp.-W03-F01:0.2638586378,WP_259388919.1-Enterobacter-huaxiensis:0.0525397251)68.8/58:0.0262194565,WP_230350752.1-Lelliottia-sp.-WAP21:0.1802229429)87.5/98:0.0455253031,WP_265185975.1-Enterobacter-mori:0.1256959252)76.8/45:0.0240188145,((WP_032636062.1-Enterobacter-kobei:0.0663234993,WP_138014015.1-Enterobacter-ludwigii:0.0882811327)89.8/99:0.0709863041,(WP_063137673.1-Enterobacter-cloacae-complex:0.0371855566,WP_165464597.1-Enterobacter-cloacae:0.0769874883)0/35:0.0215277766)89.7/99:0.0539569617)80.9/38:0.0347860153)61.9/41:0.0340435949,(((WP_096205048.1-Enterobacter:0.0429522552,WP_100165931.1-Enterobacter-mori:0.0343296325)91.7/100:0.0778613756,WP_242172213.1-Enterobacter:0.1226996847)55.4/98:0.0588708063,WP_239609625.1-Lelliottia-amnigena:0.1330634925)91.6/21:0.0689279167)75.5/18:0.0198504884,(((((WP_042716137.1-Enterobacter-sp.-B509:0.0251054580,WP_241820523.1-Enterobacter-kobei:0.0892746502)98.2/100:0.1103430110,(WP_172688853.1-Enterobacterales:0.1376677663,WP_126546226.1-Enterobacter-quasimori:0.1182416677)84.6/100:0.0452884884)89/90:0.0505395480,(WP_034915863.1-Erwinia-sp.-9145:0.2626644006,(((WP_045338355.1-Enterobacter-hormaechei:0.1703357734,(WP_182115563.1-Enterobacter-kobei:0.0453928104,WP_213381159.1-Enterobacter-cloacae:0.0780420045)88.7/100:0.0938945961)37.3/99:0.0626694373,WP_063158410.1-Enterobacter-hormaechei:0.1004947926)81.3/98:0.0377898766,WP_063869545.1-Enterobacter-hormaechei:0.0718088755)38.4/92:0.0250454982)77/84:0.0158828591)73.3/88:0.0124923349,((((((WP_141109706.1-Escherichia-albertii:0.1136752647,WP_001572827.1-Escherichia-coli:0.0725123326)12.9/93:0.0285751969,WP_203360733.1-Citrobacter-sp.-R56:0.0675178366)88/98:0.0646200936,WP_258993080.1-Scandinavium-hiltneri:0.0874148829)95.8/100:0.0966984392,(((WP_095374511.1-Escherichia-coli:0.0119460621,WP_012906377.1-Citrobacter-rodentium:0.1149849278)98.9/100:0.1087329269,WP_096122580.1-Escherichia-coli:0.1075588947)49.7/97:0.0310783419,WP_162379936.1-Citrobacter-sp.-NCU1:0.2047151747)84.5/98:0.0539784761)82.3/94:0.0405421583,WP_046888360.1-Enterobacter-cloacae:0.0762066433)79.3/92:0.0299719249,WP_126315015.1-Enterobacter-hormaechei:0.1327059876)78.2/86:0.0142225652)83.6/83:0.0205524590,WP_033487312.1-Enterobacter-cloacae-complex:0.0984049260)76.7/47:0.0197559312)83.1/20:0.0232433042,WP_208283396.1-Enterobacter-sichuanensis:0.1305243503)74.9/79:0.0139482802,WP_259000881.1-Scandinavium-goeteborgense:0.2365127718)78.9/75:0.0175230952,(WP_175320495.1-Hafnia-paralvei:0.1307498872,WP_130996599.1-Hafnia-paralvei:0.2242918705)99.8/97:0.2322080307)79.9/72:0.0164210746)0/20:0.0000022115,((((WP_109548670.1-Escherichia-coli:0.0645239291,WP_001100790.1-Escherichia:0.0195610881)95.9/100:0.0841693986,WP_181237856.1-Escherichia-marmotae:0.1118896434)70.8/52:0.0167814271,WP_107179425.1-Escherichia:0.1278877748)52.5/23:0.0189813911,(WP_061892435.1-Escherichia-coli:0.1620872289,WP_001100798.1-Enterobacteriaceae:0.1426726779)91.8/94:0.0853484533)81.7/16:0.0435339431)88.4/24:0.0319509415,((WP_162648791.1-Citrobacter-freundii:0.1439051700,(WP_060683585.1-Citrobacter-portucalensis:0.1452891000,(WP_079798993.1-Salmonella-enterica:0.1091182205,WP_032255798.1-Enterobacteriaceae:0.1024651360)76.7/98:0.0167791863)93.9/98:0.0802091356)80.9/100:0.0498128667,(WP_195994783.1-Citrobacter-portucalensis:0.2661019738,WP_048218074.1-Citrobacter-freundii-complex:0.2300674980)90.7/100:0.0894026643)96.4/99:0.0857571972)74.7/82:0.0094593861,((((WP_181680936.1-Citrobacter-sp.-RHB25-C09:0.1444977117,(WP_090049757.1-Citrobacter-amalonaticus:0.0431849219,WP_196013128.1-Citrobacter-farmeri:0.0331223467)98.7/100:0.1546084695)78.2/100:0.0486840826,WP_103775088.1-Citrobacter-amalonaticus:0.2353415456)90.1/100:0.1031716324,WP_121265736.1-Enterobacter-sp.-R1-2018:0.3531457104)86.4/100:0.0938258994,WP_249031739.1-Tenebrionicola-larvae:0.3286637293)81.4/95:0.0521726216)78.3/79:0.0118745087)90.5/91:0.0391304130,(WP_032659356.1-Enterobacter-hormaechei:0.5629850379,(WP_220393175.1-Citrobacter-freundii:0.3495541845,WP_219093375.1-Escherichia-coli:0.1975983735)76.8/98:0.0358559713)89.3/75:0.0544290183)85.2/71:0.0368076095,((WP_170897747.1-Escherichia-coli:0.2106516826,(WP_001100781.1-Escherichia-coli:0.2255415492,WP_208464084.1-Escherichia-coli:0.1423926655)67.4/85:0.0275508380)96.2/79:0.1038614572,(WP_042093344.1-Escherichia-coli:0.2611458262,WP_089706542.1-Escherichia-coli:0.1101941907)74.8/65:0.0508872218)77/64:0.0266477296)82.1/64:0.0327470156,WP_001542972.1-Escherichia-coli:0.1790095200)98.5/94:0.1490092434,((WP_241647798.1-Rosenbergiella-sp.-JB07:0.3500978921,WP_065370581.1-Kosakonia-sacchari:0.2398233872)88.4/100:0.0513781573,WP_181599127.1-Citrobacter-sp.-RHB36-C18:0.2864612389)78.3/93:0.0454551599)87.6/85:0.0816887605)38.3/60:0.0219894813,(WP_160868038.1-Pantoea-sp.-Taur:0.4252577347,WP_008499438.1-Enterobacter-cloacae-complex:0.2352557515)87.5/93:0.0710965130)81.8/79:0.0347302018,(WP_090085547.1-Escherichia-coli:0.0585695046,(WP_103283990.1-Citrobacter:0.2120388891,WP_234083964.1-Enterobacter-quasiroggenkampii:0.1655679522)45.6/94:0.0296589677)99.9/100:0.2248921533)79.7/90:0.0335653760,((((((((((((WP_182931197.1-Enterobacter-cloacae:0.2132203027,(WP_048284323.1-Pluralibacter-gergoviae:0.1785230481,WP_211776196.1-Klebsiella-pneumoniae:0.0724622033)53.5/65:0.0217449826)20/59:0.0112244187,WP_047723323.1-Klebsiella-michiganensis:0.0610241680)71.1/60:0.0139636965,(WP_095100910.1-Raoultella-sp.-10-1:0.0858899815,WP_065799756.1-Klebsiella-pneumoniae:0.0079895255)85.1/97:0.0374557769)48.3/87:0.0212970359,WP_149270444.1-Klebsiella-aerogenes:0.0450760208)74.3/34:0.0110094571,(((WP_234537246.1-Raoultella-ornithinolytica:0.0478464006,WP_113990968.1-Raoultella:0.0316608026)96/100:0.0500547947,WP_104011323.1-Klebsiella-michiganensis:0.0531221385)84.7/40:0.0243904003,(WP_234579544.1-Raoultella-terrigena:0.0387343149,WP_186601692.1-Klebsiella-quasipneumoniae:0.0395217989)75/41:0.0077537542)75.6/89:0.0106963985)77.4/46:0.0100292908,(WP_213760001.1-Klebsiella-aerogenes:0.0802500292,WP_223370324.1-Klebsiella-pneumoniae:0.0407557726)56.8/97:0.0159424667)71.9/47:0.0074250138,WP_221694709.1-Klebsiella-aerogenes:0.1228058831)70.5/91:0.0201315194,WP_128320364.1-Raoultella-planticola:0.0645433878)58.5/75:0.0094522872,WP_046852082.1-Raoultella-planticola:0.0588040195)0/74:0.0000025706,(WP_236948387.1-Klebsiella-sp.-2019SCSN059:0.0483146166,((WP_114692711.1-Klebsiella-pneumoniae:0.0458823477,WP_259891680.1-Klebsiella-variicola:0.0374386441)0/36:0.0000027330,WP_123755035.1-Raoultella-sp.-BIGb0399:0.0451226091)77.4/94:0.0112902573)79.2/90:0.0099858721)95.2/90:0.0792640304,(((WP_095100919.1-Raoultella-sp.-10-1:0.0751981250,WP_212709529.1-Tatumella-sp.-JGM130:0.0116891787)92.5/100:0.0788690800,WP_139968923.1-Klebsiella-pneumoniae:0.1611402793)82.6/100:0.0479968669,WP_236948353.1-Klebsiella-sp.-2019SCSN059:0.1139669619)78.4/95:0.0406233298)88.4/97:0.0641427688,WP_123822891.1-Klebsiella:0.0495490998)99.1/100:0.1317285128)85.9/99:0.0553822332,(WP_072663757.1-Enterobacteriaceae:0.1155481233,WP_021241446.1-Enterobacter-roggenkampii:0.2176041853)64.5/98:0.0287144411)97.1/100:0.1213248702)93.5/100:0.0679725419,XP_011503919.1-Ceratosolen-solmsi-marchali:0.1084637066);

**Nematode RML-3, with bacterial and eukaryotic homologs**

**Shown in Figure S2**

ML tree method: IQ-TREE

Sequences: 2787

Substitution model: WAG

(WP_063501760.1-Escherichia-coli:0.0311515439,(WP_044173167.1-Phytobacter-massiliensis:0.0302561146,((((((((WP_203451401.1-Lelliottia-sp.-RWM.1:0.0537806620,WP_185654835.1-Citrobacter-braakii:0.0928492029)89.6/100:0.0571094182,WP_183170804.1-Atlantibacter-sp.-RC6:0.1630048519)98.1/100:0.0789925121,(WP_160868038.1-Pantoea-sp.-Taur:0.2037159947,WP_008499438.1-Enterobacter-cloacae-complex:0.1484119631)80.4/84:0.0482763661)57.6/83:0.0144460737,((WP_090085547.1-Escherichia-coli:0.0512416863,WP_103283990.1-Citrobacter:0.1391955133)10.7/40:0.0126213097,WP_234083964.1-Enterobacter-quasiroggenkampii:0.0941953165)99.9/100:0.1264769264)86.1/94:0.0350396477,((((((((((((WP_249615874.1-Craterilacuibacter-sp.-RT1T:0.2675506095,WP_014088374.1-Pseudogulbenkiania-sp.-NH8B:0.1578133254)26/45:0.0224072457,WP_008952446.1-Pseudogulbenkiania-ferrooxidans:0.1660848395)25.3/36:0.0461919379,(WP_163316497.1-Crenobacter-sedimenti:0.1007001195,WP_160797365.1-Craterilacuibacter-sinensis:0.1094132134)99/100:0.1239910528)93.6/93:0.0648458889,(((((((((((((((((((((((((WP_250059255.1-Halomonas-sp.-ATCH28:0.2129176201,WP_179153381.1-Chromohalobacter-salexigens:0.1510387791)92.5/100:0.0588318697,WP_250939176.1-unclassified-Larsenimonas:0.1833328452)88/93:0.0493786000,(WP_106478747.1-Phytohalomonas-tamaricis:0.1485948578,(WP_138213821.1-Pseudomonas-nitroreducens:0.0367360319,WP_017517140.1-Pseudomonas-nitroreducens:0.0152172452)100/100:0.1977010974)81.1/88:0.0562381459)89.3/94:0.0309826020,(((WP_136867012.1-Guyparkeria-sp.-SB14A:0.1828511795,((WP_139871681.1-Candidatus-Methylopumilus-universalis:0.1414374588,(WP_139929800.1-Candidatus-Methylopumilus-universalis:0.2040130928,WP_140005423.1-Candidatus-Methylopumilus-planktonicus:0.0600028095)94.6/100:0.0788455488)99.6/100:0.1486228722,WP_058575749.1-unclassified-Guyparkeria:0.1520288280)14.5/82:0.0207675561)93.6/94:0.0715311589,(WP_218634960.1-Marinobacter-arenosus:0.0841189621,WP_240716217.1-Halomonas-neptunia:0.0834521308)98.4/100:0.0965607432)71.2/82:0.0382655099,((((WP_269022943.1-Methylomonas-sp.-MP1:0.0864628972,(WP_196437371.1-Methylomonas-sp.-LL1:0.0895897700,(WP_150047608.1-Methylomonas-rhizoryzae:0.2004687622,WP_066983474.1-Methylomonas-lenta:0.0708881741)86.7/100:0.0404948636)90.1/96:0.0446598948)90.5/99:0.0455885503,(WP_140911374.1-Methylomonas-koyamae:0.1512938929,((WP_071159776.1-Methylomonas-sp.-LWB:0.1281447939,(WP_020485671.1-Methylomonas-sp.-MK1:0.1270497486,(WP_256607223.1-Methylomonas-sp.-WSC-7:0.0568393099,(WP_205450646.1-Methylomonas-sp.-EFPC1:0.0688866106,WP_064036612.1-Methylomonas-methanica:0.0000022353)53.4/100:0.0080406870)94.3/100:0.0514108362)89.5/98:0.0458787327)57.8/71:0.0067058962,WP_036273427.1-Methylomonas:0.0831388335)88.3/100:0.0319663621)79.1/100:0.0399517898)85.1/99:0.0332892102,(WP_148817865.1-Pigmentiphaga-aceris:0.2324644948,WP_064030819.1-Methylomonas-koyamae:0.1157670730)59.1/90:0.0434038166)98.5/90:0.0975227512,(WP_019964198.1-Pannonibacter-phragmitetus:0.3682701005,WP_138632053.1-Pseudoalteromonas-spongiae:0.1830299092)34.3/78:0.0383496884)89.1/81:0.0495820820)65/81:0.0248541513)98.1/90:0.0810512652,((WP_077733980.1-Paenalcaligenes-hominis:0.0321660509,WP_167661700.1-Paenalcaligenes-hominis:0.0310241054)100/100:0.1595215117,WP_216129682.1-Azospira-inquinata:0.2411097165)80.7/61:0.0470042628)0/9:0.0000026188,((((((WP_044615503.1-Gynuella-sunshinyii:0.2431424585,(WP_046292890.1-Pandoraea-oxalativorans:0.0711613563,WP_150719538.1-Pandoraea-capi:0.0814144417)100/100:0.2097447858)68.7/100:0.0524518306,WP_092579398.1-Achromobacter-sp.-NFACC18-2:0.2587477315)67.9/97:0.0433785277,WP_227683402.1-Pusillimonas-sp.-MFBS29:0.1517750708)90.6/86:0.0441322954,((((WP_102075761.1-Pollutimonas-subterranea:0.1324462560,WP_102071501.1-Pollutimonas-nitritireducens:0.1572697408)73.9/99:0.0463418866,(WP_212592286.1-Janthinobacterium-sp.-TND4EL3:0.1552354180,WP_047550623.1-Methylotenera-sp.-G11:0.1607467508)66.2/100:0.0507068850)80.8/96:0.0418769778,(((WP_220354719.1-Rhizobium-cauense:0.1759595189,(((WP_184450570.1-Rhizobium-sp.-BK538:0.1356418527,WP_233136718.1-Agrobacterium-vaccinii:0.1810073341)87.6/93:0.0379697474,WP_016557527.1-Rhizobium-grahamii:0.1220886382)75.7/17:0.0146410338,(WP_018896487.1-Rhizobium-sp.-2MFCol3.1:0.1662804620,(WP_072373837.1-Rhizobium-tibeticum:0.0800222849,WP_007537558.1-Rhizobium-mesoamericanum:0.0496506102)83.5/90:0.0427975485)80.6/7:0.0219157585)52.7/75:0.0414134427)71.1/98:0.0345277644,WP_153273632.1-Rhizobium-grahamii:0.1556633720)97.6/100:0.0854532579,WP_068803504.1-Immundisolibacter-cernigliae:0.1429213567)88.3/97:0.0396652570)48/85:0.0169588596,(WP_019141909.1-Noviherbaspirillum-massiliense:0.1024906247,WP_146330669.1-Noviherbaspirillum-sp.-UKPF54:0.1718209201)94.6/93:0.0629195896)87.7/91:0.0310319045)76/86:0.0221796130,(WP_230502293.1-Janthinobacterium-sp.-FW305-128:0.2570412563,WP_249918475.1-Pseudomonas-sp.-R3.Fl:0.1201846476)90.7/96:0.0541825437)59.5/88:0.0335003474,WP_065086507.1-Pseudomonas:0.1792767427)92.9/83:0.0315776409)80.5/5:0.0132021108,((((((((WP_225237063.1-Massilia-oculi:0.2091861601,WP_147937383.1-Massilia-arenae:0.1162191017)97.9/100:0.1130587800,(WP_192051557.1-unclassified-Massilia:0.0993762106,WP_136222466.1-Massilia-sp.-Mn16-1_5:0.0630761566)61.7/73:0.0532902248)94.4/100:0.0731607171,((WP_057247240.1-Duganella-sp.-Root198D2:0.1397347794,(((WP_029241333.1-Pseudomonas:0.0670165163,((WP_054996663.1-Pseudomonas-ficuserectae:0.0289513463,(WP_007252556.1-Pseudomonas-syringae-group:0.0372816315,WP_025390602.1-Pseudomonas-syringae:0.0236881303)84.5/100:0.0227190692)93.9/100:0.0342508185,WP_125830676.1-Pseudomonas-sp.-v388:0.0674287717)48.8/97:0.0237453915)85.9/97:0.0372127312,((WP_016973326.1-Pseudomonas-tolaasii:0.0905570055,WP_099548838.1-Pseudomonas-sp.-2995-3:0.0854870778)68.1/97:0.0414920553,(WP_177015116.1-Pseudomonas-costantinii:0.0479577564,WP_141063556.1-Pseudomonas-fluorescens:0.0816504862)87.7/100:0.0371637673)97.7/100:0.0850743800)87.3/100:0.0540634019,WP_217963608.1-Candidimonas-humi:0.1916609653)84.1/94:0.0582909049)30.6/19:0.0298838783,WP_155457065.1-Massilia-eburnea:0.1105681379)79.3/72:0.0291128363)61.3/44:0.0348298847,(WP_058422493.1-Pseudomonas:0.1335078399,WP_223462047.1-Pseudomonas-sp.-A-R-26:0.1489695633)91/98:0.0529415184)95.3/89:0.0414285190,((WP_042945119.1-Pseudomonas-fluorescens:0.0574643167,WP_181643258.1-Pseudomonas-sivasensis:0.0391559125)99.6/100:0.0945542658,((((WP_169965430.1-unclassified-Pseudomonas:0.0349998077,WP_047278503.1-Pseudomonas-lundensis:0.0235452020)99.1/100:0.0776668512,WP_205489922.1-Pseudomonas-lactucae:0.1349190483)83.5/21:0.0260486088,WP_214501190.1-Pseudomonas-sp.-RC3H12:0.2012054130)74.8/7:0.0082620127,WP_104727848.1-Pseudomonas-oleovorans:0.1195881062)30/7:0.0270539978)86.7/54:0.0246407749)86.8/53:0.0123959378,((WP_058488093.1-Pseudomonas:0.1165805237,WP_043256200.1-Pseudomonas-knackmussii:0.0945706731)47.6/63:0.0289209215,((WP_088420173.1-Pseudomonas-nitroreducens:0.0994651588,WP_254475170.1-unclassified-Pseudomonas:0.0580178976)69.9/100:0.0248971399,WP_205338011.1-Pseudomonas-sp.-PDNC002:0.0525364526)99.8/100:0.1174202381)95.4/100:0.0491295240)80.6/92:0.0169867549,((((WP_191471912.1-Pseudomonas-asiatica:0.1552275850,(WP_043318452.1-Pseudomonas-sp.-PI1:0.0401480972,WP_043268312.1-Pseudomonas:0.0280179266)86.8/100:0.0229291511)95/100:0.0444528481,(WP_089390899.1-Pseudomonas-delhiensis:0.0880915360,WP_076426141.1-Pseudomonas-alcaligenes:0.1057712312)79.6/100:0.0271390612)95.3/100:0.0551054261,WP_233297479.1-Pseudomonas:0.1204155742)77.2/97:0.0211150510,(WP_184594840.1-Pseudomonas-nitritireducens:0.1336145688,WP_192381882.1-Pseudomonas-sp.-PDM20:0.0938530580)99.1/100:0.1064533025)93.5/92:0.0502887286)73/57:0.0218777113,(((((WP_020564242.1-Methylosarcina-fibrata:0.1376412478,WP_074973890.1-Pseudomonas-otitidis:0.1474204159)92.3/17:0.0514398497,(WP_101152997.1-Pseudomonas-sp.-Choline-3u-10:0.0986441954,((WP_046617631.1-Stutzerimonas-stutzeri:0.0519809709,WP_135265705.1-Stutzerimonas-stutzeri:0.0462058233)92.5/100:0.0505186005,WP_003299494.1-Stutzerimonas-stutzeri:0.0779083227)67/100:0.0243930169)91.7/96:0.0371644416)78.1/16:0.0166755703,(WP_125865093.1-Stutzerimonas-stutzeri:0.1264389797,WP_200290630.1-Stutzerimonas-balearica:0.0924613410)95.9/96:0.0631044371)78/19:0.0132339328,((WP_180152801.1-Parapusillimonas-granuli:0.1149633555,(WP_088146851.1-Achromobacter-denitrificans:0.0803674752,WP_105555808.1-Achromobacter-sp.-MYb9:0.0670324481)97.3/100:0.0645268858)53.3/52:0.0275699038,(WP_011111299.1-Nitrosomonas-europaea:0.1303942691,WP_046857648.1-Pseudomonas-sp.-CCOS-191:0.1305222098)67.8/63:0.0292420939)94.9/41:0.0498486178)54.1/75:0.0141080357,((WP_209872004.1-Pseudomonas-sp.-BP8:0.1353324284,WP_066093302.1-Hydrogenophaga-crassostreae:0.1291568044)73/99:0.0281196426,(((WP_177138191.1-Hydrogenophaga-aromaticivorans:0.1016593039,(WP_236964528.1-Hydrogenophaga-sp.-SL48:0.0677609423,(WP_226493830.1-Hydrogenophaga-taeniospiralis:0.0406601625,WP_210824367.1-Hydrogenophaga-aromaticivorans:0.0488710466)68.2/87:0.0313849982)97.5/100:0.0663066624)53.1/26:0.0214529893,WP_133157403.1-Hydrogenophaga-pseudoflava:0.1199565887)91.4/27:0.0415061203,WP_085721622.1-Pseudomonas:0.1225191976)94.7/99:0.0485692847)93.2/88:0.0438485781)79.6/11:0.0330002640)78.9/5:0.0115995176)82.1/5:0.0139421873,((WP_076518421.1-Achromobacter-sp.-MFA1-R4:0.0731098881,(WP_104142565.1-Achromobacter-spanius:0.0195333190,WP_120157261.1-Achromobacter-sp.-B7:0.0460400234)86.2/100:0.0302980390)92.4/100:0.0498241002,WP_158658301.1-Achromobacter-sp.-AONIH1:0.2439891660)93.5/8:0.0634945109)61.4/5:0.0333346824,((WP_209394989.1-Rhodoferax-sp.-AJA081-3:0.1990167694,WP_094844257.1-Bordetella-genomosp.-11:0.1331277832)7.6/8:0.0420035934,WP_166067558.1-Diaphorobacter-sp.-HDW4B:0.1672393026)85.3/8:0.0473020767)93.4/86:0.0408126411,(WP_033955816.1-Pseudomonas-aeruginosa:0.0180740672,WP_003153917.1-Pseudomonas-aeruginosa-group:0.0350098633)100/100:0.1612504178)0/7:0.0281718006,((((WP_066098770.1-Halothiobacillus-sp.-LS2:0.1078596279,WP_070073009.1-Acidihalobacter-aeolianus:0.1455069002)95.3/100:0.0592156589,(WP_168987795.1-Azoarcus-taiwanensis:0.1752557596,(WP_018865194.1-Thioalkalivibrio-sp.-ARh3:0.1637486410,(((((WP_238617884.1-Ectothiorhodospira-shaposhnikovii:0.1072251247,WP_025282248.1-Ectothiorhodospira-haloalkaliphila:0.0941800956)93.3/100:0.0620695437,WP_238622182.1-unclassified-Ectothiorhodospira:0.0924650753)84.9/100:0.0517156857,(((((((WP_060482506.1-Pseudomonas-sp.-NBRC-111119:0.1380608330,WP_159409308.1-Pseudomonas-putida:0.0681196989)97/97:0.0525142452,(((WP_182365203.1-Pseudomonas-juntendi:0.0569113831,WP_182365213.1-Pseudomonas:0.0849327969)36.8/67:0.0125746038,((WP_110607173.1-Pseudomonas-fulva:0.0113858888,WP_088523134.1-unclassified-Pseudomonas:0.0412559183)96.7/100:0.0333270098,(WP_167339957.1-Pseudomonas-fulva:0.0279768219,WP_186476580.1-Pseudomonas-promysalinigenes:0.0357774818)71.1/98:0.0171780099)86.3/96:0.0170699673)87.2/58:0.0133797693,(WP_267821885.1-Pseudomonas-putida:0.0715112008,(WP_201214098.1-Pseudomonas-sp.-S36:0.0187325066,WP_054890059.1-Pseudomonas:0.0524196073)95.1/100:0.0360019652)72.8/7:0.0048970761)70.2/7:0.0040024653)81.8/61:0.0123833795,(WP_167277242.1-Pantoea-sp.-Tr-811:0.0172131093,WP_103449615.1-Pseudomonas-putida:0.0460999478)97.1/100:0.0420908849)71.9/74:0.0081605538,((WP_115272924.1-Pseudomonas-putida:0.0890034108,(WP_046489762.1-Pseudomonas-veronii:0.1008535523,(WP_059395284.1-Pseudomonas-putida:0.0748146619,(WP_009685980.1-Pseudomonas:0.0701585676,WP_024086460.1-Pseudomonas:0.0452968001)92.3/100:0.0336961372)88.5/100:0.0245192072)89.9/98:0.0273177661)89.2/83:0.0278559433,((WP_196129978.1-Pseudomonas-fulva:0.0788460791,WP_201164743.1-Pseudomonas-sp.-S60:0.0257840470)98.8/100:0.0681453544,(((WP_174142546.1-Pseudomonas-putida:0.1106630149,WP_027907237.1-Pseudomonas:0.0311097316)29.9/100:0.0094855513,WP_075044077.1-Pseudomonas-putida:0.0406553278)69/100:0.0214915036,(WP_023379297.1-Pseudomonas-sp.-VLB120:0.0776300559,WP_202675963.1-Pseudomonas-sp.-SK2:0.0333584624)98/100:0.0587831179)92.1/100:0.0392791246)76.1/100:0.0252633371)79.1/68:0.0125255761)90.3/90:0.0229102908,(WP_029885231.1-Pseudomonas:0.0248353045,(WP_230064532.1-Pseudomonas-sp.-CIP-10:0.0176150284,WP_197863301.1-Pseudomonas-putida:0.0348805061)97/100:0.0339233947)18.8/71:0.0084595522)96.8/79:0.0579914675,WP_060394622.1-Pseudomonas-entomophila:0.0815986435)96.3/100:0.0629682319,WP_183165449.1-Azomonas-macrocytogenes:0.0955574819)74.9/94:0.0294665109)62.2/66:0.0307487157,(WP_256068944.1-Stutzerimonas-stutzeri:0.1241743684,WP_004074545.1-Desulfobacter-postgatei:0.2102431724)39/29:0.0269451516)12.8/20:0.0290840102,WP_026685886.1-Azovibrio-restrictus:0.1518010910)21.2/48:0.0167375974)64.9/75:0.0282059317)94.7/98:0.0498274472)25.5/74:0.0180654751,(WP_136386324.1-Pseudothauera-rhizosphaerae:0.1278638729,(WP_114649035.1-Pseudothauera-hydrothermalis:0.0724158624,WP_187718084.1-Thauera-sp.-CAU-1555:0.1128466387)81.6/99:0.0482550429)99.5/100:0.1141389105)71.5/94:0.0185531768,(WP_013164002.1-Desulfurivibrio-alkaliphilus:0.1623985642,WP_263768642.1-Propionivibrio-sp.-SG131:0.1911377750)89.8/57:0.0823629381)83.5/8:0.0267713266)84.2/23:0.0283051441,(((WP_121459656.1-Azonexus-fungiphilus:0.1750012128,WP_117174481.1-Rhodoferax-lacus:0.1580680550)24/99:0.0249852214,((WP_094482496.1-Rhodoferax-sp.-TH121:0.0577861878,WP_105262671.1-Rhodoferax-sp.-TS-BS-61-7:0.0414529159)89.6/100:0.0329066732,WP_214121631.1-Curvibacter-sp.-CHRR-16:0.1947138892)97.4/100:0.0843463174)88.9/99:0.0360974457,(WP_066884254.1-Dechloromonas:0.0842105277,WP_027458229.1-Dechloromonas-agitata:0.0827499370)95.9/100:0.0642637661)92.1/99:0.0504639572)89.2/23:0.0301081082,(((((((((WP_072282603.1-Syntrophotalea-acetylenivorans:0.1081902944,WP_092056970.1-Desulfuromonas-acetexigens:0.1312556421)80.7/45:0.0407491154,WP_155876508.1-Desulfuromonas-sp.-AOP6:0.1945170013)73.1/42:0.0366484927,WP_043770486.1-Desulfobulbus-propionicus:0.1782327119)0/63:0.0451815130,WP_148894253.1-Geothermobacter-ehrlichii:0.2371945495)32.3/89:0.0302269954,WP_028317018.1-Desulfobulbus-elongatus:0.2199223321)92.6/53:0.0469118419,(WP_153584563.1-Caenimonas-koreensis:0.2092913334,(WP_007185580.1-Hydrocarboniphaga-effusa:0.0628418843,WP_007185565.1-Hydrocarboniphaga-effusa:0.0573476483)99.1/100:0.1216651276)99.1/100:0.1235242731)88.3/29:0.0358978672,(((WP_227390214.1-Halomonas-malpeensis:0.3696375835,WP_153110537.1-Propionivibrio-limicola:0.1260087713)92.3/95:0.0784716937,((WP_226989104.1-Desulfuromonas-sp.-TF:0.2651225237,WP_227459030.1-Cupriavidus-pauculus:0.1857111932)97.1/99:0.1235353277,((WP_167682126.1-Rhodocyclus-gracilis:0.2056681257,(WP_153115862.1-Rhodocyclus-tenuis:0.0379626104,WP_200385811.1-Rhodocyclus-tenuis:0.0169381452)95.6/100:0.0700291653)94.3/100:0.0693535703,WP_201212349.1-Rhodocyclus-purpureus:0.1039908830)81.9/98:0.0517648653)49.6/58:0.0586909182)31.5/25:0.0096815386,((WP_110265962.1-Sinimarinibacterium-flocculans:0.2301296418,(WP_234070476.1-Myxococcus-stipitatus:0.1804579977,WP_120555541.1-Corallococcus-aberystwythensis:0.1968829135)99.8/100:0.1734933694)95.2/70:0.0870445331,(((((((WP_169107474.1-Comamonas-suwonensis:0.0423311179,WP_219343559.1-Comamonas-sp.-Y33R10-2:0.0693462193)95.5/100:0.0437612958,(WP_003073420.1-Comamonas:0.0609468359,((WP_211457163.1-Comamonas-sp.-PE63:0.0419155348,WP_148729134.1-Comamonas-sp.-Z3:0.0382445902)40.8/71:0.0056952207,WP_034391098.1-Comamonas-thiooxydans:0.0130583777)76.4/71:0.0075935169)86.3/99:0.0241148262)70.4/99:0.0127204809,WP_218241177.1-Comamonas-fluminis:0.0891547317)99.9/100:0.1268276183,WP_012203208.1-Delftia-acidovorans:0.1429492637)49.4/99:0.0434339768,WP_058879441.1-Comamonas-kerstersii:0.0964528233)83.5/98:0.0570689835,WP_066475960.1-Comamonas-terrae:0.0971617653)99.8/100:0.1384079426,((((((((WP_036226274.1-Mesoaciditoga-lauensis:0.1438561920,WP_047753768.1-Kosmotoga-pacifica:0.1234818159)0/28:0.0241107924,(((WP_077198159.1-Thermosipho-affectus:0.0650279500,WP_012057290.1-Thermosipho-melanesiensis:0.1348712648)86.8/100:0.0347994737,(WP_127352844.1-Caldicellulosiruptor-changbaiensis:0.0484526224,WP_014043315.1-Caldicellulosiruptor-acetigenus:0.0270015674)100/100:0.2521868637)92.6/100:0.0460608451,((((((WP_117708548.1-Fusobacterium-mortiferum:0.0000023577,(WP_204716698.1-Fusobacterium-mortiferum:0.0155084994,WP_204716813.1-Fusobacterium-mortiferum:0.2884006226)90.1/94:0.0455676251)58.7/94:0.0190927209,WP_115270465.1-Fusobacterium-necrogenes:0.0573158869)97.4/98:0.0651044962,(((WP_130889691.1-Fusobacterium-ulcerans:0.0379604667,(WP_005950923.1-Fusobacterium-varium:0.0351323298,(WP_070580525.1-Fusobacterium:0.0464753077,WP_227372109.1-Fusobacterium-ulcerans:0.0226163977)16.1/100:0.0118461708)88/100:0.0131901154)50.9/100:0.0097371219,(WP_241635364.1-Fusobacterium-gastrosuis:0.0971813538,(WP_074518138.1-Fusobacterium-necrophorum:0.0623875487,(WP_211495349.1-Fusobacterium-necrophorum:0.1235974647,WP_032890075.1-Fusobacterium-nucleatum:0.3948299710)83.9/100:0.0440887974)96.5/100:0.0589093066)97.8/100:0.0633459724)84.8/98:0.0236355149,WP_261793436.1-Fusobacterium-varium:0.2150291655)93.6/99:0.0518140570)95.4/100:0.0887057947,((WP_147004474.1-Leptotrichia-wadei:0.0240877608,WP_018499460.1-Leptotrichia-wadei:0.0288072951)97.5/100:0.0774863762,((WP_068155536.1-Leptotrichia-sp.-oral-taxon-847:0.0700287541,(WP_211617428.1-Leptotrichia-sp.-oral-taxon-218:0.0133860237,WP_094080762.1-Leptotrichia-sp.-oral-taxon-498:0.0414371582)97.1/100:0.0540852968)22.1/90:0.0272410592,WP_099992503.1-Fusobacterium-pseudoperiodonticum:0.0906686121)80.3/100:0.0447259764)99.7/100:0.1636892724)89.3/99:0.0710586544,WP_012548442.1-Dictyoglomus-thermophilum:0.2440626007)73/24:0.0084671990,(WP_013388736.1-Ilyobacter-polytropus:0.3051000676,WP_173194267.1-Paenibacillus-frigoriresistens:0.1975191340)74.2/24:0.0378674404)98.5/100:0.0870896194)89.1/100:0.0471684025)95.3/100:0.0629655593,(WP_099829233.1-Mesotoga-sp.-Brook.08.105.5.1:0.1361888274,WP_099830786.1-Mesotoga-sp.-Brook.08.105.5.1:0.1323240290)98.9/100:0.1140100286)53.9/98:0.0447532511,(((((((((((((((WP_054742706.1-Cellulosilyticum-ruminicola:0.0708388134,(WP_177671917.1-Zhenhengia-yiwuensis:0.0370911739,WP_053982981.1-Niameybacter-massiliensis:0.0921963174)99.9/100:0.0879941144)91.6/100:0.0404816897,WP_069999280.1-Cellulosilyticum-sp.-I15G10I2:0.1756559610)99/100:0.0930924893,WP_072892386.1-Clostridium-fallax:0.0797468988)86.6/86:0.0269410632,(WP_073337580.1-Clostridium-grantii:0.1109323185,WP_072832886.1-Clostridium-collagenovorans:0.0293414801)79.4/82:0.0166387799)82.1/60:0.0135909557,((WP_195972826.1-Clostridium-thermobutyricum:0.0738891268,(WP_209795868.1-Clostridium-moniliforme:0.0569271661,WP_194191032.1-Clostridium-chrysemydis:0.0865869631)93.6/100:0.0411837995)87.4/98:0.0205068207,WP_040194942.1-Clostridium-culturomicium:0.0518970799)88.9/97:0.0208634780)86.3/71:0.0125853882,((((((WP_094369216.1-Romboutsia-weinsteinii:0.1243398767,WP_021359979.1-Clostridioides-difficile:0.2343622794)65.7/100:0.0350454101,(WP_074918830.1-Terrisporobacter-glycolicus:0.0601978883,WP_195929149.1-Clostridium-sp.-1001270J_160509_D11:0.0282295920)78.1/100:0.0172539729)91.1/97:0.0271840559,(((WP_092726456.1-Romboutsia-lituseburensis:0.1470222710,WP_258242885.1-Romboutsia-lituseburensis:0.0294074613)98.2/100:0.0643497331,WP_057560648.1-Paeniclostridium-sordellii:0.0569185124)50.3/100:0.0225424186,((WP_057580750.1-Paeniclostridium-sordellii:0.0940595166,WP_057538715.1-Paeniclostridium-sordellii:0.0356408506)78.7/99:0.0130345229,WP_195252007.1-Romboutsia-sp.-1001713B170207_170306_H8:0.0465350943)61/97:0.0173633897)88.6/97:0.0299015239)91/94:0.0272957428,(((WP_152890015.1-Clostridium-tarantellae:0.1109962587,WP_047001949.1-Clostridium-sp.-C8:0.0376111930)90.4/98:0.0227528343,(WP_195959433.1-Clostridium-tertium:0.0554386997,WP_195429094.1-Clostridium-sp.-D46t1_190503_E9:0.0212243985)0/76:0.0074471525)76.4/98:0.0062606238,(WP_066890539.1-Clostridium-nigeriense:0.0392954430,WP_016207773.1-Clostridium-sartagoforme:0.0328536517)89.9/61:0.0165181649)97.7/99:0.0312698207)0/28:0.0000028168,(((((WP_055265352.1-Clostridium:0.0316816400,(WP_186859336.1-Clostridium-sp.-NSJ-6:0.0511793027,(WP_055265304.1-Clostridium-disporicum:0.0522137374,(WP_154821366.1-Clostridium-saudiense:0.0074746055,WP_195245504.1-Clostridium-celatum:0.0595548882)70.2/98:0.0232678379)32.9/97:0.0276897205)59/97:0.0443848954)86.1/100:0.0596069878,WP_186834444.1-Eubacteriales:0.0989461183)95.8/100:0.0510930797,((WP_066874757.1-Clostridium-mediterraneense:0.1570945586,WP_198610231.1-Clostridium-perfringens:0.0168832060)93.1/93:0.0340874332,(((WP_235068992.1-Turicibacter-sp.-TJ11:0.0546546807,(WP_161831114.1-Turicibacter-sp.-TS3:0.0509954306,WP_195933776.1-Turicibacter-sanguinis:0.0170627195)42.8/50:0.0067840447)12.5/41:0.0092638736,WP_155223812.1-Turicibacter-sanguinis:0.0268137047)87.2/94:0.0202814368,WP_212726030.1-Turicibacter-bilis:0.0106441957)99.5/99:0.0742787983)39.2/72:0.0149912341)59/87:0.0135593574,WP_055258173.1-Sarcina-ventriculi:0.0924870830)94/98:0.0371595161,WP_055184115.1-Clostridium-paraputrificum:0.0599005535)37.9/67:0.0129817302)83.4/44:0.0183226513,(((((WP_029452249.1-Clostridium-algidicarnis:0.0850898887,WP_216183125.1-Clostridium-gasigenes:0.0190675527)59.9/99:0.0295503741,WP_216313757.1-Clostridium-gasigenes:0.0579725019)98.4/100:0.0535002017,(WP_197096717.1-Paeniclostridium-sordellii:0.0687125800,WP_238903698.1-Clostridium-sp.-YIM-B02506:0.0536306307)81.4/88:0.0162659007)71.1/57:0.0059037628,(WP_163195303.1-Clostridium-thermarum:0.1163179956,WP_066677671.1-Clostridium-septicum:0.0259513754)30.1/49:0.0138188054)80.7/50:0.0145739177,WP_095406848.1-Romboutsia-maritimum:0.1137786183)74.2/23:0.0071084405)90.7/64:0.0244323043)80.9/63:0.0182823123,(((WP_055069574.1-Clostridium-massiliamazoniense:0.1128888155,(WP_090010800.1-Clostridium-sp.-DSM-8431:0.0934289704,(WP_106059304.1-Clostridium-vincentii:0.0874625989,WP_143316579.1-Clostridium:0.0205154185)91.4/100:0.0367269343)58.8/77:0.0161668077)90.3/98:0.0223879543,(WP_261829490.1-Clostridium-sp.-TW13:0.1349760904,WP_148566871.1-Acetobacterium-paludosum:0.2653741460)70.1/94:0.0253721577)87.6/74:0.0179037875,(WP_066825124.1-Clostridium-tepidiprofundi:0.0875712581,WP_072889493.1-Tepidibacter-formicigenes:0.0621685247)90.1/99:0.0238028858)85.6/73:0.0143135209)90.9/89:0.0310744858,WP_072990516.1-Clostridium-cavendishii:0.1845623337)72.3/35:0.0085476086,WP_133628539.1-Fonticella-tunisiensis:0.0625471459)34.3/71:0.0154931249,((WP_055666468.1-Desnuesiella-massiliensis:0.1155596917,((((WP_099190526.1-Tepidibacter-mesophilus:0.1044119243,WP_226126924.1-Clostridium-algoriphilum:0.2706224662)86.6/46:0.0433961447,((WP_142413383.1-Hathewaya-massiliensis:0.1494601304,WP_035130905.1-Clostridium-sulfidigenes:0.1637504068)25.6/97:0.0237753841,WP_072903743.1-Hathewaya-proteolytica:0.2447650670)98.1/98:0.0834665450)42/47:0.0161984160,WP_095404508.1-Romboutsia-maritimum:0.2299246211)36.4/90:0.0198143193,WP_072725426.1-Tepidibacter-thalassicus:0.1455263673)89.1/70:0.0282007676)92.4/77:0.0398582818,(((WP_216288229.1-Clostridium-psychrophilum:0.1156389038,WP_216137569.1-Clostridium-estertheticum:0.0315063290)25/100:0.0054661364,WP_220704225.1-Clostridium-estertheticum:0.0332222569)91.1/100:0.0387917391,WP_220321605.1-Clostridium-tagluense:0.0501820147)99/100:0.0757229143)69.9/68:0.0198409211)94.1/88:0.0435573597,((((((((WP_169105633.1-Clostridium-sp.-SM-530-WT-3G:0.0485300936,WP_027636628.1-Clostridium-butyricum:0.0797335189)20.1/28:0.0202038109,WP_077866663.1-Clostridium-saccharobutylicum:0.1679006714)87.3/95:0.0292197191,(((WP_261829154.1-Clostridium-sp.-TW13:0.0692922332,(WP_077855754.1-Clostridium-beijerinckii:0.0310521808,WP_041894754.1-Clostridium-beijerinckii:0.0246741574)99.1/100:0.0595924653)85.1/98:0.0149822006,WP_077848125.1-Clostridium-puniceum:0.1145038677)66.5/81:0.0088940225,((WP_185652834.1-Clostridium-sp.-DJ247:0.0199620071,WP_185652641.1-Clostridium-sp.-DJ247:0.0572162820)88.5/100:0.0209287226,((WP_022750218.1-Clostridium-saccharobutylicum:0.0367279533,(WP_015391310.1-Clostridium-saccharoperbutylacetonicum:0.0549117734,(WP_077846315.1-Clostridium-puniceum:0.0419654885,WP_238915641.1-Clostridium-sp.-YIM-B02555:0.0506489309)86/98:0.0140194519)86.5/98:0.0190723298)16.2/96:0.0194673821,WP_226126241.1-Clostridium-algoriphilum:0.1151781740)60.2/90:0.0121881962)85.7/94:0.0200168764)37.5/75:0.0131083718)71.8/8:0.0094226971,WP_224033465.1-Clostridium-gelidum:0.0808195647)59.9/18:0.0058453223,((WP_265859166.1-Clostridium-botulinum:0.0969753867,(WP_009169009.1-Clostridium-sp.-DL-VIII:0.0471021898,WP_077856313.1-Clostridium-beijerinckii:0.0132847957)95.1/64:0.0462002558)48.9/59:0.0273042648,WP_154531076.1-Inconstantimicrobium-porci:0.1022676779)71.1/51:0.0225739539)94.8/95:0.0442747973,(((WP_009169246.1-unclassified-Clostridium:0.0839482407,WP_077843022.1-Clostridium-beijerinckii:0.1170843005)89.1/100:0.0344032852,WP_041896941.1-Clostridium-beijerinckii:0.1411790647)93.1/100:0.0481816261,(WP_069680896.1-Clostridium-taeniosporum:0.1625047235,WP_160359894.1-Clostridium-chromiireducens:0.1954828845)31.4/62:0.0331238722)54.4/93:0.0261785882)82.4/93:0.0266181971,WP_079421587.1-Clostridium-oryzae:0.2713130871)75.8/92:0.0510157949,WP_216390068.1-Clostridium-sp.-MSJ-8:0.1521966831)90.4/93:0.0351343427)80.2/89:0.0379682252,(((WP_227847393.1-Clostridioides-sp.-ZZV15-6598:0.0889809068,WP_227846117.1-Clostridioides-sp.-ES-S-0145-01:0.0267676115)94.8/100:0.0385659419,(WP_227842647.1-unclassified-Clostridioides:0.0285103120,WP_003420982.1-Clostridioides-difficile:0.0763419778)85.7/100:0.0176613834)91.4/100:0.0332045570,WP_038810653.1-Clostridioides-difficile:0.0210264316)100/100:0.2273997113)82.4/57:0.0298346990,(WP_009529486.1-Peptoanaerobacter-stomatis:0.1311210292,WP_068913445.1-Criibacterium-bergeronii:0.1366173794)100/100:0.1860747967)54.7/52:0.0213850927,(((((WP_068555806.1-Thermotalea-metallivorans:0.0950185979,WP_073236503.1-Desulforamulus-putei:0.0877786434)84.8/100:0.0270728108,((WP_035164300.1-Caloranaerobacter-azorensis:0.0445817676,((WP_069974893.1-Geosporobacter-ferrireducens:0.2638940099,((WP_223038819.1-Crassaminicella-profunda:0.0884121397,WP_148809648.1-Crassaminicella-thermophila:0.0675466502)40.3/100:0.0343787313,WP_218283043.1-Crassaminicella-sp.-143-21:0.2030737126)20.5/95:0.0521349951)86.3/100:0.0655028769,WP_250858284.1-Oceanirhabdus-seepicola:0.1612364128)98.1/100:0.1071566424)66.5/84:0.0311090507,WP_095133872.1-Anaeromicrobium-sediminis:0.1436703964)41.7/83:0.0277108358)75.6/72:0.0121665200,WP_090442301.1-Natronincola-peptidivorans:0.1266804073)94/91:0.0365035583,(((WP_252130283.1-Clostridium-sp.-SYSU_GA19001:0.2067425884,WP_202749105.1-Clostridium-sp.-YIM-B02515:0.1060639213)98.2/100:0.0997029759,WP_216462460.1-Clostridium-bornimense:0.1263742838)4.1/36:0.0101906032,((((WP_104410444.1-Clostridium-algidicarnis:0.0521519728,WP_185162668.1-Clostridium-algidicarnis:0.0425157183)99.8/100:0.1288462269,WP_234123145.1-Clostridium-hydrogenum:0.1481613026)87.9/72:0.0354289823,((WP_084114600.1-Clostridium-acidisoli:0.0830731141,(WP_015614219.1-Clostridium-pasteurianum:0.0190303443,(WP_010235680.1-Clostridium-arbusti:0.0109554780,WP_003443690.1-Clostridium-pasteurianum:0.0448778542)91.6/100:0.0256724445)95.1/100:0.0301049735)11.3/71:0.0078658662,WP_026884435.1-Clostridium-akagii:0.0873760463)96.2/100:0.0565138501)16.3/71:0.0185252326,((WP_090038599.1-Clostridium-frigidicarnis:0.1202215380,WP_209459870.1-Youngiibacter-multivorans:0.1582230844)91/100:0.0586179034,((WP_164955553.1-Clostridium-estertheticum:0.1486806680,WP_125002390.1-Clostridium-tagluense:0.1236182443)76.1/96:0.0151484255,(WP_226136064.1-Clostridium-estertheticum:0.0236497051,WP_216146875.1-Clostridium-frigoris:0.0608151779)99.6/100:0.0836961033)98.9/100:0.0925350629)90.2/99:0.0387470619)55.3/48:0.0244718480)94.9/91:0.0450447319)82.1/37:0.0164930113,(((WP_165412483.1-Clostridium-estertheticum:0.0566443990,WP_216302578.1-unclassified-Clostridium:0.0088340489)100/100:0.1572891353,((WP_204598070.1-Clostridium-pascui:0.1173524202,WP_216457288.1-Clostridium-simiarum:0.1103290028)48.3/90:0.0277651357,WP_027632043.1-Clostridium-hydrogeniformans:0.1113098044)69.8/97:0.0267440962)88.3/100:0.0360007555,(((((WP_066398085.1-Cytobacillus-eiseniae:0.1628228413,WP_213236459.1-Fusibacter-paucivorans:0.4195972754)86.1/54:0.0511215177,((WP_056034132.1-Paenibacillus-sp.-Leaf72:0.0495478754,WP_090798944.1-Paenibacillus-sp.-GP183:0.2015254372)40.4/9:0.0573198547,WP_197936429.1-Paenibacillus-sp.-GSMTC-2017:0.1071953414)73.8/12:0.0485928941)91.6/9:0.0392032562,(WP_076307499.1-Paenibacillus-odorifer:0.0491537427,((WP_042140168.1-Paenibacillus-sp.-FSL-P4-0081:0.0099839911,WP_036721118.1-Paenibacillus-sp.-FSL-R7-277:0.0612900338)89.3/100:0.0305699036,WP_074087222.1-Paenibacillus-sp.-P32E:0.0288452586)93.7/100:0.0685120193)99.3/100:0.0966154202)59.2/8:0.0190538716,WP_262684732.1-Paenibacillus-sp.-WQ-127069:0.1606821035)58.5/62:0.0293224726,WP_036935152.1-Pseudobacteroides-cellulosolvens:0.1347102953)94.6/62:0.0508866990)95.4/61:0.0468807867)81.3/25:0.0241202826)92.5/94:0.0366375984,(((((((((((WP_058965344.1-Fournierella-massiliensis:0.1449247152,WP_185752312.1-Butyrivibrio-sp.-AE2032:0.2381709382)30.6/94:0.0490981755,WP_155202396.1-Ruthenibacterium-lactatiformans:0.1585316766)76.6/100:0.0579690824,(WP_186886939.1-Anaerofilum-hominis:0.1382185388,WP_102049807.1-Pygmaiobacter-massiliensis:0.1776424080)79.6/98:0.0685809157)98.7/100:0.1247224160,(((WP_072827923.1-Fibrobacter-sp.-UWB12:0.0231797028,WP_073304542.1-Fibrobacter:0.1676199774)23.6/17:0.0176271142,WP_088630837.1-unclassified-Fibrobacter:0.0182342237)86/99:0.0724656586,(((WP_097034884.1-Fibrobacter-sp.-UWT3:0.0363826278,WP_073056819.1-unclassified-Fibrobacter:0.0203450351)43.2/8:0.0279345102,WP_073155408.1-Fibrobacter:0.0345305696)7.4/7:0.0265374701,WP_073188830.1-Fibrobacter-sp.-UWP2:0.0029967997)93/93:0.0850679661)99.9/100:0.1715213934)89.8/75:0.0713227132,((WP_266162911.1-Caproicibacterium-sp.-BJN0003:0.1285300622,WP_256135338.1-Bittarella-massiliensis:0.2043439692)87.8/98:0.0777399711,WP_220321633.1-Clostridium-tagluense:0.2020559128)73.7/94:0.0559806439)48.2/58:0.0379376206,(WP_105204772.1-Neobittarella-massiliensis:0.1145234840,WP_022475313.1-Huintestinicola-butyrica:0.2101859972)97.9/76:0.0755303690)79.6/89:0.0217265487,((((WP_091684654.1-Anaerocolumna-aminovalerica:0.1464107991,(WP_033168390.1-Clostridium-sp.-KNHs205:0.0776554076,((WP_073588199.1-Anaerocolumna-xylanovorans:0.0313649504,WP_185256524.1-Anaerocolumna-chitinilytica:0.1086446011)67/100:0.0367525969,WP_073279502.1-Anaerocolumna-jejuensis:0.0914781488)71.9/100:0.0332221159)99.9/100:0.1303549985)98.4/100:0.0823386554,((((WP_060930762.1-Lachnoanaerobaculum-saburreum:0.1472399377,((((((((((((((((WP_024837362.1-Lacrimispora:0.0814429402,WP_077612318.1-Clostridium-sp.-Marseille-P2415:0.0328415313)0/97:0.0000024546,(WP_024293365.1-Lachnospiraceae:0.0338633092,WP_092242699.1-Lacrimispora:0.0225650202)95.9/100:0.0337062665)82.7/97:0.0161596197,((WP_124065139.1-Clostridium-sp.-E02:0.0762927141,WP_117415169.1-Clostridium-indicum:0.0420821303)73.5/90:0.0092882229,WP_143321893.1-Clostridium-sp.-HBUAS56010:0.0259397271)81.6/89:0.0153636979)22.1/65:0.0109587128,WP_138313639.1-Hungatella-hathewayi:0.0249127897)86.6/85:0.0183107983,WP_186424215.1-Lacrimispora-celerecrescens:0.1212579482)94.1/92:0.0379157413,((WP_227052204.1-Anaerostipes-caccae:0.0330606737,WP_137330070.1-Anaerostipes-rhamnosivorans:0.0632507914)99.5/100:0.0706702267,WP_006781439.1-Hungatella-hathewayi:0.0716423236)81.1/99:0.0134726347)23.9/29:0.0196002215,(WP_215718428.1-Anaerostipes-hadrus:0.0109647665,WP_075760385.1-Anaerostipes-sp.-494a:0.0461428379)99.7/100:0.0611123724)84.7/8:0.0156350312,((((WP_231062061.1-Lacrimispora-sp.-NSJ-141:0.0869926779,WP_249304640.1-Qiania-dongpingensis:0.0508045789)95.6/99:0.0531784675,WP_118541387.1-unclassified-Clostridium:0.0622444107)26.5/54:0.0116848379,WP_215700061.1-Clostridium-sp.-MCC353:0.0595110618)66.4/20:0.0046956281,(WP_024736308.1-Enterocloster-asparagiformis:0.0397297700,WP_066545911.1-Clostridium-sp.-AT4:0.0252711531)90.8/90:0.0200170204)73.2/7:0.0055285536)94.6/94:0.0281810767,WP_118243934.1-Clostridium-sp.-AM58-1XD:0.0450875301)89.5/46:0.0168838027,(WP_087223935.1-Lachnoclostridium-sp.-An14:0.0382354323,WP_101876396.1-Lachnoclostridium-edouardi:0.0442534240)95.5/100:0.0312531278)0/5:0.0000025715,WP_118742831.1-Clostridium-sp.-AM28-20LB:0.0284236358)84.2/7:0.0112664985,WP_008399284.1-unclassified-Clostridium:0.0171830759)76.7/8:0.0077224708,((WP_045093538.1-Clostridium-sp.-FS41:0.0415876822,WP_025485467.1-Lachnospiraceae:0.0514962569)17.8/100:0.0126683595,((WP_154471253.1-Clostridium-porci:0.1078591971,WP_178313416.1-Lacrimispora-sp.-210928-DFI.3.58:0.0023868723)51.2/54:0.0144596639,WP_002577448.1-Enterocloster:0.0636899123)77.6/52:0.0098519861)90.4/93:0.0323660751)82.8/91:0.0308818562,(WP_003497689.1-_Clostridium:0.0402460429,WP_125142063.1-Clostridium-transplantifaecale:0.0404167167)79.1/100:0.0144058084)18.7/94:0.0311765599,WP_251497048.1-Otoolea-sp.-DSM-111138:0.1106552387)78.3/100:0.0345104819,WP_118555425.1-Lachnotalea-sp.-AF33-28:0.1548567835)82.5/100:0.0521740118)95.1/100:0.0577270540,WP_005539496.1-Johnsonella-ignava:0.1651436839)82.8/99:0.0493249660,((((((((((WP_035764434.1-Butyrivibrio-sp.-NC2002:0.0764359233,(((WP_026526812.1-Butyrivibrio-sp.-VCD2006:0.1299429574,WP_026491547.1-unclassified-Butyrivibrio:0.0164638805)0/75:0.0274610894,WP_044913869.1-Butyrivibrio-sp.-WCE2006:0.0257005338)93.5/100:0.0597287583,WP_091822189.1-Butyrivibrio-sp.-ob235:0.0599726133)0/27:0.0272847753)98.7/100:0.0867685811,((((((WP_022768176.1-unclassified-Butyrivibrio:0.0222778582,WP_022766545.1-Butyrivibrio-sp.-XPD2006:0.0441821644)98.6/100:0.0752280197,WP_034446055.1-Butyrivibrio-sp.-AE2032:0.0606674869)83.5/98:0.0217070482,WP_071175335.1-Butyrivibrio:0.0487560746)46.8/53:0.0346980484,WP_024867291.1-Butyrivibrio-sp.-FCS014:0.0340430276)50.2/52:0.0282286184,WP_026513668.1-Butyrivibrio-sp.-LB2008:0.0520187202)75.5/99:0.0129232553,(WP_026663187.1-Butyrivibrio-proteoclasticus:0.0644953495,(WP_026652320.1-Butyrivibrio-proteoclasticus:0.0379470194,(WP_013279951.1-Butyrivibrio-proteoclasticus:0.0231943886,WP_092060841.1-Butyrivibrio-sp.-INlla14:0.0447451827)73.7/96:0.0141979744)75.5/95:0.0071972512)1/15:0.0124279600)96.4/100:0.0616312400)99.7/100:0.1124155982,(WP_109624131.1-Murimonas-intestini:0.0659467708,WP_178047055.1-Fusicatenibacter-sp.-CLA-AA-H277:0.0844712083)86/81:0.0241246265)11.4/71:0.0098902918,(WP_230400100.1-Novisyntrophococcus-fermenticellae:0.1499393932,((WP_173751851.1-Blautia-caecimuris:0.0437516634,WP_117997894.1-Blautia-obeum:0.0119518273)99.8/100:0.0676893290,(((WP_158367551.1-Lachnospiraceae:0.0297710014,WP_235247881.1-Faecalicatena-contorta:0.0655782287)87.3/100:0.0335488283,(WP_251390531.1-Mediterraneibacter-sp.-DSM-111629:0.0360389401,WP_073962323.1-_Ruminococcus:0.0231693048)88.1/100:0.0225237199)84.9/98:0.0164053323,WP_070089756.1-Merdimonas-faecis:0.0279514203)78.1/98:0.0109385968)74.4/90:0.0068259362)84.8/83:0.0174747998)82.6/89:0.0149069613,WP_132379639.1-Muricomes-intestini:0.0836958550)87.5/92:0.0187857369,(((WP_019160740.1-Blautia:0.0546131813,WP_173738571.1-Blautia-schinkii:0.0217227186)93.4/100:0.0279025406,(WP_158421413.1-Blautia:0.0333658323,(WP_148445023.1-Eubacteriales:0.0242812184,WP_187566223.1-Blautia-faecis:0.0439745835)92.4/100:0.0239111090)60/100:0.0132468403)99.8/100:0.0854462887,(WP_099466994.1-Konateibacter-massiliensis:0.0503605696,WP_151145296.1-Candidatus-Galacturonibacter-soehngenii:0.0599792026)98.6/100:0.0770733877)81.7/90:0.0299971141)90.4/93:0.0257939409,WP_087156947.1-Lachnoclostridium-sp.-An181:0.0530035109)86.9/94:0.0241437122,WP_073994331.1-Parasporobacterium-paucivorans:0.1387128064)81.9/92:0.0126394736,(WP_158351687.1-Diplocloster-modestus:0.1804730654,WP_115483253.1-Anaerosacchariphilus-polymeriproducens:0.0759489286)21.3/56:0.0249562980)94.8/98:0.0518204178,WP_158346166.1-Lachnospiraceae:0.1731893956)97.7/100:0.0902563884)72.4/93:0.0284081914,WP_226914794.1-Intestinibacter-bartlettii:0.2253888496)88.6/83:0.0354991734)78.3/81:0.0291304969,((((WP_073273890.1-Anaerocolumna-jejuensis:0.1299835092,WP_033167869.1-Clostridium-sp.-KNHs205:0.1300911906)8.6/69:0.0399577573,(((WP_010268185.1-Paenibacillus-senegalensis:0.1624619489,WP_155699987.1-Paenibacillus-psychroresistens:0.0505838517)62.8/47:0.0301934849,WP_090040610.1-Clostridium-frigidicarnis:0.0231003001)86.7/47:0.0251552784,WP_202751253.1-Clostridium-sp.-YIM-B02515:0.0343927817)93.8/81:0.0531668267)99.8/100:0.1388362663,(WP_249305773.1-Jutongia-hominis:0.1401216030,(WP_022463459.1-Jutongia-huaianensis:0.0670909369,(WP_055944671.1-unclassified-Clostridium:0.0364133812,WP_216564272.1-Eubacterium-sp.-MSJ-13:0.0259387152)95/100:0.0436305436)66.3/100:0.0399113013)99.7/100:0.1434417710)81.4/100:0.0318409761,(WP_253721803.1-Treponema-sp.-OMZ-787:0.2826098609,WP_058352784.1-Acetivibrio-ethanolgignens:0.1714184908)69.5/98:0.0552286852)95.8/97:0.0638150257)78.9/97:0.0269142770,WP_078711972.1-Anaerorhabdus-furcosa:0.4649408173)98.3/26:0.0905410210)81.8/25:0.0402292802,WP_138788932.1-Paenibacillus-thermoaerophilus:0.3736899263)49/58:0.0272178869,WP_160686059.1-Clostridium-sp.-C2-6-12:0.1944343134)79.5/49:0.0180296727,WP_119970050.1-Clostridium-fermenticellae:0.1938359382)89.3/52:0.0353184717,((((((((WP_088538685.1-Methanobrevibacter-sp.-87.7:0.0549651630,WP_016359256.1-Methanobrevibacter-sp.-AbM4:0.0984432302)97.8/100:0.0792830435,WP_067148181.1-Methanobrevibacter-olleyae:0.1009776958)85.3/100:0.0315629138,WP_116670296.1-Methanobrevibacter-woesei:0.1845579155)99.8/97:0.1067248008,(WP_248612240.1-Methanobacterium-alcaliphilum:0.1130889185,((WP_071907175.1-Methanobacterium-congolense:0.0209394402,WP_209584978.1-Methanobacterium-aggregans:0.0371046584)99.3/100:0.0926120276,(WP_069583666.1-Methanobacterium:0.1399905328,WP_048190917.1-Methanobacterium-sp.-SMA-27:0.1097901203)94.1/97:0.0535314445)67.7/73:0.0136722205)86.8/92:0.0227287229)59.4/86:0.0117243729,(WP_011406138.1-Methanosphaera:0.1239533972,(WP_112123902.1-Methanosphaera-sp.-BMS:0.0845869329,(WP_069592847.1-Methanosphaera-sp.-WGK6:0.0297749778,WP_095608065.1-Methanosphaera-cuniculi:0.0893415030)98/100:0.0675999041)54.6/99:0.0370158846)94.7/100:0.0595189429)87.8/90:0.0287201073,(WP_013644504.1-Methanobacterium-lacus:0.1583363465,WP_054834988.1-Methanobrevibacter-arboriphilus:0.1481882380)85.2/97:0.0287047494)91.4/97:0.0398619807,(WP_112093115.1-Methanothermobacter-tenebrarum:0.2005257486,WP_013413362.1-Methanothermus-fervidus:0.1596530009)99.3/99:0.1221739028)83.9/93:0.0410407676,(WP_038266426.1-Peptoclostridium-litorale:0.2638508869,(WP_111570784.1-Halanaerobium-saccharolyticum:0.0704102254,(WP_089717478.1-Halanaerobium-congolense:0.0471188342,WP_089862761.1-Halanaerobium-salsuginis:0.0251216872)84.4/100:0.0280977771)100/100:0.1810103880)90.5/58:0.0537248218)82.5/33:0.0254519128)0/4:0.0159857906)98.3/96:0.1007080865)97.2/97:0.1049638609,((((WP_018154793.1-Methanothermococcus-thermolithotrophicus:0.1001995411,WP_048201153.1-Methanocaldococcus-bathoardescens:0.0627267860)90.2/98:0.0396974319,((WP_052562942.1-Candidatus-Brocadia-sinica:0.2655416818,WP_173298646.1-Thermanaeromonas-sp.-C210:0.2199927973)67.1/99:0.0576953951,(((WP_011869065.1-Methanococcus-maripaludis:0.0776164704,WP_013998786.1-Methanococcus-maripaludis:0.0460691953)79.9/100:0.0294765296,WP_181493664.1-Methanococcus-maripaludis:0.0619342269)82.9/100:0.0243878197,WP_181504225.1-Methanococcus-maripaludis:0.0345700509)99.9/100:0.1652267775)71.1/98:0.0502781496)96.4/100:0.0741855956,(((WP_245533116.1-Coprothermobacter-platensis:0.2973895189,(WP_119360783.1-Meiothermus-luteus:0.0812133174,(WP_208013758.1-Meiothermus-sp.-CFH-77666:0.0995457316,WP_027877894.1-Meiothermus-cerbereus:0.0530356156)83.4/100:0.0722919789)100/100:0.3686250124)93.3/74:0.1072015568,(WP_150071182.1-Thermodesulfobacterium-sp.-TA1:0.1654135086,WP_022856039.1-Thermodesulfobacterium:0.1820686657)50.8/74:0.0427108047)84.4/98:0.0742897509,(((((WP_011996389.1-Coxiella-burnetii:0.4575255200,WP_013638952.1-Desulfurobacterium-thermolithotrophum:0.0300788832)58.9/100:0.0123214545,WP_132526207.1-Phorcysia-thermohydrogeniphila:0.0695133747)86.7/100:0.0225886125,WP_089322203.1-Desulfurobacterium-atlanticum:0.1236398654)93.5/100:0.0526797213,(WP_029522925.1-Persephonella-sp.-KM09-Lau-8:0.0959731966,(WP_200674641.1-Persephonella-atlantica:0.0657548069,WP_096999769.1-Persephonella-hydrogeniphila:0.0901411347)90.7/98:0.0498629364)84.5/98:0.0547517065)30.7/94:0.0316776569,WP_028950938.1-Sulfurihydrogenibium-subterraneum:0.0912142758)98/100:0.0906246031)92.5/97:0.0797319072)80.8/99:0.0672469153,((WP_010868297.1-Pyrococcus-abyssi:0.0136021574,((((((WP_013798345.1-Methanotorris-igneus:0.0283447467,WP_013866394.1-Methanothermococcus-okinawensis:0.1492354795)69.8/100:0.0233033300,WP_007044577.1-Methanotorris-formicicus:0.0589338889)94.2/100:0.0490464530,WP_148120035.1-Methanofervidicoccus-sp.-A16:0.1395520875)20.4/59:0.0264032694,((WP_167913028.1-Thermococcus-sp.-21S7:0.0647045153,WP_048150691.1-Palaeococcus-ferrophilus:0.0228751047)97.5/100:0.0398803822,((((WP_088855609.1-Thermococcus-siculi:0.0483165166,WP_087036111.1-Thermococcus-litoralis:0.0296160206)83.9/98:0.0157941473,WP_167888701.1-Thermococcus-sp.-MV5:0.0630077384)90.4/93:0.0179497768,(WP_013468354.1-Thermococcus-barophilus:0.0453150606,WP_013906489.1-Pyrococcus-yayanosii:0.0177345714)85.9/95:0.0114992092)88.2/96:0.0117527911,WP_048053121.1-Pyrococcus-horikoshii:0.0235420203)81.9/96:0.0112918180)81.4/78:0.0158092435)75.5/60:0.0113134327,(((((((WP_237702072.1-Thermococcus-sp.-AM4:0.0291390896,WP_223211916.1-Thermococcus-henrietii:0.0549488250)96.2/100:0.0559448882,WP_209475361.1-Thermococcus-stetteri:0.0686312738)58.8/51:0.0126202983,((((WP_088865724.1-Thermococcus-barossii:0.0517552699,(((WP_139679991.1-Thermococcus-indicus:0.0231779664,WP_148882943.1-Thermococcus-aciditolerans:0.0342410829)91.9/57:0.0267232967,WP_258085050.1-Thermococcus-sp.-813A4:0.0309378211)39.1/36:0.0109961173,WP_058938040.1-Thermococcus-celericrescens:0.0301107705)75.1/36:0.0082389373)75/35:0.0063899577,WP_167711671.1-unclassified-Thermococcus:0.0225838056)86.2/5:0.0110693259,WP_088181368.1-Thermococcus-sp.-5-4:0.0467074774)74.3/4:0.0048769241,WP_232460978.1-Thermococcus-radiotolerans:0.0469811863)93.5/91:0.0238841409)79.5/30:0.0127357421,(WP_152879125.1-Thermococcus-sp.-101-C5:0.0485485111,WP_015850310.1-Thermococcus-sibiricus:0.0424790720)75.6/98:0.0065712150)83.1/41:0.0108479987,WP_253304977.1-Thermococcus-aggregans:0.0261231887)85.1/87:0.0119589964,WP_088862340.1-Thermococcus-celer:0.0784374614)55.3/53:0.0097766288,WP_068664737.1-Thermococcus-piezophilus:0.0499446001)91/80:0.0225328415)93.4/94:0.0376769744,WP_014733609.1-Pyrococcus-sp.-ST04:0.0837694429)89.5/95:0.0364886523)94.5/99:0.0601777745,WP_048103006.1-Aciduliprofundum-boonei:0.1577449185)96.7/100:0.1100852730)99.5/100:0.1320585582)81.2/93:0.0677871072,WP_236098952.1-Dethiosulfovibrio:0.3264887098)99.2/99:0.1812581764,((((((((((((WP_160564586.1-Apibacter-sp.-B2912:0.0956640407,(WP_055424523.1-Apibacter-mensalis:0.0774461520,(WP_160568957.1-unclassified-Apibacter:0.0150227916,WP_166004639.1-Apibacter-sp.-B2966:0.0407967007)99.1/100:0.0762553001)68.7/100:0.0449391213)98.3/100:0.0889598210,(WP_105245553.1-Apibacter-adventoris:0.1591275085,WP_146261858.1-Apibacter-muscae:0.0961648166)88.1/100:0.0399536214)100/100:0.1647598605,(WP_084015419.1-Moheibacter-sediminis:0.2555060790,WP_182042771.1-Moheibacter-lacus:0.1961289221)93.9/100:0.0746278684)79.1/100:0.0498947635,(WP_068596492.1-Vaginella-massiliensis:0.1288830894,((WP_230477342.1-Faecalibacter-bovis:0.0922117800,WP_125349156.1-Empedobacter-falsenii:0.0879098344)83.7/98:0.0299704181,(WP_125113420.1-Algoriella-xinjiangensis:0.0720796716,WP_092908829.1-Algoriella-xinjiangensis:0.0333129902)95/100:0.0409944614)99.1/98:0.0946920516)97.2/100:0.0915701984)91.9/100:0.0615667945,(((((WP_014783622.1-Aequorivita-sublithincola:0.3138474011,WP_199156598.1-Pedobacter-sp.-ASV2:0.0669790320)87.3/54:0.0409701771,((WP_230143398.1-unclassified-Pedobacter:0.0901183200,WP_215166022.1-Pedobacter-sp.-ISL-64:0.0775067810)92.9/100:0.0682251778,WP_133260761.1-Pedobacter-changchengzhani:0.1004169499)89.8/100:0.0456772293)75/54:0.0264063458,WP_109929147.1-Pedobacter-paludis:0.0408998266)96.8/54:0.0483945711,((WP_123203876.1-Pedobacter-jejuensis:0.0693095512,WP_188416210.1-Pedobacter-mendelii:0.0269265088)0/7:0.0136423432,WP_056087060.1-Pedobacter-sp.-Leaf176:0.0463066481)76.5/93:0.0121846808)44.4/52:0.0252692237,WP_056872627.1-Pedobacter-sp.-Leaf194:0.0573788991)100/100:0.2517210099)81.2/73:0.0833023511,WP_111126400.1-Mesonia-sp.-K7:0.4727426240)42.3/84:0.0314404366,((WP_107809536.1-Elizabethkingia-sp.-YR214:0.1163177574,WP_219971314.1-Chryseobacterium-sp.-LJ756:0.1126763567)36.4/99:0.0181693483,(WP_076395614.1-Chryseobacterium:0.0398656411,WP_088467981.1-Chryseobacterium-sp.-VAUSW3:0.1356282891)64.3/91:0.0380994409)100/100:0.1523532300)94/97:0.0707829849,((((WP_233378072.1-Maribellus-sp.-CM-23:0.0518092922,WP_247978154.1-Maribellus-sp.-YY47:0.0681485751)98.1/100:0.1052892333,(WP_167609329.1-Maribellus-sediminis:0.1353537568,(WP_045033348.1-Draconibacterium-sediminis:0.0255830404,WP_163345719.1-Draconibacterium-halophilum:0.0670415600)99.1/100:0.1160042522)93.7/100:0.0654537654)99.9/100:0.2001036737,(((WP_106543043.1-Prolixibacter-denitrificans:0.1002732067,WP_025865917.1-Prolixibacter-bellariivorans:0.0193073044)0/41:0.0021596080,WP_153632016.1-Prolixibacter-sp.-SD074:0.0490270040)99.9/100:0.1833655736,((WP_159517929.1-Sunxiuqinia-indica:0.0935269279,WP_053182762.1-Sunxiuqinia-dokdonensis:0.0808128909)79.9/100:0.0535969556,WP_133466511.1-Sunxiuqinia-elliptica:0.1187735805)99/100:0.1143549458)64.2/100:0.0539165299)65.8/99:0.0512764388,((WP_068701711.1-Paludibacter-jiangxiensis:0.2602946344,((WP_011500232.1-Methanococcoides-burtonii:0.1620498219,WP_135611779.1-Methanococcoides-sp.-AM1:0.1438991119)69/100:0.0604450799,((((WP_128503182.1-Methanosarcina-sp.-MSH10X1:0.1338737821,(WP_011033115.1-Methanosarcina-mazei:0.0709300075,WP_048134921.1-Methanosarcina-sp.-1.H.T.1A.1:0.0376141336)32/100:0.0321288202)62/99:0.0457241258,WP_048167026.1-Methanosarcina:0.0703495585)71.1/53:0.0336452754,WP_048177163.1-Methanosarcina-sp.-MTP4:0.0737587360)24.7/53:0.0376106168,(WP_048137589.1-Methanosarcina-horonobensis:0.1265230028,((WP_048184631.1-Methanosarcina-siciliae:0.0511473165,WP_048173663.1-Methanosarcina-siciliae:0.0514127734)53.9/100:0.0107339262,WP_011023680.1-Methanosarcina-acetivorans:0.0311183449)96.1/100:0.0617542135)91.3/100:0.0594095349)83.8/99:0.0697377433)98.4/100:0.1333502074)58.8/97:0.0596795272,(((WP_101263396.1-Labilibaculum-filiforme:0.0366485496,WP_245870435.1-Labilibaculum-manganireducens:0.0394211735)98.5/100:0.0735738212,(WP_152734748.1-Marinifilum-sp.-N1E240:0.0674660143,(WP_171595603.1-Marinifilum-caeruleilacunae:0.0564893751,(WP_268189532.1-Marinifilum-sp.-D737:0.0460300953,WP_054720444.1-Marinifilum-fragile:0.0294152084)83.5/100:0.0181479147)85.8/100:0.0226514396)94.2/100:0.0470024307)36.1/95:0.0349957957,WP_127342456.1-Ancylomarina-longa:0.0996535898)99.7/100:0.1472425523)74.5/86:0.0385808952)97.9/99:0.1013442255)48.4/75:0.0354586924,(((WP_205722928.1-Fulvivirga-lutea:0.3149035216,(((((((((((WP_053470876.1-unclassified-Flavobacterium:0.1341098018,WP_095952323.1-Flavobacterium-sp.-ACN6:0.0357943312)67.5/81:0.0063992593,WP_056200692.1-Flavobacterium-sp.-Root186:0.0297685821)86.4/81:0.0113289631,WP_115889823.1-Flavobacterium-cutihirudinis:0.0471814903)32.4/100:0.0053947558,WP_198857739.1-Flavobacterium-sp.-IB48:0.0359023750)79/99:0.0077580225,(WP_269237486.1-Flavobacterium-sp.-IMCC34775:0.0719813822,WP_163393939.1-Flavobacterium-limi:0.0326740005)0/100:0.0000023782)75.4/99:0.0106103148,WP_264205577.1-Flavobacterium-sp.-LS1R49:0.0221479012)89/100:0.0171661541,((WP_173968237.1-Flavobacterium-collinsii:0.0655320274,WP_264525312.1-Flavobacterium-sp.-N502536:0.0403983886)94.1/100:0.0269937315,WP_070907856.1-Flavobacterium-tructae:0.0721657897)74.1/94:0.0062473458)71.6/93:0.0068989599,WP_091091738.1-Flavobacterium-gillisiae:0.0879508007)98.5/100:0.0509727097,WP_248907802.1-Flavobacterium-sp.-K5-23:0.1022828286)81.9/100:0.0292745501,(((WP_116761981.1-Flavobacterium-laiguense:0.0384296425,WP_143377960.1-Flavobacterium-sp.-ZT3R18:0.0417445903)98.5/100:0.0624099201,((WP_144071503.1-Flavobacterium-franklandianum:0.0575914657,WP_133277350.1-Flavobacterium-nackdongense:0.0777173135)47.5/100:0.0255042035,WP_194311471.1-Flavobacterium-soyangense:0.1051228619)75.2/92:0.0229715550)83.4/92:0.0262687228,WP_103806521.1-Flavobacterium-alvei:0.0783128462)88.4/94:0.0403212490)99.4/100:0.1253127721,(WP_140000461.1-Flavobacterium-microcysteis:0.1627966391,(((WP_194640005.1-Flavobacterium-sp.-HJJ:0.0770322507,WP_073246022.1-Flavobacterium-flevense:0.0575934233)80.1/100:0.0217930577,WP_100843428.1-Flavobacterium-sp.-5:0.0818189411)74.4/94:0.0424468472,WP_140487657.1-Flavobacterium-sp.-GSA192:0.0428656732)98.3/100:0.1277647824)94.9/100:0.0814009040)95.1/100:0.0868765970)56.5/68:0.0639088957,(WP_116872776.1-Terrimonas-sp.:0.2718301613,(WP_004155558.1-Microscilla-marina:0.2299006552,WP_101358295.1-Raineya-orbicola:0.1626729057)99.8/100:0.2078582934)88.9/44:0.0860840987)76.4/44:0.0454531271,WP_057952567.1-Salinivirga-cyanobacteriivorans:0.3768144111)87.6/41:0.0638173527)95.2/55:0.0783289510,((((WP_013633636.1-Pseudopedobacter-saltans:0.1059041403,(WP_160842504.1-Pedobacter-sp.-HMF7647:0.0464399834,(WP_074492796.1-Mucilaginibacter-polytrichastri:0.1137920424,(WP_258137696.1-Mucilaginibacter-sp.-BJC16-A38:0.1305182668,(WP_259068220.1-Mucilaginibacter-sp.-X4EP1:0.0520321219,WP_144912900.1-Mucilaginibacter-frigoritolerans:0.0685040016)87.6/99:0.0711969395)87.2/94:0.1037542177)99.8/100:0.1644202260)94.1/52:0.0666221192)67.8/52:0.0458004257,WP_154287284.1-Pedobacter-puniceum:0.0909284754)99.9/99:0.1987685254,WP_163286637.1-Cryomorpha-ignava:0.3313621167)2.2/95:0.0544829030,WP_130092062.1-Brumimicrobium-glaciale:0.2670504956)96.7/100:0.1198457646)93.2/60:0.0987865887,((WP_040596643.1-Prevotella-stercorea:0.0860666909,WP_207153370.1-Prevotella-herbatica:0.0671879639)99.2/100:0.1906687892,(WP_223927321.1-Prevotella-lacticifex:0.1706868175,WP_009010825.1-Prevotella-sp.-C561:0.1239224853)99.7/100:0.2425288291)88.4/99:0.0837800038)0/17:0.0485105700,(WP_085659180.1-Prosthecochloris-sp.-HL-130-GSB:0.2900881845,WP_038563105.1-Draconibacterium-orientale:0.2258482869)94.7/100:0.0991152310)98.8/97:0.1638797773)88.1/100:0.0617622242,(KAI6244242.1-Aphelenchoides-fujianensis:0.3265619709,KAI6170547.1-Aphelenchoides-bicaudatus:0.4073081704)97/100:0.1326499127)93.6/100:0.0703299481)35.2/33:0.0466529060)93.8/68:0.0600779990)90.2/25:0.0381331477)88.6/23:0.0358078475,(WP_083560927.1-Oceanococcus-atlanticus:0.2338087595,WP_104228621.1-Solimonas-fluminis:0.1946381166)98.8/76:0.1415897917)82.1/20:0.0199505593,(((((WP_148717329.1-Chitinolyticbacter-meiyuanensis:0.2965902921,WP_123210144.1-Stagnimonas-aquatica:0.3968274087)85.9/96:0.1058094568,(WP_176062241.1-Anaeromyxobacter-diazotrophicus:0.1240264064,WP_248361410.1-Anaeromyxobacter-oryzae:0.1766607471)97.8/100:0.0939900140)89/91:0.0514353351,(((WP_018882239.1-unclassified-Thioalkalivibrio:0.2654141764,WP_125198734.1-Guyparkeria-sp.-SCN-R1:0.1301648819)87.5/99:0.0624536185,(((WP_026296529.1-Thioalkalivibrio-sp.-ALJ2:0.1127501315,WP_019569965.1-Thioalkalivibrio-sp.-ALE11:0.0660551065)85.8/97:0.0206424836,WP_028484075.1-Thioalkalivibrio-sp.-ALE17:0.0350811634)80.4/96:0.0104116532,WP_018169543.1-Thioalkalivibrio-sp.-ALMg9:0.0244220316)98.7/98:0.0764171868)39.2/75:0.0282712616,WP_250072676.1-Guyparkeria-hydrothermalis:0.1494828611)98.4/99:0.1034534210)88.1/98:0.0411923047,WP_205202385.1-Azonexus-hydrophilus:0.1592059662)61.9/75:0.0504089365,WP_173581083.1-Azonexus-fungiphilus:0.2105508940)68.8/16:0.0152648722)81.6/66:0.0198821992)91.6/41:0.0339941979,((((WP_014706296.1-Methylophaga-nitratireducenticrescens:0.0269870278,WP_136552334.1-Methylophaga-sp.-SB9B:0.0375459593)100/100:0.2702248544,(WP_108623287.1-Orrella-marina:0.2661483461,WP_243478771.1-Orrella-sp.-f23:0.2339462752)82.1/100:0.0503207294)69.9/99:0.0511338604,WP_258826205.1-Massilia-agri:0.2596153352)36.5/7:0.0339654541,((WP_204103763.1-Spirulina-major:0.2941892378,(WP_044881819.1-Neochlamydia-sp.-EPS4:0.0515974748,WP_166155173.1-unclassified-Neochlamydia:0.0000024123)100/100:0.2855653884)10.3/6:0.0235231558,(WP_009207329.1-Sulfuricella-denitrificans:0.2249431894,(WP_002810067.1-Nitrosococcus-oceani:0.0691906486,WP_013221233.1-Nitrosococcus-watsonii:0.0252281918)99.6/100:0.1513620304)37.9/6:0.0350083251)86/5:0.0451668811)87.3/6:0.0309371601)15.5/4:0.0156584979,((((((((((((WP_212718705.1-unclassified-Synechocystis:0.0233652403,WP_228015966.1-Synechocystis-salina:0.0853591007)76.2/100:0.0307603993,WP_010873214.1-unclassified-Synechocystis:0.0295121622)100/100:0.2513736290,(((WP_099797666.1-Thermostichus-lividus:0.2789072782,WP_196167318.1-Pseudomonas-monteilii:0.1761671180)51.3/81:0.0681226462,WP_153720048.1-Spiribacter-sp.-C176:0.2211366239)76.9/95:0.0338853233,WP_048706771.1-Thauera-humireducens:0.1960933363)50.7/93:0.0476254266)90.9/95:0.0504143231,(WP_228215938.1-Aromatoleum-toluclasticum:0.1761298597,WP_199264031.1-Desulfobulbus-oligotrophicus:0.1841203810)86.9/51:0.0340948497)81.3/28:0.0173312849,((WP_040200185.1-Geoalkalibacter-subterraneus:0.1799134483,WP_227181580.1-Leeia-sp.-TBRC-13508:0.2433683259)5.8/17:0.0210069627,(((WP_006216420.1-Achromobacter-piechaudii:0.1050587819,WP_219991400.1-Achromobacter-sp.-ES-001:0.0606171197)84/100:0.0281812656,((WP_175192416.1-Achromobacter-deleyi:0.0646206091,WP_100503206.1-Achromobacter-ruhlandii:0.1176489581)77.5/86:0.0315442906,WP_183017169.1-Achromobacter-sp.-UMC46:0.0477143950)78.3/86:0.0156851613)100/99:0.1335155128,WP_091467897.1-Methylophilus-rhizosphaerae:0.2391552820)88.9/63:0.0460032346)36/33:0.0310737236)95.9/54:0.0520229956,(((((WP_238511579.1-Cupriavidus-gilardii:0.2303566761,((WP_213445339.1-Variovorax-sp.-PCZ-1:0.1993508769,WP_226921657.1-Kinneretia-sp.-XES5:0.1567827076)83.9/99:0.0613740436,WP_224042884.1-Cupriavidus-respiraculi:0.2009575330)5.8/63:0.0211230123)33.1/58:0.0243746704,WP_116600700.1-Cupriavidus-alkaliphilus:0.1336403327)9.7/25:0.0241606356,(WP_018008101.1-Cupriavidus-neocaledonicus:0.0350654203,WP_183298446.1-Cupriavidus-alkaliphilus:0.0493274148)99.3/100:0.1036392704)96.8/77:0.0642250882,(((WP_116346290.1-Cupriavidus-taiwanensis:0.0862498026,WP_124685047.1-Cupriavidus-pauculus:0.1887913169)99.9/100:0.2238015301,WP_226440716.1-Quatrionicoccus-australiensis:0.2104456072)41.8/61:0.0268024102,((WP_220807781.1-Noviherbaspirillum-aridicola:0.2858740718,WP_137423090.1-Desulfoglaeba-alkanexedens:0.2094066460)78.4/96:0.0555995210,WP_150984056.1-Cupriavidus-basilensis:0.2553724916)93.5/93:0.0870000899)58.4/86:0.0411696021)70.7/48:0.0330370510,(((((((((((((WP_209862330.1-Pseudomonas-sp.-BP8:0.2864070864,WP_225786622.1-Pseudomonas:0.0299994718)85.5/93:0.0404397021,(((((WP_054891693.1-Pseudomonas:0.0626888637,WP_236238170.1-Pseudomonas-faucium:0.1065116007)51.6/95:0.0238580445,WP_158461286.1-Pseudomonas-fluorescens:0.1660789587)17.5/95:0.0281389422,(WP_200141208.1-unclassified-Pseudomonas:0.1446525303,WP_029612831.1-Pseudomonas-parafulva:0.0224752168)94.3/100:0.0574283058)83.6/98:0.0415002019,WP_110605176.1-Pseudomonas-soli:0.1826487083)2.2/26:0.0281890631,(((WP_217887155.1-Pseudomonas-xanthosomae:0.0320579763,WP_217840807.1-Pseudomonas-fakonensis:0.0326433004)87.4/100:0.0258240176,WP_186654015.1-Pseudomonas-xantholysinigenes:0.1472465441)56.6/94:0.0235985782,(WP_201176141.1-Pseudomonas-sp.-S31:0.0314251560,WP_261240966.1-unclassified-Pseudomonas:0.0333285299)98.4/100:0.0783712356)60.6/73:0.0223538514)80.5/70:0.0184827956)97.5/100:0.0654111713,((WP_181290220.1-Pseudomonas-brassicacearum:0.1233934334,((((WP_201004194.1-Pseudomonas-cichorii:0.0673046027,WP_176569640.1-Pseudomonas-eucalypticola:0.0980761255)87.5/91:0.0396191578,(WP_219063768.1-Pseudomonas-sp.-UMAB-08:0.1508830874,((WP_222540780.1-Pseudomonas-sp.-DR208:0.1004350746,((WP_065875813.1-Pseudomonas:0.0356776855,(WP_033898696.1-Pseudomonas-fluorescens-group:0.0287712349,(WP_186687394.1-Pseudomonas-sp.-SWRI18:0.0452118553,WP_169378974.1-Pseudomonas:0.0134210543)84.2/100:0.0217800214)84/100:0.0110189670)98.9/100:0.0619158260,WP_105697656.1-Pseudomonas-poae:0.0334291918)83.2/69:0.0122539735)9.9/51:0.0107244082,(WP_253509974.1-Pseudomonas-rhodesiae:0.0918166216,WP_093202821.1-Pseudomonas-yamanorum:0.0258845578)84.7/97:0.0274512694)78.8/96:0.0302866766)89.6/97:0.0406380879)86.2/89:0.0240774611,(WP_249678359.1-Pseudomonas-sp.-PIA16:0.1142076972,WP_043192328.1-Pseudomonas-rhizosphaerae:0.1553948139)89.2/58:0.0391433863)63.3/31:0.0085210762,WP_268799018.1-Pseudomonas-sp.-BSw22131:0.1342394578)88.9/64:0.0373494058)90/96:0.0375207620,WP_108096457.1-Pseudomonas-sp.-GV071:0.0850910670)86.6/100:0.0504338130)84.9/100:0.0336517016,WP_252273535.1-Pseudomonas-sp.-YIM-B01952:0.0909137141)78.9/100:0.0211462323,WP_143488038.1-Pseudomonas-mangiferae:0.1113442210)94.4/100:0.0468898291,((((((WP_058067682.1-Pseudomonas-sp.-TTU2014-080ASC:0.1169379059,WP_077525673.1-Pseudomonas-resinovorans:0.0889402878)15.7/38:0.0190086117,WP_137821430.1-Pseudomonas-sp.-D-2018:0.0685418017)96.1/60:0.0545085025,WP_016495240.1-Pseudomonas-resinovorans:0.0584848308)79.1/19:0.0110190284,(WP_003453405.1-Pseudomonas:0.0937088746,WP_044873077.1-Pseudomonas-sp.-LFM046:0.0519698494)71.9/28:0.0202784767)80.4/32:0.0121461923,(WP_151138079.1-Pseudomonas-lalkuanensis:0.0424226198,WP_028626762.1-Pseudomonas-resinovorans:0.0338872070)13.6/65:0.0116193020)76/28:0.0096893318,WP_120654068.1-Pseudomonas-sp.-DY-1:0.0770046355)86.7/60:0.0200557379)31/39:0.0116028316,(WP_192399936.1-Pseudomonas-sp.-PDM15:0.1205088689,WP_226282979.1-Pseudomonas-lalkuanensis:0.0483755942)98.5/100:0.0634806240)93.6/97:0.0341115338,((((WP_090448460.1-Pseudomonas-benzenivorans:0.1418681278,(WP_233685494.1-Pseudomonas-anguilliseptica:0.0433949828,WP_257778987.1-Pseudomonas-sp.-32.2.56:0.0089931110)99.5/100:0.0841930847)83.9/87:0.0231903369,((WP_025164100.1-Pseudomonas-taeanensis:0.0471612316,WP_147170838.1-Pseudomonas-sp.-SJZ079:0.0056084456)94.3/100:0.0330743658,(WP_208707982.1-Pseudomonas-sp.-LPB0260:0.0215750106,WP_255840290.1-Pseudomonas-benzenivorans:0.0341591832)98.3/100:0.0573717977)60.5/82:0.0055880263)63.2/88:0.0203986484,(WP_069518708.1-Pseudomonas:0.1023681947,((WP_131198269.1-Pseudomonas-dryadis:0.0767144847,(WP_131178052.1-Pseudomonas-daroniae:0.0497762190,(WP_084307530.1-Pseudomonas-flavescens:0.0558622800,(WP_249909557.1-Pseudomonas-fulva:0.0368062476,(WP_074889242.1-Pseudomonas-argentinensis:0.0222252360,WP_027910132.1-Pseudomonas-sp.-URMO17WK12_I4:0.0375941136)63/98:0.0122653996)87.9/100:0.0282960180)78.9/97:0.0235940115)97.1/100:0.0557335033)96.7/99:0.0631325670,(((((((WP_119694090.1-Pseudomonas:0.0393153578,WP_106738315.1-Pseudomonas-mendocina:0.0432114220)92/100:0.0300224575,WP_150606948.1-Pseudomonas-oleovorans:0.0468962928)92.2/91:0.0316108504,WP_024309956.1-Pseudomonas-sp.-P818:0.0333867178)84/14:0.0136950753,(WP_143497094.1-Pseudomonas-sp.-ALS1131:0.0331262253,WP_090337209.1-Pseudomonas-chengduensis:0.0186343300)0/16:0.0002280249)86.3/21:0.0166510506,WP_074939507.1-Pseudomonas-composti:0.0186922171)87.1/54:0.0153561438,WP_021444205.1-Pseudomonas-sp.-EGD-AK9:0.0428871651)86.5/55:0.0207754757,WP_159970704.1-Pseudomonas-sp.-8Z:0.0652369380)95.8/100:0.0472185851)87.2/93:0.0283574272)73.9/46:0.0074658848)89.6/74:0.0241456553,WP_238040454.1-Pseudomonas-sp.-MMS21-TM103:0.0990738050)27.8/87:0.0079245221)93.6/99:0.0483079065,WP_220803742.1-Pseudomonas-sp.-NCCP-436:0.1055523536)70.2/54:0.0299272172,WP_090225506.1-Pseudomonas-kuykendallii:0.0850755882)91.1/97:0.0429855002,((WP_183165579.1-Azomonas-macrocytogenes:0.1279833560,(((WP_102836461.1-Stutzerimonas-stutzeri:0.0228170193,(WP_256071125.1-Stutzerimonas-stutzeri:0.0286352154,WP_045423249.1-Pseudomonadaceae:0.0230664441)85.9/100:0.0113636867)97.1/100:0.0404561851,(WP_255961233.1-Stutzerimonas-stutzeri:0.0229588992,WP_025243649.1-Stutzerimonas-stutzeri:0.0400096265)0/37:0.0009119793)95.3/100:0.0433368368,(WP_122077451.1-Pseudomonadaceae:0.0282272175,((WP_037024448.1-Pseudomonas-saudiphocaensis:0.0573096433,WP_181084609.1-Stutzerimonas-stutzeri:0.0057192381)94.8/100:0.0290519136,(((WP_011911332.1-Stutzerimonas-stutzeri:0.0388452806,(WP_237255234.1-Pseudomonas-oligotrophica:0.0514391326,WP_043295157.1-Stutzerimonas-degradans:0.0188420236)63.7/99:0.0285332138)81.8/77:0.0117924703,WP_120997453.1-_Pseudomonas:0.0284938867)41.5/26:0.0079722589,(WP_230358311.1-Pseudomonas-sp.-NP21570:0.0136932478,(WP_193680295.1-Pseudomonas-lopnurensis:0.0376780638,WP_200589398.1-Stutzerimonas-stutzeri:0.0191648563)90.8/100:0.0284413033)88.6/83:0.0204315371)94.1/85:0.0274260784)76.3/100:0.0067153801)13.5/96:0.0030426015)67.6/97:0.0296099072)91/100:0.0369938508,WP_028238613.1-Stutzerimonas-azotifigens:0.0487808934)99.8/100:0.0927611758)34.3/11:0.0184821701,((WP_119020982.1-Halomonas-sp.-JS92-SW72:0.2680674657,(((((WP_200222041.1-Achromobacter-xylosoxidans:0.1171312964,WP_259251071.1-Achromobacter-sp.-JUb104:0.1284134801)94.1/100:0.0588012767,(WP_088588627.1-Achromobacter-marplatensis:0.1186951266,WP_169273406.1-Achromobacter-sp.-Bel:0.0747227245)96.1/100:0.0645946947)77.2/97:0.0488617882,((((WP_111826915.1-Achromobacter-sp.-HZ01:0.0849506957,WP_102775120.1-Achromobacter:0.0173925605)99.8/100:0.1137222157,WP_043544564.1-Achromobacter-sp.-RTa:0.0302345537)29.5/55:0.0058223102,WP_129525645.1-Achromobacter-agilis:0.0305520356)86.1/55:0.0170595900,WP_129240460.1-Achromobacter-veterisilvae:0.0343774774)95.3/99:0.0585277278)0/10:0.0061345476,WP_174717299.1-Achromobacter-denitrificans:0.0741064822)87.8/72:0.0292251803,(WP_175208175.1-Achromobacter-anxifer:0.0351219287,((WP_088158656.1-Achromobacter-xylosoxidans:0.0781146222,WP_175164188.1-Achromobacter-aegrifaciens:0.0295293101)50.4/70:0.0173895273,(((WP_251864322.1-Achromobacter-sp.-Marseille-Q4962:0.0614367939,WP_164793551.1-Achromobacter-insolitus:0.0279718547)97.5/99:0.0466348703,WP_207472676.1-Achromobacter-sp.-SD115:0.0084937330)90.9/99:0.0292950067,WP_125471167.1-Achromobacter:0.0166637630)87.3/99:0.0326126192)68.8/71:0.0239634761)96.6/100:0.0667649772)94.4/99:0.0750404079)89/95:0.0613803714,(WP_084055900.1-Desulfacinum-hydrothermale:0.2121628312,WP_123289658.1-Desulfosoma-caldarium:0.1879054084)99.5/100:0.1415977818)74.5/72:0.0235521896)57.5/15:0.0055339338,WP_136680209.1-Neptunomonas-sp.-XY-337:0.1805397927)39.6/85:0.0291098474)82.5/53:0.0130231848)88.3/71:0.0215500708,((WP_217345508.1-Noviherbaspirillum-sp.-L7-7A:0.1404742984,WP_237247350.1-Sideroxyarcus-emersonii:0.1841349790)37/97:0.0357200628,WP_047549861.1-Methylotenera-sp.-G11:0.1752060990)95.9/98:0.0542969035)72.1/71:0.0077785298,(((((WP_198111724.1-Inhella-proteolytica:0.2554875602,((WP_037482909.1-Sphaerotilus-natans:0.1391193975,(WP_028996355.1-Azohydromonas-australica:0.0606266024,WP_066337818.1-Azohydromonas-lata:0.0791261121)96.6/100:0.0869675183)56.6/100:0.0698139476,WP_241396682.1-Pelomonas-sp.-CA6:0.1101018700)24.2/96:0.0336288518)95.7/100:0.0858834710,(WP_230291920.1-Croceicoccus-sp.-Ery5:0.2997904121,WP_164156863.1-Sandarakinorhabdus-rubra:0.1470772258)90.8/100:0.0747390535)98.9/99:0.1245051026,(WP_025281494.1-Ectothiorhodospira-haloalkaliphila:0.2896004061,WP_253536546.1-Ralstonia-sp.-21YRMH01-3:0.2357074231)73.6/87:0.0501133814)43.5/8:0.0280835238,WP_154206467.1-Ralstonia-pickettii:0.1986110991)77.3/9:0.0189445339,(((WP_036252969.1-Methylobacter-sp.-BBA5.1:0.1863326140,((((WP_019918777.1-Methyloversatilis-discipulorum:0.0375865183,WP_018229738.1-Methyloversatilis-universalis:0.0474928016)99.1/100:0.0917322246,((WP_101941536.1-Uliginosibacterium-sp.-TH139:0.0757181432,WP_095523004.1-Candidatus-Dactylopiibacterium-carminicum:0.0848761770)98.2/100:0.0984536877,WP_183409942.1-Litorivivens-lipolytica:0.3006631056)21.4/51:0.0329542661)99.4/100:0.1014736889,((((((WP_129151777.1-Achromobacter-aloeverae:0.0767885975,WP_088599355.1-unclassified-Achromobacter:0.0637236176)98.8/100:0.0965654491,WP_057654143.1-Bordetella-sp.-N:0.1312931352)74.4/76:0.0506342928,WP_086067427.1-Bordetella:0.1605086543)86.1/84:0.0305715612,(WP_066641528.1-Bordetella-sp.-H567:0.0549129280,(WP_086058993.1-Bordetella-genomosp.-9:0.1420239691,WP_094856450.1-Bordetella-genomosp.-10:0.1223331102)96.3/94:0.0818337293)88.9/93:0.0734325667)99.2/100:0.1139663819,WP_067106371.1-Methylobacillus-sp.-MM3:0.1599067258)88.3/87:0.0311695519,((WP_109041087.1-Azospira-sp.-I13:0.0869262717,WP_014237023.1-Azospira-oryzae:0.1032954468)96.7/100:0.0869457357,WP_201673630.1-Ramlibacter-monticola:0.1720171212)65.8/84:0.0327864961)90.3/97:0.0407178412)89.8/47:0.0368968899,(((WP_047193087.1-Schlegelella-brevitalea:0.1937826004,WP_019561187.1-Caldimonas-manganoxidans:0.1677822955)86.7/99:0.0839602925,WP_074865046.1-Atopomonas-hussainii:0.1433104365)91/99:0.0715693226,((WP_160809027.1-Nitrosomonas-sp.-HPC101:0.2289889029,WP_101249932.1-Telmatospirillum-siberiense:0.2762571046)81.1/96:0.0398614291,WP_220807643.1-Noviherbaspirillum-aridicola:0.1863822154)90.5/95:0.0678140955)64.3/86:0.0269413627)4.7/31:0.0078205811)92.6/82:0.0398469154,(((((WP_074796726.1-Nitrosospira-briensis:0.0618638268,(WP_090882638.1-Nitrosovibrio-sp.-Nv6:0.0474480018,WP_097065525.1-Nitrosovibrio-sp.-Nv4:0.0329600652)91.7/100:0.0481360324)100/100:0.1664829952,WP_122244186.1-Vandammella-animalimorsus:0.1764401030)91.3/83:0.0438463128,((WP_223986959.1-Cupriavidus-pampae:0.0529489576,((WP_011517306.1-Cupriavidus-metallidurans:0.0447517040,WP_019451253.1-Cupriavidus-sp.-BIS7:0.0449854372)0/33:0.0000026009,WP_082079449.1-Cupriavidus-basilensis:0.0702286378)76.3/64:0.0222612785)100/100:0.1188581976,WP_169420802.1-Ramlibacter-agri:0.1610671960)64.2/93:0.0170601767)74.7/45:0.0098109284,((((WP_026436726.1-Acidovorax-sp.-JHL-9:0.1288818162,WP_086926522.1-Acidovorax-carolinensis:0.1291031509)78.3/99:0.0387472653,WP_174484170.1-methanotrophic-endosymbiont-of-Bathymodiolus-puteoserpentis:0.1483275811)64.5/85:0.0373289785,WP_137890779.1-Ramlibacter-sp.-2FC:0.0940842683)77.8/31:0.0145954376,WP_137918285.1-Hydrogenophaga-sp.-2FB:0.1254656829)70.1/19:0.0084790694)91.4/38:0.0243914267,((((WP_106935816.1-unclassified-Variovorax:0.1819995257,(((WP_126747644.1-Variovorax-sp.-DXTD-1:0.1110760236,((WP_192194832.1-Variovorax-sp.-38R:0.1565696342,WP_160553990.1-Xylophilus-rhododendri:0.1231209881)25.9/100:0.0359751401,WP_093441791.1-Variovorax-sp.-770b2:0.1093210556)91.5/99:0.0496173189)81.2/83:0.0216691536,WP_068630051.1-Variovorax-sp.-PAMC-28711:0.1791503069)69.7/80:0.0138106305,(WP_093207264.1-Variovorax-sp.-YR750:0.0865671535,WP_206177103.1-Variovorax-sp.-RKNM96:0.0851627976)89.3/96:0.0321896524)90.5/92:0.0295043344)54.3/92:0.0213376252,WP_219210863.1-Variovorax-boronicumulans:0.1413472937)87.5/98:0.0380045025,WP_056898839.1-Pseudorhodoferax-sp.-Leaf274:0.1743944787)89.4/81:0.0372967942,WP_057673948.1-Curvibacter-sp.-PAE-UM:0.1351084538)80.2/41:0.0149171815)92/93:0.0273623964)69.5/84:0.0320396321,WP_212785890.1-Ferrigenium-kumadai:0.1772553734)57.3/10:0.0195297904)97.5/91:0.0484683709)30.1/69:0.0219166707,(((WP_070114691.1-Acidithiobacillus-caldus:0.2011603009,((WP_009566950.1-Acidithiobacillus:0.0421510373,WP_215870853.1-unclassified-Acidithiobacillus:0.0592276755)73.2/100:0.0401825943,(WP_101536718.1-Acidithiobacillus-sp.-SH:0.1455308501,WP_163095688.1-Acidithiobacillus-ferrianus:0.0254586749)37.5/85:0.0315735365)98.8/100:0.1065673733)88.1/100:0.0715526039,WP_152145882.1-Alkalilimnicola-sp.-S0819:0.1929639914)99/99:0.1232972360,((WP_108647113.1-Polynucleobacter-rarus:0.0191656834,WP_084281813.1-Polynucleobacter-kasalickyi:0.0366924906)100/100:0.3256050121,((WP_215371371.1-Polynucleobacter-sp.-MWH-UH24A:0.1484722604,WP_173942040.1-Polynucleobacter-antarcticus:0.1422060246)89.2/100:0.0716936262,((((WP_128113711.1-Polynucleobacter-necessarius:0.0927000995,WP_215385554.1-Polynucleobacter-sp.-AP-Kolm-20A-A1:0.0611023191)95.7/100:0.0680799258,WP_216177372.1-Polynucleobacter-sp.-AP-Feld-500C-C5:0.1420051628)81.6/100:0.0288489123,WP_114639791.1-Polynucleobacter-necessarius:0.0969994093)92/100:0.0456750065,((WP_216254756.1-Polynucleobacter-sp.-AP-Melu-500A-A1:0.0449609724,WP_216218975.1-Polynucleobacter-sp.-AP-Latsch-80-C2:0.0423255063)99/100:0.0775265153,((((((WP_215319389.1-Polynucleobacter-paneuropaeus:0.0803758011,(WP_215352354.1-Polynucleobacter-sp.-MWH-P3-07-1:0.0372517875,WP_215380622.1-Polynucleobacter-paludilacus:0.0348697623)87.6/100:0.0337949915)97.1/100:0.0615613629,((((WP_215376847.1-Polynucleobacter-asymbioticus:0.0241684617,WP_215328794.1-Polynucleobacter-sp.-AP-Sving-400A-A2:0.0484035518)99.3/100:0.0481812104,WP_173959551.1-Polynucleobacter-arcticus:0.0271750389)88.4/100:0.0170840887,WP_216225867.1-Polynucleobacter-sp.-MG-27-Goln-C1:0.0818367807)23.7/99:0.0059662305,(WP_216168716.1-Polynucleobacter-bastaniensis:0.0706912509,WP_216178894.1-Polynucleobacter-sp.-80A-SIGWE:0.0131141034)87.8/100:0.0193773456)86.9/99:0.0216692609)71.7/21:0.0141971418,((WP_215299358.1-Polynucleobacter-sp.-AP-Nino-20-G2:0.0802280466,WP_216197047.1-Polynucleobacter-sp.-UK-Mo-2m-Kol15:0.0553947735)56.5/25:0.0134322456,WP_062307935.1-Polynucleobacter-sinensis:0.0599628910)0/19:0.0000025547)73.3/93:0.0060262404,WP_215334312.1-Polynucleobacter-sp.-AP-Ainpum-60-G11:0.0383650430)85.3/84:0.0111416781,(WP_216268685.1-Polynucleobacter-sp.-Nonnen-W13:0.0385441926,WP_251374783.1-Polynucleobacter-corsicus:0.0328283562)73.8/90:0.0053841386)90.8/98:0.0268337709,WP_076024025.1-Polynucleobacter-sphagniphilus:0.0797783008)75.4/91:0.0157159612)95.8/99:0.0499181585)83.6/100:0.0411262489)98.4/100:0.1349944130)99.2/98:0.1568447316)76.8/37:0.0368679576)0/10:0.0005199851,(WP_040097704.1-Geoalkalibacter-ferrihydriticus:0.1906163551,WP_066727622.1-Desulfuromonas-sp.-DDH964:0.1840097565)92.7/94:0.0468614725)62.5/29:0.0186253479,WP_054772520.1-Methylogaea-oryzae:0.1548407329)74.7/51:0.0293606843,(((((((((((((WP_089129802.1-Tolypothrix-sp.-NIES-4075:0.0720084316,(WP_048866388.1-Scytonema-tolypothrichoides:0.0730064946,WP_096563464.1-Scytonema-sp.-NIES-4073:0.0930300599)81.6/55:0.0255123256)29.4/54:0.0347505782,WP_096563566.1-Scytonema-sp.-NIES-4073:0.0704332042)99.9/100:0.1225290032,(((WP_009344618.1-Cylindrospermopsis-raciborskii:0.0606495367,WP_240868223.1-Cylindrospermopsis-raciborskii:0.0207126333)99.8/100:0.1006348888,WP_220611028.1-Sphaerospermopsis-torques-reginae:0.0351316910)51.4/40:0.0228745905,WP_096570274.1-Sphaerospermopsis:0.0357407905)97.5/100:0.0654117737)42/33:0.0222751278,(((WP_237989437.1-Nostoc-sp.-UHCC-0870:0.0576947295,WP_190559626.1-Anabaena-sphaerica:0.0862536022)81.9/100:0.0291306451,WP_206263219.1-Nostoc-sp.-UHCC-0702:0.0585492116)70.7/99:0.0315942200,WP_096557790.1-Nostoc-sp.-NIES-4103:0.1186557533)90.4/98:0.0408275202)25.2/86:0.0133761775,((((WP_015128325.1-Calothrix-sp.-PCC-7507:0.1014410200,WP_017653186.1-Fortiea-contorta:0.1330817526)63.6/100:0.0424697536,((WP_193199005.1-Nostoc-sp.-MG11:0.0399527896,WP_162396505.1-Nostoc-sp.-B-2019:0.0394507433)94.5/100:0.0516569077,WP_214439445.1-Atlanticothrix-silvestris:0.1113410581)81/90:0.0462027513)69.4/89:0.0237808012,(((WP_045870014.1-unclassified-Tolypothrix:0.0231722742,WP_190656031.1-Nostoc-linckia:0.0308100369)98.4/100:0.0611809470,(WP_190743070.1-Tolypothrix-sp.-FACHB-123:0.0412865248,WP_190545010.1-Calothrix:0.0291985850)97.6/100:0.0614352974)53.9/100:0.0280430597,WP_086758595.1-Nostoc-sp.-106C:0.0584950418)81.8/90:0.0266910853)86.8/99:0.0305319059,WP_016866333.1-Fischerella:0.0688776830)70.8/96:0.0162174521)99.5/100:0.0898246283,WP_015143663.1-Pleurocapsa-sp.-PCC-7327:0.1115093978)83.8/98:0.0483295240,WP_013324143.1-Gloeothece-verrucosa:0.0827498769)1.7/60:0.0126932733,((((WP_106232846.1-Pleurocapsa-sp.-CCALA-161:0.1285494459,WP_229639079.1-Waterburya-agarophytonicola:0.0253718290)78.4/100:0.0383114732,WP_096721927.1-Chondrocystis-sp.-NIES-4102:0.0620673698)97.6/100:0.0848519521,WP_006510035.1-Xenococcus-sp.-PCC-7305:0.1370907282)94.6/100:0.0607235709,(WP_015193936.1-Stanieria-cyanosphaera:0.0178962861,WP_096384422.1-Stanieria-sp.-NIES-3757:0.0767944092)100/100:0.1217009527)95.1/95:0.0526871342)77.3/84:0.0108273779,((WP_106909014.1-Microcystis-sp.-MC19:0.1906815952,WP_124977990.1-Aphanothece-sacrum:0.1482484509)69.4/33:0.0171452451,(((WP_190772446.1-unclassified-Leptolyngbya:0.1931494375,WP_264322079.1-Zarconia-navalis:0.1191478324)87.5/79:0.0389261007,(WP_012595695.1-Rippkaea-orientalis:0.1342810677,(((WP_007308862.1-Crocosphaera-watsonii:0.0690370549,WP_107668138.1-Cyanothece-sp.-BG0011:0.0276140581)83.9/79:0.0113670750,WP_009545547.1-Crocosphaera-subtropica:0.0169918434)87.1/78:0.0130890874,WP_008278255.1-Crocosphaera-chwakensis:0.0366612061)98/98:0.0810932552)48.8/30:0.0351531150)46.8/26:0.0294013764,WP_036478304.1-Myxosarcina-sp.-GI1:0.1687672220)25.1/30:0.0165784195)88/83:0.0283024052)89.4/86:0.0279093985,(WP_008052766.1-Limnospira-indica:0.1646688736,WP_006529211.1-Gloeocapsa-sp.-PCC-73106:0.1899122918)88.4/89:0.0438796668)88.3/75:0.0329838914,((((((((WP_102179481.1-Fischerella-thermalis:0.0220042715,((WP_096675774.1-Fischerella-sp.-NIES-4106:0.0439281690,WP_053458209.1-Hapalosiphon-sp.-MRB220:0.0251030376)74.3/91:0.0053062129,(WP_131119525.1-Westiellopsis-prolifica:0.0432613576,WP_016860528.1-Fischerella-muscicola:0.0191594981)87.6/96:0.0115527413)97.9/97:0.0634299955)100/100:0.1235015275,(WP_224410051.1-Oscillatoria-salina:0.1503800517,WP_250123015.1-Chroococcidiopsis-sp.-CCMEE-29:0.1121392827)90.3/99:0.0419387215)35.2/40:0.0329412052,((WP_190825375.1-Iningainema-tapete:0.1667038065,(WP_015202791.1-Crinalium-epipsammum:0.1579559499,WP_035154029.1-Calothrix-sp.-336/3:0.2442332753)87.8/100:0.0477315856)75.9/100:0.0541698831,((WP_099102643.1-Nostoc-sp.-Peltigera-malacea-cyanobiont-DB3992:0.0728985090,(WP_190950926.1-Nostoc-punctiforme:0.0266070301,((WP_221704991.1-Desmonostoc-muscorum:0.0404000199,WP_096537650.1-Nostoc-linckia:0.0111966816)75.1/100:0.0082977544,WP_190959861.1-Nostoc:0.0309397160)28.3/91:0.0192704358)97.6/100:0.0699395619)99.6/100:0.0868325010,(((((((WP_193938462.1-Cuspidothrix-issatschenkoi:0.1268957587,(WP_190370804.1-Anabaena-sp.-FACHB-1237:0.1201707967,WP_168537172.1-Anabaena-sp.-UHCC-0253:0.0277451194)10.5/86:0.0143955426)85.1/65:0.0163606513,(((WP_190392428.1-Dolichospermum-sp.-FACHB-1091:0.0092018890,(WP_028089231.1-Dolichospermum-circinale:0.0164990857,WP_137908967.1-Nostocales:0.0395605172)98.3/100:0.0526234603)99.9/100:0.0916350412,((WP_096666215.1-Dolichospermum-compactum:0.0534026651,(WP_190387835.1-Aphanizomenon:0.0519363042,WP_168466156.1-Aphanizomenon:0.0164099849)28.2/98:0.0120078560)64.8/96:0.0131861636,WP_015080416.1-Anabaena-sp.-90:0.0284120474)95.8/97:0.0281347634)41.7/65:0.0056640381,WP_168502193.1-Anabaena-sp.-UHCC-0187:0.0000021765)86/65:0.0205424238)99.9/100:0.0959488295,(WP_010998622.1-Nostocaceae:0.0523156642,WP_011320078.1-Nostocaceae:0.0479302795)82.5/100:0.0166701236)92.2/98:0.0358788153,(WP_239729951.1-Nodularia-sphaerocarpa:0.0518905046,WP_173403323.1-Nodularia-spumigena:0.0475837003)96.4/100:0.0482710119)24.2/30:0.0063356030,(WP_199337786.1-Nostoc-sp.-FACHB-280:0.0908940111,WP_199337214.1-Nostoc-sp.-FACHB-152:0.0917655010)87.8/28:0.0283805885)41.9/29:0.0103179661,(WP_015138800.1-Nostoc-sp.-PCC-7524:0.0664345695,WP_066425233.1-Anabaena-sp.-4-3:0.0713453522)97.4/100:0.0551326691)97.8/100:0.0624883720,(WP_096624129.1-Calothrix-sp.-NIES-3974:0.0607476378,WP_095721999.1-Calothrix-elsteri:0.0786346160)99/100:0.0785026608)72.4/100:0.0226275891)89.6/100:0.0327478428)61.3/38:0.0204960925)98.1/97:0.0592058707,((((WP_015170385.1-Geitlerinema-sp.-PCC-7407:0.1953503398,WP_194029353.1-Lusitaniella-coriacea:0.1648595259)88/98:0.0362485734,(((WP_233747769.1-Leptodesmis-sichuanensis:0.1633192091,WP_223049852.1-Leptothermofonsia-sichuanensis:0.0817357970)63.6/94:0.0067859958,(WP_192216758.1-Microcoleus-sp.-FACHB-SPT15:0.1268267742,WP_224345144.1-Kovacikia-minuta:0.0895277907)44.5/95:0.0246536458)91.2/97:0.0248868560,WP_073549891.1-Chroogloeocystis-siderophila:0.1335596107)68.3/89:0.0127635486)88.2/85:0.0216560231,(WP_190666400.1-Coleofasciculus-sp.-FACHB-501:0.0979774189,WP_190525322.1-Coleofasciculus-sp.-FACHB-T130:0.0431956020)97.5/100:0.0575243797)67.7/24:0.0061384250,(WP_193873404.1-unclassified-Coleofasciculus:0.1015421839,WP_254010234.1-Symplocastrum-sp.-BBK-W-15:0.1086246276)91.2/75:0.0358738451)64.1/27:0.0149312437)76.3/83:0.0179110362,(WP_008178350.1-Moorena-producens:0.1378387818,WP_190493486.1-unclassified-Microcoleus:0.1761461413)83.2/89:0.0434153808)47/87:0.0261320747,((WP_017661508.1-Baaleninema-simplex:0.2115546580,WP_007355666.1-Kamptonema:0.0857867588)36.2/79:0.0338721279,(WP_226578226.1-Microseira-wollei:0.1490689931,((WP_073595522.1-Phormidium-ambiguum:0.0575547052,WP_193982193.1-Phormidium-sp.-LEGE-05292:0.0547612399)99.5/100:0.0826433357,(WP_249064905.1-Argonema-antarcticum:0.0421524531,WP_190468950.1-unclassified-Planktothrix:0.0760714658)78.2/97:0.0114722242)87.9/95:0.0374366883)91.9/97:0.0398772733)73.4/77:0.0188260432)70.3/71:0.0119125619,(WP_017715496.1-Oscillatoria-sp.-PCC-10802:0.1272459118,(WP_190665495.1-Microcoleus-sp.-FACHB-672:0.0529688942,WP_261220767.1-Ancylothrix-sp.-D3o:0.1449341072)96.4/98:0.0583747341)86.3/81:0.0315064090)77.7/49:0.0133557028,((((((((WP_202221768.1-Okeania-sp.-KiyG1:0.0348697402,WP_094672483.1-Hydrocoleum-sp.-CS-953:0.0353486059)33.5/100:0.0124623922,WP_124146462.1-Okeania-hirsuta:0.0083527502)99.8/100:0.1208957331,WP_071516849.1-Geitlerinema-sp.-PCC-9228:0.1621162599)94.2/100:0.0685542082,(WP_009783380.1-Lyngbya:0.0702021533,WP_046277942.1-Limnoraphis-robusta:0.0371805471)94.7/100:0.0558069987)75.6/95:0.0386865720,((WP_083623900.1-Planktothrix-serta:0.0337345557,WP_072717475.1-Planktothrix:0.0230375859)85.6/100:0.0169347708,(WP_079676747.1-Planktothrix-sp.-PCC-11201:0.0210465534,(WP_026797127.1-Planktothrix:0.0221410318,WP_214559879.1-Planktothrix-agardhii:0.0347731454)71.9/100:0.0231006271)98.3/100:0.0578915169)97.1/100:0.0628815400)84.2/59:0.0297771473,WP_190877489.1-unclassified-Planktothricoides:0.1277526916)89.4/34:0.0226518273,(WP_190531554.1-Coleofasciculus-sp.-FACHB-542:0.1634233217,((((WP_236507253.1-Tychonema-sp.-BBK16:0.0343895475,WP_096828456.1-Tychonema-bourrellyi:0.0556680225)93.7/100:0.0302121997,WP_242724542.1-Microcoleus-vaginatus:0.0283190454)83.8/76:0.0113841811,WP_015174083.1-Oscillatoria-nigro-viridis:0.0168593649)72.5/77:0.0068220006,((WP_194009709.1-unclassified-Tychonema:0.0318228587,WP_265235823.1-Lyngbya-sp.-CCAP-1446/10:0.0360283632)88.5/100:0.0170735797,WP_193973976.1-Microcoleus-sp.-LEGE-07076:0.0427139612)48.6/76:0.0127484750)99.1/100:0.0777152191)62.3/75:0.0264412050)74.5/22:0.0184305073,(((WP_261197618.1-unclassified-Laspinema:0.0414077440,WP_207087219.1-Phormidium-pseudopriestleyi:0.0312116210)16.3/81:0.0134529101,WP_261204505.1-Laspinema-sp.-D2d:0.0268704819)99/100:0.0832567515,WP_200665545.1-Oxynema-sp.-CENA135:0.1731793229)78.6/38:0.0253122041)77.5/44:0.0116284610)92.9/59:0.0370715962)92.4/88:0.0465664461,WP_069965800.1-unclassified-Desertifilum:0.1769672145)84.5/89:0.0672090852,(((WP_204137747.1-Halomicronema-sp.-CCY15110:0.0415363415,WP_039724568.1-Leptolyngbya-sp.-LK:0.0616223953)99.9/100:0.1149949089,WP_023076112.1-Leptolyngbya-sp.-Heron-Island-J:0.1610964186)94.1/100:0.0645704482,WP_017324474.1-Synechococcus-sp.-PCC-7336:0.1912140052)72.6/98:0.0480553858)95.7/97:0.0655264394)79.9/18:0.0191415921)94.6/20:0.0602128949,(((((WP_015405619.1-Desulfocapsa-sulfexigens:0.1661421024,(WP_103114677.1-Geothermobacter-hydrogeniphilus:0.2048653956,WP_221248821.1-Desulfuromonas-versatilis:0.1685003465)93.8/97:0.0657449757)90.8/51:0.0417613980,(((WP_205222281.1-Desulfogranum-marinum:0.1556124868,WP_028582690.1-Desulfogranum-japonicum:0.2017849998)18.1/26:0.0422477595,(WP_205242362.1-Desulfobulbus-alkaliphilus:0.1248817790,WP_267928740.1-Desulfolithobacter-dissulfuricans:0.0966979260)84/97:0.0480229928)96.8/100:0.0646842874,((((WP_250680171.1-Geomonas-sp.-Red32:0.1875612513,WP_027714505.1-Desulfuromonas-sp.-TF:0.1371911566)47.1/12:0.0274328626,WP_053550565.1-Desulfuromonas-soudanensis:0.1985079735)65.9/12:0.0199582154,WP_072282412.1-Syntrophotalea-acetylenivorans:0.1799650474)85/23:0.0390531770,(WP_199381999.1-Geomesophilobacter-sediminis:0.1805892699,WP_205218910.1-Desulforhopalus-vacuolatus:0.2120345565)19.7/21:0.0333174713)51.2/83:0.0379218586)51.6/23:0.0228192318)93.6/62:0.0413248739,((WP_163339734.1-Desulfopila-sp.-IMCC35008:0.2071383795,(WP_136805251.1-Desulfosediminicola-flagellatus:0.1721877740,WP_073615262.1-Desulfopila-aestuarii:0.1724359182)51.3/60:0.0091982735)95.6/100:0.0736301552,(WP_228856481.1-Desulfomarina-profundi:0.2174594630,WP_183349429.1-Desulfoprunum-benzoelyticum:0.1420305465)85.1/95:0.0469251190)72/73:0.0205008997)86.3/77:0.0223280865,WP_066347484.1-Geminocystis-sp.-NIES-3708:0.2879090529)47.1/22:0.0311554447,WP_054689510.1-Desulfosarcina-cetonica:0.1858861530)79.1/51:0.0373243302)94/23:0.0563579549,((((((((((((((WP_065305794.1-Janthinobacterium-psychrotolerans:0.1037862521,WP_034778721.1-Janthinobacterium-lividum:0.0293622451)63.8/63:0.0041012169,WP_035822645.1-Janthinobacterium-sp.-RA13:0.0526277175)76.8/63:0.0071661325,WP_152282105.1-Janthinobacterium-violaceinigrum:0.0173244743)73/63:0.0056395429,WP_183742413.1-unclassified-Janthinobacterium:0.0401783371)71.1/64:0.0056824873,(((((WP_139090581.1-Janthinobacterium-lividum:0.0563597837,WP_230539136.1-Janthinobacterium-sp.-FW305-129:0.0348436813)85.8/95:0.0121223124,WP_094442213.1-Janthinobacterium-sp.-PC23-8:0.0540048354)31.5/93:0.0056822568,WP_215403374.1-Janthinobacterium-sp.-JC611:0.0212376898)77.2/98:0.0073190418,WP_102124214.1-Janthinobacterium-sp.-ROICE36:0.0287925280)77.1/71:0.0056870631,WP_096238059.1-Janthinobacterium-svalbardensis:0.0170982656)79.5/71:0.0056282229)99.4/94:0.0833319906,WP_101482628.1-unclassified-Janthinobacterium:0.1330294194)71.4/93:0.0282158135,WP_023495308.1-Methyloglobulus-morosus:0.1386414878)34.2/27:0.0114764675,(WP_099917421.1-Massilia-psychrophila:0.0994808616,(WP_205319084.1-Glaciimonas-sp.-PAMC28666:0.0643188125,WP_092396716.1-Collimonas-sp.-OK412:0.0944452805)37.2/93:0.0132263563)82/91:0.0137873112)95.4/89:0.0488595446,WP_250479419.1-unclassified-Caballeronia:0.2252139961)91.4/98:0.0359636470,(((WP_162020603.1-Undibacterium-crateris:0.1103987865,WP_186884883.1-Undibacterium:0.0377027971)0/96:0.0160808429,WP_186861877.1-Undibacterium-griseum:0.0532958205)98.6/100:0.0809228131,((((((((((((WP_207000528.1-Trinickia-sp.-DHG64:0.1006490875,WP_124149903.1-Paraburkholderia-dinghuensis:0.0827123620)53.9/100:0.0275808502,(WP_266250364.1-Paraburkholderia-megapolitana:0.1143879367,WP_026121090.1-Paraburkholderia-kururiensis:0.0977297636)58.2/100:0.0457097763)98.5/100:0.0871723327,WP_134195781.1-Paraburkholderia-rhizosphaerae:0.1639786477)88.4/22:0.0357308982,WP_121276956.1-Trinickia-fusca:0.0967075408)88.5/23:0.0306872423,WP_059889870.1-Burkholderia-stagnalis:0.1588500919)72.5/6:0.0078736579,WP_209935850.1-Paraburkholderia-sp.-LEh10:0.0511344812)90.4/7:0.0244039195,((((WP_060092900.1-Burkholderia-vietnamiensis:0.0184860521,WP_226113708.1-Burkholderia-sp.-AU33545:0.0411038796)100/100:0.1134043301,WP_136892825.1-Trinickia-sp.-7GSK02:0.0346345730)32.3/7:0.0085288265,(WP_205976236.1-Paraburkholderia-sp.-Se-20369:0.1719311760,WP_206951398.1-Trinickia-sp.-4D114:0.0635120564)98.1/80:0.0761055724)47.2/7:0.0176105353,(WP_054928771.1-Paraburkholderia-caribensis:0.0396144277,(WP_201650988.1-Paraburkholderia-sabiae:0.0392756961,WP_205970438.1-Paraburkholderia-sp.-Tr-20389:0.0281726886)84.3/99:0.0113270050)78.1/90:0.0102816478)84.2/7:0.0127572333)86.2/20:0.0157316805,((((WP_201138986.1-Paraburkholderia-domus:0.0397278747,WP_266262001.1-Paraburkholderia-madseniana:0.0195054474)99.9/100:0.0974988290,WP_009889585.1-Burkholderia-thailandensis:0.0629124303)86.5/99:0.0350828151,WP_061168975.1-Caballeronia-hypogeia:0.0686260073)90.6/21:0.0247391264,((WP_168792792.1-Paraburkholderia-aromaticivorans:0.0575801447,((WP_004544228.1-Burkholderia:0.0250572273,WP_069272519.1-Burkholderia-cepacia:0.1625747224)84.4/100:0.0208247639,WP_059583574.1-pseudomallei-group:0.0243066148)81.6/100:0.0228160485)86.8/78:0.0224745338,(((WP_175791145.1-Burkholderia-ambifaria:0.0795347248,WP_059842355.1-Burkholderia-multivorans:0.0309525613)75.8/100:0.0109972910,WP_105772332.1-Burkholderia-multivorans:0.0303235023)88.1/100:0.0178102312,(WP_006496666.1-Burkholderia-cenocepacia:0.0250266323,WP_059716874.1-Burkholderia-ubonensis:0.0311053810)92.5/100:0.0226153479)76.3/98:0.0137062125)0/72:0.0000027683)84.5/19:0.0130613641)85.2/68:0.0134629227,WP_060156927.1-Burkholderia-ubonensis:0.0738074095)93.7/87:0.0398690765,WP_059459219.1-Burkholderia-vietnamiensis:0.1888285554)90.7/90:0.0356836522,(WP_179404315.1-Burkholderia-guangdongensis:0.1344050239,((WP_063494936.1-Paraburkholderia-phytofirmans:0.0254134174,(WP_254318368.1-Paraburkholderia-sp.-CNPSo-3272:0.0054678488,WP_042262232.1-Paraburkholderia-heleia:0.0519534372)95.8/100:0.0329769308)98/100:0.0752386707,WP_038711716.1-Burkholderia-sp.-lig30:0.1724073830)82.9/90:0.0477465621)90.3/32:0.0348303717)65.7/30:0.0401809492,WP_056136218.1-Massilia-sp.-Root133:0.1357984553)99.3/100:0.1151632510)94.1/37:0.0475565426)76.1/18:0.0101078175,((((WP_115680924.1-Cupriavidus-taiwanensis:0.1882914888,WP_135702508.1-Cupriavidus-oxalaticus:0.1086175189)4.7/9:0.0237272678,(WP_018313502.1-Cupriavidus-sp.-UYPR2.512:0.1674842927,WP_174426450.1-Cupriavidus-basilensis:0.2406434431)84.6/47:0.0395272470)86.3/19:0.0351301816,WP_071014919.1-Cupriavidus-sp.-USMAHM13:0.1666363925)98/99:0.0676768025,WP_217900122.1-Noviherbaspirillum-humi:0.1549747153)84.5/34:0.0211302626)92/81:0.0423457438,(WP_237889684.1-Dissulfurimicrobium-hydrothermale:0.2463268530,(WP_250024752.1-Cupriavidus-campinensis:0.1408105594,WP_094797980.1-Bordetella-genomosp.-5:0.1223504229)74.9/99:0.0648193302)10/7:0.0198234393)66.4/4:0.0162767816,(WP_247737070.1-Thauera-aromatica:0.0485047900,WP_039805139.1-Azotobacter-chroococcum:0.1109554299)97.8/100:0.0694693129)68.7/8:0.0266988017,((WP_195874647.1-Herminiimonas-contaminans:0.0995737954,WP_041297045.1-Janthinobacterium-sp.-Marseille:0.0758631990)87.7/100:0.0367191469,((WP_076590920.1-Herminiimonas-arsenitoxidans:0.1375315055,WP_206604468.1-Herminiimonas-fonticola:0.0328233221)77.3/74:0.0191177836,WP_188420499.1-Oxalicibacterium-solurbis:0.1226256007)71.7/97:0.0251145473)99.7/100:0.1014605816)8.3/6:0.0154955677)88/17:0.0393521544,(WP_239190817.1-Candidatus-Nitrotoga-sp.-HW29:0.0396125301,((WP_239235002.1-Candidatus-Nitrotoga-sp.-BS:0.0450161369,WP_239797721.1-Candidatus-Nitrotoga-arctica:0.0224681071)86.4/65:0.0152106761,WP_239287644.1-Candidatus-Nitrotoga-sp.-1052:0.0130617883)73.3/51:0.0132884349)100/100:0.1688551243)62.8/94:0.0325480772,(WP_188630571.1-Oceanisphaera-marina:0.0748612242,(WP_104485062.1-Oceanisphaera-arctica:0.0860926610,WP_204795415.1-Oceanisphaera-litoralis:0.0375528889)86.1/93:0.0363276943)100/100:0.1524246889)81.2/94:0.0235740949,(WP_020932001.1-Cycloclasticus-zancles:0.1947769891,(WP_159673755.1-Andreprevotia-sp.-IGB-42:0.1410257970,WP_084093192.1-Andreprevotia-lacus:0.0631675675)99.7/100:0.1125224120)91.3/30:0.0532366399)47.2/25:0.0165541261,((WP_213263532.1-Burkholderia-pyrrocinia:0.1021132722,WP_060279477.1-Burkholderia-cepacia-complex:0.1000474821)98.5/100:0.1054060196,(((WP_208101977.1-Burkholderia-sp.-D-99:0.0288498893,WP_205714023.1-Burkholderia-cepacia:0.0254072659)43.2/100:0.0062502433,WP_174919663.1-Burkholderia-metallica:0.0182172530)83.8/100:0.0303145353,WP_077021900.1-Burkholderia-cenocepacia:0.0707102500)99.9/100:0.1445093956)93.4/100:0.0533804408)95/79:0.0589311243,(WP_043620211.1-Chromobacterium-piscinae:0.1578308754,(WP_103900886.1-Chromobacterium-alticapitis:0.1569551331,WP_045052286.1-Chromobacterium-violaceum:0.1658720823)90.3/100:0.0516219871)94.9/100:0.0765690143)4.4/14:0.0261965945,WP_189531325.1-Paludibacterium-paludis:0.2924980474)84.6/41:0.0262858953,((((WP_132257458.1-Paucimonas-lemoignei:0.0659422133,WP_119785517.1-Noviherbaspirillum-sedimenti:0.0486762510)100/100:0.2407113038,(WP_026470910.1-Alkanindiges-illinoisensis:0.1757551816,(WP_248147450.1-Moraxella-osloensis:0.2232281578,WP_092690500.1-Acinetobacter-kyonggiensis:0.1353214088)88.6/100:0.0661176956)99.6/100:0.1469010406)84.9/100:0.0673571737,(WP_134696535.1-Aeromonas-taiwanensis:0.3631576927,(WP_094838100.1-Bordetella-genomosp.-4:0.1525101138,WP_160351730.1-Bordetella-sp.-15P40C-2:0.1603052154)97.4/100:0.1198231663)55.1/85:0.0742610562)94.7/82:0.0939152203,(WP_208005626.1-Oceanisphaera-pacifica:0.0877840876,(WP_087034853.1-Oceanisphaera-profunda:0.0639926856,WP_087034873.1-Oceanisphaera-profunda:0.0759639730)0/85:0.0402671421)96.7/100:0.0824487840)83.6/63:0.0345879423)94.1/65:0.0407362307,(((((WP_223932214.1-Aeromonas-caviae:0.0857863842,WP_161506954.1-Aeromonas-media:0.0252108383)97.6/100:0.0774344317,((WP_208149206.1-Shewanella-algae:0.2135413093,(((WP_045500143.1-Vibrio-hyugaensis:0.1287045310,WP_045419977.1-Vibrio-campbellii:0.0635622081)61.2/88:0.0310724621,WP_025533544.1-Vibrio-parahaemolyticus:0.0545423241)93.6/90:0.0392916035,WP_011788809.1-Shewanella-sp.-W3-18-1:0.0564453574)74.7/90:0.0190277166)44/95:0.0263327170,WP_219142360.1-Aeromonas-sanarellii:0.0984636513)97.1/100:0.0768921083)91.4/88:0.0477228195,((((WP_248067241.1-unclassified-Shewanella:0.0742847831,WP_148505453.1-Vibrio-cholerae:0.0781705748)97.7/100:0.0726703898,(WP_218558465.1-Shewanella-putrefaciens:0.0730761760,WP_011919582.1-Shewanella-putrefaciens:0.0196758769)99.5/100:0.0726982672)75.3/97:0.0217528978,(((((WP_050718733.1-Aeromonas-tecta:0.1181095851,WP_100859741.1-Aeromonas-lusitana:0.0329984930)74.5/97:0.0081149493,WP_042654929.1-Aeromonas-encheleia:0.0461274467)97.1/100:0.0502421922,WP_077098334.1-Aeromonas-hydrophila:0.0434961164)57.7/97:0.0136101381,WP_043167431.1-Aeromonas-hydrophila:0.0870463284)67.6/94:0.0161952459,WP_252420256.1-Aeromonas-taiwanensis:0.0559792998)98.9/100:0.0761126908)62.3/81:0.0466981047,(((WP_147247296.1-Brenneria-salicis:0.0339014835,WP_172289906.1-unclassified-Brenneria:0.0825649689)79.6/100:0.0148192882,WP_039356706.1-Pectobacterium-actinidiae:0.0824819268)97.1/80:0.0577842623,((WP_198997094.1-Pectobacterium-carotovorum:0.0169893508,(WP_165707548.1-Pectobacterium:0.0262102760,WP_174866005.1-Pectobacterium-polaris:0.0241352122)99.2/100:0.0511824173)85/80:0.0171650363,WP_145962313.1-Pectobacterium-peruviense:0.0427713782)99.2/100:0.0866278777)96/95:0.0616034084)83.6/59:0.0287037594)87.4/46:0.0269628316,((((WP_073350970.1-Aeromonas-hydrophila:0.0958561331,WP_190284026.1-Aeromonas-hydrophila:0.0324494027)85.6/100:0.0234257032,WP_241344946.1-Aeromonas-sp.-MR16:0.0558198745)89.9/100:0.0291489952,(WP_223952962.1-Aeromonas-caviae:0.1340270241,WP_244081328.1-Aeromonas-caviae:0.1219351482)96.4/99:0.0638345103)95.4/99:0.0490945008,(WP_076573677.1-Aeromonas-sp.-RU39B:0.1565462496,WP_108652490.1-Dongshaea-marina:0.1374222181)96.3/45:0.0652718805)81.7/39:0.0166957238)88.7/14:0.0243929078,WP_102949229.1-Aeromonas-caviae:0.1777251158)87.1/15:0.0283229249)76.6/7:0.0170577459)71.2/7:0.0200691450,((((((((((WP_004123988.1-Rhizobium-freirei:0.0800616990,WP_095433322.1-Rhizobium-sp.-11515TR:0.0305904104)80.4/82:0.0176448605,((WP_183712806.1-unclassified-Rhizobium:0.0323350965,WP_015341698.1-Rhizobium:0.0250196105)53.9/75:0.0112218160,WP_106999541.1-Rhizobium-sp.-JAB6:0.0443713902)0/74:0.0000023551)75.1/96:0.0170557482,WP_184706966.1-Rhizobium-lusitanum:0.0767450467)0/91:0.0000022406,(WP_174074493.1-Agrobacterium-rhizogenes:0.0778733772,((WP_183765619.1-unclassified-Rhizobium:0.0311901453,WP_120667780.1-unclassified-Rhizobium:0.0454686906)86.7/100:0.0267857530,WP_037195771.1-Rhizobium-lusitanum:0.0234547836)37.9/99:0.0117832721)81.2/72:0.0125520708)74.6/75:0.0080156165,WP_092719658.1-Rhizobium-multihospitium:0.0309074036)87.5/98:0.0212515608,WP_168880117.1-Rhizobium-sp.-P28RR-XV:0.0365760189)40.4/70:0.0107576945,WP_112341687.1-Rhizobium-tropici:0.0311403268)85.2/99:0.0145974205,(WP_199146672.1-Rhizobium-sp.-ASV8:0.1381011986,WP_145321829.1-unclassified-Rhizobium:0.0768373569)33.4/63:0.0077023212)99.8/100:0.1080671428,(WP_043579071.1-Chromobacterium:0.0522870299,WP_071113424.1-Chromobacterium-sphagni:0.0746160731)99.7/100:0.1140201759)99.7/100:0.1040171316,((WP_174873379.1-Vogesella-oryzae:0.0841049203,WP_137008594.1-Aquitalea-aquatilis:0.2227786230)42.2/63:0.0564533425,WP_147289921.1-Crenobacter-cavernae:0.1038516230)53.4/98:0.0367139222)93/7:0.0446019692)95.5/98:0.0442411222,((WP_224211331.1-Aeromonas-enteropelogenes:0.1469830282,WP_082041448.1-Aeromonas-eucrenophila:0.1581045075)80.8/77:0.0557993062,WP_252995952.1-Aeromonas-encheleia:0.1746284620)83.6/72:0.0295897882)85.2/62:0.0199954801,(((((((WP_172912938.1-Pantoea-stewartii:0.1001876686,WP_145394131.1-Pantoea-sp.-SJZ147:0.0981497417)99.8/100:0.1567087746,WP_086868392.1-Kosakonia-pseudosacchari:0.3322604663)99.1/75:0.1306774495,(((((((WP_071788415.1-Franconibacter-pulveris:0.3080923425,WP_020232007.1-Escherichia-coli:0.1416556375)88.2/94:0.0528285598,(WP_228338545.1-Enterobacter-hormaechei:0.1218925108,(WP_109845273.1-Enterobacter:0.0441780302,WP_150146763.1-Enterobacter-roggenkampii:0.0336446018)87.3/100:0.0538251410)99.5/100:0.1168085114)87.3/98:0.0585942248,((WP_232581009.1-Photobacterium-phosphoreum:0.3072878584,(WP_073514120.1-Vibrio-parahaemolyticus:0.0490546977,WP_032481483.1-Vibrio:0.0195000124)100/100:0.3007438719)93.2/99:0.1142406890,WP_153752706.1-unclassified-Citrobacter:0.3005418351)57.9/74:0.0921023831)91.5/41:0.0754608952,(WP_162556830.1-Proteus:0.0691668368,WP_161779122.1-Proteus-sp.-G2675:0.0671944263)99.2/100:0.1064132335)96.9/40:0.0713597500,WP_236366479.1-Aeromonas-veronii:0.1360259741)0/12:0.0018300792,((((((((((((((((((((((((((((((((((((WP_051234278.1-Halomonas-halocynthiae:0.3571044755,WP_110648656.1-Salinicola-peritrichatus:0.2119751971)99.4/100:0.1715437130,WP_021818206.1-Halomonas-huangheensis:0.0997938116)77.1/33:0.0369872284,(WP_248234415.1-Marinobacter-bryozoorum:0.0792851865,(WP_228223924.1-Marinobacter-nauticus:0.0430400722,WP_098418752.1-Marinobacter-sp.-LV10MA510-1:0.0942173385)93.7/100:0.0504554648)94.9/100:0.0514172625)87.3/34:0.0327700573,((((((WP_066801876.1-Moraxella-oblonga:0.1049145347,(WP_197128702.1-Psychrobacter-sp.-NZS113:0.1249837605,WP_201616712.1-Psychrobacter-immobilis:0.1277165087)98.9/100:0.1144712861)93.6/100:0.1117840903,WP_036548095.1-Nitrincola-lacisaponensis:0.2720534641)88.8/98:0.0471178488,((WP_041157945.1-Halomonas-sp.-KHS3:0.0437072186,((WP_120386672.1-Halomonas-alkaliphila:0.1101781352,WP_226251132.1-unclassified-Halomonas:0.0489558084)34.7/78:0.0212490894,WP_141441391.1-Halomonas-titanicae:0.0570127217)92.5/90:0.0532582390)99.3/100:0.0899223703,WP_228219791.1-Halomonas-meridiana:0.0746274179)96.6/99:0.0547289121)92.5/94:0.0328047311,((((((WP_030071911.1-Halomonas-alkaliantarctica:0.0402040030,WP_095603285.1-Halomonas-humidisoli:0.0224662859)80.9/44:0.0169322990,WP_253485683.1-Halomonas-sp.-XH26:0.0240370791)76.5/44:0.0106075948,WP_038485954.1-Halomonas-campaniensis:0.0182474606)98.7/100:0.0619217088,WP_162218282.1-Halomonas-alkaliphila:0.0247812975)95.6/100:0.0389919834,WP_078087027.1-unclassified-Halomonas:0.0224296391)0/96:0.0024987629,WP_096655487.1-Halomonas-nigrificans:0.0385116542)97/100:0.0683381216)92.2/74:0.0442967227,(WP_096281135.1-Halomonas-sp.-JB37:0.1578807589,(WP_027958907.1-unclassified-Halomonas:0.0354241707,WP_064236373.1-unclassified-Halomonas:0.0403173133)97.5/100:0.0665889055)90.1/100:0.0445291864)80.8/63:0.0141235411,WP_127061154.1-Halomonas-nanhaiensis:0.1081669887)96.2/77:0.0539964780)94.1/34:0.0262991767,((((((WP_158774340.1-Cobetia-sp.-L2A1:0.0589026394,WP_144727127.1-Cobetia-crustatorum:0.0264689658)19.2/87:0.0097459092,WP_088744253.1-Cobetia-sp.-QF-1:0.0458802878)89.8/99:0.0248195163,WP_158774127.1-Cobetia-sp.-L2A1:0.0225619415)30.3/99:0.0078619408,(WP_077373671.1-Cobetia:0.0412116681,((WP_213115235.1-Cobetia-sp.-MC34:0.0330036018,WP_200017180.1-Cobetia-sp.-cqz5-12:0.0473996384)94.2/100:0.0423946118,WP_225346840.1-Cobetia-amphilecti:0.0309608227)92.1/100:0.0269303384)92.2/100:0.0278849300)98.3/99:0.0943193991,(WP_251978361.1-unclassified-Salinicola:0.0658126147,WP_189443129.1-Salinicola-rhizosphaerae:0.0756979595)99.7/100:0.1187957961)98/90:0.0864845037,((WP_017842145.1-Methylotuvimicrobium-buryatense:0.2523658147,WP_073436013.1-Halomonas-cupida:0.0462470416)36.3/67:0.0379996333,WP_149282777.1-Halomonas-sp.-Y2R2:0.0500758775)94.3/71:0.0569304432)73.1/65:0.0058355760)84.8/32:0.0311010591,WP_035566812.1-Halomonas:0.1870651008)94/42:0.0432538518,(WP_193343673.1-Halomonas-elongata:0.0483926541,WP_064701650.1-Halomonas-caseinilytica:0.0167382614)99.8/100:0.0764820304)88/42:0.0339229430,((WP_159342670.1-Halomonas-meridiana:0.2076238038,WP_253554573.1-Halomonas-sp.-DN3:0.1342266484)81.4/100:0.0283577070,WP_102589198.1-Halomonas-urumqiensis:0.0873424828)70.8/99:0.0177748132)91.9/41:0.0268719233,((WP_224850395.1-Marinobacter-nauticus:0.1126981024,WP_110630630.1-Salinicola-salarius:0.0647768077)75.8/86:0.0316211056,((((WP_079551663.1-Halomonas-subglaciescola:0.0535013927,WP_195687443.1-Halomonas-sp.-KAO:0.0463365544)86.4/99:0.0150063819,WP_027961801.1-Halomonas-halodenitrificans:0.0278730791)62.5/99:0.0119598042,((WP_253557393.1-unclassified-Halomonas:0.0361181705,WP_089676106.1-Halomonas-aquamarina:0.0233717346)0/99:0.0000023836,WP_062362595.1-Halomonas-axialensis:0.0295917336)65.5/100:0.0159135402)45.1/87:0.0111062331,WP_089675781.1-Halomonas-aquamarina:0.0292916784)91.2/98:0.0267212656)85.1/82:0.0155065420)90.1/29:0.0353521090,WP_193085952.1-Halomonas-sp.-FME65:0.0519127329)97.6/69:0.0982871720,WP_081193716.1-Halomonas-sp.-BC1:0.1259474521)93.7/75:0.0889768307,WP_254581577.1-Marinobacterium-sedimentorum:0.2128871521)31.4/97:0.0420301685,WP_179929642.1-Halomonas-salicampi:0.2396436134)89.1/100:0.0541660731,(WP_015904383.1-Desulforapulum-autotrophicum:0.1966672689,WP_111960646.1-Desulfobacter-hydrogenophilus:0.2104946988)84.6/100:0.1015372871)83.5/94:0.0794725667,(((WP_151033206.1-Cellvibrio-sp.-KY-GH-1:0.0662633112,WP_049631514.1-Cellvibrio-sp.-pealriver:0.0617182557)73.9/100:0.0472944511,WP_062059887.1-Cellvibrio-sp.-OA-2007:0.0565954513)100/100:0.1753201435,WP_178968040.1-Neptunomonas-phycophila:0.2113517827)82.2/93:0.0953544106)53.4/88:0.0582789986,WP_156872703.1-Ketobacter-sp.-MCCC-1A13808:0.2422757814)81.6/95:0.0502137518,((((((WP_125841544.1-Stutzerimonas-stutzeri:0.2263677407,((WP_262731117.1-Alcanivorax-marinus:0.1219887406,(WP_106232428.1-Halomonas-ventosae:0.0722465088,WP_228234339.1-Alcanivorax-marinus:0.1386094836)72.4/100:0.0352394741)83.2/99:0.0207384272,WP_175070852.1-Halomonas-taeanensis:0.0880309529)34.1/94:0.0788283947)79.9/99:0.0607238074,(WP_019018778.1-Halomonas-lutea:0.2035883933,WP_092999033.1-Thiohalomonas-denitrificans:0.0915175963)88.8/100:0.0978265023)86/100:0.0362963168,((WP_146294265.1-Euhalothece-natronophila:0.1606205231,WP_159299499.1-Spiribacter-sp.-SSL99:0.1716313314)91.9/100:0.0555310136,WP_011630054.1-Alkalilimnicola-ehrlichii:0.1433634105)83.9/91:0.0442838442)79.5/91:0.0514255072,(WP_127477393.1-Sulfurivermis-fontis:0.2058285811,WP_028988655.1-Thermithiobacillus-tepidarius:0.1990000629)3.1/57:0.0310184414)98/99:0.1078620568,WP_069857109.1-Desulfoplanes-formicivorans:0.2015815341)87.2/82:0.0272340473,((WP_116367876.1-Parahaliea-mediterranea:0.0584052094,WP_148063811.1-Parahaliea-aestuarii:0.0846623770)100/100:0.2319394423,WP_129645432.1-Steroidobacter-agariperforans:0.2222849814)99.2/98:0.1357071046)87.3/84:0.0407035674)97.6/99:0.0923025286,((((WP_262601261.1-Endozoicomonas-euniceicola:0.0762592220,WP_066015073.1-Endozoicomonas-atrinae:0.0991163581)89.7/100:0.0624199428,WP_257287521.1-Endozoicomonas-sp.-SESOKO2:0.0895120652)93.9/100:0.0736122684,(WP_067585802.1-Endozoicomonas-ascidiicola:0.1070754880,(WP_020580625.1-Endozoicomonas-elysicola:0.0918330441,WP_257293945.1-Endozoicomonas-sp.-YOMI1:0.0910314741)90.6/100:0.0648736523)99.6/100:0.1448354695)73.2/100:0.0700630076,WP_116686321.1-Pelagibaculum-spongiae:0.2639359352)94.3/88:0.1046135552)58.1/81:0.0230813360,WP_255857717.1-Bacterioplanoides-sp.-SCSIO-12839:0.2150551316)80/94:0.0422840374,(((((((((WP_132974196.1-Thiogranum-longum:0.1511200462,WP_201344172.1-Thiohalobacter-sp.-COW1:0.1185340865)99.2/100:0.1880420716,WP_198243301.1-methane-oxidizing-endosymbiont-of-Gigantopelta-aegis:0.2208058184)89.6/91:0.1134983403,(WP_161826415.1-Petrachloros-mirabilis:0.3346850801,WP_062153084.1-Beggiatoa-leptomitoformis:0.2599550737)43.3/44:0.0718318697)85.1/49:0.0669999596,((((((WP_176021856.1-Stenotrophomonas-sp.-NA06056:0.0460889812,WP_049440708.1-Stenotrophomonas-maltophilia:0.0408961464)98.4/100:0.1105552588,WP_132999175.1-Luteimonas-arsenica:0.2029372129)83/100:0.0732032250,WP_078758981.1-Lysobacter-spongiicola:0.2072577058)72.6/100:0.0617679190,((((((((WP_019467068.1-Dyella-japonica:0.1920082073,WP_204636097.1-Dyella-kyungheensis:0.0990554819)92.4/52:0.0408457393,((WP_049621948.1-Frateuria-defendens:0.1174820844,WP_179481164.1-unclassified-Rhodanobacter:0.0734080558)84/78:0.0401125663,((((((WP_214557828.1-unclassified-Rhodanobacter:0.0319211277,WP_068095172.1-unclassified-Rhodanobacter:0.0259939118)32.2/97:0.0173447209,((WP_008437080.1-Rhodanobacter-thiooxydans:0.0373939314,WP_063107523.1-Rhodanobacter-thiooxydans:0.0261374467)37.1/100:0.0100125540,WP_211005527.1-Rhodanobacter-sp.-B2A1Ga4:0.0334641059)85.3/100:0.0164351489)98.5/100:0.0452075827,WP_072762453.1-Rhodanobacter-sp.-OK091:0.0112268477)79.8/37:0.0124267237,((WP_077439480.1-Rhodanobacter-sp.-C01:0.0872664820,WP_077445946.1-Rhodanobacter-sp.-C05:0.0397429251)34.9/72:0.0126403281,WP_158752649.1-Dyella-sp.-S184:0.0276309600)83.1/52:0.0174290525)75.3/37:0.0195091205,WP_184416890.1-Rhodanobacter-sp.-MP7CTX1:0.0409095824)88.1/90:0.0301546142,WP_015446789.1-Rhodanobacter-denitrificans:0.0524027318)60.1/95:0.0219142488)79.1/56:0.0261281449)60.8/30:0.0108787932,(((WP_063674027.1-Dyella-thiooxydans:0.1221447012,WP_253565187.1-Dyella-sp.-Sa:0.0576064810)86.1/98:0.0232445900,(WP_209622196.1-Frateuria-flava:0.0695541640,WP_091339503.1-Frateuria-terrea:0.0400349480)98.4/100:0.0540334178)83.9/98:0.0189056119,(WP_201315287.1-Dyella-sp.-EPa41:0.1012413247,(((WP_168608661.1-Oleiagrimonas-citrea:0.0525029684,WP_113064358.1-Oleiagrimonas-sp.-MCCC-1A03011:0.0222443462)75.6/100:0.0312077981,WP_043099571.1-Oleiagrimonas-soli:0.0777690150)99.9/100:0.1272747997,(WP_188800319.1-Dyella-caseinilytica:0.1286650568,WP_192554051.1-Dyella-acidiphila:0.1132611135)95.5/100:0.0631498648)80.8/87:0.0168942822)51.4/76:0.0217572365)93.8/80:0.0389750844)79.1/48:0.0190820070,((WP_233843762.1-Dyella-sp.-2HG41-7:0.0330175981,WP_188793403.1-Dyella-nitratireducens:0.0677755811)92.3/94:0.0411495624,WP_126673574.1-Dyella-dinghuensis:0.0427670429)99.4/100:0.0865722996)91.3/99:0.0340130596,(WP_111981026.1-Dyella-jiangningensis:0.1338381155,WP_266183414.1-Dyella-sp.-RBB1W86:0.0667713907)90.7/100:0.0236735191)92.3/96:0.0283901326,((WP_131151171.1-Dyella:0.1669129282,WP_130619635.1-Dyella-amyloliquefaciens:0.0553701792)85.3/82:0.0227795557,(WP_266169930.1-Dyella-sp.-RXD159:0.0677820008,(WP_114846719.1-Dyella-tabacisoli:0.0426476928,WP_090451016.1-Dyella-sp.-OK004:0.0620008177)84.8/87:0.0220476059)58.3/82:0.0126447831)0/56:0.0000020686)32.6/72:0.0108025470,WP_199044928.1-Dyella-sp.-ASV24:0.0604692814)90.4/95:0.0380223078,WP_214570707.1-unclassified-Dyella:0.0951678019)97.3/100:0.0936015913)99.9/100:0.1473243497,(((WP_256181981.1-Cloacibacillus-evryensis:0.1799311765,WP_087368268.1-Cloacibacillus-sp.-An23:0.0709259696)100/100:0.3514000347,(((WP_246588317.1-Pelorhabdus-rhamnosifermentans:0.1601563812,(((((WP_092074275.1-Dendrosporobacter-quercicolus:0.1844847877,(WP_145090815.1-Sporomusa-sp.-KB1:0.0462313662,WP_093793074.1-Sporomusa-acidovorans:0.0613742062)98.4/100:0.0917562678)87.4/28:0.0358848274,(WP_054258683.1-Propionispora-sp.-2/2-37:0.1178702929,(WP_149735423.1-Propionispora-hippei:0.0709166258,WP_091744506.1-Propionispora-vibrioides:0.1102745793)84.4/100:0.0356133064)94.8/92:0.0513221750)79.6/6:0.0258770543,((WP_132081660.1-Anaerospora-hongkongensis:0.1020522714,WP_132083235.1-Anaerospora-hongkongensis:0.1284210914)97.4/100:0.0866123321,WP_188400350.1-Sporomusa-sp.-GT1:0.2045416743)23.8/7:0.0254007702)59.2/38:0.0316890675,((WP_188399481.1-Sporomusa-sp.-GT1:0.2150339511,(WP_094603851.1-Sporomusa-silvacetica:0.1007755772,WP_144352952.1-Sporomusa-termitida:0.1192596323)86.4/100:0.0467394994)81.6/84:0.0321549648,WP_054258662.1-Propionispora-sp.-2/2-37:0.1683687222)93.6/94:0.0505001957)75.8/38:0.0158844388,(WP_007952170.1-Pelosinus-fermentans:0.0854007905,WP_007939097.1-Pelosinus:0.0948155228)98.3/100:0.0651265099)90.7/90:0.0402906079)84.9/99:0.0432898881,WP_110954047.1-Massilibacillus-massiliensis:0.1986179878)97.3/100:0.1043057134,((WP_090940141.1-Pelosinus-propionicus:0.1753463285,WP_240986382.1-Acididesulfobacillus-acetoxydans:0.1928594285)83.9/98:0.0441438286,WP_092486361.1-Desulfoscipio-geothermicus:0.1384969166)76.5/89:0.0415643232)86.9/85:0.0457390320)79.2/32:0.0360928670,((((((((WP_200343816.1-Halochromatium-glycolicum:0.1372814463,WP_201222000.1-Halochromatium-roseum:0.1057267500)98.2/100:0.1166597393,(WP_200237937.1-Thiohalocapsa-halophila:0.0472296689,WP_058558062.1-Thiohalocapsa-sp.-ML1:0.1931036385)93.5/100:0.0767739694)97.3/100:0.1088745468,(WP_182584206.1-Thiospirillum-jenense:0.2766342684,(WP_200286492.1-Rhabdochromatium-marinum:0.1065208623,WP_009150811.1-Thiorhodovibrio-sp.-970:0.0670699318)97/100:0.0946582860)94.5/100:0.0962022149)90.8/100:0.0619486006,WP_018175268.1-unclassified-Thioalkalivibrio:0.3060745439)3.2/35:0.0098794492,((((((((WP_200207675.1-Chromatium-weissei:0.1521190440,WP_200154756.1-Chromatium-okenii:0.0785154922)94.8/100:0.0801788796,(WP_200388819.1-Thiocapsa-imhoffii:0.1388598932,((WP_007191850.1-Thiocapsa-marina:0.0353615730,WP_120796078.1-Thiocapsa-rosea:0.0298494081)0/37:0.0052329482,WP_232409912.1-Thiocapsa-bogorovii:0.0183524911)96.2/98:0.0675820998)93.4/100:0.0637839383)90.9/97:0.0490395679,(WP_200158060.1-Allochromatium-vinosum:0.0622429348,WP_200376120.1-Thiocystis-violacea:0.0848727096)93/98:0.0476295605)18.5/77:0.0209833109,((WP_043749566.1-Imhoffiella-purpurea:0.0617012689,WP_007039504.1-Thiorhodococcus-drewsii:0.0531066312)86.5/100:0.0271667021,((WP_164456073.1-Thiorhodococcus-minor:0.0821452864,WP_200330387.1-Thiocystis-violacea:0.0689008882)19.5/100:0.0177589361,WP_164652045.1-Thiorhodococcus-mannitoliphagus:0.0356370043)98.1/100:0.0558679889)85.1/99:0.0184057452)84.3/93:0.0200930175,WP_201092787.1-Thiocystis-minor:0.0664813412)72/88:0.0053383454,WP_132978621.1-Thiobaca-trueperi:0.0844955497)99.3/99:0.1102613540,(WP_100918325.1-Candidatus-Thiodictyon-syntrophicum:0.0774144368,WP_020502667.1-Lamprocystis-purpurea:0.0967015571)97.7/100:0.0792337164)87/100:0.0382333620,WP_089729519.1-Candidatus-Thiosymbion-oneisti:0.2218859586)94.1/99:0.0641493108)96.5/94:0.0835548426,(WP_141730858.1-Oligoflexus-tunisiensis:0.2816715665,WP_101758446.1-Oceanicoccus-sp.-KOV_DT_Chl:0.2758442997)28.7/80:0.0762485235)89.3/79:0.0607187520,WP_141198172.1-Persicimonas-caeni:0.3409505499)92.6/95:0.0643907732,(WP_084578244.1-Sporomusa-malonica:0.3256034939,WP_131919244.1-Heliophilum-fasciatum:0.1354519849)82.6/35:0.0519953190)33.9/18:0.0190574488)87.3/86:0.0408853057)97/95:0.0757956191,((WP_207681674.1-Desulfonema-magnum:0.2956142131,WP_099324654.1-Candidatus-Kuenenia-stuttgartiensis:0.3437193736)86.6/89:0.1056553934,(WP_077329105.1-Virgibacillus-siamensis:0.3411930949,WP_230840800.1-Gloeobacter-morelensis:0.2216332813)62.7/86:0.0656644136)91.4/76:0.0779912180)91.7/80:0.0662482894)82.3/48:0.0487152801,((((((((WP_094934028.1-Enterobacter-cloacae-complex:0.2798937548,WP_086590801.1-Vibrio-parahaemolyticus:0.2083320397)61.2/83:0.0934000236,WP_004741399.1-Vibrio:0.2176567552)82.8/13:0.0368437644,WP_016537220.1-Cedecea-davisae:0.1176119913)44/13:0.0328233776,((WP_010368546.1-Pseudoalteromonas-piscicida:0.2859105444,(WP_039473409.1-Vibrio-sinaloensis:0.2249279792,WP_016795256.1-Vibrio-cyclitrophicus:0.1582539767)80.2/99:0.0365985077)95.5/95:0.0554210001,(((WP_269408820.1-Pseudoalteromonas-shioyasakiensis:0.1423250018,WP_207379479.1-Shewanella-sedimentimangrovi:0.1447989246)90.3/100:0.0498444707,(WP_061013169.1-Vibrionaceae:0.1664829838,WP_155662549.1-Aliivibrio-fischeri:0.1004714170)92.9/100:0.0517113912)81.4/99:0.0337238546,WP_171328835.1-Vibrio-splendidus:0.1745894622)93.5/62:0.0539960211)89.9/23:0.0357080016)92.9/16:0.0808642845,((WP_195504469.1-Citrobacter-farmeri:0.1969771738,WP_064554809.1-Buttiauxella-noackiae:0.2016407840)96.8/100:0.1316131658,(WP_001100815.1-Escherichia-coli:0.2132268916,WP_240304790.1-Vibrio-campbellii:0.2950394304)92.2/31:0.0799072158)19.5/11:0.0337574336)85/30:0.0543262305,WP_057065012.1-Citrobacter:0.3662602399)56.9/98:0.0727916280,WP_216735790.1-Hahella-sp.-HN01:0.1556687304)96.1/96:0.0953726660,(WP_110520589.1-Herpetosiphon-llansteffanensis:0.4285629907,(WP_239493201.1-Luteitalea-sp.-TBR-22:0.2335833304,WP_058867107.1-Chloracidobacterium-thermophilum:0.2836496528)78.9/100:0.0817245770)86.7/96:0.1168346117)92.2/85:0.0884619607)95.9/66:0.0786397815,WP_008811140.1-Parasutterella-excrementihominis:0.3335881277)16.2/58:0.0354849145,(((((((((GJNO01008138.1-Laxus-oneistus-TSA:0.5967650392,(((VDM95900.1-Thelazia-callipaeda:0.1738601509,(((((VDO27950.1-Onchocerca-flexuosa:0.1307548684,VDO31788.1-Brugia-timori:0.0968397500)82/97:0.0226114945,XP_003135693.1-Loa-loa:0.0638040453)91.5/96:0.0243212325,CAG9536712.1-Cercopithifilaria-johnstoni:0.0268483623)82/96:0.0139801211,VBB33646.1-Acanthocheilonema-viteae:0.0820735650)75.6/97:0.0096268974,VDK73328.1-Litomosoides-sigmodontis:0.0330451923)99/100:0.1166840314)99.9/100:0.2218856605,(VDD97390.1-Enterobius-vermicularis:0.2780342686,KHN87800.1-Toxocara-canis:0.1430630080)76.7/100:0.0516750604)62.3/100:0.0561435626,VDN56932.1-Dracunculus-medinensis:0.2324704436)88.8/99:0.0834888243)11.2/65:0.0173163756,(KAI1709308.1-Ditylenchus-destructor:0.1071456221,KAI1730870.1-Ditylenchus-destructor:0.0286979034)100/100:0.1540478717)89.9/92:0.0702084279,((GGJS01006545.1-Plectus-murrayi-TSA:0.2849393133,KAE9552274.1-Halicephalobus-sp.-NKZ332:0.2450493017)71/39:0.0275726150,TKR72632.1-Steinernema-carpocapsae:0.1616124842)76.9/39:0.0427720270)90.1/55:0.0615849906,(KAH7695359.1-Aphelenchus-avenae:0.2408751321,((CAD5228290.1-Bursaphelenchus-okinawaensis:0.0455853028,CAD5235627.1-Bursaphelenchus-xylophilus:0.0282627952)38.7/96:0.0509553367,((KAI6206096.1-Aphelenchoides-besseyi:0.1292205733,KAI6191010.1-Aphelenchoides-bicaudatus:0.0745904666)49.6/100:0.0500564990,KAI6234710.1-Aphelenchoides-fujianensis:0.0921891571)95.4/100:0.0863878529)97.6/100:0.0975616715)88.4/100:0.0706292282)97.6/62:0.1143336761,((CAI4229259.1-Auanema-sp.-JU1783:0.2888355732,((KHJ99538.1-Oesophagostomum-dentatum:0.0513500637,EPB80542.1-Ancylostoma-ceylanicum:0.0420201317)95.8/100:0.0646126775,((VDO65958.1-Haemonchus-placei:0.1172136029,VDL70484.1-Nippostrongylus-brasiliensis:0.0789510972)54.8/100:0.0218928671,VDO95824.1-Heligmosomoides-polygyrus:0.0658815147)99.3/100:0.1182979289)97.1/100:0.0998242350)52/80:0.0400793734,PAV65180.1-Diploscapter-pachys:0.1893012863)39.1/69:0.0136394786)55.4/51:0.0513018127,(CAD6189320.1-Caenorhabditis-auriculariae:0.2820374501,(CAB3400093.1-Caenorhabditis-bovis:0.1528410843,((NP_509046.1-Caenorhabditis-elegans:0.0322570304,XP_003117635.1-Caenorhabditis-remanei:0.0220980400)51.8/100:0.0320343809,CAI5454992.1-Caenorhabditis-angaria:0.0358735630)92.9/100:0.0539613247)87.3/100:0.0658404971)88.6/97:0.0698948397)100/100:0.4564924378,(WP_042281872.1-Candidatus-Protochlamydia-sp.-R18:0.1225371355,WP_068471358.1-Parachlamydia-sp.-C2:0.1060879477)99.9/100:0.3345881728)70.1/95:0.1219169941,(WP_008983868.1-Alishewanella-agri:0.4949675167,WP_265845290.1-Plesiomonas-shigelloides:0.4078029645)88.1/93:0.1038753105)71.3/85:0.0648211203,(WP_255135177.1-Gracilimonas-sp.-CAU-1638:0.1739100776,WP_020401568.1-Gracilimonas-tropica:0.2044481434)100/100:0.2602991412)78/75:0.0792890308)83.6/96:0.0743335483,((((((((WP_171588903.1-Pelistega-europaea:0.2336791379,WP_087653467.1-Klebsiella/Raoultella-group:0.1422731895)77.8/64:0.0194166220,(WP_154751639.1-unclassified-Erwinia:0.1641823911,WP_140470594.1-Ewingella-americana:0.1631046235)40.1/48:0.0284938425)85.4/37:0.0216826142,WP_165428828.1-Rouxiella-badensis:0.0895800735)17.2/6:0.0154967512,(((WP_040666911.1-Neisseria-wadsworthii:0.0841450348,WP_135034416.1-Neisseria:0.1139920818)96.6/99:0.0531268443,(((WP_214236207.1-Rosenbergiella-sp.-S61:0.0512694216,(WP_241647140.1-Rosenbergiella-sp.-JB07:0.0909969587,WP_214195533.1-Rosenbergiella-collisarenosi:0.0757408649)32.1/72:0.0225622305)90.3/98:0.0711267242,((WP_032441370.1-Klebsiella-pneumoniae:0.1039850638,(WP_015583911.1-Raoultella-ornithinolytica:0.0506445245,WP_265776798.1-Klebsiella-oxytoca:0.1225676517)62.5/100:0.0306450921)87/100:0.0465669515,(WP_029991521.1-Tatumella-ptyseos:0.0609482202,(WP_212707699.1-Tatumella-sp.-JGM130:0.0351513895,WP_087487466.1-Tatumella-citrea:0.0472093019)34.7/86:0.0092773958)65.2/93:0.0273842381)79.2/99:0.0318500894)86/98:0.0408955727,WP_101506252.1-Erwinia-sp.-B116:0.1063399675)89/55:0.0383235733)75.9/53:0.0073607179,(((WP_228398973.1-Insectihabitans-xujianqingii:0.0670157903,WP_074823218.1-Pragia-fontium:0.0781582762)93.9/100:0.0468934967,WP_130971788.1-Hafnia-alvei:0.1058030561)90.9/99:0.0354848742,((((((WP_193937755.1-Klebsiella-grimontii:0.1270320870,(WP_142514871.1-Klebsiella-huaxiensis:0.1172197024,(WP_024360036.1-Klebsiella:0.0388928406,WP_234284225.1-Klebsiella-oxytoca:0.0713676918)66/100:0.0282620997)85.9/97:0.0253065063)79.7/94:0.0166792950,(WP_148250181.1-Raoultella-terrigena:0.0795822675,WP_032686784.1-Raoultella-planticola:0.0232910322)94.4/100:0.0367096636)86.4/92:0.0131655482,WP_200519240.1-Klebsiella-michiganensis:0.1051143489)71.1/23:0.0056258831,((WP_131063651.1-Raoultella-planticola:0.0205046016,WP_163980548.1-Raoultella-ornithinolytica:0.0323750325)99.4/100:0.0660786415,(WP_142430937.1-Klebsiella-oxytoca:0.0946591507,((WP_239614184.1-Raoultella-ornithinolytica:0.0757397019,WP_186921353.1-Klebsiella-quasipneumoniae:0.1123172435)0/95:0.0000024288,((WP_032699180.1-Raoultella:0.0414797295,WP_228399925.1-Raoultella:0.0285742570)71/100:0.0073654454,WP_004138711.1-Klebsiella:0.0210020282)97.6/100:0.0450710548)88.7/96:0.0221061095)83.8/86:0.0200899702)72.4/47:0.0048461736)85.2/18:0.0139613332,((((((WP_136167279.1-Brenneria-sp.-CFCC-11842:0.0354526518,WP_264089117.1-unclassified-Brenneria:0.0769315288)36.4/99:0.0295467930,WP_109053612.1-Brenneria-roseae:0.0704737200)87.8/100:0.0249141381,(((((((((((((((((WP_180742451.1-Pectobacterium-sp.-CFBP8739:0.1535073621,WP_039517132.1-Pectobacterium-brasiliense:0.0131516528)83.7/72:0.0142856567,WP_039280054.1-Pectobacterium-brasiliense:0.0269407019)88.9/70:0.0182923357,WP_039346077.1-Pectobacterium-fontis:0.0239993885)75.2/63:0.0068128071,((WP_047710158.1-Pectobacterium-atrosepticum:0.0474559479,(WP_033071760.1-Pectobacterium-parmentieri:0.0248328378,WP_198482318.1-Pectobacterium-parmentieri:0.0339222633)19/63:0.0056858738)94.5/100:0.0329213909,WP_226305805.1-Pectobacterium-versatile:0.0178795893)60.3/98:0.0196925261)90.2/95:0.0212282645,WP_103972204.1-Pectobacterium:0.0119364963)77.1/62:0.0059325433,WP_205558955.1-Pectobacterium-brasiliense:0.0475093388)0/38:0.0000029317,(WP_151242707.1-Pectobacterium-carotovorum:0.0351578067,WP_116186106.1-Pectobacterium-aquaticum:0.0354859317)2.8/73:0.0058445454)91.5/91:0.0260791496,(WP_180777316.1-Pectobacterium-aroidearum:0.0486304418,WP_107169883.1-Pectobacterium:0.0297898518)74.4/99:0.0089834484)68.9/82:0.0118201968,WP_039504461.1-Pectobacterium:0.0209875799)76.3/35:0.0085023437,WP_048259384.1-Pectobacterium-peruviense:0.0353482987)73.4/28:0.0057114933,WP_138255615.1-Pectobacterium:0.0177596513)75.4/31:0.0068345545,(WP_205946867.1-Pectobacterium:0.0476995125,WP_233976643.1-Pectobacterium-versatile:0.0177228892)88.7/96:0.0168024480)69.6/69:0.0114821981,WP_109411993.1-Pectobacterium-polaris:0.0417973042)0/41:0.0000026110,((WP_249680869.1-Pectobacterium-polaris:0.0370821298,WP_125232508.1-Pectobacterium-versatile:0.0389341720)76.2/99:0.0111934472,WP_005974084.1-Pectobacterium-wasabiae:0.0242454156)73.7/100:0.0130080499)86.8/94:0.0181136411,(WP_233968669.1-Pectobacterium-polaris:0.0460313938,WP_119888241.1-Pectobacterium:0.0368454401)67.1/97:0.0154763130)86.8/94:0.0192535405,WP_132452860.1-Samsonia-erythrinae:0.0951749321)0/13:0.0092079557,WP_130637964.1-Pectobacterium-versatile:0.0567488188)80.4/100:0.0237318002)98.8/100:0.0592076706,(((((WP_096000273.1-Klebsiella-pneumoniae:0.1254201861,WP_248464278.1-Pectobacterium-versatile:0.1008021396)79.6/98:0.0207181580,((WP_168640580.1-Dickeya-sp.-CFBP-2040:0.0770160666,(((((WP_038918362.1-Dickeya:0.0262021587,WP_123251184.1-Dickeya-undicola:0.0394261962)73.2/100:0.0059873299,WP_022632763.1-Dickeya-solani:0.0223779526)93.3/100:0.0232657954,WP_236306746.1-Dickeya-zeae:0.0412025094)74.7/34:0.0058299839,(WP_027712673.1-Dickeya-chrysanthemi:0.0352679819,WP_012770533.1-Dickeya:0.0294460895)77.6/66:0.0057874313)74.7/39:0.0060902552,WP_012883965.1-Dickeya-parazeae:0.0546467542)68.3/91:0.0064612547)99.1/100:0.0712730513,(WP_148877341.1-Serratia-marcescens:0.0479116652,WP_193158894.1-Serratia-marcescens:0.0437515606)95.6/100:0.0547922540)61/99:0.0216637543)91.7/95:0.0361234864,((WP_214648782.1-Pectobacterium-carotovorum:0.0297174083,((WP_261846379.1-Pectobacterium-carotovorum:0.0469958900,WP_121590317.1-Brenneria-goodwinii:0.0058774306)95.4/73:0.0231461638,(WP_182099415.1-Pectobacterium-aroidearum:0.0290815915,WP_039482612.1-Pectobacterium:0.0233721672)74.3/68:0.0059603681)15.6/49:0.0057543183)61.1/49:0.0029980948,WP_249228983.1-Brenneria-tiliae:0.0391009360)90.1/91:0.0245635206)41.4/69:0.0101963817,WP_121576612.1-Brenneria-alni:0.0798464021)0/41:0.0000022878,WP_062791489.1-Serratia-plymuthica:0.0731267064)86.5/59:0.0184885935)86.2/75:0.0187178556,WP_095845203.1-Gibbsiella-quercinecans:0.0398331710)93.6/99:0.0375125627,WP_137713144.1-Brenneria-rubrifaciens:0.0810338252)80.6/81:0.0269561308)0/6:0.0000026973,WP_112167140.1-Klebsiella:0.1382803575)78.4/50:0.0149668416)89.3/13:0.0232962597)85.2/6:0.0164717594)87.5/14:0.0324838922,(((WP_253460089.1-Erwinia-persicina:0.1822368039,(((WP_168836617.1-Vibrio-agarilyticus:0.0851343250,(WP_076537303.1-Shewanella-sp.-UCD-KL21:0.0207556238,WP_102531614.1-Shewanella-sp.-10N.286.52.B9:0.0503966645)100/100:0.3463330838)94.7/100:0.0713849033,(WP_137671336.1-Agarivorans-sp.-Toyoura001:0.0477299700,WP_221077378.1-Agarivorans-albus:0.0366976585)95.9/100:0.0755934418)97.8/100:0.0802317183,((((((((((((WP_127021653.1-Rheinheimera-mangrovi:0.2008469470,(WP_138320019.1-Thalassotalea-litorea:0.1106343706,WP_136736185.1-Thalassotalea-mangrovi:0.1065867163)99.9/100:0.1628927829)61.9/98:0.0408709432,WP_114327629.1-Candidatus-Colwellia-aromaticivorans:0.0963141059)84.7/98:0.0362631613,WP_101108574.1-Psychromonas-sp.-psych-6C06:0.1537922104)84/98:0.0532654671,((WP_016794198.1-Vibrio-cyclitrophicus:0.0317331013,WP_102424249.1-Vibrio-cyclitrophicus:0.0379081107)92.4/100:0.0321108024,(WP_261903470.1-Vibrio-fortis:0.0670435822,WP_240207223.1-Vibrio-sp.-CyArs1:0.0294303616)91.2/100:0.0290172848)97.6/100:0.0690354387)86.2/94:0.0542825804,WP_150896168.1-Vibrio-fortis:0.1438964274)74.7/93:0.0534632353,(((WP_260491935.1-Pseudoalteromonas-flavipulchra:0.1504298722,WP_039494249.1-Pseudoalteromonas-flavipulchra:0.0598120759)24.7/88:0.0277852330,WP_055022295.1-Pseudoalteromonas-sp.-P1-26:0.0501930357)86.6/88:0.0345152285,(WP_149602089.1-Pseudoalteromonas-distincta:0.0358866179,WP_248518672.1-Pseudoalteromonas-sp.-2CM32C:0.0182320844)99.1/100:0.0877754305)98.8/100:0.1130404210)42.3/78:0.0464428997,(WP_199145406.1-Citrobacter-sp.-ASV29:0.1791759075,WP_148513713.1-Vibrio:0.1234524487)89.4/97:0.0595352848)84.1/95:0.0364481335,WP_067584343.1-Endozoicomonas-ascidiicola:0.1812064258)88.6/92:0.0473911540,(WP_132584255.1-Rheinheimera-sp.-D18:0.2270817240,((((((WP_131912015.1-Celerinatantimonas-diazotrophica:0.1671490460,((WP_092385961.1-Halopseudomonas-salegens:0.1368182133,WP_036859756.1-Porticoccus-hydrocarbonoclasticus:0.1814404718)92.1/94:0.0725505172,((WP_076722556.1-Marinobacter-lutaoensis:0.1694075597,((WP_106695101.1-Marinobacter-shengliensis:0.0508646493,(WP_092030923.1-Marinobacter-sp.-DSM-26671:0.0883427697,WP_148609564.1-Marinobacter-sp.-BW6:0.0769161574)85.3/89:0.0280069724)68.6/88:0.0081439261,WP_222473125.1-Marinobacter-daepoensis:0.0521008206)97.5/99:0.0693850866)0.9/75:0.0295115241,WP_036130839.1-Marinobacter-nitratireducens:0.1114297874)23.5/94:0.0409131982)87.4/100:0.0395545342)86.3/98:0.0323822528,(WP_027875186.1-Spongiibacter-marinus:0.1199355005,WP_204995064.1-Spongiibacter-marinus:0.0792193213)88.7/99:0.0386008766)92.3/97:0.0345152141,WP_168451636.1-Spongiibacter-thalassae:0.0400706963)92.8/96:0.0480774370,WP_062382560.1-Zhongshania-aliphaticivorans:0.0397480390)99.3/100:0.0830596238,(WP_184464382.1-Zhongshania-antarctica:0.0414367814,WP_103683243.1-Marortus-luteolus:0.0290215954)79.9/99:0.0114234234)26.2/41:0.0256611372,WP_198570926.1-Spongiibacter-nanhainus:0.0466371264)93.7/99:0.0586304025)79.2/99:0.0530421783)94.5/92:0.0524776239,((WP_145010177.1-Pseudomonas-psychrotolerans:0.0129709765,WP_077172422.1-Pseudomonas-psychrotolerans:0.0391116207)99.9/100:0.1606992633,WP_084334293.1-Pseudomonas-indica:0.0897850016)45.1/97:0.0384611958)87.5/89:0.0407074010,(WP_261844981.1-Amphritea-ceti:0.1999111555,((((((WP_269124033.1-Marinomonas-sp.-15G1-11:0.0244558208,WP_191602238.1-Marinomonas-algicola:0.0652374361)99/100:0.0954687838,WP_048693477.1-Catenovulum-maritimum:0.1419343460)90.9/100:0.0543274388,WP_228191388.1-Marinomonas-communis:0.1076274833)18.3/99:0.0236103470,WP_067216963.1-Marinomonas-gallaica:0.0504401564)42.6/95:0.0233032098,WP_067204681.1-Marinomonas-aquimarina:0.0279468400)89.8/96:0.0256210790,WP_055461762.1-Marinomonas-fungiae:0.0459279812)89.1/97:0.0398912648)90.6/99:0.0505252005)84.7/90:0.0307266111,WP_100256321.1-Reinekea-forsetii:0.3625932606)74.9/80:0.0117797292)90.2/97:0.0415823086)63.9/56:0.0355027652,(WP_246757812.1-Erwinia-phyllosphaerae:0.1318972303,WP_232759612.1-unclassified-Pantoea:0.1041323798)92.9/100:0.0499663326)83.5/86:0.0290742648,((WP_146378871.1-Cronobacter-turicensis:0.0547400754,WP_032180402.1-Escherichia-coli:0.0707860775)99.6/100:0.1138550044,(WP_126816534.1-Enterobacter-mori:0.0869739593,WP_256879438.1-Escherichia-albertii:0.0654219046)96/100:0.0584741726)73.7/41:0.0266428122)43.7/38:0.0350831249)86.2/11:0.0309201405,(WP_024553491.1-Franconibacter-helveticus:0.1892072901,((((WP_267333169.1-unclassified-Gilliamella:0.1187661682,(((((WP_086319877.1-Gilliamella-apis:0.0159300335,WP_198214961.1-Gilliamella-sp.-M0320:0.0480473520)92.7/99:0.0258648235,WP_039129858.1-Gilliamella:0.0217032016)74.2/99:0.0090538227,(WP_065737496.1-Gilliamella-apicola:0.0313497181,WP_267331332.1-unclassified-Gilliamella:0.0484256984)96/100:0.0440238387)91.4/100:0.0309076788,((WP_065579324.1-Gilliamella-apis:0.0655792229,WP_160432071.1-Gilliamella-sp.-Pas-s25:0.0904808992)61.2/47:0.0157295303,(WP_065587409.1-Gilliamella-apicola:0.0488704752,WP_065580031.1-Gilliamella-apicola:0.0552842219)90/100:0.0248358929)49.1/41:0.0129282842)35.1/79:0.0220357278,((WP_065574597.1-Gilliamella-apicola:0.0435982377,WP_065569248.1-Gilliamella-apicola:0.0335338009)74.5/98:0.0076332736,WP_065610260.1-Gilliamella-apicola:0.0321792316)98.2/100:0.0514935804)91.2/65:0.0309906811)29/29:0.0115970174,WP_086361229.1-Gilliamella:0.1025198743)26.7/35:0.0105721226,WP_267361838.1-Gilliamella-sp.-B2772:0.1074078213)91.3/99:0.0623659948,(WP_247177500.1-Escherichia-coli:0.2642622094,YP_009985672.1-Escherichia-phage-muut:0.1951136591)99.6/100:0.1956311259)65.1/67:0.0530808231)56.4/6:0.0258460627)92.6/57:0.0587365591,WP_049841125.1-Klebsiella-sp.-RIT-PI-d:0.1958316810)95.2/98:0.0910706258,WP_028357569.1-Brackiella-oedipodis:0.1198604526)93.6/100:0.0750730716)89.3/100:0.0596378433,((WP_133132051.1-Legionella-sp.-W10-070:0.2407641153,((WP_027223798.1-Legionella-pneumophila:0.0286172267,(((WP_044499121.1-Legionella-pneumophila:0.0224895442,(WP_027220106.1-Legionella-pneumophila:0.0141141690,(WP_027266778.1-Legionella-pneumophila:0.0304642286,((WP_031567420.1-Legionella-wadsworthii:0.1482346783,(((WP_232890831.1-unclassified-Legionella:0.0379872338,(WP_058459148.1-Legionella-bozemanae:0.0235307782,WP_058518697.1-Legionella-parisiensis:0.0439472644)89.4/100:0.0272552762)99.7/100:0.0797598136,(WP_058513550.1-Legionella-santicrucis:0.0998330351,(WP_058463758.1-Legionella-cincinnatiensis:0.0189089352,WP_128130294.1-Legionella-sainthelensi:0.0489897152)90.1/100:0.0310636272)50.7/97:0.0208288639)86.9/99:0.0226586475,WP_045097199.1-Legionella-fallonii:0.1013187396)71.5/99:0.0276364598)93.5/99:0.0524350371,(WP_058494182.1-Legionella-worsleiensis:0.1136064713,(WP_127106294.1-Legionella-sp.-km535:0.0339176978,WP_058474153.1-Legionella-quateirensis:0.0279141207)94.1/99:0.0402200316)94.7/100:0.0566948064)96.9/99:0.0591432948)83.5/99:0.0166979926)84.6/99:0.0146637826)87.4/98:0.0119796182,(WP_061513621.1-Legionella:0.0136471880,WP_027226296.1-Legionella-pneumophila:0.0492886704)98.9/94:0.0572174238)71.3/91:0.0038257388,WP_106142999.1-Legionella-pneumophila:0.0336730678)94/97:0.0272885471)82/98:0.0120140394,WP_265717942.1-Legionella-sp.-PATHC032:0.0221718849)100/100:0.1931819665)85.5/100:0.0867903971,(WP_205000141.1-Parashewanella-hymeniacidonis:0.0558628537,WP_129946484.1-Shewanella-sp.-OPT22:0.0370256176)100/100:0.3541492337)68.5/92:0.0669535879)98.2/100:0.1112914226)44.6/70:0.0554374099,WP_011189464.1-Desulfotalea-psychrophila:0.2432321666)94.4/94:0.0793675746,((((((((((((((((((WP_167868984.1-Shewanella-sp.-WXL01:0.2828022733,(WP_138986566.1-Neptunomonas-concharum:0.2395500074,WP_047049276.1-Vibrio-mexicanus:0.0767622629)41.5/98:0.0353394564)89.1/98:0.0314353191,WP_264874461.1-Vibrio-agarivorans:0.0764666919)68.8/60:0.0115507431,WP_020335761.1-Vibrio-natriegens:0.0863184437)92.4/60:0.0236864471,((((((WP_222525605.1-Ferrimonas-balearica:0.1000125245,WP_013344594.1-Ferrimonas-balearica:0.1045299300)99/100:0.1216582466,WP_047874860.1-Photobacterium-aphoticum:0.1387542661)78.2/28:0.0196745605,(WP_193630602.1-Shewanella-benthica:0.0828586292,WP_096084688.1-Agaribacterium-haliotis:0.2420803967)94.1/29:0.0669936724)53.6/85:0.0247848072,WP_012143346.1-Shewanella-sediminis:0.1461406387)83.3/73:0.0172106396,(((((WP_167678613.1-Shewanella-aestuarii:0.1115935275,WP_137222113.1-Shewanella-sp.-MEBiC00475:0.0273637093)82.9/100:0.0304686483,WP_188955407.1-Shewanella-ulleungensis:0.0516572813)94.2/100:0.0572694249,((WP_138577097.1-unclassified-Pseudoalteromonas:0.0838943749,(WP_264390127.1-Pseudoalteromonas-sp.-A3:0.0209203197,(WP_096739047.1-Pseudoalteromonas-atlantica:0.0341839038,WP_248462211.1-Pseudoalteromonas-sp.-2CM37A:0.0159634368)92.4/100:0.0195583618)76.1/65:0.0102479206)80/33:0.0290657143,((WP_182693438.1-Pseudoalteromonas-sp.-SR45-5:0.0559453068,WP_256848444.1-Pseudoalteromonas-carrageenovora:0.0153089508)76.5/54:0.0084158485,WP_126005793.1-Pseudoalteromonas-sp.-Xi13:0.0280270853)74.4/31:0.0083647931)64.6/27:0.0401836405)18.4/99:0.0241285210,WP_155694061.1-Psychrosphaera-haliotis:0.1070372245)80.2/95:0.0223136939,(WP_022943097.1-Psychromonas-hadalis:0.1094549388,(WP_113818410.1-Psychromonas-sp.-B3M02:0.0436074453,(WP_025564776.1-Psychromonas-sp.-SP041:0.0393926965,WP_166372414.1-Psychromonas-sp.-SA13A:0.0132758415)93.9/100:0.0472745142)78.6/98:0.0270050197)90.2/93:0.0360479086)86.1/93:0.0217489624)79.9/26:0.0166905222,((WP_050648448.1-Vibrio-sp.-J2-4:0.1039353039,WP_205390674.1-Vibrio:0.0711927883)1.5/20:0.0117852569,WP_140076235.1-Vibrio-parahaemolyticus:0.1045828066)80.4/31:0.0186337617)79.6/37:0.0086791073)85.8/33:0.0117594522,WP_164649367.1-Vibrio-astriarenae:0.0654717536)92.7/33:0.0246079545,(WP_084884043.1-Vibrio-sp.-qd031:0.0931824718,((WP_269599241.1-Salinivibrio-kushneri:0.0501513290,WP_136483218.1-Vibrio-sp.-H11:0.0635447877)93.5/100:0.0534412569,WP_159656123.1-Vibrio-atypicus:0.0864869965)72.7/59:0.0257403120)88/40:0.0203788106)79.5/60:0.0108389731,WP_093326971.1-Thalassotalea-agarivorans:0.0952505381)32.9/19:0.0345149607,(WP_016959527.1-Enterovibrio-norvegicus:0.0521536694,WP_229491984.1-Enterovibrio-norvegicus:0.0981312932)93.9/99:0.0498774167)92.9/99:0.0690611027,(((WP_142941196.1-Aliikangiella-marina:0.2040901379,(WP_081148784.1-Cognaticolwellia-beringensis:0.0341000975,WP_007634918.1-Paraglaciecola-psychrophila:0.0376015450)100/100:0.1483771401)87.9/97:0.0568607192,WP_068546663.1-Thalassotalea-crassostreae:0.1366534272)42.9/93:0.0287165612,(((WP_248714967.1-Agarivorans-sp.-TSD2052:0.0413735625,WP_152781274.1-Agarivorans-sp.-B2Z047:0.0748484519)99.8/100:0.1269750095,(WP_220103444.1-Neiella-holothuriorum:0.0862393863,WP_087505717.1-Neiella-marina:0.1128908347)97.1/100:0.0618784282)49.5/50:0.0558136763,(((WP_028862352.1-Psychromonas-aquimarina:0.1359730704,WP_187408568.1-Saccharophagus-sp.-K07:0.1204755307)70.6/91:0.0239336742,WP_188151970.1-Teredinibacter-waterburyi:0.2041933039)23.8/92:0.0368889567,WP_251261538.1-Echinimonas-agarilytica:0.1170654571)91.2/96:0.0466484778)33/63:0.0331983015)84.8/45:0.0430085380)86.4/95:0.0339382879,((WP_250459773.1-Microbulbifer-sp.-GX-H0434:0.1122175489,WP_152451766.1-Microbulbifer-sp.-THAF38:0.1010557716)90/100:0.0498503748,WP_009839692.1-Pseudoalteromonas-tunicata:0.3402718426)54.7/68:0.0744327738)82.2/71:0.0407965559,(WP_124026287.1-Alteromonas-sediminis:0.1165769823,WP_100655941.1-Alteromonas-flava:0.1686409905)99.4/97:0.1147066159)70.2/8:0.0091662196,((WP_118961222.1-Colwellia-sp.-RSH04:0.2402841401,(WP_216048042.1-Aliiglaciecola-lipolytica:0.1271832044,WP_057181266.1-Colwellia:0.1559586249)41.3/90:0.0154047622)39.9/88:0.0411666020,(WP_131095783.1-Alteromonas-sp.-KUL42:0.0772699722,((((((((WP_100972994.1-Alteromonas-sp.-MB-3u-76:0.0854237039,WP_269520955.1-Alteromonas-sp.-BMJM2:0.0388566629)87.6/98:0.0241662739,((WP_071816068.1-Alteromonas-sp.-V450:0.0796176858,WP_198816918.1-Alteromonas-sp.-IB21:0.0139265103)79.4/100:0.0151337212,WP_163085646.1-Alteromonas-profundi:0.0500904961)57.2/49:0.0148425388)68.1/51:0.0124637276,WP_163106912.1-Alteromonas-genovensis:0.0316783714)82.5/48:0.0136925942,WP_057795858.1-Alteromonas-stellipolaris:0.0258096698)63.4/41:0.0061253713,(((WP_156078883.1-Alteromonas:0.0449237497,WP_202262575.1-unclassified-Alteromonas:0.0283442958)84/81:0.0133534799,WP_187447473.1-Alteromonas-sp.-BZK5:0.0172222370)74.2/75:0.0051928482,((WP_063457834.1-Alteromonas-stellipolaris:0.0281162018,(WP_202940767.1-Alteromonas-macleodii:0.0196262928,WP_061095937.1-Alteromonas-macleodii:0.0331634869)81.1/74:0.0098677969)80.4/89:0.0055326014,((WP_044058190.1-Alteromonas-australica:0.0340863691,WP_159580202.1-Alteromonas-sp.-KUL106:0.0169090686)89/100:0.0112257815,(WP_014977514.1-Alteromonas-macleodii:0.0283489265,WP_239220085.1-Alteromonas-sp.-MmMcT2-2:0.0225248315)0/96:0.0000028985)52.5/87:0.0056105410)0/92:0.0000025851)95.6/93:0.0224685786)75.5/77:0.0089034842,(WP_130638173.1-Alteromonas-sp.-KUL49:0.0455613870,WP_115592514.1-Alteromonas-aestuariivivens:0.0379277932)89.9/96:0.0202393226)90.2/97:0.0207381861,(((((WP_018984642.1-Salinimonas-chungwhensis:0.0732557977,WP_265616768.1-Alteromonas-sp.-ASW11-7:0.0614853423)89.4/99:0.0298195058,WP_119499891.1-Alteromonas-sp.-RKMC-009:0.0503194811)19.3/73:0.0131963967,WP_265607972.1-Alteromonas-sp.-ASW11-130:0.0645556782)81/64:0.0112747268,(WP_216028209.1-Alteromonas-sp.-C1M14:0.0569950649,WP_216026947.1-Aestuariibacter-sp.-A3R04:0.0636242618)0/32:0.0000021144)89.6/53:0.0173236886,(WP_070176406.1-Alteromonas-lipolytica:0.0247196056,WP_073325432.1-Marisediminitalea-aggregata:0.0473200088)96.3/100:0.0370636120)93.6/94:0.0253348179)91.4/92:0.0173972664,(WP_262993134.1-Alteromonas-sp.-ASW11-19:0.0501280242,(WP_218354448.1-Alteromonas-lipotrueiana:0.0608813467,WP_195811108.1-Salinimonas-marina:0.0480386948)88/99:0.0478473425)79.1/95:0.0139105026)94.4/89:0.0328050252)79.4/98:0.0189925982)81.6/88:0.0433052084)1.6/28:0.0211577816,(WP_102798355.1-Bowmanella-denitrificans:0.0619946161,WP_214344498.1-Bowmanella-yangjiangensis:0.0269971990)100/100:0.1331639150)96.8/94:0.0687531446,WP_186505417.1-Neptunicella-marina:0.1373610892)88/96:0.0294823853,(WP_160153838.1-Microbulbifer-sp.-ALW1:0.1886885053,((WP_263600466.1-Microbulbifer-agarilyticus:0.1087074650,WP_078085574.1-Microbulbifer-mangrovi:0.0707949054)17.2/53:0.0098345188,WP_029249715.1-Microbulbifer-agarilyticus:0.0865003859)93.2/98:0.0587648932)97.1/99:0.0801171919)90.8/94:0.0457387841,((WP_138624780.1-Pseudoalteromonas-sp.-S4492:0.1200153298,WP_130050482.1-Pseudoalteromonas-sp.-MEBiC-03485:0.1370583225)74.8/99:0.0462617209,(WP_189610147.1-Saccharospirillum-salsuginis:0.1737047002,WP_111640782.1-Marinimicrobium-alkaliphilum:0.2244712367)90.2/98:0.0612754533)87.1/99:0.0675272605)91.6/99:0.0532576785,(((WP_085356321.1-Neisseria-dumasiana:0.1596007680,(WP_127202078.1-Neisseria-meningitidis:0.0989984526,WP_237092367.1-Neisseria-sp.-ZJ104:0.1070493588)14.9/77:0.0333111547)61.9/85:0.0099278895,WP_101133177.1-Neisseria-meningitidis:0.0630925310)79.6/88:0.0428153473,WP_005552781.1-Aggregatibacter-actinomycetemcomitans:0.1164718899)99.9/100:0.1134909010)88.2/99:0.0296466457,WP_031579865.1-Ruminobacter-sp.-RM87:0.2729639008)94.2/97:0.0517611592)88.5/66:0.0278056231,((WP_144135068.1-Kosakonia-radicincitans:0.1998329161,((WP_088208149.1-Enterobacter-cancerogenus:0.0785282324,WP_121932958.1-Enterobacter-hormaechei:0.0807088805)35.1/100:0.0346348609,WP_023330866.1-Enterobacter:0.0441898836)97.1/100:0.0780519747)69.4/98:0.0478310978,(WP_120453320.1-Kalamiella-piersonii:0.0856269280,((((WP_214216788.1-Rosenbergiella-epipactidis:0.0462699624,WP_241623482.1-Rosenbergiella-epipactidis:0.0215911905)99.6/100:0.1001315977,(WP_192411534.1-Pantoea-sp.-PNT02:0.0644937007,(((WP_125311258.1-Pantoea-dispersa:0.0669620602,(WP_069025179.1-Pantoea-agglomerans:0.0107868801,WP_140917215.1-Pantoea-deleyi:0.0387089261)92.6/90:0.0273224344)87/61:0.0172196743,WP_097116658.1-Pantoea-sp.-GL120224-02:0.0093760017)91.5/59:0.0259005741,WP_150019269.1-unclassified-Pantoea:0.0505200552)61.9/95:0.0127625134)96.3/100:0.0585950356)88.2/97:0.0330044572,WP_187508308.1-Pantoea-agglomerans:0.1273452297)83.8/84:0.0318396748,(WP_269294270.1-Pantoea-sp.-LMR881:0.0345852191,WP_215846277.1-Pantoea-sp.-EnvD:0.0662031947)98/100:0.0621414531)72/82:0.0162313767)77.4/79:0.0312489670)96.2/98:0.0856815046)77/24:0.0124371543,WP_105076797.1-Pantoea-ananatis:0.2619437322)83.3/82:0.0278207646,(((WP_188026786.1-Tolumonas-osonensis:0.0161682540,WP_024870911.1-Tolumonas-lignilytica:0.1039203441)99.9/100:0.1135488751,(WP_187463805.1-Photobacterium-sp.-BZF1:0.0478550513,(WP_039467548.1-Photobacterium-gaetbulicola:0.0549564602,WP_146157089.1-Photobacterium-rosenbergii:0.1239401700)93.7/73:0.0640068869)25.5/37:0.0176336732)90.1/66:0.0434596576,WP_047883806.1-Photobacterium-ganghwense:0.0944742868)88.1/85:0.0410916051)62.8/39:0.0081687636,((WP_075764060.1-Photobacterium-proteolyticum:0.0788422838,(WP_222242368.1-Pseudoalteromonas-piscicida:0.2149325445,((WP_069589493.1-Salinivibrio-sp.-DV:0.0783478503,WP_077662143.1-Salinivibrio-proteolyticus:0.0151445462)90.7/97:0.0402240995,((WP_077659434.1-Salinivibrio-kushneri:0.0974719847,WP_077659452.1-Salinivibrio-kushneri:0.0230269617)86.3/55:0.0235364571,WP_269578450.1-Salinivibrio-kushneri:0.0648277948)44.2/55:0.0195686624)99.6/99:0.1487619782)90.4/99:0.0710320756)42.9/93:0.0244934073,WP_239411815.1-Photobacterium-sp.-OFAV2-7:0.1120588878)98.9/85:0.0609241601)95/76:0.0371478796,(((((WP_020431327.1-Vibrio-fluvialis:0.0675611870,WP_055453311.1-Vibrio-fluvialis:0.0370050637)90.2/100:0.0309742619,(WP_017004373.1-Enterovibrio-norvegicus:0.1481465438,(WP_114765998.1-Vibrio-rhodolitus:0.0942901199,WP_182108830.1-Vibrio-marinisediminis:0.0630656254)92/100:0.0557777341)87/81:0.0442644660)82.8/58:0.0243873927,((WP_107253996.1-Photobacterium-indicum:0.1058720807,(WP_138627230.1-Pseudoalteromonas-sp.-S4741:0.1237177439,WP_269377025.1-Vibrio-sinaloensis:0.1233757504)85.6/100:0.0364959748)85/81:0.0454367282,(WP_193290728.1-Vibrio-parahaemolyticus:0.1712830129,((WP_075989065.1-Vibrio-fluvialis:0.0337750100,WP_044366945.1-Vibrio-fluvialis:0.0248164028)93.4/100:0.0529073565,(WP_076657081.1-unclassified-Vibrio:0.0540327256,WP_239929047.1-Vibrio-kanaloae:0.0271770417)95.8/100:0.0578734158)95.3/100:0.0806835452)63.6/18:0.0305139464)51.6/28:0.0220385417)72.5/15:0.0175025647,((WP_200225058.1-Rahnella-aceris:0.0902937164,WP_195814554.1-Rahnella:0.0877033921)84.4/100:0.0401276513,WP_169403689.1-Rouxiella-aceris:0.0644554600)100/100:0.1331159577)79.2/26:0.0186932723,(WP_117235674.1-Vibrio-maerlii:0.1029262840,(WP_103882085.1-Vibrio-hangzhouensis:0.0449079614,WP_265676699.1-Vibrio-sp.-ZSDZ65:0.0903519331)89.5/99:0.0316805278)79.7/93:0.0381752048)75.1/27:0.0109339242)93.7/84:0.0321856917,(((WP_261600599.1-Photobacterium-sp.-TY1-4:0.0799043797,((WP_170899585.1-Vibrio-sp.-3-2-1:0.0970260965,WP_017221010.1-Moritella-dasanensis:0.0829573074)93.2/100:0.0373682004,WP_107284330.1-Photobacterium-lipolyticum:0.0698166961)91.5/99:0.0371611766)64.5/93:0.0126803361,WP_159197271.1-Escherichia-coli:0.0906787716)86.8/86:0.0242863206,(((((WP_211647400.1-Photobacterium-sp.-GJ3:0.0461258746,WP_120511428.1-Photobacterium-salinisoli:0.0289943541)91.7/100:0.0422943801,(WP_172533652.1-Vibrio-aestuarianus:0.2027124212,WP_046218796.1-Photobacterium-halotolerans:0.1211085961)74.3/76:0.0436008251)80.9/70:0.0465445536,(((WP_112404745.1-Vibrio-diazotrophicus:0.0336552986,WP_202661182.1-Vibrio-fluvialis:0.0520057073)92.2/100:0.0457413967,((((WP_263100201.1-Kosakonia-sp.-ML.JS2a:0.0236219681,WP_200550470.1-Kosakonia-sp.-LAM2021:0.0758227369)99.4/100:0.0932789745,((WP_015854950.1-Musicola-paradisiaca:0.0301221901,WP_192457620.1-Musicola-keenii:0.0792556786)99.9/100:0.1394682198,((WP_232612305.1-Photobacterium-phosphoreum:0.1827666754,WP_136989485.1-Vibrio-sp.-F13:0.1988985340)82.8/100:0.0528847979,WP_023404414.1-Vibrio-halioticoli:0.1636157681)84.4/65:0.0467654748)43.4/6:0.0331728626)83.2/6:0.0200311555,WP_094109047.1-Lonsdalea-iberica:0.0953754949)9.9/6:0.0188435236,(WP_026742149.1-Lonsdalea-quercina:0.0277717534,WP_094117837.1-Lonsdalea-britannica:0.0288047907)98.6/100:0.0695051872)94.8/84:0.0677733251)70.3/68:0.0404927086,(WP_047463293.1-Vibrio-fluvialis:0.0897493451,WP_148560072.1-Vibrio-mimicus:0.1206879586)94.1/99:0.0528305617)22.2/46:0.0283412634)69.1/21:0.0091758571,WP_124938567.1-Vibrio-viridaestus:0.1068146078)74.8/22:0.0161783547,((WP_107294031.1-Photobacterium-phosphoreum:0.0307032051,WP_107276976.1-Photobacterium-sp.-NCIMB-13483:0.0533428124)99.5/100:0.1044721825,WP_088133934.1-Vibrio-gazogenes:0.1873117767)0/14:0.0091984631)80.4/79:0.0100562796)88.6/91:0.0217704451)72.7/10:0.0060367915,(WP_268679810.1-Vibrio-barjaei:0.0877628246,(WP_070968050.1-Vibrio-sonorensis:0.1592453376,(WP_123015137.1-Vibrio-zhugei:0.0348307402,WP_077313846.1-Vibrio-palustris:0.0205766634)97.2/100:0.0686999162)95.5/98:0.0635705547)83.3/55:0.0189155719)89/58:0.0220726813,(((((((WP_239941760.1-Vibrio-mediterranei:0.0754255915,((WP_062462511.1-Vibrio-mediterranei:0.0547533794,(WP_171392669.1-Vibrio:0.0768997261,(WP_120356561.1-Alginatibacterium-sediminis:0.2061501795,WP_234495031.1-Vibrio-maritimus:0.1239878335)73.7/99:0.0389135931)86.9/92:0.0296485716)84/89:0.0399850206,WP_252012765.1-Vibrio-sp.-SCSIO-43140:0.1018662278)43.3/81:0.0248623808)86.9/79:0.0322763176,((WP_252012797.1-Vibrio-sp.-SCSIO-43140:0.1072360320,(WP_102532726.1-Vibrio-sp.-10N.286.48.B7:0.0671880421,WP_250617847.1-Vibrio-sp.-S4B1:0.0588945534)78.7/86:0.0333503274)48.8/32:0.0079132666,WP_101064017.1-Moritella-sp.-Urea-trap-13:0.1637024793)47.7/29:0.0251357557)83.3/95:0.0212985733,WP_221070344.1-Vibrio-alfacsensis:0.0883374946)92.4/99:0.0429734389,(((WP_190336189.1-Vibrio-sp.-S9_S30:0.0650221025,WP_022612811.1-Vibrio-nigripulchritudo:0.0848640079)98.3/100:0.0641005086,((WP_182744595.1-Shewanella-sp.-SR44-4:0.1503210059,WP_152797327.1-Salinivibrio-sp.-VYel1:0.1362066493)82.8/100:0.0273836699,WP_039484180.1-Vibrio-sinaloensis:0.1197200409)81.8/100:0.0267521711)81.1/56:0.0196636410,WP_045898906.1-Enterobacteriaceae:0.1161927093)80.8/37:0.0161323396)75.7/46:0.0178547800,((WP_069212356.1-Vibrio-cholerae:0.0503458723,WP_202641945.1-Vibrio-fluvialis:0.0859067863)97.6/100:0.0893027950,WP_068898547.1-Enterovibrio-pacificus:0.1207889484)5.7/29:0.0172370917)84.9/44:0.0271270864,((WP_161119826.1-Photobacterium-lucens:0.0918357206,WP_065194916.1-Photobacterium-phosphoreum:0.1190540756)56.4/8:0.0359246473,((WP_094127815.1-Vibrio-sp.-V09_P4A23P171:0.0774424116,WP_199867560.1-Vibrio-anguillarum:0.0748042304)92/100:0.0492880751,WP_211689658.1-Moritella-sp.-28:0.2113155427)81.4/51:0.0234564297)15.6/6:0.0136333969)25.1/6:0.0194754693,(((WP_192015074.1-Photobacterium-arenosum:0.0691400417,WP_088876367.1-Vibrio-mediterranei:0.1617674121)83/94:0.0399922414,(WP_028024862.1-Enterovibrio-calviensis:0.1411827661,WP_017007824.1-Enterovibrio-norvegicus:0.1377973279)3.6/90:0.0250555260)61.3/94:0.0217875226,((WP_101201417.1-Paraglaciecola-sp.-MB-3u-78:0.1330348641,WP_101040106.1-Psychromonas-sp.-MB-3u-54:0.1365122595)58.8/100:0.0490468965,WP_036979211.1-Providencia-alcalifaciens:0.1534429521)85.1/96:0.0348753098)89.5/88:0.0372151163)86.8/30:0.0222866404)92.9/37:0.0394501733,(((((WP_065208126.1-Photobacterium-phosphoreum:0.0919835923,(WP_232582982.1-Photobacterium-phosphoreum:0.0367132386,((WP_256383785.1-Photobacterium-toruni:0.0341480024,WP_232578536.1-Photobacterium-carnosum:0.0448747435)0/70:0.0000022084,WP_107228404.1-Photobacterium-phosphoreum:0.0742350743)95.8/100:0.0372007118)94.9/100:0.0451606308)71.4/39:0.0153689446,WP_060997345.1-Photobacterium-aquimaris:0.0880371979)75.4/53:0.0122637440,(WP_107293979.1-Photobacterium-phosphoreum:0.0923871957,(WP_087853896.1-Photobacterium-andalusiense:0.0466084734,WP_107274737.1-Photobacterium-phosphoreum:0.0676849736)97.9/100:0.0594424048)87.6/55:0.0253167765)60.2/26:0.0112004197,(((WP_232575020.1-Photobacterium-carnosum:0.0740776283,WP_232585319.1-Photobacterium-iliopiscarium:0.0850469969)78/54:0.0218339155,((WP_045045404.1-Photobacterium-kishitanii:0.0305907279,(WP_045030671.1-Photobacterium-phosphoreum:0.0419053753,WP_107264793.1-Photobacterium-phosphoreum:0.0203671954)80.1/95:0.0162269627)94.4/98:0.0418789974,(WP_107295906.1-Photobacterium-phosphoreum:0.1047281220,WP_107228425.1-Photobacterium-phosphoreum:0.0951960999)92.8/62:0.0522807717)82.1/51:0.0273321468)66.1/52:0.0250655957,WP_232600381.1-Photobacterium-carnosum:0.0640553691)71.4/30:0.0252652211)86.8/91:0.0257095414,((((WP_151182432.1-Photobacterium-damselae:0.0946965919,(WP_053987023.1-Photobacterium-leiognathi:0.0658026301,WP_086367281.1-Photobacterium-damselae:0.0514592740)16.2/85:0.0117450339)79.4/84:0.0206291596,(WP_106261939.1-Photobacterium-damselae:0.0828350160,WP_115426223.1-Photobacterium-damselae:0.0319139950)72.3/91:0.0255141976)86.2/86:0.0185869223,((((WP_238087945.1-Photobacterium-damselae:0.0537928326,(WP_065172005.1-Photobacterium-damselae:0.0596083922,WP_107185088.1-Photobacterium-leiognathi:0.0596728381)44.1/98:0.0164041245)91/100:0.0287453573,WP_151170663.1-Photobacterium-damselae:0.0202089423)78.2/96:0.0194265763,WP_045063675.1-Photobacterium-leiognathi:0.0611173818)81.2/98:0.0170992873,WP_237041371.1-Photobacterium-damselae:0.0679469850)50.1/71:0.0116119552)91.5/96:0.0326130164,(WP_237042700.1-Photobacterium-damselae:0.0438881951,(WP_013365656.1-_Enterobacter:0.1362501902,(((WP_258988101.1-Scandinavium-hiltneri:0.0649431215,WP_181620050.1-Enterobacter-sp.-RHBSTW-00994:0.0446272099)96.5/100:0.0526540068,WP_193129600.1-Klebsiella-aerogenes:0.0671496095)94.5/100:0.0534737162,WP_214579582.1-Enterobacter-asburiae:0.0669557371)89.7/100:0.0445940546)100/100:0.1488532066)0/49:0.0199795799)85.4/48:0.0212320785)80/63:0.0120558370)85.5/37:0.0160428908,(((((WP_208891742.1-Vibrio-anguillarum:0.1051251046,WP_148533341.1-Vibrio-cholerae:0.0399727303)80.5/94:0.0195603066,(WP_045460498.1-Vibrio-hyugaensis:0.1190903135,WP_107294203.1-Photobacterium-sp.-GB-50:0.1023076263)47/89:0.0443228052)81.8/80:0.0220406426,(((WP_000864156.1-Vibrio-cholerae:0.0363022788,WP_000864152.1-Vibrio-cholerae:0.0777440905)89.6/100:0.0394823611,(WP_188234612.1-Vibrio-sp.-Y2-5:0.0529716844,WP_057556016.1-Vibrio-cholerae:0.0150545994)100/100:0.1472314156)80.8/50:0.0228039845,(WP_095464062.1-Vibrio-cholerae:0.0511895427,WP_149560528.1-Vibrio-cholerae:0.0731328396)88.6/96:0.0372423084)82.9/18:0.0221982209)85.7/19:0.0210310156,(WP_102786270.1-Vibrio-cholerae:0.0619187738,WP_095478449.1-Vibrio:0.0695474506)80.9/21:0.0239561250)74.9/55:0.0065065293,(WP_175242188.1-Vibrio-cholerae:0.0538799384,WP_199351799.1-Vibrio-cholerae:0.0667186780)92.3/85:0.0436148384)90/58:0.0194677793)77/77:0.0055597674,((((((((((((((((WP_011771697.1-Psychromonas-ingrahamii:0.0793209572,WP_182690147.1-Psychromonas-sp.-SR45-3:0.1073620573)98.7/100:0.0780801072,(((WP_045569365.1-Vibrio:0.0633622825,WP_039437343.1-Vibrio-navarrensis:0.0342019258)98.2/100:0.0733205020,WP_176299354.1-Vibrio-sp.-05-20-BW147:0.1136864769)92.4/100:0.0498406045,WP_140233673.1-Shewanella-polaris:0.1556522060)91.2/100:0.0654796248)58.1/92:0.0243233357,(WP_044365158.1-Vibrio-fluvialis:0.1923735467,WP_088905011.1-Shewanella-marisflavi:0.1532998066)50.7/93:0.0377861457)90.8/92:0.0402080135,(((WP_238813939.1-Vibrio-vulnificus:0.0852759216,(WP_182754472.1-Shewanella-sp.-SG41-3:0.0857473944,(WP_192022372.1-Shewanella-sp.-WPAGA9:0.1367153294,WP_248975945.1-Shewanella-kaireitica:0.1142556782)94.5/99:0.0591624840)80.9/79:0.0235516563)65.5/39:0.0269377019,((WP_061023752.1-Vibrio-splendidus:0.0438915196,(WP_220768829.1-Shewanella-sp.-MBTL60-007:0.0828071941,(WP_110456309.1-Shewanella-algidipiscicola:0.1338398636,WP_204999274.1-Parashewanella-hymeniacidonis:0.1051366288)87.7/86:0.0494308331)47.1/80:0.0153144887)93.7/92:0.0453263838,(WP_248991952.1-Shewanella-profunda:0.0901033981,WP_059744187.1-Shewanella-frigidimarina:0.1422278693)86/80:0.0302975127)2/56:0.0302804365)76.8/52:0.0217927169,(WP_011638275.1-Shewanella-frigidimarina:0.0816224217,(WP_220774251.1-Shewanella-glacialipiscicola:0.0525042636,WP_182751750.1-Shewanella-sp.-SG44-2:0.0893320717)96.2/100:0.0764276976)94.1/97:0.0512786756)75.2/48:0.0111552789)38.6/45:0.0151807531,((((WP_102457123.1-Vibrio-breoganii:0.0792756425,(WP_108083222.1-Vibrio-sp.-ZF-223:0.1050114362,(WP_102532836.1-Vibrio-sp.-10N.286.48.B7:0.0667374852,WP_254101081.1-Alteromonas-sp.-LMIT007:0.1072381885)72.2/99:0.0293647061)92/97:0.0411219682)90.3/91:0.0303085311,(WP_172380549.1-Vibrio-sp.-Vb339:0.0856915869,(WP_102443327.1-Vibrio-breoganii:0.0294014209,WP_261923504.1-Shewanella-sp.-NFH-SH190041:0.1779606671)92.4/97:0.0471366987)93.4/98:0.0473705986)88/83:0.0252457326,WP_077647682.1-Salinivibrio-sp.-IB643:0.1004769446)72.8/69:0.0069330888,(((((WP_017027407.1-Vibrio-breoganii:0.0930702596,(((((WP_095430409.1-Vibrio-vulnificus:0.0518774371,WP_114994956.1-Grimontia-hollisae:0.1414543878)97.1/100:0.0667355475,WP_118894008.1-Vibrio-vulnificus:0.0786655786)88.3/99:0.0357380036,(WP_239850080.1-Shewanella-sp.-Isolate8:0.0859207934,WP_211678393.1-Moritella-sp.-36:0.1851303292)69.2/97:0.0210797888)41.3/54:0.0237422155,WP_239872404.1-Shewanella-insulae:0.0814124285)63.3/94:0.0282612459,WP_102448490.1-Vibrio-splendidus:0.0932050672)86.8/95:0.0227789919)32/51:0.0121683380,WP_193262143.1-Vibrio-parahaemolyticus:0.1872509332)37.7/49:0.0123215494,(WP_232600401.1-Photobacterium-carnosum:0.0999452545,(WP_138589904.1-Pseudoalteromonas-aurantia:0.0958829015,((WP_218529053.1-Shewanella-sp.-NIFS-20-20:0.0935800258,WP_225483056.1-Vibrio-vulnificus:0.0652451923)92.5/100:0.0458294595,WP_012549058.1-Aliivibrio-salmonicida:0.1361548525)70.2/97:0.0358229610)87.6/97:0.0391057686)84.1/86:0.0248731850)11.3/45:0.0069120404,WP_203433235.1-Shewanella-sp.-LZH-2:0.1548658684)42.6/49:0.0116716080,WP_017449116.1-Vibrio-parahaemolyticus:0.1023662260)77.4/51:0.0144646583)95/55:0.0401944223)90.6/87:0.0229972270,(((((WP_155660100.1-Aliivibrio-fischeri:0.0653354456,WP_095457151.1-Vibrio:0.0765672907)91/100:0.0331157304,(WP_252020469.1-Vibrio-sp.-SCSIO-43136:0.0862235726,WP_243584106.1-Vibrio-splendidus:0.1226348406)75.2/100:0.0303238604)93.3/100:0.0481827246,(WP_108452928.1-Enterobacter-cloacae:0.2477618904,WP_166696422.1-Providencia-rettgeri:0.0579109366)82.1/98:0.0698565808)96.1/98:0.0558475062,(WP_176284769.1-Photobacterium-damselae:0.1132539927,WP_220058598.1-Shewanella-sp.-FJAT-53555:0.1752691695)70.5/82:0.0326765903)72.9/19:0.0071671737,(((((WP_232603643.1-Photobacterium-carnosum:0.0381940781,WP_232615715.1-Photobacterium-carnosum:0.1187631034)90.5/98:0.0554219798,WP_130314193.1-Vibrio-vulnificus:0.1092338488)59.6/90:0.0312607542,WP_193154481.1-Vibrio-navarrensis:0.1207817359)88.3/95:0.0335262300,(WP_045031875.1-Photobacterium-phosphoreum:0.0206780138,WP_107213458.1-Photobacterium-phosphoreum:0.0299937277)100/100:0.1017055784)77.1/16:0.0140728955,(WP_031827242.1-Vibrio-parahaemolyticus:0.1572047535,WP_194091044.1-Vibrio-hibernica:0.1461066210)94.6/95:0.0547156562)68.8/21:0.0167702911)90.5/80:0.0266642820)86.3/13:0.0159382570,(((((WP_076542412.1-Shewanella-sp.-UCD-KL21:0.1786195617,WP_007466453.1-Photobacterium-marinum:0.1708475910)88.1/97:0.0721425159,WP_237309237.1-Vibrio:0.1651432922)42.8/94:0.0426325239,WP_133040538.1-Shewanella-fodinae:0.1813074592)63.8/93:0.0378438444,WP_161680902.1-Proteus:0.2130117141)79.2/90:0.0500730026,WP_234235026.1-Shewanella-sp.-AS16:0.1525992810)0.1/11:0.0029217424)83.3/63:0.0160284686,(((((((WP_038175370.1-Vibrio-pacinii:0.1007933633,((WP_020332409.1-Vibrio-fluvialis:0.0477358638,WP_096629284.1-Salinivibrio-sp.-YCSC6:0.1188631989)68.6/96:0.0427925681,WP_171749982.1-Vibrio-sp.-RE88:0.1248892515)81.9/87:0.0206056346)85.6/44:0.0326380674,(((WP_067416136.1-Enterovibrio-coralii:0.0816225713,WP_102038478.1-Shewanella-sp.-GutCb:0.1849040054)61/53:0.0084221342,((WP_197173106.1-Pseudoalteromonas-sp.-NZS11_1:0.1271749280,WP_182748079.1-Pseudoalteromonas-sp.-SR41-1:0.0981692856)92.1/100:0.0546559111,WP_149593442.1-Pseudoalteromonas-sp.-FUC4:0.0985071183)90.1/84:0.0374177883)92.7/59:0.0338994051,(WP_102395415.1-Enterovibrio-norvegicus:0.0881033750,WP_112353352.1-Shewanella-benthica:0.1645723199)84.4/51:0.0330293906)0/4:0.0027122300)82.2/10:0.0241464564,(((((WP_046223474.1-Vibrio-sp.-ECSMB14106:0.0672076286,(WP_102481006.1-Vibrio-breoganii:0.0897597867,WP_258489575.1-Vibrio-alginolyticus:0.1119004720)77.4/91:0.0289935184)57.4/72:0.0384867000,WP_102562373.1-Vibrio-breoganii:0.1112401019)64.3/16:0.0075779452,((WP_261874894.1-Vibrio-rarus:0.1193409359,WP_017031451.1-Vibrio-breoganii:0.1017073563)0/18:0.0000022339,WP_102533425.1-Vibrio-breoganii:0.1131226013)37.1/5:0.0196116474)97.1/75:0.0445814253,((((((WP_102546929.1-Vibrio-breoganii:0.1101552735,WP_102282524.1-Vibrio-cyclitrophicus:0.1129596163)39.2/40:0.0257005976,WP_102461639.1-Vibrio-breoganii:0.0976308295)88.8/58:0.0359813504,(WP_022590465.1-Vibrio-nigripulchritudo:0.1235338457,WP_045977433.1-Vibrio-neptunius:0.1423186725)96.1/99:0.0669311719)0/22:0.0021547454,((WP_102455158.1-Vibrio-breoganii:0.0928709183,((WP_188011373.1-Vibrio-vulnificus:0.1366206613,WP_102451772.1-Vibrio-breoganii:0.0982927448)94.1/100:0.0531513543,WP_166378370.1-Pseudoalteromonas-sp.-Z9A4:0.1402400549)7.2/55:0.0304371423)91.5/98:0.0472334233,(WP_135455667.1-Vibrio-echinoideorum:0.0544492953,WP_218382835.1-Photobacterium-sp.-WH24:0.1532230984)81.9/88:0.0331789966)88.4/28:0.0271019739)93.1/41:0.0257698020,(((WP_261835819.1-Vibrio-ishigakensis:0.0950360242,WP_017221048.1-Moritella-dasanensis:0.0896437419)8/70:0.0436256853,WP_141346804.1-Vibrio-inusitatus:0.1336005792)90.7/55:0.0423231584,(WP_102446429.1-Vibrio-breoganii:0.1231460889,WP_102329601.1-Vibrio-breoganii:0.1444072038)47.6/23:0.0315153158)53/15:0.0136116329)76.2/73:0.0142359911,WP_102941572.1-Vibrio-diazotrophicus:0.1139646887)73.4/82:0.0065762317)66.3/63:0.0238211008,((WP_223418564.1-Shewanella-inventionis:0.0924700352,WP_011497090.1-Shewanella-denitrificans:0.1463982082)93.6/76:0.0491030951,(WP_017063118.1-Vibrio:0.0915737357,WP_102285079.1-Vibrio-lentus:0.0957020852)89.3/99:0.0374966452)60.6/63:0.0225026651)85.3/66:0.0251667819)81.3/90:0.0230405616,(WP_217542775.1-Vibrio-metschnikovii:0.0551626295,WP_113593871.1-Vibrio:0.0558441614)97.8/100:0.0767349099)73.4/6:0.0303205976,(WP_225486226.1-Vibrio-vulnificus:0.1328172373,((WP_068331709.1-Photobacterium-jeanii:0.1245127213,(WP_031414678.1-Vibrio-parahaemolyticus:0.0784433148,WP_248468589.1-Shewanella-sp.-1CM18E:0.1303332994)85.3/100:0.0424967167)89.9/100:0.0475563880,(WP_086585081.1-Vibrio-parahaemolyticus:0.0195142989,WP_217557493.1-Vibrio-metschnikovii:0.0541471972)73.1/89:0.0497472677)29.6/95:0.0239207398)33.2/6:0.0078141876)56.7/11:0.0172876236,((((WP_233983986.1-Vibrio-fluvialis:0.0397284544,((WP_208844179.1-Pseudoalteromonas-xiamenensis:0.0868698755,WP_072616325.1-Vibrio-vulnificus:0.1279820762)13/60:0.0397696221,WP_099031861.1-Pseudoalteromonas-sp.-GCY:0.0893776173)92.3/93:0.0550296480)83/52:0.0272421681,((WP_238819535.1-Vibrio-furnissii:0.0616031510,WP_152138288.1-Plesiomonas-shigelloides:0.1095914317)69.5/91:0.0316168393,WP_233976434.1-Vibrio:0.1190263298)59.6/45:0.0267209453)11/58:0.0128001065,(WP_220051788.1-Shewanella-sp.-FJAT-51860:0.1080001839,WP_029788868.1-Vibrio:0.1294403548)91.9/86:0.0410908867)87.5/40:0.0306159229,((WP_165312521.1-Vibrio-ziniensis:0.0663335675,WP_206974914.1-Vibrio-parahaemolyticus:0.0997779400)91.2/99:0.0402099021,WP_143680610.1-Vibrio-furnissii:0.0887928310)78.2/96:0.0184260608)85.6/39:0.0156226106)88.3/12:0.0222083171,((((WP_193255073.1-Vibrio-navarrensis:0.0405332491,WP_258469755.1-Vibrio-alginolyticus:0.0749496770)94.1/100:0.0427919677,(WP_011716450.1-Shewanella-sp.-ANA-3:0.1754178904,WP_206625521.1-Vibrio-vulnificus:0.0942961203)87.4/95:0.0500781878)74.8/93:0.0394891445,WP_172561824.1-Vibrio-furnissii:0.0753294526)82.2/21:0.0220632261,(WP_158107962.1-Vibrio-furnissii:0.0676425663,(WP_185890410.1-Vibrio-vulnificus:0.1359657241,WP_106651046.1-Shewanella-sp.-Pdp11:0.1140474043)64.9/22:0.0229727188)93.5/26:0.0359296734)79.7/35:0.0126151280)89.2/20:0.0306756200)90.3/41:0.0354165011,((((((((((((WP_102563398.1-Vibrio-sp.-10N.286.49.B3:0.0696897428,(WP_120030433.1-Vibrio-sinensis:0.0729790443,((WP_171342182.1-Vibrio-cyclitrophicus:0.1053701778,WP_161972844.1-Enterovibrio-baiacu:0.0965816034)42.8/93:0.0455926102,(WP_259522409.1-Shewanella-baltica:0.0787519483,WP_173243873.1-Shewanella-sp.-DC2-4:0.1038573850)84.7/100:0.0662640343)85.8/60:0.0374197825)83.5/56:0.0252036561)71.3/43:0.0103787088,WP_235060641.1-Vibrio-sp.-SS-MA-C1-2:0.1607481698)72.1/57:0.0323335070,(((((((WP_041510674.1-Shewanella:0.0384107675,WP_187575439.1-Shewanella-algae:0.0506958843)15.7/32:0.0164930448,(WP_208158397.1-Shewanella-algae:0.0641172292,WP_208157589.1-Shewanella-algae:0.0313937545)95.4/100:0.0420609224)96/100:0.0408014357,WP_102475754.1-Shewanella-sp.-10N.286.52.C2:0.1408827880)57.5/38:0.0074459471,WP_020328057.1-Vibrio-fluvialis:0.0954308049)70.5/98:0.0355067413,(((WP_064489977.1-Vibrio-parahaemolyticus:0.0383947072,WP_201976964.1-Aeromonas-enteropelogenes:0.0917092485)94.3/100:0.0494952347,WP_193250619.1-Vibrio-parahaemolyticus:0.1161170315)83.1/20:0.0189950349,(WP_171138527.1-Vibrio-plantisponsor:0.0514678498,WP_102966424.1-Vibrio:0.0673647243)99.5/100:0.1041091621)48.4/24:0.0225016549)74.2/64:0.0444842370,(WP_039841338.1-Vibrio-harveyi-group:0.0587258651,WP_193167845.1-Vibrio-navarrensis:0.2182117629)82.9/86:0.0545808365)75.9/28:0.0298589139,(((WP_033540599.1-Shewanella:0.1178318831,WP_031815865.1-Vibrio-parahaemolyticus:0.0892280004)82.8/97:0.0325325007,WP_258516089.1-Vibrio-alginolyticus:0.0675628085)90.8/75:0.0421757138,((WP_103194250.1-Vibrio-vulnificus:0.0376090760,WP_158146950.1-Vibrio-fluvialis:0.0935719920)94.2/100:0.0487601636,WP_233927909.1-Vibrio-fluvialis:0.1054978783)94.5/97:0.0480392802)71.4/28:0.0122559030)74.4/44:0.0250631032)0/8:0.0160488143,(WP_069533349.1-Vibrio-parahaemolyticus:0.0887125980,WP_229630939.1-Vibrio-coralliilyticus:0.1745143387)84.5/78:0.0286039096)87.6/42:0.0381554603,((((WP_102986644.1-Vibrio-diazotrophicus:0.0759955536,WP_224657569.1-Vibrio-vulnificus:0.1169161812)19.8/8:0.0105398954,WP_238097025.1-Vibrio-cincinnatiensis:0.1511574614)93.3/95:0.0422192018,((WP_211915530.1-Vibrio-sp.-B1REV9:0.0751883384,((WP_192928291.1-Vibrio-vulnificus:0.0376075546,WP_132698543.1-Vibrio-harveyi-group:0.0772725766)98.6/100:0.0786045572,WP_017420706.1-Vibrio-vulnificus:0.0664768302)77.3/100:0.0214763033)43.7/51:0.0298835432,(WP_182742106.1-Shewanella-sp.-SG44-6:0.2031352234,WP_017024716.1-Vibrio-rumoiensis:0.0775624555)91.8/96:0.0583350749)81.9/94:0.0175467653)75.8/89:0.0198001582,WP_213972370.1-Vibrio-alginolyticus:0.1610738941)22.6/15:0.0175429535)89.7/59:0.0309128904,(WP_014230611.1-Vibrio-sp.-EJY3:0.0689986986,(WP_243025268.1-Vibrio-cincinnatiensis:0.3058038696,(WP_047110243.1-Vibrio-vulnificus:0.1695038978,WP_102313365.1-Vibrio-cyclitrophicus:0.1190450460)40.4/92:0.0491012097)40.9/98:0.0463936761)91.8/7:0.0367480840)89.4/7:0.0248505445,((WP_048661380.1-Vibrio-crassostreae:0.0499056409,WP_193846925.1-Vibrio-sp.-OPT10:0.0758332615)90/100:0.0355879913,WP_238130349.1-Vibrio-cincinnatiensis:0.1167351785)79.1/96:0.0302226943)74.8/5:0.0065540143,(WP_065600047.1-Aliivibrio-sp.-1S128:0.0714518809,WP_130066657.1-Aliivibrio-finisterrensis:0.1143865197)90.9/93:0.0536994425)0/5:0.0012368506,(((WP_135455718.1-Vibrio-echinoideorum:0.0905397074,(WP_161672404.1-Vibrio-toranzoniae:0.0455274957,WP_017040654.1-Vibrio-genomosp.-F10:0.1348229830)93.7/89:0.0432739969)0/40:0.0000029856,((WP_137027437.1-Vibrio-kanaloae:0.0249434715,WP_108117598.1-Vibrio-splendidus:0.0558892460)83.9/100:0.0231858617,WP_137025093.1-Vibrio-kanaloae:0.1052298697)88.9/90:0.0364416374)83.2/92:0.0156854581,((((WP_155682326.1-Aliivibrio-fischeri:0.0723412526,(WP_102385704.1-Vibrio-cyclitrophicus:0.0824185297,WP_239937663.1-Vibrio-chagasii:0.1094986527)87.8/98:0.0380428482)74.1/56:0.0111819986,(WP_108189654.1-Vibrio-splendidus:0.1026502185,(WP_063660043.1-Aliivibrio-fischeri:0.0597303221,WP_243579549.1-Vibrio-splendidus:0.1430107833)80.6/94:0.0335763939)73.7/60:0.0392026984)39.1/56:0.0192898403,(WP_102482947.1-Vibrio-splendidus:0.0672477252,WP_102394203.1-Vibrio-sp.-10N.222.54.B6:0.0955379553)87.4/89:0.0341362682)69.4/53:0.0241084073,WP_076646094.1-Vibrio-sp.-10N.222.47.A9:0.1306876814)30.8/48:0.0140531470)15.1/18:0.0121217850)90.4/7:0.0282507339,(((WP_017020702.1-Aliivibrio-logei:0.1235883844,(WP_005597034.1-Vibrio-scophthalmi:0.1226878874,(WP_108176621.1-Vibrio-splendidus:0.0373789517,WP_239918548.1-Vibrio-sp.-Isolate22:0.0303305906)74.7/100:0.0202312132)74.8/100:0.0262460395)83.6/97:0.0250882766,WP_140342839.1-Vibrio-parahaemolyticus:0.1558610116)69.1/7:0.0146069278,(WP_016784063.1-Vibrio:0.0469888550,WP_005395829.1-Vibrio-diabolicus:0.1882975417)88.8/7:0.0344602808)91.4/7:0.0328934381)96/91:0.0413681169,(((((((((((((WP_268674123.1-Vibrio-scophthalmi:0.0902459666,WP_017085924.1-Vibrio-splendidus:0.0821166792)81.4/99:0.0366668285,(WP_102353859.1-Vibrio-cyclitrophicus:0.0850241708,WP_171315631.1-Vibrio-splendidus:0.0894517845)90.5/99:0.0487064961)58.8/93:0.0389591090,(WP_102298256.1-Vibrio-cyclitrophicus:0.0554837010,WP_135445143.1-Vibrio-tasmaniensis:0.1566193490)0/20:0.0024986567)87.8/52:0.0251080991,((((WP_012551417.1-Aliivibrio-salmonicida:0.0613604560,WP_012276595.1-Shewanella-halifaxensis:0.0530749107)45.7/100:0.0236132645,WP_135385147.1-Vibrio-tasmaniensis:0.1175375389)84.5/62:0.0487377535,WP_102571993.1-Vibrio:0.0676932729)89.5/47:0.0412547139,WP_265133188.1-Vibrio-splendidus:0.0662818090)37.3/12:0.0071274206)75.6/37:0.0198606113,(WP_045069930.1-Photobacterium-leiognathi:0.1185912203,WP_248388778.1-Vibrio-cyclitrophicus:0.0971281621)88/73:0.0345586346)81.9/62:0.0208343165,WP_004733311.1-Vibrio-splendidus:0.0840864257)75.9/72:0.0100407699,(((WP_059021773.1-Vibrio-coralliirubri:0.0374500701,WP_135380937.1-Vibrio-tasmaniensis:0.0791420345)83.3/99:0.0175613200,WP_102276040.1-Vibrio-lentus:0.0826253817)61.5/94:0.0221592249,(WP_019826652.1-Vibrio-splendidus:0.0943629139,WP_182796407.1-Vibrio-sp.-SG41-7:0.1166292966)68.9/81:0.0182275591)88/91:0.0301027581)86.7/76:0.0242572869,((((WP_155682312.1-Aliivibrio-fischeri:0.0565686978,(WP_012532661.1-Aliivibrio-fischeri:0.0944114476,WP_155677763.1-Aliivibrio-fischeri:0.1013439769)16.3/78:0.0301467948)21.7/84:0.0339156101,((WP_108162474.1-Vibrio-splendidus:0.0262387016,WP_004742905.1-Vibrio-tubiashii:0.0464502914)99.6/100:0.0770115718,(WP_016768872.1-Vibrio-cyclitrophicus:0.0870819335,WP_017069285.1-Vibrio-crassostreae:0.0654074083)84.3/76:0.0270794416)91.3/95:0.0481279408)33.6/25:0.0238089908,(WP_130049361.1-Aliivibrio-finisterrensis:0.0575764482,WP_260256886.1-Vibrio-sp.-NC2:0.0801927641)82.3/73:0.0206658064)75.1/44:0.0103578206,((WP_258428294.1-Vibrio-sp.-RM-69-4:0.0712270102,WP_102428030.1-Vibrio-sp.-10N.261.54.E10:0.1196011423)83.1/60:0.0286084737,(WP_193834777.1-Vibrio-sp.-OPT20:0.0386860347,(WP_029865567.1-Vibrio-parahaemolyticus:0.0564892602,WP_213967907.1-Vibrio-alginolyticus:0.0275134732)98.6/100:0.0734671913)90.8/100:0.0319960108)92.9/67:0.0353918932)73.9/77:0.0162494678)88.1/95:0.0344963547,WP_077679840.1-Vibrio-kanaloae:0.0876264187)21/85:0.0095463884,(WP_145570128.1-Vibrio-tasmaniensis:0.0874405732,WP_144217665.1-Shewanella-algae:0.1628916452)94.3/95:0.0627739053)80/82:0.0138872399,((WP_063656223.1-Aliivibrio-fischeri:0.0847843163,WP_155675904.1-Aliivibrio-fischeri:0.0695060867)74.4/88:0.0284145853,WP_017105169.1-Vibrio:0.1058741166)90.9/98:0.0350504677)0/37:0.0046834792,((WP_137008959.1-Vibrio-kanaloae:0.0953571247,WP_135353160.1-Aliivibrio-fischeri:0.0624796476)89.8/99:0.0403353575,WP_248638627.1-Vibrio-crassostreae:0.0906221117)87.9/96:0.0367014143)86.2/80:0.0209149210,(((WP_171343915.1-Vibrio-cyclitrophicus:0.0673951126,(((((((WP_162023628.1-Shewanella-sp.-Arc9-LZ:0.0529338053,WP_248997855.1-Shewanella-marinintestina:0.0475327650)94.3/100:0.0439888018,((WP_009847952.1-Vibrio:0.1303803686,WP_233894267.1-Vibrio-kanaloae:0.0945904289)73.6/100:0.0364352392,(WP_048661352.1-Vibrio-crassostreae:0.0626317109,WP_114634227.1-Vibrio-splendidus:0.0528576316)93.6/100:0.0670857007)94.3/100:0.0564594524)84/99:0.0378714780,WP_017094106.1-Vibrio-splendidus:0.0506043312)87.5/97:0.0214515823,WP_065296243.1-Vibrio-natriegens:0.0979136976)80.4/97:0.0211855179,WP_009843931.1-Vibrio-sp.-AND4:0.0590328628)88.2/97:0.0256893693,((((WP_172851426.1-Vibrio-coralliilyticus:0.1123486959,WP_138522257.1-Pseudoalteromonas-galatheae:0.1857827004)17.6/92:0.0252022531,WP_025523961.1-Vibrio-parahaemolyticus:0.1360678447)69.2/97:0.0321014966,(WP_213886549.1-Vibrio-alginolyticus:0.1600255271,WP_158173598.1-Vibrio-alginolyticus:0.0424304067)55.5/96:0.0329743734)97.4/98:0.0672342898,(((WP_041156219.1-Vibrio-mytili:0.1176758306,(WP_047480300.1-Vibrio-campbellii:0.0497424685,WP_045494469.1-Vibrio-hyugaensis:0.0322106019)98.8/100:0.0916839598)94.9/100:0.0829159266,WP_102523418.1-Vibrio-tapetis:0.1169609692)24.3/99:0.0204572140,WP_069440067.1-Pseudoalteromonas-tetraodonis:0.1916101919)94.1/91:0.0528352289)59.3/88:0.0140270081)86.7/95:0.0360444747,WP_248368760.1-Vibrio-sp.-1CM8B:0.1011008468)28.5/50:0.0306560691)74.8/46:0.0079009092,((((WP_048899071.1-Photobacterium-swingsii:0.1043322597,WP_108186198.1-Vibrio-splendidus:0.1195891600)13.3/65:0.0322414458,(WP_046209502.1-Vibrio:0.0949704327,WP_061039794.1-Vibrio-coralliirubri:0.0958991089)91.9/100:0.0439005383)69.7/45:0.0242757980,(WP_139685731.1-Vibrio-tasmaniensis:0.0890860107,WP_248386338.1-Vibrio-sp.-ED004:0.1377395627)12.6/64:0.0388541507)83.9/58:0.0189618850,(WP_102248986.1-Vibrio-tasmaniensis:0.1342145765,WP_135445135.1-Vibrio-tasmaniensis:0.0769394856)98.8/99:0.0771978791)71.3/26:0.0062270029)92.3/90:0.0301473938,((WP_161753058.1-Vibrio-toranzoniae:0.1023321445,WP_102554090.1-Vibrio-splendidus:0.0808315978)85.6/91:0.0273370814,(((WP_102503160.1-Vibrio-splendidus:0.1061440465,WP_121852788.1-Parashewanella-spongiae:0.1536283196)91.3/95:0.0599210107,(WP_102420589.1-Vibrio-cyclitrophicus:0.0786318542,(WP_102457030.1-Vibrio-cyclitrophicus:0.0820930419,WP_132940416.1-Vibrio-crassostreae:0.0948962409)37.5/75:0.0124400379)88.6/81:0.0326327610)82.4/85:0.0256145616,(WP_102459739.1-Vibrio-lentus:0.0716777080,WP_102363333.1-Vibrio-cyclitrophicus:0.1015066678)81.7/92:0.0319980445)91/89:0.0486527827)82.5/75:0.0154069784)89.2/73:0.0254480292)79.4/26:0.0108359161)46.9/58:0.0115178314,((WP_243584123.1-Vibrio-splendidus:0.1008679545,WP_268809473.1-Vibrio-parahaemolyticus:0.1117112238)77.4/29:0.0144055697,(((WP_102462022.1-Vibrio-splendidus:0.0412200690,WP_187463684.1-Photobacterium-sp.-BZF1:0.0512806976)98.9/100:0.0839093769,(WP_163929979.1-Paraferrimonas-sp.-SM1919:0.0871279059,WP_087845049.1-Photobacterium-malacitanum:0.1065033995)88.3/89:0.0349664874)47.7/37:0.0182517163,(WP_172855536.1-Vibrio-coralliilyticus:0.1692913081,WP_259574191.1-Shewanella-baltica:0.1417787345)91/80:0.0533350950)85.1/30:0.0254452475)89.2/75:0.0253479312)77.5/91:0.0098016987)68.7/53:0.0095974212,((((WP_161129723.1-Photobacterium-lucens:0.0924474678,WP_206634697.1-Vibrio-vulnificus:0.1259643991)88.1/90:0.0314144724,(((WP_188008910.1-Grimontia-hollisae:0.0897080611,WP_065173140.1-Photobacterium-kishitanii:0.1237498925)53.9/94:0.0418957314,WP_058428281.1-Pseudoalteromonas:0.1628893021)31.9/71:0.0183004850,WP_129120576.1-Veronia-nyctiphanis:0.1244928686)63.7/68:0.0137501418)83.7/83:0.0364091561,WP_180567593.1-Oceanospirillales:0.1303151117)21.1/62:0.0094675653,WP_103155729.1-Vibrio-vulnificus:0.0942505079)94.6/86:0.0424702839)84.8/82:0.0121142829,(((WP_249572008.1-Vibrio-cholerae:0.0687676537,(WP_255122387.1-Vibrio-cholerae:0.0749730750,WP_088132129.1-Vibrio-cholerae:0.0884944046)41/98:0.0289782288)87.3/92:0.0403980221,((WP_000864153.1-Vibrio-cholerae:0.0367068470,WP_251878143.1-Grimontia-sp.-020920N:0.1109710606)93.7/96:0.0507094254,WP_045618469.1-Vibrio-vulnificus:0.0970045939)4.3/27:0.0171883712)59.5/62:0.0183160354,(((WP_230432956.1-Plesiomonas-shigelloides:0.0797272206,WP_222763364.1-Vibrio-fluvialis:0.0724316244)96.3/99:0.0587518634,((WP_148519618.1-Vibrio-cholerae:0.0216207368,(WP_167570263.1-Aeromonas-veronii:0.0508582738,WP_000864158.1-Vibrio-metoecus:0.0281329809)94.5/100:0.0259982371)87.2/100:0.0154413261,WP_148520854.1-Vibrio-cholerae:0.0275625269)95/100:0.0502723693)77.8/53:0.0188448549,(WP_148560094.1-Vibrio-mimicus:0.1072937500,WP_084988554.1-Vibrio:0.1155007256)90.8/63:0.0402987013)77.2/43:0.0162393314)86/46:0.0195978093)87/55:0.0162172664,((WP_267741605.1-Vibrio-cholerae:0.0924221969,(WP_172567459.1-Vibrio-fluvialis:0.0699915570,WP_199354891.1-Vibrio-cholerae:0.0377290377)95.1/100:0.0582572207)30.6/82:0.0187479304,WP_102137258.1-Vibrio-cholerae:0.0881370917)94.9/96:0.0529689291)86.3/60:0.0166415082,((WP_017008437.1-Enterovibrio-norvegicus:0.0945220891,WP_261894433.1-Vibrio-porteresiae:0.1530137589)23.6/90:0.0193112313,WP_251878087.1-Grimontia-sp.-020920N:0.0912056375)82.6/67:0.0263097049)55.4/88:0.0152912182,(WP_237041393.1-Photobacterium-damselae:0.1267531309,((WP_038178862.1-Vibrio-rhizosphaerae:0.0881712160,WP_107187026.1-Photobacterium-kishitanii:0.1499911748)11/31:0.0391923246,WP_006646466.1-Photobacterium-sp.-SKA34:0.1293634468)58.8/95:0.0157890800)90.9/83:0.0393655522)65.8/40:0.0155237905,WP_000864154.1-Vibrio-cholerae:0.0569233089)70.9/38:0.0064348893,(WP_080158029.1-Photobacterium-piscicola:0.1412487135,(((((WP_065176337.1-Photobacterium-aquimaris:0.0546159458,WP_107199726.1-Photobacterium-phosphoreum:0.0638213979)90.2/100:0.0409354372,WP_023931546.1-Photobacterium-leiognathi:0.0881646033)95.5/100:0.0519153742,WP_107213932.1-Photobacterium-phosphoreum:0.0661144272)92.9/97:0.0530019177,WP_065173166.1-Photobacterium-kishitanii:0.0929650892)78.7/35:0.0272242479,WP_047876260.1-Photobacterium-aphoticum:0.1094276042)37/36:0.0223087675)91.5/76:0.0399556588)74.8/73:0.0062071352)97.3/90:0.0459000023,(WP_045039948.1-Photobacterium-iliopiscarium:0.1149923455,(WP_050928752.1-Yersinia-enterocolitica:0.1688803000,WP_012883389.1-Dickeya-parazeae:0.1686291928)89.2/100:0.0434905559)78/92:0.0446527314)24.4/53:0.0156114435,(((WP_261298106.1-Shewanella-sp.-SP1S2-7:0.0782659819,WP_167678467.1-Shewanella-aestuarii:0.0746225097)99.7/100:0.1631575549,WP_248994245.1-Shewanella-gaetbuli:0.1709916664)83.9/100:0.1028659287,WP_212599444.1-Shewanella-sp.-M16:0.1453166277)98.9/98:0.1260255007)33.6/21:0.0086911414,((WP_035339894.1-Dickeya:0.0160101840,WP_125258312.1-Dickeya-lacustris:0.0716755414)100/100:0.1395753980,WP_042868619.1-Dickeya:0.0785943959)97.2/100:0.0619208328)99.7/98:0.0904091612)94.9/41:0.0843426482,WP_097364011.1-Edwardsiella-tarda:0.2278858663)38.7/11:0.0198742284)28.9/83:0.0352698129,WP_225633057.1-Aeromonas-enteropelogenes:0.1505069770)87.6/86:0.0310669150,WP_061279316.1-Kluyvera-cryocrescens:0.2690024620)91.7/96:0.0375753545,((((((((((((((WP_236369955.1-Aeromonas-veronii:0.0384621942,WP_113736183.1-Aeromonas-caviae:0.0322800479)63.8/100:0.0121797817,((WP_042019775.1-Aeromonas-sobria:0.0438016634,WP_139495680.1-Aeromonas-veronii:0.0203085432)97.2/100:0.0441102656,(WP_098969895.1-Aeromonas-sp.-CU5:0.0386257010,WP_159163188.1-Aeromonas-veronii:0.0393389899)93.6/100:0.0312700963)91/100:0.0264186665)85.2/82:0.0210558154,((((WP_263173612.1-Shewanella-sp.-SM73:0.1124829830,((WP_263191039.1-Shewanella-sp.-SM87:0.0565962039,WP_263255112.1-Shewanella-sp.-SM32:0.0399184280)81/95:0.0153190493,(WP_263178828.1-Shewanella-sp.-SM101:0.0346198207,WP_109248233.1-Shewanella-algae:0.0594579588)90/100:0.0279600850)88/98:0.0338610510)94.6/100:0.0476828582,(WP_220777726.1-Shewanella-hafniensis:0.0225249830,WP_055647506.1-Shewanella-sp.-Sh95:0.0282923012)91.9/100:0.0367600915)88/100:0.0323199683,(WP_082969903.1-Aeromonas-dhakensis:0.0884091140,WP_104454419.1-Aeromonas-sp.-ASNIH2:0.0449135666)74/62:0.0085160338)62.9/81:0.0309685533,WP_201035193.1-Aeromonas-media:0.0900617605)95.2/98:0.0441017011)97.8/100:0.0507063569,WP_143237783.1-Aeromonas-caviae:0.0394204247)82.4/73:0.0125368138,(((WP_113739100.1-Aeromonas-veronii:0.0430926405,(WP_139459904.1-Aeromonas-veronii:0.0300564000,WP_041208446.1-Aeromonas-jandaei:0.1221881298)99.4/98:0.0729554786)64.1/26:0.0092486325,WP_226032029.1-Aeromonas-hydrophila:0.0289474139)83.2/50:0.0114108898,WP_214003767.1-Aeromonas-caviae:0.0402742168)73.3/68:0.0055019191)87.7/73:0.0165047269,WP_252615985.1-Aeromonas-hydrophila:0.0612846880)74.6/74:0.0064568633,(WP_080988732.1-Aeromonas-caviae:0.0371010769,WP_108536035.1-Aeromonas-sp.-HMWF014:0.0287220616)91.1/100:0.0230442811)94.7/100:0.0455835831,(WP_100646402.1-Aeromonas-veronii:0.0567830348,((WP_252601295.1-Aeromonas-dhakensis:0.0207594394,WP_139432874.1-Aeromonas-dhakensis:0.1148482387)89.6/100:0.0251084970,WP_106553824.1-Aeromonas-hydrophila:0.0499123692)33.1/100:0.0128941763)99.9/100:0.0994137010)88.5/99:0.0308167215,(((WP_268010485.1-Aeromonas-dhakensis:0.0572670294,WP_098985072.1-Aeromonas-sp.-CA23:0.0381022572)99/100:0.1044600841,(WP_181268965.1-Aeromonas-veronii:0.0298315974,(WP_129146711.1-Aeromonas-caviae:0.0216046395,(WP_081843667.1-Aeromonas-taiwanensis:0.0161893722,WP_139744464.1-Aeromonas-media:0.0371883666)83.6/100:0.0197061864)68.2/96:0.0166387903)83.5/96:0.0492880429)89.7/100:0.0661355431,WP_200561028.1-unclassified-Erwinia:0.2323653735)91.1/98:0.0600353603)88.4/98:0.0365974197,(WP_059168599.1-Aeromonas-salmonicida:0.0577918718,(WP_033115905.1-Aeromonas-lacus:0.0369994900,((WP_223920301.1-Aeromonas-caviae:0.0282822745,WP_226058924.1-Shewanella-sp.-SR1:0.0984657785)82.8/100:0.0156834899,(WP_058394234.1-Aeromonas-salmonicida:0.0243774475,WP_257690725.1-Aeromonas-veronii:0.0406306370)80.2/100:0.0176215481)84.9/100:0.0181013019)91.8/99:0.0330822876)87.2/99:0.0279556796)81.5/99:0.0169909420,(((WP_223945983.1-Aeromonas-caviae:0.0888972945,(((WP_265458754.1-Aeromonas-salmonicida:0.0679737734,WP_125597700.1-Aeromonas-salmonicida:0.0208317707)10.4/29:0.0017503650,WP_163128050.1-Aeromonas-allosaccharophila:0.0707136071)0/24:0.0000023833,(WP_180911080.1-Aeromonas-caviae:0.0340211495,WP_264818242.1-Aeromonas-caviae:0.0225344293)92.2/100:0.0171189383)75.3/66:0.0073577744)76.2/27:0.0078838907,(WP_040098646.1-Aeromonas:0.0756316800,((WP_263309349.1-Aeromonas-caviae:0.0340860325,(WP_041217344.1-Aeromonas-hydrophila:0.0342163587,WP_255912544.1-Aeromonas-sp.-JL9:0.0284277629)76.2/87:0.0057413678)79.9/86:0.0086766515,WP_257686696.1-Aeromonas-veronii:0.0336891230)72.6/96:0.0081993213)76.5/99:0.0089142921)74/28:0.0069317774,(((WP_139750984.1-Aeromonas-media:0.0262752975,WP_224430864.1-Aeromonas-rivuli:0.0244823382)82.9/100:0.0256886194,WP_041996113.1-Aeromonas-bivalvium:0.0393944757)89.5/100:0.0269227167,WP_265776268.1-Aeromonas-veronii:0.1015873114)76.4/95:0.0180337282)96/69:0.0418676769)96.1/98:0.0550929895,WP_265468300.1-Aeromonas-salmonicida:0.1030563568)19.5/32:0.0172767127,(((WP_216990617.1-Aeromonas-sp.-FDAARGOS-1409:0.1198903406,((((((WP_148612677.1-Aeromonas-sobria:0.0373405763,(WP_201355655.1-Aeromonas-hydrophila:0.0377796485,WP_201951905.1-Aeromonas-veronii:0.0287344970)42/98:0.0057147484)42.4/52:0.0109428067,WP_171268810.1-Aeromonas-media:0.0204321842)99.9/100:0.0893756263,((((WP_212539243.1-Aeromonas-salmonicida:0.0293778154,((WP_187590672.1-Aeromonas-hydrophila:0.0977107891,WP_017782281.1-Aeromonas-dhakensis:0.0292761210)28.3/100:0.0125117157,WP_060586794.1-Aeromonas-schubertii:0.0430489223)99.8/100:0.0766017472)89.5/100:0.0246149259,WP_201904077.1-Aeromonas-veronii:0.0495702679)76.1/99:0.0171514857,WP_206874356.1-Aeromonas-veronii:0.0085549025)89.6/85:0.0217018630,(WP_042063659.1-Aeromonas-allosaccharophila:0.0310334080,WP_256684552.1-Aeromonas-salmonicida:0.0263379213)69.1/84:0.0117004434)87.1/99:0.0275748202)10.6/57:0.0180034132,((((((((WP_247163170.1-Escherichia-coli:0.0173986535,WP_038644959.1-Pantoea:0.0679805922)99.2/100:0.0739599577,(WP_252422582.1-Aeromonas-taiwanensis:0.0173451473,WP_263719954.1-Aeromonas-caviae:0.0752293937)56.2/80:0.0358117644)90.5/98:0.0327533538,WP_033131297.1-Aeromonas-aquatica:0.0224717018)93.2/84:0.0374143596,WP_241344952.1-Aeromonas-sp.-MR16:0.0126774268)91.5/84:0.0267575021,WP_160838676.1-Aeromonas-caviae:0.0586670051)83.5/100:0.0218543820,(WP_116547128.1-Plesiomonas-shigelloides:0.0720030664,(((WP_139496062.1-Escherichia-coli:0.0565808850,WP_215269322.1-Aeromonas-dhakensis:0.0350462050)44.1/97:0.0055705846,WP_201996661.1-Aeromonas-veronii:0.0330919221)75.4/83:0.0079661672,WP_235463836.1-Aeromonas-australiensis:0.0152179204)78.3/86:0.0195256690)85.1/93:0.0237952621)74.2/94:0.0425325906,WP_167569591.1-Aeromonas-veronii:0.0736657717)81.1/84:0.0232755686,WP_049048056.1-Aeromonas-hydrophila:0.1004216908)98.1/100:0.0701751336)86/38:0.0311759034,WP_042649704.1-Aeromonas-media:0.0817107610)49/48:0.0162982423,(WP_163135231.1-Aeromonas-rivipollensis:0.0219874004,WP_241279715.1-Aeromonas-sp.-MR19:0.0331141067)100/100:0.1608753998)34.9/54:0.0270287381)20/13:0.0130318989,((WP_236563281.1-Aeromonas-veronii:0.1162308370,(((WP_201942221.1-Aeromonas-veronii:0.1071179883,((((WP_268854930.1-Aeromonas-enteropelogenes:0.0932344333,WP_139692403.1-Aeromonas-sobria:0.0234158499)73/31:0.0060497510,(WP_203131090.1-Aeromonas-jandaei:0.0412853473,WP_223380193.1-Aeromonas-enteropelogenes:0.0335317303)85.3/27:0.0114339284)74.3/78:0.0056784418,(WP_042657830.1-Aeromonas:0.0375802180,WP_111901368.1-Aeromonas-veronii:0.0329987310)76.7/87:0.0107569850)78.9/90:0.0106109216,WP_159143398.1-Aeromonas-veronii:0.0168341149)87/97:0.0269441164)78.4/99:0.0403774939,(WP_236321701.1-Aeromonas-veronii:0.0526431018,WP_257119864.1-Aeromonas-hydrophila:0.0250787525)92/100:0.0487959971)15.3/39:0.0157746698,(WP_257707542.1-Aeromonas-caviae:0.0466777375,WP_182951616.1-Aeromonas-sp.-WP2-W18-CRE-05:0.0521698678)98.2/100:0.0688131310)90/92:0.0423703534)68.5/12:0.0253548744,((WP_213320225.1-Aeromonas-caviae:0.0502414279,WP_203764723.1-Aeromonas-caviae:0.0635754034)96.8/100:0.0851441163,WP_069554789.1-Aeromonas-sp.-DNP9:0.1046221696)88.3/43:0.0351156167)84.3/18:0.0162571377)75.7/7:0.0061346119,((WP_064338398.1-Aeromonas-veronii:0.1242055452,(((WP_042864139.1-Aeromonas-caviae:0.0326345474,WP_225159881.1-Aeromonas-caviae:0.0364983534)88.4/92:0.0317656380,(WP_244056308.1-Aeromonas-caviae:0.1106045138,WP_202046159.1-Aeromonas-media:0.0802119061)96.8/91:0.0814310968)71.1/94:0.0343342692,WP_045790994.1-Aeromonas-hydrophila:0.1216276720)81/99:0.0389906706)69.5/91:0.0343712633,WP_253626250.1-Aeromonas-hydrophila:0.1227356960)88.3/83:0.0288995613)87/17:0.0217387707)86.4/70:0.0160916689,((((((WP_234664799.1-Aeromonas-allosaccharophila:0.1148472386,((WP_257692500.1-Aeromonas-caviae:0.1067596754,WP_113719982.1-Aeromonas-salmonicida:0.1016699440)26.1/58:0.0416112662,((WP_227095090.1-Aeromonas-veronii:0.0807292999,WP_152109808.1-Plesiomonas-shigelloides:0.0484132035)86.4/100:0.0363102172,WP_168366848.1-Dickeya-zeae:0.1411377240)87.8/100:0.0350114294)68.3/62:0.0540642839)89.6/88:0.0302552757,WP_268037199.1-Aeromonas-dhakensis:0.0670106030)21.3/17:0.0108414534,((WP_139733908.1-Aeromonas-veronii:0.0139702594,((WP_234676856.1-Aeromonas-rivipollensis:0.0153353376,WP_269382839.1-Aeromonas-hydrophila:0.0611425206)82.2/89:0.0128341028,WP_222134508.1-Vibrio-cholerae:0.1154432060)77.2/74:0.0124545124)58.5/74:0.0232628316,WP_234579942.1-Aeromonas-salmonicida:0.0472152582)92.3/100:0.0504682615)48.5/13:0.0131422903,(WP_268421127.1-Aeromonas-media:0.0478003727,(WP_111910157.1-Aeromonas:0.0186907587,WP_128343177.1-Aeromonas-caviae:0.0502632730)12.8/53:0.0084997616)96.9/100:0.0492013925)90.4/32:0.0259650761,WP_139406182.1-Aeromonas-veronii:0.1996733749)76.7/41:0.0188129430,(((WP_245131329.1-Aeromonas-veronii:0.0970191541,WP_210550341.1-Aeromonas-sp.-SrichE-2G:0.1949588679)73.4/43:0.0244160811,((WP_101347551.1-Aeromonas-sobria:0.0511004598,WP_201955411.1-Aeromonas-veronii:0.0484781037)98.2/100:0.0598853352,(WP_136501435.1-Aeromonas-veronii:0.1176585989,(WP_268029297.1-Aeromonas-hydrophila:0.0861664544,WP_201942265.1-Aeromonas-veronii:0.1028032520)98.4/100:0.0884623052)80.9/98:0.0334521842)80.8/79:0.0203779338)19.9/46:0.0269927238,(((WP_050666161.1-Aeromonas-schubertii:0.0519829650,WP_059168611.1-Aeromonas-salmonicida:0.0859560667)24.1/79:0.0244298997,WP_210600017.1-Aeromonas-sp.-MrichA-1:0.0328997824)90.7/98:0.0278816805,WP_225159845.1-Aeromonas-caviae:0.1084692758)83.8/75:0.0323420005)89.1/73:0.0286313062)87.3/69:0.0265018093)92.6/77:0.0282508338)86.1/52:0.0180850046,((((WP_189493202.1-Formosimonas-limnophila:0.1619353806,WP_133620003.1-Hydromonas-duriensis:0.1732352744)98.9/99:0.1099305458,(WP_213789144.1-Enterobacter-hormaechei:0.0196358588,((WP_148242178.1-Enterobacter-asburiae:0.0443065192,WP_237994110.1-Enterobacter-sp.-DRP3:0.0166223343)74.2/100:0.0051505197,WP_237424876.1-Enterobacter-cloacae-complex-sp.-ECC445:0.0390694649)86.4/100:0.0259957926)99.8/100:0.1610349475)97.4/96:0.0819781009,(((WP_151054911.1-Cellvibrio-sp.-KY-YJ-3:0.1406487150,WP_078044787.1-Cellvibrio-sp.-PSBB023:0.1495071807)36.5/71:0.0164608800,(WP_012489013.1-Cellvibrio-japonicus:0.1162350144,WP_087465811.1-Cellvibrio-sp.-PSBB006:0.1329447438)85/71:0.0329773480)97.6/100:0.0683164623,((WP_048221111.1-Citrobacter:0.1036568908,WP_263807164.1-Salmonella-enterica:0.1264262927)98.9/100:0.1194810939,WP_172410167.1-Escherichia-coli:0.2223243992)82.8/56:0.0329626413)87.6/54:0.0295446007)60.1/31:0.0132992126,((WP_047682696.1-Xenorhabdus:0.0519006463,WP_099124588.1-Xenorhabdus-stockiae:0.0577372595)98.7/100:0.0854788808,(WP_074024273.1-Xenorhabdus-eapokensis:0.0489681444,WP_217470569.1-Photorhabdus-akhurstii:0.0744585357)96.7/100:0.0778653490)98.6/99:0.0810736152)84.6/55:0.0218407110)74.4/57:0.0087607493)93.5/71:0.0403358678,((WP_035600513.1-Edwardsiella-tarda:0.1938940417,WP_125117341.1-Aeromonas-caviae:0.2004158254)0/83:0.0280827218,(WP_202782010.1-Enterobacter-asburiae:0.0717732212,WP_144817260.1-Pseudescherichia-sp.-L3:0.0282555129)100/100:0.1378062809)94.6/100:0.0519780526)74.6/66:0.0096843407,(((((((WP_085037002.1-Cronobacter-sakazakii:0.1562473665,WP_076770125.1-Kosakonia-cowanii:0.0967788362)99/100:0.1257295694,WP_216981205.1-Lelliottia-amnigena:0.1933173143)62.9/100:0.0603950067,(WP_227534785.1-Erwinia-rhapontici:0.0991513493,WP_017800228.1-Erwinia-toletana:0.1257107775)95.5/100:0.0790222443)0/59:0.0233622469,WP_255851625.1-Erwinia-persicina:0.1200633548)98.3/99:0.0826358502,(((WP_167863639.1-Erwinia-rhapontici:0.1046254658,WP_039385357.1-Pantoea-eucrina:0.1459004073)86.6/100:0.0371880766,WP_118663751.1-Erwinia-persicina:0.1155637435)87.1/100:0.0271241595,WP_062742814.1-Erwinia-persicina:0.1395325676)56.3/100:0.0484070053)75.5/89:0.0219729976,((WP_171149131.1-Erwinia-sp.-JH02:0.1073629845,WP_206164314.1-Citrobacter:0.1968761271)85.7/84:0.0427423802,WP_191149682.1-Erwinia-aphidicola:0.0986459285)78.9/84:0.0260472332)81.2/83:0.0181194704,(((((((((((((WP_151711904.1-Enterobacter-ludwigii:0.0253588748,(WP_032941356.1-Citrobacter-freundii:0.1568732348,WP_058678686.1-Enterobacter:0.0649096076)91.3/100:0.0498213915)64.7/100:0.0393459825,WP_223560970.1-unclassified-Enterobacter:0.0250370177)99.8/100:0.1886066755,WP_044720521.1-Escherichia-coli:0.2032759522)98.1/100:0.1457042583,WP_230863267.1-Enterobacter-cloacae:0.2414187984)88.8/100:0.1079725493,((WP_029399168.1-Escherichia-coli:0.1787027884,WP_050862535.1-Enterobacter:0.2294233897)21.5/100:0.0395370200,(WP_034496584.1-Buttiauxella-agrestis:0.2438289290,WP_164716793.1-Cedecea-lapagei:0.2228394932)91.5/100:0.0786508740)82.5/100:0.0537679946)93.7/98:0.0799969511,((WP_108569539.1-unclassified-Pantoea:0.1430681266,WP_028018520.1-Enterobacter-cloacae-complex:0.3416479434)93.2/97:0.0827103322,WP_032659356.1-Enterobacter-hormaechei:0.2254591339)91.1/47:0.0528456105)84.8/38:0.0306847964,(WP_234505032.1-Kluyvera-intermedia:0.2760462317,WP_029881813.1-Enterobacter:0.1663013158)88.5/38:0.0409731707)74.5/6:0.0088705456,((WP_039304623.1-Cedecea-neteri:0.2329308444,(WP_106902182.1-Citrobacter-freundii:0.1551408865,(WP_126291239.1-Enterobacter-hormaechei:0.1146118757,((WP_062932819.1-Enterobacter-hormaechei:0.0296015409,WP_252042312.1-Enterobacter-chengduensis:0.0317946119)66/100:0.0134235632,((WP_049008289.1-Enterobacter-cloacae:0.0520433498,WP_048999124.1-Enterobacter:0.0312816884)34.6/21:0.0063207852,WP_063144508.1-Enterobacter-cloacae:0.0355840547)82.1/21:0.0125627926)88.4/96:0.0480487703)93.9/100:0.0812330759)86.9/95:0.0441480754)47.5/72:0.0454583989,WP_159282060.1-Rahnella-variigena:0.2571001208)85.5/95:0.0440156276)12.7/5:0.0125546707,WP_250341216.1-Pantoea-eucrina:0.1974429523)59.1/52:0.0242997802,(((((WP_257124601.1-Aeromonas-veronii:0.2017662581,WP_043158915.1-Aeromonas-hydrophila:0.2645745272)84.1/100:0.0711246724,WP_038258850.1-Xenorhabdus-bovienii:0.1898098573)86.2/77:0.0412929882,((WP_064374050.1-Pantoea:0.2156251055,(WP_036421240.1-Morganella-morganii:0.1100006081,(WP_264998074.1-Morganella-morganii:0.0582609157,WP_186849509.1-Morganella-morganii:0.0687261473)91.4/100:0.0608999316)92.6/100:0.0656968113)89.2/96:0.0512780986,(WP_038223662.1-Xenorhabdus-bovienii:0.1314526679,(WP_161748440.1-Proteus-sp.-G2658:0.1304284579,(WP_049199273.1-Proteus-mirabilis:0.0603458574,WP_157954980.1-Proteus-terrae:0.0766399588)12.4/47:0.0449278020)98.7/100:0.0905531879)90.5/100:0.0400682940)93/74:0.0431713919)7.7/18:0.0115204031,WP_086113286.1-Xenorhabdus-beddingii:0.1937149776)87.9/59:0.0277185554,(WP_211007190.1-Pectobacterium-brasiliense:0.3013250376,(((((((WP_159241675.1-Proteus-terrae:0.1314700527,WP_195697765.1-Providencia-rettgeri:0.1277945586)92.5/86:0.0575926876,WP_238563424.1-Providencia-rettgeri:0.0878137187)29.9/50:0.0240995900,WP_165880752.1-Providencia-rettgeri:0.1250457124)90.1/96:0.0282851951,WP_231068620.1-Providencia-huaxiensis:0.1082964663)85/89:0.0187250566,(WP_154625743.1-Providencia-stuartii:0.0965466095,WP_004924376.1-Providencia-stuartii:0.1091482742)95.6/86:0.0550037868)8.7/48:0.0140343988,(WP_187128479.1-Providencia-sp.-JUb39:0.1141171298,WP_193115284.1-Escherichia-coli:0.1036662580)95.7/91:0.0588444534)33.8/55:0.0130112964,(WP_169342312.1-Providencia-stuartii:0.1368381909,WP_164565644.1-Providencia-rettgeri:0.1517244249)89.5/82:0.0491434360)85/93:0.0317231870)88.4/80:0.0368559679)93.3/66:0.0405666272)95.7/97:0.0418174145,((((((WP_090123257.1-Kosakonia-arachidis:0.0732810772,WP_020454293.1-Kosakonia-sp.-SOY2:0.0748757957)87.1/100:0.0409485024,WP_017458471.1-Kosakonia-sacchari:0.1213394865)44/100:0.0347109584,(WP_200002836.1-Citrobacter-freundii:0.1272335454,WP_080329661.1-Enterobacter-cloacae-complex:0.1112700693)99.5/100:0.1237844600)7.7/91:0.0590845225,WP_090460354.1-Enterobacter-sp.-kpr-6:0.0798600591)99.4/100:0.1172336676,(WP_070133611.1-Candidatus-Erwinia-dacicola:0.1517655279,WP_140919905.1-Insectihabitans-xujianqingii:0.1406522753)39.6/95:0.0312841495)86.7/93:0.0367412618,(((WP_194857598.1-Citrobacter-freundii:0.2221852465,((WP_141032926.1-Escherichia-coli:0.1115639943,WP_196390204.1-Enterobacter-mori:0.0389660201)31.7/87:0.0295831799,(WP_252032207.1-Enterobacter-asburiae:0.0427122667,WP_111966211.1-Enterobacter-cloacae-complex:0.0256334240)26.6/93:0.0206125360)98.4/96:0.1182188152)99.9/96:0.1466323061,(WP_105274811.1-Escherichia:0.1537968339,(WP_064512195.1-Buttiauxella-gaviniae:0.1615441908,WP_061356323.1-Escherichia-coli:0.3915493562)87/98:0.0632026701)89.9/97:0.0455859872)70.5/91:0.0211475542,(WP_100272167.1-Citrobacter-portucalensis:0.1467676882,WP_269535377.1-Citrobacter-sedlakii:0.1803082698)80.4/95:0.0260978546)69.5/87:0.0348244037)76.5/78:0.0183442439)85.8/83:0.0208033762,(WP_168756429.1-Aeromonas-hydrophila:0.3119060714,((WP_162648791.1-Citrobacter-freundii:0.0776649411,(WP_060683585.1-Citrobacter-portucalensis:0.0961619692,(WP_079798993.1-Salmonella-enterica:0.0779277193,WP_032255798.1-Enterobacteriaceae:0.0666945763)70.2/99:0.0074780049)98.7/99:0.0585435679)89.6/100:0.0459065779,(WP_195994783.1-Citrobacter-portucalensis:0.1523874227,WP_048218074.1-Citrobacter-freundii-complex:0.1092081834)90.4/100:0.0430857740)90.8/100:0.0329196363)90/92:0.0273199323)0/29:0.0000020967,((((((((WP_157959988.1-Citrobacter:0.3190701561,(WP_167816599.1-Enterobacter-cloacae:0.2006779247,(WP_262183194.1-Enterobacter-mori:0.1106424367,(WP_063412371.1-Enterobacter-cloacae:0.0370765472,WP_047027451.1-Enterobacter-kobei:0.0396088140)98.1/100:0.0866357083)96.8/100:0.0812379169)69.7/100:0.0595370302)25.6/33:0.0390703429,WP_086539147.1-Citrobacter-freundii:0.1954028266)12.5/12:0.0181367346,WP_100029735.1-Escherichia-coli:0.0895174292)88.8/34:0.0288581123,(WP_001297344.1-Escherichia-coli:0.1236957838,WP_138098269.1-Jejubacter-calystegiae:0.1137736029)91.6/98:0.0533128759)89.4/41:0.0359155851,((WP_119938542.1-Enterobacter-huaxiensis:0.0792255337,WP_050871166.1-Enterobacter-cloacae-complex:0.0301370624)96.2/100:0.0487115142,(WP_228937168.1-Escherichia-marmotae:0.0276168101,WP_040079227.1-Escherichia-coli:0.0296787181)81.7/100:0.0107745392)87.7/72:0.0286456961)85.7/62:0.0188941973,((((((((WP_127791218.1-Escherichia-coli:0.1246073968,WP_059222105.1-Escherichia-albertii:0.0895415542)0/58:0.0037488930,WP_187209312.1-Escherichia-coli:0.0589875506)82.4/63:0.0197828133,(WP_218070502.1-Escherichia-coli:0.0758322881,(WP_146314246.1-Escherichia-coli:0.0324725072,WP_096967326.1-Escherichia-coli:0.0184294025)96.9/100:0.0612631733)42.6/25:0.0199406646)80.8/78:0.0195194249,(((((WP_034458444.1-Buttiauxella-noackiae:0.1122581152,WP_253383865.1-unidentified-bacterial-endosymbiont:0.1874482430)94.8/98:0.0754233576,WP_096972986.1-Escherichia-coli:0.0575157881)72.9/81:0.0148798061,WP_053290779.1-Escherichia-coli:0.0808421287)84.9/98:0.0323695054,(WP_097421351.1-Escherichia-coli:0.0749854084,(WP_137653986.1-Escherichia-albertii:0.0291682874,WP_001100785.1-Escherichia-coli:0.0283744143)93.1/100:0.0435472856)82/98:0.0297034255)81.8/78:0.0194072571,WP_181465273.1-Escherichia-fergusonii:0.0596067832)82.3/65:0.0117272579)92.2/40:0.0260189993,((((WP_187216555.1-Escherichia-coli:0.1330841230,((((WP_049002539.1-Citrobacter-freundii:0.0857692154,(WP_249576357.1-Enterobacter-vonholyi:0.0457896078,WP_161617217.1-Yokenella-regensburgei:0.0342993294)97.8/100:0.0540020822)70.7/100:0.0234995064,((WP_079834758.1-Salmonella-enterica:0.0333763262,WP_058109100.1-Salmonella-enterica:0.0224408805)1.7/66:0.0249185160,(WP_079793206.1-Salmonella-enterica:0.0787955197,WP_001100807.1-Salmonella:0.0688063737)98.5/100:0.0823720921)97.8/100:0.0628657253)91.5/100:0.0406151499,WP_032303308.1-Escherichia:0.0393002031)95.7/100:0.0393880825,WP_061892435.1-Escherichia-coli:0.1002571283)73.8/89:0.0068182439)86.3/64:0.0177583149,WP_239609625.1-Lelliottia-amnigena:0.0972402331)77.9/56:0.0156120194,WP_107179425.1-Escherichia:0.0601217069)17.4/26:0.0115462974,((WP_109548670.1-Escherichia-coli:0.0429432315,WP_001100790.1-Escherichia:0.0103140888)98.1/100:0.0672365746,WP_181237856.1-Escherichia-marmotae:0.0584834772)88.1/96:0.0211139228)77.1/29:0.0105736684)75.4/39:0.0070704370,((((((WP_193360946.1-Enterobacter-cloacae-complex-sp.-P39RS:0.0445978358,WP_182945676.1-Enterobacter-cloacae:0.0169889197)98.6/100:0.0710211309,WP_265185975.1-Enterobacter-mori:0.0529954134)75.8/96:0.0201140635,((WP_032636062.1-Enterobacter-kobei:0.0522381114,WP_138014015.1-Enterobacter-ludwigii:0.0413202988)58.3/100:0.0312352390,(WP_063137673.1-Enterobacter-cloacae-complex:0.0307832855,WP_165464597.1-Enterobacter-cloacae:0.0434259490)84.4/100:0.0313350134)74.7/96:0.0202623593)83.8/93:0.0238389759,(((WP_096205048.1-Enterobacter:0.0258765265,WP_100165931.1-Enterobacter-mori:0.0271062144)94.9/100:0.0482113699,WP_242172213.1-Enterobacter:0.0665423880)92.1/99:0.0427411074,(((((((WP_042716137.1-Enterobacter-sp.-B509:0.0184114444,WP_241820523.1-Enterobacter-kobei:0.0548027402)88.8/100:0.0483336447,WP_045339085.1-Enterobacter-cloacae-complex:0.1255607303)84.5/95:0.0218215847,WP_063869545.1-Enterobacter-hormaechei:0.0435583067)88/94:0.0252045809,WP_063158410.1-Enterobacter-hormaechei:0.0604128450)80.5/90:0.0165622546,(WP_045338355.1-Enterobacter-hormaechei:0.0913809553,(WP_208283396.1-Enterobacter-sichuanensis:0.0761354329,(WP_182115563.1-Enterobacter-kobei:0.0406004875,WP_213381159.1-Enterobacter-cloacae:0.0435409456)91.3/100:0.0402014282)85.4/100:0.0356564571)83.8/98:0.0308877187)51.3/87:0.0138715428,WP_126315015.1-Enterobacter-hormaechei:0.0920336549)0/58:0.0002417719,WP_033487312.1-Enterobacter-cloacae-complex:0.0610006750)68.7/75:0.0217762133)89.3/90:0.0314439425)90.5/93:0.0254026037,(((((WP_214593796.1-Enterobacter-mori:0.0367375990,WP_086532312.1-Enterobacter-ludwigii:0.0607715968)99.4/100:0.0821765158,((WP_047742849.1-Enterobacter:0.0609503761,WP_040022520.1-Enterobacter-cloacae:0.0647180013)99/100:0.0837812903,(WP_129995405.1-Enterobacter-cloacae:0.0436617200,WP_042099316.1-Escherichia-coli:0.0640491894)9.7/95:0.0161651031)96/100:0.0679737333)94.6/99:0.0467298467,(WP_259047621.1-Klebsiella-sp.-BIGb0407:0.0962053360,(WP_121265736.1-Enterobacter-sp.-R1-2018:0.1708047216,WP_249031739.1-Tenebrionicola-larvae:0.1825467282)82.6/52:0.0251075911)76.8/52:0.0196916141)87.6/53:0.0219530243,((WP_105591414.1-Pantoea-coffeiphila:0.0438068468,WP_123334936.1-Erwinia-sp.-JUb26:0.0350456942)100/100:0.1615910705,WP_015386517.1-Cronobacter-sakazakii:0.1127300041)65.5/38:0.0343620670)91/88:0.0381201986,WP_259000881.1-Scandinavium-goeteborgense:0.1183584278)41.3/31:0.0067894475)90.5/76:0.0226560257,((WP_171269898.1-Aeromonas-media:0.0661349007,WP_216983019.1-Aeromonas-sp.-FDAARGOS-1415:0.0855961474)99.9/100:0.1185884068,WP_075180894.1-Pantoea-sp.-1.19:0.1128081497)98.1/96:0.0724295417)74.7/55:0.0066906292)96/52:0.0324463089,(((((((((WP_115348269.1-Hafnia-alvei:0.1194126330,WP_046458830.1-Obesumbacterium-proteus:0.1905098365)3.5/94:0.0343352247,WP_061059890.1-Hafnia-paralvei:0.1391532771)78.8/96:0.0298333703,(WP_130961887.1-Hafnia-paralvei:0.1050455622,(WP_175320495.1-Hafnia-paralvei:0.0537644979,WP_130996599.1-Hafnia-paralvei:0.1373225242)97.9/100:0.0801910033)82.1/97:0.0401105247)92.4/99:0.0503145315,(WP_120162698.1-Rahnella-variigena:0.0480071512,WP_183847446.1-Rahnella-inusitata:0.0455320372)100/100:0.1869258041)91.3/91:0.0404448529,(((WP_058662478.1-Enterobacteriaceae:0.0420980492,WP_061709645.1-Pseudenterobacter-timonensis:0.0299213087)99.8/100:0.1309933768,WP_112164393.1-Rahnella-inusitata:0.1156088471)93.5/100:0.0628168693,WP_157953065.1-Limnobaculum-parvum:0.1171879631)93.9/96:0.0416138216)79.9/74:0.0151577046,WP_047609274.1-Rahnella-aquatilis:0.1244777441)85.9/68:0.0337412713,(WP_181624464.1-Citrobacter-freundii:0.1506600057,WP_001515521.1-Escherichia-coli:0.1308265597)93.3/99:0.0509391389)89.1/93:0.0556149384,(WP_174633339.1-Escherichia-coli:0.0743562807,WP_072664248.1-Escherichia:0.0685044003)94.6/99:0.0545013297)82.3/63:0.0187121427,(((((WP_170897747.1-Escherichia-coli:0.1218814584,WP_001542972.1-Escherichia-coli:0.0940559906)38/92:0.0363626564,WP_001100781.1-Escherichia-coli:0.1313217201)78.1/60:0.0203787086,(WP_042093344.1-Escherichia-coli:0.1328735530,WP_089706542.1-Escherichia-coli:0.0491501434)91.8/60:0.0403862920)75.2/71:0.0085766219,WP_262938691.1-Escherichia-albertii:0.0797159870)95/97:0.0297388685,(((((((((WP_232904305.1-Buttiauxella-sp.-W03-F01:0.1305748176,WP_259388919.1-Enterobacter-huaxiensis:0.0409212967)63.6/74:0.0292408920,WP_230350752.1-Lelliottia-sp.-WAP21:0.0939571227)93.5/100:0.0397481438,((((WP_141109706.1-Escherichia-albertii:0.0715585051,WP_001572827.1-Escherichia-coli:0.0495442120)9.8/71:0.0239022580,WP_203360733.1-Citrobacter-sp.-R56:0.0473691629)89.4/100:0.0381105805,WP_258993080.1-Scandinavium-hiltneri:0.0590648874)84.2/100:0.0282222054,WP_046888360.1-Enterobacter-cloacae:0.0765303174)79.3/74:0.0197966368)24.4/85:0.0111109909,WP_034915863.1-Erwinia-sp.-9145:0.1285883260)66.7/85:0.0212276639,(WP_172688853.1-Enterobacterales:0.0947618374,WP_126546226.1-Enterobacter-quasimori:0.0571511547)87.9/91:0.0403437456)68.6/97:0.0347904998,WP_105963664.1-Escherichia-coli:0.0701615305)0/57:0.0000028523,((WP_025210617.1-Escherichia-coli:0.0783798043,WP_085454021.1-Escherichia-coli:0.0885224563)10.8/4:0.0292776556,WP_001527244.1-Escherichia-coli:0.0607666594)83.9/32:0.0224220767)28/10:0.0121133104,(((WP_063610377.1-Escherichia-coli:0.0607364990,WP_001100798.1-Enterobacteriaceae:0.0834177121)84.2/42:0.0237357647,(WP_088541377.1-Escherichia-coli:0.0223345984,WP_222691870.1-Escherichia-marmotae:0.0491614338)93.7/100:0.0348531086)85.9/25:0.0224664071,(WP_157704420.1-Escherichia-coli:0.0785439646,WP_042200020.1-Enterobacteriaceae:0.0327786478)92.7/100:0.0423311874)77.5/13:0.0153745867)85.9/81:0.0157487531,((WP_095374511.1-Escherichia-coli:0.0131237711,WP_012906377.1-Citrobacter-rodentium:0.0708986295)98.9/100:0.0625783425,(WP_096122580.1-Escherichia-coli:0.0536136609,WP_162379936.1-Citrobacter-sp.-NCU1:0.1347749559)78/99:0.0139006022)92.5/99:0.0268430314)50.4/6:0.0093045371)90.4/28:0.0343149834)28.7/5:0.0034479178)76.5/85:0.0109045478,(((WP_208464084.1-Escherichia-coli:0.0959030894,(WP_262931739.1-Escherichia-albertii:0.0297381882,WP_160188772.1-Escherichia-coli:0.0378617013)97.9/100:0.0665131011)69.5/90:0.0140498186,(WP_021540003.1-Escherichia-coli:0.0163795355,WP_047657564.1-Enterobacteriaceae:0.0348951503)99.5/100:0.0711627375)90.1/34:0.0293543866,(WP_021577651.1-Escherichia-coli:0.0979431993,(WP_220393175.1-Citrobacter-freundii:0.1973228139,WP_219093375.1-Escherichia-coli:0.1110189831)85.3/89:0.0417488462)74.1/30:0.0124115668)85.4/87:0.0176905552)97.1/96:0.0412890077)89.3/40:0.0177341338,(((((WP_202667543.1-Lelliottia-amnigena:0.1335467252,WP_153879942.1-Citrobacter-freundii:0.1070007692)49.8/85:0.0259388419,((WP_103946175.1-Lelliottia-aquatilis:0.0804673492,WP_235147774.1-Enterobacter-cloacae:0.1970918327)72/84:0.0081562672,WP_123427447.1-Lelliottia:0.0493580599)98.3/100:0.0675224226)88.8/86:0.0388258024,(WP_137650031.1-Escherichia-albertii:0.1060241463,WP_001100786.1-Escherichia-coli:0.0968956877)0/18:0.0035416823)73.9/56:0.0258829475,(((WP_095844187.1-Escherichia-coli:0.0459351069,(WP_086118650.1-Escherichia-coli:0.0226977267,WP_089538038.1-Escherichia-coli:0.0282166339)81.1/100:0.0113652083)90.7/100:0.0558606638,WP_179154141.1-Citrobacter-freundii:0.2098631523)95.8/98:0.0760636510,((((WP_001100804.1-Enterobacteriaceae:0.1098589497,((WP_001610415.1-Escherichia-coli:0.0337416068,WP_138183825.1-Escherichia-sp.-E4385:0.0174660252)99.6/100:0.1103269095,(WP_089585421.1-Escherichia-coli:0.0314989899,WP_104705897.1-Escherichia-coli:0.0305728418)99.5/100:0.1114667910)77.4/98:0.0425351228)94.3/83:0.0490860434,((WP_107192407.1-Escherichia-albertii:0.0136086080,WP_181673375.1-Escherichia-fergusonii:0.0391550281)99.2/100:0.0957419980,((WP_032214194.1-Escherichia-coli:0.0399281868,(WP_059217428.1-Escherichia-albertii:0.0240640620,(WP_073278747.1-Escherichia-coli:0.0643361099,WP_087893724.1-Enterobacteriaceae:0.0313865647)99.9/100:0.0991397709)74.7/88:0.0148207915)79.2/88:0.0188772640,WP_085459683.1-Escherichia-coli:0.0630491347)99.4/100:0.0871411683)45.5/80:0.0258246870)92/47:0.0390799908,WP_267387874.1-Citrobacter-portucalensis:0.1829179655)78.5/42:0.0122910145,(((WP_016236761.1-Enterobacteriaceae:0.0758221895,WP_024239504.1-Escherichia-coli:0.0881483613)95.7/100:0.0626606142,WP_033870520.1-Enterobacteriaceae:0.1451266327)87.3/77:0.0326629545,(WP_248249449.1-Enterobacter-roggenkampii:0.1317302489,WP_032936727.1-Citrobacter-freundii:0.0697319886)99.7/99:0.1095752719)90.7/73:0.0310187537)75.8/40:0.0192393791)91.3/88:0.0455194562)79.4/88:0.0168532701,((WP_181680936.1-Citrobacter-sp.-RHB25-C09:0.0739735485,WP_103775088.1-Citrobacter-amalonaticus:0.1653016158)41/69:0.0226045746,(WP_090049757.1-Citrobacter-amalonaticus:0.0299400651,WP_196013128.1-Citrobacter-farmeri:0.0212164244)99.5/100:0.0851018526)100/100:0.1232664930)84/47:0.0202170984)40.9/65:0.0048477826,WP_073446180.1-Escherichia-coli:0.1867985028)86.5/92:0.0150951936)93/93:0.0374119839)93.4/80:0.0395103903)14.2/47:0.0180235094,(((WP_097405671.1-Kosakonia:0.0484507360,WP_161518249.1-Kosakonia-sp.-MH5:0.0451251909)73/100:0.0468271553,WP_128482629.1-Kosakonia-cowanii:0.1001211589)99.9/100:0.1296924795,((((WP_238085128.1-Pseudescherichia-vulneris:0.0186104738,WP_032652659.1-Enterobacter-cloacae-complex:0.0803718402)38.5/87:0.0112092119,WP_129363899.1-Enterobacter-cloacae:0.0260857501)63.6/97:0.0159116445,(WP_257661844.1-Citrobacter-portucalensis:0.0954810666,WP_193134185.1-Citrobacter-freundii:0.0668961414)87.4/100:0.0252943548)95.7/100:0.0642322915,(WP_042284276.1-Citrobacter-sedlakii:0.1585086298,(WP_024193259.1-Escherichia-coli:0.0830213839,WP_099433281.1-Citrobacter-amalonaticus:0.0310747091)90.8/100:0.0617861533)96.8/100:0.0769624071)94.2/100:0.0720878445)96/84:0.0599941103)96.6/99:0.0695484521,(WP_241647798.1-Rosenbergiella-sp.-JB07:0.2115114393,WP_181599127.1-Citrobacter-sp.-RHB36-C18:0.1562844654)0/20:0.0126035948)21.9/28:0.0343042355,WP_065370581.1-Kosakonia-sacchari:0.1292667931)95.9/99:0.0489505651)64.1/89:0.0125910139,((((((((((((((WP_182931197.1-Enterobacter-cloacae:0.1134485223,WP_211776196.1-Klebsiella-pneumoniae:0.0522341858)71.3/96:0.0110355471,WP_047723323.1-Klebsiella-michiganensis:0.0384952260)75.1/84:0.0078454459,((WP_139968923.1-Klebsiella-pneumoniae:0.0758286380,WP_065799756.1-Klebsiella-pneumoniae:0.0054672253)0/96:0.0000025584,WP_095100910.1-Raoultella-sp.-10-1:0.0613288583)92.4/99:0.0329750513)76.6/67:0.0055761898,WP_104011323.1-Klebsiella-michiganensis:0.0390931630)76/83:0.0101637802,(((WP_048284323.1-Pluralibacter-gergoviae:0.1007599417,WP_221694709.1-Klebsiella-aerogenes:0.0558076083)44.1/92:0.0142135292,WP_213760001.1-Klebsiella-aerogenes:0.0624758535)74.7/40:0.0068259208,(WP_223370324.1-Klebsiella-pneumoniae:0.0405289678,WP_234579544.1-Raoultella-terrigena:0.0261630097)44.7/16:0.0118932555)48.4/7:0.0088599556)26.5/68:0.0144274908,WP_149270444.1-Klebsiella-aerogenes:0.0273387237)86.3/92:0.0194579321,WP_046852082.1-Raoultella-planticola:0.0330821774)0.9/11:0.0106492366,WP_128320364.1-Raoultella-planticola:0.0550975689)0/6:0.0021031497,(WP_114692711.1-Klebsiella-pneumoniae:0.0334746112,WP_123755035.1-Raoultella-sp.-BIGb0399:0.0332620337)74.1/14:0.0055698431)77/12:0.0056755543,WP_236948387.1-Klebsiella-sp.-2019SCSN059:0.0324695506)85/12:0.0108414348,WP_259891680.1-Klebsiella-variicola:0.0170840043)76/67:0.0056117554,((WP_234537246.1-Raoultella-ornithinolytica:0.0451076833,WP_113990968.1-Raoultella:0.0112205476)94/99:0.0330336050,WP_186601692.1-Klebsiella-quasipneumoniae:0.0414535136)74.9/30:0.0084394307)97.9/96:0.0618497653,((WP_095100919.1-Raoultella-sp.-10-1:0.0406454692,WP_212709529.1-Tatumella-sp.-JGM130:0.0168822498)98.8/100:0.0555407759,WP_236948353.1-Klebsiella-sp.-2019SCSN059:0.0599817303)66/96:0.0161131936)94/99:0.0410899035,WP_123822891.1-Klebsiella:0.0292597006)98.5/100:0.0608002458)80.6/97:0.0365687961,WP_021241446.1-Enterobacter-roggenkampii:0.1355410131)24.4/34:0.0302305643,WP_072663757.1-Enterobacteriaceae:0.0622838477)98.6/100:0.0763638025)95/100:0.0508287050,XP_011503919.1-Ceratosolen-solmsi-marchali:0.0721402728);

**Nematode RML-3, with bacterial and eukaryotic homologs**

**Shown in Figure S2**

ML tree method: IQ-TREE

Sequences: 2787

Substitution model: WAG+I+G4

(WP_063501760.1-Escherichia-coli:0.0390233249,(WP_044173167.1-Phytobacter-massiliensis:0.0437034213,((((((WP_203451401.1-Lelliottia-sp.-RWM.1:0.0621849622,WP_185654835.1-Citrobacter-braakii:0.1166770134)89.3/100:0.0710266097,WP_183170804.1-Atlantibacter-sp.-RC6:0.2228878737)96.6/98:0.1050177903,((WP_160868038.1-Pantoea-sp.-Taur:0.3118646958,WP_008499438.1-Enterobacter-cloacae-complex:0.1947661791)87.2/96:0.0556326825,(WP_090085547.1-Escherichia-coli:0.0445848661,(WP_103283990.1-Citrobacter:0.1731460393,WP_234083964.1-Enterobacter-quasiroggenkampii:0.1274709311)47.5/100:0.0277154292)100/100:0.1708982142)0/62:0.0035146479)83.4/98:0.0390532646,(((((((((((((((((WP_249615874.1-Craterilacuibacter-sp.-RT1T:0.4569254426,(WP_163316497.1-Crenobacter-sedimenti:0.1499783763,WP_160797365.1-Craterilacuibacter-sinensis:0.1339620926)97.5/100:0.1889488098)55.4/100:0.0215847198,WP_014088374.1-Pseudogulbenkiania-sp.-NH8B:0.1986163401)87.1/100:0.0711450284,WP_008952446.1-Pseudogulbenkiania-ferrooxidans:0.2609199498)83.3/100:0.0514322457,((WP_174873379.1-Vogesella-oryzae:0.1581570198,WP_147289921.1-Crenobacter-cavernae:0.1420312216)0/33:0.0182771499,WP_137008594.1-Aquitalea-aquatilis:0.3743729333)30.8/76:0.0407720856)77/89:0.0362871309,(((((((((((((((WP_004123988.1-Rhizobium-freirei:0.1011661088,WP_095433322.1-Rhizobium-sp.-11515TR:0.0379845816)75.5/97:0.0164163950,WP_015341698.1-Rhizobium:0.0434310281)74.9/13:0.0071817267,WP_106999541.1-Rhizobium-sp.-JAB6:0.0493251945)0/13:0.0016763437,WP_183712806.1-unclassified-Rhizobium:0.0457779776)90.6/100:0.0201445650,((WP_183765619.1-unclassified-Rhizobium:0.0346418346,WP_120667780.1-unclassified-Rhizobium:0.0599774958)61.7/100:0.0299311883,WP_037195771.1-Rhizobium-lusitanum:0.0331181503)84.4/99:0.0158521042)0/10:0.0000020615,WP_184706966.1-Rhizobium-lusitanum:0.0998167831)40.3/71:0.0075687597,WP_092719658.1-Rhizobium-multihospitium:0.0344738556)64.2/77:0.0125211357,WP_174074493.1-Agrobacterium-rhizogenes:0.1010352919)86/97:0.0230985747,WP_168880117.1-Rhizobium-sp.-P28RR-XV:0.0425151218)74.7/97:0.0092270235,WP_112341687.1-Rhizobium-tropici:0.0392636474)88/97:0.0171421414,WP_199146672.1-Rhizobium-sp.-ASV8:0.1900130942)38/100:0.0054509130,WP_145321829.1-unclassified-Rhizobium:0.0947959114)99.5/100:0.1491948295,(WP_043579071.1-Chromobacterium:0.0618669582,WP_071113424.1-Chromobacterium-sphagni:0.1010904446)99.8/100:0.1589893649)95.6/100:0.0881976085,((((((((((((((((((((((WP_250059255.1-Halomonas-sp.-ATCH28:0.3091328701,WP_179153381.1-Chromohalobacter-salexigens:0.2205217865)87.7/100:0.0801727306,WP_250939176.1-unclassified-Larsenimonas:0.2568673532)83/99:0.0654558866,WP_106478747.1-Phytohalomonas-tamaricis:0.2403204490)89.7/93:0.0813119297,(((WP_139871681.1-Candidatus-Methylopumilus-universalis:0.2132057156,(WP_139929800.1-Candidatus-Methylopumilus-universalis:0.2797369442,WP_140005423.1-Candidatus-Methylopumilus-planktonicus:0.0721611148)89.2/100:0.1011568064)99.3/100:0.2487882185,WP_058575749.1-unclassified-Guyparkeria:0.2036982394)87.3/96:0.1214757707,(WP_218634960.1-Marinobacter-arenosus:0.1321611709,WP_240716217.1-Halomonas-neptunia:0.0897628463)95.5/100:0.1394255057)0/19:0.0120006837)91.2/15:0.0963878676,(((((WP_269022943.1-Methylomonas-sp.-MP1:0.0967263434,(WP_196437371.1-Methylomonas-sp.-LL1:0.1249117624,(WP_150047608.1-Methylomonas-rhizoryzae:0.3011164552,WP_066983474.1-Methylomonas-lenta:0.0824446996)86.4/98:0.0579818747)85.3/97:0.0624979210)87.9/94:0.0524886110,(WP_140911374.1-Methylomonas-koyamae:0.1983954593,((WP_071159776.1-Methylomonas-sp.-LWB:0.1656758340,(WP_020485671.1-Methylomonas-sp.-MK1:0.1746613830,((WP_256607223.1-Methylomonas-sp.-WSC-7:0.0731829927,WP_205450646.1-Methylomonas-sp.-EFPC1:0.0794364354)11.3/100:0.0057786134,WP_064036612.1-Methylomonas-methanica:0.0000023732)91.8/100:0.0703944902)92.4/100:0.0674639924)0/13:0.0000026635,WP_036273427.1-Methylomonas:0.1110952302)85.8/100:0.0510405497)76.8/99:0.0526568964)88.2/94:0.0547703877,(WP_148817865.1-Pigmentiphaga-aceris:0.3513656627,WP_064030819.1-Methylomonas-koyamae:0.1587151104)30.3/49:0.0512021543)95.3/94:0.1064387448,(WP_019964198.1-Pannonibacter-phragmitetus:0.5906854103,WP_138632053.1-Pseudoalteromonas-spongiae:0.2474106665)83.6/38:0.0717531096)83.1/18:0.0438225396,(WP_138213821.1-Pseudomonas-nitroreducens:0.0335076808,WP_017517140.1-Pseudomonas-nitroreducens:0.0277493605)100/100:0.2815083292)76.9/13:0.0253889156)93.5/81:0.0625092370,(((WP_177138191.1-Hydrogenophaga-aromaticivorans:0.1214036894,WP_133157403.1-Hydrogenophaga-pseudoflava:0.1728558381)75.7/99:0.0159461693,(WP_236964528.1-Hydrogenophaga-sp.-SL48:0.0844908085,(WP_226493830.1-Hydrogenophaga-taeniospiralis:0.0427950936,WP_210824367.1-Hydrogenophaga-aromaticivorans:0.0660313853)62.6/100:0.0418769148)98.7/100:0.0909424309)91.9/99:0.0725059131,(WP_158658301.1-Achromobacter-sp.-AONIH1:0.4065522136,WP_085721622.1-Pseudomonas:0.1124982929)77.7/72:0.0518485032)89.6/71:0.0563382823)0/21:0.0000023049,((((WP_136867012.1-Guyparkeria-sp.-SB14A:0.3965744529,WP_102075761.1-Pollutimonas-subterranea:0.1365999825)59.6/53:0.0449536662,WP_102071501.1-Pollutimonas-nitritireducens:0.2775822409)81.7/52:0.0514880022,(WP_066098770.1-Halothiobacillus-sp.-LS2:0.1474630854,WP_070073009.1-Acidihalobacter-aeolianus:0.2246298097)94.2/99:0.1133799330)79.3/80:0.0261800719,((((WP_230502293.1-Janthinobacterium-sp.-FW305-128:0.4474502161,(((((((WP_220354719.1-Rhizobium-cauense:0.2452591201,WP_016557527.1-Rhizobium-grahamii:0.1338907203)87.5/65:0.0508483762,((WP_184450570.1-Rhizobium-sp.-BK538:0.2079704188,(WP_072373837.1-Rhizobium-tibeticum:0.0988148864,WP_007537558.1-Rhizobium-mesoamericanum:0.0596164243)88.3/100:0.0514829411)60.3/99:0.0230025229,WP_018896487.1-Rhizobium-sp.-2MFCol3.1:0.2202022888)80.8/99:0.0191363023)78.5/66:0.0337458082,WP_233136718.1-Agrobacterium-vaccinii:0.2849800028)72.7/100:0.0412199215,WP_153273632.1-Rhizobium-grahamii:0.2121502055)98/100:0.1372158724,WP_068803504.1-Immundisolibacter-cernigliae:0.1756694160)83.7/98:0.0639036063,WP_074973890.1-Pseudomonas-otitidis:0.1821328002)76.7/91:0.0300582243,((WP_180152801.1-Parapusillimonas-granuli:0.1499400067,(WP_088146851.1-Achromobacter-denitrificans:0.1079828988,WP_105555808.1-Achromobacter-sp.-MYb9:0.0866950202)95.9/100:0.0922699497)67.5/94:0.0482662419,(WP_011111299.1-Nitrosomonas-europaea:0.1651812218,WP_046857648.1-Pseudomonas-sp.-CCOS-191:0.1971283394)49.4/88:0.0305754416)83.1/44:0.0405926870)71.7/28:0.0175917213)72.2/11:0.0234538236,(WP_019141909.1-Noviherbaspirillum-massiliense:0.1465910493,WP_146330669.1-Noviherbaspirillum-sp.-UKPF54:0.2513456428)89.4/98:0.0567071732)83.7/30:0.0399050867,WP_047550623.1-Methylotenera-sp.-G11:0.2440835478)60.9/16:0.0176874204,WP_217963608.1-Candidimonas-humi:0.2829755553)53.6/9:0.0097613344)94.7/88:0.0441528585)23.2/6:0.0223224260,((((WP_044615503.1-Gynuella-sunshinyii:0.4000163998,WP_092579398.1-Achromobacter-sp.-NFACC18-2:0.4117376936)57.7/96:0.0781954833,WP_227683402.1-Pusillimonas-sp.-MFBS29:0.2045442182)89.6/96:0.0829961842,(((((WP_006216420.1-Achromobacter-piechaudii:0.1206947153,WP_219991400.1-Achromobacter-sp.-ES-001:0.0857683088)91.4/100:0.0513628285,WP_183017169.1-Achromobacter-sp.-UMC46:0.0497994195)87.6/99:0.0380502191,WP_100503206.1-Achromobacter-ruhlandii:0.1456251877)76.7/99:0.0338919902,WP_175192416.1-Achromobacter-deleyi:0.0546864034)100/100:0.2542018995,WP_065086507.1-Pseudomonas:0.2155237809)85.1/89:0.0377148934)74.5/70:0.0283532746,(WP_076518421.1-Achromobacter-sp.-MFA1-R4:0.0871238381,(WP_104142565.1-Achromobacter-spanius:0.0357898318,WP_120157261.1-Achromobacter-sp.-B7:0.0414664944)87.1/96:0.0361083035)99.5/95:0.1242126818)62.3/51:0.0195061379)89.7/42:0.0371621502,((WP_209872004.1-Pseudomonas-sp.-BP8:0.1869776846,WP_094844257.1-Bordetella-genomosp.-11:0.2646147767)2.2/58:0.0423624391,WP_066093302.1-Hydrogenophaga-crassostreae:0.1966862554)77.9/67:0.0263117430)65.5/30:0.0153205745,((WP_125865093.1-Stutzerimonas-stutzeri:0.1881119968,WP_200290630.1-Stutzerimonas-balearica:0.1113143466)77.4/97:0.0649213996,(WP_101152997.1-Pseudomonas-sp.-Choline-3u-10:0.1114682579,((WP_046617631.1-Stutzerimonas-stutzeri:0.0751568843,WP_135265705.1-Stutzerimonas-stutzeri:0.0454484363)92.4/100:0.0838404111,WP_003299494.1-Stutzerimonas-stutzeri:0.0866600620)86.3/100:0.0399616918)91.5/100:0.0597617025)85.8/86:0.0388952724)72.3/77:0.0273470857,(((WP_042945119.1-Pseudomonas-fluorescens:0.0738062219,WP_181643258.1-Pseudomonas-sivasensis:0.0494894504)97.7/100:0.0999436659,WP_205489922.1-Pseudomonas-lactucae:0.1885874538)58.7/89:0.0287121467,((WP_169965430.1-unclassified-Pseudomonas:0.0403625811,WP_047278503.1-Pseudomonas-lundensis:0.0332545567)99/100:0.1156344186,WP_104727848.1-Pseudomonas-oleovorans:0.1698121353)73.6/86:0.0329312475)88.6/96:0.0418309071)84.7/81:0.0211045590,WP_214501190.1-Pseudomonas-sp.-RC3H12:0.2805029630)76.7/75:0.0087885960,((((WP_191471912.1-Pseudomonas-asiatica:0.2023443891,(WP_043318452.1-Pseudomonas-sp.-PI1:0.0435458746,WP_043268312.1-Pseudomonas:0.0355288382)62.2/100:0.0238645872)94.6/100:0.0662815971,(WP_089390899.1-Pseudomonas-delhiensis:0.1182654678,WP_076426141.1-Pseudomonas-alcaligenes:0.1553282263)59/99:0.0208912126)96.4/100:0.0745445338,WP_233297479.1-Pseudomonas:0.1671339066)61.8/99:0.0193438109,(WP_184594840.1-Pseudomonas-nitritireducens:0.1825703540,WP_192381882.1-Pseudomonas-sp.-PDM20:0.1135015783)98.2/100:0.1509478638)85.2/96:0.0327584437)71.3/62:0.0151220477,((WP_077733980.1-Paenalcaligenes-hominis:0.0508550875,WP_167661700.1-Paenalcaligenes-hominis:0.0258573944)100/100:0.2413483799,WP_020564242.1-Methylosarcina-fibrata:0.2317240182)35.7/68:0.0300646696)76/72:0.0207422428,((((WP_029241333.1-Pseudomonas:0.0690082166,((WP_054996663.1-Pseudomonas-ficuserectae:0.0428777535,(WP_007252556.1-Pseudomonas-syringae-group:0.0479677390,WP_025390602.1-Pseudomonas-syringae:0.0253552156)23.5/99:0.0203883148)93.1/100:0.0463030597,WP_125830676.1-Pseudomonas-sp.-v388:0.0769498439)88.3/100:0.0446934434)89.3/100:0.0650536626,((WP_016973326.1-Pseudomonas-tolaasii:0.1165319960,WP_099548838.1-Pseudomonas-sp.-2995-3:0.1068805161)79.1/100:0.0362830633,(WP_177015116.1-Pseudomonas-costantinii:0.0557623915,WP_141063556.1-Pseudomonas-fluorescens:0.0970634827)93.2/100:0.0616460978)95.3/100:0.0911499099)95.3/100:0.1027864438,(WP_033955816.1-Pseudomonas-aeruginosa:0.0285053869,WP_003153917.1-Pseudomonas-aeruginosa-group:0.0380786261)99.9/100:0.2193999940)93.1/97:0.0840547505,((WP_058488093.1-Pseudomonas:0.1595485446,WP_043256200.1-Pseudomonas-knackmussii:0.1255657117)30.6/80:0.0309367794,((WP_088420173.1-Pseudomonas-nitroreducens:0.1272273438,WP_254475170.1-unclassified-Pseudomonas:0.0718262543)74.3/100:0.0294885301,WP_205338011.1-Pseudomonas-sp.-PDNC002:0.0665731194)100/100:0.1540931855)81.8/94:0.0422998567)26.9/58:0.0135692682)69.1/51:0.0111829058,(((((WP_225237063.1-Massilia-oculi:0.2908603382,WP_147937383.1-Massilia-arenae:0.1570830861)95.5/99:0.1399817572,(WP_192051557.1-unclassified-Massilia:0.1299513741,WP_136222466.1-Massilia-sp.-Mn16-1_5:0.0805148747)86.3/100:0.0836113581)90.7/99:0.0785654412,((WP_057247240.1-Duganella-sp.-Root198D2:0.1713104921,WP_212592286.1-Janthinobacterium-sp.-TND4EL3:0.2832983221)85.8/100:0.0643282796,WP_155457065.1-Massilia-eburnea:0.1257406222)90.1/100:0.0656216565)43.5/97:0.0420410728,(WP_058422493.1-Pseudomonas:0.1551475789,WP_223462047.1-Pseudomonas-sp.-A-R-26:0.2418969380)90.9/99:0.0817004079)71.3/92:0.0118205511,WP_249918475.1-Pseudomonas-sp.-R3.Fl:0.2275021014)84.6/90:0.0283400407)93.6/93:0.0600178282,((WP_176062241.1-Anaeromyxobacter-diazotrophicus:0.1380964260,WP_248361410.1-Anaeromyxobacter-oryzae:0.3173306742)94.9/100:0.1467905041,WP_160809027.1-Nitrosomonas-sp.-HPC101:0.4145315052)95.1/92:0.1391290208)10.9/54:0.0058072600,((((WP_129151777.1-Achromobacter-aloeverae:0.0919579378,WP_088599355.1-unclassified-Achromobacter:0.0747657418)96.6/100:0.0971032931,WP_057654143.1-Bordetella-sp.-N:0.1938197028)76.8/97:0.0468273717,WP_086067427.1-Bordetella:0.2320062631)90.7/100:0.0639974704,(WP_066641528.1-Bordetella-sp.-H567:0.0898921642,(WP_086058993.1-Bordetella-genomosp.-9:0.1937140655,WP_094856450.1-Bordetella-genomosp.-10:0.1563988701)92.8/100:0.0954831212)89.9/100:0.0743212046)97.6/98:0.1281348675)0/61:0.0233101693,((((((((((WP_188630571.1-Oceanisphaera-marina:0.1350115287,WP_204795415.1-Oceanisphaera-litoralis:0.0533225302)14.6/85:0.0127866524,WP_104485062.1-Oceanisphaera-arctica:0.0996533411)99.9/100:0.1978375017,((WP_072282603.1-Syntrophotalea-acetylenivorans:0.1431572506,((WP_092056970.1-Desulfuromonas-acetexigens:0.1481538013,WP_043770486.1-Desulfobulbus-propionicus:0.2545129115)42.5/94:0.0660872459,WP_148894253.1-Geothermobacter-ehrlichii:0.3456618173)80.2/82:0.0348778526)62.3/80:0.0126154526,(WP_028317018.1-Desulfobulbus-elongatus:0.3653989000,WP_155876508.1-Desulfuromonas-sp.-AOP6:0.2252657133)92.1/89:0.1091708693)92.7/98:0.1074327130)71.5/97:0.0451380817,WP_072282412.1-Syntrophotalea-acetylenivorans:0.2169666475)92.2/84:0.0666613863,(WP_053550565.1-Desulfuromonas-soudanensis:0.2892367665,WP_027714505.1-Desulfuromonas-sp.-TF:0.2370496978)76.6/87:0.0427662303)78.8/65:0.0175595975,(((((WP_205222281.1-Desulfogranum-marinum:0.2570907857,WP_205242362.1-Desulfobulbus-alkaliphilus:0.1435313036)87.6/57:0.0615534732,WP_267928740.1-Desulfolithobacter-dissulfuricans:0.1284918172)97.3/79:0.1010846257,((WP_250680171.1-Geomonas-sp.-Red32:0.1897949074,WP_199381999.1-Geomesophilobacter-sediminis:0.2200698246)83.3/100:0.0565322077,WP_205218910.1-Desulforhopalus-vacuolatus:0.3136566189)93/93:0.0659338121)70.6/18:0.0132169161,(WP_036478304.1-Myxosarcina-sp.-GI1:0.1883879661,WP_066347484.1-Geminocystis-sp.-NIES-3708:0.4073091861)94.6/93:0.1382750442)33.7/20:0.0370705304,WP_028582690.1-Desulfogranum-japonicum:0.3496531185)81.7/68:0.0354375234)82.7/56:0.0294846234,(WP_015405619.1-Desulfocapsa-sulfexigens:0.2549818062,(WP_137423090.1-Desulfoglaeba-alkanexedens:0.3526351516,(WP_040200185.1-Geoalkalibacter-subterraneus:0.1917164979,WP_040097704.1-Geoalkalibacter-ferrihydriticus:0.3113996097)82.5/83:0.0598993402)52.3/41:0.0384626524)85/21:0.0647259860)82.6/24:0.0302922608,(WP_103114677.1-Geothermobacter-hydrogeniphilus:0.3298970853,WP_221248821.1-Desulfuromonas-versatilis:0.2490788605)82/99:0.0961585414)85.4/24:0.0419895216,(((WP_163339734.1-Desulfopila-sp.-IMCC35008:0.3016579310,WP_136805251.1-Desulfosediminicola-flagellatus:0.2588420900)0/85:0.0000028837,WP_073615262.1-Desulfopila-aestuarii:0.2471719614)96.6/100:0.1069861973,(WP_228856481.1-Desulfomarina-profundi:0.3496462269,WP_183349429.1-Desulfoprunum-benzoelyticum:0.1697895506)52.4/100:0.0621752479)86.1/55:0.0618044720)90.8/82:0.0440977123,(((((((WP_212718705.1-unclassified-Synechocystis:0.0300428825,WP_228015966.1-Synechocystis-salina:0.1112216931)87.3/100:0.0383755904,WP_010873214.1-unclassified-Synechocystis:0.0361181612)100/100:0.4207340185,(WP_084055900.1-Desulfacinum-hydrothermale:0.3003159005,WP_123289658.1-Desulfosoma-caldarium:0.2875138968)95.9/100:0.1789955491)80.3/57:0.0459787364,((((WP_116346290.1-Cupriavidus-taiwanensis:0.1566348732,WP_124685047.1-Cupriavidus-pauculus:0.2179602974)99.9/100:0.3970345096,WP_119020982.1-Halomonas-sp.-JS92-SW72:0.3802674982)83.3/87:0.0630450070,((WP_200222041.1-Achromobacter-xylosoxidans:0.1521470448,WP_259251071.1-Achromobacter-sp.-JUb104:0.1725854630)94.7/100:0.0800864655,(((((WP_111826915.1-Achromobacter-sp.-HZ01:0.1211951253,WP_102775120.1-Achromobacter:0.0179170619)99.9/100:0.1481531943,(WP_129525645.1-Achromobacter-agilis:0.0316576881,(WP_129240460.1-Achromobacter-veterisilvae:0.0473375324,WP_043544564.1-Achromobacter-sp.-RTa:0.0296000629)89.1/100:0.0203795199)75.8/100:0.0158483405)94.5/100:0.0689449474,(((WP_175208175.1-Achromobacter-anxifer:0.0585214637,(((WP_251864322.1-Achromobacter-sp.-Marseille-Q4962:0.0751370213,WP_164793551.1-Achromobacter-insolitus:0.0337351444)97.8/92:0.0589800412,WP_207472676.1-Achromobacter-sp.-SD115:0.0097427495)93.6/90:0.0372929324,WP_125471167.1-Achromobacter:0.0190007079)94.7/94:0.0432824543)81.6/73:0.0139123935,WP_175164188.1-Achromobacter-aegrifaciens:0.0455984866)88.8/75:0.0335340808,WP_088158656.1-Achromobacter-xylosoxidans:0.0677527199)99.5/98:0.1081309768)70.1/74:0.0102201787,WP_174717299.1-Achromobacter-denitrificans:0.0924990035)91.5/98:0.0413558708,(WP_088588627.1-Achromobacter-marplatensis:0.1668629390,WP_169273406.1-Achromobacter-sp.-Bel:0.0912999306)97.2/100:0.1009852746)58.1/63:0.0230626198)95.2/100:0.1020159814)90.5/76:0.0773352941,(((WP_099797666.1-Thermostichus-lividus:0.4370972710,WP_196167318.1-Pseudomonas-monteilii:0.2849092738)77.6/92:0.0884381247,WP_153720048.1-Spiribacter-sp.-C176:0.3570094201)79.9/87:0.0500024135,WP_048706771.1-Thauera-humireducens:0.2817254371)86.1/55:0.0570209232)71.7/56:0.0166638241)89.2/98:0.0356217464,((((((WP_238511579.1-Cupriavidus-gilardii:0.3094115563,WP_224042884.1-Cupriavidus-respiraculi:0.2647823321)73.4/100:0.0500433793,(WP_018008101.1-Cupriavidus-neocaledonicus:0.0498941911,WP_183298446.1-Cupriavidus-alkaliphilus:0.0552728596)99.7/100:0.1533861074)46.9/95:0.0162063485,WP_116600700.1-Cupriavidus-alkaliphilus:0.1853876079)81.9/93:0.0558479808,((((WP_115680924.1-Cupriavidus-taiwanensis:0.2422327844,WP_135702508.1-Cupriavidus-oxalaticus:0.1536550376)78.9/40:0.0301012767,(WP_018313502.1-Cupriavidus-sp.-UYPR2.512:0.2499438591,WP_174426450.1-Cupriavidus-basilensis:0.3575777862)27.3/13:0.0477077104)73/19:0.0258648333,WP_071014919.1-Cupriavidus-sp.-USMAHM13:0.2521311481)65.3/70:0.0496267821,(WP_150984056.1-Cupriavidus-basilensis:0.4023824843,WP_250479419.1-unclassified-Caballeronia:0.3353797558)77.5/88:0.0495859773)85.5/73:0.0364290241)91.7/96:0.0562881524,((WP_250024752.1-Cupriavidus-campinensis:0.1612956920,WP_094797980.1-Bordetella-genomosp.-5:0.2235899982)55.1/100:0.0992450905,WP_136680209.1-Neptunomonas-sp.-XY-337:0.2168469673)73.1/99:0.0120269697)87.6/69:0.0447682225,(((((((((((((WP_209862330.1-Pseudomonas-sp.-BP8:0.4466388421,(WP_201176141.1-Pseudomonas-sp.-S31:0.0285186366,WP_261240966.1-unclassified-Pseudomonas:0.0494157420)90.2/100:0.0706510036)69.5/88:0.0431706272,WP_225786622.1-Pseudomonas:0.0751145479)83.6/86:0.0347118044,((WP_054891693.1-Pseudomonas:0.0822794763,WP_236238170.1-Pseudomonas-faucium:0.1429196978)62.8/97:0.0374513219,(WP_158461286.1-Pseudomonas-fluorescens:0.2386711084,(WP_200141208.1-unclassified-Pseudomonas:0.1990503764,WP_029612831.1-Pseudomonas-parafulva:0.0188032120)93.3/100:0.0730165359)7.9/68:0.0317817844)94.4/96:0.0640589450)37.8/57:0.0102752035,(((WP_217887155.1-Pseudomonas-xanthosomae:0.0308780755,WP_217840807.1-Pseudomonas-fakonensis:0.0402850794)81/100:0.0167402105,WP_186654015.1-Pseudomonas-xantholysinigenes:0.2145401926)72.2/99:0.0239748800,WP_110605176.1-Pseudomonas-soli:0.2632866129)19.1/53:0.0236475065)99.2/100:0.0915979380,((WP_181290220.1-Pseudomonas-brassicacearum:0.1630319451,((((WP_201004194.1-Pseudomonas-cichorii:0.0799777968,WP_176569640.1-Pseudomonas-eucalypticola:0.1261958531)89.7/71:0.0497054009,(WP_219063768.1-Pseudomonas-sp.-UMAB-08:0.2048670422,((WP_222540780.1-Pseudomonas-sp.-DR208:0.1185661829,(WP_253509974.1-Pseudomonas-rhodesiae:0.1101771470,WP_093202821.1-Pseudomonas-yamanorum:0.0292526715)90/100:0.0394334042)15.2/66:0.0109443851,((WP_065875813.1-Pseudomonas:0.0434678049,(WP_033898696.1-Pseudomonas-fluorescens-group:0.0339968366,(WP_186687394.1-Pseudomonas-sp.-SWRI18:0.0537512618,WP_169378974.1-Pseudomonas:0.0167964327)86.7/100:0.0255266354)76.2/100:0.0117887398)99.2/100:0.0737502949,WP_105697656.1-Pseudomonas-poae:0.0411674896)72.9/75:0.0116042649)28.7/92:0.0392400094)89.5/100:0.0460432096)88.3/71:0.0300842094,WP_043192328.1-Pseudomonas-rhizosphaerae:0.2575230861)0/32:0.0072431441,(WP_249678359.1-Pseudomonas-sp.-PIA16:0.1754165116,WP_268799018.1-Pseudomonas-sp.-BSw22131:0.1649435263)58.7/49:0.0141619583)89.6/97:0.0499488351)84.9/100:0.0442843077,WP_108096457.1-Pseudomonas-sp.-GV071:0.1129080819)75.7/97:0.0706895593)79.2/100:0.0371252804,WP_252273535.1-Pseudomonas-sp.-YIM-B01952:0.1067541989)87/99:0.0348820196,WP_143488038.1-Pseudomonas-mangiferae:0.1374468318)91.3/99:0.0426753358,(((WP_058067682.1-Pseudomonas-sp.-TTU2014-080ASC:0.1543052412,(WP_192399936.1-Pseudomonas-sp.-PDM15:0.1481645588,WP_226282979.1-Pseudomonas-lalkuanensis:0.0677410308)82.2/100:0.0476892069)92.7/99:0.0457077280,WP_003453405.1-Pseudomonas:0.1237650097)76.1/69:0.0099942352,(((WP_077525673.1-Pseudomonas-resinovorans:0.1032848163,WP_137821430.1-Pseudomonas-sp.-D-2018:0.0987114266)97.8/99:0.0760020239,WP_028626762.1-Pseudomonas-resinovorans:0.0375475897)74.2/62:0.0096847941,((WP_016495240.1-Pseudomonas-resinovorans:0.0640127986,WP_044873077.1-Pseudomonas-sp.-LFM046:0.0622864637)90.1/98:0.0359011877,(WP_120654068.1-Pseudomonas-sp.-DY-1:0.0850115647,WP_151138079.1-Pseudomonas-lalkuanensis:0.0426446929)69.8/77:0.0198967103)0/52:0.0000028118)82.7/77:0.0148564637)89/87:0.0378002201)83.4/87:0.0352568646,(((((WP_090448460.1-Pseudomonas-benzenivorans:0.2039707583,((WP_025164100.1-Pseudomonas-taeanensis:0.0575599490,WP_147170838.1-Pseudomonas-sp.-SJZ079:0.0064395273)92.1/100:0.0450135563,(WP_208707982.1-Pseudomonas-sp.-LPB0260:0.0246887624,WP_255840290.1-Pseudomonas-benzenivorans:0.0422207393)95/100:0.0644894832)80.6/93:0.0209450742)21.3/49:0.0146248077,((WP_131198269.1-Pseudomonas-dryadis:0.1234956019,(WP_131178052.1-Pseudomonas-daroniae:0.0639611749,(WP_084307530.1-Pseudomonas-flavescens:0.0689259915,(WP_249909557.1-Pseudomonas-fulva:0.0448726951,(WP_074889242.1-Pseudomonas-argentinensis:0.0252869111,WP_027910132.1-Pseudomonas-sp.-URMO17WK12_I4:0.0453130639)41.1/100:0.0138435175)85.1/100:0.0330255057)72.2/100:0.0262285093)87.1/100:0.0422892966)92.2/100:0.0690906459,(WP_233685494.1-Pseudomonas-anguilliseptica:0.0611891485,WP_257778987.1-Pseudomonas-sp.-32.2.56:0.0000020071)99.7/100:0.1121253876)67.6/62:0.0351036871)82.8/57:0.0197988628,WP_069518708.1-Pseudomonas:0.1323185616)74.7/59:0.0084563448,(((((((WP_119694090.1-Pseudomonas:0.0517722036,WP_106738315.1-Pseudomonas-mendocina:0.0490079201)92.6/100:0.0407322748,WP_150606948.1-Pseudomonas-oleovorans:0.0510562367)95.3/91:0.0435317394,WP_143497094.1-Pseudomonas-sp.-ALS1131:0.0363622756)12.8/8:0.0045077241,(WP_024309956.1-Pseudomonas-sp.-P818:0.0536894727,WP_090337209.1-Pseudomonas-chengduensis:0.0211656198)0/22:0.0000027400)41.1/89:0.0181575523,WP_074939507.1-Pseudomonas-composti:0.0208502228)86.5/95:0.0178767642,WP_021444205.1-Pseudomonas-sp.-EGD-AK9:0.0537525440)86.3/95:0.0237885548,WP_159970704.1-Pseudomonas-sp.-8Z:0.0817313585)96.1/100:0.0626640218)87.5/68:0.0323204451,WP_238040454.1-Pseudomonas-sp.-MMS21-TM103:0.1148089999)87.3/95:0.0337748455)89/95:0.0471200498,(WP_220803742.1-Pseudomonas-sp.-NCCP-436:0.1468483789,WP_090225506.1-Pseudomonas-kuykendallii:0.1200153561)24.7/74:0.0231730193)93.7/90:0.0506888027,(WP_018865194.1-Thioalkalivibrio-sp.-ARh3:0.2688286082,WP_025281494.1-Ectothiorhodospira-haloalkaliphila:0.3869066778)91.6/100:0.0862126175)83.5/91:0.0277608354,((WP_183165579.1-Azomonas-macrocytogenes:0.1658916312,((((WP_102836461.1-Stutzerimonas-stutzeri:0.0277462300,(WP_256071125.1-Stutzerimonas-stutzeri:0.0343833454,WP_045423249.1-Pseudomonadaceae:0.0277796807)81.7/100:0.0130552590)98.1/100:0.0511622788,WP_255961233.1-Stutzerimonas-stutzeri:0.0272256479)0/91:0.0012834279,WP_025243649.1-Stutzerimonas-stutzeri:0.0467073098)97.2/98:0.0560954179,(WP_122077451.1-Pseudomonadaceae:0.0327646619,((WP_037024448.1-Pseudomonas-saudiphocaensis:0.0670024103,WP_181084609.1-Stutzerimonas-stutzeri:0.0071120171)69.3/100:0.0249963657,((WP_011911332.1-Stutzerimonas-stutzeri:0.0212101554,WP_237255234.1-Pseudomonas-oligotrophica:0.0827113092)67/11:0.0217432653,((((WP_230358311.1-Pseudomonas-sp.-NP21570:0.0349680169,WP_200589398.1-Stutzerimonas-stutzeri:0.0323633678)18.5/28:0.0156518578,WP_193680295.1-Pseudomonas-lopnurensis:0.0454828965)81.5/94:0.0203622586,WP_043295157.1-Stutzerimonas-degradans:0.0556003635)70.6/12:0.0116997935,WP_120997453.1-_Pseudomonas:0.0237112820)90.6/12:0.0318777482)60.9/99:0.0212320119)85.3/98:0.0174498476)0/94:0.0000021918)84/98:0.0360737816)84/100:0.0436975846,WP_028238613.1-Stutzerimonas-azotifigens:0.0603737553)99.6/100:0.1199775742)86.8/91:0.0310066987,((((WP_213445339.1-Variovorax-sp.-PCZ-1:0.3824556851,WP_226921657.1-Kinneretia-sp.-XES5:0.1678497214)85.4/97:0.1062180655,WP_091467897.1-Methylophilus-rhizosphaerae:0.3368218846)63.1/56:0.0274792934,(WP_228215938.1-Aromatoleum-toluclasticum:0.2715372325,WP_226440716.1-Quatrionicoccus-australiensis:0.2947754764)86.5/55:0.0627329256)71.5/51:0.0378477227,WP_199264031.1-Desulfobulbus-oligotrophicus:0.3177545833)82.9/89:0.0383632810)73.6/52:0.0160214887)77.9/44:0.0213236473)68.1/57:0.0344536772,((((((((((WP_198111724.1-Inhella-proteolytica:0.4551165299,WP_241396682.1-Pelomonas-sp.-CA6:0.1301446940)73.8/99:0.0402312712,(WP_037482909.1-Sphaerotilus-natans:0.1974456958,(WP_028996355.1-Azohydromonas-australica:0.0813585220,WP_066337818.1-Azohydromonas-lata:0.0965715847)94.4/100:0.1086649989)63.7/100:0.0922006155)92.6/99:0.1195800498,(WP_047193087.1-Schlegelella-brevitalea:0.2516887073,WP_019561187.1-Caldimonas-manganoxidans:0.2349382488)72.1/95:0.1098851453)76.7/100:0.0917919417,((((((WP_169107474.1-Comamonas-suwonensis:0.0492113733,WP_219343559.1-Comamonas-sp.-Y33R10-2:0.0865035656)93.7/100:0.0530122814,(WP_003073420.1-Comamonas:0.0723228888,((WP_211457163.1-Comamonas-sp.-PE63:0.0491199164,WP_148729134.1-Comamonas-sp.-Z3:0.0450123308)55.7/72:0.0066762882,WP_034391098.1-Comamonas-thiooxydans:0.0150470718)74.4/72:0.0082860854)87.4/97:0.0272712141)70.4/97:0.0147218479,WP_218241177.1-Comamonas-fluminis:0.1123116978)99.1/100:0.1537242622,WP_012203208.1-Delftia-acidovorans:0.2046272177)12.7/85:0.0593107602,WP_058879441.1-Comamonas-kerstersii:0.1244115692)93.6/100:0.0886495372,WP_066475960.1-Comamonas-terrae:0.1137587827)99.6/100:0.1937252625)84.2/89:0.0430402217,(((((WP_220807781.1-Noviherbaspirillum-aridicola:0.5007891903,(KAI6244242.1-Aphelenchoides-fujianensis:0.5522656086,KAI6170547.1-Aphelenchoides-bicaudatus:0.6662214666)96.2/99:0.1981666201)48.8/95:0.0748467077,(((((((WP_018154793.1-Methanothermococcus-thermolithotrophicus:0.1530896291,WP_048201153.1-Methanocaldococcus-bathoardescens:0.0526245783)93/96:0.0615665253,(((WP_010868297.1-Pyrococcus-abyssi:0.0634433790,((((WP_013798345.1-Methanotorris-igneus:0.0391722064,WP_013866394.1-Methanothermococcus-okinawensis:0.1848537121)79.7/94:0.0258145268,WP_007044577.1-Methanotorris-formicicus:0.0643071131)98.6/100:0.0855484773,((((((((((((WP_237702072.1-Thermococcus-sp.-AM4:0.0281215679,WP_223211916.1-Thermococcus-henrietii:0.0636496643)98.7/100:0.0755299350,WP_058938040.1-Thermococcus-celericrescens:0.0314021242)73.4/76:0.0138845486,(WP_139679991.1-Thermococcus-indicus:0.0397767650,WP_148882943.1-Thermococcus-aciditolerans:0.0317522689)80.2/100:0.0287049620)74.7/75:0.0076426518,WP_088865724.1-Thermococcus-barossii:0.0555126642)86.3/87:0.0134624083,WP_232460978.1-Thermococcus-radiotolerans:0.0512086290)0/86:0.0000024603,WP_167711671.1-unclassified-Thermococcus:0.0256891899)62.6/96:0.0163430679,WP_258085050.1-Thermococcus-sp.-813A4:0.0304202164)85.4/94:0.0193613147,WP_088181368.1-Thermococcus-sp.-5-4:0.0529061501)93.9/100:0.0300107548,(WP_152879125.1-Thermococcus-sp.-101-C5:0.0535253075,WP_015850310.1-Thermococcus-sibiricus:0.0492952061)73.6/99:0.0073379582)86.9/99:0.0133384094,WP_253304977.1-Thermococcus-aggregans:0.0395493768)83.9/86:0.0138982308,WP_088862340.1-Thermococcus-celer:0.1002092507)60.6/64:0.0067901567,WP_068664737.1-Thermococcus-piezophilus:0.0573403476)85.8/89:0.0213449184)54.6/84:0.0172292490,((((WP_048103006.1-Aciduliprofundum-boonei:0.2643483214,WP_148120035.1-Methanofervidicoccus-sp.-A16:0.1438197124)88.7/100:0.0365105829,(WP_013468354.1-Thermococcus-barophilus:0.0468161619,WP_013906489.1-Pyrococcus-yayanosii:0.0267637499)69.5/76:0.0191951798)77/78:0.0086252816,WP_048053121.1-Pyrococcus-horikoshii:0.0330112174)0/54:0.0000023185,(((WP_167913028.1-Thermococcus-sp.-21S7:0.0772598219,WP_048150691.1-Palaeococcus-ferrophilus:0.0273571390)92.2/100:0.0295513939,(WP_167888701.1-Thermococcus-sp.-MV5:0.0539849204,WP_209475361.1-Thermococcus-stetteri:0.0738472823)89.6/84:0.0381784720)22.6/60:0.0140633582,(WP_088855609.1-Thermococcus-siculi:0.0556700526,WP_087036111.1-Thermococcus-litoralis:0.0375058312)93.3/96:0.0389120380)81.7/63:0.0133202132)83.8/84:0.0324791043)62.7/98:0.0352678029)55.3/98:0.0403390374,WP_014733609.1-Pyrococcus-sp.-ST04:0.0570785423)100/100:0.2803479940,(((WP_245533116.1-Coprothermobacter-platensis:0.5743614816,WP_022856039.1-Thermodesulfobacterium:0.1893192142)52.1/100:0.0848477226,WP_150071182.1-Thermodesulfobacterium-sp.-TA1:0.2125678419)92.3/100:0.1299390022,(((WP_029522925.1-Persephonella-sp.-KM09-Lau-8:0.1266016713,(WP_200674641.1-Persephonella-atlantica:0.0725501404,WP_096999769.1-Persephonella-hydrogeniphila:0.1219553078)84.9/100:0.0538537589)93.7/100:0.0602720781,WP_028950938.1-Sulfurihydrogenibium-subterraneum:0.1365931720)26.5/90:0.0207276217,((WP_132526207.1-Phorcysia-thermohydrogeniphila:0.0864597681,WP_013638952.1-Desulfurobacterium-thermolithotrophum:0.0515185638)83/100:0.0201100613,WP_089322203.1-Desulfurobacterium-atlanticum:0.1574573356)94.2/100:0.0634513234)96.3/100:0.1119506641)83.8/98:0.0526616762)98.3/98:0.1435495981)28.9/74:0.0232074966,(((WP_011869065.1-Methanococcus-maripaludis:0.0925054754,WP_013998786.1-Methanococcus-maripaludis:0.0565359486)74.6/100:0.0371525639,WP_181493664.1-Methanococcus-maripaludis:0.0742229587)58.5/99:0.0182463290,WP_181504225.1-Methanococcus-maripaludis:0.0477737072)100/100:0.2580473452)68.4/84:0.0802593281,WP_052562942.1-Candidatus-Brocadia-sinica:0.3927942349)72.7/88:0.0895250755,WP_173298646.1-Thermanaeromonas-sp.-C210:0.2474399246)94.3/99:0.1852104027,((WP_119360783.1-Meiothermus-luteus:0.1969198096,WP_027877894.1-Meiothermus-cerbereus:0.0579414903)81.5/100:0.0633813486,WP_208013758.1-Meiothermus-sp.-CFH-77666:0.0740683362)100/100:0.5608556929)58.3/100:0.0318337658,WP_011996389.1-Coxiella-burnetii:0.6149275631)98/100:0.2543452311)55.5/94:0.0688010561,(WP_226989104.1-Desulfuromonas-sp.-TF:0.3982684213,WP_227459030.1-Cupriavidus-pauculus:0.3172894385)83.6/100:0.0768686456)81.1/82:0.0600744027,(WP_227390214.1-Halomonas-malpeensis:0.5772895942,(WP_083560927.1-Oceanococcus-atlanticus:0.3663220164,WP_104228621.1-Solimonas-fluminis:0.2462358424)96.5/100:0.1518488663)82.5/95:0.0570584366)93.5/97:0.0688997761,WP_227181580.1-Leeia-sp.-TBRC-13508:0.3457106380)95/81:0.0827781673)89.7/66:0.0540793684,((((WP_132974196.1-Thiogranum-longum:0.1702758138,WP_201344172.1-Thiohalobacter-sp.-COW1:0.2038094926)99.7/100:0.3145613155,(WP_110265962.1-Sinimarinibacterium-flocculans:0.3918486673,(WP_234070476.1-Myxococcus-stipitatus:0.2424756637,WP_120555541.1-Corallococcus-aberystwythensis:0.2861950421)98.7/100:0.2506220684)89.7/98:0.1203242455)25/89:0.0602762416,((WP_230291920.1-Croceicoccus-sp.-Ery5:0.4695054602,WP_164156863.1-Sandarakinorhabdus-rubra:0.2027523131)88.8/97:0.0949407299,((WP_070114691.1-Acidithiobacillus-caldus:0.3125380212,(((WP_009566950.1-Acidithiobacillus:0.0439747962,WP_215870853.1-unclassified-Acidithiobacillus:0.0796309730)94.8/100:0.0690624530,WP_163095688.1-Acidithiobacillus-ferrianus:0.0479827138)45.5/95:0.0392478656,WP_101536718.1-Acidithiobacillus-sp.-SH:0.1525023503)90.5/100:0.0956374484)91.2/100:0.1496863179,WP_152145882.1-Alkalilimnicola-sp.-S0819:0.3178861917)90.6/100:0.1134981434)75.8/97:0.0270330204)88/88:0.0606317827,WP_122244186.1-Vandammella-animalimorsus:0.2835323705)56.1/67:0.0289062392)84.3/65:0.0403583877,WP_154206467.1-Ralstonia-pickettii:0.2220213623)25.7/56:0.0566536677,WP_253536546.1-Ralstonia-sp.-21YRMH01-3:0.4783785093)0/68:0.0005047905,(((WP_036252969.1-Methylobacter-sp.-BBA5.1:0.2700713952,((((WP_019918777.1-Methyloversatilis-discipulorum:0.0456568309,WP_018229738.1-Methyloversatilis-universalis:0.0600555350)54.9/100:0.0726511324,WP_183409942.1-Litorivivens-lipolytica:0.5096739890)48.2/100:0.0558900648,(WP_101941536.1-Uliginosibacterium-sp.-TH139:0.1108421450,WP_095523004.1-Candidatus-Dactylopiibacterium-carminicum:0.0925381355)97.5/100:0.1584354535)94.2/100:0.1062972858,WP_074865046.1-Atopomonas-hussainii:0.2667627696)53/93:0.0457581468)87.6/91:0.0387952380,(((WP_109041087.1-Azospira-sp.-I13:0.1089327231,WP_014237023.1-Azospira-oryzae:0.1365364782)99/100:0.1465787114,WP_201673630.1-Ramlibacter-monticola:0.2048189750)68.2/18:0.0505058139,(WP_067106371.1-Methylobacillus-sp.-MM3:0.2229575658,WP_212785890.1-Ferrigenium-kumadai:0.2287158473)71.1/21:0.0207143077)93.1/99:0.0612582687)75.9/69:0.0107186547,(((((((((WP_046292890.1-Pandoraea-oxalativorans:0.0705246254,WP_150719538.1-Pandoraea-capi:0.1178354657)100/100:0.3851477567,WP_101249932.1-Telmatospirillum-siberiense:0.3742221008)73.4/98:0.0635939412,WP_220807643.1-Noviherbaspirillum-aridicola:0.2963859927)94.7/68:0.1214557175,((WP_074796726.1-Nitrosospira-briensis:0.0694696684,(WP_090882638.1-Nitrosovibrio-sp.-Nv6:0.0603490594,WP_097065525.1-Nitrosovibrio-sp.-Nv4:0.0335261315)92.1/100:0.0635878120)99.9/100:0.2691590047,(WP_223986959.1-Cupriavidus-pampae:0.0555537423,((WP_011517306.1-Cupriavidus-metallidurans:0.0506968194,WP_019451253.1-Cupriavidus-sp.-BIS7:0.0539323778)0/90:0.0023278131,WP_082079449.1-Cupriavidus-basilensis:0.0834905557)84.1/90:0.0298906793)99.7/100:0.1583140909)19.3/40:0.0221061440)47.1/38:0.0131884410,WP_174484170.1-methanotrophic-endosymbiont-of-Bathymodiolus-puteoserpentis:0.2052968586)86.1/71:0.0208909589,WP_137918285.1-Hydrogenophaga-sp.-2FB:0.1824311062)0/24:0.0000024288,((WP_026436726.1-Acidovorax-sp.-JHL-9:0.2168363747,(WP_086926522.1-Acidovorax-carolinensis:0.1770979044,WP_166067558.1-Diaphorobacter-sp.-HDW4B:0.2221681092)59/68:0.0335491067)79.3/96:0.0213437871,WP_169420802.1-Ramlibacter-agri:0.2537538937)90.3/51:0.0313065290)74.7/17:0.0071329877,((((((((WP_106935816.1-unclassified-Variovorax:0.2485358487,WP_068630051.1-Variovorax-sp.-PAMC-28711:0.2143855029)58.5/97:0.0616891313,WP_206177103.1-Variovorax-sp.-RKNM96:0.0977677922)51.2/80:0.0295612468,WP_093207264.1-Variovorax-sp.-YR750:0.1092695758)44.2/45:0.0341005715,WP_126747644.1-Variovorax-sp.-DXTD-1:0.1649515046)81.7/84:0.0481054523,(WP_093441791.1-Variovorax-sp.-770b2:0.1285706960,WP_160553990.1-Xylophilus-rhododendri:0.1892348146)70.6/98:0.0583840669)73.9/79:0.0295257074,WP_192194832.1-Variovorax-sp.-38R:0.2152704946)60.6/55:0.0388660283,WP_219210863.1-Variovorax-boronicumulans:0.1897553808)94.4/95:0.0570888208,WP_137890779.1-Ramlibacter-sp.-2FC:0.1127467803)77.6/16:0.0154846271)87.9/50:0.0197884421,(WP_057673948.1-Curvibacter-sp.-PAE-UM:0.1715032149,WP_056898839.1-Pseudorhodoferax-sp.-Leaf274:0.2718785577)72.3/60:0.0294749228)91/77:0.0356844138)92.2/84:0.0333711992)97.3/77:0.0590281502,(WP_217345508.1-Noviherbaspirillum-sp.-L7-7A:0.2375838280,WP_047549861.1-Methylotenera-sp.-G11:0.2028100317)91.2/100:0.0659346533)75.6/39:0.0142954517)54/55:0.0447846603,(((((((((((((((((((((((WP_089129802.1-Tolypothrix-sp.-NIES-4075:0.0817775898,WP_096563566.1-Scytonema-sp.-NIES-4073:0.1165969536)77.3/92:0.0427708049,WP_048866388.1-Scytonema-tolypothrichoides:0.0893910339)69.8/92:0.0331132742,WP_096563464.1-Scytonema-sp.-NIES-4073:0.0841286900)100/100:0.1918965351,((WP_015128325.1-Calothrix-sp.-PCC-7507:0.1202223572,WP_017653186.1-Fortiea-contorta:0.1775377854)48.6/100:0.0528914824,((WP_193199005.1-Nostoc-sp.-MG11:0.0431856301,WP_162396505.1-Nostoc-sp.-B-2019:0.0499312549)94.5/100:0.0539451763,WP_214439445.1-Atlanticothrix-silvestris:0.1428434955)76.9/100:0.0672199948)7.1/81:0.0158618388)58.4/86:0.0151523630,(((WP_045870014.1-unclassified-Tolypothrix:0.0274064049,WP_190656031.1-Nostoc-linckia:0.0355699363)98.6/100:0.0741106202,(WP_190743070.1-Tolypothrix-sp.-FACHB-123:0.0486151236,WP_190545010.1-Calothrix:0.0343722543)97.3/100:0.0745771641)76.5/100:0.0310017763,WP_086758595.1-Nostoc-sp.-106C:0.0707872959)88.4/100:0.0428395834)52.6/80:0.0236897334,WP_016866333.1-Fischerella:0.0856729572)61.9/83:0.0176329100,(((((WP_009344618.1-Cylindrospermopsis-raciborskii:0.0743956319,WP_240868223.1-Cylindrospermopsis-raciborskii:0.0230854319)99.5/100:0.1328303953,WP_096570274.1-Sphaerospermopsis:0.0492108844)0/55:0.0093860981,WP_220611028.1-Sphaerospermopsis-torques-reginae:0.0549574349)98.6/100:0.0927522039,((WP_237989437.1-Nostoc-sp.-UHCC-0870:0.0754583328,WP_190559626.1-Anabaena-sphaerica:0.1007562093)5.5/65:0.0238361203,WP_206263219.1-Nostoc-sp.-UHCC-0702:0.0856022692)84.4/98:0.0304207591)33/100:0.0244666606,WP_096557790.1-Nostoc-sp.-NIES-4103:0.1594458244)3/35:0.0376263478)99.7/100:0.1413153610,((((((((WP_106232846.1-Pleurocapsa-sp.-CCALA-161:0.1664531536,WP_229639079.1-Waterburya-agarophytonicola:0.0300068842)88.4/100:0.0406626668,WP_096721927.1-Chondrocystis-sp.-NIES-4102:0.0815951695)97.9/100:0.1272291688,WP_006510035.1-Xenococcus-sp.-PCC-7305:0.1774295015)91.5/100:0.1017343345,WP_264322079.1-Zarconia-navalis:0.1874681818)78.8/80:0.0503472364,(WP_012595695.1-Rippkaea-orientalis:0.1838958721,(((WP_007308862.1-Crocosphaera-watsonii:0.0858127321,WP_107668138.1-Cyanothece-sp.-BG0011:0.0333397271)84.6/99:0.0140889317,WP_009545547.1-Crocosphaera-subtropica:0.0205998859)82.2/96:0.0151669696,WP_008278255.1-Crocosphaera-chwakensis:0.0440348583)97.1/97:0.1019344694)39/52:0.0377985515)88.2/98:0.0357555138,(WP_106909014.1-Microcystis-sp.-MC19:0.2695822838,WP_124977990.1-Aphanothece-sacrum:0.2130804108)40.9/71:0.0091944984)87.7/98:0.0320689315,WP_013324143.1-Gloeothece-verrucosa:0.1129890588)70.2/62:0.0258194123,WP_015143663.1-Pleurocapsa-sp.-PCC-7327:0.1549007555)44.6/78:0.0326288916)65.7/82:0.0189943283,(WP_015193936.1-Stanieria-cyanosphaera:0.0283875911,WP_096384422.1-Stanieria-sp.-NIES-3757:0.0867618276)99.9/100:0.1728218538)94.8/99:0.0530514459,WP_194029353.1-Lusitaniella-coriacea:0.2608754179)50.5/88:0.0222741408,(WP_224410051.1-Oscillatoria-salina:0.2236471433,WP_069965800.1-unclassified-Desertifilum:0.2646535552)81.9/46:0.0398477231)42.9/40:0.0088743118,(WP_226578226.1-Microseira-wollei:0.1977981292,((WP_073595522.1-Phormidium-ambiguum:0.0690415846,WP_193982193.1-Phormidium-sp.-LEGE-05292:0.0713934447)99/100:0.1066944591,(WP_249064905.1-Argonema-antarcticum:0.0556801226,WP_190468950.1-unclassified-Planktothrix:0.0885152689)12.6/80:0.0170210252)79.2/96:0.0346584420)94.5/97:0.0622718180)75.3/70:0.0132813848,((((((WP_102179481.1-Fischerella-thermalis:0.0324578778,((WP_096675774.1-Fischerella-sp.-NIES-4106:0.0498469245,WP_053458209.1-Hapalosiphon-sp.-MRB220:0.0333054192)73.3/100:0.0067422457,(WP_131119525.1-Westiellopsis-prolifica:0.0493703678,WP_016860528.1-Fischerella-muscicola:0.0269039220)87.7/100:0.0141339602)96.3/100:0.0658060966)99.9/100:0.1859441581,(WP_015202791.1-Crinalium-epipsammum:0.2111658790,WP_035154029.1-Calothrix-sp.-336/3:0.3515993058)81.1/97:0.0522480961)48.2/29:0.0319434356,WP_190825375.1-Iningainema-tapete:0.2534409049)68.4/31:0.0173170107,((WP_099102643.1-Nostoc-sp.-Peltigera-malacea-cyanobiont-DB3992:0.0874756455,((WP_190950926.1-Nostoc-punctiforme:0.0500814243,(WP_221704991.1-Desmonostoc-muscorum:0.0474879529,WP_096537650.1-Nostoc-linckia:0.0147694688)75.9/96:0.0072278368)58.9/89:0.0255132815,WP_190959861.1-Nostoc:0.0156915524)96.8/96:0.0815430747)99.8/100:0.1150961210,(((((((WP_193938462.1-Cuspidothrix-issatschenkoi:0.1628726673,((WP_096666215.1-Dolichospermum-compactum:0.0594255283,(WP_190387835.1-Aphanizomenon:0.0620289184,WP_168466156.1-Aphanizomenon:0.0196698820)73.4/100:0.0181146677)83.1/97:0.0153561266,WP_015080416.1-Anabaena-sp.-90:0.0286801809)85.2/97:0.0134083178)80.5/23:0.0255776325,((WP_190392428.1-Dolichospermum-sp.-FACHB-1091:0.0056967662,(WP_028089231.1-Dolichospermum-circinale:0.0193141218,WP_137908967.1-Nostocales:0.0488930210)97.8/100:0.0677575332)99.9/100:0.1276396143,WP_168502193.1-Anabaena-sp.-UHCC-0187:0.0000022444)77.4/92:0.0071920666)78.1/19:0.0145275886,(WP_190370804.1-Anabaena-sp.-FACHB-1237:0.1535837798,WP_168537172.1-Anabaena-sp.-UHCC-0253:0.0407172759)89.7/93:0.0318091196)100/100:0.1195408320,(WP_010998622.1-Nostocaceae:0.0624055572,WP_011320078.1-Nostocaceae:0.0592233455)77.9/100:0.0181941921)95.2/99:0.0505442956,((WP_015138800.1-Nostoc-sp.-PCC-7524:0.0745241719,WP_066425233.1-Anabaena-sp.-4-3:0.0898900286)97.3/100:0.0690201701,(WP_199337214.1-Nostoc-sp.-FACHB-152:0.1120334076,(WP_239729951.1-Nodularia-sphaerocarpa:0.0691210984,WP_173403323.1-Nodularia-spumigena:0.0469981809)93.9/100:0.0459428004)86.5/87:0.0232761985)69/85:0.0076482906)79.3/91:0.0166578138,WP_199337786.1-Nostoc-sp.-FACHB-280:0.1295258605)94.9/100:0.0599297075,(WP_096624129.1-Calothrix-sp.-NIES-3974:0.0757056748,WP_095721999.1-Calothrix-elsteri:0.0979572920)99.3/100:0.1169644239)79/100:0.0242616950)95.5/100:0.0502293572)88.6/99:0.0354261759,WP_006529211.1-Gloeocapsa-sp.-PCC-73106:0.3380193105)92.9/76:0.0478545023,((WP_008178350.1-Moorena-producens:0.2409262696,(((WP_250123015.1-Chroococcidiopsis-sp.-CCMEE-29:0.2242872767,((WP_233747769.1-Leptodesmis-sichuanensis:0.2242564771,(WP_192216758.1-Microcoleus-sp.-FACHB-SPT15:0.1696828238,WP_224345144.1-Kovacikia-minuta:0.1136084651)56.9/100:0.0258772040)77.7/88:0.0165463803,(WP_223049852.1-Leptothermofonsia-sichuanensis:0.0746036506,WP_073549891.1-Chroogloeocystis-siderophila:0.1741378315)79.3/87:0.0315737947)91.6/90:0.0394280708)78.8/61:0.0087439154,WP_190531554.1-Coleofasciculus-sp.-FACHB-542:0.2041787638)0/53:0.0000024412,(WP_193873404.1-unclassified-Coleofasciculus:0.1207561400,WP_254010234.1-Symplocastrum-sp.-BBK-W-15:0.1463094176)92.2/80:0.0413272613)23.7/60:0.0134099019)25.1/76:0.0086807480,(WP_190666400.1-Coleofasciculus-sp.-FACHB-501:0.1202263801,WP_190525322.1-Coleofasciculus-sp.-FACHB-T130:0.0600440179)97.4/100:0.0761653782)91.1/99:0.0336035050)77.1/46:0.0086919291)91.6/64:0.0333897389,((((((((((WP_202221768.1-Okeania-sp.-KiyG1:0.0441318727,WP_094672483.1-Hydrocoleum-sp.-CS-953:0.0430922169)66.1/93:0.0170526986,WP_124146462.1-Okeania-hirsuta:0.0047282695)99.1/100:0.1580321849,WP_071516849.1-Geitlerinema-sp.-PCC-9228:0.2291189628)71.7/92:0.0649658696,WP_008052766.1-Limnospira-indica:0.2414546463)77/93:0.0487528431,(WP_009783380.1-Lyngbya:0.0774672288,WP_046277942.1-Limnoraphis-robusta:0.0531275828)93.6/100:0.0665856424)86.7/85:0.0489946584,((WP_083623900.1-Planktothrix-serta:0.0353499033,WP_072717475.1-Planktothrix:0.0317857677)36.9/93:0.0214734830,(WP_079676747.1-Planktothrix-sp.-PCC-11201:0.0259121221,(WP_026797127.1-Planktothrix:0.0271461952,WP_214559879.1-Planktothrix-agardhii:0.0409761403)21.5/99:0.0276698514)97.1/100:0.0707815490)96.2/100:0.0633038914)89.6/87:0.0419479862,WP_190877489.1-unclassified-Planktothricoides:0.1494772661)76.3/62:0.0353039667,WP_200665545.1-Oxynema-sp.-CENA135:0.2234142389)68.4/73:0.0177858410,(WP_261197618.1-unclassified-Laspinema:0.0409003752,(WP_207087219.1-Phormidium-pseudopriestleyi:0.0353981028,WP_261204505.1-Laspinema-sp.-D2d:0.0419928107)71/100:0.0223801745)100/100:0.1078531641)89.5/42:0.0297640069,(WP_190665495.1-Microcoleus-sp.-FACHB-672:0.0731622586,WP_261220767.1-Ancylothrix-sp.-D3o:0.1807260925)97.2/96:0.0896772158)83.8/43:0.0176247230)81.6/21:0.0187838181,((WP_190493486.1-unclassified-Microcoleus:0.2745935859,WP_190772446.1-unclassified-Leptolyngbya:0.2820315694)0/19:0.0000025978,(WP_007355666.1-Kamptonema:0.0971921161,(((((WP_236507253.1-Tychonema-sp.-BBK16:0.0403296424,WP_096828456.1-Tychonema-bourrellyi:0.0674412120)92.1/100:0.0354685123,WP_242724542.1-Microcoleus-vaginatus:0.0336812160)83.2/100:0.0133116268,WP_015174083.1-Oscillatoria-nigro-viridis:0.0195702434)88.9/100:0.0208729903,(WP_194009709.1-unclassified-Tychonema:0.0378139034,WP_265235823.1-Lyngbya-sp.-CCAP-1446/10:0.0431868610)89.2/100:0.0207958016)41.3/98:0.0114737817,WP_193973976.1-Microcoleus-sp.-LEGE-07076:0.0400038617)99/100:0.1089493556)92.6/81:0.0563839545)36.8/18:0.0191416919)90.6/56:0.0405711734,WP_017715496.1-Oscillatoria-sp.-PCC-10802:0.1591704550)74.4/78:0.0325634375,WP_017661508.1-Baaleninema-simplex:0.3105266746)91.9/59:0.0596167010,(WP_015170385.1-Geitlerinema-sp.-PCC-7407:0.2234954622,(WP_204103763.1-Spirulina-major:0.4414248268,(WP_258826205.1-Massilia-agri:0.4126577469,(WP_002810067.1-Nitrosococcus-oceani:0.0460191225,WP_013221233.1-Nitrosococcus-watsonii:0.0722519238)100/100:0.2458111078)76.6/24:0.0400279902)78.3/17:0.0286933535)87.8/37:0.0421932104)75.3/11:0.0162505384,(((WP_204137747.1-Halomicronema-sp.-CCY15110:0.0503714909,WP_039724568.1-Leptolyngbya-sp.-LK:0.0808020708)99.5/100:0.1669405539,WP_023076112.1-Leptolyngbya-sp.-Heron-Island-J:0.2193186115)94.6/100:0.0999501954,WP_017324474.1-Synechococcus-sp.-PCC-7336:0.2640908152)92.8/99:0.0729026822)88.5/32:0.0347271021,WP_054689510.1-Desulfosarcina-cetonica:0.3245512436)77.9/9:0.0215280701,WP_066727622.1-Desulfuromonas-sp.-DDH964:0.3700418591)39.1/5:0.0303086867,((((WP_148717329.1-Chitinolyticbacter-meiyuanensis:0.5209927763,WP_123210144.1-Stagnimonas-aquatica:0.6876490509)93.6/92:0.2086241986,(WP_108623287.1-Orrella-marina:0.3249948629,WP_243478771.1-Orrella-sp.-f23:0.4074234038)86.2/99:0.0849851176)87.1/59:0.0942608668,((((WP_205202385.1-Azonexus-hydrophilus:0.2307733370,WP_173581083.1-Azonexus-fungiphilus:0.2488855699)55.8/84:0.0288271022,WP_121459656.1-Azonexus-fungiphilus:0.2656511319)92.9/89:0.0739024220,((((((WP_136386324.1-Pseudothauera-rhizosphaerae:0.1966926920,(WP_114649035.1-Pseudothauera-hydrothermalis:0.0989165775,WP_187718084.1-Thauera-sp.-CAU-1555:0.1524241868)77.8/90:0.0409415602)97.1/100:0.1536157813,((((((WP_238617884.1-Ectothiorhodospira-shaposhnikovii:0.1405898561,WP_025282248.1-Ectothiorhodospira-haloalkaliphila:0.1224051030)92.6/100:0.0730949156,WP_238622182.1-unclassified-Ectothiorhodospira:0.1209035654)91.7/100:0.0783517435,((((((WP_060482506.1-Pseudomonas-sp.-NBRC-111119:0.2035841014,((WP_196129978.1-Pseudomonas-fulva:0.0850562378,WP_201164743.1-Pseudomonas-sp.-S60:0.0446760854)98.3/100:0.0788287745,(((WP_174142546.1-Pseudomonas-putida:0.1376573461,WP_027907237.1-Pseudomonas:0.0420263437)35.7/100:0.0120828935,WP_075044077.1-Pseudomonas-putida:0.0454523482)76.4/100:0.0330246825,(WP_023379297.1-Pseudomonas-sp.-VLB120:0.0997383716,WP_202675963.1-Pseudomonas-sp.-SK2:0.0362869754)98/100:0.0702727006)89.9/100:0.0394962270)86.8/100:0.0241295104)75.3/79:0.0163179910,(((WP_159409308.1-Pseudomonas-putida:0.1440771742,(((WP_182365203.1-Pseudomonas-juntendi:0.0731032477,WP_182365213.1-Pseudomonas:0.1082426996)72.7/77:0.0179805162,((WP_110607173.1-Pseudomonas-fulva:0.0137874200,WP_088523134.1-unclassified-Pseudomonas:0.0505137497)95.8/100:0.0353500751,(WP_167339957.1-Pseudomonas-fulva:0.0322574357,WP_186476580.1-Pseudomonas-promysalinigenes:0.0424931593)83.1/99:0.0258280049)80.9/73:0.0158144317)89.2/63:0.0167967561,(WP_201214098.1-Pseudomonas-sp.-S36:0.0221358113,WP_054890059.1-Pseudomonas:0.0651425869)95.6/99:0.0449437019)63/65:0.0041792295)73.1/64:0.0110670825,(WP_167277242.1-Pantoea-sp.-Tr-811:0.0204674940,WP_103449615.1-Pseudomonas-putida:0.0560477401)98.3/100:0.0552823280)68.3/58:0.0081123849,(WP_115272924.1-Pseudomonas-putida:0.1301626611,WP_267821885.1-Pseudomonas-putida:0.0529525038)88.9/71:0.0346438507)75/81:0.0144610030)85.2/81:0.0277389295,(WP_046489762.1-Pseudomonas-veronii:0.1095375310,(WP_059395284.1-Pseudomonas-putida:0.0945547688,(WP_009685980.1-Pseudomonas:0.0769922711,WP_024086460.1-Pseudomonas:0.0650824288)77.1/100:0.0424236271)88.2/99:0.0445188850)69.7/93:0.0555761240)79.2/91:0.0133841498,(WP_029885231.1-Pseudomonas:0.0247561332,(WP_230064532.1-Pseudomonas-sp.-CIP-10:0.0213166784,WP_197863301.1-Pseudomonas-putida:0.0420064214)97.8/100:0.0460229922)75.5/93:0.0070297433)95.2/91:0.0659842600,WP_060394622.1-Pseudomonas-entomophila:0.1122259067)97/98:0.0857635025,WP_183165449.1-Azomonas-macrocytogenes:0.1176156491)79.5/96:0.0329660400)78.9/85:0.0352030950,WP_004074545.1-Desulfobacter-postgatei:0.3149437116)29.4/73:0.0291085001,WP_256068944.1-Stutzerimonas-stutzeri:0.1614793073)89.5/92:0.0804994565,WP_026685886.1-Azovibrio-restrictus:0.1429537476)83/64:0.0687924385)52.8/67:0.0402984254,WP_168987795.1-Azoarcus-taiwanensis:0.2258099009)85.6/91:0.0416705457,(((WP_167682126.1-Rhodocyclus-gracilis:0.3192541817,(WP_153115862.1-Rhodocyclus-tenuis:0.0508308323,WP_200385811.1-Rhodocyclus-tenuis:0.0190079306)92.5/100:0.0733610743)93/100:0.0819985339,WP_201212349.1-Rhodocyclus-purpureus:0.1524283475)95.9/100:0.1220995645,WP_153110537.1-Propionivibrio-limicola:0.2600588115)84.7/92:0.0775510003)81.7/42:0.0356471657,WP_216129682.1-Azospira-inquinata:0.4123178886)81.3/49:0.0330201422,(WP_013164002.1-Desulfurivibrio-alkaliphilus:0.2731936392,WP_263768642.1-Propionivibrio-sp.-SG131:0.2295261507)91.1/56:0.0947216123)75.2/98:0.0155422230)86.1/61:0.0408048695,((WP_117174481.1-Rhodoferax-lacus:0.2375236704,((WP_094482496.1-Rhodoferax-sp.-TH121:0.0890335513,WP_105262671.1-Rhodoferax-sp.-TS-BS-61-7:0.0438728200)87.2/99:0.0471593788,WP_214121631.1-Curvibacter-sp.-CHRR-16:0.2627089782)92.3/100:0.1129696070)49.5/85:0.0360100367,(WP_066884254.1-Dechloromonas:0.1000954028,WP_027458229.1-Dechloromonas-agitata:0.1205950100)96.6/100:0.1002909682)91.9/94:0.0651340227)73.4/62:0.0117579391)0/30:0.0000021300,WP_209394989.1-Rhodoferax-sp.-AJA081-3:0.3862137991)89.2/55:0.0495041506)82.6/10:0.0309160567,(((WP_014706296.1-Methylophaga-nitratireducenticrescens:0.0000024152,WP_136552334.1-Methylophaga-sp.-SB9B:0.0764371742)100/100:0.4715626121,(WP_153584563.1-Caenimonas-koreensis:0.2983507256,(WP_007185580.1-Hydrocarboniphaga-effusa:0.0773635822,WP_007185565.1-Hydrocarboniphaga-effusa:0.0747793439)98.7/100:0.1744145224)91/100:0.1301139629)58.4/61:0.0610272799,WP_054772520.1-Methylogaea-oryzae:0.2170412150)18.9/14:0.0307223525)68.6/22:0.0135205000)80.7/35:0.0360399184)87.7/66:0.0368283220)89.2/85:0.0664628026,((((((WP_239190817.1-Candidatus-Nitrotoga-sp.-HW29:0.0568045446,WP_239797721.1-Candidatus-Nitrotoga-arctica:0.0283387692)48.1/91:0.0075298263,WP_239287644.1-Candidatus-Nitrotoga-sp.-1052:0.0217261903)71.6/75:0.0080288178,WP_239235002.1-Candidatus-Nitrotoga-sp.-BS:0.0514923646)99.5/100:0.2138313596,(WP_044881819.1-Neochlamydia-sp.-EPS4:0.0622903661,WP_166155173.1-unclassified-Neochlamydia:0.0000024290)100/100:0.4070633131)75.5/97:0.0508994415,((WP_237889684.1-Dissulfurimicrobium-hydrothermale:0.2789871475,WP_009207329.1-Sulfuricella-denitrificans:0.3161392604)55.8/66:0.0377224996,WP_237247350.1-Sideroxyarcus-emersonii:0.2098726346)69.8/93:0.0640690986)80.5/89:0.0265653449,(((((((((((WP_065305794.1-Janthinobacterium-psychrotolerans:0.1349284575,(WP_139090581.1-Janthinobacterium-lividum:0.0806347404,WP_035822645.1-Janthinobacterium-sp.-RA13:0.0530870558)74.5/99:0.0082863815)59.6/99:0.0053315672,WP_034778721.1-Janthinobacterium-lividum:0.0361911086)82/100:0.0125631505,WP_152282105.1-Janthinobacterium-violaceinigrum:0.0208824860)77.3/100:0.0067781895,WP_183742413.1-unclassified-Janthinobacterium:0.0487631635)75.8/100:0.0064254433,((((WP_094442213.1-Janthinobacterium-sp.-PC23-8:0.0677291631,WP_230539136.1-Janthinobacterium-sp.-FW305-129:0.0541933434)6.8/100:0.0069131162,WP_215403374.1-Janthinobacterium-sp.-JC611:0.0246498002)77.4/100:0.0097005305,WP_102124214.1-Janthinobacterium-sp.-ROICE36:0.0350004697)75.5/100:0.0071276225,WP_096238059.1-Janthinobacterium-svalbardensis:0.0206230061)77.4/100:0.0070437442)99.4/100:0.1088111448,(WP_101482628.1-unclassified-Janthinobacterium:0.1980397086,(WP_099917421.1-Massilia-psychrophila:0.1404225778,(WP_205319084.1-Glaciimonas-sp.-PAMC28666:0.0758622478,WP_092396716.1-Collimonas-sp.-OK412:0.1231070003)0/85:0.0079055370)87.5/100:0.0345179989)50.2/92:0.0201077199)82.4/98:0.0186419198,WP_023495308.1-Methyloglobulus-morosus:0.1671418332)98.3/99:0.0952702592,WP_217900122.1-Noviherbaspirillum-humi:0.2132067123)71.5/89:0.0147293022,(((WP_162020603.1-Undibacterium-crateris:0.1582144379,WP_186884883.1-Undibacterium:0.0383413490)32.8/96:0.0124726990,WP_186861877.1-Undibacterium-griseum:0.0734763907)99.3/100:0.1122437691,(((((WP_207000528.1-Trinickia-sp.-DHG64:0.1386569914,WP_124149903.1-Paraburkholderia-dinghuensis:0.1016830900)65.4/100:0.0540793289,(WP_266250364.1-Paraburkholderia-megapolitana:0.1771886959,WP_026121090.1-Paraburkholderia-kururiensis:0.0954178853)66.9/100:0.0620546348)93.9/100:0.0940949883,((((((((((((((WP_059459219.1-Burkholderia-vietnamiensis:0.2326711481,WP_121276956.1-Trinickia-fusca:0.1062502851)53.6/100:0.0391477030,WP_134195781.1-Paraburkholderia-rhizosphaerae:0.2692015955)84.9/100:0.0311323376,WP_136892825.1-Trinickia-sp.-7GSK02:0.0433418265)83.2/100:0.0187580393,WP_201650988.1-Paraburkholderia-sabiae:0.0419257713)28.3/53:0.0130594726,WP_205970438.1-Paraburkholderia-sp.-Tr-20389:0.0422974340)69.7/53:0.0158760674,WP_054928771.1-Paraburkholderia-caribensis:0.0310377363)82/100:0.0198899935,(WP_205976236.1-Paraburkholderia-sp.-Se-20369:0.2466615020,WP_206951398.1-Trinickia-sp.-4D114:0.0651169852)98.3/95:0.1178070815)75.6/70:0.0180447831,WP_209935850.1-Paraburkholderia-sp.-LEh10:0.0907826706)59.8/88:0.0181119780,WP_061168975.1-Caballeronia-hypogeia:0.0962001054)56.1/16:0.0137030224,((WP_060156927.1-Burkholderia-ubonensis:0.0886214853,WP_059889870.1-Burkholderia-stagnalis:0.1976119186)89.3/96:0.0308010362,(WP_006496666.1-Burkholderia-cenocepacia:0.0274826123,WP_059716874.1-Burkholderia-ubonensis:0.0397192651)88.8/100:0.0208141089)75.1/80:0.0075067792)77.7/16:0.0108672975,((WP_175791145.1-Burkholderia-ambifaria:0.1013123990,WP_059842355.1-Burkholderia-multivorans:0.0380381242)77.6/77:0.0117344149,WP_105772332.1-Burkholderia-multivorans:0.0351953908)93.3/81:0.0290730740)44.2/86:0.0153974304,(WP_060092900.1-Burkholderia-vietnamiensis:0.0258494108,WP_226113708.1-Burkholderia-sp.-AU33545:0.0460898115)99.9/100:0.1609318986)67.6/28:0.0090969848,(WP_168792792.1-Paraburkholderia-aromaticivorans:0.0659087541,((WP_004544228.1-Burkholderia:0.0279194660,WP_069272519.1-Burkholderia-cepacia:0.2335033917)78.9/96:0.0174371437,WP_059583574.1-pseudomallei-group:0.0405388845)93.3/96:0.0274707849)82.8/61:0.0240575376)85.5/71:0.0209663842,((WP_201138986.1-Paraburkholderia-domus:0.0470683088,WP_266262001.1-Paraburkholderia-madseniana:0.0254267388)98.9/100:0.1146579085,WP_009889585.1-Burkholderia-thailandensis:0.0924440975)92.7/100:0.0476102806)67.6/99:0.0604104330)68.6/97:0.0381860710,(WP_179404315.1-Burkholderia-guangdongensis:0.1834168833,((WP_063494936.1-Paraburkholderia-phytofirmans:0.0305730522,(WP_254318368.1-Paraburkholderia-sp.-CNPSo-3272:0.0063495207,WP_042262232.1-Paraburkholderia-heleia:0.0633211194)92.5/100:0.0407933877)98.6/100:0.0974978329,WP_038711716.1-Burkholderia-sp.-lig30:0.2389935134)83.8/99:0.0539879661)79.3/96:0.0318544215)86/95:0.0501697564,WP_056136218.1-Massilia-sp.-Root133:0.1989822596)99/100:0.1279416058)90.1/99:0.0467616224)74.8/79:0.0192495683,((WP_195874647.1-Herminiimonas-contaminans:0.1332464628,WP_041297045.1-Janthinobacterium-sp.-Marseille:0.0962891820)78.5/100:0.0289880533,(WP_076590920.1-Herminiimonas-arsenitoxidans:0.1674187188,(WP_206604468.1-Herminiimonas-fonticola:0.0453884143,WP_188420499.1-Oxalicibacterium-solurbis:0.1756320343)63.7/100:0.0143804128)85.9/100:0.0374401798)99.4/100:0.1122100077)82.6/77:0.0359043761,(WP_247737070.1-Thauera-aromatica:0.0455351794,WP_039805139.1-Azotobacter-chroococcum:0.1606990221)98.1/99:0.1176373786)92.4/99:0.0655465665)81.3/51:0.0214866015)98.5/100:0.1118143987,(((WP_213263532.1-Burkholderia-pyrrocinia:0.1384923247,WP_060279477.1-Burkholderia-cepacia-complex:0.1369140034)97.2/100:0.1330330397,(((WP_208101977.1-Burkholderia-sp.-D-99:0.0336466255,WP_205714023.1-Burkholderia-cepacia:0.0296579228)71.4/100:0.0077870504,WP_174919663.1-Burkholderia-metallica:0.0215792571)85.7/100:0.0426350511,WP_077021900.1-Burkholderia-cenocepacia:0.0820580230)99.5/100:0.2041765648)90.1/100:0.0509365823,(WP_159673755.1-Andreprevotia-sp.-IGB-42:0.2014026926,WP_084093192.1-Andreprevotia-lacus:0.0639373351)99.6/100:0.1934655054)57.4/98:0.0250881574)93.4/100:0.0780088364,(WP_189531325.1-Paludibacterium-paludis:0.4468599898,(WP_043620211.1-Chromobacterium-piscinae:0.1957839353,(WP_103900886.1-Chromobacterium-alticapitis:0.2243329096,WP_045052286.1-Chromobacterium-violaceum:0.2367611037)88.4/100:0.0848133384)88.1/100:0.0907287840)84.8/100:0.0501871890)87.9/99:0.0283095900)0/92:0.0011786792,WP_020932001.1-Cycloclasticus-zancles:0.3264936090)94.2/80:0.0850902048)86/80:0.0462454019,(((WP_147247296.1-Brenneria-salicis:0.0359799153,WP_172289906.1-unclassified-Brenneria:0.1030226173)86.8/100:0.0365819832,WP_039356706.1-Pectobacterium-actinidiae:0.0793354096)93.9/100:0.0860944824,((WP_198997094.1-Pectobacterium-carotovorum:0.0188807209,(WP_165707548.1-Pectobacterium:0.0324685851,WP_174866005.1-Pectobacterium-polaris:0.0261398780)98.9/100:0.0633691963)61.4/100:0.0170266733,WP_145962313.1-Pectobacterium-peruviense:0.0541888165)97.7/100:0.1103794926)96.3/100:0.0980318388)64.3/89:0.0331946803,(((((((WP_236369955.1-Aeromonas-veronii:0.0472017449,(((WP_042019775.1-Aeromonas-sobria:0.0522453251,WP_139495680.1-Aeromonas-veronii:0.0237575292)96.9/100:0.0550136239,(WP_098969895.1-Aeromonas-sp.-CU5:0.0457114750,WP_159163188.1-Aeromonas-veronii:0.0489810034)90.4/100:0.0356369395)91.2/100:0.0362777126,WP_113736183.1-Aeromonas-caviae:0.0407931855)43.6/96:0.0182379985)0/95:0.0040733317,(((((((WP_080988732.1-Aeromonas-caviae:0.0373150247,WP_108536035.1-Aeromonas-sp.-HMWF014:0.0424553685)92.2/100:0.0351583881,WP_252615985.1-Aeromonas-hydrophila:0.0655877918)40.1/87:0.0184841362,WP_214003767.1-Aeromonas-caviae:0.0476358316)75.4/76:0.0076667291,WP_143237783.1-Aeromonas-caviae:0.0637389431)37.9/66:0.0089201877,WP_226032029.1-Aeromonas-hydrophila:0.0461944703)85/72:0.0194072150,WP_113739100.1-Aeromonas-veronii:0.0465851555)0/65:0.0000023478,(WP_139459904.1-Aeromonas-veronii:0.0388558094,WP_041208446.1-Aeromonas-jandaei:0.1529779578)99.4/92:0.1011446661)93.5/100:0.0752389852)95.1/100:0.0557610567,WP_201035193.1-Aeromonas-media:0.1184974083)68.3/93:0.0388650325,((WP_263173612.1-Shewanella-sp.-SM73:0.1474028805,((WP_263191039.1-Shewanella-sp.-SM87:0.0682203152,WP_263255112.1-Shewanella-sp.-SM32:0.0471711819)76.9/95:0.0155135569,(WP_263178828.1-Shewanella-sp.-SM101:0.0419347338,WP_109248233.1-Shewanella-algae:0.0689552361)91.8/100:0.0361581322)86/100:0.0399633930)93.4/100:0.0637542966,(WP_220777726.1-Shewanella-hafniensis:0.0268466019,WP_055647506.1-Shewanella-sp.-Sh95:0.0321734817)92.3/100:0.0443564209)77.7/98:0.0248713808)72.5/93:0.0135653641,WP_082969903.1-Aeromonas-dhakensis:0.1115307692)92.2/99:0.0474475913,WP_104454419.1-Aeromonas-sp.-ASNIH2:0.0132150153)98.6/99:0.0944910599,(((((WP_073350970.1-Aeromonas-hydrophila:0.0405047788,WP_082041448.1-Aeromonas-eucrenophila:0.2831752757)95.1/97:0.0840607151,WP_190284026.1-Aeromonas-hydrophila:0.0428581655)85.2/97:0.0295579800,WP_241344946.1-Aeromonas-sp.-MR16:0.0679855398)93.4/100:0.0832896359,(WP_223952962.1-Aeromonas-caviae:0.1808872378,WP_244081328.1-Aeromonas-caviae:0.1785482014)24.7/53:0.0513953233)66.9/55:0.0524012029,((WP_208149206.1-Shewanella-algae:0.2973136349,(((WP_045500143.1-Vibrio-hyugaensis:0.1682747065,WP_045419977.1-Vibrio-campbellii:0.0773320941)44.2/99:0.0355572132,WP_025533544.1-Vibrio-parahaemolyticus:0.0693220953)92.1/100:0.0443071957,WP_011788809.1-Shewanella-sp.-W3-18-1:0.0713760657)66.7/100:0.0221952003)92.6/100:0.0575111908,WP_219142360.1-Aeromonas-sanarellii:0.0935223977)99.4/100:0.1480082055)63.4/61:0.0195557613)88.7/96:0.0326863039)84.2/98:0.0197489578,((WP_076573677.1-Aeromonas-sp.-RU39B:0.1381222387,WP_134696535.1-Aeromonas-taiwanensis:0.6246236959)92.7/98:0.1062829088,WP_108652490.1-Dongshaea-marina:0.1889847800)91.8/98:0.0684611431)85.9/96:0.0241396892,(((((WP_132257458.1-Paucimonas-lemoignei:0.0734307376,WP_119785517.1-Noviherbaspirillum-sedimenti:0.0653149790)100/100:0.3416817988,(WP_026470910.1-Alkanindiges-illinoisensis:0.2294142566,(WP_248147450.1-Moraxella-osloensis:0.3212776831,WP_092690500.1-Acinetobacter-kyonggiensis:0.1627606160)88.6/100:0.0920053022)98.6/100:0.2213520291)90/100:0.0881143695,(WP_094838100.1-Bordetella-genomosp.-4:0.2013291149,WP_160351730.1-Bordetella-sp.-15P40C-2:0.2176374925)99.3/100:0.2473573615)94.4/99:0.1138054840,(WP_208005626.1-Oceanisphaera-pacifica:0.0708802772,(WP_087034853.1-Oceanisphaera-profunda:0.0711352953,WP_087034873.1-Oceanisphaera-profunda:0.1122395533)93/100:0.0804411207)94.9/100:0.0972532580)66.8/96:0.0364267044,WP_200561028.1-unclassified-Erwinia:0.3372114451)87/76:0.0471897788)58.1/62:0.0225509770,(((WP_234664799.1-Aeromonas-allosaccharophila:0.1710817841,(((((((WP_201942221.1-Aeromonas-veronii:0.1690564772,WP_159143398.1-Aeromonas-veronii:0.0135582669)73.6/73:0.0108904394,(WP_203131090.1-Aeromonas-jandaei:0.0619373833,((WP_042657830.1-Aeromonas:0.0451717270,WP_111901368.1-Aeromonas-veronii:0.0384672669)38.3/98:0.0145286440,WP_139692403.1-Aeromonas-sobria:0.0364327261)0/95:0.0000020775)65.3/94:0.0107649159)40.6/68:0.0130131463,WP_268854930.1-Aeromonas-enteropelogenes:0.1147268052)56.1/74:0.0075012091,WP_223380193.1-Aeromonas-enteropelogenes:0.0322179311)99.2/100:0.0992672028,(WP_236321701.1-Aeromonas-veronii:0.0695307502,(WP_257119864.1-Aeromonas-hydrophila:0.0225546209,((((((WP_248067241.1-unclassified-Shewanella:0.1744942390,(WP_218558465.1-Shewanella-putrefaciens:0.0709026110,WP_011919582.1-Shewanella-putrefaciens:0.0385643619)96.9/100:0.1151074645)96.6/94:0.1086958022,WP_043167431.1-Aeromonas-hydrophila:0.0950735344)75.2/91:0.0131432070,WP_252420256.1-Aeromonas-taiwanensis:0.0851358640)76/93:0.0106676279,WP_077098334.1-Aeromonas-hydrophila:0.0608883046)94.4/93:0.0494965302,(WP_050718733.1-Aeromonas-tecta:0.1519638665,WP_100859741.1-Aeromonas-lusitana:0.0400716417)73.2/97:0.0129280076)60/94:0.0375951738,WP_042654929.1-Aeromonas-encheleia:0.0353588086)96.5/100:0.0716069165)67.7/59:0.0070110612)85.2/86:0.0358552088)4.5/35:0.0151205026,(WP_257707542.1-Aeromonas-caviae:0.0514346800,WP_182951616.1-Aeromonas-sp.-WP2-W18-CRE-05:0.0658276289)96.6/100:0.0796886240)95.5/98:0.0710785602,WP_069554789.1-Aeromonas-sp.-DNP9:0.1177948301)81.7/39:0.0487399352)40.7/15:0.0387775881,((((WP_216990617.1-Aeromonas-sp.-FDAARGOS-1409:0.1231813944,WP_236563281.1-Aeromonas-veronii:0.1486279041)49/74:0.0351558200,(((WP_059168599.1-Aeromonas-salmonicida:0.0686280969,(WP_033115905.1-Aeromonas-lacus:0.0471433377,((WP_223920301.1-Aeromonas-caviae:0.0348273897,WP_226058924.1-Shewanella-sp.-SR1:0.1242668993)81.3/98:0.0194672403,(WP_058394234.1-Aeromonas-salmonicida:0.0307205368,WP_257690725.1-Aeromonas-veronii:0.0498498404)77.3/98:0.0199058990)83.9/96:0.0196631356)87.8/98:0.0407842534)84.5/98:0.0412749324,((WP_268010485.1-Aeromonas-dhakensis:0.0783522361,WP_098985072.1-Aeromonas-sp.-CA23:0.0377456311)99.3/100:0.1340809967,(WP_181268965.1-Aeromonas-veronii:0.0325808538,(WP_129146711.1-Aeromonas-caviae:0.0246256977,(WP_081843667.1-Aeromonas-taiwanensis:0.0191316957,WP_139744464.1-Aeromonas-media:0.0430101159)86.7/100:0.0228528543)87.2/100:0.0211112788)77/100:0.0612272275)99.7/100:0.1839876516)80.7/81:0.0217750622,(((WP_223945983.1-Aeromonas-caviae:0.1140451909,(((WP_265458754.1-Aeromonas-salmonicida:0.0814714382,WP_125597700.1-Aeromonas-salmonicida:0.0246731924)0/98:0.0015347604,WP_163128050.1-Aeromonas-allosaccharophila:0.0881670102)0/93:0.0000021247,(WP_180911080.1-Aeromonas-caviae:0.0399915807,WP_264818242.1-Aeromonas-caviae:0.0267317581)91.6/100:0.0201057728)75/93:0.0083714963)75.3/93:0.0062808887,(WP_040098646.1-Aeromonas:0.0953951096,((WP_263309349.1-Aeromonas-caviae:0.0389452805,(WP_041217344.1-Aeromonas-hydrophila:0.0339170922,WP_148505453.1-Vibrio-cholerae:0.1969659925)77/96:0.0094285132)75.4/96:0.0108149323,(WP_257686696.1-Aeromonas-veronii:0.0347387472,WP_255912544.1-Aeromonas-sp.-JL9:0.0328626908)74.9/95:0.0082155166)77.3/99:0.0143561398)76.1/99:0.0123517084)73.3/98:0.0090373561,(((WP_139750984.1-Aeromonas-media:0.0308808386,WP_224430864.1-Aeromonas-rivuli:0.0283091127)80/100:0.0261519181,WP_041996113.1-Aeromonas-bivalvium:0.0530685367)89.6/100:0.0334397973,WP_265776268.1-Aeromonas-veronii:0.1308882528)83.4/94:0.0207144670)91.7/93:0.0420381629)87.8/76:0.0390551807)82/44:0.0291198109,((WP_100646402.1-Aeromonas-veronii:0.0575914521,((WP_252601295.1-Aeromonas-dhakensis:0.0240882399,WP_139432874.1-Aeromonas-dhakensis:0.1435417243)89/100:0.0316936030,WP_106553824.1-Aeromonas-hydrophila:0.0593951569)80.3/100:0.0289281009)98.8/100:0.1504828426,WP_224211331.1-Aeromonas-enteropelogenes:0.2763313658)77.8/70:0.0279863647)74.6/42:0.0085929846,((WP_213320225.1-Aeromonas-caviae:0.0641740776,WP_203764723.1-Aeromonas-caviae:0.0803469517)96.4/100:0.0982218777,(WP_257692500.1-Aeromonas-caviae:0.1688161508,WP_113719982.1-Aeromonas-salmonicida:0.1319653131)86.6/96:0.0696029857)90.8/62:0.0581828729)22.8/14:0.0119269713)84.3/68:0.0245205163,(WP_265468300.1-Aeromonas-salmonicida:0.1522682225,((((((WP_245131329.1-Aeromonas-veronii:0.1380633274,(WP_136501435.1-Aeromonas-veronii:0.1756767850,(WP_268029297.1-Aeromonas-hydrophila:0.1091806595,WP_201942265.1-Aeromonas-veronii:0.1339881343)95.2/100:0.1012214221)63.9/97:0.0446968825)0/65:0.0072578263,((WP_101347551.1-Aeromonas-sobria:0.0748965336,WP_201955411.1-Aeromonas-veronii:0.0480268004)94.8/100:0.0639327383,WP_210550341.1-Aeromonas-sp.-SrichE-2G:0.2839837560)66.8/86:0.0289544763)88.5/79:0.0337640525,(((WP_050666161.1-Aeromonas-schubertii:0.0635107824,WP_059168611.1-Aeromonas-salmonicida:0.1162998713)72.5/100:0.0335911365,WP_210600017.1-Aeromonas-sp.-MrichA-1:0.0373957938)88.3/100:0.0450028528,(WP_225159845.1-Aeromonas-caviae:0.1287466612,(WP_163135231.1-Aeromonas-rivipollensis:0.0309952874,WP_241279715.1-Aeromonas-sp.-MR19:0.0314828887)100/100:0.2248466554)57.7/83:0.0337064531)83.6/78:0.0285951242)84.7/96:0.0283381884,((((((WP_148612677.1-Aeromonas-sobria:0.0482025670,WP_201355655.1-Aeromonas-hydrophila:0.0474283860)0/60:0.0057461302,WP_201951905.1-Aeromonas-veronii:0.0379024532)75.2/98:0.0093864719,WP_171268810.1-Aeromonas-media:0.0236453063)100/100:0.1537271481,(WP_042063659.1-Aeromonas-allosaccharophila:0.0363494588,WP_256684552.1-Aeromonas-salmonicida:0.0343655756)53.1/96:0.0158176565)59/98:0.0089162149,(((WP_212539243.1-Aeromonas-salmonicida:0.0343085309,((WP_187590672.1-Aeromonas-hydrophila:0.1221921390,WP_017782281.1-Aeromonas-dhakensis:0.0337926374)50.7/100:0.0131508182,WP_060586794.1-Aeromonas-schubertii:0.0513408760)99.6/100:0.0934265355)88.9/100:0.0280703606,WP_201904077.1-Aeromonas-veronii:0.0603728019)85.6/100:0.0202709205,WP_206874356.1-Aeromonas-veronii:0.0105625383)75.9/99:0.0095197854)98.2/97:0.0885609544,WP_042649704.1-Aeromonas-media:0.0990264824)36.6/69:0.0490536642)78.5/57:0.0216371641,(WP_253626250.1-Aeromonas-hydrophila:0.1548024624,(WP_139406182.1-Aeromonas-veronii:0.2562640709,WP_061279316.1-Kluyvera-cryocrescens:0.3975143277)78.3/55:0.0368457176)78.9/55:0.0235326544)74.7/75:0.0119615012,(((WP_268421127.1-Aeromonas-media:0.0664499560,WP_111910157.1-Aeromonas:0.0209770458)73.5/96:0.0122765807,WP_128343177.1-Aeromonas-caviae:0.0499819042)94.3/100:0.0555379285,(WP_268037199.1-Aeromonas-dhakensis:0.0968018873,(WP_139733908.1-Aeromonas-veronii:0.0179007293,((WP_234579942.1-Aeromonas-salmonicida:0.0726662221,(WP_234676856.1-Aeromonas-rivipollensis:0.0153150453,WP_269382839.1-Aeromonas-hydrophila:0.0814807815)52.8/100:0.0077479802)52.5/93:0.0118970938,WP_222134508.1-Vibrio-cholerae:0.1507080815)0/93:0.0030654120)97.3/100:0.0742534659)74.5/97:0.0218545797)87.3/100:0.0476378609)90.9/76:0.0350239655)0/56:0.0000021561)96.3/77:0.0498726985)0/72:0.0000020432,((WP_047682696.1-Xenorhabdus:0.0585139479,WP_099124588.1-Xenorhabdus-stockiae:0.0737648716)97.9/100:0.1367067377,(WP_074024273.1-Xenorhabdus-eapokensis:0.0730481186,WP_217470569.1-Photorhabdus-akhurstii:0.0761705931)88.6/100:0.0737391040)98.9/100:0.1112016492)83.5/60:0.0290811779,(((WP_151054911.1-Cellvibrio-sp.-KY-YJ-3:0.1827537807,WP_087465811.1-Cellvibrio-sp.-PSBB006:0.1957273778)75.4/22:0.0239997473,(WP_078044787.1-Cellvibrio-sp.-PSBB023:0.2183980653,WP_012489013.1-Cellvibrio-japonicus:0.1626841901)71.5/22:0.0234205525)99.7/100:0.1193708481,((((WP_097405671.1-Kosakonia:0.0482602445,WP_161518249.1-Kosakonia-sp.-MH5:0.0647403665)90.7/100:0.0645987100,WP_128482629.1-Kosakonia-cowanii:0.1107431844)98.4/100:0.1308741588,WP_206164314.1-Citrobacter:0.2115062735)65.9/96:0.0550450458,(WP_048221111.1-Citrobacter:0.1186593643,WP_263807164.1-Salmonella-enterica:0.1919011295)99.3/100:0.1970846039)84/86:0.0474482495)87.1/68:0.0298586769)95.4/81:0.0556583615,((((((((((((((((((((WP_085037002.1-Cronobacter-sakazakii:0.1924759615,WP_076770125.1-Kosakonia-cowanii:0.1540723909)98.9/100:0.1862801644,((WP_227534785.1-Erwinia-rhapontici:0.1447799300,WP_017800228.1-Erwinia-toletana:0.1464780975)91.2/100:0.0916417028,WP_255851625.1-Erwinia-persicina:0.1776031851)86.9/100:0.0806722167)48/75:0.0223737700,WP_216981205.1-Lelliottia-amnigena:0.3084034371)87.8/99:0.0635804477,(WP_089585421.1-Escherichia-coli:0.0406862640,WP_104705897.1-Escherichia-coli:0.0324671889)93.5/100:0.0930558505)95.6/99:0.1026225108,(WP_001610415.1-Escherichia-coli:0.0629048683,WP_138183825.1-Escherichia-sp.-E4385:0.0000024782)99/100:0.1567478399)94.8/96:0.0735847358,WP_001100804.1-Enterobacteriaceae:0.1123785564)87/85:0.0480876895,(WP_137650031.1-Escherichia-albertii:0.0954615369,(WP_001100786.1-Escherichia-coli:0.1111886766,(WP_248249449.1-Enterobacter-roggenkampii:0.1476969202,WP_032936727.1-Citrobacter-freundii:0.1279755483)99.3/100:0.1575957535)85.2/96:0.0350997947)66/88:0.0256575504)24.6/28:0.0361961371,(WP_157959988.1-Citrobacter:0.4138238640,WP_073446180.1-Escherichia-coli:0.1512342324)87.7/86:0.1092884021)72.8/43:0.0277538688,(((((WP_107192407.1-Escherichia-albertii:0.0205604746,WP_181673375.1-Escherichia-fergusonii:0.0428580285)98.2/100:0.1057996818,((WP_032214194.1-Escherichia-coli:0.0477817343,(WP_059217428.1-Escherichia-albertii:0.0319694882,(WP_073278747.1-Escherichia-coli:0.0839159901,WP_087893724.1-Enterobacteriaceae:0.0302532188)99.7/100:0.1352072660)77.7/97:0.0149696063)64.6/97:0.0077175474,WP_085459683.1-Escherichia-coli:0.0866814334)98.9/100:0.1550320243)93.8/100:0.0669846051,WP_179154141.1-Citrobacter-freundii:0.3490750093)0/27:0.0000020605,WP_033870520.1-Enterobacteriaceae:0.1837243575)86.4/47:0.0217886107,WP_267387874.1-Citrobacter-portucalensis:0.2678827215)28.2/43:0.0316416659)80.3/34:0.0232913226,(WP_095844187.1-Escherichia-coli:0.0515183231,(WP_086118650.1-Escherichia-coli:0.0260240867,WP_089538038.1-Escherichia-coli:0.0330934536)81.9/100:0.0170166771)100/100:0.1505138004)76/30:0.0149801998,(WP_100272167.1-Citrobacter-portucalensis:0.2703541683,WP_138098269.1-Jejubacter-calystegiae:0.1841486240)0/11:0.0041422605)80.6/11:0.0087179560,(((WP_189493202.1-Formosimonas-limnophila:0.2349288307,WP_133620003.1-Hydromonas-duriensis:0.2538793364)99.6/100:0.2232066783,(WP_213789144.1-Enterobacter-hormaechei:0.0003320692,((WP_148242178.1-Enterobacter-asburiae:0.0508277761,WP_237994110.1-Enterobacter-sp.-DRP3:0.0187722422)73.3/100:0.0058219958,WP_237424876.1-Enterobacter-cloacae-complex-sp.-ECC445:0.0456673817)88.2/100:0.0517274667)98.3/100:0.1996519610)91.9/99:0.1227809936,(WP_016236761.1-Enterobacteriaceae:0.0886938497,WP_024239504.1-Escherichia-coli:0.1216147224)99.8/100:0.1464136379)0/9:0.0000022317)96.6/91:0.0482676654,(((((((((((WP_151711904.1-Enterobacter-ludwigii:0.0279104119,(WP_032941356.1-Citrobacter-freundii:0.2022257201,WP_058678686.1-Enterobacter:0.0764122576)90.2/100:0.0583721736)81.3/100:0.0710687240,WP_223560970.1-unclassified-Enterobacter:0.0000021877)98.6/100:0.2582836892,WP_044720521.1-Escherichia-coli:0.3611640592)98/100:0.2536969424,WP_064374050.1-Pantoea:0.2574754746)85.5/99:0.0541716340,(WP_036421240.1-Morganella-morganii:0.1480897165,(WP_264998074.1-Morganella-morganii:0.0715942086,WP_186849509.1-Morganella-morganii:0.0851089716)93.9/100:0.0804576434)94/100:0.1072160346)89.2/97:0.0534474446,((((WP_257124601.1-Aeromonas-veronii:0.3503552993,WP_039304623.1-Cedecea-neteri:0.3940123205)65.3/55:0.0389429230,((WP_043158915.1-Aeromonas-hydrophila:0.4778894034,WP_038258850.1-Xenorhabdus-bovienii:0.2276959227)37.1/71:0.0372991278,WP_086113286.1-Xenorhabdus-beddingii:0.2359984185)47.7/75:0.0581069148)0/36:0.0154919562,(WP_211007190.1-Pectobacterium-brasiliense:0.4479716552,(((((((WP_159241675.1-Proteus-terrae:0.1765963771,WP_195697765.1-Providencia-rettgeri:0.1757981986)92.3/95:0.0753912677,WP_238563424.1-Providencia-rettgeri:0.1121748608)74/37:0.0210238644,WP_165880752.1-Providencia-rettgeri:0.1774013871)84.5/97:0.0274420912,WP_231068620.1-Providencia-huaxiensis:0.1455968208)25.7/41:0.0245223633,WP_004924376.1-Providencia-stuartii:0.2046786629)0/69:0.0000022663,((WP_187128479.1-Providencia-sp.-JUb39:0.1507368783,WP_193115284.1-Escherichia-coli:0.1485834356)87.7/100:0.0449801513,WP_154625743.1-Providencia-stuartii:0.1567717670)85.8/100:0.0272322168)80.3/97:0.0284535618,(WP_169342312.1-Providencia-stuartii:0.1670837641,WP_164565644.1-Providencia-rettgeri:0.2016458742)74.8/98:0.0521694134)78.6/99:0.0416829767)92.6/83:0.0748446670)76.4/63:0.0160975925,(WP_038223662.1-Xenorhabdus-bovienii:0.1994093228,((WP_161748440.1-Proteus-sp.-G2658:0.1999077421,WP_049199273.1-Proteus-mirabilis:0.0652956270)44.5/100:0.0449171537,WP_157954980.1-Proteus-terrae:0.0641746407)96.4/100:0.1192407559)89.9/98:0.0709406532)83.3/74:0.0227645004)71.8/73:0.0155938228,(((((WP_090123257.1-Kosakonia-arachidis:0.0839122498,WP_020454293.1-Kosakonia-sp.-SOY2:0.1001022643)86.2/100:0.0429292314,WP_017458471.1-Kosakonia-sacchari:0.1579481676)76.3/100:0.0479433254,(WP_200002836.1-Citrobacter-freundii:0.1694569000,WP_080329661.1-Enterobacter-cloacae-complex:0.1456737740)98.2/100:0.1721725740)78.5/100:0.0899469605,WP_090460354.1-Enterobacter-sp.-kpr-6:0.0784085395)96.6/99:0.1729865405,WP_168756429.1-Aeromonas-hydrophila:0.4575571818)79.2/81:0.0533639528)84.2/85:0.0251423765,(((WP_234505032.1-Kluyvera-intermedia:0.3954674245,WP_230863267.1-Enterobacter-cloacae:0.4379727670)0/26:0.0696385709,(WP_029399168.1-Escherichia-coli:0.3055218889,((WP_034496584.1-Buttiauxella-agrestis:0.3339761454,WP_164716793.1-Cedecea-lapagei:0.3581195019)88.7/100:0.1066067269,WP_050862535.1-Enterobacter:0.3710209032)32/84:0.0170421889)93.4/98:0.1194589265)84.7/95:0.0509719492,(WP_194857598.1-Citrobacter-freundii:0.3603688720,(WP_141032926.1-Escherichia-coli:0.1054093460,(WP_196390204.1-Enterobacter-mori:0.0516900738,(WP_252032207.1-Enterobacter-asburiae:0.0496372298,WP_111966211.1-Enterobacter-cloacae-complex:0.0312743774)91.8/95:0.0529016394)42.6/88:0.0414133324)93.9/95:0.1275179394)98.8/100:0.2227056545)73.8/85:0.0177527949)90.6/85:0.0364658835,((WP_159282060.1-Rahnella-variigena:0.3696063205,(WP_106902182.1-Citrobacter-freundii:0.2085924513,(WP_126291239.1-Enterobacter-hormaechei:0.1482993252,((((WP_062932819.1-Enterobacter-hormaechei:0.0305981805,WP_252042312.1-Enterobacter-chengduensis:0.0420180767)76.2/100:0.0254155672,WP_048999124.1-Enterobacter:0.0373552619)76.3/99:0.0086843296,WP_063144508.1-Enterobacter-cloacae:0.0370584596)86.1/99:0.0253838139,WP_049008289.1-Enterobacter-cloacae:0.0416774447)84.6/99:0.0459996939)92.3/100:0.0905658163)48.8/100:0.0593569701)76.1/94:0.0211688939,WP_035600513.1-Edwardsiella-tarda:0.2832075111)90.9/93:0.0492220550)73.9/72:0.0196959034,WP_269535377.1-Citrobacter-sedlakii:0.2527472901)88.6/33:0.0451284608,((WP_021540003.1-Escherichia-coli:0.0000024541,WP_047657564.1-Enterobacteriaceae:0.0605371305)99.7/100:0.0983946493,(((WP_105274811.1-Escherichia:0.1778852295,WP_064512195.1-Buttiauxella-gaviniae:0.2930878302)87.6/99:0.0587174507,WP_172410167.1-Escherichia-coli:0.2833838029)64.9/86:0.0611733035,((WP_202782010.1-Enterobacter-asburiae:0.0802418691,WP_144817260.1-Pseudescherichia-sp.-L3:0.0409082126)100/100:0.2348643440,WP_140919905.1-Insectihabitans-xujianqingii:0.2369733075)75.8/34:0.0296873716)77.5/33:0.0235489700)85/32:0.0168168836)76.9/83:0.0142178764)80.5/58:0.0126477988,(((WP_001297344.1-Escherichia-coli:0.1660696181,(((WP_047742849.1-Enterobacter:0.0773326847,WP_040022520.1-Enterobacter-cloacae:0.0754885600)99.3/100:0.1215574729,WP_129995405.1-Enterobacter-cloacae:0.0616715968)76/86:0.0388663053,WP_042099316.1-Escherichia-coli:0.0421426738)95.7/86:0.0894179015)89.6/85:0.0605865923,WP_072664248.1-Escherichia:0.1435572186)87.8/24:0.0459267412,((((WP_119938542.1-Enterobacter-huaxiensis:0.0959526221,WP_050871166.1-Enterobacter-cloacae-complex:0.0392433937)84.4/86:0.0284747096,(WP_167816599.1-Enterobacter-cloacae:0.2582067041,(WP_262183194.1-Enterobacter-mori:0.1327589182,(WP_063412371.1-Enterobacter-cloacae:0.0389041076,WP_047027451.1-Enterobacter-kobei:0.0509518592)98.1/100:0.1176226159)96.1/100:0.1396317927)96.6/99:0.1130777259)84.1/40:0.0231215766,(WP_228937168.1-Escherichia-marmotae:0.0315041276,WP_040079227.1-Escherichia-coli:0.0373432630)86.7/100:0.0160324461)88.7/40:0.0321668116,(WP_100029735.1-Escherichia-coli:0.1248131726,WP_086539147.1-Citrobacter-freundii:0.2874707028)88.6/37:0.0452309578)82/62:0.0159258020)74.6/11:0.0079565118)75.2/12:0.0086546306,(((WP_162648791.1-Citrobacter-freundii:0.1050696896,(WP_060683585.1-Citrobacter-portucalensis:0.1169879206,(WP_079798993.1-Salmonella-enterica:0.0922257997,WP_032255798.1-Enterobacteriaceae:0.0782251512)74.2/90:0.0125807105)94.1/90:0.0721027424)86.4/100:0.0504080407,(WP_195994783.1-Citrobacter-portucalensis:0.2174565493,WP_048218074.1-Citrobacter-freundii-complex:0.1485881722)88/99:0.0597769679)92/88:0.0542236036,WP_061892435.1-Escherichia-coli:0.1448821892)85.6/77:0.0298082446)88.1/22:0.0271150926,(((WP_115348269.1-Hafnia-alvei:0.1429431033,(WP_046458830.1-Obesumbacterium-proteus:0.2586564044,WP_125117341.1-Aeromonas-caviae:0.3306011238)71.6/99:0.0398445105)64.9/45:0.0562936011,(WP_130961887.1-Hafnia-paralvei:0.1805924572,WP_061059890.1-Hafnia-paralvei:0.1716981555)34.6/100:0.0187515086)92.8/100:0.0827476665,((WP_181624464.1-Citrobacter-freundii:0.2015303836,WP_001515521.1-Escherichia-coli:0.1684275843)95/99:0.0736509631,(((((WP_058662478.1-Enterobacteriaceae:0.0435495667,WP_061709645.1-Pseudenterobacter-timonensis:0.0454609917)99.7/100:0.1853939242,WP_112164393.1-Rahnella-inusitata:0.1518990633)90.7/94:0.0775718756,WP_047609274.1-Rahnella-aquatilis:0.1535151553)61/73:0.0182997042,(WP_120162698.1-Rahnella-variigena:0.0556399986,WP_183847446.1-Rahnella-inusitata:0.0644180727)100/100:0.3305147329)44.9/70:0.0326523422,WP_157953065.1-Limnobaculum-parvum:0.2063046061)74.2/93:0.0262081105)79.9/77:0.0420052492)94.3/98:0.0607702330)0/36:0.0000028235,((((((WP_108569539.1-unclassified-Pantoea:0.1922369298,WP_028018520.1-Enterobacter-cloacae-complex:0.6397819257)83.1/100:0.1444812913,WP_250341216.1-Pantoea-eucrina:0.1462678181)86.6/99:0.0982843574,(WP_032659356.1-Enterobacter-hormaechei:0.2796329712,WP_029881813.1-Enterobacter:0.2683782310)53.6/94:0.0375257240)85.4/91:0.0384541897,WP_042093344.1-Escherichia-coli:0.2010560780)0/48:0.0000023038,WP_089706542.1-Escherichia-coli:0.0772139076)60.5/83:0.0424106878,((((WP_170897747.1-Escherichia-coli:0.1300522858,WP_001100781.1-Escherichia-coli:0.1759266095)64.3/91:0.0535494619,WP_001542972.1-Escherichia-coli:0.1265797392)85/96:0.0364172243,(WP_220393175.1-Citrobacter-freundii:0.2653710391,WP_219093375.1-Escherichia-coli:0.1494958500)85.6/96:0.0495492745)53.9/72:0.0315428882,WP_262938691.1-Escherichia-albertii:0.1119060407)29.5/59:0.0056350835)94/81:0.0419717308)81.9/45:0.0097580038,(WP_187216555.1-Escherichia-coli:0.1674917419,WP_059222105.1-Escherichia-albertii:0.1133414615)87/89:0.0401496506)0/36:0.0020848695,((((((((((((((WP_103946175.1-Lelliottia-aquatilis:0.1254858194,WP_123427447.1-Lelliottia:0.0433559576)46.8/95:0.0342751180,WP_235147774.1-Enterobacter-cloacae:0.2538150508)89.7/100:0.0543495518,((WP_214593796.1-Enterobacter-mori:0.0545184930,WP_086532312.1-Enterobacter-ludwigii:0.0607751777)99.1/100:0.1406426368,WP_045339085.1-Enterobacter-cloacae-complex:0.1516964327)64.3/72:0.0348444485)0.3/27:0.0305773967,((WP_096205048.1-Enterobacter:0.0267352715,WP_100165931.1-Enterobacter-mori:0.0360729336)93.9/100:0.0589798635,WP_242172213.1-Enterobacter:0.0937283047)90.5/98:0.0449496334)7.1/21:0.0096658212,WP_126315015.1-Enterobacter-hormaechei:0.0944200026)48/77:0.0179490260,((WP_042716137.1-Enterobacter-sp.-B509:0.0245026243,WP_241820523.1-Enterobacter-kobei:0.0617423020)93.3/100:0.0740349062,WP_172688853.1-Enterobacterales:0.1403448762)42.1/87:0.0186367974)87.1/85:0.0364588007,(((WP_045338355.1-Enterobacter-hormaechei:0.1243380363,(WP_182115563.1-Enterobacter-kobei:0.0328942875,WP_213381159.1-Enterobacter-cloacae:0.0655539448)91.7/100:0.0707901513)63.2/61:0.0507309322,WP_063158410.1-Enterobacter-hormaechei:0.0710490676)69.9/64:0.0147896171,WP_063869545.1-Enterobacter-hormaechei:0.0628142437)64.4/71:0.0188822637)42.6/52:0.0157401447,WP_033487312.1-Enterobacter-cloacae-complex:0.0622941790)91.2/92:0.0481137763,(((WP_193360946.1-Enterobacter-cloacae-complex-sp.-P39RS:0.0572005440,WP_182945676.1-Enterobacter-cloacae:0.0183144974)98.3/100:0.0945193194,WP_265185975.1-Enterobacter-mori:0.0696333801)81.3/91:0.0229944026,((WP_032636062.1-Enterobacter-kobei:0.0659635177,WP_138014015.1-Enterobacter-ludwigii:0.0508318988)71.7/100:0.0375777507,(WP_063137673.1-Enterobacter-cloacae-complex:0.0374856066,WP_165464597.1-Enterobacter-cloacae:0.0528915887)82/100:0.0426886198)82.8/100:0.0343405029)75.1/86:0.0180050948)90.6/88:0.0368080754,((((((WP_181680936.1-Citrobacter-sp.-RHB25-C09:0.1156090355,(WP_090049757.1-Citrobacter-amalonaticus:0.0298411640,WP_196013128.1-Citrobacter-farmeri:0.0308993946)98.4/100:0.1130310128)76.4/100:0.0450022230,WP_103775088.1-Citrobacter-amalonaticus:0.1842987150)96.2/100:0.1335766359,WP_121265736.1-Enterobacter-sp.-R1-2018:0.2067000248)87.4/100:0.0483389184,WP_259047621.1-Klebsiella-sp.-BIGb0407:0.1577116534)30/99:0.0082840909,WP_249031739.1-Tenebrionicola-larvae:0.2640216054)88.3/99:0.0562773953,WP_259000881.1-Scandinavium-goeteborgense:0.1327325920)83.2/95:0.0325272307)77.8/89:0.0136151656,WP_075180894.1-Pantoea-sp.-1.19:0.2496030672)75.2/69:0.0126923373,((((((((WP_127791218.1-Escherichia-coli:0.1290072083,WP_174633339.1-Escherichia-coli:0.1097987541)28.2/91:0.0160477088,WP_025210617.1-Escherichia-coli:0.0941861347)46.9/95:0.0301436560,WP_001527244.1-Escherichia-coli:0.0860815662)75.4/93:0.0235492913,(WP_085454021.1-Escherichia-coli:0.1289288085,(WP_088541377.1-Escherichia-coli:0.0295410774,WP_222691870.1-Escherichia-marmotae:0.0567517599)85.9/100:0.0278193033)17.6/93:0.0233324728)89.5/98:0.0374447007,((((WP_049002539.1-Citrobacter-freundii:0.1166828436,(WP_249576357.1-Enterobacter-vonholyi:0.0608173008,WP_161617217.1-Yokenella-regensburgei:0.0425996704)95.1/100:0.0602808659)82.7/100:0.0279488677,((WP_079834758.1-Salmonella-enterica:0.0014124296,(WP_079793206.1-Salmonella-enterica:0.1159416807,WP_001100807.1-Salmonella:0.0749197066)99.4/100:0.1468297986)75.5/100:0.0375284628,WP_058109100.1-Salmonella-enterica:0.0311810444)99.7/100:0.0923735616)92.9/100:0.0437507069,WP_032303308.1-Escherichia:0.0552178121)91.6/100:0.0600559255,WP_253383865.1-unidentified-bacterial-endosymbiont:0.3267036596)80.1/91:0.0195885029)76.1/88:0.0156825169,((((((((WP_208464084.1-Escherichia-coli:0.1562095157,WP_096122580.1-Escherichia-coli:0.0682259470)69.6/80:0.0055778424,(WP_095374511.1-Escherichia-coli:0.0123977615,WP_012906377.1-Citrobacter-rodentium:0.0873870919)99.1/100:0.0803479006)87.5/80:0.0275819476,WP_162379936.1-Citrobacter-sp.-NCU1:0.1634534753)86.5/99:0.0228919838,((((WP_141109706.1-Escherichia-albertii:0.0940293086,WP_001572827.1-Escherichia-coli:0.0547157692)76.7/100:0.0382517271,WP_203360733.1-Citrobacter-sp.-R56:0.0426581749)88.6/100:0.0465796987,WP_258993080.1-Scandinavium-hiltneri:0.0771766556)95.3/100:0.0650638584,WP_046888360.1-Enterobacter-cloacae:0.0789359711)70.8/87:0.0162203525)82.9/83:0.0148445092,((WP_232904305.1-Buttiauxella-sp.-W03-F01:0.1721813703,WP_259388919.1-Enterobacter-huaxiensis:0.0592934387)82.5/100:0.0359761942,WP_230350752.1-Lelliottia-sp.-WAP21:0.1279535593)91.3/98:0.0506494926)82.5/81:0.0139887885,WP_208283396.1-Enterobacter-sichuanensis:0.1086190195)62.6/20:0.0125745730,(WP_034915863.1-Erwinia-sp.-9145:0.1615464389,(WP_191149682.1-Erwinia-aphidicola:0.1642256289,WP_126546226.1-Enterobacter-quasimori:0.0829589520)79.4/100:0.0322683637)82.8/87:0.0256790465)77.2/26:0.0218283054,WP_105963664.1-Escherichia-coli:0.0667340569)73.7/26:0.0072793938)84.2/24:0.0125863378,(((WP_063610377.1-Escherichia-coli:0.0820587497,WP_001100798.1-Enterobacteriaceae:0.0948427398)83.7/96:0.0353930485,WP_218070502.1-Escherichia-coli:0.0607984097)89.5/83:0.0371954876,(WP_239609625.1-Lelliottia-amnigena:0.1493521508,WP_015386517.1-Cronobacter-sakazakii:0.1593781012)79.1/24:0.0262833277)88.8/22:0.0320585493)75.1/63:0.0076310742,((((WP_175320495.1-Hafnia-paralvei:0.1040913283,WP_130996599.1-Hafnia-paralvei:0.1586216776)97.7/100:0.1097130801,((((WP_171269898.1-Aeromonas-media:0.1109378007,WP_216983019.1-Aeromonas-sp.-FDAARGOS-1415:0.0734975169)98.5/100:0.1474372788,WP_168366848.1-Dickeya-zeae:0.1095564761)84.1/79:0.0433673456,(WP_227095090.1-Aeromonas-veronii:0.0848864956,WP_152109808.1-Plesiomonas-shigelloides:0.0711597152)98.4/100:0.1233731737)87/75:0.0332844867,((WP_064338398.1-Aeromonas-veronii:0.1620493994,(((WP_042864139.1-Aeromonas-caviae:0.0422822099,WP_225159881.1-Aeromonas-caviae:0.0397162482)96.5/100:0.0718969906,(WP_252995952.1-Aeromonas-encheleia:0.2418009502,(WP_244056308.1-Aeromonas-caviae:0.1162243995,WP_202046159.1-Aeromonas-media:0.1321957533)93.4/88:0.0748005539)81/76:0.0355037994)80.7/72:0.0331211515,(WP_045790994.1-Aeromonas-hydrophila:0.1594524127,(((((((WP_247163170.1-Escherichia-coli:0.0189252953,WP_038644959.1-Pantoea:0.0806105762)98.8/100:0.0931127185,(WP_252422582.1-Aeromonas-taiwanensis:0.0198609670,WP_263719954.1-Aeromonas-caviae:0.0893970682)77.8/84:0.0434774807)90.3/99:0.0409568636,WP_033131297.1-Aeromonas-aquatica:0.0227790323)95.6/92:0.0452550547,WP_241344952.1-Aeromonas-sp.-MR16:0.0131553390)88.6/91:0.0278966823,WP_160838676.1-Aeromonas-caviae:0.0775541253)0/71:0.0009710998,(WP_167569591.1-Aeromonas-veronii:0.1309932411,(WP_116547128.1-Plesiomonas-shigelloides:0.0941302609,(((WP_139496062.1-Escherichia-coli:0.0686228893,WP_215269322.1-Aeromonas-dhakensis:0.0410921221)18.7/95:0.0066977521,WP_201996661.1-Aeromonas-veronii:0.0383630226)63.3/77:0.0050366866,WP_235463836.1-Aeromonas-australiensis:0.0222391710)41.4/75:0.0228588895)31/88:0.0072532564)59.3/68:0.0445048760)76.8/64:0.0393203498,WP_049048056.1-Aeromonas-hydrophila:0.1535931459)96.9/100:0.0850723432)72.1/66:0.0131260411)88.1/86:0.0397502294)53.6/88:0.0160468292,(WP_223932214.1-Aeromonas-caviae:0.0116876227,WP_161506954.1-Aeromonas-media:0.1260587936)99.7/100:0.1442339001)69.3/78:0.0174931727)94.8/99:0.0691280991)90.2/93:0.0407256368,WP_034458444.1-Buttiauxella-noackiae:0.1878699783)90.1/81:0.0461089103,(((((WP_262931739.1-Escherichia-albertii:0.0229649193,WP_160188772.1-Escherichia-coli:0.0577086500)99.6/100:0.1289633533,WP_053290779.1-Escherichia-coli:0.1280017070)75.6/91:0.0155849049,WP_096972986.1-Escherichia-coli:0.0759645267)74.1/85:0.0185333072,(WP_157704420.1-Escherichia-coli:0.0809045077,WP_042200020.1-Enterobacteriaceae:0.0665539361)80.5/99:0.0259554827)85.1/85:0.0226593427,((WP_187209312.1-Escherichia-coli:0.0985284680,(WP_105591414.1-Pantoea-coffeiphila:0.0670497630,WP_123334936.1-Erwinia-sp.-JUb26:0.0329903047)100/100:0.2276412216)62.9/80:0.0148185338,(WP_137653986.1-Escherichia-albertii:0.0356767291,WP_001100785.1-Escherichia-coli:0.0312561242)97.6/100:0.0732177699)69.4/79:0.0206421494)77.1/45:0.0159404066)74/45:0.0068149518)90.7/84:0.0218647836)78.9/63:0.0095748408,(((((WP_109548670.1-Escherichia-coli:0.0381377089,WP_001100790.1-Escherichia:0.0238259844)90.9/100:0.0772577155,WP_097421351.1-Escherichia-coli:0.0965952480)62.7/97:0.0316503534,((WP_146314246.1-Escherichia-coli:0.0389471292,WP_096967326.1-Escherichia-coli:0.0202343896)98.9/100:0.0834941150,WP_181465273.1-Escherichia-fergusonii:0.0641567234)83.5/100:0.0332432490)76/91:0.0143719762,WP_181237856.1-Escherichia-marmotae:0.0722026771)85.3/91:0.0221504209,WP_107179425.1-Escherichia:0.0989978749)0/70:0.0010326751)89/77:0.0233450709,WP_021577651.1-Escherichia-coli:0.1380098220)0/8:0.0000022687)90.5/95:0.0317014526,((((((WP_238085128.1-Pseudescherichia-vulneris:0.0203087951,WP_032652659.1-Enterobacter-cloacae-complex:0.0945092082)41/63:0.0135729723,WP_129363899.1-Enterobacter-cloacae:0.0288603743)78.7/63:0.0163229838,(WP_257661844.1-Citrobacter-portucalensis:0.1209780523,WP_193134185.1-Citrobacter-freundii:0.0807725788)84.7/100:0.0295780449)95.1/77:0.0912538227,(WP_042284276.1-Citrobacter-sedlakii:0.2165570726,(WP_024193259.1-Escherichia-coli:0.1009698547,WP_099433281.1-Citrobacter-amalonaticus:0.0319436137)82.8/100:0.0794519704)92.2/100:0.0811171099)97.8/100:0.1296388370,WP_070133611.1-Candidatus-Erwinia-dacicola:0.2404773389)81/79:0.0333630970,((((WP_167863639.1-Erwinia-rhapontici:0.1418918406,WP_039385357.1-Pantoea-eucrina:0.1858001354)70.5/100:0.0659570884,WP_062742814.1-Erwinia-persicina:0.1837960304)72/90:0.0903165464,WP_118663751.1-Erwinia-persicina:0.0962364438)92.7/98:0.0882653558,WP_153879942.1-Citrobacter-freundii:0.1659748341)46.7/65:0.0334198692)84.1/58:0.0374694880)79.8/69:0.0287822295)87.7/91:0.0259156615,WP_202667543.1-Lelliottia-amnigena:0.2077492219)76.5/90:0.0097145549,((((((((WP_172912938.1-Pantoea-stewartii:0.1357633600,WP_145394131.1-Pantoea-sp.-SJZ147:0.1148799411)97.9/100:0.2218691465,WP_061356323.1-Escherichia-coli:0.6725744069)28.3/97:0.0552767208,WP_153752706.1-unclassified-Citrobacter:0.5396879964)13.1/91:0.0648310324,WP_086868392.1-Kosakonia-pseudosacchari:0.6166933654)90.9/95:0.1033007572,WP_097364011.1-Edwardsiella-tarda:0.3536594967)46.7/85:0.0224776985,((WP_071788415.1-Franconibacter-pulveris:0.4415177131,(((WP_232581009.1-Photobacterium-phosphoreum:0.5070978892,(WP_073514120.1-Vibrio-parahaemolyticus:0.0622649946,WP_032481483.1-Vibrio:0.0137763544)99.8/100:0.5463696393)97/89:0.3433904002,(WP_228338545.1-Enterobacter-hormaechei:0.1420625076,(WP_109845273.1-Enterobacter:0.0488882824,WP_150146763.1-Enterobacter-roggenkampii:0.0418085304)86.2/100:0.0735294909)89.4/100:0.1117967437)75.5/78:0.0730894182,WP_020232007.1-Escherichia-coli:0.2462716534)89.6/91:0.0641191028)94.6/98:0.0935225767,((((((((((((((((((((((((((((WP_051234278.1-Halomonas-halocynthiae:0.6073426809,WP_110648656.1-Salinicola-peritrichatus:0.3137910553)97.6/100:0.2425833380,(WP_081193716.1-Halomonas-sp.-BC1:0.2495290064,WP_035566812.1-Halomonas:0.2040627891)64.5/99:0.0401468618)80.2/88:0.0325682306,((WP_224850395.1-Marinobacter-nauticus:0.1621162233,WP_110630630.1-Salinicola-salarius:0.0740113037)81.5/100:0.0275462400,((WP_193085952.1-Halomonas-sp.-FME65:0.1262202222,((WP_159342670.1-Halomonas-meridiana:0.3034587691,WP_253554573.1-Halomonas-sp.-DN3:0.1763430090)66.2/100:0.0204002928,WP_102589198.1-Halomonas-urumqiensis:0.1037674746)94.1/100:0.0449613094)0/87:0.0000026739,(((WP_079551663.1-Halomonas-subglaciescola:0.0681512229,(WP_195687443.1-Halomonas-sp.-KAO:0.0313359144,(WP_193343673.1-Halomonas-elongata:0.0662123334,WP_064701650.1-Halomonas-caseinilytica:0.0110947753)99.8/100:0.1447823469)92.1/98:0.0339754001)89.6/98:0.0209271599,WP_027961801.1-Halomonas-halodenitrificans:0.0241288368)89.1/97:0.0211825850,((WP_089675781.1-Halomonas-aquamarina:0.0525597390,WP_062362595.1-Halomonas-axialensis:0.0215421578)58.4/28:0.0143668818,(WP_253557393.1-unclassified-Halomonas:0.0467779963,WP_089676106.1-Halomonas-aquamarina:0.0284337255)0/98:0.0000029240)83/28:0.0121450536)95.1/97:0.0406873798)88.2/90:0.0209197379)97.4/100:0.0671698246)85.7/92:0.0323124327,((WP_248234415.1-Marinobacter-bryozoorum:0.1056697713,(WP_228223924.1-Marinobacter-nauticus:0.0569651333,WP_098418752.1-Marinobacter-sp.-LV10MA510-1:0.1275268363)88.4/100:0.0681382483)93.1/100:0.0788919551,(WP_021818206.1-Halomonas-huangheensis:0.1359894187,((WP_017842145.1-Methylotuvimicrobium-buryatense:0.3756835126,WP_073436013.1-Halomonas-cupida:0.0561051024)14/97:0.0240815813,WP_149282777.1-Halomonas-sp.-Y2R2:0.0752280183)94.7/100:0.0842107604)75.9/100:0.0432839040)88.5/94:0.0449888656)90.8/97:0.0402404326,WP_127061154.1-Halomonas-nanhaiensis:0.1504664172)22.9/18:0.0186164967,(WP_096281135.1-Halomonas-sp.-JB37:0.2298534639,(WP_027958907.1-unclassified-Halomonas:0.0361230356,WP_064236373.1-unclassified-Halomonas:0.0565754237)94.9/100:0.1002146847)78.2/95:0.0437448210)91.8/66:0.0498684732,((((((WP_030071911.1-Halomonas-alkaliantarctica:0.0486003561,WP_095603285.1-Halomonas-humidisoli:0.0263725697)79.4/47:0.0194333975,WP_253485683.1-Halomonas-sp.-XH26:0.0294148518)58.9/47:0.0114861691,WP_038485954.1-Halomonas-campaniensis:0.0221603016)98.5/100:0.0790713290,WP_162218282.1-Halomonas-alkaliphila:0.0286536713)95.7/100:0.0464441138,WP_078087027.1-unclassified-Halomonas:0.0256093042)73/100:0.0121171681,WP_096655487.1-Halomonas-nigrificans:0.0371621177)98.4/100:0.1019322556)68.6/38:0.0258005625,(((WP_066801876.1-Moraxella-oblonga:0.1545657898,(WP_197128702.1-Psychrobacter-sp.-NZS113:0.1814036943,WP_201616712.1-Psychrobacter-immobilis:0.1498570013)98.2/100:0.1674032164)93.5/100:0.1735504516,WP_036548095.1-Nitrincola-lacisaponensis:0.3915327591)70.9/68:0.0590794713,((WP_041157945.1-Halomonas-sp.-KHS3:0.0671051047,(WP_120386672.1-Halomonas-alkaliphila:0.1347036162,(WP_141441391.1-Halomonas-titanicae:0.0781185969,WP_226251132.1-unclassified-Halomonas:0.0646608838)46.7/44:0.0120635773)61.9/43:0.0551746326)98.5/99:0.1108596963,WP_228219791.1-Halomonas-meridiana:0.0851236453)92.8/62:0.0733841329)88.9/58:0.0393059645)0/2:0.0195573855,((((WP_158774340.1-Cobetia-sp.-L2A1:0.0730854961,(WP_144727127.1-Cobetia-crustatorum:0.0345665079,WP_088744253.1-Cobetia-sp.-QF-1:0.0620863960)0/26:0.0051044092)89/100:0.0319305929,WP_158774127.1-Cobetia-sp.-L2A1:0.0229001331)77.1/99:0.0115983420,(WP_077373671.1-Cobetia:0.0477696895,((WP_213115235.1-Cobetia-sp.-MC34:0.0359858632,WP_200017180.1-Cobetia-sp.-cqz5-12:0.0614211142)91.7/100:0.0560403150,WP_225346840.1-Cobetia-amphilecti:0.0441677494)83.9/100:0.0311389839)94.2/100:0.0324441229)96.3/99:0.0940207789,(WP_251978361.1-unclassified-Salinicola:0.0757358173,WP_189443129.1-Salinicola-rhizosphaerae:0.1073204549)99.2/100:0.1685885615)95.1/91:0.0959905279)96.2/91:0.1000123259,WP_179929642.1-Halomonas-salicampi:0.3727144685)67.2/98:0.0224153010,WP_254581577.1-Marinobacterium-sedimentorum:0.3895621228)91.4/100:0.0675844742,(WP_015904383.1-Desulforapulum-autotrophicum:0.3026259204,WP_111960646.1-Desulfobacter-hydrogenophilus:0.2821214748)95.3/74:0.1681523091)86.3/70:0.0422308047,(((((((((((((WP_018882239.1-unclassified-Thioalkalivibrio:0.4228676897,WP_125198734.1-Guyparkeria-sp.-SCN-R1:0.1938297775)81.7/100:0.0623423452,WP_250072676.1-Guyparkeria-hydrothermalis:0.2123612119)23.6/99:0.0307831476,(((WP_026296529.1-Thioalkalivibrio-sp.-ALJ2:0.1380852740,WP_019569965.1-Thioalkalivibrio-sp.-ALE11:0.0828942842)84.2/100:0.0256962041,WP_028484075.1-Thioalkalivibrio-sp.-ALE17:0.0435600119)73.9/100:0.0099993815,WP_018169543.1-Thioalkalivibrio-sp.-ALMg9:0.0324610684)96.9/100:0.0984110883)100/100:0.2589046457,WP_092999033.1-Thiohalomonas-denitrificans:0.0673726138)48.3/99:0.0530735336,WP_019018778.1-Halomonas-lutea:0.2913993629)95.2/99:0.1389900495,(WP_125841544.1-Stutzerimonas-stutzeri:0.3558438935,((WP_262731117.1-Alcanivorax-marinus:0.1817490667,(WP_106232428.1-Halomonas-ventosae:0.0812370537,WP_228234339.1-Alcanivorax-marinus:0.1877945616)87.9/100:0.0413271499)79.3/100:0.0253990607,WP_175070852.1-Halomonas-taeanensis:0.1148019310)11.7/95:0.0860390259)66.4/96:0.0800297213)87.5/99:0.0561970955,((WP_146294265.1-Euhalothece-natronophila:0.2356457642,WP_159299499.1-Spiribacter-sp.-SSL99:0.2426221025)90.8/100:0.0919393272,WP_011630054.1-Alkalilimnicola-ehrlichii:0.1817088608)90.6/95:0.0645528733)85.2/96:0.0484747443,WP_127477393.1-Sulfurivermis-fontis:0.3313537319)82.8/96:0.0388988468,WP_028988655.1-Thermithiobacillus-tepidarius:0.2736974539)92.8/100:0.1030287586,WP_069857109.1-Desulfoplanes-formicivorans:0.3455115554)64.8/96:0.0470227478,WP_129645432.1-Steroidobacter-agariperforans:0.5692996683)88.1/93:0.0891773118,WP_156872703.1-Ketobacter-sp.-MCCC-1A13808:0.4055279567)46.1/80:0.0567516033,(((WP_151033206.1-Cellvibrio-sp.-KY-GH-1:0.0790149996,WP_049631514.1-Cellvibrio-sp.-pealriver:0.0792085542)76.1/100:0.0851596158,WP_062059887.1-Cellvibrio-sp.-OA-2007:0.0385426063)99.6/100:0.3009888674,WP_178968040.1-Neptunomonas-phycophila:0.3026959785)96.6/88:0.1839699114)12.3/18:0.0361680010)98.8/93:0.1508756191,((((((((((WP_167868984.1-Shewanella-sp.-WXL01:0.4030854145,(WP_222525605.1-Ferrimonas-balearica:0.1460044904,WP_013344594.1-Ferrimonas-balearica:0.1336264315)82.8/100:0.0941644658)11.6/95:0.0596197740,WP_138986566.1-Neptunomonas-concharum:0.3356253178)81.6/25:0.0359297524,(WP_016959527.1-Enterovibrio-norvegicus:0.0755799729,WP_229491984.1-Enterovibrio-norvegicus:0.1117848971)87.3/99:0.0484144288)91.3/23:0.0560699294,(((((WP_047049276.1-Vibrio-mexicanus:0.1783860969,WP_264874461.1-Vibrio-agarivorans:0.0894679436)75.8/94:0.0226728322,WP_020335761.1-Vibrio-natriegens:0.1103482873)84.3/80:0.0249364325,((((WP_050648448.1-Vibrio-sp.-J2-4:0.1387656506,WP_205390674.1-Vibrio:0.0832031717)75.4/40:0.0166232050,(WP_047874860.1-Photobacterium-aphoticum:0.2422287913,WP_140076235.1-Vibrio-parahaemolyticus:0.1229690134)10/4:0.0141983788)78/14:0.0148207499,(((((WP_167678613.1-Shewanella-aestuarii:0.1453430822,WP_137222113.1-Shewanella-sp.-MEBiC00475:0.0314720817)50.2/93:0.0381688585,WP_188955407.1-Shewanella-ulleungensis:0.0669920759)92.6/95:0.0598959011,(WP_155694061.1-Psychrosphaera-haliotis:0.1322427387,((WP_138577097.1-unclassified-Pseudoalteromonas:0.1009949796,(WP_264390127.1-Pseudoalteromonas-sp.-A3:0.0241739971,(WP_096739047.1-Pseudoalteromonas-atlantica:0.0391349202,WP_248462211.1-Pseudoalteromonas-sp.-2CM37A:0.0190075525)92.3/95:0.0226425071)75.5/86:0.0126435939)86.3/68:0.0303620669,((WP_182693438.1-Pseudoalteromonas-sp.-SR45-5:0.0671950737,WP_256848444.1-Pseudoalteromonas-carrageenovora:0.0181386981)74.5/70:0.0111160157,WP_126005793.1-Pseudoalteromonas-sp.-Xi13:0.0317390331)78.6/49:0.0154691998)92.4/68:0.0521103007)41.1/56:0.0252011291)89/96:0.0525734192,(WP_113818410.1-Psychromonas-sp.-B3M02:0.0625737018,(WP_025564776.1-Psychromonas-sp.-SP041:0.0467338536,WP_166372414.1-Psychromonas-sp.-SA13A:0.0140826696)79.3/100:0.0442397728)92/100:0.0487821046)54/63:0.0164896666,WP_022943097.1-Psychromonas-hadalis:0.1704051939)85.5/79:0.0272449094)12/12:0.0113546518,WP_012143346.1-Shewanella-sediminis:0.2348828768)76.7/39:0.0081991412)87.1/80:0.0180370284,WP_164649367.1-Vibrio-astriarenae:0.0946117875)20.4/81:0.0087436293,((WP_193630602.1-Shewanella-benthica:0.1903487217,WP_159656123.1-Vibrio-atypicus:0.0552012320)89.6/94:0.0566757888,(WP_269599241.1-Salinivibrio-kushneri:0.0670828632,WP_136483218.1-Vibrio-sp.-H11:0.0694816525)93.3/100:0.0701102429)92.7/86:0.0519415855)74.6/28:0.0092790005)27.9/7:0.0046144400,WP_084884043.1-Vibrio-sp.-qd031:0.1413010553)86.3/22:0.0248566222,WP_093326971.1-Thalassotalea-agarivorans:0.1093139556)95.1/36:0.0768359392,(((((((WP_160153838.1-Microbulbifer-sp.-ALW1:0.2776330830,((WP_263600466.1-Microbulbifer-agarilyticus:0.1393304872,WP_078085574.1-Microbulbifer-mangrovi:0.0809504322)70.2/96:0.0120829447,WP_029249715.1-Microbulbifer-agarilyticus:0.1057372216)86.4/96:0.0709577912)95.5/100:0.1187084170,((WP_138624780.1-Pseudoalteromonas-sp.-S4492:0.2220660715,WP_130050482.1-Pseudoalteromonas-sp.-MEBiC-03485:0.1386939581)77.8/100:0.0908765344,(WP_189610147.1-Saccharospirillum-salsuginis:0.2720661789,WP_111640782.1-Marinimicrobium-alkaliphilum:0.3345088075)84.4/100:0.0645043994)92.4/100:0.0941256932)84.2/95:0.0456467274,WP_118961222.1-Colwellia-sp.-RSH04:0.3384459940)42.9/94:0.0214878161,WP_186505417.1-Neptunicella-marina:0.2298759393)87.4/94:0.0517712001,(WP_102798355.1-Bowmanella-denitrificans:0.0780554117,WP_214344498.1-Bowmanella-yangjiangensis:0.0347993230)99.9/100:0.2054049092)40.8/80:0.0389167520,(WP_216048042.1-Aliiglaciecola-lipolytica:0.1833064212,WP_057181266.1-Colwellia:0.2196899587)89.4/99:0.0587727841)83.4/19:0.0337358079,((WP_131095783.1-Alteromonas-sp.-KUL42:0.1035055670,((((((((WP_100972994.1-Alteromonas-sp.-MB-3u-76:0.1053974018,WP_269520955.1-Alteromonas-sp.-BMJM2:0.0450360693)73.7/96:0.0243889580,(((WP_130638173.1-Alteromonas-sp.-KUL49:0.0660659479,WP_115592514.1-Alteromonas-aestuariivivens:0.0470800606)27.3/98:0.0189200959,WP_163085646.1-Alteromonas-profundi:0.0369663845)92.4/20:0.0324774415,(WP_071816068.1-Alteromonas-sp.-V450:0.0933124520,WP_198816918.1-Alteromonas-sp.-IB21:0.0207254464)78.3/100:0.0134402276)80.1/18:0.0185125822)75/17:0.0077396012,(((((((WP_156078883.1-Alteromonas:0.0624161042,WP_014977514.1-Alteromonas-macleodii:0.0248268058)13.2/64:0.0099448319,(WP_044058190.1-Alteromonas-australica:0.0403675904,WP_159580202.1-Alteromonas-sp.-KUL106:0.0199502383)87.6/98:0.0130950553)0/42:0.0000027560,WP_239220085.1-Alteromonas-sp.-MmMcT2-2:0.0266104526)82.8/54:0.0065878188,(WP_063457834.1-Alteromonas-stellipolaris:0.0329054571,(WP_202940767.1-Alteromonas-macleodii:0.0232455203,WP_061095937.1-Alteromonas-macleodii:0.0402151076)79.5/95:0.0115092211)78.5/95:0.0065784977)0/55:0.0000023574,(WP_202262575.1-unclassified-Alteromonas:0.0495870172,WP_187447473.1-Alteromonas-sp.-BZK5:0.0211172647)73.7/64:0.0050322206)92.5/68:0.0281189249,WP_057795858.1-Alteromonas-stellipolaris:0.0373085348)73.4/27:0.0088352479,WP_163106912.1-Alteromonas-genovensis:0.0408907295)77.2/27:0.0084459659)98.5/97:0.0587487925,(WP_070176406.1-Alteromonas-lipolytica:0.0340079523,WP_073325432.1-Marisediminitalea-aggregata:0.0504657115)82.4/100:0.0190843195)87.9/19:0.0345511349,WP_216026947.1-Aestuariibacter-sp.-A3R04:0.0611388610)79.2/19:0.0215568458,((WP_265616768.1-Alteromonas-sp.-ASW11-7:0.0992553672,WP_119499891.1-Alteromonas-sp.-RKMC-009:0.0588976772)5.4/71:0.0232030701,WP_265607972.1-Alteromonas-sp.-ASW11-130:0.0863254283)70.3/42:0.0085765699)42.6/16:0.0255384990,WP_216028209.1-Alteromonas-sp.-C1M14:0.0482760017)83.6/93:0.0390299719,(WP_262993134.1-Alteromonas-sp.-ASW11-19:0.1063129211,((WP_218354448.1-Alteromonas-lipotrueiana:0.0675218183,WP_195811108.1-Salinimonas-marina:0.0673010607)84.6/91:0.0221954772,WP_018984642.1-Salinimonas-chungwhensis:0.0972942390)62.7/90:0.0091961982)83/97:0.0256669689)83.7/97:0.0330658245)85.8/97:0.0473623152,(WP_124026287.1-Alteromonas-sediminis:0.1889547446,WP_100655941.1-Alteromonas-flava:0.1949952111)99.8/100:0.1600645402)37/19:0.0088962685)95.1/100:0.0708907341)58/16:0.0378359328,((WP_142941196.1-Aliikangiella-marina:0.3087929741,(WP_081148784.1-Cognaticolwellia-beringensis:0.0422290023,WP_007634918.1-Paraglaciecola-psychrophila:0.0432750521)99.6/100:0.1974096714)73.9/97:0.0513934235,WP_068546663.1-Thalassotalea-crassostreae:0.2124295942)91.7/96:0.0716315297)56.3/9:0.0303315775,((WP_220103444.1-Neiella-holothuriorum:0.1031363322,WP_087505717.1-Neiella-marina:0.1463303170)98.1/100:0.0946196674,((WP_028862352.1-Psychromonas-aquimarina:0.1877767285,WP_187408568.1-Saccharophagus-sp.-K07:0.1465995898)86.4/99:0.0515682693,(WP_251261538.1-Echinimonas-agarilytica:0.1495049405,(WP_096084688.1-Agaribacterium-haliotis:0.3343879698,WP_188151970.1-Teredinibacter-waterburyi:0.2084923582)94.8/70:0.1245306122)40.4/36:0.0182484193)94/67:0.0728269409)80.8/40:0.0498437436)94.3/37:0.0751849731,(WP_248714967.1-Agarivorans-sp.-TSD2052:0.0580328508,WP_152781274.1-Agarivorans-sp.-B2Z047:0.0863091335)99.3/100:0.1075863757)60.9/20:0.0112099997)98.1/54:0.1385970374,(((((WP_108647113.1-Polynucleobacter-rarus:0.0068694954,WP_084281813.1-Polynucleobacter-kasalickyi:0.0593726107)100/100:0.6269954955,((WP_215371371.1-Polynucleobacter-sp.-MWH-UH24A:0.1849370564,WP_173942040.1-Polynucleobacter-antarcticus:0.1860652768)81.4/100:0.0885254230,((((WP_128113711.1-Polynucleobacter-necessarius:0.1027913211,WP_215385554.1-Polynucleobacter-sp.-AP-Kolm-20A-A1:0.0768693569)95.2/100:0.0779329402,WP_216177372.1-Polynucleobacter-sp.-AP-Feld-500C-C5:0.1714742501)73.4/99:0.0329674531,WP_114639791.1-Polynucleobacter-necessarius:0.1207828581)86.5/71:0.0418409464,((WP_216254756.1-Polynucleobacter-sp.-AP-Melu-500A-A1:0.0571403413,WP_216218975.1-Polynucleobacter-sp.-AP-Latsch-80-C2:0.0420409458)99.8/100:0.0995201999,(((((WP_215319389.1-Polynucleobacter-paneuropaeus:0.0996881795,(WP_215352354.1-Polynucleobacter-sp.-MWH-P3-07-1:0.0472536461,WP_215380622.1-Polynucleobacter-paludilacus:0.0363131799)33.4/100:0.0360337119)96.1/99:0.0607440923,WP_062307935.1-Polynucleobacter-sinensis:0.0544536980)79.4/96:0.0174672340,WP_216197047.1-Polynucleobacter-sp.-UK-Mo-2m-Kol15:0.0713110407)76/96:0.0076632296,((((((WP_215376847.1-Polynucleobacter-asymbioticus:0.0272912063,WP_215328794.1-Polynucleobacter-sp.-AP-Sving-400A-A2:0.0548670243)98.5/100:0.0549146109,WP_173959551.1-Polynucleobacter-arcticus:0.0302125076)92.1/100:0.0246842894,WP_216225867.1-Polynucleobacter-sp.-MG-27-Goln-C1:0.0859757610)45.1/98:0.0064024677,(WP_216168716.1-Polynucleobacter-bastaniensis:0.0805803557,WP_216178894.1-Polynucleobacter-sp.-80A-SIGWE:0.0147703256)85.1/100:0.0160918452)88.9/91:0.0234329815,WP_215299358.1-Polynucleobacter-sp.-AP-Nino-20-G2:0.0926505278)87.6/43:0.0219038080,(WP_216268685.1-Polynucleobacter-sp.-Nonnen-W13:0.0497528607,(WP_215334312.1-Polynucleobacter-sp.-AP-Ainpum-60-G11:0.0460892947,WP_251374783.1-Polynucleobacter-corsicus:0.0358026561)51.9/41:0.0059822365)85.6/68:0.0134135844)52.6/33:0.0063465494)90/97:0.0268232496,WP_076024025.1-Polynucleobacter-sphagniphilus:0.0940429523)76.6/46:0.0121981602)95.4/100:0.0768999201)10.3/40:0.0469445648)89/100:0.1226324310)97.4/100:0.2262103439,(WP_116367876.1-Parahaliea-mediterranea:0.0995059386,WP_148063811.1-Parahaliea-aestuarii:0.0921507980)99.4/100:0.2423951604)98.2/96:0.2476419609,((((WP_262601261.1-Endozoicomonas-euniceicola:0.1057968094,WP_066015073.1-Endozoicomonas-atrinae:0.1264000771)93.2/100:0.0849546686,WP_257287521.1-Endozoicomonas-sp.-SESOKO2:0.1206691625)89.5/100:0.0761978308,(WP_067585802.1-Endozoicomonas-ascidiicola:0.1522116832,(WP_020580625.1-Endozoicomonas-elysicola:0.1045925301,WP_257293945.1-Endozoicomonas-sp.-YOMI1:0.1306531733)91.4/100:0.0797341575)99.2/100:0.2141270344)75.5/94:0.0795966188,WP_116686321.1-Pelagibaculum-spongiae:0.4002080174)94.7/99:0.1321175597)72.4/42:0.0325378145,(((((((((WP_036226274.1-Mesoaciditoga-lauensis:0.1480708378,((((WP_077198159.1-Thermosipho-affectus:0.0714440303,WP_012057290.1-Thermosipho-melanesiensis:0.1758767419)9.5/89:0.0214393849,(WP_127352844.1-Caldicellulosiruptor-changbaiensis:0.0593081460,WP_014043315.1-Caldicellulosiruptor-acetigenus:0.0263318707)100/100:0.3602790807)89.8/100:0.0665877652,((((((WP_117708548.1-Fusobacterium-mortiferum:0.0000028763,(WP_204716698.1-Fusobacterium-mortiferum:0.0217325084,WP_204716813.1-Fusobacterium-mortiferum:0.3644114019)6.6/14:0.0468244303)68.6/14:0.0210456685,WP_115270465.1-Fusobacterium-necrogenes:0.0638518559)96.4/95:0.0763377132,(((WP_130889691.1-Fusobacterium-ulcerans:0.0396498687,((WP_005950923.1-Fusobacterium-varium:0.0313465489,WP_070580525.1-Fusobacterium:0.0537990659)8.1/99:0.0111132094,WP_227372109.1-Fusobacterium-ulcerans:0.0325947444)85.7/100:0.0149955743)76.2/100:0.0094763874,(WP_241635364.1-Fusobacterium-gastrosuis:0.1104981760,(WP_074518138.1-Fusobacterium-necrophorum:0.0715442885,(WP_211495349.1-Fusobacterium-necrophorum:0.1276623411,WP_032890075.1-Fusobacterium-nucleatum:0.5825230625)88.2/83:0.0645500492)93.9/83:0.0688777605)97.4/83:0.0659426889)87.6/82:0.0267816142,WP_261793436.1-Fusobacterium-varium:0.2468627912)91.8/94:0.0535216783)97.3/100:0.1184475337,((WP_147004474.1-Leptotrichia-wadei:0.0245687185,WP_018499460.1-Leptotrichia-wadei:0.0319188263)95.1/100:0.0681152115,(WP_068155536.1-Leptotrichia-sp.-oral-taxon-847:0.0655635392,(WP_099992503.1-Fusobacterium-pseudoperiodonticum:0.1168674061,(WP_211617428.1-Leptotrichia-sp.-oral-taxon-218:0.0136919434,WP_094080762.1-Leptotrichia-sp.-oral-taxon-498:0.0454161221)96.3/100:0.0546115967)22.7/77:0.0331150534)92.2/100:0.0654536713)99.4/100:0.2102028814)87.3/92:0.0779022414,(WP_013388736.1-Ilyobacter-polytropus:0.4286324164,WP_012548442.1-Dictyoglomus-thermophilum:0.2803229959)0/10:0.0371220393)74.4/85:0.0154994282,WP_173194267.1-Paenibacillus-frigoriresistens:0.2794603954)98.6/100:0.0994700216)88.3/92:0.0517003349,WP_047753768.1-Kosmotoga-pacifica:0.1548699679)92.4/92:0.0601188997)95.7/100:0.0955225415,(WP_099829233.1-Mesotoga-sp.-Brook.08.105.5.1:0.1498485270,WP_099830786.1-Mesotoga-sp.-Brook.08.105.5.1:0.1758737266)98.1/100:0.1443227775)84.6/93:0.0635393525,(((((((((((((((((WP_054742706.1-Cellulosilyticum-ruminicola:0.0837170002,(WP_177671917.1-Zhenhengia-yiwuensis:0.0459781284,WP_053982981.1-Niameybacter-massiliensis:0.1062004291)98/100:0.1051441121)94.1/100:0.0606870270,WP_069999280.1-Cellulosilyticum-sp.-I15G10I2:0.2109677441)99.7/100:0.1401569425,((WP_072725426.1-Tepidibacter-thalassicus:0.2213450650,WP_072889493.1-Tepidibacter-formicigenes:0.0513699638)89.8/90:0.0443352117,WP_066825124.1-Clostridium-tepidiprofundi:0.0976838628)87.8/87:0.0254683590)76/83:0.0083093424,((((((WP_094369216.1-Romboutsia-weinsteinii:0.1817042770,(WP_074918830.1-Terrisporobacter-glycolicus:0.0643039957,WP_195929149.1-Clostridium-sp.-1001270J_160509_D11:0.0398547160)86.6/100:0.0269985166)90.3/99:0.0364133038,((WP_092726456.1-Romboutsia-lituseburensis:0.1717663417,WP_258242885.1-Romboutsia-lituseburensis:0.0278809874)98.4/98:0.1000037796,(WP_057560648.1-Paeniclostridium-sordellii:0.0856841504,((WP_057580750.1-Paeniclostridium-sordellii:0.1121008416,WP_057538715.1-Paeniclostridium-sordellii:0.0415218132)82.4/99:0.0144620087,WP_195252007.1-Romboutsia-sp.-1001713B170207_170306_H8:0.0546993286)83.6/99:0.0148652025)32.1/91:0.0134795319)83.9/96:0.0220575567)88.3/97:0.0286538480,(((((WP_029452249.1-Clostridium-algidicarnis:0.0999874059,WP_216183125.1-Clostridium-gasigenes:0.0220393249)76.8/100:0.0399834913,WP_216313757.1-Clostridium-gasigenes:0.0625013493)94/100:0.0511597202,WP_197096717.1-Paeniclostridium-sordellii:0.0870078638)0/15:0.0126834132,WP_066677671.1-Clostridium-septicum:0.0529256089)84.8/49:0.0188370526,((WP_095406848.1-Romboutsia-maritimum:0.1160320610,WP_238903698.1-Clostridium-sp.-YIM-B02506:0.0755928215)67.1/56:0.0059985583,((((WP_195959433.1-Clostridium-tertium:0.0705184393,WP_195429094.1-Clostridium-sp.-D46t1_190503_E9:0.0177359170)48/100:0.0177271913,WP_047001949.1-Clostridium-sp.-C8:0.0580172818)73.4/100:0.0209510822,WP_066890539.1-Clostridium-nigeriense:0.0487730532)74.3/87:0.0065347059,WP_016207773.1-Clostridium-sartagoforme:0.0274834665)94/100:0.0233400910)90.4/82:0.0294084200)87.3/95:0.0184080206)81.2/98:0.0170973122,WP_163195303.1-Clostridium-thermarum:0.1211557306)66.9/97:0.0088502336,(WP_055666468.1-Desnuesiella-massiliensis:0.0756335917,WP_095404508.1-Romboutsia-maritimum:0.3366876695)98.7/94:0.1269574464)66/78:0.0166784596,(((((((WP_055265352.1-Clostridium:0.0270639917,(WP_186859336.1-Clostridium-sp.-NSJ-6:0.0556965522,(WP_055265304.1-Clostridium-disporicum:0.0590951173,(WP_154821366.1-Clostridium-saudiense:0.0080712069,WP_195245504.1-Clostridium-celatum:0.0669885609)86.6/100:0.0256827474)75.5/99:0.0331572807)88.5/99:0.0554442267)89.4/100:0.0684138161,WP_186834444.1-Eubacteriales:0.1223694810)86.8/100:0.0343382650,(WP_066874757.1-Clostridium-mediterraneense:0.1465851571,WP_152890015.1-Clostridium-tarantellae:0.1389615886)84.2/100:0.0627937974)14.2/95:0.0318210097,(WP_198610231.1-Clostridium-perfringens:0.0500072136,((WP_235068992.1-Turicibacter-sp.-TJ11:0.0601421951,((WP_161831114.1-Turicibacter-sp.-TS3:0.0554168304,WP_155223812.1-Turicibacter-sanguinis:0.0248485529)69.9/93:0.0163887458,WP_195933776.1-Turicibacter-sanguinis:0.0252803333)53.9/93:0.0066109007)89.5/97:0.0268375550,WP_212726030.1-Turicibacter-bilis:0.0120626627)99.6/100:0.0937880854)64.5/95:0.0262887634)86.4/100:0.0309547442,WP_055258173.1-Sarcina-ventriculi:0.0787936951)76.1/98:0.0120008587,WP_055069574.1-Clostridium-massiliamazoniense:0.1428236571)92.6/96:0.0315838368,WP_133628539.1-Fonticella-tunisiensis:0.0719321730)0/90:0.0000021225)93.5/87:0.0320540415)88.2/93:0.0131785586,(((((WP_195972826.1-Clostridium-thermobutyricum:0.0818036034,(WP_209795868.1-Clostridium-moniliforme:0.0634584728,WP_194191032.1-Clostridium-chrysemydis:0.1096273528)95.1/99:0.0486614047)88.9/99:0.0226800883,WP_040194942.1-Clostridium-culturomicium:0.0645724223)84.7/97:0.0154084524,(WP_055184115.1-Clostridium-paraputrificum:0.0398953665,(WP_090010800.1-Clostridium-sp.-DSM-8431:0.1101200762,(WP_106059304.1-Clostridium-vincentii:0.1127075722,WP_143316579.1-Clostridium:0.0157793766)94.4/100:0.0391326960)94.2/93:0.0445926847)92.7/90:0.0263887842)78.8/73:0.0141960767,((WP_073337580.1-Clostridium-grantii:0.1366453519,WP_072832886.1-Clostridium-collagenovorans:0.0324301306)82.9/85:0.0225716825,WP_072892386.1-Clostridium-fallax:0.1089084029)82.9/96:0.0169218526)95.3/76:0.0352265797,WP_261829490.1-Clostridium-sp.-TW13:0.2056157534)0/78:0.0003701043)78.2/82:0.0087854685,WP_148566871.1-Acetobacterium-paludosum:0.3983652976)20/67:0.0050965679,(((WP_216288229.1-Clostridium-psychrophilum:0.1396521126,WP_216137569.1-Clostridium-estertheticum:0.0351384487)0/98:0.0010094379,WP_220704225.1-Clostridium-estertheticum:0.0398405432)93.2/100:0.0516121264,WP_220321605.1-Clostridium-tagluense:0.0575777153)99.5/100:0.0990912976)76.9/91:0.0261077781,WP_072990516.1-Clostridium-cavendishii:0.2220296556)64.9/89:0.0413225371,(((((((WP_169105633.1-Clostridium-sp.-SM-530-WT-3G:0.0492203733,WP_077866663.1-Clostridium-saccharobutylicum:0.2144529953)11.9/86:0.0156819426,WP_027636628.1-Clostridium-butyricum:0.0913459667)91.2/96:0.0378292389,(((((WP_261829154.1-Clostridium-sp.-TW13:0.0774232546,(WP_077855754.1-Clostridium-beijerinckii:0.0366722068,WP_041894754.1-Clostridium-beijerinckii:0.0276609622)99.6/100:0.0792332996)67.6/100:0.0085585134,WP_224033465.1-Clostridium-gelidum:0.1006890873)17.6/98:0.0117172066,WP_077848125.1-Clostridium-puniceum:0.1410362457)84.6/95:0.0181762653,(WP_185652834.1-Clostridium-sp.-DJ247:0.0209560123,WP_185652641.1-Clostridium-sp.-DJ247:0.0691495463)93.2/100:0.0285608986)0/31:0.0000026768,((WP_022750218.1-Clostridium-saccharobutylicum:0.0371764520,(WP_015391310.1-Clostridium-saccharoperbutylacetonicum:0.0628270407,(WP_077846315.1-Clostridium-puniceum:0.0482548128,WP_238915641.1-Clostridium-sp.-YIM-B02555:0.0595205193)82.2/100:0.0147721496)90.6/100:0.0277282526)71.9/91:0.0212390779,(WP_226126241.1-Clostridium-algoriphilum:0.1302939394,((WP_154531076.1-Inconstantimicrobium-porci:0.1424422316,WP_009169009.1-Clostridium-sp.-DL-VIII:0.0138842698)63/88:0.0488640445,WP_077856313.1-Clostridium-beijerinckii:0.0049933949)97.5/88:0.1029599966)0/33:0.0000029538)86.1/30:0.0140021279)51.3/36:0.0192033771)80.2/36:0.0195324124,WP_265859166.1-Clostridium-botulinum:0.1143401712)92.1/95:0.0405243134,WP_160359894.1-Clostridium-chromiireducens:0.2678498435)0/23:0.0000029042,(((WP_009169246.1-unclassified-Clostridium:0.1072505126,WP_077843022.1-Clostridium-beijerinckii:0.1396786931)83.6/100:0.0328255629,WP_041896941.1-Clostridium-beijerinckii:0.1785490654)92.5/100:0.0578126687,WP_069680896.1-Clostridium-taeniosporum:0.1988517257)86.7/94:0.0335452942)98.7/100:0.1034303746,WP_216390068.1-Clostridium-sp.-MSJ-8:0.1773192500)90.5/98:0.0612202174)87.1/82:0.0390711241,((WP_079421587.1-Clostridium-oryzae:0.3513631443,(((((WP_227847393.1-Clostridioides-sp.-ZZV15-6598:0.1088654217,WP_227846117.1-Clostridioides-sp.-ES-S-0145-01:0.0294214663)95.1/100:0.0435984886,(WP_227842647.1-unclassified-Clostridioides:0.0322577187,WP_003420982.1-Clostridioides-difficile:0.0880865412)86.3/100:0.0209612627)91.5/100:0.0377638942,WP_038810653.1-Clostridioides-difficile:0.0228787627)99.3/100:0.2051587120,WP_213236459.1-Fusibacter-paucivorans:0.7219740160)74.8/47:0.0272648705,WP_226126924.1-Clostridium-algoriphilum:0.3385810015)82.8/46:0.0382735978)78.6/43:0.0248168609,WP_021359979.1-Clostridioides-difficile:0.3225652574)34.8/73:0.0444783469)70.7/88:0.0272691802,WP_099190526.1-Tepidibacter-mesophilus:0.1986142854)83.8/91:0.0288459295,((WP_112093115.1-Methanothermobacter-tenebrarum:0.2354186530,WP_013413362.1-Methanothermus-fervidus:0.2265889381)95.6/100:0.1441893508,((WP_142413383.1-Hathewaya-massiliensis:0.1646882574,WP_035130905.1-Clostridium-sulfidigenes:0.2405895978)96.7/100:0.1017579870,WP_072903743.1-Hathewaya-proteolytica:0.3010824208)88.5/99:0.0803644498)89/98:0.0378741075)93.2/79:0.0421548459,(((((((((WP_058965344.1-Fournierella-massiliensis:0.2041537039,WP_185752312.1-Butyrivibrio-sp.-AE2032:0.3446582717)86.5/100:0.0730736998,WP_155202396.1-Ruthenibacterium-lactatiformans:0.2078995585)91.1/100:0.0901504473,(WP_186886939.1-Anaerofilum-hominis:0.2023696677,WP_102049807.1-Pygmaiobacter-massiliensis:0.2412682026)89.7/100:0.0862720750)98.1/100:0.1700932476,(WP_266162911.1-Caproicibacterium-sp.-BJN0003:0.1946559063,WP_256135338.1-Bittarella-massiliensis:0.2845180496)87.7/100:0.1148477381)59.2/89:0.0397236973,((((((((WP_072827923.1-Fibrobacter-sp.-UWB12:0.0312405565,WP_073304542.1-Fibrobacter:0.2193982270)16.6/57:0.0191370338,WP_088630837.1-unclassified-Fibrobacter:0.0182524110)100/100:0.1916187926,WP_073188830.1-Fibrobacter-sp.-UWP2:0.0000029485)53.9/63:0.0317200415,WP_073155408.1-Fibrobacter:0.0428995104)71.3/62:0.0334030538,WP_097034884.1-Fibrobacter-sp.-UWT3:0.0429549767)68.8/71:0.0095795954,WP_073056819.1-unclassified-Fibrobacter:0.0141907521)99.6/100:0.2122386836,WP_022475313.1-Huintestinicola-butyrica:0.3024253271)5.6/53:0.0504454055,WP_105204772.1-Neobittarella-massiliensis:0.2109820421)93.6/98:0.0845144968)96.3/90:0.1086210547,(WP_111570784.1-Halanaerobium-saccharolyticum:0.0801589319,(WP_089717478.1-Halanaerobium-congolense:0.0525088092,WP_089862761.1-Halanaerobium-salsuginis:0.0287512760)44.6/77:0.0316084727)100/100:0.2774808902)0/32:0.0038830218,WP_160686059.1-Clostridium-sp.-C2-6-12:0.2731512162)53.9/61:0.0211283038,((((WP_220321633.1-Clostridium-tagluense:0.3354688343,(WP_252130283.1-Clostridium-sp.-SYSU_GA19001:0.2973994384,WP_202749105.1-Clostridium-sp.-YIM-B02515:0.1144757803)93.9/100:0.0867243520)81.4/70:0.0382987356,(((WP_165412483.1-Clostridium-estertheticum:0.0663377606,WP_216302578.1-unclassified-Clostridium:0.0112814580)100/100:0.2062477546,((WP_204598070.1-Clostridium-pascui:0.1400261052,WP_027632043.1-Clostridium-hydrogeniformans:0.1512418020)21.9/81:0.0123411963,WP_216457288.1-Clostridium-simiarum:0.1487532799)69.9/87:0.0316731803)10.6/62:0.0172910713,(((WP_066398085.1-Cytobacillus-eiseniae:0.3084681048,(WP_076307499.1-Paenibacillus-odorifer:0.0614002347,((WP_042140168.1-Paenibacillus-sp.-FSL-P4-0081:0.0119630657,WP_036721118.1-Paenibacillus-sp.-FSL-R7-277:0.0740976226)87.5/100:0.0361391925,WP_074087222.1-Paenibacillus-sp.-P32E:0.0351528212)91.7/100:0.0867678556)97/100:0.1075123913)0.4/45:0.0217100261,(((WP_056034132.1-Paenibacillus-sp.-Leaf72:0.0905852353,WP_197936429.1-Paenibacillus-sp.-GSMTC-2017:0.1527111348)91.2/100:0.0761754017,WP_090798944.1-Paenibacillus-sp.-GP183:0.2261052832)80/60:0.0469014993,WP_262684732.1-Paenibacillus-sp.-WQ-127069:0.1792078170)77.4/57:0.0445293017)87.6/99:0.0529964448,WP_036935152.1-Pseudobacteroides-cellulosolvens:0.2081442395)95.5/100:0.1026042684)86.5/85:0.0430884629)88.9/80:0.0302523143,((WP_104410444.1-Clostridium-algidicarnis:0.0403280992,WP_185162668.1-Clostridium-algidicarnis:0.0667454394)99.9/100:0.1720408736,((WP_090038599.1-Clostridium-frigidicarnis:0.1443038347,WP_209459870.1-Youngiibacter-multivorans:0.2165796475)86.4/100:0.0537200291,(WP_164955553.1-Clostridium-estertheticum:0.1645367924,(WP_125002390.1-Clostridium-tagluense:0.1761715600,(WP_226136064.1-Clostridium-estertheticum:0.0295800936,WP_216146875.1-Clostridium-frigoris:0.0682214585)97.2/100:0.1061122682)50.9/98:0.0459145769)93/98:0.0845351511)71.6/94:0.0343617869)85.6/90:0.0340015678)87.9/69:0.0340277293,((WP_234123145.1-Clostridium-hydrogenum:0.2116860600,((WP_084114600.1-Clostridium-acidisoli:0.0944636757,(WP_015614219.1-Clostridium-pasteurianum:0.0206141447,(WP_010235680.1-Clostridium-arbusti:0.0117233593,WP_003443690.1-Clostridium-pasteurianum:0.0506055462)91.2/100:0.0289870428)93.2/100:0.0360970470)73.7/100:0.0114645882,WP_026884435.1-Clostridium-akagii:0.0968676499)80.8/100:0.0652628461)72.8/77:0.0332181022,WP_216462460.1-Clostridium-bornimense:0.1710642266)30.1/53:0.0279971923)95/82:0.0574456085)76/69:0.0109481383,(((WP_013644504.1-Methanobacterium-lacus:0.1851156169,WP_054834988.1-Methanobrevibacter-arboriphilus:0.2106122866)65/99:0.0547148622,WP_248612240.1-Methanobacterium-alcaliphilum:0.1419536894)51.7/43:0.0178405459,((WP_011406138.1-Methanosphaera:0.1690223903,(WP_112123902.1-Methanosphaera-sp.-BMS:0.1017174242,(WP_069592847.1-Methanosphaera-sp.-WGK6:0.0325863861,WP_095608065.1-Methanosphaera-cuniculi:0.1055157272)98/100:0.0816496701)22.2/99:0.0281233923)96.7/99:0.0709848311,((WP_071907175.1-Methanobacterium-congolense:0.0233550540,WP_209584978.1-Methanobacterium-aggregans:0.0434685033)99.7/100:0.1288952716,(WP_069583666.1-Methanobacterium:0.1746978497,WP_048190917.1-Methanobacterium-sp.-SMA-27:0.1430637954)87.5/97:0.0548138295)75.5/96:0.0267277217)84.1/42:0.0168464445)99.7/97:0.0914246675)81.6/59:0.0241847449)80.9/39:0.0159802065,((((((WP_068555806.1-Thermotalea-metallivorans:0.1392252652,WP_073236503.1-Desulforamulus-putei:0.0999033377)78.3/100:0.0357733525,(WP_035164300.1-Caloranaerobacter-azorensis:0.0550716987,WP_090442301.1-Natronincola-peptidivorans:0.1873381107)44.2/93:0.0329548290)50.8/94:0.0326179766,WP_095133872.1-Anaeromicrobium-sediminis:0.1596190515)74.6/99:0.0277171667,(((WP_088538685.1-Methanobrevibacter-sp.-87.7:0.0661527049,WP_016359256.1-Methanobrevibacter-sp.-AbM4:0.1194335065)95.6/100:0.1104790723,WP_067148181.1-Methanobrevibacter-olleyae:0.1013119896)91.3/100:0.1003047132,WP_116670296.1-Methanobrevibacter-woesei:0.2273055410)98.1/98:0.1208868705)80.3/76:0.0194927939,(((((WP_091684654.1-Anaerocolumna-aminovalerica:0.1956402318,(WP_033168390.1-Clostridium-sp.-KNHs205:0.0975803161,((WP_073588199.1-Anaerocolumna-xylanovorans:0.0330401982,WP_185256524.1-Anaerocolumna-chitinilytica:0.1440023974)83.2/100:0.0494753508,WP_073279502.1-Anaerocolumna-jejuensis:0.1162895462)36.9/100:0.0383217043)99.3/100:0.1689884465)93.9/100:0.0928720100,((((WP_073273890.1-Anaerocolumna-jejuensis:0.1454532007,WP_033167869.1-Clostridium-sp.-KNHs205:0.1674706014)57.5/100:0.0527277509,(WP_010268185.1-Paenibacillus-senegalensis:0.1762988173,(WP_155699987.1-Paenibacillus-psychroresistens:0.0763981141,(WP_202751253.1-Clostridium-sp.-YIM-B02515:0.0620201150,WP_090040610.1-Clostridium-frigidicarnis:0.0291888116)33.4/93:0.0159920793)9.5/56:0.0320757509)56.3/96:0.0588462275)98.9/100:0.1540095302,((WP_249305773.1-Jutongia-hominis:0.2264860932,(WP_055944671.1-unclassified-Clostridium:0.0408597962,WP_216564272.1-Eubacterium-sp.-MSJ-13:0.0299391467)90.9/100:0.0406185038)6.4/65:0.0251386932,WP_022463459.1-Jutongia-huaianensis:0.0700115855)100/100:0.2338637069)72.9/100:0.0280096877,WP_058352784.1-Acetivibrio-ethanolgignens:0.3220033109)97.1/100:0.1290235972)83.6/91:0.0357552804,(((WP_253721803.1-Treponema-sp.-OMZ-787:0.5289380060,WP_005539496.1-Johnsonella-ignava:0.1494149243)85.3/92:0.0655874849,(WP_060930762.1-Lachnoanaerobaculum-saburreum:0.1946757941,(((((((((((((((WP_024837362.1-Lacrimispora:0.1023450055,WP_077612318.1-Clostridium-sp.-Marseille-P2415:0.0380285545)0/96:0.0000029632,(WP_024293365.1-Lachnospiraceae:0.0395779333,WP_092242699.1-Lacrimispora:0.0269128902)94.6/100:0.0394155270)90.5/96:0.0207164440,((WP_124065139.1-Clostridium-sp.-E02:0.0910733053,WP_117415169.1-Clostridium-indicum:0.0502100392)73.8/97:0.0093168098,WP_143321893.1-Clostridium-sp.-HBUAS56010:0.0324206458)86/97:0.0161941629)81.6/99:0.0162323403,WP_138313639.1-Hungatella-hathewayi:0.0197499587)90.2/99:0.0338501280,WP_186424215.1-Lacrimispora-celerecrescens:0.1516624455)84.9/100:0.0273329043,WP_154471253.1-Clostridium-porci:0.1197318871)56.8/96:0.0140433560,WP_002577448.1-Enterocloster:0.0767920256)41.1/86:0.0112009596,WP_178313416.1-Lacrimispora-sp.-210928-DFI.3.58:0.0130349875)81.7/81:0.0161283514,WP_024736308.1-Enterocloster-asparagiformis:0.0522052187)70.3/65:0.0199282612,(WP_045093538.1-Clostridium-sp.-FS41:0.0475391033,WP_025485467.1-Lachnospiraceae:0.0735039031)94.2/97:0.0245416218)0/87:0.0000020048,(((((WP_231062061.1-Lacrimispora-sp.-NSJ-141:0.1070235978,WP_249304640.1-Qiania-dongpingensis:0.0648156183)92.2/100:0.0652805503,((WP_215718428.1-Anaerostipes-hadrus:0.0133676206,WP_075760385.1-Anaerostipes-sp.-494a:0.0562503699)98/100:0.0594349651,((WP_227052204.1-Anaerostipes-caccae:0.0348788582,WP_137330070.1-Anaerostipes-rhamnosivorans:0.0854225074)99.5/100:0.1005796907,WP_006781439.1-Hungatella-hathewayi:0.0903515096)87.2/100:0.0288443155)87.1/100:0.0271518806)59.4/76:0.0047590974,WP_066545911.1-Clostridium-sp.-AT4:0.0551357114)72/73:0.0065292577,WP_118541387.1-unclassified-Clostridium:0.0719753901)76.1/76:0.0072285182,WP_215700061.1-Clostridium-sp.-MCC353:0.0671535622)92.7/93:0.0249710124)20.5/60:0.0288340382,((((WP_087223935.1-Lachnoclostridium-sp.-An14:0.0449038546,WP_101876396.1-Lachnoclostridium-edouardi:0.0543650395)89.3/100:0.0291827051,WP_118243934.1-Clostridium-sp.-AM58-1XD:0.0662661610)73.6/46:0.0092503630,WP_118742831.1-Clostridium-sp.-AM28-20LB:0.0325621258)85.9/46:0.0132392600,WP_008399284.1-unclassified-Clostridium:0.0203497953)0/49:0.0000025084)87.4/94:0.0421540083,(WP_003497689.1-_Clostridium:0.0480132388,WP_125142063.1-Clostridium-transplantifaecale:0.0474515214)83.4/100:0.0182832803)37.6/96:0.0387300983,WP_251497048.1-Otoolea-sp.-DSM-111138:0.1356263553)28.9/81:0.0343411583,WP_118555425.1-Lachnotalea-sp.-AF33-28:0.2178109469)82.5/98:0.0744715253)93.5/100:0.0796553537)76.6/91:0.0577935582,((((((((((((WP_035764434.1-Butyrivibrio-sp.-NC2002:0.0951500848,(((WP_026526812.1-Butyrivibrio-sp.-VCD2006:0.1587676329,WP_026491547.1-unclassified-Butyrivibrio:0.0177158935)55.7/100:0.0369760196,WP_044913869.1-Butyrivibrio-sp.-WCE2006:0.0247691323)93.2/100:0.0719785526,WP_091822189.1-Butyrivibrio-sp.-ob235:0.0686529457)78.5/99:0.0382049209)98.1/100:0.1083622592,(((((((((WP_022768176.1-unclassified-Butyrivibrio:0.0252029207,WP_022766545.1-Butyrivibrio-sp.-XPD2006:0.0554190614)99.3/100:0.0919376747,WP_034446055.1-Butyrivibrio-sp.-AE2032:0.0797160032)74.2/100:0.0221708408,WP_071175335.1-Butyrivibrio:0.0717008996)88/70:0.0518589747,WP_024867291.1-Butyrivibrio-sp.-FCS014:0.0298566247)71.7/63:0.0371605976,WP_026513668.1-Butyrivibrio-sp.-LB2008:0.0645959126)74.8/99:0.0184019612,WP_026663187.1-Butyrivibrio-proteoclasticus:0.0831233500)85/83:0.0157755724,WP_026652320.1-Butyrivibrio-proteoclasticus:0.0507732556)63.8/28:0.0116139511,WP_013279951.1-Butyrivibrio-proteoclasticus:0.0230004146)21.9/39:0.0234241089,WP_092060841.1-Butyrivibrio-sp.-INlla14:0.0443915750)96.7/99:0.0762219290)97.7/100:0.0969973577,((WP_019160740.1-Blautia:0.0642740579,WP_173738571.1-Blautia-schinkii:0.0254526650)88.6/100:0.0316803710,(WP_158421413.1-Blautia:0.0408401051,(WP_148445023.1-Eubacteriales:0.0287356800,WP_187566223.1-Blautia-faecis:0.0514955279)93.7/100:0.0288366375)70.6/96:0.0147458990)98.2/100:0.0804733547)96.8/97:0.0592271200,WP_178047055.1-Fusicatenibacter-sp.-CLA-AA-H277:0.1068289328)45.8/85:0.0090638756,WP_109624131.1-Murimonas-intestini:0.0794375169)87.6/91:0.0297313043,(WP_230400100.1-Novisyntrophococcus-fermenticellae:0.2081888132,((WP_173751851.1-Blautia-caecimuris:0.0508265510,WP_117997894.1-Blautia-obeum:0.0138602465)99.6/100:0.0829480132,(((WP_158367551.1-Lachnospiraceae:0.0367409368,WP_235247881.1-Faecalicatena-contorta:0.0830002664)74.3/100:0.0398668040,(WP_251390531.1-Mediterraneibacter-sp.-DSM-111629:0.0450020259,WP_073962323.1-_Ruminococcus:0.0275046409)80.8/100:0.0261225768)85.1/99:0.0190489528,WP_070089756.1-Merdimonas-faecis:0.0338708605)80.5/99:0.0132586847)73.7/88:0.0077848965)85.1/87:0.0236379115)81.4/98:0.0189415049,WP_132379639.1-Muricomes-intestini:0.1015677032)96.8/100:0.0452034579,WP_087156947.1-Lachnoclostridium-sp.-An181:0.0532223191)81.4/95:0.0200925175,(WP_073994331.1-Parasporobacterium-paucivorans:0.1477080752,(WP_099466994.1-Konateibacter-massiliensis:0.0792886841,WP_151145296.1-Candidatus-Galacturonibacter-soehngenii:0.0530723838)96/100:0.0952501025)35.4/77:0.0589456582)70.7/93:0.0190825251,WP_115483253.1-Anaerosacchariphilus-polymeriproducens:0.1082349742)15.3/83:0.0370450149,WP_158351687.1-Diplocloster-modestus:0.2395119263)35.8/83:0.0487735932,WP_158346166.1-Lachnospiraceae:0.2368889615)98.3/99:0.1194542237)86.8/91:0.0316633247)86.9/98:0.0496277149,WP_226914794.1-Intestinibacter-bartlettii:0.2717674516)89.9/100:0.0976409584,((WP_069974893.1-Geosporobacter-ferrireducens:0.3940781304,(WP_223038819.1-Crassaminicella-profunda:0.0901121893,(WP_148809648.1-Crassaminicella-thermophila:0.0546830535,WP_218283043.1-Crassaminicella-sp.-143-21:0.2993853796)88.3/100:0.0407149139)85.2/100:0.0639543755)91.8/100:0.0816725937,WP_250858284.1-Oceanirhabdus-seepicola:0.1963388098)96.7/100:0.1259126549)89.4/95:0.0604715838)76.5/70:0.0140716619,WP_119970050.1-Clostridium-fermenticellae:0.2907639257)84.8/35:0.0179134149)86/34:0.0178543470,(WP_009529486.1-Peptoanaerobacter-stomatis:0.1563204945,WP_068913445.1-Criibacterium-bergeronii:0.1837349515)99.9/100:0.2381826586)12.3/41:0.0086237795,WP_038266426.1-Peptoclostridium-litorale:0.4607408501)45.8/88:0.0654504061,((WP_138788932.1-Paenibacillus-thermoaerophilus:0.4270735041,WP_078711972.1-Anaerorhabdus-furcosa:0.5143619107)72.8/99:0.0960819288,((((((((((WP_160564586.1-Apibacter-sp.-B2912:0.1104842807,(WP_055424523.1-Apibacter-mensalis:0.0898213517,(WP_160568957.1-unclassified-Apibacter:0.0151895006,WP_166004639.1-Apibacter-sp.-B2966:0.0484128910)98.8/100:0.0909555651)73.1/98:0.0554735830)98.4/100:0.1097510291,(WP_105245553.1-Apibacter-adventoris:0.1929118341,WP_146261858.1-Apibacter-muscae:0.1168776661)82.7/98:0.0451460014)100/100:0.2225401018,(WP_084015419.1-Moheibacter-sediminis:0.3448974357,WP_182042771.1-Moheibacter-lacus:0.2626451993)96.6/100:0.1175305478)79/80:0.0465895337,(WP_068596492.1-Vaginella-massiliensis:0.1650213519,((WP_230477342.1-Faecalibacter-bovis:0.1107863976,WP_125349156.1-Empedobacter-falsenii:0.1012598409)85/97:0.0345770639,(WP_125113420.1-Algoriella-xinjiangensis:0.0856642341,WP_092908829.1-Algoriella-xinjiangensis:0.0349673627)92.7/100:0.0484174453)97.7/97:0.1168599449)97/100:0.1362572381)91.6/98:0.1074113671,(((((WP_014783622.1-Aequorivita-sublithincola:0.5182356971,WP_199156598.1-Pedobacter-sp.-ASV2:0.1003378763)64.4/89:0.0266879190,((WP_230143398.1-unclassified-Pedobacter:0.1200287593,WP_215166022.1-Pedobacter-sp.-ISL-64:0.0887224159)93.6/100:0.0830748764,WP_133260761.1-Pedobacter-changchengzhani:0.1284589011)88.8/97:0.0598816042)87.2/89:0.0417156389,WP_109929147.1-Pedobacter-paludis:0.0385725445)97.1/87:0.0604613638,(WP_123203876.1-Pedobacter-jejuensis:0.0798692451,(WP_188416210.1-Pedobacter-mendelii:0.0315952737,WP_056087060.1-Pedobacter-sp.-Leaf176:0.0543958850)18.9/66:0.0170777684)81.6/99:0.0189001399)55.2/82:0.0241829566,WP_056872627.1-Pedobacter-sp.-Leaf194:0.0748646020)99.9/100:0.3630701839)95/100:0.1368964653,(WP_107809536.1-Elizabethkingia-sp.-YR214:0.1340289222,(WP_219971314.1-Chryseobacterium-sp.-LJ756:0.1470088975,(WP_076395614.1-Chryseobacterium:0.0465098939,WP_088467981.1-Chryseobacterium-sp.-VAUSW3:0.1763412099)91.2/74:0.0539756666)66/74:0.0156111005)99.7/100:0.1963738020)69.2/63:0.0204231760,WP_111126400.1-Mesonia-sp.-K7:0.7418403763)96.8/98:0.1171369145,((((WP_205722928.1-Fulvivirga-lutea:0.4510476136,((((((((((WP_053470876.1-unclassified-Flavobacterium:0.1675691981,WP_056200692.1-Flavobacterium-sp.-Root186:0.0382402575)0/63:0.0000023060,(WP_115889823.1-Flavobacterium-cutihirudinis:0.0457951112,WP_095952323.1-Flavobacterium-sp.-ACN6:0.0388376991)43.3/64:0.0187015146)23.3/100:0.0087330382,WP_198857739.1-Flavobacterium-sp.-IB48:0.0483946226)66.8/90:0.0066129621,(WP_269237486.1-Flavobacterium-sp.-IMCC34775:0.0840115895,WP_163393939.1-Flavobacterium-limi:0.0351977374)66.7/100:0.0075625230)76.2/89:0.0076385383,WP_264205577.1-Flavobacterium-sp.-LS1R49:0.0258834904)88.3/93:0.0167787992,((WP_173968237.1-Flavobacterium-collinsii:0.0780258051,WP_264525312.1-Flavobacterium-sp.-N502536:0.0449307824)91.2/99:0.0323077680,WP_070907856.1-Flavobacterium-tructae:0.0894665566)74.7/93:0.0078728845)22.1/89:0.0068195904,WP_091091738.1-Flavobacterium-gillisiae:0.1105070362)97.5/100:0.0634510505,WP_248907802.1-Flavobacterium-sp.-K5-23:0.1267402590)84.6/99:0.0350074635,(((WP_116761981.1-Flavobacterium-laiguense:0.0453991431,WP_143377960.1-Flavobacterium-sp.-ZT3R18:0.0480851029)98.4/100:0.0749050241,((WP_144071503.1-Flavobacterium-franklandianum:0.0674939190,WP_133277350.1-Flavobacterium-nackdongense:0.0917967706)38.8/97:0.0294643396,WP_194311471.1-Flavobacterium-soyangense:0.1306441860)16.5/95:0.0261471552)84.9/92:0.0327238622,WP_103806521.1-Flavobacterium-alvei:0.0886628932)90.6/99:0.0440956777)96.9/99:0.1307541070,(WP_140000461.1-Flavobacterium-microcysteis:0.2376889887,(((WP_194640005.1-Flavobacterium-sp.-HJJ:0.0979133156,WP_073246022.1-Flavobacterium-flevense:0.0692785626)69.1/100:0.0240308307,WP_140487657.1-Flavobacterium-sp.-GSA192:0.0944016415)52.8/73:0.0381902403,WP_100843428.1-Flavobacterium-sp.-5:0.0742908377)99.2/100:0.1752906337)94.1/100:0.1045276276)95.4/100:0.1455551753)86.9/97:0.0969548545,WP_057952567.1-Salinivirga-cyanobacteriivorans:0.5668817337)75.2/92:0.0396413971,(WP_116872776.1-Terrimonas-sp.:0.4190794688,(WP_004155558.1-Microscilla-marina:0.3541695925,WP_101358295.1-Raineya-orbicola:0.2342100289)99.3/100:0.3197037204)90.9/95:0.0924909807)93.4/99:0.0954188311,((((((WP_233378072.1-Maribellus-sp.-CM-23:0.0638318815,WP_247978154.1-Maribellus-sp.-YY47:0.0772540513)95.7/100:0.1179262782,(WP_167609329.1-Maribellus-sediminis:0.1709508670,(WP_045033348.1-Draconibacterium-sediminis:0.0263480289,WP_163345719.1-Draconibacterium-halophilum:0.0825023990)99.1/100:0.1556264783)93.2/100:0.1057080641)99.8/100:0.3064544013,(WP_106543043.1-Prolixibacter-denitrificans:0.1194214651,(WP_025865917.1-Prolixibacter-bellariivorans:0.0221832554,WP_153632016.1-Prolixibacter-sp.-SD074:0.0582751691)0/97:0.0000025933)99.8/100:0.2179071292)64.4/100:0.0651916888,((WP_159517929.1-Sunxiuqinia-indica:0.1055891281,WP_053182762.1-Sunxiuqinia-dokdonensis:0.1082885225)83.4/100:0.0518566878,WP_133466511.1-Sunxiuqinia-elliptica:0.1589389971)98.8/100:0.1717876470)97.2/99:0.1055712441,(WP_068701711.1-Paludibacter-jiangxiensis:0.3895122952,(((WP_101263396.1-Labilibaculum-filiforme:0.0406814255,WP_245870435.1-Labilibaculum-manganireducens:0.0472185853)96.5/100:0.0825685685,(WP_152734748.1-Marinifilum-sp.-N1E240:0.0800245191,(WP_171595603.1-Marinifilum-caeruleilacunae:0.0649624152,(WP_268189532.1-Marinifilum-sp.-D737:0.0511970794,WP_054720444.1-Marinifilum-fragile:0.0335010271)89.3/100:0.0211473645)86.3/100:0.0237420074)93.5/100:0.0582450236)39.8/100:0.0404274057,WP_127342456.1-Ancylomarina-longa:0.1233090030)99.5/100:0.2201693504)4.4/73:0.0105867996)84.1/81:0.0274062754,((WP_011500232.1-Methanococcoides-burtonii:0.2242162918,WP_135611779.1-Methanococcoides-sp.-AM1:0.1717786393)86.5/100:0.0933356043,((((WP_128503182.1-Methanosarcina-sp.-MSH10X1:0.1513358330,(WP_011033115.1-Methanosarcina-mazei:0.0680994655,WP_048134921.1-Methanosarcina-sp.-1.H.T.1A.1:0.0576390240)61.1/94:0.0318727521)91.5/100:0.0509272281,(WP_048137589.1-Methanosarcina-horonobensis:0.1470724696,((WP_048184631.1-Methanosarcina-siciliae:0.0606893703,WP_048173663.1-Methanosarcina-siciliae:0.0589266134)27.4/99:0.0124197321,WP_011023680.1-Methanosarcina-acetivorans:0.0364333876)96.7/100:0.0932336668)96.2/100:0.0882950725)56.5/61:0.0260847973,WP_048167026.1-Methanosarcina:0.0930946413)82.9/61:0.0327995045,WP_048177163.1-Methanosarcina-sp.-MTP4:0.0814041980)78/91:0.0853280861)99.7/100:0.1978644622)93.2/97:0.0926419869)83.3/96:0.0706282769)89.4/99:0.1173523192,(((WP_085659180.1-Prosthecochloris-sp.-HL-130-GSB:0.4689964299,WP_038563105.1-Draconibacterium-orientale:0.2891650750)95.6/100:0.1643551511,(WP_040596643.1-Prevotella-stercorea:0.1180062032,WP_207153370.1-Prevotella-herbatica:0.0756617225)99.5/100:0.2636622179)76.9/99:0.0698193752,(WP_223927321.1-Prevotella-lacticifex:0.2294284266,WP_009010825.1-Prevotella-sp.-C561:0.1646663768)100/100:0.4177773671)91.8/96:0.1841600571)35.5/94:0.0940773770,(((((WP_013633636.1-Pseudopedobacter-saltans:0.1325903290,WP_154287284.1-Pedobacter-puniceum:0.1581620279)91.3/99:0.0756726518,WP_160842504.1-Pedobacter-sp.-HMF7647:0.0684386089)93.7/99:0.1344244702,(WP_074492796.1-Mucilaginibacter-polytrichastri:0.1971626836,(WP_258137696.1-Mucilaginibacter-sp.-BJC16-A38:0.2085291764,(WP_259068220.1-Mucilaginibacter-sp.-X4EP1:0.0649787410,WP_144912900.1-Mucilaginibacter-frigoritolerans:0.0822229207)72.2/99:0.0624698144)94.1/100:0.1140577363)25.8/86:0.0953304428)98.3/100:0.2119166464,WP_163286637.1-Cryomorpha-ignava:0.5770647422)66.5/100:0.1078853609,WP_130092062.1-Brumimicrobium-glaciale:0.4232011475)91.7/100:0.0955157380)97.2/100:0.1686554101)94.5/99:0.1262985579)65.1/80:0.0616357940)86.8/80:0.0962671233,WP_236098952.1-Dethiosulfovibrio:0.6560909338)88.7/99:0.1282416140,(((((((((((((((GJNO01008138.1-Laxus-oneistus-TSA:1.2551284590,(KAI1709308.1-Ditylenchus-destructor:0.1231023031,KAI1730870.1-Ditylenchus-destructor:0.0290601879)94.9/100:0.1618536935)0/92:0.0000026797,(((VDM95900.1-Thelazia-callipaeda:0.2108723477,(VDO27950.1-Onchocerca-flexuosa:0.1210975424,(VDO31788.1-Brugia-timori:0.1074377668,(XP_003135693.1-Loa-loa:0.0682612202,((VBB33646.1-Acanthocheilonema-viteae:0.0877413365,VDK73328.1-Litomosoides-sigmodontis:0.0498526402)70.5/73:0.0125753715,CAG9536712.1-Cercopithifilaria-johnstoni:0.0280012918)87.9/73:0.0227430678)51.6/63:0.0211649783)82.7/63:0.0311750479)96.8/99:0.1186954557)99.7/100:0.3054913533,(VDD97390.1-Enterobius-vermicularis:0.3634513527,KHN87800.1-Toxocara-canis:0.1658984426)86.4/100:0.0711796043)48.8/84:0.0525813343,VDN56932.1-Dracunculus-medinensis:0.3300200359)88.9/100:0.1264560881)75.6/96:0.0672416189,(GGJS01006545.1-Plectus-murrayi-TSA:0.3998177293,(KAE9552274.1-Halicephalobus-sp.-NKZ332:0.3156672393,TKR72632.1-Steinernema-carpocapsae:0.2144539407)59.9/47:0.0123987533)89.7/59:0.0594880814)96.3/96:0.1079691348,((CAD5228290.1-Bursaphelenchus-okinawaensis:0.0370595424,((KAI6206096.1-Aphelenchoides-besseyi:0.1537774936,KAI6191010.1-Aphelenchoides-bicaudatus:0.0861827250)73/100:0.0433887981,KAI6234710.1-Aphelenchoides-fujianensis:0.1106505451)99.7/100:0.1469450738)59.6/81:0.0221614566,CAD5235627.1-Bursaphelenchus-xylophilus:0.0220969036)99.5/100:0.1471468375)30.7/78:0.0350171294,KAH7695359.1-Aphelenchus-avenae:0.3040218457)98.1/97:0.1286266866,((KHJ99538.1-Oesophagostomum-dentatum:0.0523165050,EPB80542.1-Ancylostoma-ceylanicum:0.0496804903)97.4/100:0.0848599528,((VDO65958.1-Haemonchus-placei:0.1333379636,VDL70484.1-Nippostrongylus-brasiliensis:0.0883876977)84.9/100:0.0251330671,VDO95824.1-Heligmosomoides-polygyrus:0.0756106344)99.3/100:0.1313528868)98.2/100:0.1402114074)67.3/87:0.0318785762,CAI4229259.1-Auanema-sp.-JU1783:0.4209983090)2.4/48:0.0143847993,PAV65180.1-Diploscapter-pachys:0.2479771126)94.2/92:0.1195739285,(CAD6189320.1-Caenorhabditis-auriculariae:0.3460005524,(CAB3400093.1-Caenorhabditis-bovis:0.1915081758,((NP_509046.1-Caenorhabditis-elegans:0.0314988598,XP_003117635.1-Caenorhabditis-remanei:0.0297248275)51.2/100:0.0412322048,CAI5454992.1-Caenorhabditis-angaria:0.0333046720)90.5/100:0.0630328657)94.2/100:0.0982901086)62.7/100:0.0338209909)100/100:0.8643581533,(WP_042281872.1-Candidatus-Protochlamydia-sp.-R18:0.1003255625,WP_068471358.1-Parachlamydia-sp.-C2:0.1773117287)100/100:0.7075383018)64.3/36:0.1992309644,WP_207681674.1-Desulfonema-magnum:0.3977368790)78.9/28:0.1375496501,WP_077329105.1-Virgibacillus-siamensis:0.5011535213)20.9/24:0.0922006104,(WP_161826415.1-Petrachloros-mirabilis:0.5402658926,WP_099324654.1-Candidatus-Kuenenia-stuttgartiensis:0.5281309237)90/78:0.1192359289)72.7/34:0.0418503513,(WP_110520589.1-Herpetosiphon-llansteffanensis:0.8025891372,(WP_239493201.1-Luteitalea-sp.-TBR-22:0.4240785166,WP_058867107.1-Chloracidobacterium-thermophilum:0.3819487343)68.8/90:0.1260828241)86.6/50:0.1745873600)89.2/55:0.0790712052,WP_230840800.1-Gloeobacter-morelensis:0.4553199075)76.3/93:0.0624531968)85.7/45:0.0645912599,((((WP_255135177.1-Gracilimonas-sp.-CAU-1638:0.2916401934,WP_020401568.1-Gracilimonas-tropica:0.2804833520)99.9/100:0.5432085472,WP_008811140.1-Parasutterella-excrementihominis:0.4557339828)90.9/66:0.1495535869,(WP_198243301.1-methane-oxidizing-endosymbiont-of-Gigantopelta-aegis:0.3858693814,WP_062153084.1-Beggiatoa-leptomitoformis:0.4882527144)84/92:0.1196868675)75.8/21:0.0402580165,(((((WP_176021856.1-Stenotrophomonas-sp.-NA06056:0.0521783506,WP_049440708.1-Stenotrophomonas-maltophilia:0.0469932338)95.8/100:0.1279948787,WP_132999175.1-Luteimonas-arsenica:0.2650239196)93.8/98:0.1025334961,WP_078758981.1-Lysobacter-spongiicola:0.2911736835)48.5/94:0.0692871112,(((((((WP_019467068.1-Dyella-japonica:0.2648404256,((((((WP_063674027.1-Dyella-thiooxydans:0.1546321217,WP_253565187.1-Dyella-sp.-Sa:0.0716012440)82/47:0.0277997810,(WP_209622196.1-Frateuria-flava:0.0834287362,WP_091339503.1-Frateuria-terrea:0.0477466633)97.4/100:0.0656687446)77.9/47:0.0246592527,WP_201315287.1-Dyella-sp.-EPa41:0.1407486816)74.9/47:0.0130358192,(WP_188800319.1-Dyella-caseinilytica:0.1663569300,WP_192554051.1-Dyella-acidiphila:0.1354653034)98.9/100:0.1077738241)79.7/91:0.0187256575,((((WP_214557828.1-unclassified-Rhodanobacter:0.0409514144,WP_068095172.1-unclassified-Rhodanobacter:0.0282354591)76/100:0.0225681501,(WP_008437080.1-Rhodanobacter-thiooxydans:0.0507141684,(WP_063107523.1-Rhodanobacter-thiooxydans:0.0378331172,WP_211005527.1-Rhodanobacter-sp.-B2A1Ga4:0.0367122660)33/37:0.0072594360)9.8/37:0.0146015367)89.3/100:0.0365343051,((((WP_077439480.1-Rhodanobacter-sp.-C01:0.1159934773,WP_158752649.1-Dyella-sp.-S184:0.0265634515)37.1/33:0.0132714986,WP_077445946.1-Rhodanobacter-sp.-C05:0.0565838071)44.7/32:0.0184635451,WP_184416890.1-Rhodanobacter-sp.-MP7CTX1:0.0678718809)31.1/92:0.0157338725,WP_072762453.1-Rhodanobacter-sp.-OK091:0.0094051242)82/91:0.0335799593)89/100:0.0423017241,WP_015446789.1-Rhodanobacter-denitrificans:0.0638302491)90.8/100:0.0366424908)74.7/64:0.0295595203,(WP_049621948.1-Frateuria-defendens:0.1192822465,WP_179481164.1-unclassified-Rhodanobacter:0.1187725660)75.1/88:0.0518207573)64.2/63:0.0115084363)69/26:0.0250528733,(((WP_168608661.1-Oleiagrimonas-citrea:0.0600583874,WP_113064358.1-Oleiagrimonas-sp.-MCCC-1A03011:0.0280306208)82.6/100:0.0609639020,WP_043099571.1-Oleiagrimonas-soli:0.0714861653)99.9/100:0.1593923161,((WP_233843762.1-Dyella-sp.-2HG41-7:0.0463850311,WP_126673574.1-Dyella-dinghuensis:0.0850751501)10.3/29:0.0151961466,WP_188793403.1-Dyella-nitratireducens:0.0720000683)92/100:0.0647793911)86.7/96:0.0460344704)16.5/27:0.0243960695,WP_204636097.1-Dyella-kyungheensis:0.1588284581)84.2/90:0.0340135983,(WP_111981026.1-Dyella-jiangningensis:0.1686505826,WP_266183414.1-Dyella-sp.-RBB1W86:0.0809162721)35.6/91:0.0263583975)89.1/95:0.0264414942,((WP_130619635.1-Dyella-amyloliquefaciens:0.0843968689,WP_199044928.1-Dyella-sp.-ASV24:0.0738618220)71.1/89:0.0099423491,(WP_266169930.1-Dyella-sp.-RXD159:0.0808630008,(WP_114846719.1-Dyella-tabacisoli:0.0492838708,WP_090451016.1-Dyella-sp.-OK004:0.0736488197)85.8/100:0.0287250532)38/96:0.0130120403)79.3/95:0.0076209136)72.9/87:0.0101847191,WP_131151171.1-Dyella:0.2199802455)18.4/81:0.0356872722,WP_214570707.1-unclassified-Dyella:0.1148860968)98.1/100:0.1289874642)99.4/100:0.2005413514,((((WP_256181981.1-Cloacibacillus-evryensis:0.2247037812,WP_087368268.1-Cloacibacillus-sp.-An23:0.1060984528)100/100:0.5829441250,((WP_084578244.1-Sporomusa-malonica:0.4656022117,WP_131919244.1-Heliophilum-fasciatum:0.2411429307)73.4/95:0.0386850186,((WP_090940141.1-Pelosinus-propionicus:0.2493211254,WP_240986382.1-Acididesulfobacillus-acetoxydans:0.2653589067)84.9/100:0.0539656396,WP_092486361.1-Desulfoscipio-geothermicus:0.1703057856)89.5/98:0.0505805546)86.1/95:0.0584542657)76.3/78:0.0174872019,((WP_246588317.1-Pelorhabdus-rhamnosifermentans:0.2125524003,(((WP_092074275.1-Dendrosporobacter-quercicolus:0.2903214126,(((WP_054258683.1-Propionispora-sp.-2/2-37:0.1590540552,WP_149735423.1-Propionispora-hippei:0.0835478406)0/43:0.0243748352,WP_091744506.1-Propionispora-vibrioides:0.1505068568)47.4/90:0.0356099201,WP_188400350.1-Sporomusa-sp.-GT1:0.3034111644)65.2/87:0.0379414199)81.9/65:0.0369516604,(((WP_007952170.1-Pelosinus-fermentans:0.1140939866,WP_007939097.1-Pelosinus:0.1153942953)94.2/100:0.0739889597,((WP_132081660.1-Anaerospora-hongkongensis:0.0908178526,WP_132083235.1-Anaerospora-hongkongensis:0.2067003779)97.3/100:0.1127169627,WP_188399481.1-Sporomusa-sp.-GT1:0.3066516962)32.1/81:0.0277429767)63.8/80:0.0544402964,((WP_094603851.1-Sporomusa-silvacetica:0.1378341008,WP_144352952.1-Sporomusa-termitida:0.1373750426)86.9/100:0.0588377646,WP_054258662.1-Propionispora-sp.-2/2-37:0.2454361671)81.1/93:0.0583402895)70.7/81:0.0209139658)85.5/68:0.0362463456,(WP_145090815.1-Sporomusa-sp.-KB1:0.0482252367,WP_093793074.1-Sporomusa-acidovorans:0.0789311648)97.7/100:0.1163480243)75.1/96:0.0372997458)79.4/99:0.0430144351,WP_110954047.1-Massilibacillus-massiliensis:0.2688646114)97.9/100:0.1421168258)84.4/41:0.0415731643,(((((((WP_200343816.1-Halochromatium-glycolicum:0.1777663886,WP_201222000.1-Halochromatium-roseum:0.1330631270)97.8/100:0.1503120862,(WP_200237937.1-Thiohalocapsa-halophila:0.0549454709,WP_058558062.1-Thiohalocapsa-sp.-ML1:0.2669004934)96.2/100:0.1111459691)96.4/100:0.1489613605,(WP_182584206.1-Thiospirillum-jenense:0.4066414913,(WP_200286492.1-Rhabdochromatium-marinum:0.1451201648,WP_009150811.1-Thiorhodovibrio-sp.-970:0.0673903278)94.9/100:0.1260142388)81.3/100:0.1248415486)93.9/100:0.0963477821,((((((WP_200207675.1-Chromatium-weissei:0.2176304669,WP_200154756.1-Chromatium-okenii:0.1035001582)98.1/100:0.1468636705,(WP_200158060.1-Allochromatium-vinosum:0.0665549148,WP_200376120.1-Thiocystis-violacea:0.1183309246)81.6/100:0.0460162263)60.4/88:0.0175569199,(((WP_043749566.1-Imhoffiella-purpurea:0.0800687765,WP_007039504.1-Thiorhodococcus-drewsii:0.0659650725)75.9/100:0.0306419837,((WP_164456073.1-Thiorhodococcus-minor:0.0983974905,WP_200330387.1-Thiocystis-violacea:0.0920098775)78.3/100:0.0223274563,WP_164652045.1-Thiorhodococcus-mannitoliphagus:0.0474644848)94.2/100:0.0677195900)86.8/100:0.0242211894,(WP_201092787.1-Thiocystis-minor:0.0800848833,WP_132978621.1-Thiobaca-trueperi:0.1090399276)81.8/100:0.0214402123)86.2/99:0.0263293524)88.7/94:0.0423815764,(WP_200388819.1-Thiocapsa-imhoffii:0.1745527338,((WP_007191850.1-Thiocapsa-marina:0.0434736808,WP_120796078.1-Thiocapsa-rosea:0.0374324081)68.4/99:0.0068296506,WP_232409912.1-Thiocapsa-bogorovii:0.0226740876)96.8/100:0.0893082723)95.1/100:0.0896977539)94.8/100:0.0930335587,(WP_100918325.1-Candidatus-Thiodictyon-syntrophicum:0.0892000035,WP_020502667.1-Lamprocystis-purpurea:0.1131689039)95.2/100:0.1026152693)49.2/91:0.0630096418,WP_089729519.1-Candidatus-Thiosymbion-oneisti:0.3318415580)87.9/99:0.0620099098)0/22:0.0125305159,WP_018175268.1-unclassified-Thioalkalivibrio:0.4512109901)85.8/98:0.0896619443,WP_101758446.1-Oceanicoccus-sp.-KOV_DT_Chl:0.5109357001)76.7/81:0.0496819996,(WP_141198172.1-Persicimonas-caeni:0.4973283798,WP_141730858.1-Oligoflexus-tunisiensis:0.4476456088)58.5/65:0.0611332963)92.2/89:0.0933288640)82.3/52:0.0438790197)97.4/96:0.1185974453)79.3/26:0.0423286905)90.1/62:0.0870314285,((((((((((((((((((((WP_008983868.1-Alishewanella-agri:1.1630800790,WP_265845290.1-Plesiomonas-shigelloides:0.5349048367)98.6/99:0.4600820110,WP_087653467.1-Klebsiella/Raoultella-group:0.0765881897)50/62:0.0773767619,WP_028357569.1-Brackiella-oedipodis:0.2329099272)82.4/31:0.0469047269,WP_171588903.1-Pelistega-europaea:0.3178813382)71.9/53:0.0289754012,WP_154751639.1-unclassified-Erwinia:0.2485717442)49.6/54:0.0331570118,(((((WP_214236207.1-Rosenbergiella-sp.-S61:0.0339880401,(WP_241647140.1-Rosenbergiella-sp.-JB07:0.1288952531,WP_214195533.1-Rosenbergiella-collisarenosi:0.1068068147)86.1/100:0.0491006976)99.7/99:0.1370445223,WP_087487466.1-Tatumella-citrea:0.0555988751)64.7/86:0.0055281356,WP_212707699.1-Tatumella-sp.-JGM130:0.0513484326)70.4/86:0.0101756077,WP_029991521.1-Tatumella-ptyseos:0.0796861858)77.7/93:0.0175583071,(WP_032441370.1-Klebsiella-pneumoniae:0.1159182970,(WP_015583911.1-Raoultella-ornithinolytica:0.0558157445,WP_265776798.1-Klebsiella-oxytoca:0.1604582276)81.4/100:0.0554684049)90.7/95:0.0525617079)95.5/96:0.0734719388)82.9/51:0.0408697220,(WP_140470594.1-Ewingella-americana:0.2502704502,WP_186921353.1-Klebsiella-quasipneumoniae:0.1686051538)40.5/71:0.0087032865)82/53:0.0334894959,(((WP_024553491.1-Franconibacter-helveticus:0.3110261765,WP_096000273.1-Klebsiella-pneumoniae:0.1480407756)78.5/90:0.0492547186,WP_248464278.1-Pectobacterium-versatile:0.1757614769)79.8/86:0.0312835240,(WP_193937755.1-Klebsiella-grimontii:0.1891422921,(WP_142514871.1-Klebsiella-huaxiensis:0.1459218094,(WP_024360036.1-Klebsiella:0.0473272012,WP_234284225.1-Klebsiella-oxytoca:0.0890005999)90.5/100:0.0456128543)88.7/100:0.0316851809)74/92:0.0107855725)85.5/79:0.0190999743)58.9/25:0.0155135803,((WP_040666911.1-Neisseria-wadsworthii:0.1075336257,WP_135034416.1-Neisseria:0.1553578982)93.5/100:0.0621287733,WP_101506252.1-Erwinia-sp.-B116:0.1589245994)86.3/58:0.0372067010)0/11:0.0000021475,(((((WP_228398973.1-Insectihabitans-xujianqingii:0.0895289131,WP_074823218.1-Pragia-fontium:0.0948819742)98.1/100:0.0821194394,((((WP_136167279.1-Brenneria-sp.-CFCC-11842:0.0460220559,WP_264089117.1-unclassified-Brenneria:0.0997577210)74.7/100:0.0380823041,WP_109053612.1-Brenneria-roseae:0.0924110054)87.3/100:0.0351982415,((((((((WP_180742451.1-Pectobacterium-sp.-CFBP8739:0.1954588232,WP_180777316.1-Pectobacterium-aroidearum:0.0106557228)94.6/93:0.0694910323,WP_039504461.1-Pectobacterium:0.0241579003)70.4/49:0.0129762437,(WP_233968669.1-Pectobacterium-polaris:0.0576474361,WP_048259384.1-Pectobacterium-peruviense:0.0144121594)90.3/84:0.0286405051)77/47:0.0087418213,WP_138255615.1-Pectobacterium:0.0223201717)86.7/46:0.0141530271,(WP_109411993.1-Pectobacterium-polaris:0.0501518526,((WP_249680869.1-Pectobacterium-polaris:0.0467059017,WP_125232508.1-Pectobacterium-versatile:0.0496789781)78.1/100:0.0159158856,WP_005974084.1-Pectobacterium-wasabiae:0.0281263873)87.3/100:0.0161766636)76.6/85:0.0080855758)86.2/40:0.0144889980,((((((((((WP_047710158.1-Pectobacterium-atrosepticum:0.0585085033,WP_198482318.1-Pectobacterium-parmentieri:0.0378288884)14/86:0.0093867152,WP_033071760.1-Pectobacterium-parmentieri:0.0313262745)93.3/99:0.0425444801,WP_226305805.1-Pectobacterium-versatile:0.0206592520)86.3/96:0.0245644025,((WP_039280054.1-Pectobacterium-brasiliense:0.0351618070,WP_039517132.1-Pectobacterium-brasiliense:0.0339441750)86.9/73:0.0224296604,WP_039346077.1-Pectobacterium-fontis:0.0312680453)73.6/54:0.0071707431)92.2/97:0.0278231906,WP_103972204.1-Pectobacterium:0.0161126381)76.8/15:0.0075934646,(WP_205558955.1-Pectobacterium-brasiliense:0.0531187240,WP_151242707.1-Pectobacterium-carotovorum:0.0451084423)73.4/8:0.0061332918)74.7/11:0.0071663244,WP_116186106.1-Pectobacterium-aquaticum:0.0465415187)86.6/95:0.0208090414,WP_119888241.1-Pectobacterium:0.0460254453)70.1/87:0.0137853653,WP_205946867.1-Pectobacterium:0.0651962702)62.4/73:0.0083085277,WP_233976643.1-Pectobacterium-versatile:0.0286981324)0/64:0.0000025419)85.3/37:0.0151724340,WP_107169883.1-Pectobacterium:0.0660400338)91.9/98:0.0297954548,(WP_132452860.1-Samsonia-erythrinae:0.1244830854,WP_130637964.1-Pectobacterium-versatile:0.0566334740)32.5/100:0.0216679261)80.5/99:0.0209370158)95.1/100:0.0779102275,(((((((WP_168640580.1-Dickeya-sp.-CFBP-2040:0.1143600838,(WP_027712673.1-Dickeya-chrysanthemi:0.0446645023,WP_012770533.1-Dickeya:0.0366318651)74.7/100:0.0072761368)75.9/81:0.0071644529,(WP_012883965.1-Dickeya-parazeae:0.0709906369,WP_236306746.1-Dickeya-zeae:0.0379173027)72.8/85:0.0138273378)74.4/96:0.0069276686,((WP_038918362.1-Dickeya:0.0322724716,WP_123251184.1-Dickeya-undicola:0.0490277943)37.6/99:0.0068150340,WP_022632763.1-Dickeya-solani:0.0276700421)88.8/99:0.0225163144)98.2/100:0.0973651919,(WP_148877341.1-Serratia-marcescens:0.0669516220,WP_193158894.1-Serratia-marcescens:0.0515957441)96.3/100:0.0751607197)85.8/100:0.0423896917,WP_062791489.1-Serratia-plymuthica:0.0603274685)87/99:0.0341458998,(WP_121576612.1-Brenneria-alni:0.0961022066,WP_095845203.1-Gibbsiella-quercinecans:0.0802368554)14.8/77:0.0057615912)85.3/97:0.0213144970,((WP_214648782.1-Pectobacterium-carotovorum:0.0368580022,((WP_261846379.1-Pectobacterium-carotovorum:0.0604809584,WP_121590317.1-Brenneria-goodwinii:0.0067307362)94.3/100:0.0299087719,(WP_182099415.1-Pectobacterium-aroidearum:0.0363827374,WP_039482612.1-Pectobacterium:0.0292229975)72/100:0.0066831742)23.5/93:0.0072837294)0/92:0.0000022641,WP_249228983.1-Brenneria-tiliae:0.0530582787)86.1/94:0.0283836764)82.7/94:0.0220961394)90.5/97:0.0443149060)80.6/97:0.0217319304,WP_130971788.1-Hafnia-alvei:0.1395765363)78.1/86:0.0158417774,(((WP_112167140.1-Klebsiella:0.1312603862,WP_200519240.1-Klebsiella-michiganensis:0.0993731842)91.8/100:0.0569370440,WP_137713144.1-Brenneria-rubrifaciens:0.1158945399)78.9/96:0.0219417191,WP_165428828.1-Rouxiella-badensis:0.1138534597)0/84:0.0000024196)86.4/95:0.0154269598,(((WP_131063651.1-Raoultella-planticola:0.0234544272,WP_163980548.1-Raoultella-ornithinolytica:0.0383478055)99/100:0.0865412965,(WP_148250181.1-Raoultella-terrigena:0.0953398272,WP_032686784.1-Raoultella-planticola:0.0317011833)93.3/100:0.0476456082)22.2/78:0.0162019360,(WP_142430937.1-Klebsiella-oxytoca:0.1348501412,(WP_239614184.1-Raoultella-ornithinolytica:0.1075628158,((WP_032699180.1-Raoultella:0.0556243375,WP_228399925.1-Raoultella:0.0329052690)72/99:0.0139344048,WP_004138711.1-Klebsiella:0.0193961406)92.7/100:0.0449003432)69/92:0.0216607990)78.6/88:0.0169961120)45.5/78:0.0110283367)74.9/83:0.0094878403)91.9/44:0.0480891818,((((WP_267333169.1-unclassified-Gilliamella:0.1588414026,WP_267361838.1-Gilliamella-sp.-B2772:0.1462674232)34.9/99:0.0091640676,WP_086361229.1-Gilliamella:0.1338191704)83.5/95:0.0304639272,(((((WP_086319877.1-Gilliamella-apis:0.0183995941,WP_198214961.1-Gilliamella-sp.-M0320:0.0575945551)93/100:0.0302643806,WP_039129858.1-Gilliamella:0.0258174627)72.9/98:0.0096671791,(WP_065737496.1-Gilliamella-apicola:0.0382319798,WP_267331332.1-unclassified-Gilliamella:0.0600027372)98.1/100:0.0572733704)93.2/99:0.0387437792,(WP_065579324.1-Gilliamella-apis:0.0915375113,(WP_065587409.1-Gilliamella-apicola:0.0604850806,WP_065580031.1-Gilliamella-apicola:0.0658135336)53.7/100:0.0240788871)63.5/30:0.0221283843)70/50:0.0155527195,WP_160432071.1-Gilliamella-sp.-Pas-s25:0.1209446110)80.4/93:0.0226096758)52.2/68:0.0189586388,((WP_065574597.1-Gilliamella-apicola:0.0549463763,WP_065569248.1-Gilliamella-apicola:0.0380171682)46.8/99:0.0074081482,WP_065610260.1-Gilliamella-apicola:0.0421197481)93.4/99:0.0500685961)99.9/100:0.1726901610)32.4/81:0.0395972401,(WP_246757812.1-Erwinia-phyllosphaerae:0.1885992451,WP_232759612.1-unclassified-Pantoea:0.1445740123)91.6/99:0.0847514206)76.2/66:0.0108157827,((WP_126816534.1-Enterobacter-mori:0.1173979551,WP_256879438.1-Escherichia-albertii:0.0880909519)69.2/100:0.0558267468,WP_049841125.1-Klebsiella-sp.-RIT-PI-d:0.3467761929)90.1/84:0.0414443362)45.6/13:0.0125018607,(WP_146378871.1-Cronobacter-turicensis:0.0742704137,WP_032180402.1-Escherichia-coli:0.0864439133)99.8/100:0.1894638516)96.1/75:0.0776501665,((WP_253460089.1-Erwinia-persicina:0.2498399189,WP_100256321.1-Reinekea-forsetii:0.5474960171)83.1/27:0.0689347018,((WP_168836617.1-Vibrio-agarilyticus:0.1289858124,(WP_076537303.1-Shewanella-sp.-UCD-KL21:0.0214050894,WP_102531614.1-Shewanella-sp.-10N.286.52.B9:0.0560649749)100/100:0.5263149481)89.3/99:0.0970024371,(WP_137671336.1-Agarivorans-sp.-Toyoura001:0.0547406491,WP_221077378.1-Agarivorans-albus:0.0491400944)93.5/100:0.0886750121)93.6/99:0.0962024539)75.9/21:0.0195246824)82.4/84:0.0296112729,(WP_261844981.1-Amphritea-ceti:0.3250808408,((((((WP_269124033.1-Marinomonas-sp.-15G1-11:0.0281870918,WP_191602238.1-Marinomonas-algicola:0.0905173317)98.9/100:0.1438550201,WP_048693477.1-Catenovulum-maritimum:0.2096527267)89.4/100:0.0723712649,WP_228191388.1-Marinomonas-communis:0.1587876178)0/88:0.0192516910,WP_067216963.1-Marinomonas-gallaica:0.0726927998)82.6/96:0.0267223915,WP_067204681.1-Marinomonas-aquimarina:0.0270859133)84.6/96:0.0219553980,WP_055461762.1-Marinomonas-fungiae:0.0665189785)88.6/99:0.0520507856)94.4/100:0.0906358755)56.4/80:0.0347899291,((WP_145010177.1-Pseudomonas-psychrotolerans:0.0070141468,WP_077172422.1-Pseudomonas-psychrotolerans:0.0573580569)100/100:0.2419550551,WP_084334293.1-Pseudomonas-indica:0.1208347557)92.9/100:0.0891397656)91.6/95:0.0678469692,(((((WP_138320019.1-Thalassotalea-litorea:0.1467440388,WP_136736185.1-Thalassotalea-mangrovi:0.1406734429)99.9/100:0.2722884418,WP_114327629.1-Candidatus-Colwellia-aromaticivorans:0.1307431926)78/100:0.0428485803,WP_101108574.1-Psychromonas-sp.-psych-6C06:0.2431923545)78/98:0.0758298906,(WP_199145406.1-Citrobacter-sp.-ASV29:0.2997664013,WP_148513713.1-Vibrio:0.1953270487)88.7/98:0.0873615454)51.2/50:0.0486638038,(((WP_016794198.1-Vibrio-cyclitrophicus:0.0513050434,WP_102424249.1-Vibrio-cyclitrophicus:0.0425187354)90.6/100:0.0438669714,(WP_261903470.1-Vibrio-fortis:0.0760440484,WP_240207223.1-Vibrio-sp.-CyArs1:0.0488506609)88.7/100:0.0339270577)92.4/100:0.0810994200,(WP_247177500.1-Escherichia-coli:0.4054026576,YP_009985672.1-Escherichia-phage-muut:0.2470866150)99.4/100:0.2971841958)92.1/58:0.0919800897)63.7/37:0.0131499481)80.9/61:0.0290728382,((((((((((WP_094934028.1-Enterobacter-cloacae-complex:0.5424915174,WP_016795256.1-Vibrio-cyclitrophicus:0.1628465593)75.2/80:0.0660953945,WP_039473409.1-Vibrio-sinaloensis:0.3026426543)82.7/70:0.0570091748,(WP_010368546.1-Pseudoalteromonas-piscicida:0.3679268692,WP_086590801.1-Vibrio-parahaemolyticus:0.3416474552)80.2/78:0.0598558828)91.7/92:0.0861664302,(((WP_269408820.1-Pseudoalteromonas-shioyasakiensis:0.1845989324,WP_207379479.1-Shewanella-sedimentimangrovi:0.1908772800)93.6/99:0.0722483101,(WP_061013169.1-Vibrionaceae:0.2177914105,WP_155662549.1-Aliivibrio-fischeri:0.1258461183)89.7/99:0.0841053825)0/28:0.0000021659,WP_171328835.1-Vibrio-splendidus:0.2476501405)79.1/36:0.0363194999)37.5/44:0.0495964011,WP_016537220.1-Cedecea-davisae:0.1855807607)48.2/94:0.0250798279,((WP_057065012.1-Citrobacter:0.5827673162,WP_001100815.1-Escherichia-coli:0.2552941136)92.7/100:0.1656322996,(WP_195504469.1-Citrobacter-farmeri:0.2817018430,WP_064554809.1-Buttiauxella-noackiae:0.2703114089)95.8/100:0.1671054503)12.2/91:0.0550651055)77.3/94:0.0419292481,(WP_240304790.1-Vibrio-campbellii:0.4280276905,WP_004741399.1-Vibrio:0.2240637802)94.1/97:0.1228366187)88.2/98:0.0717534975,WP_216735790.1-Hahella-sp.-HN01:0.3023138150)98.3/99:0.1602824089,((WP_127021653.1-Rheinheimera-mangrovi:0.3518647342,(WP_131912015.1-Celerinatantimonas-diazotrophica:0.1836594223,(((WP_027875186.1-Spongiibacter-marinus:0.1564104705,WP_204995064.1-Spongiibacter-marinus:0.0975034477)82.4/100:0.0456156616,((((WP_184464382.1-Zhongshania-antarctica:0.0484538250,WP_198570926.1-Spongiibacter-nanhainus:0.0903050435)74.3/96:0.0062878323,WP_103683243.1-Marortus-luteolus:0.0335224192)98.8/99:0.1105538455,WP_062382560.1-Zhongshania-aliphaticivorans:0.0572877246)80.4/96:0.0468446232,WP_168451636.1-Spongiibacter-thalassae:0.0462444700)90.1/96:0.0395321877)86.3/96:0.0496757942,((WP_092385961.1-Halopseudomonas-salegens:0.1925161890,WP_036859756.1-Porticoccus-hydrocarbonoclasticus:0.2665251516)81/98:0.0979837233,((WP_076722556.1-Marinobacter-lutaoensis:0.2280432711,(((WP_106695101.1-Marinobacter-shengliensis:0.0637012532,WP_148609564.1-Marinobacter-sp.-BW6:0.1193884859)50/41:0.0103865198,WP_222473125.1-Marinobacter-daepoensis:0.0724542010)43.3/40:0.0286669003,WP_092030923.1-Marinobacter-sp.-DSM-26671:0.1265117611)88.1/100:0.0654807086)81.3/98:0.0519612689,WP_036130839.1-Marinobacter-nitratireducens:0.1370468898)53.8/98:0.0500207564)82/97:0.0551620918)86.8/93:0.0618855831)81.5/96:0.0520192663)62.9/84:0.0524553043,WP_132584255.1-Rheinheimera-sp.-D18:0.3746851132)0/75:0.0022557104)93.3/86:0.0615275397,(WP_150896168.1-Vibrio-fortis:0.2420071607,WP_067584343.1-Endozoicomonas-ascidiicola:0.2512704873)64.7/77:0.0253525030)84.6/82:0.0404419829)16.3/23:0.0125648166,((WP_260491935.1-Pseudoalteromonas-flavipulchra:0.2082141194,(WP_039494249.1-Pseudoalteromonas-flavipulchra:0.0962182909,WP_055022295.1-Pseudoalteromonas-sp.-P1-26:0.0717275733)34.3/93:0.0420159024)76.2/99:0.0289823093,(WP_149602089.1-Pseudoalteromonas-distincta:0.0425877669,WP_248518672.1-Pseudoalteromonas-sp.-2CM32C:0.0268872104)97.2/100:0.1167929684)99.7/99:0.2194528009)99.1/97:0.2295116748)45.7/88:0.0706649561,((((((((WP_133132051.1-Legionella-sp.-W10-070:0.6070761826,(WP_058463758.1-Legionella-cincinnatiensis:0.0241064092,WP_128130294.1-Legionella-sainthelensi:0.0536777825)84.6/97:0.0353430195)0/16:0.0000023802,WP_058513550.1-Legionella-santicrucis:0.1135660505)71.4/16:0.0244478485,(WP_232890831.1-unclassified-Legionella:0.0448522848,(WP_058459148.1-Legionella-bozemanae:0.0264849255,WP_058518697.1-Legionella-parisiensis:0.0511082608)85/100:0.0296104252)99.5/100:0.0982923871)82.8/16:0.0250280486,WP_045097199.1-Legionella-fallonii:0.1260755608)74/16:0.0240066065,WP_031567420.1-Legionella-wadsworthii:0.1967412885)92.2/17:0.0681003618,((((((WP_027223798.1-Legionella-pneumophila:0.0330052428,WP_265717942.1-Legionella-sp.-PATHC032:0.0398051704)92.8/89:0.0309143307,WP_106142999.1-Legionella-pneumophila:0.0375829556)70.9/85:0.0058912621,(WP_061513621.1-Legionella:0.0142231599,WP_027226296.1-Legionella-pneumophila:0.0580189987)99.4/96:0.0674244419)87.4/100:0.0139339969,WP_044499121.1-Legionella-pneumophila:0.0261620841)84.7/100:0.0167227008,WP_027220106.1-Legionella-pneumophila:0.0166247856)82.7/100:0.0204425209,WP_027266778.1-Legionella-pneumophila:0.0342702523)96.6/100:0.0803199385)94.7/17:0.0682554108,(WP_127106294.1-Legionella-sp.-km535:0.0400877434,WP_058474153.1-Legionella-quateirensis:0.0315586315)90.5/100:0.0451920654)0/7:0.0558078246,WP_058494182.1-Legionella-worsleiensis:0.0849036692)100/100:0.3924548816)89.1/47:0.1196816927,(WP_205000141.1-Parashewanella-hymeniacidonis:0.0658137606,WP_129946484.1-Shewanella-sp.-OPT22:0.0492311211)100/100:0.5919701308)93/97:0.1377017942)93.3/50:0.0904045433)76.3/80:0.0457953075,(WP_020332409.1-Vibrio-fluvialis:0.0559104336,WP_096629284.1-Salinivibrio-sp.-YCSC6:0.1801729525)48.1/91:0.0514683366)55.1/56:0.0146972585,WP_102038478.1-Shewanella-sp.-GutCb:0.2202124521)77.2/55:0.0159413820,(((((((((((((WP_255857717.1-Bacterioplanoides-sp.-SCSIO-12839:0.4014040222,((WP_250459773.1-Microbulbifer-sp.-GX-H0434:0.0937367096,WP_152451766.1-Microbulbifer-sp.-THAF38:0.1861166555)93.4/100:0.0910935223,WP_009839692.1-Pseudoalteromonas-tunicata:0.5810188290)81.2/88:0.0401404585)82.1/78:0.0361844351,((WP_085356321.1-Neisseria-dumasiana:0.1621354198,((WP_101133177.1-Neisseria-meningitidis:0.0552265228,WP_005552781.1-Aggregatibacter-actinomycetemcomitans:0.2019926891)36.9/95:0.0197429839,(WP_127202078.1-Neisseria-meningitidis:0.1167059447,WP_237092367.1-Neisseria-sp.-ZJ104:0.1522712929)67.6/100:0.0489187240)90.8/95:0.0827771441)94.8/100:0.1355073583,WP_031579865.1-Ruminobacter-sp.-RM87:0.4643930029)75.7/92:0.0880185993)87.5/95:0.0664683557,(WP_188026786.1-Tolumonas-osonensis:0.0181299190,WP_024870911.1-Tolumonas-lignilytica:0.1310453184)92.7/100:0.0683433327)90.3/18:0.0576251595,(WP_187463805.1-Photobacterium-sp.-BZF1:0.0485123136,(WP_039467548.1-Photobacterium-gaetbulicola:0.0768403567,WP_146157089.1-Photobacterium-rosenbergii:0.1657978245)95.7/96:0.0978441073)86.4/73:0.0470803373)0/6:0.0278901411,WP_047883806.1-Photobacterium-ganghwense:0.1645135051)17.2/8:0.0178548312,WP_105076797.1-Pantoea-ananatis:0.4026711004)82.2/86:0.0608370941,(((WP_076542412.1-Shewanella-sp.-UCD-KL21:0.3759055973,WP_007466453.1-Photobacterium-marinum:0.1429978267)83.6/34:0.1273268173,WP_144135068.1-Kosakonia-radicincitans:0.3067719274)46.3/3:0.0235256608,((((WP_088208149.1-Enterobacter-cancerogenus:0.1127350599,WP_121932958.1-Enterobacter-hormaechei:0.0921841135)91.3/100:0.0550078698,WP_023330866.1-Enterobacter:0.0352634488)99.8/100:0.1695118761,WP_120453320.1-Kalamiella-piersonii:0.1053108399)76.4/22:0.0236460412,((((WP_214216788.1-Rosenbergiella-epipactidis:0.0571126807,WP_241623482.1-Rosenbergiella-epipactidis:0.0231429111)99.1/100:0.1333477180,(WP_192411534.1-Pantoea-sp.-PNT02:0.0624745159,((WP_125311258.1-Pantoea-dispersa:0.0826253238,((WP_150019269.1-unclassified-Pantoea:0.0706601927,WP_140917215.1-Pantoea-deleyi:0.0434762985)43.2/92:0.0063845701,WP_069025179.1-Pantoea-agglomerans:0.0067635865)96.2/94:0.0359333427)82.5/91:0.0242353518,WP_097116658.1-Pantoea-sp.-GL120224-02:0.0000020819)90.3/94:0.0585671886)94.8/98:0.0928208128)84.7/99:0.0450907857,WP_187508308.1-Pantoea-agglomerans:0.1791870676)81.4/27:0.0439678377,(WP_269294270.1-Pantoea-sp.-LMR881:0.0464783968,WP_215846277.1-Pantoea-sp.-EnvD:0.0757473602)94.3/100:0.0546000831)79.7/20:0.0194488776)84.4/11:0.0326258265)90.1/44:0.0667689486)75/21:0.0132678446,((WP_120356561.1-Alginatibacterium-sediminis:0.3413659515,((WP_075989065.1-Vibrio-fluvialis:0.0379497980,WP_044366945.1-Vibrio-fluvialis:0.0342223982)46.5/93:0.0234787971,(WP_076657081.1-unclassified-Vibrio:0.0654756548,WP_239929047.1-Vibrio-kanaloae:0.0351389526)98.9/100:0.1108680891)85.9/100:0.0695055259)46.8/46:0.0552111547,WP_182744595.1-Shewanella-sp.-SR44-4:0.2668853950)79.1/43:0.0479846387)82.7/23:0.0250414081,(WP_020431327.1-Vibrio-fluvialis:0.0876242023,WP_055453311.1-Vibrio-fluvialis:0.0612093662)90/100:0.0627923623)84.7/30:0.0229822689,(((((WP_239941760.1-Vibrio-mediterranei:0.1070338935,(WP_062462511.1-Vibrio-mediterranei:0.1045818426,(WP_221070344.1-Vibrio-alfacsensis:0.0910043582,WP_011189464.1-Desulfotalea-psychrophila:0.4409467620)85.2/95:0.0743132661)43.9/79:0.0199245534)74.2/83:0.0155771202,(((WP_252012797.1-Vibrio-sp.-SCSIO-43140:0.1259741545,WP_252012765.1-Vibrio-sp.-SCSIO-43140:0.1222839398)68.8/63:0.0141216920,(WP_171392669.1-Vibrio:0.0901552719,WP_234495031.1-Vibrio-maritimus:0.1926132842)92.6/91:0.0574821293)72.7/76:0.0210267096,((WP_102532726.1-Vibrio-sp.-10N.286.48.B7:0.0785616639,WP_250617847.1-Vibrio-sp.-S4B1:0.0836733934)76.3/99:0.0335209223,WP_101064017.1-Moritella-sp.-Urea-trap-13:0.2415427715)76.3/19:0.0277832935)76.7/26:0.0360895391)88.9/44:0.0513605870,WP_268679810.1-Vibrio-barjaei:0.1069942908)81/28:0.0478224234,WP_102533425.1-Vibrio-breoganii:0.1500280977)18.1/9:0.0357750497,((((WP_261600599.1-Photobacterium-sp.-TY1-4:0.0839454376,(WP_114765998.1-Vibrio-rhodolitus:0.1208037863,WP_182108830.1-Vibrio-marinisediminis:0.0892025476)93.5/100:0.0867806645)90.5/74:0.0486513649,((WP_170899585.1-Vibrio-sp.-3-2-1:0.1152233537,WP_017221010.1-Moritella-dasanensis:0.1203765806)88.9/100:0.0460998781,WP_107284330.1-Photobacterium-lipolyticum:0.0909443008)94.9/99:0.0588627397)0/17:0.0000028222,WP_159197271.1-Escherichia-coli:0.1213701684)85.1/42:0.0278762968,((((((WP_211647400.1-Photobacterium-sp.-GJ3:0.0451450796,WP_120511428.1-Photobacterium-salinisoli:0.0488714174)91.5/100:0.0573003630,WP_046218796.1-Photobacterium-halotolerans:0.2019306684)86.9/58:0.0464409215,((WP_112404745.1-Vibrio-diazotrophicus:0.0481177045,WP_202661182.1-Vibrio-fluvialis:0.0566855624)96.6/100:0.0763952058,((((WP_263100201.1-Kosakonia-sp.-ML.JS2a:0.0292394513,WP_200550470.1-Kosakonia-sp.-LAM2021:0.0899073806)99.9/100:0.1410819437,(WP_094109047.1-Lonsdalea-iberica:0.0924035787,(WP_026742149.1-Lonsdalea-quercina:0.0348251996,WP_094117837.1-Lonsdalea-britannica:0.0363619148)99/100:0.0945482404)78.4/95:0.0203395208)52.4/31:0.0135332741,(WP_015854950.1-Musicola-paradisiaca:0.0466499407,WP_192457620.1-Musicola-keenii:0.0768272418)100/100:0.2320717626)94.6/100:0.0807998535,(WP_038178862.1-Vibrio-rhizosphaerae:0.1565654499,((WP_006646466.1-Photobacterium-sp.-SKA34:0.1602560794,WP_172533652.1-Vibrio-aestuarianus:0.2805170784)36.4/96:0.0543534329,WP_107187026.1-Photobacterium-kishitanii:0.1632155495)29.5/29:0.0296161507)0/23:0.0157252743)82/53:0.0291341880)81.1/53:0.0236885044)0/74:0.0000027736,WP_124938567.1-Vibrio-viridaestus:0.1461324043)60.7/76:0.0193347055,(((WP_237041393.1-Photobacterium-damselae:0.0985334022,WP_080158029.1-Photobacterium-piscicola:0.1847499087)96.9/96:0.1114681918,(WP_047463293.1-Vibrio-fluvialis:0.1070470311,WP_148560072.1-Vibrio-mimicus:0.1627354413)0/82:0.0334236578)86.4/62:0.0396561202,(WP_123015137.1-Vibrio-zhugei:0.0412208103,WP_077313846.1-Vibrio-palustris:0.0241899795)99.8/100:0.1828154651)67.4/30:0.0196612131)36.9/19:0.0160061412,WP_088133934.1-Vibrio-gazogenes:0.2464913293)55.2/21:0.0186141843)71.8/20:0.0210036799)79.6/17:0.0247049555)75/11:0.0090294539,((WP_261894433.1-Vibrio-porteresiae:0.1953434395,(WP_103882085.1-Vibrio-hangzhouensis:0.0641454514,WP_265676699.1-Vibrio-sp.-ZSDZ65:0.1158311003)81.5/100:0.0316011727)60/92:0.0226873153,WP_117235674.1-Vibrio-maerlii:0.1173148267)91.8/85:0.0423104175)77.9/14:0.0111926626,(((WP_200225058.1-Rahnella-aceris:0.1079842251,WP_195814554.1-Rahnella:0.1083480299)74.2/100:0.0373310193,WP_169403689.1-Rouxiella-aceris:0.0954399388)99.1/100:0.1384910366,WP_022590465.1-Vibrio-nigripulchritudo:0.2648287426)76.3/41:0.0279846012)92.6/40:0.0381469003,(WP_171749982.1-Vibrio-sp.-RE88:0.1000428672,WP_243025268.1-Vibrio-cincinnatiensis:0.5007383850)89.7/74:0.0777208263)39/7:0.0111320538)47.1/11:0.0138809293,((WP_038175370.1-Vibrio-pacinii:0.1416082556,WP_193262143.1-Vibrio-parahaemolyticus:0.2791907050)61.8/34:0.0237339064,(((WP_075764060.1-Photobacterium-proteolyticum:0.1617282581,WP_239411815.1-Photobacterium-sp.-OFAV2-7:0.1295513356)56/99:0.0418113019,((WP_069589493.1-Salinivibrio-sp.-DV:0.0929646134,WP_077662143.1-Salinivibrio-proteolyticus:0.0152968697)91.2/100:0.0481081255,((WP_077659434.1-Salinivibrio-kushneri:0.1284817593,WP_077659452.1-Salinivibrio-kushneri:0.0282738265)87.2/100:0.0301230582,WP_269578450.1-Salinivibrio-kushneri:0.0779379231)37.6/100:0.0221194064)100/100:0.2628128649)77.8/91:0.0348025316,((WP_261923504.1-Shewanella-sp.-NFH-SH190041:0.2430483347,((WP_041156219.1-Vibrio-mytili:0.1525728836,(WP_047480300.1-Vibrio-campbellii:0.0567369634,WP_045494469.1-Vibrio-hyugaensis:0.0445694667)98.2/100:0.1137976431)94.4/100:0.1009623396,WP_102523418.1-Vibrio-tapetis:0.1828500250)36.8/74:0.0443166565)79.8/73:0.0418215241,((WP_197173106.1-Pseudoalteromonas-sp.-NZS11_1:0.1235680084,WP_182748079.1-Pseudoalteromonas-sp.-SR41-1:0.1799009616)79.8/99:0.0470863502,WP_166378370.1-Pseudoalteromonas-sp.-Z9A4:0.1652063956)90.7/98:0.0633556598)85.5/54:0.0404532328)45.7/36:0.0263920832)60/46:0.0223627900)56.6/27:0.0286875912,((((((((((((((((WP_011771697.1-Psychromonas-ingrahamii:0.1238353282,WP_182690147.1-Psychromonas-sp.-SR45-3:0.1193427256)94.3/100:0.1127470831,WP_161680902.1-Proteus:0.3403682856)16.7/86:0.0256778210,WP_014230611.1-Vibrio-sp.-EJY3:0.1481996400)60.8/77:0.0101727384,((((WP_237042700.1-Photobacterium-damselae:0.1118145464,WP_220058598.1-Shewanella-sp.-FJAT-53555:0.2937155668)67.4/86:0.0082178023,(WP_013365656.1-_Enterobacter:0.2086653511,(((WP_258988101.1-Scandinavium-hiltneri:0.0716539092,WP_181620050.1-Enterobacter-sp.-RHBSTW-00994:0.0583289724)95.6/100:0.0678435753,WP_193129600.1-Klebsiella-aerogenes:0.0714180264)92.4/100:0.0601542282,WP_214579582.1-Enterobacter-asburiae:0.0949869773)20/100:0.0209153644)100/100:0.1835264629)81.9/91:0.0328617455,WP_129120576.1-Veronia-nyctiphanis:0.2273226489)12.3/83:0.0299479154,(WP_261874894.1-Vibrio-rarus:0.0998135891,WP_138522257.1-Pseudoalteromonas-galatheae:0.2982587483)92.8/98:0.1031566854)72.8/82:0.0105267354)80.1/95:0.0196171884,(WP_102457123.1-Vibrio-breoganii:0.1440297384,(WP_077647682.1-Salinivibrio-sp.-IB643:0.1173293210,WP_152797327.1-Salinivibrio-sp.-VYel1:0.2286387994)65/97:0.0267766568)83.6/95:0.0234487510)91.3/90:0.0377651040,((((((((((((WP_193255073.1-Vibrio-navarrensis:0.0310510242,WP_258469755.1-Vibrio-alginolyticus:0.1168761318)95/100:0.0685841173,(((WP_162023628.1-Shewanella-sp.-Arc9-LZ:0.0698568811,WP_248997855.1-Shewanella-marinintestina:0.0525368438)88.2/100:0.0482662708,WP_017094106.1-Vibrio-splendidus:0.1105722331)23/66:0.0199206894,((WP_009847952.1-Vibrio:0.1870317266,WP_233894267.1-Vibrio-kanaloae:0.1305802479)65.5/100:0.0357059807,(WP_048661352.1-Vibrio-crassostreae:0.0715085985,WP_114634227.1-Vibrio-splendidus:0.0652835657)92.6/100:0.0955818722)91.6/100:0.0688690329)87.3/100:0.0457126385)82.8/98:0.0234000830,WP_065296243.1-Vibrio-natriegens:0.1328008228)68.6/96:0.0444269829,((((WP_017027407.1-Vibrio-breoganii:0.1487428265,WP_108189654.1-Vibrio-splendidus:0.0973400741)93.9/97:0.0861474183,(WP_239872404.1-Shewanella-insulae:0.1147248335,(WP_239850080.1-Shewanella-sp.-Isolate8:0.0513556027,WP_069440067.1-Pseudoalteromonas-tetraodonis:0.3076414090)89.1/100:0.0604321105)7.6/92:0.0075018150)52.3/95:0.0379400389,(WP_211689658.1-Moritella-sp.-28:0.2990775678,(WP_011716450.1-Shewanella-sp.-ANA-3:0.2789583299,WP_206625521.1-Vibrio-vulnificus:0.0764704285)45.1/98:0.0451859877)77.5/86:0.0583379085)86.2/81:0.0309566639,(WP_106651046.1-Shewanella-sp.-Pdp11:0.1430172839,WP_238097025.1-Vibrio-cincinnatiensis:0.2028665621)0/78:0.0000024353)86.5/93:0.0279935754)90.2/97:0.0469498404,(WP_086585081.1-Vibrio-parahaemolyticus:0.0223633425,WP_217557493.1-Vibrio-metschnikovii:0.0630198387)94.1/99:0.0556583867)84.6/92:0.0307948054,((WP_068331709.1-Photobacterium-jeanii:0.1615169253,(WP_031414678.1-Vibrio-parahaemolyticus:0.1186667404,WP_248468589.1-Shewanella-sp.-1CM18E:0.1630669806)80/99:0.0502401045)39.4/94:0.0409342138,WP_077679840.1-Vibrio-kanaloae:0.0912475444)50.3/95:0.0123452909)79.9/91:0.0186624181,(WP_217542775.1-Vibrio-metschnikovii:0.0525161737,WP_113593871.1-Vibrio:0.0870994654)98.6/99:0.1064974092)24.9/66:0.0177718639,((WP_165312521.1-Vibrio-ziniensis:0.1113670158,((WP_172561824.1-Vibrio-furnissii:0.1176179687,(WP_047110243.1-Vibrio-vulnificus:0.2271901672,WP_102313365.1-Vibrio-cyclitrophicus:0.1510479113)91.9/100:0.0648746909)73.8/91:0.0153669263,(WP_192928291.1-Vibrio-vulnificus:0.0265018650,WP_132698543.1-Vibrio-harveyi-group:0.1155489509)99.8/100:0.1278786879)12.9/65:0.0049704867)78.6/98:0.0202299566,(WP_238819535.1-Vibrio-furnissii:0.1091857120,WP_143680610.1-Vibrio-furnissii:0.1091550042)82.1/96:0.0303027563)88/90:0.0261279836)76.9/82:0.0098785143,((((WP_233983986.1-Vibrio-fluvialis:0.0907630341,((WP_185890410.1-Vibrio-vulnificus:0.1785343862,WP_225486226.1-Vibrio-vulnificus:0.1214897612)70.8/98:0.0226414682,WP_099031861.1-Pseudoalteromonas-sp.-GCY:0.1241568734)51.4/98:0.0394861066)84.3/96:0.0284180440,WP_069533349.1-Vibrio-parahaemolyticus:0.1765674074)0/85:0.0000026668,(((WP_208844179.1-Pseudoalteromonas-xiamenensis:0.1383298424,WP_072616325.1-Vibrio-vulnificus:0.1694427100)75.6/99:0.0417084456,WP_233976434.1-Vibrio:0.1514258733)81.5/97:0.0220927504,WP_229630939.1-Vibrio-coralliilyticus:0.2133570907)76.4/91:0.0129898304)80.4/96:0.0134451065,((((WP_102563398.1-Vibrio-sp.-10N.286.49.B3:0.1092233186,(WP_064489977.1-Vibrio-parahaemolyticus:0.0688242468,WP_201976964.1-Aeromonas-enteropelogenes:0.0959479416)91/100:0.0507347704)92.6/99:0.0520776560,(WP_211915530.1-Vibrio-sp.-B1REV9:0.0964292336,WP_017420706.1-Vibrio-vulnificus:0.1052996709)63.2/96:0.0230325975)85.4/93:0.0339626548,(WP_193167845.1-Vibrio-navarrensis:0.3089980107,WP_144217665.1-Shewanella-algae:0.1772933209)80.1/80:0.0485878895)73.2/71:0.0101369525,((WP_039841338.1-Vibrio-harveyi-group:0.1589745846,(WP_020328057.1-Vibrio-fluvialis:0.1364863543,(((WP_145570128.1-Vibrio-tasmaniensis:0.1409475209,WP_224657569.1-Vibrio-vulnificus:0.1552445531)47.8/21:0.0292194024,WP_102475754.1-Shewanella-sp.-10N.286.52.C2:0.2157753051)37.2/78:0.0282851890,WP_120030433.1-Vibrio-sinensis:0.1069708616)28.5/34:0.0510688260)0/30:0.0000023798)84.1/71:0.0303066473,(((WP_033540599.1-Shewanella:0.1380878057,WP_031815865.1-Vibrio-parahaemolyticus:0.1306667707)55.2/88:0.0378666256,(WP_193250619.1-Vibrio-parahaemolyticus:0.1845094751,WP_258516089.1-Vibrio-alginolyticus:0.0904973301)33.7/80:0.0128854024)89.1/98:0.0449822750,((WP_103194250.1-Vibrio-vulnificus:0.0420238417,WP_158146950.1-Vibrio-fluvialis:0.1197844943)92.1/100:0.0637724940,WP_233927909.1-Vibrio-fluvialis:0.1323455652)93.5/100:0.0625632953)85.2/94:0.0276642102)24.6/48:0.0155234537)91.6/97:0.0356288760)86.3/93:0.0200167195)45.1/83:0.0170622751,(WP_206974914.1-Vibrio-parahaemolyticus:0.1073051150,WP_058428281.1-Pseudoalteromonas:0.1988268737)91.3/93:0.0884359704)77.7/99:0.0121336078,(WP_213972370.1-Vibrio-alginolyticus:0.1605305369,WP_017024716.1-Vibrio-rumoiensis:0.1655585039)89.9/100:0.0484890538)75.2/97:0.0092776859,WP_158107962.1-Vibrio-furnissii:0.1476468459)63.7/96:0.0060215417)83.1/84:0.0184017314,((((WP_138589904.1-Pseudoalteromonas-aurantia:0.1716441609,(WP_218529053.1-Shewanella-sp.-NIFS-20-20:0.1209696173,WP_225483056.1-Vibrio-vulnificus:0.0797429330)88.4/100:0.0522215409)44/98:0.0316949471,WP_012549058.1-Aliivibrio-salmonicida:0.1690758610)85.1/100:0.0555695089,WP_016784063.1-Vibrio:0.0847745751)41.9/88:0.0165528856,WP_140342839.1-Vibrio-parahaemolyticus:0.2191010106)88.6/91:0.0381273332)82.7/76:0.0153896150,((((WP_065600047.1-Aliivibrio-sp.-1S128:0.0771923886,((WP_259522409.1-Shewanella-baltica:0.1238412600,WP_173243873.1-Shewanella-sp.-DC2-4:0.1306113710)98.1/100:0.1267979087,WP_130066657.1-Aliivibrio-finisterrensis:0.1427676401)80.5/95:0.0453185123)94.7/93:0.0762991500,WP_203433235.1-Shewanella-sp.-LZH-2:0.2415424841)0/68:0.0000029905,(WP_017020702.1-Aliivibrio-logei:0.1600028904,((WP_045031875.1-Photobacterium-phosphoreum:0.0243272402,WP_107213458.1-Photobacterium-phosphoreum:0.0357423023)99.2/100:0.1489448367,(WP_172380549.1-Vibrio-sp.-Vb339:0.1388413085,WP_102443327.1-Vibrio-breoganii:0.0720914068)86.5/100:0.0504484815)78.1/88:0.0416035104)84.6/90:0.0432986150)78.4/61:0.0165120217,(WP_102448490.1-Vibrio-splendidus:0.1142279735,WP_005395829.1-Vibrio-diabolicus:0.2386153276)83.6/87:0.0491873357)77.8/74:0.0117724241)73.8/72:0.0072157931,(((((WP_135455718.1-Vibrio-echinoideorum:0.0972431309,WP_048899071.1-Photobacterium-swingsii:0.1570131348)43.7/94:0.0173441654,(WP_161672404.1-Vibrio-toranzoniae:0.0487518794,WP_017040654.1-Vibrio-genomosp.-F10:0.1896329272)88.3/94:0.0537285542)78.3/85:0.0157255047,((WP_137027437.1-Vibrio-kanaloae:0.0254780086,WP_108117598.1-Vibrio-splendidus:0.0734968771)89.5/100:0.0408983633,(WP_017449116.1-Vibrio-parahaemolyticus:0.1540497196,WP_137025093.1-Vibrio-kanaloae:0.1301749429)74.7/59:0.0205379282)90.4/58:0.0386396910)63.2/53:0.0125528990,WP_076646094.1-Vibrio-sp.-10N.222.47.A9:0.1779881840)77.4/79:0.0196214413,(WP_155682326.1-Aliivibrio-fischeri:0.1289201765,(WP_102482947.1-Vibrio-splendidus:0.0936081927,WP_102394203.1-Vibrio-sp.-10N.222.54.B6:0.1231536316)0.4/65:0.0236970319)68.9/80:0.0352485724)82.6/79:0.0170188738)73/84:0.0083885388,(WP_063660043.1-Aliivibrio-fischeri:0.1098134188,((WP_171342182.1-Vibrio-cyclitrophicus:0.1402551509,WP_161972844.1-Enterovibrio-baiacu:0.1334837568)89.3/95:0.0745269259,WP_243579549.1-Vibrio-splendidus:0.1770192640)35.1/61:0.0313250849)93.1/84:0.0640105741)86/66:0.0195362923,(WP_161753058.1-Vibrio-toranzoniae:0.1460212902,WP_139685731.1-Vibrio-tasmaniensis:0.1587311207)38.5/48:0.0240317537)74.1/56:0.0084467809,(((((((((WP_268674123.1-Vibrio-scophthalmi:0.1218487096,WP_102420589.1-Vibrio-cyclitrophicus:0.1197685895)38.5/68:0.0232098664,WP_017085924.1-Vibrio-splendidus:0.1063150590)88.8/71:0.0465355892,(WP_102457030.1-Vibrio-cyclitrophicus:0.1185633433,WP_102298256.1-Vibrio-cyclitrophicus:0.0694233335)87.2/98:0.0304229171)77.4/68:0.0224407897,(((WP_108162474.1-Vibrio-splendidus:0.0287039762,WP_004742905.1-Vibrio-tubiashii:0.0549503351)98.6/100:0.1118662550,(WP_102503160.1-Vibrio-splendidus:0.1822181364,WP_017069285.1-Vibrio-crassostreae:0.0642759437)42.1/82:0.0298060834)85.1/95:0.0459447598,WP_016768872.1-Vibrio-cyclitrophicus:0.0910415466)88.5/63:0.0450533153)87.5/62:0.0277904039,(WP_248368760.1-Vibrio-sp.-1CM8B:0.1354571454,(WP_102385704.1-Vibrio-cyclitrophicus:0.0775671573,WP_239937663.1-Vibrio-chagasii:0.1812090172)92.4/99:0.0650941315)71.1/20:0.0148976331)87.3/69:0.0374725056,((((((((((WP_108083222.1-Vibrio-sp.-ZF-223:0.1278033679,((WP_102532836.1-Vibrio-sp.-10N.286.48.B7:0.1120524059,WP_254101081.1-Alteromonas-sp.-LMIT007:0.1423355845)87.7/100:0.0593787134,WP_121852788.1-Parashewanella-spongiae:0.1932349242)53.7/99:0.0189781926)93.5/96:0.0627938550,WP_259574191.1-Shewanella-baltica:0.2014027638)68.6/42:0.0107764728,((WP_243584123.1-Vibrio-splendidus:0.1286415233,WP_029788868.1-Vibrio:0.2188497425)75.6/97:0.0300706900,(WP_102462022.1-Vibrio-splendidus:0.0591913681,WP_187463684.1-Photobacterium-sp.-BZF1:0.0649213217)99.1/100:0.1124261572)87.6/95:0.0452093283)86/45:0.0270252919,(WP_223418564.1-Shewanella-inventionis:0.1372452278,WP_011497090.1-Shewanella-denitrificans:0.2180343953)91.3/93:0.1005717807)0/60:0.0000023383,(((WP_163929979.1-Paraferrimonas-sp.-SM1919:0.0989665336,(WP_213886549.1-Vibrio-alginolyticus:0.1728965739,WP_158173598.1-Vibrio-alginolyticus:0.0938666016)96.1/100:0.1088771954)88.3/96:0.0560353247,WP_087845049.1-Photobacterium-malacitanum:0.1191951029)12.1/74:0.0150573354,(WP_172855536.1-Vibrio-coralliilyticus:0.2307058525,WP_234235026.1-Shewanella-sp.-AS16:0.1714959215)85.5/84:0.0520648434)77.5/76:0.0305765772)90.9/74:0.0451452471,WP_193154481.1-Vibrio-navarrensis:0.1770766251)0/29:0.0000026258,WP_088905011.1-Shewanella-marisflavi:0.2374106344)83.6/53:0.0252324504,WP_130314193.1-Vibrio-vulnificus:0.1522895606)83.3/82:0.0263566108,(WP_102459739.1-Vibrio-lentus:0.1206835426,WP_102363333.1-Vibrio-cyclitrophicus:0.1281006598)81.7/80:0.0384891723)67.3/62:0.0062369918,WP_102554090.1-Vibrio-splendidus:0.1130664934)88.9/79:0.0334820244)86.7/73:0.0455908564,((WP_063656223.1-Aliivibrio-fischeri:0.1190396717,WP_155675904.1-Aliivibrio-fischeri:0.0754821870)82.9/93:0.0283338664,WP_017105169.1-Vibrio:0.1364621073)84.5/94:0.0298499422)73.4/26:0.0072835604,((((((((((WP_059021773.1-Vibrio-coralliirubri:0.0549704381,WP_135380937.1-Vibrio-tasmaniensis:0.1013594041)87.2/92:0.0370046028,(WP_238130349.1-Vibrio-cincinnatiensis:0.1710089040,WP_102276040.1-Vibrio-lentus:0.0740184715)80.7/90:0.0506496916)84.8/88:0.0334397695,(((WP_005597034.1-Vibrio-scophthalmi:0.1313112091,(WP_108176621.1-Vibrio-splendidus:0.0511342198,WP_239918548.1-Vibrio-sp.-Isolate22:0.0262752254)97/100:0.0684008415)68.1/99:0.0127669524,WP_135445143.1-Vibrio-tasmaniensis:0.1905556390)90/91:0.0380360379,WP_248388778.1-Vibrio-cyclitrophicus:0.1379939987)47.2/84:0.0296525362)49.5/37:0.0196350437,WP_265133188.1-Vibrio-splendidus:0.0993901986)80.6/16:0.0326859789,(((WP_017063118.1-Vibrio:0.0809976106,WP_132940416.1-Vibrio-crassostreae:0.1504884849)80.6/98:0.0314018136,(WP_004733311.1-Vibrio-splendidus:0.1122497220,(WP_009843931.1-Vibrio-sp.-AND4:0.0890660122,(WP_102353859.1-Vibrio-cyclitrophicus:0.1338553277,WP_171315631.1-Vibrio-splendidus:0.1051471581)85.5/100:0.0446045527)43.6/100:0.0546264740)0/86:0.0000029893)87.4/95:0.0244245531,WP_182796407.1-Vibrio-sp.-SG41-7:0.1971690733)10.2/13:0.0076623189)65.6/22:0.0256797914,(WP_193834777.1-Vibrio-sp.-OPT20:0.0419627157,(WP_029865567.1-Vibrio-parahaemolyticus:0.0612861383,WP_213967907.1-Vibrio-alginolyticus:0.0359532177)97.3/100:0.0906429742)97.2/100:0.0881445199)56.2/11:0.0146759457,((WP_130049361.1-Aliivibrio-finisterrensis:0.0696344917,WP_019826652.1-Vibrio-splendidus:0.1423346347)50.8/74:0.0280100658,WP_260256886.1-Vibrio-sp.-NC2:0.1100315453)79.7/67:0.0202618394)75.4/14:0.0099557826,(WP_012532661.1-Aliivibrio-fischeri:0.1352030030,(WP_258428294.1-Vibrio-sp.-RM-69-4:0.1072980980,WP_102428030.1-Vibrio-sp.-10N.261.54.E10:0.1438504893)69.2/89:0.0370834192)88/11:0.0388472975)83/14:0.0160339444,(WP_155682312.1-Aliivibrio-fischeri:0.0653048658,(WP_155677763.1-Aliivibrio-fischeri:0.1470186334,(WP_046209502.1-Vibrio:0.1219772925,(((WP_061039794.1-Vibrio-coralliirubri:0.0670793010,(WP_044365158.1-Vibrio-fluvialis:0.2792110437,WP_025523961.1-Vibrio-parahaemolyticus:0.1509984539)90.4/100:0.0615666524)77.7/100:0.0537954992,WP_065194916.1-Photobacterium-phosphoreum:0.1664512044)85.1/100:0.0537165792,WP_248386338.1-Vibrio-sp.-ED004:0.2325878335)74.3/14:0.0159076386)89.6/94:0.0549940208)42.2/4:0.0313618717)83.6/5:0.0284695083)89.8/4:0.0345998270,((WP_171343915.1-Vibrio-cyclitrophicus:0.0714841302,(WP_048661380.1-Vibrio-crassostreae:0.0883085072,WP_193846925.1-Vibrio-sp.-OPT10:0.0857167843)85.3/99:0.0391429729)78/95:0.0242046563,(WP_102248986.1-Vibrio-tasmaniensis:0.1873127250,WP_135445135.1-Vibrio-tasmaniensis:0.0788886603)97.5/99:0.1010366338)90.8/77:0.0409713744)81.8/11:0.0187509095)84.7/40:0.0134015859,(((WP_137008959.1-Vibrio-kanaloae:0.1216457873,WP_135353160.1-Aliivibrio-fischeri:0.0857453521)83.9/100:0.0374562165,WP_248638627.1-Vibrio-crassostreae:0.1319652939)87.1/96:0.0278105724,((((WP_012551417.1-Aliivibrio-salmonicida:0.0875750040,WP_012276595.1-Shewanella-halifaxensis:0.0544731875)77.5/100:0.0357822739,WP_135385147.1-Vibrio-tasmaniensis:0.1520991775)89.3/100:0.0603266185,WP_102571993.1-Vibrio:0.1008073578)81.6/97:0.0638544088,(WP_102285079.1-Vibrio-lentus:0.1214851038,WP_235060641.1-Vibrio-sp.-SS-MA-C1-2:0.2178985223)22.6/90:0.0239668000)84.2/94:0.0350496493)26.3/60:0.0121379490)89.2/78:0.0258759852)89.7/95:0.0189152725,(((((((((((((((((WP_041510674.1-Shewanella:0.0313988425,(WP_187575439.1-Shewanella-algae:0.0675149985,WP_101201417.1-Paraglaciecola-sp.-MB-3u-78:0.2683508333)73.1/24:0.0175964151)91.9/24:0.0517956644,(WP_208158397.1-Shewanella-algae:0.0811887725,WP_208157589.1-Shewanella-algae:0.0372539232)85.5/100:0.0291550209)82.4/24:0.0414500623,(WP_045460498.1-Vibrio-hyugaensis:0.1422487800,WP_220051788.1-Shewanella-sp.-FJAT-51860:0.1476730982)88.7/95:0.0660368077)87.8/46:0.0679136005,(WP_068898547.1-Enterovibrio-pacificus:0.1699045202,WP_039484180.1-Vibrio-sinaloensis:0.1748004297)65.2/19:0.0345892453)0/14:0.0000024691,((WP_094127815.1-Vibrio-sp.-V09_P4A23P171:0.1164292547,WP_199867560.1-Vibrio-anguillarum:0.0704526141)96.2/100:0.0766993813,WP_045898906.1-Enterobacteriaceae:0.1364144796)79.6/99:0.0356841790)39.3/24:0.0235212730,WP_107294203.1-Photobacterium-sp.-GB-50:0.1688102440)44.1/90:0.0242822995,(WP_208891742.1-Vibrio-anguillarum:0.1477010980,WP_148533341.1-Vibrio-cholerae:0.0606995392)84.3/95:0.0408621763)0/57:0.0011899525,(WP_000864156.1-Vibrio-cholerae:0.0515166156,WP_000864152.1-Vibrio-cholerae:0.1049363994)96.4/97:0.0726010444)81.1/82:0.0209752215,((WP_095464062.1-Vibrio-cholerae:0.0960373728,WP_149560528.1-Vibrio-cholerae:0.0825097499)85.2/99:0.0340128392,WP_095478449.1-Vibrio:0.0789809937)53.8/86:0.0332836481)74.1/46:0.0122080882,(WP_000864154.1-Vibrio-cholerae:0.0777197893,WP_102786270.1-Vibrio-cholerae:0.0742321478)76.8/95:0.0197194933)75.8/45:0.0150706788,((WP_069212356.1-Vibrio-cholerae:0.0244756929,WP_202641945.1-Vibrio-fluvialis:0.1331866609)99.4/92:0.1165851023,WP_175242188.1-Vibrio-cholerae:0.0885961535)82.7/23:0.0297148765)90.8/30:0.0402352833,((((WP_102986644.1-Vibrio-diazotrophicus:0.1469808341,(WP_107294031.1-Photobacterium-phosphoreum:0.0408423321,WP_107276976.1-Photobacterium-sp.-NCIMB-13483:0.0635233596)97.4/100:0.0998517081)84.8/98:0.0384701972,(WP_087853896.1-Photobacterium-andalusiense:0.0286924379,WP_107274737.1-Photobacterium-phosphoreum:0.1090231168)95.5/100:0.0643970723)84.7/67:0.0390340628,((((((((WP_065208126.1-Photobacterium-phosphoreum:0.1243198845,(WP_107293979.1-Photobacterium-phosphoreum:0.0878772860,WP_107228425.1-Photobacterium-phosphoreum:0.1674999069)89.1/100:0.0824792158)28.1/78:0.0114132042,WP_060997345.1-Photobacterium-aquimaris:0.1156769243)71.4/74:0.0149517539,(WP_232582982.1-Photobacterium-phosphoreum:0.0557498754,((WP_256383785.1-Photobacterium-toruni:0.0426167147,WP_232578536.1-Photobacterium-carnosum:0.0514926203)0/71:0.0000023436,WP_107228404.1-Photobacterium-phosphoreum:0.0919040646)91.3/90:0.0391464570)96.7/100:0.0577181474)82.4/92:0.0278456604,(WP_232603643.1-Photobacterium-carnosum:0.1107190565,WP_232615715.1-Photobacterium-carnosum:0.0776403563)93.1/99:0.0555921081)89.3/93:0.0297265864,WP_232600381.1-Photobacterium-carnosum:0.0526871159)40.6/82:0.0044111854,WP_232575020.1-Photobacterium-carnosum:0.1365720286)85.7/23:0.0318020662,(WP_107213932.1-Photobacterium-phosphoreum:0.0967010803,WP_065173166.1-Photobacterium-kishitanii:0.1534381969)91.2/75:0.0529056259)9/12:0.0099954347,(((WP_045045404.1-Photobacterium-kishitanii:0.0540641697,WP_107264793.1-Photobacterium-phosphoreum:0.0275145152)9.1/30:0.0133691276,WP_045030671.1-Photobacterium-phosphoreum:0.0397971495)95.5/100:0.0752517174,WP_232585319.1-Photobacterium-iliopiscarium:0.1122023752)79/38:0.0326871856)53.8/71:0.0243098046)72.4/43:0.0275816783,(WP_199351799.1-Vibrio-cholerae:0.0936370313,(WP_045039948.1-Photobacterium-iliopiscarium:0.1541016481,((WP_065176337.1-Photobacterium-aquimaris:0.0636217653,WP_107199726.1-Photobacterium-phosphoreum:0.0812292049)88/100:0.0447548657,WP_023931546.1-Photobacterium-leiognathi:0.1157291109)87/81:0.0599114040)85.9/94:0.0511839191)90.4/29:0.0577599845)74.4/28:0.0113078473)80/78:0.0127616384,(((WP_161119826.1-Photobacterium-lucens:0.1287912969,(WP_188234612.1-Vibrio-sp.-Y2-5:0.0605147001,WP_057556016.1-Vibrio-cholerae:0.0238824338)100/100:0.1946653911)70.3/95:0.0120069407,(WP_171138527.1-Vibrio-plantisponsor:0.0554274844,WP_102966424.1-Vibrio:0.1008186722)99.7/100:0.1565216468)84.2/85:0.0235132346,((((((WP_151182432.1-Photobacterium-damselae:0.1359164441,(WP_106261939.1-Photobacterium-damselae:0.0873499511,WP_115426223.1-Photobacterium-damselae:0.0630155751)19.9/72:0.0171488032)44.3/68:0.0172071686,WP_053987023.1-Photobacterium-leiognathi:0.1189648628)34.8/64:0.0202397630,WP_086367281.1-Photobacterium-damselae:0.0666613921)83.6/72:0.0226261125,(((WP_238087945.1-Photobacterium-damselae:0.0694574313,WP_017004373.1-Enterovibrio-norvegicus:0.2086690762)0/80:0.0000020862,(WP_065172005.1-Photobacterium-damselae:0.0868289443,WP_107185088.1-Photobacterium-leiognathi:0.0713088648)84.4/96:0.0218835110)84.7/96:0.0215423474,WP_151170663.1-Photobacterium-damselae:0.0585128055)84.2/74:0.0281785718)0/43:0.0000028812,WP_237041371.1-Photobacterium-damselae:0.0994828208)5.7/45:0.0052771069,WP_045063675.1-Photobacterium-leiognathi:0.1040619715)91.5/96:0.0358902595)86.7/81:0.0257631786)87.7/95:0.0222006793,WP_047876260.1-Photobacterium-aphoticum:0.1948438670)79/94:0.0160118549,(((((((((WP_232600401.1-Photobacterium-carnosum:0.1032906299,(WP_102481006.1-Vibrio-breoganii:0.1265880378,WP_258489575.1-Vibrio-alginolyticus:0.1346274244)93.5/98:0.0977566378)24.1/69:0.0170780439,WP_176284769.1-Photobacterium-damselae:0.1560799601)91.4/87:0.0518866263,WP_088132129.1-Vibrio-cholerae:0.0993088125)0/45:0.0314565440,(WP_255122387.1-Vibrio-cholerae:0.1222982141,(WP_188008910.1-Grimontia-hollisae:0.1239905669,WP_065173140.1-Photobacterium-kishitanii:0.1559159700)87.3/100:0.0650863786)73.6/9:0.0235950302)74.1/20:0.0261391667,((WP_249572008.1-Vibrio-cholerae:0.1113125260,WP_103155729.1-Vibrio-vulnificus:0.1214020110)0/33:0.0117457679,(WP_152138288.1-Plesiomonas-shigelloides:0.1664770626,WP_045618469.1-Vibrio-vulnificus:0.1304191555)45.5/18:0.0141398727)75.4/22:0.0198839293)84.1/19:0.0247397919,(WP_000864153.1-Vibrio-cholerae:0.0181612899,WP_251878143.1-Grimontia-sp.-020920N:0.1705461282)96.5/98:0.0820846527)76/30:0.0123543397,(WP_161129723.1-Photobacterium-lucens:0.1370304790,(WP_084988554.1-Vibrio:0.1374083417,WP_206634697.1-Vibrio-vulnificus:0.1697595444)29.2/96:0.0157883355)90.6/98:0.0528761396)75.1/30:0.0095503681,(WP_230432956.1-Plesiomonas-shigelloides:0.1027016281,WP_222763364.1-Vibrio-fluvialis:0.1072275459)92.7/97:0.0635896398)82.1/51:0.0239863280,((WP_148519618.1-Vibrio-cholerae:0.0254875445,(WP_167570263.1-Aeromonas-veronii:0.0616207977,WP_000864158.1-Vibrio-metoecus:0.0334889126)91.2/100:0.0293395688)77.3/100:0.0112057056,WP_148520854.1-Vibrio-cholerae:0.0387825622)99/100:0.0722028326)92/85:0.0466992545)0/35:0.0000024709,WP_045069930.1-Photobacterium-leiognathi:0.1648871524)76.6/42:0.0114811726,((((WP_067416136.1-Enterovibrio-coralii:0.0881512652,WP_149593442.1-Pseudoalteromonas-sp.-FUC4:0.1811073946)83.1/100:0.0316573608,WP_017008437.1-Enterovibrio-norvegicus:0.1274865240)0/35:0.0207687770,WP_251878087.1-Grimontia-sp.-020920N:0.1464838988)51.3/79:0.0160963455,(((WP_267741605.1-Vibrio-cholerae:0.1211332684,(WP_172567459.1-Vibrio-fluvialis:0.0967004129,WP_199354891.1-Vibrio-cholerae:0.0361524425)93.8/100:0.0863927884)0/8:0.0243170559,WP_102137258.1-Vibrio-cholerae:0.1067624069)53.6/93:0.0525394759,WP_148560094.1-Vibrio-mimicus:0.1885357014)75.8/91:0.0312707643)78.6/43:0.0095234072)80.5/50:0.0135269973)72.8/54:0.0114246146,((WP_107253996.1-Photobacterium-indicum:0.1309824826,WP_107295906.1-Photobacterium-phosphoreum:0.1733871805)80.3/100:0.0598835322,WP_102395415.1-Enterovibrio-norvegicus:0.1068661522)89.2/100:0.0464970949)86/89:0.0213281035,WP_180567593.1-Oceanospirillales:0.1949011827)74/14:0.0097043100,((((((WP_046223474.1-Vibrio-sp.-ECSMB14106:0.1002191195,WP_102562373.1-Vibrio-breoganii:0.1296096230)81.5/98:0.0369518953,WP_218382835.1-Photobacterium-sp.-WH24:0.1818416774)75.3/17:0.0339489044,(((WP_102546929.1-Vibrio-breoganii:0.1620024845,(WP_141346804.1-Vibrio-inusitatus:0.1538098147,WP_102282524.1-Vibrio-cyclitrophicus:0.1467695324)73.8/100:0.0408493222)0/65:0.0203606842,WP_102461639.1-Vibrio-breoganii:0.1233130844)76.9/69:0.0221739128,(WP_102455158.1-Vibrio-breoganii:0.1836428226,WP_135455667.1-Vibrio-echinoideorum:0.0937644037)78.5/69:0.0289179763)70.6/81:0.0404096898)24.1/7:0.0147293082,(WP_017031451.1-Vibrio-breoganii:0.0485333172,WP_248975945.1-Shewanella-kaireitica:0.1886807119)95.9/99:0.0849107035)84.2/61:0.0338049107,WP_193290728.1-Vibrio-parahaemolyticus:0.2524569882)59.7/43:0.0178752845,((WP_190336189.1-Vibrio-sp.-S9_S30:0.0745309161,WP_022612811.1-Vibrio-nigripulchritudo:0.1360458012)96.2/98:0.0905466481,((WP_070968050.1-Vibrio-sonorensis:0.2096010032,(WP_261835819.1-Vibrio-ishigakensis:0.1339200980,WP_017221048.1-Moritella-dasanensis:0.1264692938)82.6/100:0.0586621643)74.7/72:0.0202226653,(((WP_188011373.1-Vibrio-vulnificus:0.2418173904,WP_102329601.1-Vibrio-breoganii:0.1612318674)75.2/26:0.0413433711,WP_102446429.1-Vibrio-breoganii:0.1695507442)33.2/20:0.0223531124,WP_102451772.1-Vibrio-breoganii:0.2188468266)61.7/42:0.0302275709)57.4/29:0.0174107563)76.9/47:0.0315195269)0/24:0.0086844307)90.1/21:0.0378981720)83.4/28:0.0310617201,WP_102941572.1-Vibrio-diazotrophicus:0.1385702428)84.5/37:0.0268350766,(((WP_050928752.1-Yersinia-enterocolitica:0.2596365465,((WP_035339894.1-Dickeya:0.0228254736,WP_125258312.1-Dickeya-lacustris:0.0847984595)99.9/100:0.1686724830,WP_042868619.1-Dickeya:0.1028796186)93.2/100:0.1037705189)77.2/91:0.0309461467,WP_012883389.1-Dickeya-parazeae:0.2182528816)89.1/97:0.0643029360,WP_045977433.1-Vibrio-neptunius:0.2395901247)83.9/52:0.0307499629)81.4/45:0.0169346141,((((WP_238813939.1-Vibrio-vulnificus:0.1011620440,(WP_118894008.1-Vibrio-vulnificus:0.1174488665,WP_112353352.1-Shewanella-benthica:0.2115407348)85.8/21:0.0381533666)71/20:0.0263932975,((((((WP_061023752.1-Vibrio-splendidus:0.0636012553,(WP_220768829.1-Shewanella-sp.-MBTL60-007:0.1213125049,((WP_110456309.1-Shewanella-algidipiscicola:0.1566091893,WP_204999274.1-Parashewanella-hymeniacidonis:0.1509528018)33.4/94:0.0312325973,WP_194091044.1-Vibrio-hibernica:0.2136633056)82.3/94:0.0511366229)18/45:0.0089587768)89.6/88:0.0476838016,WP_248991952.1-Shewanella-profunda:0.1358715084)72.6/90:0.0238460086,((WP_011638275.1-Shewanella-frigidimarina:0.1512812788,(WP_220774251.1-Shewanella-glacialipiscicola:0.0296636893,WP_182751750.1-Shewanella-sp.-SG44-2:0.1558274566)89.5/100:0.0914628729)89.8/100:0.0548399826,WP_059744187.1-Shewanella-frigidimarina:0.1798262562)85.2/95:0.0250742715)86.4/94:0.0361764920,WP_182754472.1-Shewanella-sp.-SG41-3:0.1578958636)0/50:0.0032260671,((((WP_045569365.1-Vibrio:0.0781916391,WP_039437343.1-Vibrio-navarrensis:0.0439166078)99/100:0.1130237286,WP_176299354.1-Vibrio-sp.-05-20-BW147:0.1301452394)89.8/100:0.0641899606,WP_140233673.1-Shewanella-polaris:0.2194582619)80.7/51:0.0514604820,WP_192022372.1-Shewanella-sp.-WPAGA9:0.1758901718)90.5/60:0.0665894464)24.2/46:0.0191987419,(WP_095430409.1-Vibrio-vulnificus:0.0563488565,WP_114994956.1-Grimontia-hollisae:0.1943674678)97.8/100:0.1028887902)87.2/20:0.0335816784)79.3/84:0.0229461036,((WP_172851426.1-Vibrio-coralliilyticus:0.1549318990,WP_182742106.1-Shewanella-sp.-SG44-6:0.2557362769)66/97:0.0658261864,WP_108186198.1-Vibrio-splendidus:0.1098442857)91.4/96:0.0688317447)80.7/91:0.0315410325,(((WP_155660100.1-Aliivibrio-fischeri:0.0888580543,WP_095457151.1-Vibrio:0.0953607039)90.7/96:0.0368909983,(WP_252020469.1-Vibrio-sp.-SCSIO-43136:0.1124459765,WP_243584106.1-Vibrio-splendidus:0.1568271692)74.3/100:0.0319036878)91.9/96:0.0533700259,(WP_031827242.1-Vibrio-parahaemolyticus:0.3123545815,WP_166696422.1-Providencia-rettgeri:0.1310848134)0.6/27:0.0235082287)91.1/84:0.0465280047)67.4/61:0.0253208856)87/58:0.0332970254,(((WP_192015074.1-Photobacterium-arenosum:0.0896479028,WP_088876367.1-Vibrio-mediterranei:0.2274746823)79.3/89:0.0342751090,(WP_036979211.1-Providencia-alcalifaciens:0.1584458324,WP_101040106.1-Psychromonas-sp.-MB-3u-54:0.2118026457)93.4/96:0.0862988553)82.8/64:0.0427613686,((((WP_268809473.1-Vibrio-parahaemolyticus:0.1290387741,WP_023404414.1-Vibrio-halioticoli:0.2144913611)67.4/97:0.0342567552,WP_028024862.1-Enterovibrio-calviensis:0.1852533725)76.9/81:0.0241784266,WP_017007824.1-Enterovibrio-norvegicus:0.1800630145)0/76:0.0000022703,(WP_211678393.1-Moritella-sp.-36:0.2362688289,(WP_138627230.1-Pseudoalteromonas-sp.-S4741:0.1728576499,WP_269377025.1-Vibrio-sinaloensis:0.1660311108)63.6/96:0.0357978651)90.6/77:0.0672276673)86.5/59:0.0262324048)0/44:0.0000026628)90.5/78:0.0662573341,((WP_108452928.1-Enterobacter-cloacae:0.3002644117,WP_237309237.1-Vibrio:0.2391110467)13.5/75:0.0402121130,WP_133040538.1-Shewanella-fodinae:0.2456502486)91.5/80:0.0686660775)77.2/65:0.0289376817,((((WP_261298106.1-Shewanella-sp.-SP1S2-7:0.0989753653,WP_167678467.1-Shewanella-aestuarii:0.0942500773)98.8/100:0.2115978282,WP_248994245.1-Shewanella-gaetbuli:0.2751184319)81.6/100:0.0841471199,WP_222242368.1-Pseudoalteromonas-piscicida:0.3077329944)76.2/93:0.1022026402,WP_212599444.1-Shewanella-sp.-M16:0.3311719571)80.9/75:0.0523479692)0/36:0.0000022680,(WP_232612305.1-Photobacterium-phosphoreum:0.2837093160,WP_136989485.1-Vibrio-sp.-F13:0.2455177311)91.7/96:0.0891340289)99.5/100:0.1266681465,((WP_162556830.1-Proteus:0.0727565563,WP_161779122.1-Proteus-sp.-G2675:0.0938315293)99.6/100:0.1957199589,WP_236366479.1-Aeromonas-veronii:0.2171491521)80.5/98:0.0462527251)87.7/98:0.0298087567)94.2/97:0.0601262073)78.6/92:0.0194270451,WP_225633057.1-Aeromonas-enteropelogenes:0.2349385168)88.8/91:0.0388956445,WP_102949229.1-Aeromonas-caviae:0.2947318159)91.6/97:0.0529668508)82.9/92:0.0264804497,WP_171149131.1-Erwinia-sp.-JH02:0.1643795910)98.7/100:0.0872116814,((WP_241647798.1-Rosenbergiella-sp.-JB07:0.2703366393,WP_065370581.1-Kosakonia-sacchari:0.1819459244)86.3/98:0.0392204317,WP_181599127.1-Citrobacter-sp.-RHB36-C18:0.2253265963)69/96:0.0195935232)92.2/99:0.0582745486)82.6/97:0.0292000543,((((((((((WP_182931197.1-Enterobacter-cloacae:0.1612017973,WP_211776196.1-Klebsiella-pneumoniae:0.0669894166)63.6/86:0.0096980154,WP_047723323.1-Klebsiella-michiganensis:0.0423465001)53.6/86:0.0155999202,WP_104011323.1-Klebsiella-michiganensis:0.0406091055)79.1/81:0.0146087021,(((WP_048284323.1-Pluralibacter-gergoviae:0.1467278620,WP_065799756.1-Klebsiella-pneumoniae:0.0104398575)12.1/63:0.0068296523,WP_095100910.1-Raoultella-sp.-10-1:0.0651789327)92.3/84:0.0259634357,WP_149270444.1-Klebsiella-aerogenes:0.0381613299)77/64:0.0097008318)76.4/71:0.0093349284,WP_234579544.1-Raoultella-terrigena:0.0415787871)54.3/70:0.0063694023,WP_221694709.1-Klebsiella-aerogenes:0.0971160174)76.6/76:0.0074574425,(WP_213760001.1-Klebsiella-aerogenes:0.0653419478,WP_223370324.1-Klebsiella-pneumoniae:0.0339507890)80.6/91:0.0130239384)87.5/91:0.0137025335,(((((WP_095100919.1-Raoultella-sp.-10-1:0.0587026322,WP_212709529.1-Tatumella-sp.-JGM130:0.0117694117)95.6/100:0.0716851611,WP_139968923.1-Klebsiella-pneumoniae:0.1211205241)78.9/97:0.0265580570,WP_236948353.1-Klebsiella-sp.-2019SCSN059:0.1008009770)94.3/96:0.0758590777,(WP_128320364.1-Raoultella-planticola:0.0609335435,(WP_236948387.1-Klebsiella-sp.-2019SCSN059:0.0391887837,((WP_114692711.1-Klebsiella-pneumoniae:0.0388780562,WP_259891680.1-Klebsiella-variicola:0.0320111999)0/92:0.0000024046,WP_123755035.1-Raoultella-sp.-BIGb0399:0.0380585133)83.2/95:0.0113721348)76.1/98:0.0071305743)58.4/71:0.0085311205)0/16:0.0031957614,WP_046852082.1-Raoultella-planticola:0.0396171399)77.6/19:0.0071049024)75.5/19:0.0066752131,((WP_234537246.1-Raoultella-ornithinolytica:0.0395332292,WP_113990968.1-Raoultella:0.0253610820)98.4/100:0.0464112542,WP_186601692.1-Klebsiella-quasipneumoniae:0.0262212127)75.2/36:0.0067314834)99.6/99:0.1114256909,WP_123822891.1-Klebsiella:0.0250098904)99.6/100:0.1100025074)84.5/100:0.0340696901,(WP_072663757.1-Enterobacteriaceae:0.0752993786,WP_021241446.1-Enterobacter-roggenkampii:0.1867280663)40.1/99:0.0283117526)97.8/100:0.0998969321)94.8/100:0.0533878724,XP_011503919.1-Ceratosolen-solmsi-marchali:0.0868395184);

**Nematode RML-3, with bacterial and eukaryotic homologs**

**Shown in Figure S2**

ML tree method: FastTree

Sequences: 2787

Substitution model: JTT

(((((('WP_046223474.1-Vibrio-sp.-ECSMB14106':0.09899000000000013,('WP_102546929.1-Vibrio-breoganii':0.12068999999999974,'WP_102461639.1-Vibrio-breoganii':0.1266099999999999)0.228:0.058170000000000055)0.871:0.06113999999999997,'WP_218382835.1-Photobacterium-sp.-WH24':0.1732499999999999)0.823:0.04364999999999997,'WP_135455667.1-Vibrio-echinoideorum':0.09006000000000025)0.824:0.037760000000000016,('WP_065296243.1-Vibrio-natriegens':0.11907000000000023,('WP_166378370.1-Pseudoalteromonas-sp.-Z9A4':0.16768000000000027,'WP_102455158.1-Vibrio-breoganii':0.12288000000000032)0.73:0.06312999999999969)0.451:0.024799999999999933)0.848:0.031140000000000168,(('WP_075764060.1-Photobacterium-proteolyticum':0.1619600000000001,'WP_239411815.1-Photobacterium-sp.-OFAV2-7':0.12441000000000013)0.938:0.07406999999999986,('WP_197173106.1-Pseudoalteromonas-sp.-NZS11_1':0.1687700000000003,('WP_149593442.1-Pseudoalteromonas-sp.-FUC4':0.1382399999999997,('WP_261923504.1-Shewanella-sp.-NFH-SH190041':0.20307999999999993,'WP_102523418.1-Vibrio-tapetis':0.16268999999999956)0.616:0.038140000000000285)0.494:0.05149000000000026)0.645:0.01391999999999971)0.284:0.01827000000000023)0.907:0.036909999999999776,(('WP_022612811.1-Vibrio-nigripulchritudo':0.1143799999999997,('WP_190336189.1-Vibrio-sp.-S9_S30':0.06235999999999997,(('WP_077662143.1-Salinivibrio-proteolyticus':0.019689999999999763,'WP_069589493.1-Salinivibrio-sp.-DV':0.07521999999999984)0.348:0.021279999999999966,('WP_077659452.1-Salinivibrio-kushneri':0.02933999999999992,('WP_077659434.1-Salinivibrio-kushneri':0.1159100000000004,'WP_269578450.1-Salinivibrio-kushneri':0.06849000000000016)0.431:0.020899999999999697)0.917:0.04906999999999995)1.0:0.2787299999999999)0.457:0.020309999999999828)0.945:0.06420000000000048,(('WP_141346804.1-Vibrio-inusitatus':0.1895500000000001,('WP_017221048.1-Moritella-dasanensis':0.1245400000000001,'WP_261835819.1-Vibrio-ishigakensis':0.1293000000000002)0.717:0.036680000000000046)0.622:0.01808999999999994,('WP_102446429.1-Vibrio-breoganii':0.17212000000000005,('WP_165312521.1-Vibrio-ziniensis':0.11045999999999978,'WP_172561824.1-Vibrio-furnissii':0.09702999999999973)0.765:0.02825000000000033)0.862:0.04310999999999998)0.81:0.02845000000000031)0.856:0.01806999999999981,((('WP_171749982.1-Vibrio-sp.-RE88':0.1647000000000003,'WP_102941572.1-Vibrio-diazotrophicus':0.13458000000000014)0.752:0.03351999999999977,(((('WP_048899071.1-Photobacterium-swingsii':0.12131999999999987,'WP_103155729.1-Vibrio-vulnificus':0.12298999999999971)0.872:0.03824000000000005,('WP_248386338.1-Vibrio-sp.-ED004':0.16789999999999994,('WP_108186198.1-Vibrio-splendidus':0.14753000000000016,'WP_046209502.1-Vibrio':0.14124000000000025)0.808:0.03872000000000009)0.645:0.013659999999999783)0.761:0.026550000000000296,'WP_139685731.1-Vibrio-tasmaniensis':0.14271000000000011)0.423:0.06657000000000002,(((((('WP_102329601.1-Vibrio-breoganii':0.18401000000000023,(('WP_017027407.1-Vibrio-breoganii':0.11046999999999985,('WP_172380549.1-Vibrio-sp.-Vb339':0.12954,'WP_102443327.1-Vibrio-breoganii':0.06557000000000013)0.938:0.07507999999999981)0.447:0.024720000000000297,(('WP_096629284.1-Salinivibrio-sp.-YCSC6':0.14805000000000046,('WP_020332409.1-Vibrio-fluvialis':0.06952999999999987,'WP_193262143.1-Vibrio-parahaemolyticus':0.25402000000000013)0.221:0.019360000000000266)0.946:0.08359000000000005,('WP_014230611.1-Vibrio-sp.-EJY3':0.11349000000000009,'WP_005395829.1-Vibrio-diabolicus':0.20108000000000015)0.637:0.046820000000000306)0.758:0.015229999999999855)0.789:0.023470000000000102)0.504:0.011900000000000244,'WP_203433235.1-Shewanella-sp.-LZH-2':0.22405000000000008)0.792:0.030819999999999848,((((('WP_102533425.1-Vibrio-breoganii':0.12888999999999973,('WP_254101081.1-Alteromonas-sp.-LMIT007':0.15354,'WP_102532836.1-Vibrio-sp.-10N.286.48.B7':0.08563999999999972)0.966:0.07662999999999975)0.747:0.01067000000000018,'WP_102562373.1-Vibrio-breoganii':0.12462000000000018)0.793:0.04021000000000008,((('WP_185890410.1-Vibrio-vulnificus':0.1666300000000005,'WP_238130349.1-Vibrio-cincinnatiensis':0.13894000000000029)0.751:0.02071999999999985,(((('WP_233983986.1-Vibrio-fluvialis':0.06325000000000003,'WP_099031861.1-Pseudoalteromonas-sp.-GCY':0.1431300000000002)0.832:0.03803000000000001,('WP_239872404.1-Shewanella-insulae':0.14210000000000012,'WP_208844179.1-Pseudoalteromonas-xiamenensis':0.12056999999999984)0.196:0.02059000000000033)0.842:0.032629999999999715,'WP_233976434.1-Vibrio':0.14009000000000027)0.822:0.030950000000000255,('WP_045460498.1-Vibrio-hyugaensis':0.18742999999999999,'WP_220051788.1-Shewanella-sp.-FJAT-51860':0.09711999999999987)0.892:0.05418000000000012)0.672:0.025439999999999685)0.276:0.014149999999999885,('WP_143680610.1-Vibrio-furnissii':0.11779000000000028,('WP_218529053.1-Shewanella-sp.-NIFS-20-20':0.13319999999999954,'WP_225483056.1-Vibrio-vulnificus':0.04878999999999989)0.978:0.0812400000000002)0.886:0.039839999999999876)0.597:0.032110000000000305)0.422:0.026429999999999954,(('WP_193255073.1-Vibrio-navarrensis':0.022769999999999957,'WP_258469755.1-Vibrio-alginolyticus':0.11095999999999995)0.974:0.07873999999999981,('WP_065600047.1-Aliivibrio-sp.-1S128':0.11261999999999972,'WP_130066657.1-Aliivibrio-finisterrensis':0.13386999999999993)0.789:0.03943999999999992)0.883:0.04369000000000023)0.788:0.031870000000000065,('WP_152797327.1-Salinivibrio-sp.-VYel1':0.21302999999999983,(('WP_182742106.1-Shewanella-sp.-SG44-6':0.26731000000000016,'WP_102282524.1-Vibrio-cyclitrophicus':0.1429800000000001)0.843:0.06203000000000003,'WP_121852788.1-Parashewanella-spongiae':0.1951299999999998)0.607:0.060419999999999696)0.827:0.03805000000000014)0.366:0.021339999999999915)0.291:0.000550,((('WP_204999274.1-Parashewanella-hymeniacidonis':0.19799999999999995,('WP_102457123.1-Vibrio-breoganii':0.12073,'WP_077647682.1-Salinivibrio-sp.-IB643':0.09454999999999991)0.442:0.03994000000000009)0.341:0.016080000000000094,(('WP_059744187.1-Shewanella-frigidimarina':0.20355000000000034,('WP_011638275.1-Shewanella-frigidimarina':0.11589999999999989,('WP_182751750.1-Shewanella-sp.-SG44-2':0.13190999999999997,'WP_220774251.1-Shewanella-glacialipiscicola':0.03932000000000002)0.964:0.10132999999999992)0.875:0.041790000000000216)0.403:0.028209999999999624,'WP_248991952.1-Shewanella-profunda':0.11672999999999956)0.852:0.03304000000000018)0.768:0.02466999999999997,(('WP_238813939.1-Vibrio-vulnificus':0.09559000000000051,'WP_158107962.1-Vibrio-furnissii':0.1334900000000001)0.666:0.01020999999999983,'WP_182754472.1-Shewanella-sp.-SG41-3':0.15632000000000001)0.413:0.006590000000000096)0.932:0.02690999999999999)0.845:0.01916999999999991,('WP_031414678.1-Vibrio-parahaemolyticus':0.16036000000000028,(('WP_217557493.1-Vibrio-metschnikovii':0.049980000000000135,'WP_086585081.1-Vibrio-parahaemolyticus':0.03221000000000007)0.966:0.07703999999999978,('WP_140342839.1-Vibrio-parahaemolyticus':0.16983999999999977,('WP_217542775.1-Vibrio-metschnikovii':0.05484,'WP_113593871.1-Vibrio':0.06583999999999968)0.927:0.07357999999999976)0.797:0.03390000000000004)0.8:0.02819000000000038)0.691:0.04281999999999986)0.543:0.014890000000000292,(((((((((('WP_171138527.1-Vibrio-plantisponsor':0.051730000000000054,'WP_102966424.1-Vibrio':0.08845999999999998)0.995:0.12417999999999996,'WP_251878087.1-Grimontia-sp.-020920N':0.09318999999999988)0.86:0.0400400000000003,('WP_017004373.1-Enterovibrio-norvegicus':0.21597,('WP_023404414.1-Vibrio-halioticoli':0.2195800000000001,('WP_045039948.1-Photobacterium-iliopiscarium':0.12865000000000038,('WP_023931546.1-Photobacterium-leiognathi':0.10801000000000016,('WP_107199726.1-Photobacterium-phosphoreum':0.07597999999999949,'WP_065176337.1-Photobacterium-aquimaris':0.05708000000000002)0.815:0.0394300000000003)0.927:0.07394000000000034)0.401:0.03644999999999987)0.355:0.028750000000000053)0.66:0.02721000000000018)0.805:0.018919999999999604,(((('WP_068898547.1-Enterovibrio-pacificus':0.17859999999999987,('WP_069212356.1-Vibrio-cholerae':0.03421999999999992,'WP_202641945.1-Vibrio-fluvialis':0.09931000000000001)0.908:0.08118999999999987)0.842:0.0429400000000002,((('WP_045063675.1-Photobacterium-leiognathi':0.06084999999999985,(('WP_086367281.1-Photobacterium-damselae':0.07900000000000018,((('WP_115426223.1-Photobacterium-damselae':0.053830000000000044,'WP_151182432.1-Photobacterium-damselae':0.1307100000000001)0.763:0.020770000000000177,'WP_106261939.1-Photobacterium-damselae':0.07254000000000005)0.662:0.03114999999999979,'WP_237041371.1-Photobacterium-damselae':0.0747399999999998)0.471:0.030200000000000227)0.684:0.0016999999999995907,('WP_151170663.1-Photobacterium-damselae':0.040280000000000094,('WP_238087945.1-Photobacterium-damselae':0.06408000000000014,('WP_107185088.1-Photobacterium-leiognathi':0.0741499999999995,'WP_065172005.1-Photobacterium-damselae':0.07327999999999957)0.823:0.01967000000000052)0.666:0.021160000000000068)0.725:0.02811999999999948)0.837:0.02607000000000026)0.933:0.04573999999999989,('WP_065208126.1-Photobacterium-phosphoreum':0.14697999999999967,(('WP_020431327.1-Vibrio-fluvialis':0.09512999999999971,'WP_055453311.1-Vibrio-fluvialis':0.03161000000000014)0.988:0.09597999999999995,('WP_221070344.1-Vibrio-alfacsensis':0.14732999999999974,(((('WP_268679810.1-Vibrio-barjaei':0.09465999999999974,('WP_103882085.1-Vibrio-hangzhouensis':0.07966000000000006,'WP_265676699.1-Vibrio-sp.-ZSDZ65':0.09672999999999998)0.727:0.05516999999999994)0.901:0.04550999999999972,'WP_250617847.1-Vibrio-sp.-S4B1':0.06834000000000007)0.153:0.016450000000000298,'WP_102532726.1-Vibrio-sp.-10N.286.48.B7':0.10436000000000023)0.875:0.03133000000000008,((('WP_252012765.1-Vibrio-sp.-SCSIO-43140':0.11668000000000012,('WP_062462511.1-Vibrio-mediterranei':0.07889999999999997,('WP_234495031.1-Vibrio-maritimus':0.1888099999999997,'WP_171392669.1-Vibrio':0.06588999999999956)0.893:0.05215000000000014)0.241:0.035350000000000215)0.824:0.020509999999999806,'WP_239941760.1-Vibrio-mediterranei':0.09475999999999996)0.809:0.0301499999999999,'WP_252012797.1-Vibrio-sp.-SCSIO-43140':0.10874000000000006)0.697:0.012270000000000003)0.174:0.008239999999999803)0.915:0.03679000000000032)0.776:0.016229999999999745)0.747:0.007509999999999906)0.844:0.026120000000000143,(('WP_028024862.1-Enterovibrio-calviensis':0.22475999999999985,('WP_107294031.1-Photobacterium-phosphoreum':0.031179999999999986,'WP_107276976.1-Photobacterium-sp.-NCIMB-13483':0.061590000000000256)0.938:0.0801099999999999)0.584:0.03934000000000015,(('WP_087853896.1-Photobacterium-andalusiense':0.03122999999999987,'WP_107274737.1-Photobacterium-phosphoreum':0.09635000000000016)0.99:0.08164999999999978,(('WP_232600381.1-Photobacterium-carnosum':0.07343999999999973,(('WP_232585319.1-Photobacterium-iliopiscarium':0.07102999999999948,('WP_232582982.1-Photobacterium-phosphoreum':0.04944999999999977,('WP_107228404.1-Photobacterium-phosphoreum':0.08485000000000031,('WP_256383785.1-Photobacterium-toruni':0.04233999999999982,'WP_232578536.1-Photobacterium-carnosum':0.04848000000000008)0.002:0.000550)0.923:0.03678999999999988)0.956:0.06099999999999994)0.915:0.033339999999999925,(('WP_045045404.1-Photobacterium-kishitanii':0.03653999999999957,('WP_045030671.1-Photobacterium-phosphoreum':0.04875000000000007,'WP_107264793.1-Photobacterium-phosphoreum':0.021179999999999755)0.552:0.016750000000000043)0.977:0.06744000000000039,('WP_053987023.1-Photobacterium-leiognathi':0.10216000000000003,'WP_232575020.1-Photobacterium-carnosum':0.10793000000000008)0.666:0.02601000000000031)0.784:0.020609999999999573)0.859:0.0328400000000002)0.772:0.022070000000000256,'WP_107213932.1-Photobacterium-phosphoreum':0.12988)0.725:0.015379999999999505)0.834:0.026830000000000354)0.685:0.009729999999999794)0.79:0.020199999999999996)0.825:0.026190000000000158,((('WP_107294203.1-Photobacterium-sp.-GB-50':0.1772100000000001,('WP_148533341.1-Vibrio-cholerae':0.052799999999999514,'WP_208891742.1-Vibrio-anguillarum':0.13912999999999975)0.722:0.025060000000000304)0.803:0.022130000000000205,(('WP_102786270.1-Vibrio-cholerae':0.08549999999999969,('WP_000864156.1-Vibrio-cholerae':0.05060999999999982,'WP_000864152.1-Vibrio-cholerae':0.09628999999999976)0.876:0.031159999999999854)0.205:0.016690000000000538,('WP_161119826.1-Photobacterium-lucens':0.10856999999999983,(('WP_061039794.1-Vibrio-coralliirubri':0.054689999999999905,('WP_044365158.1-Vibrio-fluvialis':0.29556000000000004,'WP_025523961.1-Vibrio-parahaemolyticus':0.1350100000000003)0.573:0.06357999999999997)0.974:0.12959000000000032,('WP_182690147.1-Psychromonas-sp.-SR45-3':0.1409400000000005,'WP_011771697.1-Psychromonas-ingrahamii':0.08062000000000014)0.967:0.09895999999999994)0.133:0.02774999999999972)0.897:0.0636300000000003)0.855:0.030450000000000088)0.745:0.009249999999999758,(('WP_175242188.1-Vibrio-cholerae':0.0749200000000001,('WP_149560528.1-Vibrio-cholerae':0.08353999999999973,'WP_199351799.1-Vibrio-cholerae':0.08763000000000032)0.184:0.042910000000000004)0.018:0.03337999999999974,(('WP_095464062.1-Vibrio-cholerae':0.08231999999999973,'WP_117235674.1-Vibrio-maerlii':0.16904999999999948)0.612:0.03066000000000013,'WP_095478449.1-Vibrio':0.07141999999999982)0.784:0.026049999999999685)0.792:0.023030000000000328)0.818:0.03624000000000027)0.877:0.02768999999999977,((((('WP_045898906.1-Enterobacteriaceae':0.13442000000000043,(('WP_022590465.1-Vibrio-nigripulchritudo':0.21773000000000042,('WP_088876367.1-Vibrio-mediterranei':0.25412999999999997,'WP_192015074.1-Photobacterium-arenosum':0.05286999999999997)0.867:0.0510900000000003)0.794:0.024529999999999497,(('WP_036979211.1-Providencia-alcalifaciens':0.15656000000000025,'WP_101040106.1-Psychromonas-sp.-MB-3u-54':0.21926000000000023)0.163:0.044159999999999755,('WP_102451772.1-Vibrio-breoganii':0.16195000000000004,'WP_188011373.1-Vibrio-vulnificus':0.16222000000000003)0.819:0.12534)0.775:0.02491999999999983)0.895:0.05334000000000039)0.748:0.025559999999999583,('WP_211678393.1-Moritella-sp.-36':0.26182000000000016,(('WP_239929047.1-Vibrio-kanaloae':0.03367999999999993,'WP_076657081.1-unclassified-Vibrio':0.05696000000000012)0.977:0.08760000000000012,('WP_075989065.1-Vibrio-fluvialis':0.03139000000000003,'WP_044366945.1-Vibrio-fluvialis':0.029739999999999878)0.871:0.03802000000000039)0.927:0.09299999999999997)0.907:0.07613999999999965)0.701:0.014490000000000336,((((('WP_000864154.1-Vibrio-cholerae':0.09224999999999994,'WP_038178862.1-Vibrio-rhizosphaerae':0.11886000000000019)0.903:0.0526399999999998,((('WP_077313846.1-Vibrio-palustris':0.025009999999999977,'WP_123015137.1-Vibrio-zhugei':0.03601999999999972)0.999:0.15931000000000006,('WP_046218796.1-Photobacterium-halotolerans':0.1602999999999999,'WP_172533652.1-Vibrio-aestuarianus':0.26468999999999987)0.243:0.011239999999999917)0.266:0.03567000000000009,(('WP_211647400.1-Photobacterium-sp.-GJ3':0.053160000000000096,'WP_120511428.1-Photobacterium-salinisoli':0.033580000000000165)0.971:0.062269999999999825,('WP_101064017.1-Moritella-sp.-Urea-trap-13':0.21128999999999998,('WP_169403689.1-Rouxiella-aceris':0.057630000000000514,('WP_200225058.1-Rahnella-aceris':0.10095999999999972,'WP_195814554.1-Rahnella':0.10082999999999975)0.873:0.042030000000000456)0.979:0.11660999999999966)0.821:0.046330000000000204)0.783:0.02198000000000011)0.166:0.002349999999999852)0.728:0.044820000000000526,((('WP_193290728.1-Vibrio-parahaemolyticus':0.2431399999999999,'WP_261894433.1-Vibrio-porteresiae':0.1657099999999998)0.863:0.05740999999999996,'WP_124938567.1-Vibrio-viridaestus':0.12617000000000012)0.715:0.035490000000000244,(('WP_261600599.1-Photobacterium-sp.-TY1-4':0.09637999999999991,('WP_182108830.1-Vibrio-marinisediminis':0.0872200000000003,'WP_114765998.1-Vibrio-rhodolitus':0.11073999999999984)0.913:0.06808999999999976)0.485:0.032600000000000406,'WP_159197271.1-Escherichia-coli':0.11582000000000026)0.839:0.03221000000000007)0.585:0.017360000000000042)0.21:0.013269999999999449,'WP_088133934.1-Vibrio-gazogenes':0.22708999999999957)0.797:0.016370000000000218,(('WP_202661182.1-Vibrio-fluvialis':0.05636000000000019,'WP_112404745.1-Vibrio-diazotrophicus':0.04275000000000029)0.936:0.05631999999999948,(('WP_015854950.1-Musicola-paradisiaca':0.04389000000000021,'WP_192457620.1-Musicola-keenii':0.05936999999999992)1.0:0.2062599999999999,(('WP_200550470.1-Kosakonia-sp.-LAM2021':0.08064000000000027,'WP_263100201.1-Kosakonia-sp.-ML.JS2a':0.033600000000000296)0.998:0.14230999999999971,('WP_094109047.1-Lonsdalea-iberica':0.09802,('WP_026742149.1-Lonsdalea-quercina':0.0337799999999997,'WP_094117837.1-Lonsdalea-britannica':0.0282300000000002)0.961:0.0771799999999998)0.847:0.029819999999999958)0.678:0.02051000000000025)0.93:0.06640999999999986)0.712:0.035210000000000186)0.758:0.01656000000000013)0.629:0.013129999999999864,((('WP_068331709.1-Photobacterium-jeanii':0.1919500000000003,(('WP_238097025.1-Vibrio-cincinnatiensis':0.19248999999999983,'WP_118894008.1-Vibrio-vulnificus':0.11789000000000005)0.234:0.017300000000000537,'WP_230432956.1-Plesiomonas-shigelloides':0.12008000000000019)0.611:0.016610000000000014)0.522:0.022459999999999702,('WP_132698543.1-Vibrio-harveyi-group':0.1083700000000003,'WP_192928291.1-Vibrio-vulnificus':0.022740000000000204)0.995:0.12989000000000006)0.838:0.0240800000000001,(('WP_017007824.1-Enterovibrio-norvegicus':0.15154000000000023,('WP_114994956.1-Grimontia-hollisae':0.16680000000000028,'WP_095430409.1-Vibrio-vulnificus':0.05750999999999973)0.939:0.09179000000000004)0.926:0.06790999999999991,(('WP_107253996.1-Photobacterium-indicum':0.16086,('WP_269377025.1-Vibrio-sinaloensis':0.15955999999999992,'WP_138627230.1-Pseudoalteromonas-sp.-S4741':0.16669)0.798:0.034170000000000034)0.771:0.03727999999999998,('WP_182748079.1-Pseudoalteromonas-sp.-SR41-1':0.15280000000000005,('WP_107284330.1-Photobacterium-lipolyticum':0.05418000000000012,('WP_017221010.1-Moritella-dasanensis':0.1091899999999999,'WP_170899585.1-Vibrio-sp.-3-2-1':0.10766000000000009)0.969:0.0840500000000004)0.929:0.05702999999999969)0.433:0.028200000000000003)0.829:0.028259999999999952)0.671:0.026149999999999896)0.904:0.04861000000000004)0.882:0.019640000000000324,('WP_107187026.1-Photobacterium-kishitanii':0.18727000000000027,'WP_006646466.1-Photobacterium-sp.-SKA34':0.15717999999999988)0.755:0.027810000000000112)0.778:0.014619999999999855)0.568:0.010399999999999743)0.827:0.018629999999999924,((('WP_199867560.1-Vibrio-anguillarum':0.10026000000000002,'WP_094127815.1-Vibrio-sp.-V09_P4A23P171':0.08565000000000023)0.937:0.07284999999999986,(('WP_011716450.1-Shewanella-sp.-ANA-3':0.26959,'WP_268809473.1-Vibrio-parahaemolyticus':0.08273999999999981)0.874:0.0754400000000004,('WP_258516089.1-Vibrio-alginolyticus':0.0945100000000001,('WP_033540599.1-Shewanella':0.15377000000000018,'WP_031815865.1-Vibrio-parahaemolyticus':0.11246000000000045)0.752:0.037189999999999834)0.862:0.052850000000000286)0.446:0.03552999999999962)0.637:0.04255999999999993,(('WP_107295906.1-Photobacterium-phosphoreum':0.13099999999999978,('WP_107293979.1-Photobacterium-phosphoreum':0.08888000000000007,'WP_107228425.1-Photobacterium-phosphoreum':0.13506000000000018)0.84:0.043320000000000025)0.896:0.07135000000000025,('WP_065194916.1-Photobacterium-phosphoreum':0.12453000000000003,'WP_193250619.1-Vibrio-parahaemolyticus':0.15761999999999965)0.281:0.05071000000000048)0.712:0.0226999999999995)0.793:0.024329999999999963)0.888:0.022720000000000518,(('WP_065173166.1-Photobacterium-kishitanii':0.17998000000000003,((('WP_188008910.1-Grimontia-hollisae':0.1623199999999998,'WP_065173140.1-Photobacterium-kishitanii':0.08006000000000002)0.933:0.07316999999999974,('WP_232600401.1-Photobacterium-carnosum':0.12460999999999967,'WP_232615715.1-Photobacterium-carnosum':0.09297999999999984)0.796:0.02536000000000005)0.383:0.017260000000000275,'WP_060997345.1-Photobacterium-aquimaris':0.12495999999999974)0.865:0.04569000000000001)0.849:0.0367099999999998,(((('WP_161129723.1-Photobacterium-lucens':0.11257000000000028,('WP_084988554.1-Vibrio':0.12802999999999987,'WP_206634697.1-Vibrio-vulnificus':0.15589999999999993)0.838:0.0517000000000003)0.855:0.040119999999999933,((('WP_199354891.1-Vibrio-cholerae':0.03459999999999974,'WP_172567459.1-Vibrio-fluvialis':0.08775000000000022)0.981:0.09324000000000021,('WP_267741605.1-Vibrio-cholerae':0.12467000000000006,'WP_102137258.1-Vibrio-cholerae':0.07274000000000003)0.018:0.02546000000000026)0.746:0.015839999999999854,((((('WP_161972844.1-Enterovibrio-baiacu':0.09144000000000041,'WP_259522409.1-Shewanella-baltica':0.15573000000000015)0.851:0.044019999999999726,'WP_067416136.1-Enterovibrio-coralii':0.09830999999999968)0.48:0.016760000000000108,'WP_017008437.1-Enterovibrio-norvegicus':0.1221000000000001)0.921:0.059539999999999704,'WP_222763364.1-Vibrio-fluvialis':0.13459999999999983)0.713:0.013190000000000257,('WP_238819535.1-Vibrio-furnissii':0.07833000000000023,'WP_152138288.1-Plesiomonas-shigelloides':0.10457000000000027)0.946:0.05610999999999988)0.491:0.03492000000000006)0.892:0.06559000000000026)0.174:0.009119999999999795,('WP_148560094.1-Vibrio-mimicus':0.15504000000000007,('WP_237041393.1-Photobacterium-damselae':0.12134,'WP_080158029.1-Photobacterium-piscicola':0.15741000000000005)0.828:0.05238999999999994)0.645:0.03872000000000009)0.843:0.023680000000000145,((('WP_148520854.1-Vibrio-cholerae':0.029879999999999907,('WP_148519618.1-Vibrio-cholerae':0.022950000000000248,('WP_000864158.1-Vibrio-metoecus':0.030389999999999695,'WP_167570263.1-Aeromonas-veronii':0.05608999999999975)0.918:0.027270000000000127)0.865:0.016189999999999927)0.97:0.07956999999999992,('WP_237042700.1-Photobacterium-damselae':0.06129000000000007,('WP_013365656.1-[Enterobacter':0.20147000000000048,('WP_214579582.1-Enterobacter-asburiae':0.08492999999999995,('WP_193129600.1-Klebsiella-aerogenes':0.0676899999999998,('WP_258988101.1-Scandinavium-hiltneri':0.06806000000000001,'WP_181620050.1-Enterobacter-sp.-RHBSTW-00994':0.05103999999999953)0.932:0.050530000000000186)0.924:0.05649000000000015)0.116:0.025290000000000035)0.998:0.21756999999999982)0.894:0.0702799999999999)0.68:0.005119999999999791,('WP_255122387.1-Vibrio-cholerae':0.1055600000000001,('WP_249572008.1-Vibrio-cholerae':0.09406000000000025,'WP_088132129.1-Vibrio-cholerae':0.09941999999999984)0.386:0.03486000000000011)0.689:0.028499999999999748)0.866:0.02943000000000051)0.774:0.012939999999999507)0.732:0.011530000000000484)0.737:0.01110999999999951,((('WP_000864153.1-Vibrio-cholerae':0.062100000000000044,'WP_251878143.1-Grimontia-sp.-020920N':0.1099199999999998)0.888:0.05586000000000002,('WP_045618469.1-Vibrio-vulnificus':0.11790000000000012,'WP_047876260.1-Photobacterium-aphoticum':0.1617700000000002)0.129:0.034849999999999604)0.807:0.039280000000000204,'WP_102395415.1-Enterovibrio-norvegicus':0.14053000000000004)0.872:0.05284999999999984)0.734:0.009800000000000253,(('WP_148560072.1-Vibrio-mimicus':0.18649999999999967,'WP_176284769.1-Photobacterium-damselae':0.13473999999999986)0.883:0.052380000000000315,('WP_045069930.1-Photobacterium-leiognathi':0.1272000000000002,('WP_102481006.1-Vibrio-breoganii':0.13131999999999966,'WP_258489575.1-Vibrio-alginolyticus':0.10215999999999958)0.836:0.04576000000000047)0.837:0.04990999999999968)0.793:0.023249999999999993)0.273:0.010419999999999874,(((('WP_194091044.1-Vibrio-hibernica':0.22043999999999997,('WP_041156219.1-Vibrio-mytili':0.14802000000000026,('WP_045494469.1-Vibrio-hyugaensis':0.040699999999999736,'WP_047480300.1-Vibrio-campbellii':0.054119999999999724)0.947:0.1010500000000003)0.978:0.15273000000000003)0.321:0.03811999999999971,'WP_101201417.1-Paraglaciecola-sp.-MB-3u-78':0.17457000000000011)0.841:0.032520000000000326,(('WP_261874894.1-Vibrio-rarus':0.16470999999999947,((((('WP_243579549.1-Vibrio-splendidus':0.19679999999999964,'WP_192022372.1-Shewanella-sp.-WPAGA9':0.16737000000000002)0.818:0.04701000000000022,('WP_140233673.1-Shewanella-polaris':0.21659999999999968,('WP_176299354.1-Vibrio-sp.-05-20-BW147':0.14297000000000004,('WP_039437343.1-Vibrio-navarrensis':0.040000000000000036,'WP_045569365.1-Vibrio':0.06984000000000012)0.969:0.0947)0.922:0.05575999999999981)0.108:0.01949000000000023)0.825:0.042609999999999815,('WP_057556016.1-Vibrio-cholerae':0.04679999999999973,'WP_188234612.1-Vibrio-sp.-Y2-5':0.02898999999999985)1.0:0.1915499999999999)0.164:0.020640000000000214,'WP_039484180.1-Vibrio-sinaloensis':0.18162000000000011)0.392:0.011109999999999953,(('WP_012549058.1-Aliivibrio-salmonicida':0.1538400000000002,('WP_011497090.1-Shewanella-denitrificans':0.21426999999999996,'WP_223418564.1-Shewanella-inventionis':0.07231999999999994)0.981:0.12287000000000026)0.0:0.000540,'WP_129120576.1-Veronia-nyctiphanis':0.18456000000000028)0.907:0.0379299999999998)0.846:0.02945999999999982)0.752:0.01409000000000038,('WP_017031451.1-Vibrio-breoganii':0.07808999999999999,'WP_248975945.1-Shewanella-kaireitica':0.16505999999999998)0.954:0.06701000000000024)0.44:0.01269000000000009)0.946:0.061859999999999804,((((('WP_042868619.1-Dickeya':0.10067000000000004,('WP_035339894.1-Dickeya':0.018739999999999757,'WP_125258312.1-Dickeya-lacustris':0.08545000000000025)1.0:0.16525999999999996)0.954:0.07559000000000005,((((('WP_129946484.1-Shewanella-sp.-OPT22':0.05118,'WP_205000141.1-Parashewanella-hymeniacidonis':0.05553000000000008)1.0:0.5061799999999996,('WP_133132051.1-Legionella-sp.-W10-070':0.23520999999999992,('WP_058513550.1-Legionella-santicrucis':0.10104999999999986,((('WP_232890831.1-unclassified-Legionella':0.039309999999999956,('WP_058459148.1-Legionella-bozemanae':0.02157000000000009,'WP_058518697.1-Legionella-parisiensis':0.043930000000000025)0.864:0.026959999999999873)0.996:0.0871900000000001,(('WP_031567420.1-Legionella-wadsworthii':0.18455999999999984,(('WP_027266778.1-Legionella-pneumophila':0.032480000000000064,('WP_027220106.1-Legionella-pneumophila':0.014310000000000045,('WP_044499121.1-Legionella-pneumophila':0.023050000000000015,(('WP_265717942.1-Legionella-sp.-PATHC032':0.033949999999999925,'WP_027223798.1-Legionella-pneumophila':0.02972000000000019)0.829:0.027190000000000047,(('WP_061513621.1-Legionella':0.015730000000000022,'WP_027226296.1-Legionella-pneumophila':0.04773000000000005)0.994:0.0609099999999998,'WP_106142999.1-Legionella-pneumophila':0.033949999999999925)0.0:0.000540)0.874:0.011540000000000106)0.898:0.019890000000000185)0.812:0.016300000000000203)0.972:0.07168999999999981,('WP_058494182.1-Legionella-worsleiensis':0.13310999999999984,('WP_127106294.1-Legionella-sp.-km535':0.03865000000000007,'WP_058474153.1-Legionella-quateirensis':0.0284399999999998)0.898:0.03718000000000021)0.953:0.06828000000000012)0.934:0.06336999999999993)0.676:0.016669999999999963,'WP_045097199.1-Legionella-fallonii':0.11526000000000014)0.873:0.024849999999999817)0.415:0.020529999999999937,('WP_058463758.1-Legionella-cincinnatiensis':0.01957999999999993,'WP_128130294.1-Legionella-sainthelensi':0.04972999999999983)0.943:0.03354999999999997)0.564:0.000540)0.997:0.30934000000000017)0.985:0.24929999999999986)0.839:0.06827000000000005,((((((((((('WP_084055900.1-Desulfacinum-hydrothermale':0.29197000000000006,'WP_123289658.1-Desulfosoma-caldarium':0.25680000000000014)0.991:0.21016999999999975,(('WP_048706771.1-Thauera-humireducens':0.20700999999999992,('WP_018865194.1-Thioalkalivibrio-sp.-ARh3':0.20683000000000007,('WP_116346290.1-Cupriavidus-taiwanensis':0.1596000000000002,'WP_124685047.1-Cupriavidus-pauculus':0.16964000000000024)0.999:0.3646799999999999)0.757:0.06454000000000004)0.924:0.08332000000000006,(('WP_119020982.1-Halomonas-sp.-JS92-SW72':0.22603000000000018,'WP_227390214.1-Halomonas-malpeensis':0.5604200000000001)0.948:0.17222999999999988,(((('WP_111826915.1-Achromobacter-sp.-HZ01':0.10993999999999993,'WP_102775120.1-Achromobacter':0.018520000000000092)0.997:0.12832,('WP_129525645.1-Achromobacter-agilis':0.02958000000000016,('WP_129240460.1-Achromobacter-veterisilvae':0.040140000000000065,'WP_043544564.1-Achromobacter-sp.-RTa':0.026419999999999888)0.864:0.01617999999999986)0.766:0.013959999999999972)0.978:0.05875999999999992,('WP_174717299.1-Achromobacter-denitrificans':0.07518000000000002,('WP_175208175.1-Achromobacter-anxifer':0.03960000000000008,((('WP_207472676.1-Achromobacter-sp.-SD115':0.010310000000000041,('WP_164793551.1-Achromobacter-insolitus':0.029739999999999878,'WP_251864322.1-Achromobacter-sp.-Marseille-Q4962':0.06633000000000022)0.978:0.052059999999999995)0.897:0.029770000000000074,'WP_125471167.1-Achromobacter':0.018219999999999903)0.947:0.041180000000000216,('WP_088158656.1-Achromobacter-xylosoxidans':0.08580999999999994,'WP_175164188.1-Achromobacter-aegrifaciens':0.03354999999999997)0.756:0.013170000000000126)0.383:0.020509999999999806)0.997:0.10765000000000002)0.748:0.011379999999999946)0.787:0.0277099999999999,(('WP_088588627.1-Achromobacter-marplatensis':0.13773999999999997,'WP_169273406.1-Achromobacter-sp.-Bel':0.07494999999999985)0.954:0.07577999999999996,('WP_200222041.1-Achromobacter-xylosoxidans':0.13999000000000006,'WP_259251071.1-Achromobacter-sp.-JUb104':0.16122000000000014)0.934:0.06816000000000022)0.729:0.04068000000000005)0.941:0.11940000000000017)0.173:0.06089000000000011)0.671:0.02049999999999974)0.42:0.023090000000000277,(('WP_212718705.1-unclassified-Synechocystis':0.000530,('WP_228015966.1-Synechocystis-salina':0.09143999999999997,'WP_010873214.1-unclassified-Synechocystis':0.06369999999999987)0.053:0.02425999999999995)1.0:0.3875000000000002,(('WP_199264031.1-Desulfobulbus-oligotrophicus':0.25184000000000006,'WP_153720048.1-Spiribacter-sp.-C176':0.25726000000000004)0.042:0.05683000000000016,('WP_099797666.1-Thermostichus-lividus':0.3659499999999998,'WP_196167318.1-Pseudomonas-monteilii':0.28937999999999997)0.871:0.09236000000000022)0.835:0.051369999999999916)0.89:0.06653999999999982)0.791:0.02126000000000028,('WP_221248821.1-Desulfuromonas-versatilis':0.2679299999999998,'WP_103114677.1-Geothermobacter-hydrogeniphilus':0.2721999999999998)0.643:0.10430000000000028)0.427:0.0289299999999999,(((((('WP_027458229.1-Dechloromonas-agitata':0.11052000000000017,'WP_066884254.1-Dechloromonas':0.09043000000000001)0.975:0.09283999999999981,('WP_117174481.1-Rhodoferax-lacus':0.2232099999999999,('WP_214121631.1-Curvibacter-sp.-CHRR-16':0.2400500000000001,('WP_105262671.1-Rhodoferax-sp.-TS-BS-61-7':0.028140000000000054,'WP_094482496.1-Rhodoferax-sp.-TH121':0.08591999999999977)0.894:0.0464500000000001)0.956:0.11073000000000022)0.617:0.0354000000000001)0.914:0.057949999999999946,((('WP_028317018.1-Desulfobulbus-elongatus':0.2911600000000001,('WP_148894253.1-Geothermobacter-ehrlichii':0.2937400000000001,(('WP_155876508.1-Desulfuromonas-sp.-AOP6':0.26785000000000014,('WP_072282603.1-Syntrophotalea-acetylenivorans':0.14415999999999984,'WP_092056970.1-Desulfuromonas-acetexigens':0.1665899999999998)0.833:0.06191000000000013)0.817:0.04142000000000001,'WP_043770486.1-Desulfobulbus-propionicus':0.22513000000000005)0.721:0.06203000000000003)0.772:0.03657000000000021)0.786:0.05506000000000011,(('WP_153584563.1-Caenimonas-koreensis':0.23864000000000019,('WP_007185580.1-Hydrocarboniphaga-effusa':0.0719599999999998,'WP_007185565.1-Hydrocarboniphaga-effusa':0.06205999999999978)0.987:0.17559000000000013)0.982:0.13361,(('WP_201212349.1-Rhodocyclus-purpureus':0.16903000000000024,('WP_167682126.1-Rhodocyclus-gracilis':0.2645599999999999,('WP_153115862.1-Rhodocyclus-tenuis':0.039120000000000044,'WP_200385811.1-Rhodocyclus-tenuis':0.021729999999999805)0.957:0.08426)0.47:0.03617999999999988)0.936:0.11380999999999997,(('WP_220807781.1-Noviherbaspirillum-aridicola':0.4921500000000001,'WP_017324474.1-Synechococcus-sp.-PCC-7336':0.31103999999999976)0.319:0.06199000000000021,('WP_226989104.1-Desulfuromonas-sp.-TF':0.3778600000000001,'WP_227459030.1-Cupriavidus-pauculus':0.3105500000000001)0.808:0.11543999999999999)0.873:0.06123000000000012)0.717:0.02032999999999996)0.786:0.026040000000000063)0.926:0.07134000000000018,(('WP_153110537.1-Propionivibrio-limicola':0.22022999999999993,('WP_136386324.1-Pseudothauera-rhizosphaerae':0.1361699999999999,('WP_187718084.1-Thauera-sp.-CAU-1555':0.13668000000000013,'WP_114649035.1-Pseudothauera-hydrothermalis':0.08641999999999994)0.604:0.0650599999999999)0.989:0.15465000000000018)0.86:0.0779399999999999,(((((('WP_134696535.1-Aeromonas-taiwanensis':0.5776599999999998,(('WP_160351730.1-Bordetella-sp.-15P40C-2':0.1601499999999998,'WP_094838100.1-Bordetella-genomosp.-4':0.21984000000000004)0.978:0.20362999999999998,('WP_019964198.1-Pannonibacter-phragmitetus':0.52075,('WP_026470910.1-Alkanindiges-illinoisensis':0.12356999999999996,('WP_248147450.1-Moraxella-osloensis':0.32669000000000015,'WP_092690500.1-Acinetobacter-kyonggiensis':0.19030000000000014)0.891:0.10448000000000013)0.998:0.3515999999999999)0.737:0.06558999999999982)0.713:0.0549599999999999)0.874:0.13065999999999978,'WP_148717329.1-Chitinolyticbacter-meiyuanensis':0.5497700000000001)0.534:0.04565999999999981,('WP_243478771.1-Orrella-sp.-f23':0.39378,'WP_108623287.1-Orrella-marina':0.28306999999999993)0.946:0.1190199999999999)0.587:0.06354999999999977,'WP_121459656.1-Azonexus-fungiphilus':0.18511999999999995)0.862:0.06888000000000005,'WP_205202385.1-Azonexus-hydrophilus':0.2035499999999999)0.478:0.057010000000000005,'WP_173581083.1-Azonexus-fungiphilus':0.22224999999999984)0.857:0.037659999999999805)0.39:0.012410000000000032)0.542:0.020610000000000017)0.865:0.045270000000000366,(('WP_227683402.1-Pusillimonas-sp.-MFBS29':0.2595299999999998,('WP_044615503.1-Gynuella-sunshinyii':0.41764,'WP_092579398.1-Achromobacter-sp.-NFACC18-2':0.3659699999999999)0.7:0.07718999999999987)0.829:0.05094000000000021,(((('WP_064030819.1-Methylomonas-koyamae':0.20787999999999984,((('WP_140911374.1-Methylomonas-koyamae':0.20523999999999987,('WP_071159776.1-Methylomonas-sp.-LWB':0.15225,('WP_036273427.1-Methylomonas':0.11072000000000015,('WP_020485671.1-Methylomonas-sp.-MK1':0.18644000000000016,('WP_256607223.1-Methylomonas-sp.-WSC-7':0.0651600000000001,('WP_205450646.1-Methylomonas-sp.-EFPC1':0.07886999999999977,'WP_064036612.1-Methylomonas-methanica':0.000540)0.728:0.006839999999999957)0.85:0.04551999999999978)0.884:0.057399999999999896)0.679:0.01432000000000011)0.793:0.03448999999999991)0.851:0.04823999999999984,('WP_269022943.1-Methylomonas-sp.-MP1':0.1005600000000002,(('WP_066983474.1-Methylomonas-lenta':0.09082999999999997,'WP_150047608.1-Methylomonas-rhizoryzae':0.27143000000000006)0.819:0.05705999999999989,'WP_196437371.1-Methylomonas-sp.-LL1':0.11308000000000007)0.853:0.048290000000000166)0.917:0.06450999999999985)0.8:0.02856000000000014,('WP_148817865.1-Pigmentiphaga-aceris':0.30298000000000025,('WP_004074545.1-Desulfobacter-postgatei':0.23892999999999986,(('WP_168987795.1-Azoarcus-taiwanensis':0.2544599999999999,('WP_256068944.1-Stutzerimonas-stutzeri':0.1045799999999999,('WP_013164002.1-Desulfurivibrio-alkaliphilus':0.25511000000000017,'WP_026685886.1-Azovibrio-restrictus':0.15113999999999983)0.884:0.05464000000000002)0.15:0.04016999999999982)0.843:0.044459999999999944,(('WP_183165449.1-Azomonas-macrocytogenes':0.13938000000000006,('WP_238622182.1-unclassified-Ectothiorhodospira':0.11246,('WP_025282248.1-Ectothiorhodospira-haloalkaliphila':0.10368999999999984,'WP_238617884.1-Ectothiorhodospira-shaposhnikovii':0.13093999999999983)0.949:0.07474999999999987)0.913:0.07026999999999983)0.788:0.030199999999999783,('WP_060482506.1-Pseudomonas-sp.-NBRC-111119':0.15416000000000007,(('WP_115272924.1-Pseudomonas-putida':0.10514000000000001,('WP_046489762.1-Pseudomonas-veronii':0.12101000000000006,('WP_059395284.1-Pseudomonas-putida':0.09730999999999979,('WP_009685980.1-Pseudomonas':0.07911000000000001,'WP_024086460.1-Pseudomonas':0.044340000000000046)0.878:0.027610000000000134)0.888:0.028200000000000003)0.874:0.02939999999999987)0.843:0.028830000000000133,((('WP_201164743.1-Pseudomonas-sp.-S60':0.03901000000000021,'WP_196129978.1-Pseudomonas-fulva':0.07261000000000006)0.987:0.08478999999999992,(('WP_202675963.1-Pseudomonas-sp.-SK2':0.03244999999999987,'WP_023379297.1-Pseudomonas-sp.-VLB120':0.08931999999999984)0.99:0.06862000000000013,('WP_075044077.1-Pseudomonas-putida':0.04222999999999999,('WP_174142546.1-Pseudomonas-putida':0.12548000000000004,'WP_027907237.1-Pseudomonas':0.032810000000000006)0.652:0.009749999999999925)0.703:0.02237)0.905:0.03499000000000008)0.907:0.02856000000000014,((('WP_267821885.1-Pseudomonas-putida':0.06885999999999992,((('WP_186476580.1-Pseudomonas-promysalinigenes':0.038590000000000124,'WP_167339957.1-Pseudomonas-fulva':0.027169999999999916)0.859:0.01957999999999993,(('WP_201214098.1-Pseudomonas-sp.-S36':0.011810000000000098,'WP_054890059.1-Pseudomonas':0.061189999999999856)0.972:0.04837999999999987,('WP_110607173.1-Pseudomonas-fulva':0.012179999999999858,'WP_088523134.1-unclassified-Pseudomonas':0.043950000000000156)0.905:0.02251000000000003)0.849:0.01476999999999995)0.813:0.013339999999999907,('WP_182365213.1-Pseudomonas':0.09565000000000001,('WP_159409308.1-Pseudomonas-putida':0.13176999999999994,'WP_182365203.1-Pseudomonas-juntendi':0.05494000000000021)0.478:0.014800000000000146)0.745:0.00735000000000019)0.894:0.026250000000000107)0.741:0.000950,('WP_103449615.1-Pseudomonas-putida':0.054300000000000015,'WP_167277242.1-Pantoea-sp.-Tr-811':0.011870000000000047)0.985:0.05685999999999991)0.497:0.0142199999999999,('WP_029885231.1-Pseudomonas':0.0265200000000001,('WP_060394622.1-Pseudomonas-entomophila':0.14912999999999998,('WP_197863301.1-Pseudomonas-putida':0.035600000000000076,'WP_230064532.1-Pseudomonas-sp.-CIP-10':0.019930000000000003)0.434:0.0259999999999998)0.472:0.012030000000000207)0.968:0.04296999999999995)0.816:0.000540)0.83:0.019179999999999975)0.184:0.03401999999999994)0.976:0.09025000000000016)0.485:0.013520000000000199)0.756:0.026370000000000005)0.39:0.0571299999999999)0.918:0.06855000000000011)0.598:0.02915000000000001)0.72:0.017990000000000173,('WP_138632053.1-Pseudoalteromonas-spongiae':0.2587099999999998,('WP_151054911.1-Cellvibrio-sp.-KY-YJ-3':0.1721499999999998,('WP_078044787.1-Cellvibrio-sp.-PSBB023':0.2160099999999998,('WP_087465811.1-Cellvibrio-sp.-PSBB006':0.1675399999999998,'WP_012489013.1-Cellvibrio-japonicus':0.1529799999999999)0.488:0.05177999999999994)0.103:0.054790000000000116)0.911:0.10607999999999995)0.967:0.11378999999999984)0.927:0.08348000000000022,((('WP_240716217.1-Halomonas-neptunia':0.10240999999999989,'WP_218634960.1-Marinobacter-arenosus':0.10424999999999995)0.914:0.08241999999999994,('WP_058575749.1-unclassified-Guyparkeria':0.25712,(('WP_250059255.1-Halomonas-sp.-ATCH28':0.2814199999999998,'WP_179153381.1-Chromohalobacter-salexigens':0.19228999999999985)0.85:0.06599000000000022,'WP_250939176.1-unclassified-Larsenimonas':0.2585600000000001)0.929:0.10057999999999989)0.857:0.06315999999999988)0.617:0.06868000000000007,(('WP_139871681.1-Candidatus-Methylopumilus-universalis':0.21274999999999977,('WP_140005423.1-Candidatus-Methylopumilus-planktonicus':0.07853000000000021,'WP_139929800.1-Candidatus-Methylopumilus-universalis':0.26932)0.891:0.07934999999999981)0.979:0.15832999999999986,('WP_018896487.1-Rhizobium-sp.-2MFCol3.1':0.15592000000000006,(('WP_233136718.1-Agrobacterium-vaccinii':0.23905999999999983,('WP_184450570.1-Rhizobium-sp.-BK538':0.16986999999999997,'WP_153273632.1-Rhizobium-grahamii':0.22848999999999986)0.075:0.03491)0.081:0.022050000000000125,(('WP_220354719.1-Rhizobium-cauense':0.20816999999999997,'WP_016557527.1-Rhizobium-grahamii':0.12003999999999992)0.688:0.037519999999999776,('WP_007537558.1-Rhizobium-mesoamericanum':0.05604999999999993,'WP_072373837.1-Rhizobium-tibeticum':0.07824000000000009)0.895:0.04978000000000016)0.821:0.029180000000000206)0.876:0.040280000000000094)0.988:0.13179000000000007)0.901:0.08143000000000011)0.886:0.08415999999999979)0.953:0.06918999999999986,(('WP_230502293.1-Janthinobacterium-sp.-FW305-128':0.37094000000000005,'WP_249918475.1-Pseudomonas-sp.-R3.Fl':0.17074999999999996)0.877:0.07227000000000006,(((((('WP_047550623.1-Methylotenera-sp.-G11':0.19368000000000007,'WP_212592286.1-Janthinobacterium-sp.-TND4EL3':0.23137000000000008)0.485:0.06391999999999998,'WP_217963608.1-Candidimonas-humi':0.2627099999999998)0.644:0.03143999999999991,(('WP_102071501.1-Pollutimonas-nitritireducens':0.2533599999999998,('WP_136867012.1-Guyparkeria-sp.-SB14A':0.37629,'WP_102075761.1-Pollutimonas-subterranea':0.14110999999999985)0.751:0.028890000000000082)0.806:0.029360000000000053,('WP_066098770.1-Halothiobacillus-sp.-LS2':0.1208499999999999,'WP_070073009.1-Acidihalobacter-aeolianus':0.2035800000000001)0.964:0.10270000000000001)0.773:0.012890000000000068)0.918:0.03705999999999987,('WP_180152801.1-Parapusillimonas-granuli':0.13771999999999984,(('WP_057247240.1-Duganella-sp.-Root198D2':0.22894000000000014,'WP_155457065.1-Massilia-eburnea':0.08238000000000012)0.944:0.08097000000000021,(('WP_192051557.1-unclassified-Massilia':0.10877999999999988,'WP_136222466.1-Massilia-sp.-Mn16-1_5':0.09078000000000008)0.907:0.0737000000000001,('WP_225237063.1-Massilia-oculi':0.2938700000000001,'WP_147937383.1-Massilia-arenae':0.10257000000000005)0.976:0.13321000000000005)0.903:0.07098000000000004)0.845:0.023190000000000044)0.897:0.052589999999999915)0.713:0.00658000000000003,('WP_058422493.1-Pseudomonas':0.19984000000000002,'WP_223462047.1-Pseudomonas-sp.-A-R-26':0.18125999999999998)0.628:0.07898000000000005)0.936:0.043070000000000164,((((('WP_020564242.1-Methylosarcina-fibrata':0.19162999999999997,(('WP_046857648.1-Pseudomonas-sp.-CCOS-191':0.15768000000000004,(('WP_068803504.1-Immundisolibacter-cernigliae':0.19984000000000002,'WP_074973890.1-Pseudomonas-otitidis':0.15626000000000007)0.287:0.040059999999999985,('WP_019141909.1-Noviherbaspirillum-massiliense':0.14973999999999998,'WP_146330669.1-Noviherbaspirillum-sp.-UKPF54':0.2190500000000002)0.868:0.06389999999999985)0.888:0.045949999999999935)0.28:0.016719999999999846,'WP_011111299.1-Nitrosomonas-europaea':0.18533)0.896:0.0541100000000001)0.862:0.03128999999999982,(('WP_101152997.1-Pseudomonas-sp.-Choline-3u-10':0.1048,('WP_003299494.1-Stutzerimonas-stutzeri':0.07641000000000009,('WP_046617631.1-Stutzerimonas-stutzeri':0.06865999999999994,'WP_135265705.1-Stutzerimonas-stutzeri':0.041770000000000085)0.952:0.07954999999999979)0.854:0.03083999999999998)0.893:0.04488000000000003,('WP_125865093.1-Stutzerimonas-stutzeri':0.16898999999999997,'WP_200290630.1-Stutzerimonas-balearica':0.10339000000000009)0.951:0.0700099999999999)0.786:0.016570000000000196)0.874:0.022159999999999958,((('WP_076518421.1-Achromobacter-sp.-MFA1-R4':0.06864999999999988,('WP_104142565.1-Achromobacter-spanius':0.026959999999999873,'WP_120157261.1-Achromobacter-sp.-B7':0.0452300000000001)0.934:0.04188999999999998)0.99:0.10570000000000013,((('WP_219210863.1-Variovorax-boronicumulans':0.15164,('WP_192194832.1-Variovorax-sp.-38R':0.1933600000000002,(('WP_126747644.1-Variovorax-sp.-DXTD-1':0.1439499999999998,('WP_093207264.1-Variovorax-sp.-YR750':0.10372000000000003,'WP_206177103.1-Variovorax-sp.-RKNM96':0.0943900000000002)0.324:0.0398900000000002)0.802:0.03216000000000019,('WP_160553990.1-Xylophilus-rhododendri':0.15926000000000018,'WP_093441791.1-Variovorax-sp.-770b2':0.13924000000000003)0.156:0.0381800000000001)0.907:0.052119999999999944)0.695:0.013430000000000053)0.962:0.09417000000000009,'WP_166067558.1-Diaphorobacter-sp.-HDW4B':0.14461000000000013)0.165:0.04465999999999992,'WP_094844257.1-Bordetella-genomosp.-11':0.22332000000000019)0.652:0.06184000000000012)0.837:0.023149999999999782,('WP_209872004.1-Pseudomonas-sp.-BP8':0.17384000000000022,'WP_066093302.1-Hydrogenophaga-crassostreae':0.1670600000000002)0.798:0.05155999999999983)0.883:0.02876000000000012)0.749:0.008840000000000181,(('WP_184594840.1-Pseudomonas-nitritireducens':0.16782999999999992,'WP_192381882.1-Pseudomonas-sp.-PDM20':0.10985000000000023)0.995:0.14305000000000012,('WP_233297479.1-Pseudomonas':0.1506099999999999,(('WP_191471912.1-Pseudomonas-asiatica':0.17933999999999983,('WP_043318452.1-Pseudomonas-sp.-PI1':0.044620000000000104,'WP_043268312.1-Pseudomonas':0.027120000000000033)0.573:0.02417999999999987)0.948:0.05874999999999986,('WP_076426141.1-Pseudomonas-alcaligenes':0.1341800000000002,'WP_089390899.1-Pseudomonas-delhiensis':0.11237999999999992)0.377:0.020399999999999974)0.957:0.06267999999999985)0.821:0.028249999999999886)0.891:0.047210000000000196)0.084:0.017710000000000115,(('WP_249615874.1-Craterilacuibacter-sp.-RT1T':0.4926499999999998,('WP_077733980.1-Paenalcaligenes-hominis':0.03779000000000021,'WP_167661700.1-Paenalcaligenes-hominis':0.0283199999999999)0.977:0.16443999999999992)0.894:0.09148999999999985,(((((('WP_105555808.1-Achromobacter-sp.-MYb9':0.07419999999999982,'WP_088146851.1-Achromobacter-denitrificans':0.09619)0.987:0.12361999999999984,('WP_085721622.1-Pseudomonas':0.15300000000000002,(('WP_236964528.1-Hydrogenophaga-sp.-SL48':0.06847000000000003,('WP_226493830.1-Hydrogenophaga-taeniospiralis':0.03860999999999981,'WP_210824367.1-Hydrogenophaga-aromaticivorans':0.05702000000000007)0.812:0.048109999999999875)0.966:0.0791599999999999,('WP_177138191.1-Hydrogenophaga-aromaticivorans':0.1248499999999999,'WP_133157403.1-Hydrogenophaga-pseudoflava':0.1469999999999998)0.072:0.015600000000000058)0.942:0.06752000000000002)0.16:0.018289999999999917)0.717:0.010250000000000092,('WP_106478747.1-Phytohalomonas-tamaricis':0.26742,('WP_017517140.1-Pseudomonas-nitroreducens':0.010050000000000114,'WP_138213821.1-Pseudomonas-nitroreducens':0.044459999999999944)0.997:0.1916199999999999)0.918:0.07505000000000006)0.936:0.04923000000000011,'WP_104727848.1-Pseudomonas-oleovorans':0.15730000000000022)0.696:0.02461000000000002,((('WP_169965430.1-unclassified-Pseudomonas':0.04249999999999998,'WP_047278503.1-Pseudomonas-lundensis':0.023320000000000007)0.997:0.1324700000000001,('WP_205489922.1-Pseudomonas-lactucae':0.14622000000000002,('WP_042945119.1-Pseudomonas-fluorescens':0.061249999999999805,'WP_181643258.1-Pseudomonas-sivasensis':0.048939999999999984)0.999:0.12064999999999992)0.165:0.02082000000000006)0.695:0.013640000000000096,'WP_065086507.1-Pseudomonas':0.26907999999999976)0.734:0.013259999999999827)0.837:0.02245000000000008,('WP_214501190.1-Pseudomonas-sp.-RC3H12':0.2693599999999998,((('WP_029241333.1-Pseudomonas':0.05717999999999979,('WP_125830676.1-Pseudomonas-sp.-v388':0.06495000000000006,('WP_054996663.1-Pseudomonas-ficuserectae':0.038899999999999935,('WP_025390602.1-Pseudomonas-syringae':0.019320000000000004,'WP_007252556.1-Pseudomonas-syringae-group':0.04416999999999982)0.361:0.019239999999999924)0.94:0.04499999999999993)0.886:0.04856000000000016)0.93:0.08122000000000007,(('WP_141063556.1-Pseudomonas-fluorescens':0.08858999999999995,'WP_177015116.1-Pseudomonas-costantinii':0.04508000000000001)0.881:0.055709999999999926,('WP_016973326.1-Pseudomonas-tolaasii':0.10463999999999984,'WP_099548838.1-Pseudomonas-sp.-2995-3':0.08818999999999999)0.829:0.03980000000000006)0.833:0.061659999999999826)0.992:0.15965000000000007,(('WP_043256200.1-Pseudomonas-knackmussii':0.11997999999999998,'WP_058488093.1-Pseudomonas':0.1470699999999998)0.74:0.04773000000000005,('WP_205338011.1-Pseudomonas-sp.-PDNC002':0.051349999999999785,('WP_088420173.1-Pseudomonas-nitroreducens':0.12681999999999993,'WP_254475170.1-unclassified-Pseudomonas':0.06709999999999994)0.845:0.028490000000000126)0.997:0.14109000000000016)0.845:0.037760000000000016)0.806:0.03006999999999982)0.678:0.01817000000000002)0.723:0.010190000000000143)0.749:0.009240000000000137)0.882:0.0319799999999999)0.451:0.008319999999999883)0.812:0.01662000000000008)0.188:0.012939999999999952)0.931:0.04642000000000035)0.759:0.019769999999999843,(('WP_040097704.1-Geoalkalibacter-ferrihydriticus':0.30552,'WP_054772520.1-Methylogaea-oryzae':0.16564999999999985)0.779:0.049900000000000055,('WP_209394989.1-Rhodoferax-sp.-AJA081-3':0.2898399999999999,(('WP_263768642.1-Propionivibrio-sp.-SG131':0.2533599999999998,('WP_003153917.1-Pseudomonas-aeruginosa-group':0.04518999999999984,'WP_033955816.1-Pseudomonas-aeruginosa':0.007410000000000139)0.994:0.1954600000000002)0.86:0.04922000000000004,'WP_158658301.1-Achromobacter-sp.-AONIH1':0.4072800000000001)0.419:0.01269000000000009)0.845:0.0447700000000002)0.855:0.02715999999999985)0.808:0.02049999999999974,((((((('WP_019918777.1-Methyloversatilis-discipulorum':0.04449000000000014,'WP_018229738.1-Methyloversatilis-universalis':0.04992999999999981)0.982:0.13832999999999984,('WP_095523004.1-Candidatus-Dactylopiibacterium-carminicum':0.09623999999999988,'WP_101941536.1-Uliginosibacterium-sp.-TH139':0.0964299999999998)0.97:0.11534999999999984)0.992:0.16010999999999997,(('WP_036252969.1-Methylobacter-sp.-BBA5.1':0.25951999999999975,'WP_212785890.1-Ferrigenium-kumadai':0.21801000000000004)0.784:0.032239999999999824,'WP_174484170.1-methanotrophic-endosymbiont-of-Bathymodiolus-puteoserpentis':0.21550000000000002)0.561:0.03215000000000012)0.135:0.000920,(((('WP_223986959.1-Cupriavidus-pampae':0.048249999999999904,('WP_082079449.1-Cupriavidus-basilensis':0.07576999999999989,('WP_011517306.1-Cupriavidus-metallidurans':0.04675999999999991,'WP_019451253.1-Cupriavidus-sp.-BIS7':0.0485199999999999)0.599:0.000540)0.872:0.029570000000000096)1.0:0.1696200000000001,((('WP_169420802.1-Ramlibacter-agri':0.1811600000000002,'WP_057673948.1-Curvibacter-sp.-PAE-UM':0.15920000000000023)0.851:0.038879999999999804,('WP_056898839.1-Pseudorhodoferax-sp.-Leaf274':0.23186000000000018,('WP_068630051.1-Variovorax-sp.-PAMC-28711':0.21963,'WP_106935816.1-unclassified-Variovorax':0.19560000000000022)0.653:0.055379999999999985)0.862:0.05046000000000017)0.728:0.010359999999999925,('WP_137918285.1-Hydrogenophaga-sp.-2FB':0.15504000000000007,('WP_137890779.1-Ramlibacter-sp.-2FC':0.10110999999999981,('WP_086926522.1-Acidovorax-carolinensis':0.16459000000000001,'WP_026436726.1-Acidovorax-sp.-JHL-9':0.18135000000000012)0.889:0.06013000000000002)0.815:0.031019999999999825)0.635:0.00943000000000005)0.187:0.0022400000000000198)0.0:0.000550,(('WP_074796726.1-Nitrosospira-briensis':0.06706000000000012,('WP_097065525.1-Nitrosovibrio-sp.-Nv4':0.02815000000000012,'WP_090882638.1-Nitrosovibrio-sp.-Nv6':0.05259999999999998)0.871:0.046619999999999884)0.998:0.20988000000000007,('WP_122244186.1-Vandammella-animalimorsus':0.15460999999999991,'WP_220807643.1-Noviherbaspirillum-aridicola':0.29768000000000017)0.808:0.07565)0.817:0.050060000000000215)0.922:0.048210000000000086,(('WP_253536546.1-Ralstonia-sp.-21YRMH01-3':0.3606699999999998,'WP_154206467.1-Ralstonia-pickettii':0.14373000000000014)0.801:0.08200999999999992,((('WP_066475960.1-Comamonas-terrae':0.1180699999999999,('WP_058879441.1-Comamonas-kerstersii':0.10571999999999981,('WP_012203208.1-Delftia-acidovorans':0.18040000000000012,('WP_218241177.1-Comamonas-fluminis':0.10158000000000023,(('WP_169107474.1-Comamonas-suwonensis':0.03988000000000014,'WP_219343559.1-Comamonas-sp.-Y33R10-2':0.08137999999999979)0.916:0.04352,('WP_003073420.1-Comamonas':0.06585999999999981,('WP_034391098.1-Comamonas-thiooxydans':0.014660000000000117,('WP_148729134.1-Comamonas-sp.-Z3':0.039750000000000174,'WP_211457163.1-Comamonas-sp.-PE63':0.04527999999999999)0.492:0.006069999999999798)0.748:0.008529999999999927)0.887:0.026670000000000194)0.788:0.018079999999999874)0.994:0.15264999999999995)0.133:0.0477599999999998)0.904:0.07501999999999986)0.994:0.23145000000000016,(('WP_047193087.1-Schlegelella-brevitalea':0.25493999999999994,('WP_074865046.1-Atopomonas-hussainii':0.2526600000000001,'WP_019561187.1-Caldimonas-manganoxidans':0.17031000000000018)0.406:0.08395000000000019)0.279:0.03948000000000018,(('WP_230291920.1-Croceicoccus-sp.-Ery5':0.42673000000000005,'WP_164156863.1-Sandarakinorhabdus-rubra':0.16265000000000018)0.96:0.11208000000000018,(('WP_037482909.1-Sphaerotilus-natans':0.17386999999999997,('WP_028996355.1-Azohydromonas-australica':0.07877,'WP_066337818.1-Azohydromonas-lata':0.07769000000000004)0.957:0.0941200000000002)0.891:0.07787999999999995,('WP_241396682.1-Pelomonas-sp.-CA6':0.11891000000000007,'WP_198111724.1-Inhella-proteolytica':0.39159999999999995)0.435:0.020319999999999894)0.962:0.10848000000000013)0.88:0.06271000000000004)0.276:0.04391999999999996)0.831:0.056239999999999846,(('WP_025281494.1-Ectothiorhodospira-haloalkaliphila':0.39449999999999985,('WP_176062241.1-Anaeromyxobacter-diazotrophicus':0.15812999999999988,'WP_248361410.1-Anaeromyxobacter-oryzae':0.22953)0.959:0.12795999999999985)0.75:0.04468000000000005,('WP_123210144.1-Stagnimonas-aquatica':0.7505700000000002,'WP_160809027.1-Nitrosomonas-sp.-HPC101':0.3205100000000001)0.5:0.07177999999999995)0.723:0.0921599999999998)0.773:0.029939999999999856)0.887:0.059410000000000185)0.893:0.03794999999999993)0.929:0.045869999999999855,(('WP_109041087.1-Azospira-sp.-I13':0.0983299999999998,'WP_014237023.1-Azospira-oryzae':0.11004000000000014)0.972:0.09859000000000018,'WP_201673630.1-Ramlibacter-monticola':0.23353000000000002)0.026:0.04136000000000006)0.717:0.02787000000000006,((('WP_066641528.1-Bordetella-sp.-H567':0.06976000000000004,('WP_086058993.1-Bordetella-genomosp.-9':0.15666000000000002,'WP_094856450.1-Bordetella-genomosp.-10':0.14747999999999983)0.954:0.09771999999999981)0.851:0.06267999999999985,('WP_086067427.1-Bordetella':0.19286000000000003,('WP_057654143.1-Bordetella-sp.-N':0.15198,('WP_088599355.1-unclassified-Achromobacter':0.0654300000000001,'WP_129151777.1-Achromobacter-aloeverae':0.08883000000000019)0.983:0.1146799999999999)0.701:0.05575999999999981)0.904:0.04985999999999979)0.981:0.10349999999999993,(('WP_091467897.1-Methylophilus-rhizosphaerae':0.28222999999999976,(('WP_100503206.1-Achromobacter-ruhlandii':0.13095999999999997,(('WP_219991400.1-Achromobacter-sp.-ES-001':0.07568000000000019,'WP_006216420.1-Achromobacter-piechaudii':0.1102599999999998)0.944:0.051120000000000054,'WP_183017169.1-Achromobacter-sp.-UMC46':0.04720000000000013)0.882:0.031260000000000066)0.615:0.04752000000000001,'WP_175192416.1-Achromobacter-deleyi':0.04120999999999997)0.992:0.2161200000000001)0.61:0.06296999999999997,'WP_067106371.1-Methylobacillus-sp.-MM3':0.1506400000000001)0.89:0.0639299999999996)0.783:0.024180000000000312)0.959:0.04903999999999975,('WP_073615262.1-Desulfopila-aestuarii':0.21369000000000016,('WP_136805251.1-Desulfosediminicola-flagellatus':0.24212999999999996,'WP_163339734.1-Desulfopila-sp.-IMCC35008':0.27901)0.794:0.04186999999999985)0.99:0.13575999999999988)0.0:0.000540)0.927:0.028400000000000425)0.779:0.012109999999999843,((('WP_104485062.1-Oceanisphaera-arctica':0.09353999999999996,('WP_188630571.1-Oceanisphaera-marina':0.12333000000000016,'WP_204795415.1-Oceanisphaera-litoralis':0.03555000000000019)0.094:0.017440000000000122)1.0:0.17743999999999982,((((('WP_077021900.1-Burkholderia-cenocepacia':0.07642999999999978,('WP_174919663.1-Burkholderia-metallica':0.019680000000000142,('WP_208101977.1-Burkholderia-sp.-D-99':0.031210000000000182,'WP_205714023.1-Burkholderia-cepacia':0.026639999999999997)0.773:0.006909999999999972)0.848:0.03602000000000016)0.996:0.19727000000000006,('WP_213263532.1-Burkholderia-pyrrocinia':0.1225299999999998,'WP_060279477.1-Burkholderia-cepacia-complex':0.13325999999999993)0.983:0.1472699999999998)0.879:0.05827999999999989,('WP_189531325.1-Paludibacterium-paludis':0.49404000000000003,('WP_043620211.1-Chromobacterium-piscinae':0.15691999999999995,('WP_103900886.1-Chromobacterium-alticapitis':0.22665999999999986,'WP_045052286.1-Chromobacterium-violaceum':0.20685999999999982)0.886:0.08375000000000021)0.886:0.10485999999999995)0.856:0.07485000000000008)0.842:0.03849999999999998,('WP_020932001.1-Cycloclasticus-zancles':0.27212000000000014,('WP_084093192.1-Andreprevotia-lacus':0.07041999999999993,'WP_159673755.1-Andreprevotia-sp.-IGB-42':0.1701100000000002)0.982:0.13580999999999976)0.796:0.047190000000000065)0.822:0.052280000000000104,('WP_150719538.1-Pandoraea-capi':0.07956999999999992,'WP_046292890.1-Pandoraea-oxalativorans':0.08402999999999983)1.0:0.4065599999999998)0.798:0.03465000000000007)0.941:0.07982000000000022,(((((('WP_239235002.1-Candidatus-Nitrotoga-sp.-BS':0.049599999999999866,('WP_239287644.1-Candidatus-Nitrotoga-sp.-1052':0.014170000000000016,'WP_239190817.1-Candidatus-Nitrotoga-sp.-HW29':0.05046000000000017)0.508:0.014200000000000212)0.528:0.007330000000000059,'WP_239797721.1-Candidatus-Nitrotoga-arctica':0.017710000000000115)0.999:0.24175000000000013,'WP_204103763.1-Spirulina-major':0.40679)0.727:0.05565000000000042,((('WP_006529211.1-Gloeocapsa-sp.-PCC-73106':0.29063000000000017,((('WP_069965800.1-unclassified-Desertifilum':0.17876000000000003,('WP_073595522.1-Phormidium-ambiguum':0.07385999999999981,'WP_193982193.1-Phormidium-sp.-LEGE-05292':0.05738999999999983)0.983:0.09278999999999993)0.949:0.05812000000000017,('WP_190772446.1-unclassified-Leptolyngbya':0.24384000000000006,(((((('WP_194029353.1-Lusitaniella-coriacea':0.17871000000000015,('WP_009207329.1-Sulfuricella-denitrificans':0.23926999999999987,('WP_013221233.1-Nitrosococcus-watsonii':0.02690999999999999,'WP_002810067.1-Nitrosococcus-oceani':0.08610000000000007)0.999:0.2540800000000001)0.936:0.10400000000000009)0.361:0.022499999999999964,'WP_015170385.1-Geitlerinema-sp.-PCC-7407':0.2390500000000002)0.939:0.06422999999999979,('WP_008178350.1-Moorena-producens':0.21850000000000014,(('WP_190666400.1-Coleofasciculus-sp.-FACHB-501':0.10604000000000013,'WP_190525322.1-Coleofasciculus-sp.-FACHB-T130':0.055010000000000225)0.979:0.06535000000000002,((('WP_073549891.1-Chroogloeocystis-siderophila':0.1673,(('WP_233747769.1-Leptodesmis-sichuanensis':0.20943999999999985,('WP_224345144.1-Kovacikia-minuta':0.09907999999999983,'WP_192216758.1-Microcoleus-sp.-FACHB-SPT15':0.15660000000000007)0.724:0.026510000000000034)0.36:0.002969999999999917,'WP_223049852.1-Leptothermofonsia-sichuanensis':0.09270999999999985)0.687:0.03196000000000021)0.817:0.0242,(((('WP_099102643.1-Nostoc-sp.-Peltigera-malacea-cyanobiont-DB3992':0.07555999999999985,('WP_190959861.1-Nostoc':0.017830000000000013,('WP_190950926.1-Nostoc-punctiforme':0.042609999999999815,('WP_096537650.1-Nostoc-linckia':0.016140000000000043,'WP_221704991.1-Desmonostoc-muscorum':0.04330999999999996)0.735:0.006250000000000089)0.54:0.022079999999999878)0.979:0.07508999999999988)0.985:0.10374999999999979,(('WP_095721999.1-Calothrix-elsteri':0.08517000000000019,'WP_096624129.1-Calothrix-sp.-NIES-3974':0.07113000000000014)0.984:0.09651999999999994,('WP_199337786.1-Nostoc-sp.-FACHB-280':0.11893999999999982,((('WP_010998622.1-Nostocaceae':0.054440000000000044,'WP_011320078.1-Nostocaceae':0.055629999999999846)0.802:0.017100000000000115,((((('WP_190392428.1-Dolichospermum-sp.-FACHB-1091':0.012249999999999872,('WP_137908967.1-Nostocales':0.04410999999999987,'WP_028089231.1-Dolichospermum-circinale':0.017669999999999852)0.983:0.05459000000000014)0.986:0.07520000000000016,'WP_190370804.1-Anabaena-sp.-FACHB-1237':0.13942999999999994)0.915:0.033339999999999925,(('WP_096666215.1-Dolichospermum-compactum':0.05583999999999989,('WP_190387835.1-Aphanizomenon':0.05514999999999981,'WP_168466156.1-Aphanizomenon':0.017160000000000064)0.563:0.013030000000000097)0.674:0.013990000000000169,'WP_015080416.1-Anabaena-sp.-90':0.030240000000000045)0.928:0.02381000000000011)0.377:0.005599999999999827,'WP_168502193.1-Anabaena-sp.-UHCC-0187':0.005739999999999856)0.744:0.011270000000000113,('WP_168537172.1-Anabaena-sp.-UHCC-0253':0.03738000000000019,'WP_193938462.1-Cuspidothrix-issatschenkoi':0.13354)0.895:0.02868000000000004)0.998:0.11039000000000021)0.937:0.04436000000000018,(('WP_066425233.1-Anabaena-sp.-4-3':0.08655999999999997,'WP_015138800.1-Nostoc-sp.-PCC-7524':0.0728200000000001)0.99:0.06156000000000006,('WP_199337214.1-Nostoc-sp.-FACHB-152':0.10838999999999999,('WP_173403323.1-Nodularia-spumigena':0.04502000000000006,'WP_239729951.1-Nodularia-sphaerocarpa':0.060939999999999994)0.914:0.04049000000000014)0.884:0.021549999999999958)0.498:0.007420000000000204)0.841:0.01778000000000013)0.957:0.0679599999999998)0.83:0.027140000000000164)0.935:0.05575999999999981,(('WP_102179481.1-Fischerella-thermalis':0.021479999999999944,(('WP_131119525.1-Westiellopsis-prolifica':0.045479999999999965,'WP_016860528.1-Fischerella-muscicola':0.02389999999999981)0.848:0.012589999999999879,('WP_096675774.1-Fischerella-sp.-NIES-4106':0.046250000000000124,'WP_053458209.1-Hapalosiphon-sp.-MRB220':0.02967999999999993)0.687:0.006009999999999849)0.95:0.0690900000000001)1.0:0.18896999999999986,(('WP_015202791.1-Crinalium-epipsammum':0.20164999999999988,'WP_035154029.1-Calothrix-sp.-336/3':0.3395199999999998)0.87:0.05881999999999987,'WP_190825375.1-Iningainema-tapete':0.2170700000000001)0.522:0.026590000000000114)0.854:0.02599000000000018)0.882:0.03652999999999995,('WP_224410051.1-Oscillatoria-salina':0.19186000000000014,'WP_250123015.1-Chroococcidiopsis-sp.-CCMEE-29':0.13539000000000012)0.728:0.06555)0.565:0.048189999999999955)0.744:0.010339999999999794,('WP_193873404.1-unclassified-Coleofasciculus':0.12327000000000021,'WP_254010234.1-Symplocastrum-sp.-BBK-W-15':0.1200000000000001)0.811:0.038310000000000066)0.117:0.000540)0.465:0.009030000000000094)0.752:0.010130000000000194)0.919:0.03424000000000005,(('WP_190531554.1-Coleofasciculus-sp.-FACHB-542':0.17453999999999992,('WP_226578226.1-Microseira-wollei':0.1614399999999998,('WP_190468950.1-unclassified-Planktothrix':0.0941200000000002,'WP_249064905.1-Argonema-antarcticum':0.04850999999999983)0.854:0.03917999999999999)0.884:0.03227000000000002)0.748:0.014969999999999928,'WP_190493486.1-unclassified-Microcoleus':0.25081)0.805:0.01758000000000015)0.841:0.017220000000000013,((('WP_007355666.1-Kamptonema':0.09818000000000016,((('WP_242724542.1-Microcoleus-vaginatus':0.02978000000000014,('WP_096828456.1-Tychonema-bourrellyi':0.06000000000000005,'WP_236507253.1-Tychonema-sp.-BBK16':0.036230000000000206)0.916:0.03155000000000019)0.857:0.011940000000000062,'WP_015174083.1-Oscillatoria-nigro-viridis':0.017739999999999867)0.763:0.008160000000000167,('WP_193973976.1-Microcoleus-sp.-LEGE-07076':0.0452300000000001,('WP_194009709.1-unclassified-Tychonema':0.03377000000000008,'WP_265235823.1-Lyngbya-sp.-CCAP-1446/10':0.03805000000000014)0.888:0.017430000000000057)0.277:0.011410000000000142)0.991:0.09240000000000004)0.463:0.05710000000000015,(('WP_200665545.1-Oxynema-sp.-CENA135':0.2094100000000001,('WP_261197618.1-unclassified-Laspinema':0.03636000000000017,('WP_261204505.1-Laspinema-sp.-D2d':0.03636000000000017,'WP_207087219.1-Phormidium-pseudopriestleyi':0.032560000000000144)0.699:0.01731999999999978)0.988:0.08248999999999995)0.459:0.026899999999999924,((('WP_009783380.1-Lyngbya':0.07851000000000008,'WP_046277942.1-Limnoraphis-robusta':0.038879999999999804)0.971:0.08845999999999998,(('WP_072717475.1-Planktothrix':0.029790000000000205,'WP_083623900.1-Planktothrix-serta':0.030689999999999884)0.06:0.016140000000000043,('WP_079676747.1-Planktothrix-sp.-PCC-11201':0.02317999999999998,('WP_214559879.1-Planktothrix-agardhii':0.03641000000000005,'WP_026797127.1-Planktothrix':0.025030000000000108)0.222:0.025479999999999947)0.976:0.07089999999999996)0.966:0.06029000000000018)0.886:0.037310000000000176,'WP_190877489.1-unclassified-Planktothricoides':0.14921000000000006)0.827:0.026349999999999874)0.406:0.02984000000000009)0.768:0.012630000000000141,('WP_017715496.1-Oscillatoria-sp.-PCC-10802':0.1426599999999998,('WP_190665495.1-Microcoleus-sp.-FACHB-672':0.053830000000000044,'WP_261220767.1-Ancylothrix-sp.-D3o':0.17628999999999984)0.968:0.0636899999999998)0.922:0.035989999999999966)0.764:0.006810000000000205)0.903:0.028189999999999937,('WP_017661508.1-Baaleninema-simplex':0.22861999999999982,('WP_008052766.1-Limnospira-indica':0.19164000000000003,('WP_071516849.1-Geitlerinema-sp.-PCC-9228':0.2208000000000001,('WP_202221768.1-Okeania-sp.-KiyG1':0.027810000000000112,('WP_124146462.1-Okeania-hirsuta':0.016210000000000058,'WP_094672483.1-Hydrocoleum-sp.-CS-953':0.043769999999999865)0.262:0.014959999999999862)0.991:0.14284000000000008)0.458:0.06465999999999994)0.835:0.04430999999999985)0.937:0.06713000000000013)0.859:0.020239999999999814)0.143:0.007050000000000001)0.767:0.014310000000000045,((('WP_106909014.1-Microcystis-sp.-MC19':0.1895500000000001,('WP_166155173.1-unclassified-Neochlamydia':0.000540,'WP_044881819.1-Neochlamydia-sp.-EPS4':0.05566000000000004)1.0:0.41023999999999994)0.315:0.035359999999999836,('WP_015193936.1-Stanieria-cyanosphaera':0.017450000000000188,'WP_096384422.1-Stanieria-sp.-NIES-3757':0.08760000000000012)0.999:0.1974800000000001)0.887:0.05879000000000012,(((('WP_012595695.1-Rippkaea-orientalis':0.1659299999999999,('WP_008278255.1-Crocosphaera-chwakensis':0.0424199999999999,('WP_009545547.1-Crocosphaera-subtropica':0.019709999999999894,('WP_007308862.1-Crocosphaera-watsonii':0.0823100000000001,'WP_107668138.1-Cyanothece-sp.-BG0011':0.031950000000000145)0.829:0.013170000000000126)0.784:0.012469999999999981)0.984:0.11594000000000015)0.803:0.03988000000000014,('WP_264322079.1-Zarconia-navalis':0.1838099999999998,('WP_036478304.1-Myxosarcina-sp.-GI1':0.1671100000000001,('WP_006510035.1-Xenococcus-sp.-PCC-7305':0.1466400000000001,('WP_096721927.1-Chondrocystis-sp.-NIES-4102':0.08243,('WP_106232846.1-Pleurocapsa-sp.-CCALA-161':0.16005000000000003,'WP_229639079.1-Waterburya-agarophytonicola':0.01758000000000015)0.883:0.038390000000000146)0.99:0.13486999999999982)0.916:0.06547999999999998)0.522:0.03910999999999998)0.564:0.02226999999999979)0.807:0.019750000000000156,'WP_124977990.1-Aphanothece-sacrum':0.20630999999999977)0.412:0.025450000000000195,('WP_013324143.1-Gloeothece-verrucosa':0.11037000000000008,((('WP_096563566.1-Scytonema-sp.-NIES-4073':0.07832000000000017,('WP_089129802.1-Tolypothrix-sp.-NIES-4075':0.09004000000000012,('WP_096563464.1-Scytonema-sp.-NIES-4073':0.10996000000000006,'WP_048866388.1-Scytonema-tolypothrichoides':0.07782)0.66:0.02472999999999992)0.57:0.042469999999999786)0.997:0.1506599999999998,(('WP_220611028.1-Sphaerospermopsis-torques-reginae':0.048779999999999824,(('WP_009344618.1-Cylindrospermopsis-raciborskii':0.06985999999999981,'WP_240868223.1-Cylindrospermopsis-raciborskii':0.0194700000000001)0.998:0.12907000000000002,'WP_096570274.1-Sphaerospermopsis':0.04458999999999991)0.14:0.00898000000000021)0.987:0.09250999999999987,(('WP_016866333.1-Fischerella':0.07467999999999986,(('WP_086758595.1-Nostoc-sp.-106C':0.0571299999999999,(('WP_190545010.1-Calothrix':0.03047999999999984,'WP_190743070.1-Tolypothrix-sp.-FACHB-123':0.044540000000000024)0.978:0.06779000000000002,('WP_190656031.1-Nostoc-linckia':0.03230999999999984,'WP_045870014.1-unclassified-Tolypothrix':0.025819999999999954)0.982:0.07041999999999993)0.49:0.034199999999999786)0.784:0.0366200000000001,(('WP_214439445.1-Atlanticothrix-silvestris':0.1354000000000002,('WP_193199005.1-Nostoc-sp.-MG11':0.0488900000000001,'WP_162396505.1-Nostoc-sp.-B-2019':0.03800999999999988)0.947:0.05915999999999988)0.637:0.0487700000000002,('WP_017653186.1-Fortiea-contorta':0.16793000000000013,'WP_015128325.1-Calothrix-sp.-PCC-7507':0.12543000000000015)0.139:0.03523999999999994)0.823:0.028350000000000097)0.555:0.02760000000000007)0.889:0.030790000000000095,('WP_096557790.1-Nostoc-sp.-NIES-4103':0.13445000000000018,('WP_206263219.1-Nostoc-sp.-UHCC-0702':0.0570400000000002,('WP_190559626.1-Anabaena-sphaerica':0.09269999999999978,'WP_237989437.1-Nostoc-sp.-UHCC-0870':0.06512999999999991)0.873:0.04106999999999994)0.901:0.03733999999999993)0.78:0.04281999999999986)0.02:0.020000000000000018)0.188:0.02491000000000021)0.993:0.11524000000000001,'WP_015143663.1-Pleurocapsa-sp.-PCC-7327':0.10994999999999999)0.72:0.05967000000000011)0.831:0.027839999999999865)0.727:0.0189499999999998)0.816:0.03546000000000005)0.525:0.04817999999999989)0.368:0.043070000000000164,('WP_023076112.1-Leptolyngbya-sp.-Heron-Island-J':0.2166100000000002,('WP_039724568.1-Leptolyngbya-sp.-LK':0.06641999999999992,'WP_204137747.1-Halomicronema-sp.-CCY15110':0.0487700000000002)0.994:0.16303)0.965:0.11631999999999998)0.861:0.0394899999999998,('WP_237889684.1-Dissulfurimicrobium-hydrothermale':0.2794500000000002,'WP_237247350.1-Sideroxyarcus-emersonii':0.18821999999999983)0.963:0.12400999999999973)0.018:0.002380000000000493)0.883:0.0335700000000001,((('WP_137423090.1-Desulfoglaeba-alkanexedens':0.3186,'WP_066347484.1-Geminocystis-sp.-NIES-3708':0.40741000000000005)0.397:0.08461000000000007,((('WP_054689510.1-Desulfosarcina-cetonica':0.30242000000000013,('WP_250680171.1-Geomonas-sp.-Red32':0.1723300000000001,'WP_199381999.1-Geomesophilobacter-sediminis':0.20106000000000002)0.855:0.06380000000000008)0.5:0.019480000000000164,('WP_072282412.1-Syntrophotalea-acetylenivorans':0.2468300000000001,('WP_053550565.1-Desulfuromonas-soudanensis':0.27647999999999984,'WP_027714505.1-Desulfuromonas-sp.-TF':0.20179999999999998)0.72:0.03110999999999997)0.593:0.029049999999999798)0.843:0.04942000000000002,'WP_205218910.1-Desulforhopalus-vacuolatus':0.31861000000000006)0.641:0.017570000000000086)0.579:0.039979999999999905,(('WP_228856481.1-Desulfomarina-profundi':0.31599999999999984,'WP_183349429.1-Desulfoprunum-benzoelyticum':0.15196999999999994)0.905:0.08131999999999984,(('WP_028582690.1-Desulfogranum-japonicum':0.24059000000000008,('WP_205222281.1-Desulfogranum-marinum':0.1931400000000001,('WP_267928740.1-Desulfolithobacter-dissulfuricans':0.15275999999999978,'WP_205242362.1-Desulfobulbus-alkaliphilus':0.1299800000000002)0.565:0.05979000000000001)0.787:0.059579999999999966)0.93:0.06801000000000013,('WP_040200185.1-Geoalkalibacter-subterraneus':0.28749,'WP_015405619.1-Desulfocapsa-sulfexigens':0.17621999999999982)0.925:0.08150000000000013)0.181:0.015839999999999854)0.79:0.06321000000000021)0.908:0.058470000000000244)0.76:0.011489999999999778,((('WP_066727622.1-Desulfuromonas-sp.-DDH964':0.3518699999999999,('WP_047549861.1-Methylotenera-sp.-G11':0.19207000000000019,'WP_217345508.1-Noviherbaspirillum-sp.-L7-7A':0.21274000000000015)0.708:0.022390000000000132)0.843:0.0460799999999999,('WP_136680209.1-Neptunomonas-sp.-XY-337':0.20489999999999986,((('WP_028238613.1-Stutzerimonas-azotifigens':0.06870999999999983,('WP_183165579.1-Azomonas-macrocytogenes':0.15872999999999982,(('WP_025243649.1-Stutzerimonas-stutzeri':0.043079999999999785,('WP_255961233.1-Stutzerimonas-stutzeri':0.025679999999999925,('WP_102836461.1-Stutzerimonas-stutzeri':0.024950000000000028,('WP_045423249.1-Pseudomonadaceae':0.025069999999999926,'WP_256071125.1-Stutzerimonas-stutzeri':0.031210000000000182)0.881:0.012410000000000032)0.97:0.04760000000000009)0.683:0.003210000000000157)0.981:0.05235000000000012,('WP_122077451.1-Pseudomonadaceae':0.030050000000000132,(((('WP_043295157.1-Stutzerimonas-degradans':0.03583999999999987,'WP_237255234.1-Pseudomonas-oligotrophica':0.04343999999999992)0.93:0.03501999999999983,'WP_011911332.1-Stutzerimonas-stutzeri':0.025599999999999845)0.325:0.018749999999999822,('WP_120997453.1-[Pseudomonas':0.03399000000000019,('WP_230358311.1-Pseudomonas-sp.-NP21570':0.015019999999999811,('WP_193680295.1-Pseudomonas-lopnurensis':0.04249999999999998,'WP_200589398.1-Stutzerimonas-stutzeri':0.0201699999999998)0.562:0.03055999999999992)0.837:0.02644000000000002)0.319:0.017269999999999897)0.536:0.018990000000000062,('WP_037024448.1-Pseudomonas-saudiphocaensis':0.06162000000000001,'WP_181084609.1-Stutzerimonas-stutzeri':0.006400000000000183)0.77:0.02430000000000021)0.801:0.0136099999999999)0.751:0.000550)0.76:0.033560000000000034)0.648:0.03465000000000007)0.98:0.09112000000000009,('WP_228215938.1-Aromatoleum-toluclasticum':0.21932000000000018,(('WP_115680924.1-Cupriavidus-taiwanensis':0.23206000000000016,('WP_135702508.1-Cupriavidus-oxalaticus':0.1337600000000001,('WP_018313502.1-Cupriavidus-sp.-UYPR2.512':0.23448000000000002,('WP_071014919.1-Cupriavidus-sp.-USMAHM13':0.17722999999999978,'WP_150984056.1-Cupriavidus-basilensis':0.34391000000000016)0.939:0.11385000000000023)0.694:0.031130000000000102)0.244:0.009860000000000202)0.938:0.050899999999999945,(('WP_116600700.1-Cupriavidus-alkaliphilus':0.14467000000000008,('WP_183298446.1-Cupriavidus-alkaliphilus':0.049290000000000056,'WP_018008101.1-Cupriavidus-neocaledonicus':0.046660000000000146)0.997:0.13919999999999977)0.176:0.029739999999999878,(('WP_213445339.1-Variovorax-sp.-PCZ-1':0.27669999999999995,'WP_226921657.1-Kinneretia-sp.-XES5':0.21381000000000006)0.907:0.08951999999999982,('WP_224042884.1-Cupriavidus-respiraculi':0.2691699999999999,'WP_238511579.1-Cupriavidus-gilardii':0.25908)0.553:0.028140000000000054)0.516:0.03869999999999996)0.969:0.07794999999999996)0.643:0.01097999999999999)0.881:0.03546000000000005)0.937:0.060309999999999864,('WP_226440716.1-Quatrionicoccus-australiensis':0.37188,(('WP_143488038.1-Pseudomonas-mangiferae':0.12444999999999995,('WP_252273535.1-Pseudomonas-sp.-YIM-B01952':0.10821000000000014,((('WP_268799018.1-Pseudomonas-sp.-BSw22131':0.16427000000000014,(('WP_249678359.1-Pseudomonas-sp.-PIA16':0.14494000000000007,'WP_181290220.1-Pseudomonas-brassicacearum':0.16130000000000022)0.731:0.011159999999999837,'WP_043192328.1-Pseudomonas-rhizosphaerae':0.21105000000000018)0.823:0.0218799999999999)0.796:0.028890000000000082,(('WP_176569640.1-Pseudomonas-eucalypticola':0.1293700000000002,'WP_201004194.1-Pseudomonas-cichorii':0.059419999999999806)0.907:0.043099999999999916,((('WP_105697656.1-Pseudomonas-poae':0.040300000000000225,('WP_065875813.1-Pseudomonas':0.04126999999999992,('WP_033898696.1-Pseudomonas-fluorescens-group':0.03243000000000018,('WP_186687394.1-Pseudomonas-sp.-SWRI18':0.049430000000000085,'WP_169378974.1-Pseudomonas':0.014639999999999986)0.825:0.023820000000000174)0.415:0.009459999999999802)0.986:0.0676899999999998)0.735:0.01200000000000001,('WP_222540780.1-Pseudomonas-sp.-DR208':0.11018000000000017,('WP_253509974.1-Pseudomonas-rhodesiae':0.10686000000000018,'WP_093202821.1-Pseudomonas-yamanorum':0.0265200000000001)0.888:0.0354899999999998)0.222:0.007080000000000197)0.45:0.039200000000000124,'WP_219063768.1-Pseudomonas-sp.-UMAB-08':0.18252999999999986)0.9:0.042159999999999975)0.761:0.023149999999999782)0.971:0.09550999999999998,(((('WP_236238170.1-Pseudomonas-faucium':0.13551000000000002,'WP_054891693.1-Pseudomonas':0.07038000000000011)0.593:0.03508000000000022,('WP_158461286.1-Pseudomonas-fluorescens':0.21035000000000004,('WP_029612831.1-Pseudomonas-parafulva':0.018549999999999844,'WP_200141208.1-unclassified-Pseudomonas':0.18043999999999993)0.955:0.0676000000000001)0.346:0.029830000000000023)0.915:0.05249000000000015,(('WP_201176141.1-Pseudomonas-sp.-S31':0.02314000000000016,'WP_261240966.1-unclassified-Pseudomonas':0.044480000000000075)0.989:0.09270000000000023,((('WP_217840807.1-Pseudomonas-fakonensis':0.033259999999999845,'WP_217887155.1-Pseudomonas-xanthosomae':0.033949999999999925)0.835:0.025570000000000093,'WP_186654015.1-Pseudomonas-xantholysinigenes':0.19130000000000003)0.586:0.014650000000000052,'WP_110605176.1-Pseudomonas-soli':0.24692999999999987)0.126:0.027559999999999807)0.222:0.0189499999999998)0.812:0.030269999999999797,('WP_225786622.1-Pseudomonas':0.035089999999999844,'WP_209862330.1-Pseudomonas-sp.-BP8':0.4333800000000001)0.771:0.028690000000000104)0.979:0.0791599999999999)0.902:0.05184999999999995)0.875:0.02958000000000016)0.837:0.029999999999999805,((('WP_003453405.1-Pseudomonas':0.10734000000000021,((('WP_016495240.1-Pseudomonas-resinovorans':0.0499900000000002,'WP_044873077.1-Pseudomonas-sp.-LFM046':0.06590000000000007)0.851:0.025869999999999838,(('WP_077525673.1-Pseudomonas-resinovorans':0.10998000000000019,'WP_137821430.1-Pseudomonas-sp.-D-2018':0.07188000000000017)0.169:0.021399999999999864,'WP_058067682.1-Pseudomonas-sp.-TTU2014-080ASC':0.13518000000000008)0.936:0.05000999999999989)0.741:0.009479999999999933,('WP_028626762.1-Pseudomonas-resinovorans':0.035719999999999974,('WP_151138079.1-Pseudomonas-lalkuanensis':0.04074,'WP_120654068.1-Pseudomonas-sp.-DY-1':0.07405999999999979)0.838:0.01822999999999997)0.795:0.012789999999999857)0.268:0.01866000000000012)0.808:0.016669999999999963,('WP_226282979.1-Pseudomonas-lalkuanensis':0.06301999999999985,'WP_192399936.1-Pseudomonas-sp.-PDM15':0.13878000000000013)0.967:0.06528)0.717:0.016430000000000167,(('WP_220803742.1-Pseudomonas-sp.-NCCP-436':0.12069999999999981,'WP_090225506.1-Pseudomonas-kuykendallii':0.12071999999999994)0.853:0.043629999999999836,('WP_238040454.1-Pseudomonas-sp.-MMS21-TM103':0.10277000000000003,((('WP_159970704.1-Pseudomonas-sp.-8Z':0.0796199999999998,('WP_021444205.1-Pseudomonas-sp.-EGD-AK9':0.044890000000000096,((('WP_150606948.1-Pseudomonas-oleovorans':0.04292000000000007,('WP_119694090.1-Pseudomonas':0.04671000000000003,'WP_106738315.1-Pseudomonas-mendocina':0.04192000000000018)0.944:0.04469000000000012)0.926:0.03517999999999999,(('WP_024309956.1-Pseudomonas-sp.-P818':0.04892999999999992,'WP_090337209.1-Pseudomonas-chengduensis':0.01847999999999983)0.609:0.006800000000000139,'WP_143497094.1-Pseudomonas-sp.-ALS1131':0.031179999999999986)0.634:0.012820000000000054)0.116:0.001180000000000181,'WP_074939507.1-Pseudomonas-composti':0.023379999999999956)0.891:0.014479999999999826)0.389:0.016280000000000072)0.945:0.06152999999999986,('WP_131198269.1-Pseudomonas-dryadis':0.09886999999999979,('WP_131178052.1-Pseudomonas-daroniae':0.06069000000000013,('WP_084307530.1-Pseudomonas-flavescens':0.06692000000000009,('WP_249909557.1-Pseudomonas-fulva':0.04239000000000015,('WP_027910132.1-Pseudomonas-sp.-URMO17WK12:I4':0.04070000000000018,'WP_074889242.1-Pseudomonas-argentinensis':0.025049999999999795)0.39:0.013100000000000112)0.826:0.026720000000000077)0.653:0.02613000000000021)0.951:0.06323999999999996)0.97:0.07645999999999997)0.288:0.016140000000000043,(((('WP_255840290.1-Pseudomonas-benzenivorans':0.032810000000000006,'WP_208707982.1-Pseudomonas-sp.-LPB0260':0.026390000000000136)0.98:0.050839999999999996,('WP_147170838.1-Pseudomonas-sp.-SJZ079':0.0022199999999998887,'WP_025164100.1-Pseudomonas-taeanensis':0.055620000000000225)0.964:0.057539999999999925)0.798:0.014759999999999884,('WP_090448460.1-Pseudomonas-benzenivorans':0.15888999999999998,(('WP_233685494.1-Pseudomonas-anguilliseptica':0.050510000000000055,'WP_257778987.1-Pseudomonas-sp.-32.2.56':0.004859999999999864)0.997:0.10636000000000001,'WP_108096457.1-Pseudomonas-sp.-GV071':0.13287000000000004)0.837:0.023390000000000022)0.725:0.008599999999999941)0.928:0.026260000000000172,'WP_069518708.1-Pseudomonas':0.1121500000000002)0.684:0.008550000000000058)0.868:0.02760000000000007)0.853:0.027590000000000003)0.826:0.049209999999999976)0.632:0.0407099999999998)0.937:0.05642000000000014)0.127:0.01756000000000002)0.797:0.031019999999999825)0.787:0.029139999999999944)0.852:0.033479999999999954,((('WP_250479419.1-unclassified-Caballeronia':0.3405800000000001,('WP_104228621.1-Solimonas-fluminis':0.2727499999999998,'WP_083560927.1-Oceanococcus-atlanticus':0.31373000000000006)0.978:0.21105000000000018)0.227:0.014489999999999892,('WP_094797980.1-Bordetella-genomosp.-5':0.1652,(('WP_174426450.1-Cupriavidus-basilensis':0.3748499999999999,'WP_227181580.1-Leeia-sp.-TBRC-13508':0.27954999999999997)0.768:0.05912000000000006,'WP_250024752.1-Cupriavidus-campinensis':0.19309999999999983)0.259:0.04217000000000004)0.652:0.06477999999999984)0.576:0.04683999999999999,(((('WP_195874647.1-Herminiimonas-contaminans':0.1156299999999999,'WP_041297045.1-Janthinobacterium-sp.-Marseille':0.09215000000000018)0.764:0.021869999999999834,('WP_076590920.1-Herminiimonas-arsenitoxidans':0.1501199999999998,('WP_188420499.1-Oxalicibacterium-solurbis':0.15694000000000008,'WP_206604468.1-Herminiimonas-fonticola':0.04421000000000008)0.553:0.012420000000000098)0.873:0.03379999999999983)0.994:0.11177999999999999,((('WP_162020603.1-Undibacterium-crateris':0.13188000000000022,('WP_186861877.1-Undibacterium-griseum':0.07692999999999994,'WP_186884883.1-Undibacterium':0.03799999999999981)0.543:0.019870000000000054)0.986:0.10111999999999988,((('WP_179404315.1-Burkholderia-guangdongensis':0.14434000000000013,('WP_063494936.1-Paraburkholderia-phytofirmans':0.034809999999999786,('WP_042262232.1-Paraburkholderia-heleia':0.05990999999999991,'WP_254318368.1-Paraburkholderia-sp.-CNPSo-3272':0.000530)0.764:0.03399999999999981)0.996:0.1098300000000001)0.502:0.026619999999999866,(('WP_038711716.1-Burkholderia-sp.-lig30':0.21939999999999982,'WP_059459219.1-Burkholderia-vietnamiensis':0.2503799999999998)0.861:0.03905000000000003,((('WP_206951398.1-Trinickia-sp.-4D114':0.07818000000000014,'WP_205976236.1-Paraburkholderia-sp.-Se-20369':0.18893000000000004)0.97:0.08218000000000014,('WP_059889870.1-Burkholderia-stagnalis':0.19600000000000017,((('WP_061168975.1-Caballeronia-hypogeia':0.08993999999999991,(((('WP_134195781.1-Paraburkholderia-rhizosphaerae':0.27637,('WP_009889585.1-Burkholderia-thailandensis':0.0779399999999999,('WP_201138986.1-Paraburkholderia-domus':0.041850000000000165,'WP_266262001.1-Paraburkholderia-madseniana':0.020389999999999908)0.995:0.09920000000000018)0.627:0.025049999999999795)0.593:0.03428999999999993,'WP_069272519.1-Burkholderia-cepacia':0.22678999999999983)0.344:0.008929999999999882,(('WP_060092900.1-Burkholderia-vietnamiensis':0.022600000000000176,'WP_226113708.1-Burkholderia-sp.-AU33545':0.03785000000000016)1.0:0.15874999999999995,(('WP_105772332.1-Burkholderia-multivorans':0.02568999999999999,('WP_059842355.1-Burkholderia-multivorans':0.037739999999999885,'WP_175791145.1-Burkholderia-ambifaria':0.08729999999999993)0.793:0.010219999999999896)0.91:0.022969999999999935,('WP_168792792.1-Paraburkholderia-aromaticivorans':0.06316999999999995,('WP_059583574.1-pseudomallei-group':0.0375700000000001,'WP_004544228.1-Burkholderia':0.0396399999999999)0.88:0.021539999999999893)0.923:0.03285999999999989)0.655:0.009459999999999802)0.0:0.000530)0.899:0.013549999999999951,('WP_006496666.1-Burkholderia-cenocepacia':0.01780999999999988,'WP_059716874.1-Burkholderia-ubonensis':0.042400000000000215)0.916:0.024840000000000195)0.641:0.000540)0.877:0.022139999999999826,'WP_060156927.1-Burkholderia-ubonensis':0.08273999999999981)0.875:0.01252999999999993,((('WP_136892825.1-Trinickia-sp.-7GSK02':0.05277999999999983,('WP_054928771.1-Paraburkholderia-caribensis':0.04294999999999982,('WP_201650988.1-Paraburkholderia-sabiae':0.036989999999999856,'WP_205970438.1-Paraburkholderia-sp.-Tr-20389':0.03523999999999994)0.736:0.00633999999999979)0.819:0.013830000000000009)0.128:0.006899999999999906,'WP_121276956.1-Trinickia-fusca':0.15718999999999994)0.882:0.01778000000000013,'WP_209935850.1-Paraburkholderia-sp.-LEh10':0.08293000000000017)0.744:0.007249999999999979)0.717:0.01377000000000006)0.865:0.031870000000000065)0.263:0.03377000000000008,(('WP_266250364.1-Paraburkholderia-megapolitana':0.1549999999999998,'WP_026121090.1-Paraburkholderia-kururiensis':0.09539000000000009)0.789:0.05262000000000011,('WP_207000528.1-Trinickia-sp.-DHG64':0.1177100000000002,'WP_124149903.1-Paraburkholderia-dinghuensis':0.09631999999999996)0.802:0.046030000000000015)0.969:0.09362000000000004)0.821:0.03268000000000004)0.638:0.01548000000000016)0.833:0.04610000000000003,'WP_056136218.1-Massilia-sp.-Root133':0.1909200000000002)0.978:0.10306000000000015)0.923:0.04798999999999998,('WP_217900122.1-Noviherbaspirillum-humi':0.20940000000000003,((('WP_101482628.1-unclassified-Janthinobacterium':0.17559000000000013,('WP_099917421.1-Massilia-psychrophila':0.12428999999999979,('WP_092396716.1-Collimonas-sp.-OK412':0.1136499999999998,'WP_205319084.1-Glaciimonas-sp.-PAMC28666':0.0643600000000002)0.422:0.013069999999999915)0.866:0.029970000000000052)0.466:0.016039999999999832,('WP_065305794.1-Janthinobacterium-psychrotolerans':0.1394899999999999,(('WP_096238059.1-Janthinobacterium-svalbardensis':0.0177299999999998,('WP_102124214.1-Janthinobacterium-sp.-ROICE36':0.030429999999999957,('WP_215403374.1-Janthinobacterium-sp.-JC611':0.02295999999999987,('WP_094442213.1-Janthinobacterium-sp.-PC23-8':0.05667,('WP_230539136.1-Janthinobacterium-sp.-FW305-129':0.036639999999999784,'WP_139090581.1-Janthinobacterium-lividum':0.06186999999999987)0.862:0.012840000000000185)0.475:0.005809999999999871)0.805:0.006959999999999855)0.781:0.006130000000000191)0.793:0.006209999999999827,((('WP_035822645.1-Janthinobacterium-sp.-RA13':0.057319999999999816,'WP_034778721.1-Janthinobacterium-lividum':0.03664999999999985)0.736:0.00633999999999979,'WP_152282105.1-Janthinobacterium-violaceinigrum':0.018060000000000187)0.76:0.006180000000000074,'WP_183742413.1-unclassified-Janthinobacterium':0.042139999999999844)0.732:0.005999999999999783)0.849:0.000540)0.991:0.09511000000000003)0.804:0.01487000000000016,'WP_023495308.1-Methyloglobulus-morosus':0.15919000000000016)0.986:0.08049000000000017)0.671:0.008080000000000087)0.837:0.022069999999999812)0.077:0.01725000000000021,('WP_247737070.1-Thauera-aromatica':0.05295000000000005,'WP_039805139.1-Azotobacter-chroococcum':0.13847999999999994)0.98:0.10487000000000002)0.949:0.07111)0.851:0.03528999999999982)0.811:0.0158400000000003)0.898:0.018750000000000266)0.808:0.018049999999999677)0.865:0.04414000000000007,('WP_258826205.1-Massilia-agri':0.33157999999999976,((('WP_014706296.1-Methylophaga-nitratireducenticrescens':0.002689999999999859,'WP_136552334.1-Methylophaga-sp.-SB9B':0.06538999999999984)1.0:0.33607999999999993,('WP_070114691.1-Acidithiobacillus-caldus':0.28573000000000004,('WP_101536718.1-Acidithiobacillus-sp.-SH':0.1371199999999999,('WP_163095688.1-Acidithiobacillus-ferrianus':0.036680000000000046,('WP_009566950.1-Acidithiobacillus':0.04536000000000007,'WP_215870853.1-unclassified-Acidithiobacillus':0.06397999999999993)0.968:0.07521000000000022)0.691:0.03257000000000021)0.937:0.10524999999999984)0.991:0.20372999999999974)0.057:0.052050000000000374,'WP_216129682.1-Azospira-inquinata':0.32604000000000033)0.491:0.03170999999999946)0.768:0.040240000000000276)0.942:0.07792000000000021,(('WP_028988655.1-Thermithiobacillus-tepidarius':0.23918000000000017,('WP_127477393.1-Sulfurivermis-fontis':0.27626000000000017,(('WP_011630054.1-Alkalilimnicola-ehrlichii':0.14547,('WP_146294265.1-Euhalothece-natronophila':0.22646999999999995,'WP_159299499.1-Spiribacter-sp.-SSL99':0.19700000000000006)0.898:0.07432000000000016)0.908:0.05964999999999998,(('WP_019018778.1-Halomonas-lutea':0.2724099999999998,(((('WP_028484075.1-Thioalkalivibrio-sp.-ALE17':0.036519999999999886,('WP_019569965.1-Thioalkalivibrio-sp.-ALE11':0.06999999999999984,'WP_026296529.1-Thioalkalivibrio-sp.-ALJ2':0.1205099999999999)0.855:0.02373000000000003)0.735:0.007420000000000204,'WP_018169543.1-Thioalkalivibrio-sp.-ALMg9':0.028420000000000112)0.955:0.09213000000000005,('WP_250072676.1-Guyparkeria-hydrothermalis':0.1905800000000002,('WP_125198734.1-Guyparkeria-sp.-SCN-R1':0.1753499999999999,'WP_018882239.1-unclassified-Thioalkalivibrio':0.38214000000000015)0.816:0.046949999999999825)0.039:0.025599999999999845)0.997:0.20667999999999997,'WP_092999033.1-Thiohalomonas-denitrificans':0.05989999999999984)0.403:0.04295999999999989)0.953:0.12185999999999986,('WP_125841544.1-Stutzerimonas-stutzeri':0.33899999999999997,(('WP_106232428.1-Halomonas-ventosae':0.07464000000000004,('WP_262731117.1-Alcanivorax-marinus':0.13518000000000008,'WP_228234339.1-Alcanivorax-marinus':0.14731000000000005)0.875:0.043340000000000156)0.817:0.023439999999999905,'WP_175070852.1-Halomonas-taeanensis':0.1076999999999999)0.402:0.07582999999999984)0.865:0.07054999999999989)0.867:0.053910000000000124)0.893:0.056680000000000064)0.45:0.04479999999999995)0.774:0.07055000000000033,((('WP_178968040.1-Neptunomonas-phycophila':0.2688299999999999,('WP_062059887.1-Cellvibrio-sp.-OA-2007':0.04524999999999979,('WP_151033206.1-Cellvibrio-sp.-KY-GH-1':0.07167000000000012,'WP_049631514.1-Cellvibrio-sp.-pealriver':0.0715300000000001)0.796:0.07677999999999985)1.0:0.29532000000000025)0.876:0.12378,(('WP_015904383.1-Desulforapulum-autotrophicum':0.22956999999999983,'WP_111960646.1-Desulfobacter-hydrogenophilus':0.3076599999999998)0.686:0.0847500000000001,(('WP_254581577.1-Marinobacterium-sedimentorum':0.3117700000000001,('WP_051234278.1-Halomonas-halocynthiae':0.6274500000000001,'WP_110648656.1-Salinicola-peritrichatus':0.26774999999999993)0.939:0.1771600000000002)0.192:0.08538999999999985,('WP_179929642.1-Halomonas-salicampi':0.33511999999999986,(('WP_017842145.1-Methylotuvimicrobium-buryatense':0.2787799999999998,'WP_036548095.1-Nitrincola-lacisaponensis':0.34875999999999996)0.933:0.11079000000000017,(((('WP_228219791.1-Halomonas-meridiana':0.10671000000000008,('WP_120386672.1-Halomonas-alkaliphila':0.0915499999999998,(('WP_141441391.1-Halomonas-titanicae':0.054159999999999986,'WP_041157945.1-Halomonas-sp.-KHS3':0.07498000000000005)0.648:0.02544000000000013,'WP_226251132.1-unclassified-Halomonas':0.05859000000000014)0.892:0.04411999999999994)0.853:0.06932999999999989)0.973:0.09482999999999997,(('WP_251978361.1-unclassified-Salinicola':0.06904000000000021,'WP_189443129.1-Salinicola-rhizosphaerae':0.09170999999999996)0.994:0.1623899999999998,(('WP_158774127.1-Cobetia-sp.-L2A1':0.019039999999999946,('WP_088744253.1-Cobetia-sp.-QF-1':0.05505000000000004,('WP_144727127.1-Cobetia-crustatorum':0.02768000000000015,'WP_158774340.1-Cobetia-sp.-L2A1':0.041669999999999874)0.272:0.010040000000000049)0.901:0.026269999999999794)0.804:0.0116900000000002,('WP_077373671.1-Cobetia':0.04095000000000004,('WP_225346840.1-Cobetia-amphilecti':0.038759999999999906,('WP_200017180.1-Cobetia-sp.-cqz5-12':0.0584699999999998,'WP_213115235.1-Cobetia-sp.-MC34':0.027890000000000192)0.926:0.054520000000000124)0.87:0.031480000000000175)0.907:0.02823999999999982)0.958:0.09224999999999994)0.457:0.043769999999999865)0.805:0.039899999999999824,((('WP_127061154.1-Halomonas-nanhaiensis':0.15393999999999997,(('WP_078087027.1-unclassified-Halomonas':0.02145000000000019,('WP_162218282.1-Halomonas-alkaliphila':0.0318299999999998,('WP_030071911.1-Halomonas-alkaliantarctica':0.04410999999999987,(('WP_095603285.1-Halomonas-humidisoli':0.03308,'WP_038485954.1-Halomonas-campaniensis':0.023169999999999913)0.302:0.010940000000000172,'WP_253485683.1-Halomonas-sp.-XH26':0.02461000000000002)0.745:0.012669999999999959)0.984:0.06386000000000003)0.928:0.03600000000000003)0.782:0.01291999999999982,'WP_096655487.1-Halomonas-nigrificans':0.031789999999999985)0.996:0.11587000000000014)0.33:0.013180000000000192,('WP_096281135.1-Halomonas-sp.-JB37':0.2013799999999999,('WP_064236373.1-unclassified-Halomonas':0.05393999999999988,'WP_027958907.1-unclassified-Halomonas':0.029240000000000155)0.954:0.09452000000000016)0.772:0.04701000000000022)0.922:0.0312800000000002,(('WP_035566812.1-Halomonas':0.21587999999999985,('WP_081193716.1-Halomonas-sp.-BC1':0.2421000000000002,(('WP_102589198.1-Halomonas-urumqiensis':0.10591000000000017,((('WP_159342670.1-Halomonas-meridiana':0.2974000000000001,'WP_253554573.1-Halomonas-sp.-DN3':0.15565000000000007)0.811:0.04532999999999987,(('WP_193343673.1-Halomonas-elongata':0.05567000000000011,'WP_064701650.1-Halomonas-caseinilytica':0.013539999999999885)0.997:0.12253999999999987,('WP_110630630.1-Salinicola-salarius':0.06944000000000017,'WP_224850395.1-Marinobacter-nauticus':0.15045999999999982)0.533:0.02942)0.784:0.01537999999999995)0.0:0.000550,((('WP_089676106.1-Halomonas-aquamarina':0.02444999999999986,'WP_253557393.1-unclassified-Halomonas':0.04089000000000009)0.0:0.000540,('WP_062362595.1-Halomonas-axialensis':0.018400000000000194,'WP_089675781.1-Halomonas-aquamarina':0.045809999999999906)0.603:0.012360000000000149)0.776:0.006390000000000118,('WP_027961801.1-Halomonas-halodenitrificans':0.02364999999999995,('WP_195687443.1-Halomonas-sp.-KAO':0.02619999999999978,'WP_079551663.1-Halomonas-subglaciescola':0.05954000000000015)0.916:0.023229999999999862)0.87:0.01902999999999988)0.954:0.03230999999999984)0.612:0.009079999999999977)0.79:0.017370000000000108,'WP_193085952.1-Halomonas-sp.-FME65':0.09098000000000006)0.883:0.037349999999999994)0.334:0.03141999999999978)0.921:0.043950000000000156,(('WP_021818206.1-Halomonas-huangheensis':0.13343000000000016,('WP_073436013.1-Halomonas-cupida':0.07298999999999989,'WP_149282777.1-Halomonas-sp.-Y2R2':0.0682999999999998)0.123:0.04190000000000005)0.905:0.059109999999999996,('WP_248234415.1-Marinobacter-bryozoorum':0.08661999999999992,('WP_228223924.1-Marinobacter-nauticus':0.050839999999999996,'WP_098418752.1-Marinobacter-sp.-LV10MA510-1':0.11392999999999986)0.897:0.06620999999999988)0.929:0.07208000000000014)0.14:0.026489999999999903)0.855:0.023559999999999803)0.841:0.030870000000000175)0.744:0.021440000000000126,('WP_066801876.1-Moraxella-oblonga':0.11036000000000001,('WP_197128702.1-Psychrobacter-sp.-NZS113':0.16259000000000023,'WP_201616712.1-Psychrobacter-immobilis':0.15046999999999988)0.975:0.15115999999999996)0.995:0.23529)0.146:0.03485000000000005)0.914:0.07283000000000017)0.627:0.013619999999999965)0.951:0.18001000000000023)0.922:0.12058999999999997)0.747:0.10293000000000019,'WP_152145882.1-Alkalilimnicola-sp.-S0819':0.30945)0.516:0.06476000000000015)0.906:0.07057000000000002)0.634:0.02762999999999982,(((('WP_084281813.1-Polynucleobacter-kasalickyi':0.03285999999999989,'WP_108647113.1-Polynucleobacter-rarus':0.025339999999999918)1.0:0.6244800000000001,(('WP_215371371.1-Polynucleobacter-sp.-MWH-UH24A':0.16572999999999993,'WP_173942040.1-Polynucleobacter-antarcticus':0.17581999999999987)0.716:0.07621000000000011,((('WP_215299358.1-Polynucleobacter-sp.-AP-Nino-20-G2':0.07572000000000001,((('WP_215319389.1-Polynucleobacter-paneuropaeus':0.08763999999999994,('WP_215380622.1-Polynucleobacter-paludilacus':0.03576999999999986,'WP_215352354.1-Polynucleobacter-sp.-MWH-P3-07-1':0.03835999999999995)0.437:0.03175000000000017)0.976:0.06863000000000019,((('WP_173959551.1-Polynucleobacter-arcticus':0.026819999999999844,('WP_215376847.1-Polynucleobacter-asymbioticus':0.023229999999999862,'WP_215328794.1-Polynucleobacter-sp.-AP-Sving-400A-A2':0.048350000000000115)0.979:0.05006999999999984)0.9:0.017669999999999852,'WP_216225867.1-Polynucleobacter-sp.-MG-27-Goln-C1':0.08118999999999987)0.409:0.005710000000000104,('WP_216168716.1-Polynucleobacter-bastaniensis':0.07536999999999994,'WP_216178894.1-Polynucleobacter-sp.-80A-SIGWE':0.012169999999999792)0.881:0.020000000000000018)0.857:0.021319999999999784)0.665:0.013339999999999907,(('WP_062307935.1-Polynucleobacter-sinensis':0.054930000000000145,'WP_216197047.1-Polynucleobacter-sp.-UK-Mo-2m-Kol15':0.06626000000000021)0.733:0.006959999999999855,('WP_215334312.1-Polynucleobacter-sp.-AP-Ainpum-60-G11':0.03846000000000016,('WP_216268685.1-Polynucleobacter-sp.-Nonnen-W13':0.0413199999999998,'WP_251374783.1-Polynucleobacter-corsicus':0.03270000000000017)0.41:0.01413999999999982)0.756:0.007340000000000124)0.168:0.000530)0.823:0.025650000000000173)0.873:0.027089999999999836,('WP_076024025.1-Polynucleobacter-sphagniphilus':0.09037000000000006,('WP_216218975.1-Polynucleobacter-sp.-AP-Latsch-80-C2':0.036960000000000104,'WP_216254756.1-Polynucleobacter-sp.-AP-Melu-500A-A1':0.050920000000000076)0.993:0.08713999999999977)0.617:0.0048400000000001775)0.947:0.058230000000000004,('WP_114639791.1-Polynucleobacter-necessarius':0.1078800000000002,('WP_216177372.1-Polynucleobacter-sp.-AP-Feld-500C-C5':0.16342999999999996,('WP_128113711.1-Polynucleobacter-necessarius':0.09651999999999994,'WP_215385554.1-Polynucleobacter-sp.-AP-Kolm-20A-A1':0.06219999999999981)0.945:0.060229999999999784)0.834:0.030990000000000073)0.904:0.04991000000000012)0.248:0.05017000000000005)0.889:0.13588000000000022)0.965:0.19113999999999987,('WP_148063811.1-Parahaliea-aestuarii':0.08741999999999983,'WP_116367876.1-Parahaliea-mediterranea':0.04986999999999986)0.994:0.21845999999999988)0.043:0.11729000000000012,'WP_129645432.1-Steroidobacter-agariperforans':0.37944999999999984)0.959:0.13356000000000012)0.81:0.09597999999999995,(('WP_101249932.1-Telmatospirillum-siberiense':0.4307099999999999,('WP_189493202.1-Formosimonas-limnophila':0.2847700000000004,'WP_133620003.1-Hydromonas-duriensis':0.18168000000000006)0.95:0.12163999999999975)0.933:0.13615000000000022,(((('WP_255857717.1-Bacterioplanoides-sp.-SCSIO-12839':0.3227100000000003,('WP_116686321.1-Pelagibaculum-spongiae':0.35367000000000015,(('WP_067585802.1-Endozoicomonas-ascidiicola':0.13101999999999991,('WP_257293945.1-Endozoicomonas-sp.-YOMI1':0.1162700000000001,'WP_020580625.1-Endozoicomonas-elysicola':0.08427000000000007)0.933:0.09011000000000013)0.983:0.18990000000000018,('WP_257287521.1-Endozoicomonas-sp.-SESOKO2':0.11438000000000015,('WP_066015073.1-Endozoicomonas-atrinae':0.11825999999999981,'WP_262601261.1-Endozoicomonas-euniceicola':0.09602000000000022)0.606:0.04673999999999978)0.945:0.11142999999999992)0.845:0.1072700000000002)0.758:0.14127000000000045)0.846:0.07214999999999971,'WP_069857109.1-Desulfoplanes-formicivorans':0.2208199999999998)0.464:0.03758999999999979,'WP_156872703.1-Ketobacter-sp.-MCCC-1A13808':0.4350299999999998)0.905:0.0678700000000001,'WP_183409942.1-Litorivivens-lipolytica':0.4790599999999996)0.813:0.04393000000000047)0.268:0.023639999999999883)0.893:0.06408000000000014,((('WP_198243301.1-methane-oxidizing-endosymbiont-of-Gigantopelta-aegis':0.4105299999999996,('WP_201344172.1-Thiohalobacter-sp.-COW1':0.14589000000000008,'WP_132974196.1-Thiogranum-longum':0.15505000000000013)0.973:0.23907999999999996)0.305:0.10572000000000026,('WP_161826415.1-Petrachloros-mirabilis':0.47436999999999996,(('WP_234070476.1-Myxococcus-stipitatus':0.2796500000000002,'WP_120555541.1-Corallococcus-aberystwythensis':0.1965300000000001)0.997:0.35946999999999996,'WP_062153084.1-Beggiatoa-leptomitoformis':0.36746999999999996)0.073:0.09079000000000015)0.599:0.06357999999999997)0.86:0.14921000000000006,((((('WP_008811140.1-Parasutterella-excrementihominis':0.5424500000000001,(((('WP_110954047.1-Massilibacillus-massiliensis':0.25340000000000007,((('WP_054258662.1-Propionispora-sp.-2/2-37':0.1885300000000001,('WP_094603851.1-Sporomusa-silvacetica':0.10794000000000015,'WP_144352952.1-Sporomusa-termitida':0.13005999999999984)0.941:0.07794999999999996)0.827:0.028859999999999886,('WP_188400350.1-Sporomusa-sp.-GT1':0.2826599999999999,((((('WP_007952170.1-Pelosinus-fermentans':0.10578000000000021,'WP_007939097.1-Pelosinus':0.09961999999999982)0.909:0.06089000000000011,('WP_132083235.1-Anaerospora-hongkongensis':0.19510000000000005,'WP_132081660.1-Anaerospora-hongkongensis':0.07288000000000006)0.953:0.10767000000000015)0.928:0.05408999999999997,('WP_054258683.1-Propionispora-sp.-2/2-37':0.13493000000000022,('WP_091744506.1-Propionispora-vibrioides':0.12042999999999981,'WP_149735423.1-Propionispora-hippei':0.07948999999999984)0.474:0.03724000000000016)0.899:0.04903000000000013)0.835:0.018400000000000194,('WP_246588317.1-Pelorhabdus-rhamnosifermentans':0.18812000000000006,('WP_145090815.1-Sporomusa-sp.-KB1':0.0394899999999998,'WP_093793074.1-Sporomusa-acidovorans':0.07044000000000006)0.983:0.0977800000000002)0.911:0.053449999999999775)0.263:0.012129999999999974,'WP_092074275.1-Dendrosporobacter-quercicolus':0.25389000000000017)0.751:0.030489999999999906)0.688:0.014190000000000147)0.887:0.050129999999999786,'WP_188399481.1-Sporomusa-sp.-GT1':0.24569000000000019)0.9:0.07595000000000018)0.917:0.07663999999999982,((('WP_084578244.1-Sporomusa-malonica':0.41303,'WP_131919244.1-Heliophilum-fasciatum':0.2508300000000001)0.868:0.06584000000000012,('WP_092486361.1-Desulfoscipio-geothermicus':0.1326900000000002,('WP_240986382.1-Acididesulfobacillus-acetoxydans':0.25669999999999993,'WP_090940141.1-Pelosinus-propionicus':0.23356999999999983)0.897:0.07188999999999979)0.691:0.041059999999999874)0.847:0.06801000000000013,(((('WP_266183414.1-Dyella-sp.-RBB1W86':0.07301000000000002,'WP_111981026.1-Dyella-jiangningensis':0.14502000000000015)0.918:0.038040000000000074,((('WP_131151171.1-Dyella':0.18968000000000007,'WP_130619635.1-Dyella-amyloliquefaciens':0.056379999999999875)0.822:0.024869999999999948,'WP_199044928.1-Dyella-sp.-ASV24':0.07195000000000018)0.741:0.005990000000000162,('WP_266169930.1-Dyella-sp.-RXD159':0.07351999999999981,('WP_114846719.1-Dyella-tabacisoli':0.04350999999999994,'WP_090451016.1-Dyella-sp.-OK004':0.06524000000000019)0.809:0.024490000000000123)0.798:0.01167000000000007)0.825:0.011270000000000113)0.604:0.01142999999999983,((('WP_019467068.1-Dyella-japonica':0.23701000000000016,(('WP_126673574.1-Dyella-dinghuensis':0.054549999999999876,('WP_188793403.1-Dyella-nitratireducens':0.06655999999999995,'WP_233843762.1-Dyella-sp.-2HG41-7':0.034969999999999946)0.418:0.028020000000000156)0.93:0.05975000000000019,('WP_043099571.1-Oleiagrimonas-soli':0.05288000000000004,('WP_113064358.1-Oleiagrimonas-sp.-MCCC-1A03011':0.020059999999999967,'WP_168608661.1-Oleiagrimonas-citrea':0.05805000000000016)0.866:0.05791000000000013)0.996:0.13709000000000016)0.853:0.046170000000000044)0.746:0.01438000000000006,((((('WP_253565187.1-Dyella-sp.-Sa':0.0727199999999999,('WP_091339503.1-Frateuria-terrea':0.040979999999999794,'WP_209622196.1-Frateuria-flava':0.0693999999999999)0.981:0.060639999999999805)0.585:0.019629999999999814,('WP_063674027.1-Dyella-thiooxydans':0.12306000000000017,'WP_049621948.1-Frateuria-defendens':0.14068000000000014)0.492:0.015039999999999942)0.679:0.02856000000000014,(('WP_188800319.1-Dyella-caseinilytica':0.14265000000000017,'WP_192554051.1-Dyella-acidiphila':0.12058999999999997)0.963:0.07956999999999992,'WP_201315287.1-Dyella-sp.-EPa41':0.11463000000000001)0.06:0.007239999999999913)0.958:0.04908000000000001,('WP_179481164.1-unclassified-Rhodanobacter':0.12273999999999985,((('WP_072762453.1-Rhodanobacter-sp.-OK091':0.009069999999999911,(('WP_068095172.1-unclassified-Rhodanobacter':0.029349999999999987,'WP_214557828.1-unclassified-Rhodanobacter':0.030629999999999935)0.3:0.018440000000000012,('WP_008437080.1-Rhodanobacter-thiooxydans':0.03905000000000003,('WP_063107523.1-Rhodanobacter-thiooxydans':0.028340000000000032,'WP_211005527.1-Rhodanobacter-sp.-B2A1Ga4':0.035639999999999894)0.432:0.006260000000000154)0.734:0.019320000000000004)0.96:0.04809000000000019)0.732:0.013329999999999842,('WP_184416890.1-Rhodanobacter-sp.-MP7CTX1':0.04551999999999978,'WP_158752649.1-Dyella-sp.-S184':0.02845999999999993)0.886:0.021119999999999806)0.66:0.01969999999999983,(('WP_015446789.1-Rhodanobacter-denitrificans':0.061109999999999776,'WP_077445946.1-Rhodanobacter-sp.-C05':0.03338000000000019)0.27:0.018870000000000164,'WP_077439480.1-Rhodanobacter-sp.-C01':0.09994000000000014)0.0:0.000540)0.94:0.049819999999999975)0.558:0.024719999999999853)0.791:0.020849999999999813,'WP_204636097.1-Dyella-kyungheensis':0.14754999999999985)0.61:0.012350000000000083)0.889:0.03745999999999983,'WP_214570707.1-unclassified-Dyella':0.13135999999999992)0.611:0.02273000000000014)0.964:0.10311000000000003,('WP_078758981.1-Lysobacter-spongiicola':0.22589000000000015,('WP_132999175.1-Luteimonas-arsenica':0.22109999999999985,('WP_176021856.1-Stenotrophomonas-sp.-NA06056':0.04117000000000015,'WP_049440708.1-Stenotrophomonas-maltophilia':0.04436000000000018)0.982:0.1366299999999998)0.961:0.10748999999999986)0.378:0.06842000000000015)1.0:0.27310999999999996)0.28:0.015350000000000197)0.692:0.03263999999999978,(((((((('WP_201222000.1-Halochromatium-roseum':0.11256000000000022,'WP_200343816.1-Halochromatium-glycolicum':0.1499600000000001)0.977:0.1178300000000001,('WP_200237937.1-Thiohalocapsa-halophila':0.06570000000000009,'WP_058558062.1-Thiohalocapsa-sp.-ML1':0.22047000000000017)0.944:0.10355999999999987)0.968:0.14197000000000015,('WP_182584206.1-Thiospirillum-jenense':0.37192999999999987,('WP_200286492.1-Rhabdochromatium-marinum':0.12017999999999995,'WP_009150811.1-Thiorhodovibrio-sp.-970':0.06266999999999978)0.938:0.09336000000000011)0.469:0.09953000000000012)0.912:0.07970999999999995,'WP_018175268.1-unclassified-Thioalkalivibrio':0.4022700000000001)0.76:0.014819999999999833,('WP_089729519.1-Candidatus-Thiosymbion-oneisti':0.30771000000000015,((('WP_200388819.1-Thiocapsa-imhoffii':0.15364000000000022,('WP_232409912.1-Thiocapsa-bogorovii':0.019730000000000025,('WP_120796078.1-Thiocapsa-rosea':0.03270000000000017,'WP_007191850.1-Thiocapsa-marina':0.03878000000000004)0.693:0.005939999999999834)0.955:0.07529999999999992)0.95:0.07629999999999981,((('WP_200207675.1-Chromatium-weissei':0.18969000000000014,'WP_200154756.1-Chromatium-okenii':0.08827000000000007)0.99:0.1269,('WP_200158060.1-Allochromatium-vinosum':0.0524,'WP_200376120.1-Thiocystis-violacea':0.10246999999999984)0.844:0.039470000000000116)0.662:0.01089999999999991,(('WP_132978621.1-Thiobaca-trueperi':0.09112000000000009,'WP_201092787.1-Thiocystis-minor':0.06684000000000001)0.847:0.02112999999999987,(('WP_164652045.1-Thiorhodococcus-mannitoliphagus':0.041770000000000085,('WP_164456073.1-Thiorhodococcus-minor':0.08570000000000011,'WP_200330387.1-Thiocystis-violacea':0.08057999999999987)0.813:0.020459999999999923)0.96:0.056700000000000195,('WP_043749566.1-Imhoffiella-purpurea':0.06915000000000004,'WP_007039504.1-Thiorhodococcus-drewsii':0.0566399999999998)0.842:0.028410000000000046)0.887:0.021889999999999965)0.865:0.02682999999999991)0.871:0.028469999999999995)0.959:0.0789200000000001,('WP_100918325.1-Candidatus-Thiodictyon-syntrophicum':0.07813999999999988,'WP_020502667.1-Lamprocystis-purpurea':0.09879999999999978)0.957:0.08631000000000011)0.608:0.04784999999999995)0.872:0.05005000000000015)0.969:0.10280999999999985,'WP_101758446.1-Oceanicoccus-sp.-KOV_DT_Chl':0.49297000000000013)0.115:0.034889999999999866,'WP_141730858.1-Oligoflexus-tunisiensis':0.4312900000000002)0.498:0.03033999999999981,'WP_141198172.1-Persicimonas-caeni':0.48828000000000005)0.882:0.10355999999999987)0.885:0.07120999999999977,('WP_256181981.1-Cloacibacillus-evryensis':0.22642000000000007,'WP_087368268.1-Cloacibacillus-sp.-An23':0.0778500000000002)1.0:0.49105)0.714:0.10017000000000031)0.755:0.040939999999999976,('WP_255135177.1-Gracilimonas-sp.-CAU-1638':0.29715999999999987,'WP_020401568.1-Gracilimonas-tropica':0.2619500000000001)1.0:0.5736800000000004)0.801:0.04132999999999987,((('WP_094934028.1-Enterobacter-cloacae-complex':0.4448400000000001,'WP_057065012.1-Citrobacter':0.5900400000000001)0.88:0.12055000000000016,(('WP_240304790.1-Vibrio-campbellii':0.4006400000000001,'WP_001100815.1-Escherichia-coli':0.3355999999999999)0.944:0.1394000000000002,(((('WP_039473409.1-Vibrio-sinaloensis':0.27041000000000004,'WP_016795256.1-Vibrio-cyclitrophicus':0.2259199999999999)0.791:0.05118,('WP_086590801.1-Vibrio-parahaemolyticus':0.3136700000000001,'WP_010368546.1-Pseudoalteromonas-piscicida':0.3862399999999999)0.694:0.05384000000000011)0.875:0.06391999999999998,'WP_004741399.1-Vibrio':0.3419699999999999)0.378:0.04117000000000015,((('WP_195504469.1-Citrobacter-farmeri':0.2404799999999998,'WP_064554809.1-Buttiauxella-noackiae':0.2784399999999998)0.988:0.19444000000000017,(('WP_207379479.1-Shewanella-sedimentimangrovi':0.1730900000000002,'WP_269408820.1-Pseudoalteromonas-shioyasakiensis':0.19370999999999983)0.932:0.06298000000000004,('WP_171328835.1-Vibrio-splendidus':0.24415999999999993,('WP_061013169.1-Vibrionaceae':0.23546000000000022,'WP_155662549.1-Aliivibrio-fischeri':0.11663999999999985)0.803:0.07486000000000015)0.578:0.0032199999999997786)0.873:0.04545000000000021)0.876:0.046940000000000204,('WP_216735790.1-Hahella-sp.-HN01':0.3551000000000002,'WP_016537220.1-Cedecea-davisae':0.14905999999999997)0.746:0.021380000000000177)0.744:0.06031999999999993)0.756:0.026569999999999983)0.428:0.01587000000000005)0.982:0.11021999999999998,(('WP_100256321.1-Reinekea-forsetii':0.4116200000000001,(('WP_008983868.1-Alishewanella-agri':1.0070899999999998,(('WP_042281872.1-Candidatus-Protochlamydia-sp.-R18':0.10305000000000009,'WP_068471358.1-Parachlamydia-sp.-C2':0.16954000000000002)0.989:0.4990999999999999,('GJNO01008138.1-Laxus-oneistus-TSA':1.0036199999999997,('KAF8386991.1-Pristionchus-pacificus':0.46340999999999966,('CAD2157554.1-Meloidogyne-enterolobii':0.42584000000000044,(((((('TKR72632.1-Steinernema-carpocapsae':0.20731999999999973,('KAE9552274.1-Halicephalobus-sp.-NKZ332':0.2513300000000003,('KAH7695359.1-Aphelenchus-avenae':0.3082399999999996,(('CAD5228290.1-Bursaphelenchus-okinawaensis':0.042019999999999946,'CAD5235627.1-Bursaphelenchus-xylophilus':0.032569999999999766)0.086:0.04800000000000004,('KAI6234710.1-Aphelenchoides-fujianensis':0.09445000000000014,('KAI6206096.1-Aphelenchoides-besseyi':0.1392300000000004,'KAI6191010.1-Aphelenchoides-bicaudatus':0.08000999999999969)0.6:0.04734000000000016)0.965:0.10236)0.955:0.10569999999999968)0.875:0.1060800000000004)0.868:0.06624000000000008)0.754:0.03064,'GGJS01006545.1-Plectus-murrayi-TSA':0.38854000000000033)0.869:0.031890000000000196,('KAI1730870.1-Ditylenchus-destructor':0.001420000000000421,'KAI1709308.1-Ditylenchus-destructor':0.1598499999999996)1.0:0.2222299999999997)0.847:0.04203000000000001,('VDN56932.1-Dracunculus-medinensis':0.3216399999999999,(('VDD97390.1-Enterobius-vermicularis':0.4025699999999999,'KHN87800.1-Toxocara-canis':0.1356200000000003)0.846:0.06341999999999981,('HAHQ01030060.1-Ascaridia-galli-TSA':0.1266499999999997,('VDM95900.1-Thelazia-callipaeda':0.1812100000000001,('VDO27950.1-Onchocerca-flexuosa':0.11066999999999982,('VDO31788.1-Brugia-timori':0.10092999999999996,(('CAG9536712.1-Cercopithifilaria-johnstoni':0.026810000000000223,('VDK73328.1-Litomosoides-sigmodontis':0.046319999999999695,'VBB33646.1-Acanthocheilonema-viteae':0.08288000000000029)0.756:0.011510000000000353)0.856:0.020870000000000388,'XP_003135693.1-Loa-loa':0.06705999999999968)0.407:0.022520000000000095)0.803:0.029040000000000177)0.985:0.1484399999999999)0.998:0.3083600000000004)0.745:0.06484000000000023)0.673:0.05960999999999972)0.94:0.09534999999999982)0.933:0.08101000000000003,((('PAV65180.1-Diploscapter-pachys':0.22831999999999963,(('EPB80542.1-Ancylostoma-ceylanicum':0.03734000000000037,'KHJ99538.1-Oesophagostomum-dentatum':0.055600000000000094)0.949:0.0873400000000002,('VDO95824.1-Heligmosomoides-polygyrus':0.07312999999999992,('VDO65958.1-Haemonchus-placei':0.12917999999999985,'VDL70484.1-Nippostrongylus-brasiliensis':0.0791700000000004)0.874:0.023579999999999934)0.993:0.12464999999999993)0.964:0.12517999999999985)0.859:0.04748000000000019,'CAI4229259.1-Auanema-sp.-JU1783':0.41327000000000025)0.671:0.04252000000000011,('CAD6189320.1-Caenorhabditis-auriculariae':0.35134000000000043,('CAB3400093.1-Caenorhabditis-bovis':0.2096499999999999,('CAI5454992.1-Caenorhabditis-angaria':0.020470000000000432,('NP_509046.1-Caenorhabditis-elegans':0.028999999999999915,'XP_003117635.1-Caenorhabditis-remanei':0.027149999999999785)0.734:0.048499999999999766)0.839:0.03817000000000004)0.931:0.06388000000000016)0.977:0.12889000000000017)0.886:0.08259999999999978)0.863:0.06654999999999989,'XP_024501126.1-Strongyloides-ratti':0.2585600000000001)0.828:0.05204999999999993)0.283:0.04768999999999979)0.538:0.14754000000000023)1.0:0.6928399999999999)0.683:0.24689000000000005)0.534:0.14677000000000007,'WP_265845290.1-Plesiomonas-shigelloides':0.7423600000000001)0.819:0.07731000000000021)0.755:0.16279999999999983,((('WP_171588903.1-Pelistega-europaea':0.34911000000000003,((('WP_024553491.1-Franconibacter-helveticus':0.23667000000000016,'WP_140470594.1-Ewingella-americana':0.23050000000000015)0.889:0.0760200000000002,((('WP_028357569.1-Brackiella-oedipodis':0.2277,'WP_087653467.1-Klebsiella/Raoultella-group':0.14078999999999997)0.837:0.07475999999999994,('WP_253460089.1-Erwinia-persicina':0.26763000000000003,'WP_154751639.1-unclassified-Erwinia':0.2136)0.642:0.03204999999999991)0.495:0.03135999999999983,((('WP_165428828.1-Rouxiella-badensis':0.10355999999999987,(('WP_112167140.1-Klebsiella':0.1180699999999999,'WP_200519240.1-Klebsiella-michiganensis':0.10316999999999998)0.921:0.04579000000000022,'WP_137713144.1-Brenneria-rubrifaciens':0.11443000000000003)0.724:0.01324000000000014)0.365:0.01554000000000011,((('WP_130971788.1-Hafnia-alvei':0.11350000000000016,('WP_074823218.1-Pragia-fontium':0.09288000000000007,'WP_228398973.1-Insectihabitans-xujianqingii':0.08055000000000012)0.948:0.06271000000000004)0.955:0.04972999999999983,(('WP_148250181.1-Raoultella-terrigena':0.08589000000000002,'WP_032686784.1-Raoultella-planticola':0.02624000000000004)0.985:0.059699999999999864,(('WP_142430937.1-Klebsiella-oxytoca':0.1265200000000002,(('WP_004138711.1-Klebsiella':0.03708,('WP_032699180.1-Raoultella':0.04906999999999995,'WP_228399925.1-Raoultella':0.03141000000000016)0.209:0.000810)0.975:0.041850000000000165,('WP_239614184.1-Raoultella-ornithinolytica':0.08850999999999987,'WP_186921353.1-Klebsiella-quasipneumoniae':0.15983999999999998)0.737:0.013279999999999959)0.778:0.028189999999999937)0.789:0.018899999999999917,('WP_163980548.1-Raoultella-ornithinolytica':0.033339999999999925,'WP_131063651.1-Raoultella-planticola':0.023859999999999992)0.992:0.07028999999999996)0.019:0.009669999999999845)0.0:0.000540)0.839:0.010170000000000012,(('WP_193937755.1-Klebsiella-grimontii':0.17672,('WP_142514871.1-Klebsiella-huaxiensis':0.1458400000000002,('WP_234284225.1-Klebsiella-oxytoca':0.08158000000000021,'WP_024360036.1-Klebsiella':0.04850000000000021)0.299:0.032709999999999795)0.863:0.04010999999999987)0.752:0.013630000000000031,('WP_248464278.1-Pectobacterium-versatile':0.14148000000000005,(((('WP_095845203.1-Gibbsiella-quercinecans':0.060960000000000125,'WP_062791489.1-Serratia-plymuthica':0.06823999999999986)0.833:0.0188600000000001,(('WP_193158894.1-Serratia-marcescens':0.047719999999999985,'WP_148877341.1-Serratia-marcescens':0.06311)0.984:0.09214000000000011,(('WP_109053612.1-Brenneria-roseae':0.09222000000000019,('WP_136167279.1-Brenneria-sp.-CFCC-11842':0.03981000000000012,'WP_264089117.1-unclassified-Brenneria':0.09573000000000009)0.65:0.03639999999999999)0.718:0.015839999999999854,(('WP_130637964.1-Pectobacterium-versatile':0.04838999999999993,'WP_132452860.1-Samsonia-erythrinae':0.12126999999999999)0.17:0.014879999999999782,(((('WP_180777316.1-Pectobacterium-aroidearum':0.01908999999999983,'WP_180742451.1-Pectobacterium-sp.-CFBP8739':0.1718700000000002)0.838:0.0627399999999998,((('WP_109411993.1-Pectobacterium-polaris':0.05301,('WP_233968669.1-Pectobacterium-polaris':0.05179,'WP_048259384.1-Pectobacterium-peruviense':0.012939999999999952)0.666:0.01305999999999985)0.851:0.01352999999999982,'WP_039504461.1-Pectobacterium':0.0336599999999998)0.752:0.004770000000000163,'WP_138255615.1-Pectobacterium':0.021380000000000177)0.765:0.00788000000000011)0.794:0.006009999999999849,((('WP_119888241.1-Pectobacterium':0.0485199999999999,'WP_233976643.1-Pectobacterium-versatile':0.03882999999999992)0.548:0.005879999999999885,((('WP_226305805.1-Pectobacterium-versatile':0.019979999999999887,('WP_033071760.1-Pectobacterium-parmentieri':0.027270000000000127,('WP_198482318.1-Pectobacterium-parmentieri':0.03483999999999998,'WP_047710158.1-Pectobacterium-atrosepticum':0.05455999999999994)0.265:0.008550000000000058)0.94:0.03869999999999996)0.759:0.021910000000000096,('WP_039346077.1-Pectobacterium-fontis':0.02829000000000015,('WP_039517132.1-Pectobacterium-brasiliense':0.030520000000000103,'WP_039280054.1-Pectobacterium-brasiliense':0.032119999999999926)0.867:0.01872000000000007)0.733:0.005609999999999893)0.908:0.0194700000000001,'WP_103972204.1-Pectobacterium':0.02079999999999993)0.77:0.006709999999999994)0.83:0.012350000000000083,(('WP_151242707.1-Pectobacterium-carotovorum':0.03489999999999993,('WP_107169883.1-Pectobacterium':0.052360000000000184,'WP_205558955.1-Pectobacterium-brasiliense':0.04395999999999978)0.865:0.021780000000000133)0.23:0.000550,('WP_205946867.1-Pectobacterium':0.054749999999999854,'WP_116186106.1-Pectobacterium-aquaticum':0.029790000000000205)0.831:0.019909999999999872)0.787:0.006609999999999783)0.744:0.013440000000000119)0.805:0.011620000000000186,('WP_005974084.1-Pectobacterium-wasabiae':0.03824000000000005,('WP_249680869.1-Pectobacterium-polaris':0.03801999999999994,'WP_125232508.1-Pectobacterium-versatile':0.04278000000000004)0.43:0.007060000000000066)0.875:0.01686999999999994)0.942:0.03377000000000008)0.897:0.04591000000000012)0.958:0.07140000000000013)0.798:0.01839000000000013)0.699:0.015179999999999971,(('WP_168640580.1-Dickeya-sp.-CFBP-2040':0.08896000000000015,('WP_236306746.1-Dickeya-zeae':0.03993000000000002,(('WP_012883965.1-Dickeya-parazeae':0.07069999999999999,('WP_027712673.1-Dickeya-chrysanthemi':0.03893000000000013,('WP_022632763.1-Dickeya-solani':0.02513999999999994,('WP_123251184.1-Dickeya-undicola':0.04511999999999983,'WP_038918362.1-Dickeya':0.02793000000000001)0.465:0.006380000000000052)0.914:0.025700000000000056)0.794:0.011070000000000135)0.71:0.005879999999999885,'WP_012770533.1-Dickeya':0.033809999999999896)0.739:0.007150000000000212)0.0:0.000540)0.999:0.11325999999999992,((('WP_214648782.1-Pectobacterium-carotovorum':0.03308,(('WP_261846379.1-Pectobacterium-carotovorum':0.05519000000000007,'WP_121590317.1-Brenneria-goodwinii':0.005850000000000133)0.929:0.027270000000000127,('WP_182099415.1-Pectobacterium-aroidearum':0.03283000000000014,'WP_039482612.1-Pectobacterium':0.0264000000000002)0.738:0.006990000000000052)0.339:0.006510000000000016)0.733:0.005879999999999885,'WP_249228983.1-Brenneria-tiliae':0.042910000000000004)0.892:0.025339999999999918,'WP_121576612.1-Brenneria-alni':0.0874299999999999)0.861:0.01731999999999978)0.691:0.004669999999999952)0.785:0.03142999999999985,'WP_096000273.1-Klebsiella-pneumoniae':0.16448000000000018)0.584:0.015140000000000153)0.941:0.048880000000000035)0.866:0.017549999999999955)0.649:0.0037799999999998946)0.915:0.031239999999999934,(((('WP_214236207.1-Rosenbergiella-sp.-S61':0.04986999999999986,('WP_214195533.1-Rosenbergiella-collisarenosi':0.10107,'WP_241647140.1-Rosenbergiella-sp.-JB07':0.12071999999999994)0.381:0.03083999999999998)0.967:0.09645999999999999,(('WP_029991521.1-Tatumella-ptyseos':0.07568000000000019,('WP_212707699.1-Tatumella-sp.-JGM130':0.04371999999999998,'WP_087487466.1-Tatumella-citrea':0.05820999999999987)0.635:0.014009999999999856)0.788:0.03191999999999995,('WP_032441370.1-Klebsiella-pneumoniae':0.11256999999999984,('WP_265776798.1-Klebsiella-oxytoca':0.14577000000000018,'WP_015583911.1-Raoultella-ornithinolytica':0.05388999999999999)0.379:0.03697000000000017)0.778:0.056599999999999984)0.764:0.03760999999999992)0.624:0.06130000000000013,'WP_101506252.1-Erwinia-sp.-B116':0.15361000000000002)0.64:0.024430000000000174,('WP_040666911.1-Neisseria-wadsworthii':0.1043400000000001,'WP_135034416.1-Neisseria':0.14529000000000014)0.955:0.07189999999999985)0.709:0.010409999999999808)0.889:0.025700000000000056)0.475:0.005539999999999878)0.904:0.02679000000000009,('WP_049841125.1-Klebsiella-sp.-RIT-PI-d':0.3534600000000001,(('WP_146378871.1-Cronobacter-turicensis':0.06637000000000004,'WP_032180402.1-Escherichia-coli':0.08428000000000013)1.0:0.19457000000000013,('WP_256879438.1-Escherichia-albertii':0.07013999999999987,'WP_126816534.1-Enterobacter-mori':0.11771999999999982)0.842:0.06929000000000007)0.362:0.0454699999999999)0.727:0.015449999999999964)0.186:0.00890000000000013)0.885:0.036880000000000024,(('WP_246757812.1-Erwinia-phyllosphaerae':0.1956699999999998,'WP_232759612.1-unclassified-Pantoea':0.12224000000000013)0.944:0.08748999999999985,(('YP_009985672.1-Escherichia-phage-muut':0.21137000000000006,'WP_247177500.1-Escherichia-coli':0.41889999999999983)0.996:0.2750300000000001,(('WP_160432071.1-Gilliamella-sp.-Pas-s25':0.10931999999999986,(('WP_065587409.1-Gilliamella-apicola':0.053469999999999906,'WP_065580031.1-Gilliamella-apicola':0.06034000000000006)0.884:0.028589999999999893,('WP_065579324.1-Gilliamella-apis':0.08228999999999997,(('WP_065737496.1-Gilliamella-apicola':0.0341499999999999,'WP_267331332.1-unclassified-Gilliamella':0.05806000000000022)0.984:0.05580000000000007,('WP_039129858.1-Gilliamella':0.026639999999999997,('WP_198214961.1-Gilliamella-sp.-M0320':0.05310000000000015,'WP_086319877.1-Gilliamella-apis':0.01869999999999994)0.917:0.03080000000000016)0.411:0.000540)0.451:0.03358999999999979)0.719:0.029570000000000096)0.819:0.01974000000000009)0.525:0.017079999999999984,(('WP_065610260.1-Gilliamella-apicola':0.03177999999999992,('WP_065569248.1-Gilliamella-apicola':0.03813999999999984,'WP_065574597.1-Gilliamella-apicola':0.053630000000000067)0.627:0.01081999999999983)0.989:0.062129999999999797,('WP_267333169.1-unclassified-Gilliamella':0.13792999999999989,('WP_086361229.1-Gilliamella':0.12401000000000018,'WP_267361838.1-Gilliamella-sp.-B2772':0.1442000000000001)0.296:0.025619999999999976)0.868:0.027340000000000142)0.809:0.01772000000000018)0.962:0.09794999999999998)0.86:0.044340000000000046)0.723:0.028099999999999792)0.913:0.05740999999999996,(((((((('WP_248518672.1-Pseudoalteromonas-sp.-2CM32C':0.025949999999999918,'WP_149602089.1-Pseudoalteromonas-distincta':0.037450000000000205)0.966:0.10291000000000006,('WP_055022295.1-Pseudoalteromonas-sp.-P1-26':0.06343999999999994,('WP_260491935.1-Pseudoalteromonas-flavipulchra':0.21692,'WP_039494249.1-Pseudoalteromonas-flavipulchra':0.07464000000000004)0.14:0.02466999999999997)0.833:0.04651000000000005)0.977:0.15524000000000004,('WP_150896168.1-Vibrio-fortis':0.1961599999999999,'WP_067584343.1-Endozoicomonas-ascidiicola':0.2577400000000001)0.423:0.040830000000000144)0.711:0.06415000000000015,(('WP_016794198.1-Vibrio-cyclitrophicus':0.04002999999999979,'WP_102424249.1-Vibrio-cyclitrophicus':0.04388000000000014)0.841:0.03856999999999999,('WP_261903470.1-Vibrio-fortis':0.07967000000000013,'WP_240207223.1-Vibrio-sp.-CyArs1':0.03277000000000019)0.873:0.03398000000000012)0.972:0.11804000000000014)0.696:0.03949999999999987,(('WP_101108574.1-Psychromonas-sp.-psych-6C06':0.2360899999999999,(('WP_138320019.1-Thalassotalea-litorea':0.15009000000000006,'WP_136736185.1-Thalassotalea-mangrovi':0.1231399999999998)0.999:0.28424000000000005,'WP_114327629.1-Candidatus-Colwellia-aromaticivorans':0.11400999999999994)0.84:0.04776999999999987)0.843:0.04903000000000013,(('WP_127021653.1-Rheinheimera-mangrovi':0.2996699999999999,('WP_132584255.1-Rheinheimera-sp.-D18':0.3178399999999999,(((('WP_036130839.1-Marinobacter-nitratireducens':0.1090399999999998,((('WP_148609564.1-Marinobacter-sp.-BW6':0.10851999999999995,'WP_106695101.1-Marinobacter-shengliensis':0.056789999999999896)0.0:0.000540,('WP_222473125.1-Marinobacter-daepoensis':0.05279999999999996,'WP_092030923.1-Marinobacter-sp.-DSM-26671':0.11947999999999981)0.488:0.03161000000000014)0.962:0.0966499999999999,'WP_076722556.1-Marinobacter-lutaoensis':0.21607999999999983)0.27:0.035499999999999865)0.719:0.04178000000000015,('WP_036859756.1-Porticoccus-hydrocarbonoclasticus':0.23091000000000017,'WP_092385961.1-Halopseudomonas-salegens':0.17288000000000014)0.91:0.09357999999999977)0.909:0.0663800000000001,(('WP_204995064.1-Spongiibacter-marinus':0.08885999999999994,'WP_027875186.1-Spongiibacter-marinus':0.1500300000000001)0.675:0.03476999999999997,((('WP_103683243.1-Marortus-luteolus':0.029100000000000126,('WP_198570926.1-Spongiibacter-nanhainus':0.08368999999999982,'WP_184464382.1-Zhongshania-antarctica':0.04402000000000017)0.729:0.004910000000000192)0.99:0.10875000000000012,'WP_062382560.1-Zhongshania-aliphaticivorans':0.0466899999999999)0.772:0.04592000000000018,'WP_168451636.1-Spongiibacter-thalassae':0.04161999999999999)0.908:0.045179999999999776)0.486:0.03302000000000005)0.886:0.06835000000000013,'WP_131912015.1-Celerinatantimonas-diazotrophica':0.18019000000000007)0.877:0.056700000000000195)0.757:0.034800000000000164)0.97:0.11791000000000018,('WP_148513713.1-Vibrio':0.1818599999999999,'WP_199145406.1-Citrobacter-sp.-ASV29':0.29715999999999987)0.419:0.06613999999999987)0.764:0.039690000000000225)0.786:0.02414000000000005)0.975:0.0916800000000002,((('WP_067216963.1-Marinomonas-gallaica':0.05292999999999992,(('WP_048693477.1-Catenovulum-maritimum':0.19921999999999995,('WP_191602238.1-Marinomonas-algicola':0.0732400000000002,'WP_269124033.1-Marinomonas-sp.-15G1-11':0.036150000000000126)0.98:0.12473000000000001)0.917:0.08590000000000009,('WP_067204681.1-Marinomonas-aquimarina':0.023600000000000065,'WP_055461762.1-Marinomonas-fungiae':0.07611999999999997)0.551:0.02010999999999985)0.731:0.02389000000000019)0.894:0.04349999999999987,'WP_228191388.1-Marinomonas-communis':0.11667000000000005)0.89:0.0607000000000002,'WP_261844981.1-Amphritea-ceti':0.38004000000000016)0.886:0.06097000000000019)0.0:0.0011399999999999189,('WP_084334293.1-Pseudomonas-indica':0.11317000000000021,('WP_077172422.1-Pseudomonas-psychrotolerans':0.05087000000000019,'WP_145010177.1-Pseudomonas-psychrotolerans':0.00627000000000022)0.999:0.21826999999999996)0.937:0.0859700000000001)0.91:0.03713999999999995,(('WP_168836617.1-Vibrio-agarilyticus':0.10088000000000008,('WP_076537303.1-Shewanella-sp.-UCD-KL21':0.0240800000000001,'WP_102531614.1-Shewanella-sp.-10N.286.52.B9':0.04756999999999989)1.0:0.53254)0.9:0.10578999999999983,('WP_221077378.1-Agarivorans-albus':0.028960000000000097,'WP_137671336.1-Agarivorans-sp.-Toyoura001':0.0660099999999999)0.902:0.08555000000000001)0.99:0.14118999999999993)0.825:0.02679000000000009)0.944:0.11242000000000019)0.948:0.1402899999999998)0.932:0.14293000000000022)0.938:0.08787000000000011,(((('WP_239493201.1-Luteitalea-sp.-TBR-22':0.38165000000000004,'WP_058867107.1-Chloracidobacterium-thermophilum':0.3340299999999998)0.933:0.22390000000000043,('WP_207681674.1-Desulfonema-magnum':0.46720000000000006,'WP_099324654.1-Candidatus-Kuenenia-stuttgartiensis':0.46214999999999984)0.878:0.13763000000000014)0.845:0.06571999999999978,('WP_077329105.1-Virgibacillus-siamensis':0.40951000000000004,'WP_110265962.1-Sinimarinibacterium-flocculans':0.39339000000000013)0.955:0.1676899999999999)0.576:0.04314000000000018,'WP_230840800.1-Gloeobacter-morelensis':0.45013000000000014)0.798:0.03122999999999987)0.532:0.03540999999999972,(('WP_236098952.1-Dethiosulfovibrio':0.5873999999999997,((('WP_038266426.1-Peptoclostridium-litorale':0.37936999999999976,(('WP_186886939.1-Anaerofilum-hominis':0.19240000000000013,'WP_102049807.1-Pygmaiobacter-massiliensis':0.2120000000000002)0.811:0.05101999999999984,('WP_155202396.1-Ruthenibacterium-lactatiformans':0.20258999999999983,('WP_185752312.1-Butyrivibrio-sp.-AE2032':0.33804999999999996,'WP_058965344.1-Fournierella-massiliensis':0.18396999999999997)0.809:0.0812400000000002)0.951:0.11920999999999982)0.976:0.16852)0.868:0.06513999999999998,(((('WP_220321633.1-Clostridium-tagluense':0.3053699999999999,('WP_256135338.1-Bittarella-massiliensis':0.30310999999999977,'WP_266162911.1-Caproicibacterium-sp.-BJN0003':0.15341000000000005)0.899:0.11735000000000007)0.811:0.0890200000000001,(('WP_253721803.1-Treponema-sp.-OMZ-787':0.4739300000000002,('WP_058352784.1-Acetivibrio-ethanolgignens':0.30782,(('WP_249305773.1-Jutongia-hominis':0.17289999999999983,('WP_022463459.1-Jutongia-huaianensis':0.08611000000000013,('WP_216564272.1-Eubacterium-sp.-MSJ-13':0.02994999999999992,'WP_055944671.1-unclassified-Clostridium':0.0336599999999998)0.902:0.03617999999999988)0.524:0.047190000000000065)0.997:0.21560999999999986,('WP_033167869.1-Clostridium-sp.-KNHs205':0.15054999999999996,('WP_073273890.1-Anaerocolumna-jejuensis':0.15522999999999998,('WP_010268185.1-Paenibacillus-senegalensis':0.1673499999999999,('WP_155699987.1-Paenibacillus-psychroresistens':0.07159000000000004,('WP_202751253.1-Clostridium-sp.-YIM-B02515':0.05993000000000004,'WP_090040610.1-Clostridium-frigidicarnis':0.02519999999999989)0.367:0.011600000000000055)0.071:0.02776999999999985)0.935:0.06266000000000016)0.087:0.023169999999999913)0.999:0.16788999999999987)0.299:0.0195599999999998)0.084:0.027740000000000098)0.89:0.07873000000000019,(('WP_091684654.1-Anaerocolumna-aminovalerica':0.19857000000000014,(('WP_073279502.1-Anaerocolumna-jejuensis':0.10600999999999994,('WP_073588199.1-Anaerocolumna-xylanovorans':0.028500000000000192,'WP_185256524.1-Anaerocolumna-chitinilytica':0.1319300000000001)0.747:0.04585999999999979)0.683:0.048049999999999926,'WP_033168390.1-Clostridium-sp.-KNHs205':0.0880399999999999)0.995:0.18147000000000002)0.966:0.10727999999999982,('WP_226914794.1-Intestinibacter-bartlettii':0.31304,(('WP_005539496.1-Johnsonella-ignava':0.22454000000000018,('WP_060930762.1-Lachnoanaerobaculum-saburreum':0.19669999999999987,('WP_118555425.1-Lachnotalea-sp.-AF33-28':0.21891999999999978,(((('WP_006781439.1-Hungatella-hathewayi':0.08042000000000016,('WP_227052204.1-Anaerostipes-caccae':0.0318299999999998,'WP_137330070.1-Anaerostipes-rhamnosivorans':0.08904999999999985)0.995:0.11132000000000009)0.828:0.02195999999999998,('WP_075760385.1-Anaerostipes-sp.-494a':0.05396000000000001,'WP_215718428.1-Anaerostipes-hadrus':0.01244999999999985)0.992:0.07060999999999984)0.706:0.01092000000000004,((((('WP_186424215.1-Lacrimispora-celerecrescens':0.15250999999999992,((('WP_077612318.1-Clostridium-sp.-Marseille-P2415':0.040910000000000224,('WP_092242699.1-Lacrimispora':0.027089999999999836,'WP_024293365.1-Lachnospiraceae':0.035639999999999894)0.956:0.03554000000000013)0.876:0.021030000000000104,('WP_143321893.1-Clostridium-sp.-HBUAS56010':0.03232999999999997,(('WP_024837362.1-Lacrimispora':0.08949000000000007,'WP_124065139.1-Clostridium-sp.-E02':0.06841000000000008)0.846:0.02303999999999995,'WP_117415169.1-Clostridium-indicum':0.04612999999999978)0.734:0.009630000000000027)0.221:0.008779999999999788)0.232:0.010400000000000187,'WP_138313639.1-Hungatella-hathewayi':0.03232999999999997)0.853:0.018269999999999786)0.908:0.038759999999999906,((('WP_002577448.1-Enterocloster':0.06403999999999987,('WP_178313416.1-Lacrimispora-sp.-210928-DFI.3.58':0.01908999999999983,('WP_045093538.1-Clostridium-sp.-FS41':0.04530000000000012,'WP_025485467.1-Lachnospiraceae':0.06153999999999993)0.903:0.01988000000000012)0.735:0.007750000000000146)0.79:0.02925999999999984,'WP_066545911.1-Clostridium-sp.-AT4':0.029209999999999958)0.727:0.005399999999999849,'WP_024736308.1-Enterocloster-asparagiformis':0.04512999999999989)0.864:0.013959999999999972)0.835:0.013250000000000206,'WP_118541387.1-unclassified-Clostridium':0.07569999999999988)0.689:0.004570000000000185,('WP_215700061.1-Clostridium-sp.-MCC353':0.051979999999999915,('WP_249304640.1-Qiania-dongpingensis':0.055620000000000225,'WP_231062061.1-Lacrimispora-sp.-NSJ-141':0.10576000000000008)0.958:0.06674999999999986)0.783:0.02281000000000022)0.763:0.007080000000000197,((('WP_008399284.1-unclassified-Clostridium':0.03686999999999996,('WP_118742831.1-Clostridium-sp.-AM28-20LB':0.017920000000000158,('WP_003497689.1-[Clostridium':0.053469999999999906,'WP_125142063.1-Clostridium-transplantifaecale':0.040039999999999853)0.969:0.05000999999999989)0.175:0.019010000000000193)0.382:0.000550,('WP_101876396.1-Lachnoclostridium-edouardi':0.052999999999999936,'WP_087223935.1-Lachnoclostridium-sp.-An14':0.043070000000000164)0.931:0.028369999999999784)0.906:0.0189499999999998,'WP_118243934.1-Clostridium-sp.-AM58-1XD':0.04380000000000006)0.933:0.026330000000000187)0.806:0.013030000000000097)0.559:0.03092000000000006,('WP_154471253.1-Clostridium-porci':0.10849999999999982,'WP_251497048.1-Otoolea-sp.-DSM-111138':0.15307000000000004)0.691:0.028430000000000177)0.879:0.05058000000000007)0.437:0.04560999999999993)0.951:0.09992000000000001)0.357:0.049739999999999895,('WP_158346166.1-Lachnospiraceae':0.2521,('WP_158351687.1-Diplocloster-modestus':0.20820000000000016,(((('WP_087156947.1-Lachnoclostridium-sp.-An181':0.07233,('WP_115483253.1-Anaerosacchariphilus-polymeriproducens':0.12158999999999986,((('WP_109624131.1-Murimonas-intestini':0.08434000000000008,'WP_178047055.1-Fusicatenibacter-sp.-CLA-AA-H277':0.10780000000000012)0.562:0.01648000000000005,(((((('WP_251390531.1-Mediterraneibacter-sp.-DSM-111629':0.03738999999999981,'WP_073962323.1-[Ruminococcus':0.033009999999999984)0.854:0.026559999999999917,('WP_235247881.1-Faecalicatena-contorta':0.08644000000000007,'WP_158367551.1-Lachnospiraceae':0.029370000000000118)0.762:0.040160000000000196)0.754:0.011029999999999873,'WP_070089756.1-Merdimonas-faecis':0.03266999999999998)0.572:0.01576000000000022,'WP_230400100.1-Novisyntrophococcus-fermenticellae':0.2062599999999999)0.342:0.000530,'WP_132379639.1-Muricomes-intestini':0.12069000000000019)0.206:0.031540000000000123,('WP_173751851.1-Blautia-caecimuris':0.04651000000000005,'WP_117997894.1-Blautia-obeum':0.012389999999999901)0.99:0.06922000000000006)0.769:0.017980000000000107)0.902:0.02436999999999978,(('WP_148445023.1-Eubacteriales':0.027670000000000083,'WP_187566223.1-Blautia-faecis':0.05237999999999987)0.764:0.011730000000000018,('WP_158421413.1-Blautia':0.03461999999999987,('WP_173738571.1-Blautia-schinkii':0.021809999999999885,'WP_019160740.1-Blautia':0.06185000000000018)0.945:0.04079999999999995)0.314:0.025049999999999795)0.998:0.11059000000000019)0.867:0.029240000000000155)0.562:0.022170000000000023)0.912:0.04341999999999979,('WP_099466994.1-Konateibacter-massiliensis':0.07133000000000012,'WP_151145296.1-Candidatus-Galacturonibacter-soehngenii':0.060309999999999864)0.976:0.08335999999999988)0.316:0.029879999999999907,'WP_073994331.1-Parasporobacterium-paucivorans':0.15779999999999994)0.405:0.03168000000000015,((('WP_091822189.1-Butyrivibrio-sp.-ob235':0.07136999999999993,(('WP_026526812.1-Butyrivibrio-sp.-VCD2006':0.15702999999999978,'WP_026491547.1-unclassified-Butyrivibrio':0.012379999999999836)0.34:0.027750000000000163,'WP_044913869.1-Butyrivibrio-sp.-WCE2006':0.034440000000000026)0.927:0.05975999999999981)0.713:0.03744000000000014,'WP_035764434.1-Butyrivibrio-sp.-NC2002':0.08370999999999995)0.979:0.13051000000000013,((((('WP_022766545.1-Butyrivibrio-sp.-XPD2006':0.05168000000000017,'WP_022768176.1-unclassified-Butyrivibrio':0.0242)0.997:0.10160999999999998,'WP_034446055.1-Butyrivibrio-sp.-AE2032':0.06117000000000017)0.872:0.046860000000000124,'WP_024867291.1-Butyrivibrio-sp.-FCS014':0.03312999999999988)0.874:0.03039000000000014,('WP_071175335.1-Butyrivibrio':0.06559999999999988,'WP_026513668.1-Butyrivibrio-sp.-LB2008':0.05030000000000001)0.674:0.02947999999999995)0.813:0.024230000000000196,('WP_026663187.1-Butyrivibrio-proteoclasticus':0.07256000000000018,('WP_026652320.1-Butyrivibrio-proteoclasticus':0.041630000000000056,('WP_092060841.1-Butyrivibrio-sp.-INlla14':0.05030000000000001,'WP_013279951.1-Butyrivibrio-proteoclasticus':0.02790999999999988)0.838:0.017189999999999817)0.772:0.011029999999999873)0.219:0.016490000000000116)0.903:0.060229999999999784)0.992:0.13972000000000007)0.579:0.032560000000000144)0.801:0.036840000000000206)0.978:0.14017999999999997)0.74:0.02574999999999994)0.867:0.052810000000000024)0.031:0.02546999999999988)0.976:0.12141000000000002)0.669:0.009459999999999802,(((((('WP_073188830.1-Fibrobacter-sp.-UWP2':0.02425999999999995,'WP_073155408.1-Fibrobacter':0.04342999999999986)0.475:0.030569999999999986,'WP_073056819.1-unclassified-Fibrobacter':0.023950000000000138)0.485:0.014899999999999913,'WP_097034884.1-Fibrobacter-sp.-UWT3':0.022170000000000023)0.554:0.07181000000000015,('WP_072827923.1-Fibrobacter-sp.-UWB12':0.022479999999999833,('WP_073304542.1-Fibrobacter':0.2090399999999999,'WP_088630837.1-unclassified-Fibrobacter':0.034440000000000026)0.137:0.009469999999999867)0.97:0.11447999999999992)0.996:0.22875999999999985,'WP_022475313.1-Huintestinicola-butyrica':0.2618900000000002)0.268:0.03537999999999997,'WP_105204772.1-Neobittarella-massiliensis':0.16319000000000017)0.97:0.08731)0.937:0.06020000000000003,(((((((((('WP_147004474.1-Leptotrichia-wadei':0.02425999999999995,'WP_018499460.1-Leptotrichia-wadei':0.030289999999999928)0.908:0.07698999999999989,('WP_068155536.1-Leptotrichia-sp.-oral-taxon-847':0.06335999999999986,('WP_099992503.1-Fusobacterium-pseudoperiodonticum':0.12054000000000009,('WP_094080762.1-Leptotrichia-sp.-oral-taxon-498':0.042079999999999895,'WP_211617428.1-Leptotrichia-sp.-oral-taxon-218':0.013349999999999973)0.95:0.052589999999999915)0.235:0.03361999999999998)0.869:0.05859000000000014)0.994:0.2343799999999998,((('WP_117708548.1-Fusobacterium-mortiferum':0.000540,('WP_204716698.1-Fusobacterium-mortiferum':0.026040000000000063,'WP_204716813.1-Fusobacterium-mortiferum':0.36053999999999986)0.327:0.04125999999999985)0.851:0.02068000000000003,'WP_115270465.1-Fusobacterium-necrogenes':0.062120000000000175)0.964:0.08334999999999981,((('WP_130889691.1-Fusobacterium-ulcerans':0.03927000000000014,('WP_227372109.1-Fusobacterium-ulcerans':0.0329600000000001,('WP_005950923.1-Fusobacterium-varium':0.03135000000000021,'WP_070580525.1-Fusobacterium':0.05412999999999979)0.055:0.01114000000000015)0.895:0.015080000000000204)0.732:0.013259999999999827,('WP_241635364.1-Fusobacterium-gastrosuis':0.11188999999999982,('WP_211495349.1-Fusobacterium-necrophorum':0.19132000000000016,'WP_074518138.1-Fusobacterium-necrophorum':0.07453999999999983)0.943:0.07141999999999982)0.973:0.06850000000000023)0.84:0.026029999999999998,('WP_261793436.1-Fusobacterium-varium':0.15345999999999993,'WP_032890075.1-Fusobacterium-nucleatum':0.6374200000000001)0.97:0.11574000000000018)0.893:0.05035999999999996)0.969:0.12715999999999994)0.894:0.06850000000000023,('WP_013388736.1-Ilyobacter-polytropus':0.41228,'WP_173194267.1-Paenibacillus-frigoriresistens':0.30139000000000005)0.46:0.04261999999999988)0.621:0.01741999999999999,'WP_012548442.1-Dictyoglomus-thermophilum':0.37554)0.871:0.03864000000000001,((('WP_014043315.1-Caldicellulosiruptor-acetigenus':0.024350000000000094,'WP_127352844.1-Caldicellulosiruptor-changbaiensis':0.05628000000000011)1.0:0.33260999999999985,('WP_077198159.1-Thermosipho-affectus':0.08212999999999981,'WP_012057290.1-Thermosipho-melanesiensis':0.16781999999999986)0.591:0.04157000000000011)0.916:0.07651999999999992,('WP_047753768.1-Kosmotoga-pacifica':0.15010000000000012,(('WP_099829233.1-Mesotoga-sp.-Brook.08.105.5.1':0.1468600000000002,'WP_099830786.1-Mesotoga-sp.-Brook.08.105.5.1':0.17330999999999985)0.999:0.24136999999999986,'WP_036226274.1-Mesoaciditoga-lauensis':0.16942000000000013)0.386:0.021710000000000118)0.318:0.02851999999999988)0.968:0.08874000000000004)0.966:0.09866999999999981,('WP_213236459.1-Fusibacter-paucivorans':0.5346800000000003,'WP_148566871.1-Acetobacterium-paludosum':0.30952)0.965:0.12494000000000005)0.87:0.0607000000000002,'WP_138788932.1-Paenibacillus-thermoaerophilus':0.5480199999999997)0.769:0.030889999999999862,'WP_119970050.1-Clostridium-fermenticellae':0.25290999999999997)0.859:0.0319799999999999,((((('WP_011406138.1-Methanosphaera':0.1610800000000001,('WP_112123902.1-Methanosphaera-sp.-BMS':0.10204000000000013,('WP_095608065.1-Methanosphaera-cuniculi':0.10424999999999995,'WP_069592847.1-Methanosphaera-sp.-WGK6':0.02798999999999996)0.962:0.07779999999999987)0.324:0.039530000000000065)0.96:0.0771799999999998,(((('WP_048190917.1-Methanobacterium-sp.-SMA-27':0.12847000000000008,'WP_069583666.1-Methanobacterium':0.17072999999999983)0.926:0.059250000000000025,('WP_071907175.1-Methanobacterium-congolense':0.02220000000000022,'WP_209584978.1-Methanobacterium-aggregans':0.043849999999999945)0.996:0.11075999999999997)0.727:0.01471,'WP_248612240.1-Methanobacterium-alcaliphilum':0.13416000000000006)0.875:0.0224899999999999,('WP_116670296.1-Methanobrevibacter-woesei':0.23621999999999987,('WP_067148181.1-Methanobrevibacter-olleyae':0.12625999999999982,('WP_088538685.1-Methanobrevibacter-sp.-87.7':0.05478000000000005,'WP_016359256.1-Methanobrevibacter-sp.-AbM4':0.11805000000000021)0.97:0.09811000000000014)0.761:0.026619999999999866)0.998:0.1401300000000001)0.702:0.01562999999999981)0.751:0.017049999999999788,('WP_054834988.1-Methanobrevibacter-arboriphilus':0.1819700000000002,'WP_013644504.1-Methanobacterium-lacus':0.20391000000000004)0.902:0.03753999999999991)0.992:0.07024000000000008,((((('WP_226126924.1-Clostridium-algoriphilum':0.35643,(('WP_072990516.1-Clostridium-cavendishii':0.21948999999999996,((('WP_163195303.1-Clostridium-thermarum':0.13852999999999982,(('WP_073337580.1-Clostridium-grantii':0.1395200000000001,(('WP_072892386.1-Clostridium-fallax':0.11712999999999996,(('WP_152890015.1-Clostridium-tarantellae':0.13011000000000017,'WP_195972826.1-Clostridium-thermobutyricum':0.0823999999999998)0.494:0.032360000000000166,('WP_194191032.1-Clostridium-chrysemydis':0.1078800000000002,'WP_209795868.1-Clostridium-moniliforme':0.057720000000000216)0.933:0.04258000000000006)0.632:0.03681000000000001)0.75:0.007319999999999993,('WP_072832886.1-Clostridium-collagenovorans':0.03283000000000014,'WP_040194942.1-Clostridium-culturomicium':0.09431000000000012)0.589:0.00611000000000006)0.771:0.010029999999999983)0.927:0.030710000000000015,((('WP_095406848.1-Romboutsia-maritimum':0.07617999999999991,('WP_220321605.1-Clostridium-tagluense':0.04139999999999988,('WP_216288229.1-Clostridium-psychrophilum':0.11485000000000012,('WP_216137569.1-Clostridium-estertheticum':0.03064,'WP_220704225.1-Clostridium-estertheticum':0.03708)0.827:0.02035000000000009)0.892:0.04707999999999979)0.999:0.11091000000000006)0.868:0.024249999999999883,((('WP_047001949.1-Clostridium-sp.-C8':0.05740999999999996,('WP_195959433.1-Clostridium-tertium':0.06524999999999981,'WP_195429094.1-Clostridium-sp.-D46t1_190503_E9':0.018520000000000092)0.725:0.015899999999999803)0.769:0.02350999999999992,'WP_066890539.1-Clostridium-nigeriense':0.0424500000000001)0.526:0.006209999999999827,'WP_016207773.1-Clostridium-sartagoforme':0.029770000000000074)0.953:0.03166000000000002)0.908:0.020919999999999828,(('WP_066677671.1-Clostridium-septicum':0.0485199999999999,(('WP_029452249.1-Clostridium-algidicarnis':0.09584999999999999,'WP_216183125.1-Clostridium-gasigenes':0.021479999999999944)0.685:0.034600000000000186,('WP_216313757.1-Clostridium-gasigenes':0.03966000000000003,'WP_066874757.1-Clostridium-mediterraneense':0.19060999999999995)0.845:0.030629999999999935)0.967:0.061389999999999834)0.762:0.010110000000000063,('WP_238903698.1-Clostridium-sp.-YIM-B02506':0.06208999999999998,'WP_197096717.1-Paeniclostridium-sordellii':0.0829200000000001)0.642:0.00566000000000022)0.406:0.018440000000000012)0.851:0.015699999999999825)0.774:0.019120000000000026)0.408:0.00287000000000015,(('WP_055258173.1-Sarcina-ventriculi':0.07869999999999999,((('WP_161831114.1-Turicibacter-sp.-TS3':0.03986000000000001,('WP_212726030.1-Turicibacter-bilis':0.03941000000000017,('WP_195933776.1-Turicibacter-sanguinis':0.025490000000000013,('WP_235068992.1-Turicibacter-sp.-TJ11':0.05650000000000022,'WP_155223812.1-Turicibacter-sanguinis':0.031239999999999934)0.313:0.005830000000000002)0.0:0.000650)0.326:0.02350999999999992)0.982:0.08853,('WP_186834444.1-Eubacteriales':0.10869000000000018,((('WP_055265304.1-Clostridium-disporicum':0.054380000000000095,('WP_195245504.1-Clostridium-celatum':0.06363999999999992,'WP_154821366.1-Clostridium-saudiense':0.007490000000000219)0.786:0.02809000000000017)0.587:0.030409999999999826,'WP_186859336.1-Clostridium-sp.-NSJ-6':0.05425000000000013)0.813:0.050679999999999836,'WP_055265352.1-Clostridium':0.033279999999999976)0.859:0.06620999999999988)0.965:0.06320000000000014)0.785:0.024760000000000115,'WP_198610231.1-Clostridium-perfringens':0.05394999999999994)0.936:0.03141999999999978)0.931:0.033949999999999925,((('WP_066825124.1-Clostridium-tepidiprofundi':0.11194999999999977,'WP_072889493.1-Tepidibacter-formicigenes':0.06781000000000015)0.921:0.03954000000000013,('WP_055069574.1-Clostridium-massiliamazoniense':0.13603999999999994,((('WP_261829490.1-Clostridium-sp.-TW13':0.1947000000000001,('WP_090010800.1-Clostridium-sp.-DSM-8431':0.07225999999999999,'WP_216390068.1-Clostridium-sp.-MSJ-8':0.26363000000000003)0.378:0.030819999999999848)0.125:0.018510000000000026,('WP_106059304.1-Clostridium-vincentii':0.09619,'WP_143316579.1-Clostridium':0.021990000000000176)0.885:0.0429400000000002)0.821:0.030629999999999935,'WP_055184115.1-Clostridium-paraputrificum':0.04235999999999995)0.721:0.028389999999999915)0.439:0.01338000000000017)0.845:0.029389999999999805,((('WP_092726456.1-Romboutsia-lituseburensis':0.16541000000000006,'WP_258242885.1-Romboutsia-lituseburensis':0.029980000000000118)0.976:0.10063999999999984,('WP_057560648.1-Paeniclostridium-sordellii':0.08029999999999982,('WP_195252007.1-Romboutsia-sp.-1001713B170207_170306_H8':0.0532499999999998,('WP_057580750.1-Paeniclostridium-sordellii':0.10790999999999995,'WP_057538715.1-Paeniclostridium-sordellii':0.03765000000000018)0.792:0.014489999999999892)0.847:0.015019999999999811)0.364:0.013759999999999994)0.825:0.025399999999999867,('WP_094369216.1-Romboutsia-weinsteinii':0.1909200000000002,('WP_074918830.1-Terrisporobacter-glycolicus':0.06989000000000001,'WP_195929149.1-Clostridium-sp.-1001270J_160509_D11':0.030549999999999855)0.845:0.029040000000000177)0.865:0.02571000000000012)0.965:0.051810000000000134)0.832:0.013840000000000074)0.741:0.008030000000000204)0.824:0.01680999999999999,'WP_133628539.1-Fonticella-tunisiensis':0.05502999999999991)0.549:0.025100000000000122)0.193:0.030349999999999877,(('WP_072725426.1-Tepidibacter-thalassicus':0.14002999999999988,'WP_095404508.1-Romboutsia-maritimum':0.3217699999999999)0.869:0.05024000000000006,('WP_055666468.1-Desnuesiella-massiliensis':0.1415099999999998,('WP_069999280.1-Cellulosilyticum-sp.-I15G10I2':0.19101999999999997,('WP_054742706.1-Cellulosilyticum-ruminicola':0.08063999999999982,('WP_177671917.1-Zhenhengia-yiwuensis':0.04726000000000008,'WP_053982981.1-Niameybacter-massiliensis':0.10133999999999999)0.996:0.11398999999999981)0.951:0.07704999999999984)0.984:0.13158000000000003)0.666:0.007559999999999789)0.569:0.02143000000000006)0.895:0.052569999999999784)0.726:0.03393999999999986,((('WP_227846117.1-Clostridioides-sp.-ES-S-0145-01':0.029570000000000096,'WP_227847393.1-Clostridioides-sp.-ZZV15-6598':0.10851999999999995)0.969:0.04115000000000002,('WP_227842647.1-unclassified-Clostridioides':0.02992999999999979,'WP_003420982.1-Clostridioides-difficile':0.08648999999999996)0.863:0.020140000000000047)0.903:0.03666999999999998,'WP_038810653.1-Clostridioides-difficile':0.020999999999999908)1.0:0.2862)0.493:0.026499999999999968,((((('WP_041896941.1-Clostridium-beijerinckii':0.17927999999999988,('WP_077843022.1-Clostridium-beijerinckii':0.13710999999999984,'WP_009169246.1-unclassified-Clostridium':0.10069999999999979)0.829:0.02336999999999989)0.897:0.04667000000000021,'WP_069680896.1-Clostridium-taeniosporum':0.21299)0.916:0.04740000000000011,((((('WP_265859166.1-Clostridium-botulinum':0.10150000000000015,('WP_077856313.1-Clostridium-beijerinckii':0.000540,('WP_154531076.1-Inconstantimicrobium-porci':0.13526000000000016,'WP_009169009.1-Clostridium-sp.-DL-VIII':0.016910000000000203)0.726:0.046339999999999826)0.952:0.0644499999999999)0.952:0.04970000000000008,(('WP_077848125.1-Clostridium-puniceum':0.1386099999999999,((('WP_041894754.1-Clostridium-beijerinckii':0.02760000000000007,'WP_077855754.1-Clostridium-beijerinckii':0.03277999999999981)0.993:0.07606000000000002,'WP_261829154.1-Clostridium-sp.-TW13':0.07477)0.827:0.017819999999999947,(('WP_022750218.1-Clostridium-saccharobutylicum':0.047699999999999854,('WP_015391310.1-Clostridium-saccharoperbutylacetonicum':0.052890000000000104,('WP_238915641.1-Clostridium-sp.-YIM-B02555':0.058250000000000135,'WP_077846315.1-Clostridium-puniceum':0.04634999999999989)0.866:0.020140000000000047)0.758:0.024849999999999817)0.724:0.014419999999999877,'WP_224033465.1-Clostridium-gelidum':0.09791000000000016)0.241:0.006510000000000016)0.133:0.007490000000000219)0.783:0.007280000000000175,('WP_185652834.1-Clostridium-sp.-DJ247':0.020199999999999996,'WP_185652641.1-Clostridium-sp.-DJ247':0.06511000000000022)0.914:0.02699000000000007)0.0:0.000540)0.491:0.016189999999999927,'WP_226126241.1-Clostridium-algoriphilum':0.13640000000000008)0.749:0.008490000000000109,('WP_027636628.1-Clostridium-butyricum':0.08244000000000007,'WP_169105633.1-Clostridium-sp.-SM-530-WT-3G':0.06564000000000014)0.881:0.02984000000000009)0.883:0.03542999999999985,('WP_077866663.1-Clostridium-saccharobutylicum':0.18203000000000014,'WP_160359894.1-Clostridium-chromiireducens':0.24216999999999977)0.535:0.029650000000000176)0.77:0.025100000000000122)0.974:0.09026999999999985,'WP_079421587.1-Clostridium-oryzae':0.35652000000000017)0.766:0.04081000000000001,'WP_021359979.1-Clostridioides-difficile':0.3388300000000002)0.68:0.030349999999999877)0.605:0.028579999999999828,'WP_099190526.1-Tepidibacter-mesophilus':0.18452000000000002)0.788:0.03515999999999986,(('WP_072903743.1-Hathewaya-proteolytica':0.2982800000000001,('WP_142413383.1-Hathewaya-massiliensis':0.1890299999999998,'WP_035130905.1-Clostridium-sulfidigenes':0.21686000000000005)0.964:0.09990999999999994)0.879:0.07023000000000001,('WP_112093115.1-Methanothermobacter-tenebrarum':0.23408000000000007,'WP_013413362.1-Methanothermus-fervidus':0.22355999999999998)0.932:0.14369999999999994)0.863:0.042300000000000004)0.951:0.061329999999999885)0.787:0.01670000000000016,((('WP_068913445.1-Criibacterium-bergeronii':0.16109000000000018,'WP_009529486.1-Peptoanaerobacter-stomatis':0.17259000000000002)0.996:0.20797,('WP_111570784.1-Halanaerobium-saccharolyticum':0.0731799999999998,('WP_089717478.1-Halanaerobium-congolense':0.05088999999999988,'WP_089862761.1-Halanaerobium-salsuginis':0.02414000000000005)0.84:0.03324000000000016)0.998:0.2258)0.93:0.0892599999999999,(((('WP_216462460.1-Clostridium-bornimense':0.15302000000000016,('WP_234123145.1-Clostridium-hydrogenum':0.2164299999999999,('WP_026884435.1-Clostridium-akagii':0.09331999999999985,('WP_084114600.1-Clostridium-acidisoli':0.09253,('WP_015614219.1-Clostridium-pasteurianum':0.023109999999999964,('WP_003443690.1-Clostridium-pasteurianum':0.04878999999999989,'WP_010235680.1-Clostridium-arbusti':0.011519999999999975)0.889:0.024869999999999948)0.947:0.03756000000000004)0.504:0.009020000000000028)0.867:0.07695000000000007)0.847:0.048430000000000195)0.177:0.0019000000000000128,('WP_252130283.1-Clostridium-sp.-SYSU_GA19001':0.29115,'WP_202749105.1-Clostridium-sp.-YIM-B02515':0.12916000000000016)0.984:0.13329000000000013)0.866:0.0379299999999998,(('WP_036935152.1-Pseudobacteroides-cellulosolvens':0.16694999999999993,(((('WP_090798944.1-Paenibacillus-sp.-GP183':0.25032999999999994,('WP_056034132.1-Paenibacillus-sp.-Leaf72':0.07945000000000002,'WP_197936429.1-Paenibacillus-sp.-GSMTC-2017':0.1573199999999999)0.592:0.06511999999999984)0.365:0.03869999999999996,'WP_066398085.1-Cytobacillus-eiseniae':0.28263000000000016)0.701:0.044179999999999886,('WP_076307499.1-Paenibacillus-odorifer':0.052509999999999835,('WP_074087222.1-Paenibacillus-sp.-P32E':0.0383300000000002,('WP_042140168.1-Paenibacillus-sp.-FSL-P4-0081':0.009539999999999882,'WP_036721118.1-Paenibacillus-sp.-FSL-R7-277':0.06867000000000001)0.907:0.03569999999999984)0.958:0.09250999999999987)0.975:0.1037300000000001)0.837:0.043260000000000076,'WP_262684732.1-Paenibacillus-sp.-WQ-127069':0.18089999999999984)0.875:0.05528000000000022)0.969:0.11836000000000002,((('WP_209459870.1-Youngiibacter-multivorans':0.21069000000000004,'WP_090038599.1-Clostridium-frigidicarnis':0.13153000000000015)0.957:0.08625000000000016,('WP_164955553.1-Clostridium-estertheticum':0.18523999999999985,('WP_125002390.1-Clostridium-tagluense':0.18398000000000003,('WP_216146875.1-Clostridium-frigoris':0.05987000000000009,'WP_226136064.1-Clostridium-estertheticum':0.030819999999999848)0.976:0.09823000000000004)0.803:0.04261999999999988)0.853:0.061630000000000074)0.925:0.03509999999999991,(('WP_104410444.1-Clostridium-algidicarnis':0.041990000000000194,'WP_185162668.1-Clostridium-algidicarnis':0.06186999999999987)1.0:0.17731999999999992,(('WP_204598070.1-Clostridium-pascui':0.10660999999999987,('WP_216457288.1-Clostridium-simiarum':0.14073000000000002,'WP_027632043.1-Clostridium-hydrogeniformans':0.14059)0.676:0.034479999999999844)0.835:0.027029999999999887,('WP_165412483.1-Clostridium-estertheticum':0.05920000000000014,'WP_216302578.1-unclassified-Clostridium':0.0205700000000002)1.0:0.1794699999999998)0.728:0.03514000000000017)0.832:0.03291999999999984)0.819:0.022380000000000067)0.67:0.03011000000000008)0.876:0.0470600000000001,((('WP_250858284.1-Oceanirhabdus-seepicola':0.2199399999999998,('WP_069974893.1-Geosporobacter-ferrireducens':0.4003700000000001,('WP_223038819.1-Crassaminicella-profunda':0.08384999999999998,('WP_148809648.1-Crassaminicella-thermophila':0.061040000000000205,'WP_218283043.1-Crassaminicella-sp.-143-21':0.3008600000000001)0.839:0.028280000000000083)0.85:0.05892000000000008)0.922:0.0849899999999999)0.902:0.07466000000000017,'WP_160686059.1-Clostridium-sp.-C2-6-12':0.2530000000000001)0.912:0.07711999999999986,(('WP_090442301.1-Natronincola-peptidivorans':0.1826699999999999,('WP_095133872.1-Anaeromicrobium-sediminis':0.1821600000000001,('WP_068555806.1-Thermotalea-metallivorans':0.1254200000000001,'WP_073236503.1-Desulforamulus-putei':0.10768999999999984)0.558:0.023620000000000196)0.799:0.026499999999999968)0.258:0.010079999999999867,'WP_035164300.1-Caloranaerobacter-azorensis':0.07196999999999987)0.856:0.04646000000000017)0.776:0.01617999999999986)0.914:0.03654000000000002)0.693:0.007919999999999927)0.862:0.036290000000000155)0.851:0.03204999999999991)0.838:0.03952)0.955:0.13226999999999967,((('WP_130092062.1-Brumimicrobium-glaciale':0.46629999999999994,('WP_163286637.1-Cryomorpha-ignava':0.55715,(('WP_160842504.1-Pedobacter-sp.-HMF7647':0.06902000000000008,('WP_013633636.1-Pseudopedobacter-saltans':0.12182000000000004,'WP_154287284.1-Pedobacter-puniceum':0.18353999999999981)0.921:0.07588000000000017)0.965:0.17393999999999998,('WP_074492796.1-Mucilaginibacter-polytrichastri':0.1811799999999999,('WP_144912900.1-Mucilaginibacter-frigoritolerans':0.05720999999999998,('WP_258137696.1-Mucilaginibacter-sp.-BJC16-A38':0.24236000000000013,'WP_259068220.1-Mucilaginibacter-sp.-X4EP1':0.0406200000000001)0.152:0.04292000000000007)0.961:0.11494999999999989)0.668:0.0956999999999999)0.992:0.24791000000000007)0.147:0.08179999999999987)0.941:0.12165999999999988,((('WP_085659180.1-Prosthecochloris-sp.-HL-130-GSB':0.4442400000000002,'WP_038563105.1-Draconibacterium-orientale':0.2995700000000001)0.955:0.16616000000000009,(('WP_040596643.1-Prevotella-stercorea':0.08043999999999984,'WP_207153370.1-Prevotella-herbatica':0.09269000000000016)0.99:0.2340399999999998,('WP_223927321.1-Prevotella-lacticifex':0.22502999999999984,'WP_009010825.1-Prevotella-sp.-C561':0.15161999999999987)1.0:0.43057999999999996)0.346:0.07298999999999989)0.956:0.15817000000000014,((((('WP_101358295.1-Raineya-orbicola':0.2492000000000001,'WP_004155558.1-Microscilla-marina':0.34750000000000014)0.989:0.2804500000000001,'WP_116872776.1-Terrimonas-sp.':0.43360999999999983)0.911:0.10293000000000019,(('WP_205722928.1-Fulvivirga-lutea':0.45615000000000006,(((('WP_091091738.1-Flavobacterium-gillisiae':0.10338000000000003,(((('WP_269237486.1-Flavobacterium-sp.-IMCC34775':0.07799999999999985,'WP_163393939.1-Flavobacterium-limi':0.034479999999999844)0.934:0.000550,(('WP_115889823.1-Flavobacterium-cutihirudinis':0.045319999999999805,('WP_095952323.1-Flavobacterium-sp.-ACN6':0.04041000000000006,'WP_056200692.1-Flavobacterium-sp.-Root186':0.037290000000000045)0.385:0.011660000000000004)0.78:0.007150000000000212,'WP_198857739.1-Flavobacterium-sp.-IB48':0.03445000000000009)0.882:0.011760000000000215)0.864:0.011489999999999778,'WP_264205577.1-Flavobacterium-sp.-LS1R49':0.023540000000000116)0.854:0.012550000000000061,('WP_053470876.1-unclassified-Flavobacterium':0.16591999999999985,('WP_070907856.1-Flavobacterium-tructae':0.08655999999999997,('WP_173968237.1-Flavobacterium-collinsii':0.07475999999999994,'WP_264525312.1-Flavobacterium-sp.-N502536':0.042320000000000135)0.895:0.027590000000000003)0.59:0.008389999999999898)0.699:0.005930000000000213)0.704:0.007000000000000117)0.992:0.06191000000000013,'WP_248907802.1-Flavobacterium-sp.-K5-23':0.1190199999999999)0.905:0.035969999999999835,((('WP_116761981.1-Flavobacterium-laiguense':0.03864000000000001,'WP_143377960.1-Flavobacterium-sp.-ZT3R18':0.04919999999999991)0.987:0.06660000000000021,('WP_194311471.1-Flavobacterium-soyangense':0.12123000000000017,('WP_133277350.1-Flavobacterium-nackdongense':0.0838000000000001,'WP_144071503.1-Flavobacterium-franklandianum':0.06389000000000022)0.169:0.023480000000000167)0.539:0.030130000000000212)0.788:0.02751999999999999,'WP_103806521.1-Flavobacterium-alvei':0.08743999999999996)0.898:0.04537000000000013)0.99:0.16066000000000003,('WP_140000461.1-Flavobacterium-microcysteis':0.2190500000000002,('WP_140487657.1-Flavobacterium-sp.-GSA192':0.04288999999999987,('WP_100843428.1-Flavobacterium-sp.-5':0.10090999999999983,('WP_194640005.1-Flavobacterium-sp.-HJJ':0.09043000000000001,'WP_073246022.1-Flavobacterium-flevense':0.06701999999999986)0.807:0.021710000000000118)0.42:0.05481999999999987)0.996:0.1884800000000002)0.959:0.10978000000000021)0.933:0.11673)0.929:0.1141700000000001,'WP_057952567.1-Salinivirga-cyanobacteriivorans':0.59497)0.55:0.0396399999999999)0.932:0.07927000000000017,((((('WP_048137589.1-Methanosarcina-horonobensis':0.1483699999999999,('WP_011023680.1-Methanosarcina-acetivorans':0.03373999999999988,('WP_048173663.1-Methanosarcina-siciliae':0.05384000000000011,'WP_048184631.1-Methanosarcina-siciliae':0.05465000000000009)0.091:0.010959999999999859)0.946:0.06721999999999984)0.912:0.06646999999999981,('WP_048177163.1-Methanosarcina-sp.-MTP4':0.0820599999999998,('WP_048167026.1-Methanosarcina':0.0725699999999998,('WP_128503182.1-Methanosarcina-sp.-MSH10X1':0.1607099999999999,('WP_011033115.1-Methanosarcina-mazei':0.08409000000000022,'WP_048134921.1-Methanosarcina-sp.-1.H.T.1A.1':0.04131000000000018)0.132:0.03573000000000004)0.452:0.04746000000000006)0.679:0.04477999999999982)0.406:0.05494000000000021)0.808:0.08042999999999978,('WP_011500232.1-Methanococcoides-burtonii':0.2189000000000001,'WP_135611779.1-Methanococcoides-sp.-AM1':0.1876500000000001)0.743:0.057679999999999954)0.975:0.1615000000000002,'WP_068701711.1-Paludibacter-jiangxiensis':0.40431000000000017)0.267:0.04825999999999997,(('WP_127342456.1-Ancylomarina-longa':0.12504999999999988,(('WP_101263396.1-Labilibaculum-filiforme':0.04023999999999983,'WP_245870435.1-Labilibaculum-manganireducens':0.047559999999999825)0.972:0.08122000000000007,('WP_152734748.1-Marinifilum-sp.-N1E240':0.07771000000000017,('WP_171595603.1-Marinifilum-caeruleilacunae':0.06103000000000014,('WP_268189532.1-Marinifilum-sp.-D737':0.04838999999999993,'WP_054720444.1-Marinifilum-fragile':0.03067999999999982)0.891:0.02021000000000006)0.854:0.02403999999999984)0.919:0.050349999999999895)0.509:0.046009999999999884)0.997:0.1956699999999998,(('WP_133466511.1-Sunxiuqinia-elliptica':0.15538000000000007,('WP_159517929.1-Sunxiuqinia-indica':0.10575000000000001,'WP_053182762.1-Sunxiuqinia-dokdonensis':0.09918000000000005)0.836:0.042180000000000106)0.998:0.18409999999999993,(('WP_106543043.1-Prolixibacter-denitrificans':0.0989399999999998,('WP_153632016.1-Prolixibacter-sp.-SD074':0.05394999999999994,'WP_025865917.1-Prolixibacter-bellariivorans':0.019429999999999836)0.646:0.012849999999999806)0.995:0.20574000000000003,(('WP_233378072.1-Maribellus-sp.-CM-23':0.05419000000000018,'WP_247978154.1-Maribellus-sp.-YY47':0.07609999999999983)0.925:0.11240000000000006,('WP_167609329.1-Maribellus-sediminis':0.16443999999999992,('WP_045033348.1-Draconibacterium-sediminis':0.024239999999999817,'WP_163345719.1-Draconibacterium-halophilum':0.07804000000000011)0.99:0.15165000000000006)0.944:0.09771999999999981)0.999:0.2807599999999999)0.667:0.061900000000000066)0.961:0.09567000000000014)0.527:0.029119999999999813)0.961:0.10875000000000012)0.768:0.06706000000000012,((('WP_014783622.1-Aequorivita-sublithincola':0.3082600000000002,('WP_199156598.1-Pedobacter-sp.-ASV2':0.09033999999999986,((('WP_056872627.1-Pedobacter-sp.-Leaf194':0.09272999999999998,('WP_123203876.1-Pedobacter-jejuensis':0.07336000000000009,('WP_188416210.1-Pedobacter-mendelii':0.027120000000000033,'WP_056087060.1-Pedobacter-sp.-Leaf176':0.05003000000000002)0.466:0.014949999999999797)0.393:0.016770000000000174)0.974:0.05629000000000017,'WP_109929147.1-Pedobacter-paludis':0.03980000000000006)0.905:0.03503999999999996,('WP_133260761.1-Pedobacter-changchengzhani':0.12141000000000002,('WP_215166022.1-Pedobacter-sp.-ISL-64':0.08312000000000008,'WP_230143398.1-unclassified-Pedobacter':0.1137999999999999)0.914:0.07045000000000012)0.902:0.0583800000000001)0.689:0.02970000000000006)0.979:0.21826999999999996)0.978:0.20812000000000008,(('WP_084015419.1-Moheibacter-sediminis':0.34694,'WP_182042771.1-Moheibacter-lacus':0.2576100000000001)0.914:0.12245999999999979,((('WP_160564586.1-Apibacter-sp.-B2912':0.11277999999999988,('WP_055424523.1-Apibacter-mensalis':0.08679000000000014,('WP_160568957.1-unclassified-Apibacter':0.013440000000000119,'WP_166004639.1-Apibacter-sp.-B2966':0.04736999999999991)0.985:0.09014999999999995)0.547:0.0537399999999999)0.961:0.09894999999999987,('WP_105245553.1-Apibacter-adventoris':0.2063299999999999,'WP_146261858.1-Apibacter-muscae':0.1312500000000001)0.643:0.05555000000000021)1.0:0.25339,('WP_068596492.1-Vaginella-massiliensis':0.16671000000000014,(('WP_092908829.1-Algoriella-xinjiangensis':0.03208000000000011,'WP_125113420.1-Algoriella-xinjiangensis':0.08603000000000005)0.924:0.04985999999999979,('WP_125349156.1-Empedobacter-falsenii':0.10060000000000002,'WP_230477342.1-Faecalibacter-bovis':0.1055600000000001)0.804:0.0328400000000002)0.978:0.11660000000000004)0.986:0.1675399999999998)0.678:0.024719999999999853)0.968:0.14457999999999993)0.931:0.11296000000000017,('WP_111126400.1-Mesonia-sp.-K7':0.7160100000000003,(('WP_076395614.1-Chryseobacterium':0.04639000000000015,('WP_107809536.1-Elizabethkingia-sp.-YR214':0.14429999999999987,'WP_219971314.1-Chryseobacterium-sp.-LJ756':0.14022000000000023)0.894:0.053970000000000073)0.706:0.06013000000000002,'WP_088467981.1-Chryseobacterium-sp.-VAUSW3':0.11581000000000019)0.999:0.22470999999999997)0.803:0.000540)0.971:0.14669)0.935:0.12412000000000001)0.305:0.058949999999999836)0.934:0.12359000000000009,'WP_078711972.1-Anaerorhabdus-furcosa':0.5388000000000002)0.942:0.10559999999999992)0.934:0.1150500000000001)0.123:0.012360000000000149,((('WP_011996389.1-Coxiella-burnetii':0.6595200000000001,('WP_208013758.1-Meiothermus-sp.-CFH-77666':0.06747000000000014,('WP_027877894.1-Meiothermus-cerbereus':0.05501999999999985,'WP_119360783.1-Meiothermus-luteus':0.1811799999999999)0.217:0.05729000000000006)1.0:0.5762999999999998)0.86:0.11194999999999977,(((((('WP_011869065.1-Methanococcus-maripaludis':0.08508000000000004,'WP_013998786.1-Methanococcus-maripaludis':0.05372000000000021)0.873:0.04084000000000021,'WP_181493664.1-Methanococcus-maripaludis':0.0644499999999999)0.44:0.017220000000000013,'WP_181504225.1-Methanococcus-maripaludis':0.0525199999999999)0.999:0.25922,('WP_052562942.1-Candidatus-Brocadia-sinica':0.38951,'WP_173298646.1-Thermanaeromonas-sp.-C210':0.29357999999999995)0.818:0.08074000000000003)0.301:0.061960000000000015,('WP_048201153.1-Methanocaldococcus-bathoardescens':0.05221000000000009,'WP_018154793.1-Methanothermococcus-thermolithotrophicus':0.14802999999999988)0.826:0.03921000000000019)0.881:0.09722000000000008,(('WP_048103006.1-Aciduliprofundum-boonei':0.23875000000000002,(((((((('WP_087036111.1-Thermococcus-litoralis':0.0367099999999998,'WP_088855609.1-Thermococcus-siculi':0.048130000000000006)0.941:0.040589999999999904,('WP_013468354.1-Thermococcus-barophilus':0.052979999999999805,'WP_013906489.1-Pyrococcus-yayanosii':0.017999999999999794)0.892:0.017659999999999787)0.387:0.000540,('WP_068664737.1-Thermococcus-piezophilus':0.05932000000000004,((('WP_167888701.1-Thermococcus-sp.-MV5':0.07338999999999984,'WP_209475361.1-Thermococcus-stetteri':0.04789999999999983)0.838:0.027699999999999836,('WP_253304977.1-Thermococcus-aggregans':0.014520000000000088,('WP_015850310.1-Thermococcus-sibiricus':0.045459999999999834,'WP_152879125.1-Thermococcus-sp.-101-C5':0.052649999999999864)0.887:0.02088000000000001)0.847:0.02145000000000019)0.776:0.021419999999999995,(('WP_014733609.1-Pyrococcus-sp.-ST04':0.11777000000000015,('WP_223211916.1-Thermococcus-henrietii':0.05347999999999997,'WP_237702072.1-Thermococcus-sp.-AM4':0.03569999999999984)0.972:0.05171999999999999)0.215:0.011089999999999822,('WP_058938040.1-Thermococcus-celericrescens':0.027960000000000207,((('WP_088865724.1-Thermococcus-barossii':0.061799999999999855,('WP_232460978.1-Thermococcus-radiotolerans':0.049370000000000136,('WP_167711671.1-unclassified-Thermococcus':0.01977999999999991,'WP_088181368.1-Thermococcus-sp.-5-4':0.0685699999999998)0.745:0.006079999999999863)0.0:0.000550)0.755:0.012340000000000018,'WP_258085050.1-Thermococcus-sp.-813A4':0.03384000000000009)0.753:0.009999999999999787,('WP_139679991.1-Thermococcus-indicus':0.03108999999999984,'WP_148882943.1-Thermococcus-aciditolerans':0.03697000000000017)0.928:0.025370000000000115)0.549:0.00774000000000008)0.843:0.02112999999999987)0.929:0.038269999999999804)0.165:0.0011299999999998533)0.978:0.049350000000000005)0.671:0.01053000000000015,'WP_048053121.1-Pyrococcus-horikoshii':0.027039999999999953)0.817:0.012309999999999821,('WP_048150691.1-Palaeococcus-ferrophilus':0.024630000000000152,'WP_167913028.1-Thermococcus-sp.-21S7':0.07412000000000019)0.966:0.04549000000000003)0.79:0.010540000000000216,(('WP_013866394.1-Methanothermococcus-okinawensis':0.17627000000000015,'WP_013798345.1-Methanotorris-igneus':0.030980000000000008)0.866:0.027639999999999887,'WP_007044577.1-Methanotorris-formicicus':0.06262999999999996)0.997:0.08709999999999996)0.887:0.023330000000000073,('WP_148120035.1-Methanofervidicoccus-sp.-A16':0.16178999999999988,'WP_088862340.1-Thermococcus-celer':0.08705000000000007)0.313:0.000550)0.613:0.054079999999999906,'WP_010868297.1-Pyrococcus-abyssi':0.02906999999999993)0.915:0.046780000000000044)0.999:0.23484000000000016,((('WP_028950938.1-Sulfurihydrogenibium-subterraneum':0.14399000000000006,('WP_029522925.1-Persephonella-sp.-KM09-Lau-8':0.11704999999999988,('WP_200674641.1-Persephonella-atlantica':0.06368000000000018,'WP_096999769.1-Persephonella-hydrogeniphila':0.11258999999999997)0.904:0.05296000000000012)0.933:0.05384000000000011)0.475:0.017329999999999846,('WP_089322203.1-Desulfurobacterium-atlanticum':0.1524899999999998,('WP_013638952.1-Desulfurobacterium-thermolithotrophum':0.045700000000000074,'WP_132526207.1-Phorcysia-thermohydrogeniphila':0.08322999999999992)0.891:0.02737999999999996)0.897:0.06425999999999998)0.96:0.11655000000000015,('WP_150071182.1-Thermodesulfobacterium-sp.-TA1':0.26011000000000006,('WP_022856039.1-Thermodesulfobacterium':0.20509999999999984,'WP_245533116.1-Coprothermobacter-platensis':0.5583999999999998)0.116:0.08214999999999995)0.627:0.10002999999999984)0.908:0.08115000000000006)0.642:0.07886000000000015)0.771:0.04703999999999997)0.96:0.14484999999999992,('WP_110520589.1-Herpetosiphon-llansteffanensis':0.68011,('KAI6170547.1-Aphelenchoides-bicaudatus':0.6699099999999998,'KAI6244242.1-Aphelenchoides-fujianensis':0.4925100000000002)0.907:0.13919999999999977)0.948:0.19689000000000023)0.864:0.12365000000000004)0.958:0.12054999999999971)0.699:0.04186000000000023)0.983:0.12496000000000018)0.88:0.06813999999999965)0.982:0.18296000000000046,('WP_102949229.1-Aeromonas-caviae':0.3345799999999999,((('WP_236366479.1-Aeromonas-veronii':0.23045000000000027,('WP_161779122.1-Proteus-sp.-G2675':0.07103999999999999,'WP_162556830.1-Proteus':0.08227999999999991)0.991:0.15207999999999977)0.825:0.041979999999999684,('WP_225633057.1-Aeromonas-enteropelogenes':0.24228000000000005,('WP_097364011.1-Edwardsiella-tarda':0.3669199999999999,(('WP_043158915.1-Aeromonas-hydrophila':0.43423999999999996,(((('WP_234505032.1-Kluyvera-intermedia':0.3965200000000002,((('WP_195697765.1-Providencia-rettgeri':0.17635999999999985,'WP_159241675.1-Proteus-terrae':0.17859999999999987)0.801:0.07039000000000017,'WP_154625743.1-Providencia-stuartii':0.13892000000000015)0.848:0.030659999999999688,(('WP_193115284.1-Escherichia-coli':0.13929000000000036,'WP_187128479.1-Providencia-sp.-JUb39':0.14913000000000043)0.89:0.0538599999999998,((('WP_165880752.1-Providencia-rettgeri':0.18667999999999996,'WP_238563424.1-Providencia-rettgeri':0.12159999999999993)0.149:0.02104999999999979,'WP_231068620.1-Providencia-huaxiensis':0.15166999999999975)0.828:0.029290000000000482,(('WP_164565644.1-Providencia-rettgeri':0.18453000000000008,'WP_169342312.1-Providencia-stuartii':0.16491999999999996)0.78:0.05500000000000016,'WP_004924376.1-Providencia-stuartii':0.17131000000000007)0.706:0.01980000000000004)0.771:0.010009999999999852)0.834:0.021630000000000038)0.883:0.06425999999999998)0.896:0.06539000000000028,(('WP_157954980.1-Proteus-terrae':0.051209999999999756,('WP_161748440.1-Proteus-sp.-G2658':0.21384999999999987,'WP_049199273.1-Proteus-mirabilis':0.05201999999999973)0.907:0.052810000000000024)1.0:0.15907000000000027,((((((((('WP_129363899.1-Enterobacter-cloacae':0.028099999999999792,('WP_238085128.1-Pseudescherichia-vulneris':0.025269999999999904,'WP_032652659.1-Enterobacter-cloacae-complex':0.08808999999999978)0.429:0.007540000000000102)0.612:0.014279999999999848,('WP_257661844.1-Citrobacter-portucalensis':0.10477999999999987,'WP_193134185.1-Citrobacter-freundii':0.07333999999999996)0.854:0.026100000000000012)0.946:0.07611999999999997,('WP_042284276.1-Citrobacter-sedlakii':0.21327000000000007,('WP_024193259.1-Escherichia-coli':0.09582999999999986,'WP_099433281.1-Citrobacter-amalonaticus':0.03149999999999986)0.789:0.07414999999999994)0.937:0.0838599999999996)0.988:0.13873000000000024,(('WP_123427447.1-Lelliottia':0.05566000000000004,('WP_235147774.1-Enterobacter-cloacae':0.2628400000000002,'WP_103946175.1-Lelliottia-aquatilis':0.09014000000000033)0.766:0.0189499999999998)0.978:0.09146999999999972,('WP_062742814.1-Erwinia-persicina':0.17628999999999984,('WP_118663751.1-Erwinia-persicina':0.13580999999999976,('WP_167863639.1-Erwinia-rhapontici':0.1187999999999998,'WP_039385357.1-Pantoea-eucrina':0.1881400000000002)0.884:0.06495000000000006)0.734:0.04866999999999999)0.907:0.0676899999999998)0.605:0.02776000000000023)0.856:0.03994999999999971,((('WP_017800228.1-Erwinia-toletana':0.16400000000000015,'WP_227534785.1-Erwinia-rhapontici':0.12544999999999984)0.896:0.07289000000000012,('WP_216981205.1-Lelliottia-amnigena':0.26876999999999995,('WP_076770125.1-Kosakonia-cowanii':0.1288999999999998,'WP_085037002.1-Cronobacter-sakazakii':0.19575999999999993)0.896:0.14307999999999987)0.907:0.12450000000000028)0.537:0.03849999999999998,'WP_255851625.1-Erwinia-persicina':0.17886000000000024)0.984:0.1133999999999995)0.871:0.03678000000000026,((('WP_070133611.1-Candidatus-Erwinia-dacicola':0.1995199999999997,'WP_140919905.1-Insectihabitans-xujianqingii':0.1950599999999998)0.067:0.04090000000000016,(('WP_090460354.1-Enterobacter-sp.-kpr-6':0.07270999999999983,(('WP_080329661.1-Enterobacter-cloacae-complex':0.12908000000000008,'WP_200002836.1-Citrobacter-freundii':0.16122000000000014)0.985:0.17361000000000004,('WP_017458471.1-Kosakonia-sacchari':0.13504000000000005,('WP_020454293.1-Kosakonia-sp.-SOY2':0.09552000000000005,'WP_090123257.1-Kosakonia-arachidis':0.06208999999999998)0.902:0.04662999999999995)0.85:0.04348999999999981)0.896:0.09254000000000007)0.991:0.1592399999999996,(('WP_105274811.1-Escherichia':0.16342999999999996,'WP_064512195.1-Buttiauxella-gaviniae':0.3042499999999997)0.934:0.10016000000000025,'WP_029881813.1-Enterobacter':0.2730999999999999)0.706:0.016039999999999832)0.087:0.010420000000000318)0.909:0.06614000000000031,((((((((((('WP_244081328.1-Aeromonas-caviae':0.17064000000000012,'WP_223952962.1-Aeromonas-caviae':0.16588000000000003)0.896:0.07914999999999983,(((('WP_011919582.1-Shewanella-putrefaciens':0.03019000000000016,'WP_218558465.1-Shewanella-putrefaciens':0.06885999999999992)0.996:0.10898999999999992,('WP_226058924.1-Shewanella-sp.-SR1':0.1662300000000001,('WP_248067241.1-unclassified-Shewanella':0.09823000000000004,'WP_148505453.1-Vibrio-cholerae':0.08964999999999979)0.896:0.06106999999999996)0.218:0.0312800000000002)0.197:0.006520000000000081,((('WP_252420256.1-Aeromonas-taiwanensis':0.07999,('WP_077098334.1-Aeromonas-hydrophila':0.05358000000000018,('WP_042654929.1-Aeromonas-encheleia':0.05229999999999979,('WP_050718733.1-Aeromonas-tecta':0.1365599999999998,'WP_100859741.1-Aeromonas-lusitana':0.03575000000000017)0.746:0.008539999999999992)0.964:0.05296000000000012)0.787:0.015080000000000204)0.721:0.007039999999999935,'WP_043167431.1-Aeromonas-hydrophila':0.09507999999999983)0.707:0.05779999999999985,('WP_163316497.1-Crenobacter-sedimenti':0.13088999999999995,'WP_160797365.1-Craterilacuibacter-sinensis':0.13027999999999995)1.0:0.25021000000000004)0.507:0.055070000000000174)0.978:0.10322999999999993,(((('WP_145962313.1-Pectobacterium-peruviense':0.051959999999999784,('WP_198997094.1-Pectobacterium-carotovorum':0.021220000000000017,('WP_174866005.1-Pectobacterium-polaris':0.025399999999999867,'WP_165707548.1-Pectobacterium':0.030409999999999826)0.985:0.06029000000000018)0.678:0.014800000000000146)0.993:0.13805999999999985,('WP_039356706.1-Pectobacterium-actinidiae':0.07520999999999978,('WP_147247296.1-Brenneria-salicis':0.039470000000000116,'WP_172289906.1-unclassified-Brenneria':0.08970000000000011)0.866:0.031379999999999963)0.59:0.06489000000000011)0.795:0.025450000000000195,((('WP_043579071.1-Chromobacterium':0.04836000000000018,'WP_071113424.1-Chromobacterium-sphagni':0.10117999999999983)0.992:0.14188,(('WP_184706966.1-Rhizobium-lusitanum':0.046580000000000066,(((('WP_015341698.1-Rhizobium':0.03422999999999998,'WP_183712806.1-unclassified-Rhizobium':0.03069999999999995)0.871:0.012230000000000185,(('WP_095433322.1-Rhizobium-sp.-11515TR':0.031130000000000102,'WP_004123988.1-Rhizobium-freirei':0.10090000000000021)0.837:0.0253000000000001,'WP_106999541.1-Rhizobium-sp.-JAB6':0.043950000000000156)0.179:0.000550)0.94:0.019829999999999792,('WP_037195771.1-Rhizobium-lusitanum':0.03332999999999986,('WP_183765619.1-unclassified-Rhizobium':0.03064,'WP_120667780.1-unclassified-Rhizobium':0.0519400000000001)0.429:0.020910000000000206)0.791:0.010219999999999896)0.328:0.000540,('WP_092719658.1-Rhizobium-multihospitium':0.031849999999999934,(('WP_112341687.1-Rhizobium-tropici':0.0425399999999998,'WP_168880117.1-Rhizobium-sp.-P28RR-XV':0.039550000000000196)0.782:0.011249999999999982,('WP_145321829.1-unclassified-Rhizobium':0.10614999999999997,'WP_174074493.1-Agrobacterium-rhizogenes':0.08318000000000003)0.187:0.0026700000000001722)0.871:0.021220000000000017)0.502:0.007070000000000132)0.873:0.04213000000000022)0.268:0.05867000000000022,'WP_199146672.1-Rhizobium-sp.-ASV8':0.10053000000000001)0.995:0.17395000000000005)0.946:0.11057000000000006,(('WP_008952446.1-Pseudogulbenkiania-ferrooxidans':0.20953,'WP_014088374.1-Pseudogulbenkiania-sp.-NH8B':0.2145999999999999)0.83:0.04903000000000013,(('WP_174873379.1-Vogesella-oryzae':0.13544,'WP_147289921.1-Crenobacter-cavernae':0.1563699999999999)0.624:0.01941000000000015,'WP_137008594.1-Aquitalea-aquatilis':0.33732000000000006)0.809:0.062489999999999934)0.877:0.06164000000000014)0.931:0.0767199999999999)0.941:0.07119000000000009,(('WP_208149206.1-Shewanella-algae':0.28614000000000006,(((('WP_045419977.1-Vibrio-campbellii':0.058230000000000004,'WP_045500143.1-Vibrio-hyugaensis':0.18815999999999988)0.829:0.04618000000000011,'WP_025533544.1-Vibrio-parahaemolyticus':0.06251000000000007)0.948:0.04775000000000018,'WP_011788809.1-Shewanella-sp.-W3-18-1':0.05522999999999989)0.915:0.03860999999999981,('WP_219142360.1-Aeromonas-sanarellii':0.1510600000000002,(('WP_202046159.1-Aeromonas-media':0.06462000000000012,'WP_244056308.1-Aeromonas-caviae':0.15987000000000018)0.948:0.07894999999999985,('WP_225159881.1-Aeromonas-caviae':0.031039999999999957,('WP_042864139.1-Aeromonas-caviae':0.025720000000000187,'WP_045790994.1-Aeromonas-hydrophila':0.2168000000000001)0.738:0.017430000000000057)0.946:0.05995999999999979)0.95:0.06111999999999984)0.74:0.010489999999999888)0.679:0.011210000000000164)0.967:0.08838999999999997,('WP_223932214.1-Aeromonas-caviae':0.08788000000000018,'WP_161506954.1-Aeromonas-media':0.04104000000000019)0.964:0.08605000000000018)0.908:0.05806000000000022)0.833:0.03297999999999979)0.732:0.01462999999999992)0.91:0.04972000000000021,(('WP_076573677.1-Aeromonas-sp.-RU39B':0.22067000000000014,'WP_108652490.1-Dongshaea-marina':0.1903999999999999)0.912:0.07880999999999982,('WP_208005626.1-Oceanisphaera-pacifica':0.07936999999999994,('WP_087034853.1-Oceanisphaera-profunda':0.06501000000000001,'WP_087034873.1-Oceanisphaera-profunda':0.1051899999999999)0.938:0.058949999999999836)0.995:0.14417999999999997)0.705:0.010499999999999954)0.95:0.048130000000000006,('WP_253626250.1-Aeromonas-hydrophila':0.08626000000000023,('WP_073350970.1-Aeromonas-hydrophila':0.03874999999999984,('WP_190284026.1-Aeromonas-hydrophila':0.03238999999999992,'WP_241344946.1-Aeromonas-sp.-MR16':0.09262000000000015)0.967:0.07474999999999987)0.995:0.11147000000000018)0.639:0.052810000000000024)0.741:0.02369000000000021,((((('WP_042649704.1-Aeromonas-media':0.10606999999999989,(('WP_206874356.1-Aeromonas-veronii':0.00926999999999989,((('WP_060586794.1-Aeromonas-schubertii':0.047960000000000225,('WP_187590672.1-Aeromonas-hydrophila':0.10907,'WP_017782281.1-Aeromonas-dhakensis':0.02878999999999987)0.226:0.009700000000000042)0.995:0.08246999999999982,'WP_212539243.1-Aeromonas-salmonicida':0.030450000000000088)0.894:0.022850000000000037,'WP_201904077.1-Aeromonas-veronii':0.05187000000000008)0.78:0.017040000000000166)0.763:0.008950000000000014,(('WP_256684552.1-Aeromonas-salmonicida':0.02933999999999992,'WP_042063659.1-Aeromonas-allosaccharophila':0.031200000000000117)0.535:0.014440000000000008,('WP_171268810.1-Aeromonas-media':0.021069999999999922,('WP_148612677.1-Aeromonas-sobria':0.03855999999999993,('WP_201951905.1-Aeromonas-veronii':0.02961999999999998,'WP_201355655.1-Aeromonas-hydrophila':0.039280000000000204)0.29:0.00541999999999998)0.526:0.010889999999999844)1.0:0.13104000000000005)0.501:0.007460000000000022)0.94:0.06563000000000008)0.868:0.040830000000000144,('WP_139406182.1-Aeromonas-veronii':0.22327999999999992,('WP_241279715.1-Aeromonas-sp.-MR19':0.030219999999999914,'WP_163135231.1-Aeromonas-rivipollensis':0.026949999999999807)1.0:0.1950400000000001)0.895:0.05993000000000004)0.758:0.02118999999999982,((('WP_201955411.1-Aeromonas-veronii':0.05049999999999999,'WP_101347551.1-Aeromonas-sobria':0.06689999999999996)0.923:0.0693999999999999,('WP_268029297.1-Aeromonas-hydrophila':0.09760999999999997,'WP_201942265.1-Aeromonas-veronii':0.12412000000000001)0.986:0.14312000000000014)0.878:0.04364999999999997,((('WP_210600017.1-Aeromonas-sp.-MrichA-1':0.047639999999999905,('WP_050666161.1-Aeromonas-schubertii':0.053220000000000045,'WP_059168611.1-Aeromonas-salmonicida':0.11204999999999998)0.684:0.02145000000000019)0.855:0.04027000000000003,(('WP_069554789.1-Aeromonas-sp.-DNP9':0.13657999999999992,('WP_203764723.1-Aeromonas-caviae':0.08598999999999979,'WP_213320225.1-Aeromonas-caviae':0.04682999999999993)0.946:0.08684000000000003)0.673:0.05271999999999988,('WP_113719982.1-Aeromonas-salmonicida':0.12880000000000003,('WP_257692500.1-Aeromonas-caviae':0.15759999999999996,('WP_234664799.1-Aeromonas-allosaccharophila':0.12089000000000016,'WP_136501435.1-Aeromonas-veronii':0.17594999999999983)0.293:0.03031000000000006)0.548:0.034720000000000084)0.726:0.04815000000000014)0.837:0.0412300000000001)0.846:0.02967999999999993,('WP_225159845.1-Aeromonas-caviae':0.12024999999999997,(('WP_167569591.1-Aeromonas-veronii':0.08352000000000004,(('WP_116547128.1-Plesiomonas-shigelloides':0.0826699999999998,('WP_235463836.1-Aeromonas-australiensis':0.01432000000000011,('WP_201996661.1-Aeromonas-veronii':0.03157999999999994,('WP_139496062.1-Escherichia-coli':0.057110000000000216,'WP_215269322.1-Aeromonas-dhakensis':0.03717000000000015)0.21:0.005850000000000133)0.933:0.03202000000000016)0.471:0.012849999999999806)0.642:0.014969999999999928,((((('WP_247163170.1-Escherichia-coli':0.015330000000000066,'WP_038644959.1-Pantoea':0.06946999999999992)0.994:0.09465000000000012,'WP_033131297.1-Aeromonas-aquatica':0.04709999999999992)0.293:0.014940000000000175,('WP_252422582.1-Aeromonas-taiwanensis':0.019849999999999923,'WP_263719954.1-Aeromonas-caviae':0.07588999999999979)0.888:0.025290000000000035)0.929:0.04295999999999989,'WP_160838676.1-Aeromonas-caviae':0.058089999999999975)0.898:0.030409999999999826,'WP_241344952.1-Aeromonas-sp.-MR16':0.025450000000000195)0.961:0.05088999999999988)0.62:0.045889999999999986)0.89:0.052059999999999995,'WP_049048056.1-Aeromonas-hydrophila':0.09649000000000019)0.99:0.11495999999999995)0.747:0.029959999999999987)0.545:0.029900000000000038)0.762:0.017830000000000013)0.355:0.024210000000000065,(('WP_268037199.1-Aeromonas-dhakensis':0.10099999999999998,('WP_128343177.1-Aeromonas-caviae':0.044099999999999806,('WP_268421127.1-Aeromonas-media':0.06246000000000018,'WP_111910157.1-Aeromonas':0.01896999999999993)0.796:0.013469999999999871)0.968:0.04620999999999986)0.025:0.011649999999999938,('WP_139733908.1-Aeromonas-veronii':0.012179999999999858,('WP_222134508.1-Vibrio-cholerae':0.1338499999999998,(('WP_234676856.1-Aeromonas-rivipollensis':0.019589999999999996,'WP_269382839.1-Aeromonas-hydrophila':0.09816000000000003)0.782:0.007140000000000146,'WP_234579942.1-Aeromonas-salmonicida':0.06850000000000023)0.618:0.00818999999999992)0.774:0.010600000000000165)0.987:0.07561000000000018)0.92:0.04226999999999981)0.81:0.01664999999999983,('WP_265468300.1-Aeromonas-salmonicida':0.10462000000000016,((((('WP_163128050.1-Aeromonas-allosaccharophila':0.08199999999999985,('WP_125597700.1-Aeromonas-salmonicida':0.02149000000000001,'WP_265458754.1-Aeromonas-salmonicida':0.07249000000000017)0.738:0.0035199999999999676)0.999:0.000550,(('WP_180911080.1-Aeromonas-caviae':0.03629999999999978,'WP_264818242.1-Aeromonas-caviae':0.024239999999999817)0.783:0.009949999999999903,'WP_245131329.1-Aeromonas-veronii':0.1616200000000001)0.41:0.008150000000000102)0.099:0.005970000000000031,(('WP_223945983.1-Aeromonas-caviae':0.0842299999999998,('WP_059168599.1-Aeromonas-salmonicida':0.04953999999999992,('WP_033115905.1-Aeromonas-lacus':0.04748000000000019,('WP_223920301.1-Aeromonas-caviae':0.05188000000000015,('WP_257690725.1-Aeromonas-veronii':0.04041000000000006,'WP_058394234.1-Aeromonas-salmonicida':0.03198999999999996)0.793:0.01656000000000013)0.794:0.019839999999999858)0.931:0.045640000000000125)0.974:0.07520999999999978)0.865:0.03318999999999983,(('WP_040098646.1-Aeromonas':0.08693000000000017,('WP_257686696.1-Aeromonas-veronii':0.03752000000000022,('WP_263309349.1-Aeromonas-caviae':0.03708,('WP_255912544.1-Aeromonas-sp.-JL9':0.03074000000000021,'WP_041217344.1-Aeromonas-hydrophila':0.03788999999999998)0.703:0.006279999999999841)0.752:0.007699999999999818)0.767:0.011579999999999924)0.752:0.009360000000000035,(('WP_041996113.1-Aeromonas-bivalvium':0.04691000000000001,('WP_139750984.1-Aeromonas-media':0.02760000000000007,'WP_224430864.1-Aeromonas-rivuli':0.024700000000000166)0.762:0.021069999999999922)0.926:0.03577999999999992,'WP_265776268.1-Aeromonas-veronii':0.12399000000000004)0.866:0.021139999999999937)0.0:0.000540)0.536:0.006870000000000154)0.932:0.0724800000000001,((('WP_201942221.1-Aeromonas-veronii':0.14678000000000013,((((('WP_042657830.1-Aeromonas':0.04524999999999979,'WP_111901368.1-Aeromonas-veronii':0.028690000000000104)0.783:0.011629999999999807,'WP_159143398.1-Aeromonas-veronii':0.02389000000000019)0.329:0.004919999999999813,'WP_203131090.1-Aeromonas-jandaei':0.049100000000000144)0.743:0.006769999999999943,'WP_139692403.1-Aeromonas-sobria':0.029119999999999813)0.725:0.006689999999999863,('WP_223380193.1-Aeromonas-enteropelogenes':0.036189999999999944,'WP_268854930.1-Aeromonas-enteropelogenes':0.10225000000000017)0.642:0.006359999999999921)0.849:0.03623999999999983)0.641:0.04997000000000007,('WP_236321701.1-Aeromonas-veronii':0.06721000000000021,'WP_257119864.1-Aeromonas-hydrophila':0.019379999999999953)0.898:0.05412000000000017)0.088:0.018569999999999975,('WP_257707542.1-Aeromonas-caviae':0.04726000000000008,'WP_182951616.1-Aeromonas-sp.-WP2-W18-CRE-05':0.05790000000000006)0.977:0.06630000000000003)0.89:0.06070999999999982)0.438:0.02416000000000018,('WP_236563281.1-Aeromonas-veronii':0.13911000000000007,'WP_216990617.1-Aeromonas-sp.-FDAARGOS-1409':0.11012999999999984)0.349:0.0284399999999998)0.916:0.044259999999999966)0.966:0.07385999999999981)0.761:0.008960000000000079)0.666:0.0049000000000001265,((('WP_263807164.1-Salmonella-enterica':0.1902499999999998,'WP_048221111.1-Citrobacter':0.09643999999999986)0.991:0.22431,('WP_206164314.1-Citrobacter':0.1760299999999999,('WP_128482629.1-Kosakonia-cowanii':0.10245999999999977,('WP_161518249.1-Kosakonia-sp.-MH5':0.06034000000000006,'WP_097405671.1-Kosakonia':0.040690000000000115)0.931:0.06550000000000011)0.984:0.1494399999999998)0.732:0.03772999999999982)0.033:0.07310999999999979,(('WP_047682696.1-Xenorhabdus':0.045560000000000045,'WP_099124588.1-Xenorhabdus-stockiae':0.07620000000000005)0.99:0.14776999999999996,('WP_217470569.1-Photorhabdus-akhurstii':0.061549999999999994,'WP_074024273.1-Xenorhabdus-eapokensis':0.07362000000000002)0.814:0.05274000000000001)0.956:0.09515999999999991)0.923:0.06994999999999996)0.877:0.05154000000000014,(((('WP_252601295.1-Aeromonas-dhakensis':0.024129999999999985,'WP_139432874.1-Aeromonas-dhakensis':0.1267999999999998)0.837:0.02566999999999986,'WP_106553824.1-Aeromonas-hydrophila':0.06660000000000021)0.864:0.033090000000000064,'WP_100646402.1-Aeromonas-veronii':0.04320000000000013)0.998:0.14835999999999983,(('WP_224211331.1-Aeromonas-enteropelogenes':0.15221000000000018,'WP_082041448.1-Aeromonas-eucrenophila':0.2503500000000001)0.776:0.06490000000000018,('WP_210550341.1-Aeromonas-sp.-SrichE-2G':0.2868200000000001,((((((('WP_159163188.1-Aeromonas-veronii':0.04090999999999978,'WP_098969895.1-Aeromonas-sp.-CU5':0.04667000000000021)0.907:0.043989999999999974,('WP_139495680.1-Aeromonas-veronii':0.035130000000000106,'WP_042019775.1-Aeromonas-sobria':0.059779999999999944)0.755:0.039919999999999956)0.944:0.053770000000000095,'WP_113736183.1-Aeromonas-caviae':0.03135000000000021)0.755:0.010969999999999924,'WP_236369955.1-Aeromonas-veronii':0.04344999999999999)0.775:0.01576999999999984,(('WP_082969903.1-Aeromonas-dhakensis':0.10691000000000006,('WP_104454419.1-Aeromonas-sp.-ASNIH2':0.04197000000000006,'WP_201035193.1-Aeromonas-media':0.12702999999999998)0.475:0.017650000000000166)0.905:0.039060000000000095,(('WP_055647506.1-Shewanella-sp.-Sh95':0.026539999999999786,'WP_220777726.1-Shewanella-hafniensis':0.02972000000000019)0.903:0.03182000000000018,('WP_263173612.1-Shewanella-sp.-SM73':0.14406000000000008,(('WP_109248233.1-Shewanella-algae':0.06976000000000004,'WP_263178828.1-Shewanella-sp.-SM101':0.035820000000000185)0.936:0.046030000000000015,('WP_263191039.1-Shewanella-sp.-SM87':0.06485999999999992,'WP_263255112.1-Shewanella-sp.-SM32':0.040859999999999896)0.719:0.009539999999999882)0.835:0.035559999999999814)0.957:0.0717500000000002)0.725:0.01167000000000007)0.982:0.06654999999999989)0.806:0.04566999999999988,(('WP_139459904.1-Aeromonas-veronii':0.04104999999999981,'WP_041208446.1-Aeromonas-jandaei':0.13518000000000008)0.985:0.08232999999999979,('WP_113739100.1-Aeromonas-veronii':0.04922000000000004,('WP_226032029.1-Aeromonas-hydrophila':0.04298000000000002,((('WP_252615985.1-Aeromonas-hydrophila':0.04869000000000012,('WP_108536035.1-Aeromonas-sp.-HMWF014':0.04051999999999989,'WP_080988732.1-Aeromonas-caviae':0.03376000000000001)0.919:0.0342699999999998)0.471:0.01770000000000005,'WP_214003767.1-Aeromonas-caviae':0.0435500000000002)0.76:0.007519999999999971,'WP_143237783.1-Aeromonas-caviae':0.05832000000000015)0.753:0.007620000000000182)0.723:0.010229999999999961)0.742:0.008849999999999802)0.795:0.029490000000000016)0.889:0.05780999999999992,('WP_200561028.1-unclassified-Erwinia':0.3634599999999999,(('WP_268010485.1-Aeromonas-dhakensis':0.06488000000000005,'WP_098985072.1-Aeromonas-sp.-CA23':0.03645999999999994)0.97:0.09744999999999981,('WP_181268965.1-Aeromonas-veronii':0.03387999999999991,('WP_129146711.1-Aeromonas-caviae':0.021440000000000126,('WP_139744464.1-Aeromonas-media':0.035829999999999806,'WP_081843667.1-Aeromonas-taiwanensis':0.018510000000000026)0.882:0.021999999999999797)0.815:0.015699999999999825)0.93:0.0828199999999999)0.884:0.08755999999999986)0.671:0.08117999999999981)0.878:0.04775000000000018)0.768:0.02915000000000001)0.749:0.02265000000000006)0.742:0.03932000000000002)0.237:0.016249999999999876,('WP_064338398.1-Aeromonas-veronii':0.13962999999999992,(('WP_168366848.1-Dickeya-zeae':0.1675399999999998,('WP_152109808.1-Plesiomonas-shigelloides':0.07010000000000005,'WP_227095090.1-Aeromonas-veronii':0.08165999999999984)0.943:0.09173000000000009)0.835:0.04593999999999987,'WP_252995952.1-Aeromonas-encheleia':0.2161200000000001)0.724:0.04286000000000012)0.907:0.05116999999999994)0.928:0.06757999999999997,(('WP_144817260.1-Pseudescherichia-sp.-L3':0.03377000000000008,'WP_202782010.1-Enterobacter-asburiae':0.0758700000000001)0.998:0.17739999999999956,('WP_125117341.1-Aeromonas-caviae':0.28017000000000003,'WP_035600513.1-Edwardsiella-tarda':0.26081999999999983)0.828:0.03982999999999981)0.845:0.04422000000000015)0.933:0.06825999999999999,('WP_100272167.1-Citrobacter-portucalensis':0.22268000000000043,('WP_269535377.1-Citrobacter-sedlakii':0.2125499999999998,'WP_172410167.1-Escherichia-coli':0.3209399999999998)0.386:0.05510000000000037)0.825:0.05235999999999974)0.74:0.016520000000000312,(((((('WP_115348269.1-Hafnia-alvei':0.14217000000000013,('WP_032659356.1-Enterobacter-hormaechei':0.2745099999999998,('WP_132257458.1-Paucimonas-lemoignei':0.0867300000000002,'WP_119785517.1-Noviherbaspirillum-sedimenti':0.04531000000000018)1.0:0.44087999999999994)0.886:0.08042999999999978)0.827:0.0359799999999999,(('WP_183847446.1-Rahnella-inusitata':0.04917000000000016,'WP_120162698.1-Rahnella-variigena':0.06129000000000007)1.0:0.3302100000000001,('WP_061059890.1-Hafnia-paralvei':0.18501000000000012,'WP_130961887.1-Hafnia-paralvei':0.16529999999999978)0.385:0.025349999999999984)0.391:0.011620000000000186)0.912:0.07085000000000008,('WP_175320495.1-Hafnia-paralvei':0.0885600000000002,'WP_130996599.1-Hafnia-paralvei':0.18084999999999996)0.956:0.09062999999999999)0.946:0.07450000000000001,(('WP_162648791.1-Citrobacter-freundii':0.11198999999999959,('WP_060683585.1-Citrobacter-portucalensis':0.11629999999999985,('WP_079798993.1-Salmonella-enterica':0.08450999999999986,'WP_032255798.1-Enterobacteriaceae':0.07631999999999994)0.528:0.004090000000000149)0.925:0.06384999999999996)0.867:0.055600000000000094,('WP_048218074.1-Citrobacter-freundii-complex':0.15476000000000045,'WP_195994783.1-Citrobacter-portucalensis':0.2001400000000002)0.852:0.053879999999999484)0.84:0.025120000000000253)0.719:0.019179999999999975,(('WP_181624464.1-Citrobacter-freundii':0.21108000000000038,'WP_001515521.1-Escherichia-coli':0.1576200000000001)0.963:0.10327999999999982,('WP_047609274.1-Rahnella-aquatilis':0.17449999999999966,('WP_157953065.1-Limnobaculum-parvum':0.20826999999999973,('WP_112164393.1-Rahnella-inusitata':0.14266999999999985,('WP_061709645.1-Pseudenterobacter-timonensis':0.04457999999999984,'WP_058662478.1-Enterobacteriaceae':0.03765000000000018)0.997:0.17243999999999993)0.573:0.05801999999999996)0.761:0.054079999999999906)0.799:0.024090000000000167)0.936:0.05279999999999996)0.489:0.015519999999999978,(((((('WP_249031739.1-Tenebrionicola-larvae':0.26690000000000014,('WP_123334936.1-Erwinia-sp.-JUb26':0.033110000000000195,'WP_105591414.1-Pantoea-coffeiphila':0.060550000000000104)0.99:0.1660400000000002)0.232:0.06384999999999996,((('WP_103775088.1-Citrobacter-amalonaticus':0.16571999999999987,('WP_181680936.1-Citrobacter-sp.-RHB25-C09':0.09941999999999984,('WP_196013128.1-Citrobacter-farmeri':0.02923000000000009,'WP_090049757.1-Citrobacter-amalonaticus':0.026819999999999844)0.989:0.11265999999999998)0.858:0.04289999999999994)0.984:0.13806999999999992,'WP_121265736.1-Enterobacter-sp.-R1-2018':0.2183600000000001)0.142:0.01933000000000007,'WP_191149682.1-Erwinia-aphidicola':0.1429999999999998)0.938:0.06570000000000009)0.621:0.013189999999999813,(('WP_253383865.1-unidentified-bacterial-endosymbiont':0.2984600000000004,('WP_171269898.1-Aeromonas-media':0.07439000000000018,'WP_216983019.1-Aeromonas-sp.-FDAARGOS-1415':0.09865000000000013)0.972:0.14842000000000022)0.591:0.04380000000000006,'WP_075180894.1-Pantoea-sp.-1.19':0.16425)0.894:0.04826999999999959)0.749:0.018780000000000463,'WP_034458444.1-Buttiauxella-noackiae':0.2471700000000001)0.778:0.02737999999999996,(((('WP_208283396.1-Enterobacter-sichuanensis':0.09469000000000038,(('WP_046888360.1-Enterobacter-cloacae':0.06418000000000035,('WP_258993080.1-Scandinavium-hiltneri':0.07316000000000011,(('WP_203360733.1-Citrobacter-sp.-R56':0.06269999999999998,'WP_001572827.1-Escherichia-coli':0.055629999999999846)0.519:0.017409999999999926,'WP_141109706.1-Escherichia-albertii':0.0747399999999998)0.826:0.04444999999999988)0.964:0.07046000000000019)0.756:0.01980000000000004,((('WP_012906377.1-Citrobacter-rodentium':0.08408000000000015,'WP_095374511.1-Escherichia-coli':0.009469999999999867)0.982:0.08678000000000008,('WP_096122580.1-Escherichia-coli':0.061350000000000016,'WP_162379936.1-Citrobacter-sp.-NCU1':0.16383999999999999)0.66:0.019789999999999974)0.93:0.044340000000000046,('WP_259000881.1-Scandinavium-goeteborgense':0.13648999999999978,'WP_259047621.1-Klebsiella-sp.-BIGb0407':0.18961000000000006)0.602:0.02491000000000021)0.859:0.033740000000000325)0.829:0.017759999999999998)0.209:0.003809999999999647,((('WP_063869545.1-Enterobacter-hormaechei':0.06215999999999999,('WP_241820523.1-Enterobacter-kobei':0.06355999999999984,'WP_042716137.1-Enterobacter-sp.-B509':0.016199999999999992)0.996:0.08655999999999997)0.49:0.013069999999999915,('WP_126546226.1-Enterobacter-quasimori':0.07658999999999994,'WP_172688853.1-Enterobacterales':0.11187000000000014)0.924:0.04891999999999985)0.91:0.03825000000000012,('WP_033487312.1-Enterobacter-cloacae-complex':0.043239999999999945,('WP_126315015.1-Enterobacter-hormaechei':0.10664000000000007,((('WP_213381159.1-Enterobacter-cloacae':0.06755000000000022,'WP_182115563.1-Enterobacter-kobei':0.024070000000000036)0.814:0.058149999999999924,'WP_045338355.1-Enterobacter-hormaechei':0.12969000000000008)0.54:0.03205999999999998,'WP_063158410.1-Enterobacter-hormaechei':0.07945000000000002)0.852:0.029139999999999944)0.474:0.02660000000000018)0.697:0.02613999999999983)0.806:0.02234999999999987)0.751:0.02336000000000027,(((('WP_232904305.1-Buttiauxella-sp.-W03-F01':0.13870000000000005,'WP_034915863.1-Erwinia-sp.-9145':0.14270000000000005)0.297:0.03017000000000003,('WP_230350752.1-Lelliottia-sp.-WAP21':0.15808,'WP_259388919.1-Enterobacter-huaxiensis':0.03471000000000002)0.763:0.026539999999999786)0.89:0.03529999999999989,((('WP_138014015.1-Enterobacter-ludwigii':0.07175999999999982,'WP_032636062.1-Enterobacter-kobei':0.049960000000000004)0.873:0.03824000000000005,('WP_063137673.1-Enterobacter-cloacae-complex':0.031789999999999985,'WP_165464597.1-Enterobacter-cloacae':0.0535000000000001)0.62:0.02942)0.879:0.03763000000000005,('WP_265185975.1-Enterobacter-mori':0.07900000000000018,('WP_193360946.1-Enterobacter-cloacae-complex-sp.-P39RS':0.05271999999999988,'WP_182945676.1-Enterobacter-cloacae':0.01982000000000017)0.969:0.07555000000000023)0.787:0.025069999999999926)0.783:0.018909999999999982)0.762:0.022029999999999994,(('WP_242172213.1-Enterobacter':0.07291999999999987,('WP_100165931.1-Enterobacter-mori':0.029490000000000016,'WP_096205048.1-Enterobacter':0.029189999999999827)0.944:0.07127000000000017)0.859:0.056929999999999925,'WP_239609625.1-Lelliottia-amnigena':0.07957999999999998)0.95:0.06653999999999982)0.065:0.011730000000000018)0.79:0.034889999999999866,((('WP_250341216.1-Pantoea-eucrina':0.2492099999999997,(((('WP_079834758.1-Salmonella-enterica':0.006089999999999929,('WP_001100807.1-Salmonella':0.07458000000000009,'WP_079793206.1-Salmonella-enterica':0.1160000000000001)0.994:0.1415700000000002)0.531:0.03049999999999997,'WP_058109100.1-Salmonella-enterica':0.030850000000000044)0.989:0.0879500000000002,('WP_049002539.1-Citrobacter-freundii':0.1073599999999999,('WP_161617217.1-Yokenella-regensburgei':0.037819999999999965,'WP_249576357.1-Enterobacter-vonholyi':0.057640000000000136)0.951:0.06262999999999996)0.85:0.025349999999999984)0.899:0.04147999999999996,'WP_032303308.1-Escherichia':0.06358999999999959)0.778:0.025409999999999933)0.841:0.03108000000000022,'WP_127791218.1-Escherichia-coli':0.14798)0.72:0.022619999999999862,'WP_021577651.1-Escherichia-coli':0.1708099999999999)0.108:0.012739999999999974)0.916:0.045880000000000365)0.799:0.011889999999999734,'WP_015386517.1-Cronobacter-sakazakii':0.20582000000000011)0.761:0.010639999999999983)0.942:0.03392000000000017)0.75:0.009830000000000005)0.76:0.0040399999999998215)0.991:0.000540,(((((('WP_137650031.1-Escherichia-albertii':0.1266099999999999,((('WP_033870520.1-Enterobacteriaceae':0.19548999999999994,('WP_138098269.1-Jejubacter-calystegiae':0.15013999999999994,('WP_016236761.1-Enterobacteriaceae':0.08318000000000003,'WP_024239504.1-Escherichia-coli':0.10648999999999997)0.945:0.07957999999999998)0.555:0.06131999999999982)0.835:0.054079999999999906,(('WP_001610415.1-Escherichia-coli':0.03994000000000009,'WP_138183825.1-Escherichia-sp.-E4385':0.021419999999999995)0.988:0.16850999999999994,('WP_104705897.1-Escherichia-coli':0.03022999999999998,'WP_089585421.1-Escherichia-coli':0.03954000000000013)0.993:0.15504000000000007)0.86:0.06264000000000003)0.24:0.05469999999999997,'WP_001100804.1-Enterobacteriaceae':0.1521300000000001)0.736:0.03737999999999975)0.842:0.040620000000000545,('WP_073446180.1-Escherichia-coli':0.19835000000000003,'WP_219093375.1-Escherichia-coli':0.17032999999999987)0.264:0.013120000000000243)0.9:0.03742000000000001,('WP_001542972.1-Escherichia-coli':0.12395999999999985,'WP_153879942.1-Citrobacter-freundii':0.16229000000000005)0.899:0.05082000000000031)0.861:0.023699999999999832,((('WP_171149131.1-Erwinia-sp.-JH02':0.18618000000000023,'WP_202667543.1-Lelliottia-amnigena':0.1927500000000002)0.9:0.060329999999999995,((('WP_042093344.1-Escherichia-coli':0.20828999999999986,('WP_100029735.1-Escherichia-coli':0.13422,'WP_001100786.1-Escherichia-coli':0.14118999999999993)0.733:0.041160000000000085)0.707:0.015859999999999985,(('WP_040079227.1-Escherichia-coli':0.0360699999999996,'WP_228937168.1-Escherichia-marmotae':0.026839999999999975)0.857:0.014930000000000554,('WP_050871166.1-Enterobacter-cloacae-complex':0.027160000000000295,'WP_119938542.1-Enterobacter-huaxiensis':0.1073000000000004)0.92:0.04962)0.837:0.023969999999999825)0.926:0.042859999999999676,(('WP_208464084.1-Escherichia-coli':0.0629900000000001,('WP_170897747.1-Escherichia-coli':0.14270000000000005,'WP_001100781.1-Escherichia-coli':0.16325000000000012)0.872:0.09503000000000039)0.946:0.08872999999999998,('WP_001297344.1-Escherichia-coli':0.1642800000000002,('WP_042099316.1-Escherichia-coli':0.042910000000000004,('WP_129995405.1-Enterobacter-cloacae':0.05480000000000018,(('WP_045339085.1-Enterobacter-cloacae-complex':0.1578400000000002,('WP_214593796.1-Enterobacter-mori':0.04971000000000014,'WP_086532312.1-Enterobacter-ludwigii':0.06197000000000008)0.939:0.09167000000000014)0.919:0.08761000000000019,('WP_040022520.1-Enterobacter-cloacae':0.06395000000000017,'WP_047742849.1-Enterobacter':0.07040999999999986)0.988:0.10821000000000014)0.788:0.02301999999999982)0.881:0.03507000000000016)0.977:0.09446999999999983)0.849:0.06117000000000017)0.457:0.03212999999999955)0.734:0.015890000000000182)0.678:0.010389999999999677,((('WP_174633339.1-Escherichia-coli':0.09480999999999984,'WP_072664248.1-Escherichia':0.08499999999999996)0.906:0.07762000000000002,('WP_262931739.1-Escherichia-albertii':0.03609000000000018,'WP_160188772.1-Escherichia-coli':0.040280000000000094)0.964:0.08793999999999969)0.861:0.038899999999999935,((('WP_096972986.1-Escherichia-coli':0.09313000000000038,'WP_089706542.1-Escherichia-coli':0.11426000000000025)0.865:0.045789999999999775,(('WP_181237856.1-Escherichia-marmotae':0.06925000000000026,('WP_001100798.1-Enterobacteriaceae':0.11797999999999975,'WP_061892435.1-Escherichia-coli':0.10374999999999979)0.901:0.06204000000000054)0.155:0.01760999999999946,('WP_109548670.1-Escherichia-coli':0.04770999999999992,'WP_001100790.1-Escherichia':0.014499999999999957)0.96:0.0651799999999998)0.911:0.034820000000000295)0.718:0.015439999999999898,(((('WP_105963664.1-Escherichia-coli':0.06924999999999981,('WP_063610377.1-Escherichia-coli':0.09109999999999996,'WP_053290779.1-Escherichia-coli':0.1139199999999998)0.0:0.000540)0.894:0.0295399999999999,(('WP_218070502.1-Escherichia-coli':0.0673199999999996,('WP_107179425.1-Escherichia':0.11161000000000021,'WP_187209312.1-Escherichia-coli':0.08377999999999997)0.813:0.0313399999999997)0.885:0.04173000000000027,(('WP_085454021.1-Escherichia-coli':0.11702000000000012,('WP_088541377.1-Escherichia-coli':0.022229999999999972,'WP_222691870.1-Escherichia-marmotae':0.05996999999999986)0.68:0.04300000000000015)0.826:0.02597000000000005,(('WP_187216555.1-Escherichia-coli':0.18942000000000014,'WP_059222105.1-Escherichia-albertii':0.09235999999999978)0.82:0.04894000000000043,('WP_096967326.1-Escherichia-coli':0.02507999999999999,'WP_146314246.1-Escherichia-coli':0.03329000000000004)0.91:0.06463000000000019)0.856:0.038679999999999826)0.74:0.010569999999999968)0.563:0.008480000000000043)0.868:0.02126000000000028,'WP_181465273.1-Escherichia-fergusonii':0.06425999999999998)0.782:0.02041000000000004,((('WP_025210617.1-Escherichia-coli':0.13626000000000005,'WP_001527244.1-Escherichia-coli':0.05317000000000016)0.933:0.0631999999999997,('WP_157704420.1-Escherichia-coli':0.09769000000000005,'WP_042200020.1-Enterobacteriaceae':0.0423300000000002)0.85:0.032239999999999824)0.804:0.021990000000000176,(('WP_097421351.1-Escherichia-coli':0.0891099999999998,('WP_001100785.1-Escherichia-coli':0.03326999999999991,'WP_137653986.1-Escherichia-albertii':0.028880000000000017)0.92:0.053609999999999935)0.892:0.03974000000000011,'WP_262938691.1-Escherichia-albertii':0.14697000000000005)0.733:0.01648999999999967)0.796:0.01819000000000015)0.899:0.045299999999999674)0.534:0.009840000000000071)0.887:0.015349999999999753)0.766:0.007830000000000226)0.487:0.010790000000000077,('WP_021540003.1-Escherichia-coli':0.012410000000000032,'WP_047657564.1-Enterobacteriaceae':0.04567000000000032)0.998:0.10299999999999976)0.891:0.019689999999999763,((('WP_267387874.1-Citrobacter-portucalensis':0.24712000000000023,('WP_248249449.1-Enterobacter-roggenkampii':0.16456999999999988,'WP_032936727.1-Citrobacter-freundii':0.09440999999999988)0.998:0.18815999999999988)0.67:0.051979999999999915,(('WP_213789144.1-Enterobacter-hormaechei':0.000660,('WP_237424876.1-Enterobacter-cloacae-complex-sp.-ECC445':0.04234999999999989,('WP_148242178.1-Enterobacter-asburiae':0.046940000000000204,'WP_237994110.1-Enterobacter-sp.-DRP3':0.01750000000000007)0.733:0.0053399999999999)0.882:0.04820000000000002)1.0:0.3055000000000003,(('WP_095844187.1-Escherichia-coli':0.06268999999999991,('WP_086118650.1-Escherichia-coli':0.025450000000000195,'WP_089538038.1-Escherichia-coli':0.03190999999999988)0.593:0.006359999999999921)0.939:0.10342999999999991,('WP_179154141.1-Citrobacter-freundii':0.25550000000000006,'WP_046458830.1-Obesumbacterium-proteus':0.27986999999999984)0.22:0.07534000000000018)0.874:0.05690000000000017)0.814:0.050049999999999706)0.376:0.0284399999999998,(('WP_086539147.1-Citrobacter-freundii':0.26205999999999996,'WP_220393175.1-Citrobacter-freundii':0.30046000000000017)0.81:0.038590000000000124,(('WP_181673375.1-Escherichia-fergusonii':0.04377999999999993,'WP_107192407.1-Escherichia-albertii':0.016570000000000196)0.999:0.1753800000000001,(('WP_262183194.1-Enterobacter-mori':0.1445599999999998,('WP_047027451.1-Enterobacter-kobei':0.04565999999999981,'WP_063412371.1-Enterobacter-cloacae':0.04137000000000013)0.955:0.09122999999999992)0.994:0.17469,('WP_167816599.1-Enterobacter-cloacae':0.27303999999999995,(('WP_087893724.1-Enterobacteriaceae':0.04305000000000003,'WP_073278747.1-Escherichia-coli':0.06519999999999992)0.902:0.07593000000000005,('WP_059217428.1-Escherichia-albertii':0.031629999999999825,('WP_032214194.1-Escherichia-coli':0.04294999999999982,'WP_085459683.1-Escherichia-coli':0.09208000000000016)0.208:0.016300000000000203)0.934:0.07161000000000017)0.943:0.10381000000000018)0.207:0.07618999999999998)0.213:0.034580000000000055)0.416:0.005920000000000147)0.877:0.030579999999999607)0.81:0.019379999999999953)0.776:0.006800000000000139)0.963:0.05194999999999972,((('WP_108569539.1-unclassified-Pantoea':0.33794999999999975,'WP_157959988.1-Citrobacter':0.34582999999999986)0.879:0.11288999999999971,('WP_168756429.1-Aeromonas-hydrophila':0.37019,'WP_061279316.1-Kluyvera-cryocrescens':0.35272999999999977)0.9:0.10577999999999976)0.72:0.026390000000000136,(('WP_194857598.1-Citrobacter-freundii':0.31352,(('WP_252032207.1-Enterobacter-asburiae':0.04502000000000006,'WP_111966211.1-Enterobacter-cloacae-complex':0.027099999999999902)0.843:0.03207000000000004,('WP_196390204.1-Enterobacter-mori':0.039039999999999964,'WP_141032926.1-Escherichia-coli':0.1376900000000001)0.047:0.026489999999999903)0.966:0.16179999999999994)0.985:0.1934800000000001,(('WP_230863267.1-Enterobacter-cloacae':0.35965999999999987,('WP_044720521.1-Escherichia-coli':0.3955899999999999,(('WP_151711904.1-Enterobacter-ludwigii':0.034009999999999874,('WP_058678686.1-Enterobacter':0.06985000000000019,'WP_032941356.1-Citrobacter-freundii':0.19417000000000018)0.913:0.057479999999999976)0.719:0.05489999999999995,'WP_223560970.1-unclassified-Enterobacter':0.012119999999999909)0.985:0.2284600000000001)0.957:0.2345600000000001)0.812:0.08052999999999999,((('WP_164716793.1-Cedecea-lapagei':0.33626999999999985,'WP_034496584.1-Buttiauxella-agrestis':0.3339500000000002)0.861:0.09857000000000005,'WP_050862535.1-Enterobacter':0.38207000000000013)0.823:0.040610000000000035,'WP_029399168.1-Escherichia-coli':0.26812000000000014)0.921:0.07634000000000007)0.95:0.10758000000000001)0.664:0.028739999999999988)0.711:0.009509999999999685)0.89:0.024880000000000457,(('WP_036421240.1-Morganella-morganii':0.14788999999999985,('WP_264998074.1-Morganella-morganii':0.07759999999999989,'WP_186849509.1-Morganella-morganii':0.08069999999999977)0.928:0.06722000000000028)0.898:0.11175000000000024,'WP_064374050.1-Pantoea':0.3052400000000004)0.936:0.08166999999999991)0.797:0.019909999999999872)0.174:0.0015100000000001224)0.845:0.030389999999999695,('WP_159282060.1-Rahnella-variigena':0.32504,('WP_039304623.1-Cedecea-neteri':0.36214000000000013,('WP_106902182.1-Citrobacter-freundii':0.21764000000000028,('WP_126291239.1-Enterobacter-hormaechei':0.1453000000000002,('WP_049008289.1-Enterobacter-cloacae':0.040759999999999685,('WP_063144508.1-Enterobacter-cloacae':0.03353000000000028,(('WP_252042312.1-Enterobacter-chengduensis':0.038439999999999586,'WP_062932819.1-Enterobacter-hormaechei':0.02893999999999952)0.825:0.024110000000000298,'WP_048999124.1-Enterobacter':0.035009999999999764)0.74:0.007589999999999986)0.848:0.02220999999999984)0.772:0.03874000000000022)0.931:0.07806000000000024)0.436:0.05948999999999982)0.742:0.033500000000000085)0.902:0.047169999999999934)0.717:0.01794999999999991,(('WP_038223662.1-Xenorhabdus-bovienii':0.20978999999999992,('WP_086113286.1-Xenorhabdus-beddingii':0.22419999999999973,'WP_038258850.1-Xenorhabdus-bovienii':0.21860999999999997)0.707:0.0524300000000002)0.821:0.030139999999999834,('WP_257124601.1-Aeromonas-veronii':0.2820800000000001,'WP_211007190.1-Pectobacterium-brasiliense':0.4701400000000002)0.926:0.11262999999999979)0.65:0.016739999999999977)0.936:0.06631000000000009)0.789:0.03330000000000011,('WP_153752706.1-unclassified-Citrobacter':0.54731,((('WP_145394131.1-Pantoea-sp.-SJZ147':0.12688999999999995,'WP_172912938.1-Pantoea-stewartii':0.1174599999999999)0.978:0.22706999999999988,('WP_061356323.1-Escherichia-coli':0.5937099999999997,'WP_086868392.1-Kosakonia-pseudosacchari':0.52779)0.397:0.15690999999999988)0.868:0.11233999999999966,(('WP_181599127.1-Citrobacter-sp.-RHB36-C18':0.20313000000000025,('WP_065370581.1-Kosakonia-sacchari':0.19311999999999996,'WP_241647798.1-Rosenbergiella-sp.-JB07':0.2559999999999998)0.856:0.04151000000000016)0.849:0.04898999999999987,('WP_160868038.1-Pantoea-sp.-Taur':0.29352,(('WP_008499438.1-Enterobacter-cloacae-complex':0.23641999999999985,(((('WP_123822891.1-Klebsiella':0.03440000000000021,(('WP_212709529.1-Tatumella-sp.-JGM130':0.015260000000000051,'WP_095100919.1-Raoultella-sp.-10-1':0.05096999999999996)0.972:0.07902999999999993,'WP_236948353.1-Klebsiella-sp.-2019SCSN059':0.09135999999999989)0.842:0.03593000000000002)0.594:0.03058000000000005,(((('WP_139968923.1-Klebsiella-pneumoniae':0.11151,('WP_048284323.1-Pluralibacter-gergoviae':0.11769000000000007,'WP_211776196.1-Klebsiella-pneumoniae':0.06400000000000006)0.313:0.01462999999999992)0.867:0.029389999999999805,'WP_213760001.1-Klebsiella-aerogenes':0.051899999999999835)0.724:0.008859999999999868,('WP_221694709.1-Klebsiella-aerogenes':0.09000999999999992,((('WP_128320364.1-Raoultella-planticola':0.05419000000000018,'WP_223370324.1-Klebsiella-pneumoniae':0.0360100000000001)0.515:0.006380000000000052,(('WP_186601692.1-Klebsiella-quasipneumoniae':0.017139999999999933,('WP_113990968.1-Raoultella':0.026839999999999975,'WP_234537246.1-Raoultella-ornithinolytica':0.03514000000000017)0.977:0.046969999999999956)0.874:0.01205999999999996,(('WP_047723323.1-Klebsiella-michiganensis':0.05101000000000022,'WP_104011323.1-Klebsiella-michiganensis':0.02566999999999986)0.939:0.02925999999999984,'WP_234579544.1-Raoultella-terrigena':0.02218000000000009)0.763:0.006180000000000074)0.89:0.012480000000000047)0.789:0.009949999999999903,(('WP_149270444.1-Klebsiella-aerogenes':0.0253899999999998,('WP_095100910.1-Raoultella-sp.-10-1':0.05455999999999994,'WP_065799756.1-Klebsiella-pneumoniae':0.018299999999999983)0.881:0.028669999999999973)0.353:0.010229999999999961,('WP_046852082.1-Raoultella-planticola':0.03304000000000018,('WP_123755035.1-Raoultella-sp.-BIGb0399':0.028820000000000068,('WP_114692711.1-Klebsiella-pneumoniae':0.03376000000000001,('WP_259891680.1-Klebsiella-variicola':0.022889999999999855,'WP_236948387.1-Klebsiella-sp.-2019SCSN059':0.039000000000000146)0.402:0.00566000000000022)0.613:0.005090000000000039)0.759:0.02159000000000022)0.854:0.01244999999999985)0.82:0.011859999999999982)0.614:0.0010599999999998388)0.736:0.012589999999999879)0.859:0.03351000000000015,'WP_182931197.1-Enterobacter-cloacae':0.14308999999999994)0.852:0.03498000000000001)0.991:0.09039999999999981,(('WP_044173167.1-Phytobacter-massiliensis':0.03750000000000009,('WP_063501760.1-Escherichia-coli':0.03349000000000002,'XP_011503919.1-Ceratosolen-solmsi-marchali':0.07902999999999993)0.937:0.052529999999999966)0.973:0.07571000000000039,('WP_072663757.1-Enterobacteriaceae':0.08651000000000009,'WP_021241446.1-Enterobacter-roggenkampii':0.1774600000000004)0.742:0.021279999999999966)0.953:0.058829999999999494)0.799:0.020929999999999893,('WP_090085547.1-Escherichia-coli':0.036909999999999776,('WP_234083964.1-Enterobacter-quasiroggenkampii':0.1349800000000001,'WP_103283990.1-Citrobacter':0.16394999999999982)0.587:0.02668999999999988)0.999:0.1654)0.821:0.031140000000000168)0.806:0.029450000000000198,('WP_183170804.1-Atlantibacter-sp.-RC6':0.2183299999999999,('WP_203451401.1-Lelliottia-sp.-RWM.1':0.0571600000000001,'WP_185654835.1-Citrobacter-braakii':0.11021999999999998)0.876:0.05967999999999973)0.956:0.08175000000000043)0.824:0.027789999999999537)0.798:0.03108000000000022)0.995:0.15505999999999975)0.773:0.022050000000000125)0.825:0.03882999999999992)0.906:0.05384000000000011)0.334:0.0041599999999997195)0.903:0.04499999999999993)0.838:0.022210000000000285,('WP_028018520.1-Enterobacter-cloacae-complex':0.7413099999999999,(('WP_071788415.1-Franconibacter-pulveris':0.4230400000000003,('WP_232581009.1-Photobacterium-phosphoreum':0.5278399999999999,('WP_073514120.1-Vibrio-parahaemolyticus':0.05787999999999993,'WP_032481483.1-Vibrio':0.013799999999999812)0.997:0.51545)0.972:0.3182800000000001)0.187:0.032529999999999504,('WP_020232007.1-Escherichia-coli':0.2613699999999999,('WP_228338545.1-Enterobacter-hormaechei':0.13911000000000007,('WP_150146763.1-Enterobacter-roggenkampii':0.04032999999999998,'WP_109845273.1-Enterobacter':0.04486999999999952)0.73:0.06099000000000032)0.988:0.16690999999999967)0.906:0.07437000000000005)0.813:0.044150000000000134)0.902:0.08349999999999991)0.807:0.030289999999999928)0.872:0.0535300000000003)0.662:0.05830999999999964,('WP_011189464.1-Desulfotalea-psychrophila':0.40531000000000006,((('WP_138986566.1-Neptunomonas-concharum':0.2869099999999998,(((((('WP_269599241.1-Salinivibrio-kushneri':0.06040000000000001,'WP_136483218.1-Vibrio-sp.-H11':0.05909999999999993)0.956:0.06390999999999991,('WP_159656123.1-Vibrio-atypicus':0.04783000000000026,'WP_193630602.1-Shewanella-benthica':0.1804300000000003)0.896:0.05286999999999997)0.913:0.044720000000000315,('WP_164649367.1-Vibrio-astriarenae':0.08681000000000028,(((('WP_047874860.1-Photobacterium-aphoticum':0.20587999999999962,'WP_205390674.1-Vibrio':0.07029999999999959)0.828:0.0269200000000005,('WP_022943097.1-Psychromonas-hadalis':0.13101999999999947,'WP_020335761.1-Vibrio-natriegens':0.09648999999999974)0.859:0.04196000000000044)0.753:0.008859999999999868,(('WP_047049276.1-Vibrio-mexicanus':0.14590000000000014,'WP_012143346.1-Shewanella-sediminis':0.18057000000000034)0.288:0.03866999999999976,'WP_140076235.1-Vibrio-parahaemolyticus':0.10611000000000015)0.75:0.017209999999999948)0.743:0.011290000000000244,((('WP_113818410.1-Psychromonas-sp.-B3M02':0.0453800000000002,('WP_166372414.1-Psychromonas-sp.-SA13A':0.013040000000000163,'WP_025564776.1-Psychromonas-sp.-SP041':0.04366000000000003)0.89:0.051750000000000185)0.956:0.05815999999999999,('WP_155694061.1-Psychrosphaera-haliotis':0.1114200000000003,(('WP_188955407.1-Shewanella-ulleungensis':0.05359999999999987,('WP_167678613.1-Shewanella-aestuarii':0.1298600000000003,'WP_137222113.1-Shewanella-sp.-MEBiC00475':0.02352000000000043)0.905:0.03797999999999968)0.867:0.031709999999999905,(('WP_126005793.1-Pseudoalteromonas-sp.-Xi13':0.028410000000000046,('WP_138577097.1-unclassified-Pseudoalteromonas':0.08701999999999988,('WP_264390127.1-Pseudoalteromonas-sp.-A3':0.022410000000000263,('WP_248462211.1-Pseudoalteromonas-sp.-2CM37A':0.016439999999999788,'WP_096739047.1-Pseudoalteromonas-atlantica':0.03438000000000008)0.919:0.018829999999999902)0.793:0.013970000000000038)0.95:0.041630000000000056)0.011:0.0034199999999997566,('WP_256848444.1-Pseudoalteromonas-carrageenovora':0.016550000000000065,'WP_182693438.1-Pseudoalteromonas-sp.-SR45-5':0.06040000000000001)0.758:0.008239999999999803)0.98:0.07108000000000025)0.597:0.04056000000000015)0.609:0.030359999999999943)0.915:0.02426999999999957,('WP_050648448.1-Vibrio-sp.-J2-4':0.1098300000000001,'WP_264874461.1-Vibrio-agarivorans':0.09976999999999991)0.875:0.02876999999999974)0.718:0.010170000000000456)0.91:0.02835999999999972)0.055:0.007700000000000262)0.443:0.007229999999999848,('WP_084884043.1-Vibrio-sp.-qd031':0.12128999999999968,'WP_093326971.1-Thalassotalea-agarivorans':0.11868999999999996)0.749:0.012310000000000265)0.938:0.0498700000000003,(('WP_229491984.1-Enterovibrio-norvegicus':0.1072099999999998,'WP_016959527.1-Enterovibrio-norvegicus':0.06350999999999996)0.927:0.058470000000000244,('WP_167868984.1-Shewanella-sp.-WXL01':0.3714200000000001,('WP_013344594.1-Ferrimonas-balearica':0.12368999999999986,'WP_222525605.1-Ferrimonas-balearica':0.14644000000000013)0.874:0.09962999999999989)0.925:0.0789500000000003)0.194:0.008680000000000021)0.883:0.03662999999999972,((((((('WP_160153838.1-Microbulbifer-sp.-ALW1':0.2331500000000002,('WP_029249715.1-Microbulbifer-agarilyticus':0.09392999999999985,('WP_263600466.1-Microbulbifer-agarilyticus':0.12643000000000004,'WP_078085574.1-Microbulbifer-mangrovi':0.07279999999999998)0.726:0.012309999999999821)0.914:0.0863299999999998)0.968:0.09452999999999978,(('WP_111640782.1-Marinimicrobium-alkaliphilum':0.31396999999999986,'WP_189610147.1-Saccharospirillum-salsuginis':0.27278)0.85:0.06252000000000013,('WP_130050482.1-Pseudoalteromonas-sp.-MEBiC-03485':0.11821999999999999,'WP_138624780.1-Pseudoalteromonas-sp.-S4492':0.2302599999999999)0.821:0.09127000000000018)0.909:0.08366999999999969)0.808:0.04366000000000003,'WP_118961222.1-Colwellia-sp.-RSH04':0.3397899999999998)0.53:0.022619999999999862,'WP_186505417.1-Neptunicella-marina':0.20331999999999972)0.94:0.07681000000000004,((('WP_214344498.1-Bowmanella-yangjiangensis':0.03041999999999989,'WP_102798355.1-Bowmanella-denitrificans':0.06868000000000007)1.0:0.17971000000000004,(('WP_131095783.1-Alteromonas-sp.-KUL42':0.08684000000000003,(('WP_262993134.1-Alteromonas-sp.-ASW11-19':0.060430000000000206,('WP_218354448.1-Alteromonas-lipotrueiana':0.07267999999999963,'WP_195811108.1-Salinimonas-marina':0.052959999999999674)0.85:0.044310000000000294)0.797:0.016180000000000305,((('WP_073325432.1-Marisediminitalea-aggregata':0.049249999999999794,'WP_070176406.1-Alteromonas-lipolytica':0.024599999999999955)0.951:0.04039999999999999,((('WP_216028209.1-Alteromonas-sp.-C1M14':0.044319999999999915,('WP_018984642.1-Salinimonas-chungwhensis':0.08596000000000004,'WP_265616768.1-Alteromonas-sp.-ASW11-7':0.06150000000000011)0.802:0.03840999999999983)0.82:0.015960000000000196,('WP_216026947.1-Aestuariibacter-sp.-A3R04':0.06455999999999973,'WP_265607972.1-Alteromonas-sp.-ASW11-130':0.06943999999999972)0.756:0.011600000000000499)0.782:0.009710000000000107,'WP_119499891.1-Alteromonas-sp.-RKMC-009':0.06652000000000013)0.709:0.011649999999999494)0.941:0.03078000000000003,('WP_163085646.1-Alteromonas-profundi':0.036290000000000155,(((('WP_202940767.1-Alteromonas-macleodii':0.017730000000000246,('WP_061095937.1-Alteromonas-macleodii':0.040069999999999606,((((('WP_156078883.1-Alteromonas':0.053749999999999964,'WP_014977514.1-Alteromonas-macleodii':0.021990000000000176)0.751:0.008550000000000058,'WP_239220085.1-Alteromonas-sp.-MmMcT2-2':0.02336999999999989)0.775:0.000540,('WP_044058190.1-Alteromonas-australica':0.03555000000000019,'WP_159580202.1-Alteromonas-sp.-KUL106':0.017450000000000188)0.791:0.010909999999999975)0.435:0.005830000000000002,('WP_202262575.1-unclassified-Alteromonas':0.04267000000000021,'WP_187447473.1-Alteromonas-sp.-BZK5':0.017879999999999896)0.752:0.005479999999999929)0.776:0.0048400000000001775,'WP_063457834.1-Alteromonas-stellipolaris':0.029279999999999973)0.721:0.00641999999999987)0.079:0.00651000000000046)0.9:0.021090000000000053,((('WP_100972994.1-Alteromonas-sp.-MB-3u-76':0.08387000000000011,'WP_269520955.1-Alteromonas-sp.-BMJM2':0.045970000000000066)0.638:0.025510000000000144,('WP_198816918.1-Alteromonas-sp.-IB21':0.01800999999999986,'WP_071816068.1-Alteromonas-sp.-V450':0.08661999999999992)0.824:0.01725000000000021)0.85:0.019279999999999742,'WP_163106912.1-Alteromonas-genovensis':0.027169999999999916)0.839:0.01456000000000035)0.164:0.004889999999999617,'WP_057795858.1-Alteromonas-stellipolaris':0.03537999999999997)0.888:0.01700999999999997,('WP_130638173.1-Alteromonas-sp.-KUL49':0.056219999999999715,'WP_115592514.1-Alteromonas-aestuariivivens':0.0362199999999997)0.544:0.020459999999999923)0.742:0.007360000000000255)0.894:0.022059999999999746)0.864:0.01702000000000048)0.901:0.03521999999999981)0.875:0.03025000000000011,('WP_216048042.1-Aliiglaciecola-lipolytica':0.18023000000000033,'WP_057181266.1-Colwellia':0.20671000000000017)0.872:0.036029999999999784)0.877:0.03523999999999994)0.724:0.023309999999999942,(('WP_100655941.1-Alteromonas-flava':0.2314400000000001,'WP_124026287.1-Alteromonas-sediminis':0.1422500000000002)0.972:0.10986999999999991,'WP_009839692.1-Pseudoalteromonas-tunicata':0.5775200000000003)0.68:0.03892999999999969)0.626:0.004369999999999763)0.986:0.10931000000000024,((('WP_220103444.1-Neiella-holothuriorum':0.08498999999999945,'WP_087505717.1-Neiella-marina':0.14656999999999965)0.963:0.07842000000000038,(('WP_028862352.1-Psychromonas-aquimarina':0.17436000000000007,'WP_187408568.1-Saccharophagus-sp.-K07':0.1417900000000003)0.849:0.041979999999999684,(('WP_096084688.1-Agaribacterium-haliotis':0.3414999999999999,'WP_188151970.1-Teredinibacter-waterburyi':0.1885300000000001)0.948:0.13988000000000023,'WP_251261538.1-Echinimonas-agarilytica':0.12626999999999988)0.162:0.020469999999999988)0.947:0.07540000000000013)0.809:0.06662999999999997,('WP_152781274.1-Agarivorans-sp.-B2Z047':0.07542999999999989,'WP_248714967.1-Agarivorans-sp.-TSD2052':0.05763000000000007)0.999:0.16469000000000023)0.483:0.03644999999999987)0.232:0.011749999999999705,(('WP_152451766.1-Microbulbifer-sp.-THAF38':0.13709999999999978,'WP_250459773.1-Microbulbifer-sp.-GX-H0434':0.12334000000000023)0.99:0.13392999999999988,('WP_142941196.1-Aliikangiella-marina':0.28157999999999994,('WP_068546663.1-Thalassotalea-crassostreae':0.2033999999999998,('WP_007634918.1-Paraglaciecola-psychrophila':0.030980000000000008,'WP_081148784.1-Cognaticolwellia-beringensis':0.04422000000000015)1.0:0.1912499999999997)0.691:0.02992000000000017)0.885:0.0526399999999998)0.854:0.04657)0.913:0.04708000000000023)0.877:0.06259999999999977)0.949:0.08598000000000017,(((('WP_007466453.1-Photobacterium-marinum':0.25102,((('WP_269294270.1-Pantoea-sp.-LMR881':0.04538999999999982,'WP_215846277.1-Pantoea-sp.-EnvD':0.06454999999999966)0.966:0.06298000000000048,('WP_120453320.1-Kalamiella-piersonii':0.10173999999999994,((('WP_241623482.1-Rosenbergiella-epipactidis':0.02322000000000024,'WP_214216788.1-Rosenbergiella-epipactidis':0.05152000000000001)0.993:0.13034999999999997,('WP_192411534.1-Pantoea-sp.-PNT02':0.059800000000000075,('WP_097116658.1-Pantoea-sp.-GL120224-02':0.000550,('WP_125311258.1-Pantoea-dispersa':0.07543000000000033,('WP_069025179.1-Pantoea-agglomerans':0.005899999999999572,('WP_150019269.1-unclassified-Pantoea':0.061590000000000256,'WP_140917215.1-Pantoea-deleyi':0.03835999999999995)0.433:0.005249999999999755)0.954:0.032410000000000494)0.884:0.020139999999999603)0.796:0.047590000000000465)0.957:0.08591999999999977)0.852:0.040470000000000006,'WP_187508308.1-Pantoea-agglomerans':0.17755999999999972)0.782:0.038040000000000074)0.048:0.02018000000000031)0.348:0.021059999999999857,(('WP_023330866.1-Enterobacter':0.04150000000000009,('WP_088208149.1-Enterobacter-cancerogenus':0.10255999999999998,'WP_121932958.1-Enterobacter-hormaechei':0.08562000000000003)0.9:0.04695000000000027)0.934:0.09246999999999961,'WP_144135068.1-Kosakonia-radicincitans':0.2934199999999998)0.891:0.07215000000000016)0.779:0.03147000000000011)0.564:0.03448000000000029,('WP_211689658.1-Moritella-sp.-28':0.24685000000000024,('WP_085356321.1-Neisseria-dumasiana':0.19757000000000025,(('WP_005552781.1-Aggregatibacter-actinomycetemcomitans':0.1903999999999999,'WP_101133177.1-Neisseria-meningitidis':0.04289999999999994)0.824:0.03438000000000008,('WP_237092367.1-Neisseria-sp.-ZJ104':0.15661999999999976,'WP_127202078.1-Neisseria-meningitidis':0.09018999999999977)0.228:0.041850000000000165)0.39:0.02153000000000027)0.995:0.20455999999999985)0.93:0.10909000000000013)0.372:0.03634000000000004,'WP_031579865.1-Ruminobacter-sp.-RM87':0.4612600000000002)0.664:0.010529999999999706,(('WP_047883806.1-Photobacterium-ganghwense':0.1580699999999995,('WP_187463805.1-Photobacterium-sp.-BZF1':0.04255999999999993,('WP_146157089.1-Photobacterium-rosenbergii':0.15598,'WP_039467548.1-Photobacterium-gaetbulicola':0.07466999999999979)0.926:0.0880399999999999)0.346:0.03697999999999979)0.658:0.03622000000000014,('WP_105076797.1-Pantoea-ananatis':0.40886999999999984,('WP_188026786.1-Tolumonas-osonensis':0.020190000000000374,'WP_024870911.1-Tolumonas-lignilytica':0.11760999999999999)0.912:0.06710999999999956)0.692:0.03603000000000023)0.936:0.04881000000000002)0.839:0.03363000000000005)0.844:0.056459999999999955,'WP_182744595.1-Shewanella-sp.-SR44-4':0.2181099999999998)0.902:0.10319000000000011)0.12:0.03090999999999955)0.798:0.030600000000000183)0.84:0.03628000000000009,('WP_012883389.1-Dickeya-parazeae':0.1893100000000003,'WP_050928752.1-Yersinia-enterocolitica':0.23646000000000011)0.795:0.0432699999999997)0.923:0.05292999999999992,(('WP_166696422.1-Providencia-rettgeri':0.14423999999999992,(('WP_155660100.1-Aliivibrio-fischeri':0.07541999999999982,'WP_095457151.1-Vibrio':0.08183000000000007)0.945:0.04614999999999991,('WP_243584106.1-Vibrio-splendidus':0.14694999999999947,'WP_252020469.1-Vibrio-sp.-SCSIO-43136':0.09762000000000004)0.83:0.03361999999999998)0.907:0.042540000000000244)0.927:0.05644999999999989,('WP_108452928.1-Enterobacter-cloacae':0.29807000000000006,('WP_220058598.1-Shewanella-sp.-FJAT-53555':0.21272000000000002,('WP_212599444.1-Shewanella-sp.-M16':0.1951400000000003,('WP_248994245.1-Shewanella-gaetbuli':0.20938999999999997,('WP_261298106.1-Shewanella-sp.-SP1S2-7':0.09170999999999996,'WP_167678467.1-Shewanella-aestuarii':0.09306999999999999)0.996:0.23566999999999982)0.964:0.15876)0.931:0.1233200000000001)0.811:0.05746000000000029)0.856:0.044079999999999675)0.794:0.02283999999999997)0.709:0.02513999999999994,(((('WP_133040538.1-Shewanella-fodinae':0.2599499999999999,('WP_088905011.1-Shewanella-marisflavi':0.21769999999999978,'WP_234235026.1-Shewanella-sp.-AS16':0.1508799999999999)0.891:0.06695999999999991)0.73:0.0293300000000003,(('WP_222242368.1-Pseudoalteromonas-piscicida':0.27661000000000024,'WP_243025268.1-Vibrio-cincinnatiensis':0.4372600000000002)0.436:0.06325999999999965,('WP_031827242.1-Vibrio-parahaemolyticus':0.21708999999999978,('WP_232612305.1-Photobacterium-phosphoreum':0.31834000000000007,'WP_136989485.1-Vibrio-sp.-F13':0.22551999999999994)0.919:0.10352999999999968)0.719:0.035050000000000026)0.812:0.03398000000000012)0.778:0.011849999999999916,((('WP_172851426.1-Vibrio-coralliilyticus':0.1542300000000001,'WP_138522257.1-Pseudoalteromonas-galatheae':0.25050000000000017)0.519:0.03186999999999962,('WP_158173598.1-Vibrio-alginolyticus':0.0943000000000005,'WP_213886549.1-Vibrio-alginolyticus':0.15652000000000044)0.849:0.03524999999999956)0.946:0.09026000000000023,(('WP_237309237.1-Vibrio':0.2565299999999997,'WP_161680902.1-Proteus':0.28634000000000004)0.878:0.07767000000000035,('WP_076542412.1-Shewanella-sp.-UCD-KL21':0.30528999999999984,'WP_120356561.1-Alginatibacterium-sediminis':0.30989999999999984)0.409:0.05111000000000043)0.732:0.03135999999999983)0.572:0.01471)0.742:0.009930000000000216,(('WP_045031875.1-Photobacterium-phosphoreum':0.020790000000000308,'WP_107213458.1-Photobacterium-phosphoreum':0.0342699999999998)0.999:0.14471999999999996,('WP_193154481.1-Vibrio-navarrensis':0.13938999999999968,('WP_232603643.1-Photobacterium-carnosum':0.09905000000000008,'WP_130314193.1-Vibrio-vulnificus':0.11644999999999994)0.855:0.04292000000000007)0.841:0.03404000000000007)0.844:0.03901000000000021)0.834:0.023829999999999796)0.67:0.019730000000000025)0.768:0.01181999999999972)0.84:0.010950000000000237,(((((('WP_005597034.1-Vibrio-scophthalmi':0.15160000000000018,('WP_259574191.1-Shewanella-baltica':0.2119399999999998,'WP_243584123.1-Vibrio-splendidus':0.1208499999999999)0.061:0.056090000000000195)0.664:0.021709999999999674,('WP_135445143.1-Vibrio-tasmaniensis':0.1856800000000005,((('WP_017420706.1-Vibrio-vulnificus':0.09627000000000052,'WP_211915530.1-Vibrio-sp.-B1REV9':0.10539000000000032)0.968:0.0827899999999997,'WP_182796407.1-Vibrio-sp.-SG41-7':0.1347999999999998)0.778:0.0195599999999998,(('WP_019826652.1-Vibrio-splendidus':0.10960000000000036,('WP_102482947.1-Vibrio-splendidus':0.11532999999999971,'WP_102571993.1-Vibrio':0.10000999999999971)0.4:0.029220000000000468)0.846:0.02815999999999974,('WP_260256886.1-Vibrio-sp.-NC2':0.10594000000000037,'WP_130049361.1-Aliivibrio-finisterrensis':0.06829000000000018)0.8:0.03296999999999972)0.812:0.023299999999999876)0.0:0.000550)0.421:0.017089999999999606)0.904:0.030110000000000525,(((('WP_102285079.1-Vibrio-lentus':0.11529999999999996,(('WP_132940416.1-Vibrio-crassostreae':0.1069,'WP_102457030.1-Vibrio-cyclitrophicus':0.12327000000000021)0.834:0.03750000000000009,'WP_017063118.1-Vibrio':0.09748000000000001)0.146:0.02143000000000006)0.851:0.041589999999999794,'WP_155682312.1-Aliivibrio-fischeri':0.11122999999999994)0.691:0.00865999999999989,'WP_009843931.1-Vibrio-sp.-AND4':0.08309999999999995)0.945:0.058820000000000316,(('WP_102298256.1-Vibrio-cyclitrophicus':0.09467999999999988,'WP_265133188.1-Vibrio-splendidus':0.07703999999999978)0.699:0.02847000000000044,(('WP_248388778.1-Vibrio-cyclitrophicus':0.13833000000000029,(('WP_087845049.1-Photobacterium-malacitanum':0.12280999999999986,'WP_163929979.1-Paraferrimonas-sp.-SM1919':0.11867000000000028)0.844:0.048740000000000006,(('WP_102462022.1-Vibrio-splendidus':0.04268000000000027,'WP_187463684.1-Photobacterium-sp.-BZF1':0.0702799999999999)0.985:0.09925000000000006,'WP_171342182.1-Vibrio-cyclitrophicus':0.1492899999999997)0.15:0.02507999999999999)0.829:0.0351600000000003)0.234:0.02110999999999974,('WP_102448490.1-Vibrio-splendidus':0.1079399999999997,'WP_063660043.1-Aliivibrio-fischeri':0.1217299999999999)0.875:0.04797999999999991)0.79:0.019010000000000193)0.628:0.011489999999999778)0.867:0.022550000000000292)0.792:0.018029999999999546,(((('WP_017105169.1-Vibrio':0.1491800000000003,('WP_135380937.1-Vibrio-tasmaniensis':0.1088300000000002,'WP_059021773.1-Vibrio-coralliirubri':0.0470600000000001)0.802:0.032420000000000115)0.66:0.022969999999999935,(('WP_102276040.1-Vibrio-lentus':0.08908000000000005,'WP_102385704.1-Vibrio-cyclitrophicus':0.1129500000000001)0.905:0.04709999999999992,('WP_102353859.1-Vibrio-cyclitrophicus':0.09411000000000014,('WP_004733311.1-Vibrio-splendidus':0.10302999999999995,'WP_171315631.1-Vibrio-splendidus':0.10775999999999986)0.894:0.04651000000000005)0.898:0.06006)0.859:0.04268000000000027)0.827:0.02444999999999986,(('WP_063656223.1-Aliivibrio-fischeri':0.09659999999999958,'WP_155675904.1-Aliivibrio-fischeri':0.09339999999999993)0.87:0.04286000000000012,((((('WP_076646094.1-Vibrio-sp.-10N.222.47.A9':0.07824999999999971,'WP_155682326.1-Aliivibrio-fischeri':0.1335499999999996)0.181:0.05350000000000055,'WP_102363333.1-Vibrio-cyclitrophicus':0.09163000000000032)0.46:0.047559999999999825,('WP_102554090.1-Vibrio-splendidus':0.10759000000000007,'WP_102459739.1-Vibrio-lentus':0.11939000000000011)0.634:0.04404999999999992)0.806:0.026539999999999786,(('WP_248638627.1-Vibrio-crassostreae':0.09244999999999992,'WP_239937663.1-Vibrio-chagasii':0.19832000000000027)0.28:0.03456999999999999,('WP_016768872.1-Vibrio-cyclitrophicus':0.09724999999999984,('WP_017040654.1-Vibrio-genomosp.-F10':0.17947000000000024,'WP_206625521.1-Vibrio-vulnificus':0.13751000000000024)0.605:0.039839999999999876)0.879:0.05018000000000011)0.778:0.017129999999999868)0.468:0.022090000000000387,('WP_135353160.1-Aliivibrio-fischeri':0.07373999999999992,'WP_137008959.1-Vibrio-kanaloae':0.12805)0.926:0.06464000000000025)0.738:0.015069999999999695)0.773:0.010629999999999917)0.605:0.017039999999999722,((('WP_171343915.1-Vibrio-cyclitrophicus':0.08380999999999972,('WP_048661380.1-Vibrio-crassostreae':0.07024000000000008,'WP_193846925.1-Vibrio-sp.-OPT10':0.0891700000000002)0.788:0.023099999999999454)0.735:0.021570000000000533,('WP_161753058.1-Vibrio-toranzoniae':0.15758000000000028,('WP_016784063.1-Vibrio':0.09450000000000003,('WP_239918548.1-Vibrio-sp.-Isolate22':0.02491000000000021,'WP_108176621.1-Vibrio-splendidus':0.04797000000000029)0.964:0.08103999999999978)0.333:0.04391000000000034)0.762:0.026250000000000107)0.761:0.013799999999999812,(('WP_017449116.1-Vibrio-parahaemolyticus':0.17771999999999988,('WP_108117598.1-Vibrio-splendidus':0.06519999999999992,'WP_137027437.1-Vibrio-kanaloae':0.023130000000000095)0.345:0.016459999999999475)0.808:0.0281100000000003,(('WP_137025093.1-Vibrio-kanaloae':0.1069899999999997,'WP_077679840.1-Vibrio-kanaloae':0.1085799999999999)0.881:0.052160000000000206,((('WP_135455718.1-Vibrio-echinoideorum':0.10298000000000052,'WP_102475754.1-Shewanella-sp.-10N.286.52.C2':0.1632100000000003)0.834:0.031039999999999957,(('WP_239850080.1-Shewanella-sp.-Isolate8':0.12807999999999975,(((('WP_224657569.1-Vibrio-vulnificus':0.11464000000000052,'WP_145570128.1-Vibrio-tasmaniensis':0.1615700000000002)0.613:0.0367999999999995,'WP_102986644.1-Vibrio-diazotrophicus':0.16007999999999978)0.374:0.026860000000000106,('WP_064489977.1-Vibrio-parahaemolyticus':0.04797000000000029,'WP_201976964.1-Aeromonas-enteropelogenes':0.10509000000000013)0.987:0.0911799999999996)0.683:0.014140000000000263,(('WP_102563398.1-Vibrio-sp.-10N.286.49.B3':0.10035000000000016,('WP_120030433.1-Vibrio-sinensis':0.06865000000000032,'WP_173243873.1-Shewanella-sp.-DC2-4':0.16608)0.682:0.03208999999999973)0.829:0.04089000000000009,('WP_020328057.1-Vibrio-fluvialis':0.05620000000000003,'WP_102394203.1-Vibrio-sp.-10N.222.54.B6':0.15012999999999987)0.908:0.05078000000000005)0.847:0.04081000000000001)0.617:0.03462999999999994)0.741:0.03892999999999969,(('WP_208158397.1-Shewanella-algae':0.07354000000000038,'WP_208157589.1-Shewanella-algae':0.03298000000000023)0.933:0.03803999999999963,('WP_041510674.1-Shewanella':0.04074000000000044,'WP_187575439.1-Shewanella-algae':0.05771000000000015)0.877:0.03498999999999963)0.413:0.01980000000000004)0.911:0.04337000000000035)0.743:0.018079999999999874,('WP_161672404.1-Vibrio-toranzoniae':0.07394999999999996,('WP_017094106.1-Vibrio-splendidus':0.09968000000000021,((('WP_114634227.1-Vibrio-splendidus':0.06072999999999995,'WP_048661352.1-Vibrio-crassostreae':0.060939999999999994)0.87:0.08749999999999991,('WP_009847952.1-Vibrio':0.16529999999999978,'WP_233894267.1-Vibrio-kanaloae':0.1291500000000001)0.803:0.038429999999999964)0.916:0.0739399999999999,('WP_248997855.1-Shewanella-marinintestina':0.0525199999999999,'WP_162023628.1-Shewanella-sp.-Arc9-LZ':0.06522000000000006)0.879:0.05099000000000009)0.495:0.02583000000000002)0.878:0.05215999999999976)0.566:0.023540000000000116)0.928:0.046640000000000015)0.0:0.000540)0.933:0.03302000000000005)0.948:0.0461999999999998)0.673:0.011849999999999916)0.786:0.012110000000000287,((((('WP_108189654.1-Vibrio-splendidus':0.16796999999999995,'WP_012532661.1-Aliivibrio-fischeri':0.09184999999999999)0.886:0.04994999999999994,('WP_069440067.1-Pseudoalteromonas-tetraodonis':0.24917000000000034,'WP_155677763.1-Aliivibrio-fischeri':0.08142000000000049)0.786:0.02483999999999975)0.914:0.04997000000000007,(((('WP_061023752.1-Vibrio-splendidus':0.062429999999999986,('WP_110456309.1-Shewanella-algidipiscicola':0.20709,'WP_220768829.1-Shewanella-sp.-MBTL60-007':0.1131899999999999)0.058:0.012849999999999806)0.882:0.03224000000000027,'WP_138589904.1-Pseudoalteromonas-aurantia':0.14778000000000002)0.872:0.03778999999999977,'WP_102420589.1-Vibrio-cyclitrophicus':0.1181000000000001)0.23:0.013840000000000074,('WP_017069285.1-Vibrio-crassostreae':0.08173999999999992,'WP_102503160.1-Vibrio-splendidus':0.14733000000000018)0.797:0.026259999999999728)0.819:0.03459000000000012)0.711:0.04571000000000014,('WP_268674123.1-Vibrio-scophthalmi':0.10708999999999991,'WP_017085924.1-Vibrio-splendidus':0.11161999999999983)0.901:0.053710000000000147)0.382:0.021809999999999885,(('WP_258428294.1-Vibrio-sp.-RM-69-4':0.1048,(('WP_225486226.1-Vibrio-vulnificus':0.15839000000000025,'WP_102428030.1-Vibrio-sp.-10N.261.54.E10':0.1214599999999999)0.93:0.06525000000000025,('WP_193834777.1-Vibrio-sp.-OPT20':0.0577700000000001,('WP_029865567.1-Vibrio-parahaemolyticus':0.06469999999999976,'WP_213967907.1-Vibrio-alginolyticus':0.017669999999999852)0.986:0.0711400000000002)0.036:0.020250000000000323)0.745:0.011859999999999538)0.931:0.05184999999999995,('WP_248368760.1-Vibrio-sp.-1CM8B':0.12745000000000006,(((('WP_102248986.1-Vibrio-tasmaniensis':0.1348900000000004,'WP_135445135.1-Vibrio-tasmaniensis':0.1095900000000003)0.957:0.0942999999999996,('WP_047110243.1-Vibrio-vulnificus':0.22445000000000004,'WP_102313365.1-Vibrio-cyclitrophicus':0.13797999999999977)0.941:0.0975299999999999)0.359:0.019680000000000142,'WP_072616325.1-Vibrio-vulnificus':0.1596000000000002)0.758:0.016449999999999854,('WP_029788868.1-Vibrio':0.2193000000000005,('WP_108162474.1-Vibrio-splendidus':0.044150000000000134,'WP_004742905.1-Vibrio-tubiashii':0.032259999999999955)0.939:0.0703100000000001)0.813:0.04587999999999948)0.748:0.028990000000000293)0.47:0.03092999999999968)0.782:0.01805000000000012)0.772:0.013020000000000032)0.929:0.020809999999999995,((('WP_144217665.1-Shewanella-algae':0.12754999999999983,('WP_172855536.1-Vibrio-coralliilyticus':0.17643999999999993,'WP_045977433.1-Vibrio-neptunius':0.2349000000000001)0.865:0.06034999999999968)0.79:0.03733000000000031,((('WP_012551417.1-Aliivibrio-salmonicida':0.03415000000000035,'WP_012276595.1-Shewanella-halifaxensis':0.041400000000000325)0.95:0.08418999999999954,'WP_135385147.1-Vibrio-tasmaniensis':0.12088999999999972)0.744:0.04427000000000003,'WP_108083222.1-Vibrio-sp.-ZF-223':0.15808)0.475:0.0146200000000003)0.935:0.052059999999999995,(('WP_206974914.1-Vibrio-parahaemolyticus':0.1111399999999998,('WP_058428281.1-Pseudoalteromonas':0.14993000000000034,'WP_017020702.1-Aliivibrio-logei':0.18176000000000014)0.81:0.03291999999999984)0.913:0.05780999999999992,(((('WP_213972370.1-Vibrio-alginolyticus':0.15039999999999987,'WP_017024716.1-Vibrio-rumoiensis':0.16210999999999975)0.859:0.038770000000000415,('WP_047463293.1-Vibrio-fluvialis':0.11882999999999999,('WP_193167845.1-Vibrio-navarrensis':0.24401000000000028,'WP_039841338.1-Vibrio-harveyi-group':0.13036000000000048)0.3:0.06745999999999963)0.842:0.038440000000000474)0.8:0.024699999999999722,('WP_229630939.1-Vibrio-coralliilyticus':0.2237300000000002,'WP_235060641.1-Vibrio-sp.-SS-MA-C1-2':0.19408000000000047)0.857:0.038349999999999884)0.819:0.024290000000000145,(('WP_248468589.1-Shewanella-sp.-1CM18E':0.12563999999999975,('WP_069533349.1-Vibrio-parahaemolyticus':0.10546000000000033,'WP_106651046.1-Shewanella-sp.-Pdp11':0.15957000000000043)0.599:0.03205999999999953)0.709:0.023680000000000145,(('WP_233927909.1-Vibrio-fluvialis':0.1318299999999999,('WP_158146950.1-Vibrio-fluvialis':0.1169399999999996,'WP_103194250.1-Vibrio-vulnificus':0.04488000000000003)0.908:0.05790000000000006)0.882:0.06120999999999999,('WP_102038478.1-Shewanella-sp.-GutCb':0.20759000000000016,'WP_112353352.1-Shewanella-benthica':0.1489900000000004)0.902:0.08087999999999962)0.366:0.05913000000000013)0.628:0.05127000000000015)0.302:0.008519999999999861)0.185:0.012350000000000083)0.781:0.022499999999999964)0.273:0.000530)0.192:0.026340000000000252)0.786:0.013809999999999878)0.756:0.015649999999999942)0.857:0.014920000000000044,('WP_038175370.1-Vibrio-pacinii':0.13198,('WP_070968050.1-Vibrio-sonorensis':0.18706999999999985,'WP_180567593.1-Oceanospirillales':0.1250599999999995)0.848:0.04824000000000028)0.333:0.022850000000000037)0.744:0.006699999999999928);

**Nematode RML-4, with bacterial, eukaryotic, and viral homologs**

**Shown in Figure S6**

ML tree method: IQ-TREE

Sequences: 406

Substitution model: LG+G4

(WP_119768922.1-Herbaspirillum-sp.-K1R23-30:4.4305569198,(((((CAB4124302.1-uncultured-Caudovirales-phage:3.0078064339,WP_149500027.1-Pirellulaceae:0.5332484990)66.7/81:0.0685935114,WP_075082891.1-Mariniblastus-fucicola:0.6493191655)88.8/81:0.1897016897,(((WP_174581714.1-Candidatus-Methylacidithermus-pantelleriae:0.5109308086,(WP_142659230.1-Methylacidimicrobium-tartarophylax:0.0722827936,WP_142525354.1-Methylacidimicrobium-cyclopophantes:0.1989601493)100/100:0.3110556824)99.4/99:0.2954561165,WP_006980331.1-Chthoniobacter-flavus:0.3203429543)82.9/35:0.0787997719,((WP_162030150.1-unclassified-Lentimonas:0.3939141366,((WP_107834426.1-Opitutus-sp.-ER46:0.2135601273,WP_096056354.1-Nibricoccus-aquaticus:0.1961736804)8.7/24:0.0349380384,(WP_069962141.1-Lacunisphaera-limnophila:0.1199035807,WP_129048794.1-Oleiharenicola-lentus:0.0755535795)96.9/100:0.1231223761)97.9/99:0.1398647781)89/92:0.1403904016,((WP_146599621.1-Novipirellula-aureliae:0.3729321716,WP_146593448.1-Novipirellula-galeiformis:0.2515394302)95.7/100:0.1216689037,(((WP_146512969.1-Rubripirellula-amarantea:0.3900950181,((WP_007324562.1-Rhodopirellula-baltica:0.0838690330,WP_008655232.1-Rhodopirellula-europaea:0.0359889342)65.2/100:0.0146601726,WP_007336828.1-Rhodopirellula-baltica:0.0577145408)99.8/100:0.2722681597)81.6/99:0.1196978234,WP_153559344.1-Roseimaritima-sp.-JC651:0.2114353238)88/99:0.0790283222,WP_145097213.1-Rosistilla-carotiformis:0.4613976552)84.2/99:0.0645243327)95.5/100:0.1125753485)68.8/52:0.0525560723)8.9/12:0.0609121119)26/36:0.1276723975,(WP_105335905.1-Blastopirellula-marina:0.1034478421,((WP_114369629.1-Blastopirellula-cremea:0.0988161453,WP_105331861.1-Blastopirellula:0.0869646403)74.2/76:0.0296920122,WP_105358014.1-Blastopirellula-marina:0.0636744568)28/44:0.0324386702)100/100:0.5196675600)87.4/78:0.4711788264,WP_075077632.1-Terrimicrobium-sacchariphilum:0.8221926562)79.7/92:0.7338543568,((((((((((((((XP_002629938.1-Caenorhabditis-briggsae:0.3127649454,XP_013307549.1-Necator-americanus:0.2973681421)99.7/100:0.3835608189,(XP_024500051.1-Strongyloides-ratti:0.3797759863,(XP_020306694.1-Loa-loa:0.1274022046,XP_001899896.1-Brugia-malayi:0.0471350750)100/100:0.5428237105)39.4/99:0.1786450248)100/100:0.6334680232,(XP_024507764.1-Strongyloides-ratti:0.7199732377,XP_013291821.1-Necator-americanus:0.4653972896)99.2/100:0.4779083856)99.8/100:0.4920542884,((QBK88046.1-Marseillevirus-LCMAC202:0.5822726780,(QPI16522.1-Virus-NIOZ-UU159:0.5000424857,ARF10663.1-Hokovirus-HKV1:0.4944368440)30.3/33:0.0406699140)84.3/40:0.0563412371,(AYV84745.1-Hyperionvirus-sp.:0.4987018990,QFG74443.1-Megaviridae-environmental-sample:0.4918693120)82.3/96:0.1095762191)90/99:0.0894463989)86.7/97:0.0715290131,((XP_028886860.1-Trypanosoma-theileri:0.0918779149,(XP_811455.1-Trypanosoma-cruzi-strain-CL-Brener:0.1073126976,(XP_029242692.1-Trypanosoma-rangeli:0.0989043754,XP_029232362.1-Trypanosoma-conorhini:0.0752106958)85.8/100:0.0743343317)98.5/100:0.2028726265)100/100:0.5220394441,((QIG60112.1-Dishui-Lake-large-algae-virus-1:0.7234907103,ARF12483.1-Klosneuvirus-KNV1:0.5189957104)11.5/86:0.0542915048,((7JID_A-Acanthamoeba-polyphaga-mimivirus:0.5883557612,ARF09734.1-Indivirus-ILV1:0.3714876876)59.8/99:0.0927793190,(QKF93545.1-Fadolivirus-1:0.3468958037,ARF12675.1-Klosneuvirus-KNV1:0.4178686382)55.9/99:0.0632021681)93.9/100:0.0841618155)89.4/99:0.0778392276)91.7/92:0.0729078721)83.1/92:0.0898917089,(QBK86789.1-Marseillevirus-LCMAC103:0.7844900709,QBK93652.1-Pithovirus-LCPAC404:0.7660344720)14.7/95:0.0677103764)100/100:0.8573882055,(((XP_001580918.1-Trichomonas-vaginalis-G3:0.5444808053,XP_001317111.1-Trichomonas-vaginalis-G3:0.8055347075)100/100:1.4126512560,(YP_001648294.1-putative-NAD-dependent-epimerase-dehydratase-Ostreococcus-tauri-virus-OtV5_1:0.2210681488,YP_004061822.1-hypothetical-protein-OlV1_190-Ostreococcus-lucimarinus-virus-1_1:0.3157250255)100/100:1.6321061880)99.3/100:0.7307985919,(XP_013421209.1-Lingula-anatina:0.1255718150,XP_013399968.1-Lingula-anatina:0.1206639548)99.5/100:0.4249739678)77.1/90:0.1769766786)71.5/90:0.0442768432,(QBK89631.1-Pithovirus-LCPAC001:0.9364430111,QBK92300.1-Pithovirus-LCPAC304:0.6103189514)94.5/100:0.2159226542)95.5/92:0.0868066092,(XP_014148374.1-Sphaeroforma-arctica-JP610:0.8123879784,(XP_012895844.1-Blastocystis-hominis:0.4570252288,XP_013762229.1-Thecamonas-trahens-ATCC-50062:0.5134559992)89/99:0.1031083309)87.9/62:0.0704522278)71.3/26:0.0211201958,((XP_012756676.1-Acytostelium-subglobosum-LB1:0.3100835498,XP_020434016.1-Heterostelium-album-PN500:0.4178618836)100/100:0.4701414996,XP_001738724.1-Entamoeba-dispar-SAW760:0.5839937001)63.2/27:0.1054023136)77.9/51:0.1210443678,((((XP_003057320.1-Micromonas-pusilla-CCMP1545:0.0734063067,XP_002500589.1-Micromonas-commoda:0.1234301577)100/100:0.4413490755,(XP_007514013.1-Bathycoccus-prasinos:0.2593140886,XP_003084264.2-Ostreococcus-tauri:0.3200189366)57/98:0.1685097121)99.2/100:0.2467450881,((((XP_005648979.1-Coccomyxa-subellipsoidea-C-169:0.2759579398,((XP_002955639.1-Volvox-carteri-f.-nagariensis:0.0769387390,XP_001695032.1-Chlamydomonas-reinhardtii:0.1190721721)98.1/100:0.1062368864,XP_013901107.1-Monoraphidium-neglectum:0.1359961861)65.6/56:0.0229804576)82.2/45:0.0285229050,XP_005851985.1-Chlorella-variabilis:0.1365863996)33.9/50:0.0405843432,XP_011395817.1-Auxenochlorella-protothecoides:0.1528132986)100/100:0.3403581510,(((((((((((((((((((((((((XP_024966115.1-Cynara-cardunculus-var.-scolymus:0.1834439986,XP_024966589.1-Cynara-cardunculus-var.-scolymus:0.0096773114)89.3/100:0.0362210486,(XP_022013454.1-Helianthus-annuus:0.0794711975,XP_023765550.1-Lactuca-sativa:0.0651961910)7.5/91:0.0208752078)94.7/100:0.0434152822,(XP_022028279.1-Helianthus-annuus:0.0219025161,XP_023756522.1-Lactuca-sativa:0.0961621031)89.1/100:0.0245173296)88.5/100:0.0194395448,(XP_015630775.1-Oryza-sativa-Japonica-Group:0.0308901567,XP_035817787.1-Zea-mays:0.0497255075)99.8/100:0.0944833512)86/96:0.0224304781,XP_028106852.1-Camellia-sinensis:0.0444009931)58/61:0.0102437099,((XP_009590265.1-Nicotiana-tomentosiformis:0.0402184879,(XP_015055792.1-Solanum-pennellii:0.0577736130,XP_009610279.1-Nicotiana-tomentosiformis:0.0193693626)91.9/100:0.0239947002)68.1/95:0.0219120448,XP_004244296.1-Solanum-lycopersicum:0.0459227262)74.9/82:0.0066225795)65/77:0.0071369425,(XP_011098163.1-Sesamum-indicum:0.0214212505,XP_022869935.1-Olea-europaea-var.-sylvestris:0.0721322226)82.5/100:0.0101633602)70.1/82:0.0098961538,((((((XP_026399575.1-Papaver-somniferum:0.2230597395,XP_026394899.1-Papaver-somniferum:0.0579797607)95.5/97:0.0533964866,(XP_012850628.1-Erythranthe-guttata:0.0552433133,(XP_017247541.1-Daucus-carota-subsp.-sativus:0.0400661669,XP_017252226.1-Daucus-carota-subsp.-sativus:0.0528379613)94.4/100:0.0433409073)39.1/85:0.0139561439)82.1/83:0.0189523383,XP_027127598.1-Coffea-arabica:0.0539120201)93/96:0.0334264934,XP_019193709.1-Ipomoea-nil:0.0789164381)71.4/89:0.0085356421,XP_019153924.1-Ipomoea-nil:0.0629546973)77.6/85:0.0070332643,((XP_019158937.1-Ipomoea-nil:0.0451046023,XP_031093021.1-Ipomoea-triloba:0.0362281344)100/100:0.1550579920,(XP_022848522.1-Olea-europaea-var.-sylvestris:0.0383974662,XP_028062202.1-Camellia-sinensis:0.1090931439)56.1/90:0.0230565815)66.6/80:0.0153764160)0/67:0.0000028579)90.9/87:0.0143231595,((((XP_028763810.1-Prosopis-alba:0.0731356529,((XP_019455512.1-Lupinus-angustifolius:0.0720628716,XP_020226307.1-Cajanus-cajan:0.0177149057)40.6/38:0.0052038748,XP_014523158.1-Vigna-radiata-var.-radiata:0.0611330771)93.9/100:0.0262339357)33.1/35:0.0108623478,((XP_015954107.1-Arachis-duranensis:0.0525573763,(XP_003531412.1-Glycine-max:0.0426945962,XP_019449231.1-Lupinus-angustifolius:0.0501108543)73.4/97:0.0044788795)0/90:0.0000029104,(XP_004509959.1-Cicer-arietinum:0.0696321982,XP_024630471.1-Medicago-truncatula:0.0374372328)99.2/100:0.0498604673)78.7/94:0.0184874605)77.4/95:0.0080639705,((XP_012836512.1-Erythranthe-guttata:0.0919570878,XP_011082806.1-Sesamum-indicum:0.0115018162)95/95:0.0317745350,XP_028755472.1-Prosopis-alba:0.0491041123)86.2/95:0.0124246527)84.4/93:0.0129502867,XP_038690143.1-Tripterygium-wilfordii:0.0636906396)92.8/94:0.0236553304)86/88:0.0106707587,(((((((XP_035549685.1-Juglans-regia:0.2297260230,XP_040986737.1-Juglans-microcarpa-x-Juglans-regia:0.0364312593)0/78:0.0000022691,(((XP_034907656.1-Populus-alba:0.0378327328,XP_034886774.1-Populus-alba:0.0356649592)96.5/100:0.0440100674,(XP_021607919.1-Manihot-esculenta:0.0309965472,XP_021595297.1-Manihot-esculenta:0.0862788404)90/100:0.0295072704)85.3/100:0.0195166364,XP_015582142.2-Ricinus-communis:0.0308488285)85.3/100:0.0101237606)77.8/96:0.0052234424,((XP_030460928.1-Syzygium-oleosum:0.0438332356,XP_010060763.2-Eucalyptus-grandis:0.0421278653)98.1/100:0.0462591532,(((((XP_017626421.1-Gossypium-arboreum:0.0806389834,XP_039006232.1-Hibiscus-syriacus:0.0727378161)66.7/96:0.0090529045,XP_017642129.1-Gossypium-arboreum:0.0645127902)53.1/46:0.0110683415,XP_017641827.1-Gossypium-arboreum:0.0375547790)74.6/35:0.0082401221,XP_021288242.1-Herrania-umbratica:0.0486194517)71.2/36:0.0077031298,XP_022734139.1-Durio-zibethinus:0.0522683428)91.1/85:0.0233271711)84.5/33:0.0121317175)76/32:0.0048450251,XP_021911421.1-Carica-papaya:0.0643196860)69.3/38:0.0059102258,XP_006442344.1-Citrus-clementina:0.0620903238)79.1/36:0.0111210421,(((((((XP_010502125.1-Camelina-sativa:0.0096275786,XP_013644676.1-Brassica-napus:0.0757386228)79.1/83:0.0171499238,XP_010535018.1-Tarenaya-hassleriana:0.0686072906)35.6/60:0.0148706726,XP_010552134.1-Tarenaya-hassleriana:0.0369130608)99.8/100:0.0635612171,XP_031405972.1-Punica-granatum:0.0613523352)74.9/86:0.0088417329,((XP_009376726.1-Pyrus-x-bretschneideri:0.0459654766,(XP_024158479.1-Rosa-chinensis:0.0314567814,XP_008226345.1-Prunus-mume:0.0447342288)66/100:0.0205597984)97.1/100:0.0490685577,XP_034679245.1-Vitis-riparia:0.0477627564)87.4/99:0.0201214775)85.5/85:0.0104102805,(XP_030490411.1-Cannabis-sativa:0.0887249617,XP_010093448.1-Morus-notabilis:0.0520909049)83.4/97:0.0146782591)75.4/84:0.0053449837,((XP_031254775.1-Pistacia-vera:0.0691995671,((((XP_010537694.1-Tarenaya-hassleriana:0.0551520893,(XP_010472689.1-Camelina-sativa:0.0329776094,XP_009149137.1-Brassica-rapa:0.0577828084)86.8/99:0.0215279088)92.5/99:0.0301567823,((XP_010671686.1-Beta-vulgaris-subsp.-vulgaris:0.0520614400,XP_010675264.1-Beta-vulgaris-subsp.-vulgaris:0.0312596630)34.2/97:0.0111391523,(XP_021842589.1-Spinacia-oleracea:0.0654693254,XP_021726937.1-Chenopodium-quinoa:0.0233913197)24.8/99:0.0129674635)92/100:0.0262829477)92/98:0.0260803105,XP_022943017.1-Cucurbita-moschata:0.0599076773)85.2/87:0.0149908749,XP_040960845.1-Gossypium-hirsutum:0.0593600252)75.1/38:0.0048119244)83.6/38:0.0120622330,XP_023887068.1-Quercus-suber:0.0453937414)80.3/35:0.0099752635)77.6/11:0.0048896685)13.9/20:0.0108859437,XP_030445623.1-Syzygium-oleosum:0.0689066287)76/91:0.0067377878)0/86:0.0000026370,(XP_031494579.1-Nymphaea-colorata:0.0664556641,XP_031498803.1-Nymphaea-colorata:0.1185548609)89.5/97:0.0413755799)90/95:0.0185651857,XP_010273412.1-Nelumbo-nucifera:0.0598969823)70.4/92:0.0119022090,XP_009421279.1-Musa-acuminata-subsp.-malaccensis:0.0504697304)92.4/92:0.0239211844,(((((((XP_019176650.1-Ipomoea-nil:0.0328194920,XP_031123431.1-Ipomoea-triloba:0.0738337255)83.8/100:0.0256249853,(XP_019176466.1-Ipomoea-nil:0.0549315094,XP_031122529.1-Ipomoea-triloba:0.0750175714)98.5/100:0.0866964144)100/100:0.3986631626,XP_030932284.1-Quercus-lobata:0.1593942419)98.2/97:0.1097247301,XP_020106036.1-Ananas-comosus:0.0594932948)89.6/92:0.0316007550,(XP_020099497.1-Ananas-comosus:0.0404701251,XP_020691264.1-Dendrobium-catenatum:0.1110122216)33/82:0.0098925923)46.5/62:0.0087886518,(XP_010915435.1-Elaeis-guineensis:0.0585678741,(XP_020088693.1-Ananas-comosus:0.0421842468,XP_020097680.1-Ananas-comosus:0.1317507655)82.4/84:0.0151591379)78.4/69:0.0126006475)0/85:0.0000028662,((((((XP_009394163.1-Musa-acuminata-subsp.-malaccensis:0.0633698943,(XP_009395981.1-Musa-acuminata-subsp.-malaccensis:0.0487277881,XP_009394841.1-Musa-acuminata-subsp.-malaccensis:0.0456603943)41.1/96:0.0051180571)91.2/96:0.0265027588,XP_020683487.1-Dendrobium-catenatum:0.1138614179)50.7/99:0.0132446674,XP_039118740.1-Dioscorea-cayenensis-subsp.-rotundata:0.0765691687)84.1/89:0.0128783747,((((XP_039836094.1-Panicum-virgatum:0.0236986703,XP_039834344.1-Panicum-virgatum:0.0575542574)94.5/100:0.0332789208,XP_040376729.1-Oryza-brachyantha:0.0489625251)74.5/100:0.0136521040,XP_003571828.1-Brachypodium-distachyon:0.0408850162)80.8/100:0.0171268423,(XP_020245360.1-Asparagus-officinalis:0.0392611060,XP_020265382.1-Asparagus-officinalis:0.0838057362)95/99:0.0364429288)86.4/83:0.0115835288)80.3/72:0.0106510922,(XP_039116054.1-Dioscorea-cayenensis-subsp.-rotundata:0.0640833140,XP_039122405.1-Dioscorea-cayenensis-subsp.-rotundata:0.0409386566)90.6/82:0.0187929060)81.3/90:0.0094109980,(XP_038976488.1-Phoenix-dactylifera:0.0293623339,XP_029120148.1-Elaeis-guineensis:0.1485001443)89.6/100:0.0242477348)88.1/89:0.0152415259)81.3/89:0.0108277972)57.8/82:0.0095817216,((XP_031393943.1-Punica-granatum:0.0922917545,XP_030466822.1-Syzygium-oleosum:0.0616134599)76.6/100:0.0107992901,XP_024042451.1-Citrus-clementina:0.0524520360)79.8/93:0.0232414762)79.7/76:0.0142679219,((((((XP_009356708.1-Pyrus-x-bretschneideri:0.0994890829,XP_009377674.1-Pyrus-x-bretschneideri:0.0220816758)94.4/100:0.0470320398,XP_024181512.1-Rosa-chinensis:0.0372421313)93.8/100:0.0321875551,XP_021898354.1-Carica-papaya:0.1769216433)74.5/93:0.0070384573,XP_010109352.1-Morus-notabilis:0.0555873372)67.6/87:0.0101065633,XP_030482186.1-Cannabis-sativa:0.0897116164)89.9/91:0.0234603510,(((XP_022147970.1-Momordica-charantia:0.2327591207,(((XP_017255528.1-Daucus-carota-subsp.-sativus:0.2430707208,XP_039058813.1-Hibiscus-syriacus:0.0649752127)98.2/96:0.0906998449,(XP_039044917.1-Hibiscus-syriacus:0.0340677281,XP_017604897.1-Gossypium-arboreum:0.0492277059)87.3/100:0.0217865999)73.7/96:0.0077295710,XP_021274443.1-Herrania-umbratica:0.0644239071)88.3/96:0.0306205932)81.9/95:0.0165636668,XP_023917946.1-Quercus-suber:0.0577089039)0/32:0.0067477990,XP_018836873.1-Juglans-regia:0.0519496256)83.5/95:0.0224940124)25.4/75:0.0096192328)93.5/86:0.0337534752,XP_027063112.1-Coffea-arabica:0.1243004890)9.6/37:0.0065508090,XP_006845495.1-Amborella-trichopoda:0.0957281856)10.9/74:0.0258121683,(((((((XP_006483733.3-Citrus-sinensis:0.2755075435,XP_031276519.1-Pistacia-vera:0.1796245929)85.8/93:0.0411103554,(XP_021903120.1-Carica-papaya:0.1599172806,XP_007011619.2-Theobroma-cacao:0.1176628073)75.9/52:0.0140092362)53.7/49:0.0078311120,(((XP_021596157.1-Manihot-esculenta:0.0893611581,XP_021673389.1-Hevea-brasiliensis:0.0328341699)91/100:0.0290742843,XP_012076356.1-Jatropha-curcas:0.0570789433)97.7/100:0.0754821086,(XP_026382559.1-Papaver-somniferum:0.1110118321,XP_034929454.1-Populus-alba:0.0854628262)61.3/98:0.0342574903)91.5/93:0.0390175930)87/50:0.0196883625,((((((((XP_038696288.1-Tripterygium-wilfordii:0.0378998721,XP_038712281.1-Tripterygium-wilfordii:0.0459594123)100/100:0.1116666188,XP_028112986.1-Camellia-sinensis:0.1173524313)53.4/86:0.0210665911,XP_015867255.1-Ziziphus-jujuba:0.0926232263)0/16:0.0139523888,(((XP_028802307.1-Prosopis-alba:0.1602194511,XP_025657523.1-Arachis-hypogaea:0.0735292936)49/97:0.0196273375,(XP_019435147.1-Lupinus-angustifolius:0.1396917834,(((XP_004498563.1-Cicer-arietinum:0.0478845758,XP_003588651.1-Medicago-truncatula:0.0552863996)97.5/100:0.0453622241,XP_027333604.1-Abrus-precatorius:0.0751420860)73/51:0.0068692156,XP_003545554.1-Glycine-max:0.0326762803)36.7/50:0.0065354609)35.8/80:0.0172924926)87.6/83:0.0198451389,XP_014504303.2-Vigna-radiata-var.-radiata:0.0287391581)99.7/100:0.0775257005)82.2/94:0.0131090288,(XP_031130453.1-Ipomoea-triloba:0.1610118122,(((XP_034199488.1-Prunus-dulcis:0.0329022333,XP_008393937.2-Malus-domestica:0.0499469490)97.7/100:0.0452738656,XP_024168007.1-Rosa-chinensis:0.0359949781)79.2/100:0.0179211704,XP_004292875.1-Fragaria-vesca-subsp.-vesca:0.0475056200)70.3/91:0.0176733693)87.5/87:0.0324168502)91.6/83:0.0239191604,XP_002271970.1-Vitis-vinifera:0.0759948874)0/35:0.0000021617,XP_010268536.1-Nelumbo-nucifera:0.1063461376)70.8/49:0.0066321584,XP_011649755.1-Cucumis-sativus:0.0998021621)56.5/53:0.0162759777)77.3/88:0.0082681379,((XP_024986874.1-Cynara-cardunculus-var.-scolymus:0.1444851418,XP_022036513.1-Helianthus-annuus:0.0838905715)41.4/92:0.0298248096,XP_023751163.1-Lactuca-sativa:0.0923303431)100/100:0.1719011959)69.9/85:0.0091181940,XP_023913627.1-Quercus-suber:0.0948075338)93.4/97:0.0518123475,(XP_016499075.1-Nicotiana-tabacum:0.0609106813,(XP_006355193.1-Solanum-tuberosum:0.0528507369,XP_016554776.1-Capsicum-annuum:0.0472805080)17.3/99:0.0174418937)99.9/100:0.1056192917)77.1/85:0.0434656840)95.1/97:0.0561145305,XP_002964674.1-Selaginella-moellendorffii:0.2109016150)87.3/97:0.0418053892,(XP_024378587.1-Physcomitrium-patens:0.1334890365,(XP_024365289.1-Physcomitrium-patens:0.0441414032,(XP_024377136.1-Physcomitrium-patens:0.0457966104,XP_024375765.1-Physcomitrium-patens:0.0377903154)81.2/100:0.0177930819)98.9/100:0.0834021462)96.9/100:0.0732883870)14.1/89:0.0409559907,(((((((((((((((((((XP_020261993.1-Asparagus-officinalis:0.3808860576,(XP_039854304.1-Panicum-virgatum:0.0362862038,(XP_037451911.1-Triticum-dicoccoides:0.0188072207,XP_015627488.1-Oryza-sativa-Japonica-Group:0.0608562000)98.2/100:0.0523926345)1/77:0.0173935748)61.9/93:0.0163323768,((XP_008776517.2-Phoenix-dactylifera:0.0490961819,XP_008776518.3-Phoenix-dactylifera:0.0325032504)73.9/79:0.0051160067,XP_010933908.1-Elaeis-guineensis:0.0425038216)90/100:0.0229388397)78.5/44:0.0296205211,(XP_020085231.1-Ananas-comosus:0.1428882738,(XP_009389578.1-Musa-acuminata-subsp.-malaccensis:0.0403704554,XP_009400458.1-Musa-acuminata-subsp.-malaccensis:0.0807247401)73.9/47:0.0086063414)69.6/46:0.0086685244)53.8/51:0.0211875474,(XP_020702727.1-Dendrobium-catenatum:0.0457956254,XP_020599681.1-Phalaenopsis-equestris:0.0980166199)95.4/100:0.0484859076)95.1/97:0.0474156135,(XP_010053354.2-Eucalyptus-grandis:0.0365248851,XP_030476172.1-Syzygium-oleosum:0.0457175187)95.6/100:0.0433935463)89.2/87:0.0286960994,XP_010272587.1-Nelumbo-nucifera:0.0629528153)75.4/77:0.0109757874,XP_028063311.1-Camellia-sinensis:0.2743441575)11.6/47:0.0035447790,((((XP_040988144.1-Juglans-microcarpa-x-Juglans-regia:0.0636090793,XP_040990363.1-Juglans-microcarpa-x-Juglans-regia:0.0645474756)47.8/86:0.0253853398,(XP_021901871.1-Carica-papaya:0.0742220576,(((XP_010533730.1-Tarenaya-hassleriana:0.0422581270,XP_010553490.1-Tarenaya-hassleriana:0.0384975517)95.4/100:0.0366411288,XP_013742606.1-Brassica-napus:0.0509509527)68.8/99:0.0089618882,XP_006391796.1-Eutrema-salsugineum:0.0392133089)96.5/100:0.0379410864)87.2/95:0.0303075695)23.1/78:0.0103671810,((XP_038687038.1-Tripterygium-wilfordii:0.0361535591,XP_038678971.1-Tripterygium-wilfordii:0.0480185400)94/100:0.0309513584,(((XP_012087027.1-Jatropha-curcas:0.0415398625,XP_021630406.1-Manihot-esculenta:0.0393531688)67.9/83:0.0134817994,XP_021605124.1-Manihot-esculenta:0.0295180751)77.4/86:0.0155108914,XP_002509687.1-Ricinus-communis:0.0470696090)93/100:0.0266222510)74.8/82:0.0099153701)80.5/69:0.0094803948,(((XP_021844479.1-Spinacia-oleracea:0.0562412349,XP_010676576.1-Beta-vulgaris-subsp.-vulgaris:0.0717222308)84.7/100:0.0227862031,XP_039138369.1-Dioscorea-cayenensis-subsp.-rotundata:0.1211530896)96.5/99:0.0429586193,XP_034888031.1-Populus-alba:0.0392313934)88.3/78:0.0196430038)76.6/61:0.0150773894)76.7/53:0.0085485701,XP_010108054.1-Morus-notabilis:0.0440567300)81.5/77:0.0101088550,((XP_026380000.1-Papaver-somniferum:0.1402243024,XP_028107302.1-Camellia-sinensis:0.0208634292)83.7/75:0.0361697153,XP_028098644.1-Camellia-sinensis:0.0614124104)94.2/95:0.0363492249)82.9/63:0.0103104432,((XP_027080882.1-Coffea-arabica:0.0548945957,((((XP_024994198.1-Cynara-cardunculus-var.-scolymus:0.0660400939,((XP_024985704.1-Cynara-cardunculus-var.-scolymus:0.0462454723,XP_023752083.1-Lactuca-sativa:0.0460384259)0/76:0.0000020168,XP_022022173.1-Helianthus-annuus:0.0930268922)99.1/99:0.0558707236)87.3/97:0.0292233728,XP_023752812.1-Lactuca-sativa:0.0566194793)59.8/83:0.0251654601,XP_022002185.1-Helianthus-annuus:0.0749621402)47.5/82:0.0093048124,XP_024958708.1-Cynara-cardunculus-var.-scolymus:0.0250993527)99.5/100:0.0763463367)89.1/94:0.0234699883,XP_031124697.1-Ipomoea-triloba:0.0276738471)79.9/78:0.0098765875)74.8/44:0.0052713741,(XP_031125635.1-Ipomoea-triloba:0.0555209620,((XP_004245662.1-Solanum-lycopersicum:0.0665071991,XP_016567886.1-Capsicum-annuum:0.0448010348)59.4/100:0.0268206175,XP_019247171.1-Nicotiana-attenuata:0.0579187450)94.8/100:0.0445375388)91.4/99:0.0243734332)60/42:0.0053583347,((XP_008238760.2-Prunus-mume:0.0438558741,XP_008373757.2-Malus-domestica:0.0507638317)95/100:0.0377100636,((XP_019421934.1-Lupinus-angustifolius:0.0756980045,XP_015964276.1-Arachis-duranensis:0.0341220974)93.7/95:0.0308615957,XP_012838043.1-Erythranthe-guttata:0.0707030580)0/63:0.0000028649)90.9/94:0.0196554436)0/57:0.0000028994,(((XP_017230733.1-Daucus-carota-subsp.-sativus:0.0745894883,XP_017235998.1-Daucus-carota-subsp.-sativus:0.0669128959)98.7/97:0.0634331460,XP_034709549.1-Vitis-riparia:0.0359558987)82.5/73:0.0131149862,(XP_022854995.1-Olea-europaea-var.-sylvestris:0.0324742279,XP_011072055.1-Sesamum-indicum:0.0442181139)89.6/98:0.0219669576)90.6/57:0.0248473965)78.7/78:0.0157256546,(XP_031284170.1-Pistacia-vera:0.0624105832,(XP_023876864.1-Quercus-suber:0.0534418785,XP_015867767.1-Ziziphus-jujuba:0.0521121994)61.6/95:0.0048991618)90.8/99:0.0181064928)0/75:0.0000027627,XP_006440508.1-Citrus-clementina:0.0517031533)53.4/28:0.0150798234,((((((XP_020217507.1-Cajanus-cajan:0.0382075714,(XP_027913194.1-Vigna-unguiculata:0.0611119863,XP_007158451.1-Phaseolus-vulgaris:0.0420553881)95.5/100:0.0483667268)27.5/39:0.0140638355,XP_028246958.1-Glycine-soja:0.0474655024)99.3/100:0.0718727374,(XP_004515790.1-Cicer-arietinum:0.0356559189,XP_013457661.1-Medicago-truncatula:0.0447699513)97.4/100:0.0504346205)80.1/11:0.0134381595,(XP_015964685.1-Arachis-duranensis:0.1313835103,XP_008437195.1-Cucumis-melo:0.0559907949)56.6/11:0.0275735363)79.6/66:0.0066352559,XP_028762773.1-Prosopis-alba:0.0747412431)71.7/47:0.0042759501,(XP_024171332.1-Rosa-chinensis:0.0820160159,XP_031397838.1-Punica-granatum:0.0529532627)87.3/94:0.0161784564)74.2/75:0.0056395146)82.9/51:0.0117566794,((XP_039009698.1-Hibiscus-syriacus:0.0838807734,XP_012471878.1-Gossypium-raimondii:0.0289810891)92.3/99:0.0325808938,((XP_021277974.1-Herrania-umbratica:0.0533944178,XP_022738126.1-Durio-zibethinus:0.0264832564)82.4/97:0.0127481849,XP_022719777.1-Durio-zibethinus:0.0350992336)80/97:0.0165420451)98.9/100:0.0672668961)15.2/39:0.0063591384,((XP_027336418.1-Abrus-precatorius:0.0434659876,XP_028211060.1-Glycine-soja:0.0657312926)88.2/100:0.0321122817,XP_007155573.1-Phaseolus-vulgaris:0.0714227493)96.2/97:0.0445706897)0/54:0.0192888632)0/43:0.0000026691,XP_006857961.1-Amborella-trichopoda:0.0844742384)71.7/83:0.0394066239,XP_031495988.1-Nymphaea-colorata:0.0997273943)82.1/94:0.0663463083,XP_030500851.1-Cannabis-sativa:0.0895216274)99/100:0.1848794812)87.5/98:0.0901203445)95.7/98:0.1706075847,(XP_009039633.1-Aureococcus-anophagefferens:0.5588458717,XP_005826217.1-Guillardia-theta-CCMP2712:0.5125497558)92.6/71:0.1616216856)67.2/26:0.0788718410)42.4/10:0.0469206704,(XP_005537979.1-Cyanidioschyzon-merolae-strain-10D:0.2590308563,XP_005707655.1-Galdieria-sulphuraria:0.3489297635)100/100:0.4151894534)85.2/16:0.0593088051,((((XP_003663944.1-Thermothelomyces-thermophilus-ATCC-42464:0.5633989497,(XP_007806273.1-Endocarpon-pusillum-Z07020:0.1541497300,XP_031903416.1-Aspergillus-alliaceus:0.2251664506)99.8/100:0.4607643059)100/100:0.4406618250,XP_025386083.1-Aspergillus-eucalypticola-CBS-122712:0.6928065115)99.5/100:0.3776083271,((((((((((XP_007929610.1-Pseudocercospora-fijiensis-CIRAD86:0.2391298347,(XP_033662844.1-Zasmidium-cellare-ATCC-36951:0.2526513629,XP_023626149.1-Ramularia-collo-cygni:0.2484864666)79.4/100:0.0945169007)99.7/100:0.2934336909,XP_007677597.1-Baudoinia-panamericana-UAMH-10762:0.4279129752)99.4/100:0.3422282703,XP_026600440.1-Aspergillus-mulundensis:0.6177090062)89.3/99:0.0956357194,(XP_033538296.1-Eremomyces-bilateralis-CBS-781.70:0.3930868099,(XP_008714317.1-Cyphellophora-europaea-CBS-101466:0.1569103036,XP_018001010.1-Phialophora-attinorum:0.0732852567)99.8/100:0.2728588702)66.4/96:0.0558630603)78.3/94:0.0554963449,XP_007682126.1-Bipolaris-oryzae-ATCC-44560:0.4626513938)29.6/89:0.0164744487,((((((XP_006966750.1-Trichoderma-reesei-QM6a:0.0539915266,XP_024752113.1-Trichoderma-citrinoviride:0.0674385336)94.5/100:0.0443221600,(XP_013959583.1-Trichoderma-virens-Gv29-8:0.0306503587,XP_024774633.1-Trichoderma-harzianum-CBS-226.95:0.0451252399)91.4/100:0.0255864224)91.7/100:0.0302892405,(XP_013937979.1-Trichoderma-atroviride-IMI-206040:0.0334388464,XP_024760396.1-Trichoderma-asperellum-CBS-433.97:0.0424334218)75/100:0.0141691712)99.9/100:0.1764867724,XP_018174549.1-Purpureocillium-lilacinum:0.2295378921)28.1/49:0.0667314698,((XP_018180926.1-Purpureocillium-lilacinum:0.2393542514,((XP_031031860.1-Fusarium-oxysporum-NRRL-32931:0.0281540164,XP_018254302.1-Fusarium-oxysporum-f.-sp.-lycopersici-4287:0.0657798214)97.5/100:0.0602603247,(XP_036532672.1-Fusarium-subglutinans:0.0520458537,(XP_018761705.1-Fusarium-verticillioides-7600:0.0652035477,XP_041690159.1-Fusarium-mangiferae:0.0380609362)60.1/100:0.0174280973)55.9/99:0.0152151654)99.9/100:0.1689109146)39.1/53:0.0563908973,(XP_011320564.1-Fusarium-graminearum-PH-1:0.2780921273,XP_003044390.1-Fusarium-vanettenii-77-13-4:0.1290896639)68.8/86:0.0597399750)94.2/96:0.0836989019)93.1/100:0.0472396683,((XP_016587658.1-Sporothrix-schenckii-1099-18:0.0917658974,XP_014171746.1-Grosmannia-clavigera-kw1407:0.1088898250)97.7/100:0.0866108670,(XP_006695147.1-Chaetomium-thermophilum-var.-thermophilum-DSM-1495:0.1173705777,(((XP_003654153.1-Thermothielavioides-terrestris-NRRL-8126:0.1003196834,XP_003665385.1-Thermothelomyces-thermophilus-ATCC-42464:0.0786143510)50.5/73:0.0271502631,XP_001226132.1-Chaetomium-globosum-CBS-148.51:0.0802615325)94.6/95:0.0472939076,((((XP_028467777.1-Sodiomyces-alkalinus-F11:0.1696897363,XP_016641259.1-Scedosporium-apiospermum:0.0545653534)94.4/99:0.0535265899,((XP_009226599.1-Gaeumannomyces-tritici-R3-111a-1:0.0753739910,XP_030983137.1-Pyricularia-grisea:0.0855872837)96.8/100:0.0611160042,(((((XP_018126611.1-Pseudogymnoascus-verrucosus:0.1506731058,(((XP_008080873.1-Glarea-lozoyensis-ATCC-20868:0.1285707592,(XP_031871468.1-Venustampulla-echinocandica:0.0162530494,XP_031003915.1-Lachnellula-hyalina:0.0839523464)22.9/92:0.0265824234)85.2/91:0.0395264535,((XP_007297111.1-Marssonina-brunnea-f.-sp.-multigermtubi-MB_m1:0.1275812690,XP_018073143.1-Mollisia-scopiformis:0.0216175332)91.6/97:0.0484376317,XP_024728048.1-Hyaloscypha-bicolor-E:0.0551793581)67/85:0.0214518865)84.7/86:0.0261201793,XP_024719770.1-Amorphotheca-resinae-ATCC-22711:0.0341535808)83.9/88:0.0265852584)45.7/91:0.0275723699,XP_038767199.1-Botrytis-porri:0.1038326432)98.1/100:0.0614695078,((XP_033441339.1-Daldinia-childiae:0.1410371068,(XP_007833569.1-Pestalotiopsis-fici-W106-1:0.0920851573,XP_040711048.1-Pseudomassariella-vexata:0.0903406797)75.4/77:0.0225129065)98.9/100:0.0878799816,(XP_003002732.1-Verticillium-alfalfae-VaMs.102:0.1016151034,((XP_038746351.1-Colletotrichum-karsti:0.0392065956,XP_022477045.1-Colletotrichum-orchidophilum:0.0375013976)86.1/100:0.0173927394,(XP_008095879.1-Colletotrichum-graminicola-M1.001:0.0400803542,XP_018159045.1-Colletotrichum-higginsianum-IMI-349063:0.0367835672)91.9/59:0.0233078537)87.4/58:0.0302357007)71.9/72:0.0446266488)48.8/91:0.0093263644)92.8/92:0.0332087685,XP_030996958.1-Phialemoniopsis-curvata:0.0819487563)79.6/74:0.0344536741,(XP_040773122.1-Cryphonectria-parasitica-EP155:0.1201514411,XP_007912887.1-Phaeoacremonium-minimum-UCRPA7:0.0190722093)95.9/100:0.0528815281)95.3/85:0.0496598595)0/26:0.0131933157)56/86:0.0339305196,XP_001905352.1-Podospora-anserina-S-mat_:0.0516912332)90.3/86:0.0262476815,XP_001728253.1-Neurospora-crassa-OR74A:0.1030460438)87.6/86:0.0226377261)83/81:0.0288954181)42.7/88:0.0162692984)82.6/92:0.0394153161)90.2/100:0.0565505075)59.8/78:0.0310692068,((XP_013342528.1-Aureobasidium-subglaciale-EXF-2481:0.1970839126,XP_013423002.1-Aureobasidium-namibiae-CBS-147.97:0.1045340635)0/83:0.0081175285,XP_040880347.1-Aureobasidium-melanogenum-CBS-110374:0.0524147303)99.9/100:0.1442526957)88.4/92:0.0489617019,XP_002836423.1-Tuber-melanosporum-Mel28:0.2039624230)64.2/92:0.0698733664,XP_011117055.1-Orbilia-oligospora-ATCC-24927:0.2218051628)94.3/100:0.1082079894,(XP_016609714.1-Spizellomyces-punctatus-DAOM-BR117:0.2876807297,XP_040727330.1-Protomyces-lactucae-debilis:0.3713011475)25.1/34:0.0589892488)95.8/93:0.1675195030)51.8/88:0.1068629648,(XP_012204866.1-Saprolegnia-parasitica-CBS-223.65:0.0484928306,XP_008610980.1-Saprolegnia-diclina-VS20:0.0468143771)100/100:1.1616012960)89.5/18:0.0854344837)88.6/89:0.1422184396,((YP_009329336.1-NAD-dependent-epimerase-dehydratase-putative-Cedratvirus-A11_1:0.0960915562,QIN54504.1-Cedratvirus-kamchatka:0.0652472931)65.7/92:0.0183847019,SPN78831.1-Brazilian-cedratvirus-IHUMI:0.0120497027)99.8/100:0.7071937161)100/100:1.2832790950);
